# A *Trypanosoma brucei* ORFeome-based Gain-of-Function Library reveals novel genes associated with melarsoprol resistance

**DOI:** 10.1101/849042

**Authors:** M Carter, HS Kim, S Gomez, S Gritz, S Larson, D Schulz, GA Hovel-Miner

**Affiliations:** The George Washington University, Department of Microbiology, Immunology, and Tropical Medicine, 2300 Eye St., Washington, D.C, 20037; The Public Health Research Institute at the International Center for Public Health New Jersey Medical School – Rutgers, The State University of New Jersey, 225 Warren Street, Newark, NJ 07103-3535; Harvey Mudd College, Department of Biology, 1250 N. Dartmouth Ave., Claremont, CA 91711

## Abstract

*Trypanosoma brucei* is an early branching protozoan that causes Human and Animal African Trypanosomiasis. Forward genetics approaches are powerful tools for uncovering novel aspects of Trypanosomatid biology, pathogenesis, and therapeutic approaches against trypanosomiasis. Here we have generated a *T. brucei* ORFeome consisting of over 90% of the targeted genome and used it to make an inducible Gain-of-Function library for broadly applicable forward genetic screening. Using a critical drug of last resort, melarsoprol, we conducted a proof of principle genetic screen. Hits arising from this screen support the significance of trypanothione, a key player in redox metabolism, as a target of melarsoprol and implicate novel proteins of the flagellum and mitochondria in drug resistance. This study has produced two powerful new genetic tools for kinetoplastida research, which are expected to promote major advances in kinetoplastida biology and therapeutic development in the years to come.

## INTRODUCTION

Trypanosomatids are a major parasitic lineage that include the African trypanosomes, American trypanosomes, and Leishmania spp. (family *Trypanosomatidae*, order *Kinetoplastida*), which collectively cause death and disease in millions of people living in tropical and sub-tropical regions (1). There are no vaccines against this family of parasites and the limited number of anti-Trypanosomatid drugs present ongoing challenges of host toxicity, complex treatment regimens, and burgeoning drug resistance (2). These evolutionarily ancient eukaryotes have highly divergent genomes from well-established model organisms with more than 50% of open reading frames (ORFs) annotated as hypothetical (3). Trypanosomatid parasites appear to have diverged from a shared ancestor around 100 million years ago, yet have retained a high proportion of orthologous gene clusters and chromosomal synteny. Of the 9,068 genes in the *Trypanosoma brucei* (African trypanosome) genome, 6158 are orthologous with both *Trypanosoma cruzi* (American trypanosome) and *Leishmania major* (3). Many of the nonsyntenic regions of the genome appear to be directly associated with organism specific aspects of parasitism, such as surface antigen variation, leaving many biological functions highly conserved among kinetoplastida parasites (3). Trypanosomatids share essential subcellular structures, such as the flagellum, glycosomes (specialized organelles of glycolysis), and the kinetoplast (a network of mitochondrial DNA). While reverse genetics based on well-established models can promote discrete advances, forward genetics approaches have the potential to uncover important aspects of kinetoplastida biology shared among orthologous genes.

*T. brucei,* the causative agent of Human African trypanosomiasis (HAT), has traditionally been the most genetically tractable of the Trypanosomatid parasites. Discoveries made in *T. brucei* have provided critical understanding to the biology of other kinetoplastida as well as providing insights into eukaryotic evolution (4–7) and uncovered broadly significant biological processes, such as GPI-anchors and mitochondrial RNA-editing (8–10). For the past decade a whole-genome interfering-RNA (RNAi) knock-down library has been the primary forward-genetics tool in *T. brucei* resulting in identification of essential genes and genes associated with drug resistance and pathogenesis, to name a few (11–13). A strength of the RNAi library and associated RNA-Interference Targeted (RIT)-seq approaches is the identification of genes that result in a loss-of-fitness phenotype (14). However, knockdown strategies may fail when screening for a cellular event that requires a particular gene to be turned on rather than turned off. Furthermore, they preclude identification of essential genes associated with key processes (14). A Gain-of-Function library approach may be more effective in the identification of genes whose expression is critical to the initiation of the event. For example, Gain-of-Function screens have been employed to identify drug targets, as well as resistance mechanisms (15–17), and in areas of basic biology have been critical for discoveries in the areas of chromosome segregation and cell cycle, signal transduction, transcriptional regulation, cell polarity, and stem cell biology (18). Thus, Gain-of-Function screens are a powerful tool that can be used as a complement to loss-of-function screening to uncover new biology in diverse areas.

Traditional methods of inducible gene library formation by cDNA synthesis and cloning are not viable for *T. brucei* as most gene expression regulation in kinetoplastida occurs post-transcriptionally, with 5’ and 3’UTRs playing a major role in determining steady state levels of their associated transcripts (19). Existing overexpression libraries for *T. brucei*, generated by whole-genome fragmentation methods have demonstrated the usefulness of this approach, for example, in the identification of the benzoxaborole drug target (polyadenylation specificity factor, CPSF3), despite their somewhat limited ability to control individual ORF expression (20–23). In addition, generation of an ORFeome provides a tool that can be applied to the downstream generation of multiple whole-genome methodologies including yeast 2-hybrid libraries, tagging libraries, inducible expression libraries for Gain-of-Function studies, and complementation for loss-of-function mutant phenotypes (24–27). A *T. brucei* ORFeome would permit production of an inducible Gain-of-Function library, which is expected to be an ideal design for the identification of novel gene functions based on observable phenotypes in large scale screens.

African trypanosomiasis is transmitted by the tsetse vector and characterized by a stage one bloodstream infection (though other tissues are implicated (28)) and a stage two central nervous systems (CNS) infection, which is invariably fatal if left untreated. HAT is predominantly caused by *T. b. gambiense*, whose stage two infections can be treated by NECT (nifurtimox/eflornithine combination therapy) (2) and fexinidazole (29). To a lesser extent HAT is caused by *T. b. rhodesiense,* which rapidly progresses to stage two infection and host death. Melarsoprol, an arsenical compound, remains the only drug available to treat stage two *T. b. rhodesiense* infections despite treatment challenges, high levels of host toxicity, and increasing reports of drug resistance (30). Following melarsoprol uptake by the parasite-specific P2 adenosine transporter (*AT1* gene) and aquaglyceroporin transporter (*AQP2 gene*), intracellularly metabolized melarsoprol interacts with trypanothione (30). Trypanothione is a kinetoplastida specific form of glutathione required for thiol based redox metabolism (31). While the inactivation of trypanothione by melarsoprol has been demonstrated, and effects on cellular redox metabolism are known, it is has remained unclear if melarsoprol has more than one intracellular target and if inactivation of trypanothione is responsible for cell lysis (30).

In this study we sought to produce a whole-genome *T. brucei* ORFeome and use it to create a Gain-of-Function library for genetic screens. Following PCR amplification and cloning, the resulting ORFeome includes more than 90% of the targeted *T. brucei* ORFs (>6500 of 7245 ORFs targeted). The ORFeome was then introduced into bloodstream *T. brucei* (*Lister427*) to produce the *T. brucei* Gain-of-Function (GoF) Library. Because of melarsoprol’s clinical significance and unknown aspects of its functionality, it was selected for a proof of principle GoF library genetic screen. Following the isolation of melarsoprol survivor populations from the induced GoF library, we identified 57 genes that were significantly overrepresented, which included an anticipated biosynthetic precursor of trypanothione. In addition, we identified sets of genes associated with gene expression, proteins predicted to localize to the mitochondria, and others that localize to regions of the flagellum, whose association with melarsoprol have not been reported previously. Collectively, these findings support the usefulness of the broadly applicable *T. brucei* ORFeome developed herein, the essential functionality of the resulting GoF library, and provide new insights into mechanisms of melarsoprol-based cell killing and genes whose induced expression can promote drug tolerance. Thus, we have generated two genetic tools and a novel dataset, which are all posed to provide new insights into Trypanosomatid biology, pathogenesis, and aid in the development of novel therapeutics.

## RESULTS

### Generation of a Trypanosoma brucei ORFeome

*T. brucei* and others in the class kinetoplastida primarily regulate gene expression post-transcriptionally based on transcript stability mediated by 5’ and 3’ UTRs (19). Thus, previous production of overexpression libraries, based on whole-genome fragmentation and cloning (20, 21), cannot ensure that individual ORFs are similarly activated upon induction. To mitigate these problems, we produced a *T. brucei* ORFeome using PCR amplification, which ensures the placement of each ORF under the same inducible gene regulation system. ORF start and stop sites for all *T. brucei* genes were obtained from available *TREU927* ribosomal profiling data for 9200 genes (32). The targeted *T. brucei* ORFeome filtered out 1956 ORFs that were unsuitable in size (<100 bp or > 4500 bp), an undesired product (annotated as ribosomal genes, VSGs, ESAGS, pseudogenes, or hypothetical unlikely), or if they were known multidrug resistant channels (including MRPA whose overexpression is known to cause melarsoprol resistance (33)). PCR primers for the resulting 7245 targeted ORFs were designed *in silico* with matched melting temperatures and *attB1* cloning sites were added to both forward and reverse primers for subsequent Gateway cloning. The resulting 7245 primer pairs for ORFeome amplification were synthesized and resuspended in 21 separate 384-well plates that were organized by ORF size and gene annotations as either ‘known’ or ‘hypothetical’ (Table 1 and SUP. 1).

**Table 1.** 384-well oligo plates. Plate names, ORF size ranges, and number of oligo pairs per plate are indicated.

| Plate name | Min length (bp) | Max length (bp) | # ORFs per plate |
| --- | --- | --- | --- |
| 0_hypothetical | 102 | 375 | 373 |
| 0_known | 147 | 591 | 315 |
| 1_hypothetical | 375 | 522 | 374 |
| 1_known | 594 | 849 | 365 |
| 2_hypothetical | 522 | 666 | 372 |
| 2_known | 849 | 1056 | 363 |
| 3_hypothetical | 669 | 822 | 382 |
| 3_known | 1056 | 1287 | 367 |
| 4_hypothetical | 822 | 993 | 372 |
| 4_known | 1287 | 1524 | 348 |
| 5_hypothetical | 993 | 1155 | 365 |
| 5_known | 1524 | 1857 | 342 |
| 6_hypothetical | 1158 | 1365 | 375 |
| 6_known | 1857 | 2337 | 362 |
| 7_hypothetical | 1368 | 1635 | 378 |
| 7_known | 2340 | 3504 | 376 |
| 8_hypothetical | 1635 | 1953 | 381 |
| 9_hypothetical | 1953 | 2508 | 381 |
| 10_hypothetical | 2508 | 3501 | 381 |
| hypothetical_last | 3504 | 4488 | 135 |
| known_last | 3507 | 4497 | 138 |

Each gene of the ORFeome was PCR amplified in 384-well format from *Lister427* genomic DNA. The general quality of the PCR reactions was assessed by the addition of SYBR green and measurement of the resulting Relative Fluorescence Units (RFU) (FIG. 1A). Based on the SYBR green assessment, the initial PCR reactions resulted in the successful amplification of 94% of the ORFeome (6820/7245 ORFs) (FIG. 1B & SUP. 2). Most PCR plates resulted in ≥90% amplification and only three of the 384-well plates between 80-90%. To increase our coverage, we attempted to reamplify 429 failed PCR reactions and succeeded in producing 228 products resulting in a final total of 7039 PCR products amplified (97.2% of the targeted genome).

**Figure 1.**
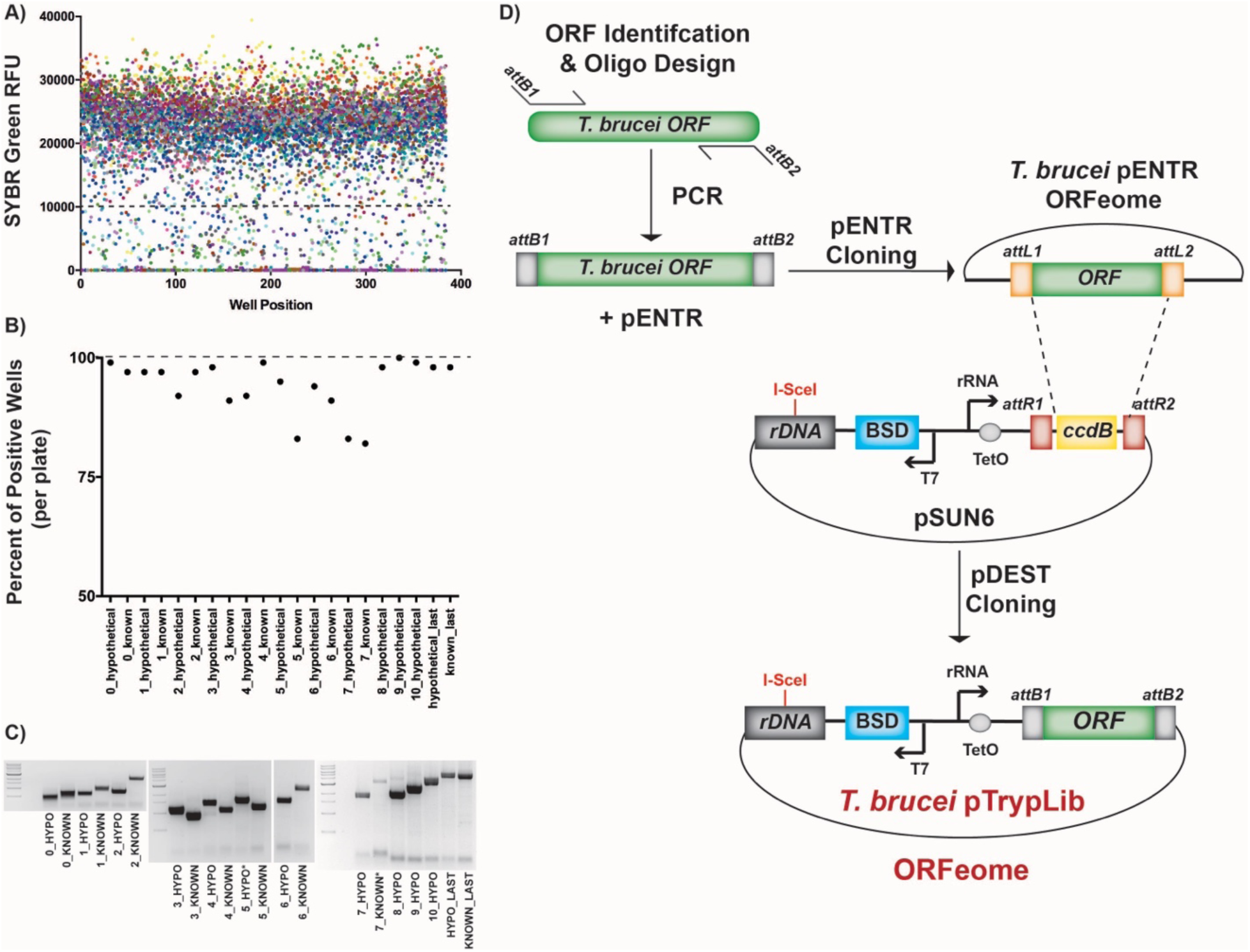
Generating a *T. brucei* ORFeome. A) Assessment of PCR amplification by SYBR Green Relative Fluorescence Units (RFU). Each dot represents an individual PCR reaction, each color represents one of the 21, 384-well plates, graph is of SYBR Relative Fluorescence Units (RFU) vs. 384-well plate position. B) Percent of PCR positive wells (SYBR assessment) for each of the 21, 384-well plates, from the first time amplified. C) Agarose gel bands from each of the 21, 384-well PCR plates pooled prior to gel extraction and cloning, from the first time amplified, compared to 10 Kb ladder DNA ladder. D) ORFeome cloning strategy: *attB1* site addition to *T. brucei* ORFs during PCR amplification, Gateway cloning into pENTR to generate the pENTR ORFeome, and Gateway cloning into *T. brucei* specific pDEST type vector (pSUN6, shown in detail in SUP. 3) to generate the complete pTrypLib ORFeome.

The PCR reactions from each 384-well plate were pooled (10 ul from each well) into 21 corresponding PCR product pools, irrespective of the SYBR result, which maintained the product size range associated with each plate (Table 1). Each resulting size-sorted PCR pool was run on agarose gels and gel purified prior to gateway cloning (FIG. 1C). Thus, the DNA bands visualized on gels in Figure 1C represent hundreds of similarly sized PCR products arising from the same PCR plate, which were gel extracted prior to gateway cloning. Gateway cloning occurred in two steps: 1) introduction of ORFs into a general pENTR vector, which can function as a useful repository for the ORFeome, and 2) transfer of ORFs into a pDEST vector designed for ORF integration into the *T. brucei* rDNA spacer region under the rRNA promoter and tetracycline inducible gene regulation (FIG. 1D and SUP. 3 pSUN6 vector map). For the purpose of this study, we refer to the entire set of cloned ORFs as the pENTR ORFeome and the resulting set of pDEST (pSUN6 derived) libraries ready for transfection into the *T. brucei* genome as the pTrypLib ORFeome.

### Sequencing, assessment, and final coverage of the T. brucei ORFeome

The resulting *T. brucei* pENTR and pTrypLib ORFeome harboring plasmids were each pooled and prepared for Illumina sequencing by tagmentation. In order to assess which of the 7245 targeted ORFs were not present in the pENTR and pTrypLib ORFeome harboring plasmids, we aligned the sequencing reads to the trypanosome genome, removed PCR duplicates, and counted the number of reads corresponding to each targeted gene. Because we knew that some of the targeted genes were highly similar or duplicated, we aligned the reads under two modes, one that required unique alignments and one that allowed multiple alignments. Both datasets were then assessed to determine how many genes were ‘missing’ from each library; defined as any targeted gene with zero aligned reads. The analysis of uniquely aligned reads determined that 1,845 genes were missing from the pENTR library and 2,593 ORFs were missing from the pTrypLib ORFeome. For reads that were multiply aligned, 1,656 ORFs were missing from the pENTR library and 2,420 ORFs were missing from the pTrypLib ORFeome (Figure 2A – pENTR_1, pDEST_1).

**Figure 2.**
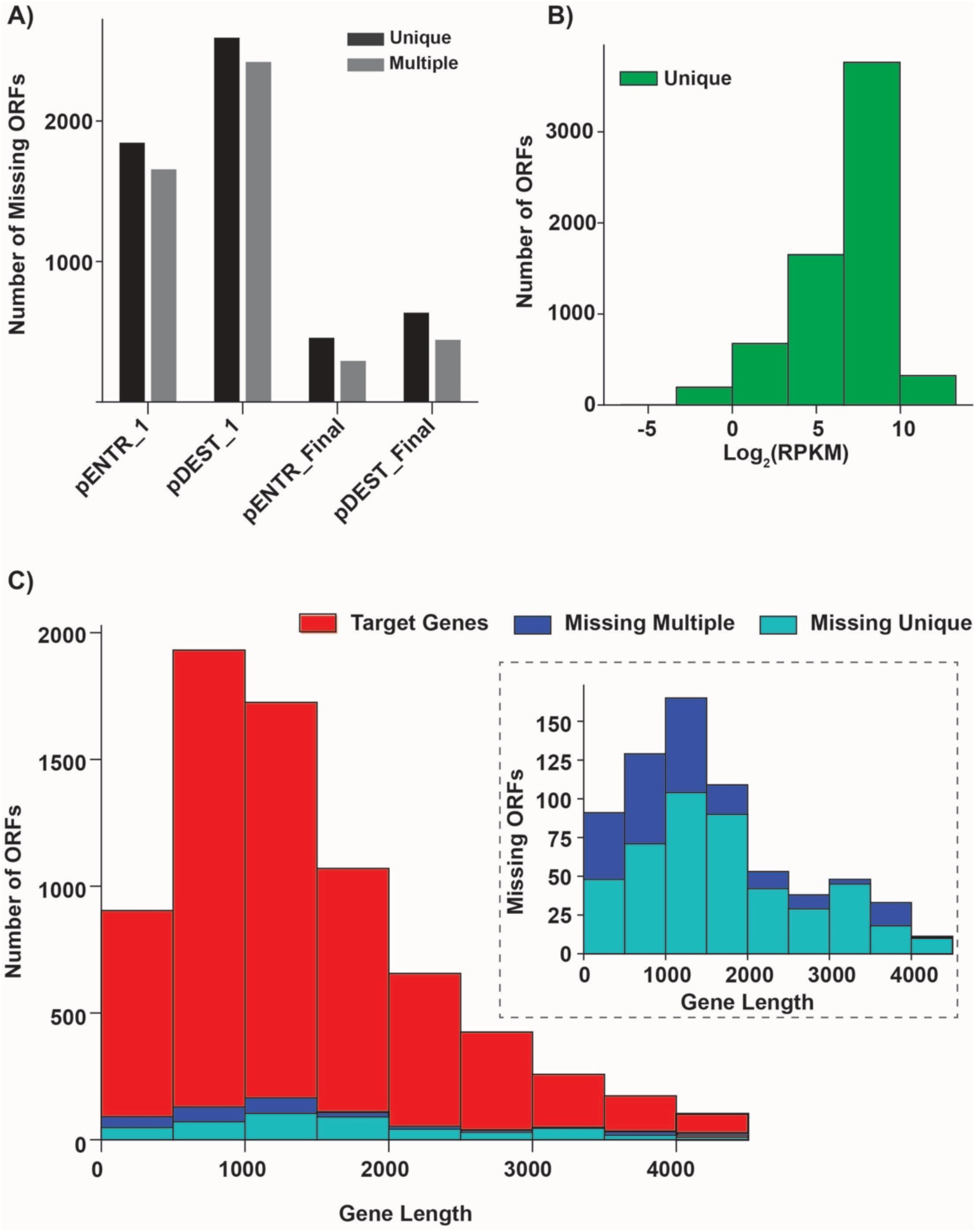
Assessment of pENTR and pDEST (pTrypLib) plasmid libraries. A) Bar graph showing the number of targeted ORFs with zero detectable aligned reads from the first round of cloning (pENTR_1 and pDEST_1) and after both rounds of cloning (pENTR_Final and pDEST_ Final) using analysis generated from both uniquely and multiply aligned reads. B) Histogram showing the distribution of normalized read counts for each ORF in the pooled pDEST (pTrypLib) plasmid libraries (Uniquely aligned reads shown, SUP. 6 both uniquely and multiply aligned reads). C) Histograms showing the distribution of ORF lengths for the target gene list (red) and the set of ORFs with zero detectable aligned reads after both rounds of cloning (labeled as missing). Analyses from unique (dark blue) and multiply (light blue) aligned reads are shown. Inset graph, target ORF lengths have been left out to better visualize the lengths of the missing ORFs.

Our analysis indicated that many of the ‘missing’ ORFs were positive by SYBR analysis following PCR and showed appropriately sized bands when randomly selected missing ORFs were run on agarose gels (approx. 100 PCR products visualized, data not shown). Thus, we predicted that our DNA input was exceeding the functionality of the gateway system in these cloning stages and that the largest losses arose during the formation of the pENTR ORFeome. To address this limitation, PCR products representing each missing ORF were isolated from the original PCR plates (using a Perkin-Emer Janus Automated Workstation to select 2450 PCR products) and used to produce 8 additional size-sorted ORF sub-pools (SUP. 4 – Table of cloning pools including ‘MISS_1-8’), which were gel purified and gateway cloned. These 8 new plasmid pools were then combined with the previous pools and sequenced by tagmentation, as described above.

Our final ORFeome analysis indicated that 457 and 292 ORFs were missing after both rounds of cloning in the pENTR vector using unique or multiply aligned reads, respectively. Gateway cloning into the final pDEST vector to form the pTrypLib ORFeome resulted in 636 and 442 missing ORFs using unique or multiple alignments, respectively (FIG. 2A – pDEST_ Final and SUP. 5 – pENTR and pDEST missing ORFs). Thus, the final pTrypLib ORFeome contains between 6,609 and 6,803 (depending on the type of analysis used) *T. brucei* ORFs, resulting in the successful inclusion of 91-94% of the targeted ORFs.

The resulting sequencing data were used to assess both the overall coverage of ORFs in the library and if specific ORF characteristics (length or annotation) resulted in biases during ORFeome generation. First, we analyzed the coverage of each ORF in the pTrypLib ORFeome. For ORFs that were present in the plasmid libraries (based on reads aligned), we analyzed the count distribution in the sequenced pDEST plasmids for both uniquely and multiply aligned reads. This analysis showed that most ORFs resulted in log_2_(RPKM) values between 0 and 10 (FIG. 2B and SUP. 6). Thus, the number of poorly represented ORFs (RPKM < 1) is 195 for uniquely aligned reads and 369 for multiply aligned reads, representing 3% or 5% of all ORFs in the library, respectively. The log_2_(RPKM) value was then plotted against each corresponding ORF length to determine if gene length affected their representation in the ORFeome (SUP. 6). No strong correlation was observed between ORF length and coverage in the pTrypLib ORFeome, with a best fit line showing a small negative slope for both unique and multiply aligned reads (−.000673 and −.000778, respectively). Thus, in general, shorter ORFs are not significantly more highly represented than longer ORFs.

Next, we sought to determine if there were patterns in the properties of genes missing from the pTrypLib ORFeome. Histograms of ORF length were generated for targeted genes, and compared to histograms of ORF lengths that were missing from the pDEST library. The distribution of ORF lengths in the targeted library was similar to those missing from the the pDEST library. These results indicated the successful cloning of the ORFeome into pDEST and that the resulting libraries were not skewed toward larger or smaller ORFs (FIG. 2C). Thus, we have generated both pENTR ORFeome and pTrypLib ORFeome genetic tools that will broadly support progress in Trypanosomatid research.

### Gain-of-Function library generation from the inducible pTrypLib ORFeome

The pTrypLib ORFeome contains more than 6,500 *T. brucei* ORFs ready for transfection into the genome using targeting to the rDNA spacer regions by homologous recombination. To increase the frequency of transfection and ensure integration into a single rDNA spacer we employed the established method of generating a Landing Pad (LP) parental cell line that targets recombination to a single iteration of the rDNA spacer repeats. The stable LP cell line is then transfected with pRPa-Sce* harboring the ISceI enzyme and restriction site directed for recombination in the targeted rDNA spacer (FIG. 3A) (12). Thus, following ISceI DNA double-stranded break induction by doxycycline (Dox), transfected ORFs are recombined into the targeted rDNA spacer resulting in more consistent transcription induction among library incorporated ORFs.

**Figure 3.**
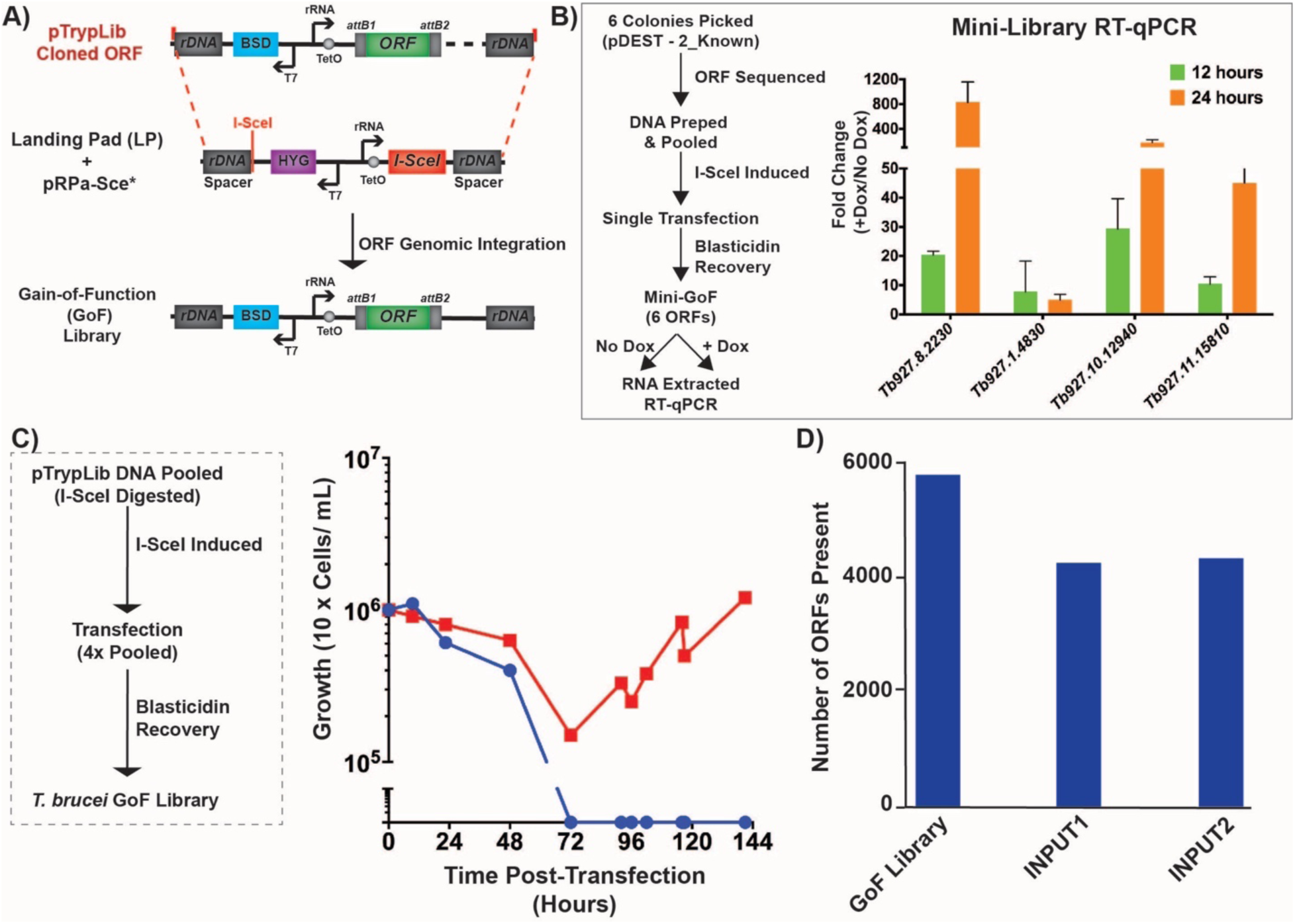
Generation and validation of the *T. brucei* GoF library. A) Transfection of pTrypLib ORFeome into parental Landing Pad (LP) cell line harboring pRPa-Sce* plasmid for I-SceI induced enzymatic cleavage of a single rDNA spacer site (12). B) Generation of a Mini-library GoF library, 6 ORFs. Procedure shown in flow diagram on the left. Transcript level of four GoF library encoded ORFs (exogenous) measured by RT-qPCR following 12 and 24 hours of doxycycline induction compared to uninduced cells (No Dox). C) Generation of the pTrypLib ORFeome based GoF library. Procedure shown in flow diagram on the left. Graph indicates the recovery of GoF library harboring cells (red line) compared to mock transfection (blue line) in blasticidin. D) Assessment of the number of ORFeome genes present in the GoF library following initial transfection (GoF library), in INPUT1 (following freeze thaw and minimal propagation), and INPUT2, which used an alternative NGS protocol.

Validating inducible ORF transcription from a whole-ORFeome transfection would be challenging to detect due to the overall complexity of the resulting cell populations (harboring approx. 6,500 ORFs in multiple copies). Thus, prior to transfecting the TrypLib ORFeome we sought to verify that the transcription of individual ORFs within this library would be inducible and highly transcribed. Toward this goal we generated a subset ‘mini-library’, in which one pDEST bacterial pool was plated (SUP. 2, Plate name – “2_Known”, ORF range 849-1056 bp) and six random individual colonies isolated, plasmid DNA prepared, and sequenced to identify the ORF harbored in each pDEST vector (FIG. 3B). The resulting plasmids were pooled equimolarly, transfected into *T. brucei*, recovered as a single population of parasites (the ORFeome ‘mini-library’), and then grown with or without Dox induction for 12 or 24 hours prior to RNA extraction. The relative level of four ORFs were determined using RT-qPCR with a forward oligo specific to pDEST (*attB1*site) and a reverse oligo specific to individual ORF. Because these oligo pairs can amply the ORFs in the ORFeome library (exogenous) and not endogenous ORF, we can measure the inducibility of our ORFeome library. Levels of ORF transcripts were compared between uninduced and Dox induced culture. All ORFs showed increased transcript levels following Dox induction at 12 and 24 hours, 3 of the 4 ORFs resulted in approximately 10-30-fold increased after 12 hours and 50-600-fold increased after 24 hours (FIG. 3B). Thus, the overall strategy of ORFeome exogenous transcription induction from pTrypLib cloned ORFs is viable.

We then generated a doxycycline inducible *T. brucei* Gain-of-Function (GoF) library from the pTrypLib ORFeome by transfection of 360 million cells (LP parent following pRPa-Sce incorporation of ISceI cassette) and selected in blasticidin. After 72 hours of selection mock transfected cells were dead and approx. 60 million cells survived transfection (FIG. 3C – red line), which were then propagated to 3 billion cells over 3 days to generate the *T. brucei* GoF library. Assessment of the resulting GoF libraries by Illumina sequencing requires a specialized procedure to ensure that only the introduced pTrypLib ORFs are sequenced and not the endogenous gene copies. Illumina sequencing libraries are prepared using a gateway *attB1* site specific forward oligo, during PCR enrichment, and a similar custom sequencing primer complementary to the *attB1* site upstream of the introduced ORF. Thus, the resulting sequencing reads primarily correspond to the 5’ ends of the introduced ORF (SUP. 7A – Sequencing strategy). Immediately following transfection and recovery in blasticidin, the *T. brucei* GoF library consisted of 5,818 ORFs and then between 4,269 and 4,360 ORFs when thawed and propagated for 3 days. The GoF ‘INPUT’ library, arising from 3 days of library propagation, represents the full complement of ORFs present before a screening condition is applied (FIG. 3D). It is unclear if the apparent loss of 1,459 genes arose through an artifact associated with a relatively low number of NGS reads returned from those samples or a true loss of content between library transfection and the subsequent thawing of frozen library.

### Identification of overrepresented GoF library ORFs in a melarsoprol survivor screen

Melarsoprol remains the only drug available for the treatment of second stage *T. b. rhodesiense* infection, yet it is highly toxic and difficult to administer (30). In addition, there have been increased reports of melarsoprol resistance and treatment failures in sub-Saharan Africa since the 1990s, only some of which have been explained by transporter mutations (34–37). Since melarsoprol has been widely studied, with both known and unknown genes associated with its cytotoxicity and treatment failures, it was selected for a *T. brucei* GoF library proof of principle genetic screen.

To identify ORFs whose induced expression promote survival in the presence of lethal doses of melarsoprol, we tested three concentrations of drug on the LP parental cell line. Similar to previous reports, we observed that *T. brucei* LP cells died after 3 days in 35 nM, 5 days in 26 nM, and 7 days in 17 nM melarsoprol (17 nM is approximately two times the standard EC50) (FIG. 4A) (11). We predicted that the progression of cell killing might be a critical factor in design of the GoF Library melarsoprol resistance screen. That is cells that die too quickly might not have the opportunity to successfully activate ORFs that confer resistance. In fact, when a genetic screen was attempted in 35 nM melarsoprol, the death of the culture was delayed by one day in both uninduced and induced GoF library cells when compared with parental landing pad, yet no survivor population emerged (FIG. 4B – Red dashed and dotted lines overlap). In contrast, when cells harboring the GoF library were treated with 17 nM melarosoprol, uninduced cells (no Dox), took almost 4 days longer to die than the LP parental line (preliminary test, data not shown). Thus, a GoF forward-genetic screen was prepared in 17 nM melarsoprol.

**Figure 4.**
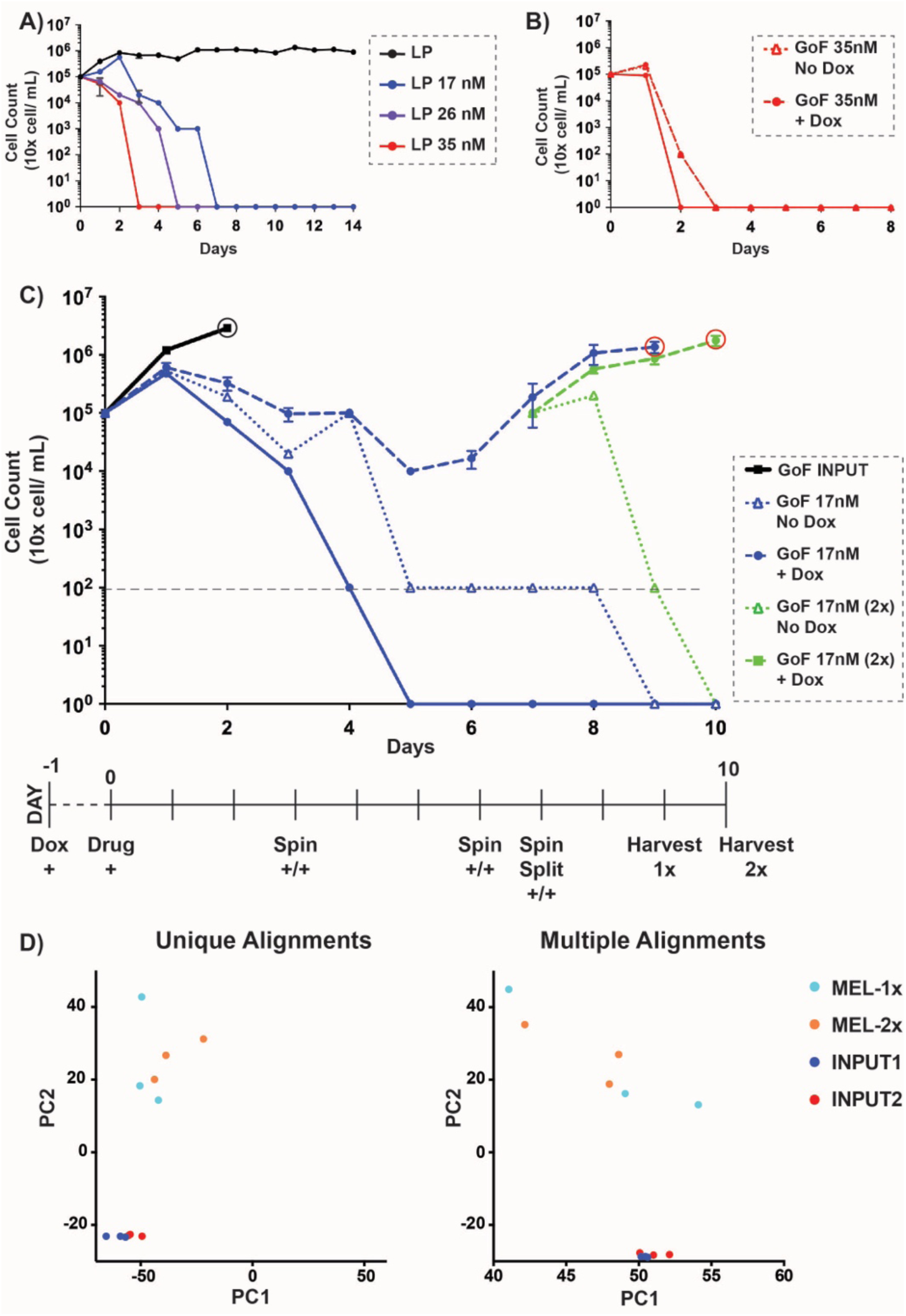
Isolation of melarsoprol survivor populations from a GoF screen. A) Growth of Landing Pad (LP) parental cell line in 17 nM (blue line), 26 nM (purple line), 35 nM (red line), or no (black line) melarsoprol treatment over 14 days. B) GoF library screen in 35 nM melarsoprol treatment: LP cell line (solid red line), uninduced GoF library (red dotted line), induced GoF library (red dashed line); dotted and dashed lines overlap. C) GoF genetic screen in 17 nM melarsoprol is shown as a time course at the bottom and graph of cell growth: INPUT, untreated GoF library harboring cells grown for 3 days (black line), LP parental cell line (solid blue line), uninduced GoF library (blue triangles on dotted line), induced GoF library (blue circles on dashed line), harvested on Day 9 (red circle on blue line) to produce MEL-1x. On day 7, biological triplicates from induced GoF library (blue circles on dashed line) were split into two sets of triplicate samples, both with 17 nM melarsoprol, one of which was not further induced (green triangles on dotted line). The other continued to be induced (green squares on dashed green line) and was harvested on Day 10 to produce MEL-2x (red circle indicates harvest). D) Principle Component Analysis (PCA) comparing INPUT libraries (1 and 2) with libraries arising following 2 weeks of continuous melarsoprol selection (MEL-1x and MEL-2x).

The experimental steps for the 17 nM melarsoprol GoF genetic screen, including 24 hour pre-induction of all Dox treated cultures (Day -1), are shown at the bottom of Figure 4C. A freezer vial containing 100 million GoF library harboring cells was thawed and propagated for less than 3 days before splitting it into triplicate samples for the following conditions: GoF ‘INPUT’ (grown for 3 days without melarsoprol selection), GoF 17 nM No Dox (uninduced library maintained in melarsoprol), and GoF 17 nM +Dox (Dox induced library maintained in melarsoprol) (FIG. 4C). The parental cell line (LP) was maintained in melarsoprol alongside GoF library cultures and were below the limit of detection (FIG. 4 – grey dashed line at 1×10^2^ cells/mL) on day 4 with complete cell death observed from day 5 until the completion of the screen on day 10. In contrast, uninduced (No Dox) GoF Library harboring cells showed partially delayed melarsoprol killing, with counts below detection at day 5, but some observable surviving cells in culture until day 8, after which all cells appeared dead (FIG. 4C – Blue dotted line). The melarsoprol treated, Dox induced GoF library was distinct from LP parent by day 3, and by day 5 populations of survivors emerged and grew similarly well in all three induced GoF (+Dox) biological replicates (FIG. 4C – Blue dashed line). At day 7 the biological replicates of induced (+Dox) GoF library melarsoprol survivor cultures were split into media with and without Dox during continued treatment with 17nM melarsoprol (FIG. 4C – Green lines). The uninduced (no Dox) cultures died rapidly over the next 2 days (dotted green line), whereas the induced GoF library, once again, survived and grew robustly (dashed green line) in the presence of melarsoprol. Thus, survival in the presence of melarsoprol was dependent on GoF library induction. Biological triplicate cultures of melarsoprol survivors were harvested from both GoF ‘MEL-1x’ and GoF ‘MEL-2x’ (resulting from the day 7 split) on day 9 and day 10 respectively (FIG. 4C – blue and green dashed lines, red circles indicate the date of harvest). Genomic DNA from biological triplicate cultures of untreated INPUT (day 3, black circle), GoF ‘MEL-1x’, and GoF ‘MEL-2x’ cultures (9 cultures total grown to ∼1 million cells per mL 200 mL each) was prepared for NGS analysis.

To identify the pTrypLib ORF incorporated into the GoF library harboring cells recovered from each replicate and condition (INPUT, GoF MEL-1x, and GoF MEL-2x), genomic DNA was fragmented and prepared for ORFeome specific Illumina sequencing (shown in SUP. 6). During PCR enrichment INPUT samples were prepared in two ways: long elongation time (INPUT1, 75 seconds) and shorter elongation time (INPUT2, 20 seconds). Sequencing data was obtained in biological triplicate from the two sets of INPUT libraries (no melarsoprol treatment) and the two sets of melarsoprol selected parasites (MEL-1x and MEL-2x). We performed principle component analysis on the sequencing data from both unique and multiple alignment datasets (FIG. 4D). The PCA analysis shows two clearly separated clusters for untreated and melarsoprol treated samples, with most biological replicates clustering proximal to one another. Melarsoprol treated samples were distinct from INPUT and show more variation between samples (FIG. 1D). In order to check whether ORF length influenced the level of coverage in the genomic INPUT libraries, we analyzed whether ORF representation in the INPUT libraries correlated with ORF length. We observed, at best, a weak negative association between gene length and normalized read count (slopes of −.00042 and −.00045 for unique and multiple alignment analysis, respectively), indicating that ORF representation in the library is largely independent of ORF length (SUP. 8)

We reasoned that any gene whose induction contributed to melarsoprol resistance should be overrepresented in libraries generated from melarsoprol survivor populations. The designation ‘overrepresented’ indicates that the normalized read count for a particular ORF in the presence of melarsoprol is higher in libraries arising from doxycycline induced parasites harboring the GoF library (MEL-1x and MEL-2x) when compared with the untreated GoF library (INPUT), which suggests that overrepresented genes have enhanced survival in the presence of drug. To determine the fold change that represents a valid difference between melarsoprol treated and untreated conditions, we compared each of the three biological replicates of INPUT2 to one another and counted the number of ORFs with a 1.5, 2.0, or 4.0-fold in normalized read count (FIG. 5A – INPUT2 replicates compared pairwise). By evaluating the biological variation between similarly treated replicates we found that while many ORFs varied in normalized read count by greater than 1.5-fold between replicates (more than 300), very few ORFs varied by greater than 4-fold (FIG. 5A, similar results obtained from INPUT 1, data not shown). Thus, we used a 4-fold change in normalized read count between melarsoprol treated and untreated samples as the minimum threshold for identifying an ORF as overrepresented (a ‘hit’) in this study.

**Figure 5.**
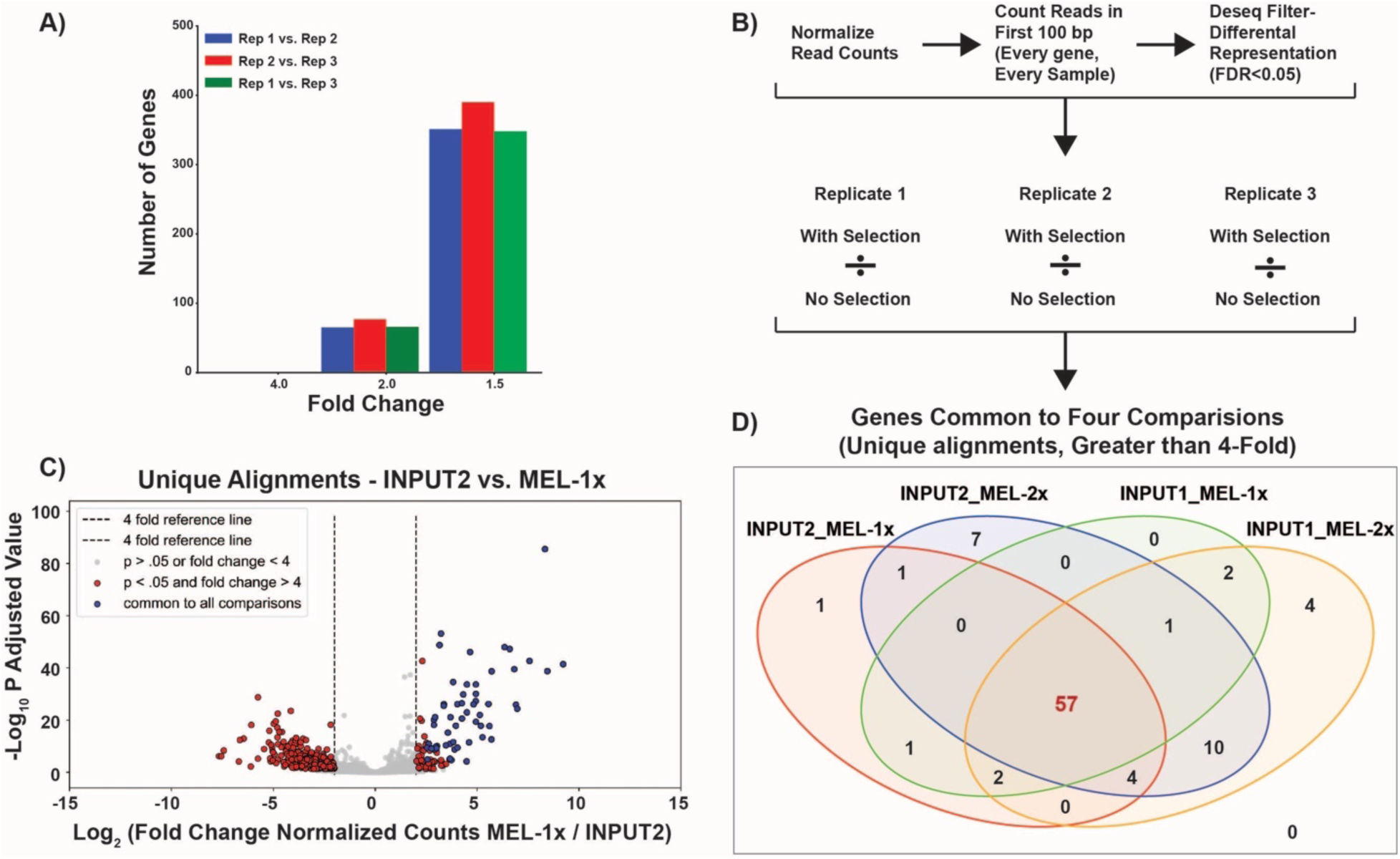
Identification of significantly overrepresented ORFs in melarsoprol GoF survivor populations. A) The number of genes with a fold change > 1.5, 2, or 4-fold is given for all three replicates of INPUT2: Replicate 1 vs. Replicate 2, Replicate 2 vs. Replicate 3, and Replicate 1 vs. Replicate 3. B) Hit calling pipeline to identify genes overrepresented in melarsoprol survivor populations. Following read count normalization and alignment to the first 100 bp of each gene in every sample, DESeq was used to filter for differential representation using a P adjusted cutoff of <0.05. Each melarsoprol treated replicate was compared with each untreated INPUT replicate and those with 4-fold or greater change of melarsoprol selected over unselected in all three comparisons were called as preliminary hits. Preliminary hits were determined for INPUT1 vs. MEL-1x, INPUT2 vs. MEL-1x, INPUT1 vs. MEL-2x, and INPUT2 vs. MEL-2x. The final list of 57 hits represent genes common to all four comparisons. C) Volcano plot showing the −Log_10_(P adjusted value) vs. Log_2_ (Fold Change in normalized counts for the comparison of melarsoprol selected MEL-1x/INPUT2) for each ORF in the targeted library. Red dots represent fold changes >4 with a P adjusted significance of <0.05 and blue dots represent overrepresented ORFs common to all four comparisons described in B. D) Venn Diagram illustrates the significantly overrepresented genes common to each comparison and shared among all comparisons, resulting in 57 overrepresented hits identified in melarsoprol survivor populations compared to input populations.

In order to identify ORFs overrepresented in melarsoprol survivor populations, we aligned reads from each of the 12 samples (3 reps of each: INPUT1, INPUT2, MEL-1x, and MEL-2x) to the genome, and then calculated the number of reads that fell within the first 100bp of each of the ORFeome targeted ORFs (SUP. 7B – Example read alignments). DESeq was used to normalize read counts between libraries (38). We performed this analysis for uniquely aligned reads and multiple alignments. DESeq was used to identify genes that were ‘differentially expressed’ between melarsoprol selected samples and INPUT samples with a p adjusted value of < 0.05 (FIG. 5B) (38). In this context, ‘differential expression’ simply refers to an ORF being overrepresented or underrepresented in the melarsoprol selected samples. To be considered as a potential hit, we required a minimum normalized read count of 5 in INPUT samples to avoid ORFs being called as overrepresented that were simply poorly covered. To identify putative hits, we then asked which of the differentially represented ORFs with and p adjusted value < 0.05 had a normalized read count > 4-fold higher in 3 biological replicates of melarsoprol treated populations (MEL-1x and MEL-2x) compared to untreated populations (INPUT1 and INPUT2). Four different comparisons were analyzed using this pipeline: INPUT1 vs. Mel-1x, INPUT1 vs. Mel-2x, INPUT2 vs. Mel-1x, and INPUT2 vs. Mel-2x (FIG. 5B). Figure 5C shows a volcano plot of DESeq generated significance values compared with fold change for the comparison between INPUT2 and MEL-1x. After hits had been called for each individual comparison, we identified the hits common between all 4 comparisons for both uniquely and multiply aligned reads (FIG. 5D and SUP. 9 – Tables of all comparisons). These analyses resulted in the identification of 57 overrepresented ORFs (uniquely aligned) in the GoF melarsoprol survivor populations compared to INPUT populations. In the comparison of INPUT2 vs. MEL-1x depicted in a volcano plot, we observe that these 57 ORFs common to all the comparisons were among the most highly overrepresented genes and with some of the lowest P adjusted values determined by DESeq (FIG. 5C – blue dots). Similar results were obtained for all comparisons between melarsoprol selected and input samples. We conclude that our analysis is reasonably robust at identifying the most consistent, statistically significant, highly overrepresented genes in melarsoprol treated samples compared to untreated samples.

The 57 genes overrepresented in melarsoprol survivor populations consist of 25 genes annotated as conserved hypothetical and among the genes with annotations most have only putative functional assignments (SUP. 10). Among the top 5 overrepresented hits we identified that the gene encoding γ-glutamylcysteine synthetase (GSH1, *Tb927.10.12370*) was 191-fold higher among melarsoprol survivors when compared with input control. GSH1 is required for the biosynthesis of trypanothione, a known target of melarsoprol, and its overexpression has been demonstrated to promote melarsoprol resistance in laboratory studies (33). This finding supports the role of trypanothione as one of the major intracellular targets of melarsoprol and the usefulness of the GoF library in the direct identification of drug targets. Using available functional annotations (tritrypdb.org), we found that 15 of the 57 overrepresented hits are either known or implicated in gene expression regulation. It is not clear at this time if this is a true aspect of melarsoprol functionality or whether this is an artifact arising from the nature of the GoF library; namely increased transcript stability and translation of diverse proteins having a nonspecific effect on cell viability in the presence of drug. Only through further GoF library validation will we know if gene expression is truly associated with melarsoprol resistance. We then utilized publicly available resources (tritrypdb.org and tryptag.org) to develop additional categories based on subcellular localization (from proteomic and microscopic studies) and published protein functions. We found that the majority of overrepresented genes encoded proteins that localize to the mitochondria and kinetoplast (10 genes, shown in Table 2) or the flagellum (9 genes, shown in Table 3) (SUP. 10 – Hits listed with categories). The prevalence of mitochondrial and flagellar proteins arising from the melarsoprol GoF screening may implicate previously unrecognized melarsoprol drug targets or pathways of drug tolerance.

**Table 2.** Melarsoprol GoF hits with Mitochondrial localization.

| GENE ID | CATEGORY | DESCRIPTION | LOCALIZATION (TrypTag.org) | FOLD CHANGE | P ADJUSTED DESEQ |
| --- | --- | --- | --- | --- | --- |
| Tb927.11.590 | Mitochondrial* | hypothetical protein, conserved | Strong mitochondrial signal, kinetoplast | 350 | 1.81E-39 |
| Tb927.10.12050 | Mitochondrial* | hypothetical protein, conserved | kinetoplast, mitochondrion | 31 | 2.16E-34 |
| Tb927.9.7200 | Mitochondrial* | hypothetical protein, conserved | kinetoplast, mitochondrion | 31 | 5.61E-28 |
| Tb927.9.7080 | Mitochondrial & ER<sup>o</sup> | hypothetical protein, conserved | cytoplasm (patchy, points) | 23 | 1.14E-23 |
| Tb927.2.5210 | Mitochondrial# | 3-oxoacyl-ACP reductase, putative | mitochondrion, kinetoplast (strong) | 23 | 2.16E-34 |
| Tb927.11.2910 | Mitochondrial* | phosphoglycerate mutase, putative (iPGAM) | mitochondrion (75%), kinetoplast (75%) | 23 | 6.79E-05 |
| Tb927.9.5220 | Mitochondrial<sup>o</sup> | conserved protein | endocytic, cytoplasm(weak) | 19 | 2.60E-21 |
| Tb927.10.6850 | Mitochondrial# | Mitochondrial ribosomal protein S18, putative | cytoplasm(reticulated) | 16 | 3.68E-10 |
| Tb927.8.7790 | Mitochondrial<sup>o</sup> | zinc finger domain, LSD1 subclass, putative | ND | 13 | 7.88E-22 |
| Tb927.11.5600 | Mitochondrial# | Archaic Translocase of outer membrane 14 kDa subunit | mitochondrion | 8 | 9.21E-11 |
| Tb927.3.5190 | Mitochondrial+ | hypothetical protein, conserved | mitochondrion | 7 | 8.48E-21 |

Key: + From localization data, <sup>o</sup> Proteomic data, \* both localization and proteomics, or # published

**Table 3.** Melarsoprol GoF hits with Flagellar Localization.

| GENE ID | CATEGORY | DESCRIPTION | LOCALIZATION | FOLD CHANGE | P ADJUSTED DESEQ |
| --- | --- | --- | --- | --- | --- |
| Tb927.11.11435 | Flagellar+ | dynein light chain lc6 | Strong flagellum axoneme signal | 120 | 1.30E-26 |
| Tb927.9.15020 | Flagellar+ | hypothetical protein, conserved | Weak axoneme signal | 31 | 4.45E-20 |
| Tb927.11.1810 | Flagellar+ | Ring finger domain containing protein, putative | flagellar pocket (ring) | 16 | 1.50E-18 |
| Tb927.10.390 | Flagellar* | hypothetical protein, conserved | Flagellum Matrix Proteome (BSF) | 14 | 2.78E-12 |
| Tb927.5.4150 | Flagellar* | hypothetical protein, conserved | paraflagellar rod | 13 | 6.29E-06 |
| Tb927.7.710 | Flagellar* | heat shock 70 kDa protein, putative (HSP70) | cell tip (anterior), cytoplasm, flagellar cytoplasm | 10 | 3.15E-26 |
| Tb927.10.9060 | Flagellar + | hypothetical protein, conserved | basal body | 8 | 5.86E-10 |
| Tb927.1.1020 | Flagellar + | leucine-rich repeat-containing protein | hook complex | 8 | 1.61E-10 |
| Tb927.6.3980 | Flagellar + | hypothetical protein, conserved | Flagellum, axoneme | 6 | 3.88E-10 |

Key: + From localization data, <sup>o</sup> Proteomic data, \* both localization and proteomics, or # published

### Overexpression of genes identified in melarsoprol GoF screen result in increased drug tolerance

GoF library induction in the presence of melarsoprol resulted in the isolation of survivor populations harboring a consistent set of overrepresented genes. The growth environment arising from the induced expression of many genes is likely distinct from cell populations overexpressing only one gene and may influence population survival through the secretion of unknown factors or other events. To determine if the overexpression of individual genes identified among the 57 overrepresented hits can contribute to melarsoprol resistance, we cloned a subset of genes in to the pLEW100v5-BSD vector, which permits Dox inducible gene expression in a manner similar to the GoF library itself. Bloodstream form *T. brucei* (SM) were transfected with linearized plasmids harboring individual genes for overexpression and, following their recovery were, assayed for their effects on melarsoprol sensitivity by alamarBlue assays. The gene encoding γ-glutamylcysteine synthetase (*Tb927.10.12370*, GCS1), which is responsible for the biosynthesis of glutathione used to form trypanothione, was among the most overrepresented genes (190-fold), is essential, and its overexpression has been linked to resistance in previous studies (39, 40). As anticipated induced overexpression of *GSH1* resulted in an increase in the EC50 of melarsoprol of almost 2-fold (FIG. 6A – SM = EC50 7.5 nM and OE GSH1 = 14.5 nM) suggesting that increased expression of components of this pathway can produce more trypanothione and diminish the efficacy of melarsoprol. The trypanothione pathway (FIG. 6B) and its implications for melarsoprol resistance are covered in more detail in the discussion.

**Figure 6.**
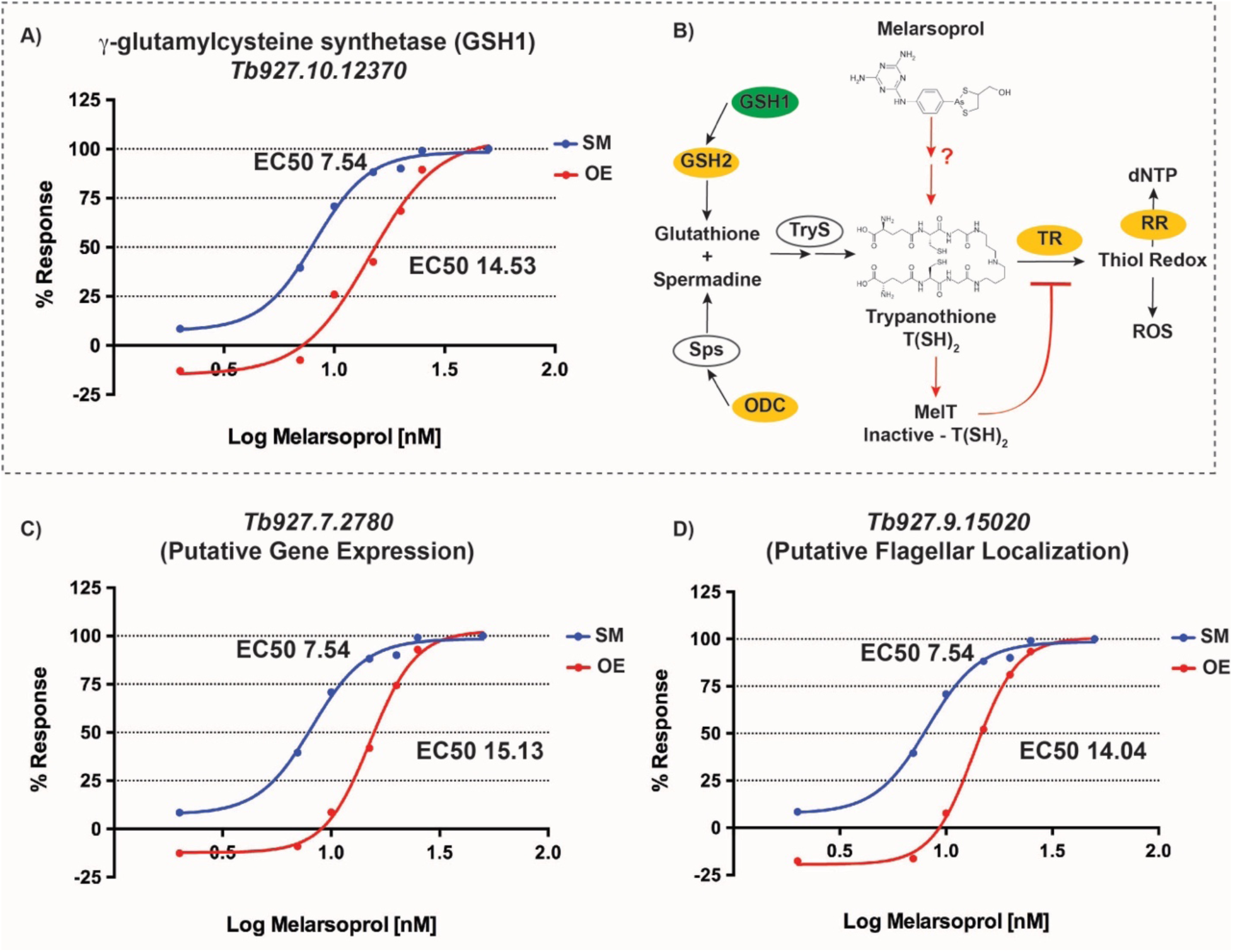
Melarsoprol sensitivity following gene overexpression. AlamarBlue assays were used to measure the EC50 over a range of melarsoprol concentrations (2, 7, 10, 15, 20, 25, and 50 nM) for both parental cell line (SM, blue line) and following Dox induction of genes identified in the melarsoporol GoF screen (A, C, D red lines labeled OE). A) Effect of γ-glutamylcysteine synthetase (*Tb927.10.12370*, GSH1) overexpression (red line) on the EC50 of melarsoprol (shown in text on the graph). B) The associated trypanothione (T(SH)_2_) pathway with GSH1 is also depicted, in which the green oval is the overrepresented hit GSH, yellow ovals indicate pathway components that were modestly effected but did not meet the statistical cutoffs for GoF hit calling (SUP. 11), and genes encoding enzymes in white ovals were not changed in the GoF screen. The effects of melarsoprol on T(SH)_2_ are also depicted as well as the outcomes of this pathway (panel B). Data panels C) and D) show the effects of *Tb927.7.2780* and *Tb927.9.15020* overexpression on the EC50 of melarsoprol, respectively. All alamarBlue assays were done in biological triplicate for both uninduced (No Dox, not shown) and induced (+Dox) conditions. Data shown in panel A, C, and D (red lines) are single experimental representative of all Dox induced biological replicates.

We also found that the overexpression of the hypothetical conserved gene *Tb927.7.2780* (overrep. 322-fold) resulted in a 2-fold shift in EC50 (FIG. 6C – OE = EC50 15.1). The protein encoded by *Tb927.7.2780* is a predicted post-transcriptional activator associated with mRNA stability or increased translation (41) and is annotated as an essential gene based on RIT-seq analysis (42). Whether melarsoprol resistance associated with *Tb927.7.2780* arises from direct effects on gene expression of known pathways, such as trypanothione biosynthesis enzymes, or more general effects on global gene regulation remains to be determined. A similar shift in EC50 was also observed upon overexpression of *Tb927.9.15020* (FIG. 6C – OE = EC50 14.0), which encodes a hypothetical protein of unknown function (overrep. 30.1-fold) that we categorized as flagellar (axonemal) based on TrypTag.org localization data (images shared ahead of online deposition). It is unclear at this time how flagellar proteins may participate in melarsoprol resistance, discussed in more detail below. Multiple genes encoding mitochondrially localized proteins were cloned for similar overexpression analyses, but resulted in assay challenges arising from either growth defects, inability to reduce alamarBlue, or both. Because cell viability in alamarBlue assays is reported based on the reduction of the active reagent (resazurin) by mitochondrial redox metabolism, it is not a surprise that the effects mitochondrial proteins could not be assayed in this manner and may indicate their association with the trypanothione redox pathway itself. The role of mitochondrial proteins in melarsoprol tolerance will be the subject of future investigations. Together these results show that genes identified in melarsoprol GoF screening can promote drug resistance upon overexpression, support the significance of the trypanothione pathway, and implicate new aspects of kinetoplastida biology in melarsoprol resistance.

## DISCUSSION

The forward genetics tools generated here address an urgent need to extend genomic functional characterization in *T. brucei* and its kinetoplastida relatives. More than 30 years of genetic and biochemical studies in Trypanosomatids, 10 of which included the extensive use of an RNAi-based loss-of-function library, have produced key discoveries in parasitology and basic biology. Yet, with the functions of more than 50% of Trypanosomatid encoded genes largely unknown, many mysteries remain unsolved and very few functional pathways have been delineated. To unlock potential discoveries in Trypanosomatids, here we have generated two powerful tools for forward genetics and unbiased studies based on the *T. brucei* genome (*TREU927*): an ORFeome consisting of over 6500 ORFs and an inducible Gain-of-Function library harbored in *T. brucei* parasites (*Lister427*) for genetic screening. To validate the functionality of the *T. brucei* GoF library, we conducted a genetic screen using the troublesome drug melarsoprol and identified both an anticipated target as well as novel genes that may reveal new facets for melarsoprol’s mechanism of action.

Melarsoprol has a complex history dating back to early uses of arsenic treatment of HAT, treatments which are of specific significance because of their ability to cross the blood-brain barrier and treat second stage infections (30). *In vivo* melarsoprol is rapidly metabolized to other trypanocidal metabolites, including melarsen oxide, which interacts with trypanothione to from MelT, a potent inhibitor of trypanothione reductase (TR) (11). Yet, the precise mechanism of melarsoprol cell killing has remained unclear and the complex forms of metabolized melarsoprol may have additional intracellular targets. Melarsoprol drug resistance and treatment failures have been on the rise since the 1990s and while most resistance has been connected to mutations in aquaglyceroporin transporters, numerous treatment failures remain unexplained (30). The trypanothione biosynthetic precursor, γ-glutamylcysteine synthetase (GCS1), was among the most overrepresented genes in melarsoprol survivors and is known to result in melarsoprol resistance upon laboratory overexpression (39, 40). Upon deeper excavation of the melarsoprol GoF screen data, we observed that a number of genes upstream and downstream of T(SH)_2_ with enriched reads in melarsoprol survivors, but did not meet the criteria to be called overrepresented (SUP. 11). For instance, both of the T(SH)_2_ biosynthetic precursors ornithine decarboxylase (ODC, *Tb927.11.13730*) and glutathione synthetase (GSH2, *Tb927.7.4000*) were partially overrepresented in some cultures of melarsoprol survivors (but not all). Downstream from T(SH)_2_, we saw increased levels of trypanothione reductase (TR, *Tb927.10.10390*), a trypanothione dependent peroxidase (*Tb927.7.1140*), and ribonucleotide reductase (RR, *Tb927.11.7840*), in some survivor cultures. The generation of dNTPs by the action of RR has known effects on kDNA replication (43) and melarsoprol treatment was recently shown to stall cell cycle (44), which might suggest inhibition of DNA synthesis as a mechanism of cell killing. Collectively these findings may further support the trypanothione pathway as a primary target of melarsoprol-based cell killing.

Among the 57 significantly overrepresented genes approximately 45% are annotated as hypothetical. The majority of overrepresented genes with functional predictions are associated with gene expression (15 genes) with the largest groups associated post-transcriptional regulation (6 genes) or splicing factors (5 genes). Notably, one of the overrepresented genes associated with post-transcriptional gene expression regulation (*Tb927.10.1490*, MKT1L) was also a high significance gene in melarsoprol RNAi screening (45). For overrepresented genes without functional annotations we focused on the cellular localization of the encoded proteins (tritrypdb.org and tryptag.org) and identified a preponderance of proteins that localize to either the mitochondria (11 genes) or flagellum (9 genes) (FIG. 7). The flagellar proteins have diverse localizations in procyclic form (insect stage) parasites, that include localization to the flagellar pocket and along the body of the organelle (axonemal and paraflagellar rod, specifically). Since the melarsoprol transporter AQP2 is specifically localized to the flagellar pocket in bloodstream form parasites (46), one could speculate that overexpression of a protein that interferes with drug transport in this region might enhance resistance. Two of the overrepresented flagellar genes, one axonemal (*Tb927.5.4150*) and one localized to the hook complex (*Tb927.1.1020*), were associated with a loss of fitness in RIT-seq screens, which is indicative of their essentiality. It will be the subject of future studies to determine how flagellar proteins might contribute to drug resistance or if they are viable drug targets.

**Figure 7.**
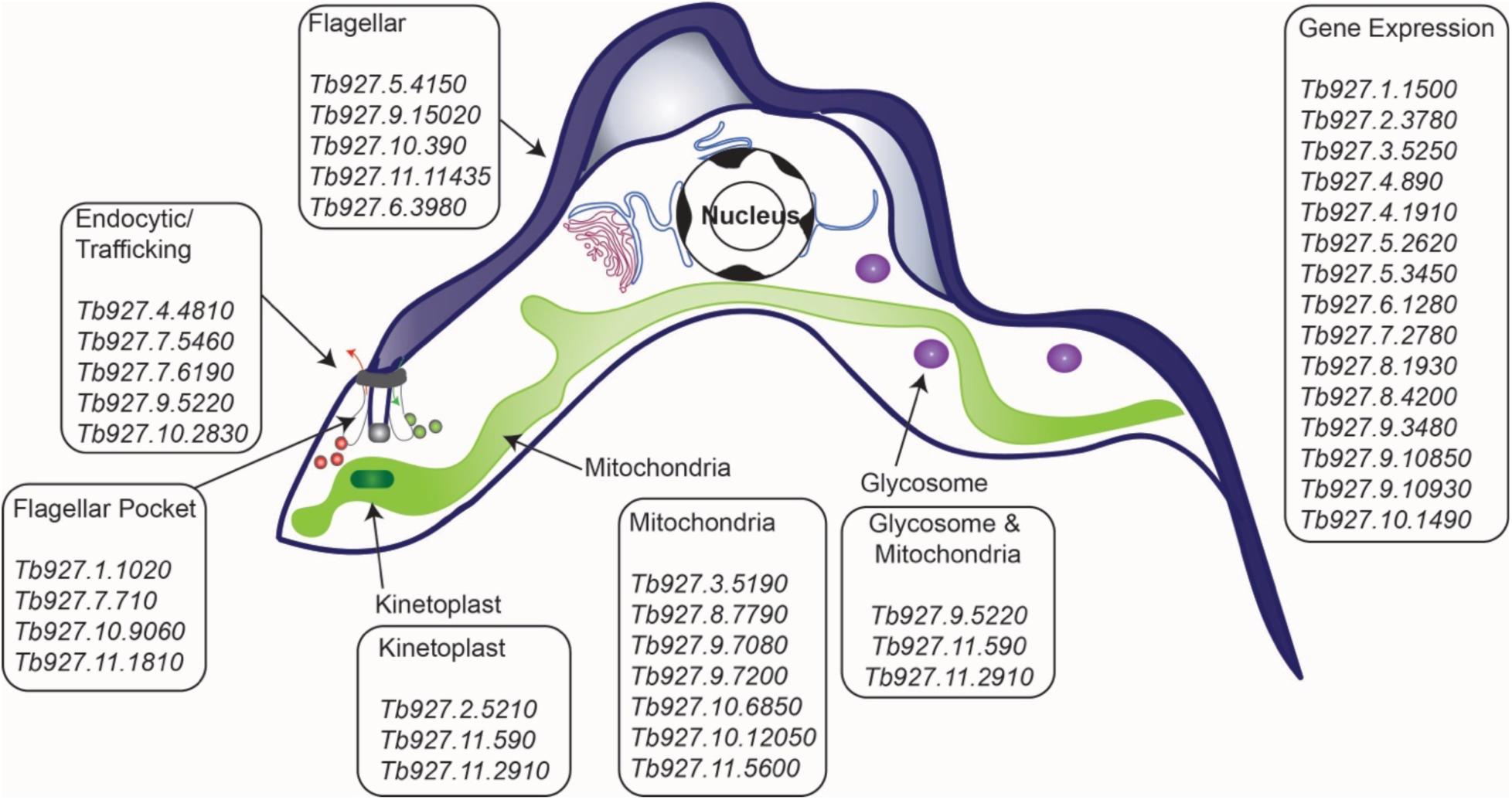
Categories of overrepresented genes in melarsoprol GoF screen survivor populations. An illustrative diagram of a *T. brucei* cell is shown with relevant features highlighted. Boxes of gene numbers indicate major functional or localization categories for the genes overrepresented in melarsoprol survivor populations.

Connections between T(SH)_2_ and mitochondrial redox reactions have been widely established (31) and the anti-Trypanosomatid drugs pentamidine and nifurtimox have been linked to specific mitochondrial functions (44, 47). Yet, the data arising from the GoF screen described herein represent the first genetic links between melarsoprol and the mitochondria. Furthermore, since the majority of the GoF proteins that localize to the mitochondria are hypothetical or have only putative functionality, the 11 mitochondrial genes identified here have the potential to uncover novel biology in the mitochondria and kinetoplast. Among them, the best studied gene encodes β-ketoacyl-ACP reductase (*Tb927.2.5210*, 23-fold overrep.), a key member of the Type II Fatty Acid Biosynthesis (FAS) pathway in the mitochondria. Interestingly, lipoic acid, whose biosynthesis arises from Type II FAS, is an alternative redox active low molecular mass thiol alternative to T(SH)_2_ (31, 48). Further studies will be required to determine if overexpression of β-ketoacyl-ACP reductase and other novel mitochondrial proteins promote melarsoprol resistance either through their redox potential or other functions in the mitochondria.

The functionality of the *T. brucei* ORFeome could be extended to generate additional genetic tools, such as, yeast two-hybrid libraries, tagging libraries, and in dominant negative genetic screening approaches (22,26,49,50). Based on the conservation of orthologous gene clusters among kinetoplastida (3), we also predict the ORFeome could be used in other Trypanosomatids to generate orthologous Gain-of-Function libraries and other tools. The vast majority of genes overrepresented in melarsoprol survivor populations (∼80%, see SUP. 13) are conserved among sequenced kinetoplastida genomes. This supports the use of these tools to broadly expand our understanding of gene functions in this family of parasites. Notably, approximately 25% of all genes overrepresented in melarsoprol survivors were identified as resulting in loss-of-fitness in RIT-seq screening (26-23% depending on life cycle stage) (SUP. 12 – RIT-seq compared), which validates the usefulness of the GoF library tool in identifying essential genes, an acknowledged limitation of RNAi based screening (14, 51). Additionally, we found genes overrepresented in the melarsoprol GoF screen that were also identified in nifurtimox and melarsoprol RNAi based screens (SUP. 12 – RIT-seq compared) (51). Thus, we see the GoF library as a powerful new tool, complementary to RNAi knock-down approaches, to expand our understanding of drug targets and pathways of resistance. In future drug resistance screens, it will be useful to determine if certain sets of genes are commonly overrepresented in survivor populations and contrast that with genes that appear uniquely associated with an individual treatment. The tools and discoveries arising from this study are expected to support broad advances in basic biology, pathogenesis, pathways of drug resistance, and the discovery of new therapeutic targets in Trypanosomatid parasites.

## MATERIALS AND METHODS

### Oligo design and PCR amplification of the ORFeome

In order to design oligos for the ORFeome, we used gene annotations generated from ribosome profiling data (32). We first filtered out genes that were either < 100 bp in size or > 4,500bp in size. We then filtered out any genes annotated as pseudogenes, hypothetical protein unlikely, VSG, ISG, ESAG, GRESAG, or ribosomal protein. Following this procedure, we used custom python scripts to design forward and reverse primers. If the first 30bp of the ORF had a Tm >55, this primer was used. Otherwise, a nucleotide was added one at a time until the primer had a Tm of greater than 55. A reverse primer was designed using the same procedure for the reverse complement of the last 30 nucleotides. Many genes in the *T. brucei* genome have identical beginnings and ends, and we did not wish to order duplicate sets of primers that were identical to each other. For this reason, only one primer pair was kept for duplicated or highly similar genes, and a record was made of any pair that was eliminated based on duplication. All excluded ORFs are listed in SUP. 14. Finally, the ATT cloning sequences 5’-GGGGACAAGTTTGTACAAAAAAGCAGGCT and 5’-GGGGACCACTTTGTACAAGAAAGCTGGGT were added to the forward and reverse primers, respectively.

Each 384-well plate containing mixed forward and reverse primer pairs were diluted to 10 mM in primer dilution plates. PCR plates were filled with a master mix for KOD Hotstart (Novagen #71086) PCR reactions containing Lister427 gDNA according to manufacturer’s specifications and primer pairs specific to each ORF were transferred into the associated well number for each plate by TECAN Freedom EVO 150 pipetting instrument. PCR reaction conditions followed manufacturers specifications for 25 μL reactions with annealing temperature of 55°C and elongation times ranging from 10-60 seconds depending on the anticipated lengths of products in each plate. Following PCR reaction completion SYBR Green I (Invitrogen S-7563) was added to each well at 1:10 PCR volume for a final concentration of 10x SYBR and relative fluorescence units (RFU) were measured on a BioTek Synergy H4 Hybrid Multi-Mode Microplate reader in comparison to a standard dilution of known DNA concentration. The resulting PCR reactions from all wells of each 384-well plate were then pooled prior to agarose gel separation and DNA extraction. PCR reactions called ‘negative’ based on SYBR (less than 10,000 RFU) were revisited by isolating primer pairs from original 384-well plates, using a Perkin-Emer Janus Automated Workstation, into two new PCR plates (SUP. 2 – NEG_PICKS_#1 and NEG_PICKS_#2), which were then amplified using KOD Hotstart specifications and extracted from agarose gels as described. The resulting extracted DNAs from each PCR product pool were utilized in subsequent gateway cloning reactions.

### Gateway cloning and plasmids

The pENTR library was generated by cloning each size-sorted PCR product pool into pENTR 221 Gateway Entry vector according to manufacturer’s specifications (https://www.thermofisher.com/us/en/home/life-science/cloning/gateway-cloning.html) and transformed into One Shot *ccdB* Survival Competent Cells by electroporation. The resulting transformants were plated on large LB plates, assessed for efficiency of transformation, and bacterial colonies isolated from plates into LB liquid, which was split for maxi preps of plasmid and storage at −80°C in glycerol stocks. A *T. brucei* specific pDEST vector, pSUN6 (map shown in SUP. 3) was generated by introducing a *ccdB* gateway cassette into a pLEW type vector (52) for incorporation into the *T. brucei* genome based on rDNA spacer homology, blasticidin selection, and ORF transcription from an rDNA spacer promoter repressed by two tetracycline operators. Pools of pENTR plasmids harboring size-sorted ORF populations were combined with pSUN6 in LR clonase reactions and transfected into *ccdB* Survival Competent Cells following manufacturers guidelines as described. The resulting transformants were plated on large LB plates, assessed for efficiency of transformation, and bacteria and DNA isolated as described for pENTR steps above. The resulting extracted DNAs from both pENTR and pDEST ORFeome gateway cloning steps were assessed by NGS, described below. Following the initial assessment of ‘missing’ ORFs from both pENTR and pDEST cloning reactions, ‘missing’ PCR products were isolated from original plates, using a Perkin-Emer Janus Automated Workstation, to generate 8 new pools of size sorted PCR reactions (SUP. 4), which underwent the same series of pENTR and pDEST (pSUN6) cloning reactions described above and subjected to NGS analysis. The final set of ORFeome NGS validated (below) pDEST vectors are pooled to generate a single pTrypLib ORFeome for introduction into the *T. brucei* genome.

### T. brucei cell lines and transfections

Cell lines were generated from *Lister427* bloodstream-form trypanosomes derived from the ‘single marker’ (SM) line (53) and maintained in HMI-9 medium (54) under appropriate drug selection when indicated. A landing pad (LP) cell line was generated using plasmids gifted to us by the Alsford Lab and validated for inducible gene expression, prior to transfection with pRPaSce* as described (12, 51). LP parasites harboring the I-SceI restriction site and enzyme in the targeted rDNA spacer are doxycycline induced to permit ISceI cutting prior to pTrypLib ORFeome transfection by AMAXA Nucleofector (55). To generate the *T. brucei* GoF library described here, four 100ml flasks grown to ∼1 million cells/mL were AMAXA transfected in four separate transfection reactions, which were then pooled into a single cell population in 500 mL of HMI-9 containing blasticidin and recovered in a large roller flask (FIG. 3C). An additional four transfections were completed in parallel with TE (Mock) to compare outgrowth with GoF Library transfection. The resulting blasticidin recovered GoF Library population was expanded to an 800 mL culture at ∼1 million cells per mL and saved in aliquots of 100 million cells per vial for future genetic screens. Cells were also sampled prior to freezing for NSG analysis (‘GoF library’, described below) and after freeze thaw (INPUT1).

Single gene overexpression cell lines were generated by cloning ORFs of interest into pLEW100v5-BSD (https://www.addgene.org/27658/), which following validation were digested with NotI and transfected into SM by AMAXA.

### Quantitative PCR assessment of individual ORF induction

Individual cloned ORFs were selected randomly from pDEST colonies plated on LB originating from the pool ‘2_known’, ORFs confirmed by traditional DNA sequencing, and DNAs arising from 6 individual ORF harboring pDEST vectors were transfected into LP harboring pRPaSce* by AMAXA as described above. This generated a ‘Mini-Library’ following transfection and recovery, which was split into No Dox and +Dox conditions for 24 hours, RNA extracted, and cDNA prepared by Superscript III (ThermoFisher #18080044), prior to qPCR analysis. Quantitative PCR data was produced on a Bio-Rad CFX96 Real-Time PCR Detection System with iTaq Universal SYBR Green Supermix (Bio-Rad, #1725121) reagent using a forward primer unique to cloned ORFeome genes (5’-GGGGACAAGTTTGTACAAAAAAGCAGGCT) and reverse primer to each gene analyzed: *Tb927.8.2230* (PRIMER: 5’-CACGGTTTTTGCCCATTCGT), *Tb927.1.4830* (PRIMER: 5’-ATTTTTGCCGAAGCGCTTGA), *Tb927.10.12940* (PRIMER: 5’-CCGTGATTCCCTGTCGACAT), and *Tb927.11.15810* (PRIMER: 5’-CACCACCCGATGTACGGTAG). With these primer combinations only transcripts arising from the exogenous ORFeome copy were measured with and without Dox induction (FIG. 3B).

### Next-generation Illumina sequencing of the T. brucei ORFeome and Gain-of-Function libraries

Libraries for the pENTR and pDEST (pTrypLib) ORF plasmid libraries were prepared for next generation Illumina sequencing using tagmentation kits from Illumina (Nextera XT kit) according to the manufacturer’s instructions. Sequencing was performed either on an Illumina HiSeq 2500 or an Illumina MiSeq. Reads were trimmed for quality using trim galore (http://www.bioinformatics.babraham.ac.uk/projects/trim_galore/) using this command: trim_galore --nextera --stringency 3. Reads were aligned to the Tb927v5.1 genome using bowtie (56) requiring unique alignments: bowtie --best --strata -t -v 2 -a -m 1, or allowing multiple alignments: bowtie --best --strata -t -v 2 -a -m 10. RPKM values were calculated using SeqMonk from Babraham Bioinformatics (http://www.bioinformatics.babraham.ac.uk/projects/seqmonk).

For quantification of introduced ORFs following transfection into the *T. brucei* to generate the GoF library, genomic DNA was isolated from 100-200 million transfected parasites and fragmented by sonicating water bath (Bioruptor). The NEBNext Ultra II DNA library prep kit was then used to prepare libraries for high throughput sequencing according to the manufacturer’s instructions with the following exception: a custom universal primer containing the *ATT* cloning site was used for 5’ indexing and PCR enrichment (5’-AATGATACGGCGACCACCGAGATATATAACAAGTTTGTACAAAAAAGCAGGCTATG). A custom sequencing oligo (5’-GGGACAAGTTTGTAC AAAAAAGCAGGCTATG) is loaded on to the Illumina sequencing platform to sequence only GoF library containing reads. Analysis and raw counts were calculated using SeqMonk from Babraham Bioinformatics (http://www.bioinformatics.babraham.ac.uk/projects/seqmonk). Sequencing reads were trimmed for quality and aligned to the genome using the same parameters described above. Raw counts were calculated for reads aligning exclusively to the first 100 bp of the ORF using SeqMonk from Babraham Bioinformatics. DESeq2 was used to calculate the normalized read counts between sequencing libraries (38).

### Melarsoprol GoF genetic screening

For melarsoprol selective screening, GoF library cells were seeded for each condition at 1×10^5^ cell/ml, induced with doxycycline (1µg/mL) 24 hours (when appropriate), and grown in HMI-9 medium containing Dox (when appropriate) plus melarsoprol at 17 nM or 35 nM melarsoprol (BoC sciences, CAS 494-79-1). Melarsoprol stocks were diluted in DMSO.

### Bioinformatic analysis platforms for ORF identification and hit calling pipeline

In order to call which genes were overrepresented in the melarsoprol selected libraries, we exclusively counted reads that fell within the first 100bp of the gene. DeSeq (38) was then used to identify genes that were ‘differentially expressed’ in the melarsoprol selected libraries with a multiple testing corrected p value of less than 0.05. Using normalized read counts, we then calculated the fold change between each pair of melarsoprol selected and unselected replicates (3 total) for every statistically significant gene called by DESeq, provided that gene had a normalized read count of 5 or greater in the minimally propagated samples. If all 3 replicates showed a > 4 fold change for a particular gene, that gene was considered significantly overrepresented. Because we had two sets of replicates for unselected libraries and two sets of replicates for melarsoprol selected libraries, we ran the pipeline described above for all 4 comparisons of selected and unselected sets of replicates. Only those genes that were called as hits in all 4 comparisons were reported in the final hit list.

### EC50 determination by alamarblue

For EC50 determination, induced and uninduced cells were plated across a melarsoprol dilution series and assessed viability after 72 hours using Alamarblue (ThermoFisher) as previously described (57). All experiments were performed in biological triplicate.

## ACKNOWLEDGEMENTS

The authors would like to extend our sincere thanks to the individuals and consortiums whose support made this work possible. Dr. Marilyn Parsons (now of Seattle Children’s Hospital) provided prepublication access to the TREU927 ribosomal profiling data that provided the gene starts and stops for all ORFeome targeted ORFs. Drs. Christine Clayton and Esteban Erben (Heidelberg University) provided their essential insights into methods for successful ORFeome gateway cloning. Data provided directly from the tryptag.org consortium was critical in the categorization of hits arising from melarsoprol GoF screening. Similarly, tritrypdb.org was an essential resource throughout all stages of the work described herein. We thank Eugenia Silva-Herzog for her contribution to the generation of ‘mini-library’ overexpression data. Finally, we would like to thank Dr. F. Nina Papavasiliou, whose support and generosity have been invaluable.

## LIST OF ACRONYMS AND ABBREVIATIONS

ORF: Open reading frame
RIT: RNA-Interference Targeted
GOF: Gain-of-Function
RFU: Relative fluorescence units
DOX: doxycycline
SM: Single marker
LP: Landing pad
NGS: Next generation sequencing
GSH1: γ-glutamylcysteine synthetase
TR: Trypanothione reductase
T(SH)_2_: Trypanothione
GSH2: glutathione synthetase
ODC: Ornithine decarboxylase
TXN: Tryparedoxin
RR: Ribonucleotide reductase

## SUPPLEMENT LEGENDS

**Supplement 1. Table of oligo pairs from all 21, 384-well plates.** The sequences of all 7245 oligo pairs used to generate the *T. brucei* ORFeome are included with respect to their plate “name” and well position in each plate.

**Supplement 2. Table of ORFeome amplification tracking.** Table indicates the percent of PCR amplifications that were scored as positive (based on SYBR assessment) and the number of missing ORFs in reference to each oligo plate. Sub-tables show the reamplification of “NEG_PICKS” in two addition PCR plates, which recovered an additional 228 positive PCR products and the final total of 7044 positive PCR reactions for an overall success rate of 97.2%.

**Supplement 3. Map of *T. brucei* specific pDEST vector, pSUN6.** Critical features of plasmid map are indicated, including: rDNA spacer homology, T7 terminators, Blasticidin resistance cassette, rRNA promoter, two tetracycline operators, attR site for pENTR recombination, and the 3’ GPEET and 5’ Aldolase (long) UTRs.

**Supplement 4. Table of all final ORF pools used to make the pENTR ORFeome.** Following the first pENTR_1 and pDEST_1 ORFeome assessments, ORFs missing from libraries were identified and reselected from original 21 PCR plates to form an additional 8, size sorted pools. The original 21 pools, 2 pools of PCR reamplifications (NEG_PICKS) and 8 ‘missing’ ORF pools (1-8_MISS) collectively result in 31 pools for pENTR cloning. The size ranges and total number of ORFs per pool are indicated.

**Supplement 5. Tables of all missing genes from final pENTR and pDEST (pTrypLib) ORFeomes.** Consists of four sub-tables for all the ORFs identified as missing from either pENTR or pDEST by analysis of uniquely or multiply aligned reads: MISSING_pENTR_UNIQUE, MISSING_pENTR_MULTIPLE, MISSING_pDEST_UNIQUE, AND MISSING_pDEST_MULTIPLE.

**Supplement 6. Assessment of the pTrypLib ORFeome.** A) Histograms showing the distribution of normalized read counts in the pTrypLib ORFeome in analyses using uniquely aligning reads (left) and allowing multiple alignments (right). B) The length of each ORF in the pTrypLib Orfeome was plotted against the normalized read count of the ORF. A best fit line was calculated using linear regression (shown in white). For uniquely aligned reads this was y = 7.623751 - 0.000673x and for multiply aligned reads this was y = 7.306181 -0.000778x.

**Supplement 7. Sequencing strategy and results from transfected libraries.** A) Library preps from pTrypLib transfected parasites proceed by first fragmenting genomic DNA, ligating illumina adaptors, and amplifying library introduced ORFs using a primer complementary to complementary to the *attB1* Gateway cloning sequence and the standard Illumina barcoded reverse primer. Sequencing proceeds using a custom forward primer complementary to the *attB1* Gateway cloning sequence. B) Red rectangles in the first row represent annotated genes from a section of the *T. brucei* chromosome 5. Bars in subsequent rows represent reads that align to the genes in the first row. Most reads align to the first 100bp of the gene, as expected from the library prep and sequencing strategy.

**Supplement 8. Assessment of coverage in the GoF INPUT library.** A) Histograms showing the distribution of normalized read counts in the sequencing libraries from INPUT 1 in analyses using uniquely aligning reads (left) and allowing multiple alignments (right). B) The length of each ORF in the pTrypLib Orfeome was plotted against the normalized read count for the INPUT1 sequencing library. A best fit line was calculated using linear regression (shown in white). For uniquely aligned reads this was y = 6.046291 -0.000423x and for multiply aligned reads this was y = 6.083779 - 0.000452x.

**Supplement 9. Tables of all data pertaining to genes overrepresented in melarsoprol survivor populations.**

**Supplement 10. Table of categorization and localization of genes overrepresented in melarsoprol survivor populations.**

**Supplement 11. Table of trypanothione pathway genes and modest effects observed in melarsoprol GoF survivor populations.**

**Supplement 12. Table of melarsoprol GoF hits compared with RIT-seq genetic screens.**

**Supplement 13. Table conservation of the 57 genes overrepresented in melarsoprol survivors among other kinetoplastida.**

**Supplement 14. Table of genes not included in ORFeome design.**

## REFERENCES

1. Barrett MP, Burchmore RJS, Stich A, Lazzari JO, Frasch AC, Cazzulo JJ, Krishna S. The trypanosomiases. Lancet. 2003 Nov 1;362(9394):1469–80.

2. Fairlamb AH, Gow NAR, Matthews KR, Waters AP. Drug resistance in eukaryotic microorganisms. Nature Microbiology. 2016 Jun 24;1(7):555–15. PMCID: PMC5215055

3. El-Sayed NM, Myler PJ, Blandin G, Berriman M, Crabtree J, Aggarwal G, Caler E, Renauld H, Worthey EA, Hertz-Fowler C, Ghedin E, Peacock C, Bartholomeu DC, Haas BJ, Tran A-N, Wortman JR, Alsmark UCM, Angiuoli S, Anupama A, Badger J, Bringaud F, Cadag E, Carlton JM, Cerqueira GC, Creasy T, Delcher AL, Djikeng A, Embley TM, Hauser C, Ivens AC, Kummerfeld SK, Pereira-Leal JB, Nilsson D, Peterson J, Salzberg SL, Shallom J, Silva JC, Sundaram J, Westenberger S, White O, Melville SE, Donelson JE, Andersson B, Stuart KD, Hall N. Comparative genomics of trypanosomatid parasitic protozoa. Science (New York, N.Y.). American Association for the Advancement of Science; 2005 Jul 15;309(5733):404–9.

4. Akiyoshi B, Gull K. Evolutionary cell biology of chromosome segregation: insights from trypanosomes. Open Biol. 2013 May;3(5):130023. PMCID: PMC3866873

5. Matthews KR. 25 years of African trypanosome research: From description to molecular dissection and new drug discovery. Molecular & Biochemical Parasitology. 2015 Mar;200(1-2):30–40. PMCID: PMC4509711

6. Mani J, Meisinger C, Schneider A. Peeping at TOMs-Diverse Entry Gates to Mitochondria Provide Insights into the Evolution of Eukaryotes. Mol. Biol. Evol. 2016 Feb;33(2):337–51.

7. Jackson AP, Otto TD, Aslett M, Armstrong SD, Bringaud F, Schlacht A, Hartley C, Sanders M, Wastling JM, Dacks JB, Acosta-Serrano A, Field MC, Ginger ML, Berriman M. Kinetoplastid Phylogenomics Reveals the Evolutionary Innovations Associated with the Origins of Parasitism. Curr. Biol. 2016 Jan 25;26(2):161–72. PMCID: PMC4728078

8. Mayor S, Menon AK, Cross GA. Glycolipid precursors for the membrane anchor of Trypanosoma brucei variant surface glycoproteins. II. Lipid structures of phosphatidylinositol-specific phospholipase C sensitive and resistant glycolipids. J. Biol. Chem. 1990 Apr 15;265(11):6174–81.

9. Tschudi C, Ullu E. Trypanosomatid protozoa provide paradigms of eukaryotic biology. Infect Agents Dis. 1994 Aug;3(4):181–6.

10. Simpson L, Thiemann OH, Savill NJ, Alfonzo JD, Maslov DA. Evolution of RNA editing in trypanosome mitochondria. Proc Natl Acad Sci USA. National Academy of Sciences; 2000 Jun 20;97(13):6986–93. PMCID: PMC34374

11. Alsford S, Kelly JM, Baker N, Horn D. Genetic dissection of drug resistance in trypanosomes. Parasitology. George Washington Libraries; 2013 Apr 3;140(12):1478–91.

12. Glover L, Alsford S, Baker N, Turner DJ, Sanchez-Flores A, Hutchinson S, Hertz-Fowler C, Berriman M, Horn D. Genome-scale RNAi screens for high-throughput phenotyping in bloodstream-form African trypanosomes. Nat Protoc. 2015 Jan;10(1):106–33.

13. Currier RB, Cooper A, Burrell-Saward H, Macleod A, Alsford S. Decoding the network of Trypanosoma brucei proteins that determines sensitivity to apolipoprotein-L1. Raper J, editor. PLoS Pathog. 2018 Jan 18;14(1):e1006855–26.

14. Alsford S, Turner DJ, Obado SO, Sanchez-Flores A, Glover L, Berriman M, Hertz-Fowler C, Horn D. High-throughput phenotyping using parallel sequencing of RNA interference targets in the African trypanosome. Genome Research. Cold Spring Harbor Lab; 2011 Jun 1;21(6):915–24. PMCID: PMC3106324

15. Normark S, Edlund T, Grundström T, Bergström S, Wolf-Watz H. Escherichia coli K-12 mutants hyperproducing chromosomal beta-lactamase by gene repetitions. J. Bacteriol. American Society for Microbiology (ASM); 1977 Dec;132(3):912–22. PMCID: PMC235595

16. Rine J, Hansen W, Hardeman E, Davis RW. Targeted selection of recombinant clones through gene dosage effects. Proc Natl Acad Sci USA. National Academy of Sciences; 1983 Nov;80(22):6750–4. PMCID: PMC390063

17. Sandegren L, Andersson DI. Bacterial gene amplification: implications for the evolution of antibiotic resistance. Nat. Rev. Microbiol. Nature Publishing Group; 2009 Aug;7(8):578–88.

18. Prelich G. Gene overexpression: uses, mechanisms, and interpretation. Genetics. Genetics; 2012 Mar;190(3):841–54. PMCID: PMC3296252

19. Clayton CE. Gene expression in Kinetoplastids. Curr. Opin. Microbiol. 2016 Aug;32:46–51.

20. Begolo D, Erben E, Clayton C. Drug target identification using a trypanosome overexpression library. Antimicrob. Agents Chemother. American Society for Microbiology Journals; 2014 Oct;58(10):6260–4. PMCID: PMC4187942

21. Wall RJ, Rico E, Lukac I, Zuccotto F, Elg S, Gilbert IH, Freund Y, Alley MRK, Field MC, Wyllie S, Horn D. Clinical and veterinary trypanocidal benzoxaboroles target CPSF3. Proc. Natl. Acad. Sci. U.S.A. National Academy of Sciences; 2018 Sep 18;115(38):9616–21. PMCID: PMC6156652

22. Rual J-F, Hill DE, Vidal M. ORFeome projects: gateway between genomics and omics. Curr Opin Chem Biol. 2004 Feb;8(1):20–5.

23. Li Q-R, Carvunis A-R, Yu H, Han J-DJ, Zhong Q, Simonis N, Tam S, Hao T, Klitgord NJ, Dupuy D, Mou D, Wapinski I, Regev A, Hill DE, Cusick ME, Vidal M. Revisiting the Saccharomyces cerevisiae predicted ORFeome. Genome Research. Cold Spring Harbor Lab; 2008 Aug;18(8):1294–303. PMCID: PMC2493439

24. Gallegos ME, Balakrishnan S, Chandramouli P, Arora S, Azameera A, Babushekar A, Bargoma E, Bokhari A, Chava SK, Das P, Desai M, Decena D, Saramma SDD, Dey B, Doss A-L, Gor N, Gudiputi L, Guo C, Hande S, Jensen M, Jones S, Jones N, Jorgens D, Karamchedu P, Kamrani K, Kolora LD, Kristensen L, Kwan K, Lau H, Maharaj P, Mander N, Mangipudi K, Menakuru H, Mody V, Mohanty S, Mukkamala S, Mundra SA, Nagaraju S, Narayanaswamy R, Ndungu-Case C, Noorbakhsh M, Patel J, Patel P, Pendem SV, Ponakala A, Rath M, Robles MC, Rokkam D, Roth C, Sasidharan P, et al. The C. elegans rab family: identification, classification and toolkit construction. D’Adamo P, editor. PLoS ONE. 2012;7(11):e49387. PMCID: PMC3504004

25. Mehla J, Caufield JH, Uetz P. Mapping Protein–Protein Interactions Using Yeast Two-Hybrid Assays. Cold Spring Harb Protoc. 2015 May 1;2015(5):pdb.prot086157–12.

26. Bischof J, Duffraisse M, Furger E, Ajuria L, Giraud G, Vanderperre S, Paul R, Björklund M, Ahr D, Ahmed AW, Spinelli L, Brun C, Basler K, Merabet S. Generation of a versatile BiFC ORFeome library for analyzing protein-protein interactions in live Drosophila. Elife. eLife Sciences Publications Limited; 2018 Sep 24;7:1251. PMCID: PMC6177257

27. King J, Foster J, Davison JM, Rawls JF, Breton G. Zebrafish Transcription Factor ORFeome for Gene Discovery and Regulatory Network Elucidation. Zebrafish. Mary Ann Liebert, Inc. 140 Huguenot Street, 3rd Floor New Rochelle, NY 10801 USA; 2018 Apr;15(2):202–5. PMCID: PMC6112160

28. Trindade S, Rijo-Ferreira F, Carvalho T, Pinto-Neves D, Guegan F, Aresta-Branco F, Bento F, Young SA, Pinto A, Van Den Abbeele J, Ribeiro RM, Dias S, Smith TK, Figueiredo LM. Trypanosoma brucei Parasites Occupy and Functionally Adapt to the Adipose Tissue in Mice. Cell Host Microbe. Elsevier Inc; 2016 Jun 8;19(6):837–48.

29. Deeks ED. Fexinidazole: First Global Approval. Drugs. 2019 Feb;79(2):215–20.

30. Fairlamb AH, Horn D. Melarsoprol Resistance in African Trypanosomiasis. Elsevier Ltd; 2018 May 10;:1–12.

31. Krauth-Siegel RL, Comini MA. Redox control in trypanosomatids, parasitic protozoa with trypanothione-based thiol metabolism. Biochimica et Biophysica Acta (BBA) - General Subjects. 2008 Nov;1780(11):1236–48.

32. Parsons M, Ramasamy G, Vasconcelos EJR, Jensen BC, Myler PJ. Advancing Trypanosoma brucei genome annotation through ribosome profiling and spliced leader mapping. Molecular & Biochemical Parasitology. 2015 Aug;202(2):1–10. PMCID: PMC4644489

33. Shahi SK, Krauth-Siegel RL, Clayton CE. Overexpression of the putative thiol conjugate transporter TbMRPA causes melarsoprol resistance in Trypanosoma brucei. Mol. Microbiol. John Wiley & Sons, Ltd (10.1111); 2002 Mar;43(5):1129–38.

34. Graf FE, Ludin P, Wenzler T, Kaiser M, Brun R, Pyana PP, Büscher P, de Koning HP, Horn D, Mäser P. Aquaporin 2 Mutations in Trypanosoma brucei gambiense Field Isolates Correlate with Decreased Susceptibility to Pentamidine and Melarsoprol. Matovu E, editor. PLoS Negl Trop Dis. 2013 Oct 10;7(10):e2475–7. PMCID: PMC3794916

35. Baker N, de Koning HP, Mäser P, Horn D. Drug resistance in Africantrypanosomiasis: the melarsoprol andpentamidine story. Trends Parasitol. Elsevier Ltd; 2013 Mar 1;29(3):110–8.

36. Munday JC, Eze AA, Baker N, Glover L, Clucas C, Aguinaga Andrés D, Natto MJ, Teka IA, McDonald J, Lee RS, Graf FE, Ludin P, Burchmore RJS, Turner CMR, Tait A, Macleod A, Mäser P, Barrett MP, Horn D, de Koning HP. Trypanosoma brucei aquaglyceroporin 2 is a high-affinity transporter for pentamidine and melaminophenyl arsenic drugs and the main genetic determinant of resistance to these drugs. J. Antimicrob. Chemother. 2014 Mar;69(3):651–63. PMCID: PMC3922157

37. Graf FE, Ludin P, Arquint C, Schmidt RS, Schaub N, Renggli CK, Munday JC, Krezdorn J, Baker N, Horn D, Balmer O, Caccone A, Koning HP, Mäser P. Comparative genomics of drug resistance in Trypanosoma brucei rhodesiense. Cellular and Molecular Life Sciences. Springer International Publishing; 2016 Mar 13;73(17):3387–400. PMCID: PMC4967103

38. Love MI, Huber W, Anders S. Moderated estimation of fold change and dispersion for RNA-seq data with DESeq2. Genome Biol. BioMed Central; 2014;15(12):550–21. PMCID: PMC4302049

39. Lueder DV, Phillips MA. Characterization of Trypanosoma brucei gamma-glutamylcysteine synthetase, an essential enzyme in the biosynthesis of trypanothione (diglutathionylspermidine). J. Biol. Chem. American Society for Biochemistry and Molecular Biology; 1996 Jul 19;271(29):17485–90.

40. Huynh TT, Huynh VT, Harmon MA, Phillips MA. Gene Knockdown of γ-Glutamylcysteine Synthetase by RNA iin the Parasitic Protozoa Trypanosoma bruceiDemonstrates That It Is an Essential Enzyme. J. Biol. Chem. 2003 Oct 3;278(41):39794–800.

41. Singh A, Minia I, Droll D, Fadda A, Clayton C, Erben E. Trypanosome MKT1 and the RNA-binding protein ZC3H11: interactions and potential roles in post-transcriptional regulatory networks. Nucleic Acids Res. 2014 Jan 25;42(7):4652–68.

42. Alsford S, Turner DJ, Obado SO, Sanchez-Flores A, Glover L, Berriman M, Hertz-Fowler C, Horn D. High-throughput phenotyping using parallel sequencing of RNA interference targets in the African trypanosome. Genome Research. 2011 Jun 1;21(6):915–24. PMCID: PMC3106324

43. Motyka SA, Drew ME, Yildirir G, Englund PT. Overexpression of a cytochrome b5 reductase-like protein causes kinetoplast DNA loss in Trypanosoma brucei. J. Biol. Chem. American Society for Biochemistry and Molecular Biology; 2006 Jul 7;281(27):18499–506.

44. Thomas JA, Baker N, Hutchinson S, Dominicus C, Trenaman A, Glover L, Alsford S, Horn D. Insights into antitrypanosomal drug mode-of-action from cytology-based profiling. Clos J, editor. PLoS Negl Trop Dis. Public Library of Science; 2018 Nov 26;12(11):e0006980–19.

45. Alsford S, Eckert S, Baker N, Glover L, Sanchez-Flores A, Leung KF, Turner DJ, Field MC, Berriman M, Horn D. High-throughput decoding of antitrypanosomal drug efficacy and resistance. Nature. Nature Publishing Group; 2019 Apr 24;482(7384):232–6.

46. Baker N, Glover L, Munday JC, Aguinaga Andrés D, Barrett MP, de Koning HP, Horn D. Aquaglyceroporin 2 controls susceptibility to melarsoprol and pentamidine in African trypanosomes. Proc. Natl. Acad. Sci. U.S.A. National Academy of Sciences; 2012 Jul 3;109(27):10996–1001. PMCID: PMC3390834

47. Fidalgo LM, Gille L. Mitochondria and Trypanosomatids: Targets and Drugs. Pharm Res. 2011 Sep 21;28(11):2758–70.

48. Stephens JL, Lee SH, Paul KS, Englund PT. Mitochondrial fatty acid synthesis in Trypanosoma brucei. J. Biol. Chem. 2007 Feb 16;282(7):4427–36.

49. Grant IM, Balcha D, Hao T, Shen Y, Trivedi P, Patrushev I, Fortriede JD, Karpinka JB, Liu L, Zorn AM, Stukenberg PT, Hill DE, Gilchrist MJ. The Xenopus ORFeome: A resource that enables functional genomics. Dev. Biol. 2015 Dec 15;408(2):345–57. PMCID: PMC4684507

50. Mehla J, Caufield JH, Uetz P. Mapping Protein–Protein Interactions Using Yeast Two-Hybrid Assays. Cold Spring Harb Protoc. 2015 May 1;2015(5):pdb.prot086157–12.

51. Alsford S, Eckert S, Baker N, Glover L, Sanchez-Flores A, Leung KF, Turner DJ, Field MC, Berriman M, Horn D. High-throughput decoding of antitrypanosomal drug efficacy and resistance. Nature. Nature Publishing Group; 2012 Jan 25;482(7384):232–6. PMCID: PMC3303116

52. Siegel TN, Tan KSW, Cross GAM. Systematic study of sequence motifs for RNA trans splicing in Trypanosoma brucei. Mol. Cell. Biol. 2005 Nov;25(21):9586–94. PMCID: PMC1265811

53. Wirtz E, Leal S, Ochatt C, Cross GA. A tightly regulated inducible expression system for conditional gene knock-outs and dominant-negative genetics in Trypanosoma brucei. Molecular & Biochemical Parasitology. 1999 Mar 15;99(1):89–101.

54. Hirumi H, Hirumi K. Axenic culture of African trypanosome bloodstream forms. Parasitol. Today (Regul. Ed.). 1994 Feb;10(2):80–4.

55. Burkard G, Fragoso CM, Roditi I. Highly efficient stable transformation of bloodstream forms of Trypanosoma brucei. Molecular & Biochemical Parasitology. 2007 Jun;153(2):220–3.

56. Langmead B, Trapnell C, Pop M, Salzberg SL. Ultrafast and memory-efficient alignment of short DNA sequences to the human genome. Genome Biol. BioMed Central; 2009;10(3):R25–10. PMCID: PMC2690996

57. Baker N, Alsford S, Horn D. Genome-wide RNAi screens in African trypanosomes identify the nifurtimox activator NTR and the eflornithine transporter AAT6. Molecular & Biochemical Parasitology. Elsevier B.V; 2011 Mar 1;176(1):55–7. PMCID: PMC3032052

