## Supplementary material for "A *Trypanosoma brucei* ORFeome-based Gain-of-Function Library reveals novel genes associated with melarsoprol resistance": SUP. 1

| Plate Name | Well Postion | Forward primer name | Forward primer sequence | Reverse primer name | Reverse primer sequence |
|---|---|---|---|---|---|
| 0_known | A1 | Tb927.9.4961_for | GGGGACAAGTTTGTACAAAAAAGCAGGCTATGCCACCGAAGAGCAACG | Tb927.9.4961_rev | GGGGACCACTTTGTACAAGAAAGCTGGGTTCAACGCACAGACTTGGGTTTG |
| 0_known | A2 | Tb927.10.14615_for | GGGGACAAGTTTGTACAAAAAAGCAGGCTATGCAGCAGGCGGACTATTATG | Tb927.10.14615_rev | GGGGACCACTTTGTACAAGAAAGCTGGGTTTACCATGAATTAACGACCAACTTAAATAT |
| 0_known | A3 | Tb927.10.4740_for | GGGGACAAGTTTGTACAAAAAAGCAGGCTATGCACCTTCAAGTTTATCTCGTGG | Tb927.10.4740_rev | GGGGACCACTTTGTACAAGAAAGCTGGGTTTAAAGTGGCTTTGGGCGTTTTTC |
| 0_known | A4 | Tb927.3.960_for | GGGGACAAGTTTGTACAAAAAAGCAGGCTATGGATTTCCTGGATGAAACCGTG | Tb927.3.960_rev | GGGGACCACTTTGTACAAGAAAGCTGGGTTCAAGCGCCGAGTACAACATTATT |
| 0_known | A5 | Tb927.11.1870_for | GGGGACAAGTTTGTACAAAAAAGCAGGCTATGGCGAAGGCATCTGCTG | Tb927.11.1870_rev | GGGGACCACTTTGTACAAGAAAGCTGGGTTCACGCCTTCTTAGCGGCG |
| 0_known | A6 | Tb927.8.7430_for | GGGGACAAGTTTGTACAAAAAAGCAGGCTATGGCCGACGAGGAACCA | Tb927.8.7430_rev | GGGGACCACTTTGTACAAGAAAGCTGGGTTTACACCAACGTTGGCCACAG |
| 0_known | A8 | Tb927.11.3705_for | GGGGACAAGTTTGTACAAAAAAGCAGGCTATGATCATTCCTGTTCGGTGTTTCA | Tb927.11.3705_rev | GGGGACCACTTTGTACAAGAAAGCTGGGTTTACACCGCAACACTGGAGTCA |
| 0_known | A9 | Tb927.10.13700_for | GGGGACAAGTTTGTACAAAAAAGCAGGCTATGCCGAGACCGTGCATTC | Tb927.10.13700_rev | GGGGACCACTTTGTACAAGAAAGCTGGGTTCATTTAGTTATATCAGACACCACCTGC |
| 0_known | A10 | Tb927.3.2650_for | GGGGACAAGTTTGTACAAAAAAGCAGGCTATGTCGAGTGACGAAGTTGCTG | Tb927.3.2650_rev | GGGGACCACTTTGTACAAGAAAGCTGGGTCTAAAAACTCCGGACACTCCCTCG |
| 0_known | A11 | Tb927.9.10250_for | GGGGACAAGTTTGTACAAAAAAGCAGGCTATGGATGCAAATGTACCGGCG | Tb927.9.10250_rev | GGGGACCACTTTGTACAAGAAAGCTGGGTCTATTCCTTTGGTACTTCCCGAATGTAC |
| 0_known | A12 | Tb927.11.1880_for | GGGGACAAGTTTGTACAAAAAAGCAGGCTATGGCGAAGACAACTGCTGC | Tb927.11.1880_rev | GGGGACCACTTTGTACAAGAAAGCTGGGTTCACGCCTTCTTAGCGGCG |
| 0_known | A15 | Tb927.4.2540_for | GGGGACAAGTTTGTACAAAAAAGCAGGCTATGCTTCTTAAGGTGAAGACTGTAAGC | Tb927.4.2540_rev | GGGGACCACTTTGTACAAGAAAGCTGGGTTCAGCAACCCGCGCGCAG |
| 0_known | A16 | Tb927.11.1800_for | GGGGACAAGTTTGTACAAAAAAGCAGGCTATGAACAACACAACCGCTACTGT | Tb927.11.1800_rev | GGGGACCACTTTGTACAAGAAAGCTGGGTTCACGCCTTCTTAGCGGCG |
| 0_known | A17 | Tb927.7.7380_for | GGGGACAAGTTTGTACAAAAAAGCAGGCTATGGATCCCTTCGTGTTTCTGC | Tb927.7.7380_rev | GGGGACCACTTTGTACAAGAAAGCTGGGTTCAGTTAACCTGATACGATTGGGGA |
| 0_known | A18 | Tb927.9.10170_for | GGGGACAAGTTTGTACAAAAAAGCAGGCTATGGATATAAAAGTGAAGAGTGCCCAC | Tb927.9.10170_rev | GGGGACCACTTTGTACAAGAAAGCTGGGTTTAGAATTCCCATGTCTGTCGGCA |
| 0_known | A19 | Tb927.11.14310_for | GGGGACAAGTTTGTACAAAAAAGCAGGCTATGCCCGCCAAGAAGCGC | Tb927.11.14310_rev | GGGGACCACTTTGTACAAGAAAGCTGGGTTTACACTTCAAGACCGACAATATCCAC |
| 0_known | A20 | Tb927.9.13070_for | GGGGACAAGTTTGTACAAAAAAGCAGGCTATGGACTACGAAGAACCAGGACA | Tb927.9.13070_rev | GGGGACCACTTTGTACAAGAAAGCTGGGTTTATTTCCCCTTACTCTTTTTCTTATCTTT |
| 0_known | A21 | Tb927.1.1170_for | GGGGACAAGTTTGTACAAAAAAGCAGGCTATGCTTTCGTATACAGTGAAGGAGGA | Tb927.1.1170_rev | GGGGACCACTTTGTACAAGAAAGCTGGGTTCAAATAGTGTCATATGTTGTGGGTTGC |
| 0_known | A22 | Tb927.3.1250_for | GGGGACAAGTTTGTACAAAAAAGCAGGCTATGCTGATTCCAGTCCGTTGC | Tb927.3.1250_rev | GGGGACCACTTTGTACAAGAAAGCTGGGTTTACTGCCGGCGTGGCAC |
| 0_known | A23 | Tb927.9.3530_for | GGGGACAAGTTTGTACAAAAAAGCAGGCTATGGCTTCGGTGAATATGGAGGA | Tb927.9.3530_rev | GGGGACCACTTTGTACAAGAAAGCTGGGTCTAAGAAGTGGTGGAAGCCACAC |
| 0_known | A24 | Tb927.6.2700_for | GGGGACAAGTTTGTACAAAAAAGCAGGCTATGAGCGTCACAACAAAGCAGA | Tb927.6.2700_rev | GGGGACCACTTTGTACAAGAAAGCTGGGTCTATATGCCAATAGGGTGAATGACACC |
| 0_known | B1 | Tb927.10.8700_for | GGGGACAAGTTTGTACAAAAAAGCAGGCTATGGTGTACATTGAGAACGGCAG | Tb927.10.8700_rev | GGGGACCACTTTGTACAAGAAAGCTGGGTTCATCCACTCGCGCAACCG |
| 0_known | B2 | Tb927.11.11435_for | GGGGACAAGTTTGTACAAAAAAGCAGGCTATGACGCAGCTCCCCTCAG | Tb927.11.11435_rev | GGGGACCACTTTGTACAAGAAAGCTGGGTTCATGACGACTTGAAAAGAAGCACA |
| 0_known | B3 | Tb927.11.711_for | GGGGACAAGTTTGTACAAAAAAGCAGGCTATGTCTACTGATCGGAAAGCTATTATTAAG | Tb927.11.711_rev | GGGGACCACTTTGTACAAGAAAGCTGGGTCTATCCTGACTTGAACAGAAGAATTGCA |
| 0_known | B4 | Tb927.5.4440_for | GGGGACAAGTTTGTACAAAAAAGCAGGCTATGTCTAAGCACCATGAATTCAACATTAAC | Tb927.5.4440_rev | GGGGACCACTTTGTACAAGAAAGCTGGGTTTACGTCACTGTCTTCCACAAAAGG |
| 0_known | B5 | Tb927.11.320_for | GGGGACAAGTTTGTACAAAAAAGCAGGCTATGATGAGTGATCGCAAGACCAATG | Tb927.11.320_rev | GGGGACCACTTTGTACAAGAAAGCTGGGTTCAACCTGTTTTCCACAGGAGAATG |
| 0_known | B6 | Tb927.9.13820_for | GGGGACAAGTTTGTACAAAAAAGCAGGCTATGGCCACCACATACGAAGAATTT | Tb927.9.13820_rev | GGGGACCACTTTGTACAAGAAAGCTGGGTTCATTTTCCGGGGAACTGGGC |
| 0_known | B8 | Tb927.9.13775_for | GGGGACAAGTTTGTACAAAAAAGCAGGCTATGGCCACTACATACGAAGAATTTGC | Tb927.9.13775_rev | GGGGACCACTTTGTACAAGAAAGCTGGGTTCATTTTCCGGGGAACTGGGC |
| 0_known | B9 | Tb927.9.13785_for | GGGGACAAGTTTGTACAAAAAAGCAGGCTATGGCCACCACATACGAAGAATTT | Tb927.9.13785_rev | GGGGACCACTTTGTACAAGAAAGCTGGGTTCACTTTCCGGGGAACTGGG |
| 0_known | B11 | Tb927.5.4030_for | GGGGACAAGTTTGTACAAAAAAGCAGGCTATGTCGGCACAGCAGGCA | Tb927.5.4030_rev | GGGGACCACTTTGTACAAGAAAGCTGGGTTTACTGGAATGGGTTCTGTTGAAGGA |
| 0_known | B12 | Tb927.4.2010_for | GGGGACAAGTTTGTACAAAAAAGCAGGCTATGGTGGAGGAAACGCTTGATG | Tb927.4.2010_rev | GGGGACCACTTTGTACAAGAAAGCTGGGTCTACAGGTTCGACGGGGGC |
| 0_known | B13 | Tb927.11.1370_for | GGGGACAAGTTTGTACAAAAAAGCAGGCTATGCCCTCTATCGCTTCGATGA | Tb927.11.1370_rev | GGGGACCACTTTGTACAAGAAAGCTGGGTTTAGCTCAGCAAACCATCGAGTTT |
| 0_known | B14 | Tb927.8.5380_for | GGGGACAAGTTTGTACAAAAAAGCAGGCTATGTCTCAAAATGAAGAAGCCCCTG | Tb927.8.5380_rev | GGGGACCACTTTGTACAAGAAAGCTGGGTTTAAACCCTGCCCACACGGT |
| 0_known | B15 | Tb927.11.16390_for | GGGGACAAGTTTGTACAAAAAAGCAGGCTATGTCTGGCCGGGACTACTTC | Tb927.11.16390_rev | GGGGACCACTTTGTACAAGAAAGCTGGGTCTATTGTGTCTTGGGCCGCTT |
| 0_known | B16 | Tb927.9.14480_for | GGGGACAAGTTTGTACAAAAAAGCAGGCTATGTCTGAGATAGAGGAAAGCTTCCA | Tb927.9.14480_rev | GGGGACCACTTTGTACAAGAAAGCTGGGTTCAAACATCTGAAAATCGTTGAATTACAAT |
| 0_known | B17 | Tb927.10.12000_for | GGGGACAAGTTTGTACAAAAAAGCAGGCTATGACAAACACCGTCAAACACTCA | Tb927.10.12000_rev | GGGGACCACTTTGTACAAGAAAGCTGGGTTTACTCTTCTTCTTCTTGCGTCACG |
| 0_known | B18 | Tb927.7.1340_for | GGGGACAAGTTTGTACAAAAAAGCAGGCTATGCTCCGCGTTTCCATTCC | Tb927.7.1340_rev | GGGGACCACTTTGTACAAGAAAGCTGGGTTCAATTCTGGATAACACCAAGCAACG |
| 0_known | B19 | Tb927.5.4260_for | GGGGACAAGTTTGTACAAAAAAGCAGGCTATGGCGAAGGGTAAGAAGAGTGG | Tb927.5.4260_rev | GGGGACCACTTTGTACAAGAAAGCTGGGTTTATGCATAACCGTACAGAATCTTGCC |
| 0_known | B24 | Tb927.2.2670_for | GGGGACAAGTTTGTACAAAAAAGCAGGCTATGGCAAAGGGCAAGAGAGTCG | Tb927.2.2670_rev | GGGGACCACTTTGTACAAGAAAGCTGGGTTCAGTCGTACCCGTAAAGAACCT |
| 0_known | C7 | Tb927.11.9700_for | GGGGACAAGTTTGTACAAAAAAGCAGGCTATGCCCTCTATTACTCAGGAGACG | Tb927.11.9700_rev | GGGGACCACTTTGTACAAGAAAGCTGGGTCTATGAAGGCACTTCGCTCGC |
| 0_known | C8 | Tb927.6.3490_for | GGGGACAAGTTTGTACAAAAAAGCAGGCTATGCACATTTCAACACCCACTCT | Tb927.6.3490_rev | GGGGACCACTTTGTACAAGAAAGCTGGGTTCAATCCTCCAACCTTACGCTGT |
| 0_known | C9 | Tb927.7.3120_for | GGGGACAAGTTTGTACAAAAAAGCAGGCTATGCCCGCGGCGGAGTCC | Tb927.7.3120_rev | GGGGACCACTTTGTACAAGAAAGCTGGGTTCACTCATTTGATCGCTCCGTCC |
| 0_known | C10 | Tb927.5.3745_for | GGGGACAAGTTTGTACAAAAAAGCAGGCTATGAGTAATACAAATTCGTCTGGGTGC | Tb927.5.3745_rev | GGGGACCACTTTGTACAAGAAAGCTGGGTTTACGAACGTGCAATACGTTCAGC |
| 0_known | C11 | Tb927.9.3370_for | GGGGACAAGTTTGTACAAAAAAGCAGGCTATGTCGGTAGTGGATGTTTATAGCGT | Tb927.9.3370_rev | GGGGACCACTTTGTACAAGAAAGCTGGGTTTAGTTGTCTTTGATGATATCTCTTAATTT |
| 0_known | C12 | Tb927.1.1770_for | GGGGACAAGTTTGTACAAAAAAGCAGGCTATGAATAACGCATTGGATCCGGC | Tb927.1.1770_rev | GGGGACCACTTTGTACAAGAAAGCTGGGTTCAGCCCCTTTCAATTTCGGC |
| 0_known | C13 | Tb927.10.4620_for | GGGGACAAGTTTGTACAAAAAAGCAGGCTATGCCGCTTCAGTACGACATC | Tb927.10.4620_rev | GGGGACCACTTTGTACAAGAAAGCTGGGTTCAGTTAATTGCGACGAGTTCCATT |
| 0_known | C14 | Tb927.11.5840_for | GGGGACAAGTTTGTACAAAAAAGCAGGCTATGAATGCAAACGACGAGATGAGC | Tb927.11.5840_rev | GGGGACCACTTTGTACAAGAAAGCTGGGTTCAGTTCAGCGAGTGGATCTCTAA |
| 0_known | C15 | Tb927.10.10320_for | GGGGACAAGTTTGTACAAAAAAGCAGGCTATGAAAGAGAAGTTGACTCTAGCTGATG | Tb927.10.10320_rev | GGGGACCACTTTGTACAAGAAAGCTGGGTCTATGCCACAACACCGTACACA |
| 0_known | C16 | Tb927.9.5220_for | GGGGACAAGTTTGTACAAAAAAGCAGGCTATGGGCGGCACCGGAGAG | Tb927.9.5220_rev | GGGGACCACTTTGTACAAGAAAGCTGGGTCTAAGTTCGCATAATGTGCTTGTTTTCG |
| 0_known | C17 | Tb927.8.2310_for | GGGGACAAGTTTGTACAAAAAAGCAGGCTATGCCTCCCAAGTATGACAATGTCC | Tb927.8.2310_rev | GGGGACCACTTTGTACAAGAAAGCTGGGTTCAGTTGTCCTGACATCGCAGT |
| 0_known | C18 | Tb927.2.4540_for | GGGGACAAGTTTGTACAAAAAAGCAGGCTATGGGCCACCAAAATATGCTTCA | Tb927.2.4540_rev | GGGGACCACTTTGTACAAGAAAGCTGGGTTCAATCGCGTTTCCGCTTGG |
| 0_known | C19 | Tb927.3.2030_for | GGGGACAAGTTTGTACAAAAAAGCAGGCTATGCCACAAACTGTAGATCAGGTTG | Tb927.3.2030_rev | GGGGACCACTTTGTACAAGAAAGCTGGGTTCATCGCAGTATCTCGAATGTATCAAAA |
| 0_known | C20 | Tb927.10.9620_for | GGGGACAAGTTTGTACAAAAAAGCAGGCTATGCGGCTGCCGCATCCG | Tb927.10.9620_rev | GGGGACCACTTTGTACAAGAAAGCTGGGTTTACACGAGGGGTAACAAAAAAAAAAAGA |
| 0_known | C21 | Tb927.1.4570_for | GGGGACAAGTTTGTACAAAAAAGCAGGCTATGTTTGTTTTTATATGCCGTGACTTTGT | Tb927.1.4570_rev | GGGGACCACTTTGTACAAGAAAGCTGGGTTTAGTGTGTAATGATGTCATGCAACTGG |
| 0_known | C22 | Tb927.2.5850_for | GGGGACAAGTTTGTACAAAAAAGCAGGCTATGTCTGGTGAGGAACCCACTAAA | Tb927.2.5850_rev | GGGGACCACTTTGTACAAGAAAGCTGGGTCTATACAGTGCTCTGCGGCAAC |
| 0_known | C23 | Tb927.10.10600_for | GGGGACAAGTTTGTACAAAAAAGCAGGCTATGAAAGAATTAAAAGAACTTCTGGATCAG | Tb927.10.10600_rev | GGGGACCACTTTGTACAAGAAAGCTGGGTTTATTTTTTTTCGTTTGATTTGTTTTGCGC |
| 0_known | C24 | Tb927.10.10570_for | GGGGACAAGTTTGTACAAAAAAGCAGGCTATGGCCACTCCTAAGAGCACA | Tb927.10.10570_rev | GGGGACCACTTTGTACAAGAAAGCTGGGTTTAGCTGGAAGCGTGTGACAC |
| 0_known | D1 | Tb927.7.3420_for | GGGGACAAGTTTGTACAAAAAAGCAGGCTATGTCGCGAAATGACTGCGT | Tb927.7.3420_rev | GGGGACCACTTTGTACAAGAAAGCTGGGTTCAAACAACATCCAACAGCGTGA |
| 0_known | D7 | Tb927.10.4690_for | GGGGACAAGTTTGTACAAAAAAGCAGGCTATGACGTCCAACGTGAAGGAAC | Tb927.10.4690_rev | GGGGACCACTTTGTACAAGAAAGCTGGGTCTACAACCCCTTGAAGACGACAATT |
| 0_known | D10 | Tb927.10.10451_for | GGGGACAAGTTTGTACAAAAAAGCAGGCTATGGCCACTCCTAGGAGCAC | Tb927.10.10451_rev | GGGGACCACTTTGTACAAGAAAGCTGGGTTTAGCTGGAAGCGTGTGACAC |
| 0_known | D18 | Tb927.11.9040_for | GGGGACAAGTTTGTACAAAAAAGCAGGCTATGTCCCGTGTCCCTATTAGCG | Tb927.11.9040_rev | GGGGACCACTTTGTACAAGAAAGCTGGGTTCACCAATCGTCATCCATGCG |
| 0_known | D19 | Tb927.11.7040_for | GGGGACAAGTTTGTACAAAAAAGCAGGCTATGCGGCGGTGTGTTTATTATTGC | Tb927.11.7040_rev | GGGGACCACTTTGTACAAGAAAGCTGGGTTCAGTTGTTCGTCAAAGTTCGCT |
| 0_known | D20 | Tb927.5.3210_for | GGGGACAAGTTTGTACAAAAAAGCAGGCTATGGACGAACCCACTCATAACTCC | Tb927.5.3210_rev | GGGGACCACTTTGTACAAGAAAGCTGGGTTCACGCCATGCACCAAAGAC |
| 0_known | D22 | Tb927.8.5120_for | GGGGACAAGTTTGTACAAAAAAGCAGGCTATGCCACCAAAGGAGCGTG | Tb927.8.5120_rev | GGGGACCACTTTGTACAAGAAAGCTGGGTTTAGTCCTTTAATGTCTCGAGGTATGCG |
| 0_known | D23 | Tb927.6.4140_for | GGGGACAAGTTTGTACAAAAAAGCAGGCTATGCCAAACCTCGTTGTCACTG | Tb927.6.4140_rev | GGGGACCACTTTGTACAAGAAAGCTGGGTCTAAGAGCGAGTGAAATAAAACTTAATGCA |
| 0_known | D24 | Tb927.7.4060_for | GGGGACAAGTTTGTACAAAAAAGCAGGCTATGTCCGATATCACGTACGAGAACG | Tb927.7.4060_rev | GGGGACCACTTTGTACAAGAAAGCTGGGTTCATGACTGAGGAATAGGATTGGCT |
| 0_known | E1 | Tb927.11.7740_for | GGGGACAAGTTTGTACAAAAAAGCAGGCTATGGTGAACGAGTACGATGAAGC | Tb927.11.7740_rev | GGGGACCACTTTGTACAAGAAAGCTGGGTCTAAATAAACATAGTTTCTGTAGAGAAAGT |
| 0_known | E2 | Tb927.4.890_for | GGGGACAAGTTTGTACAAAAAAGCAGGCTATGAACACGGAGGGGCTCC | Tb927.4.890_rev | GGGGACCACTTTGTACAAGAAAGCTGGGTTTACTTCTTTGGCTTCTTACGGGAAC |
| 0_known | E3 | Tb927.8.690_for | GGGGACAAGTTTGTACAAAAAAGCAGGCTATGTCCGAAAAGCTGAGAGCAG | Tb927.8.690_rev | GGGGACCACTTTGTACAAGAAAGCTGGGTTCACGCGAGCCTCTTGATAATG |
| 0_known | E4 | Tb927.4.4320_for | GGGGACAAGTTTGTACAAAAAAGCAGGCTATGTTTTCTGTGTGTTACGTGACGA | Tb927.4.4320_rev | GGGGACCACTTTGTACAAGAAAGCTGGGTTTAGCGGCAACTGCAACCG |
| 0_known | E5 | Tb927.10.12090_for | GGGGACAAGTTTGTACAAAAAAGCAGGCTATGCCACCGCGGGCTCGA | Tb927.10.12090_rev | GGGGACCACTTTGTACAAGAAAGCTGGGTTCACCGTTGAACACGTTGTGG |
| 0_known | E6 | Tb927.11.13140_for | GGGGACAAGTTTGTACAAAAAAGCAGGCTATGCTGCGTCGCGCTGGC | Tb927.11.13140_rev | GGGGACCACTTTGTACAAGAAAGCTGGGTCTACCAGCCGCGATGGTAGA |
| 0_known | E8 | Tb927.8.6950_for | GGGGACAAGTTTGTACAAAAAAGCAGGCTATGGAATTTAACGCATCAACTACCAACG | Tb927.8.6950_rev | GGGGACCACTTTGTACAAGAAAGCTGGGTTCAAGATCTGATCTCTTGCACGACA |
| 0_known | E10 | Tb927.11.3660_for | GGGGACAAGTTTGTACAAAAAAGCAGGCTATGGCAGATAACCACGAAGAACC | Tb927.11.3660_rev | GGGGACCACTTTGTACAAGAAAGCTGGGTCTAATCCAGTTTAACGAAGAAGACAGTCA |
| 0_known | E12 | Tb927.7.5900_for | GGGGACAAGTTTGTACAAAAAAGCAGGCTATGAAGTATAACTTCAAGGATTCACATTCT | Tb927.7.5900_rev | GGGGACCACTTTGTACAAGAAAGCTGGGTTTATCCAAATGCCGCCTCACC |
| 0_known | E14 | Tb927.10.12100_for | GGGGACAAGTTTGTACAAAAAAGCAGGCTATGCCACCGCGGGCTCGA | Tb927.10.12100_rev | GGGGACCACTTTGTACAAGAAAGCTGGGTTCAACGTTGCACACGTTGTGG |
| 0_known | E17 | Tb927.6.4810_for | GGGGACAAGTTTGTACAAAAAAGCAGGCTATGGTTCAACGCATCCGCG | Tb927.6.4810_rev | GGGGACCACTTTGTACAAGAAAGCTGGGTTCACGAGCCGGCAGCGGG |
| 0_known | E18 | Tb927.11.4450_for | GGGGACAAGTTTGTACAAAAAAGCAGGCTATGTCAGAAAAAGGTGGTGACGAC | Tb927.11.4450_rev | GGGGACCACTTTGTACAAGAAAGCTGGGTCTACTCCTTTTTATCAGCGTTTTCCAAC |
| 0_known | E19 | Tb927.3.3480_for | GGGGACAAGTTTGTACAAAAAAGCAGGCTATGGGTGAGCCAAAACAGACG | Tb927.3.3480_rev | GGGGACCACTTTGTACAAGAAAGCTGGGTCTACCCCGCTTGCGTGTC |
| 0_known | E20 | Tb927.10.7470_for | GGGGACAAGTTTGTACAAAAAAGCAGGCTATGTCCGATGAGCGAATTCTCCT | Tb927.10.7470_rev | GGGGACCACTTTGTACAAGAAAGCTGGGTTCAATCCTCGGCCTTGACCTC |
| 0_known | E21 | Tb927.7.1470_for | GGGGACAAGTTTGTACAAAAAAGCAGGCTATGATGCGCCGCCTTGCC | Tb927.7.1470_rev | GGGGACCACTTTGTACAAGAAAGCTGGGTTTACGAGAAGAGCATCAAGAACGC |
| 0_known | E22 | Tb927.6.4340_for | GGGGACAAGTTTGTACAAAAAAGCAGGCTATGTCTGTCAAATCCGAGCGGC | Tb927.6.4340_rev | GGGGACCACTTTGTACAAGAAAGCTGGGTCTATCGGAGGGGAGTGTCCAAA |
| 0_known | E23 | Tb927.10.13110_for | GGGGACAAGTTTGTACAAAAAAGCAGGCTATGGCTGCAATTGGAGGCG | Tb927.10.13110_rev | GGGGACCACTTTGTACAAGAAAGCTGGGTCTACGCACTAACACCATTTGCATTG |
| 0_known | E24 | Tb927.10.1570_for | GGGGACAAGTTTGTACAAAAAAGCAGGCTATGATGCGCCGTCTCGCT | Tb927.10.1570_rev | GGGGACCACTTTGTACAAGAAAGCTGGGTCTACGAGAAGAGCATCAGAAACGC |
| 0_known | F1 | Tb927.11.5280_for | GGGGACAAGTTTGTACAAAAAAGCAGGCTATGATGCGCCGCCTTGCC | Tb927.11.5280_rev | GGGGACCACTTTGTACAAGAAAGCTGGGTCTACGAGAAGAGCATCAGAAACGC |
| 0_known | F2 | Tb927.11.13650_for | GGGGACAAGTTTGTACAAAAAAGCAGGCTATGGAGGGGTTTGTATCGCTGA | Tb927.11.13650_rev | GGGGACCACTTTGTACAAGAAAGCTGGGTTCAGCTTGAGAATAGAAAGTATATAATAAT |
| 0_known | F3 | Tb927.10.14110_for | GGGGACAAGTTTGTACAAAAAAGCAGGCTATGGCCTCGTTCGGCTCT | Tb927.10.14110_rev | GGGGACCACTTTGTACAAGAAAGCTGGGTTTACGACACTGCGCTTTCACG |
| 0_known | F4 | Tb927.3.3720_for | GGGGACAAGTTTGTACAAAAAAGCAGGCTATGCAGCCGCAGAGCTCC | Tb927.3.3720_rev | GGGGACCACTTTGTACAAGAAAGCTGGGTTCACCGGGACTTCAGCATTATATATATCA |
| 0_known | F5 | Tb927.7.5910_for | GGGGACAAGTTTGTACAAAAAAGCAGGCTATGAGTAAAAAAGATAGCAAGTACAAAATG | Tb927.7.5910_rev | GGGGACCACTTTGTACAAGAAAGCTGGGTTTAGCATCCAAATGTCGCCTCAC |
| 0_known | F6 | Tb927.7.4070_for | GGGGACAAGTTTGTACAAAAAAGCAGGCTATGTCCGATATCACGTACGAGAACG | Tb927.7.4070_rev | GGGGACCACTTTGTACAAGAAAGCTGGGTTTAGGTGGGAACGGATTTCCACT |
| 0_known | F7 | Tb927.11.11975_for | GGGGACAAGTTTGTACAAAAAAGCAGGCTATGGACCGGAGACTCATCGAAA | Tb927.11.11975_rev | GGGGACCACTTTGTACAAGAAAGCTGGGTTTACATATCCACAGATGGCTCCTTAGAG |
| 0_known | F8 | Tb927.8.6450_for | GGGGACAAGTTTGTACAAAAAAGCAGGCTATGTCCCACAACCTATTTACTGAGGA | Tb927.8.6450_rev | GGGGACCACTTTGTACAAGAAAGCTGGGTTTATGCGGTGGCCTCGACG |
| 0_known | F11 | Tb927.3.3100_for | GGGGACAAGTTTGTACAAAAAAGCAGGCTATGGGCAAAGAAAACATGAAGGGC | Tb927.3.3100_rev | GGGGACCACTTTGTACAAGAAAGCTGGGTTCACTGCTTATCCTCCACAAAAATAATGT |
| 0_known | F12 | Tb927.6.480_for | GGGGACAAGTTTGTACAAAAAAGCAGGCTATGGCACCTCGTTCCCTTTATCT | Tb927.6.480_rev | GGGGACCACTTTGTACAAGAAAGCTGGGTTTAGAATGCGGCAACGAGACC |
| 0_known | F13 | Tb927.1.2230_for | GGGGACAAGTTTGTACAAAAAAGCAGGCTATGGGTTGTGGTGGCTCAAAG | Tb927.1.2230_rev | GGGGACCACTTTGTACAAGAAAGCTGGGTCTATTGCCTGTCTCCATCAGTAATGG |
| 0_known | F14 | Tb927.10.8320_for | GGGGACAAGTTTGTACAAAAAAGCAGGCTATGTTCTCCTGTGCGCTGAGA | Tb927.10.8320_rev | GGGGACCACTTTGTACAAGAAAGCTGGGTCTAGTGGTGACCCAGATCCACA |
| 0_known | F15 | Tb927.8.4280_for | GGGGACAAGTTTGTACAAAAAAGCAGGCTATGTCCTTTCAAGAGATTGGAACGG | Tb927.8.4280_rev | GGGGACCACTTTGTACAAGAAAGCTGGGTTTAGTGGCCACCGCCGCC |
| 0_known | F16 | Tb927.9.12820_for | GGGGACAAGTTTGTACAAAAAAGCAGGCTATGCAAACAGATTTTGGAGATGCCA | Tb927.9.12820_rev | GGGGACCACTTTGTACAAGAAAGCTGGGTTTAATATTGATAGAGGCCGAATACCGTCA |
| 0_known | F17 | Tb927.10.6670_for | GGGGACAAGTTTGTACAAAAAAGCAGGCTATGATGGAGGACGATGAGGGTTT | Tb927.10.6670_rev | GGGGACCACTTTGTACAAGAAAGCTGGGTTTACTCCTTTGCATCATTCGCATTTAGC |
| 0_known | F18 | Tb927.9.10280_for | GGGGACAAGTTTGTACAAAAAAGCAGGCTATGATCTTCCCCAGTGCAGCA | Tb927.9.10280_rev | GGGGACCACTTTGTACAAGAAAGCTGGGTTCACTTGGCTACCCACTCATTAGG |
| 0_known | F19 | Tb927.11.7707_for | GGGGACAAGTTTGTACAAAAAAGCAGGCTATGGTGAACGAGTACCATGAAGC | Tb927.11.7707_rev | GGGGACCACTTTGTACAAGAAAGCTGGGTCTACTCCCAGTACAGCGCATACA |
| 0_known | F20 | Tb927.11.4460_for | GGGGACAAGTTTGTACAAAAAAGCAGGCTATGACGACAGGAAAGAGCGACC | Tb927.11.4460_rev | GGGGACCACTTTGTACAAGAAAGCTGGGTTTACGCTGTAGCCTTCTCCTTCT |
| 0_known | F21 | Tb927.8.5180_for | GGGGACAAGTTTGTACAAAAAAGCAGGCTATGGATGCGTTGACGACCAG | Tb927.8.5180_rev | GGGGACCACTTTGTACAAGAAAGCTGGGTTCAATCCTGAGGTACGGAAGGTT |
| 0_known | F22 | Tb927.10.11300_for | GGGGACAAGTTTGTACAAAAAAGCAGGCTATGTCTCACTACCTTTTGGTCACTTTC | Tb927.10.11300_rev | GGGGACCACTTTGTACAAGAAAGCTGGGTTCAGCTATCCTCAGCAAGAAGTTTATTTAG |
| 0_known | F23 | Tb927.10.8950_for | GGGGACAAGTTTGTACAAAAAAGCAGGCTATGCTTCGTGTTTCTCGATTATCCC | Tb927.10.8950_rev | GGGGACCACTTTGTACAAGAAAGCTGGGTCTACTTTCGCATCTGTTTCTGCTGT |
| 0_known | F24 | Tb927.10.15150_for | GGGGACAAGTTTGTACAAAAAAGCAGGCTATGTTTTTCTGTCCATTCTGTGGAACG | Tb927.10.15150_rev | GGGGACCACTTTGTACAAGAAAGCTGGGTCTAATCCTGCTTCCACTGGAAGC |
| 0_known | G1 | Tb927.10.13650_for | GGGGACAAGTTTGTACAAAAAAGCAGGCTATGGATAGTGAGGAAGGTGAGTTCG | Tb927.10.13650_rev | GGGGACCACTTTGTACAAGAAAGCTGGGTTTATTGATCTCGCTTTTCCCATGCT |
| 0_known | G2 | Tb927.11.13960_for | GGGGACAAGTTTGTACAAAAAAGCAGGCTATGAGTGTTAGGAAGCCTATAACGCC | Tb927.11.13960_rev | GGGGACCACTTTGTACAAGAAAGCTGGGTTTACTTCCCGCGGTTACGCT |
| 0_known | G4 | Tb927.1.4550_for | GGGGACAAGTTTGTACAAAAAAGCAGGCTATGTTTGTTTTTATATGCCGTGACTTTGT | Tb927.1.4550_rev | GGGGACCACTTTGTACAAGAAAGCTGGGTTCATGCAACTGGAAGAGCAGGG |
| 0_known | G6 | Tb927.10.8970_for | GGGGACAAGTTTGTACAAAAAAGCAGGCTATGCTTCGTGTTTCTCGATTATCCC | Tb927.10.8970_rev | GGGGACCACTTTGTACAAGAAAGCTGGGTTCACTTACTTCGCATCTGTTTCTGC |
| 0_known | G7 | Tb927.11.16150_for | GGGGACAAGTTTGTACAAAAAAGCAGGCTATGTCGACAAGTGAGGGCAATG | Tb927.11.16150_rev | GGGGACCACTTTGTACAAGAAAGCTGGGTTTATTGTTTAGACAAAGTTAGCTGTGCAC |
| 0_known | G10 | Tb927.3.1780_for | GGGGACAAGTTTGTACAAAAAAGCAGGCTATGCTTTCCCAATATCTCAGGCG | Tb927.3.1780_rev | GGGGACCACTTTGTACAAGAAAGCTGGGTCTATCCACCTTTAACTGAAGCTGTTGC |
| 0_known | G11 | Tb927.1.2260_for | GGGGACAAGTTTGTACAAAAAAGCAGGCTATGGGTTGCGGTGGCTCA | Tb927.1.2260_rev | GGGGACCACTTTGTACAAGAAAGCTGGGTTTATTTCCTGCCAGATCCCCTGC |
| 0_known | G12 | Tb927.11.12100_for | GGGGACAAGTTTGTACAAAAAAGCAGGCTATGTTCAACCAGTTTTCGAAGATGCC | Tb927.11.12100_rev | GGGGACCACTTTGTACAAGAAAGCTGGGTTTAAGCAGTAATACACGTCGGTGG |
| 0_known | G13 | Tb927.3.720_for | GGGGACAAGTTTGTACAAAAAAGCAGGCTATGCAGGGCTATTTTGCAGTAAATCC | Tb927.3.720_rev | GGGGACCACTTTGTACAAGAAAGCTGGGTTTATGCCATGGGCGGAACATC |
| 0_known | G14 | Tb927.6.3290_for | GGGGACAAGTTTGTACAAAAAAGCAGGCTATGGATGATGATAAACTTGTGATGTTTGAC | Tb927.6.3290_rev | GGGGACCACTTTGTACAAGAAAGCTGGGTCTACTCACGCGAGGCGTG |
| 0_known | G15 | Tb927.1.2160_for | GGGGACAAGTTTGTACAAAAAAGCAGGCTATGGGTTGCATTCAGTCGACAG | Tb927.1.2160_rev | GGGGACCACTTTGTACAAGAAAGCTGGGTTTATAGACGATATTGTTCTTCAGATCCTGC |
| 0_known | G16 | Tb927.10.15580_for | GGGGACAAGTTTGTACAAAAAAGCAGGCTATGGCGGGCACCACCGCT | Tb927.10.15580_rev | GGGGACCACTTTGTACAAGAAAGCTGGGTTTACATTGACGTTTGTCCCAGCTT |
| 0_known | G17 | Tb927.10.4950_for | GGGGACAAGTTTGTACAAAAAAGCAGGCTATGGAAGGTGTAGCTGATATTCCCT | Tb927.10.4950_rev | GGGGACCACTTTGTACAAGAAAGCTGGGTTCATAGGAGTAGCTTTTTCTTTCGATGC |
| 0_known | G18 | Tb927.10.10420_for | GGGGACAAGTTTGTACAAAAAAGCAGGCTATGCTGCGCCGCATTTTTG | Tb927.10.10420_rev | GGGGACCACTTTGTACAAGAAAGCTGGGTTCAGAGAAGATCCTTCTCCTTTAGAAGC |
| 0_known | G19 | Tb927.11.200_for | GGGGACAAGTTTGTACAAAAAAGCAGGCTATGATACAGTCGGATATTGTTTCGTATGTT | Tb927.11.200_rev | GGGGACCACTTTGTACAAGAAAGCTGGGTTCAGTGCTGAACGTCCAGCA |
| 0_known | G20 | Tb927.9.15420_for | GGGGACAAGTTTGTACAAAAAAGCAGGCTATGGTCAAGCCGTTTGTGCC | Tb927.9.15420_rev | GGGGACCACTTTGTACAAGAAAGCTGGGTTTACACCTGGACCTTCCTCAACTT |
| 0_known | G21 | Tb927.11.16200_for | GGGGACAAGTTTGTACAAAAAAGCAGGCTATGGATCGCAAGCTTGTTGAGATG | Tb927.11.16200_rev | GGGGACCACTTTGTACAAGAAAGCTGGGTTCATGATGTTTGAGCGGGCC |
| 0_known | G22 | Tb927.11.2200_for | GGGGACAAGTTTGTACAAAAAAGCAGGCTATGTCCACCTCACATGGCATGC | Tb927.11.2200_rev | GGGGACCACTTTGTACAAGAAAGCTGGGTCTATTCAACATCCAAACCTTTGCTCTG |
| 0_known | G23 | Tb927.11.7280_for | GGGGACAAGTTTGTACAAAAAAGCAGGCTATGGAGTCCACTTTGACGCAGT | Tb927.11.7280_rev | GGGGACCACTTTGTACAAGAAAGCTGGGTTTAATTGTCTTGATGGTCGCTGCT |
| 0_known | G24 | Tb927.1.2450_for | GGGGACAAGTTTGTACAAAAAAGCAGGCTATGTCGAGGACCAAGGAAACCG | Tb927.1.2450_rev | GGGGACCACTTTGTACAAGAAAGCTGGGTTTATGCACGTTCACCGCGT |
| 0_known | H2 | Tb927.11.6640_for | GGGGACAAGTTTGTACAAAAAAGCAGGCTATGACAACGTCACAACTCCCAAA | Tb927.11.6640_rev | GGGGACCACTTTGTACAAGAAAGCTGGGTCTAGGTAGCAAAAATGTAGCACAATAAAAG |
| 0_known | H4 | Tb927.1.3820_for | GGGGACAAGTTTGTACAAAAAAGCAGGCTATGGCGACTGGTCGACTAAAAAAC | Tb927.1.3820_rev | GGGGACCACTTTGTACAAGAAAGCTGGGTTTACTGGCAGCCGAGTGCT |
| 0_known | H9 | Tb927.7.2830_for | GGGGACAAGTTTGTACAAAAAAGCAGGCTATGGCAACACCCAAACAGGC | Tb927.7.2830_rev | GGGGACCACTTTGTACAAGAAAGCTGGGTCTAGACGCTTGGCGTCGC |

| | | | | | |
|---|---|---|---|---|---|
| 0_known | H14 | Tb927.7.3520_for | GGGGACAAGTTTGTACAAAAAAGCAGGCTATGTCTGCAACCACAAAACAAGC | Tb927.7.3520_rev | GGGGACCACTTTGTACAAGAAAGCTGGGTCTAATGCAGAGCTTTCGACCTCTT |
| 0_known | H16 | Tb927.7.3320_for | GGGGACAAGTTTGTACAAAAAAGCAGGCTATGCCTTCACACTACCGGTACC | Tb927.7.3320_rev | GGGGACCACTTTGTACAAGAAAGCTGGGTTCAACCAAATGTATTTTCCACACCATACG |
| 0_known | H18 | Tb927.7.2820_for | GGGGACAAGTTTGTACAAAAAAGCAGGCTATGGCAACACCCAAGCAGG | Tb927.7.2820_rev | GGGGACCACTTTGTACAAGAAAGCTGGGTCTAGACGCTTGGCGTCGC |
| 0_known | H19 | Tb927.11.8890_for | GGGGACAAGTTTGTACAAAAAAGCAGGCTATGAGTTTTCACCCTCGGGATGC | Tb927.11.8890_rev | GGGGACCACTTTGTACAAGAAAGCTGGGTCTAGTGCCCCTCAGGGTCC |
| 0_known | I1 | Tb927.5.580_for | GGGGACAAGTTTGTACAAAAAAGCAGGCTATGCAGCAGGTTCCACCCG | Tb927.5.580_rev | GGGGACCACTTTGTACAAGAAAGCTGGGTTTAAGATTGCTGCTGACGTCTTTGG |
| 0_known | I2 | Tb927.7.5760_for | GGGGACAAGTTTGTACAAAAAAGCAGGCTATGTCGGAATACGCAGATTGCAC | Tb927.7.5760_rev | GGGGACCACTTTGTACAAGAAAGCTGGGTTTACTCCTTTTCACTTAACCAGCGATAAT |
| 0_known | I3 | Tb927.7.5790_for | GGGGACAAGTTTGTACAAAAAAGCAGGCTATGAGGAACTCATTTGCACTTCTGC | Tb927.7.5790_rev | GGGGACCACTTTGTACAAGAAAGCTGGGTTCACATGTTGCTTTTAACAAATTCCTTCA |
| 0_known | I4 | Tb927.3.5180_for | GGGGACAAGTTTGTACAAAAAAGCAGGCTATGGCCATGTCTGGTGTTTCTG | Tb927.3.5180_rev | GGGGACCACTTTGTACAAGAAAGCTGGGTTTACCGGTTCGACTTCACTTTGC |
| 0_known | I5 | Tb927.11.3310_for | GGGGACAAGTTTGTACAAAAAAGCAGGCTATGGTTGAGGTACCGCGTAATTTC | Tb927.11.3310_rev | GGGGACCACTTTGTACAAGAAAGCTGGGTTCAATACGTTGAACCCTCCTCCG |
| 0_known | I6 | Tb927.9.9840_for | GGGGACAAGTTTGTACAAAAAAGCAGGCTATGCGTCGCGCACTGTTT | Tb927.9.9840_rev | GGGGACCACTTTGTACAAGAAAGCTGGGTCTATTCTATAAATTTGTTGTATTCATCGGC |
| 0_known | I7 | Tb927.11.1495_for | GGGGACAAGTTTGTACAAAAAAGCAGGCTATGTCCTTTTTCCAGTTGGATTGGG | Tb927.11.1495_rev | GGGGACCACTTTGTACAAGAAAGCTGGGTTTAGCTGTTTCCCCCAGATACGT |
| 0_known | I8 | Tb927.11.14950_for | GGGGACAAGTTTGTACAAAAAAGCAGGCTATGGCCTTCAACCAACGCTTT | Tb927.11.14950_rev | GGGGACCACTTTGTACAAGAAAGCTGGGTCTACTGCTGCAGATGGTTATTATCTAACC |
| 0_known | I9 | Tb927.9.11380_for | GGGGACAAGTTTGTACAAAAAAGCAGGCTATGGGAAAGGACAAGGCCAACG | Tb927.9.11380_rev | GGGGACCACTTTGTACAAGAAAGCTGGGTCTACACGATAGCTGGAGCGTGG |
| 0_known | I10 | Tb927.10.15350_for | GGGGACAAGTTTGTACAAAAAAGCAGGCTATGGCGCAAATGAAGAAAATAACACC | Tb927.10.15350_rev | GGGGACCACTTTGTACAAGAAAGCTGGGTTTAGTTACGCTCGCCTCGGA |
| 0_known | I11 | Tb927.9.11410_for | GGGGACAAGTTTGTACAAAAAAGCAGGCTATGGGAAAGGACAAGGCCAACG | Tb927.9.11410_rev | GGGGACCACTTTGTACAAGAAAGCTGGGTCTAGACGATGGCTGGAGCGT |
| 0_known | I12 | Tb927.1.2150_for | GGGGACAAGTTTGTACAAAAAAGCAGGCTATGGGTCTATTTCTGAGCAAACAAGG | Tb927.1.2150_rev | GGGGACCACTTTGTACAAGAAAGCTGGGTTTAAAACGATGTCGCCTTGAAGGG |
| 0_known | I13 | Tb927.10.6070_for | GGGGACAAGTTTGTACAAAAAAGCAGGCTATGGCTGATAACATGCAGATGTCAA | Tb927.10.6070_rev | GGGGACCACTTTGTACAAGAAAGCTGGGTTTATTTCACAGGGCAGGCACG |
| 0_known | I14 | Tb927.10.10260_for | GGGGACAAGTTTGTACAAAAAAGCAGGCTATGGCACCTCGTTCCCTTTATCT | Tb927.10.10260_rev | GGGGACCACTTTGTACAAGAAAGCTGGGTTTAGAATGCGGCAACGAGAGC |
| 0_known | I15 | Tb927.9.11230_for | GGGGACAAGTTTGTACAAAAAAGCAGGCTATGTCACGAGCATCTATAATGAAGGAAG | Tb927.9.11230_rev | GGGGACCACTTTGTACAAGAAAGCTGGGTCTACGCACTCTCATGCTCTGTAAAG |
| 0_known | I16 | Tb927.3.3330_for | GGGGACAAGTTTGTACAAAAAAGCAGGCTATGTGGGACCCTTTTCGTGATATTG | Tb927.3.3330_rev | GGGGACCACTTTGTACAAGAAAGCTGGGTTTACTGAATCGTTACGGATGTTGCA |
| 0_known | I17 | Tb927.7.4480_for | GGGGACAAGTTTGTACAAAAAAGCAGGCTATGTCCAACTGCATTTTCTGTAAGATTG | Tb927.7.4480_rev | GGGGACCACTTTGTACAAGAAAGCTGGGTTCACAACTTCTCGAGAGCCTCA |
| 0_known | I18 | Tb927.11.7900_for | GGGGACAAGTTTGTACAAAAAAGCAGGCTATGATTCGCGCTTCCATAGTTAAGC | Tb927.11.7900_rev | GGGGACCACTTTGTACAAGAAAGCTGGGTCTAAAAGTCATCGCTGAAGCTCTGG |
| 0_known | I19 | Tb927.11.1000_for | GGGGACAAGTTTGTACAAAAAAGCAGGCTATGGATTTATTAGGAAGCCAAATCGCT | Tb927.11.1000_rev | GGGGACCACTTTGTACAAGAAAGCTGGGTCTACAGCTGCATACTCGCCTCT |
| 0_known | I20 | Tb927.10.12420_for | GGGGACAAGTTTGTACAAAAAAGCAGGCTATGGACTCTTTATACAGCTGGAATGATTC | Tb927.10.12420_rev | GGGGACCACTTTGTACAAGAAAGCTGGGTTTACACATTTCGGTTACCCCGGG |
| 0_known | I21 | Tb927.11.7350_for | GGGGACAAGTTTGTACAAAAAAGCAGGCTATGCCGCCAACTAAAGGAGGA | Tb927.11.7350_rev | GGGGACCACTTTGTACAAGAAAGCTGGGTCTACAGTACCTTGGAGTACGCCT |
| 0_known | I22 | Tb927.3.3760_for | GGGGACAAGTTTGTACAAAAAAGCAGGCTATGTCTGGCCTCACTAAGTATCTTCC | Tb927.3.3760_rev | GGGGACCACTTTGTACAAGAAAGCTGGGTTCAGTTGGGCCACGGAAAGT |
| 0_known | I23 | Tb927.3.3780_for | GGGGACAAGTTTGTACAAAAAAGCAGGCTATGTCTGGCCTCGCTAAGTATCT | Tb927.3.3780_rev | GGGGACCACTTTGTACAAGAAAGCTGGGTTCAGTTGGGCCACGGAAAGT |
| 0_known | I24 | Tb927.4.3510_for | GGGGACAAGTTTGTACAAAAAAGCAGGCTATGTCTGAGCGTGATGAGGATGA | Tb927.4.3510_rev | GGGGACCACTTTGTACAAGAAAGCTGGGTTCAGATGTTTGTGTAGCGCTCG |
| 0_known | J2 | Tb927.1.1210_for | GGGGACAAGTTTGTACAAAAAAGCAGGCTATGGGTTACACACGGGAGCG | Tb927.1.1210_rev | GGGGACCACTTTGTACAAGAAAGCTGGGTTTACTTCATTCTTGCACGCCCC |
| 0_known | J3 | Tb927.11.12680_for | GGGGACAAGTTTGTACAAAAAAGCAGGCTATGGCCGCAACAAACTCACC | Tb927.11.12680_rev | GGGGACCACTTTGTACAAGAAAGCTGGGTTCATACCAGAACACCCCTGCTG |
| 0_known | J4 | Tb927.10.8280_for | GGGGACAAGTTTGTACAAAAAAGCAGGCTATGGAGGCGCTTCGCGTT | Tb927.10.8280_rev | GGGGACCACTTTGTACAAGAAAGCTGGGTCTATCCGTCAAACATCCTTTTTATTTTGGT |
| 0_known | J5 | Tb927.11.11100_for | GGGGACAAGTTTGTACAAAAAAGCAGGCTATGACCGATATAGTGGACCTGGAAC | Tb927.11.11100_rev | GGGGACCACTTTGTACAAGAAAGCTGGGTCTACAAATCCAGCCTCATTTGTCTCT |
| 0_known | J6 | Tb927.9.6870_for | GGGGACAAGTTTGTACAAAAAAGCAGGCTATGTCTCGAGTGTACGTTGGTAACA | Tb927.9.6870_rev | GGGGACCACTTTGTACAAGAAAGCTGGGTTCACGGGGAACAAGGCCC |
| 0_known | J7 | Tb927.3.860_for | GGGGACAAGTTTGTACAAAAAAGCAGGCTATGCAGCGGTCTCAACTTGC | Tb927.3.860_rev | GGGGACCACTTTGTACAAGAAAGCTGGGTTTACTTGGCCATTGGGTTCTGAGA |
| 0_known | J8 | Tb927.5.1000_for | GGGGACAAGTTTGTACAAAAAAGCAGGCTATGGCTCTTCGTCGCATTCAAA | Tb927.5.1000_rev | GGGGACCACTTTGTACAAGAAAGCTGGGTTCACATAGCATACTGCCGCGT |
| 0_known | J9 | Tb927.11.13040_for | GGGGACAAGTTTGTACAAAAAAGCAGGCTATGGCCGATCAACTCTCCAACG | Tb927.11.13040_rev | GGGGACCACTTTGTACAAGAAAGCTGGGTCTATTTGCTCATCATCATTTTCACGAACT |
| 0_known | J11 | Tb927.11.7930_for | GGGGACAAGTTTGTACAAAAAAGCAGGCTATGGCTTCCTCGAAAGTAATATTGGAAG | Tb927.11.7930_rev | GGGGACCACTTTGTACAAGAAAGCTGGGTCTAACTTCCGGGGCGTTTCATC |
| 0_known | J13 | Tb927.7.3410_for | GGGGACAAGTTTGTACAAAAAAGCAGGCTATGGCGGCGCTTACTGATG | Tb927.7.3410_rev | GGGGACCACTTTGTACAAGAAAGCTGGGTCTATTTGCCACGCATCTGCATCA |
| 0_known | J15 | Tb927.11.13780_for | GGGGACAAGTTTGTACAAAAAAGCAGGCTATGTCGTGGCAGTCCTACGT | Tb927.11.13780_rev | GGGGACCACTTTGTACAAGAAAGCTGGGTTCAGTAGTCCAAGCTAATCAAATACTCCG |
| 0_known | J16 | Tb927.9.8000_for | GGGGACAAGTTTGTACAAAAAAGCAGGCTATGACGCTGACACCTCGTATTATTAAAG | Tb927.9.8000_rev | GGGGACCACTTTGTACAAGAAAGCTGGGTTCACGCCTTGTTGGCGTATTT |
| 0_known | J17 | Tb927.11.10420_for | GGGGACAAGTTTGTACAAAAAAGCAGGCTATGAGTGCACCAATGGCTCG | Tb927.11.10420_rev | GGGGACCACTTTGTACAAGAAAGCTGGGTTTATGAGTTGAGCTGCCACTCCG |
| 0_known | J18 | Tb927.11.3090_for | GGGGACAAGTTTGTACAAAAAAGCAGGCTATGGAGGAAAGGGAGGAGGGA | Tb927.11.3090_rev | GGGGACCACTTTGTACAAGAAAGCTGGGTTTACGAATTTTCGCTGCAGATACGA |
| 0_known | J19 | Tb927.10.7190_for | GGGGACAAGTTTGTACAAAAAAGCAGGCTATGAGAGTTGTCGTTCAGCGTG | Tb927.10.7190_rev | GGGGACCACTTTGTACAAGAAAGCTGGGTTCACTTATTTCGGCTGTCCAGC |
| 0_known | J20 | Tb927.11.13290_for | GGGGACAAGTTTGTACAAAAAAGCAGGCTATGACAACACTTCTCGACCCCC | Tb927.11.13290_rev | GGGGACCACTTTGTACAAGAAAGCTGGGTTTAGCGTTGAGCCAACCCCA |
| 0_known | J21 | Tb927.11.16430_for | GGGGACAAGTTTGTACAAAAAAGCAGGCTATGAGTTCTTCATGGCTGTCCTTCG | Tb927.11.16430_rev | GGGGACCACTTTGTACAAGAAAGCTGGGTTCATAGCTGCTCATCTGTCGCA |
| 0_known | J22 | Tb927.5.1360_for | GGGGACAAGTTTGTACAAAAAAGCAGGCTATGAGAAAGATATATATTGCGGGTCCTGC | Tb927.5.1360_rev | GGGGACCACTTTGTACAAGAAAGCTGGGTTTACTTACTGGGGAAATTTGCCAGG |
| 0_known | J23 | Tb927.6.4530_for | GGGGACAAGTTTGTACAAAAAAGCAGGCTATGTCCTCTTCTGCTCTGTTGTTCG | Tb927.6.4530_rev | GGGGACCACTTTGTACAAGAAAGCTGGGTCTAATACGAATGCAACCGCCCC |
| 0_known | J24 | Tb927.11.16130_for | GGGGACAAGTTTGTACAAAAAAGCAGGCTATGCCCAGCGAGCGGACC | Tb927.11.16130_rev | GGGGACCACTTTGTACAAGAAAGCTGGGTTTATGCCCTTTCATAAATTTGCTTGACG |
| 0_known | K1 | Tb927.5.1880_for | GGGGACAAGTTTGTACAAAAAAGCAGGCTATGACAGACCGACCTCCGAC | Tb927.5.1880_rev | GGGGACCACTTTGTACAAGAAAGCTGGGTCTAACCCGCCCTCTCCTCG |
| 0_known | K2 | Tb927.11.7940_for | GGGGACAAGTTTGTACAAAAAAGCAGGCTATGTCCAGCGCATTTGACGAT | Tb927.11.7940_rev | GGGGACCACTTTGTACAAGAAAGCTGGGTTCAGCGAAAAATCTCTGGGGTCA |
| 0_known | K3 | Tb927.8.6090_for | GGGGACAAGTTTGTACAAAAAAGCAGGCTATGTCTACAAACGGTAATTCCGTAACA | Tb927.8.6090_rev | GGGGACCACTTTGTACAAGAAAGCTGGGTTCAACACGAGGCGTTTGCC |
| 0_known | K4 | Tb927.11.8970_for | GGGGACAAGTTTGTACAAAAAAGCAGGCTATGACGCGCAAGGTGGCT | Tb927.11.8970_rev | GGGGACCACTTTGTACAAGAAAGCTGGGTCTAACAACCATTCGTGTTAGATTCGGC |
| 0_known | K5 | Tb927.8.2560_for | GGGGACAAGTTTGTACAAAAAAGCAGGCTATGACGGAGTTGACACGCCT | Tb927.8.2560_rev | GGGGACCACTTTGTACAAGAAAGCTGGGTCTAACGGCGTACAGCAGCTG |
| 0_known | K6 | Tb927.10.3420_for | GGGGACAAGTTTGTACAAAAAAGCAGGCTATGAACTTAAATTGTGAAGGACGTGACG | Tb927.10.3420_rev | GGGGACCACTTTGTACAAGAAAGCTGGGTTTATAAATATATATAAATATTTTTTTTTAA |
| 0_known | K7 | Tb927.11.15920_for | GGGGACAAGTTTGTACAAAAAAGCAGGCTATGAATGGCGGCGCCATT | Tb927.11.15920_rev | GGGGACCACTTTGTACAAGAAAGCTGGGTTCACTCATCGCACAACTGTGAAA |
| 0_known | K8 | Tb927.11.3680_for | GGGGACAAGTTTGTACAAAAAAGCAGGCTATGGCAAATGCTGTTAGCGAAAAG | Tb927.11.3680_rev | GGGGACCACTTTGTACAAGAAAGCTGGGTCTACATAACACCGCCGCCC |
| 0_known | K9 | Tb927.11.3380_for | GGGGACAAGTTTGTACAAAAAAGCAGGCTATGGCAGACGGTAAGGCAGA | Tb927.11.3380_rev | GGGGACCACTTTGTACAAGAAAGCTGGGTCTATGTGGTGGTTTTGGAAAGGGC |
| 0_known | K10 | Tb927.9.6130_for | GGGGACAAGTTTGTACAAAAAAGCAGGCTATGTCATTTCCAAAGGAATGGACCG | Tb927.9.6130_rev | GGGGACCACTTTGTACAAGAAAGCTGGGTTTAAGAGATGCCCGCCAGTTGC |
| 0_known | K11 | Tb927.11.15000_for | GGGGACAAGTTTGTACAAAAAAGCAGGCTATGGTCCGGCGGAATAATAAGTCG | Tb927.11.15000_rev | GGGGACCACTTTGTACAAGAAAGCTGGGTTTACTCTCCACGAGCACGCT |
| 0_known | K12 | Tb927.4.4620_for | GGGGACAAGTTTGTACAAAAAAGCAGGCTATGATCCGTCGCACGGCA | Tb927.4.4620_rev | GGGGACCACTTTGTACAAGAAAGCTGGGTTCACAGATCTTCAGTCGTCATTGTTAG |
| 0_known | K13 | Tb927.10.280_for | GGGGACAAGTTTGTACAAAAAAGCAGGCTATGCCCTTTGTCGATCACAATAAATACA | Tb927.10.280_rev | GGGGACCACTTTGTACAAGAAAGCTGGGTCTACTCATCATATACGGGTTTCTTCCCC |
| 0_known | K14 | Tb927.11.12360_for | GGGGACAAGTTTGTACAAAAAAGCAGGCTATGCCGTACCCAGACGTCC | Tb927.11.12360_rev | GGGGACCACTTTGTACAAGAAAGCTGGGTTCAACCCTGTGGCAGTGTTGG |
| 0_known | K15 | Tb927.10.14210_for | GGGGACAAGTTTGTACAAAAAAGCAGGCTATGAGGGCGCAGTCAGACC | Tb927.10.14210_rev | GGGGACCACTTTGTACAAGAAAGCTGGGTTTAGACTGACCGGAAAGAATCGGC |
| 0_known | K16 | Tb927.3.4000_for | GGGGACAAGTTTGTACAAAAAAGCAGGCTATGATAAGATATCTTTTGCTTATAAGTCGC | Tb927.3.4000_rev | GGGGACCACTTTGTACAAGAAAGCTGGGTTTAGCGTGCCTTTGAGGAATTTGT |
| 0_known | K17 | Tb927.7.1170_for | GGGGACAAGTTTGTACAAAAAAGCAGGCTATGATTACAGAGGGCGCCGC | Tb927.7.1170_rev | GGGGACCACTTTGTACAAGAAAGCTGGGTCTACTCACGCTCCCGTTTTCC |
| 0_known | K18 | Tb927.10.14810_for | GGGGACAAGTTTGTACAAAAAAGCAGGCTATGGCTCGCGGTCAAATCATG | Tb927.10.14810_rev | GGGGACCACTTTGTACAAGAAAGCTGGGTTCACTTCTTCGGCTGGGGTT |
| 0_known | K19 | Tb927.8.920_for | GGGGACAAGTTTGTACAAAAAAGCAGGCTATGCCGTGCCCCATCAGC | Tb927.8.920_rev | GGGGACCACTTTGTACAAGAAAGCTGGGTCTACTCCCCATCATTGAAAGGTGGT |
| 0_known | K20 | Tb927.1.1630_for | GGGGACAAGTTTGTACAAAAAAGCAGGCTATGGGTGGAACAAAGAAAAAGGCA | Tb927.1.1630_rev | GGGGACCACTTTGTACAAGAAAGCTGGGTTCAAAACAGCGGCTCATCGTAC |
| 0_known | K21 | Tb927.8.1440_for | GGGGACAAGTTTGTACAAAAAAGCAGGCTATGTTAAAGGTAGGCCAGAGACACA | Tb927.8.1440_rev | GGGGACCACTTTGTACAAGAAAGCTGGGTCTACTCGACACGGGGCACA |
| 0_known | K22 | Tb927.3.5270_for | GGGGACAAGTTTGTACAAAAAAGCAGGCTATGAGTAGTGAGAGATATGTGGATGGTG | Tb927.3.5270_rev | GGGGACCACTTTGTACAAGAAAGCTGGGTTCATAAAGAGTCGAGCATTTCTAAAACCT |
| 0_known | K23 | Tb927.4.2710_for | GGGGACAAGTTTGTACAAAAAAGCAGGCTATGACGGCAATTGCGATGAAACG | Tb927.4.2710_rev | GGGGACCACTTTGTACAAGAAAGCTGGGTCTAAAACGGACATTGAGCTCGCT |
| 0_known | K24 | Tb927.10.8300_for | GGGGACAAGTTTGTACAAAAAAGCAGGCTATGGAGGCGCTTCGCGTT | Tb927.10.8300_rev | GGGGACCACTTTGTACAAGAAAGCTGGGTTTAAAGGCGTCTAAGAAAAAGCACAGT |
| 0_known | L1 | Tb927.11.9790_for | GGGGACAAGTTTGTACAAAAAAGCAGGCTATGGCGGACCTCTTAACGGT | Tb927.11.9790_rev | GGGGACCACTTTGTACAAGAAAGCTGGGTTTAATCGGTTGAAAACGCTTCCTTCT |
| 0_known | L2 | Tb927.10.5120_for | GGGGACAAGTTTGTACAAAAAAGCAGGCTATGCTAGCTCTTACATCACGCCG | Tb927.10.5120_rev | GGGGACCACTTTGTACAAGAAAGCTGGGTTTACGATGGCACACCACTTTGG |
| 0_known | L3 | Tb927.11.6010_for | GGGGACAAGTTTGTACAAAAAAGCAGGCTATGCGCGGTGCCGATCAG | Tb927.11.6010_rev | GGGGACCACTTTGTACAAGAAAGCTGGGTTTATACGAGCTCCCTTTTTTCAAGCA |
| 0_known | L4 | Tb927.8.6680_for | GGGGACAAGTTTGTACAAAAAAGCAGGCTATGATTAAGGCTGTTATAATAATTAATATG | Tb927.8.6680_rev | GGGGACCACTTTGTACAAGAAAGCTGGGTTTAGTAATGATAGCCACTAGATGATGATCC |
| 0_known | L5 | Tb927.5.1730_for | GGGGACAAGTTTGTACAAAAAAGCAGGCTATGTTTGGTTCCCGGAAGGC | Tb927.5.1730_rev | GGGGACCACTTTGTACAAGAAAGCTGGGTTTATTCTTCGGCTTTCTTGGCCAC |
| 0_known | L6 | Tb927.8.2100_for | GGGGACAAGTTTGTACAAAAAAGCAGGCTATGCTAAGTGACGATACCTGTCAAC | Tb927.8.2100_rev | GGGGACCACTTTGTACAAGAAAGCTGGGTTCAAGAGCAACCACCCGTATAGG |
| 0_known | L7 | Tb927.11.16870_for | GGGGACAAGTTTGTACAAAAAAGCAGGCTATGTCATGGCGAGCGCGT | Tb927.11.16870_rev | GGGGACCACTTTGTACAAGAAAGCTGGGTTCACGGCATTATTTTAAAACCGACATTT |
| 0_known | L8 | Tb927.9.5650_for | GGGGACAAGTTTGTACAAAAAAGCAGGCTATGGATCCGGCGGTTAGGG | Tb927.9.5650_rev | GGGGACCACTTTGTACAAGAAAGCTGGGTCTAACTTTCCACTTTTTTGGGTTTAAGAAC |
| 0_known | L9 | Tb927.3.1410_for | GGGGACAAGTTTGTACAAAAAAGCAGGCTATGCCGCGTCCCTTTGGT | Tb927.3.1410_rev | GGGGACCACTTTGTACAAGAAAGCTGGGTCTAGGCAGGAATATAGAACTTACGTCTGC |
| 0_known | L10 | Tb927.10.8710_for | GGGGACAAGTTTGTACAAAAAAGCAGGCTATGGACGGAGATGATGGTTCGC | Tb927.10.8710_rev | GGGGACCACTTTGTACAAGAAAGCTGGGTTTACTTCTTAGGGGTGAGAAGCATACG |
| 0_known | L11 | Tb927.11.740_for | GGGGACAAGTTTGTACAAAAAAGCAGGCTATGTCTGACGATGAGGGACAGTT | Tb927.11.740_rev | GGGGACCACTTTGTACAAGAAAGCTGGGTTTATCGCTCAGCTGCATTCTTGAA |
| 0_known | L12 | Tb927.10.12530_for | GGGGACAAGTTTGTACAAAAAAGCAGGCTATGGACATAGATTCCCCCTATAAAATTTTG | Tb927.10.12530_rev | GGGGACCACTTTGTACAAGAAAGCTGGGTTCACTGCTGGGTTTGTCGC |
| 0_known | L13 | Tb927.7.1120_for | GGGGACAAGTTTGTACAAAAAAGCAGGCTATGTCTACAATCTTTGACTTTGAGGTGC | Tb927.7.1120_rev | GGGGACCACTTTGTACAAGAAAGCTGGGTTCACGCGCTCTGCGTGCT |
| 0_known | L14 | Tb927.4.4980_for | GGGGACAAGTTTGTACAAAAAAGCAGGCTATGCTTTCCGCAACTTTCGGC | Tb927.4.4980_rev | GGGGACCACTTTGTACAAGAAAGCTGGGTTCACCGCATTTCATTCGTTGTTTC |
| 0_known | L15 | Tb927.5.1570_for | GGGGACAAGTTTGTACAAAAAAGCAGGCTATGGCAAAGAAGAACAAGCACGT | Tb927.5.1570_rev | GGGGACCACTTTGTACAAGAAAGCTGGGTCTAGAAAACATAAAGGCCGTAGGGG |
| 0_known | L16 | Tb927.6.4950_for | GGGGACAAGTTTGTACAAAAAAGCAGGCTATGCAAGCGGAAGAGACCACA | Tb927.6.4950_rev | GGGGACCACTTTGTACAAGAAAGCTGGGTCTACCACGCCTTGATTTTAAAATGCA |
| 0_known | L17 | Tb927.4.3460_for | GGGGACAAGTTTGTACAAAAAAGCAGGCTATGCGTCGCTTGCAGAAGG | Tb927.4.3460_rev | GGGGACCACTTTGTACAAGAAAGCTGGGTTCACGAGGTATCTTTGTTTTTTTTATTCAA |
| 0_known | L18 | Tb927.7.1130_for | GGGGACAAGTTTGTACAAAAAAGCAGGCTATGTCAGCTGCTTCGTCAATCTTT | Tb927.7.1130_rev | GGGGACCACTTTGTACAAGAAAGCTGGGTTCACGCGCTCTGCGTGCT |
| 0_known | L19 | Tb927.8.680_for | GGGGACAAGTTTGTACAAAAAAGCAGGCTATGTTCTCTTCGACATTTAGGCGTT | Tb927.8.680_rev | GGGGACCACTTTGTACAAGAAAGCTGGGTTCACTCCAACGTAGCGACTGA |
| 0_known | L20 | Tb927.11.13690.1_for | GGGGACAAGTTTGTACAAAAAAGCAGGCTATGGAGTCGATTTTTTGCGTGGA | Tb927.11.13690.1_rev | GGGGACCACTTTGTACAAGAAAGCTGGGTCTATTGCTCATATCCTGAGCTGAAAAACA |
| 0_known | L21 | Tb927.4.3650_for | GGGGACAAGTTTGTACAAAAAAGCAGGCTATGTCCGTCTTGGTCAAGGGG | Tb927.4.3650_rev | GGGGACCACTTTGTACAAGAAAGCTGGGTTTATACAACATCAGTTTCTTGTGCCCT |
| 0_known | L22 | Tb927.5.4350_for | GGGGACAAGTTTGTACAAAAAAGCAGGCTATGTACCGAAAAAATGTATGTGTGGTAATA | Tb927.5.4350_rev | GGGGACCACTTTGTACAAGAAAGCTGGGTTTATATGGAAGACCTTGTTTTCAAAAATGC |
| 0_known | L23 | Tb927.8.6760_for | GGGGACAAGTTTGTACAAAAAAGCAGGCTATGAAGATCTTCAGGGATATCTTAACAAAT | Tb927.8.6760_rev | GGGGACCACTTTGTACAAGAAAGCTGGGTTCACACGCGTTCGCCCTT |
| 0_known | L24 | Tb927.7.2590_for | GGGGACAAGTTTGTACAAAAAAGCAGGCTATGAGTGGTCAGCAAGGTGAGG | Tb927.7.2590_rev | GGGGACCACTTTGTACAAGAAAGCTGGGTCTAAGCCACCGTCTGCGC |
| 0_known | M2 | Tb927.7.1180_for | GGGGACAAGTTTGTACAAAAAAGCAGGCTATGCAACGATACAAGGAGGGCA | Tb927.7.1180_rev | GGGGACCACTTTGTACAAGAAAGCTGGGTTCACCTCGCTTCCAACACAAC |
| 0_known | M3 | Tb927.1.630_for | GGGGACAAGTTTGTACAAAAAAGCAGGCTATGAGCATACAACCAATTGTACAACTTTTG | Tb927.1.630_rev | GGGGACCACTTTGTACAAGAAAGCTGGGTTCATCTGGGTTCAAGTGCTTCTCT |
| 0_known | M4 | Tb927.8.5880_for | GGGGACAAGTTTGTACAAAAAAGCAGGCTATGCCCAAGAATATGGGAAAGGGC | Tb927.8.5880_rev | GGGGACCACTTTGTACAAGAAAGCTGGGTTCAGAGGTCGTCAAAGTTTGCC |
| 0_known | M5 | Tb927.1.4490_for | GGGGACAAGTTTGTACAAAAAAGCAGGCTATGACTTCCGCGTACCGCA | Tb927.1.4490_rev | GGGGACCACTTTGTACAAGAAAGCTGGGTTCAATATTTGCTTTTCATGCGACCC |
| 0_known | M6 | Tb927.5.1030_for | GGGGACAAGTTTGTACAAAAAAGCAGGCTATGCTGCGCACTTCCCTG | Tb927.5.1030_rev | GGGGACCACTTTGTACAAGAAAGCTGGGTTCACACCTTTTGCTGCTGCTT |
| 0_known | M7 | Tb927.9.12410_for | GGGGACAAGTTTGTACAAAAAAGCAGGCTATGCGACAAACTGGAAGTTTTCAGC | Tb927.9.12410_rev | GGGGACCACTTTGTACAAGAAAGCTGGGTTCACTCATCGGGTAGATCCTCGA |
| 0_known | M8 | Tb927.11.8910_for | GGGGACAAGTTTGTACAAAAAAGCAGGCTATGATTCGCAAGACTCCACCAC | Tb927.11.8910_rev | GGGGACCACTTTGTACAAGAAAGCTGGGTTCAAAAGTATCCTATGGTCCAATTGTAGC |
| 0_known | M9 | Tb927.11.9140_for | GGGGACAAGTTTGTACAAAAAAGCAGGCTATGAGGAGTAGCCGGGGTATTTT | Tb927.11.9140_rev | GGGGACCACTTTGTACAAGAAAGCTGGGTTTACTGTGTACCGTGTTCACCCG |
| 0_known | M10 | Tb927.11.510_for | GGGGACAAGTTTGTACAAAAAAGCAGGCTATGTCCCAGCAACCGCAGT | Tb927.11.510_rev | GGGGACCACTTTGTACAAGAAAGCTGGGTTTACTGACGGGATATCGGAGGC |
| 0_known | M11 | Tb927.6.2130_for | GGGGACAAGTTTGTACAAAAAAGCAGGCTATGTCTTTAGTTGAAAGTCAGAAGGACATG | Tb927.6.2130_rev | GGGGACCACTTTGTACAAGAAAGCTGGGTTTAGTTAGAGGTACCACCACGCTG |
| 0_known | M12 | Tb927.7.1140_for | GGGGACAAGTTTGTACAAAAAAGCAGGCTATGCTGCGTTCATCTCGGAAAA | Tb927.7.1140_rev | GGGGACCACTTTGTACAAGAAAGCTGGGTTCAAAGTCGCGCACTCCCG |
| 0_known | M13 | Tb927.8.5780_for | GGGGACAAGTTTGTACAAAAAAGCAGGCTATGGACGTCAACTGTACATTGATTGA | Tb927.8.5780_rev | GGGGACCACTTTGTACAAGAAAGCTGGGTTTACATGATGGCGCAACCTCC |
| 0_known | M14 | Tb927.3.3750_for | GGGGACAAGTTTGTACAAAAAAGCAGGCTATGTCGACTCAACTTATTTCCGGTG | Tb927.3.3750_rev | GGGGACCACTTTGTACAAGAAAGCTGGGTTTACCGACGCAGCCACGA |
| 0_known | M15 | Tb927.10.14260_for | GGGGACAAGTTTGTACAAAAAAGCAGGCTATGACGACAATTGAGTTCATATTTCTCGT | Tb927.10.14260_rev | GGGGACCACTTTGTACAAGAAAGCTGGGTCTACAGTCCTTTTGATTCTTTGTCCATCA |
| 0_known | M16 | Tb927.11.880_for | GGGGACAAGTTTGTACAAAAAAGCAGGCTATGTCATACAGGCCCCACCAC | Tb927.11.880_rev | GGGGACCACTTTGTACAAGAAAGCTGGGTTTATAGTTGGCCGCACGAGTCA |
| 0_known | M17 | Tb927.8.550_for | GGGGACAAGTTTGTACAAAAAAGCAGGCTATGAACCCAAATGCTGTTGCTACT | Tb927.8.550_rev | GGGGACCACTTTGTACAAGAAAGCTGGGTTCACCAGTAGAGACGGTGTGC |
| 0_known | M18 | Tb927.4.4680_for | GGGGACAAGTTTGTACAAAAAAGCAGGCTATGATCAAGAATTATTTTGATCTCTCTAGC | Tb927.4.4680_rev | GGGGACCACTTTGTACAAGAAAGCTGGGTCTAGAAGGCGCTACGGTACCA |
| 0_known | M19 | Tb927.7.890_for | GGGGACAAGTTTGTACAAAAAAGCAGGCTATGGCTGCTTTACGCTACTTGC | Tb927.7.890_rev | GGGGACCACTTTGTACAAGAAAGCTGGGTTCAGTGATGAGGGGTTATGGTGTG |
| 0_known | M20 | Tb927.3.3450_for | GGGGACAAGTTTGTACAAAAAAGCAGGCTATGGGCTTGCTTACGCTTCTG | Tb927.3.3450_rev | GGGGACCACTTTGTACAAGAAAGCTGGGTCTATTTCTTTATGCTTTTAATGGCCCACAT |
| 0_known | M21 | Tb927.10.370_for | GGGGACAAGTTTGTACAAAAAAGCAGGCTATGGAAGTTGCTGCGGTTGC | Tb927.10.370_rev | GGGGACCACTTTGTACAAGAAAGCTGGGTTTAAAACTGCAGAACATGGGCATTATCC |
| 0_known | M22 | Tb927.9.13100_for | GGGGACAAGTTTGTACAAAAAAGCAGGCTATGACAACAGGTGGAAAGCAAGAA | Tb927.9.13100_rev | GGGGACCACTTTGTACAAGAAAGCTGGGTTTAACACACCTCTGCATGCTGTC |
| 0_known | M23 | Tb927.7.6360_for | GGGGACAAGTTTGTACAAAAAAGCAGGCTATGTCTCTTACAGGTGATGATGCGG | Tb927.7.6360_rev | GGGGACCACTTTGTACAAGAAAGCTGGGTCTACGAGCCCCTCTTCGATTTCT |
| 0_known | M24 | Tb927.9.11720_for | GGGGACAAGTTTGTACAAAAAAGCAGGCTATGCGGCGACTGATATCATCAC | Tb927.9.11720_rev | GGGGACCACTTTGTACAAGAAAGCTGGGTTCAGCTTGACACCTCACCAACA |
| 0_known | N1 | Tb927.10.4840_for | GGGGACAAGTTTGTACAAAAAAGCAGGCTATGTACCGTGAGAGCCTGCT | Tb927.10.4840_rev | GGGGACCACTTTGTACAAGAAAGCTGGGTTTAGGGCCGCGAGCGCTT |
| 0_known | N2 | Tb927.3.3470_for | GGGGACAAGTTTGTACAAAAAAGCAGGCTATGTTCGACTACCTTCTTTCCCTCT | Tb927.3.3470_rev | GGGGACCACTTTGTACAAGAAAGCTGGGTTTAGGATGACTTAGCCCCACTGC |
| 0_known | N3 | Tb927.7.3900_for | GGGGACAAGTTTGTACAAAAAAGCAGGCTATGGCAAGCGTAATTGAAAAGGGG | Tb927.7.3900_rev | GGGGACCACTTTGTACAAGAAAGCTGGGTTCAAGAAAATCGTGGTCCATCATTGA |
| 0_known | N4 | Tb927.2.2900_for | GGGGACAAGTTTGTACAAAAAAGCAGGCTATGAGTGAAGAAGAACTCTTGTATTACACT | Tb927.2.2900_rev | GGGGACCACTTTGTACAAGAAAGCTGGGTTCAGTTAGTCGATAAGGCTTGAAAGAAC |
| 0_known | N5 | Tb927.3.1000_for | GGGGACAAGTTTGTACAAAAAAGCAGGCTATGCGGCGCACATGTTGC | Tb927.3.1000_rev | GGGGACCACTTTGTACAAGAAAGCTGGGTCTATTCGGTTTCTCCAGCCCCT |
| 0_known | N6 | Tb927.9.3000_for | GGGGACAAGTTTGTACAAAAAAGCAGGCTATGCAATGCTCATCATTGAAGACCG | Tb927.9.3000_rev | GGGGACCACTTTGTACAAGAAAGCTGGGTTCAGAAAGTTCCTGGACACCTTGA |
| 0_known | N7 | Tb927.9.13680_for | GGGGACAAGTTTGTACAAAAAAGCAGGCTATGGGTCAATGGTTGGCTTCCG | Tb927.9.13680_rev | GGGGACCACTTTGTACAAGAAAGCTGGGTCTACTTCATACTCTTCTTAATGTTTGCTGA |
| 0_known | N8 | Tb927.9.13710_for | GGGGACAAGTTTGTACAAAAAAGCAGGCTATGGGTCAGTGGTTGGCTTCC | Tb927.9.13710_rev | GGGGACCACTTTGTACAAGAAAGCTGGGTCTACTTCATACTCTTCTTAATGTTTGCTGA |
| 0_known | N9 | Tb927.7.2510_for | GGGGACAAGTTTGTACAAAAAAGCAGGCTATGTCGTCGAAGGACATTATTGGAC | Tb927.7.2510_rev | GGGGACCACTTTGTACAAGAAAGCTGGGTTCAACTCTTGGGCACAAACTTTGG |
| 0_known | N11 | Tb927.6.4990_for | GGGGACAAGTTTGTACAAAAAAGCAGGCTATGTTTCGCACGTTCGGTCG | Tb927.6.4990_rev | GGGGACCACTTTGTACAAGAAAGCTGGGTCTAGTGGTGTTTTAGGGCAGCAA |
| 0_known | N12 | Tb927.9.13650_for | GGGGACAAGTTTGTACAAAAAAGCAGGCTATGGGTCAATGGTTGGCTTCCG | Tb927.9.13650_rev | GGGGACCACTTTGTACAAGAAAGCTGGGTCTACTTCATACTCTTCTTAATGTTCACTGA |

| | | | | | |
|---|---|---|---|---|---|
| 0_known | N13 | Tb927.11.6130_for | GGGGACAAGTTTGTACAAAAAAGCAGGCTATGGCGGAAGAGGTGTCGG | Tb927.11.6130_rev | GGGGACCACTTTGTACAAGAAAGCTGGGTTTACGCCTCTTCGCACCAGC |
| 0_known | N15 | Tb927.11.580_for | GGGGACAAGTTTGTACAAAAAAGCAGGCTATGATAGTACCGTCAAATTTTGGGTACG | Tb927.11.580_rev | GGGGACCACTTTGTACAAGAAAGCTGGGTTTAGTCGTTTTCATAGAAAAGGATAGTTGG |
| 0_known | N16 | Tb927.4.2260_for | GGGGACAAGTTTGTACAAAAAAGCAGGCTATGTCCTCATCTCGTCCTGCC | Tb927.4.2260_rev | GGGGACCACTTTGTACAAGAAAGCTGGGTTCAGTAGTCATCATCTTCCAGCATGA |
| 0_known | N17 | Tb927.6.3650_for | GGGGACAAGTTTGTACAAAAAAGCAGGCTATGCTTAAGGGGATCCGTAGCC | Tb927.6.3650_rev | GGGGACCACTTTGTACAAGAAAGCTGGGTTTACGGGGCACAAGTGCTAAGT |
| 0_known | N18 | Tb927.9.11740_for | GGGGACAAGTTTGTACAAAAAAGCAGGCTATGTCGGTTGTGATCTCAACATCAC | Tb927.9.11740_rev | GGGGACCACTTTGTACAAGAAAGCTGGGTTCATACATTTAGCTTGATGTGGCCC |
| 0_known | N19 | Tb927.10.4540_for | GGGGACAAGTTTGTACAAAAAAGCAGGCTATGGAGAGTCAGTGGAACGGTT | Tb927.10.4540_rev | GGGGACCACTTTGTACAAGAAAGCTGGGTTTACCCACCAAGCAAAAACGGC |
| 0_known | N20 | Tb927.9.3480_for | GGGGACAAGTTTGTACAAAAAAGCAGGCTATGTATAATGGCATCGCTCCGATC | Tb927.9.3480_rev | GGGGACCACTTTGTACAAGAAAGCTGGGTTTACTCTCCCTTGATACGCTTCACC |
| 0_known | N21 | Tb927.10.4270_for | GGGGACAAGTTTGTACAAAAAAGCAGGCTATGGAGTACATGCATCATATTCAGGGC | Tb927.10.4270_rev | GGGGACCACTTTGTACAAGAAAGCTGGGTTTAAAGATTCTCCTTAAGGTATTCGTTGAC |
| 0_known | N22 | Tb927.3.5550_for | GGGGACAAGTTTGTACAAAAAAGCAGGCTATGGTAAACTTGCGTCTTCAGGTG | Tb927.3.5550_rev | GGGGACCACTTTGTACAAGAAAGCTGGGTTTACCTCATTTGGGTGAGTTGTTCAA |
| 0_known | N23 | Tb927.8.6440_for | GGGGACAAGTTTGTACAAAAAAGCAGGCTATGGAAGACAATCTTGCGGCAG | Tb927.8.6440_rev | GGGGACCACTTTGTACAAGAAAGCTGGGTTCAACATACAATTAAAGTACTACCCTTGAA |
| 0_known | N24 | Tb927.7.1390_for | GGGGACAAGTTTGTACAAAAAAGCAGGCTATGACTGTGAACATCGTTGCGC | Tb927.7.1390_rev | GGGGACCACTTTGTACAAGAAAGCTGGGTTTAGCTCCAACTTGGGTACGGG |
| 0_known | O1 | Tb927.9.3590_for | GGGGACAAGTTTGTACAAAAAAGCAGGCTATGCGGCGACTGAGCGGT | Tb927.9.3590_rev | GGGGACCACTTTGTACAAGAAAGCTGGGTTTACAGGTCCCGACAAGTGATTCC |
| 0_known | O2 | Tb927.11.380_for | GGGGACAAGTTTGTACAAAAAAGCAGGCTATGGCAGAGCTTCATGTTATTGGAG | Tb927.11.380_rev | GGGGACCACTTTGTACAAGAAAGCTGGGTTCAAGCACAACGGACTCCTAATCG |
| 0_known | O3 | Tb927.10.6980_for | GGGGACAAGTTTGTACAAAAAAGCAGGCTATGTCGTCTTTAGACTCTGTTAGAGAAGA | Tb927.10.6980_rev | GGGGACCACTTTGTACAAGAAAGCTGGGTCTAAATCGACTTTCGTAGTGTTTCCATCA |
| 0_known | O4 | Tb927.5.3580_for | GGGGACAAGTTTGTACAAAAAAGCAGGCTATGGCAGAGGTGCATGTATGTAGG | Tb927.5.3580_rev | GGGGACCACTTTGTACAAGAAAGCTGGGTTCACAGTTCCACAATGTCAGTGTTT |
| 0_known | O5 | Tb927.6.3840_for | GGGGACAAGTTTGTACAAAAAAGCAGGCTATGTCCTGTTTCCTTTGCAACCA | Tb927.6.3840_rev | GGGGACCACTTTGTACAAGAAAGCTGGGTCTAATTGTTCTTCTTTGTCAACTGGTCCA |
| 0_known | O6 | Tb927.11.13900_for | GGGGACAAGTTTGTACAAAAAAGCAGGCTATGGAAGGGCCCAAGTTAAAGGG | Tb927.11.13900_rev | GGGGACCACTTTGTACAAGAAAGCTGGGTTTAGTACACATGGGCTCGATTCATTAGC |
| 0_known | O7 | Tb927.3.610_for | GGGGACAAGTTTGTACAAAAAAGCAGGCTATGACCACTTTTAGGGATTTCAACTTAGTG | Tb927.3.610_rev | GGGGACCACTTTGTACAAGAAAGCTGGGTTTAATTCCACTCCAATTCATCAGGACG |
| 0_known | O8 | Tb927.7.1670_for | GGGGACAAGTTTGTACAAAAAAGCAGGCTATGGCCGCTGAGGCTACTG | Tb927.7.1670_rev | GGGGACCACTTTGTACAAGAAAGCTGGGTCTACTTCGCCTCCTCCCCTTC |
| 0_known | O9 | Tb927.10.4250_for | GGGGACAAGTTTGTACAAAAAAGCAGGCTATGGGTTTACTATCGATCATCCGCA | Tb927.10.4250_rev | GGGGACCACTTTGTACAAGAAAGCTGGGTCTACCCCGCAAAATACATTCTACTGG |
| 0_known | O10 | Tb927.10.3980_for | GGGGACAAGTTTGTACAAAAAAGCAGGCTATGAAAAGGTGGAGGAGAAGGCG | Tb927.10.3980_rev | GGGGACCACTTTGTACAAGAAAGCTGGGTTCATTCCCTTCTTCCCTCATAAAAATAAAA |
| 0_known | O11 | Tb927.6.1970_for | GGGGACAAGTTTGTACAAAAAAGCAGGCTATGGCAGAATACCTCATTGATCTGACG | Tb927.6.1970_rev | GGGGACCACTTTGTACAAGAAAGCTGGGTTTACGAACGGCCGCGCTT |
| 0_known | O12 | Tb927.8.7000_for | GGGGACAAGTTTGTACAAAAAAGCAGGCTATGAACACGCCCTCTGTTCAATG | Tb927.8.7000_rev | GGGGACCACTTTGTACAAGAAAGCTGGGTCTATCCGCTGTCGCATGGAC |
| 0_known | O13 | Tb927.1.840_for | GGGGACAAGTTTGTACAAAAAAGCAGGCTATGCGCCGGATGTGTGTG | Tb927.1.840_rev | GGGGACCACTTTGTACAAGAAAGCTGGGTTTAAGCGTCAATTTCACTTTTGATTTTGGC |
| 0_known | O14 | Tb927.9.12200_for | GGGGACAAGTTTGTACAAAAAAGCAGGCTATGACTCGCGCTGGATTTAAGG | Tb927.9.12200_rev | GGGGACCACTTTGTACAAGAAAGCTGGGTTTACTGCTCAATCACCTTAGTTGTCAG |
| 0_known | O16 | Tb927.5.1710_for | GGGGACAAGTTTGTACAAAAAAGCAGGCTATGATGCGCCGTGTTTATGCC | Tb927.5.1710_rev | GGGGACCACTTTGTACAAGAAAGCTGGGTTTACTCTACCTTTACATCAAACAATCCTGC |
| 0_known | O17 | Tb927.5.2880_for | GGGGACAAGTTTGTACAAAAAAGCAGGCTATGGGGTCAAAGGAGGACCAAG | Tb927.5.2880_rev | GGGGACCACTTTGTACAAGAAAGCTGGGTTCAGAAAAGGTTTAAATCTTTCCACTCTTT |
| 0_known | O18 | Tb927.9.8100_for | GGGGACAAGTTTGTACAAAAAAGCAGGCTATGAGCGCTAACGATGTTCAAAACC | Tb927.9.8100_rev | GGGGACCACTTTGTACAAGAAAGCTGGGTTCAGTTGGGATCCATCGTCAGT |
| 0_known | O19 | Tb927.9.5960_for | GGGGACAAGTTTGTACAAAAAAGCAGGCTATGCCTGCTGCACCGCTT | Tb927.9.5960_rev | GGGGACCACTTTGTACAAGAAAGCTGGGTTTAATACTTGGAGGCGACTTTTGCA |
| 0_known | O21 | Tb927.9.13200_for | GGGGACAAGTTTGTACAAAAAAGCAGGCTATGAGGCTAACGGCTCTTTTGG | Tb927.9.13200_rev | GGGGACCACTTTGTACAAGAAAGCTGGGTTTATGACATCATTAACATCGCGGAAACT |
| 0_known | O22 | Tb927.8.2200_for | GGGGACAAGTTTGTACAAAAAAGCAGGCTATGTCACAGCAGGTTGGAGAGG | Tb927.8.2200_rev | GGGGACCACTTTGTACAAGAAAGCTGGGTCTAAGATCCAACCATCACCATCCAGG |
| 0_known | O23 | Tb927.9.7650_for | GGGGACAAGTTTGTACAAAAAAGCAGGCTATGGGTAACGTGCTGAGTTGGT | Tb927.9.7650_rev | GGGGACCACTTTGTACAAGAAAGCTGGGTTCAATGCGTATCGGGTAAGTGACT |
| 0_known | O24 | Tb927.11.16160_for | GGGGACAAGTTTGTACAAAAAAGCAGGCTATGACAAATCCAGCAGAACAGGC | Tb927.11.16160_rev | GGGGACCACTTTGTACAAGAAAGCTGGGTTTAGTCATTACCCTGATCCGCCG |
| 0_known | P1 | Tb927.8.5060_for | GGGGACAAGTTTGTACAAAAAAGCAGGCTATGGGACAATCCAAAACGAAGCT | Tb927.8.5060_rev | GGGGACCACTTTGTACAAGAAAGCTGGGTTCAGCCCCTCTTCGTCCCA |
| 0_known | P2 | Tb927.5.970_for | GGGGACAAGTTTGTACAAAAAAGCAGGCTATGGCTGTACAACCGGGAAAGG | Tb927.5.970_rev | GGGGACCACTTTGTACAAGAAAGCTGGGTCTACATGTTCAGCTGGGAGAATGC |
| 0_known | P3 | Tb927.10.8580_for | GGGGACAAGTTTGTACAAAAAAGCAGGCTATGGGCCTGTTGGAGTTCCT | Tb927.10.8580_rev | GGGGACCACTTTGTACAAGAAAGCTGGGTTTATACTTGGCCGTCTGGCCT |
| 0_known | P4 | Tb927.9.9060_for | GGGGACAAGTTTGTACAAAAAAGCAGGCTATGCCTGTCTGCAACAACGAC | Tb927.9.9060_rev | GGGGACCACTTTGTACAAGAAAGCTGGGTTCACTTCTTGGAAAGAGTCTTTTGAACT |
| 0_known | P5 | Tb927.7.6230_for | GGGGACAAGTTTGTACAAAAAAGCAGGCTATGGGGGCTTTGGTTTCGC | Tb927.7.6230_rev | GGGGACCACTTTGTACAAGAAAGCTGGGTTCACGAGTCTGCAGTGACGA |
| 0_known | P6 | Tb927.11.14490_for | GGGGACAAGTTTGTACAAAAAAGCAGGCTATGTTCTACAAAATGAAATTGGAGAGGAAT | Tb927.11.14490_rev | GGGGACCACTTTGTACAAGAAAGCTGGGTCTACAGTGGTCCTAAATAATTCCCTACCA |
| 0_known | P7 | Tb927.4.2040_for | GGGGACAAGTTTGTACAAAAAAGCAGGCTATGCCTTCATATCCGCCACGT | Tb927.4.2040_rev | GGGGACCACTTTGTACAAGAAAGCTGGGTTTACTCCTCATTGCCACCACCC |
| 0_known | P8 | Tb927.11.9450_for | GGGGACAAGTTTGTACAAAAAAGCAGGCTATGCGCATTTCTGAGCGAGC | Tb927.11.9450_rev | GGGGACCACTTTGTACAAGAAAGCTGGGTTCATTCCAATACCCCACAATCCACA |
| 0_known | P9 | Tb927.10.200_for | GGGGACAAGTTTGTACAAAAAAGCAGGCTATGAAGACACAAAACTTGCTTTTTGCT | Tb927.10.200_rev | GGGGACCACTTTGTACAAGAAAGCTGGGTTCACTGACACGGTCCGTAGTT |
| 0_known | P10 | Tb927.9.7230_for | GGGGACAAGTTTGTACAAAAAAGCAGGCTATGGGACAACTTATTAGTGGCCTTTG | Tb927.9.7230_rev | GGGGACCACTTTGTACAAGAAAGCTGGGTTCACGCACTGCTCGCAGG |
| 0_known | P11 | Tb927.1.1400_for | GGGGACAAGTTTGTACAAAAAAGCAGGCTATGTGTGACACAGGCGTTCAG | Tb927.1.1400_rev | GGGGACCACTTTGTACAAGAAAGCTGGGTTCAACTCCAAAAGGCCAGTTCC |
| 0_known | P12 | Tb927.8.5030_for | GGGGACAAGTTTGTACAAAAAAGCAGGCTATGTCCCAACAGCGGCAGG | Tb927.8.5030_rev | GGGGACCACTTTGTACAAGAAAGCTGGGTCTATTGTTCAACCTTAAAGGTTTCCTTCTC |
| 0_known | P13 | Tb927.10.2120_for | GGGGACAAGTTTGTACAAAAAAGCAGGCTATGAGTGATAGTTTTGCTTTGGATGATGC | Tb927.10.2120_rev | GGGGACCACTTTGTACAAGAAAGCTGGGTTCATTCACGTTTCGTGACCGAA |
| 0_known | P14 | Tb927.5.4500_for | GGGGACAAGTTTGTACAAAAAAGCAGGCTATGGGGCTGTTTTCGTGGTTT | Tb927.5.4500_rev | GGGGACCACTTTGTACAAGAAAGCTGGGTCTAAGAATTTTGAAGGTACTGTGATATCCA |
| 0_known | P15 | Tb927.8.2090_for | GGGGACAAGTTTGTACAAAAAAGCAGGCTATGAAGGTGGTGGTCAATGAAATGT | Tb927.8.2090_rev | GGGGACCACTTTGTACAAGAAAGCTGGGTTCACTCCAAATCATTCAAGATACTACTCTG |
| 0_known | P16 | Tb927.8.6910_for | GGGGACAAGTTTGTACAAAAAAGCAGGCTATGCCAGTCAAGAGGTCCAACA | Tb927.8.6910_rev | GGGGACCACTTTGTACAAGAAAGCTGGGTTTATTGCTTCGCCTTGGGGAC |
| 0_known | P17 | Tb927.7.3450_for | GGGGACAAGTTTGTACAAAAAAGCAGGCTATGCTCTGCCCTCCAGATGTT | Tb927.7.3450_rev | GGGGACCACTTTGTACAAGAAAGCTGGGTTCACAACTCTGCCTCTTCATCTGC |
| 0_known | P18 | Tb927.10.5020_for | GGGGACAAGTTTGTACAAAAAAGCAGGCTATGGAGGAGGAGAGTCACGC | Tb927.10.5020_rev | GGGGACCACTTTGTACAAGAAAGCTGGGTTTATTTTCCCTTTTTTTTCTTCTCCTTTTC |
| 0_known | P19 | Tb927.10.15590_for | GGGGACAAGTTTGTACAAAAAAGCAGGCTATGTATGGTGTTGAGTGTTTTCGGC | Tb927.10.15590_rev | GGGGACCACTTTGTACAAGAAAGCTGGGTTTACAAGTCTCCCTTCCCCGC |
| 0_known | P20 | Tb927.8.1080_for | GGGGACAAGTTTGTACAAAAAAGCAGGCTATGAGCACCACCAACCGTGG | Tb927.8.1080_rev | GGGGACCACTTTGTACAAGAAAGCTGGGTCTAATAGAGCGAAGTTTTCTTCATGATGC |
| 0_known | P21 | Tb927.2.1680_for | GGGGACAAGTTTGTACAAAAAAGCAGGCTATGTTGTCGCGCACCTTTTTTG | Tb927.2.1680_rev | GGGGACCACTTTGTACAAGAAAGCTGGGTTTATCCCTGTTCAATTTGGCCACAA |
| 0_known | P22 | Tb927.9.3170_for | GGGGACAAGTTTGTACAAAAAAGCAGGCTATGAAGCGCTTTGTCACACCA | Tb927.9.3170_rev | GGGGACCACTTTGTACAAGAAAGCTGGGTCTAGTTACTGATTTTGTTCCCATCCTTGA |
| 0_known | P24 | Tb927.2.4580_for | GGGGACAAGTTTGTACAAAAAAGCAGGCTATGGCTGCGACGACCGAG | Tb927.2.4580_rev | GGGGACCACTTTGTACAAGAAAGCTGGGTTTATTTAGTGAATTCCTCCTCTGGTGC |
| 0_hypothetical | A1 | Tb927.8.1305_for | GGGGACAAGTTTGTACAAAAAAGCAGGCTATGTTGTGGCAAGCCAAAAGAAC | Tb927.8.1305_rev | GGGGACCACTTTGTACAAGAAAGCTGGGTCTAGACTACAACGGCGGGCA |
| 0_hypothetical | A2 | Tb927.3.515_for | GGGGACAAGTTTGTACAAAAAAGCAGGCTATGAAGCACTGGTGGAAGAGTGG | Tb927.3.515_rev | GGGGACCACTTTGTACAAGAAAGCTGGGTTTATAGCATGTCGCCCACTGGC |
| 0_hypothetical | A3 | Tb927.10.7675_for | GGGGACAAGTTTGTACAAAAAAGCAGGCTATGCTCCCGCTTTTCGCATG | Tb927.10.7675_rev | GGGGACCACTTTGTACAAGAAAGCTGGGTTTAATACGCCACGGCGACTTC |
| 0_hypothetical | A4 | Tb927.9.15905_for | GGGGACAAGTTTGTACAAAAAAGCAGGCTATGATTCATTTCTGCTGTTGTGATAGCA | Tb927.9.15905_rev | GGGGACCACTTTGTACAAGAAAGCTGGGTTTACATTCCTTCCTTCATAACCTTACCG |
| 0_hypothetical | A5 | Tb927.10.16121_for | GGGGACAAGTTTGTACAAAAAAGCAGGCTATGCCGGAAGTGAAGATGATGTTG | Tb927.10.16121_rev | GGGGACCACTTTGTACAAGAAAGCTGGGTTTACTGCATCGATGCGTTGTGC |
| 0_hypothetical | A6 | Tb927.8.6035_for | GGGGACAAGTTTGTACAAAAAAGCAGGCTATGGGGACCATCAGCCGC | Tb927.8.6035_rev | GGGGACCACTTTGTACAAGAAAGCTGGGTTTACTTGGAGCGCTGGCGC |
| 0_hypothetical | A9 | Tb927.11.5855_for | GGGGACAAGTTTGTACAAAAAAGCAGGCTATGAAGGAAATAATAACCTTAAGGGAATTT | Tb927.11.5855_rev | GGGGACCACTTTGTACAAGAAAGCTGGGTTTAAATGGTCAAGACGCTAAGTGTGT |
| 0_hypothetical | A10 | Tb927.9.19145_for | GGGGACAAGTTTGTACAAAAAAGCAGGCTATGATTCATTTCTGCTGTTGTGATAGCA | Tb927.9.19145_rev | GGGGACCACTTTGTACAAGAAAGCTGGGTTTACATTCTTTCCTTCATAACCTTACCGAA |
| 0_hypothetical | A11 | Tb927.9.4755_for | GGGGACAAGTTTGTACAAAAAAGCAGGCTATGGTGCCACGCCGAGAG | Tb927.9.4755_rev | GGGGACCACTTTGTACAAGAAAGCTGGGTTTACTGCGGCAAATGATGGAAGG |
| 0_hypothetical | A12 | Tb927.7.6855_for | GGGGACAAGTTTGTACAAAAAAGCAGGCTATGAAAGGCACGTTTTGCTTTGG | Tb927.7.6855_rev | GGGGACCACTTTGTACAAGAAAGCTGGGTTCAACCCTTCGTATACATTTTCCCCA |
| 0_hypothetical | A13 | Tb927.2.725_for | GGGGACAAGTTTGTACAAAAAAGCAGGCTATGGAGTCGTTTCTCCACACCT | Tb927.2.725_rev | GGGGACCACTTTGTACAAGAAAGCTGGGTTCAAGAGTGAGCTCTGGCCTT |
| 0_hypothetical | A14 | Tb927.2.1045_for | GGGGACAAGTTTGTACAAAAAAGCAGGCTATGGAGACGTTTCTCCACATCTGC | Tb927.2.1045_rev | GGGGACCACTTTGTACAAGAAAGCTGGGTTCAAGAGTGAGCTCTGGCCTT |
| 0_hypothetical | A15 | Tb927.2.4925_for | GGGGACAAGTTTGTACAAAAAAGCAGGCTATGGGCACGACCGAGATAAAATG | Tb927.2.4925_rev | GGGGACCACTTTGTACAAGAAAGCTGGGTTTATCTTTTGTCCACGTTACGTGCA |
| 0_hypothetical | A16 | Tb927.7.6425_for | GGGGACAAGTTTGTACAAAAAAGCAGGCTATGTTGCAGTTCCGTAGTTGTTCT | Tb927.7.6425_rev | GGGGACCACTTTGTACAAGAAAGCTGGGTTCAAGAATGACACGCAGAAATAATAAAAGG |
| 0_hypothetical | A17 | Tb927.11.6565_for | GGGGACAAGTTTGTACAAAAAAGCAGGCTATGCTTCAGTTGAGCATACCATGTAG | Tb927.11.6565_rev | GGGGACCACTTTGTACAAGAAAGCTGGGTTTACTGCTCATAGAAAACAGCTCCCC |
| 0_hypothetical | A18 | Tb927.1.2760_for | GGGGACAAGTTTGTACAAAAAAGCAGGCTATGTGGGTTTCCATCTACCGATGT | Tb927.1.2760_rev | GGGGACCACTTTGTACAAGAAAGCTGGGTTTAATCCTCTTTAACTTTTACCACCTCCG |
| 0_hypothetical | A19 | Tb927.8.2685_for | GGGGACAAGTTTGTACAAAAAAGCAGGCTATGGAATTAGACAGGGTTGTTTGGATT | Tb927.8.2685_rev | GGGGACCACTTTGTACAAGAAAGCTGGGTCTAGGCGTTTAAATTCACATCAGGTAAAAA |
| 0_hypothetical | A20 | Tb927.11.16915_for | GGGGACAAGTTTGTACAAAAAAGCAGGCTATGCTTCATTGTTTAAGCTGTATCGAATTG | Tb927.11.16915_rev | GGGGACCACTTTGTACAAGAAAGCTGGGTTCACATTACTTCTACCGTTGTTTCCTTC |
| 0_hypothetical | A21 | Tb927.11.8775_for | GGGGACAAGTTTGTACAAAAAAGCAGGCTATGCGTGAACTGCCATTTTCCA | Tb927.11.8775_rev | GGGGACCACTTTGTACAAGAAAGCTGGGTTCACTCGTCATAAAGTTTCGTATCGC |
| 0_hypothetical | A22 | Tb927.10.9465_for | GGGGACAAGTTTGTACAAAAAAGCAGGCTATGCACAATCTTAACACCATCCAACA | Tb927.10.9465_rev | GGGGACCACTTTGTACAAGAAAGCTGGGTCTACTTATTGCCGGCACGAAGAC |
| 0_hypothetical | A23 | Tb927.10.1991_for | GGGGACAAGTTTGTACAAAAAAGCAGGCTATGTCTGTGCTACATGAAGTTGTGAT | Tb927.10.1991_rev | GGGGACCACTTTGTACAAGAAAGCTGGGTCTATTTGTTGTCACCACTCTCACATATGG |
| 0_hypothetical | A24 | Tb927.8.8215_for | GGGGACAAGTTTGTACAAAAAAGCAGGCTATGGCAAACACGGCTCTGC | Tb927.8.8215_rev | GGGGACCACTTTGTACAAGAAAGCTGGGTTTACAGACTTTCACCGACACTATGAGC |
| 0_hypothetical | B1 | Tb927.5.1145_for | GGGGACAAGTTTGTACAAAAAAGCAGGCTATGGGAGAAAAGGATTCTGTGGCG | Tb927.5.1145_rev | GGGGACCACTTTGTACAAGAAAGCTGGGTTCACATCCACAGTAAGTTATGCACTTG |
| 0_hypothetical | B2 | Tb927.7.5945_for | GGGGACAAGTTTGTACAAAAAAGCAGGCTATGAGAGCATGGCATTCATTGTTCA | Tb927.7.5945_rev | GGGGACCACTTTGTACAAGAAAGCTGGGTCTACTGCCTGTCAAATGCCGAG |
| 0_hypothetical | B3 | Tb927.1.5180_for | GGGGACAAGTTTGTACAAAAAAGCAGGCTATGCGAGCAGGAGAAACGAC | Tb927.1.5180_rev | GGGGACCACTTTGTACAAGAAAGCTGGGTTTACGTAACCCGCACAATATGTGG |
| 0_hypothetical | B4 | Tb927.10.9466_for | GGGGACAAGTTTGTACAAAAAAGCAGGCTATGTGTGTAGTACATTGGAGGTGTACT | Tb927.10.9466_rev | GGGGACCACTTTGTACAAGAAAGCTGGGTTTACCCTACAGCAACGAAAATAAACGA |
| 0_hypothetical | B5 | Tb927.11.4105_for | GGGGACAAGTTTGTACAAAAAAGCAGGCTATGGATCCCATCATATGGGAGGATAAC | Tb927.11.4105_rev | GGGGACCACTTTGTACAAGAAAGCTGGGTCTATTTGCTCCGCTGGTATTCATGC |
| 0_hypothetical | B6 | Tb927.10.8550_for | GGGGACAAGTTTGTACAAAAAAGCAGGCTATGCACGTTGGCCTCGCG | Tb927.10.8550_rev | GGGGACCACTTTGTACAAGAAAGCTGGGTTCACAACAGAAGACACGTCCGG |
| 0_hypothetical | B7 | Tb927.11.101_for | GGGGACAAGTTTGTACAAAAAAGCAGGCTATGAAGGGTAAGTGTACCTCTTCTGG | Tb927.11.101_rev | GGGGACCACTTTGTACAAGAAAGCTGGGTTTATATGGTTTTGACCAGCCTCATCG |
| 0_hypothetical | B8 | Tb927.10.9025_for | GGGGACAAGTTTGTACAAAAAAGCAGGCTATGATTTACGACATTACCACTCCGC | Tb927.10.9025_rev | GGGGACCACTTTGTACAAGAAAGCTGGGTCTACAGCCCAGTACTTGCAGC |
| 0_hypothetical | B9 | Tb927.9.4895_for | GGGGACAAGTTTGTACAAAAAAGCAGGCTATGAAGCAGGTCTGTGCGC | Tb927.9.4895_rev | GGGGACCACTTTGTACAAGAAAGCTGGGTTCATCCTCTGAAGATTGAACAAATTGTGC |
| 0_hypothetical | B10 | Tb927.11.14705_for | GGGGACAAGTTTGTACAAAAAAGCAGGCTATGGCAGGGGCGACTCTAG | Tb927.11.14705_rev | GGGGACCACTTTGTACAAGAAAGCTGGGTTCACTGTATCATTGACTGTGGTCGA |
| 0_hypothetical | B11 | Tb927.6.4010_for | GGGGACAAGTTTGTACAAAAAAGCAGGCTATGTCCACACGTCAAGGTGGT | Tb927.6.4010_rev | GGGGACCACTTTGTACAAGAAAGCTGGGTTTATTTCTTTCCGCCCTTCATTTTCGC |
| 0_hypothetical | B12 | Tb927.1.3850_for | GGGGACAAGTTTGTACAAAAAAGCAGGCTATGAGATCTGGGAGGAAGGCTG | Tb927.1.3850_rev | GGGGACCACTTTGTACAAGAAAGCTGGGTTTAAGCACTAATGGAGCTGAGCTTG |
| 0_hypothetical | B13 | Tb927.2.4785_for | GGGGACAAGTTTGTACAAAAAAGCAGGCTATGGCCTCTTGGACAACCTACC | Tb927.2.4785_rev | GGGGACCACTTTGTACAAGAAAGCTGGGTTTACGACCACTTTTTGCCCAACT |
| 0_hypothetical | B14 | Tb927.11.2235_for | GGGGACAAGTTTGTACAAAAAAGCAGGCTATGTTCCCCGGCATGCAGC | Tb927.11.2235_rev | GGGGACCACTTTGTACAAGAAAGCTGGGTTTAAATGCCGTTAAGTAATACAGCGCT |
| 0_hypothetical | B15 | Tb927.11.161_for | GGGGACAAGTTTGTACAAAAAAGCAGGCTATGTTTACTCCATCGCTGGCG | Tb927.11.161_rev | GGGGACCACTTTGTACAAGAAAGCTGGGTTTACGATTTCATCATGCGCCGG |
| 0_hypothetical | B16 | Tb927.9.3290_for | GGGGACAAGTTTGTACAAAAAAGCAGGCTATGTCGCGCGGCAACCAA | Tb927.9.3290_rev | GGGGACCACTTTGTACAAGAAAGCTGGGTTCAACAACCAGTCTTCCGCG |
| 0_hypothetical | B17 | Tb927.2.960_for | GGGGACAAGTTTGTACAAAAAAGCAGGCTATGGTTAGAGAAACTTTGATTTGCCTTGA | Tb927.2.960_rev | GGGGACCACTTTGTACAAGAAAGCTGGGTTCAAAGCGCTGCCCAATTGG |
| 0_hypothetical | B18 | Tb927.9.770_for | GGGGACAAGTTTGTACAAAAAAGCAGGCTATGTTCTTTGGTGGGGAAGGAAAG | Tb927.9.770_rev | GGGGACCACTTTGTACAAGAAAGCTGGGTTTACCCTTCTGTAACTTTCCCTTCCC |
| 0_hypothetical | B19 | Tb927.7.7475_for | GGGGACAAGTTTGTACAAAAAAGCAGGCTATGTGCAAGCACCTCGGC | Tb927.7.7475_rev | GGGGACCACTTTGTACAAGAAAGCTGGGTTCACCCAATCACACTCGCTAATTC |
| 0_hypothetical | B20 | Tb927.9.9115_for | GGGGACAAGTTTGTACAAAAAAGCAGGCTATGCACGCGGAGTGGGAC | Tb927.9.9115_rev | GGGGACCACTTTGTACAAGAAAGCTGGGTCTAGTCTTCCTTTCGGCCAATTGC |
| 0_hypothetical | B21 | Tb927.4.170_for | GGGGACAAGTTTGTACAAAAAAGCAGGCTATGGTTAGAGAAACTTTGATTTGCTTTGAC | Tb927.4.170_rev | GGGGACCACTTTGTACAAGAAAGCTGGGTTCAAAGCGCTGTCCAAATGGT |
| 0_hypothetical | B22 | Tb927.7.6852_for | GGGGACAAGTTTGTACAAAAAAGCAGGCTATGGAAATCCAAAACTTTCCTTTCTCCC | Tb927.7.6852_rev | GGGGACCACTTTGTACAAGAAAGCTGGGTCTATACTTTGTGAATCATAAATTTGGGTGA |
| 0_hypothetical | B23 | Tb927.11.13785_for | GGGGACAAGTTTGTACAAAAAAGCAGGCTATGCACTTATGCCGGGGCG | Tb927.11.13785_rev | GGGGACCACTTTGTACAAGAAAGCTGGGTTCACATCTGGCAACGAGTATTACG |
| 0_hypothetical | B24 | Tb927.10.4340_for | GGGGACAAGTTTGTACAAAAAAGCAGGCTATGGGGCACTTCGGGGGA | Tb927.10.4340_rev | GGGGACCACTTTGTACAAGAAAGCTGGGTTCATGTTTTTTGATGGCGATGTATAGATGA |
| 0_hypothetical | C1 | Tb927.10.8585_for | GGGGACAAGTTTGTACAAAAAAGCAGGCTATGGAATATAGGATTAAAGTGCGGAACG | Tb927.10.8585_rev | GGGGACCACTTTGTACAAGAAAGCTGGGTTCAATTACCAACCGAATTGCTGTACT |
| 0_hypothetical | C2 | Tb927.11.15255_for | GGGGACAAGTTTGTACAAAAAAGCAGGCTATGAGCATGCGTATGACCCCT | Tb927.11.15255_rev | GGGGACCACTTTGTACAAGAAAGCTGGGTTTAAATATTGTTGTTACTTGCTTCCCTAAT |
| 0_hypothetical | C3 | Tb927.11.12045_for | GGGGACAAGTTTGTACAAAAAAGCAGGCTATGGATCCTCGACGAGGGGG | Tb927.11.12045_rev | GGGGACCACTTTGTACAAGAAAGCTGGGTTTACTTTGTTCCTGAAGGCGCTG |
| 0_hypothetical | C4 | Tb927.10.4350_for | GGGGACAAGTTTGTACAAAAAAGCAGGCTATGGTAGACAAAGCAAAAACTCGAGA | Tb927.10.4350_rev | GGGGACCACTTTGTACAAGAAAGCTGGGTTCAGTCGTGCTTTTTTGAGCATGT |
| 0_hypothetical | C5 | Tb927.10.4125_for | GGGGACAAGTTTGTACAAAAAAGCAGGCTATGGTAATGTTTTTGGCCACTGCC | Tb927.10.4125_rev | GGGGACCACTTTGTACAAGAAAGCTGGGTTCACGTCCAACCAGCCGA |
| 0_hypothetical | C6 | Tb927.7.1025_for | GGGGACAAGTTTGTACAAAAAAGCAGGCTATGGATTATCAGTTCTTGTGGGATGTG | Tb927.7.1025_rev | GGGGACCACTTTGTACAAGAAAGCTGGGTTTAGTCTCCCTTGGTACGGGTTTT |
| 0_hypothetical | C7 | Tb927.7.2985_for | GGGGACAAGTTTGTACAAAAAAGCAGGCTATGATGTACGTGCTTGTAGCCG | Tb927.7.2985_rev | GGGGACCACTTTGTACAAGAAAGCTGGGTTTATTCCTCCTTCCGAGCACGA |
| 0_hypothetical | C9 | Tb927.11.2245_for | GGGGACAAGTTTGTACAAAAAAGCAGGCTATGCTTCTAGGAGGTTTTGTTCCCC | Tb927.11.2245_rev | GGGGACCACTTTGTACAAGAAAGCTGGGTTTAGTGATGGTCGTCATGGTGGG |
| 0_hypothetical | C10 | Tb927.3.4005_for | GGGGACAAGTTTGTACAAAAAAGCAGGCTATGGAAGGGAAACGGGACTGC | Tb927.3.4005_rev | GGGGACCACTTTGTACAAGAAAGCTGGGTCTAACGACCGCGCTTCCC |
| 0_hypothetical | C11 | Tb927.10.1615_for | GGGGACAAGTTTGTACAAAAAAGCAGGCTATGCCGACGGGACACGTG | Tb927.10.1615_rev | GGGGACCACTTTGTACAAGAAAGCTGGGTTCACGCTGTCTCCGCCAA |
| 0_hypothetical | C12 | Tb927.3.650_for | GGGGACAAGTTTGTACAAAAAAGCAGGCTATGATGAACCACCAGCGGCC | Tb927.3.650_rev | GGGGACCACTTTGTACAAGAAAGCTGGGTTTACAAGTCGTCATCGTAGTTGGGT |
| 0_hypothetical | C13 | Tb927.8.4175_for | GGGGACAAGTTTGTACAAAAAAGCAGGCTATGGGTTTGGAGAATGTGCCTACG | Tb927.8.4175_rev | GGGGACCACTTTGTACAAGAAAGCTGGGTCTAATAGCGGACCCTACACGTATCC |
| 0_hypothetical | C14 | Tb927.10.7985_for | GGGGACAAGTTTGTACAAAAAAGCAGGCTATGTTTTCTCTGACCCAAACCTGG | Tb927.10.7985_rev | GGGGACCACTTTGTACAAGAAAGCTGGGTTCACGGCATGTCACGGTACG |
| 0_hypothetical | C15 | Tb927.10.9445_for | GGGGACAAGTTTGTACAAAAAAGCAGGCTATGAAAAAAGGCAACCGTCGCG | Tb927.10.9445_rev | GGGGACCACTTTGTACAAGAAAGCTGGGTCTATAATCCGCATGCCTTCTCGG |
| 0_hypothetical | C16 | Tb927.10.8350_for | GGGGACAAGTTTGTACAAAAAAGCAGGCTATGACATCCTTGGGGCTGCT | Tb927.10.8350_rev | GGGGACCACTTTGTACAAGAAAGCTGGGTCTACCAGTCGGGGGCGCC |
| 0_hypothetical | C17 | Tb927.11.15665_for | GGGGACAAGTTTGTACAAAAAAGCAGGCTATGCTCAGGATCAAAATCGCCG | Tb927.11.15665_rev | GGGGACCACTTTGTACAAGAAAGCTGGGTTTATGTTGAAAACTTTTGCGGGTCG |
| 0_hypothetical | C18 | Tb927.3.2476_for | GGGGACAAGTTTGTACAAAAAAGCAGGCTATGCCTGAGACGCCACAGC | Tb927.3.2476_rev | GGGGACCACTTTGTACAAGAAAGCTGGGTTTACGAGAGCGCTGCCTCA |
| 0_hypothetical | C19 | Tb927.7.4641_for | GGGGACAAGTTTGTACAAAAAAGCAGGCTATGAAAAGAGCCGCACCGTC | Tb927.7.4641_rev | GGGGACCACTTTGTACAAGAAAGCTGGGTTCATTTGCATCGAAGCTCCTCTTG |
| 0_hypothetical | C20 | Tb927.10.8115_for | GGGGACAAGTTTGTACAAAAAAGCAGGCTATGTCAGAAGCCGTCGATAGCT | Tb927.10.8115_rev | GGGGACCACTTTGTACAAGAAAGCTGGGTCTATGATAGAGGGTGCCTGATTGATATGG |
| 0_hypothetical | C21 | Tb927.8.4695_for | GGGGACAAGTTTGTACAAAAAAGCAGGCTATGCGGTTTTCTCGGAGGTTATTT | Tb927.8.4695_rev | GGGGACCACTTTGTACAAGAAAGCTGGGTTCACTTCGCCGGCCCGCC |
| 0_hypothetical | C22 | Tb927.11.16575_for | GGGGACAAGTTTGTACAAAAAAGCAGGCTATGCCCGCCCTTACTATCGC | Tb927.11.16575_rev | GGGGACCACTTTGTACAAGAAAGCTGGGTTTACTCCTCGTCGGCTTCCG |
| 0_hypothetical | C23 | Tb927.2.3035_for | GGGGACAAGTTTGTACAAAAAAGCAGGCTATGTTGGAGTCTGACACGCAGT | Tb927.2.3035_rev | GGGGACCACTTTGTACAAGAAAGCTGGGTTTAAGAAGTGGAAACCCACATAAATGCC |
| 0_hypothetical | C24 | Tb927.9.6890_for | GGGGACAAGTTTGTACAAAAAAGCAGGCTATGGGCCCACGCATGGCC | Tb927.9.6890_rev | GGGGACCACTTTGTACAAGAAAGCTGGGTTTATTCATCACCTCCCTGACATTCCA |
| 0_hypothetical | D1 | Tb927.11.8185_for | GGGGACAAGTTTGTACAAAAAAGCAGGCTATGTCATCAGTAGGAAGTAACCGTAATTC | Tb927.11.8185_rev | GGGGACCACTTTGTACAAGAAAGCTGGGTCTAGCTAATGCGCTTCACGCC |
| 0_hypothetical | D2 | Tb927.7.7350_for | GGGGACAAGTTTGTACAAAAAAGCAGGCTATGATGTCCCTCATGCTTGGACC | Tb927.7.7350_rev | GGGGACCACTTTGTACAAGAAAGCTGGGTTCATCTGACTGCTCCGCAGC |
| 0_hypothetical | D3 | Tb927.11.15200_for | GGGGACAAGTTTGTACAAAAAAGCAGGCTATGTCTGACAATGCTTCACCTCCT | Tb927.11.15200_rev | GGGGACCACTTTGTACAAGAAAGCTGGGTTCATCTCATCTCCTCTCGTGGATTG |
| 0_hypothetical | D4 | Tb927.11.8130_for | GGGGACAAGTTTGTACAAAAAAGCAGGCTATGGACTTTTTATTCGGCACCATGT | Tb927.11.8130_rev | GGGGACCACTTTGTACAAGAAAGCTGGGTCTATTCTTTTTTGATGTTTCGTGTTGGTAT |
| 0_hypothetical | D5 | Tb927.2.1895_for | GGGGACAAGTTTGTACAAAAAAGCAGGCTATGTTCAACCCCAAGAAGATGTTGG | Tb927.2.1895_rev | GGGGACCACTTTGTACAAGAAAGCTGGGTTTACTTCGCTCCACCGGCA |

| | | | | | |
|---|---|---|---|---|---|
| 0_hypothetical | D6 | Tb927.11.12425_for | GGGGACAAGTTTGTACAAAAAAGCAGGCTATGGCAGGCAGTTCTGAATATTCC | Tb927.11.12425_rev | GGGGACCACTTTGTACAAGAAAGCTGGGTCTAGACAACTTTAAAGGCGTTTGTCCC |
| 0_hypothetical | D7 | Tb927.11.485_for | GGGGACAAGTTTGTACAAAAAAGCAGGCTATGGAAAATATTACAAATGTCTCCATTGCC | Tb927.11.485_rev | GGGGACCACTTTGTACAAGAAAGCTGGGTCTACAGATTTGATACGCCGCTGT |
| 0_hypothetical | D8 | Tb927.10.8375_for | GGGGACAAGTTTGTACAAAAAAGCAGGCTATGAGGGCTCTTTTGCTCTTTTTTGC | Tb927.10.8375_rev | GGGGACCACTTTGTACAAGAAAGCTGGGTTCACCTGTCAGGATGACCCAG |
| 0_hypothetical | D9 | Tb927.6.5095_for | GGGGACAAGTTTGTACAAAAAAGCAGGCTATGATTCGACGGACAGTAGTGACG | Tb927.6.5095_rev | GGGGACCACTTTGTACAAGAAAGCTGGGTTTAGTGGTGTGATTGGACAGCG |
| 0_hypothetical | D10 | Tb927.11.17015_for | GGGGACAAGTTTGTACAAAAAAGCAGGCTATGCAAAACGCACACAAGAGAGA | Tb927.11.17015_rev | GGGGACCACTTTGTACAAGAAAGCTGGGTCTACTCCGAAGTATTGTTATGGCCC |
| 0_hypothetical | D11 | Tb927.11.14595_for | GGGGACAAGTTTGTACAAAAAAGCAGGCTATGGTGCTTGCGTTAACATCACC | Tb927.11.14595_rev | GGGGACCACTTTGTACAAGAAAGCTGGGTTCAGCGTGCCCACCACCG |
| 0_hypothetical | D12 | Tb927.8.7255_for | GGGGACAAGTTTGTACAAAAAAGCAGGCTATGACGCATAAGGTCCGCCT | Tb927.8.7255_rev | GGGGACCACTTTGTACAAGAAAGCTGGGTTCAATCAGCGAAAAACATCAATGCTT |
| 0_hypothetical | D13 | Tb927.10.1780_for | GGGGACAAGTTTGTACAAAAAAGCAGGCTATGCATCCACTTGCTCGCC | Tb927.10.1780_rev | GGGGACCACTTTGTACAAGAAAGCTGGGTTCAGTAGACCCGCGTTGTTCC |
| 0_hypothetical | D14 | Tb927.10.4140_for | GGGGACAAGTTTGTACAAAAAAGCAGGCTATGAGCGTATTCTTCGACATCGAG | Tb927.10.4140_rev | GGGGACCACTTTGTACAAGAAAGCTGGGTTCAATACACAAACAGCATTCGCAC |
| 0_hypothetical | D15 | Tb927.5.2560_for | GGGGACAAGTTTGTACAAAAAAGCAGGCTATGTCGAATCCTAACTTCTGGACAAC | Tb927.5.2560_rev | GGGGACCACTTTGTACAAGAAAGCTGGGTTTATACCGCTTCCTCTTCCAGCA |
| 0_hypothetical | D16 | Tb927.5.2325_for | GGGGACAAGTTTGTACAAAAAAGCAGGCTATGAGTGGCATTCGAGTGGCG | Tb927.5.2325_rev | GGGGACCACTTTGTACAAGAAAGCTGGGTTCATCCGTTCCTGGCATCCC |
| 0_hypothetical | D17 | Tb927.6.250_for | GGGGACAAGTTTGTACAAAAAAGCAGGCTATGAAAAGAATGAAATACACATTTCGATGT | Tb927.6.250_rev | GGGGACCACTTTGTACAAGAAAGCTGGGTTTAGTTACGGCAAGGAATTGCTGAA |
| 0_hypothetical | D18 | Tb927.10.6415_for | GGGGACAAGTTTGTACAAAAAAGCAGGCTATGCTGGGTCGTCGGTTCT | Tb927.10.6415_rev | GGGGACCACTTTGTACAAGAAAGCTGGGTTCAGCCATTTAGACAGTGGGGA |
| 0_hypothetical | D19 | Tb927.1.2780_for | GGGGACAAGTTTGTACAAAAAAGCAGGCTATGATACCAGTTCATGCGCTGAG | Tb927.1.2780_rev | GGGGACCACTTTGTACAAGAAAGCTGGGTCTAGCGACCCACACTGATTTCC |
| 0_hypothetical | D20 | Tb927.3.3010_for | GGGGACAAGTTTGTACAAAAAAGCAGGCTATGCGGCTTCTTGCACGAC | Tb927.3.3010_rev | GGGGACCACTTTGTACAAGAAAGCTGGGTTCAGTGTGAACTCTCCTTGTCACG |
| 0_hypothetical | D21 | Tb927.1.1175_for | GGGGACAAGTTTGTACAAAAAAGCAGGCTATGATGAACAGGCCGCCGC | Tb927.1.1175_rev | GGGGACCACTTTGTACAAGAAAGCTGGGTTCAGTTTGTACTATGCTCTGTTGGGA |
| 0_hypothetical | D22 | Tb927.3.3240_for | GGGGACAAGTTTGTACAAAAAAGCAGGCTATGAAAAAAAGCGTTTCCTTGAAGGAG | Tb927.3.3240_rev | GGGGACCACTTTGTACAAGAAAGCTGGGTTCACTCCATGTCATACATGACATCCA |
| 0_hypothetical | D23 | Tb927.11.7045_for | GGGGACAAGTTTGTACAAAAAAGCAGGCTATGCATGTGGAACCCAGAGAAAC | Tb927.11.7045_rev | GGGGACCACTTTGTACAAGAAAGCTGGGTTCATCGCCGTCTGCCTTTACT |
| 0_hypothetical | D24 | Tb927.3.630_for | GGGGACAAGTTTGTACAAAAAAGCAGGCTATGTCGATTTCACCCGCAGAGA | Tb927.3.630_rev | GGGGACCACTTTGTACAAGAAAGCTGGGTCTACGCATCCTTTTTTCCCCGC |
| 0_hypothetical | E1 | Tb927.10.5050_for | GGGGACAAGTTTGTACAAAAAAGCAGGCTATGATCCGCCGTAGTTGTGC | Tb927.10.5050_rev | GGGGACCACTTTGTACAAGAAAGCTGGGTTCAGTAATCCTTGGTGTGGATGGG |
| 0_hypothetical | E2 | Tb927.9.1365_for | GGGGACAAGTTTGTACAAAAAAGCAGGCTATGGAAGCTGATGCCAATACATTTAACC | Tb927.9.1365_rev | GGGGACCACTTTGTACAAGAAAGCTGGGTTCAAACGCCTGTCCGTCGC |
| 0_hypothetical | E3 | Tb927.9.8655_for | GGGGACAAGTTTGTACAAAAAAGCAGGCTATGTGTACGTGTGACGCAGAGG | Tb927.9.8655_rev | GGGGACCACTTTGTACAAGAAAGCTGGGTTTACTCCCCTACGGGGGCT |
| 0_hypothetical | E4 | Tb927.11.165_for | GGGGACAAGTTTGTACAAAAAAGCAGGCTATGCCTTTAAGCCCTACAAGGATGA | Tb927.11.165_rev | GGGGACCACTTTGTACAAGAAAGCTGGGTTCACTCAACCACACCGCCC |
| 0_hypothetical | E5 | Tb927.1.2000_for | GGGGACAAGTTTGTACAAAAAAGCAGGCTATGCGAATGTTTTGCGTCGTC | Tb927.1.2000_rev | GGGGACCACTTTGTACAAGAAAGCTGGGTTCAGGTGGTGATACCACCATACG |
| 0_hypothetical | E6 | Tb927.9.3010_for | GGGGACAAGTTTGTACAAAAAAGCAGGCTATGAGTTCTAAGAGCAGTGTGGGA | Tb927.9.3010_rev | GGGGACCACTTTGTACAAGAAAGCTGGGTCTATGATCCAGCGGCCGAAGT |
| 0_hypothetical | E7 | Tb927.11.3040_for | GGGGACAAGTTTGTACAAAAAAGCAGGCTATGGGACGTAGGCTGCGG | Tb927.11.3040_rev | GGGGACCACTTTGTACAAGAAAGCTGGGTTTAATGCAGAAACCGCATGCGA |
| 0_hypothetical | E8 | Tb927.3.5655_for | GGGGACAAGTTTGTACAAAAAAGCAGGCTATGCTAGTTAGCGCCAGCGA | Tb927.3.5655_rev | GGGGACCACTTTGTACAAGAAAGCTGGGTTCACCTGAAGATGACGGTACATTTTTT |
| 0_hypothetical | E9 | Tb927.10.260_for | GGGGACAAGTTTGTACAAAAAAGCAGGCTATGGTCACTCAAACTGAAATGGAGG | Tb927.10.260_rev | GGGGACCACTTTGTACAAGAAAGCTGGGTCTACGGTAATGGCTTCGCCG |
| 0_hypothetical | E10 | Tb927.10.11585_for | GGGGACAAGTTTGTACAAAAAAGCAGGCTATGGAGTGTGTATTTGTCTGTAGAAGGA | Tb927.10.11585_rev | GGGGACCACTTTGTACAAGAAAGCTGGGTTCACATGTGTGTTTGTTTGTCTGTTTG |
| 0_hypothetical | E11 | Tb927.11.7260_for | GGGGACAAGTTTGTACAAAAAAGCAGGCTATGAGACAACCCTTTCCGACTACT | Tb927.11.7260_rev | GGGGACCACTTTGTACAAGAAAGCTGGGTTTATGACGAGTCTGCTTTTGCGT |
| 0_hypothetical | E12 | Tb927.7.4010_for | GGGGACAAGTTTGTACAAAAAAGCAGGCTATGAGTAACACACGCCAGCG | Tb927.7.4010_rev | GGGGACCACTTTGTACAAGAAAGCTGGGTTCATTCACTCCGTTGACGCTTC |
| 0_hypothetical | E13 | Tb927.2.1290_for | GGGGACAAGTTTGTACAAAAAAGCAGGCTATGTTCCCTTTCGTGTTCCCTTTT | Tb927.2.1290_rev | GGGGACCACTTTGTACAAGAAAGCTGGGTTTAGAATGTATTGCCATTTTTTTCGTTGTT |
| 0_hypothetical | E14 | Tb927.4.1770_for | GGGGACAAGTTTGTACAAAAAAGCAGGCTATGTTTCGGGTTTCCAGACGG | Tb927.4.1770_rev | GGGGACCACTTTGTACAAGAAAGCTGGGTTTAGGTCTTCCATCCGGCATTCT |
| 0_hypothetical | E15 | Tb927.7.2200_for | GGGGACAAGTTTGTACAAAAAAGCAGGCTATGCGCCTGGCTGTAAAGC | Tb927.7.2200_rev | GGGGACCACTTTGTACAAGAAAGCTGGGTCTACAACTTCAACATTTGCTTCACAGC |
| 0_hypothetical | E16 | Tb927.10.4090_for | GGGGACAAGTTTGTACAAAAAAGCAGGCTATGTTTTTGCGACCTATCGCGC | Tb927.10.4090_rev | GGGGACCACTTTGTACAAGAAAGCTGGGTTCAATCGTATACTTTGAGCGCAACT |
| 0_hypothetical | E17 | Tb927.10.7205_for | GGGGACAAGTTTGTACAAAAAAGCAGGCTATGTCTAATGCAACTCCCGTTACTG | Tb927.10.7205_rev | GGGGACCACTTTGTACAAGAAAGCTGGGTCTAAACTGGACCTCTGGGATGATTCC |
| 0_hypothetical | E18 | Tb927.1.305_for | GGGGACAAGTTTGTACAAAAAAGCAGGCTATGTATGAAGGTTTCTGTGCAAGAAGC | Tb927.1.305_rev | GGGGACCACTTTGTACAAGAAAGCTGGGTTCAAACATAAATATTACCCATCAACAGTTG |
| 0_hypothetical | E19 | Tb927.10.4960_for | GGGGACAAGTTTGTACAAAAAAGCAGGCTATGGGGTACGACAGACGATCG | Tb927.10.4960_rev | GGGGACCACTTTGTACAAGAAAGCTGGGTTTACGGCGGCGCCCGCCG |
| 0_hypothetical | E21 | Tb927.11.10075_for | GGGGACAAGTTTGTACAAAAAAGCAGGCTATGACTGACGTTAACAGATATCTTCAAAGT | Tb927.11.10075_rev | GGGGACCACTTTGTACAAGAAAGCTGGGTTTACTGAAAGGGCAAACCACACG |
| 0_hypothetical | E22 | Tb927.10.7375_for | GGGGACAAGTTTGTACAAAAAAGCAGGCTATGCTAGGTATTACGCCGCAAC | Tb927.10.7375_rev | GGGGACCACTTTGTACAAGAAAGCTGGGTCTATAGGGGTGTAATACCCCACTCTG |
| 0_hypothetical | E23 | Tb927.4.2330_for | GGGGACAAGTTTGTACAAAAAAGCAGGCTATGTTTAGGAGGACATTCTTTACACCAAT | Tb927.4.2330_rev | GGGGACCACTTTGTACAAGAAAGCTGGGTCTACGGTTTAATTTGACGGCAGTCC |
| 0_hypothetical | E24 | Tb927.8.2391_for | GGGGACAAGTTTGTACAAAAAAGCAGGCTATGAAGTACCGACAATGTTCTCTGAC | Tb927.8.2391_rev | GGGGACCACTTTGTACAAGAAAGCTGGGTTTATAGGCCCACATTATGAAGGGAAGG |
| 0_hypothetical | F1 | Tb927.8.595_for | GGGGACAAGTTTGTACAAAAAAGCAGGCTATGGGACATTCAGCAAAAATTACTAGAGG | Tb927.8.595_rev | GGGGACCACTTTGTACAAGAAAGCTGGGTCTACGCCTCTGGTCCCTTTACC |
| 0_hypothetical | F2 | Tb927.9.2980_for | GGGGACAAGTTTGTACAAAAAAGCAGGCTATGAGTTTTGGCCTTGAATACAGCG | Tb927.9.2980_rev | GGGGACCACTTTGTACAAGAAAGCTGGGTTTAGGCAGTTGCGGTTAACCG |
| 0_hypothetical | F3 | Tb927.7.2105_for | GGGGACAAGTTTGTACAAAAAAGCAGGCTATGATGTCTAGGCGGATTTGTATTCG | Tb927.7.2105_rev | GGGGACCACTTTGTACAAGAAAGCTGGGTTCATCGCTTAACAAAGACGCGG |
| 0_hypothetical | F4 | Tb927.1.490_for | GGGGACAAGTTTGTACAAAAAAGCAGGCTATGCAATGTGCTTTGAGTGGGG | Tb927.1.490_rev | GGGGACCACTTTGTACAAGAAAGCTGGGTTTAATGTTGGACGCTGATGTGAGG |
| 0_hypothetical | F5 | Tb927.4.180_for | GGGGACAAGTTTGTACAAAAAAGCAGGCTATGCAAGGTGCTTTGAGTGGG | Tb927.4.180_rev | GGGGACCACTTTGTACAAGAAAGCTGGGTTTAATGTTGGACGCTGATGTGAGG |
| 0_hypothetical | F6 | Tb927.11.12365_for | GGGGACAAGTTTGTACAAAAAAGCAGGCTATGGAGGGTAATTTCTACGTTTCTGAC | Tb927.11.12365_rev | GGGGACCACTTTGTACAAGAAAGCTGGGTCTACTGTTTCGACCCCGAATCAC |
| 0_hypothetical | F7 | Tb927.9.1390_for | GGGGACAAGTTTGTACAAAAAAGCAGGCTATGCACGCCGCACCTTATG | Tb927.9.1390_rev | GGGGACCACTTTGTACAAGAAAGCTGGGTTCAAGCAATCACAACATCACGCA |
| 0_hypothetical | F8 | Tb927.11.5390_for | GGGGACAAGTTTGTACAAAAAAGCAGGCTATGAATCAGTCTAGTTCATTATGGGGCG | Tb927.11.5390_rev | GGGGACCACTTTGTACAAGAAAGCTGGGTCTACTTTCCTTTGCTTCCTGTCTTACC |
| 0_hypothetical | F9 | Tb927.10.1290_for | GGGGACAAGTTTGTACAAAAAAGCAGGCTATGTGGCGCGAAAAGTGGG | Tb927.10.1290_rev | GGGGACCACTTTGTACAAGAAAGCTGGGTTTAGCGCCTCAACAAACGTTTTG |
| 0_hypothetical | F10 | Tb927.7.4685_for | GGGGACAAGTTTGTACAAAAAAGCAGGCTATGAAGGGACGAGGCGGAG | Tb927.7.4685_rev | GGGGACCACTTTGTACAAGAAAGCTGGGTTCACTTTGTCGCTTCGTTTCGG |
| 0_hypothetical | F11 | Tb927.7.7495_for | GGGGACAAGTTTGTACAAAAAAGCAGGCTATGCTGCAGAACCAGCAGTC | Tb927.7.7495_rev | GGGGACCACTTTGTACAAGAAAGCTGGGTTTACGCAAACCGCCGCAG |
| 0_hypothetical | F12 | Tb927.1.1025_for | GGGGACAAGTTTGTACAAAAAAGCAGGCTATGTTTCGCTCGTCGATGGTAG | Tb927.1.1025_rev | GGGGACCACTTTGTACAAGAAAGCTGGGTCTAAAGAACAAGTGATCGACCATCGT |
| 0_hypothetical | F13 | Tb927.9.6440_for | GGGGACAAGTTTGTACAAAAAAGCAGGCTATGTTATACGACCGCAGCATAGC | Tb927.9.6440_rev | GGGGACCACTTTGTACAAGAAAGCTGGGTTCACCCGCTTTCCTCCGG |
| 0_hypothetical | F14 | Tb927.8.2185_for | GGGGACAAGTTTGTACAAAAAAGCAGGCTATGACTCGAATACCTGGGGTCATC | Tb927.8.2185_rev | GGGGACCACTTTGTACAAGAAAGCTGGGTTCACCGAAACATCCTGTCGTCG |
| 0_hypothetical | F15 | Tb927.9.9100_for | GGGGACAAGTTTGTACAAAAAAGCAGGCTATGTCATACCAATCCAGCGAAGC | Tb927.9.9100_rev | GGGGACCACTTTGTACAAGAAAGCTGGGTTTAGTGCTCTTCATTCAGCCGTTT |
| 0_hypothetical | F16 | Tb927.4.5210_for | GGGGACAAGTTTGTACAAAAAAGCAGGCTATGAGTGACTCTGTTCGCAAGAAG | Tb927.4.5210_rev | GGGGACCACTTTGTACAAGAAAGCTGGGTTCACTCTTCCTTTTTCAGAAGCTGC |
| 0_hypothetical | F17 | Tb927.3.700_for | GGGGACAAGTTTGTACAAAAAAGCAGGCTATGTTACGCCGTAGCGTACCG | Tb927.3.700_rev | GGGGACCACTTTGTACAAGAAAGCTGGGTTTATGGATGAAGCGAATGCGGATC |
| 0_hypothetical | F18 | Tb927.7.6465_for | GGGGACAAGTTTGTACAAAAAAGCAGGCTATGTGGAGAATATCCAATGTGCTCCG | Tb927.7.6465_rev | GGGGACCACTTTGTACAAGAAAGCTGGGTTCATCCTTTGTCGCACATGTTTAGC |
| 0_hypothetical | F19 | Tb927.9.1520_for | GGGGACAAGTTTGTACAAAAAAGCAGGCTATGGCTCCCGCAGTAATTGAGA | Tb927.9.1520_rev | GGGGACCACTTTGTACAAGAAAGCTGGGTTTACTCATCAAAACAGATCTGGAAATGTGT |
| 0_hypothetical | F20 | Tb927.9.15829_for | GGGGACAAGTTTGTACAAAAAAGCAGGCTATGAATTCCTCTTGTGTAAATTCAGTTATG | Tb927.9.15829_rev | GGGGACCACTTTGTACAAGAAAGCTGGGTTTATATATTTGTTACTTGGCGTTTTAACAG |
| 0_hypothetical | F21 | Tb927.3.1985_for | GGGGACAAGTTTGTACAAAAAAGCAGGCTATGAGCTCACTATTGTCTGCAGC | Tb927.3.1985_rev | GGGGACCACTTTGTACAAGAAAGCTGGGTCTAGGAAATAAAGACAAGGTTGAGCAACC |
| 0_hypothetical | F22 | Tb927.4.780_for | GGGGACAAGTTTGTACAAAAAAGCAGGCTATGTATGCCCTGGACTACAGGAAT | Tb927.4.780_rev | GGGGACCACTTTGTACAAGAAAGCTGGGTTCAGTTATGCGTAAATATTTTCCAACCGA |
| 0_hypothetical | F23 | Tb927.1.2200_for | GGGGACAAGTTTGTACAAAAAAGCAGGCTATGCGTTTGTATCTACAGGGGAGA | Tb927.1.2200_rev | GGGGACCACTTTGTACAAGAAAGCTGGGTCTATTTCTCCGCCTCGAGGTTCA |
| 0_hypothetical | F24 | Tb927.9.3780_for | GGGGACAAGTTTGTACAAAAAAGCAGGCTATGGTGTCTGCAACGGCAG | Tb927.9.3780_rev | GGGGACCACTTTGTACAAGAAAGCTGGGTTCACACTGCCTTTCGGACAAG |
| 0_hypothetical | G1 | Tb927.9.3550_for | GGGGACAAGTTTGTACAAAAAAGCAGGCTATGCGCATTTTCGTACACGTACG | Tb927.9.3550_rev | GGGGACCACTTTGTACAAGAAAGCTGGGTCTACGCATCTGTGACAACCTCC |
| 0_hypothetical | G2 | Tb927.9.8815_for | GGGGACAAGTTTGTACAAAAAAGCAGGCTATGCTTCGTCGGTCGTACTCC | Tb927.9.8815_rev | GGGGACCACTTTGTACAAGAAAGCTGGGTTTAGTACTTCCACCACAGGAACGG |
| 0_hypothetical | G3 | Tb927.8.6190_for | GGGGACAAGTTTGTACAAAAAAGCAGGCTATGTTTCGCATGACTCGGCTG | Tb927.8.6190_rev | GGGGACCACTTTGTACAAGAAAGCTGGGTTCATTCCACCTTAGTCGTGAGTGA |
| 0_hypothetical | G4 | Tb927.11.155_for | GGGGACAAGTTTGTACAAAAAAGCAGGCTATGTTACGGAGGTTACACGTCCC | Tb927.11.155_rev | GGGGACCACTTTGTACAAGAAAGCTGGGTTTAGCGGCCGTCTCCACC |
| 0_hypothetical | G5 | Tb927.7.1171_for | GGGGACAAGTTTGTACAAAAAAGCAGGCTATGGCCACTACTCGGGCG | Tb927.7.1171_rev | GGGGACCACTTTGTACAAGAAAGCTGGGTTCAGATCACTAATCTGGTTGAACCGC |
| 0_hypothetical | G6 | Tb927.7.4680_for | GGGGACAAGTTTGTACAAAAAAGCAGGCTATGGGAAAGCAAAAGAAGCGGC | Tb927.7.4680_rev | GGGGACCACTTTGTACAAGAAAGCTGGGTTCAACGATTCGTCACTCGTGC |
| 0_hypothetical | G7 | Tb927.8.6890_for | GGGGACAAGTTTGTACAAAAAAGCAGGCTATGATTCGTTCACTTGGATATCGCC | Tb927.8.6890_rev | GGGGACCACTTTGTACAAGAAAGCTGGGTCTACGCCTTGAAGAATAAGGCAACA |
| 0_hypothetical | G8 | Tb927.7.7560_for | GGGGACAAGTTTGTACAAAAAAGCAGGCTATGTTCCCTTTCGTGTTCCCTTTT | Tb927.7.7560_rev | GGGGACCACTTTGTACAAGAAAGCTGGGTTCATGAGTTCTGCAATGATGAGGGA |
| 0_hypothetical | G9 | Tb927.9.10850_for | GGGGACAAGTTTGTACAAAAAAGCAGGCTATGGGCGATCCTTTTGTGCG | Tb927.9.10850_rev | GGGGACCACTTTGTACAAGAAAGCTGGGTTCAAGTATTATGCAAATCCCTCTTCCG |
| 0_hypothetical | G10 | Tb927.2.4955_for | GGGGACAAGTTTGTACAAAAAAGCAGGCTATGGACTTTATGAATACAAGCGAGACAG | Tb927.2.4955_rev | GGGGACCACTTTGTACAAGAAAGCTGGGTTCACACCTCGGTGCCCCG |
| 0_hypothetical | G11 | Tb927.10.5410_for | GGGGACAAGTTTGTACAAAAAAGCAGGCTATGAGATTGCCAAATGTAAGTTATTGCG | Tb927.10.5410_rev | GGGGACCACTTTGTACAAGAAAGCTGGGTCTAATAAGCAGGGAGAGTGACGCC |
| 0_hypothetical | G12 | Tb927.9.17760_for | GGGGACAAGTTTGTACAAAAAAGCAGGCTATGAGAAACTTTGAGGACAGGGGG | Tb927.9.17760_rev | GGGGACCACTTTGTACAAGAAAGCTGGGTTTATAAAAACACCAACGTGTGCCGA |
| 0_hypothetical | G13 | Tb927.10.7150_for | GGGGACAAGTTTGTACAAAAAAGCAGGCTATGTTTGGGCGTATGTTTGATATGTGT | Tb927.10.7150_rev | GGGGACCACTTTGTACAAGAAAGCTGGGTCTACATGCATGAGCGCCGG |
| 0_hypothetical | G14 | Tb927.2.3420_for | GGGGACAAGTTTGTACAAAAAAGCAGGCTATGGTTTCCAGGGACTCCCG | Tb927.2.3420_rev | GGGGACCACTTTGTACAAGAAAGCTGGGTTCATTCTTCTGTAAAGGGCGGGA |
| 0_hypothetical | G15 | Tb927.11.600_for | GGGGACAAGTTTGTACAAAAAAGCAGGCTATGTCGGCTAAGGCAGCTCC | Tb927.11.600_rev | GGGGACCACTTTGTACAAGAAAGCTGGGTCTACTTAGTGAAAGCTTCAGCAAGCT |
| 0_hypothetical | G16 | Tb927.11.435_for | GGGGACAAGTTTGTACAAAAAAGCAGGCTATGAATTTTGTTAAGCGTTTGGTTTACTCA | Tb927.11.435_rev | GGGGACCACTTTGTACAAGAAAGCTGGGTTTAGTTACGCGGGACGCCA |
| 0_hypothetical | G17 | Tb927.11.12960_for | GGGGACAAGTTTGTACAAAAAAGCAGGCTATGAATTCCTACTCGCGGCTCT | Tb927.11.12960_rev | GGGGACCACTTTGTACAAGAAAGCTGGGTTTAGGTTCTCGTTTGCCCAATGTATT |
| 0_hypothetical | G18 | Tb927.11.12200_for | GGGGACAAGTTTGTACAAAAAAGCAGGCTATGAGCAGTCATTTCTCCGGTAGG | Tb927.11.12200_rev | GGGGACCACTTTGTACAAGAAAGCTGGGTTCACTCCACTGGTGGTCTCTTTT |
| 0_hypothetical | G19 | Tb927.11.365_for | GGGGACAAGTTTGTACAAAAAAGCAGGCTATGACCCTTAGCAGCGATGCG | Tb927.11.365_rev | GGGGACCACTTTGTACAAGAAAGCTGGGTTTATGGCTGACAACTGCGATACC |
| 0_hypothetical | G20 | Tb927.9.8905_for | GGGGACAAGTTTGTACAAAAAAGCAGGCTATGGCGTTGTTTTCGTGCTTTC | Tb927.9.8905_rev | GGGGACCACTTTGTACAAGAAAGCTGGGTTCACCAACGATCGCTCTTGC |
| 0_hypothetical | G21 | Tb927.10.7925_for | GGGGACAAGTTTGTACAAAAAAGCAGGCTATGGGAACGAAGGAAAATGCAACA | Tb927.10.7925_rev | GGGGACCACTTTGTACAAGAAAGCTGGGTCTATAGCCGATTTCCTTTCTGCACA |
| 0_hypothetical | G22 | Tb927.11.2461_for | GGGGACAAGTTTGTACAAAAAAGCAGGCTATGGCCGTGCCTCGTTGG | Tb927.11.2461_rev | GGGGACCACTTTGTACAAGAAAGCTGGGTTCAGTCGCTGCCGGTGTT |
| 0_hypothetical | G23 | Tb927.3.1720_for | GGGGACAAGTTTGTACAAAAAAGCAGGCTATGGCTGACGTTTTGGATCGTG | Tb927.3.1720_rev | GGGGACCACTTTGTACAAGAAAGCTGGGTCTACCAGAATTTCCACCAGAATTTAGTCT |
| 0_hypothetical | G24 | Tb927.6.4580_for | GGGGACAAGTTTGTACAAAAAAGCAGGCTATGCTTCGACGGAGCATCCT | Tb927.6.4580_rev | GGGGACCACTTTGTACAAGAAAGCTGGGTTCAACTATCTTCATCAACCCCAAAAATACC |
| 0_hypothetical | H1 | Tb927.9.7282_for | GGGGACAAGTTTGTACAAAAAAGCAGGCTATGTGTGGAGGCGACGCT | Tb927.9.7282_rev | GGGGACCACTTTGTACAAGAAAGCTGGGTCTATAGATTCCTGGCTGCCTTGAAAC |
| 0_hypothetical | H2 | Tb927.11.5365_for | GGGGACAAGTTTGTACAAAAAAGCAGGCTATGTTCCGGCAGCGGCTT | Tb927.11.5365_rev | GGGGACCACTTTGTACAAGAAAGCTGGGTTTAAGCCGATATTCGCGTTTTTCCA |
| 0_hypothetical | H3 | Tb927.1.5160_for | GGGGACAAGTTTGTACAAAAAAGCAGGCTATGGCGAGTCACAACGAATATTTGG | Tb927.1.5160_rev | GGGGACCACTTTGTACAAGAAAGCTGGGTTCAAGTCAGCCCTTGTCTAACATACA |
| 0_hypothetical | H4 | Tb927.8.2470_for | GGGGACAAGTTTGTACAAAAAAGCAGGCTATGCCCATGCTTCAACCGTAC | Tb927.8.2470_rev | GGGGACCACTTTGTACAAGAAAGCTGGGTCTATGCAAAGATTTCCCCCAGAACT |
| 0_hypothetical | H5 | Tb927.11.7475_for | GGGGACAAGTTTGTACAAAAAAGCAGGCTATGTCCGTTGTTTGTGCATTCAAAG | Tb927.11.7475_rev | GGGGACCACTTTGTACAAGAAAGCTGGGTCTACTTTTTCTTGATTGGCTTCTTTTGCC |
| 0_hypothetical | H6 | Tb927.10.7300_for | GGGGACAAGTTTGTACAAAAAAGCAGGCTATGTCGACTGTGGCGGAGC | Tb927.10.7300_rev | GGGGACCACTTTGTACAAGAAAGCTGGGTTCATCGCTTTTTGCATTTCCCATC |
| 0_hypothetical | H7 | Tb927.7.1860_for | GGGGACAAGTTTGTACAAAAAAGCAGGCTATGCCTGGGCGAAGAGGG | Tb927.7.1860_rev | GGGGACCACTTTGTACAAGAAAGCTGGGTTCAGCATCCTACCACAGTACCG |
| 0_hypothetical | H9 | Tb927.6.3140_for | GGGGACAAGTTTGTACAAAAAAGCAGGCTATGTCGGAGACAATACCCGTTTCA | Tb927.6.3140_rev | GGGGACCACTTTGTACAAGAAAGCTGGGTTTACATATCGTCCTCATCCTTTTCTGGA |
| 0_hypothetical | H10 | Tb927.11.16615_for | GGGGACAAGTTTGTACAAAAAAGCAGGCTATGTTTCTCCCCACGCATTTGT | Tb927.11.16615_rev | GGGGACCACTTTGTACAAGAAAGCTGGGTTTACTTCGCCTTCCTCGCAAGA |
| 0_hypothetical | H11 | Tb927.8.5150_for | GGGGACAAGTTTGTACAAAAAAGCAGGCTATGGCGTTTCAAACACCGTGG | Tb927.8.5150_rev | GGGGACCACTTTGTACAAGAAAGCTGGGTTTACCACAGTTTCCAAATCGAGTGG |
| 0_hypothetical | H12 | Tb927.5.3340_for | GGGGACAAGTTTGTACAAAAAAGCAGGCTATGCAGAGCCAAATGATGCTCA | Tb927.5.3340_rev | GGGGACCACTTTGTACAAGAAAGCTGGGTCTACTCCATACCGAGGGATCCCG |
| 0_hypothetical | H13 | Tb927.1.800_for | GGGGACAAGTTTGTACAAAAAAGCAGGCTATGAATTCACACATCTTCCATTTCCCA | Tb927.1.800_rev | GGGGACCACTTTGTACAAGAAAGCTGGGTCTACGGCCTCAAGTGTGTAGTGG |
| 0_hypothetical | H14 | Tb927.10.13341_for | GGGGACAAGTTTGTACAAAAAAGCAGGCTATGCTTCAGGCGCAAGGTTC | Tb927.10.13341_rev | GGGGACCACTTTGTACAAGAAAGCTGGGTTTAATGAGTGTTAACATCCATTTTCTCCTG |
| 0_hypothetical | H15 | Tb927.6.2610_for | GGGGACAAGTTTGTACAAAAAAGCAGGCTATGACATCTCCCATCCTACGAGTTG | Tb927.6.2610_rev | GGGGACCACTTTGTACAAGAAAGCTGGGTTTACAAGCGAGCCCCCATATCA |
| 0_hypothetical | H16 | Tb927.11.665_for | GGGGACAAGTTTGTACAAAAAAGCAGGCTATGCTGCGTCGATCTCTCATCG | Tb927.11.665_rev | GGGGACCACTTTGTACAAGAAAGCTGGGTCTAATACCACCGCCACCGTTC |
| 0_hypothetical | H17 | Tb927.7.1520_for | GGGGACAAGTTTGTACAAAAAAGCAGGCTATGGGTGGGGATGACCAAATATTTC | Tb927.7.1520_rev | GGGGACCACTTTGTACAAGAAAGCTGGGTCTAACGCTGGCACTTCGTGC |
| 0_hypothetical | H18 | Tb927.6.3910_for | GGGGACAAGTTTGTACAAAAAAGCAGGCTATGTCGTTTGATATGGAAGTACTGCG | Tb927.6.3910_rev | GGGGACCACTTTGTACAAGAAAGCTGGGTTCAAAGAAGGTAAGGTGCATTTGATAATAT |
| 0_hypothetical | H19 | Tb927.4.830_for | GGGGACAAGTTTGTACAAAAAAGCAGGCTATGGAAAAGGAGAACAAAAGCATCTGT | Tb927.4.830_rev | GGGGACCACTTTGTACAAGAAAGCTGGGTTTACGACACAATGCCTTCCTCCT |
| 0_hypothetical | H20 | Tb927.11.9295_for | GGGGACAAGTTTGTACAAAAAAGCAGGCTATGGGAGATTCTGAAGCCACTGC | Tb927.11.9295_rev | GGGGACCACTTTGTACAAGAAAGCTGGGTTCATGTGTGGACGTCAAAAGACAG |
| 0_hypothetical | H21 | Tb927.11.14495_for | GGGGACAAGTTTGTACAAAAAAGCAGGCTATGCAACATGACAATGTTGGAGACC | Tb927.11.14495_rev | GGGGACCACTTTGTACAAGAAAGCTGGGTTCATGTGATTAGTGCGCTCTTACG |
| 0_hypothetical | H22 | Tb927.5.3895_for | GGGGACAAGTTTGTACAAAAAAGCAGGCTATGCAAAGCAGTTGTATGCGC | Tb927.5.3895_rev | GGGGACCACTTTGTACAAGAAAGCTGGGTTCAAAATGTCACAACTTCATCAGAATCTTC |
| 0_hypothetical | H23 | Tb927.7.4515_for | GGGGACAAGTTTGTACAAAAAAGCAGGCTATGGAACAGCCGCAGCAAC | Tb927.7.4515_rev | GGGGACCACTTTGTACAAGAAAGCTGGGTCTATGGTTGAACGGGTTTTGGCA |
| 0_hypothetical | H24 | Tb927.10.10640_for | GGGGACAAGTTTGTACAAAAAAGCAGGCTATGCCTCGTGCATATTACAAACCC | Tb927.10.10640_rev | GGGGACCACTTTGTACAAGAAAGCTGGGTTCAAGCAGGAGGGGCCCG |
| 0_hypothetical | I1 | Tb927.10.8291_for | GGGGACAAGTTTGTACAAAAAAGCAGGCTATGCATCTTGATGAAATGTTGAAAGAGATT | Tb927.10.8291_rev | GGGGACCACTTTGTACAAGAAAGCTGGGTTCAGTCATCAACTTCAGTTTCAAAAGGC |
| 0_hypothetical | I2 | Tb927.11.11385_for | GGGGACAAGTTTGTACAAAAAAGCAGGCTATGGCTGAAAAAATAGCAAGCGAC | Tb927.11.11385_rev | GGGGACCACTTTGTACAAGAAAGCTGGGTTTATCCGTGAAATATCGCCGTCG |
| 0_hypothetical | I3 | Tb927.8.6990_for | GGGGACAAGTTTGTACAAAAAAGCAGGCTATGCAGGGTTCAACAGAGAAAACTAC | Tb927.8.6990_rev | GGGGACCACTTTGTACAAGAAAGCTGGGTCTAAAAAAAAAGACCGCAGCACGT |
| 0_hypothetical | I4 | Tb927.10.2340_for | GGGGACAAGTTTGTACAAAAAAGCAGGCTATGTCTAAACAAGAACTTCAACTCACTGA | Tb927.10.2340_rev | GGGGACCACTTTGTACAAGAAAGCTGGGTCTAAAGTATTAAAAGCACCACCAGAATAAG |
| 0_hypothetical | I5 | Tb927.10.14350_for | GGGGACAAGTTTGTACAAAAAAGCAGGCTATGTGGCGCATTTCTCCGC | Tb927.10.14350_rev | GGGGACCACTTTGTACAAGAAAGCTGGGTTCACGCGGTACCGCTTTTTG |
| 0_hypothetical | I6 | Tb927.4.5335_for | GGGGACAAGTTTGTACAAAAAAGCAGGCTATGCAGGTTGCAACGGAGAAA | Tb927.4.5335_rev | GGGGACCACTTTGTACAAGAAAGCTGGGTCTAAAAAAATAGACCGCAGCACGTC |
| 0_hypothetical | I7 | Tb927.8.2735_for | GGGGACAAGTTTGTACAAAAAAGCAGGCTATGAAGCGCTTCATACAAAGCCG | Tb927.8.2735_rev | GGGGACCACTTTGTACAAGAAAGCTGGGTTCACGAGTCCCTGTACCGTC |
| 0_hypothetical | I9 | Tb927.7.6770_for | GGGGACAAGTTTGTACAAAAAAGCAGGCTATGAGTTCTGCCCAGTTTGAAGC | Tb927.7.6770_rev | GGGGACCACTTTGTACAAGAAAGCTGGGTCTACTGAGGTGGCTTCCACGG |
| 0_hypothetical | I10 | Tb927.11.8165_for | GGGGACAAGTTTGTACAAAAAAGCAGGCTATGTCGCAAAAAGGTTTGAAGGGT | Tb927.11.8165_rev | GGGGACCACTTTGTACAAGAAAGCTGGGTTCATCCCTTCTTGTCGCCCTT |
| 0_hypothetical | I11 | Tb927.1.2985_for | GGGGACAAGTTTGTACAAAAAAGCAGGCTATGGCCGGGGAAGCGGGA | Tb927.1.2985_rev | GGGGACCACTTTGTACAAGAAAGCTGGGTTCATTGGTTCCTTTTCTCACGAACTT |
| 0_hypothetical | I12 | Tb927.6.590_for | GGGGACAAGTTTGTACAAAAAAGCAGGCTATGACTGAAAACATCGAAGCGGT | Tb927.6.590_rev | GGGGACCACTTTGTACAAGAAAGCTGGGTTTAATACGGATTGTTGCGGTCCTC |
| 0_hypothetical | I13 | Tb927.11.15780_for | GGGGACAAGTTTGTACAAAAAAGCAGGCTATGACAACAAGTGGAGTTGAATCGG | Tb927.11.15780_rev | GGGGACCACTTTGTACAAGAAAGCTGGGTCTACAGACCTAACTCCTTTAGGCACG |
| 0_hypothetical | I14 | Tb927.11.16540_for | GGGGACAAGTTTGTACAAAAAAGCAGGCTATGACGAGTGAGCGCGTGG | Tb927.11.16540_rev | GGGGACCACTTTGTACAAGAAAGCTGGGTTCACTTCGTGAGCTCCTGTATCA |
| 0_hypothetical | I15 | Tb927.11.2190_for | GGGGACAAGTTTGTACAAAAAAGCAGGCTATGAGCGAGCAACGAGTTTCAC | Tb927.11.2190_rev | GGGGACCACTTTGTACAAGAAAGCTGGGTTTAATTGATTTTCGACTCCTTGGGTTTCA |
| 0_hypothetical | I16 | Tb927.8.3521_for | GGGGACAAGTTTGTACAAAAAAGCAGGCTATGACCAATGAGCTCATGCGTC | Tb927.8.3521_rev | GGGGACCACTTTGTACAAGAAAGCTGGGTCTACACGCTACCTCTATTTATACCCTGA |
| 0_hypothetical | I17 | Tb927.9.2280_for | GGGGACAAGTTTGTACAAAAAAGCAGGCTATGGGACGCAGCAACAACC | Tb927.9.2280_rev | GGGGACCACTTTGTACAAGAAAGCTGGGTTCAACGCTTACCGCCTTTCTTC |
| 0_hypothetical | I18 | Tb927.9.13310_for | GGGGACAAGTTTGTACAAAAAAGCAGGCTATGGGGTTCTCTCTTGCTGGG | Tb927.9.13310_rev | GGGGACCACTTTGTACAAGAAAGCTGGGTTTATCCAAACAAAAGTGTGAACACAATCAC |

| | | | | | |
|---|---|---|---|---|---|
| 0_hypothetical | I19 | Tb927.9.17690_for | GGGGACAAGTTTGTACAAAAAAGCAGGCTATGCCGAGGACATTCGAAGTTC | Tb927.9.17690_rev | GGGGACCACTTTGTACAAGAAAGCTGGGTTCAATTCAAAAGGATTTGGTTCTTTCCTTT |
| 0_hypothetical | I20 | Tb927.7.7390_for | GGGGACAAGTTTGTACAAAAAAGCAGGCTATGAACATGCAAATCCTCCACGAA | Tb927.7.7390_rev | GGGGACCACTTTGTACAAGAAAGCTGGGTTCATCTAGTGCCCTTCATCTGCG |
| 0_hypothetical | I21 | Tb927.10.10030_for | GGGGACAAGTTTGTACAAAAAAGCAGGCTATGTCCGCCGAGGCCGTA | Tb927.10.10030_rev | GGGGACCACTTTGTACAAGAAAGCTGGGTCTACGAGTTCTCCTCCGGAGAG |
| 0_hypothetical | I22 | Tb927.8.8235_for | GGGGACAAGTTTGTACAAAAAAGCAGGCTATGTTTTATGATATCAAAGATGCTGCCTCA | Tb927.8.8235_rev | GGGGACCACTTTGTACAAGAAAGCTGGGTTCAAAAAGCCTTTAGGAAATCTGAGTTTCT |
| 0_hypothetical | I23 | Tb927.10.11030_for | GGGGACAAGTTTGTACAAAAAAGCAGGCTATGATGTTCGGTCGCCCCG | Tb927.10.11030_rev | GGGGACCACTTTGTACAAGAAAGCTGGGTCTAGAATTCCTCTTCGTCCTCCTCC |
| 0_hypothetical | I24 | Tb927.10.4190_for | GGGGACAAGTTTGTACAAAAAAGCAGGCTATGCGGGAGAACGGTGGG | Tb927.10.4190_rev | GGGGACCACTTTGTACAAGAAAGCTGGGTTCACACAACGGTTGGACTCCT |
| 0_hypothetical | J1 | Tb927.9.8650_for | GGGGACAAGTTTGTACAAAAAAGCAGGCTATGGACCCAGCACTGCAGC | Tb927.9.8650_rev | GGGGACCACTTTGTACAAGAAAGCTGGGTTTAGTACCTCTGGCCACCCTTTT |
| 0_hypothetical | J2 | Tb927.5.3090_for | GGGGACAAGTTTGTACAAAAAAGCAGGCTATGTCCTCCGGTTTCCATTTTCATG | Tb927.5.3090_rev | GGGGACCACTTTGTACAAGAAAGCTGGGTTTATTTGAATGGATTCTTGTCGTCCTCTT |
| 0_hypothetical | J3 | Tb927.9.16060_for | GGGGACAAGTTTGTACAAAAAAGCAGGCTATGCTATACGTAAGCGCTGGAGA | Tb927.9.16060_rev | GGGGACCACTTTGTACAAGAAAGCTGGGTCTAGGAGCGACACCGTAGTTTATCT |
| 0_hypothetical | J4 | Tb927.10.11530_for | GGGGACAAGTTTGTACAAAAAAGCAGGCTATGTACCTTGCGGTTTTCCAGG | Tb927.10.11530_rev | GGGGACCACTTTGTACAAGAAAGCTGGGTTCATCCGGTGTATGTTAATATGACACG |
| 0_hypothetical | J5 | Tb927.3.5740_for | GGGGACAAGTTTGTACAAAAAAGCAGGCTATGTGTGATGGCAGGAACTCGC | Tb927.3.5740_rev | GGGGACCACTTTGTACAAGAAAGCTGGGTCTAGGATAGCGTGACGGTTTTTACC |
| 0_hypothetical | J6 | Tb927.3.1600_for | GGGGACAAGTTTGTACAAAAAAGCAGGCTATGCAGCCACCTCAAACGAATC | Tb927.3.1600_rev | GGGGACCACTTTGTACAAGAAAGCTGGGTCTATTCGTTCTCCGACAAAACTTCTTTC |
| 0_hypothetical | J7 | Tb927.5.4575_for | GGGGACAAGTTTGTACAAAAAAGCAGGCTATGACGGACACGTTTGAAGGTATTT | Tb927.5.4575_rev | GGGGACCACTTTGTACAAGAAAGCTGGGTTTAGGGGGTTTTCCCCTGGTC |
| 0_hypothetical | J8 | Tb927.4.1830_for | GGGGACAAGTTTGTACAAAAAAGCAGGCTATGTCGAACCACAACCACATAACA | Tb927.4.1830_rev | GGGGACCACTTTGTACAAGAAAGCTGGGTTCATCCGCCATGCAGTGTG |
| 0_hypothetical | J9 | Tb927.9.7920_for | GGGGACAAGTTTGTACAAAAAAGCAGGCTATGTGTACGTGCTGCGTGG | Tb927.9.7920_rev | GGGGACCACTTTGTACAAGAAAGCTGGGTCTATACTGATAATTTATTGAGTTGGTTGCT |
| 0_hypothetical | J10 | Tb927.4.2660_for | GGGGACAAGTTTGTACAAAAAAGCAGGCTATGTTTATTGCCCGAACCCTTCC | Tb927.4.2660_rev | GGGGACCACTTTGTACAAGAAAGCTGGGTTTAATTCTTAAAGGGTGACCAGGGCT |
| 0_hypothetical | J11 | Tb927.11.10050_for | GGGGACAAGTTTGTACAAAAAAGCAGGCTATGGCATTTCGCCGCTTGG | Tb927.11.10050_rev | GGGGACCACTTTGTACAAGAAAGCTGGGTCTAACGAAATGGTGTAGGGAATCCC |
| 0_hypothetical | J12 | Tb927.8.4895_for | GGGGACAAGTTTGTACAAAAAAGCAGGCTATGACGTTTAAATTCTTTTACAATGTGCCG | Tb927.8.4895_rev | GGGGACCACTTTGTACAAGAAAGCTGGGTCTAAGGTCGAAGTAAATACTTATCATCGCC |
| 0_hypothetical | J13 | Tb927.11.9860_for | GGGGACAAGTTTGTACAAAAAAGCAGGCTATGCAACAAAGTCTCATTCCTGGAC | Tb927.11.9860_rev | GGGGACCACTTTGTACAAGAAAGCTGGGTTTATCGCTGAGCAAAGCTGAGC |
| 0_hypothetical | J14 | Tb927.5.4050_for | GGGGACAAGTTTGTACAAAAAAGCAGGCTATGAAAGCAACAACTAAATCAGGAGATAGC | Tb927.5.4050_rev | GGGGACCACTTTGTACAAGAAAGCTGGGTTCACGAACCCCCACGCGC |
| 0_hypothetical | J15 | Tb927.4.3600_for | GGGGACAAGTTTGTACAAAAAAGCAGGCTATGAGGCGTATGGATCCCGATG | Tb927.4.3600_rev | GGGGACCACTTTGTACAAGAAAGCTGGGTCTAGACATTGGCAGGTGCATCG |
| 0_hypothetical | J16 | Tb927.8.1490_for | GGGGACAAGTTTGTACAAAAAAGCAGGCTATGCGTCGTTCATTCTCACGC | Tb927.8.1490_rev | GGGGACCACTTTGTACAAGAAAGCTGGGTTCAGAAATCGTGGCAGCGTC |
| 0_hypothetical | J17 | Tb927.6.240_for | GGGGACAAGTTTGTACAAAAAAGCAGGCTATGATAAGGGAGCATATGGTACAATACAGT | Tb927.6.240_rev | GGGGACCACTTTGTACAAGAAAGCTGGGTTCATGTATTGAGCTTTAAAGGTCGTCG |
| 0_hypothetical | J19 | Tb927.7.4800_for | GGGGACAAGTTTGTACAAAAAAGCAGGCTATGTTGCAACGTACGACCTCG | Tb927.7.4800_rev | GGGGACCACTTTGTACAAGAAAGCTGGGTCTACATGGGCCTCCCGCT |
| 0_hypothetical | J20 | Tb927.11.15080_for | GGGGACAAGTTTGTACAAAAAAGCAGGCTATGGCGAACGTAATTGCCTCAG | Tb927.11.15080_rev | GGGGACCACTTTGTACAAGAAAGCTGGGTTTAAAATGAATAAGGTGCAAATCCGAAGAG |
| 0_hypothetical | J21 | Tb927.10.6200_for | GGGGACAAGTTTGTACAAAAAAGCAGGCTATGCTCCGACGCGTTGCT | Tb927.10.6200_rev | GGGGACCACTTTGTACAAGAAAGCTGGGTTTACACTGCCCTGTCAGGAACA |
| 0_hypothetical | J22 | Tb927.6.4130_for | GGGGACAAGTTTGTACAAAAAAGCAGGCTATGCCACCTGTGGTTAAGCG | Tb927.6.4130_rev | GGGGACCACTTTGTACAAGAAAGCTGGGTTTACGGCAATGGGTTCATCATGTC |
| 0_hypothetical | J23 | Tb927.3.2880_for | GGGGACAAGTTTGTACAAAAAAGCAGGCTATGGTTTACACTCGTTGGAAGTGTG | Tb927.3.2880_rev | GGGGACCACTTTGTACAAGAAAGCTGGGTCTACACGACTCGACTTAACTCCTCC |
| 0_hypothetical | J24 | Tb927.8.2170_for | GGGGACAAGTTTGTACAAAAAAGCAGGCTATGAGGCGATGTGCTTGTAAGATG | Tb927.8.2170_rev | GGGGACCACTTTGTACAAGAAAGCTGGGTTCACACCTTTTCTTCAGCACCAC |
| 0_hypothetical | K1 | Tb927.10.15320_for | GGGGACAAGTTTGTACAAAAAAGCAGGCTATGAAACGACTGGCGTCTTTTGG | Tb927.10.15320_rev | GGGGACCACTTTGTACAAGAAAGCTGGGTTCACATGTTTGCCACAGGTTCTG |
| 0_hypothetical | K2 | Tb927.7.7550_for | GGGGACAAGTTTGTACAAAAAAGCAGGCTATGACATTAATTGAAATTCTAGGCCACTAC | Tb927.7.7550_rev | GGGGACCACTTTGTACAAGAAAGCTGGGTTTACTGACCCAGTGCCTGGAG |
| 0_hypothetical | K3 | Tb927.2.2590_for | GGGGACAAGTTTGTACAAAAAAGCAGGCTATGTACCAGAGGACACGATTTTTGTG | Tb927.2.2590_rev | GGGGACCACTTTGTACAAGAAAGCTGGGTTCACTCAGTAGTTAGGCCGCG |
| 0_hypothetical | K4 | Tb927.11.15855_for | GGGGACAAGTTTGTACAAAAAAGCAGGCTATGTTTTGCTCCACGCCACTG | Tb927.11.15855_rev | GGGGACCACTTTGTACAAGAAAGCTGGGTTTATCCAGGTGGTTCGGGGAG |
| 0_hypothetical | K5 | Tb927.11.2825_for | GGGGACAAGTTTGTACAAAAAAGCAGGCTATGCTTACGACTGGACCGTTGA | Tb927.11.2825_rev | GGGGACCACTTTGTACAAGAAAGCTGGGTTCATTCATCTACCAATTCTGTAAAGAAACC |
| 0_hypothetical | K6 | Tb927.4.510_for | GGGGACAAGTTTGTACAAAAAAGCAGGCTATGGAGGTGGCTGCAATGAGT | Tb927.4.510_rev | GGGGACCACTTTGTACAAGAAAGCTGGGTCTAGCTTTGCTCCAGAGCATCAG |
| 0_hypothetical | K7 | Tb927.4.720_for | GGGGACAAGTTTGTACAAAAAAGCAGGCTATGCTGCGGCGACTGGTC | Tb927.4.720_rev | GGGGACCACTTTGTACAAGAAAGCTGGGTTTAAATGCTTTTTGGGAACACCAACTC |
| 0_hypothetical | K8 | Tb927.3.5350_for | GGGGACAAGTTTGTACAAAAAAGCAGGCTATGCCCGCACGTGCATCC | Tb927.3.5350_rev | GGGGACCACTTTGTACAAGAAAGCTGGGTTTAAGGACGAGCGCCACGG |
| 0_hypothetical | K9 | Tb927.8.4380_for | GGGGACAAGTTTGTACAAAAAAGCAGGCTATGTTTATGATGCTCAACGGTACCG | Tb927.8.4380_rev | GGGGACCACTTTGTACAAGAAAGCTGGGTTCAAGCAGTGGAGGAAACATACTCC |
| 0_hypothetical | K10 | Tb927.8.6430_for | GGGGACAAGTTTGTACAAAAAAGCAGGCTATGGTAAAGTTGACTCATGAAGATACGC | Tb927.8.6430_rev | GGGGACCACTTTGTACAAGAAAGCTGGGTTTAGAAACCCTTTGCAGTTTTCAATTCC |
| 0_hypothetical | K11 | Tb927.9.740_for | GGGGACAAGTTTGTACAAAAAAGCAGGCTATGCAAGGTGCTTTGAGTGGG | Tb927.9.740_rev | GGGGACCACTTTGTACAAGAAAGCTGGGTTCATAAAATATGTATCATCTCACAAGACAC |
| 0_hypothetical | K12 | Tb927.6.1240_for | GGGGACAAGTTTGTACAAAAAAGCAGGCTATGACTCGAAAACAAAAGCAGCG | Tb927.6.1240_rev | GGGGACCACTTTGTACAAGAAAGCTGGGTTTATGCCTTTGGCGTCTGGTG |
| 0_hypothetical | K13 | Tb927.3.2380_for | GGGGACAAGTTTGTACAAAAAAGCAGGCTATGGAAAAAAATGCGCATGGGC | Tb927.3.2380_rev | GGGGACCACTTTGTACAAGAAAGCTGGGTTCATTGCCGTGTTGCCCC |
| 0_hypothetical | K14 | Tb927.11.605_for | GGGGACAAGTTTGTACAAAAAAGCAGGCTATGAGGAGGCTGAGCGCA | Tb927.11.605_rev | GGGGACCACTTTGTACAAGAAAGCTGGGTTTACTTCACGCTTACTCCCTCGA |
| 0_hypothetical | K15 | Tb927.6.4020_for | GGGGACAAGTTTGTACAAAAAAGCAGGCTATGGTGCGCGCCCCTCCG | Tb927.6.4020_rev | GGGGACCACTTTGTACAAGAAAGCTGGGTTTACAAACTTCCCTTCAAATGGCAAGC |
| 0_hypothetical | K16 | Tb927.10.1810_for | GGGGACAAGTTTGTACAAAAAAGCAGGCTATGGAGGAAGCGGCTGCT | Tb927.10.1810_rev | GGGGACCACTTTGTACAAGAAAGCTGGGTCTACACTCCTAACTTCTGGTACGACC |
| 0_hypothetical | K17 | Tb927.7.5830_for | GGGGACAAGTTTGTACAAAAAAGCAGGCTATGAAGCCCGCCACTATTCCA | Tb927.7.5830_rev | GGGGACCACTTTGTACAAGAAAGCTGGGTTCAAACATCTGTTGTGGTGCACA |
| 0_hypothetical | K18 | Tb927.10.90_for | GGGGACAAGTTTGTACAAAAAAGCAGGCTATGTGCCTACTGGGTCGTTCA | Tb927.10.90_rev | GGGGACCACTTTGTACAAGAAAGCTGGGTTCATTCCAGTGGTGGAGAAACGG |
| 0_hypothetical | K19 | Tb927.10.3410_for | GGGGACAAGTTTGTACAAAAAAGCAGGCTATGCAACACTTTGTTGGTGGTACA | Tb927.10.3410_rev | GGGGACCACTTTGTACAAGAAAGCTGGGTTTAAGTGGCAAATACATTTACACTTAACTC |
| 0_hypothetical | K20 | Tb927.11.17010_for | GGGGACAAGTTTGTACAAAAAAGCAGGCTATGAGCTTAGGGAAAACTATTTATGTTGAC | Tb927.11.17010_rev | GGGGACCACTTTGTACAAGAAAGCTGGGTCTAGAATATATAAATTGCGTCATATGCTAC |
| 0_hypothetical | K21 | Tb927.11.12040_for | GGGGACAAGTTTGTACAAAAAAGCAGGCTATGGCTTCTTGCAGAACTGGGC | Tb927.11.12040_rev | GGGGACCACTTTGTACAAGAAAGCTGGGTCTAATAAGTGTACTCTTTACCAGTGTACCC |
| 0_hypothetical | K22 | Tb927.11.5950_for | GGGGACAAGTTTGTACAAAAAAGCAGGCTATGTTCTCGAAATCCGTCGCAC | Tb927.11.5950_rev | GGGGACCACTTTGTACAAGAAAGCTGGGTTCAAACGGTTGCCTGCTCC |
| 0_hypothetical | K23 | Tb927.10.8160_for | GGGGACAAGTTTGTACAAAAAAGCAGGCTATGCCCGTCCTAGTTGCGG | Tb927.10.8160_rev | GGGGACCACTTTGTACAAGAAAGCTGGGTCTACAGGCTGATGTACAAGAAGAATGC |
| 0_hypothetical | L1 | Tb927.2.2140_for | GGGGACAAGTTTGTACAAAAAAGCAGGCTATGTCATATGTTGAAAATGTGATGGCAC | Tb927.2.2140_rev | GGGGACCACTTTGTACAAGAAAGCTGGGTTTACTTCCCCTGCGCATCTGC |
| 0_hypothetical | L2 | Tb927.11.13800_for | GGGGACAAGTTTGTACAAAAAAGCAGGCTATGGATTCTGAAGTCTGCGATGAC | Tb927.11.13800_rev | GGGGACCACTTTGTACAAGAAAGCTGGGTCTACGTGACGTATTCGGAGCCA |
| 0_hypothetical | L3 | Tb927.11.7702_for | GGGGACAAGTTTGTACAAAAAAGCAGGCTATGAGTGGAAATGTGAAGTTCAGCA | Tb927.11.7702_rev | GGGGACCACTTTGTACAAGAAAGCTGGGTTCATGCGGGAGAAATCATTTCAACA |
| 0_hypothetical | L4 | Tb927.1.110_for | GGGGACAAGTTTGTACAAAAAAGCAGGCTATGAAAACAAAACGTTACAATGTTTTCCTA | Tb927.1.110_rev | GGGGACCACTTTGTACAAGAAAGCTGGGTTTAGGGGGAGCGGGTTCG |
| 0_hypothetical | L5 | Tb927.11.2960_for | GGGGACAAGTTTGTACAAAAAAGCAGGCTATGAGTGCTCTTCCTGATGCTAGC | Tb927.11.2960_rev | GGGGACCACTTTGTACAAGAAAGCTGGGTTTAAAGTGCTTTCATTTCAGATTCTTCCCC |
| 0_hypothetical | L6 | Tb927.9.9610_for | GGGGACAAGTTTGTACAAAAAAGCAGGCTATGCAAAAGGTCGGCGTCAA | Tb927.9.9610_rev | GGGGACCACTTTGTACAAGAAAGCTGGGTCTACGGGCTGCACAGGTGA |
| 0_hypothetical | L7 | Tb927.1.40_for | GGGGACAAGTTTGTACAAAAAAGCAGGCTATGAAACTGAAAGAGCTAATAGGTTGGC | Tb927.1.40_rev | GGGGACCACTTTGTACAAGAAAGCTGGGTTTAAAGTGGATTACAAATTTCATTGAAAAT |
| 0_hypothetical | L8 | Tb927.9.2055_for | GGGGACAAGTTTGTACAAAAAAGCAGGCTATGTTCAGTTGGAAGGATGTTCTCTC | Tb927.9.2055_rev | GGGGACCACTTTGTACAAGAAAGCTGGGTTTAATCCAACTGCGTTTCACCGC |
| 0_hypothetical | L9 | Tb927.6.811_for | GGGGACAAGTTTGTACAAAAAAGCAGGCTATGACTGTGTACTACTATTACGGGGAG | Tb927.6.811_rev | GGGGACCACTTTGTACAAGAAAGCTGGGTTTACATGACGTCTTCTTCATTTGGCC |
| 0_hypothetical | L10 | Tb927.4.3430_for | GGGGACAAGTTTGTACAAAAAAGCAGGCTATGGGTCAGGACCAATCCATGG | Tb927.4.3430_rev | GGGGACCACTTTGTACAAGAAAGCTGGGTTTAACGACTAAGTTCCTTTCTCGCC |
| 0_hypothetical | L11 | Tb927.6.1710_for | GGGGACAAGTTTGTACAAAAAAGCAGGCTATGTTACGACGGCTAGCTGCG | Tb927.6.1710_rev | GGGGACCACTTTGTACAAGAAAGCTGGGTTTAAACCTTAAGAAATTCCTGCGAGAGC |
| 0_hypothetical | L12 | Tb927.11.4670_for | GGGGACAAGTTTGTACAAAAAAGCAGGCTATGCCGGGCATTGAGCGT | Tb927.11.4670_rev | GGGGACCACTTTGTACAAGAAAGCTGGGTTTACGCTGAACATCCGTCAAGAATATC |
| 0_hypothetical | L13 | Tb927.9.1920_for | GGGGACAAGTTTGTACAAAAAAGCAGGCTATGCACAGTCGACGCATTTTTG | Tb927.9.1920_rev | GGGGACCACTTTGTACAAGAAAGCTGGGTTTACCGCATCTTCGGAAAAAAACAAG |
| 0_hypothetical | L14 | Tb927.9.7890_for | GGGGACAAGTTTGTACAAAAAAGCAGGCTATGTGTACGTGCAGCGTGG | Tb927.9.7890_rev | GGGGACCACTTTGTACAAGAAAGCTGGGTCTACAAACATTTGCTCCTTTCCCATACT |
| 0_hypothetical | L15 | Tb927.1.10_for | GGGGACAAGTTTGTACAAAAAAGCAGGCTATGATCGCGGGTTTCTGTTCG | Tb927.1.10_rev | GGGGACCACTTTGTACAAGAAAGCTGGGTCTATTGACTTCCAGCACTTTGTATAGAGTT |
| 0_hypothetical | L16 | Tb927.5.460_for | GGGGACAAGTTTGTACAAAAAAGCAGGCTATGGTTGTATTGAAGGGTGGCG | Tb927.5.460_rev | GGGGACCACTTTGTACAAGAAAGCTGGGTTCATCCTCCTTCATTACCATCACTGC |
| 0_hypothetical | L17 | Tb927.3.5560_for | GGGGACAAGTTTGTACAAAAAAGCAGGCTATGATTGACGATGTAAGAATTTGTCATCTC | Tb927.3.5560_rev | GGGGACCACTTTGTACAAGAAAGCTGGGTTCAGGAGGAACCGCTGCG |
| 0_hypothetical | L18 | Tb927.10.7390_for | GGGGACAAGTTTGTACAAAAAAGCAGGCTATGCTTCGCCACTCCAAGC | Tb927.10.7390_rev | GGGGACCACTTTGTACAAGAAAGCTGGGTTTAGCGTGTGCGTACCGTCG |
| 0_hypothetical | L19 | Tb927.10.15900_for | GGGGACAAGTTTGTACAAAAAAGCAGGCTATGTTTGGACGACGCGTTTACG | Tb927.10.15900_rev | GGGGACCACTTTGTACAAGAAAGCTGGGTTCAGTGAAGCACTTTCTTTACCCC |
| 0_hypothetical | L20 | Tb927.10.12610_for | GGGGACAAGTTTGTACAAAAAAGCAGGCTATGGCGCGGTGCATCATT | Tb927.10.12610_rev | GGGGACCACTTTGTACAAGAAAGCTGGGTTCACCCCTCGCGTGCTGA |
| 0_hypothetical | L21 | Tb927.9.1120_for | GGGGACAAGTTTGTACAAAAAAGCAGGCTATGGGGATTAGTTTTTGTGTGTCACT | Tb927.9.1120_rev | GGGGACCACTTTGTACAAGAAAGCTGGGTTTATATATCCTCTTCTCCAACAATTTCTCT |
| 0_hypothetical | L22 | Tb927.8.3440_for | GGGGACAAGTTTGTACAAAAAAGCAGGCTATGAGTGTTGGGAAATTTAAACGGTCA | Tb927.8.3440_rev | GGGGACCACTTTGTACAAGAAAGCTGGGTTTACCAGCGGCACATTTGACG |
| 0_hypothetical | L23 | Tb927.6.3200_for | GGGGACAAGTTTGTACAAAAAAGCAGGCTATGCTTCGTAGGAAAGGATACATTGG | Tb927.6.3200_rev | GGGGACCACTTTGTACAAGAAAGCTGGGTTCAGTGTGGAGAGGAATACCGATC |
| 0_hypothetical | L24 | Tb927.6.1600_for | GGGGACAAGTTTGTACAAAAAAGCAGGCTATGCTGCGTCGCCGGATA | Tb927.6.1600_rev | GGGGACCACTTTGTACAAGAAAGCTGGGTTCATTTGCGCCAAGGTTCCTT |
| 0_hypothetical | M1 | Tb927.11.1330_for | GGGGACAAGTTTGTACAAAAAAGCAGGCTATGTCCGCGAAGATGTGCC | Tb927.11.1330_rev | GGGGACCACTTTGTACAAGAAAGCTGGGTTCACTCCTTCAGCTCAAAGTCCA |
| 0_hypothetical | M2 | Tb927.11.15180_for | GGGGACAAGTTTGTACAAAAAAGCAGGCTATGGCGAAGAAACATCGTGTGA | Tb927.11.15180_rev | GGGGACCACTTTGTACAAGAAAGCTGGGTTCAGTACTTCTTCCTCAAATGTTTGACC |
| 0_hypothetical | M3 | Tb927.9.10000_for | GGGGACAAGTTTGTACAAAAAAGCAGGCTATGTCGAAGTTAGAGCAGCGC | Tb927.9.10000_rev | GGGGACCACTTTGTACAAGAAAGCTGGGTTTACAGTTCGTGCATCATGTATATCCG |
| 0_hypothetical | M4 | Tb927.7.4250_for | GGGGACAAGTTTGTACAAAAAAGCAGGCTATGACTCGGCAATCCGATGTG | Tb927.7.4250_rev | GGGGACCACTTTGTACAAGAAAGCTGGGTTCAGGAGGTTCCACTCGATGC |
| 0_hypothetical | M6 | Tb927.8.3600_for | GGGGACAAGTTTGTACAAAAAAGCAGGCTATGCGCGGGAGAGGACGC | Tb927.8.3600_rev | GGGGACCACTTTGTACAAGAAAGCTGGGTTTAGCCGCAGATCGGTGCC |
| 0_hypothetical | M7 | Tb927.9.10930_for | GGGGACAAGTTTGTACAAAAAAGCAGGCTATGGAACATATTCCTGCTGAAGCATC | Tb927.9.10930_rev | GGGGACCACTTTGTACAAGAAAGCTGGGTTCACAAACTCTCAGAGAACGACGG |
| 0_hypothetical | M8 | Tb927.10.7170_for | GGGGACAAGTTTGTACAAAAAAGCAGGCTATGTTCCCCATTCCGCCAGC | Tb927.10.7170_rev | GGGGACCACTTTGTACAAGAAAGCTGGGTCTAAACGCGCAAACTGACTGTG |
| 0_hypothetical | M9 | Tb927.4.440_for | GGGGACAAGTTTGTACAAAAAAGCAGGCTATGACTGGTTGGAACCAAAAAATGGC | Tb927.4.440_rev | GGGGACCACTTTGTACAAGAAAGCTGGGTTTATCGTCTATCCGGCTGAATATCGC |
| 0_hypothetical | M10 | Tb927.9.140_for | GGGGACAAGTTTGTACAAAAAAGCAGGCTATGCCGCCAGGGTCCAGT | Tb927.9.140_rev | GGGGACCACTTTGTACAAGAAAGCTGGGTCTAAAACCCCCTGTTTTTTTTTGATGAATC |
| 0_hypothetical | M11 | Tb927.11.11135_for | GGGGACAAGTTTGTACAAAAAAGCAGGCTATGGGTGGGCCACCAAGG | Tb927.11.11135_rev | GGGGACCACTTTGTACAAGAAAGCTGGGTTCAATTCATTAGATCCAGATCACCGACA |
| 0_hypothetical | M12 | Tb927.10.13770_for | GGGGACAAGTTTGTACAAAAAAGCAGGCTATGGTCCTGCGAGGCGTG | Tb927.10.13770_rev | GGGGACCACTTTGTACAAGAAAGCTGGGTTCACTCCCGAAGCGAGAAGT |
| 0_hypothetical | M13 | Tb927.10.10270_for | GGGGACAAGTTTGTACAAAAAAGCAGGCTATGCAACACCCGGGTCGC | Tb927.10.10270_rev | GGGGACCACTTTGTACAAGAAAGCTGGGTCTATGATTGAATCAAATATTGCCATGCCTC |
| 0_hypothetical | M14 | Tb927.9.2110_for | GGGGACAAGTTTGTACAAAAAAGCAGGCTATGTCACGCAATACAAAGGAATTTAACC | Tb927.9.2110_rev | GGGGACCACTTTGTACAAGAAAGCTGGGTTCAGTTCGCTTCGGGATTCTTCT |
| 0_hypothetical | M15 | Tb927.8.640_for | GGGGACAAGTTTGTACAAAAAAGCAGGCTATGGAGGGGAAGAGTGAAGCC | Tb927.8.640_rev | GGGGACCACTTTGTACAAGAAAGCTGGGTTCACATATCGGATTTCGGTGCCT |
| 0_hypothetical | M16 | Tb927.10.13810_for | GGGGACAAGTTTGTACAAAAAAGCAGGCTATGTCTAGTCGCTTTTTCCAAAAGTATTTT | Tb927.10.13810_rev | GGGGACCACTTTGTACAAGAAAGCTGGGTTTAATTCGATGTTCCGGACGACG |
| 0_hypothetical | M17 | Tb927.9.670_for | GGGGACAAGTTTGTACAAAAAAGCAGGCTATGTGGTGTGCCGCGCCA | Tb927.9.670_rev | GGGGACCACTTTGTACAAGAAAGCTGGGTTCAGGAATTGTTTCTGTTCCCTCTCA |
| 0_hypothetical | M18 | Tb927.8.4910_for | GGGGACAAGTTTGTACAAAAAAGCAGGCTATGCCCGCCAGACCCAAT | Tb927.8.4910_rev | GGGGACCACTTTGTACAAGAAAGCTGGGTCTAACGGAGGGTTAAACGAGCC |
| 0_hypothetical | M19 | Tb927.10.1580_for | GGGGACAAGTTTGTACAAAAAAGCAGGCTATGACAGCGCCCGTCTCT | Tb927.10.1580_rev | GGGGACCACTTTGTACAAGAAAGCTGGGTCTAACTCCCGGAAGATTCTCTATCGC |
| 0_hypothetical | M20 | Tb927.4.1810_for | GGGGACAAGTTTGTACAAAAAAGCAGGCTATGCTCTCCTCCACCGCC | Tb927.4.1810_rev | GGGGACCACTTTGTACAAGAAAGCTGGGTTTATTTCTTCAACATCTTTTCCGCACC |
| 0_hypothetical | M21 | Tb927.9.16410_for | GGGGACAAGTTTGTACAAAAAAGCAGGCTATGTCAGTTTGGAACGACATTCTTTGG | Tb927.9.16410_rev | GGGGACCACTTTGTACAAGAAAGCTGGGTTTACCCCCACATCTCTCTGGTTG |
| 0_hypothetical | M22 | Tb927.4.3450_for | GGGGACAAGTTTGTACAAAAAAGCAGGCTATGCTACGGCGACTTGGTGC | Tb927.4.3450_rev | GGGGACCACTTTGTACAAGAAAGCTGGGTTTAAATGTTGAGTGAGCGAGTAATGGC |
| 0_hypothetical | M23 | Tb927.9.10430_for | GGGGACAAGTTTGTACAAAAAAGCAGGCTATGCTCACTATCACCGCTCTTCT | Tb927.9.10430_rev | GGGGACCACTTTGTACAAGAAAGCTGGGTTTATTTCCGTCCCCTTTGAACCCT |
| 0_hypothetical | M24 | Tb927.8.2530_for | GGGGACAAGTTTGTACAAAAAAGCAGGCTATGGGCATAGAAACGTGGAAGGT | Tb927.8.2530_rev | GGGGACCACTTTGTACAAGAAAGCTGGGTTTACGCCGTGCCTTCCCA |
| 0_hypothetical | N1 | Tb927.8.1665_for | GGGGACAAGTTTGTACAAAAAAGCAGGCTATGCTGACTTATTGCCTTTTGGGA | Tb927.8.1665_rev | GGGGACCACTTTGTACAAGAAAGCTGGGTCTATACATCTCGCGTGTTGAAAAAAGC |
| 0_hypothetical | N2 | Tb927.1.4970_for | GGGGACAAGTTTGTACAAAAAAGCAGGCTATGGCAGACCGAGCTTTAACG | Tb927.1.4970_rev | GGGGACCACTTTGTACAAGAAAGCTGGGTTTAAACTTGAACGTGTTCCACAGAGC |
| 0_hypothetical | N3 | Tb927.9.15080_for | GGGGACAAGTTTGTACAAAAAAGCAGGCTATGAGGCCCCTCCGCGCG | Tb927.9.15080_rev | GGGGACCACTTTGTACAAGAAAGCTGGGTTCATGTTCTTTTTAATCCTTCAATGTCCAC |
| 0_hypothetical | N4 | Tb927.10.2970_for | GGGGACAAGTTTGTACAAAAAAGCAGGCTATGCTGCCCCTCCGTGTAC | Tb927.10.2970_rev | GGGGACCACTTTGTACAAGAAAGCTGGGTCTATTGCTTCTCGTTCGTTAACTGCT |
| 0_hypothetical | N5 | Tb927.10.16060_for | GGGGACAAGTTTGTACAAAAAAGCAGGCTATGTCCGAGGACACGTTTTTTCG | Tb927.10.16060_rev | GGGGACCACTTTGTACAAGAAAGCTGGGTTCATTTCTGTTTGTCTATATCCGAAGACG |
| 0_hypothetical | N6 | Tb927.2.1790_for | GGGGACAAGTTTGTACAAAAAAGCAGGCTATGAAGCACATGCCATTCAGAGAG | Tb927.2.1790_rev | GGGGACCACTTTGTACAAGAAAGCTGGGTTTAGCTCCGCCTGTGAAAACAAC |
| 0_hypothetical | N7 | Tb927.5.2200_for | GGGGACAAGTTTGTACAAAAAAGCAGGCTATGGCCACCTGCCCTAAACA | Tb927.5.2200_rev | GGGGACCACTTTGTACAAGAAAGCTGGGTTTACTCCACGGCCTCAAACACC |
| 0_hypothetical | N8 | Tb927.9.5800_for | GGGGACAAGTTTGTACAAAAAAGCAGGCTATGTTAGTCCACTTGGATAAATGGTGG | Tb927.9.5800_rev | GGGGACCACTTTGTACAAGAAAGCTGGGTTTAAACAAACCTCCCTTCGCTTCG |
| 0_hypothetical | N9 | Tb927.4.390_for | GGGGACAAGTTTGTACAAAAAAGCAGGCTATGCGAGACGAACAAAAGGAATATGC | Tb927.4.390_rev | GGGGACCACTTTGTACAAGAAAGCTGGGTTCAAAGGGCATGGAGAATAGTTTTCTT |
| 0_hypothetical | N10 | Tb927.7.4210_for | GGGGACAAGTTTGTACAAAAAAGCAGGCTATGTTAAGAAGCACTGCTGCGT | Tb927.7.4210_rev | GGGGACCACTTTGTACAAGAAAGCTGGGTTCAGACAATACCACCGAGTTTATTCTCC |
| 0_hypothetical | N11 | Tb927.3.585_for | GGGGACAAGTTTGTACAAAAAAGCAGGCTATGGCCGCAGCCGTCAAA | Tb927.3.585_rev | GGGGACCACTTTGTACAAGAAAGCTGGGTTTAATCCCGAGCGTCACCGA |
| 0_hypothetical | N12 | Tb927.4.1240_for | GGGGACAAGTTTGTACAAAAAAGCAGGCTATGTCCGTACGTTCCGATACTTTTTT | Tb927.4.1240_rev | GGGGACCACTTTGTACAAGAAAGCTGGGTTTAATTGAGCTCATTGACGGGGAATATAAC |
| 0_hypothetical | N13 | Tb927.10.11520_for | GGGGACAAGTTTGTACAAAAAAGCAGGCTATGCAACCCCCAACCCCA | Tb927.10.11520_rev | GGGGACCACTTTGTACAAGAAAGCTGGGTCTAGACCCCTCCCCCCTGC |
| 0_hypothetical | N14 | Tb927.8.2340_for | GGGGACAAGTTTGTACAAAAAAGCAGGCTATGAACCGGTTCTGCCGGC | Tb927.8.2340_rev | GGGGACCACTTTGTACAAGAAAGCTGGGTTCACTTGCACGGCCACCA |
| 0_hypothetical | N15 | Tb927.7.2780_for | GGGGACAAGTTTGTACAAAAAAGCAGGCTATGTCTAAAGCTCCTTCGCAACC | Tb927.7.2780_rev | GGGGACCACTTTGTACAAGAAAGCTGGGTCTATGCCGCCGCTGCCGG |
| 0_hypothetical | N16 | Tb927.10.12110_for | GGGGACAAGTTTGTACAAAAAAGCAGGCTATGTCGCAACGGGGTTCCG | Tb927.10.12110_rev | GGGGACCACTTTGTACAAGAAAGCTGGGTTCAAGCAGATGAAGCTGCAGC |
| 0_hypothetical | N17 | Tb927.9.13225_for | GGGGACAAGTTTGTACAAAAAAGCAGGCTATGGCAATGAGCGGGGATGG | Tb927.9.13225_rev | GGGGACCACTTTGTACAAGAAAGCTGGGTCTATGCACTCTCTCCGGAGCG |
| 0_hypothetical | N18 | Tb927.9.3540_for | GGGGACAAGTTTGTACAAAAAAGCAGGCTATGTCTTCGCTTCCCGGGG | Tb927.9.3540_rev | GGGGACCACTTTGTACAAGAAAGCTGGGTCTAATACCCGTTGTGGCTCCG |
| 0_hypothetical | N19 | Tb927.9.3840_for | GGGGACAAGTTTGTACAAAAAAGCAGGCTATGCAACGCCGAGTGCAG | Tb927.9.3840_rev | GGGGACCACTTTGTACAAGAAAGCTGGGTTCATACGCTCTTCTCCACAGCC |
| 0_hypothetical | N20 | Tb927.10.13515_for | GGGGACAAGTTTGTACAAAAAAGCAGGCTATGAACCCCCCAGAAGAATCAGAG | Tb927.10.13515_rev | GGGGACCACTTTGTACAAGAAAGCTGGGTCTATTTTGGTTGCGCTGGGGG |
| 0_hypothetical | N21 | Tb927.11.16950_for | GGGGACAAGTTTGTACAAAAAAGCAGGCTATGGTGTGGTACATTCCCTTTGTTT | Tb927.11.16950_rev | GGGGACCACTTTGTACAAGAAAGCTGGGTTCAATCATGCCGAAGGGTATAGGC |
| 0_hypothetical | N22 | Tb927.7.5010_for | GGGGACAAGTTTGTACAAAAAAGCAGGCTATGTCTCATGTCACAGCTGACCT | Tb927.7.5010_rev | GGGGACCACTTTGTACAAGAAAGCTGGGTTCACTCCGCAAGAAACTTAGCG |
| 0_hypothetical | N23 | Tb927.11.9425_for | GGGGACAAGTTTGTACAAAAAAGCAGGCTATGGACATTCTGCAGAACATTGCA | Tb927.11.9425_rev | GGGGACCACTTTGTACAAGAAAGCTGGGTTTAGTTCCACATTTGTTCTGCAACCC |
| 0_hypothetical | N24 | Tb927.11.16710_for | GGGGACAAGTTTGTACAAAAAAGCAGGCTATGGGGGTTCGCATGTTCTGC | Tb927.11.16710_rev | GGGGACCACTTTGTACAAGAAAGCTGGGTCTACAGTGGGTTGTTGGATCCAC |
| 0_hypothetical | O1 | Tb927.10.11200_for | GGGGACAAGTTTGTACAAAAAAGCAGGCTATGGGCACGATTTTTAGCAACGT | Tb927.10.11200_rev | GGGGACCACTTTGTACAAGAAAGCTGGGTCTAATTGGCAACATGGACCTCTGC |
| 0_hypothetical | O2 | Tb927.11.10220_for | GGGGACAAGTTTGTACAAAAAAGCAGGCTATGCCCCGTGGCGGCACT | Tb927.11.10220_rev | GGGGACCACTTTGTACAAGAAAGCTGGGTCTATACCCCCAGATAGCCCCG |
| 0_hypothetical | O3 | Tb927.10.500_for | GGGGACAAGTTTGTACAAAAAAGCAGGCTATGCATTTATGTTTTTCCTTTCTTTCTTTC | Tb927.10.500_rev | GGGGACCACTTTGTACAAGAAAGCTGGGTTTACACATTGCGTATATATAATAGAAAGAA |
| 0_hypothetical | O4 | Tb927.11.9665_for | GGGGACAAGTTTGTACAAAAAAGCAGGCTATGGAACCTCGCGAGATGAGT | Tb927.11.9665_rev | GGGGACCACTTTGTACAAGAAAGCTGGGTCTAAAAGTCCTCCCTGCGCC |
| 0_hypothetical | O5 | Tb927.11.5600_for | GGGGACAAGTTTGTACAAAAAAGCAGGCTATGACTGAAGCACTCTACAACTTCATC | Tb927.11.5600_rev | GGGGACCACTTTGTACAAGAAAGCTGGGTTTAAATGCCATGACGTTCACGATACT |
| 0_hypothetical | O6 | Tb927.11.160_for | GGGGACAAGTTTGTACAAAAAAGCAGGCTATGAGATCCACCGTCTTTGGTGG | Tb927.11.160_rev | GGGGACCACTTTGTACAAGAAAGCTGGGTCTATCGCAGTAGAACGCACCAAA |
| 0_hypothetical | O7 | Tb927.2.160_for | GGGGACAAGTTTGTACAAAAAAGCAGGCTATGTTTCAATTGGTTTCTACGGCCG | Tb927.2.160_rev | GGGGACCACTTTGTACAAGAAAGCTGGGTCTACATTTCCTTAAACCGGAAGCTAACC |

| | | | | | |
|---|---|---|---|---|---|
| 0_hypothetical | O8 | Tb927.10.4240_for | GGGGACAAGTTTGTACAAAAAAGCAGGCTATGATGAGAAACTCCCGGTTTCTGT | Tb927.10.4240_rev | GGGGACCACTTTGTACAAGAAAGCTGGGTTTACCACCAAGCTGCGTGC |
| 0_hypothetical | O9 | Tb927.11.9850_for | GGGGACAAGTTTGTACAAAAAAGCAGGCTATGTCACTTGTGTTGGCAGATGAA | Tb927.11.9850_rev | GGGGACCACTTTGTACAAGAAAGCTGGGTTTAACCCTCTGGCAGAAGCGC |
| 0_hypothetical | O10 | Tb927.1.400_for | GGGGACAAGTTTGTACAAAAAAGCAGGCTATGCCTCCCCTTGCTCGC | Tb927.1.400_rev | GGGGACCACTTTGTACAAGAAAGCTGGGTCTACAGGATGGTGGAAACGAAGC |
| 0_hypothetical | O11 | Tb927.8.3120_for | GGGGACAAGTTTGTACAAAAAAGCAGGCTATGTACTTTCTTTTGCTAGCTGGGATC | Tb927.8.3120_rev | GGGGACCACTTTGTACAAGAAAGCTGGGTTCATGATGATGATGCGATCCCACA |
| 0_hypothetical | O12 | Tb927.10.14370_for | GGGGACAAGTTTGTACAAAAAAGCAGGCTATGAGCGAGGAGGTGCAGC | Tb927.10.14370_rev | GGGGACCACTTTGTACAAGAAAGCTGGGTCTAAGATTCGTGCGAAGTCTTAGCC |
| 0_hypothetical | O13 | Tb927.11.16520_for | GGGGACAAGTTTGTACAAAAAAGCAGGCTATGAATTATTCAACCGGGTACTCTGC | Tb927.11.16520_rev | GGGGACCACTTTGTACAAGAAAGCTGGGTTTAAGTGCTACTTTCCCCTGTCCA |
| 0_hypothetical | O14 | Tb927.7.2690_for | GGGGACAAGTTTGTACAAAAAAGCAGGCTATGCACCGAAGGACATTCTTTCTT | Tb927.7.2690_rev | GGGGACCACTTTGTACAAGAAAGCTGGGTTCACCGTACCAGCGTATTGTGA |
| 0_hypothetical | O15 | Tb927.1.280_for | GGGGACAAGTTTGTACAAAAAAGCAGGCTATGTTTCAATTGGTTTCTGCGGC | Tb927.1.280_rev | GGGGACCACTTTGTACAAGAAAGCTGGGTCTACACTTCCTTAAACCGGAAGCTAAC |
| 0_hypothetical | O16 | Tb927.8.6310_for | GGGGACAAGTTTGTACAAAAAAGCAGGCTATGGCGGCACCACTAGCA | Tb927.8.6310_rev | GGGGACCACTTTGTACAAGAAAGCTGGGTTTACTTCTTCCCCAGTATGTCCCG |
| 0_hypothetical | O17 | Tb927.1.470_for | GGGGACAAGTTTGTACAAAAAAGCAGGCTATGTTTCAATTGGTTTCTGTGGCCG | Tb927.1.470_rev | GGGGACCACTTTGTACAAGAAAGCTGGGTCTACACTTCCTTAAACCGGAAGCTAAC |
| 0_hypothetical | O18 | Tb927.1.3810_for | GGGGACAAGTTTGTACAAAAAAGCAGGCTATGGCTCGTACTTTCGTCGTTTT | Tb927.1.3810_rev | GGGGACCACTTTGTACAAGAAAGCTGGGTCTAACCTGCGTAAACAGATGGTGC |
| 0_hypothetical | O19 | Tb927.10.2670_for | GGGGACAAGTTTGTACAAAAAAGCAGGCTATGGAGCAAAAGGAAAATGAGATCACT | Tb927.10.2670_rev | GGGGACCACTTTGTACAAGAAAGCTGGGTCTAGGCCGGAAGTACAATGGTTAATC |
| 0_hypothetical | O20 | Tb927.2.5720_for | GGGGACAAGTTTGTACAAAAAAGCAGGCTATGTTGCTGTCGCTGGCG | Tb927.2.5720_rev | GGGGACCACTTTGTACAAGAAAGCTGGGTTCATTCCCCCTGAAAGAAGCGG |
| 0_hypothetical | O21 | Tb927.9.15750_for | GGGGACAAGTTTGTACAAAAAAGCAGGCTATGAGCAACCTGACAGTGAGCA | Tb927.9.15750_rev | GGGGACCACTTTGTACAAGAAAGCTGGGTTCACAATCTCACCGTTGACCAGC |
| 0_hypothetical | O22 | Tb927.6.570_for | GGGGACAAGTTTGTACAAAAAAGCAGGCTATGGAACTGAACGAGCTGCAAC | Tb927.6.570_rev | GGGGACCACTTTGTACAAGAAAGCTGGGTTCATGTCCAAAGACACCTTTCATTTACT |
| 0_hypothetical | O23 | Tb927.4.2960_for | GGGGACAAGTTTGTACAAAAAAGCAGGCTATGAATGCGCGCGATGAACT | Tb927.4.2960_rev | GGGGACCACTTTGTACAAGAAAGCTGGGTCTAATGTCTGGCATACCGTTCTTTCA |
| 0_hypothetical | P1 | Tb927.8.2150_for | GGGGACAAGTTTGTACAAAAAAGCAGGCTATGACATGTAATGTGACGCCAGAG | Tb927.8.2150_rev | GGGGACCACTTTGTACAAGAAAGCTGGGTCTATGTTTCCACCTCGACGTCCT |
| 0_hypothetical | P3 | Tb927.8.7790_for | GGGGACAAGTTTGTACAAAAAAGCAGGCTATGCCTCCTGCCCCGCGC | Tb927.8.7790_rev | GGGGACCACTTTGTACAAGAAAGCTGGGTTTACAGATCATCATCATCGTCGAGTTTC |
| 0_hypothetical | P4 | Tb927.11.2930_for | GGGGACAAGTTTGTACAAAAAAGCAGGCTATGCTCCGCCGTTGCGTT | Tb927.11.2930_rev | GGGGACCACTTTGTACAAGAAAGCTGGGTCTAAGCGAGGTCGGTATCCGC |
| 0_hypothetical | P5 | Tb927.4.4630_for | GGGGACAAGTTTGTACAAAAAAGCAGGCTATGAGCGCTGATCCTGAAGTACC | Tb927.4.4630_rev | GGGGACCACTTTGTACAAGAAAGCTGGGTTTAAAAGAACTTGTACCATGCTGAGGC |
| 0_hypothetical | P6 | Tb927.4.4540_for | GGGGACAAGTTTGTACAAAAAAGCAGGCTATGCCGTCCATAACGCGC | Tb927.4.4540_rev | GGGGACCACTTTGTACAAGAAAGCTGGGTCTAGGGGTCTTCATCATCATCGAGT |
| 0_hypothetical | P7 | Tb927.11.10040_for | GGGGACAAGTTTGTACAAAAAAGCAGGCTATGATGCGCCGAACGCCG | Tb927.11.10040_rev | GGGGACCACTTTGTACAAGAAAGCTGGGTTTAACTATCTCCTTCGCCAGGGAG |
| 0_hypothetical | P8 | Tb927.2.1720_for | GGGGACAAGTTTGTACAAAAAAGCAGGCTATGGACAGCAACGTCACTTACG | Tb927.2.1720_rev | GGGGACCACTTTGTACAAGAAAGCTGGGTCTAATTCACTGGGGTCAGTTCAAAGC |
| 0_hypothetical | P9 | Tb927.7.2380_for | GGGGACAAGTTTGTACAAAAAAGCAGGCTATGGACAAGGACCTCGCGG | Tb927.7.2380_rev | GGGGACCACTTTGTACAAGAAAGCTGGGTTTACTCATCTGGAGAACCCTCCG |
| 0_hypothetical | P10 | Tb927.11.11840_for | GGGGACAAGTTTGTACAAAAAAGCAGGCTATGAACGAGCCTGATGCCATG | Tb927.11.11840_rev | GGGGACCACTTTGTACAAGAAAGCTGGGTTCACACCCATGCAGCTGTTG |
| 0_hypothetical | P11 | Tb927.10.13920_for | GGGGACAAGTTTGTACAAAAAAGCAGGCTATGTGGTGCTTTTCGTCGCC | Tb927.10.13920_rev | GGGGACCACTTTGTACAAGAAAGCTGGGTTCATCCTTTTCTCATTGCATGATTGCC |
| 0_hypothetical | P12 | Tb927.11.410_for | GGGGACAAGTTTGTACAAAAAAGCAGGCTATGTCGCGCAGTACAGTTTTTGG | Tb927.11.410_rev | GGGGACCACTTTGTACAAGAAAGCTGGGTTCACCGCTCGTAGCGCCA |
| 0_hypothetical | P13 | Tb927.4.120_for | GGGGACAAGTTTGTACAAAAAAGCAGGCTATGTTGTGGTGGAAACAGTGTGC | Tb927.4.120_rev | GGGGACCACTTTGTACAAGAAAGCTGGGTCTAAGTGGAACGCCAGGACG |
| 0_hypothetical | P14 | Tb927.5.1800_for | GGGGACAAGTTTGTACAAAAAAGCAGGCTATGGTGAACCTGCATGAGCTG | Tb927.5.1800_rev | GGGGACCACTTTGTACAAGAAAGCTGGGTTCAATCGAGTTTGCTCCCCATTG |
| 0_hypothetical | P15 | Tb927.8.1120_for | GGGGACAAGTTTGTACAAAAAAGCAGGCTATGAGCGGGGACGGCGTT | Tb927.8.1120_rev | GGGGACCACTTTGTACAAGAAAGCTGGGTCTAAACTTTCCCCAGAAACTTCCTGT |
| 0_hypothetical | P16 | Tb927.2.900_for | GGGGACAAGTTTGTACAAAAAAGCAGGCTATGCTGTGGCGGAAGCAG | Tb927.2.900_rev | GGGGACCACTTTGTACAAGAAAGCTGGGTCTAAGTGGAACGCCAGGACG |
| 0_hypothetical | P17 | Tb927.2.1250_for | GGGGACAAGTTTGTACAAAAAAGCAGGCTATGCTGTGGCGGAAGCAG | Tb927.2.1250_rev | GGGGACCACTTTGTACAAGAAAGCTGGGTCTAAGTGGAACATCGGGACGTG |
| 0_hypothetical | P18 | Tb927.10.490_for | GGGGACAAGTTTGTACAAAAAAGCAGGCTATGAAACAAATGTTTTATTGCAGCAAAGG | Tb927.10.490_rev | GGGGACCACTTTGTACAAGAAAGCTGGGTTCATTCACTCACATGGAAACAAAAGTACA |
| 0_hypothetical | P19 | Tb927.8.4690_for | GGGGACAAGTTTGTACAAAAAAGCAGGCTATGCTCGATAAAATCAAGACTATTGTCACT | Tb927.8.4690_rev | GGGGACCACTTTGTACAAGAAAGCTGGGTTCATTTAAACCGCGACGAAACGC |
| 0_hypothetical | P20 | Tb927.5.4370_for | GGGGACAAGTTTGTACAAAAAAGCAGGCTATGTCGCCTCTCGTATCGTCC | Tb927.5.4370_rev | GGGGACCACTTTGTACAAGAAAGCTGGGTTTAACATGGCTTCATCGGGTTGC |
| 0_hypothetical | P21 | Tb927.7.2210_for | GGGGACAAGTTTGTACAAAAAAGCAGGCTATGCGTCGCTCAGTGTTTTTCA | Tb927.7.2210_rev | GGGGACCACTTTGTACAAGAAAGCTGGGTTCACGCACAGGTTACTCCGT |
| 0_hypothetical | P22 | Tb927.11.800_for | GGGGACAAGTTTGTACAAAAAAGCAGGCTATGTCGCTGCTGAAGGGAAAG | Tb927.11.800_rev | GGGGACCACTTTGTACAAGAAAGCTGGGTCTATTCGTCTTCAAGATACACTTGACTACC |
| 0_hypothetical | P23 | Tb927.2.5260_for | GGGGACAAGTTTGTACAAAAAAGCAGGCTATGTCTGGTGGTGTTAATCCCACT | Tb927.2.5260_rev | GGGGACCACTTTGTACAAGAAAGCTGGGTTTACTCACGCCGACTCTTCTTTCG |
| 0_hypothetical | P24 | Tb927.3.945_for | GGGGACAAGTTTGTACAAAAAAGCAGGCTATGATGAGGCCATCGACGGG | Tb927.3.945_rev | GGGGACCACTTTGTACAAGAAAGCTGGGTCTAGAGGAATTTGGGTGGCTTCTG |
| 1_hypothetical | A1 | Tb927.10.120_for | GGGGACAAGTTTGTACAAAAAAGCAGGCTATGTTTTCCAAACGCTGTTTTGCA | Tb927.10.120_rev | GGGGACCACTTTGTACAAGAAAGCTGGGTTCATCGATATCCTGTATTTCTCCAAGAATC |
| 1_hypothetical | A2 | Tb927.8.1750_for | GGGGACAAGTTTGTACAAAAAAGCAGGCTATGTCATCCCCCGCTCCATC | Tb927.8.1750_rev | GGGGACCACTTTGTACAAGAAAGCTGGGTTTAGATATCATCAGTAAGATCCAACTTCAC |
| 1_hypothetical | A3 | Tb927.10.14650_for | GGGGACAAGTTTGTACAAAAAAGCAGGCTATGACTGGACTTACGGTGCCG | Tb927.10.14650_rev | GGGGACCACTTTGTACAAGAAAGCTGGGTTCATTCACCAAGACGAATATTCAATAACGG |
| 1_hypothetical | A4 | Tb927.10.15140_for | GGGGACAAGTTTGTACAAAAAAGCAGGCTATGTCATTGAGGGAGATTGAACTTGC | Tb927.10.15140_rev | GGGGACCACTTTGTACAAGAAAGCTGGGTCTACATCCCCATGTTCTTCATCGTC |
| 1_hypothetical | A6 | Tb927.8.5870_for | GGGGACAAGTTTGTACAAAAAAGCAGGCTATGTCACCATGGAAAAAGGAGGAGG | Tb927.8.5870_rev | GGGGACCACTTTGTACAAGAAAGCTGGGTCTACACTCCACGCAGGTACGA |
| 1_hypothetical | A7 | Tb927.7.840_for | GGGGACAAGTTTGTACAAAAAAGCAGGCTATGCTTCGCCGTAGCAGTG | Tb927.7.840_rev | GGGGACCACTTTGTACAAGAAAGCTGGGTTCACAACGGAGCGGGCGT |
| 1_hypothetical | A8 | Tb927.2.3315_for | GGGGACAAGTTTGTACAAAAAAGCAGGCTATGGCTAGCATGCTTCATGAAACT | Tb927.2.3315_rev | GGGGACCACTTTGTACAAGAAAGCTGGGTCTAGTAACTCTTACCCTTTCCGATTTCC |
| 1_hypothetical | A9 | Tb927.10.6430_for | GGGGACAAGTTTGTACAAAAAAGCAGGCTATGTTTCTAACGAGGCAGCACC | Tb927.10.6430_rev | GGGGACCACTTTGTACAAGAAAGCTGGGTCTAAAACCTGTCACCGGAGATGG |
| 1_hypothetical | A10 | Tb927.2.3275_for | GGGGACAAGTTTGTACAAAAAAGCAGGCTATGGCTGGCGTGCTTCATG | Tb927.2.3275_rev | GGGGACCACTTTGTACAAGAAAGCTGGGTCTAGTAACTCTTACCCTTTCCGATTTCC |
| 1_hypothetical | A11 | Tb927.8.5070_for | GGGGACAAGTTTGTACAAAAAAGCAGGCTATGGATAAGAGCAGCAAAAACTCCA | Tb927.8.5070_rev | GGGGACCACTTTGTACAAGAAAGCTGGGTTCAAGCCCCGACCTGCGC |
| 1_hypothetical | A15 | Tb927.10.16080_for | GGGGACAAGTTTGTACAAAAAAGCAGGCTATGTTCAGTCGCCGCGTG | Tb927.10.16080_rev | GGGGACCACTTTGTACAAGAAAGCTGGGTCTACATTTCGCTCGTCCGTCC |
| 1_hypothetical | A16 | Tb927.10.1670_for | GGGGACAAGTTTGTACAAAAAAGCAGGCTATGGGAAGTGGGGGTTCCC | Tb927.10.1670_rev | GGGGACCACTTTGTACAAGAAAGCTGGGTCTAGAAAATTGGGTCCCAGCGG |
| 1_hypothetical | A17 | Tb927.2.2770_for | GGGGACAAGTTTGTACAAAAAAGCAGGCTATGCCCGTTATTGTTGTAAACCTCG | Tb927.2.2770_rev | GGGGACCACTTTGTACAAGAAAGCTGGGTTTAAGCCCTACGTGAAAAGAAGAAGC |
| 1_hypothetical | A19 | Tb927.9.3410_for | GGGGACAAGTTTGTACAAAAAAGCAGGCTATGTTGCGCATCCGCCGA | Tb927.9.3410_rev | GGGGACCACTTTGTACAAGAAAGCTGGGTCTATCCATTCAGCCAACGCTTCA |
| 1_hypothetical | A20 | Tb927.10.750_for | GGGGACAAGTTTGTACAAAAAAGCAGGCTATGTTAAATAGCTGTTTGTGCAAAATCTTC | Tb927.10.750_rev | GGGGACCACTTTGTACAAGAAAGCTGGGTTCATGCACAACTCACTGAGCTG |
| 1_hypothetical | A21 | Tb927.3.2420_for | GGGGACAAGTTTGTACAAAAAAGCAGGCTATGGCTGGTGCAAAGGAAGTG | Tb927.3.2420_rev | GGGGACCACTTTGTACAAGAAAGCTGGGTTCAATTTCGCTTTTTCTCTCCTTTTGAC |
| 1_hypothetical | A22 | Tb927.9.17640_for | GGGGACAAGTTTGTACAAAAAAGCAGGCTATGCCACCATTGCAGTCCTG | Tb927.9.17640_rev | GGGGACCACTTTGTACAAGAAAGCTGGGTTCAGCCACATTTTGAGAAATCATTAAATCC |
| 1_hypothetical | A23 | Tb927.9.4550_for | GGGGACAAGTTTGTACAAAAAAGCAGGCTATGTTTGTAATGAAGTTACTGGATGCACT | Tb927.9.4550_rev | GGGGACCACTTTGTACAAGAAAGCTGGGTTCAAGAGTTTACCACATTGACCGATTC |
| 1_hypothetical | A24 | Tb927.9.15690_for | GGGGACAAGTTTGTACAAAAAAGCAGGCTATGCGTTCTACCGTCATTTGGAAT | Tb927.9.15690_rev | GGGGACCACTTTGTACAAGAAAGCTGGGTTCACCGTGTGTAACAAGTGAAGTTC |
| 1_hypothetical | B1 | Tb927.3.2475_for | GGGGACAAGTTTGTACAAAAAAGCAGGCTATGCCACCAAAGAAGAGCAGC | Tb927.3.2475_rev | GGGGACCACTTTGTACAAGAAAGCTGGGTCTATCGCTTGGAAGGCGCC |
| 1_hypothetical | B2 | Tb927.9.7270_for | GGGGACAAGTTTGTACAAAAAAGCAGGCTATGAGTAATAAAACATATTTGTGGTCCAAA | Tb927.9.7270_rev | GGGGACCACTTTGTACAAGAAAGCTGGGTTCACCTTCTCCGTGTGTACCG |
| 1_hypothetical | B3 | Tb927.11.15060.2_for | GGGGACAAGTTTGTACAAAAAAGCAGGCTATGGATTTAGCTGGTGACTTAGTGAAG | Tb927.11.15060.2_rev | GGGGACCACTTTGTACAAGAAAGCTGGGTTCATCGACCACGCTTCACCT |
| 1_hypothetical | B4 | Tb927.9.7080_for | GGGGACAAGTTTGTACAAAAAAGCAGGCTATGCCGGATTACACGCAGTTC | Tb927.9.7080_rev | GGGGACCACTTTGTACAAGAAAGCTGGGTTCACACGGGCCGACCGCG |
| 1_hypothetical | B5 | Tb927.9.11940_for | GGGGACAAGTTTGTACAAAAAAGCAGGCTATGCGTTCAGCTGTTAATCTCGG | Tb927.9.11940_rev | GGGGACCACTTTGTACAAGAAAGCTGGGTCTAAAAGAGTGCTGGAAACTTTGGGC |
| 1_hypothetical | B6 | Tb927.6.4060_for | GGGGACAAGTTTGTACAAAAAAGCAGGCTATGAGCGGTATCCGGTACCG | Tb927.6.4060_rev | GGGGACCACTTTGTACAAGAAAGCTGGGTCTAGTTGGGCATCAACTGCTCATAG |
| 1_hypothetical | B7 | Tb927.10.1640_for | GGGGACAAGTTTGTACAAAAAAGCAGGCTATGCACTTCTTTTCAAGTTACCGCT | Tb927.10.1640_rev | GGGGACCACTTTGTACAAGAAAGCTGGGTTTACCCTTCAAAGCACTCAGAAACG |
| 1_hypothetical | B8 | Tb927.5.510_for | GGGGACAAGTTTGTACAAAAAAGCAGGCTATGCGGCGCTCAATCATTCG | Tb927.5.510_rev | GGGGACCACTTTGTACAAGAAAGCTGGGTTCACTTCAATGCTTCAGGCCTTG |
| 1_hypothetical | B9 | Tb927.3.1525_for | GGGGACAAGTTTGTACAAAAAAGCAGGCTATGAGTTATATAATTGGCCATGTAATGAAA | Tb927.3.1525_rev | GGGGACCACTTTGTACAAGAAAGCTGGGTTTATTGCGATTCCCGTTCCTCCG |
| 1_hypothetical | B10 | Tb927.5.600_for | GGGGACAAGTTTGTACAAAAAAGCAGGCTATGGGCCGTGGATTCGGG | Tb927.5.600_rev | GGGGACCACTTTGTACAAGAAAGCTGGGTCTATGAACTCGCATGTCCCCG |
| 1_hypothetical | B11 | Tb927.10.15920_for | GGGGACAAGTTTGTACAAAAAAGCAGGCTATGTTTGGACGACGCGTTTACG | Tb927.10.15920_rev | GGGGACCACTTTGTACAAGAAAGCTGGGTTTATTCTCGGGTAGTTTGAAGGCG |
| 1_hypothetical | B12 | Tb927.8.1730_for | GGGGACAAGTTTGTACAAAAAAGCAGGCTATGGAACCGGCAGAGCATCG | Tb927.8.1730_rev | GGGGACCACTTTGTACAAGAAAGCTGGGTTCACGAGTCGCTGACATCCG |
| 1_hypothetical | B13 | Tb927.5.2670_for | GGGGACAAGTTTGTACAAAAAAGCAGGCTATGCCTCTATGGTCCTTTGTGGA | Tb927.5.2670_rev | GGGGACCACTTTGTACAAGAAAGCTGGGTTTACCCACACTTCAACCTTCCCT |
| 1_hypothetical | B14 | Tb927.7.4820_for | GGGGACAAGTTTGTACAAAAAAGCAGGCTATGAGGCTGCTCACTCATAATTTTCTATG | Tb927.7.4820_rev | GGGGACCACTTTGTACAAGAAAGCTGGGTCTAGTGTTCAGCAGCAATCATATCAGG |
| 1_hypothetical | B15 | Tb927.8.3730_for | GGGGACAAGTTTGTACAAAAAAGCAGGCTATGAGTGACTTGCAACCTATGAGC | Tb927.8.3730_rev | GGGGACCACTTTGTACAAGAAAGCTGGGTTTAACTCTTATTAGCTATTACCTGTCCAAC |
| 1_hypothetical | B16 | Tb927.7.5615_for | GGGGACAAGTTTGTACAAAAAAGCAGGCTATGTTTCTACATAATCGCGGCCTT | Tb927.7.5615_rev | GGGGACCACTTTGTACAAGAAAGCTGGGTTCACATGACCGCTAAAAGAATCATGA |
| 1_hypothetical | B17 | Tb927.11.8740_for | GGGGACAAGTTTGTACAAAAAAGCAGGCTATGGGAAGGATAGGGGACACTCC | Tb927.11.8740_rev | GGGGACCACTTTGTACAAGAAAGCTGGGTTTAGAAACCTGCAATTGCAACAGACA |
| 1_hypothetical | B18 | Tb927.4.5080_for | GGGGACAAGTTTGTACAAAAAAGCAGGCTATGGTCAGCCTTTTGGCATTGG | Tb927.4.5080_rev | GGGGACCACTTTGTACAAGAAAGCTGGGTCTACCTTTCTTTCTTCTCAAGGGCG |
| 1_hypothetical | B19 | Tb927.6.2220_for | GGGGACAAGTTTGTACAAAAAAGCAGGCTATGAGCGTACCTGAGTGGGC | Tb927.6.2220_rev | GGGGACCACTTTGTACAAGAAAGCTGGGTTCAGTTGACCGAAGTACATTGGGA |
| 1_hypothetical | B20 | Tb927.10.11770_for | GGGGACAAGTTTGTACAAAAAAGCAGGCTATGCTTTCAAGACAATTAGTAACGAGGT | Tb927.10.11770_rev | GGGGACCACTTTGTACAAGAAAGCTGGGTTCACACCTCCACAAAAAGCACC |
| 1_hypothetical | B21 | Tb927.10.8220_for | GGGGACAAGTTTGTACAAAAAAGCAGGCTATGCTTTCAAAACCAGTTGCACC | Tb927.10.8220_rev | GGGGACCACTTTGTACAAGAAAGCTGGGTTCATAAGTGCCCCAAGAGAACCA |
| 1_hypothetical | B22 | Tb927.11.7500_for | GGGGACAAGTTTGTACAAAAAAGCAGGCTATGGGCAAAGTAAGAGATACTCCGA | Tb927.11.7500_rev | GGGGACCACTTTGTACAAGAAAGCTGGGTTTAGAAACCTGCAATTGCAACAGACA |
| 1_hypothetical | B23 | Tb927.11.15680_for | GGGGACAAGTTTGTACAAAAAAGCAGGCTATGAGTGAGGTAGCCGATTTCAGT | Tb927.11.15680_rev | GGGGACCACTTTGTACAAGAAAGCTGGGTTCAGTTGGAAAGGCGTACGGT |
| 1_hypothetical | B24 | Tb927.3.1945_for | GGGGACAAGTTTGTACAAAAAAGCAGGCTATGGCTGCGTTGTTACCATTAGTT | Tb927.3.1945_rev | GGGGACCACTTTGTACAAGAAAGCTGGGTCTACAAATCATAATTACCTTCTTCATCTTG |
| 1_hypothetical | C2 | Tb927.7.1160_for | GGGGACAAGTTTGTACAAAAAAGCAGGCTATGCTGTGCCGTAGCCCG | Tb927.7.1160_rev | GGGGACCACTTTGTACAAGAAAGCTGGGTTCATAAGTCCTCCCCGTTCCATT |
| 1_hypothetical | C3 | Tb927.7.6420_for | GGGGACAAGTTTGTACAAAAAAGCAGGCTATGAGGTACATGGTTTTTGCTGCC | Tb927.7.6420_rev | GGGGACCACTTTGTACAAGAAAGCTGGGTCTAGGGGCAAGGGGAAAGGT |
| 1_hypothetical | C4 | Tb927.10.3160_for | GGGGACAAGTTTGTACAAAAAAGCAGGCTATGCGTTACTATCCCACTGGTCT | Tb927.10.3160_rev | GGGGACCACTTTGTACAAGAAAGCTGGGTCTACATCTCTATGCTCAACCCCCG |
| 1_hypothetical | C5 | Tb927.11.14165_for | GGGGACAAGTTTGTACAAAAAAGCAGGCTATGTCCATAAGTCAAGTGGATCTCGA | Tb927.11.14165_rev | GGGGACCACTTTGTACAAGAAAGCTGGGTTCAAAGGTTTGACGGATCATCTGAATG |
| 1_hypothetical | C6 | Tb927.11.13570_for | GGGGACAAGTTTGTACAAAAAAGCAGGCTATGAGTGAAGGGCGCTTGGA | Tb927.11.13570_rev | GGGGACCACTTTGTACAAGAAAGCTGGGTTTAAACGTGCTCGGTGTCCATG |
| 1_hypothetical | C7 | Tb927.10.14050_for | GGGGACAAGTTTGTACAAAAAAGCAGGCTATGCTGTGGGGTTCTAGAAAGGG | Tb927.10.14050_rev | GGGGACCACTTTGTACAAGAAAGCTGGGTCTACAAACAAGCTGACAGACAAAAAAATGG |
| 1_hypothetical | C8 | Tb11.02.5105_for | GGGGACAAGTTTGTACAAAAAAGCAGGCTATGGACGTGGTTGAAAAATTTCTTTCTC | Tb11.02.5105_rev | GGGGACCACTTTGTACAAGAAAGCTGGGTTCATGAAACCCCAAGCACGTG |
| 1_hypothetical | C9 | Tb927.11.11078_for | GGGGACAAGTTTGTACAAAAAAGCAGGCTATGCCTCACCGCTCCACT | Tb927.11.11078_rev | GGGGACCACTTTGTACAAGAAAGCTGGGTCTACCGTCCCTCTGATTGTTCTTCA |
| 1_hypothetical | C10 | Tb927.9.8580_for | GGGGACAAGTTTGTACAAAAAAGCAGGCTATGGTGTCATGGTTTTGGACCAC | Tb927.9.8580_rev | GGGGACCACTTTGTACAAGAAAGCTGGGTTTACTCACTAACGCCCTCAAACAGC |
| 1_hypothetical | C11 | Tb927.6.2590_for | GGGGACAAGTTTGTACAAAAAAGCAGGCTATGATTCGCCGATTAGGCAGTG | Tb927.6.2590_rev | GGGGACCACTTTGTACAAGAAAGCTGGGTCTAAAAAGGCCCAAAGCGCTTATTT |
| 1_hypothetical | C12 | Tb927.9.1640_for | GGGGACAAGTTTGTACAAAAAAGCAGGCTATGTGGCGTACTGCTGTCTTTTG | Tb927.9.1640_rev | GGGGACCACTTTGTACAAGAAAGCTGGGTCTACATGTCATCCATTCGTTCCTCTTC |
| 1_hypothetical | C13 | Tb927.11.15375_for | GGGGACAAGTTTGTACAAAAAAGCAGGCTATGGAGGTATCGGCGGATACAC | Tb927.11.15375_rev | GGGGACCACTTTGTACAAGAAAGCTGGGTTCACGATTCTTTCGTTCTTGTTCCT |
| 1_hypothetical | C14 | Tb927.10.320_for | GGGGACAAGTTTGTACAAAAAAGCAGGCTATGGACCCTGCCGATCCC | Tb927.10.320_rev | GGGGACCACTTTGTACAAGAAAGCTGGGTCTATGCAACCGCAGGTCTCTG |
| 1_hypothetical | C15 | Tb927.3.3140_for | GGGGACAAGTTTGTACAAAAAAGCAGGCTATGTGCCACGGGGGTCGC | Tb927.3.3140_rev | GGGGACCACTTTGTACAAGAAAGCTGGGTTCACCTCTCAACTCTGCGGTT |
| 1_hypothetical | C16 | Tb927.9.10650_for | GGGGACAAGTTTGTACAAAAAAGCAGGCTATGGAACAACAACAACAACAACCAG | Tb927.9.10650_rev | GGGGACCACTTTGTACAAGAAAGCTGGGTTCACATCCTGTAATCTCTTCTGTTTTCG |
| 1_hypothetical | C17 | Tb927.8.5900_for | GGGGACAAGTTTGTACAAAAAAGCAGGCTATGTTGATGGTTGTCGGACCTGA | Tb927.8.5900_rev | GGGGACCACTTTGTACAAGAAAGCTGGGTCTAACAGGGGGAGAAGCGTTGG |
| 1_hypothetical | C18 | Tb927.4.2170_for | GGGGACAAGTTTGTACAAAAAAGCAGGCTATGTCTCCCAAACTGTATGTTTGCA | Tb927.4.2170_rev | GGGGACCACTTTGTACAAGAAAGCTGGGTTCAATTGCTACTGTCTGCTCCATAAC |
| 1_hypothetical | C19 | Tb927.11.15360_for | GGGGACAAGTTTGTACAAAAAAGCAGGCTATGGGGGGTAGTTCGAGTCG | Tb927.11.15360_rev | GGGGACCACTTTGTACAAGAAAGCTGGGTTTATTCGGCCACCTCGTCTTCC |
| 1_hypothetical | C20 | Tb927.10.7720_for | GGGGACAAGTTTGTACAAAAAAGCAGGCTATGTGGCTCCGCTCCAGAA | Tb927.10.7720_rev | GGGGACCACTTTGTACAAGAAAGCTGGGTTTACTTCCCCCCTAGAACAGCAC |
| 1_hypothetical | C21 | Tb927.11.9580.1_for | GGGGACAAGTTTGTACAAAAAAGCAGGCTATGGGCGAGGAGGTACTGTC | Tb927.11.9580.1_rev | GGGGACCACTTTGTACAAGAAAGCTGGGTCTACCGCTGGACAATTTCCGC |
| 1_hypothetical | C22 | Tb927.8.3360_for | GGGGACAAGTTTGTACAAAAAAGCAGGCTATGCTGTGGGGTGCGGTT | Tb927.8.3360_rev | GGGGACCACTTTGTACAAGAAAGCTGGGTCTACATTACCAACCGCCGTCG |
| 1_hypothetical | C23 | Tb927.4.4120_for | GGGGACAAGTTTGTACAAAAAAGCAGGCTATGTACTTCGAAGAGGAAGAGAATAATGAT | Tb927.4.4120_rev | GGGGACCACTTTGTACAAGAAAGCTGGGTCTATACCCAACTGCTATACCCCGT |
| 1_hypothetical | C24 | Tb927.9.12280_for | GGGGACAAGTTTGTACAAAAAAGCAGGCTATGGGTAGAAATGTAACTGCGGTTAC | Tb927.9.12280_rev | GGGGACCACTTTGTACAAGAAAGCTGGGTTTATTTTTTGCTCCTTGATTTTTTCCTGCC |
| 1_hypothetical | D2 | Tb927.11.6700_for | GGGGACAAGTTTGTACAAAAAAGCAGGCTATGGTCAGTCGTTCGGGGT | Tb927.11.6700_rev | GGGGACCACTTTGTACAAGAAAGCTGGGTTCAGTCAACCCATCGCCGC |
| 1_hypothetical | D3 | Tb927.11.8420_for | GGGGACAAGTTTGTACAAAAAAGCAGGCTATGGCTGGCATAAACACGGTTG | Tb927.11.8420_rev | GGGGACCACTTTGTACAAGAAAGCTGGGTTTAGGGTGTGTATTCTTCGTCATCTTCA |
| 1_hypothetical | D4 | Tb927.11.8750_for | GGGGACAAGTTTGTACAAAAAAGCAGGCTATGAGCGCACAAGGGGATCC | Tb927.11.8750_rev | GGGGACCACTTTGTACAAGAAAGCTGGGTTCAATCGTCATTGTCATCACTAGAATCAG |
| 1_hypothetical | D5 | Tb927.11.13590_for | GGGGACAAGTTTGTACAAAAAAGCAGGCTATGATGAAAAATACCGTGCGGCA | Tb927.11.13590_rev | GGGGACCACTTTGTACAAGAAAGCTGGGTTTACGTGGTGCTTTTTTCCCTTTGA |
| 1_hypothetical | D6 | Tb927.8.5320_for | GGGGACAAGTTTGTACAAAAAAGCAGGCTATGCCGTTGGGCACTGATG | Tb927.8.5320_rev | GGGGACCACTTTGTACAAGAAAGCTGGGTTCAGTCAGTGGCCCGTTCA |
| 1_hypothetical | D7 | Tb927.11.16450_for | GGGGACAAGTTTGTACAAAAAAGCAGGCTATGGTTTCCAAAACGCGGCG | Tb927.11.16450_rev | GGGGACCACTTTGTACAAGAAAGCTGGGTCTACTTCACTTTCTTTTTCAACTTTAGTTT |
| 1_hypothetical | D8 | Tb927.10.12300_for | GGGGACAAGTTTGTACAAAAAAGCAGGCTATGGCTAACTTTGTGCGCAAAGG | Tb927.10.12300_rev | GGGGACCACTTTGTACAAGAAAGCTGGGTCTAGTACCCCGGCGGACG |
| 1_hypothetical | D9 | Tb927.2.4445_for | GGGGACAAGTTTGTACAAAAAAGCAGGCTATGTCGACTGAGGCTACCAATGA | Tb927.2.4445_rev | GGGGACCACTTTGTACAAGAAAGCTGGGTCTAGCGGCGCTGTTCCTG |
| 1_hypothetical | D10 | Tb927.4.2160_for | GGGGACAAGTTTGTACAAAAAAGCAGGCTATGACCGAGTCGTACCGATCG | Tb927.4.2160_rev | GGGGACCACTTTGTACAAGAAAGCTGGGTCTACACTTCAAGCATCTCGCAATCA |
| 1_hypothetical | D11 | Tb927.10.440_for | GGGGACAAGTTTGTACAAAAAAGCAGGCTATGTCCAACCGAGGCTTTACCC | Tb927.10.440_rev | GGGGACCACTTTGTACAAGAAAGCTGGGTTCAAAGAGTTGCATCCGTAGCAG |
| 1_hypothetical | D12 | Tb927.4.1420_for | GGGGACAAGTTTGTACAAAAAAGCAGGCTATGTTGTGCCGGTCATCACTG | Tb927.4.1420_rev | GGGGACCACTTTGTACAAGAAAGCTGGGTTCAACTAAGTTTCGCGCCCAC |
| 1_hypothetical | D13 | Tb927.3.2370_for | GGGGACAAGTTTGTACAAAAAAGCAGGCTATGAACAACAAGGCACCGGG | Tb927.3.2370_rev | GGGGACCACTTTGTACAAGAAAGCTGGGTTTATTTGCTTCGGTATGTCTGGATCTC |
| 1_hypothetical | D14 | Tb927.11.2070_for | GGGGACAAGTTTGTACAAAAAAGCAGGCTATGGTGCGCTCGCTTGTTG | Tb927.11.2070_rev | GGGGACCACTTTGTACAAGAAAGCTGGGTTCATACGTCACGGGACATCTCA |
| 1_hypothetical | D15 | Tb927.9.10510_for | GGGGACAAGTTTGTACAAAAAAGCAGGCTATGCACACACTGCTGGTTACC | Tb927.9.10510_rev | GGGGACCACTTTGTACAAGAAAGCTGGGTTCAACCTTTGGCATCTTCCCTATAAGC |
| 1_hypothetical | D16 | Tb927.6.4360_for | GGGGACAAGTTTGTACAAAAAAGCAGGCTATGCGGGAGAGTAATATACACATGTCG | Tb927.6.4360_rev | GGGGACCACTTTGTACAAGAAAGCTGGGTTTACCCCTCTGTGTTTGGAATGGG |
| 1_hypothetical | D17 | Tb927.10.3040_for | GGGGACAAGTTTGTACAAAAAAGCAGGCTATGTTTTCACGAAGCTTTGTCGGA | Tb927.10.3040_rev | GGGGACCACTTTGTACAAGAAAGCTGGGTCTAGTACTTGAAGATGATGCTCTCAATAGG |
| 1_hypothetical | D18 | Tb927.11.1650_for | GGGGACAAGTTTGTACAAAAAAGCAGGCTATGCTCCGGTGTTCGACTTTTT | Tb927.11.1650_rev | GGGGACCACTTTGTACAAGAAAGCTGGGTTCATTCCTCACACTCTTCCTCTAAAAGA |
| 1_hypothetical | D19 | Tb927.5.4390_for | GGGGACAAGTTTGTACAAAAAAGCAGGCTATGTCTGGAGTTGTTATAAAGAAGCCC | Tb927.5.4390_rev | GGGGACCACTTTGTACAAGAAAGCTGGGTCTACTCTCTGGAGGTAATCATTCGGT |
| 1_hypothetical | D20 | Tb927.8.3090_for | GGGGACAAGTTTGTACAAAAAAGCAGGCTATGCAAGTCTCTCCACGTGTGC | Tb927.8.3090_rev | GGGGACCACTTTGTACAAGAAAGCTGGGTCTAACTTGAAAACGGCTTGTTTTTATCCC |
| 1_hypothetical | D21 | Tb927.4.3780_for | GGGGACAAGTTTGTACAAAAAAGCAGGCTATGTTTAGAAACCGTTTTTTCATTAGGCTT | Tb927.4.3780_rev | GGGGACCACTTTGTACAAGAAAGCTGGGTTCAACTTTCATCGATACCTTTTGAAAGGC |
| 1_hypothetical | D22 | Tb927.11.13660_for | GGGGACAAGTTTGTACAAAAAAGCAGGCTATGAGCCACCCTAAGATGAAGCT | Tb927.11.13660_rev | GGGGACCACTTTGTACAAGAAAGCTGGGTTTAGATCCCAACTCTTGCGGGA |
| 1_hypothetical | D23 | Tb927.10.4790_for | GGGGACAAGTTTGTACAAAAAAGCAGGCTATGTCCCACGATGGGGATGATG | Tb927.10.4790_rev | GGGGACCACTTTGTACAAGAAAGCTGGGTTTACTTCCCCTTCTGCAACTTTAACAG |
| 1_hypothetical | D24 | Tb927.11.7430_for | GGGGACAAGTTTGTACAAAAAAGCAGGCTATGAGCCGTGCCAACGTG | Tb927.11.7430_rev | GGGGACCACTTTGTACAAGAAAGCTGGGTTTAACATTGGAGTACCTGCGCG |
| 1_hypothetical | E1 | Tb927.10.7660_for | GGGGACAAGTTTGTACAAAAAAGCAGGCTATGTACAACTGCAGTTCCAAGTTTAGG | Tb927.10.7660_rev | GGGGACCACTTTGTACAAGAAAGCTGGGTTCACCGAAACGGGAAAAAGGC |
| 1_hypothetical | E2 | Tb927.7.2600_for | GGGGACAAGTTTGTACAAAAAAGCAGGCTATGCCTACGGTGCAGCCA | Tb927.7.2600_rev | GGGGACCACTTTGTACAAGAAAGCTGGGTTTAGTCCTCACCACCAAACTCCTG |

| | | | | | |
|---|---|---|---|---|---|
| 1_hypothetical | E3 | Tb927.10.2830_for | GGGGACAAGTTTGTACAAAAAAGCAGGCTATGCATCCCCGTGTTGCTAAAC | Tb927.10.2830_rev | GGGGACCACTTTGTACAAGAAAGCTGGGTCTATTTTTCAGAAAAATCACCCTCACCCC |
| 1_hypothetical | E4 | Tb927.3.2910_for | GGGGACAAGTTTGTACAAAAAAGCAGGCTATGGGAAGGCAGTCCCAGAAG | Tb927.3.2910_rev | GGGGACCACTTTGTACAAGAAAGCTGGGTTTATCTGACGGAGGACATCATTTTCTCC |
| 1_hypothetical | E5 | Tb927.9.720_for | GGGGACAAGTTTGTACAAAAAAGCAGGCTATGCTTAAAATTGCAGCAGTCATTTTGT | Tb927.9.720_rev | GGGGACCACTTTGTACAAGAAAGCTGGGTCTATTGTTTTTTTGTTCTTTTTGACTTTTG |
| 1_hypothetical | E6 | Tb927.3.3880_for | GGGGACAAGTTTGTACAAAAAAGCAGGCTATGAGCAACCGGGAGCCA | Tb927.3.3880_rev | GGGGACCACTTTGTACAAGAAAGCTGGGTCTACTTCTGTGACGAAGCGGC |
| 1_hypothetical | E7 | Tb927.10.4810_for | GGGGACAAGTTTGTACAAAAAAGCAGGCTATGCAGGTAATTGACATGTTTGCTCT | Tb927.10.4810_rev | GGGGACCACTTTGTACAAGAAAGCTGGGTTTATTTACGGAATGACACTACACGATCAA |
| 1_hypothetical | E8 | Tb927.11.13610_for | GGGGACAAGTTTGTACAAAAAAGCAGGCTATGGGTCAGAGCACATCTTACGA | Tb927.11.13610_rev | GGGGACCACTTTGTACAAGAAAGCTGGGTTTAGAGGTCCTCCTCTTGCTCTCC |
| 1_hypothetical | E9 | Tb927.10.5800_for | GGGGACAAGTTTGTACAAAAAAGCAGGCTATGGCGGGACCTTCCAATGC | Tb927.10.5800_rev | GGGGACCACTTTGTACAAGAAAGCTGGGTCTAAAGTTGGTTTTCACACCACGGA |
| 1_hypothetical | E10 | Tb927.10.140_for | GGGGACAAGTTTGTACAAAAAAGCAGGCTATGAATGAGAAGCAACATTTTCTTCCTATG | Tb927.10.140_rev | GGGGACCACTTTGTACAAGAAAGCTGGGTTCAGCCCCACGTATTCCAGAG |
| 1_hypothetical | E11 | Tb927.11.9500_for | GGGGACAAGTTTGTACAAAAAAGCAGGCTATGGTCTCCGCAAATAAATCCACAA | Tb927.11.9500_rev | GGGGACCACTTTGTACAAGAAAGCTGGGTCTACGATGCCATGTCTTTGTTGTCC |
| 1_hypothetical | E12 | Tb927.3.4170_for | GGGGACAAGTTTGTACAAAAAAGCAGGCTATGTCCAGCAATAATCCCTCCGA | Tb927.3.4170_rev | GGGGACCACTTTGTACAAGAAAGCTGGGTCTAACGAATGCGTGCAAAATCCG |
| 1_hypothetical | E13 | Tb927.3.3680_for | GGGGACAAGTTTGTACAAAAAAGCAGGCTATGCCGCCGGTTACGGGC | Tb927.3.3680_rev | GGGGACCACTTTGTACAAGAAAGCTGGGTTTATGTCGATTTGGGGTCGCG |
| 1_hypothetical | E14 | Tb927.9.2290_for | GGGGACAAGTTTGTACAAAAAAGCAGGCTATGCCCTTAAATAATGTACAGGTGGAG | Tb927.9.2290_rev | GGGGACCACTTTGTACAAGAAAGCTGGGTTTATTCTACCGTATGCAAGGGAAGATCG |
| 1_hypothetical | E15 | Tb927.8.5360_for | GGGGACAAGTTTGTACAAAAAAGCAGGCTATGGTTGTGGTCGTGGGCG | Tb927.8.5360_rev | GGGGACCACTTTGTACAAGAAAGCTGGGTTCACTGTGGTTGCTCATGCC |
| 1_hypothetical | E16 | Tb927.1.3750_for | GGGGACAAGTTTGTACAAAAAAGCAGGCTATGTTGATTGGCATGAAATATTCTGCTC | Tb927.1.3750_rev | GGGGACCACTTTGTACAAGAAAGCTGGGTCTACCATCGTACATATGTTTTTTTTTTGG |
| 1_hypothetical | E17 | Tb927.10.8670_for | GGGGACAAGTTTGTACAAAAAAGCAGGCTATGCTGCGCCTCATTGCA | Tb927.10.8670_rev | GGGGACCACTTTGTACAAGAAAGCTGGGTTCACCTTTTCTGCAAACGCCA |
| 1_hypothetical | E18 | Tb927.5.2840_for | GGGGACAAGTTTGTACAAAAAAGCAGGCTATGCTTCGGCGGTTCGTTG | Tb927.5.2840_rev | GGGGACCACTTTGTACAAGAAAGCTGGGTTCAACCACGCCACTTCTTACTGT |
| 1_hypothetical | E19 | Tb927.7.7440_for | GGGGACAAGTTTGTACAAAAAAGCAGGCTATGTCACGAAAAGCTCTTACAAGACC | Tb927.7.7440_rev | GGGGACCACTTTGTACAAGAAAGCTGGGTTTACAAAACACCACGTCTGTGGTT |
| 1_hypothetical | E20 | Tb927.10.14100_for | GGGGACAAGTTTGTACAAAAAAGCAGGCTATGCTCACGCAAGCGGGT | Tb927.10.14100_rev | GGGGACCACTTTGTACAAGAAAGCTGGGTTCAGCAGCCTGTGATCCCTC |
| 1_hypothetical | E21 | Tb927.7.4925_for | GGGGACAAGTTTGTACAAAAAAGCAGGCTATGTTATTACTGCAGACACTGATGTTAATG | Tb927.7.4925_rev | GGGGACCACTTTGTACAAGAAAGCTGGGTCTAAGACACCTTCAAGCCAACAATCA |
| 1_hypothetical | E22 | Tb927.5.4510_for | GGGGACAAGTTTGTACAAAAAAGCAGGCTATGCTCAGGTTAAATAAATTTTTTTCTCAT | Tb927.5.4510_rev | GGGGACCACTTTGTACAAGAAAGCTGGGTTCACTTTTTAAACTTCGATCGACCCTT |
| 1_hypothetical | E23 | Tb927.11.650_for | GGGGACAAGTTTGTACAAAAAAGCAGGCTATGTCAGGCCCGGTTGTGG | Tb927.11.650_rev | GGGGACCACTTTGTACAAGAAAGCTGGGTCTATATTGCAGGAACTTTTCGTGTTTCAA |
| 1_hypothetical | E24 | Tb927.6.2280_for | GGGGACAAGTTTGTACAAAAAAGCAGGCTATGTCTGCACTTGCGGAGGC | Tb927.6.2280_rev | GGGGACCACTTTGTACAAGAAAGCTGGGTTTACTGTTTCTGCTGCTGCTGTT |
| 1_hypothetical | F2 | Tb927.11.8850_for | GGGGACAAGTTTGTACAAAAAAGCAGGCTATGCGGGCACTATACCAGTTCG | Tb927.11.8850_rev | GGGGACCACTTTGTACAAGAAAGCTGGGTTCACAAGAATGAATTGCTATCTTGTGTTTT |
| 1_hypothetical | F3 | Tb927.10.5550_for | GGGGACAAGTTTGTACAAAAAAGCAGGCTATGCGGCGCTCGTTCGTC | Tb927.10.5550_rev | GGGGACCACTTTGTACAAGAAAGCTGGGTCTACTTCGGATGTATTTTGCACACATC |
| 1_hypothetical | F4 | Tb927.10.4420_for | GGGGACAAGTTTGTACAAAAAAGCAGGCTATGAGAAAAAAGAGGGAAACAAAAAAGAAA | Tb927.10.4420_rev | GGGGACCACTTTGTACAAGAAAGCTGGGTTTAAAAGAAAGAAAGGAAAGAAAGAAAAAA |
| 1_hypothetical | F5 | Tb927.10.7820_for | GGGGACAAGTTTGTACAAAAAAGCAGGCTATGAGGCTAACCAACACGAAACC | Tb927.10.7820_rev | GGGGACCACTTTGTACAAGAAAGCTGGGTTTACTCTTTCTCCTTCCGAGAAAATGAC |
| 1_hypothetical | F6 | Tb927.10.5100_for | GGGGACAAGTTTGTACAAAAAAGCAGGCTATGTTTGTATTTGTGGTGTGTCCACG | Tb927.10.5100_rev | GGGGACCACTTTGTACAAGAAAGCTGGGTTTATTTAAAAAAAAATAAAAGAGAAGAGAA |
| 1_hypothetical | F7 | Tb927.5.4490_for | GGGGACAAGTTTGTACAAAAAAGCAGGCTATGGTACTGATGTCAAAACGAGATGC | Tb927.5.4490_rev | GGGGACCACTTTGTACAAGAAAGCTGGGTCTAACCAAATACCCTTTGCGCAGA |
| 1_hypothetical | F8 | Tb927.9.12870_for | GGGGACAAGTTTGTACAAAAAAGCAGGCTATGACGTTGTATTCTATTTACATCTTCAAT | Tb927.9.12870_rev | GGGGACCACTTTGTACAAGAAAGCTGGGTCTAAAGGATCTTCTGAGAGTGGAAAAACT |
| 1_hypothetical | F9 | Tb927.9.12890_for | GGGGACAAGTTTGTACAAAAAAGCAGGCTATGACGAAACTGGTGCGCAA | Tb927.9.12890_rev | GGGGACCACTTTGTACAAGAAAGCTGGGTTTAATCATGCAGATCCTCATTCCGG |
| 1_hypothetical | F10 | Tb927.8.4535_for | GGGGACAAGTTTGTACAAAAAAGCAGGCTATGGAGGGGCTTGGATTAGAAACG | Tb927.8.4535_rev | GGGGACCACTTTGTACAAGAAAGCTGGGTTTAACACAACTCAAGCTCAAAAGCTGG |
| 1_hypothetical | F11 | Tb927.8.3450_for | GGGGACAAGTTTGTACAAAAAAGCAGGCTATGCGGGGCTCTACGGATC | Tb927.8.3450_rev | GGGGACCACTTTGTACAAGAAAGCTGGGTCTAAGAGGTCCCTACCACCTCCC |
| 1_hypothetical | F12 | Tb927.9.14990_for | GGGGACAAGTTTGTACAAAAAAGCAGGCTATGTGGGCTTCTGGCTCTCG | Tb927.9.14990_rev | GGGGACCACTTTGTACAAGAAAGCTGGGTCTAGTACGATGCTTGCTTAACACAACC |
| 1_hypothetical | F13 | Tb927.9.1590_for | GGGGACAAGTTTGTACAAAAAAGCAGGCTATGCCAGTACCGGTTGTGGA | Tb927.9.1590_rev | GGGGACCACTTTGTACAAGAAAGCTGGGTTTACAGTTTTCCAATTGCATGGTACCA |
| 1_hypothetical | F14 | Tb927.4.2420_for | GGGGACAAGTTTGTACAAAAAAGCAGGCTATGACTCTTGCGGATACGCTACA | Tb927.4.2420_rev | GGGGACCACTTTGTACAAGAAAGCTGGGTTCATGAAACAGTCGGCTTCATACG |
| 1_hypothetical | F15 | Tb927.1.380_for | GGGGACAAGTTTGTACAAAAAAGCAGGCTATGTTTCAATTGGTTTCTGTGGCCG | Tb927.1.380_rev | GGGGACCACTTTGTACAAGAAAGCTGGGTTTACTGACCCAGTGCCTGAAGC |
| 1_hypothetical | F16 | Tb927.6.120_for | GGGGACAAGTTTGTACAAAAAAGCAGGCTATGTTTCAATTGGTTTCTGCGGC | Tb927.6.120_rev | GGGGACCACTTTGTACAAGAAAGCTGGGTTTACTGACCCAGTGCCTGAAGC |
| 1_hypothetical | F18 | Tb927.6.4570_for | GGGGACAAGTTTGTACAAAAAAGCAGGCTATGCTGAGATTATGTAGAAGACTATCATAC | Tb927.6.4570_rev | GGGGACCACTTTGTACAAGAAAGCTGGGTTCAGTACGACTCCTTTGTGTGAATG |
| 1_hypothetical | F19 | Tb927.2.680_for | GGGGACAAGTTTGTACAAAAAAGCAGGCTATGTTTCAATTGGTTTCTGTGGCCG | Tb927.2.680_rev | GGGGACCACTTTGTACAAGAAAGCTGGGTTTACTGACCCAGTGCCTGGAG |
| 1_hypothetical | F20 | Tb927.6.2310_for | GGGGACAAGTTTGTACAAAAAAGCAGGCTATGCTTTTCATGGTTTACACGTCGC | Tb927.6.2310_rev | GGGGACCACTTTGTACAAGAAAGCTGGGTCTACAGCAGCAAATTAGAAATACACAAACC |
| 1_hypothetical | F21 | Tb927.8.4630_for | GGGGACAAGTTTGTACAAAAAAGCAGGCTATGATTCTCGGTCTCGGTGATGG | Tb927.8.4630_rev | GGGGACCACTTTGTACAAGAAAGCTGGGTTTAGCGATACCATGTTGGGTGC |
| 1_hypothetical | F22 | Tb927.11.13790_for | GGGGACAAGTTTGTACAAAAAAGCAGGCTATGTTATGGAGAAATATCTCTCGATGCC | Tb927.11.13790_rev | GGGGACCACTTTGTACAAGAAAGCTGGGTTCACAACAGCTTCCGACGTCC |
| 1_hypothetical | F23 | Tb927.10.5860_for | GGGGACAAGTTTGTACAAAAAAGCAGGCTATGGAGTTGAATGAGCAGGAACTG | Tb927.10.5860_rev | GGGGACCACTTTGTACAAGAAAGCTGGGTTCAATTTTCCTTGGCAACACTTGC |
| 1_hypothetical | F24 | Tb927.10.7460_for | GGGGACAAGTTTGTACAAAAAAGCAGGCTATGGCGACATTGAATGTTAGGTGT | Tb927.10.7460_rev | GGGGACCACTTTGTACAAGAAAGCTGGGTCTAGGCCCCCACGCTAACG |
| 1_hypothetical | G1 | Tb927.6.2600_for | GGGGACAAGTTTGTACAAAAAAGCAGGCTATGAGGAAACAATTAAATATGCAGGAGGA | Tb927.6.2600_rev | GGGGACCACTTTGTACAAGAAAGCTGGGTTTAACTAAAAAATGTGCCCATGCTCTG |
| 1_hypothetical | G2 | Tb927.10.12010_for | GGGGACAAGTTTGTACAAAAAAGCAGGCTATGTTACTCGATTTCATTTTCAGTCACCT | Tb927.10.12010_rev | GGGGACCACTTTGTACAAGAAAGCTGGGTTTATTCCACCTTTTCATCAACCTTCTCTG |
| 1_hypothetical | G3 | Tb927.5.645_for | GGGGACAAGTTTGTACAAAAAAGCAGGCTATGTTAGTGTCCAAACAAATAAGCCCT | Tb927.5.645_rev | GGGGACCACTTTGTACAAGAAAGCTGGGTTCAGCTGAAGGTTCGACACAAC |
| 1_hypothetical | G4 | Tb927.4.1470_for | GGGGACAAGTTTGTACAAAAAAGCAGGCTATGCGGCACTTTGGACCTTC | Tb927.4.1470_rev | GGGGACCACTTTGTACAAGAAAGCTGGGTTCAGTACTTTGTACGCATGTTGAACA |
| 1_hypothetical | G5 | Tb927.6.1420_for | GGGGACAAGTTTGTACAAAAAAGCAGGCTATGAGCAAACTAACACCGCGC | Tb927.6.1420_rev | GGGGACCACTTTGTACAAGAAAGCTGGGTTTACAAACCAGTAACAGCAAACTGTTTTG |
| 1_hypothetical | G6 | Tb927.11.13620_for | GGGGACAAGTTTGTACAAAAAAGCAGGCTATGTTTTTCGTAACGCCGCGT | Tb927.11.13620_rev | GGGGACCACTTTGTACAAGAAAGCTGGGTTCACCCTTTGCAACCATCCG |
| 1_hypothetical | G7 | Tb927.11.7090_for | GGGGACAAGTTTGTACAAAAAAGCAGGCTATGTCCGAAGAGTTGCGGTGG | Tb927.11.7090_rev | GGGGACCACTTTGTACAAGAAAGCTGGGTTCAAATGCTTTGCTTAGCGATGC |
| 1_hypothetical | G8 | Tb927.8.1040_for | GGGGACAAGTTTGTACAAAAAAGCAGGCTATGTCTTCGTCTCAGCAATTGGG | Tb927.8.1040_rev | GGGGACCACTTTGTACAAGAAAGCTGGGTTCAGTGAAACCGAGTTCCACAATAAC |
| 1_hypothetical | G9 | Tb927.11.14660_for | GGGGACAAGTTTGTACAAAAAAGCAGGCTATGAACGATATTTACGCACGGCG | Tb927.11.14660_rev | GGGGACCACTTTGTACAAGAAAGCTGGGTTTACTCACCTTCGATAAGAGGTTCTTCC |
| 1_hypothetical | G10 | Tb927.4.550_for | GGGGACAAGTTTGTACAAAAAAGCAGGCTATGTACGGCAACAAAGGACCTC | Tb927.4.550_rev | GGGGACCACTTTGTACAAGAAAGCTGGGTTCACTCCTCAGTTTTAAACGTTTTCTTCG |
| 1_hypothetical | G11 | Tb927.4.2320_for | GGGGACAAGTTTGTACAAAAAAGCAGGCTATGGTTGTGTTTCACCAATTGATACTTCC | Tb927.4.2320_rev | GGGGACCACTTTGTACAAGAAAGCTGGGTTTACGCATGTTTCTTCCCTGTTGA |
| 1_hypothetical | G12 | Tb927.2.3610_for | GGGGACAAGTTTGTACAAAAAAGCAGGCTATGTCTTCCACGAAGTGCGC | Tb927.2.3610_rev | GGGGACCACTTTGTACAAGAAAGCTGGGTCTAGTAGTTGGGATTTACCTTCGTTACC |
| 1_hypothetical | G13 | Tb927.8.1945_for | GGGGACAAGTTTGTACAAAAAAGCAGGCTATGTTTTCGTTACGGGAGATACTTTCG | Tb927.8.1945_rev | GGGGACCACTTTGTACAAGAAAGCTGGGTCTAGTAGGTATCTACCCAGGCGTTG |
| 1_hypothetical | G14 | Tb927.9.11370_for | GGGGACAAGTTTGTACAAAAAAGCAGGCTATGTCGCAATTATGGGTAAAGCGC | Tb927.9.11370_rev | GGGGACCACTTTGTACAAGAAAGCTGGGTTCAATATGAATCTCTCCTGTCGATATTGAT |
| 1_hypothetical | G15 | Tb927.3.1690_for | GGGGACAAGTTTGTACAAAAAAGCAGGCTATGGTACTTTTTTCTACATACCGCAGC | Tb927.3.1690_rev | GGGGACCACTTTGTACAAGAAAGCTGGGTTCAATGATGTGCCTGTGCTCC |
| 1_hypothetical | G16 | Tb927.8.3070_for | GGGGACAAGTTTGTACAAAAAAGCAGGCTATGGCCTATGTTTCTCCAGCACG | Tb927.8.3070_rev | GGGGACCACTTTGTACAAGAAAGCTGGGTCTACTTCGGTAACCGCTGCTG |
| 1_hypothetical | G17 | Tb927.9.8380_for | GGGGACAAGTTTGTACAAAAAAGCAGGCTATGTCAATGTTCGGTGCCCG | Tb927.9.8380_rev | GGGGACCACTTTGTACAAGAAAGCTGGGTTCATTTTCCCACGGAATTATTATTACGCG |
| 1_hypothetical | G18 | Tb927.11.6190_for | GGGGACAAGTTTGTACAAAAAAGCAGGCTATGACACCCGTACTTCATGATGC | Tb927.11.6190_rev | GGGGACCACTTTGTACAAGAAAGCTGGGTCTAAGCAATACCCACGGAAACACC |
| 1_hypothetical | G19 | Tb927.3.1960_for | GGGGACAAGTTTGTACAAAAAAGCAGGCTATGGAAGGACCACCTCCCG | Tb927.3.1960_rev | GGGGACCACTTTGTACAAGAAAGCTGGGTTCAACACGTCTCTCCTTTAGCCA |
| 1_hypothetical | G20 | Tb927.11.6470_for | GGGGACAAGTTTGTACAAAAAAGCAGGCTATGAGCCTCAAGGACTGTGGT | Tb927.11.6470_rev | GGGGACCACTTTGTACAAGAAAGCTGGGTTCAATCATCCTCGATGTCTGCCT |
| 1_hypothetical | G21 | Tb927.10.15790_for | GGGGACAAGTTTGTACAAAAAAGCAGGCTATGGCAAATGGCGTAACGGC | Tb927.10.15790_rev | GGGGACCACTTTGTACAAGAAAGCTGGGTTCATGTTTTGTTCGTTTCGCCG |
| 1_hypothetical | G22 | Tb927.10.14120_for | GGGGACAAGTTTGTACAAAAAAGCAGGCTATGCAAGTAAGTGACTTTGCGTGT | Tb927.10.14120_rev | GGGGACCACTTTGTACAAGAAAGCTGGGTCTATGACAGGCCGCGCTTT |
| 1_hypothetical | G23 | Tb927.1.1760_for | GGGGACAAGTTTGTACAAAAAAGCAGGCTATGGTGTGGTTACACTGGACAAC | Tb927.1.1760_rev | GGGGACCACTTTGTACAAGAAAGCTGGGTTTACTTCAGGGAAAGAAAAGTGATGCT |
| 1_hypothetical | G24 | Tb927.10.12070_for | GGGGACAAGTTTGTACAAAAAAGCAGGCTATGGACGGGGGTTTAAAATCCCT | Tb927.10.12070_rev | GGGGACCACTTTGTACAAGAAAGCTGGGTTTAAAAGGTAGGAAAAACAAAAGAAACAAA |
| 1_hypothetical | H1 | Tb927.11.16510_for | GGGGACAAGTTTGTACAAAAAAGCAGGCTATGGCACCTCCAAAGGCTTCA | Tb927.11.16510_rev | GGGGACCACTTTGTACAAGAAAGCTGGGTTTACTCCGTGATGAGCTTTTCACTTC |
| 1_hypothetical | H2 | Tb927.10.1220_for | GGGGACAAGTTTGTACAAAAAAGCAGGCTATGACATCTCATCCCGCTGTTCT | Tb927.10.1220_rev | GGGGACCACTTTGTACAAGAAAGCTGGGTTTAATCCAGGGCTTTCTGGGGG |
| 1_hypothetical | H3 | Tb927.2.4840_for | GGGGACAAGTTTGTACAAAAAAGCAGGCTATGCTACTTCTTCCGCCTTTCCT | Tb927.2.4840_rev | GGGGACCACTTTGTACAAGAAAGCTGGGTTCAATCGACATGAACCCGACCA |
| 1_hypothetical | H4 | Tb927.10.10080_for | GGGGACAAGTTTGTACAAAAAAGCAGGCTATGCCACCTAAGAAGGACGAGG | Tb927.10.10080_rev | GGGGACCACTTTGTACAAGAAAGCTGGGTTCACGAAGATCTGTCTGCCCTC |
| 1_hypothetical | H5 | Tb927.6.2510_for | GGGGACAAGTTTGTACAAAAAAGCAGGCTATGTTAAGACGTTTGTTTACGCGC | Tb927.6.2510_rev | GGGGACCACTTTGTACAAGAAAGCTGGGTTTATGATGCACATTTCAGTAACTTCTCCC |
| 1_hypothetical | H6 | Tb927.10.11370_for | GGGGACAAGTTTGTACAAAAAAGCAGGCTATGTTTCGTCCCACCACCGC | Tb927.10.11370_rev | GGGGACCACTTTGTACAAGAAAGCTGGGTCTAGTACCACGTCCGCGGT |
| 1_hypothetical | H7 | Tb927.10.40_for | GGGGACAAGTTTGTACAAAAAAGCAGGCTATGTATCGACGGGCAAGTTCTCG | Tb927.10.40_rev | GGGGACCACTTTGTACAAGAAAGCTGGGTCTATTTGAAACTCCTCAGACAAGCCA |
| 1_hypothetical | H8 | Tb927.10.11280_for | GGGGACAAGTTTGTACAAAAAAGCAGGCTATGGTTGCCCGTTTCCTCTGG | Tb927.10.11280_rev | GGGGACCACTTTGTACAAGAAAGCTGGGTCTAACAAAGCAGCTGGCGATCA |
| 1_hypothetical | H9 | Tb927.10.1610_for | GGGGACAAGTTTGTACAAAAAAGCAGGCTATGTCATATAGTCTCCGTAAACTGAGGC | Tb927.10.1610_rev | GGGGACCACTTTGTACAAGAAAGCTGGGTTTAGTGAACACCAAGTCCCCACT |
| 1_hypothetical | H10 | Tb927.9.10060_for | GGGGACAAGTTTGTACAAAAAAGCAGGCTATGTCTCGGAAAGCCGGAGA | Tb927.9.10060_rev | GGGGACCACTTTGTACAAGAAAGCTGGGTTCACTTGTATGCTACGAGTGCTGG |
| 1_hypothetical | H11 | Tb927.10.10040_for | GGGGACAAGTTTGTACAAAAAAGCAGGCTATGTATAAATTAGAATTGTTGTGTACTTGG | Tb927.10.10040_rev | GGGGACCACTTTGTACAAGAAAGCTGGGTTTATGCACCTGGGAGCGGA |
| 1_hypothetical | H12 | Tb927.10.12460_for | GGGGACAAGTTTGTACAAAAAAGCAGGCTATGGACACATTATTGATGATTGGGGC | Tb927.10.12460_rev | GGGGACCACTTTGTACAAGAAAGCTGGGTTCACGAATTGAATGTATGTATCCGCG |
| 1_hypothetical | H13 | Tb927.11.12330_for | GGGGACAAGTTTGTACAAAAAAGCAGGCTATGGTGCGTTGTCAGTCGTC | Tb927.11.12330_rev | GGGGACCACTTTGTACAAGAAAGCTGGGTTTATTGCAACTTGTCCAATTTGACGAC |
| 1_hypothetical | H14 | Tb927.4.3060_for | GGGGACAAGTTTGTACAAAAAAGCAGGCTATGGAAATCATTGATCGGCCCC | Tb927.4.3060_rev | GGGGACCACTTTGTACAAGAAAGCTGGGTTCAATCACGTGAGAAGCGGGA |
| 1_hypothetical | H15 | Tb927.9.7980_for | GGGGACAAGTTTGTACAAAAAAGCAGGCTATGGCTCGCACTCGTCGT | Tb927.9.7980_rev | GGGGACCACTTTGTACAAGAAAGCTGGGTTTATTGAGTTGGTTGCTGCTCACT |
| 1_hypothetical | H16 | Tb927.10.6990_for | GGGGACAAGTTTGTACAAAAAAGCAGGCTATGACTGGCAATTCGCATTGTGT | Tb927.10.6990_rev | GGGGACCACTTTGTACAAGAAAGCTGGGTCTATGATTGTTTAACAGCCGTAATCCCG |
| 1_hypothetical | H17 | Tb927.10.10110_for | GGGGACAAGTTTGTACAAAAAAGCAGGCTATGCCCCTAAGATGCACTAAACCA | Tb927.10.10110_rev | GGGGACCACTTTGTACAAGAAAGCTGGGTTCAATTCCACGGTTGAAACAACAAC |
| 1_hypothetical | H18 | Tb927.9.7810_for | GGGGACAAGTTTGTACAAAAAAGCAGGCTATGTCGAATCTCCCACCCTTAACT | Tb927.9.7810_rev | GGGGACCACTTTGTACAAGAAAGCTGGGTCTATGTTTTATTCTCCCCTTCATCCACG |
| 1_hypothetical | H19 | Tb927.10.1160_for | GGGGACAAGTTTGTACAAAAAAGCAGGCTATGGTGCGGTACGCCAAAC | Tb927.10.1160_rev | GGGGACCACTTTGTACAAGAAAGCTGGGTCTAAGCTTCTGCTAATTGTTCCATCTGA |
| 1_hypothetical | H20 | Tb927.3.4480_for | GGGGACAAGTTTGTACAAAAAAGCAGGCTATGGCATTAGGGGGTTCACTCC | Tb927.3.4480_rev | GGGGACCACTTTGTACAAGAAAGCTGGGTTCAAATCGCGATCCACTTCAATGA |
| 1_hypothetical | H21 | Tb927.8.7210_for | GGGGACAAGTTTGTACAAAAAAGCAGGCTATGTCGAAGAAGCAAGGCGC | Tb927.8.7210_rev | GGGGACCACTTTGTACAAGAAAGCTGGGTTTATTTTAGAACACGATCCACTACCCATC |
| 1_hypothetical | H22 | Tb927.10.390_for | GGGGACAAGTTTGTACAAAAAAGCAGGCTATGTCCGTGCGCATTCGC | Tb927.10.390_rev | GGGGACCACTTTGTACAAGAAAGCTGGGTCTACAGCATGGTGCAACAACCA |
| 1_hypothetical | H23 | Tb927.4.5190_for | GGGGACAAGTTTGTACAAAAAAGCAGGCTATGACAAAAAGACATAGCGGACGC | Tb927.4.5190_rev | GGGGACCACTTTGTACAAGAAAGCTGGGTTTACTTTAGCACACGCTCCACC |
| 1_hypothetical | H24 | Tb927.11.14995_for | GGGGACAAGTTTGTACAAAAAAGCAGGCTATGCGCGGCAATCGTCTG | Tb927.11.14995_rev | GGGGACCACTTTGTACAAGAAAGCTGGGTTTAGTACAGCGACTGCGTCCT |
| 1_hypothetical | I1 | Tb927.9.10360_for | GGGGACAAGTTTGTACAAAAAAGCAGGCTATGACGGACCGTTTAACGGC | Tb927.9.10360_rev | GGGGACCACTTTGTACAAGAAAGCTGGGTTCACTCACAATTAATATGCCTCTCAACTT |
| 1_hypothetical | I2 | Tb927.2.5140_for | GGGGACAAGTTTGTACAAAAAAGCAGGCTATGAAGCGTGTAGCTCAGAGTGC | Tb927.2.5140_rev | GGGGACCACTTTGTACAAGAAAGCTGGGTTCACAATTCTTTGAGAGCCTTTTCTTGA |
| 1_hypothetical | I3 | Tb927.6.4180_for | GGGGACAAGTTTGTACAAAAAAGCAGGCTATGAGTGCGCCGCGTGTC | Tb927.6.4180_rev | GGGGACCACTTTGTACAAGAAAGCTGGGTTTAGGCCAAGTACCGAGCACC |
| 1_hypothetical | I4 | Tb927.4.2740_for | GGGGACAAGTTTGTACAAAAAAGCAGGCTATGGAAGCTGTCTTCTACGCATTTG | Tb927.4.2740_rev | GGGGACCACTTTGTACAAGAAAGCTGGGTTTACTTTTGCGAGGAGGTTGTGC |
| 1_hypothetical | I5 | Tb927.9.16070_for | GGGGACAAGTTTGTACAAAAAAGCAGGCTATGTTCGTCCGTCATTTCAAAGAAGT | Tb927.9.16070_rev | GGGGACCACTTTGTACAAGAAAGCTGGGTTCATGTGAGCATATTGGCAATGTCA |
| 1_hypothetical | I6 | Tb927.11.12730_for | GGGGACAAGTTTGTACAAAAAAGCAGGCTATGAGCTGCCTTCATGGTGGC | Tb927.11.12730_rev | GGGGACCACTTTGTACAAGAAAGCTGGGTTTAACCCGGTTTGTGGCAACA |
| 1_hypothetical | I7 | Tb927.11.12700_for | GGGGACAAGTTTGTACAAAAAAGCAGGCTATGCACTTAACCTTAGGTGTGTTAACG | Tb927.11.12700_rev | GGGGACCACTTTGTACAAGAAAGCTGGGTTCATTTTTCTGTGAAGCGCATGTAC |
| 1_hypothetical | I8 | Tb927.4.1370_for | GGGGACAAGTTTGTACAAAAAAGCAGGCTATGAACTGCAGCTCCACATTGG | Tb927.4.1370_rev | GGGGACCACTTTGTACAAGAAAGCTGGGTCTAGAAAAACGGGTAAGGCGGTG |
| 1_hypothetical | I9 | Tb927.2.4700_for | GGGGACAAGTTTGTACAAAAAAGCAGGCTATGACAGGAAAAGATGGAAAAGTACTGG | Tb927.2.4700_rev | GGGGACCACTTTGTACAAGAAAGCTGGGTTCAAGCGGTAGCGCATTCC |
| 1_hypothetical | I10 | Tb927.5.690_for | GGGGACAAGTTTGTACAAAAAAGCAGGCTATGGCGTTTGGAAATGCGTCT | Tb927.5.690_rev | GGGGACCACTTTGTACAAGAAAGCTGGGTTCATATATCACCACCCGCTGCA |
| 1_hypothetical | I11 | Tb927.7.5780_for | GGGGACAAGTTTGTACAAAAAAGCAGGCTATGATATCACTACATTTGTCCTCGCAC | Tb927.7.5780_rev | GGGGACCACTTTGTACAAGAAAGCTGGGTTTATGAGTTTTGAAAGACAAAACTGTCCAG |
| 1_hypothetical | I12 | Tb927.6.5060_for | GGGGACAAGTTTGTACAAAAAAGCAGGCTATGGCTGCCTCACAGCTTATCG | Tb927.6.5060_rev | GGGGACCACTTTGTACAAGAAAGCTGGGTTCATGCTCGCTTCGACACATTAA |
| 1_hypothetical | I13 | Tb04.3I12.100_for | GGGGACAAGTTTGTACAAAAAAGCAGGCTATGCTTGCGAATCCATCTTCCTG | Tb04.3I12.100_rev | GGGGACCACTTTGTACAAGAAAGCTGGGTTTAAAAGGACTCAAAACGTAAAAGAAAAAA |
| 1_hypothetical | I14 | Tb927.10.1870_for | GGGGACAAGTTTGTACAAAAAAGCAGGCTATGCTGCGCTACAGCCGA | Tb927.10.1870_rev | GGGGACCACTTTGTACAAGAAAGCTGGGTCTAAACAACACGGAATTCAAAGTCTTGT |
| 1_hypothetical | I15 | Tb927.7.1380_for | GGGGACAAGTTTGTACAAAAAAGCAGGCTATGAGTACGATGGATGTCCCGG | Tb927.7.1380_rev | GGGGACCACTTTGTACAAGAAAGCTGGGTTCACCTAAAGCATAGTTGAGTTTGTGT |
| 1_hypothetical | I16 | Tb927.7.3590_for | GGGGACAAGTTTGTACAAAAAAGCAGGCTATGCTTCGCCGCGCAGCT | Tb927.7.3590_rev | GGGGACCACTTTGTACAAGAAAGCTGGGTCTAATTCACTATCAAATGCAGGACAGCA |
| 1_hypothetical | I17 | Tb927.9.9810_for | GGGGACAAGTTTGTACAAAAAAGCAGGCTATGTCGCTTCAAACAACCCTAATACA | Tb927.9.9810_rev | GGGGACCACTTTGTACAAGAAAGCTGGGTCTAAAATACGTTAACTGTCATAATGGCATA |
| 1_hypothetical | I18 | Tb927.7.7090_for | GGGGACAAGTTTGTACAAAAAAGCAGGCTATGCTCGCACGTACGCTTC | Tb927.7.7090_rev | GGGGACCACTTTGTACAAGAAAGCTGGGTTTACTCCATAGCGAACACCGCT |
| 1_hypothetical | I19 | Tb927.9.13460_for | GGGGACAAGTTTGTACAAAAAAGCAGGCTATGAGTGGTGACAGCGAGCC | Tb927.9.13460_rev | GGGGACCACTTTGTACAAGAAAGCTGGGTTCATGCACCACTCATCGCCA |
| 1_hypothetical | I20 | Tb927.10.14130_for | GGGGACAAGTTTGTACAAAAAAGCAGGCTATGAAACCGTCTCAACCGTCG | Tb927.10.14130_rev | GGGGACCACTTTGTACAAGAAAGCTGGGTCTACTTTTTCCGTGTGTTGGCAGT |
| 1_hypothetical | I21 | Tb927.11.730_for | GGGGACAAGTTTGTACAAAAAAGCAGGCTATGTCGGGGTTTTCAGGGACA | Tb927.11.730_rev | GGGGACCACTTTGTACAAGAAAGCTGGGTTTAAGTCCCCGAGCCACGT |
| 1_hypothetical | I22 | Tb927.9.17670_for | GGGGACAAGTTTGTACAAAAAAGCAGGCTATGGGGATTAATTTATGTGTGTCACTCTAC | Tb927.9.17670_rev | GGGGACCACTTTGTACAAGAAAGCTGGGTTCACAAATATGTTTTTCTACTCATCGTGAG |
| 1_hypothetical | I23 | Tb927.8.1370_for | GGGGACAAGTTTGTACAAAAAAGCAGGCTATGTCGTCCAAGGTTGGTCAGG | Tb927.8.1370_rev | GGGGACCACTTTGTACAAGAAAGCTGGGTTTAGCTCCGTAGCTTGTTGAAGTCG |
| 1_hypothetical | I24 | Tb927.5.1840_for | GGGGACAAGTTTGTACAAAAAAGCAGGCTATGCTTTTTTCCCTTGACGGGC | Tb927.5.1840_rev | GGGGACCACTTTGTACAAGAAAGCTGGGTTCATACCTCCCGCCTCATTGTATC |
| 1_hypothetical | J1 | Tb927.10.4410_for | GGGGACAAGTTTGTACAAAAAAGCAGGCTATGATGGAGGTGGGCGGTT | Tb927.10.4410_rev | GGGGACCACTTTGTACAAGAAAGCTGGGTTCACGCAAAAGCCACACGC |
| 1_hypothetical | J2 | Tb927.4.1670_for | GGGGACAAGTTTGTACAAAAAAGCAGGCTATGGATAAGGCAGTTGGTAAACTAGTTAAG | Tb927.4.1670_rev | GGGGACCACTTTGTACAAGAAAGCTGGGTCTATTCCCATATAAACCACCCAACAACG |
| 1_hypothetical | J3 | Tb927.4.2100_for | GGGGACAAGTTTGTACAAAAAAGCAGGCTATGACCGTAATTCATTTATTTCTTTATGTG | Tb927.4.2100_rev | GGGGACCACTTTGTACAAGAAAGCTGGGTTTAATACCGTCCTTTTTCTTTCTGGCT |
| 1_hypothetical | J4 | Tb927.10.11350_for | GGGGACAAGTTTGTACAAAAAAGCAGGCTATGTTTCGAGTGACTGGCTTGC | Tb927.10.11350_rev | GGGGACCACTTTGTACAAGAAAGCTGGGTCTAACTTCGCATATCCTTTGCGTTCT |
| 1_hypothetical | J5 | Tb927.11.2160_for | GGGGACAAGTTTGTACAAAAAAGCAGGCTATGCAGTCGCGGCGTAAC | Tb927.11.2160_rev | GGGGACCACTTTGTACAAGAAAGCTGGGTTTAGTGATGTGCATGTACCTCACG |
| 1_hypothetical | J6 | Tb927.11.16320_for | GGGGACAAGTTTGTACAAAAAAGCAGGCTATGAATTCACAGCGGTCGGAGT | Tb927.11.16320_rev | GGGGACCACTTTGTACAAGAAAGCTGGGTTCAGGTGTTTTTATAAATCTTCCCCTTGAA |
| 1_hypothetical | J7 | Tb927.7.7010_for | GGGGACAAGTTTGTACAAAAAAGCAGGCTATGCTTCGGCACTGTACCGC | Tb927.7.7010_rev | GGGGACCACTTTGTACAAGAAAGCTGGGTCTACAACAACTCCCGTGATGGC |
| 1_hypothetical | J8 | Tb927.4.4815_for | GGGGACAAGTTTGTACAAAAAAGCAGGCTATGTCGGAAACGAACTTCACGT | Tb927.4.4815_rev | GGGGACCACTTTGTACAAGAAAGCTGGGTTTACAGTGTCTTCTCCACCTCCC |
| 1_hypothetical | J9 | Tb927.9.8570_for | GGGGACAAGTTTGTACAAAAAAGCAGGCTATGAAAGGAGGTGAAACAGCAGC | Tb927.9.8570_rev | GGGGACCACTTTGTACAAGAAAGCTGGGTTCACTGATCTTTATATACGGGAAGGTTTAC |
| 1_hypothetical | J10 | Tb927.3.940_for | GGGGACAAGTTTGTACAAAAAAGCAGGCTATGAGGATCAGTTGCTCATGCCT | Tb927.3.940_rev | GGGGACCACTTTGTACAAGAAAGCTGGGTCTACCATCGCCGCATAATGTAGG |
| 1_hypothetical | J11 | Tb927.5.296b_for | GGGGACAAGTTTGTACAAAAAAGCAGGCTATGGAATATCTCCAGGGAAGGAAGC | Tb927.5.296b_rev | GGGGACCACTTTGTACAAGAAAGCTGGGTTCATTGCTTCTGGGGCCGA |
| 1_hypothetical | J12 | Tb927.7.6020_for | GGGGACAAGTTTGTACAAAAAAGCAGGCTATGAACAATGATAAGTCAGAATTTTATATC | Tb927.7.6020_rev | GGGGACCACTTTGTACAAGAAAGCTGGGTTCATATTCCTTCTGCGGACAACTTTT |
| 1_hypothetical | J13 | Tb927.5.1170_for | GGGGACAAGTTTGTACAAAAAAGCAGGCTATGGAAAACAAAATGTCGGCTGAGC | Tb927.5.1170_rev | GGGGACCACTTTGTACAAGAAAGCTGGGTCTATGCACTCCCTGACAAAAGGG |
| 1_hypothetical | J14 | Tb927.4.2610_for | GGGGACAAGTTTGTACAAAAAAGCAGGCTATGTGTTTTACACATGTCTCTCTTGTGT | Tb927.4.2610_rev | GGGGACCACTTTGTACAAGAAAGCTGGGTTCAATACGGTTGAAAGTTGCTAGCC |

| | | | | | |
|---|---|---|---|---|---|
| 1_hypothetical | J15 | Tb927.10.3270_for | GGGGACAAGTTTGTACAAAAAAGCAGGCTATGTTGAGGCGTGTCCGTCC | Tb927.10.3270_rev | GGGGACCACTTTGTACAAGAAAGCTGGGTTCAGTTTCGGCGTCGCTCA |
| 1_hypothetical | J16 | Tb927.9.12480_for | GGGGACAAGTTTGTACAAAAAAGCAGGCTATGTCGAGTGGAAGCCATCGT | Tb927.9.12480_rev | GGGGACCACTTTGTACAAGAAAGCTGGGTCTAACCGCCATGGCGCCA |
| 1_hypothetical | J17 | Tb927.6.4620_for | GGGGACAAGTTTGTACAAAAAAGCAGGCTATGGCTGCTCAAGGGCTGT | Tb927.6.4620_rev | GGGGACCACTTTGTACAAGAAAGCTGGGTTCAGGTCTGACTCGTATAGTCGGG |
| 1_hypothetical | J18 | Tb927.2.3190_for | GGGGACAAGTTTGTACAAAAAAGCAGGCTATGGAAATTCCCATAAATGTGTCGGA | Tb927.2.3190_rev | GGGGACCACTTTGTACAAGAAAGCTGGGTTCAAAGACCTGTGACGCGTCG |
| 1_hypothetical | J19 | Tb927.10.11920_for | GGGGACAAGTTTGTACAAAAAAGCAGGCTATGCAGGCAGTGGGTGAAAC | Tb927.10.11920_rev | GGGGACCACTTTGTACAAGAAAGCTGGGTTTACCCCCCCCCACTCC |
| 1_hypothetical | J20 | Tb927.10.7960_for | GGGGACAAGTTTGTACAAAAAAGCAGGCTATGGCTCCCAAGAAGAAGGTCC | Tb927.10.7960_rev | GGGGACCACTTTGTACAAGAAAGCTGGGTTTACTTTGTTAGGAGCACCTCTCTCA |
| 1_hypothetical | J21 | Tb927.6.260_for | GGGGACAAGTTTGTACAAAAAAGCAGGCTATGTGGGGCAAGAGTTTGGATTG | Tb927.6.260_rev | GGGGACCACTTTGTACAAGAAAGCTGGGTCTAACCGGATGAGTGAGTGACGA |
| 1_hypothetical | J22 | Tb927.10.3960_for | GGGGACAAGTTTGTACAAAAAAGCAGGCTATGCTGCCGCTGTCGCGT | Tb927.10.3960_rev | GGGGACCACTTTGTACAAGAAAGCTGGGTCTAAGTACATTTTTTCGGATCTCCATTGC |
| 1_hypothetical | J23 | Tb927.11.2240_for | GGGGACAAGTTTGTACAAAAAAGCAGGCTATGGTGAGCGAAGTGATGGTTG | Tb927.11.2240_rev | GGGGACCACTTTGTACAAGAAAGCTGGGTTTACACCATAGGGCTTAGAACTCGT |
| 1_hypothetical | J24 | Tb927.10.8090_for | GGGGACAAGTTTGTACAAAAAAGCAGGCTATGCAGCGCGGATTTTGTAGG | Tb927.10.8090_rev | GGGGACCACTTTGTACAAGAAAGCTGGGTCTATTTTGTACACGAAAGCACATGTGT |
| 1_hypothetical | K1 | Tb927.10.4550_for | GGGGACAAGTTTGTACAAAAAAGCAGGCTATGGGTAGCACACCTCATTATTCTAGG | Tb927.10.4550_rev | GGGGACCACTTTGTACAAGAAAGCTGGGTTCAAGGCTTCTTAACGGTATTCATCAC |
| 1_hypothetical | K2 | Tb927.7.370_for | GGGGACAAGTTTGTACAAAAAAGCAGGCTATGCATGATTTGATCGCTGTTGTAGA | Tb927.7.370_rev | GGGGACCACTTTGTACAAGAAAGCTGGGTCTACTGCAAGGGCCGCAG |
| 1_hypothetical | K3 | Tb927.1.3350_for | GGGGACAAGTTTGTACAAAAAAGCAGGCTATGACAGATGGCCACGCAAC | Tb927.1.3350_rev | GGGGACCACTTTGTACAAGAAAGCTGGGTTCAGATATTCGCCTTATGGAAGCATAC |
| 1_hypothetical | K4 | Tb927.10.12900_for | GGGGACAAGTTTGTACAAAAAAGCAGGCTATGCTGCACTGTTCACAGTGG | Tb927.10.12900_rev | GGGGACCACTTTGTACAAGAAAGCTGGGTCTAGTGTACCAATGAGAACCTCATGTC |
| 1_hypothetical | K5 | Tb927.3.2180_for | GGGGACAAGTTTGTACAAAAAAGCAGGCTATGTTGCGGAAAACACCATTGTTC | Tb927.3.2180_rev | GGGGACCACTTTGTACAAGAAAGCTGGGTCTAGAAGAGCTCTGGCCGGC |
| 1_hypothetical | K6 | Tb927.8.1790_for | GGGGACAAGTTTGTACAAAAAAGCAGGCTATGTCTCTTTGCGCTGAGATTAACC | Tb927.8.1790_rev | GGGGACCACTTTGTACAAGAAAGCTGGGTTCAGCGGAGCAGGCGACA |
| 1_hypothetical | K7 | Tb927.9.1470_for | GGGGACAAGTTTGTACAAAAAAGCAGGCTATGGCACTGCGTGATATTGTAGG | Tb927.9.1470_rev | GGGGACCACTTTGTACAAGAAAGCTGGGTTCAAAGTCTACCTGCACGCTTTG |
| 1_hypothetical | K8 | Tb927.9.16950_for | GGGGACAAGTTTGTACAAAAAAGCAGGCTATGTGTTACTCTGATTCGTATCAAGTGTT | Tb927.9.16950_rev | GGGGACCACTTTGTACAAGAAAGCTGGGTTTATAATAAAAGAAGTAGTAATATTAATAG |
| 1_hypothetical | K9 | Tb927.3.3620_for | GGGGACAAGTTTGTACAAAAAAGCAGGCTATGTCGACATTCCAACGCCG | Tb927.3.3620_rev | GGGGACCACTTTGTACAAGAAAGCTGGGTTCAGGAGTCTGGCTCTGCG |
| 1_hypothetical | K10 | Tb927.11.13300_for | GGGGACAAGTTTGTACAAAAAAGCAGGCTATGGCAAAGGTGAGCATAGACCC | Tb927.11.13300_rev | GGGGACCACTTTGTACAAGAAAGCTGGGTTTACATCACCACTTCAGGATTGGACT |
| 1_hypothetical | K11 | Tb927.10.15860_for | GGGGACAAGTTTGTACAAAAAAGCAGGCTATGCCAAACATAAAGGGAGGGGT | Tb927.10.15860_rev | GGGGACCACTTTGTACAAGAAAGCTGGGTCTACGTATCGTTTTCCATTCCACGG |
| 1_hypothetical | K12 | Tb927.10.5160_for | GGGGACAAGTTTGTACAAAAAAGCAGGCTATGTGCGCGTGTATGTCAAGG | Tb927.10.5160_rev | GGGGACCACTTTGTACAAGAAAGCTGGGTTCACAAGCAAATATATCCCCCAATGG |
| 1_hypothetical | K13 | Tb927.11.490_for | GGGGACAAGTTTGTACAAAAAAGCAGGCTATGGCTCCACCTTCAGATATTTCGG | Tb927.11.490_rev | GGGGACCACTTTGTACAAGAAAGCTGGGTTTACGTTTCAGCACCACATCGG |
| 1_hypothetical | K14 | Tb927.10.1020_for | GGGGACAAGTTTGTACAAAAAAGCAGGCTATGCATAGGATGCCGTTCAACTC | Tb927.10.1020_rev | GGGGACCACTTTGTACAAGAAAGCTGGGTTCAGGGAGATTGAAATTGGTGTATGC |
| 1_hypothetical | K15 | Tb927.4.1760_for | GGGGACAAGTTTGTACAAAAAAGCAGGCTATGTTGCGGTTGACACGGC | Tb927.4.1760_rev | GGGGACCACTTTGTACAAGAAAGCTGGGTTCACTCACGAGAGAAAATGTTACGTC |
| 1_hypothetical | K16 | Tb927.10.130_for | GGGGACAAGTTTGTACAAAAAAGCAGGCTATGGTAATTTTCCAACGGGTTGAAGC | Tb927.10.130_rev | GGGGACCACTTTGTACAAGAAAGCTGGGTTTATGTTAGAAACCGAATGTACTGACCG |
| 1_hypothetical | K17 | Tb927.2.2070_for | GGGGACAAGTTTGTACAAAAAAGCAGGCTATGCGGTTGTACAAGTCCTCTTTT | Tb927.2.2070_rev | GGGGACCACTTTGTACAAGAAAGCTGGGTTTATTCATTGAATTGAGTTCGCATAAGGTC |
| 1_hypothetical | K18 | Tb927.7.1940_for | GGGGACAAGTTTGTACAAAAAAGCAGGCTATGGACAATATACACCACCGCCC | Tb927.7.1940_rev | GGGGACCACTTTGTACAAGAAAGCTGGGTTCAACAAGTTGCAGTTGGACCTC |
| 1_hypothetical | K19 | Tb927.11.15930_for | GGGGACAAGTTTGTACAAAAAAGCAGGCTATGACAGTTCACTTGCTCTGGATCA | Tb927.11.15930_rev | GGGGACCACTTTGTACAAGAAAGCTGGGTTCACCTCTTCACCGCGGG |
| 1_hypothetical | K20 | Tb927.10.13420_for | GGGGACAAGTTTGTACAAAAAAGCAGGCTATGAGCGTCGTCACTTCTGAAC | Tb927.10.13420_rev | GGGGACCACTTTGTACAAGAAAGCTGGGTTCATTCCCTTCCGGACACGG |
| 1_hypothetical | K21 | Tb927.4.4190_for | GGGGACAAGTTTGTACAAAAAAGCAGGCTATGGGCAAGCTCCAGGTGT | Tb927.4.4190_rev | GGGGACCACTTTGTACAAGAAAGCTGGGTTTAAGCCATGAGAGCAGGAGCT |
| 1_hypothetical | K22 | Tb927.10.9830_for | GGGGACAAGTTTGTACAAAAAAGCAGGCTATGCGCCGCACCTTCATT | Tb927.10.9830_rev | GGGGACCACTTTGTACAAGAAAGCTGGGTCTACTTAGCCTTTGCTTTCACATCTTCG |
| 1_hypothetical | K23 | Tb927.9.10200_for | GGGGACAAGTTTGTACAAAAAAGCAGGCTATGCAGGGAGAGGGAAATGAATTG | Tb927.9.10200_rev | GGGGACCACTTTGTACAAGAAAGCTGGGTTTACGCATCTTCTTCCTCGTTATCATC |
| 1_hypothetical | K24 | Tb927.8.5800_for | GGGGACAAGTTTGTACAAAAAAGCAGGCTATGAAACACAAAGATGCTCGCGG | Tb927.8.5800_rev | GGGGACCACTTTGTACAAGAAAGCTGGGTTCACTTCTTCACTCTTTTTGGCGG |
| 1_hypothetical | L1 | Tb927.4.2530_for | GGGGACAAGTTTGTACAAAAAAGCAGGCTATGGGCTTCGGTGCAGTCG | Tb927.4.2530_rev | GGGGACCACTTTGTACAAGAAAGCTGGGTTTAATTCTTCTTTTTTTCTTTCCCCGTCG |
| 1_hypothetical | L2 | Tb927.7.3690_for | GGGGACAAGTTTGTACAAAAAAGCAGGCTATGCAGAAACGAAGGGAATGTTTGG | Tb927.7.3690_rev | GGGGACCACTTTGTACAAGAAAGCTGGGTTTAAAGGAAAGGAAAAAAAAAAAAAAGAGAT |
| 1_hypothetical | L3 | Tb927.1.4730_for | GGGGACAAGTTTGTACAAAAAAGCAGGCTATGGAAGGACAAAACTCAGTTTCTCTTC | Tb927.1.4730_rev | GGGGACCACTTTGTACAAGAAAGCTGGGTCTACTCATAACAGTGCAGATCATTTGACA |
| 1_hypothetical | L4 | Tb927.9.8370_for | GGGGACAAGTTTGTACAAAAAAGCAGGCTATGCAACGCAATTGGGTATTGC | Tb927.9.8370_rev | GGGGACCACTTTGTACAAGAAAGCTGGGTTCAACAATCCAGGTAATCCGACGC |
| 1_hypothetical | L5 | Tb927.10.1480_for | GGGGACAAGTTTGTACAAAAAAGCAGGCTATGTTTGTTTTGCAGCTCATGAAGAG | Tb927.10.1480_rev | GGGGACCACTTTGTACAAGAAAGCTGGGTTTACGTGCAATCAACCTGCAGC |
| 1_hypothetical | L6 | Tb927.10.13600_for | GGGGACAAGTTTGTACAAAAAAGCAGGCTATGTTTAGTCGGTGCCAGCG | Tb927.10.13600_rev | GGGGACCACTTTGTACAAGAAAGCTGGGTCTACAATGTCTTCTGAGCGGCG |
| 1_hypothetical | L7 | Tb927.4.3830_for | GGGGACAAGTTTGTACAAAAAAGCAGGCTATGGAACCGCGTACAGTCCG | Tb927.4.3830_rev | GGGGACCACTTTGTACAAGAAAGCTGGGTTTACGCCGGTTGCGAATAGAG |
| 1_hypothetical | L8 | Tb927.5.1630_for | GGGGACAAGTTTGTACAAAAAAGCAGGCTATGTCCCTCATTACTTCACCGGG | Tb927.5.1630_rev | GGGGACCACTTTGTACAAGAAAGCTGGGTTTAGGCTAAATTTTCATCCTTCTTAAAGAA |
| 1_hypothetical | L9 | Tb927.8.5550_for | GGGGACAAGTTTGTACAAAAAAGCAGGCTATGGTCTTCTTTGTGAGAGTCACAGG | Tb927.8.5550_rev | GGGGACCACTTTGTACAAGAAAGCTGGGTCTACATTGCATTGTTGTAGTGAACAACA |
| 1_hypothetical | L10 | Tb927.11.15540_for | GGGGACAAGTTTGTACAAAAAAGCAGGCTATGTCAAAGCAGAGTAGGAAAGAGGG | Tb927.11.15540_rev | GGGGACCACTTTGTACAAGAAAGCTGGGTCTACTCATGGGAAACATCGGTGTC |
| 1_hypothetical | L11 | Tb927.4.2210_for | GGGGACAAGTTTGTACAAAAAAGCAGGCTATGAAGGCGACAATTATAACACCAGG | Tb927.4.2210_rev | GGGGACCACTTTGTACAAGAAAGCTGGGTTCAGCGCCGCCGTGCACC |
| 1_hypothetical | L12 | Tb927.10.400_for | GGGGACAAGTTTGTACAAAAAAGCAGGCTATGTTGGAGTCTTTTGAATTTGATCAGGA | Tb927.10.400_rev | GGGGACCACTTTGTACAAGAAAGCTGGGTCTAAATATCTTCACACACTGGGCAAAATC |
| 1_hypothetical | L13 | Tb927.5.3740_for | GGGGACAAGTTTGTACAAAAAAGCAGGCTATGGAGGATTCCTCAGTGAGTTCTT | Tb927.5.3740_rev | GGGGACCACTTTGTACAAGAAAGCTGGGTTTAGTAATCATTCTTTGCGGTTTTGGAC |
| 1_hypothetical | L14 | Tb927.10.3050_for | GGGGACAAGTTTGTACAAAAAAGCAGGCTATGTTGAAGGGAGATTTATTATTAGTGTGT | Tb927.10.3050_rev | GGGGACCACTTTGTACAAGAAAGCTGGGTTTATACGAAGTTGTGCTTTCGTTTTTTGG |
| 1_hypothetical | L15 | Tb927.4.1730_for | GGGGACAAGTTTGTACAAAAAAGCAGGCTATGGACACCGAGTGCCTCC | Tb927.4.1730_rev | GGGGACCACTTTGTACAAGAAAGCTGGGTCTACTTGGTGACCTTCTTGAGCATT |
| 1_hypothetical | L16 | Tb927.7.6100_for | GGGGACAAGTTTGTACAAAAAAGCAGGCTATGAGATCCGTGCATCAATCCAAG | Tb927.7.6100_rev | GGGGACCACTTTGTACAAGAAAGCTGGGTTTATTCCTCCTCCACCTCAAGCTC |
| 1_hypothetical | L17 | Tb927.7.4350_for | GGGGACAAGTTTGTACAAAAAAGCAGGCTATGAACCCGAGTCTCTCTGACAG | Tb927.7.4350_rev | GGGGACCACTTTGTACAAGAAAGCTGGGTCTAAGTTACTGGAACTGTAGCGGC |
| 1_hypothetical | L18 | Tb927.6.3920_for | GGGGACAAGTTTGTACAAAAAAGCAGGCTATGTCGGCCATTCCGCCG | Tb927.6.3920_rev | GGGGACCACTTTGTACAAGAAAGCTGGGTTTAGACGCCTACAAACCCCACG |
| 1_hypothetical | L19 | Tb927.11.7880_for | GGGGACAAGTTTGTACAAAAAAGCAGGCTATGGAACTAAACCAACTCGAATTGTTTG | Tb927.11.7880_rev | GGGGACCACTTTGTACAAGAAAGCTGGGTTCAATCACTGATGCTAAAGGTGGC |
| 1_hypothetical | L20 | Tb927.7.4780_for | GGGGACAAGTTTGTACAAAAAAGCAGGCTATGAGTAAGGGAGTTGTGCGGT | Tb927.7.4780_rev | GGGGACCACTTTGTACAAGAAAGCTGGGTCTAAGCCCTTGCACGTGCC |
| 1_hypothetical | L21 | Tb927.10.2800_for | GGGGACAAGTTTGTACAAAAAAGCAGGCTATGGTTTTTTATTCTTTTGTACTGGTGATG | Tb927.10.2800_rev | GGGGACCACTTTGTACAAGAAAGCTGGGTTCAGTTTCTCAGAAAACTGGACCAAC |
| 1_hypothetical | L22 | Tb927.11.10100_for | GGGGACAAGTTTGTACAAAAAAGCAGGCTATGCTTCGACGCACTTCACTG | Tb927.11.10100_rev | GGGGACCACTTTGTACAAGAAAGCTGGGTTTAATCACTCCGCGGTGCAAA |
| 1_hypothetical | L23 | Tb927.10.1000_for | GGGGACAAGTTTGTACAAAAAAGCAGGCTATGAAAGCCCGTTCCTCCCG | Tb927.10.1000_rev | GGGGACCACTTTGTACAAGAAAGCTGGGTTCAAAGGAGTGTTGCCGCG |
| 1_hypothetical | L24 | Tb927.5.1480_for | GGGGACAAGTTTGTACAAAAAAGCAGGCTATGCAAATGGATGAAGCGAATGAAG | Tb927.5.1480_rev | GGGGACCACTTTGTACAAGAAAGCTGGGTTTACACAGCAGAGGGAAACAGACA |
| 1_hypothetical | M1 | Tb927.7.6710_for | GGGGACAAGTTTGTACAAAAAAGCAGGCTATGCGTGTTTGTTTACGGCTTTTC | Tb927.7.6710_rev | GGGGACCACTTTGTACAAGAAAGCTGGGTCTAGGAAAAGAGGGAGAAAAGGAAGGA |
| 1_hypothetical | M2 | Tb927.8.720_for | GGGGACAAGTTTGTACAAAAAAGCAGGCTATGGAGGAACTCGTCAATCCCA | Tb927.8.720_rev | GGGGACCACTTTGTACAAGAAAGCTGGGTTCATTCAAAGGCATTTACGCATGATTC |
| 1_hypothetical | M3 | Tb927.10.6490_for | GGGGACAAGTTTGTACAAAAAAGCAGGCTATGTCTGTCGCACCAAAACGT | Tb927.10.6490_rev | GGGGACCACTTTGTACAAGAAAGCTGGGTCTACTCACCCACGCCCTTCC |
| 1_hypothetical | M4 | Tb927.2.2390_for | GGGGACAAGTTTGTACAAAAAAGCAGGCTATGGTTAATCTGCCGCTGAAGG | Tb927.2.2390_rev | GGGGACCACTTTGTACAAGAAAGCTGGGTCTATCGACGGCCACCCTTACG |
| 1_hypothetical | M5 | Tb927.11.7050_for | GGGGACAAGTTTGTACAAAAAAGCAGGCTATGGCGCTGTGGTTGAGAGT | Tb927.11.7050_rev | GGGGACCACTTTGTACAAGAAAGCTGGGTTCAAACGATATACCGAAGCACATCC |
| 1_hypothetical | M6 | Tb927.2.3920_for | GGGGACAAGTTTGTACAAAAAAGCAGGCTATGTTTTCTTGTTGCCTGCGTAGC | Tb927.2.3920_rev | GGGGACCACTTTGTACAAGAAAGCTGGGTCTACTTGCGCCTAGAATAAACAGCC |
| 1_hypothetical | M7 | Tb927.10.8140_for | GGGGACAAGTTTGTACAAAAAAGCAGGCTATGAATCGTAAAAGGCCGCGG | Tb927.10.8140_rev | GGGGACCACTTTGTACAAGAAAGCTGGGTTTATTTTTTTCGCCCAAGCGATACAAC |
| 1_hypothetical | M8 | Tb927.10.9940_for | GGGGACAAGTTTGTACAAAAAAGCAGGCTATGCGGGTACTCACGCACC | Tb927.10.9940_rev | GGGGACCACTTTGTACAAGAAAGCTGGGTCTAGTAGAGATCCCGGACATGGTATG |
| 1_hypothetical | M9 | Tb927.11.15700_for | GGGGACAAGTTTGTACAAAAAAGCAGGCTATGTCGACAAGCATCACAAATTTTGT | Tb927.11.15700_rev | GGGGACCACTTTGTACAAGAAAGCTGGGTTTACTGAGGTCGTAAGTCTTCCTCG |
| 1_hypothetical | M10 | Tb927.9.3750_for | GGGGACAAGTTTGTACAAAAAAGCAGGCTATGCCTACTCCTCGGGAAACTG | Tb927.9.3750_rev | GGGGACCACTTTGTACAAGAAAGCTGGGTTCACGAAATTTCAAAGAAATTTTGCGGT |
| 1_hypothetical | M11 | Tb927.3.3600_for | GGGGACAAGTTTGTACAAAAAAGCAGGCTATGCCCATGCTGGTCCCT | Tb927.3.3600_rev | GGGGACCACTTTGTACAAGAAAGCTGGGTTTATTTCCGGATGAGAAAACTTTCTATCCC |
| 1_hypothetical | M12 | Tb927.10.11650_for | GGGGACAAGTTTGTACAAAAAAGCAGGCTATGGGTTGTGGAGACTCGAAAGT | Tb927.10.11650_rev | GGGGACCACTTTGTACAAGAAAGCTGGGTCTACTCAGCGGGGTCATCGT |
| 1_hypothetical | M13 | Tb927.6.4790_for | GGGGACAAGTTTGTACAAAAAAGCAGGCTATGGGTGTAGCACTGCGGG | Tb927.6.4790_rev | GGGGACCACTTTGTACAAGAAAGCTGGGTTCAAAGGGTTGCACGGTACG |
| 1_hypothetical | M14 | Tb927.3.3570_for | GGGGACAAGTTTGTACAAAAAAGCAGGCTATGACTACTAGTAACGGCTTCGAATCG | Tb927.3.3570_rev | GGGGACCACTTTGTACAAGAAAGCTGGGTTCACTGCGATGACCCCTGC |
| 1_hypothetical | M15 | Tb927.1.3390_for | GGGGACAAGTTTGTACAAAAAAGCAGGCTATGCCACTTTACCGCCTTTCG | Tb927.1.3390_rev | GGGGACCACTTTGTACAAGAAAGCTGGGTCTATTTGGATTTCTTCAGACGACTGAATTC |
| 1_hypothetical | M16 | Tb927.9.6910_for | GGGGACAAGTTTGTACAAAAAAGCAGGCTATGGCTCAGTGGGTTCCTAAAACG | Tb927.9.6910_rev | GGGGACCACTTTGTACAAGAAAGCTGGGTTTAGTCCTCACCAGAAACCGCT |
| 1_hypothetical | M17 | Tb04.24M18.150_for | GGGGACAAGTTTGTACAAAAAAGCAGGCTATGTGGCATGCTGATAATGCCA | Tb04.24M18.150_rev | GGGGACCACTTTGTACAAGAAAGCTGGGTTTACTTAGGCAATTGAGAGAACGCG |
| 1_hypothetical | M18 | Tb927.11.14670_for | GGGGACAAGTTTGTACAAAAAAGCAGGCTATGAAATGGCTCCTTAGGCTCGT | Tb927.11.14670_rev | GGGGACCACTTTGTACAAGAAAGCTGGGTCTATTTTTTCCTTTCCTGGGCATCGG |
| 1_hypothetical | M19 | Tb927.10.4400_for | GGGGACAAGTTTGTACAAAAAAGCAGGCTATGAGTTTCTTCGGCAAGTTTATGAAAAA | Tb927.10.4400_rev | GGGGACCACTTTGTACAAGAAAGCTGGGTTCAAAGTGGAATGTTGTGTAGATTTGGG |
| 1_hypothetical | M20 | Tb927.2.3330_for | GGGGACAAGTTTGTACAAAAAAGCAGGCTATGTGGAGTCGCCTTTTGGGT | Tb927.2.3330_rev | GGGGACCACTTTGTACAAGAAAGCTGGGTTCAAGGAGTAGTAACGATTTGGCTTC |
| 1_hypothetical | M21 | Tb927.6.3950_for | GGGGACAAGTTTGTACAAAAAAGCAGGCTATGGAGGATCCACGGGACAATC | Tb927.6.3950_rev | GGGGACCACTTTGTACAAGAAAGCTGGGTCTAACGGCGATATTCCTCGTCAC |
| 1_hypothetical | M22 | Tb927.7.3240_for | GGGGACAAGTTTGTACAAAAAAGCAGGCTATGCTGCGCTCAACTCTGAC | Tb927.7.3240_rev | GGGGACCACTTTGTACAAGAAAGCTGGGTCTATTTCCTCCGCTCGTAAAGTTCC |
| 1_hypothetical | M23 | Tb927.11.4020_for | GGGGACAAGTTTGTACAAAAAAGCAGGCTATGCGAGCGAGGCCCTCT | Tb927.11.4020_rev | GGGGACCACTTTGTACAAGAAAGCTGGGTTTACAAGTGAAACCCCGGTTCG |
| 1_hypothetical | M24 | Tb927.11.9410_for | GGGGACAAGTTTGTACAAAAAAGCAGGCTATGCATTTACAAATAAATAAATACATATAT | Tb927.11.9410_rev | GGGGACCACTTTGTACAAGAAAGCTGGGTTCATTCTTCTTTCTGTTTGTATGTTTTGTT |
| 1_hypothetical | N1 | Tb927.6.3360_for | GGGGACAAGTTTGTACAAAAAAGCAGGCTATGATATCATTGCGGGGAAACATAGG | Tb927.6.3360_rev | GGGGACCACTTTGTACAAGAAAGCTGGGTTCAAAAGTCCAGCACTTCATCTAGTG |
| 1_hypothetical | N2 | Tb927.9.17730_for | GGGGACAAGTTTGTACAAAAAAGCAGGCTATGACAACGCTGCATGCGG | Tb927.9.17730_rev | GGGGACCACTTTGTACAAGAAAGCTGGGTTCAGTGCTGCGGGTCGCT |
| 1_hypothetical | N3 | Tb927.6.2680_for | GGGGACAAGTTTGTACAAAAAAGCAGGCTATGTACCGCACTCTCACACCG | Tb927.6.2680_rev | GGGGACCACTTTGTACAAGAAAGCTGGGTTCACAAATAACTATCCAAACCACGATAATA |
| 1_hypothetical | N4 | Tb927.8.2180_for | GGGGACAAGTTTGTACAAAAAAGCAGGCTATGCGACGCCGTGGGTCC | Tb927.8.2180_rev | GGGGACCACTTTGTACAAGAAAGCTGGGTTTAATGTCCATATCTCACCATCAGCAAAA |
| 1_hypothetical | N5 | Tb927.10.1760_for | GGGGACAAGTTTGTACAAAAAAGCAGGCTATGAACCGAAGAGTAGATTTCATATGCC | Tb927.10.1760_rev | GGGGACCACTTTGTACAAGAAAGCTGGGTCTACAGGGTGCAGAGGCGG |
| 1_hypothetical | N6 | Tb927.9.15820_for | GGGGACAAGTTTGTACAAAAAAGCAGGCTATGAATTGCATCCAGCCATCTGG | Tb927.9.15820_rev | GGGGACCACTTTGTACAAGAAAGCTGGGTTTAAAAAACCGACTTTCTGACGGCA |
| 1_hypothetical | N7 | Tb927.9.9260_for | GGGGACAAGTTTGTACAAAAAAGCAGGCTATGTCCGATGACGATGATGATGTG | Tb927.9.9260_rev | GGGGACCACTTTGTACAAGAAAGCTGGGTTTACTCGAGAACTGCTGTGTTGATAAG |
| 1_hypothetical | N8 | Tb927.11.4790_for | GGGGACAAGTTTGTACAAAAAAGCAGGCTATGCGCTTGAAAACTGCCGT | Tb927.11.4790_rev | GGGGACCACTTTGTACAAGAAAGCTGGGTTCAAGTGACCTCGAATAACACCCT |
| 1_hypothetical | N9 | Tb927.10.1010_for | GGGGACAAGTTTGTACAAAAAAGCAGGCTATGTATGAAACAGGAAGAGTAGCCACC | Tb927.10.1010_rev | GGGGACCACTTTGTACAAGAAAGCTGGGTTTAAGGGCCCTGCTTTCGCA |
| 1_hypothetical | N10 | Tb927.3.4050_for | GGGGACAAGTTTGTACAAAAAAGCAGGCTATGTTTTCGAGCTATGCAGGTTGC | Tb927.3.4050_rev | GGGGACCACTTTGTACAAGAAAGCTGGGTTCAACCACTGCCGACCGA |
| 1_hypothetical | N11 | Tb927.10.9610_for | GGGGACAAGTTTGTACAAAAAAGCAGGCTATGCAGGACGAAAGGGAAACAAG | Tb927.10.9610_rev | GGGGACCACTTTGTACAAGAAAGCTGGGTCTATTTAAATGCAAGTCGTGCTGTGTT |
| 1_hypothetical | N12 | Tb927.9.13480_for | GGGGACAAGTTTGTACAAAAAAGCAGGCTATGAAGAAGGCAAAAAAACAAGAGCC | Tb927.9.13480_rev | GGGGACCACTTTGTACAAGAAAGCTGGGTTTAAAAACAGCAGTCACAATCAAAGGG |
| 1_hypothetical | N13 | Tb927.4.3080_for | GGGGACAAGTTTGTACAAAAAAGCAGGCTATGAAGGTATCGTCGTTTATTGTTCAGC | Tb927.4.3080_rev | GGGGACCACTTTGTACAAGAAAGCTGGGTTCATGAAAAATCGGCAGACTTGAGC |
| 1_hypothetical | N14 | Tb927.11.8020_for | GGGGACAAGTTTGTACAAAAAAGCAGGCTATGGCACTTGAAGGGTCAACAC | Tb927.11.8020_rev | GGGGACCACTTTGTACAAGAAAGCTGGGTCTACACGTGCCTCAGTTCATTCA |
| 1_hypothetical | N15 | Tb927.8.5960_for | GGGGACAAGTTTGTACAAAAAAGCAGGCTATGCCCCCAAAGGATAACAGAAAAG | Tb927.8.5960_rev | GGGGACCACTTTGTACAAGAAAGCTGGGTCTACTGGGTCGAATCTCCAGATTTTG |
| 1_hypothetical | N16 | Tb927.9.13240_for | GGGGACAAGTTTGTACAAAAAAGCAGGCTATGAGTTCTGATGAGTTAGATGCTTTCC | Tb927.9.13240_rev | GGGGACCACTTTGTACAAGAAAGCTGGGTTTAACTTCGAAGCCTTGGTCGC |
| 1_hypothetical | N17 | Tb927.4.1130_for | GGGGACAAGTTTGTACAAAAAAGCAGGCTATGTATAGCAAGACACCGTGTGTTT | Tb927.4.1130_rev | GGGGACCACTTTGTACAAGAAAGCTGGGTTCATTTACTAAAGTCCTCAGCGTCAGC |
| 1_hypothetical | N18 | Tb927.11.14050_for | GGGGACAAGTTTGTACAAAAAAGCAGGCTATGGGTTTAGAAATATTTGAGATTGACACC | Tb927.11.14050_rev | GGGGACCACTTTGTACAAGAAAGCTGGGTTCAGACAAATGCCAACTTCTGATTCA |
| 1_hypothetical | N19 | Tb927.7.7060_for | GGGGACAAGTTTGTACAAAAAAGCAGGCTATGAGCGCTTTTGTAGATGTAGTTAAGA | Tb927.7.7060_rev | GGGGACCACTTTGTACAAGAAAGCTGGGTTTACACCACACCTCTCAAGCAAC |
| 1_hypothetical | N20 | Tb927.4.1040_for | GGGGACAAGTTTGTACAAAAAAGCAGGCTATGGAACAAGATTCACATCCCAGGA | Tb927.4.1040_rev | GGGGACCACTTTGTACAAGAAAGCTGGGTTTAGACAAGGTCGTATACCTTGTTCACT |
| 1_hypothetical | N21 | Tb927.8.6010_for | GGGGACAAGTTTGTACAAAAAAGCAGGCTATGGCTCTTTCAGAGAAAGCGTG | Tb927.8.6010_rev | GGGGACCACTTTGTACAAGAAAGCTGGGTCTACAACATGCTCGTCTGAGGGC |
| 1_hypothetical | N22 | Tb927.8.1430_for | GGGGACAAGTTTGTACAAAAAAGCAGGCTATGTTCTGCTTCAGTAGAGGATTGTG | Tb927.8.1430_rev | GGGGACCACTTTGTACAAGAAAGCTGGGTCTAAATCCGACTATACTGAAGAACGTTGC |
| 1_hypothetical | N23 | Tb927.8.3780_for | GGGGACAAGTTTGTACAAAAAAGCAGGCTATGCCACCCAAAAAGAAGGTGG | Tb927.8.3780_rev | GGGGACCACTTTGTACAAGAAAGCTGGGTCTAACCCTCTTTTTCATCCTTCAGTACAT |
| 1_hypothetical | N24 | Tb927.1.4160_for | GGGGACAAGTTTGTACAAAAAAGCAGGCTATGCAGATTTATGTGAAGCATCGCG | Tb927.1.4160_rev | GGGGACCACTTTGTACAAGAAAGCTGGGTTTATTTATTAACTTCCACCGCTTTCTGTTG |
| 1_hypothetical | O1 | Tb927.2.2570_for | GGGGACAAGTTTGTACAAAAAAGCAGGCTATGTCTCCGCGTACCTCTGAA | Tb927.2.2570_rev | GGGGACCACTTTGTACAAGAAAGCTGGGTTTATTCGTCATCGTCATCAGCAGAG |
| 1_hypothetical | O2 | Tb927.9.17940_for | GGGGACAAGTTTGTACAAAAAAGCAGGCTATGACATGTAGGAGTTCTCACACCT | Tb927.9.17940_rev | GGGGACCACTTTGTACAAGAAAGCTGGGTTCATTCAGATCCAAAGCCGATAAATATAAG |
| 1_hypothetical | O3 | Tb927.11.13390_for | GGGGACAAGTTTGTACAAAAAAGCAGGCTATGAGTCAGGAGGCAGGATGC | Tb927.11.13390_rev | GGGGACCACTTTGTACAAGAAAGCTGGGTCTATTGCGATTGGAGTTTATCGACTGC |
| 1_hypothetical | O4 | Tb927.10.11900_for | GGGGACAAGTTTGTACAAAAAAGCAGGCTATGACCCAAAACATTATTCGTCGCA | Tb927.10.11900_rev | GGGGACCACTTTGTACAAGAAAGCTGGGTCTAACTCTCTGTATTCAGAGCAGCCT |
| 1_hypothetical | O5 | Tb927.9.13530_for | GGGGACAAGTTTGTACAAAAAAGCAGGCTATGAGGCGGATCATGTCCCC | Tb927.9.13530_rev | GGGGACCACTTTGTACAAGAAAGCTGGGTTTACCACATCGCGTCATTTCCC |
| 1_hypothetical | O6 | Tb927.6.4490_for | GGGGACAAGTTTGTACAAAAAAGCAGGCTATGGCGCCCAGCAGTGAA | Tb927.6.4490_rev | GGGGACCACTTTGTACAAGAAAGCTGGGTTTAGTTCAGAAATAGCTTTCTTGACTCACT |
| 1_hypothetical | O7 | Tb927.8.4650_for | GGGGACAAGTTTGTACAAAAAAGCAGGCTATGCAAAGGTGTGTCCACATACTC | Tb927.8.4650_rev | GGGGACCACTTTGTACAAGAAAGCTGGGTTTACACAGCGGGCTTGAACG |
| 1_hypothetical | O8 | Tb927.9.10560_for | GGGGACAAGTTTGTACAAAAAAGCAGGCTATGGATAGCCTTGCACACAGTG | Tb927.9.10560_rev | GGGGACCACTTTGTACAAGAAAGCTGGGTTTAGTTGGAAGTACGGCTTTCATGTAAG |
| 1_hypothetical | O9 | Tb927.11.1440_for | GGGGACAAGTTTGTACAAAAAAGCAGGCTATGATGTGCAATTCCTTTGTTGCC | Tb927.11.1440_rev | GGGGACCACTTTGTACAAGAAAGCTGGGTTTACAAAGTTCCTCCGTCTCCATATTG |
| 1_hypothetical | O10 | Tb927.10.5720_for | GGGGACAAGTTTGTACAAAAAAGCAGGCTATGGGGAATTTTGGGGGTACACC | Tb927.10.5720_rev | GGGGACCACTTTGTACAAGAAAGCTGGGTCTAGGGTGTATTTGCATATTTCCTGCT |
| 1_hypothetical | O11 | Tb927.9.10230_for | GGGGACAAGTTTGTACAAAAAAGCAGGCTATGCACATTCCAACCAAGTGGG | Tb927.9.10230_rev | GGGGACCACTTTGTACAAGAAAGCTGGGTTTATTCCTCCAAATCCTTCAAGTCCG |
| 1_hypothetical | O12 | Tb927.11.15390_for | GGGGACAAGTTTGTACAAAAAAGCAGGCTATGAAGGCGTTAAAGACAAATATTACCGT | Tb927.11.15390_rev | GGGGACCACTTTGTACAAGAAAGCTGGGTTTAAGCATAAATTTCACGACAAAATTTCAT |
| 1_hypothetical | O13 | Tb927.1.2750_for | GGGGACAAGTTTGTACAAAAAAGCAGGCTATGGAGTCAGGGAAGAATGCTCA | Tb927.1.2750_rev | GGGGACCACTTTGTACAAGAAAGCTGGGTTTAGCCCCCAAAGCGGCT |
| 1_hypothetical | O14 | Tb927.9.3340_for | GGGGACAAGTTTGTACAAAAAAGCAGGCTATGTCGTACTACGACAAGTCTGACA | Tb927.9.3340_rev | GGGGACCACTTTGTACAAGAAAGCTGGGTTTAGCTGAGCTTTCGCTCGTC |
| 1_hypothetical | O15 | Tb927.11.11130_for | GGGGACAAGTTTGTACAAAAAAGCAGGCTATGTTCCACAGGGCATTCGTC | Tb927.11.11130_rev | GGGGACCACTTTGTACAAGAAAGCTGGGTTTAATCGTTGTGCCCATGTGTGT |
| 1_hypothetical | O16 | Tb927.10.8980_for | GGGGACAAGTTTGTACAAAAAAGCAGGCTATGTTGCGCTTTACACGTTTTGTT | Tb927.10.8980_rev | GGGGACCACTTTGTACAAGAAAGCTGGGTTTAAGCACTCGCCTTACCCATCG |
| 1_hypothetical | O17 | Tb927.9.9230_for | GGGGACAAGTTTGTACAAAAAAGCAGGCTATGCAGGAACTTGCGGTTTTTAAAC | Tb927.9.9230_rev | GGGGACCACTTTGTACAAGAAAGCTGGGTTCAGTTTAGGAGAACGCGGAGC |
| 1_hypothetical | O18 | Tb05.30F7.410_for | GGGGACAAGTTTGTACAAAAAAGCAGGCTATGCGGGAAAATGCAACTGCA | Tb05.30F7.410_rev | GGGGACCACTTTGTACAAGAAAGCTGGGTTCATAATACACAAATGCCTACGGTACTTC |
| 1_hypothetical | O19 | Tb927.1.1650_for | GGGGACAAGTTTGTACAAAAAAGCAGGCTATGTGGCGGGTGAAACCCT | Tb927.1.1650_rev | GGGGACCACTTTGTACAAGAAAGCTGGGTTCACTCCAAAGCGATGACTCGA |
| 1_hypothetical | O20 | Tb927.9.8710_for | GGGGACAAGTTTGTACAAAAAAGCAGGCTATGTCAGTGCTGAGGTTCTGGC | Tb927.9.8710_rev | GGGGACCACTTTGTACAAGAAAGCTGGGTCTAAGAATCATTCCTCTGATATCTGGGACT |
| 1_hypothetical | O21 | Tb927.11.1270_for | GGGGACAAGTTTGTACAAAAAAGCAGGCTATGACAAAATATGAGCTGAAGATGCAGT | Tb927.11.1270_rev | GGGGACCACTTTGTACAAGAAAGCTGGGTCTAATAATTGTAGATGGCCTGCTGAGAA |
| 1_hypothetical | O22 | Tb927.4.3710_for | GGGGACAAGTTTGTACAAAAAAGCAGGCTATGAGAGTGAACAGGACGCAGC | Tb927.4.3710_rev | GGGGACCACTTTGTACAAGAAAGCTGGGTTCAGTCACCTTTTCCATGTTTTATGTCG |
| 1_hypothetical | O23 | Tb927.5.4450_for | GGGGACAAGTTTGTACAAAAAAGCAGGCTATGTCAATGAGCAACGGAAGTAGTG | Tb927.5.4450_rev | GGGGACCACTTTGTACAAGAAAGCTGGGTTCACTTTACAGCAGTTTTTTCATACAGC |
| 1_hypothetical | O24 | Tb927.7.6400_for | GGGGACAAGTTTGTACAAAAAAGCAGGCTATGCCAGTTGCTAATGAAGCTTCC | Tb927.7.6400_rev | GGGGACCACTTTGTACAAGAAAGCTGGGTTTATATGAAAATTTGCGGGTAAGCACCT |

| | | | | | |
|---|---|---|---|---|---|
| 1_hypothetical | P1 | Tb927.10.12360_for | GGGGACAAGTTTGTACAAAAAAGCAGGCTATGAAAGCAACAAAAGCAAAATCCGG | Tb927.10.12360_rev | GGGGACCACTTTGTACAAGAAAGCTGGGTTCAAATGTTGTTGGTATCGGAAAGCA |
| 1_hypothetical | P2 | Tb927.8.5350_for | GGGGACAAGTTTGTACAAAAAAGCAGGCTATGGAAAAGCCACTGGATAACGG | Tb927.8.5350_rev | GGGGACCACTTTGTACAAGAAAGCTGGGTTCAGCCCCCGTGCGCATA |
| 1_hypothetical | P3 | Tb927.8.3160_for | GGGGACAAGTTTGTACAAAAAAGCAGGCTATGTTCCGCATCTCATTCACTTCC | Tb927.8.3160_rev | GGGGACCACTTTGTACAAGAAAGCTGGGTCTACAACAGACCTTTCAGTGGGTTG |
| 1_hypothetical | P4 | Tb927.7.5795_for | GGGGACAAGTTTGTACAAAAAAGCAGGCTATGGCATATAGCGACTATAACAAACAGG | Tb927.7.5795_rev | GGGGACCACTTTGTACAAGAAAGCTGGGTTCATCTTTTTCGCATTGTAATGACTGGA |
| 1_hypothetical | P5 | Tb927.4.3730_for | GGGGACAAGTTTGTACAAAAAAGCAGGCTATGTTTACAGGACGTTGGGTTTACAC | Tb927.4.3730_rev | GGGGACCACTTTGTACAAGAAAGCTGGGTCTAAGTGAGACACTTGCCTCCCA |
| 1_hypothetical | P6 | Tb927.11.11170_for | GGGGACAAGTTTGTACAAAAAAGCAGGCTATGAACCTAGAGCTGTATGTAACACTCG | Tb927.11.11170_rev | GGGGACCACTTTGTACAAGAAAGCTGGGTTCAGTCGTTGTTTGTGCGTGG |
| 1_hypothetical | P7 | Tb927.10.900_for | GGGGACAAGTTTGTACAAAAAAGCAGGCTATGCGTCGAGGGCACCAC | Tb927.10.900_rev | GGGGACCACTTTGTACAAGAAAGCTGGGTTCATCCACCTTTGATCGTGTTGC |
| 1_hypothetical | P8 | Tb927.10.9030_for | GGGGACAAGTTTGTACAAAAAAGCAGGCTATGAAGCGATTTTTTGACAAGCTATACC | Tb927.10.9030_rev | GGGGACCACTTTGTACAAGAAAGCTGGGTTTACTTCGTTCCAGAAGCAGAATCATT |
| 1_hypothetical | P9 | Tb927.7.6330_for | GGGGACAAGTTTGTACAAAAAAGCAGGCTATGGTGCCCTTATTTAAGACACCAAA | Tb927.7.6330_rev | GGGGACCACTTTGTACAAGAAAGCTGGGTTCAACGTAAGCATAGTGGAAATAATAACAA |
| 1_hypothetical | P10 | Tb927.10.2160_for | GGGGACAAGTTTGTACAAAAAAGCAGGCTATGGATGCGGAAGAGTACCTGC | Tb927.10.2160_rev | GGGGACCACTTTGTACAAGAAAGCTGGGTTCAATGAACTTGATCCGCACAGATG |
| 1_hypothetical | P11 | Tb927.9.12330_for | GGGGACAAGTTTGTACAAAAAAGCAGGCTATGCCACCAAAGCGTGCTG | Tb927.9.12330_rev | GGGGACCACTTTGTACAAGAAAGCTGGGTTCATTCAACTGCTGCCGGTG |
| 1_hypothetical | P12 | Tb927.7.1010_for | GGGGACAAGTTTGTACAAAAAAGCAGGCTATGCCACCGAATTCGGCTAC | Tb927.7.1010_rev | GGGGACCACTTTGTACAAGAAAGCTGGGTTCACTCCTCCGATCGGAGGG |
| 1_hypothetical | P13 | Tb927.4.2090_for | GGGGACAAGTTTGTACAAAAAAGCAGGCTATGTTACTGCCACACCCCTCG | Tb927.4.2090_rev | GGGGACCACTTTGTACAAGAAAGCTGGGTTCAAATACTGGAGGTCCCCCCC |
| 1_hypothetical | P14 | Tb927.6.2180_for | GGGGACAAGTTTGTACAAAAAAGCAGGCTATGTTACGTCGCTGCCCCT | Tb927.6.2180_rev | GGGGACCACTTTGTACAAGAAAGCTGGGTTTACTTGGGCAGCAAACCGG |
| 1_hypothetical | P15 | Tb927.8.3210_for | GGGGACAAGTTTGTACAAAAAAGCAGGCTATGACGGACACCACCAAAGGT | Tb927.8.3210_rev | GGGGACCACTTTGTACAAGAAAGCTGGGTCTACAAACGTGGGCGTTTCCT |
| 1_hypothetical | P16 | Tb927.3.5300_for | GGGGACAAGTTTGTACAAAAAAGCAGGCTATGTTATTCACCCGCTGTCTGC | Tb927.3.5300_rev | GGGGACCACTTTGTACAAGAAAGCTGGGTCTACCATTTCGGCGATGATGGC |
| 1_hypothetical | P17 | Tb927.9.910_for | GGGGACAAGTTTGTACAAAAAAGCAGGCTATGAAGCTTGCAATAGCGTTAGCA | Tb927.9.910_rev | GGGGACCACTTTGTACAAGAAAGCTGGGTTTATAGGCCGCCCGCCGT |
| 1_hypothetical | P18 | Tb927.2.5070_for | GGGGACAAGTTTGTACAAAAAAGCAGGCTATGTCGGGTGACGGTAGTGAT | Tb927.2.5070_rev | GGGGACCACTTTGTACAAGAAAGCTGGGTTCATAGACTGAATCGAGCAATCGACA |
| 1_hypothetical | P19 | Tb927.7.430_for | GGGGACAAGTTTGTACAAAAAAGCAGGCTATGCATGATTTGGTCGCTGTTGT | Tb927.7.430_rev | GGGGACCACTTTGTACAAGAAAGCTGGGTCTAACTGCAGCCAGAATGCTTTTC |
| 1_hypothetical | P21 | Tb927.10.1650_for | GGGGACAAGTTTGTACAAAAAAGCAGGCTATGCTATCGAAAATGAAATGCACTCTTAC | Tb927.10.1650_rev | GGGGACCACTTTGTACAAGAAAGCTGGGTTCAAACGAAGAAGTGAGAAAAGGGATT |
| 1_hypothetical | P22 | Tb927.7.6870_for | GGGGACAAGTTTGTACAAAAAAGCAGGCTATGTTGATTGGCATGAAATATTCTGCTC | Tb927.7.6870_rev | GGGGACCACTTTGTACAAGAAAGCTGGGTTCAAAAGGGATTTGGTTCTTTCTTTTCG |
| 1_hypothetical | P23 | Tb927.11.10980_for | GGGGACAAGTTTGTACAAAAAAGCAGGCTATGTCGAAGTACGTGAATTTAGACTCTC | Tb927.11.10980_rev | GGGGACCACTTTGTACAAGAAAGCTGGGTTCACGTAAAACGTGAAAGCACCT |
| 1_hypothetical | P24 | Tb927.10.12730_for | GGGGACAAGTTTGTACAAAAAAGCAGGCTATGACGAGTAACCGGATTTACTGTTTG | Tb927.10.12730_rev | GGGGACCACTTTGTACAAGAAAGCTGGGTTTAATGTCGACTGTAACAAATAAAATAAAA |
| 1_known | A1 | Tb927.7.570_for | GGGGACAAGTTTGTACAAAAAAGCAGGCTATGGAAAAGTACGAATTTCCCGAGG | Tb927.7.570_rev | GGGGACCACTTTGTACAAGAAAGCTGGGTTTATTTCTCTGAGGTTTCCTTCTTGTTATC |
| 1_known | A2 | Tb927.5.3040_for | GGGGACAAGTTTGTACAAAAAAGCAGGCTATGCTACGTCGAACGCGTTT | Tb927.5.3040_rev | GGGGACCACTTTGTACAAGAAAGCTGGGTTTAGGTGTGCGTCGGCACT |
| 1_known | A3 | Tb927.3.5090_for | GGGGACAAGTTTGTACAAAAAAGCAGGCTATGCCCTCCGCTGAAACACT | Tb927.3.5090_rev | GGGGACCACTTTGTACAAGAAAGCTGGGTTCAACTCCACGACTGCCAAAATT |
| 1_known | A4 | Tb927.2.2460_for | GGGGACAAGTTTGTACAAAAAAGCAGGCTATGTCCGGGCTATCTTTAGCTCA | Tb927.2.2460_rev | GGGGACCACTTTGTACAAGAAAGCTGGGTTTATACCCGCTTCCGGTGGT |
| 1_known | A5 | Tb927.8.6510_for | GGGGACAAGTTTGTACAAAAAAGCAGGCTATGCTCTCACGCATCAAGAGGC | Tb927.8.6510_rev | GGGGACCACTTTGTACAAGAAAGCTGGGTTCAGTCGATATAATGCCCACTCATCC |
| 1_known | A6 | Tb927.11.13750_for | GGGGACAAGTTTGTACAAAAAAGCAGGCTATGTCAGAGAGATATCAACAGTTAATGAAG | Tb927.11.13750_rev | GGGGACCACTTTGTACAAGAAAGCTGGGTCTAACAAGCACACGAAAACCGTTT |
| 1_known | A7 | Tb927.11.15910_for | GGGGACAAGTTTGTACAAAAAAGCAGGCTATGACCTTCAGCATTCCACCACT | Tb927.11.15910_rev | GGGGACCACTTTGTACAAGAAAGCTGGGTCTAGCTTTTCAGCAGCTGCTCC |
| 1_known | A8 | Tb927.7.4770_for | GGGGACAAGTTTGTACAAAAAAGCAGGCTATGCTTTTTTTTGACATTTCAGCGAAAAC | Tb927.7.4770_rev | GGGGACCACTTTGTACAAGAAAGCTGGGTTTACAAGACACCGCAGTCGGA |
| 1_known | A9 | Tb927.7.3750_for | GGGGACAAGTTTGTACAAAAAAGCAGGCTATGAGACCATTGCTGGTTTTTGGA | Tb927.7.3750_rev | GGGGACCACTTTGTACAAGAAAGCTGGGTTCACTGTTGGCCCCATGGG |
| 1_known | A10 | Tb927.1.1990_for | GGGGACAAGTTTGTACAAAAAAGCAGGCTATGAATAATGAGCGACCGGCG | Tb927.1.1990_rev | GGGGACCACTTTGTACAAGAAAGCTGGGTTCACAGGAGCGGAGAAAAGTTAAC |
| 1_known | A11 | Tb927.11.15030_for | GGGGACAAGTTTGTACAAAAAAGCAGGCTATGTCCTTAAAAGTGAAGAAACTAATTGAC | Tb927.11.15030_rev | GGGGACCACTTTGTACAAGAAAGCTGGGTTTATTCTTTGATGAAGCCAAGTACCTGC |
| 1_known | A12 | Tb927.9.5840_for | GGGGACAAGTTTGTACAAAAAAGCAGGCTATGTCCTGCGGTGATGCGA | Tb927.9.5840_rev | GGGGACCACTTTGTACAAGAAAGCTGGGTTTAGTTCATGCTGCTGAAGTAATCTTGG |
| 1_known | A13 | Tb927.10.13370_for | GGGGACAAGTTTGTACAAAAAAGCAGGCTATGTCTGATGAGACGCCTCAGG | Tb927.10.13370_rev | GGGGACCACTTTGTACAAGAAAGCTGGGTTTAAATCACGCTCCAATAAACACTTACCA |
| 1_known | A14 | Tb927.10.10670_for | GGGGACAAGTTTGTACAAAAAAGCAGGCTATGCTGAAGTTTCGCGAGCG | Tb927.10.10670_rev | GGGGACCACTTTGTACAAGAAAGCTGGGTTCAGTTGGTGCGAGCTGC |
| 1_known | A15 | Tb927.11.7480_for | GGGGACAAGTTTGTACAAAAAAGCAGGCTATGAAATACTCTTTTATAGGTTTACTCTCC | Tb927.11.7480_rev | GGGGACCACTTTGTACAAGAAAGCTGGGTTCATGAAGAGTCACAGAGTCCCG |
| 1_known | A17 | Tb927.1.650_for | GGGGACAAGTTTGTACAAAAAAGCAGGCTATGGACGAGATCAAGTGGCCC | Tb927.1.650_rev | GGGGACCACTTTGTACAAGAAAGCTGGGTTTATCGGAGGGGTATGTATTCCGATG |
| 1_known | A18 | Tb927.6.1820_for | GGGGACAAGTTTGTACAAAAAAGCAGGCTATGCACGCCAACTGGATTACTTG | Tb927.6.1820_rev | GGGGACCACTTTGTACAAGAAAGCTGGGTTCATAATCGGACATACGTCACCCA |
| 1_known | A19 | Tb927.9.8450_for | GGGGACAAGTTTGTACAAAAAAGCAGGCTATGACATCCACCGAGAAGCCG | Tb927.9.8450_rev | GGGGACCACTTTGTACAAGAAAGCTGGGTTCATTTCTTCAGTAGTTCCTCCCCC |
| 1_known | A20 | Tb927.11.530_for | GGGGACAAGTTTGTACAAAAAAGCAGGCTATGGAACCGGAGTGTGAAAGC | Tb927.11.530_rev | GGGGACCACTTTGTACAAGAAAGCTGGGTTTAATTCTGCTGCTGCTGCTGC |
| 1_known | A21 | Tb927.3.4240_for | GGGGACAAGTTTGTACAAAAAAGCAGGCTATGTCTGCTACCGTGGGCC | Tb927.3.4240_rev | GGGGACCACTTTGTACAAGAAAGCTGGGTTCAATTTGTCGCCCCACCCT |
| 1_known | A22 | Tb927.8.1680_for | GGGGACAAGTTTGTACAAAAAAGCAGGCTATGATGGCGAGTGCCCCC | Tb927.8.1680_rev | GGGGACCACTTTGTACAAGAAAGCTGGGTTCAACTGGGGTCACCAATGAGT |
| 1_known | A23 | Tb927.10.5840_for | GGGGACAAGTTTGTACAAAAAAGCAGGCTATGTCATCGCTAAAGGAAATCAACGG | Tb927.10.5840_rev | GGGGACCACTTTGTACAAGAAAGCTGGGTTTAGATCTTGTTCCAAGCAACAATATCCA |
| 1_known | A24 | Tb927.3.1750_for | GGGGACAAGTTTGTACAAAAAAGCAGGCTATGGCAGAGAGGTTACAAACCATTAC | Tb927.3.1750_rev | GGGGACCACTTTGTACAAGAAAGCTGGGTCTATGAACCATCTCTATAAGCCACGGA |
| 1_known | B1 | Tb927.11.1320_for | GGGGACAAGTTTGTACAAAAAAGCAGGCTATGCTTCGTCGCACGTCG | Tb927.11.1320_rev | GGGGACCACTTTGTACAAGAAAGCTGGGTTTAATCTCGAACAGAATACTTTTGAATTTC |
| 1_known | B2 | Tb927.8.7010_for | GGGGACAAGTTTGTACAAAAAAGCAGGCTATGATGCGCCGTGCCTGT | Tb927.8.7010_rev | GGGGACCACTTTGTACAAGAAAGCTGGGTCTAAGACGACGAACCATAACTATAAAAAGA |
| 1_known | B3 | Tb927.6.490_for | GGGGACAAGTTTGTACAAAAAAGCAGGCTATGTTCCGGAAGTCTTCGGAGT | Tb927.6.490_rev | GGGGACCACTTTGTACAAGAAAGCTGGGTTTACACAAATCTAAAGGCCAAGCCA |
| 1_known | B5 | Tb927.7.5460_for | GGGGACAAGTTTGTACAAAAAAGCAGGCTATGCAATTCGATGAAGATATTGTGCCC | Tb927.7.5460_rev | GGGGACCACTTTGTACAAGAAAGCTGGGTTCAGGAGTTGTTAGTTCCCGTAACG |
| 1_known | B6 | Tb927.9.12150_for | GGGGACAAGTTTGTACAAAAAAGCAGGCTATGCTCAACAGGGACGGTACC | Tb927.9.12150_rev | GGGGACCACTTTGTACAAGAAAGCTGGGTTCAGTCACGTATTCTCCTTTCTCGG |
| 1_known | B8 | Tb927.10.12330_for | GGGGACAAGTTTGTACAAAAAAGCAGGCTATGCAACCATTTGGTCAAACGC | Tb927.10.12330_rev | GGGGACCACTTTGTACAAGAAAGCTGGGTTTAGGGGGATTCGGCAGCC |
| 1_known | B9 | Tb927.11.4570_for | GGGGACAAGTTTGTACAAAAAAGCAGGCTATGTCTGTGAAGACGCTTGCC | Tb927.11.4570_rev | GGGGACCACTTTGTACAAGAAAGCTGGGTTCAACCACAGCAACCGGAAC |
| 1_known | B10 | Tb927.9.16190_for | GGGGACAAGTTTGTACAAAAAAGCAGGCTATGATTAGTAGAAAAACATATTTGTGGTTC | Tb927.9.16190_rev | GGGGACCACTTTGTACAAGAAAGCTGGGTCTAGCATGTAAATTCATGCCAAAACGC |
| 1_known | B11 | Tb927.11.14280_for | GGGGACAAGTTTGTACAAAAAAGCAGGCTATGGCTGAAGAGATTCGTACCATGA | Tb927.11.14280_rev | GGGGACCACTTTGTACAAGAAAGCTGGGTTCATAGTTGGCTAAGTAGTTTTTCAACCA |
| 1_known | B12 | Tb927.11.7100_for | GGGGACAAGTTTGTACAAAAAAGCAGGCTATGCAACAGGAGAACCGCCA | Tb927.11.7100_rev | GGGGACCACTTTGTACAAGAAAGCTGGGTTCACCGAGCGCCGATGTG |
| 1_known | B13 | Tb927.11.3080_for | GGGGACAAGTTTGTACAAAAAAGCAGGCTATGAGCACTACGAACCCACTCG | Tb927.11.3080_rev | GGGGACCACTTTGTACAAGAAAGCTGGGTTTACCGTGCACCGATGCTAATAC |
| 1_known | B14 | Tb927.9.4040_for | GGGGACAAGTTTGTACAAAAAAGCAGGCTATGTCTACATCCCGCATTACGTTG | Tb927.9.4040_rev | GGGGACCACTTTGTACAAGAAAGCTGGGTTCAGAAGCTGGAAAGGTCTGTGC |
| 1_known | B16 | Tb927.6.1380_for | GGGGACAAGTTTGTACAAAAAAGCAGGCTATGAAGGAGGAACAGAACTCCTGC | Tb927.6.1380_rev | GGGGACCACTTTGTACAAGAAAGCTGGGTCTATTCCTCTCCTGCACAACTCAC |
| 1_known | B20 | Tb927.3.4890_for | GGGGACAAGTTTGTACAAAAAAGCAGGCTATGTTACGCAGATCGGTGGTTAAC | Tb927.3.4890_rev | GGGGACCACTTTGTACAAGAAAGCTGGGTTCAAATCCTTTTCGCTAAAGCCACA |
| 1_known | B21 | Tb927.6.4720_for | GGGGACAAGTTTGTACAAAAAAGCAGGCTATGCGACCGGAGCAGTGT | Tb927.6.4720_rev | GGGGACCACTTTGTACAAGAAAGCTGGGTCTAATTCGCACGGGTTATGTTCACG |
| 1_known | B22 | Tb927.6.1210_for | GGGGACAAGTTTGTACAAAAAAGCAGGCTATGCATTCCATTCTACCATACACATTAGAT | Tb927.6.1210_rev | GGGGACCACTTTGTACAAGAAAGCTGGGTTTACCTAGCTCCCAAGACAGAAGC |
| 1_known | B23 | Tb927.10.12450_for | GGGGACAAGTTTGTACAAAAAAGCAGGCTATGCTGTATTTTACCTTGATTGTCCGA | Tb927.10.12450_rev | GGGGACCACTTTGTACAAGAAAGCTGGGTTCAAGCGGAGAGGTAAAATACGTATACC |
| 1_known | B24 | Tb927.10.3150_for | GGGGACAAGTTTGTACAAAAAAGCAGGCTATGGTATCCAACGGTAAAACCAACG | Tb927.10.3150_rev | GGGGACCACTTTGTACAAGAAAGCTGGGTTCACTTCGCCTTCCCACCC |
| 1_known | C1 | Tb927.9.14080_for | GGGGACAAGTTTGTACAAAAAAGCAGGCTATGAAGCTTTACTCCCTGGCAATTG | Tb927.9.14080_rev | GGGGACCACTTTGTACAAGAAAGCTGGGTTCACATGACGGTGCAACACC |
| 1_known | C2 | Tb927.11.7270_for | GGGGACAAGTTTGTACAAAAAAGCAGGCTATGTCCATTCTCACGTACAGCGG | Tb927.11.7270_rev | GGGGACCACTTTGTACAAGAAAGCTGGGTCTAGTCTTTGCGACCCTTGATGAG |
| 1_known | C3 | Tb927.10.7160_for | GGGGACAAGTTTGTACAAAAAAGCAGGCTATGGTGGAGTTGGTGAGGTTGA | Tb927.10.7160_rev | GGGGACCACTTTGTACAAGAAAGCTGGGTTTAGAACCTCTGCACGGGTTTTTC |
| 1_known | C4 | Tb927.11.1710_for | GGGGACAAGTTTGTACAAAAAAGCAGGCTATGATTCGACTCGCATGCCTG | Tb927.11.1710_rev | GGGGACCACTTTGTACAAGAAAGCTGGGTTTAATGGTATCGCGATGTGTCACTTAC |
| 1_known | C5 | Tb927.10.11700_for | GGGGACAAGTTTGTACAAAAAAGCAGGCTATGAGCTCTGGAAGCACGAAGC | Tb927.10.11700_rev | GGGGACCACTTTGTACAAGAAAGCTGGGTTCACTTATTTAACCCCTCCATACCATATTT |
| 1_known | C6 | Tb927.10.4710_for | GGGGACAAGTTTGTACAAAAAAGCAGGCTATGGCAGAGACGACTATCGGG | Tb927.10.4710_rev | GGGGACCACTTTGTACAAGAAAGCTGGGTCTAGCTTACAGATTGCACTCGCTC |
| 1_known | C7 | Tb927.8.3370_for | GGGGACAAGTTTGTACAAAAAAGCAGGCTATGCAGGAGAAGGAGGTGCA | Tb927.8.3370_rev | GGGGACCACTTTGTACAAGAAAGCTGGGTTCATTCTTCCGGTACAAAAAGACCC |
| 1_known | C8 | Tb927.7.6380_for | GGGGACAAGTTTGTACAAAAAAGCAGGCTATGTCGGTAGAGGCGAGTAGAC | Tb927.7.6380_rev | GGGGACCACTTTGTACAAGAAAGCTGGGTTCACTCTTTCTTTGCTTTGGTCCG |
| 1_known | C9 | Tb927.10.4590_for | GGGGACAAGTTTGTACAAAAAAGCAGGCTATGGTGATGCGTTGGGGATTTT | Tb927.10.4590_rev | GGGGACCACTTTGTACAAGAAAGCTGGGTTTATGGTCCCTCACCCGCTC |
| 1_known | C10 | Tb927.11.8470_for | GGGGACAAGTTTGTACAAAAAAGCAGGCTATGAGCGTGGGCAAACCTATCA | Tb927.11.8470_rev | GGGGACCACTTTGTACAAGAAAGCTGGGTTTACCTTTGCTGCGCAGTAAGC |
| 1_known | C11 | Tb927.5.3220_for | GGGGACAAGTTTGTACAAAAAAGCAGGCTATGTCATTGCTGCATCGGGC | Tb927.5.3220_rev | GGGGACCACTTTGTACAAGAAAGCTGGGTTTACATTCCGGTGTACCAGCACC |
| 1_known | C12 | Tb927.11.15480_for | GGGGACAAGTTTGTACAAAAAAGCAGGCTATGGCCGATACGGAGGCA | Tb927.11.15480_rev | GGGGACCACTTTGTACAAGAAAGCTGGGTTTAAGCTTTTTCTTCGGCAGCACC |
| 1_known | C13 | Tb927.11.15020_for | GGGGACAAGTTTGTACAAAAAAGCAGGCTATGGCTTTCAGCATTCCACCAC | Tb927.11.15020_rev | GGGGACCACTTTGTACAAGAAAGCTGGGTTTACAAGTCACTATGCGGTGCC |
| 1_known | C14 | Tb927.7.5570_for | GGGGACAAGTTTGTACAAAAAAGCAGGCTATGGAAAAGCATAAGGAAGGAACTGAA | Tb927.7.5570_rev | GGGGACCACTTTGTACAAGAAAGCTGGGTTTACGGAACCTCCAACTTTTCCTTC |
| 1_known | C15 | Tb927.8.890_for | GGGGACAAGTTTGTACAAAAAAGCAGGCTATGTCTACAGAGTACGATCACCTTTTTAAG | Tb927.8.890_rev | GGGGACCACTTTGTACAAGAAAGCTGGGTTTAGCAACATCCACTCTTTTGACCTC |
| 1_known | C16 | Tb927.4.3610_for | GGGGACAAGTTTGTACAAAAAAGCAGGCTATGGCCCAACCCATGCTGG | Tb927.4.3610_rev | GGGGACCACTTTGTACAAGAAAGCTGGGTTCAGCAAGAGCCAAAATACTGAAAGC |
| 1_known | C17 | Tb927.11.10240_for | GGGGACAAGTTTGTACAAAAAAGCAGGCTATGCTTCGCCGTGTCGGC | Tb927.11.10240_rev | GGGGACCACTTTGTACAAGAAAGCTGGGTTTACTCGCTAGTTTTCGCCGAC |
| 1_known | C18 | Tb927.10.8890_for | GGGGACAAGTTTGTACAAAAAAGCAGGCTATGTTCCGCGCTTGTTTGCG | Tb927.10.8890_rev | GGGGACCACTTTGTACAAGAAAGCTGGGTTCACTTTTTTTTCGTTACCTTCTTTTCTTT |
| 1_known | C19 | Tb927.10.1470_for | GGGGACAAGTTTGTACAAAAAAGCAGGCTATGGAACCAGCTTGCAAATACGA | Tb927.10.1470_rev | GGGGACCACTTTGTACAAGAAAGCTGGGTTTACAGTTTTGCCTTCACAGCGG |
| 1_known | C20 | Tb927.11.5680_for | GGGGACAAGTTTGTACAAAAAAGCAGGCTATGTCAGGAACAACTTCCATTCGAG | Tb927.11.5680_rev | GGGGACCACTTTGTACAAGAAAGCTGGGTTTACCGGCCGCGCTCCGC |
| 1_known | C21 | Tb927.10.1400_for | GGGGACAAGTTTGTACAAAAAAGCAGGCTATGGAACCAGCTTGCAAATACGA | Tb927.10.1400_rev | GGGGACCACTTTGTACAAGAAAGCTGGGTTTACCGCTTGGCTTCTCCAGC |
| 1_known | C22 | Tb927.11.14080_for | GGGGACAAGTTTGTACAAAAAAGCAGGCTATGACCATTATGATGGACGGCCC | Tb927.11.14080_rev | GGGGACCACTTTGTACAAGAAAGCTGGGTTTACTGCTCCTGATTCGCCGA |
| 1_known | C24 | Tb927.8.4030_for | GGGGACAAGTTTGTACAAAAAAGCAGGCTATGACACGATTGGCGGGTCA | Tb927.8.4030_rev | GGGGACCACTTTGTACAAGAAAGCTGGGTTTAGTCTTCCTCATCCTCTTTCCCTTC |
| 1_known | D1 | Tb927.11.15240_for | GGGGACAAGTTTGTACAAAAAAGCAGGCTATGCAGCAGCACCATTATGTTTTC | Tb927.11.15240_rev | GGGGACCACTTTGTACAAGAAAGCTGGGTTCAGCAGAAGCAGCCACTTTC |
| 1_known | D2 | Tb927.10.12190_for | GGGGACAAGTTTGTACAAAAAAGCAGGCTATGACAGATAAGGATGGTGGCACT | Tb927.10.12190_rev | GGGGACCACTTTGTACAAGAAAGCTGGGTCTACACAAAATGGGCCAATTCATTCTC |
| 1_known | D3 | Tb927.3.1090_for | GGGGACAAGTTTGTACAAAAAAGCAGGCTATGACGTACTGCCCCATTACGC | Tb927.3.1090_rev | GGGGACCACTTTGTACAAGAAAGCTGGGTTCACTCTCTGGTTGATGAGGGAG |
| 1_known | D4 | Tb927.11.4960_for | GGGGACAAGTTTGTACAAAAAAGCAGGCTATGGGTAAGGATTCCAATGATGGGA | Tb927.11.4960_rev | GGGGACCACTTTGTACAAGAAAGCTGGGTCTAATAGCGACGCCATTTGTGCT |
| 1_known | D5 | Tb927.6.5020_for | GGGGACAAGTTTGTACAAAAAAGCAGGCTATGTCGGAAAGGGACAAGCAGC | Tb927.6.5020_rev | GGGGACCACTTTGTACAAGAAAGCTGGGTTTACCAATATAAACAACACTCCGACATCC |
| 1_known | D6 | Tb927.6.2050_for | GGGGACAAGTTTGTACAAAAAAGCAGGCTATGAGTGAGTATCACATGGATCTGGG | Tb927.6.2050_rev | GGGGACCACTTTGTACAAGAAAGCTGGGTTTACTTTTTACCCTTCTTCCTGAGCG |
| 1_known | D7 | Tb927.10.6060_for | GGGGACAAGTTTGTACAAAAAAGCAGGCTATGGCGGACGAAGCTTCCA | Tb927.10.6060_rev | GGGGACCACTTTGTACAAGAAAGCTGGGTTTATTTCACAGGGCAGGCACG |
| 1_known | D8 | Tb927.6.3870_for | GGGGACAAGTTTGTACAAAAAAGCAGGCTATGGCATCAGACCCGTCGG | Tb927.6.3870_rev | GGGGACCACTTTGTACAAGAAAGCTGGGTTCAATATGGCCCACCGCGT |
| 1_known | D9 | Tb927.10.11990_for | GGGGACAAGTTTGTACAAAAAAGCAGGCTATGCGTCCGTTAACGGAGGA | Tb927.10.11990_rev | GGGGACCACTTTGTACAAGAAAGCTGGGTCTACGTCAAGCGTGCCTCG |
| 1_known | D10 | Tb927.8.4770_for | GGGGACAAGTTTGTACAAAAAAGCAGGCTATGGCGCAAGGTGACAACAG | Tb927.8.4770_rev | GGGGACCACTTTGTACAAGAAAGCTGGGTCTAACAGCACACCGCGCC |
| 1_known | D11 | Tb927.10.5760_for | GGGGACAAGTTTGTACAAAAAAGCAGGCTATGGGTCTGCATATCCTTTTATTTGGAG | Tb927.10.5760_rev | GGGGACCACTTTGTACAAGAAAGCTGGGTTTACAACTTGCTCTTTTCGATACGAAGA |
| 1_known | D12 | Tb927.11.12950_for | GGGGACAAGTTTGTACAAAAAAGCAGGCTATGCCATCTCATACCACTTCCGA | Tb927.11.12950_rev | GGGGACCACTTTGTACAAGAAAGCTGGGTTTAAAAGAAAGCTGCCACCACATCC |
| 1_known | D13 | Tb927.8.4330_for | GGGGACAAGTTTGTACAAAAAAGCAGGCTATGGAAGACATGAACCTTACGTTCAAA | Tb927.8.4330_rev | GGGGACCACTTTGTACAAGAAAGCTGGGTTTAACAGCACCCGCCACTCG |
| 1_known | D14 | Tb927.11.13940_for | GGGGACAAGTTTGTACAAAAAAGCAGGCTATGTCCACTGACCGTTCATCTCG | Tb927.11.13940_rev | GGGGACCACTTTGTACAAGAAAGCTGGGTTTAGATGTCAATTGCGTCAGGTTCA |
| 1_known | D15 | Tb927.11.9420_for | GGGGACAAGTTTGTACAAAAAAGCAGGCTATGAGCGAGGCACGCCAG | Tb927.11.9420_rev | GGGGACCACTTTGTACAAGAAAGCTGGGTTCAGATAGATGACTCTGGATTGAAGAGG |
| 1_known | D16 | Tb927.5.3560_for | GGGGACAAGTTTGTACAAAAAAGCAGGCTATGCCCATTAAATATAGTTGTGTCAATGAA | Tb927.5.3560_rev | GGGGACCACTTTGTACAAGAAAGCTGGGTTTATGACTTGCAGTTGGAAAAGTTCGG |
| 1_known | D17 | Tb927.4.2030_for | GGGGACAAGTTTGTACAAAAAAGCAGGCTATGCCTTCATATCCGTCGCAC | Tb927.4.2030_rev | GGGGACCACTTTGTACAAGAAAGCTGGGTTCAGTTGCCTTCACCCAAATCG |
| 1_known | D18 | Tb927.3.1120_for | GGGGACAAGTTTGTACAAAAAAGCAGGCTATGCAGGCTTCCTCGACGG | Tb927.3.1120_rev | GGGGACCACTTTGTACAAGAAAGCTGGGTCTAGTCGTCCTCGCCCATCG |
| 1_known | D19 | Tb927.9.1610_for | GGGGACAAGTTTGTACAAAAAAGCAGGCTATGCCGAAAGCTAAAACTACCACA | Tb927.9.1610_rev | GGGGACCACTTTGTACAAGAAAGCTGGGTTCACTGGCGGCTTCCCGC |
| 1_known | D20 | Tb927.9.11840_for | GGGGACAAGTTTGTACAAAAAAGCAGGCTATGCTCTCAGCAGCTGCTGC | Tb927.9.11840_rev | GGGGACCACTTTGTACAAGAAAGCTGGGTTTAAAAGGTATCGTTAACACGATTCATCAC |
| 1_known | D21 | Tb927.3.2000_for | GGGGACAAGTTTGTACAAAAAAGCAGGCTATGGATGAGAAGCCGCGGT | Tb927.3.2000_rev | GGGGACCACTTTGTACAAGAAAGCTGGGTTCATTGGCGAGCAGCAGAC |
| 1_known | D22 | Tb927.11.12120_for | GGGGACAAGTTTGTACAAAAAAGCAGGCTATGGCAGCTGTCACCCAGC | Tb927.11.12120_rev | GGGGACCACTTTGTACAAGAAAGCTGGGTTTACGACAGCGTAGGAGGAACG |
| 1_known | D23 | Tb927.11.11520_for | GGGGACAAGTTTGTACAAAAAAGCAGGCTATGTCTGAGTTCCAAAGGTTTGTTAAGC | Tb927.11.11520_rev | GGGGACCACTTTGTACAAGAAAGCTGGGTCTATTTGATCTTGTTCCAGTTCAAATACGT |
| 1_known | D24 | Tb927.8.4610_for | GGGGACAAGTTTGTACAAAAAAGCAGGCTATGATCACAGCAGCTTCCCCT | Tb927.8.4610_rev | GGGGACCACTTTGTACAAGAAAGCTGGGTTTAACAGCAACCACCACGCTT |
| 1_known | E1 | Tb927.10.5110_for | GGGGACAAGTTTGTACAAAAAAGCAGGCTATGCGGGTGCGTTCACTC | Tb927.10.5110_rev | GGGGACCACTTTGTACAAGAAAGCTGGGTTTAACACTCCAACTCCTGCAAAGAAA |
| 1_known | E2 | Tb927.11.10740_for | GGGGACAAGTTTGTACAAAAAAGCAGGCTATGGAAGTGGCATTCACCATTTCG | Tb927.11.10740_rev | GGGGACCACTTTGTACAAGAAAGCTGGGTTTACGCCTTTGGCCGTCG |
| 1_known | E3 | Tb927.8.5465_for | GGGGACAAGTTTGTACAAAAAAGCAGGCTATGGGTTGCTCAGGATCGAAGA | Tb927.8.5465_rev | GGGGACCACTTTGTACAAGAAAGCTGGGTTCAATTACCTTCATTTGCTCCGTTCT |
| 1_known | E4 | Tb927.8.5440_for | GGGGACAAGTTTGTACAAAAAAGCAGGCTATGGGTTGCTCAGGATCGAAGA | Tb927.8.5440_rev | GGGGACCACTTTGTACAAGAAAGCTGGGTTCAATTACCTTCATTTGCTCCGTTCT |
| 1_known | E5 | Tb927.4.430_for | GGGGACAAGTTTGTACAAAAAAGCAGGCTATGGCATCTGGAGGATCTGTTATTGG | Tb927.4.430_rev | GGGGACCACTTTGTACAAGAAAGCTGGGTTCATCGAATAATGGCTGTTTTCTCAAAGC |
| 1_known | E6 | Tb927.10.6380_for | GGGGACAAGTTTGTACAAAAAAGCAGGCTATGTCTCGTACTGACTTTTCATGTGC | Tb927.10.6380_rev | GGGGACCACTTTGTACAAGAAAGCTGGGTTCACACCATAAAAAAAGATGTGGCTAACA |
| 1_known | E7 | Tb927.11.16830_for | GGGGACAAGTTTGTACAAAAAAGCAGGCTATGTCGGACGACTTGGTAAAGATACT | Tb927.11.16830_rev | GGGGACCACTTTGTACAAGAAAGCTGGGTTCAACGCTTCAGCCGCGG |
| 1_known | E8 | Tb927.11.10000_for | GGGGACAAGTTTGTACAAAAAAGCAGGCTATGTTTTCCGGGTGGTTCAAAAGG | Tb927.11.10000_rev | GGGGACCACTTTGTACAAGAAAGCTGGGTTCAACTAACAGTCCCACGAAGCA |
| 1_known | E9 | Tb927.11.8140_for | GGGGACAAGTTTGTACAAAAAAGCAGGCTATGATGTACATTCGTACCTTCGGTG | Tb927.11.8140_rev | GGGGACCACTTTGTACAAGAAAGCTGGGTTCATCGCGCCAAATCATATCGC |
| 1_known | E10 | Tb927.9.15090_for | GGGGACAAGTTTGTACAAAAAAGCAGGCTATGGAACGGATAGCCTCCGG | Tb927.9.15090_rev | GGGGACCACTTTGTACAAGAAAGCTGGGTTTAGTCAATAAGTCGCTTCTTTCGGAA |
| 1_known | E11 | Tb927.8.4620_for | GGGGACAAGTTTGTACAAAAAAGCAGGCTATGAAAGAGGAACCCGCTTACAAG | Tb927.8.4620_rev | GGGGACCACTTTGTACAAGAAAGCTGGGTTCAGCACCTCCACTTCCTCTTT |
| 1_known | E12 | Tb927.11.500_for | GGGGACAAGTTTGTACAAAAAAGCAGGCTATGTCTCAGGTTCCACTGGCT | Tb927.11.500_rev | GGGGACCACTTTGTACAAGAAAGCTGGGTTCATTTACGGGCAGGCCGA |
| 1_known | E13 | Tb927.10.13320_for | GGGGACAAGTTTGTACAAAAAAGCAGGCTATGAGCAGTGAAAGCTACGCC | Tb927.10.13320_rev | GGGGACCACTTTGTACAAGAAAGCTGGGTCTACACTACCTGTCGGTACGTAACG |
| 1_known | E14 | Tb927.6.5140_for | GGGGACAAGTTTGTACAAAAAAGCAGGCTATGGCGCTGCAGTTGAATAAAGA | Tb927.6.5140_rev | GGGGACCACTTTGTACAAGAAAGCTGGGTTCACCACCTTGGTGTTGTACCT |
| 1_known | E15 | Tb927.9.11000_for | GGGGACAAGTTTGTACAAAAAAGCAGGCTATGGCTCCAAACCGACAGTTG | Tb927.9.11000_rev | GGGGACCACTTTGTACAAGAAAGCTGGGTCTAGCACGCACAATCGCTTTTC |
| 1_known | E16 | Tb927.4.3190_for | GGGGACAAGTTTGTACAAAAAAGCAGGCTATGCCGGCGCCGCCAGCG | Tb927.4.3190_rev | GGGGACCACTTTGTACAAGAAAGCTGGGTTCAATTCCTAAGCACTAAGAGCCCC |
| 1_known | E17 | Tb927.10.3860_for | GGGGACAAGTTTGTACAAAAAAGCAGGCTATGAACCGTCTATTCGGTCGGA | Tb927.10.3860_rev | GGGGACCACTTTGTACAAGAAAGCTGGGTTTACAGTCCCAACTGTGTTTCCAGT |
| 1_known | E18 | Tb927.10.650_for | GGGGACAAGTTTGTACAAAAAAGCAGGCTATGCCCCGTTATTACACCCCA | Tb927.10.650_rev | GGGGACCACTTTGTACAAGAAAGCTGGGTTTACGCCTCCGTGAGATCATCA |
| 1_known | E19 | Tb927.3.4650_for | GGGGACAAGTTTGTACAAAAAAGCAGGCTATGGGTCTCTGCAAAACACTTTTTGT | Tb927.3.4650_rev | GGGGACCACTTTGTACAAGAAAGCTGGGTTTATATCTTTCCTCGAAGGGCATTAGGT |

| | | | | | |
|---|---|---|---|---|---|
| 1_known | E20 | Tb927.1.3200_for | GGGGACAAGTTTGTACAAAAAAGCAGGCTATGGACGCATCCAGTGCCG | Tb927.1.3200_rev | GGGGACCACTTTGTACAAGAAAGCTGGGTTTAGCCTTCGTACTTCTTTTCTTCCATT |
| 1_known | E21 | Tb927.2.5120_for | GGGGACAAGTTTGTACAAAAAAGCAGGCTATGCTTATATCTGCCTCCTTTGTGTC | Tb927.2.5120_rev | GGGGACCACTTTGTACAAGAAAGCTGGGTCTACTTTTTGCACTTGAGGTTAGTGC |
| 1_known | E22 | Tb927.10.2530_for | GGGGACAAGTTTGTACAAAAAAGCAGGCTATGAAGGTAATTTTTCTTGGACCTCCAG | Tb927.10.2530_rev | GGGGACCACTTTGTACAAGAAAGCTGGGTTTAATTCCCATGTATCTTTGCACCGA |
| 1_known | E23 | Tb927.5.950_for | GGGGACAAGTTTGTACAAAAAAGCAGGCTATGAAACGCGTTGGGACTCTTG | Tb927.5.950_rev | GGGGACCACTTTGTACAAGAAAGCTGGGTTCATAGCTTCAGAGCAGATTTCAACTC |
| 1_known | E24 | Tb927.7.6440_for | GGGGACAAGTTTGTACAAAAAAGCAGGCTATGTCTAATGCCGAGGCCTACG | Tb927.7.6440_rev | GGGGACCACTTTGTACAAGAAAGCTGGGTTTATTTTTGCAAAAACTTTAACGTAGCAAA |
| 1_known | F1 | Tb927.6.2170_for | GGGGACAAGTTTGTACAAAAAAGCAGGCTATGCGTGCGCTTACGTTTCG | Tb927.6.2170_rev | GGGGACCACTTTGTACAAGAAAGCTGGGTCTAATCATCCGAAGCTACTCCAACTTG |
| 1_known | F2 | Tb927.8.5740_for | GGGGACAAGTTTGTACAAAAAAGCAGGCTATGTCTGGAACTTTACTCAAGGAAGAGC | Tb927.8.5740_rev | GGGGACCACTTTGTACAAGAAAGCTGGGTCTAATTCAATTTCATTGGCTCAAATGTGC |
| 1_known | F3 | Tb927.4.2670_for | GGGGACAAGTTTGTACAAAAAAGCAGGCTATGAAGCCTACAAAACCACTACTTTACA | Tb927.4.2670_rev | GGGGACCACTTTGTACAAGAAAGCTGGGTTCATTCAACTGCTTCCATGTGTTTCA |
| 1_known | F4 | Tb927.9.1720_for | GGGGACAAGTTTGTACAAAAAAGCAGGCTATGGCGGCTGAGCCCGCG | Tb927.9.1720_rev | GGGGACCACTTTGTACAAGAAAGCTGGGTTTAAAAGGAAGTATCACTGTTAAAGCAAAG |
| 1_known | F5 | Tb927.8.3790_for | GGGGACAAGTTTGTACAAAAAAGCAGGCTATGGATGCAGAAGAGAATTTCCTGAATG | Tb927.8.3790_rev | GGGGACCACTTTGTACAAGAAAGCTGGGTCTACGCACTGTTCGCAGCG |
| 1_known | F6 | Tb927.8.5330_for | GGGGACAAGTTTGTACAAAAAAGCAGGCTATGGAAGGTCCTTTCCTCAGCC | Tb927.8.5330_rev | GGGGACCACTTTGTACAAGAAAGCTGGGTCTATGAAACGCTACCGTTGGGG |
| 1_known | F7 | Tb927.11.13280_for | GGGGACAAGTTTGTACAAAAAAGCAGGCTATGCTCCGACGTATCATCAGCC | Tb927.11.13280_rev | GGGGACCACTTTGTACAAGAAAGCTGGGTCTACACAGATGTGCGAGCGAA |
| 1_known | F8 | Tb927.6.5050_for | GGGGACAAGTTTGTACAAAAAAGCAGGCTATGGCGGTTGAATCCGATGTG | Tb927.6.5050_rev | GGGGACCACTTTGTACAAGAAAGCTGGGTTTATACCATCACCGCCTTTTGGGC |
| 1_known | F9 | Tb927.10.7420_for | GGGGACAAGTTTGTACAAAAAAGCAGGCTATGAGCAAGAACGAGCGAGATAC | Tb927.10.7420_rev | GGGGACCACTTTGTACAAGAAAGCTGGGTCTACGAGCTGTCGTAATCGCTG |
| 1_known | F10 | Tb927.7.280_for | GGGGACAAGTTTGTACAAAAAAGCAGGCTATGCCCCTTCGGTTCACCT | Tb927.7.280_rev | GGGGACCACTTTGTACAAGAAAGCTGGGTCTACGCCGGTTCCTTCAGC |
| 1_known | F11 | Tb927.7.5580_for | GGGGACAAGTTTGTACAAAAAAGCAGGCTATGAAGCGTTTTAAGGCAAAGCTC | Tb927.7.5580_rev | GGGGACCACTTTGTACAAGAAAGCTGGGTTTACGGAACCTCCAACTTTTCCTTC |
| 1_known | F12 | Tb927.8.4180_for | GGGGACAAGTTTGTACAAAAAAGCAGGCTATGGTGCAAGATACAGGCAAGG | Tb927.8.4180_rev | GGGGACCACTTTGTACAAGAAAGCTGGGTTCAAACAACTGACAAATCCTTCCGT |
| 1_known | F13 | Tb927.4.3570_for | GGGGACAAGTTTGTACAAAAAAGCAGGCTATGTCGATCAAGGACGTCAATGTG | Tb927.4.3570_rev | GGGGACCACTTTGTACAAGAAAGCTGGGTCTAGATTTTATTCCAGGCCACAATGTCC |
| 1_known | F14 | Tb927.11.14290_for | GGGGACAAGTTTGTACAAAAAAGCAGGCTATGGTTCTTGTGCTTGTTGTTGGT | Tb927.11.14290_rev | GGGGACCACTTTGTACAAGAAAGCTGGGTTCATTCAGGAGGTGGAGGAGGA |
| 1_known | F16 | Tb927.10.9010_for | GGGGACAAGTTTGTACAAAAAAGCAGGCTATGATGCCTGCACCGAACG | Tb927.10.9010_rev | GGGGACCACTTTGTACAAGAAAGCTGGGTTCACCGTAATGCCAGCGC |
| 1_known | F17 | Tb927.10.8560_for | GGGGACAAGTTTGTACAAAAAAGCAGGCTATGTTGAAGAGAGGTGGGTTAAACG | Tb927.10.8560_rev | GGGGACCACTTTGTACAAGAAAGCTGGGTCTACATAACACTGCACTTTTTCTTCTTTGA |
| 1_known | F18 | Tb927.11.6030_for | GGGGACAAGTTTGTACAAAAAAGCAGGCTATGAGGAATATTAACCTCGTCGTTTTGG | Tb927.11.6030_rev | GGGGACCACTTTGTACAAGAAAGCTGGGTTCAGAGCATAGTGCATCCCGA |
| 1_known | F19 | Tb927.8.1990_for | GGGGACAAGTTTGTACAAAAAAGCAGGCTATGCTTCGCCGTTTCTCCATG | Tb927.8.1990_rev | GGGGACCACTTTGTACAAGAAAGCTGGGTCTATAGATTCTTCTCAAAGTATTCCTTCGC |
| 1_known | F20 | Tb927.7.2540_for | GGGGACAAGTTTGTACAAAAAAGCAGGCTATGGATAATGAATCTACCAATGAAGCAGC | Tb927.7.2540_rev | GGGGACCACTTTGTACAAGAAAGCTGGGTTCACTCCGCGCCTTCCGT |
| 1_known | F21 | Tb927.11.13930_for | GGGGACAAGTTTGTACAAAAAAGCAGGCTATGGTGCGGGACACGAGT | Tb927.11.13930_rev | GGGGACCACTTTGTACAAGAAAGCTGGGTTTACTCGTAATCGTCCCGCTGT |
| 1_known | F22 | Tb927.9.9750_for | GGGGACAAGTTTGTACAAAAAAGCAGGCTATGCCAGCGGACTCAGTTGT | Tb927.9.9750_rev | GGGGACCACTTTGTACAAGAAAGCTGGGTCTATGGGGCTGTGTCAGCAGG |
| 1_known | F23 | Tb927.10.12170_for | GGGGACAAGTTTGTACAAAAAAGCAGGCTATGCGATCAGCCACTGAATATACAG | Tb927.10.12170_rev | GGGGACCACTTTGTACAAGAAAGCTGGGTTCACTTGTTAGGGCTCTCCATGT |
| 1_known | F24 | Tb927.6.2420_for | GGGGACAAGTTTGTACAAAAAAGCAGGCTATGCGTCGTGCACTTGTATTCA | Tb927.6.2420_rev | GGGGACCACTTTGTACAAGAAAGCTGGGTTTACGAAACAAATTTGTTAATGCTGCTCA |
| 1_known | G1 | Tb927.1.1220_for | GGGGACAAGTTTGTACAAAAAAGCAGGCTATGCCGGCCACTGAGATGC | Tb927.1.1220_rev | GGGGACCACTTTGTACAAGAAAGCTGGGTTCATTCATCCAAATCAAACTCCTGCT |
| 1_known | G2 | Tb927.7.4850_for | GGGGACAAGTTTGTACAAAAAAGCAGGCTATGGTAGCAAGACTTAGGCAATATGAAC | Tb927.7.4850_rev | GGGGACCACTTTGTACAAGAAAGCTGGGTCTAAGCATGGTGAGGTGGGGG |
| 1_known | G3 | Tb927.10.14630_for | GGGGACAAGTTTGTACAAAAAAGCAGGCTATGCGTGGCTGCTACCTCC | Tb927.10.14630_rev | GGGGACCACTTTGTACAAGAAAGCTGGGTCTATTGCGGGGGGGATTTGTAGT |
| 1_known | G4 | Tb927.11.15230_for | GGGGACAAGTTTGTACAAAAAAGCAGGCTATGACCATGCGCGCGATG | Tb927.11.15230_rev | GGGGACCACTTTGTACAAGAAAGCTGGGTCTATACAACTCTTCTTTTGTTGAAAATACG |
| 1_known | G5 | Tb927.4.470_for | GGGGACAAGTTTGTACAAAAAAGCAGGCTATGGCACGGGGAGGAGGC | Tb927.4.470_rev | GGGGACCACTTTGTACAAGAAAGCTGGGTTCAAAAACCACGCCCGCG |
| 1_known | G6 | Tb927.10.2540_for | GGGGACAAGTTTGTACAAAAAAGCAGGCTATGCACGTGTTTCTCACAGGTC | Tb927.10.2540_rev | GGGGACCACTTTGTACAAGAAAGCTGGGTTCAACTTGACCACCAAAACAACTTGG |
| 1_known | G7 | Tb927.10.10610_for | GGGGACAAGTTTGTACAAAAAAGCAGGCTATGCTCTCCAAAGTATATTTACTTGCCTAT | Tb927.10.10610_rev | GGGGACCACTTTGTACAAGAAAGCTGGGTTCAGCTTGCCTTCTTCTTGTTGG |
| 1_known | G8 | Tb927.10.11610_for | GGGGACAAGTTTGTACAAAAAAGCAGGCTATGGTGAATCATTTAGTTGTATTGATCGGC | Tb927.10.11610_rev | GGGGACCACTTTGTACAAGAAAGCTGGGTCTACTTCTCCTCTGACATTCTTCCGT |
| 1_known | G9 | Tb927.2.3960_for | GGGGACAAGTTTGTACAAAAAAGCAGGCTATGAATAGGTCCTCCTCTACCAGTGT | Tb927.2.3960_rev | GGGGACCACTTTGTACAAGAAAGCTGGGTTTACGCCGTAATGTTGACACCAC |
| 1_known | G10 | Tb927.10.12960_for | GGGGACAAGTTTGTACAAAAAAGCAGGCTATGTCGGTGTCAGCGACACC | Tb927.10.12960_rev | GGGGACCACTTTGTACAAGAAAGCTGGGTTCAGCAGGCACACCCGCC |
| 1_known | G11 | Tb927.2.2130_for | GGGGACAAGTTTGTACAAAAAAGCAGGCTATGGAAACAAACTCAGCGCCA | Tb927.2.2130_rev | GGGGACCACTTTGTACAAGAAAGCTGGGTTTAGCAGCAACCGCCTTCCC |
| 1_known | G12 | Tb927.10.1830_for | GGGGACAAGTTTGTACAAAAAAGCAGGCTATGTCGTCTCTGCAAGATCCTTTC | Tb927.10.1830_rev | GGGGACCACTTTGTACAAGAAAGCTGGGTTCAACTAAAGACACAATAGAAGAGGAGCA |
| 1_known | G13 | Tb927.7.1790_for | GGGGACAAGTTTGTACAAAAAAGCAGGCTATGTCACAGTATGATGCGATTCTGAC | Tb927.7.1790_rev | GGGGACCACTTTGTACAAGAAAGCTGGGTTCACAGTCGTGACCTGGTAATGT |
| 1_known | G14 | Tb927.10.290_for | GGGGACAAGTTTGTACAAAAAAGCAGGCTATGTCTGAGTCTTCGTACGGTCTT | Tb927.10.290_rev | GGGGACCACTTTGTACAAGAAAGCTGGGTTTAGATTTGGTCCAGGTAATCCTTCAAC |
| 1_known | G15 | Tb927.7.3430_for | GGGGACAAGTTTGTACAAAAAAGCAGGCTATGCTACATCGTACGCGTTTAGC | Tb927.7.3430_rev | GGGGACCACTTTGTACAAGAAAGCTGGGTCTAATAACGAGGAGACAGAATGTCAAGAA |
| 1_known | G16 | Tb927.10.3710_for | GGGGACAAGTTTGTACAAAAAAGCAGGCTATGCCACCGAAACGCGCC | Tb927.10.3710_rev | GGGGACCACTTTGTACAAGAAAGCTGGGTTCAACTCACCATATGATCGCTTCCC |
| 1_known | G17 | Tb927.10.3120_for | GGGGACAAGTTTGTACAAAAAAGCAGGCTATGCTCCGTTCCCCGTGG | Tb927.10.3120_rev | GGGGACCACTTTGTACAAGAAAGCTGGGTCTACGCTAAATTCGACCGAAAAATTATGT |
| 1_known | G18 | Tb927.10.13950_for | GGGGACAAGTTTGTACAAAAAAGCAGGCTATGTCCGTTGTTAAAGTATCGCTTACG | Tb927.10.13950_rev | GGGGACCACTTTGTACAAGAAAGCTGGGTTTAAAACACCTCCGGGGGAAAGT |
| 1_known | G19 | Tb927.11.11030_for | GGGGACAAGTTTGTACAAAAAAGCAGGCTATGGGGGTGCCAACAATGGA | Tb927.11.11030_rev | GGGGACCACTTTGTACAAGAAAGCTGGGTCTAAAATGCAAACTGTTCGCACAACT |
| 1_known | G20 | Tb927.11.2260_for | GGGGACAAGTTTGTACAAAAAAGCAGGCTATGATGGCTGAATCAAGCGCG | Tb927.11.2260_rev | GGGGACCACTTTGTACAAGAAAGCTGGGTTTAGGCCTTGCTAGCGCCA |
| 1_known | G21 | Tb927.8.5470_for | GGGGACAAGTTTGTACAAAAAAGCAGGCTATGGGTTGCTCAGGATCGAAGA | Tb927.8.5470_rev | GGGGACCACTTTGTACAAGAAAGCTGGGTCTAGGCAGAACCCTCGGCC |
| 1_known | G22 | Tb927.10.1390_for | GGGGACAAGTTTGTACAAAAAAGCAGGCTATGCACTCGGGCCATCCT | Tb927.10.1390_rev | GGGGACCACTTTGTACAAGAAAGCTGGGTTTACAATTTGCTCGGGTACCGTTT |
| 1_known | G23 | Tb927.10.4700_for | GGGGACAAGTTTGTACAAAAAAGCAGGCTATGTCAAAGGAGCCGGGTCA | Tb927.10.4700_rev | GGGGACCACTTTGTACAAGAAAGCTGGGTTCATATAGCTGCGCCACCAAAC |
| 1_known | G24 | Tb927.10.7070_for | GGGGACAAGTTTGTACAAAAAAGCAGGCTATGCAGCAGCGTCTTGTTCTC | Tb927.10.7070_rev | GGGGACCACTTTGTACAAGAAAGCTGGGTTCATTTACGCTTCAGCAGACTTTCA |
| 1_known | H1 | Tb927.11.12860.2_for | GGGGACAAGTTTGTACAAAAAAGCAGGCTATGAGCAACGCAGCTCCAG | Tb927.11.12860.2_rev | GGGGACCACTTTGTACAAGAAAGCTGGGTCTAGTTCTTACGGCGCGTAACA |
| 1_known | H2 | Tb927.10.15690_for | GGGGACAAGTTTGTACAAAAAAGCAGGCTATGCAATTTGATGGACGTAGTGGT | Tb927.10.15690_rev | GGGGACCACTTTGTACAAGAAAGCTGGGTTCAGCGAAGATGAAGGTTCTGTG |
| 1_known | H3 | Tb927.7.1780_for | GGGGACAAGTTTGTACAAAAAAGCAGGCTATGTCACTTGTGGAGGTTTTGCC | Tb927.7.1780_rev | GGGGACCACTTTGTACAAGAAAGCTGGGTTTAAATCAATTTTTTTGCCTCGGCGC |
| 1_known | H4 | Tb927.10.8330_for | GGGGACAAGTTTGTACAAAAAAGCAGGCTATGGCCAGTTTACTTGCAGATGTT | Tb927.10.8330_rev | GGGGACCACTTTGTACAAGAAAGCTGGGTCTAGTTATAAGTCCTTACTTTTTCCATCCA |
| 1_known | H5 | Tb927.11.5510_for | GGGGACAAGTTTGTACAAAAAAGCAGGCTATGCCCACACGACCCCTG | Tb927.11.5510_rev | GGGGACCACTTTGTACAAGAAAGCTGGGTTCAAGCTTTGGTAGCATTACATAATGTCT |
| 1_known | H6 | Tb927.11.250_for | GGGGACAAGTTTGTACAAAAAAGCAGGCTATGACCAAAACGTGGCTACCG | Tb927.11.250_rev | GGGGACCACTTTGTACAAGAAAGCTGGGTTCACTCGCGAGACTCACACA |
| 1_known | H7 | Tb927.8.4470_for | GGGGACAAGTTTGTACAAAAAAGCAGGCTATGTTCACTTACTCTACGTTTTTACGGAG | Tb927.8.4470_rev | GGGGACCACTTTGTACAAGAAAGCTGGGTTCATCGGCCCATCACAACGT |
| 1_known | H8 | Tb927.1.1680_for | GGGGACAAGTTTGTACAAAAAAGCAGGCTATGTCGTCAAGCAAAGGTGGTG | Tb927.1.1680_rev | GGGGACCACTTTGTACAAGAAAGCTGGGTCTAGTCACTTTCGTCAAAGAATGCATTAAC |
| 1_known | H9 | Tb927.5.1080_for | GGGGACAAGTTTGTACAAAAAAGCAGGCTATGCTCGACACGAGTGGCA | Tb927.5.1080_rev | GGGGACCACTTTGTACAAGAAAGCTGGGTTTAATTTTTTTTTACTTTCTTCTTTTTTAA |
| 1_known | H10 | Tb927.3.780_for | GGGGACAAGTTTGTACAAAAAAGCAGGCTATGTCTGGCTCTGGTCATGACC | Tb927.3.780_rev | GGGGACCACTTTGTACAAGAAAGCTGGGTCTATGATGCGGGGGTTGGATG |
| 1_known | H11 | Tb927.5.1370_for | GGGGACAAGTTTGTACAAAAAAGCAGGCTATGATAGCCATTCTGTTTGGGTTCAC | Tb927.5.1370_rev | GGGGACCACTTTGTACAAGAAAGCTGGGTTCAAAACCGAAGGAACGCCG |
| 1_known | H12 | Tb927.8.2410_for | GGGGACAAGTTTGTACAAAAAAGCAGGCTATGGTTGTGCGGGAGGTGG | Tb927.8.2410_rev | GGGGACCACTTTGTACAAGAAAGCTGGGTTTAGGTTCGAGCAAACGTGATGC |
| 1_known | H13 | Tb927.10.15500_for | GGGGACAAGTTTGTACAAAAAAGCAGGCTATGCATGGAGTTGAGGTATCAGCG | Tb927.10.15500_rev | GGGGACCACTTTGTACAAGAAAGCTGGGTCTACCATATCAGATAGATAAACAGCAACGT |
| 1_known | H14 | Tb927.5.3350_for | GGGGACAAGTTTGTACAAAAAAGCAGGCTATGAGGTCTGTCATGATGCGTTG | Tb927.5.3350_rev | GGGGACCACTTTGTACAAGAAAGCTGGGTTTACTTCATAGCCTGTTCATACGTCCT |
| 1_known | H15 | Tb927.10.790_for | GGGGACAAGTTTGTACAAAAAAGCAGGCTATGCAGGGAGGAACAAAAATATATGGC | Tb927.10.790_rev | GGGGACCACTTTGTACAAGAAAGCTGGGTTTACTTCTTTTCCTCTTTTTTACCACCTAC |
| 1_known | H16 | Tb927.11.4530_for | GGGGACAAGTTTGTACAAAAAAGCAGGCTATGCAGGTGCGTCGCGCC | Tb927.11.4530_rev | GGGGACCACTTTGTACAAGAAAGCTGGGTTTACTTTTTACCTTTTCCGTGCTTACCG |
| 1_known | H17 | Tb927.3.2930_for | GGGGACAAGTTTGTACAAAAAAGCAGGCTATGTTCTACCCCAACAGCCCG | Tb927.3.2930_rev | GGGGACCACTTTGTACAAGAAAGCTGGGTTCAACCAGCGGCACCGCG |
| 1_known | H18 | Tb927.6.3320_for | GGGGACAAGTTTGTACAAAAAAGCAGGCTATGAGCACCGAAGACATTCTGATTG | Tb927.6.3320_rev | GGGGACCACTTTGTACAAGAAAGCTGGGTTTACCAGGCCACAAAGGGGC |
| 1_known | H19 | Tb927.10.13830_for | GGGGACAAGTTTGTACAAAAAAGCAGGCTATGTGGCGTAGTCAGTGCTGT | Tb927.10.13830_rev | GGGGACCACTTTGTACAAGAAAGCTGGGTTCATGCAACAAGTGAGTTGTTCATGA |
| 1_known | H20 | Tb927.6.3040_for | GGGGACAAGTTTGTACAAAAAAGCAGGCTATGAGTAGCGACAGTTCAGACTCA | Tb927.6.3040_rev | GGGGACCACTTTGTACAAGAAAGCTGGGTCTACATCACTGCGCATTTACCCT |
| 1_known | H21 | Tb927.4.3320_for | GGGGACAAGTTTGTACAAAAAAGCAGGCTATGGCTCTACACGACAACGC | Tb927.4.3320_rev | GGGGACCACTTTGTACAAGAAAGCTGGGTTCATTCGCATGTGCCGAAGTATC |
| 1_known | H22 | Tb927.2.5210_for | GGGGACAAGTTTGTACAAAAAAGCAGGCTATGCCATGGAACCTTCAAGGC | Tb927.2.5210_rev | GGGGACCACTTTGTACAAGAAAGCTGGGTTCACCACATTCCCCCATCTACAT |
| 1_known | H23 | Tb927.6.710_for | GGGGACAAGTTTGTACAAAAAAGCAGGCTATGCTACTCGTCGGTCTCACC | Tb927.6.710_rev | GGGGACCACTTTGTACAAGAAAGCTGGGTTTAGAATACGATGCGGAGGCAAC |
| 1_known | H24 | Tb927.10.6650_for | GGGGACAAGTTTGTACAAAAAAGCAGGCTATGCCGAAAATCCGCCCAG | Tb927.10.6650_rev | GGGGACCACTTTGTACAAGAAAGCTGGGTCTATTTACCCGATTTCACCTCGTAGTC |
| 1_known | I1 | Tb927.10.15870_for | GGGGACAAGTTTGTACAAAAAAGCAGGCTATGAACTACGCCGACGGTAACT | Tb927.10.15870_rev | GGGGACCACTTTGTACAAGAAAGCTGGGTTCAAGATCTCCTTGGGACAGCC |
| 1_known | I2 | Tb927.10.7850_for | GGGGACAAGTTTGTACAAAAAAGCAGGCTATGGCATGGACATGTTCCGGT | Tb927.10.7850_rev | GGGGACCACTTTGTACAAGAAAGCTGGGTCTAAATACCTCCCCTTTGGTGCTTC |
| 1_known | I4 | Tb927.11.3800_for | GGGGACAAGTTTGTACAAAAAAGCAGGCTATGCCCTGTAATGAACAGACCTCT | Tb927.11.3800_rev | GGGGACCACTTTGTACAAGAAAGCTGGGTTTACTCGTACTTCATATGCGCCAAG |
| 1_known | I5 | Tb927.8.4070_for | GGGGACAAGTTTGTACAAAAAAGCAGGCTATGACCATCTCGTCTGCGGA | Tb927.8.4070_rev | GGGGACCACTTTGTACAAGAAAGCTGGGTCTAAGGCAACGCAGACCCG |
| 1_known | I6 | Tb927.8.3380_for | GGGGACAAGTTTGTACAAAAAAGCAGGCTATGCTTCGTAAAGTCACTAGCCCA | Tb927.8.3380_rev | GGGGACCACTTTGTACAAGAAAGCTGGGTTCATGCGCCTAGTTTCCGC |
| 1_known | I7 | Tb927.11.10870_for | GGGGACAAGTTTGTACAAAAAAGCAGGCTATGCGTTACCCGGGGCGA | Tb927.11.10870_rev | GGGGACCACTTTGTACAAGAAAGCTGGGTTCACAACTCTTTTTCTCGAACTAAAATACG |
| 1_known | I10 | Tb927.10.11270_for | GGGGACAAGTTTGTACAAAAAAGCAGGCTATGGTGGATCCTTCTGTGCTGC | Tb927.10.11270_rev | GGGGACCACTTTGTACAAGAAAGCTGGGTTCAGTTGTGGACTTGCACCCC |
| 1_known | I11 | Tb927.9.11580_for | GGGGACAAGTTTGTACAAAAAAGCAGGCTATGTCTGCTCAAGCCCATACATACC | Tb927.9.11580_rev | GGGGACCACTTTGTACAAGAAAGCTGGGTTCAATAGAATTCACACACAGACGCA |
| 1_known | I12 | Tb927.4.2520_for | GGGGACAAGTTTGTACAAAAAAGCAGGCTATGAGGCGGCCCAATGGT | Tb927.4.2520_rev | GGGGACCACTTTGTACAAGAAAGCTGGGTTCATGCCGCGGGACCCTT |
| 1_known | I13 | Tb927.9.9120_for | GGGGACAAGTTTGTACAAAAAAGCAGGCTATGACTTCCACAACACAACCGAA | Tb927.9.9120_rev | GGGGACCACTTTGTACAAGAAAGCTGGGTTTATGGCACCCGCACGTG |
| 1_known | I14 | Tb927.2.4930_for | GGGGACAAGTTTGTACAAAAAAGCAGGCTATGTCTCACATTCTTTTGTTGGGTGA | Tb927.2.4930_rev | GGGGACCACTTTGTACAAGAAAGCTGGGTTCATTTCATCAACGTCATGAAGTCTGG |
| 1_known | I15 | Tb927.9.12690_for | GGGGACAAGTTTGTACAAAAAAGCAGGCTATGCATTGGGGGTATCGGCA | Tb927.9.12690_rev | GGGGACCACTTTGTACAAGAAAGCTGGGTCTAACGAATGAAAGGAATCCCCACC |
| 1_known | I16 | Tb927.9.11130_for | GGGGACAAGTTTGTACAAAAAAGCAGGCTATGGCACAAGGAGGGTTCACG | Tb927.9.11130_rev | GGGGACCACTTTGTACAAGAAAGCTGGGTCTAACTGCACCATGTCTGACAAAGG |
| 1_known | I17 | Tb927.10.6440_for | GGGGACAAGTTTGTACAAAAAAGCAGGCTATGAAAAGAGTTCTTTTACTCTTTGACGTT | Tb927.10.6440_rev | GGGGACCACTTTGTACAAGAAAGCTGGGTTTACTTCATGGCAATTATTTTTTCCACCTC |
| 1_known | I18 | Tb927.4.650_for | GGGGACAAGTTTGTACAAAAAAGCAGGCTATGCGCCGCAGTGCTGTA | Tb927.4.650_rev | GGGGACCACTTTGTACAAGAAAGCTGGGTCTATTGTCCATCAGCATCTTCGAGC |
| 1_known | I19 | Tb927.4.4080_for | GGGGACAAGTTTGTACAAAAAAGCAGGCTATGACCACAGTGGTGATGAGCG | Tb927.4.4080_rev | GGGGACCACTTTGTACAAGAAAGCTGGGTTCATGAACCATGGACGCGGG |
| 1_known | I20 | Tb927.7.3440_for | GGGGACAAGTTTGTACAAAAAAGCAGGCTATGCTCTGCCCTCCAGATGTT | Tb927.7.3440_rev | GGGGACCACTTTGTACAAGAAAGCTGGGTTTAGCACCCGCAACCTGCC |
| 1_known | I21 | Tb927.10.230_for | GGGGACAAGTTTGTACAAAAAAGCAGGCTATGTTCTCCTCCAAAACCGAGTATGA | Tb927.10.230_rev | GGGGACCACTTTGTACAAGAAAGCTGGGTTTAATTTTCCTCGGCCTGCCG |
| 1_known | I22 | Tb927.11.9810_for | GGGGACAAGTTTGTACAAAAAAGCAGGCTATGAGGGACCGCTATATCAGAGTTG | Tb927.11.9810_rev | GGGGACCACTTTGTACAAGAAAGCTGGGTTCAGAGAGTTGCTATTTTCAGACTCGT |
| 1_known | I23 | Tb927.11.850_for | GGGGACAAGTTTGTACAAAAAAGCAGGCTATGAATACGCACGCGGCC | Tb927.11.850_rev | GGGGACCACTTTGTACAAGAAAGCTGGGTCTATCCCACCATACTACGCTTGC |
| 1_known | I24 | Tb927.10.12210_for | GGGGACAAGTTTGTACAAAAAAGCAGGCTATGGTTCACCGTCAAGATGGAGC | Tb927.10.12210_rev | GGGGACCACTTTGTACAAGAAAGCTGGGTCTAACTCGTTGTGGATCCCAGATAAC |
| 1_known | J1 | Tb927.3.740_for | GGGGACAAGTTTGTACAAAAAAGCAGGCTATGATGCAACGGGATACCGC | Tb927.3.740_rev | GGGGACCACTTTGTACAAGAAAGCTGGGTTCAATGTAACCCAGTTTGTAGAATGGC |
| 1_known | J2 | Tb927.5.288b_for | GGGGACAAGTTTGTACAAAAAAGCAGGCTATGTACAGGTTCGATCATCAAGACAAG | Tb927.5.288b_rev | GGGGACCACTTTGTACAAGAAAGCTGGGTCTATAAGCGAGACTTTAAGTAGTGCTCAA |
| 1_known | J3 | Tb927.11.10060_for | GGGGACAAGTTTGTACAAAAAAGCAGGCTATGTCAGCCTTTTTAACACCCAAGA | Tb927.11.10060_rev | GGGGACCACTTTGTACAAGAAAGCTGGGTTTAAAAAAGGTACTCCGCCACAAGC |
| 1_known | J4 | Tb927.6.4930_for | GGGGACAAGTTTGTACAAAAAAGCAGGCTATGCTGGTTTCGTACTTGCGT | Tb927.6.4930_rev | GGGGACCACTTTGTACAAGAAAGCTGGGTTCACTGAAGACCGAAAGGCATG |
| 1_known | J5 | Tb927.10.4180_for | GGGGACAAGTTTGTACAAAAAAGCAGGCTATGTCTACTGTCCCTCGCCAAA | Tb927.10.4180_rev | GGGGACCACTTTGTACAAGAAAGCTGGGTTTAGCATTGCAATGACGCTAAGTCC |
| 1_known | J6 | Tb927.10.3200_for | GGGGACAAGTTTGTACAAAAAAGCAGGCTATGTATCAAGACCGTTGCATATTTTTCTCC | Tb927.10.3200_rev | GGGGACCACTTTGTACAAGAAAGCTGGGTTTATTTTAAGGGGCATTCGCGAGAG |
| 1_known | J7 | Tb927.11.14200_for | GGGGACAAGTTTGTACAAAAAAGCAGGCTATGTCTGCAATCCCTTACTTGCAA | Tb927.11.14200_rev | GGGGACCACTTTGTACAAGAAAGCTGGGTCTACATCAATGCCTCCATTAGAGTTGC |
| 1_known | J8 | Tb927.11.7020_for | GGGGACAAGTTTGTACAAAAAAGCAGGCTATGAGCTACGATCGCGCAATT | Tb927.11.7020_rev | GGGGACCACTTTGTACAAGAAAGCTGGGTTCAATCCTCTTCGGCAAGGCG |
| 1_known | J9 | Tb927.11.13240_for | GGGGACAAGTTTGTACAAAAAAGCAGGCTATGAGCCTTTTCCACAGTTACGC | Tb927.11.13240_rev | GGGGACCACTTTGTACAAGAAAGCTGGGTTCAAAACCACACTACCGGGCC |
| 1_known | J10 | Tb927.10.5300_for | GGGGACAAGTTTGTACAAAAAAGCAGGCTATGACACTGCGTACGCGC | Tb927.10.5300_rev | GGGGACCACTTTGTACAAGAAAGCTGGGTTTATGCCAGTTCATCCACCAACG |
| 1_known | J11 | Tb927.11.13580_for | GGGGACAAGTTTGTACAAAAAAGCAGGCTATGCAGAGTAGGGTGGTTTCGT | Tb927.11.13580_rev | GGGGACCACTTTGTACAAGAAAGCTGGGTTCATGAATCGTGTAATCCGCCAC |
| 1_known | J12 | Tb927.11.14840_for | GGGGACAAGTTTGTACAAAAAAGCAGGCTATGCAGTGGAAGAAAGATGACTTATTCG | Tb927.11.14840_rev | GGGGACCACTTTGTACAAGAAAGCTGGGTCTAGAAATGCAGAGGTAAACTTTCGGT |
| 1_known | J13 | Tb927.6.2330_for | GGGGACAAGTTTGTACAAAAAAGCAGGCTATGCCCGCACGTGGCTGT | Tb927.6.2330_rev | GGGGACCACTTTGTACAAGAAAGCTGGGTTTAGCGTAGTGCGTACATCTTCAC |
| 1_known | J14 | Tb927.9.9670_for | GGGGACAAGTTTGTACAAAAAAGCAGGCTATGAGCCGAGCTGGGTTTGA | Tb927.9.9670_rev | GGGGACCACTTTGTACAAGAAAGCTGGGTCTAATCAGCCTCCGCAAGTAGAGT |
| 1_known | J15 | Tb927.11.3340_for | GGGGACAAGTTTGTACAAAAAAGCAGGCTATGTCCAAGTTATTTATATCTAAACTTCCA | Tb927.11.3340_rev | GGGGACCACTTTGTACAAGAAAGCTGGGTTTACTCCAGCGCAGAAATATACGC |
| 1_known | J16 | Tb927.4.4330_for | GGGGACAAGTTTGTACAAAAAAGCAGGCTATGAAGGCGCAACGCTACG | Tb927.4.4330_rev | GGGGACCACTTTGTACAAGAAAGCTGGGTTCATAGCTTTGCCTCTCTCCTCC |
| 1_known | J17 | Tb927.11.5520_for | GGGGACAAGTTTGTACAAAAAAGCAGGCTATGTCCAAGCCACAACCCATCG | Tb927.11.5520_rev | GGGGACCACTTTGTACAAGAAAGCTGGGTTCACTGAGTGGCTTTGATGATGTC |
| 1_known | J18 | Tb927.7.2180_for | GGGGACAAGTTTGTACAAAAAAGCAGGCTATGTCACAAGTGCCAATTACCGC | Tb927.7.2180_rev | GGGGACCACTTTGTACAAGAAAGCTGGGTTTATTCAACGCTAGACAGCATTTGCT |
| 1_known | J19 | Tb927.9.15060_for | GGGGACAAGTTTGTACAAAAAAGCAGGCTATGCCAGTGTACGATGTGGCC | Tb927.9.15060_rev | GGGGACCACTTTGTACAAGAAAGCTGGGTTTAGCGCCTCCGTCGTGC |
| 1_known | J20 | Tb927.10.16070_for | GGGGACAAGTTTGTACAAAAAAGCAGGCTATGCAGACCTTACTGAGACCCC | Tb927.10.16070_rev | GGGGACCACTTTGTACAAGAAAGCTGGGTTCACAATTGATACAAGGGTGGGACA |
| 1_known | J21 | Tb927.7.2110_for | GGGGACAAGTTTGTACAAAAAAGCAGGCTATGACCAGGCCACGACTTACT | Tb927.7.2110_rev | GGGGACCACTTTGTACAAGAAAGCTGGGTTTAACAGCATAATAAATCATCGAGAGAGGT |
| 1_known | J22 | Tb927.2.3720_for | GGGGACAAGTTTGTACAAAAAAGCAGGCTATGCCCGTTTCACCGGCT | Tb927.2.3720_rev | GGGGACCACTTTGTACAAGAAAGCTGGGTTCAAATACGTCTGAGTGCTGCC |
| 1_known | J23 | Tb927.1.3280_for | GGGGACAAGTTTGTACAAAAAAGCAGGCTATGGTGCTACAGCATCTTGAACTTC | Tb927.1.3280_rev | GGGGACCACTTTGTACAAGAAAGCTGGGTCTAATTGTGTGAAATTATCCCCTCATCGA |
| 1_known | J24 | Tb927.6.2940_for | GGGGACAAGTTTGTACAAAAAAGCAGGCTATGGAAAAAGAAAAAAAAACAGTGAATATT | Tb927.6.2940_rev | GGGGACCACTTTGTACAAGAAAGCTGGGTTTATTGCGGTTCATCCGCGC |
| 1_known | K1 | Tb927.11.12080_for | GGGGACAAGTTTGTACAAAAAAGCAGGCTATGCATGGTGCTTTGGTGTTTGG | Tb927.11.12080_rev | GGGGACCACTTTGTACAAGAAAGCTGGGTTCATGCGACCCCCAATTCCT |
| 1_known | K2 | Tb927.5.4530_for | GGGGACAAGTTTGTACAAAAAAGCAGGCTATGATGGACTGCAGAGATGCTTCG | Tb927.5.4530_rev | GGGGACCACTTTGTACAAGAAAGCTGGGTTCATTGTGGTTTTTTTTCTATAGCACAACC |
| 1_known | K3 | Tb927.8.8040_for | GGGGACAAGTTTGTACAAAAAAGCAGGCTATGAAAGTGCAGGGGTACGCC | Tb927.8.8040_rev | GGGGACCACTTTGTACAAGAAAGCTGGGTTCAGGCCGCCGCCGACGC |
| 1_known | K4 | Tb927.10.7450_for | GGGGACAAGTTTGTACAAAAAAGCAGGCTATGCCTGTGATTAAGGAGTACATCAATC | Tb927.10.7450_rev | GGGGACCACTTTGTACAAGAAAGCTGGGTTTACTCGATGTTACCTAAATTCGCTTGC |
| 1_known | K5 | Tb927.11.6870_for | GGGGACAAGTTTGTACAAAAAAGCAGGCTATGGCGGGCTTTCAAATACCTG | Tb927.11.6870_rev | GGGGACCACTTTGTACAAGAAAGCTGGGTTCACTCCAATTCCTCCATCGCA |
| 1_known | K6 | Tb927.3.2190_for | GGGGACAAGTTTGTACAAAAAAGCAGGCTATGATGGAAGCGACGGAACG | Tb927.3.2190_rev | GGGGACCACTTTGTACAAGAAAGCTGGGTTCAACCCTCAGATGCGGTGA |
| 1_known | K7 | Tb927.8.730_for | GGGGACAAGTTTGTACAAAAAAGCAGGCTATGGCGCCCATGTTTCATGG | Tb927.8.730_rev | GGGGACCACTTTGTACAAGAAAGCTGGGTTTAGAACCGCCCCCGCTT |
| 1_known | K8 | Tb927.10.6640_for | GGGGACAAGTTTGTACAAAAAAGCAGGCTATGAATACTATGCAGGGAATTATGAGAGAT | Tb927.10.6640_rev | GGGGACCACTTTGTACAAGAAAGCTGGGTTCACACCAACTTCTTCTCCCGC |
| 1_known | K9 | Tb927.9.15620_for | GGGGACAAGTTTGTACAAAAAAGCAGGCTATGAGCATCACTTTTCATAACTTATGGCT | Tb927.9.15620_rev | GGGGACCACTTTGTACAAGAAAGCTGGGTTTACAGTACAGAACCGAGAAACAATCC |

| | | | | | |
|---|---|---|---|---|---|
| 1_known | K10 | Tb927.9.11310_for | GGGGACAAGTTTGTACAAAAAAGCAGGCTATGACAGGGTTCTCATTTGAAAACGT | Tb927.9.11310_rev | GGGGACCACTTTGTACAAGAAAGCTGGGTTCACTCCAATTCGGTGATTGTCAC |
| 1_known | K11 | Tb927.8.2500_for | GGGGACAAGTTTGTACAAAAAAGCAGGCTATGCACTTTCCCAGTATCGTGTTT | Tb927.8.2500_rev | GGGGACCACTTTGTACAAGAAAGCTGGGTCTACGGCTTGCGTCGCCG |
| 1_known | K13 | Tb927.5.1700_for | GGGGACAAGTTTGTACAAAAAAGCAGGCTATGGAAGGAAGTGGAAGTAACTTCTTTAC | Tb927.5.1700_rev | GGGGACCACTTTGTACAAGAAAGCTGGGTTTAAATGCCAAACTTACAATCATGTGTTTG |
| 1_known | K14 | Tb927.9.15560_for | GGGGACAAGTTTGTACAAAAAAGCAGGCTATGAGCATCACTTTTCATAACTTATGGCT | Tb927.9.15560_rev | GGGGACCACTTTGTACAAGAAAGCTGGGTTTACAGTACAGAACTGAGAAACAATCCAG |
| 1_known | K15 | Tb927.10.15130_for | GGGGACAAGTTTGTACAAAAAAGCAGGCTATGGCGTCAAAGGGTTGTTTCA | Tb927.10.15130_rev | GGGGACCACTTTGTACAAGAAAGCTGGGTTTAGAACAGGTAGAGGTAGAAGAAGAGAGC |
| 1_known | K16 | Tb927.11.9930_for | GGGGACAAGTTTGTACAAAAAAGCAGGCTATGACCGTCGCGCGCGAC | Tb927.11.9930_rev | GGGGACCACTTTGTACAAGAAAGCTGGGTCTATGCGACATCCCCAGGTTTC |
| 1_known | K17 | Tb927.2.2180_for | GGGGACAAGTTTGTACAAAAAAGCAGGCTATGGCAACAATATCGAGGCGC | Tb927.2.2180_rev | GGGGACCACTTTGTACAAGAAAGCTGGGTTCATTCTAGGGCCATGCTGCA |
| 1_known | K18 | Tb927.10.4450_for | GGGGACAAGTTTGTACAAAAAAGCAGGCTATGTCTGTAGCTGACTTAAAGGCAAAG | Tb927.10.4450_rev | GGGGACCACTTTGTACAAGAAAGCTGGGTTTAACCCCTAACAGCCCGTTGG |
| 1_known | K19 | Tb927.5.1280_for | GGGGACAAGTTTGTACAAAAAAGCAGGCTATGTCACGCTACGATATCGGCA | Tb927.5.1280_rev | GGGGACCACTTTGTACAAGAAAGCTGGGTCTACTTTGCTGCTCCATCCTTTGG |
| 1_known | K20 | Tb927.7.6660_for | GGGGACAAGTTTGTACAAAAAAGCAGGCTATGTGGCGCGTGGCATTAC | Tb927.7.6660_rev | GGGGACCACTTTGTACAAGAAAGCTGGGTCTAATAACTTAACTCACGTTTGAGGAAGGA |
| 1_known | K21 | Tb927.7.4790_for | GGGGACAAGTTTGTACAAAAAAGCAGGCTATGATTGAAGACTTCAGTGAGCACC | Tb927.7.4790_rev | GGGGACCACTTTGTACAAGAAAGCTGGGTTTAGTCTTTGCGGAGCTCAAACAT |
| 1_known | K22 | Tb927.8.1890_for | GGGGACAAGTTTGTACAAAAAAGCAGGCTATGGCAGGTAAGAAAGCTCACCC | Tb927.8.1890_rev | GGGGACCACTTTGTACAAGAAAGCTGGGTTTACCATGTCTTCTTCCAGTAGCGG |
| 1_known | K23 | Tb927.3.5280_for | GGGGACAAGTTTGTACAAAAAAGCAGGCTATGACCACGTTCAAGCGGGA | Tb927.3.5280_rev | GGGGACCACTTTGTACAAGAAAGCTGGGTCTAACGTACGCGAGCGTAGG |
| 1_known | K24 | Tb927.10.4610_for | GGGGACAAGTTTGTACAAAAAAGCAGGCTATGACGGTGAAGTATTCCATTATTGTTCC | Tb927.10.4610_rev | GGGGACCACTTTGTACAAGAAAGCTGGGTCTAAAGAAGAATGTAAAAACAGTAAAAAAT |
| 1_known | L1 | Tb927.11.12090_for | GGGGACAAGTTTGTACAAAAAAGCAGGCTATGGGCAAGCTATTTAAACCACCAT | Tb927.11.12090_rev | GGGGACCACTTTGTACAAGAAAGCTGGGTCTATATACGCTTCCGTTTTGGGGC |
| 1_known | L2 | Tb927.5.860_for | GGGGACAAGTTTGTACAAAAAAGCAGGCTATGCGCAACCGATTTGTTCGC | Tb927.5.860_rev | GGGGACCACTTTGTACAAGAAAGCTGGGTTTATGCGGTCCTCGTTCCTGC |
| 1_known | L3 | Tb927.11.290_for | GGGGACAAGTTTGTACAAAAAAGCAGGCTATGAGTTCGCCTAATTGTTGTCCTG | Tb927.11.290_rev | GGGGACCACTTTGTACAAGAAAGCTGGGTTTACGACCGGGTCAAACACGA |
| 1_known | L4 | Tb927.3.1760_for | GGGGACAAGTTTGTACAAAAAAGCAGGCTATGGTGGTGGTGGGGATGAA | Tb927.3.1760_rev | GGGGACCACTTTGTACAAGAAAGCTGGGTTTAAACACCCTTTGGGACCGC |
| 1_known | L5 | Tb927.2.5660_for | GGGGACAAGTTTGTACAAAAAAGCAGGCTATGGCGGTACTGAGCGAAGA | Tb927.2.5660_rev | GGGGACCACTTTGTACAAGAAAGCTGGGTTTAGCGAAAGCGGTGCTTCTC |
| 1_known | L6 | Tb927.9.15610_for | GGGGACAAGTTTGTACAAAAAAGCAGGCTATGAGCATCACTTTTCATAACTTATGGCT | Tb927.9.15610_rev | GGGGACCACTTTGTACAAGAAAGCTGGGTTTACAGTACAGAACCGAGAAACAGTCC |
| 1_known | L7 | Tb927.10.2350_for | GGGGACAAGTTTGTACAAAAAAGCAGGCTATGCGTAGGTCGCTTGCG | Tb927.10.2350_rev | GGGGACCACTTTGTACAAGAAAGCTGGGTTTACTTGCCAACCGCTGCG |
| 1_known | L8 | Tb927.7.3470_for | GGGGACAAGTTTGTACAAAAAAGCAGGCTATGCGGCGTGTAGTTGCTTC | Tb927.7.3470_rev | GGGGACCACTTTGTACAAGAAAGCTGGGTTCAAGTAGCGAACTGTTTAAGTAAGCG |
| 1_known | L9 | Tb927.11.14520_for | GGGGACAAGTTTGTACAAAAAAGCAGGCTATGTTACGGCATTTGTTCCGATGT | Tb927.11.14520_rev | GGGGACCACTTTGTACAAGAAAGCTGGGTTTAGTGGCTCAGGCGAGCG |
| 1_known | L10 | Tb927.6.1400_for | GGGGACAAGTTTGTACAAAAAAGCAGGCTATGTCAACGACAAAACTAGAAGGTGT | Tb927.6.1400_rev | GGGGACCACTTTGTACAAGAAAGCTGGGTCTATTCCTCTCCTGCACAACTCAC |
| 1_known | L12 | Tb927.4.1220_for | GGGGACAAGTTTGTACAAAAAAGCAGGCTATGGTGGTACCGGTGATGAATATGA | Tb927.4.1220_rev | GGGGACCACTTTGTACAAGAAAGCTGGGTTCACCCATCAGCTACTTTTACCTCG |
| 1_known | L14 | Tb927.10.11930_for | GGGGACAAGTTTGTACAAAAAAGCAGGCTATGTTATCAGGAAAGGTGGCGC | Tb927.10.11930_rev | GGGGACCACTTTGTACAAGAAAGCTGGGTCTACTGTGCTGTCCAGCCACC |
| 1_known | L15 | Tb927.3.5570_for | GGGGACAAGTTTGTACAAAAAAGCAGGCTATGGACGATCCAAGTTGGCAGC | Tb927.3.5570_rev | GGGGACCACTTTGTACAAGAAAGCTGGGTTCATACTTTATGGTACGCAACGAGC |
| 1_known | L17 | Tb927.5.3970_for | GGGGACAAGTTTGTACAAAAAAGCAGGCTATGTCCACCGGCCAACTTTC | Tb927.5.3970_rev | GGGGACCACTTTGTACAAGAAAGCTGGGTTTACACTGAAACAAACCTCGAGTCC |
| 1_known | L18 | Tb927.10.830_for | GGGGACAAGTTTGTACAAAAAAGCAGGCTATGGCGGACCTTTCCGATAATAAGT | Tb927.10.830_rev | GGGGACCACTTTGTACAAGAAAGCTGGGTTCAACGCAACAGCCGGCT |
| 1_known | L20 | Tb927.10.5290_for | GGGGACAAGTTTGTACAAAAAAGCAGGCTATGCTACGTGGCACACGGC | Tb927.10.5290_rev | GGGGACCACTTTGTACAAGAAAGCTGGGTTCACTCATCCCCATCGGGC |
| 1_known | L21 | Tb927.9.15630_for | GGGGACAAGTTTGTACAAAAAAGCAGGCTATGAGCATCACTTTTCATAATTTATGGCTA | Tb927.9.15630_rev | GGGGACCACTTTGTACAAGAAAGCTGGGTTTACAGTACAGAACCGAGAAACAGTCC |
| 1_known | L22 | Tb927.3.2840_for | GGGGACAAGTTTGTACAAAAAAGCAGGCTATGCGCCCAACATCAATTATGTTTAAA | Tb927.3.2840_rev | GGGGACCACTTTGTACAAGAAAGCTGGGTTCAGTGAAGCCAGTAACCGGG |
| 1_known | L23 | Tb927.6.4960_for | GGGGACAAGTTTGTACAAAAAAGCAGGCTATGTCATTTACGGCTAACTCTGGC | Tb927.6.4960_rev | GGGGACCACTTTGTACAAGAAAGCTGGGTTCACATGTCTTCCGTGCTCTGA |
| 1_known | L24 | Tb927.11.9530_for | GGGGACAAGTTTGTACAAAAAAGCAGGCTATGACGGACTGCATCAAGTGGT | Tb927.11.9530_rev | GGGGACCACTTTGTACAAGAAAGCTGGGTTCAGTTCACGGGTTGCTCGT |
| 1_known | M1 | Tb927.10.15950_for | GGGGACAAGTTTGTACAAAAAAGCAGGCTATGGACAATGACTTCACTGGAAACTT | Tb927.10.15950_rev | GGGGACCACTTTGTACAAGAAAGCTGGGTTTACCTCTTTGCGTACTGTGCAAT |
| 1_known | M2 | Tb927.11.14000_for | GGGGACAAGTTTGTACAAAAAAGCAGGCTATGGCCCCAAAGTCTGCTACA | Tb927.11.14000_rev | GGGGACCACTTTGTACAAGAAAGCTGGGTTCACTGCTTCCTCTTGGCATCC |
| 1_known | M3 | Tb927.10.9840_for | GGGGACAAGTTTGTACAAAAAAGCAGGCTATGGTCTCCAAGAAATCACTGTATGATG | Tb927.10.9840_rev | GGGGACCACTTTGTACAAGAAAGCTGGGTTTATTTCTTTGTCTTTTTTCGAGGTGCC |
| 1_known | M4 | Tb927.8.2000_for | GGGGACAAGTTTGTACAAAAAAGCAGGCTATGCCCAAGGTAAGCGTCCC | Tb927.8.2000_rev | GGGGACCACTTTGTACAAGAAAGCTGGGTTTATTTCTCTTCCGCCTGGGCG |
| 1_known | M5 | Tb927.6.4030_for | GGGGACAAGTTTGTACAAAAAAGCAGGCTATGCGTTTGTGCCGGCGC | Tb927.6.4030_rev | GGGGACCACTTTGTACAAGAAAGCTGGGTTTAAACGACTTGTCCAATAGCCATAAGAT |
| 1_known | M6 | Tb927.6.4820_for | GGGGACAAGTTTGTACAAAAAAGCAGGCTATGAGCAACCCTTGGAACGC | Tb927.6.4820_rev | GGGGACCACTTTGTACAAGAAAGCTGGGTTCACCTTTGAATCAAGTTTGAACGGA |
| 1_known | M7 | Tb927.9.3470_for | GGGGACAAGTTTGTACAAAAAAGCAGGCTATGTCCACCAGCAGCCGC | Tb927.9.3470_rev | GGGGACCACTTTGTACAAGAAAGCTGGGTCTATGAAATAAATTCCTCCAGTAGCGCT |
| 1_known | M8 | Tb927.10.6810_for | GGGGACAAGTTTGTACAAAAAAGCAGGCTATGTTGCCCCCTCCCCAC | Tb927.10.6810_rev | GGGGACCACTTTGTACAAGAAAGCTGGGTTCACAACTTACTGCTCGGTGAGC |
| 1_known | M9 | Tb927.11.15170_for | GGGGACAAGTTTGTACAAAAAAGCAGGCTATGGGGAAGTCCAAAGGGCC | Tb927.11.15170_rev | GGGGACCACTTTGTACAAGAAAGCTGGGTTCATACTCCGTTACACTCGCTAACC |
| 1_known | M10 | Tb927.11.9390_for | GGGGACAAGTTTGTACAAAAAAGCAGGCTATGAATGGAATGCGGGCGTATAAC | Tb927.11.9390_rev | GGGGACCACTTTGTACAAGAAAGCTGGGTTTATTCAAGTTCGCAAAGGATGCTTCG |
| 1_known | M11 | Tb927.10.4130_for | GGGGACAAGTTTGTACAAAAAAGCAGGCTATGCGTTCCTCCCAATTTTTGCT | Tb927.10.4130_rev | GGGGACCACTTTGTACAAGAAAGCTGGGTTCACTTGCCCAGGTCTGACTT |
| 1_known | M12 | Tb927.11.10270_for | GGGGACAAGTTTGTACAAAAAAGCAGGCTATGGATAGCAAACAACGAATATGGGAT | Tb927.11.10270_rev | GGGGACCACTTTGTACAAGAAAGCTGGGTTTAGAATAAAAACAACACGAAAATTGTCAT |
| 1_known | M13 | Tb927.11.6330_for | GGGGACAAGTTTGTACAAAAAAGCAGGCTATGTCATTCAAGCCAACCATCAGC | Tb927.11.6330_rev | GGGGACCACTTTGTACAAGAAAGCTGGGTTCAAAACTTTGCCAAGTTTTCAGCA |
| 1_known | M14 | Tb927.10.16140_for | GGGGACAAGTTTGTACAAAAAAGCAGGCTATGTCACTCCCAATTATGAGTGTGGT | Tb927.10.16140_rev | GGGGACCACTTTGTACAAGAAAGCTGGGTTCAATGGGTTACTTGTTTCATCTGCG |
| 1_known | M15 | Tb927.10.8050_for | GGGGACAAGTTTGTACAAAAAAGCAGGCTATGCCGCGGGTTCAGCGA | Tb927.10.8050_rev | GGGGACCACTTTGTACAAGAAAGCTGGGTTCAACGGACTAACAGATTCCACCG |
| 1_known | M16 | Tb927.11.7310_for | GGGGACAAGTTTGTACAAAAAAGCAGGCTATGCCTAGTAGGGCAGTTCTTCG | Tb927.11.7310_rev | GGGGACCACTTTGTACAAGAAAGCTGGGTCTAATGCTCGCTGCCAGAAGC |
| 1_known | M17 | Tb927.10.6030_for | GGGGACAAGTTTGTACAAAAAAGCAGGCTATGTTTAAAAACCAGTACGACACGGA | Tb927.10.6030_rev | GGGGACCACTTTGTACAAGAAAGCTGGGTTCACTCCTCAACGTCAGGGG |
| 1_known | M18 | Tb927.8.6240_for | GGGGACAAGTTTGTACAAAAAAGCAGGCTATGACAACATTGCACACTATTTCCAGC | Tb927.8.6240_rev | GGGGACCACTTTGTACAAGAAAGCTGGGTCTAATCCGCAATTTCTCCTGAGGG |
| 1_known | M19 | Tb927.11.15290_for | GGGGACAAGTTTGTACAAAAAAGCAGGCTATGGCAGTTACCCCAGACTACC | Tb927.11.15290_rev | GGGGACCACTTTGTACAAGAAAGCTGGGTCTACTTGGCAGATTGCTGGTCATG |
| 1_known | M20 | Tb927.7.2060_for | GGGGACAAGTTTGTACAAAAAAGCAGGCTATGCCTCCCCTCCCACGG | Tb927.7.2060_rev | GGGGACCACTTTGTACAAGAAAGCTGGGTTCATTGGCCACCTTCCTGCA |
| 1_known | M21 | Tb927.11.7230_for | GGGGACAAGTTTGTACAAAAAAGCAGGCTATGCCCGTGCTCCCGCCG | Tb927.11.7230_rev | GGGGACCACTTTGTACAAGAAAGCTGGGTTTAGACGAGGGCACATTCATCCT |
| 1_known | M22 | Tb927.9.10040_for | GGGGACAAGTTTGTACAAAAAAGCAGGCTATGCTGGACATCAAGATAGAAAAGTATGAC | Tb927.9.10040_rev | GGGGACCACTTTGTACAAGAAAGCTGGGTCTACGGTTCCTCATCGGCTTCC |
| 1_known | M23 | Tb927.5.2700_for | GGGGACAAGTTTGTACAAAAAAGCAGGCTATGTCAGAGCATGCAACGACC | Tb927.5.2700_rev | GGGGACCACTTTGTACAAGAAAGCTGGGTCTACTTGTAAAGAAGGTCGTAATGACCC |
| 1_known | M24 | Tb927.9.9990_for | GGGGACAAGTTTGTACAAAAAAGCAGGCTATGGCCACTCCCTCCGCC | Tb927.9.9990_rev | GGGGACCACTTTGTACAAGAAAGCTGGGTTCAAGCGTGTAAATTGTTAAGATAAAAGTC |
| 1_known | N1 | Tb927.8.2210_for | GGGGACAAGTTTGTACAAAAAAGCAGGCTATGGAAGCTCCCGCTGCA | Tb927.8.2210_rev | GGGGACCACTTTGTACAAGAAAGCTGGGTTTAGGCATGCACAAGGCTTAACC |
| 1_known | N2 | Tb927.1.860_for | GGGGACAAGTTTGTACAAAAAAGCAGGCTATGTTGGGGCTGAAGGCGA | Tb927.1.860_rev | GGGGACCACTTTGTACAAGAAAGCTGGGTTCACTGCTTAGGCGCGAAAAT |
| 1_known | N3 | Tb927.10.9250_for | GGGGACAAGTTTGTACAAAAAAGCAGGCTATGCCGCCGCCGCCACCA | Tb927.10.9250_rev | GGGGACCACTTTGTACAAGAAAGCTGGGTCTACGCCACCTCATTTGGCTT |
| 1_known | N4 | Tb927.11.180_for | GGGGACAAGTTTGTACAAAAAAGCAGGCTATGTTATCTCGCACGGTTCCTGG | Tb927.11.180_rev | GGGGACCACTTTGTACAAGAAAGCTGGGTTCAAATAACTTTAGCTTCATTGTGGAGTTT |
| 1_known | N5 | Tb927.3.4810_for | GGGGACAAGTTTGTACAAAAAAGCAGGCTATGTTTTTTGAGGATCAACTCGACGA | Tb927.3.4810_rev | GGGGACCACTTTGTACAAGAAAGCTGGGTTCAAACCAACCTCACGCGTC |
| 1_known | N6 | Tb927.10.1520_for | GGGGACAAGTTTGTACAAAAAAGCAGGCTATGCGGTCGTCGGTGCCT | Tb927.10.1520_rev | GGGGACCACTTTGTACAAGAAAGCTGGGTTCAGGAACAACAGCGGTTCCG |
| 1_known | N7 | Tb927.4.4650_for | GGGGACAAGTTTGTACAAAAAAGCAGGCTATGTTCACCCTAATAGGTCTGGGAC | Tb927.4.4650_rev | GGGGACCACTTTGTACAAGAAAGCTGGGTTTAATTAAGGTAAAAAAGACGCAAGTGCTC |
| 1_known | N8 | Tb927.7.2440_for | GGGGACAAGTTTGTACAAAAAAGCAGGCTATGAAGATATCATTTATTGGATGTGGCAAC | Tb927.7.2440_rev | GGGGACCACTTTGTACAAGAAAGCTGGGTTTAATTCAATCCCTTCTCAAGTTCCTTTGA |
| 1_known | N9 | Tb927.10.5230_for | GGGGACAAGTTTGTACAAAAAAGCAGGCTATGAGCAGTTCTGAACACGATAGTC | Tb927.10.5230_rev | GGGGACCACTTTGTACAAGAAAGCTGGGTCTATTCTTCAAGCAACTTCTTCCTCTTTTG |
| 1_known | N10 | Tb927.2.2520_for | GGGGACAAGTTTGTACAAAAAAGCAGGCTATGTTTGCACCTGCAAAATCCCT | Tb927.2.2520_rev | GGGGACCACTTTGTACAAGAAAGCTGGGTCTACGAATGGGTAATAAGGACACCG |
| 1_known | N11 | Tb927.8.3240_for | GGGGACAAGTTTGTACAAAAAAGCAGGCTATGGAAGTGGCATTCGACGTTT | Tb927.8.3240_rev | GGGGACCACTTTGTACAAGAAAGCTGGGTTTATTCCTGTTTTTGAGCTGAAACTCGA |
| 1_known | N12 | Tb927.3.4850_for | GGGGACAAGTTTGTACAAAAAAGCAGGCTATGGATTAAATGTTACGCTCCTGTGC | Tb927.3.4850_rev | GGGGACCACTTTGTACAAGAAAGCTGGGTCTACGAATTGGTGAAGAACGGTTTCC |
| 1_known | N14 | Tb927.10.15210_for | GGGGACAAGTTTGTACAAAAAAGCAGGCTATGCGACGGGGAAGTGGA | Tb927.10.15210_rev | GGGGACCACTTTGTACAAGAAAGCTGGGTTTAGTTGAACTCTAACAGTTCATCAATTAA |
| 1_known | N15 | Tb927.7.4170_for | GGGGACAAGTTTGTACAAAAAAGCAGGCTATGTTTCCATACGTCACAGACTATAGCG | Tb927.7.4170_rev | GGGGACCACTTTGTACAAGAAAGCTGGGTTCATTTCGACTTTTCGGTCCTCTTT |
| 1_known | N16 | Tb927.10.9400_for | GGGGACAAGTTTGTACAAAAAAGCAGGCTATGGGGGAGAACCGTCGAC | Tb927.10.9400_rev | GGGGACCACTTTGTACAAGAAAGCTGGGTCTACAACTCGTCCAGAAACTCCTG |
| 1_known | N17 | Tb927.11.15150_for | GGGGACAAGTTTGTACAAAAAAGCAGGCTATGGGAGCCAACTACGCGG | Tb927.11.15150_rev | GGGGACCACTTTGTACAAGAAAGCTGGGTCTAGCGGTCCTTGTGACTATTCTCC |
| 1_known | N18 | Tb927.3.3490_for | GGGGACAAGTTTGTACAAAAAAGCAGGCTATGGCAACAGAATTGAAGAAAGGACC | Tb927.3.3490_rev | GGGGACCACTTTGTACAAGAAAGCTGGGTCTACTTGGGTAGTGCTGCCATTTC |
| 1_known | N19 | Tb927.9.8680_for | GGGGACAAGTTTGTACAAAAAAGCAGGCTATGCGGCGGGTATCAGGA | Tb927.9.8680_rev | GGGGACCACTTTGTACAAGAAAGCTGGGTTCACCAGTTTGTCCACTCCGG |
| 1_known | N20 | Tb927.4.1930_for | GGGGACAAGTTTGTACAAAAAAGCAGGCTATGGCAACATGGGCCGATG | Tb927.4.1930_rev | GGGGACCACTTTGTACAAGAAAGCTGGGTTCAGTTGCGCTGTTTCGCA |
| 1_known | N21 | Tb927.8.5540_for | GGGGACAAGTTTGTACAAAAAAGCAGGCTATGATGCGCTTCTTCGTATGGATAAG | Tb927.8.5540_rev | GGGGACCACTTTGTACAAGAAAGCTGGGTTCAAAGCTGCTCGTCAGCG |
| 1_known | N22 | Tb927.2.3440_for | GGGGACAAGTTTGTACAAAAAAGCAGGCTATGGCGGCGAGGAAAGCA | Tb927.2.3440_rev | GGGGACCACTTTGTACAAGAAAGCTGGGTTTATTCATACCGCACCCACCCC |
| 1_known | N23 | Tb927.6.4050_for | GGGGACAAGTTTGTACAAAAAAGCAGGCTATGGGGTTCCCTTCCGGG | Tb927.6.4050_rev | GGGGACCACTTTGTACAAGAAAGCTGGGTTTAGTGAGAAGCACCGGGTGG |
| 1_known | N24 | Tb927.7.6350_for | GGGGACAAGTTTGTACAAAAAAGCAGGCTATGTGGCGTCGAGCTGGG | Tb927.7.6350_rev | GGGGACCACTTTGTACAAGAAAGCTGGGTCTACTTCTTTGCTGCGCCGC |
| 1_known | O1 | Tb927.9.1960_for | GGGGACAAGTTTGTACAAAAAAGCAGGCTATGCGCGTTACTTTGTGTCAGA | Tb927.9.1960_rev | GGGGACCACTTTGTACAAGAAAGCTGGGTCTACTTCCAAGTTAAAGAATAAAGATCATG |
| 1_known | O2 | Tb927.9.6940_for | GGGGACAAGTTTGTACAAAAAAGCAGGCTATGTCTTCGGCTAACTCGGAGG | Tb927.9.6940_rev | GGGGACCACTTTGTACAAGAAAGCTGGGTTCAAAACGTAAAGTCCCTTGCCTT |
| 1_known | O3 | Tb927.9.6060_for | GGGGACAAGTTTGTACAAAAAAGCAGGCTATGTCGAAACAGGAGCCGTTG | Tb927.9.6060_rev | GGGGACCACTTTGTACAAGAAAGCTGGGTTCACGCTGATGATTTCTTATCATCAGG |
| 1_known | O4 | Tb927.11.15350_for | GGGGACAAGTTTGTACAAAAAAGCAGGCTATGTCCTCCACACAGACGACG | Tb927.11.15350_rev | GGGGACCACTTTGTACAAGAAAGCTGGGTCTACTCAAGGTCTGCCGCAAG |
| 1_known | O5 | Tb927.10.13490_for | GGGGACAAGTTTGTACAAAAAAGCAGGCTATGACACCTCCTGTGCGGC | Tb927.10.13490_rev | GGGGACCACTTTGTACAAGAAAGCTGGGTTCACGCCGACTCGTTCAGA |
| 1_known | O6 | Tb927.10.6610_for | GGGGACAAGTTTGTACAAAAAAGCAGGCTATGAACTCTATGGATGATGCGGCA | Tb927.10.6610_rev | GGGGACCACTTTGTACAAGAAAGCTGGGTCTAACGGTGCATGGCTTGACG |
| 1_known | O7 | Tb927.6.1860_for | GGGGACAAGTTTGTACAAAAAAGCAGGCTATGCGTCGGATGCTGCTAGC | Tb927.6.1860_rev | GGGGACCACTTTGTACAAGAAAGCTGGGTCTATGCTACAACGTCTGATTCGTCG |
| 1_known | O8 | Tb927.11.8820_for | GGGGACAAGTTTGTACAAAAAAGCAGGCTATGGCGAAAGGAAGCGATTATGAT | Tb927.11.8820_rev | GGGGACCACTTTGTACAAGAAAGCTGGGTCTACGTCTTGTACCGCATCCAG |
| 1_known | O9 | Tb927.3.3630_for | GGGGACAAGTTTGTACAAAAAAGCAGGCTATGTTGTTTCGTACGATTCGGCA | Tb927.3.3630_rev | GGGGACCACTTTGTACAAGAAAGCTGGGTTTACTTTGTTCCTGCCACCGGC |
| 1_known | O10 | Tb927.9.5280_for | GGGGACAAGTTTGTACAAAAAAGCAGGCTATGCTCCGCGGTACCCCA | Tb927.9.5280_rev | GGGGACCACTTTGTACAAGAAAGCTGGGTTCACTTCCCACCTCCTTCCCT |
| 1_known | O11 | Tb927.11.14430_for | GGGGACAAGTTTGTACAAAAAAGCAGGCTATGTTGACCACCACCGACGT | Tb927.11.14430_rev | GGGGACCACTTTGTACAAGAAAGCTGGGTCTACACAATGCGCTGCAGATCA |
| 1_known | O12 | Tb927.6.3970_for | GGGGACAAGTTTGTACAAAAAAGCAGGCTATGTCCCTCTTTACAGTGCATTGC | Tb927.6.3970_rev | GGGGACCACTTTGTACAAGAAAGCTGGGTCTAGTGATCAATTCCACTGCTAGTCATC |
| 1_known | O13 | Tb927.10.1130_for | GGGGACAAGTTTGTACAAAAAAGCAGGCTATGCAACCTTACGGGATGTCAAC | Tb927.10.1130_rev | GGGGACCACTTTGTACAAGAAAGCTGGGTTTACGGAGTTGAAGGCATCGAAATC |
| 1_known | O14 | Tb927.11.3900_for | GGGGACAAGTTTGTACAAAAAAGCAGGCTATGTTTTGCCGTTCCGTGAAGT | Tb927.11.3900_rev | GGGGACCACTTTGTACAAGAAAGCTGGGTTCAGCGAACGGACGTCCC |
| 1_known | O15 | Tb927.10.13480_for | GGGGACAAGTTTGTACAAAAAAGCAGGCTATGGAAAATTTTGTTCTTCCACCGTACA | Tb927.10.13480_rev | GGGGACCACTTTGTACAAGAAAGCTGGGTTTACGCAGCCTCCTTTAGACACT |
| 1_known | O16 | Tb927.10.5130_for | GGGGACAAGTTTGTACAAAAAAGCAGGCTATGATCCTTAAGGAAGAGACCGTACTG | Tb927.10.5130_rev | GGGGACCACTTTGTACAAGAAAGCTGGGTTCATGTTCGGTCCCGCACT |
| 1_known | O17 | Tb927.3.5330_for | GGGGACAAGTTTGTACAAAAAAGCAGGCTATGACGGAAAAGGAAAATGGAAACAAAG | Tb927.3.5330_rev | GGGGACCACTTTGTACAAGAAAGCTGGGTTCAAGAAAAGCCAAACTGCTCCA |
| 1_known | O18 | Tb927.1.570_for | GGGGACAAGTTTGTACAAAAAAGCAGGCTATGGACGCACGTGTTCATCG | Tb927.1.570_rev | GGGGACCACTTTGTACAAGAAAGCTGGGTTCACGATCGCTTGATGTAAAGATAAAATG |
| 1_known | O19 | Tb927.4.3130_for | GGGGACAAGTTTGTACAAAAAAGCAGGCTATGTCGGTTCCACTCACAGAGG | Tb927.4.3130_rev | GGGGACCACTTTGTACAAGAAAGCTGGGTTCACATGTGACCAAATGTGCTCC |
| 1_known | O20 | Tb927.11.4700_for | GGGGACAAGTTTGTACAAAAAAGCAGGCTATGGCTCTCACTCAATCCCTAAAACT | Tb927.11.4700_rev | GGGGACCACTTTGTACAAGAAAGCTGGGTTCAAAAGTCGTTCATGAAGACCTCC |
| 1_known | O21 | Tb927.6.2890_for | GGGGACAAGTTTGTACAAAAAAGCAGGCTATGAGAGGGCCTGTTACTATATTACTACAG | Tb927.6.2890_rev | GGGGACCACTTTGTACAAGAAAGCTGGGTCTAGAGTATACGCTGCTCTGCAGT |
| 1_known | O22 | Tb927.7.880_for | GGGGACAAGTTTGTACAAAAAAGCAGGCTATGAACAAACTTACATTTACGCCACTTAC | Tb927.7.880_rev | GGGGACCACTTTGTACAAGAAAGCTGGGTTTACGACTCTTCATTTCCTGCAACA |
| 1_known | O23 | Tb927.10.4000_for | GGGGACAAGTTTGTACAAAAAAGCAGGCTATGCGCCGTTGTCCGGTT | Tb927.10.4000_rev | GGGGACCACTTTGTACAAGAAAGCTGGGTTCATTTACCAACATATCTGGGTGGTCG |
| 1_known | O24 | Tb927.11.12750_for | GGGGACAAGTTTGTACAAAAAAGCAGGCTATGTTTACTGACAACGCTGCCC | Tb927.11.12750_rev | GGGGACCACTTTGTACAAGAAAGCTGGGTTTACTGCCTTCCCGTTGCATC |
| 1_known | P1 | Tb927.11.5170_for | GGGGACAAGTTTGTACAAAAAAGCAGGCTATGTTGAATGCGGCGCTTTACG | Tb927.11.5170_rev | GGGGACCACTTTGTACAAGAAAGCTGGGTTCAGCATCGTTTCACGCCTAC |
| 1_known | P2 | Tb927.9.1680_for | GGGGACAAGTTTGTACAAAAAAGCAGGCTATGGAGGCGAAGCATGCG | Tb927.9.1680_rev | GGGGACCACTTTGTACAAGAAAGCTGGGTTCACTTGGAAATACGCAGCCG |
| 1_known | P3 | Tb927.8.4810_for | GGGGACAAGTTTGTACAAAAAAGCAGGCTATGAGCTTCAGATTTGTCCAGCG | Tb927.8.4810_rev | GGGGACCACTTTGTACAAGAAAGCTGGGTTCAATGTGCCTGCTCAGATGC |
| 1_known | P5 | Tb927.8.4830_for | GGGGACAAGTTTGTACAAAAAAGCAGGCTATGCAGACCCCTGGTGAGG | Tb927.8.4830_rev | GGGGACCACTTTGTACAAGAAAGCTGGGTTCACGTAATACTCATGATATCATCCAGAAG |
| 1_known | P6 | Tb927.11.12340_for | GGGGACAAGTTTGTACAAAAAAGCAGGCTATGCTACGAAATTACGACACACCC | Tb927.11.12340_rev | GGGGACCACTTTGTACAAGAAAGCTGGGTTTAGGAGTCAACTTCTCGCCTGC |
| 1_known | P7 | Tb927.11.8440_for | GGGGACAAGTTTGTACAAAAAAGCAGGCTATGACCTACCGTGCAGCTGT | Tb927.11.8440_rev | GGGGACCACTTTGTACAAGAAAGCTGGGTCTAGTCTTCAATATTGAACAGTTCGCGA |
| 1_known | P8 | Tb927.11.14020_for | GGGGACAAGTTTGTACAAAAAAGCAGGCTATGGCCCCAAAGTCTGCTGC | Tb927.11.14020_rev | GGGGACCACTTTGTACAAGAAAGCTGGGTTTACTTCCTCTTGGCATCCTTCTTTTC |
| 1_known | P9 | Tb927.4.3680_for | GGGGACAAGTTTGTACAAAAAAGCAGGCTATGTTTACACCGTCGACCCGA | Tb927.4.3680_rev | GGGGACCACTTTGTACAAGAAAGCTGGGTTCATGTCTTGTGCACATCGAGAAG |
| 1_known | P10 | Tb927.2.5230_for | GGGGACAAGTTTGTACAAAAAAGCAGGCTATGGAGCGGTACAAGCTGCG | Tb927.2.5230_rev | GGGGACCACTTTGTACAAGAAAGCTGGGTTTACACAACTAACCAAGGGTGCC |
| 1_known | P11 | Tb927.2.3460_for | GGGGACAAGTTTGTACAAAAAAGCAGGCTATGGAAAGAGAGCCAGCGGT | Tb927.2.3460_rev | GGGGACCACTTTGTACAAGAAAGCTGGGTTTATTCATACCGCACCCACCCC |
| 1_known | P12 | Tb927.7.5680_for | GGGGACAAGTTTGTACAAAAAAGCAGGCTATGACCGACCTTCACATGAGAGA | Tb927.7.5680_rev | GGGGACCACTTTGTACAAGAAAGCTGGGTTTAGTATTTACTGCGGGAGCGGT |
| 1_known | P13 | Tb927.10.12800_for | GGGGACAAGTTTGTACAAAAAAGCAGGCTATGACCACAGCAACTGATGTGAC | Tb927.10.12800_rev | GGGGACCACTTTGTACAAGAAAGCTGGGTTTACAGCTGCTGGTAAGGATTATGTCG |
| 1_known | P14 | Tb927.7.1720_for | GGGGACAAGTTTGTACAAAAAAGCAGGCTATGATAAAGTTCACTCTGCGGTACTTC | Tb927.7.1720_rev | GGGGACCACTTTGTACAAGAAAGCTGGGTCTAGTTTGATTGTGAGGCGCTCA |
| 1_known | P15 | Tb927.4.360_for | GGGGACAAGTTTGTACAAAAAAGCAGGCTATGGAACCCGACACGGATGC | Tb927.4.360_rev | GGGGACCACTTTGTACAAGAAAGCTGGGTCTACCTGAAGCCAAGTTCCTTTCC |
| 1_known | P16 | Tb927.11.16770_for | GGGGACAAGTTTGTACAAAAAAGCAGGCTATGCGTATCGTCATTTCCCCCG | Tb927.11.16770_rev | GGGGACCACTTTGTACAAGAAAGCTGGGTTCACTTTGGTGGTGGTGCTTC |
| 1_known | P17 | Tb927.5.4320_for | GGGGACAAGTTTGTACAAAAAAGCAGGCTATGGAAGAGGGCGGTGAAGG | Tb927.5.4320_rev | GGGGACCACTTTGTACAAGAAAGCTGGGTTTACGTGTCCTCAGAGTGTTGTTG |
| 1_known | P18 | Tb927.11.12570_for | GGGGACAAGTTTGTACAAAAAAGCAGGCTATGCAGCTAAAGGAGCGCC | Tb927.11.12570_rev | GGGGACCACTTTGTACAAGAAAGCTGGGTTCAGACCGCCAACAAATCACTTG |
| 1_known | P19 | Tb927.8.6390_for | GGGGACAAGTTTGTACAAAAAAGCAGGCTATGTTTGGCACGCCGGTT | Tb927.8.6390_rev | GGGGACCACTTTGTACAAGAAAGCTGGGTTTACGATTTCGATGAAGGTCCGGG |
| 1_known | P20 | Tb927.11.7150_for | GGGGACAAGTTTGTACAAAAAAGCAGGCTATGACTTCTCTTATCAGTCGAGTAGCA | Tb927.11.7150_rev | GGGGACCACTTTGTACAAGAAAGCTGGGTTCACCAAACAACAAGTGGATCCTTG |
| 1_known | P21 | Tb927.11.14030_for | GGGGACAAGTTTGTACAAAAAAGCAGGCTATGGCAGGTTTCACCGAGACA | Tb927.11.14030_rev | GGGGACCACTTTGTACAAGAAAGCTGGGTTCACTCCACCGATGGTGCA |
| 1_known | P22 | Tb927.1.3110_for | GGGGACAAGTTTGTACAAAAAAGCAGGCTATGTCTGCTGAGTCCATATACCGC | Tb927.1.3110_rev | GGGGACCACTTTGTACAAGAAAGCTGGGTCTACAACAGACTACTCATTTTGTTCTTCAC |
| 1_known | P23 | Tb927.6.1260_for | GGGGACAAGTTTGTACAAAAAAGCAGGCTATGTTGAGACGCCCCCCC | Tb927.6.1260_rev | GGGGACCACTTTGTACAAGAAAGCTGGGTCTACTGAGAAGTCATTAGGTTCTGGCA |
| 1_known | P24 | Tb927.11.5290_for | GGGGACAAGTTTGTACAAAAAAGCAGGCTATGGAAGTTGTTGTAAGTTTTTTAAGTGGA | Tb927.11.5290_rev | GGGGACCACTTTGTACAAGAAAGCTGGGTCTACATCCCACAGAGATTGTTTAGTGTG |
| 2_hypothetical | A1 | Tb927.10.780_for | GGGGACAAGTTTGTACAAAAAAGCAGGCTATGTATTTTTGCCACGGCTGCC | Tb927.10.780_rev | GGGGACCACTTTGTACAAGAAAGCTGGGTCTATGCTGTGGTGTTCGTGTCTG |
| 2_hypothetical | A2 | Tb927.11.7850_for | GGGGACAAGTTTGTACAAAAAAGCAGGCTATGGTGGTGTCTATGCAGAGGT | Tb927.11.7850_rev | GGGGACCACTTTGTACAAGAAAGCTGGGTCTACCCGAGCCGCCTAATAAGG |

| | | | | | |
|---|---|---|---|---|---|
| 2_hypothetical | A3 | Tb927.10.1280_for | GGGGACAAGTTTGTACAAAAAAGCAGGCTATGGGAATGAGAAAAAAGGAAACAATAAAG | Tb927.10.1280_rev | GGGGACCACTTTGTACAAGAAAGCTGGGTCTATGCCTTCTTCGATACGTCGTC |
| 2_hypothetical | A4 | Tb927.3.2350_for | GGGGACAAGTTTGTACAAAAAAGCAGGCTATGGATCCGGCGGTAGGG | Tb927.3.2350_rev | GGGGACCACTTTGTACAAGAAAGCTGGGTCTACCGTTTCCGCTTCAATTTCTCG |
| 2_hypothetical | A5 | Tb927.10.150_for | GGGGACAAGTTTGTACAAAAAAGCAGGCTATGAGCACTAGTGAACGGCGT | Tb927.10.150_rev | GGGGACCACTTTGTACAAGAAAGCTGGGTCTAGAGAGAATCAAAAAAATCATCAATATC |
| 2_hypothetical | A6 | Tb927.5.1420_for | GGGGACAAGTTTGTACAAAAAAGCAGGCTATGTTCAGATATCATGAGAGTATAAGGGAA | Tb927.5.1420_rev | GGGGACCACTTTGTACAAGAAAGCTGGGTCTACCTAACAACTTGTTCATTATGAGTTGA |
| 2_hypothetical | A7 | Tb927.5.1400_for | GGGGACAAGTTTGTACAAAAAAGCAGGCTATGTTCAGATATCATGAGAGTATAAGGGAA | Tb927.5.1400_rev | GGGGACCACTTTGTACAAGAAAGCTGGGTCTACCTAACAACTTGGTCATTATGAGTAGA |
| 2_hypothetical | A8 | Tb927.7.6840_for | GGGGACAAGTTTGTACAAAAAAGCAGGCTATGGAGAAGTTGACACTCTCGGC | Tb927.7.6840_rev | GGGGACCACTTTGTACAAGAAAGCTGGGTTTACCCGCGGGGAAGATTGT |
| 2_hypothetical | A9 | Tb927.8.5670_for | GGGGACAAGTTTGTACAAAAAAGCAGGCTATGGTAGAGAAGAAGCAAGCAAAGAAA | Tb927.8.5670_rev | GGGGACCACTTTGTACAAGAAAGCTGGGTTCACTTGTCACTGTCTGTGTCACT |
| 2_hypothetical | A10 | Tb927.11.4810_for | GGGGACAAGTTTGTACAAAAAAGCAGGCTATGAGGGCTCTCCGCCTAAC | Tb927.11.4810_rev | GGGGACCACTTTGTACAAGAAAGCTGGGTCTAGTGTGAACGCCGCGG |
| 2_hypothetical | A11 | Tb927.4.2150_for | GGGGACAAGTTTGTACAAAAAAGCAGGCTATGATTGTCGGCCAAATGAGAAGG | Tb927.4.2150_rev | GGGGACCACTTTGTACAAGAAAGCTGGGTTTACCTTTTCACACGGGAAAGGAC |
| 2_hypothetical | A12 | Tb927.7.260_for | GGGGACAAGTTTGTACAAAAAAGCAGGCTATGACGGAGCGTTCACCGG | Tb927.7.260_rev | GGGGACCACTTTGTACAAGAAAGCTGGGTTCACTTTATGACTGCTTCAATAACCCC |
| 2_hypothetical | A13 | Tb927.11.420_for | GGGGACAAGTTTGTACAAAAAAGCAGGCTATGACGTCTAAATCTATTGACTCCATGATG | Tb927.11.420_rev | GGGGACCACTTTGTACAAGAAAGCTGGGTTCAAAGCATCATATCTCTTTTCTTTAAACG |
| 2_hypothetical | A14 | Tb927.2.5040_for | GGGGACAAGTTTGTACAAAAAAGCAGGCTATGCAGAGGGACGCTGACA | Tb927.2.5040_rev | GGGGACCACTTTGTACAAGAAAGCTGGGTCTATAAACTAACCTCCGCAAGCAAGC |
| 2_hypothetical | A15 | Tb927.9.10660_for | GGGGACAAGTTTGTACAAAAAAGCAGGCTATGAGCGGAGCTGACGATGA | Tb927.9.10660_rev | GGGGACCACTTTGTACAAGAAAGCTGGGTTCAGTCCTGCTTCTTGTCGGG |
| 2_hypothetical | A16 | Tb927.8.520_for | GGGGACAAGTTTGTACAAAAAAGCAGGCTATGCGGGTGTGGCGTTTAAG | Tb927.8.520_rev | GGGGACCACTTTGTACAAGAAAGCTGGGTTTAGAAGAAAAGAAAAATAACAATTATCAA |
| 2_hypothetical | A17 | Tb927.3.5470_for | GGGGACAAGTTTGTACAAAAAAGCAGGCTATGACTGGGCGGAAGCGT | Tb927.3.5470_rev | GGGGACCACTTTGTACAAGAAAGCTGGGTTTACTTTTCACCTATGCCATTGCTATCC |
| 2_hypothetical | A18 | Tb927.4.1380_for | GGGGACAAGTTTGTACAAAAAAGCAGGCTATGACCGTTCGTGTAGCACTCC | Tb927.4.1380_rev | GGGGACCACTTTGTACAAGAAAGCTGGGTTCACAAGTCTAAAACATGTGTGAGAGG |
| 2_hypothetical | A19 | Tb927.4.5360_for | GGGGACAAGTTTGTACAAAAAAGCAGGCTATGTACGCTGTGAAACCGTGG | Tb927.4.5360_rev | GGGGACCACTTTGTACAAGAAAGCTGGGTTCACGAGCGAGGCAAGTGA |
| 2_hypothetical | A20 | Tb927.9.3850_for | GGGGACAAGTTTGTACAAAAAAGCAGGCTATGGCCAAGGATACGGCGA | Tb927.9.3850_rev | GGGGACCACTTTGTACAAGAAAGCTGGGTTTACACTCTCTTCTTGCGCTGTTG |
| 2_hypothetical | A21 | Tb927.4.3500_for | GGGGACAAGTTTGTACAAAAAAGCAGGCTATGTGCTGCGGTTGGTTGAC | Tb927.4.3500_rev | GGGGACCACTTTGTACAAGAAAGCTGGGTCTACAGGCAACAACCCGCG |
| 2_hypothetical | A22 | Tb927.1.2420_for | GGGGACAAGTTTGTACAAAAAAGCAGGCTATGAGGTGGCCGGCTGAG | Tb927.1.2420_rev | GGGGACCACTTTGTACAAGAAAGCTGGGTTCACAATTTCACAGTTACGGCTTTGA |
| 2_hypothetical | A23 | Tb927.6.2620_for | GGGGACAAGTTTGTACAAAAAAGCAGGCTATGCGCCGTGATTGTGTAAGAT | Tb927.6.2620_rev | GGGGACCACTTTGTACAAGAAAGCTGGGTTCACGAATCAGATGTTAGGTGACCC |
| 2_hypothetical | A24 | Tb927.8.4520_for | GGGGACAAGTTTGTACAAAAAAGCAGGCTATGATAGACCTCACTAAGCAGATGACG | Tb927.8.4520_rev | GGGGACCACTTTGTACAAGAAAGCTGGGTTTACACTTTTCCAGCGAGCGG |
| 2_hypothetical | B1 | Tb927.11.11180_for | GGGGACAAGTTTGTACAAAAAAGCAGGCTATGTCGGTGAAGGTTTGGGTAAAAT | Tb927.11.11180_rev | GGGGACCACTTTGTACAAGAAAGCTGGGTTCAAAAGATAGTACTCCGGCAGCT |
| 2_hypothetical | B2 | Tb927.5.2810_for | GGGGACAAGTTTGTACAAAAAAGCAGGCTATGTCACCTGCCTCACCACT | Tb927.5.2810_rev | GGGGACCACTTTGTACAAGAAAGCTGGGTTTACTTTGTGTTATCACCCTCCTTCTTG |
| 2_hypothetical | B3 | Tb927.6.3210_for | GGGGACAAGTTTGTACAAAAAAGCAGGCTATGCAACAGCCGAAAGGAATCA | Tb927.6.3210_rev | GGGGACCACTTTGTACAAGAAAGCTGGGTCTAAACTGCCTGAATGGAACCGAC |
| 2_hypothetical | B4 | Tb927.1.4710_for | GGGGACAAGTTTGTACAAAAAAGCAGGCTATGAGCGGATTGCCTGGAAGT | Tb927.1.4710_rev | GGGGACCACTTTGTACAAGAAAGCTGGGTTTACTCCGTCGACTGGTCCTC |
| 2_hypothetical | B5 | Tb927.8.4420_for | GGGGACAAGTTTGTACAAAAAAGCAGGCTATGGAAGGAAGTGGTAGCTACCG | Tb927.8.4420_rev | GGGGACCACTTTGTACAAGAAAGCTGGGTTCAGCATACAAACTCCTTCTCACCA |
| 2_hypothetical | B6 | Tb927.4.3520_for | GGGGACAAGTTTGTACAAAAAAGCAGGCTATGCTGTGTTGCGGCTGTT | Tb927.4.3520_rev | GGGGACCACTTTGTACAAGAAAGCTGGGTCTAAAGGCAACAGTGCGTGATTG |
| 2_hypothetical | B7 | Tb927.11.9405_for | GGGGACAAGTTTGTACAAAAAAGCAGGCTATGGAAGGGGAGGAGAAGGTAGA | Tb927.11.9405_rev | GGGGACCACTTTGTACAAGAAAGCTGGGTTCAGATGCGGGTCCCACAA |
| 2_hypothetical | B8 | Tb927.10.6420_for | GGGGACAAGTTTGTACAAAAAAGCAGGCTATGGGGTGCGCACCTCTT | Tb927.10.6420_rev | GGGGACCACTTTGTACAAGAAAGCTGGGTTTACTGAAAGCGCCCCTTATATGTTT |
| 2_hypothetical | B9 | Tb927.10.8370_for | GGGGACAAGTTTGTACAAAAAAGCAGGCTATGCGGAGGAAAACGACACTAAAT | Tb927.10.8370_rev | GGGGACCACTTTGTACAAGAAAGCTGGGTTTAGATAGGTTTTTTCCTGCCTTTATGTT |
| 2_hypothetical | B10 | Tb927.1.4020_for | GGGGACAAGTTTGTACAAAAAAGCAGGCTATGCATTTCATTATTTTAATGTGCTCCCG | Tb927.1.4020_rev | GGGGACCACTTTGTACAAGAAAGCTGGGTTTAAACTTCTAAACCTGATAGAAAAAAAAA |
| 2_hypothetical | B11 | Tb927.10.6020_for | GGGGACAAGTTTGTACAAAAAAGCAGGCTATGGCGAAAAAGACCACCAAGAA | Tb927.10.6020_rev | GGGGACCACTTTGTACAAGAAAGCTGGGTTCATGAACATAGCAACTCCATCAGTG |
| 2_hypothetical | B12 | Tb927.10.16050_for | GGGGACAAGTTTGTACAAAAAAGCAGGCTATGTCAGATGGACTTCCGTGGC | Tb927.10.16050_rev | GGGGACCACTTTGTACAAGAAAGCTGGGTTCACAGTACATCACACAGGTTTTCTC |
| 2_hypothetical | B13 | Tb927.5.2910_for | GGGGACAAGTTTGTACAAAAAAGCAGGCTATGTTAGTCGAACGATCGTACAAAACC | Tb927.5.2910_rev | GGGGACCACTTTGTACAAGAAAGCTGGGTTTAAGTACTTAGTCCCTCAACAAGTCGC |
| 2_hypothetical | B14 | Tb927.8.6460_for | GGGGACAAGTTTGTACAAAAAAGCAGGCTATGGAACGAATTGGTGAGTTTACCG | Tb927.8.6460_rev | GGGGACCACTTTGTACAAGAAAGCTGGGTTCAAATACCGCTTGTGGCTAACC |
| 2_hypothetical | B15 | Tb927.2.2540_for | GGGGACAAGTTTGTACAAAAAAGCAGGCTATGGTGAAGCCTCGTTCTCAGG | Tb927.2.2540_rev | GGGGACCACTTTGTACAAGAAAGCTGGGTTCATTCCCTTTTAACCCTCCCTACA |
| 2_hypothetical | B16 | Tb927.10.9550_for | GGGGACAAGTTTGTACAAAAAAGCAGGCTATGCTTGCGCAGCAAGGT | Tb927.10.9550_rev | GGGGACCACTTTGTACAAGAAAGCTGGGTTCACTTTAGGAAGGGTAGCCGG |
| 2_hypothetical | B17 | Tb927.11.13330_for | GGGGACAAGTTTGTACAAAAAAGCAGGCTATGGCTGCTTGCGATGTGAC | Tb927.11.13330_rev | GGGGACCACTTTGTACAAGAAAGCTGGGTTTAGTATGCGAACGGGTCAACTTG |
| 2_hypothetical | B18 | Tb927.10.12750_for | GGGGACAAGTTTGTACAAAAAAGCAGGCTATGTGCGTATATAATTATATCATTTATATT | Tb927.10.12750_rev | GGGGACCACTTTGTACAAGAAAGCTGGGTTCACAGAAACAAAATAATGTTAAAAAAGAA |
| 2_hypothetical | B19 | Tb927.5.3180_for | GGGGACAAGTTTGTACAAAAAAGCAGGCTATGTCAAGCAATGAAAAGGTCAGTAGT | Tb927.5.3180_rev | GGGGACCACTTTGTACAAGAAAGCTGGGTTCATTGTGATTGACGTGCGATTCC |
| 2_hypothetical | B20 | Tb927.1.5260_for | GGGGACAAGTTTGTACAAAAAAGCAGGCTATGTGCAGCTCCTCAAGCTTTT | Tb927.1.5260_rev | GGGGACCACTTTGTACAAGAAAGCTGGGTCTACATCTTAATTGCGGGAGGTGC |
| 2_hypothetical | B21 | Tb927.11.10400_for | GGGGACAAGTTTGTACAAAAAAGCAGGCTATGCTTCGGTTCACTCATAGGGC | Tb927.11.10400_rev | GGGGACCACTTTGTACAAGAAAGCTGGGTCACGAAGGGGTGCCTGTT |
| 2_hypothetical | B22 | Tb927.9.8810_for | GGGGACAAGTTTGTACAAAAAAGCAGGCTATGCCTCCCCGCAGCGCG | Tb927.9.8810_rev | GGGGACCACTTTGTACAAGAAAGCTGGGTCTAACTTTCTTGCAGGCTCGCT |
| 2_hypothetical | B23 | Tb927.9.2910_for | GGGGACAAGTTTGTACAAAAAAGCAGGCTATGTCCGGTAGGGGTCATGG | Tb927.9.2910_rev | GGGGACCACTTTGTACAAGAAAGCTGGGTTTATCCCTTCCCACGTGCCC |
| 2_hypothetical | B24 | Tb927.7.320_for | GGGGACAAGTTTGTACAAAAAAGCAGGCTATGTATACCAGTGGCCATCCGA | Tb927.7.320_rev | GGGGACCACTTTGTACAAGAAAGCTGGGTCTAAATCACGCGACTCTCCGG |
| 2_hypothetical | C1 | Tb927.10.14540_for | GGGGACAAGTTTGTACAAAAAAGCAGGCTATGTTTGAGGGGTATTTAAAGTGGTGG | Tb927.10.14540_rev | GGGGACCACTTTGTACAAGAAAGCTGGGTTCATTCCTTCCCTTGTTTTAATGTTGCG |
| 2_hypothetical | C2 | Tb927.11.13510_for | GGGGACAAGTTTGTACAAAAAAGCAGGCTATGAACCCGTCAAAGGTAGTCCA | Tb927.11.13510_rev | GGGGACCACTTTGTACAAGAAAGCTGGGTTTAGGCTGTCAGGATTCCCTCTTT |
| 2_hypothetical | C3 | Tb927.6.4330_for | GGGGACAAGTTTGTACAAAAAAGCAGGCTATGGCAGAGGGCAGTGACA | Tb927.6.4330_rev | GGGGACCACTTTGTACAAGAAAGCTGGGTTCAAGTGGTAGATTCAGTTTCAGACG |
| 2_hypothetical | C4 | Tb927.7.2740_for | GGGGACAAGTTTGTACAAAAAAGCAGGCTATGGAATACCAAGGTTCTCCACAGC | Tb927.7.2740_rev | GGGGACCACTTTGTACAAGAAAGCTGGGTCTAAAAAAAAACTTCATTAGCATATCGGCA |
| 2_hypothetical | C5 | Tb927.11.3060_for | GGGGACAAGTTTGTACAAAAAAGCAGGCTATGAACATTATACTAATAGGACTCGTTATG | Tb927.11.3060_rev | GGGGACCACTTTGTACAAGAAAGCTGGGTCTACTGGCCTTTCCTGCCCT |
| 2_hypothetical | C6 | Tb927.11.3530_for | GGGGACAAGTTTGTACAAAAAAGCAGGCTATGTTCAAACAACTCTACAAACGTGAC | Tb927.11.3530_rev | GGGGACCACTTTGTACAAGAAAGCTGGGTTCACCAACCCTGGGGAAACA |
| 2_hypothetical | C7 | Tb927.11.17860_for | GGGGACAAGTTTGTACAAAAAAGCAGGCTATGTTGTACGGTGGCACCTGC | Tb927.11.17860_rev | GGGGACCACTTTGTACAAGAAAGCTGGGTCTAATCCCGGAGGAACGTGGG |
| 2_hypothetical | C8 | Tb927.11.6520_for | GGGGACAAGTTTGTACAAAAAAGCAGGCTATGACGACTTTAGAAAAGCGGATCG | Tb927.11.6520_rev | GGGGACCACTTTGTACAAGAAAGCTGGGTTCACTCTTTCTTGACCTCATCGAATG |
| 2_hypothetical | C9 | Tb927.2.3090_for | GGGGACAAGTTTGTACAAAAAAGCAGGCTATGAGGGAGGGGGATCGAACC | Tb927.2.3090_rev | GGGGACCACTTTGTACAAGAAAGCTGGGTTCAGACTTTGCTCCACCCGC |
| 2_hypothetical | C10 | Tb927.4.4660_for | GGGGACAAGTTTGTACAAAAAAGCAGGCTATGATCATTTGGGCAGTAGGGACA | Tb927.4.4660_rev | GGGGACCACTTTGTACAAGAAAGCTGGGTTTAATTGTCCCAATACGTCCGACTTG |
| 2_hypothetical | C11 | Tb927.1.3800_for | GGGGACAAGTTTGTACAAAAAAGCAGGCTATGCGATGCACGCATGTTCG | Tb927.1.3800_rev | GGGGACCACTTTGTACAAGAAAGCTGGGTCTAACGATCGTAACGATCAGGAATGC |
| 2_hypothetical | C12 | Tb927.10.9170_for | GGGGACAAGTTTGTACAAAAAAGCAGGCTATGACGGCGCCTCTTTTCAG | Tb927.10.9170_rev | GGGGACCACTTTGTACAAGAAAGCTGGGTTCATAGAAGCCTCTGCACCATTTCC |
| 2_hypothetical | C13 | Tb927.10.13140_for | GGGGACAAGTTTGTACAAAAAAGCAGGCTATGGGCAAACGCCGAGTTG | Tb927.10.13140_rev | GGGGACCACTTTGTACAAGAAAGCTGGGTCTATGTGGTCAAGTGTGCAAACATCT |
| 2_hypothetical | C14 | Tb927.3.770_for | GGGGACAAGTTTGTACAAAAAAGCAGGCTATGGTGTTGCGCAGGTGG | Tb927.3.770_rev | GGGGACCACTTTGTACAAGAAAGCTGGGTCTAATCCTCCTCCTCCTCCGTTG |
| 2_hypothetical | C15 | Tb927.10.9790_for | GGGGACAAGTTTGTACAAAAAAGCAGGCTATGAGACTTATTTCTTTAGCAACGAGAACT | Tb927.10.9790_rev | GGGGACCACTTTGTACAAGAAAGCTGGGTTTATCGCACGCCACTCTCGT |
| 2_hypothetical | C16 | Tb927.4.4780_for | GGGGACAAGTTTGTACAAAAAAGCAGGCTATGATATGTGTAGTGCACCACACG | Tb927.4.4780_rev | GGGGACCACTTTGTACAAGAAAGCTGGGTCTACCCGAGCCGCAACAAC |
| 2_hypothetical | C17 | Tb927.11.11910_for | GGGGACAAGTTTGTACAAAAAAGCAGGCTATGGATAGTGGGACCAAAAGGGAA | Tb927.11.11910_rev | GGGGACCACTTTGTACAAGAAAGCTGGGTTTACAAGATAAGTGGAATTACTGGTGGTG |
| 2_hypothetical | C18 | Tb927.9.10830_for | GGGGACAAGTTTGTACAAAAAAGCAGGCTATGGACACGATCGAATTCCTCCA | Tb927.9.10830_rev | GGGGACCACTTTGTACAAGAAAGCTGGGTTTAACTCTTCCGAGAGGGTCGTG |
| 2_hypothetical | C19 | Tb927.8.8150_for | GGGGACAAGTTTGTACAAAAAAGCAGGCTATGGGGAAGCTTCAGGTTTGTATTTG | Tb927.8.8150_rev | GGGGACCACTTTGTACAAGAAAGCTGGGTTTATGCTGAGGCTACAGCTTCAGG |
| 2_hypothetical | C20 | Tb927.11.4720_for | GGGGACAAGTTTGTACAAAAAAGCAGGCTATGCCCGACTCGCCGACA | Tb927.11.4720_rev | GGGGACCACTTTGTACAAGAAAGCTGGGTTTATCTAGAGTCGTCTCTTGGGTGC |
| 2_hypothetical | C21 | Tb927.3.4880_for | GGGGACAAGTTTGTACAAAAAAGCAGGCTATGAGCTCCATACCGGTGGAC | Tb927.3.4880_rev | GGGGACCACTTTGTACAAGAAAGCTGGGTTTATTCAAAGGCTAGAAGAGGGTTTTGG |
| 2_hypothetical | C22 | Tb927.10.10290_for | GGGGACAAGTTTGTACAAAAAAGCAGGCTATGCACACCGGGGACAAGC | Tb927.10.10290_rev | GGGGACCACTTTGTACAAGAAAGCTGGGTTTACATTTGGGTTCTCGCGCC |
| 2_hypothetical | C23 | Tb927.5.2110_for | GGGGACAAGTTTGTACAAAAAAGCAGGCTATGTCATCGATTGCCAAACGAATGC | Tb927.5.2110_rev | GGGGACCACTTTGTACAAGAAAGCTGGGTTCAGTGGCCGCGGTGTGA |
| 2_hypothetical | C24 | Tb927.10.10370_for | GGGGACAAGTTTGTACAAAAAAGCAGGCTATGCACACCGGGGACAAGC | Tb927.10.10370_rev | GGGGACCACTTTGTACAAGAAAGCTGGGTTTACATTTGGGTTCTCGTGCCAC |
| 2_hypothetical | D1 | Tb927.7.520_for | GGGGACAAGTTTGTACAAAAAAGCAGGCTATGTCTTTGCCTGACGGAGGA | Tb927.7.520_rev | GGGGACCACTTTGTACAAGAAAGCTGGGTTTAGCCGGGGCCTTTTTCTTTT |
| 2_hypothetical | D2 | Tb927.9.12080_for | GGGGACAAGTTTGTACAAAAAAGCAGGCTATGAGTTACCTACCCGACGCG | Tb927.9.12080_rev | GGGGACCACTTTGTACAAGAAAGCTGGGTCTACTGATTTTCAGTCACACCAGTTGA |
| 2_hypothetical | D3 | Tb927.11.1690_for | GGGGACAAGTTTGTACAAAAAAGCAGGCTATGACGACGATGGAGAAACCGC | Tb927.11.1690_rev | GGGGACCACTTTGTACAAGAAAGCTGGGTTTATTTATTTTTTCTCTTTTTTGGCGAGTT |
| 2_hypothetical | D4 | Tb927.11.11930.2_for | GGGGACAAGTTTGTACAAAAAAGCAGGCTATGCGCAGCAGGAACCTG | Tb927.11.11930.2_rev | GGGGACCACTTTGTACAAGAAAGCTGGGTTTACTTCTCGGATTGAAAACGAATAGCT |
| 2_hypothetical | D5 | Tb927.10.6470_for | GGGGACAAGTTTGTACAAAAAAGCAGGCTATGCACTTGAAGGAGTGTCCACC | Tb927.10.6470_rev | GGGGACCACTTTGTACAAGAAAGCTGGGTTTACAACGTTGTAAGGGCAACAAGA |
| 2_hypothetical | D6 | Tb927.4.3480_for | GGGGACAAGTTTGTACAAAAAAGCAGGCTATGATCTTTTTCGGCGTACTCTCTTT | Tb927.4.3480_rev | GGGGACCACTTTGTACAAGAAAGCTGGGTCTACAGTTCCTCTAAATACATACACATCTC |
| 2_hypothetical | D7 | Tb927.10.8620_for | GGGGACAAGTTTGTACAAAAAAGCAGGCTATGGTATTGCACAAGTGGGCTG | Tb927.10.8620_rev | GGGGACCACTTTGTACAAGAAAGCTGGGTCTACCCTCCCCCCGAGGT |
| 2_hypothetical | D8 | Tb927.7.6030_for | GGGGACAAGTTTGTACAAAAAAGCAGGCTATGGGCCCACCTCCAAACC | Tb927.7.6030_rev | GGGGACCACTTTGTACAAGAAAGCTGGGTTCACAGCACCCCCGCCCC |
| 2_hypothetical | D9 | Tb927.8.4550_for | GGGGACAAGTTTGTACAAAAAAGCAGGCTATGCGGCGTGTTAGGGATGG | Tb927.8.4550_rev | GGGGACCACTTTGTACAAGAAAGCTGGGTCTACAGGTACTTATCTAACGTCATCGC |
| 2_hypothetical | D11 | Tb927.8.1230_for | GGGGACAAGTTTGTACAAAAAAGCAGGCTATGTCGCTTCACTGCTTCTTTGG | Tb927.8.1230_rev | GGGGACCACTTTGTACAAGAAAGCTGGGTTTAATTTCCCTTCCCTTTCTTCTCCGC |
| 2_hypothetical | D12 | Tb927.4.3330_for | GGGGACAAGTTTGTACAAAAAAGCAGGCTATGCCTGCTTCAGCCTCTACG | Tb927.4.3330_rev | GGGGACCACTTTGTACAAGAAAGCTGGGTCTAACGATGATTTGAGAGAAAGTTCATGAA |
| 2_hypothetical | D13 | Tb927.11.16350_for | GGGGACAAGTTTGTACAAAAAAGCAGGCTATGGGTATACTTCAGCACCAGGC | Tb927.11.16350_rev | GGGGACCACTTTGTACAAGAAAGCTGGGTTTACGAGTACTTATCCACCATGGCG |
| 2_hypothetical | D14 | Tb927.4.5090_for | GGGGACAAGTTTGTACAAAAAAGCAGGCTATGTACTCTTTTTGGGCCTCCCC | Tb927.4.5090_rev | GGGGACCACTTTGTACAAGAAAGCTGGGTTTAGCCGTAGCCGCGCCG |
| 2_hypothetical | D15 | Tb927.9.7250_for | GGGGACAAGTTTGTACAAAAAAGCAGGCTATGAAAAACAAAGCGGCATCTCG | Tb927.9.7250_rev | GGGGACCACTTTGTACAAGAAAGCTGGGTTTACAAACCTTTGCCTATCTTTTGCAAC |
| 2_hypothetical | D16 | Tb927.6.3080_for | GGGGACAAGTTTGTACAAAAAAGCAGGCTATGCCCGTTACTTCTTTCAATAACCG | Tb927.6.3080_rev | GGGGACCACTTTGTACAAGAAAGCTGGGTTCACTCTCCACTTTCCCTTTTAGAACC |
| 2_hypothetical | D17 | Tb927.7.7020_for | GGGGACAAGTTTGTACAAAAAAGCAGGCTATGGAGGGAAGAAGAAGCGTAAGG | Tb927.7.7020_rev | GGGGACCACTTTGTACAAGAAAGCTGGGTTTATACAACCGCGGCCAAAAAC |
| 2_hypothetical | D18 | Tb927.4.4640_for | GGGGACAAGTTTGTACAAAAAAGCAGGCTATGCGTGGAACAACAAAGCCG | Tb927.4.4640_rev | GGGGACCACTTTGTACAAGAAAGCTGGGTTTACGTAGGAAATGGAAATCTAGTTTTCCA |
| 2_hypothetical | D19 | Tb927.5.2830_for | GGGGACAAGTTTGTACAAAAAAGCAGGCTATGCGGTTCGTTTCTCGCAG | Tb927.5.2830_rev | GGGGACCACTTTGTACAAGAAAGCTGGGTCTACACAAGTGTATAACCCTTCACCTTG |
| 2_hypothetical | D20 | Tb927.4.1210_for | GGGGACAAGTTTGTACAAAAAAGCAGGCTATGCGAGAGCCTCCCTACG | Tb927.4.1210_rev | GGGGACCACTTTGTACAAGAAAGCTGGGTTTAAGAAGTGAGAGTGAATAAAGGAGACCT |
| 2_hypothetical | D21 | Tb927.8.5520_for | GGGGACAAGTTTGTACAAAAAAGCAGGCTATGCCGCATATTACTACGCCCT | Tb927.8.5520_rev | GGGGACCACTTTGTACAAGAAAGCTGGGTTCATTCGGTGGTAGACCGCA |
| 2_hypothetical | D22 | Tb927.11.8250_for | GGGGACAAGTTTGTACAAAAAAGCAGGCTATGAGCCACCCAGATGGTTCC | Tb927.11.8250_rev | GGGGACCACTTTGTACAAGAAAGCTGGGTTCACACCTCCACCTCGGC |
| 2_hypothetical | D23 | Tb927.4.4390_for | GGGGACAAGTTTGTACAAAAAAGCAGGCTATGGAGGACGTAAATAGGCAGAATCA | Tb927.4.4390_rev | GGGGACCACTTTGTACAAGAAAGCTGGGTTCAACGTAACCCAATGATTGTTTCATTC |
| 2_hypothetical | D24 | Tb927.11.300_for | GGGGACAAGTTTGTACAAAAAAGCAGGCTATGAAACGTCTACGCCGCTTC | Tb927.11.300_rev | GGGGACCACTTTGTACAAGAAAGCTGGGTTCAATTTGTGGTGCTAACGCCA |
| 2_hypothetical | E1 | Tb927.11.8040_for | GGGGACAAGTTTGTACAAAAAAGCAGGCTATGTTGAATCCGACATTTTCTCTGTACA | Tb927.11.8040_rev | GGGGACCACTTTGTACAAGAAAGCTGGGTCTAACCCCATAACTCCGCTGC |
| 2_hypothetical | E2 | Tb927.10.5090_for | GGGGACAAGTTTGTACAAAAAAGCAGGCTATGGGTCGCAGGCATGGA | Tb927.10.5090_rev | GGGGACCACTTTGTACAAGAAAGCTGGGTTTAGGCGGATGAGTCTTGTGATACG |
| 2_hypothetical | E3 | Tb927.11.13710_for | GGGGACAAGTTTGTACAAAAAAGCAGGCTATGGGTCACCTTCAGAGCACC | Tb927.11.13710_rev | GGGGACCACTTTGTACAAGAAAGCTGGGTTCATCTTTCTTTGTTGAGAAACTCTGCC |
| 2_hypothetical | E4 | Tb927.11.3640_for | GGGGACAAGTTTGTACAAAAAAGCAGGCTATGTTAAAGTGCTCCTTGTGCTATCTAG | Tb927.11.3640_rev | GGGGACCACTTTGTACAAGAAAGCTGGGTTCATGTGTAACTGTACCTATCCAAGGG |
| 2_hypothetical | E5 | Tb927.10.5630_for | GGGGACAAGTTTGTACAAAAAAGCAGGCTATGGATGATGGTGTAGAAGCCAAGC | Tb927.10.5630_rev | GGGGACCACTTTGTACAAGAAAGCTGGGTTTATTCATGTCGACGTGACAAGCG |
| 2_hypothetical | E6 | Tb927.7.3140_for | GGGGACAAGTTTGTACAAAAAAGCAGGCTATGAGCACTTTATCATCGTGGCTC | Tb927.7.3140_rev | GGGGACCACTTTGTACAAGAAAGCTGGGTTTACGCTTCGAGAAGCATCTGGT |
| 2_hypothetical | E7 | Tb927.11.9940_for | GGGGACAAGTTTGTACAAAAAAGCAGGCTATGACGGAGGGTGTTGGGC | Tb927.11.9940_rev | GGGGACCACTTTGTACAAGAAAGCTGGGTCTAGGCTATGGCACCCGCA |
| 2_hypothetical | E8 | Tb927.8.2360_for | GGGGACAAGTTTGTACAAAAAAGCAGGCTATGACTGACCCCTTCTATAGGTTACG | Tb927.8.2360_rev | GGGGACCACTTTGTACAAGAAAGCTGGGTTTAGAAGGGCAAAAAAACTTGCCG |
| 2_hypothetical | E9 | Tb927.8.2010_for | GGGGACAAGTTTGTACAAAAAAGCAGGCTATGCTGAGGGGAAGGTGGG | Tb927.8.2010_rev | GGGGACCACTTTGTACAAGAAAGCTGGGTTCACCGCTTCTGAAGAACAAGGT |
| 2_hypothetical | E10 | Tb927.8.2080_for | GGGGACAAGTTTGTACAAAAAAGCAGGCTATGATGGGTTGTGGAGGTTCTAAATTG | Tb927.8.2080_rev | GGGGACCACTTTGTACAAGAAAGCTGGGTTCACCGCACAAACTGAAACACA |
| 2_hypothetical | E11 | Tb927.9.700_for | GGGGACAAGTTTGTACAAAAAAGCAGGCTATGTATAATTTGTACATTTACTTTGTGCGA | Tb927.9.700_rev | GGGGACCACTTTGTACAAGAAAGCTGGGTTTATCCCTGTGTGACTAACCGAAAGT |
| 2_hypothetical | E12 | Tb927.11.9770_for | GGGGACAAGTTTGTACAAAAAAGCAGGCTATGGCGATCGCTAGGACCAC | Tb927.11.9770_rev | GGGGACCACTTTGTACAAGAAAGCTGGGTTCATCCCAAGCATGACGGGG |
| 2_hypothetical | E13 | Tb927.11.16220_for | GGGGACAAGTTTGTACAAAAAAGCAGGCTATGCGCCGTAATCCGTCCC | Tb927.11.16220_rev | GGGGACCACTTTGTACAAGAAAGCTGGGTTTATGCTCCATCATTGTCCGTGC |
| 2_hypothetical | E14 | Tb927.11.9970_for | GGGGACAAGTTTGTACAAAAAAGCAGGCTATGAGCTCCCTTAAGTCGGTAGC | Tb927.11.9970_rev | GGGGACCACTTTGTACAAGAAAGCTGGGTTCACGCGCCAAGACCCTT |
| 2_hypothetical | E15 | Tb927.11.570_for | GGGGACAAGTTTGTACAAAAAAGCAGGCTATGAATCGGTCGTGTTCCCGT | Tb927.11.570_rev | GGGGACCACTTTGTACAAGAAAGCTGGGTTCAACGTGTCGCAGCGCG |
| 2_hypothetical | E16 | Tb927.10.11260_for | GGGGACAAGTTTGTACAAAAAAGCAGGCTATGCTGAAAAGAACGTTAACTGTTTTAGAT | Tb927.10.11260_rev | GGGGACCACTTTGTACAAGAAAGCTGGGTTCACACTGTCTTTTTACTCTGTCGTC |
| 2_hypothetical | E17 | Tb927.11.14320_for | GGGGACAAGTTTGTACAAAAAAGCAGGCTATGACCGGTTTTGACACTGTCTG | Tb927.11.14320_rev | GGGGACCACTTTGTACAAGAAAGCTGGGTCTATTGCGCCTGCACCTCC |
| 2_hypothetical | E18 | Tb927.11.2480_for | GGGGACAAGTTTGTACAAAAAAGCAGGCTATGTGTGGGGCTCATTCGTGC | Tb927.11.2480_rev | GGGGACCACTTTGTACAAGAAAGCTGGGTTTATGAAAACGTTGCGAGCTCGT |
| 2_hypothetical | E19 | Tb927.11.3370_for | GGGGACAAGTTTGTACAAAAAAGCAGGCTATGCTCAACATCACCAGACTGC | Tb927.11.3370_rev | GGGGACCACTTTGTACAAGAAAGCTGGGTCTACCCAATGTTTTCAATACTTTGGCTATC |
| 2_hypothetical | E20 | Tb927.6.4240_for | GGGGACAAGTTTGTACAAAAAAGCAGGCTATGGTTTCACTGATGAAGTTGGCC | Tb927.6.4240_rev | GGGGACCACTTTGTACAAGAAAGCTGGGTTCACCTCACGTCCAGAAAATACATATG |
| 2_hypothetical | E21 | Tb927.4.2265_for | GGGGACAAGTTTGTACAAAAAAGCAGGCTATGACGCAAAACAAGGTGAGGAA | Tb927.4.2265_rev | GGGGACCACTTTGTACAAGAAAGCTGGGTTCATGAAGCGGCTGGTTTGAC |
| 2_hypothetical | E22 | Tb927.4.850_for | GGGGACAAGTTTGTACAAAAAAGCAGGCTATGAGAAATAATATAGAGGCGGCAGGT | Tb927.4.850_rev | GGGGACCACTTTGTACAAGAAAGCTGGGTTTACGGTTTTCCTTTAGACAATTCATATGC |
| 2_hypothetical | E23 | Tb927.10.8260_for | GGGGACAAGTTTGTACAAAAAAGCAGGCTATGCGTCAGTCATCTTTGCCTG | Tb927.10.8260_rev | GGGGACCACTTTGTACAAGAAAGCTGGGTTCACCTGTAAAAGGAGAGCACGA |
| 2_hypothetical | E24 | Tb927.6.2260_for | GGGGACAAGTTTGTACAAAAAAGCAGGCTATGCGGGCGTGGAGTAACC | Tb927.6.2260_rev | GGGGACCACTTTGTACAAGAAAGCTGGGTTTACCGATTCACTGCCACAGC |
| 2_hypothetical | F1 | Tb927.10.2660_for | GGGGACAAGTTTGTACAAAAAAGCAGGCTATGAAAAGGGATACAAACAACCACAATG | Tb927.10.2660_rev | GGGGACCACTTTGTACAAGAAAGCTGGGTTCACCACAGGGACGTGGG |
| 2_hypothetical | F2 | Tb927.5.1220_for | GGGGACAAGTTTGTACAAAAAAGCAGGCTATGATGCCCGTCGCGGGT | Tb927.5.1220_rev | GGGGACCACTTTGTACAAGAAAGCTGGGTTCAGATGCTCACCTTCTGTAACAGC |
| 2_hypothetical | F3 | Tb927.10.4660_for | GGGGACAAGTTTGTACAAAAAAGCAGGCTATGGCTGCATCAGAGGCAATG | Tb927.10.4660_rev | GGGGACCACTTTGTACAAGAAAGCTGGGTTCAACCCTTCAACGATTTTTCACTGC |
| 2_hypothetical | F4 | Tb927.7.4140_for | GGGGACAAGTTTGTACAAAAAAGCAGGCTATGCCCTCACTAGCGAACGG | Tb927.7.4140_rev | GGGGACCACTTTGTACAAGAAAGCTGGGTTCACACACCACCCAAAAGGTCG |
| 2_hypothetical | F6 | Tb927.3.820_for | GGGGACAAGTTTGTACAAAAAAGCAGGCTATGTCGGGGGCGTTCGGT | Tb927.3.820_rev | GGGGACCACTTTGTACAAGAAAGCTGGGTCTAGTAGTCGCTCGCTTGTACAGC |
| 2_hypothetical | F7 | Tb927.6.1300_for | GGGGACAAGTTTGTACAAAAAAGCAGGCTATGATGTGGAACGTGGCGC | Tb927.6.1300_rev | GGGGACCACTTTGTACAAGAAAGCTGGGTTCACAGAAACATGGAAAAATTCATTTGAGA |
| 2_hypothetical | F8 | Tb927.10.5060_for | GGGGACAAGTTTGTACAAAAAAGCAGGCTATGCCGTGGTATTTAAGCACTTTCA | Tb927.10.5060_rev | GGGGACCACTTTGTACAAGAAAGCTGGGTTCAAATGGCCCCTGTTCCTTTTC |
| 2_hypothetical | F9 | Tb927.11.7790_for | GGGGACAAGTTTGTACAAAAAAGCAGGCTATGCAAGTGACAGAAACGGTGC | Tb927.11.7790_rev | GGGGACCACTTTGTACAAGAAAGCTGGGTCTAAAGATACGGCGCCACCG |
| 2_hypothetical | F10 | Tb927.11.17016_for | GGGGACAAGTTTGTACAAAAAAGCAGGCTATGCCGATAATACGTAAGGTTCAATGT | Tb927.11.17016_rev | GGGGACCACTTTGTACAAGAAAGCTGGGTTTATCCTTTTTTGTTCGCACTGCAC |
| 2_hypothetical | F11 | Tb927.11.11300_for | GGGGACAAGTTTGTACAAAAAAGCAGGCTATGGTGTGGAAAGGATCACGTGA | Tb927.11.11300_rev | GGGGACCACTTTGTACAAGAAAGCTGGGTCTAAGCGACACGTTGGCGT |
| 2_hypothetical | F12 | Tb927.11.10080_for | GGGGACAAGTTTGTACAAAAAAGCAGGCTATGTTGCGTCTCTGCCGTG | Tb927.11.10080_rev | GGGGACCACTTTGTACAAGAAAGCTGGGTCTATGAGGGCCTTCCGTATATATTCGA |
| 2_hypothetical | F13 | Tb927.11.10410_for | GGGGACAAGTTTGTACAAAAAAGCAGGCTATGAGCATGCCATGCAATATTCGT | Tb927.11.10410_rev | GGGGACCACTTTGTACAAGAAAGCTGGGTTTAGTTTGGTAACAGAGCGAGCG |
| 2_hypothetical | F14 | Tb927.6.2070_for | GGGGACAAGTTTGTACAAAAAAGCAGGCTATGTGGCGGTTGTCGCGG | Tb927.6.2070_rev | GGGGACCACTTTGTACAAGAAAGCTGGGTTTACCATGGGTCCGTCCGC |

| | | | | | |
|---|---|---|---|---|---|
| 2_hypothetical | F15 | Tb927.11.13870_for | GGGGACAAGTTTGTACAAAAAAGCAGGCTATGGTCAACGCGACTCCCG | Tb927.11.13870_rev | GGGGACCACTTTGTACAAGAAAGCTGGGTTCAGCAAGGTAGTGAACCAAGGT |
| 2_hypothetical | F16 | Tb927.10.14760_for | GGGGACAAGTTTGTACAAAAAAGCAGGCTATGGACTTCCTAAACGAGAGTCAAAC | Tb927.10.14760_rev | GGGGACCACTTTGTACAAGAAAGCTGGGTCTATCCACTCCCTGACTGTGACA |
| 2_hypothetical | F17 | Tb927.11.6080_for | GGGGACAAGTTTGTACAAAAAAGCAGGCTATGGATAAAGATGTGGATGTGACAAAGT | Tb927.11.6080_rev | GGGGACCACTTTGTACAAGAAAGCTGGGTTCATTGTACACCTCCAGCAGACC |
| 2_hypothetical | F18 | Tb927.8.7040_for | GGGGACAAGTTTGTACAAAAAAGCAGGCTATGCGTAAGTCATTTCTTGGAGCA | Tb927.8.7040_rev | GGGGACCACTTTGTACAAGAAAGCTGGGTTCATTTCGCTGCTGCTCGC |
| 2_hypothetical | F19 | Tb927.11.4330_for | GGGGACAAGTTTGTACAAAAAAGCAGGCTATGGAGCAACCCTCCCAGATG | Tb927.11.4330_rev | GGGGACCACTTTGTACAAGAAAGCTGGGTCTAAACCATGCCCTCCGCAG |
| 2_hypothetical | F20 | Tb927.10.2360_for | GGGGACAAGTTTGTACAAAAAAGCAGGCTATGTATTTTGGTTTCGCTCATTTGCC | Tb927.10.2360_rev | GGGGACCACTTTGTACAAGAAAGCTGGGTTTACTTTATGTGGTGGAGGCGC |
| 2_hypothetical | F21 | Tb927.10.7910_for | GGGGACAAGTTTGTACAAAAAAGCAGGCTATGTTTTCCAGCGTATTACTGCGC | Tb927.10.7910_rev | GGGGACCACTTTGTACAAGAAAGCTGGGTTTACTTCCATACAAAAGTTCCCTGCG |
| 2_hypothetical | F22 | Tb927.8.2190_for | GGGGACAAGTTTGTACAAAAAAGCAGGCTATGAGGCAATGCAGCAGAACAA | Tb927.8.2190_rev | GGGGACCACTTTGTACAAGAAAGCTGGGTTCATCCTCTTGTACTCCCAACTGTT |
| 2_hypothetical | F23 | Tb927.6.1700_for | GGGGACAAGTTTGTACAAAAAAGCAGGCTATGTTGCGGTCACGTATCCCA | Tb927.6.1700_rev | GGGGACCACTTTGTACAAGAAAGCTGGGTCTAACTGTTTGGGTCGAACAAAAACTC |
| 2_hypothetical | F24 | Tb927.8.1880_for | GGGGACAAGTTTGTACAAAAAAGCAGGCTATGCTGCAAATAAACGCGTTCAA | Tb927.8.1880_rev | GGGGACCACTTTGTACAAGAAAGCTGGGTTCACTTCTTTCCACCCCCATCC |
| 2_hypothetical | G1 | Tb927.2.2200_for | GGGGACAAGTTTGTACAAAAAAGCAGGCTATGCCTGCCCCCAGTTTAGC | Tb927.2.2200_rev | GGGGACCACTTTGTACAAGAAAGCTGGGTTCAGGCTGCTGTGGCTTTTC |
| 2_hypothetical | G2 | Tb927.1.1090_for | GGGGACAAGTTTGTACAAAAAAGCAGGCTATGGTTTGGTTCTCCGCGC | Tb927.1.1090_rev | GGGGACCACTTTGTACAAGAAAGCTGGGTTTATATAATTGCATCCCCCTCGTGC |
| 2_hypothetical | G3 | Tb927.7.2980_for | GGGGACAAGTTTGTACAAAAAAGCAGGCTATGCGCCGGTCTGTATATCAGC | Tb927.7.2980_rev | GGGGACCACTTTGTACAAGAAAGCTGGGTTTATTTAAACACCTTAAAGCGTTCTTCATC |
| 2_hypothetical | G5 | Tb927.7.4630_for | GGGGACAAGTTTGTACAAAAAAGCAGGCTATGTCAGGGCGACTTATCATTTATGTG | Tb927.7.4630_rev | GGGGACCACTTTGTACAAGAAAGCTGGGTCTACGGCGGGAACGCAAA |
| 2_hypothetical | G6 | Tb927.11.16440_for | GGGGACAAGTTTGTACAAAAAAGCAGGCTATGGATGCGTTTTTGGAGTCGC | Tb927.11.16440_rev | GGGGACCACTTTGTACAAGAAAGCTGGGTTCATGTTTTTTTCTTGCTTGAAGCTGC |
| 2_hypothetical | G7 | Tb927.5.1940_for | GGGGACAAGTTTGTACAAAAAAGCAGGCTATGCCAAAAAAGAAGCCACCATGT | Tb927.5.1940_rev | GGGGACCACTTTGTACAAGAAAGCTGGGTTCACGGTTTCCCACCGCA |
| 2_hypothetical | G8 | Tb927.8.4290_for | GGGGACAAGTTTGTACAAAAAAGCAGGCTATGGGGCGAATTAGGGTGACA | Tb927.8.4290_rev | GGGGACCACTTTGTACAAGAAAGCTGGGTTCATTCCTTTCTGTGCCGGTTC |
| 2_hypothetical | G9 | Tb927.10.1880_for | GGGGACAAGTTTGTACAAAAAAGCAGGCTATGACCACGCCTGGTACTTCA | Tb927.10.1880_rev | GGGGACCACTTTGTACAAGAAAGCTGGGTTCAGTTGCGCCCCTCCGT |
| 2_hypothetical | G10 | Tb927.10.16010_for | GGGGACAAGTTTGTACAAAAAAGCAGGCTATGTGCAGCCGGCGCCCT | Tb927.10.16010_rev | GGGGACCACTTTGTACAAGAAAGCTGGGTCTATTCCGTCAGGCTGACGC |
| 2_hypothetical | G11 | Tb927.9.7710_for | GGGGACAAGTTTGTACAAAAAAGCAGGCTATGAGTGCCTTTGAGTCATTCCTCC | Tb927.9.7710_rev | GGGGACCACTTTGTACAAGAAAGCTGGGTCTACACACGAGGGGACACCT |
| 2_hypothetical | G12 | Tb927.5.4120_for | GGGGACAAGTTTGTACAAAAAAGCAGGCTATGAGGCGATGTATCCCCGC | Tb927.5.4120_rev | GGGGACCACTTTGTACAAGAAAGCTGGGTTCAAGCTGCTCCGTTCCCC |
| 2_hypothetical | G13 | Tb927.10.15620_for | GGGGACAAGTTTGTACAAAAAAGCAGGCTATGGTAAAACACCAAAATGGCGC | Tb927.10.15620_rev | GGGGACCACTTTGTACAAGAAAGCTGGGTTTATATATATATATATATATATATATAT |
| 2_hypothetical | G14 | Tb927.9.10300_for | GGGGACAAGTTTGTACAAAAAAGCAGGCTATGAGGGAACCCCCCACTACG | Tb927.9.10300_rev | GGGGACCACTTTGTACAAGAAAGCTGGGTTTATGATGACGAAGGCACTTGCAT |
| 2_hypothetical | G15 | Tb927.10.7870_for | GGGGACAAGTTTGTACAAAAAAGCAGGCTATGCCTGGCATTGCGCAA | Tb927.10.7870_rev | GGGGACCACTTTGTACAAGAAAGCTGGGTCTACGCAAATCTATCGGCAACCT |
| 2_hypothetical | G16 | Tb927.11.10620_for | GGGGACAAGTTTGTACAAAAAAGCAGGCTATGGAGGACGAACCGACTCAA | Tb927.11.10620_rev | GGGGACCACTTTGTACAAGAAAGCTGGGTTTAGTCATCGGTAACACTCTTGCCA |
| 2_hypothetical | G17 | Tb927.5.750_for | GGGGACAAGTTTGTACAAAAAAGCAGGCTATGGCTTTTCCTCATGTGAGTTATCG | Tb927.5.750_rev | GGGGACCACTTTGTACAAGAAAGCTGGGTCTACTCTAACATGTACTGACACAGGCA |
| 2_hypothetical | G18 | Tb927.7.7460_for | GGGGACAAGTTTGTACAAAAAAGCAGGCTATGGTTTTTTCCACACACGGCA | Tb927.7.7460_rev | GGGGACCACTTTGTACAAGAAAGCTGGGTTTACCTCCTCACCGTTGGTTTGA |
| 2_hypothetical | G19 | Tb927.10.8380_for | GGGGACAAGTTTGTACAAAAAAGCAGGCTATGGATACGCCCCCGATAAGTG | Tb927.10.8380_rev | GGGGACCACTTTGTACAAGAAAGCTGGGTCTAGCAGCGACTTTGCGCA |
| 2_hypothetical | G20 | Tb927.9.14230_for | GGGGACAAGTTTGTACAAAAAAGCAGGCTATGTGGAAGCAGCGTCGGA | Tb927.9.14230_rev | GGGGACCACTTTGTACAAGAAAGCTGGGTCTAACCCCTGAGGCCGCC |
| 2_hypothetical | G21 | Tb927.11.1620_for | GGGGACAAGTTTGTACAAAAAAGCAGGCTATGTGGGCGCAACAGCTTAG | Tb927.11.1620_rev | GGGGACCACTTTGTACAAGAAAGCTGGGTTTATTCCTTCCCGACGGCTTGC |
| 2_hypothetical | G22 | Tb927.11.7440_for | GGGGACAAGTTTGTACAAAAAAGCAGGCTATGAAAAGGGCAAAGGCACTTGT | Tb927.11.7440_rev | GGGGACCACTTTGTACAAGAAAGCTGGGTTTACATATCTAGGAGGACGCTCAACG |
| 2_hypothetical | G23 | Tb927.10.2620_for | GGGGACAAGTTTGTACAAAAAAGCAGGCTATGACGAATCAGTCGGACGCG | Tb927.10.2620_rev | GGGGACCACTTTGTACAAGAAAGCTGGGTTTACGGTTCGAGAGGAGGCAG |
| 2_hypothetical | G24 | Tb927.5.2010_for | GGGGACAAGTTTGTACAAAAAAGCAGGCTATGTATGTTGTTTCACTTAAAATATATATT | Tb927.5.2010_rev | GGGGACCACTTTGTACAAGAAAGCTGGGTCTAAAGGAAACGACACACACAAAACC |
| 2_hypothetical | H1 | Tb927.11.3180_for | GGGGACAAGTTTGTACAAAAAAGCAGGCTATGGGTGACTTTACCCGAGAAGA | Tb927.11.3180_rev | GGGGACCACTTTGTACAAGAAAGCTGGGTTTACATCTCAAGAACCTTCATTAGCAACA |
| 2_hypothetical | H2 | Tb927.10.2300_for | GGGGACAAGTTTGTACAAAAAAGCAGGCTATGTTTTCGAGACACTCCTTGTGG | Tb927.10.2300_rev | GGGGACCACTTTGTACAAGAAAGCTGGGTTCACGATTTTCCAGTCAAAATGCAC |
| 2_hypothetical | H3 | Tb927.7.5260_for | GGGGACAAGTTTGTACAAAAAAGCAGGCTATGGGTACCACCGCCAGTAC | Tb927.7.5260_rev | GGGGACCACTTTGTACAAGAAAGCTGGGTCTAAGCGTCCTTTACCCATGGGT |
| 2_hypothetical | H4 | Tb927.7.330_for | GGGGACAAGTTTGTACAAAAAAGCAGGCTATGGACTACAACGAAATAAGCCCAAG | Tb927.7.330_rev | GGGGACCACTTTGTACAAGAAAGCTGGGTCTACAGTTTTGAGATGGCCGCA |
| 2_hypothetical | H5 | Tb927.11.12300_for | GGGGACAAGTTTGTACAAAAAAGCAGGCTATGACATTTGTGTTCGAGGCTAGAAG | Tb927.11.12300_rev | GGGGACCACTTTGTACAAGAAAGCTGGGTTTATTGACCTGAAGGATGAAGGTGGT |
| 2_hypothetical | H6 | Tb927.7.4470_for | GGGGACAAGTTTGTACAAAAAAGCAGGCTATGACACGAGGACAAAACGATACAC | Tb927.7.4470_rev | GGGGACCACTTTGTACAAGAAAGCTGGGTTTATTTGTGCCTCGCCATGCG |
| 2_hypothetical | H7 | Tb927.8.4300_for | GGGGACAAGTTTGTACAAAAAAGCAGGCTATGAATATACATATATATATATATATATTA | Tb927.8.4300_rev | GGGGACCACTTTGTACAAGAAAGCTGGGTTTAAAAAAAAATGAATAATTTCATGCAGCA |
| 2_hypothetical | H8 | Tb927.9.920_for | GGGGACAAGTTTGTACAAAAAAGCAGGCTATGGCCAGAACCGACAAATGTC | Tb927.9.920_rev | GGGGACCACTTTGTACAAGAAAGCTGGGTTCAGCACGCATTTTGTTCCTTATCT |
| 2_hypothetical | H9 | Tb927.3.5010_for | GGGGACAAGTTTGTACAAAAAAGCAGGCTATGGCGGATAATGAAGAAGTGCC | Tb927.3.5010_rev | GGGGACCACTTTGTACAAGAAAGCTGGGTTTATGACCCCCTCCTCGACAGA |
| 2_hypothetical | H10 | Tb927.3.4130_for | GGGGACAAGTTTGTACAAAAAAGCAGGCTATGCTCCACACAACCCGCC | Tb927.3.4130_rev | GGGGACCACTTTGTACAAGAAAGCTGGGTTTAATTCACCCTCCACCGATAGTTGA |
| 2_hypothetical | H11 | Tb927.3.1260_for | GGGGACAAGTTTGTACAAAAAAGCAGGCTATGGAGGAGAAAGTGGTGACTTCG | Tb927.3.1260_rev | GGGGACCACTTTGTACAAGAAAGCTGGGTCTATGAAGAATTGTTCCTTACATCACAGTT |
| 2_hypothetical | H12 | Tb927.11.2840_for | GGGGACAAGTTTGTACAAAAAAGCAGGCTATGAGTACTGAGCCAACCCCTC | Tb927.11.2840_rev | GGGGACCACTTTGTACAAGAAAGCTGGGTTTACAAAGCGCGTTTTTTTTCCTTATCT |
| 2_hypothetical | H13 | Tb927.11.2900_for | GGGGACAAGTTTGTACAAAAAAGCAGGCTATGACAGGGATGCTCCGTTGC | Tb927.11.2900_rev | GGGGACCACTTTGTACAAGAAAGCTGGGTTCAGGAGTTAAAGTTTACATCAACCCTAC |
| 2_hypothetical | H14 | Tb927.10.15220_for | GGGGACAAGTTTGTACAAAAAAGCAGGCTATGTATCGGCTCATGGTTCCGC | Tb927.10.15220_rev | GGGGACCACTTTGTACAAGAAAGCTGGGTTTAGTAATGTTGCAAATCAAGCATGTGTT |
| 2_hypothetical | H15 | Tb927.7.5270_for | GGGGACAAGTTTGTACAAAAAAGCAGGCTATGATTCAGATTATTGCGCCGAGG | Tb927.7.5270_rev | GGGGACCACTTTGTACAAGAAAGCTGGGTTCACACGAACCTACGGAGGG |
| 2_hypothetical | H16 | Tb927.9.2400_for | GGGGACAAGTTTGTACAAAAAAGCAGGCTATGGGGTCCACAGATGGTGAA | Tb927.9.2400_rev | GGGGACCACTTTGTACAAGAAAGCTGGGTTCATCTTAGCAAATAGAGCGATGAAGTT |
| 2_hypothetical | H17 | Tb927.8.3260_for | GGGGACAAGTTTGTACAAAAAAGCAGGCTATGGGTTGGGGAAGCAGTGA | Tb927.8.3260_rev | GGGGACCACTTTGTACAAGAAAGCTGGGTTCAACCTTGCAGTTGAACCGC |
| 2_hypothetical | H18 | Tb927.1.4920_for | GGGGACAAGTTTGTACAAAAAAGCAGGCTATGGCTCCCCGTGGCGGG | Tb927.1.4920_rev | GGGGACCACTTTGTACAAGAAAGCTGGGTCTAAGTCGCTGCAAAATATATCCCCA |
| 2_hypothetical | H19 | Tb927.10.5740_for | GGGGACAAGTTTGTACAAAAAAGCAGGCTATGACAAGGCGGAACGCAAT | Tb927.10.5740_rev | GGGGACCACTTTGTACAAGAAAGCTGGGTTCATTTGCTTCGCGTCAACCC |
| 2_hypothetical | H20 | Tb927.3.4010_for | GGGGACAAGTTTGTACAAAAAAGCAGGCTATGCGGGTGTGTCGTCCC | Tb927.3.4010_rev | GGGGACCACTTTGTACAAGAAAGCTGGGTTCAGGAGTGCCTTAAATAAAGTAAGGC |
| 2_hypothetical | H21 | Tb927.6.830_for | GGGGACAAGTTTGTACAAAAAAGCAGGCTATGACCGCAACGACAACGC | Tb927.6.830_rev | GGGGACCACTTTGTACAAGAAAGCTGGGTTCACACGTTCGGATCAATCGC |
| 2_hypothetical | H22 | Tb927.5.990_for | GGGGACAAGTTTGTACAAAAAAGCAGGCTATGGATGAAAGCAACAGGGTGC | Tb927.5.990_rev | GGGGACCACTTTGTACAAGAAAGCTGGGTTTATCTTTTTCGCACAGCACGCG |
| 2_hypothetical | H23 | Tb927.10.7480_for | GGGGACAAGTTTGTACAAAAAAGCAGGCTATGCATCGCAAACCTCTGTTTGA | Tb927.10.7480_rev | GGGGACCACTTTGTACAAGAAAGCTGGGTTCAGAACAACAGTTCGGATAAAGAAGAG |
| 2_hypothetical | H24 | Tb927.11.13010_for | GGGGACAAGTTTGTACAAAAAAGCAGGCTATGGATAAAGGTATGAAGGATAAGGCTCC | Tb927.11.13010_rev | GGGGACCACTTTGTACAAGAAAGCTGGGTCTACATGAAGTTCAGGCGATGAACC |
| 2_hypothetical | I1 | Tb927.9.4990_for | GGGGACAAGTTTGTACAAAAAAGCAGGCTATGGCGGACGTTGCTGAC | Tb927.9.4990_rev | GGGGACCACTTTGTACAAGAAAGCTGGGTCTACTCCACAAGCTCTACCACCC |
| 2_hypothetical | I2 | Tb927.2.4480_for | GGGGACAAGTTTGTACAAAAAAGCAGGCTATGATATTTGGGTCTGAACCACCAC | Tb927.2.4480_rev | GGGGACCACTTTGTACAAGAAAGCTGGGTTCATTGGTGTGAATGGGAATCCTCC |
| 2_hypothetical | I3 | Tb927.10.11060_for | GGGGACAAGTTTGTACAAAAAAGCAGGCTATGCGTGAGAAGGTGGTGCC | Tb927.10.11060_rev | GGGGACCACTTTGTACAAGAAAGCTGGGTTCAACCAGTGTTACCCGAGTACG |
| 2_hypothetical | I4 | Tb927.3.2970_for | GGGGACAAGTTTGTACAAAAAAGCAGGCTATGAGCGACTCGCACGCC | Tb927.3.2970_rev | GGGGACCACTTTGTACAAGAAAGCTGGGTTCAGTCTCGGCGCCTAGAAA |
| 2_hypothetical | I5 | Tb927.10.7760_for | GGGGACAAGTTTGTACAAAAAAGCAGGCTATGGGCCAAGGCCTCACG | Tb927.10.7760_rev | GGGGACCACTTTGTACAAGAAAGCTGGGTTTAGAAGTCCATCAGCGCGTAAC |
| 2_hypothetical | I6 | Tb927.4.1520_for | GGGGACAAGTTTGTACAAAAAAGCAGGCTATGAACCCCGTCGGAGTGG | Tb927.4.1520_rev | GGGGACCACTTTGTACAAGAAAGCTGGGTTTACTTGTGAAGTCCTTCGGGTTCC |
| 2_hypothetical | I7 | Tb927.4.940_for | GGGGACAAGTTTGTACAAAAAAGCAGGCTATGTCCTTTGCTTGGACGCG | Tb927.4.940_rev | GGGGACCACTTTGTACAAGAAAGCTGGGTCTATGTCATATCTGTCCCAATCTTTCGTT |
| 2_hypothetical | I8 | Tb927.8.5840_for | GGGGACAAGTTTGTACAAAAAAGCAGGCTATGGGGGAACAGGATGGGC | Tb927.8.5840_rev | GGGGACCACTTTGTACAAGAAAGCTGGGTTTATGTATCCGCGCCGTCGC |
| 2_hypothetical | I9 | Tb927.6.2200_for | GGGGACAAGTTTGTACAAAAAAGCAGGCTATGCCCAAGGTTCTCGTTCCC | Tb927.6.2200_rev | GGGGACCACTTTGTACAAGAAAGCTGGGTCTAAAACTTACTGAACATTTGATTAATCAC |
| 2_hypothetical | I10 | Tb927.11.9830_for | GGGGACAAGTTTGTACAAAAAAGCAGGCTATGCTACGCTGTAGTTGTGCG | Tb927.11.9830_rev | GGGGACCACTTTGTACAAGAAAGCTGGGTTCAGCGCCAAAAGAAAAGCCT |
| 2_hypothetical | I11 | Tb927.2.5710_for | GGGGACAAGTTTGTACAAAAAAGCAGGCTATGCCTTGTTCGTTCCAAGAGG | Tb927.2.5710_rev | GGGGACCACTTTGTACAAGAAAGCTGGGTTTAAGGCGACAGGCGATTTACC |
| 2_hypothetical | I12 | Tb927.11.5200_for | GGGGACAAGTTTGTACAAAAAAGCAGGCTATGGACAAGAGTAAAACGATGGAGATC | Tb927.11.5200_rev | GGGGACCACTTTGTACAAGAAAGCTGGGTTCACTCGCCCCGGCCTCC |
| 2_hypothetical | I13 | Tb927.7.4190_for | GGGGACAAGTTTGTACAAAAAAGCAGGCTATGAGTCAGTTTGGTCAACTTGTGT | Tb927.7.4190_rev | GGGGACCACTTTGTACAAGAAAGCTGGGTCTATAATATGCGTGTTCCTACTACCATAAT |
| 2_hypothetical | I14 | Tb927.5.1040_for | GGGGACAAGTTTGTACAAAAAAGCAGGCTATGTCTATGGCCTACGTAATGGGAA | Tb927.5.1040_rev | GGGGACCACTTTGTACAAGAAAGCTGGGTTCACCACAGGGGACCACTG |
| 2_hypothetical | I15 | Tb927.9.3640_for | GGGGACAAGTTTGTACAAAAAAGCAGGCTATGCGACGCCTGGGCGTG | Tb927.9.3640_rev | GGGGACCACTTTGTACAAGAAAGCTGGGTTTACTCTTCAGCCCCCTCTTTCC |
| 2_hypothetical | I16 | Tb927.10.8340_for | GGGGACAAGTTTGTACAAAAAAGCAGGCTATGGGATGCTCGGCAAGCT | Tb927.10.8340_rev | GGGGACCACTTTGTACAAGAAAGCTGGGTTCAGTTGTAACGTTGCCCGC |
| 2_hypothetical | I17 | Tb927.11.11120_for | GGGGACAAGTTTGTACAAAAAAGCAGGCTATGCGTTGTGTGAGACGATCTG | Tb927.11.11120_rev | GGGGACCACTTTGTACAAGAAAGCTGGGTCTACAAATTATCATCGTGGTTCGCTGC |
| 2_hypothetical | I18 | Tb927.5.2015_for | GGGGACAAGTTTGTACAAAAAAGCAGGCTATGGCGGGTTTCAATATCGACG | Tb927.5.2015_rev | GGGGACCACTTTGTACAAGAAAGCTGGGTTCAGTGACGAGGCACCTCG |
| 2_hypothetical | I21 | Tb927.9.2390_for | GGGGACAAGTTTGTACAAAAAAGCAGGCTATGACAACACAGAACGGTGTTAACT | Tb927.9.2390_rev | GGGGACCACTTTGTACAAGAAAGCTGGGTTTATTTCCGTGGATAAAGGGCAACG |
| 2_hypothetical | I22 | Tb927.7.1630_for | GGGGACAAGTTTGTACAAAAAAGCAGGCTATGGGTCAGAAGCCGTCCAT | Tb927.7.1630_rev | GGGGACCACTTTGTACAAGAAAGCTGGGTCTATTGGTGGAAACCTACCAAGCAC |
| 2_hypothetical | I23 | Tb927.8.2260_for | GGGGACAAGTTTGTACAAAAAAGCAGGCTATGGGTTGTGGAGGTTCAGCT | Tb927.8.2260_rev | GGGGACCACTTTGTACAAGAAAGCTGGGTTCACCAGTGCATGAAACGAAACT |
| 2_hypothetical | I24 | Tb927.10.2070_for | GGGGACAAGTTTGTACAAAAAAGCAGGCTATGTCCAGAAAATCTTATCTAACCGCAG | Tb927.10.2070_rev | GGGGACCACTTTGTACAAGAAAGCTGGGTTTACCTTCGGTAGTTTCTTCGTTTTCC |
| 2_hypothetical | J2 | Tb927.10.7920_for | GGGGACAAGTTTGTACAAAAAAGCAGGCTATGAGCGACCCCCTCGTTG | Tb927.10.7920_rev | GGGGACCACTTTGTACAAGAAAGCTGGGTTTATCGCGTTTTTCCAAGTGTGACA |
| 2_hypothetical | J5 | Tb927.11.590_for | GGGGACAAGTTTGTACAAAAAAGCAGGCTATGCGGCGCACGTTAGGA | Tb927.11.590_rev | GGGGACCACTTTGTACAAGAAAGCTGGGTTTAGAAAGAAGGAAGCTTGGCACC |
| 2_hypothetical | J6 | Tb927.11.12160_for | GGGGACAAGTTTGTACAAAAAAGCAGGCTATGCGGCGTAGTCCTTGTGC | Tb927.11.12160_rev | GGGGACCACTTTGTACAAGAAAGCTGGGTCTAATTCCTTTGCCGAACTATATCCCC |
| 2_hypothetical | J7 | Tb927.8.4340_for | GGGGACAAGTTTGTACAAAAAAGCAGGCTATGGCTTCCGTCGGTAATTGC | Tb927.8.4340_rev | GGGGACCACTTTGTACAAGAAAGCTGGGTTCATTTAACGCAAGGCACTCCAC |
| 2_hypothetical | J8 | Tb927.10.9510_for | GGGGACAAGTTTGTACAAAAAAGCAGGCTATGAGAACGAATAGTCTCAACATCGG | Tb927.10.9510_rev | GGGGACCACTTTGTACAAGAAAGCTGGGTCTAATTGGAGGTTTGTGCCACAGA |
| 2_hypothetical | J9 | Tb927.10.13910_for | GGGGACAAGTTTGTACAAAAAAGCAGGCTATGCGTAGGACATTTCTCCGC | Tb927.10.13910_rev | GGGGACCACTTTGTACAAGAAAGCTGGGTTTACCACTGCTTCTTAATTTCGGGAG |
| 2_hypothetical | J10 | Tb927.9.7830_for | GGGGACAAGTTTGTACAAAAAAGCAGGCTATGACTATGAACTTTGGTAACATGACCC | Tb927.9.7830_rev | GGGGACCACTTTGTACAAGAAAGCTGGGTTTACTTCATCTCCGCATATTGAGGAGC |
| 2_hypothetical | J11 | Tb927.11.16170_for | GGGGACAAGTTTGTACAAAAAAGCAGGCTATGCGGCGCCTGTATCGC | Tb927.11.16170_rev | GGGGACCACTTTGTACAAGAAAGCTGGGTCTACAAATGGGTATTGTAACTAGGATCTGT |
| 2_hypothetical | J12 | Tb927.8.2440_for | GGGGACAAGTTTGTACAAAAAAGCAGGCTATGAGCCTTTTAAAAGCAGATCAATTGC | Tb927.8.2440_rev | GGGGACCACTTTGTACAAGAAAGCTGGGTCTACTCGCATGTATGAAGTTTTCCATCC |
| 2_hypothetical | J13 | Tb927.3.2300_for | GGGGACAAGTTTGTACAAAAAAGCAGGCTATGCTCCACACAGTGAGCGA | Tb927.3.2300_rev | GGGGACCACTTTGTACAAGAAAGCTGGGTTCAGTTATTATTACCGTCGCGCTC |
| 2_hypothetical | J14 | Tb927.10.4830_for | GGGGACAAGTTTGTACAAAAAAGCAGGCTATGTATAGTGATGGGACACGAAAGACC | Tb927.10.4830_rev | GGGGACCACTTTGTACAAGAAAGCTGGGTTTAAAAAATAATAACAGATTTGCGTTGTAG |
| 2_hypothetical | J15 | Tb927.10.4280_for | GGGGACAAGTTTGTACAAAAAAGCAGGCTATGCGGTCAACAGCTCCTCT | Tb927.10.4280_rev | GGGGACCACTTTGTACAAGAAAGCTGGGTTCAGTACTCTACCTCCTGCCCC |
| 2_hypothetical | J16 | Tb927.7.6430_for | GGGGACAAGTTTGTACAAAAAAGCAGGCTATGTCTGGGGGCCTTCCG | Tb927.7.6430_rev | GGGGACCACTTTGTACAAGAAAGCTGGGTCTATTGGTTCTCACAGCCGAGATC |
| 2_hypothetical | J17 | Tb927.9.12030_for | GGGGACAAGTTTGTACAAAAAAGCAGGCTATGGATTTAGATATCGTTGACGATGCG | Tb927.9.12030_rev | GGGGACCACTTTGTACAAGAAAGCTGGGTCTACCAGTCCAGTGAATGCGGT |
| 2_hypothetical | J18 | Tb927.8.4590_for | GGGGACAAGTTTGTACAAAAAAGCAGGCTATGGAACGTGTGAGGTACAAAGTC | Tb927.8.4590_rev | GGGGACCACTTTGTACAAGAAAGCTGGGTTCATGACCCCTTGTTCAAAGCTAC |
| 2_hypothetical | J19 | Tb927.7.4520_for | GGGGACAAGTTTGTACAAAAAAGCAGGCTATGACCGATACAATTGACGTGACG | Tb927.7.4520_rev | GGGGACCACTTTGTACAAGAAAGCTGGGTTTAAAATAGCGACATTTCAGCGTCCA |
| 2_hypothetical | J20 | Tb927.6.3270_for | GGGGACAAGTTTGTACAAAAAAGCAGGCTATGCATTTCACGTCGTCACGT | Tb927.6.3270_rev | GGGGACCACTTTGTACAAGAAAGCTGGGTCTAAGGCATTACTACTTGGTCCGC |
| 2_hypothetical | J21 | Tb927.10.9990_for | GGGGACAAGTTTGTACAAAAAAGCAGGCTATGACTGACGCCAGTAAAACAACC | Tb927.10.9990_rev | GGGGACCACTTTGTACAAGAAAGCTGGGTCTATTTGCTATTTTCAAAATGCTTCCTTAC |
| 2_hypothetical | J22 | Tb927.8.3270_for | GGGGACAAGTTTGTACAAAAAAGCAGGCTATGGAGTCCCTTGTACGTGACA | Tb927.8.3270_rev | GGGGACCACTTTGTACAAGAAAGCTGGGTTTACAATGAACATATTCGCTTCTGCTGT |
| 2_hypothetical | J23 | Tb927.3.4450_for | GGGGACAAGTTTGTACAAAAAAGCAGGCTATGCGGTGCTCTCCCCAA | Tb927.3.4450_rev | GGGGACCACTTTGTACAAGAAAGCTGGGTTTACGAATTGCGACTTATTTCTTCCGT |
| 2_hypothetical | J24 | Tb927.11.8070_for | GGGGACAAGTTTGTACAAAAAAGCAGGCTATGGATTTCCAGGGCTTACTACAACA | Tb927.11.8070_rev | GGGGACCACTTTGTACAAGAAAGCTGGGTTTATAAATCATCGGCAAGTTGTAGGCT |
| 2_hypothetical | K1 | Tb927.8.3040_for | GGGGACAAGTTTGTACAAAAAAGCAGGCTATGTTTCGCATATCCTCATGTTTTCACA | Tb927.8.3040_rev | GGGGACCACTTTGTACAAGAAAGCTGGGTTCAACGCAGTGGCGGGGT |
| 2_hypothetical | K2 | Tb927.11.14710_for | GGGGACAAGTTTGTACAAAAAAGCAGGCTATGGTCAAGAAAATGGGTAGAGACGA | Tb927.11.14710_rev | GGGGACCACTTTGTACAAGAAAGCTGGGTTCATTGTTGCTGATTTCGTTGAATTTTGT |
| 2_hypothetical | K3 | Tb927.3.2990_for | GGGGACAAGTTTGTACAAAAAAGCAGGCTATGATTCGTCATGGACTCACTGGA | Tb927.3.2990_rev | GGGGACCACTTTGTACAAGAAAGCTGGGTTCACCTCTCCACCTCACGAC |
| 2_hypothetical | K4 | Tb927.10.11950_for | GGGGACAAGTTTGTACAAAAAAGCAGGCTATGAAAAGATATATTCAACACAAGGAAAGT | Tb927.10.11950_rev | GGGGACCACTTTGTACAAGAAAGCTGGGTTCACTTAAAGTAACTTTTCATAAAGTGCTG |
| 2_hypothetical | K5 | Tb927.10.4970_for | GGGGACAAGTTTGTACAAAAAAGCAGGCTATGGAGGATATGGGCGACTTGG | Tb927.10.4970_rev | GGGGACCACTTTGTACAAGAAAGCTGGGTCTACCATTGAATCTCTTTGAGAATTGAATT |
| 2_hypothetical | K6 | Tb927.6.4500_for | GGGGACAAGTTTGTACAAAAAAGCAGGCTATGCGGCTCCTCTATGTCATCC | Tb927.6.4500_rev | GGGGACCACTTTGTACAAGAAAGCTGGGTTCATTTAACTGCCTTGGAGCGAC |
| 2_hypothetical | K7 | Tb927.4.4350_for | GGGGACAAGTTTGTACAAAAAAGCAGGCTATGTACGCACTACTGCAGCTTC | Tb927.4.4350_rev | GGGGACCACTTTGTACAAGAAAGCTGGGTTTATACAAACGACTTGCGCTCCA |
| 2_hypothetical | K8 | Tb927.11.430_for | GGGGACAAGTTTGTACAAAAAAGCAGGCTATGACACAAGGAATTTCTGAGGCTTT | Tb927.11.430_rev | GGGGACCACTTTGTACAAGAAAGCTGGGTTCATTGCCACTCTCCTTCACCT |
| 2_hypothetical | K9 | Tb927.8.3420_for | GGGGACAAGTTTGTACAAAAAAGCAGGCTATGCACGAGTTGGAGTTGCC | Tb927.8.3420_rev | GGGGACCACTTTGTACAAGAAAGCTGGGTCTCACCTACTGCACTTTGACAAGTTAAT |
| 2_hypothetical | K10 | Tb927.9.11700_for | GGGGACAAGTTTGTACAAAAAAGCAGGCTATGCAGAACTCTGCCTTCAGAGT | Tb927.9.11700_rev | GGGGACCACTTTGTACAAGAAAGCTGGGTTCAATTTAATTCAATGATGCGTGCCG |
| 2_hypothetical | K11 | Tb927.9.5370_for | GGGGACAAGTTTGTACAAAAAAGCAGGCTATGCGCTCTAAGATGGAACTGTCA | Tb927.9.5370_rev | GGGGACCACTTTGTACAAGAAAGCTGGGTTTAGCCGCATCCTCTAAGAGAAACG |
| 2_hypothetical | K12 | Tb927.10.7320_for | GGGGACAAGTTTGTACAAAAAAGCAGGCTATGGACTCAATCAATAGCGTTACGTT | Tb927.10.7320_rev | GGGGACCACTTTGTACAAGAAAGCTGGGTTCAAGCTGTACTTGCGATAATTTTCCT |
| 2_hypothetical | K13 | Tb927.7.6550_for | GGGGACAAGTTTGTACAAAAAAGCAGGCTATGAATCTAGCGAAGAAAATCATCGTGT | Tb927.7.6550_rev | GGGGACCACTTTGTACAAGAAAGCTGGGTTCAAGGTCTCGGTGATGTTGCA |
| 2_hypothetical | K14 | Tb927.11.8900_for | GGGGACAAGTTTGTACAAAAAAGCAGGCTATGAGTGTAACCCTTTCAGAGCCT | Tb927.11.8900_rev | GGGGACCACTTTGTACAAGAAAGCTGGGTCTATTGCTGATTGCAACTGGTCCT |
| 2_hypothetical | K15 | Tb927.9.12390_for | GGGGACAAGTTTGTACAAAAAAGCAGGCTATGCAAACCAATGAACGTGTCGT | Tb927.9.12390_rev | GGGGACCACTTTGTACAAGAAAGCTGGGTTTACTCTTCGTGTTGCTCACACAC |
| 2_hypothetical | K16 | Tb927.3.5410_for | GGGGACAAGTTTGTACAAAAAAGCAGGCTATGTTACGACTCTCCAGTCGGTG | Tb927.3.5410_rev | GGGGACCACTTTGTACAAGAAAGCTGGGTTCAGGAACCGCGTGTTCCT |
| 2_hypothetical | K17 | Tb927.4.520_for | GGGGACAAGTTTGTACAAAAAAGCAGGCTATGAACAAACAACTGCTTCAGACTCT | Tb927.4.520_rev | GGGGACCACTTTGTACAAGAAAGCTGGGTCTAATAATGTTTCTTGGGTGTCTTTTTCAT |
| 2_hypothetical | K18 | Tb927.4.1190_for | GGGGACAAGTTTGTACAAAAAAGCAGGCTATGGTGGATGTTCATCGAAAGGAAG | Tb927.4.1190_rev | GGGGACCACTTTGTACAAGAAAGCTGGGTCTAAAAAAATTCGAGCAGCGATGCT |
| 2_hypothetical | K19 | Tb927.6.2410_for | GGGGACAAGTTTGTACAAAAAAGCAGGCTATGAAGAGACCATGGGGTGAGTC | Tb927.6.2410_rev | GGGGACCACTTTGTACAAGAAAGCTGGGTCTAATCCGCATGCTTCGTAGATCC |
| 2_hypothetical | K20 | Tb927.9.15010_for | GGGGACAAGTTTGTACAAAAAAGCAGGCTATGCGCCGTACTGCCTTCT | Tb927.9.15010_rev | GGGGACCACTTTGTACAAGAAAGCTGGGTCTACGTACACGCCTTCAGCAAC |
| 2_hypothetical | K21 | Tb927.4.3010_for | GGGGACAAGTTTGTACAAAAAAGCAGGCTATGCTAAGAAGATGGTCTGAATGTGG | Tb927.4.3010_rev | GGGGACCACTTTGTACAAGAAAGCTGGGTTCATGCAGCAAATCGTCGTTCC |
| 2_hypothetical | K22 | Tb927.4.1290_for | GGGGACAAGTTTGTACAAAAAAGCAGGCTATGTCGACAACCCCTTACGATGA | Tb927.4.1290_rev | GGGGACCACTTTGTACAAGAAAGCTGGGTTCATTTGACTTCATCTGATTCTGCTGC |
| 2_hypothetical | K23 | Tb927.3.2850_for | GGGGACAAGTTTGTACAAAAAAGCAGGCTATGATGCTGAGCAATGCGCA | Tb927.3.2850_rev | GGGGACCACTTTGTACAAGAAAGCTGGGTTCACCCCCTAAAGAACCCAATGA |
| 2_hypothetical | K24 | Tb927.11.3430_for | GGGGACAAGTTTGTACAAAAAAGCAGGCTATGAAACCGGAAGGTGCAAAAGG | Tb927.11.3430_rev | GGGGACCACTTTGTACAAGAAAGCTGGGTCTACTTCGTTGCTTTCGGGGG |
| 2_hypothetical | L1 | Tb927.4.3040_for | GGGGACAAGTTTGTACAAAAAAGCAGGCTATGCGGCGGAGCGTCTGT | Tb927.4.3040_rev | GGGGACCACTTTGTACAAGAAAGCTGGGTTTATGGTTCATCATCTACGATTTGATCATT |
| 2_hypothetical | L2 | Tb927.9.7160_for | GGGGACAAGTTTGTACAAAAAAGCAGGCTATGGAGGACGTGAGGCCTG | Tb927.9.7160_rev | GGGGACCACTTTGTACAAGAAAGCTGGGTCTAAGACCCGTGTTCAAACATTTTCTTAC |
| 2_hypothetical | L3 | Tb927.11.3170_for | GGGGACAAGTTTGTACAAAAAAGCAGGCTATGGACTTTAACTTTCTATGGGAGATACCG | Tb927.11.3170_rev | GGGGACCACTTTGTACAAGAAAGCTGGGTCTACCGTATGAAGTAGCGCTGTAC |
| 2_hypothetical | L4 | Tb927.2.1740_for | GGGGACAAGTTTGTACAAAAAAGCAGGCTATGTCTCTAGGGAGTGCGCTT | Tb927.2.1740_rev | GGGGACCACTTTGTACAAGAAAGCTGGGTTCATCTGTCCACTTGCAGTTGTTT |
| 2_hypothetical | L5 | Tb927.4.1440_for | GGGGACAAGTTTGTACAAAAAAGCAGGCTATGCCACGCTCCCTGTTTCG | Tb927.4.1440_rev | GGGGACCACTTTGTACAAGAAAGCTGGGTTCACCGCGTCACCGGGGA |
| 2_hypothetical | L6 | Tb927.6.4080_for | GGGGACAAGTTTGTACAAAAAAGCAGGCTATGTTGCGATTTACTCGTCTTTTTCG | Tb927.6.4080_rev | GGGGACCACTTTGTACAAGAAAGCTGGGTCTAATAAAGTGCGAACATGGAGTTCATTT |

| | | | | | |
|---|---|---|---|---|---|
| 2_hypothetical | L7 | Tb927.8.8030_for | GGGGACAAGTTTGTACAAAAAAGCAGGCTATGAATAAGATGCACATATTGCCACCA | Tb927.8.8030_rev | GGGGACCACTTTGTACAAGAAAGCTGGGTTTACACAAATGATTTACGTTCCAAGAAACG |
| 2_hypothetical | L8 | Tb927.11.15420_for | GGGGACAAGTTTGTACAAAAAAGCAGGCTATGAACCAAGATTTGGACAAGGAGC | Tb927.11.15420_rev | GGGGACCACTTTGTACAAGAAAGCTGGGTCTACAACGACACAATGGTGACAGT |
| 2_hypothetical | L9 | Tb927.11.5810_for | GGGGACAAGTTTGTACAAAAAAGCAGGCTATGAGTGCACGGGGTTCCC | Tb927.11.5810_rev | GGGGACCACTTTGTACAAGAAAGCTGGGTCTATTTTGATCCTTTGGGGCGCC |
| 2_hypothetical | L10 | Tb927.7.3220_for | GGGGACAAGTTTGTACAAAAAAGCAGGCTATGCGAGCTGAAAAGGTGACAG | Tb927.7.3220_rev | GGGGACCACTTTGTACAAGAAAGCTGGGTTCACTCGGGAACATTTTGCGC |
| 2_hypothetical | L11 | Tb927.8.7500_for | GGGGACAAGTTTGTACAAAAAAGCAGGCTATGGAGAGAGTGGGTGTGGC | Tb927.8.7500_rev | GGGGACCACTTTGTACAAGAAAGCTGGGTTCAATCGATAGACATCCACGCAGA |
| 2_hypothetical | L12 | Tb927.7.5520_for | GGGGACAAGTTTGTACAAAAAAGCAGGCTATGACACCTCTTGGTTTTGATTTTACTTCA | Tb927.7.5520_rev | GGGGACCACTTTGTACAAGAAAGCTGGGTTTAAAGGAGTTTGAAATGACCTGTTACACG |
| 2_hypothetical | L13 | Tb927.11.15440_for | GGGGACAAGTTTGTACAAAAAAGCAGGCTATGTCAGCCCTTCTTCCAAGGG | Tb927.11.15440_rev | GGGGACCACTTTGTACAAGAAAGCTGGGTTCAACCCTCATTAGAAACTATGCCATAGG |
| 2_hypothetical | L14 | Tb927.9.6390_for | GGGGACAAGTTTGTACAAAAAAGCAGGCTATGTCATGTGGGAAGCCCACG | Tb927.9.6390_rev | GGGGACCACTTTGTACAAGAAAGCTGGGTTCACCGATGTTTCCTGATTATCGAC |
| 2_hypothetical | L15 | Tb927.9.10350_for | GGGGACAAGTTTGTACAAAAAAGCAGGCTATGTCTGTTAATCAGCTGCAGGAC | Tb927.9.10350_rev | GGGGACCACTTTGTACAAGAAAGCTGGGTTTATTTACCCCGTAACATGTCAAGTCG |
| 2_hypothetical | L16 | Tb927.11.9740_for | GGGGACAAGTTTGTACAAAAAAGCAGGCTATGGCTTCTGGTAGTGGTAGAGG | Tb927.11.9740_rev | GGGGACCACTTTGTACAAGAAAGCTGGGTTCACAATTGAACAATGGGAGTATAACTTAA |
| 2_hypothetical | L17 | Tb927.10.15810_for | GGGGACAAGTTTGTACAAAAAAGCAGGCTATGGGTAAGAAAAAGAACACGGCG | Tb927.10.15810_rev | GGGGACCACTTTGTACAAGAAAGCTGGGTTTAAGCTGGAGCTCCATAGACCTC |
| 2_hypothetical | L18 | Tb927.10.12930_for | GGGGACAAGTTTGTACAAAAAAGCAGGCTATGGACCAATTCACAAAACCACTGG | Tb927.10.12930_rev | GGGGACCACTTTGTACAAGAAAGCTGGGTTTAGTTGACATTTGTGCTTATCCCTACG |
| 2_hypothetical | L19 | Tb927.7.5870_for | GGGGACAAGTTTGTACAAAAAAGCAGGCTATGAAACGATGGGATGAAAAGTTAACGT | Tb927.7.5870_rev | GGGGACCACTTTGTACAAGAAAGCTGGGTTCAAGTCTTTGATGTGAGAACGGG |
| 2_hypothetical | L20 | Tb927.10.940_for | GGGGACAAGTTTGTACAAAAAAGCAGGCTATGACATTAGGGCCCACAACCA | Tb927.10.940_rev | GGGGACCACTTTGTACAAGAAAGCTGGGTTCACTCCAGTTGTCTTTCACTCCC |
| 2_hypothetical | L21 | Tb927.11.8100_for | GGGGACAAGTTTGTACAAAAAAGCAGGCTATGGCAGATGATGAACCCGAATAC | Tb927.11.8100_rev | GGGGACCACTTTGTACAAGAAAGCTGGGTTCATTCGATTTCATCAACACCCACG |
| 2_hypothetical | L22 | Tb927.3.4930_for | GGGGACAAGTTTGTACAAAAAAGCAGGCTATGAGAGAAACTGCTATTATGAACAACAGA | Tb927.3.4930_rev | GGGGACCACTTTGTACAAGAAAGCTGGGTTCAGGAAAAAAATATGTGAAGTAAAGCGTC |
| 2_hypothetical | L23 | Tb927.5.3190_for | GGGGACAAGTTTGTACAAAAAAGCAGGCTATGGGGCCGAAAAAACAACTTCG | Tb927.5.3190_rev | GGGGACCACTTTGTACAAGAAAGCTGGGTTCACGGTGGAAGCATCAATGTC |
| 2_hypothetical | L24 | Tb927.10.13980_for | GGGGACAAGTTTGTACAAAAAAGCAGGCTATGTCGATGATCGCCATGAGTTG | Tb927.10.13980_rev | GGGGACCACTTTGTACAAGAAAGCTGGGTCTACGATGGTCCGAGACTTAGGT |
| 2_hypothetical | M1 | Tb927.6.3770_for | GGGGACAAGTTTGTACAAAAAAGCAGGCTATGCTCATAGTTAAGACTGAAGGCG | Tb927.6.3770_rev | GGGGACCACTTTGTACAAGAAAGCTGGGTTTAACAGTGTTCTTTGTCTAATGGGGC |
| 2_hypothetical | M2 | Tb927.11.10250_for | GGGGACAAGTTTGTACAAAAAAGCAGGCTATGGTGGAAAAGCACATGTACGAC | Tb927.11.10250_rev | GGGGACCACTTTGTACAAGAAAGCTGGGTTTACTCAAGCCCTCTGGCCTT |
| 2_hypothetical | M3 | Tb927.6.3340_for | GGGGACAAGTTTGTACAAAAAAGCAGGCTATGGTTTACTACTTCACTCTTGCAGC | Tb927.6.3340_rev | GGGGACCACTTTGTACAAGAAAGCTGGGTTCACATGAAGTCATCTTCATCTGGAATATG |
| 2_hypothetical | M4 | Tb927.4.1580_for | GGGGACAAGTTTGTACAAAAAAGCAGGCTATGTCAAGAGGAGAACGCGGT | Tb927.4.1580_rev | GGGGACCACTTTGTACAAGAAAGCTGGGTCTAGTTATCCGAGGCGGACTCA |
| 2_hypothetical | M5 | Tb927.8.7130_for | GGGGACAAGTTTGTACAAAAAAGCAGGCTATGAACGCACTAAGCGGGAAG | Tb927.8.7130_rev | GGGGACCACTTTGTACAAGAAAGCTGGGTCTACGAGCCGTATTTCGGTAAACA |
| 2_hypothetical | M6 | Tb927.9.6620_for | GGGGACAAGTTTGTACAAAAAAGCAGGCTATGGACGCCGGAAGCGAT | Tb927.9.6620_rev | GGGGACCACTTTGTACAAGAAAGCTGGGTTTAATCCTGTAGCTTCTTTCGCTTCTC |
| 2_hypothetical | M7 | Tb927.5.740_for | GGGGACAAGTTTGTACAAAAAAGCAGGCTATGCCCTCCACCATCTGTCAC | Tb927.5.740_rev | GGGGACCACTTTGTACAAGAAAGCTGGGTTTAATGAAAGAAGAAAAAAAAGAAGAACAA |
| 2_hypothetical | M8 | Tb927.9.2160_for | GGGGACAAGTTTGTACAAAAAAGCAGGCTATGGGTCAGCGAGGCTCC | Tb927.9.2160_rev | GGGGACCACTTTGTACAAGAAAGCTGGGTTTACATACGCCGCACCGC |
| 2_hypothetical | M9 | Tb927.7.6190_for | GGGGACAAGTTTGTACAAAAAAGCAGGCTATGAGCAAAGACGGGAATTGCG | Tb927.7.6190_rev | GGGGACCACTTTGTACAAGAAAGCTGGGTTCACAAATCATCTTCACATGTGCAAC |
| 2_hypothetical | M10 | Tb927.9.8550_for | GGGGACAAGTTTGTACAAAAAAGCAGGCTATGCCTCATAACGATCCGGATGG | Tb927.9.8550_rev | GGGGACCACTTTGTACAAGAAAGCTGGGTTTATCGGCGCTTTTTCGTCTTACG |
| 2_hypothetical | M11 | Tb927.11.7330_for | GGGGACAAGTTTGTACAAAAAAGCAGGCTATGTCAACGCAGCTGGTTCG | Tb927.11.7330_rev | GGGGACCACTTTGTACAAGAAAGCTGGGTTCACAATAACATCGTGCTCTGCAC |
| 2_hypothetical | M12 | Tb927.9.2420_for | GGGGACAAGTTTGTACAAAAAAGCAGGCTATGCCGTATTACGCGGAGGA | Tb927.9.2420_rev | GGGGACCACTTTGTACAAGAAAGCTGGGTTTAGCTGTCATGTTGCTTGGTAATAACT |
| 2_hypothetical | M13 | Tb927.11.16380_for | GGGGACAAGTTTGTACAAAAAAGCAGGCTATGCTTCCAGGCAAACGTAGC | Tb927.11.16380_rev | GGGGACCACTTTGTACAAGAAAGCTGGGTTCAGCTACTCCTCTTCATGAGTTCC |
| 2_hypothetical | M14 | Tb927.7.2290_for | GGGGACAAGTTTGTACAAAAAAGCAGGCTATGAAAACCGCAAAGGTGAACAGT | Tb927.7.2290_rev | GGGGACCACTTTGTACAAGAAAGCTGGGTTCAGACTGCGAGTAAGGGAAGC |
| 2_hypothetical | M15 | Tb927.9.14140_for | GGGGACAAGTTTGTACAAAAAAGCAGGCTATGGCAACTGGAAGTCAAATCCC | Tb927.9.14140_rev | GGGGACCACTTTGTACAAGAAAGCTGGGTCTACTGCAACCCTGAAGTGCG |
| 2_hypothetical | M16 | Tb927.11.9320_for | GGGGACAAGTTTGTACAAAAAAGCAGGCTATGTATCGCTCAGTGGGAGAAGC | Tb927.11.9320_rev | GGGGACCACTTTGTACAAGAAAGCTGGGTTCATCCACCGTATGAGGGACG |
| 2_hypothetical | M17 | Tb927.11.10530_for | GGGGACAAGTTTGTACAAAAAAGCAGGCTATGCCACGCCACCACACG | Tb927.11.10530_rev | GGGGACCACTTTGTACAAGAAAGCTGGGTCTAAAATTCGCGAGTAGCTGCGA |
| 2_hypothetical | M18 | Tb927.7.4280_for | GGGGACAAGTTTGTACAAAAAAGCAGGCTATGGTGATGAAGGCTTTCCCCC | Tb927.7.4280_rev | GGGGACCACTTTGTACAAGAAAGCTGGGTTTAATCCCTTTCTTCAAGTTTCGGACG |
| 2_hypothetical | M19 | Tb927.11.970_for | GGGGACAAGTTTGTACAAAAAAGCAGGCTATGGATTGGACTGATCGTGTCCT | Tb927.11.970_rev | GGGGACCACTTTGTACAAGAAAGCTGGGTTCACCCACTGTTGTAAGGTGTTATGG |
| 2_hypothetical | M20 | Tb927.8.6900_for | GGGGACAAGTTTGTACAAAAAAGCAGGCTATGAGTCAACATGGCTTGGTTAGTG | Tb927.8.6900_rev | GGGGACCACTTTGTACAAGAAAGCTGGGTTCATGTGGCAACGGATCTGAAATC |
| 2_hypothetical | M21 | Tb927.11.11240_for | GGGGACAAGTTTGTACAAAAAAGCAGGCTATGTACAAGATAGATGGCGACGGC | Tb927.11.11240_rev | GGGGACCACTTTGTACAAGAAAGCTGGGTCTACGAAGTGCACAACGGCA |
| 2_hypothetical | M22 | Tb927.8.6630_for | GGGGACAAGTTTGTACAAAAAAGCAGGCTATGTCTCGGAATATTCACGTGAATTCA | Tb927.8.6630_rev | GGGGACCACTTTGTACAAGAAAGCTGGGTTTATGACAAATGCTTTCCAGGCGT |
| 2_hypothetical | M23 | Tb927.2.4620_for | GGGGACAAGTTTGTACAAAAAAGCAGGCTATGCCTTATCCCGATGTTTATGAACG | Tb927.2.4620_rev | GGGGACCACTTTGTACAAGAAAGCTGGGTTTATGATTTCGAGAAAAAGTATTTCCTCAG |
| 2_hypothetical | N1 | Tb927.9.9550_for | GGGGACAAGTTTGTACAAAAAAGCAGGCTATGTTGGATCACACAATTGCTATTGGT | Tb927.9.9550_rev | GGGGACCACTTTGTACAAGAAAGCTGGGTTCACTGTTGTTGCTGTTGCCA |
| 2_hypothetical | N2 | Tb927.5.3720_for | GGGGACAAGTTTGTACAAAAAAGCAGGCTATGCCAAAGCGGTCCAGGT | Tb927.5.3720_rev | GGGGACCACTTTGTACAAGAAAGCTGGGTTCACAAGACAAAGCGATCCGAAC |
| 2_hypothetical | N3 | Tb927.10.4630_for | GGGGACAAGTTTGTACAAAAAAGCAGGCTATGTTTGTTAGTGACAACGATGATTATCTC | Tb927.10.4630_rev | GGGGACCACTTTGTACAAGAAAGCTGGGTTTAACGACGCATGCAAGAGTGA |
| 2_hypothetical | N4 | Tb927.11.4140_for | GGGGACAAGTTTGTACAAAAAAGCAGGCTATGCTTTCCACTGCGTGCA | Tb927.11.4140_rev | GGGGACCACTTTGTACAAGAAAGCTGGGTTCAATCGAGGTCATATTCAAATTTAATCTT |
| 2_hypothetical | N5 | Tb927.6.4150_for | GGGGACAAGTTTGTACAAAAAAGCAGGCTATGTCTCGCCTTGGAGGAGATG | Tb927.6.4150_rev | GGGGACCACTTTGTACAAGAAAGCTGGGTCTACAGGACAAAGAATCCCCCAATTAC |
| 2_hypothetical | N6 | Tb927.11.10170_for | GGGGACAAGTTTGTACAAAAAAGCAGGCTATGTTGCGGCGTACTGTATGTG | Tb927.11.10170_rev | GGGGACCACTTTGTACAAGAAAGCTGGGTCTACTTGTTCACCTCCCCCTTCC |
| 2_hypothetical | N7 | Tb927.11.12910_for | GGGGACAAGTTTGTACAAAAAAGCAGGCTATGTCTGGTGTGTCAAGGGGG | Tb927.11.12910_rev | GGGGACCACTTTGTACAAGAAAGCTGGGTCTAGCCCTTCCACGTTGCTTT |
| 2_hypothetical | N8 | Tb927.10.2150_for | GGGGACAAGTTTGTACAAAAAAGCAGGCTATGGTTGGAACAGTATTACGCGTATATAC | Tb927.10.2150_rev | GGGGACCACTTTGTACAAGAAAGCTGGGTTCAAACAGTCTCCAGCCCGC |
| 2_hypothetical | N9 | Tb927.7.6110_for | GGGGACAAGTTTGTACAAAAAAGCAGGCTATGGAGGTGGAACTGGAGCC | Tb927.7.6110_rev | GGGGACCACTTTGTACAAGAAAGCTGGGTTTACACATGAATGCTCTTTCCCATTACA |
| 2_hypothetical | N10 | Tb927.11.14150_for | GGGGACAAGTTTGTACAAAAAAGCAGGCTATGCAGCCCATGGTGCTTC | Tb927.11.14150_rev | GGGGACCACTTTGTACAAGAAAGCTGGGTCTATTCAAGAGTCGCCAGAAGCG |
| 2_hypothetical | N11 | Tb927.11.2980_for | GGGGACAAGTTTGTACAAAAAAGCAGGCTATGGGTGATTTCGCGCTCATC | Tb927.11.2980_rev | GGGGACCACTTTGTACAAGAAAGCTGGGTTTAATCCTCAATCTCAATGTATTTCCTCAT |
| 2_hypothetical | N13 | Tb927.2.5500_for | GGGGACAAGTTTGTACAAAAAAGCAGGCTATGCTTCCGCACTCTCAGCC | Tb927.2.5500_rev | GGGGACCACTTTGTACAAGAAAGCTGGGTTTATGGCTTCCCATCGGTCGC |
| 2_hypothetical | N14 | Tb927.11.1720_for | GGGGACAAGTTTGTACAAAAAAGCAGGCTATGCAGCAGCAACAACAAGTACC | Tb927.11.1720_rev | GGGGACCACTTTGTACAAGAAAGCTGGGTTCAATTCCGCACAACACACCA |
| 2_hypothetical | N15 | Tb927.11.2110_for | GGGGACAAGTTTGTACAAAAAAGCAGGCTATGTCATTCATGTCATTGTTTCAGCG | Tb927.11.2110_rev | GGGGACCACTTTGTACAAGAAAGCTGGGTTTACTTCATAGCCGCCCGAAGA |
| 2_hypothetical | N17 | Tb927.10.10780_for | GGGGACAAGTTTGTACAAAAAAGCAGGCTATGCGCAATTCGTTGCGGT | Tb927.10.10780_rev | GGGGACCACTTTGTACAAGAAAGCTGGGTTCACTTTTCCCGCGGAACCT |
| 2_hypothetical | N18 | Tb927.9.1910_for | GGGGACAAGTTTGTACAAAAAAGCAGGCTATGCCTAACCCGTACACCGG | Tb927.9.1910_rev | GGGGACCACTTTGTACAAGAAAGCTGGGTTCAACATTTGTTAACCCTTTCGGTCG |
| 2_hypothetical | N19 | Tb927.9.2800_for | GGGGACAAGTTTGTACAAAAAAGCAGGCTATGGTTTTCTGCGTGTTTTCTTGTG | Tb927.9.2800_rev | GGGGACCACTTTGTACAAGAAAGCTGGGTTTAAATAGCAACCATGATACCTCTTTCGT |
| 2_hypothetical | N20 | Tb927.5.1270_for | GGGGACAAGTTTGTACAAAAAAGCAGGCTATGGAGCTTCTCAAGAAAACGGC | Tb927.5.1270_rev | GGGGACCACTTTGTACAAGAAAGCTGGGTCTATTCCGTTCGCACAACCTTAGA |
| 2_hypothetical | N21 | Tb927.8.5430_for | GGGGACAAGTTTGTACAAAAAAGCAGGCTATGGGCAACAACGGAAGTAGC | Tb927.8.5430_rev | GGGGACCACTTTGTACAAGAAAGCTGGGTCTATGCCGCCAGTCGAACAC |
| 2_hypothetical | N22 | Tb927.4.3180_for | GGGGACAAGTTTGTACAAAAAAGCAGGCTATGAGTACCACAGGAAGCTGGAC | Tb927.4.3180_rev | GGGGACCACTTTGTACAAGAAAGCTGGGTTTAACTGTCACTGCCTTTATTGGATTGA |
| 2_hypothetical | N23 | Tb927.5.1930_for | GGGGACAAGTTTGTACAAAAAAGCAGGCTATGCATTCCCTTTGGGTGCG | Tb927.5.1930_rev | GGGGACCACTTTGTACAAGAAAGCTGGGTCTACGCGTATCCTCTGCTGC |
| 2_hypothetical | N24 | Tb927.6.1270_for | GGGGACAAGTTTGTACAAAAAAGCAGGCTATGCCTTGCAACTGCCGTC | Tb927.6.1270_rev | GGGGACCACTTTGTACAAGAAAGCTGGGTCTAGAATATAGTGAATGTTTTCTTGAGCTC |
| 2_hypothetical | O1 | Tb927.10.1440_for | GGGGACAAGTTTGTACAAAAAAGCAGGCTATGAAGTCGTCCGACATATTCTTTGC | Tb927.10.1440_rev | GGGGACCACTTTGTACAAGAAAGCTGGGTTTACTTGTTGTACGGCAGACGG |
| 2_hypothetical | O2 | Tb927.2.3340_for | GGGGACAAGTTTGTACAAAAAAGCAGGCTATGTCTGAAAACCCCGCCCA | Tb927.2.3340_rev | GGGGACCACTTTGTACAAGAAAGCTGGGTTTACACCATTTCCACTCTCCTTGGA |
| 2_hypothetical | O3 | Tb927.5.3300_for | GGGGACAAGTTTGTACAAAAAAGCAGGCTATGTTACGCAGAGCTGTTGCA | Tb927.5.3300_rev | GGGGACCACTTTGTACAAGAAAGCTGGGTTTAGTTCGCGCTCGTCATATGC |
| 2_hypothetical | O4 | Tb927.11.13230_for | GGGGACAAGTTTGTACAAAAAAGCAGGCTATGAGAAGTAAAACGGCGGCAC | Tb927.11.13230_rev | GGGGACCACTTTGTACAAGAAAGCTGGGTCTAGTAGGCAAACCACCACCAGC |
| 2_hypothetical | O5 | Tb927.10.7860_for | GGGGACAAGTTTGTACAAAAAAGCAGGCTATGACCGTGTCAGAAGCCTCC | Tb927.10.7860_rev | GGGGACCACTTTGTACAAGAAAGCTGGGTCTACCGCGATCTGTTGGTTGC |
| 2_hypothetical | O6 | Tb927.5.3140_for | GGGGACAAGTTTGTACAAAAAAGCAGGCTATGTTGCCCATAACGCGGC | Tb927.5.3140_rev | GGGGACCACTTTGTACAAGAAAGCTGGGTTTACTCCTCTCCTGAAGGTGGGG |
| 2_hypothetical | O7 | Tb927.7.3480_for | GGGGACAAGTTTGTACAAAAAAGCAGGCTATGCTAACCGACACCCTCACA | Tb927.7.3480_rev | GGGGACCACTTTGTACAAGAAAGCTGGGTTTAAAAAAAAATGAAAAGTATATATATCAA |
| 2_hypothetical | O8 | Tb927.4.3960_for | GGGGACAAGTTTGTACAAAAAAGCAGGCTATGTCACTAGATATCTCACGATTCCTCC | Tb927.4.3960_rev | GGGGACCACTTTGTACAAGAAAGCTGGGTCTACAAACTCCACAACGATCGCA |
| 2_hypothetical | O9 | Tb927.10.6770_for | GGGGACAAGTTTGTACAAAAAAGCAGGCTATGCGTGCGCTGTTTGGC | Tb927.10.6770_rev | GGGGACCACTTTGTACAAGAAAGCTGGGTTCACAACCCCCCAGTCACG |
| 2_hypothetical | O10 | Tb927.10.13760_for | GGGGACAAGTTTGTACAAAAAAGCAGGCTATGCGGTGTCACCGAGTTTTT | Tb927.10.13760_rev | GGGGACCACTTTGTACAAGAAAGCTGGGTTTAGTTCAAGTGATCATCGGGGGA |
| 2_hypothetical | O11 | Tb927.9.8290_for | GGGGACAAGTTTGTACAAAAAAGCAGGCTATGATGCGGCGCATGTGG | Tb927.9.8290_rev | GGGGACCACTTTGTACAAGAAAGCTGGGTTCAACACTTCTCTCCCTCCTTATTATCTG |
| 2_hypothetical | O12 | Tb927.9.5390_for | GGGGACAAGTTTGTACAAAAAAGCAGGCTATGTCTATTGAGAGGCTTCGGCG | Tb927.9.5390_rev | GGGGACCACTTTGTACAAGAAAGCTGGGTTTAATCATCGCTAAGTTCATCGAAATCATC |
| 2_hypothetical | O13 | Tb927.10.12630_for | GGGGACAAGTTTGTACAAAAAAGCAGGCTATGGGCTATACGTTCTCGCGT | Tb927.10.12630_rev | GGGGACCACTTTGTACAAGAAAGCTGGGTCTACATCAAATTCTTCATCTTCTCCAGCA |
| 2_hypothetical | O14 | Tb927.7.6150_for | GGGGACAAGTTTGTACAAAAAAGCAGGCTATGTACTCTGTTCATTGGGAGGAAGA | Tb927.7.6150_rev | GGGGACCACTTTGTACAAGAAAGCTGGGTTTACACATGAATGCTCTTTCCCATTACA |
| 2_hypothetical | O15 | Tb927.9.2990_for | GGGGACAAGTTTGTACAAAAAAGCAGGCTATGGAGGAAGCGAAAGAGAATGTCC | Tb927.9.2990_rev | GGGGACCACTTTGTACAAGAAAGCTGGGTCTAATTCTCCCACGAAATTCTACTTCCC |
| 2_hypothetical | O16 | Tb927.3.2560_for | GGGGACAAGTTTGTACAAAAAAGCAGGCTATGCTGTGCCTACCAATAATACTGTTC | Tb927.3.2560_rev | GGGGACCACTTTGTACAAGAAAGCTGGGTTCAGAGAAATAGGAGCAACACTGAGG |
| 2_hypothetical | O17 | Tb927.10.6280_for | GGGGACAAGTTTGTACAAAAAAGCAGGCTATGGCTGATGCTTATCGAGAACG | Tb927.10.6280_rev | GGGGACCACTTTGTACAAGAAAGCTGGGTCTATGGACTCACAAAACTAATTAACTCTGC |
| 2_hypothetical | O18 | Tb927.5.4410_for | GGGGACAAGTTTGTACAAAAAAGCAGGCTATGGCAGGCCGTTTCCGT | Tb927.5.4410_rev | GGGGACCACTTTGTACAAGAAAGCTGGGTCTACCAGTACCTATCCACTTTCTTTGGT |
| 2_hypothetical | O19 | Tb927.11.9460_for | GGGGACAAGTTTGTACAAAAAAGCAGGCTATGACCAAAGTTTGGATGCCGT | Tb927.11.9460_rev | GGGGACCACTTTGTACAAGAAAGCTGGGTCTATGGTGTGTCGGTGGCTG |
| 2_hypothetical | O20 | Tb927.3.2550_for | GGGGACAAGTTTGTACAAAAAAGCAGGCTATGCTGTGCCTACCAATAATACTGTTC | Tb927.3.2550_rev | GGGGACCACTTTGTACAAGAAAGCTGGGTTCAGAGAAATAGGAGCAACACTGGA |
| 2_hypothetical | O21 | Tb927.8.5660_for | GGGGACAAGTTTGTACAAAAAAGCAGGCTATGATATCGCCCAGTACATTGCTG | Tb927.8.5660_rev | GGGGACCACTTTGTACAAGAAAGCTGGGTTCAGGAGTTCAAGTATAACACCAGCA |
| 2_hypothetical | O23 | Tb927.11.2400_for | GGGGACAAGTTTGTACAAAAAAGCAGGCTATGGGTTGTGGCGGTTCCA | Tb927.11.2400_rev | GGGGACCACTTTGTACAAGAAAGCTGGGTTTACTCGGATTTGTACTTCTGGAGACT |
| 2_hypothetical | O24 | Tb927.5.3110_for | GGGGACAAGTTTGTACAAAAAAGCAGGCTATGCTGAACCGGTCACGTTTG | Tb927.5.3110_rev | GGGGACCACTTTGTACAAGAAAGCTGGGTTCAGAAACCCAACCCCAGCA |
| 2_hypothetical | P1 | Tb927.9.14440_for | GGGGACAAGTTTGTACAAAAAAGCAGGCTATGTTCAGGGGAGGACAGTCC | Tb927.9.14440_rev | GGGGACCACTTTGTACAAGAAAGCTGGGTTTACCCCATATGTTCCACGGTTAGA |
| 2_hypothetical | P2 | Tb927.11.7190_for | GGGGACAAGTTTGTACAAAAAAGCAGGCTATGGGGAAGCGTGATTTGGAGA | Tb927.11.7190_rev | GGGGACCACTTTGTACAAGAAAGCTGGGTTTATCCCGCCGACGGAACG |
| 2_hypothetical | P3 | Tb927.7.6130_for | GGGGACAAGTTTGTACAAAAAAGCAGGCTATGTCAAATTCGCCGGCTAGTC | Tb927.7.6130_rev | GGGGACCACTTTGTACAAGAAAGCTGGGTTTACACATGAATGCTCTTTCCCATTACA |
| 2_hypothetical | P4 | Tb927.11.5890_for | GGGGACAAGTTTGTACAAAAAAGCAGGCTATGGCCACTGCGCGGTCG | Tb927.11.5890_rev | GGGGACCACTTTGTACAAGAAAGCTGGGTTCACAGTTCGTCAAAATACGGAATGC |
| 2_hypothetical | P5 | Tb927.8.7710_for | GGGGACAAGTTTGTACAAAAAAGCAGGCTATGGAAACCACGCACATAATCAACG | Tb927.8.7710_rev | GGGGACCACTTTGTACAAGAAAGCTGGGTTCACCAACAGTTGTTCCCAACG |
| 2_hypothetical | P6 | Tb927.9.4570_for | GGGGACAAGTTTGTACAAAAAAGCAGGCTATGACGGATTATGAGGAATTGCGG | Tb927.9.4570_rev | GGGGACCACTTTGTACAAGAAAGCTGGGTCTACCCCCGCCGTCCGCG |
| 2_hypothetical | P7 | Tb927.4.2980_for | GGGGACAAGTTTGTACAAAAAAGCAGGCTATGGGGCTCGTGCACAGT | Tb927.4.2980_rev | GGGGACCACTTTGTACAAGAAAGCTGGGTTCAGATTAATGACGCCAGCGTG |
| 2_hypothetical | P8 | Tb927.8.2290_for | GGGGACAAGTTTGTACAAAAAAGCAGGCTATGTTGATGTTTCATCTTAACGCCAATAG | Tb927.8.2290_rev | GGGGACCACTTTGTACAAGAAAGCTGGGTTCAACGCAGCCGTTTGCA |
| 2_hypothetical | P9 | Tb927.3.4410_for | GGGGACAAGTTTGTACAAAAAAGCAGGCTATGTCTTATAGTGACATTCAGAGCCTCG | Tb927.3.4410_rev | GGGGACCACTTTGTACAAGAAAGCTGGGTTCAAACCAGCCTATCGGAATCGT |
| 2_hypothetical | P10 | Tb927.4.790_for | GGGGACAAGTTTGTACAAAAAAGCAGGCTATGGCGAACATATCTCCTAATTCAGATG | Tb927.4.790_rev | GGGGACCACTTTGTACAAGAAAGCTGGGTTTAAAATAACTGGGCTCGCCTTGAT |
| 2_hypothetical | P11 | Tb927.10.14640_for | GGGGACAAGTTTGTACAAAAAAGCAGGCTATGGGCGGCGATGGCCAA | Tb927.10.14640_rev | GGGGACCACTTTGTACAAGAAAGCTGGGTTCACGTGTTTGAGCGCCC |
| 2_hypothetical | P12 | Tb927.7.5800_for | GGGGACAAGTTTGTACAAAAAAGCAGGCTATGGGCACACACATTATAGCGTATG | Tb927.7.5800_rev | GGGGACCACTTTGTACAAGAAAGCTGGGTCTACCACGGAAACATATCGCTACC |
| 2_hypothetical | P13 | Tb927.11.4870_for | GGGGACAAGTTTGTACAAAAAAGCAGGCTATGTCCGGTAGAATTTACAAGCTTTTCA | Tb927.11.4870_rev | GGGGACCACTTTGTACAAGAAAGCTGGGTCTACGATTTAGGGATAGCTGAGATGTGA |
| 2_hypothetical | P14 | Tb927.11.5460_for | GGGGACAAGTTTGTACAAAAAAGCAGGCTATGACAAATTCCCCAGATGACGATG | Tb927.11.5460_rev | GGGGACCACTTTGTACAAGAAAGCTGGGTTCAATCCCGCGTTCGCTTAAA |
| 2_hypothetical | P15 | Tb927.9.3770_for | GGGGACAAGTTTGTACAAAAAAGCAGGCTATGAGTGAGTCTAGTGCGACCAA | Tb927.9.3770_rev | GGGGACCACTTTGTACAAGAAAGCTGGGTTTAGGATTGCCGTTTCTTTTTTGAGGT |
| 2_hypothetical | P16 | Tb927.3.2220_for | GGGGACAAGTTTGTACAAAAAAGCAGGCTATGGAGGAGGACTACCATGAGTATGA | Tb927.3.2220_rev | GGGGACCACTTTGTACAAGAAAGCTGGGTTCACATAAAGTTGTCATCACCATCGG |
| 2_hypothetical | P17 | Tb927.10.9220_for | GGGGACAAGTTTGTACAAAAAAGCAGGCTATGTCTGAGGAGTCCTTGACGTT | Tb927.10.9220_rev | GGGGACCACTTTGTACAAGAAAGCTGGGTCTACCGAAATCCACTGAACCTCCC |
| 2_hypothetical | P18 | Tb927.11.13530_for | GGGGACAAGTTTGTACAAAAAAGCAGGCTATGAGTGAAGAAGATGGTGAAGGTCT | Tb927.11.13530_rev | GGGGACCACTTTGTACAAGAAAGCTGGGTCTATGTAGAGTGCGCTTCGCC |
| 2_hypothetical | P19 | Tb927.8.2300_for | GGGGACAAGTTTGTACAAAAAAGCAGGCTATGGATGATGTGAAGAGGAAGTCCG | Tb927.8.2300_rev | GGGGACCACTTTGTACAAGAAAGCTGGGTCTAAAAGCCGAACAACATTCTGTATAAGG |
| 2_hypothetical | P20 | Tb927.4.1000_for | GGGGACAAGTTTGTACAAAAAAGCAGGCTATGATGGAGCTCCCTATGGAAATGA | Tb927.4.1000_rev | GGGGACCACTTTGTACAAGAAAGCTGGGTCTACAGCTCCTGAGATCTTTCAGATGA |
| 2_hypothetical | P21 | Tb927.11.10070_for | GGGGACAAGTTTGTACAAAAAAGCAGGCTATGGTCCTCACAGGGGTCAAAC | Tb927.11.10070_rev | GGGGACCACTTTGTACAAGAAAGCTGGGTTCACTCCAACTGCCCGTTTG |
| 2_hypothetical | P22 | Tb927.3.4200_for | GGGGACAAGTTTGTACAAAAAAGCAGGCTATGGCGATCCCTTTGGGGG | Tb927.3.4200_rev | GGGGACCACTTTGTACAAGAAAGCTGGGTTCACACCAAATCATTGCGCCA |
| 2_hypothetical | P23 | Tb927.8.7310_for | GGGGACAAGTTTGTACAAAAAAGCAGGCTATGAGGAGAGTGCTGCTATTCCTG | Tb927.8.7310_rev | GGGGACCACTTTGTACAAGAAAGCTGGGTTCAAAGTAACGTGCTGTACAAAAAAGG |
| 2_hypothetical | P24 | Tb927.11.7550_for | GGGGACAAGTTTGTACAAAAAAGCAGGCTATGCTTCAAAACTGTGAGGTTGCT | Tb927.11.7550_rev | GGGGACCACTTTGTACAAGAAAGCTGGGTTTATTCAGACAAACGATCTGAGTTTGTGT |
| 2_known | A1 | Tb927.8.5230_for | GGGGACAAGTTTGTACAAAAAAGCAGGCTATGTCCTTGTCATCCACCACAATC | Tb927.8.5230_rev | GGGGACCACTTTGTACAAGAAAGCTGGGTCTACCCGCCGGAAGGGGA |
| 2_known | A2 | Tb927.9.13150_for | GGGGACAAGTTTGTACAAAAAAGCAGGCTATGGGATACAGGTTGTTGCGTG | Tb927.9.13150_rev | GGGGACCACTTTGTACAAGAAAGCTGGGTTTATGACGTGCAGGTGGCAC |
| 2_known | A3 | Tb927.10.3760_for | GGGGACAAGTTTGTACAAAAAAGCAGGCTATGTCATCGAACCGGTACACTGC | Tb927.10.3760_rev | GGGGACCACTTTGTACAAGAAAGCTGGGTCTACACAACAAGATCCGCCTCG |
| 2_known | A4 | Tb927.10.8310_for | GGGGACAAGTTTGTACAAAAAAGCAGGCTATGAAACGACAGAGGTCGGGA | Tb927.10.8310_rev | GGGGACCACTTTGTACAAGAAAGCTGGGTTTATGATAGCGGCACCCATAGTAACA |
| 2_known | A5 | Tb927.5.2690_for | GGGGACAAGTTTGTACAAAAAAGCAGGCTATGCCAATAAATAACTGTGAATTGGACG | Tb927.5.2690_rev | GGGGACCACTTTGTACAAGAAAGCTGGGTCTAGCATCCTAACACCGATTTTTTGTAGC |
| 2_known | A6 | Tb927.11.11730_for | GGGGACAAGTTTGTACAAAAAAGCAGGCTATGAAGTTTTCACGCGTTGCCA | Tb927.11.11730_rev | GGGGACCACTTTGTACAAGAAAGCTGGGTTTACTGAAACGTTATGGTAGGTGTTTCTAC |
| 2_known | A7 | Tb927.7.6200_for | GGGGACAAGTTTGTACAAAAAAGCAGGCTATGAAACCCATACCAGTAGCCATGA | Tb927.7.6200_rev | GGGGACCACTTTGTACAAGAAAGCTGGGTTCAATCGGCAGGACTGTTACCA |
| 2_known | A8 | Tb927.10.14780_for | GGGGACAAGTTTGTACAAAAAAGCAGGCTATGCCCGCAGGCATGTGT | Tb927.10.14780_rev | GGGGACCACTTTGTACAAGAAAGCTGGGTCTAGACCTTTAAGAAGGGATGTTGCAA |
| 2_known | A9 | Tb927.4.4170_for | GGGGACAAGTTTGTACAAAAAAGCAGGCTATGTACAGGTGCCGTTCCGC | Tb927.4.4170_rev | GGGGACCACTTTGTACAAGAAAGCTGGGTCTAGTCCACGCGACTGCAGT |
| 2_known | A10 | Tb927.9.3820_for | GGGGACAAGTTTGTACAAAAAAGCAGGCTATGTCGGATGTAAAAGGGGCGC | Tb927.9.3820_rev | GGGGACCACTTTGTACAAGAAAGCTGGGTTCACCTAGACATGTTGTATATCGCCC |
| 2_known | A11 | Tb927.3.1890_for | GGGGACAAGTTTGTACAAAAAAGCAGGCTATGGCCGAAACCCGAAAAGAAG | Tb927.3.1890_rev | GGGGACCACTTTGTACAAGAAAGCTGGGTCTAGAATTCTTCCATTTGTGGCTCAGG |
| 2_known | A12 | Tb927.9.11870_for | GGGGACAAGTTTGTACAAAAAAGCAGGCTATGGGCGCTAGTGAAGCAAAG | Tb927.9.11870_rev | GGGGACCACTTTGTACAAGAAAGCTGGGTTCAATAATTCGGTAGACGGCTCAGC |
| 2_known | A13 | Tb927.2.6070_for | GGGGACAAGTTTGTACAAAAAAGCAGGCTATGTTCAGCGGGCTTCAGC | Tb927.2.6070_rev | GGGGACCACTTTGTACAAGAAAGCTGGGTCTAAGAATCTGATTCTTTAGAGGCTCCAC |
| 2_known | A14 | Tb927.10.14930_for | GGGGACAAGTTTGTACAAAAAAGCAGGCTATGAACGATGCCGGTATGGGA | Tb927.10.14930_rev | GGGGACCACTTTGTACAAGAAAGCTGGGTTTAACCACCACCTCGAGTGTCA |
| 2_known | A15 | Tb927.10.11980_for | GGGGACAAGTTTGTACAAAAAAGCAGGCTATGGATACTTCCTCGGAGCTGTT | Tb927.10.11980_rev | GGGGACCACTTTGTACAAGAAAGCTGGGTTTACATTAGTGTACAACTGCTCGAATTGT |
| 2_known | A16 | Tb927.9.12290_for | GGGGACAAGTTTGTACAAAAAAGCAGGCTATGTCTCATCCCGACAATGACGC | Tb927.9.12290_rev | GGGGACCACTTTGTACAAGAAAGCTGGGTCTACACTGATGGTTGCACATCGG |
| 2_known | A17 | Tb927.8.5810_for | GGGGACAAGTTTGTACAAAAAAGCAGGCTATGACTGAGGAGTATGTCCACACG | Tb927.8.5810_rev | GGGGACCACTTTGTACAAGAAAGCTGGGTCTAATCCCTGGTGAGGGCGT |
| 2_known | A18 | Tb927.3.3770_for | GGGGACAAGTTTGTACAAAAAAGCAGGCTATGAGTGCAACAGAGGCGGT | Tb927.3.3770_rev | GGGGACCACTTTGTACAAGAAAGCTGGGTTTAGTACCTAGAGAACCGTGTCACC |
| 2_known | A19 | Tb927.7.4420_for | GGGGACAAGTTTGTACAAAAAAGCAGGCTATGTCCTCGCGTTACGACTCC | Tb927.7.4420_rev | GGGGACCACTTTGTACAAGAAAGCTGGGTTTACGGCGAGGCCAAGCG |
| 2_known | A20 | Tb927.10.12780_for | GGGGACAAGTTTGTACAAAAAAGCAGGCTATGCCCTCAAAGAAATCTATGGCAA | Tb927.10.12780_rev | GGGGACCACTTTGTACAAGAAAGCTGGGTTTATAACCCGTAAGGATTATGTCGGTAGA |

| | | | | | |
|---|---|---|---|---|---|
| 2_known | A21 | Tb927.10.14440_for | GGGGACAAGTTTGTACAAAAAAGCAGGCTATGGAGGAAAGTGTATATAAACCACTTGG | Tb927.10.14440_rev | GGGGACCACTTTGTACAAGAAAGCTGGGTTCATGTTCCTATCCCACACTCTCTG |
| 2_known | A22 | Tb927.10.4440_for | GGGGACAAGTTTGTACAAAAAAGCAGGCTATGAGGAAACCCGGGCGG | Tb927.10.4440_rev | GGGGACCACTTTGTACAAGAAAGCTGGGTCTAAACGCTCCTGCGGCG |
| 2_known | A23 | Tb927.7.5160_for | GGGGACAAGTTTGTACAAAAAAGCAGGCTATGAAGAACGCCCGCAGAGT | Tb927.7.5160_rev | GGGGACCACTTTGTACAAGAAAGCTGGGTTTACGACTTTAGCCAGTGCCCA |
| 2_known | A24 | Tb927.7.2700_for | GGGGACAAGTTTGTACAAAAAAGCAGGCTATGAAGGTATTTGCAGCTGCTGC | Tb927.7.2700_rev | GGGGACCACTTTGTACAAGAAAGCTGGGTTCAAAACTTGTAAACCATAGCCTCTGG |
| 2_known | B1 | Tb927.6.2780_for | GGGGACAAGTTTGTACAAAAAAGCAGGCTATGCGACGTAGCGTTATTCGG | Tb927.6.2780_rev | GGGGACCACTTTGTACAAGAAAGCTGGGTTTACATCAGTCTCCGCTTCTTCGC |
| 2_known | B2 | Tb927.6.4350_for | GGGGACAAGTTTGTACAAAAAAGCAGGCTATGACGTCTTCGTCAAAATCCAAGT | Tb927.6.4350_rev | GGGGACCACTTTGTACAAGAAAGCTGGGTCTACTCCATCTTGAAATCATCGCGT |
| 2_known | B3 | Tb927.7.2710_for | GGGGACAAGTTTGTACAAAAAAGCAGGCTATGAAAGCTCTGTTTGCCTCCG | Tb927.7.2710_rev | GGGGACCACTTTGTACAAGAAAGCTGGGTCTAAATTTTAAACACCATTTTCTGTATGTA |
| 2_known | B4 | Tb927.9.10950_for | GGGGACAAGTTTGTACAAAAAAGCAGGCTATGTTCGTGGATTACTACGCTGTAC | Tb927.9.10950_rev | GGGGACCACTTTGTACAAGAAAGCTGGGTTTAGATGTTAAAGAAAACCCCCCACG |
| 2_known | B5 | Tb927.9.10580_for | GGGGACAAGTTTGTACAAAAAAGCAGGCTATGTCCTCAGCAATGGCTCGC | Tb927.9.10580_rev | GGGGACCACTTTGTACAAGAAAGCTGGGTTTATTTGGCATCTTTTTCACTTTTCATATT |
| 2_known | B6 | Tb927.11.10510_for | GGGGACAAGTTTGTACAAAAAAGCAGGCTATGCTCCGTCGCAGTGGT | Tb927.11.10510_rev | GGGGACCACTTTGTACAAGAAAGCTGGGTCTATGCAGCCCCCACTTTACTG |
| 2_known | B7 | Tb927.7.1500_for | GGGGACAAGTTTGTACAAAAAAGCAGGCTATGGCCACCTCCCTTTCGG | Tb927.7.1500_rev | GGGGACCACTTTGTACAAGAAAGCTGGGTTTAATGACGTTGTTGCAGTTGCG |
| 2_known | B8 | Tb927.3.2980_for | GGGGACAAGTTTGTACAAAAAAGCAGGCTATGGTTTCCGAGGGCACTTCC | Tb927.3.2980_rev | GGGGACCACTTTGTACAAGAAAGCTGGGTTTACCGTTCCTCCATGAACTTCTTG |
| 2_known | B9 | Tb927.3.1820_for | GGGGACAAGTTTGTACAAAAAAGCAGGCTATGTCTGCCGCATTTGGTATCG | Tb927.3.1820_rev | GGGGACCACTTTGTACAAGAAAGCTGGGTCTAGCGGCGGTATTGCCC |
| 2_known | B10 | Tb927.6.3850_for | GGGGACAAGTTTGTACAAAAAAGCAGGCTATGCCATTGTTTTGTAGTTCTACCACC | Tb927.6.3850_rev | GGGGACCACTTTGTACAAGAAAGCTGGGTTCACGTGGATGGATATGGACAAACA |
| 2_known | B11 | Tb927.4.5790_for | GGGGACAAGTTTGTACAAAAAAGCAGGCTATGAGATTTGTTTCCAAGGGTGTAAGG | Tb927.4.5790_rev | GGGGACCACTTTGTACAAGAAAGCTGGGTTCACGCGTATTCCTCCTCTGC |
| 2_known | B12 | Tb927.9.15550_for | GGGGACAAGTTTGTACAAAAAAGCAGGCTATGAGCATCACTTTTAATAATTTATGGCTA | Tb927.9.15550_rev | GGGGACCACTTTGTACAAGAAAGCTGGGTTTACAGTACAGAACCGAGAAACAGTCC |
| 2_known | B14 | Tb927.10.5040_for | GGGGACAAGTTTGTACAAAAAAGCAGGCTATGGGTTTGGAGGATATTGATGTAGATGA | Tb927.10.5040_rev | GGGGACCACTTTGTACAAGAAAGCTGGGTTCAACTCGGTGGAAAGAACCAATC |
| 2_known | B15 | Tb927.11.3550_for | GGGGACAAGTTTGTACAAAAAAGCAGGCTATGTCGACAAAGCCAAAACCTTTAGT | Tb927.11.3550_rev | GGGGACCACTTTGTACAAGAAAGCTGGGTTTAATCCTTCTTCCGGTCCTCGC |
| 2_known | B16 | Tb927.11.2920_for | GGGGACAAGTTTGTACAAAAAAGCAGGCTATGTCACGATGTGGGCGC | Tb927.11.2920_rev | GGGGACCACTTTGTACAAGAAAGCTGGGTTTACCGGTCCGTTCCCTCG |
| 2_known | B17 | Tb927.9.12170_for | GGGGACAAGTTTGTACAAAAAAGCAGGCTATGGCGGGAGTTCTCACTGAA | Tb927.9.12170_rev | GGGGACCACTTTGTACAAGAAAGCTGGGTTTACTGCAAAGCATCAATGAAGCG |
| 2_known | B18 | Tb927.11.9180_for | GGGGACAAGTTTGTACAAAAAAGCAGGCTATGAGTAACGCTAACGCTGCTC | Tb927.11.9180_rev | GGGGACCACTTTGTACAAGAAAGCTGGGTTTACTGAGGAGAAAAGCATTTGGAGG |
| 2_known | B19 | Tb927.9.5190_for | GGGGACAAGTTTGTACAAAAAAGCAGGCTATGCTTGAGGCTCAGGTTCTGC | Tb927.9.5190_rev | GGGGACCACTTTGTACAAGAAAGCTGGGTTTACTCGGCGTCGTCCACC |
| 2_known | B20 | Tb927.7.1900_for | GGGGACAAGTTTGTACAAAAAAGCAGGCTATGGAGAAGTACCTCCTTGGCC | Tb927.7.1900_rev | GGGGACCACTTTGTACAAGAAAGCTGGGTTCAGTACTCGTTGAAATATGTATGACAAAG |
| 2_known | B21 | Tb927.2.1890_for | GGGGACAAGTTTGTACAAAAAAGCAGGCTATGAACAAACAAAGTTTGTATGAAGGTTTC | Tb927.2.1890_rev | GGGGACCACTTTGTACAAGAAAGCTGGGTTTAAATGTCGATACCTGTCGACAAATCA |
| 2_known | B22 | Tb927.10.4930_for | GGGGACAAGTTTGTACAAAAAAGCAGGCTATGGGGGAAACGCTGTCCA | Tb927.10.4930_rev | GGGGACCACTTTGTACAAGAAAGCTGGGTTCAACGCTCACTACAAAAGAACGC |
| 2_known | B23 | Tb927.7.1880_for | GGGGACAAGTTTGTACAAAAAAGCAGGCTATGGTTGGCGAGAGGACCG | Tb927.7.1880_rev | GGGGACCACTTTGTACAAGAAAGCTGGGTTCATTCCCACTCCCCACACTC |
| 2_known | C1 | Tb927.11.11510_for | GGGGACAAGTTTGTACAAAAAAGCAGGCTATGGCAAACTGTACGACTCAGGC | Tb927.11.11510_rev | GGGGACCACTTTGTACAAGAAAGCTGGGTTTACGCGGTTCTCCCAAGCC |
| 2_known | C2 | Tb927.11.440_for | GGGGACAAGTTTGTACAAAAAAGCAGGCTATGGGACGCAATAAGAAAAAGAGCA | Tb927.11.440_rev | GGGGACCACTTTGTACAAGAAAGCTGGGTCTAGTTTCCGTAACATTTGACCTTCTTCT |
| 2_known | C3 | Tb927.8.1030_for | GGGGACAAGTTTGTACAAAAAAGCAGGCTATGCGGACAGAGGGCCTC | Tb927.8.1030_rev | GGGGACCACTTTGTACAAGAAAGCTGGGTTCACCTCAATTCACGTTTGGAAAAAGA |
| 2_known | C4 | Tb927.11.16600_for | GGGGACAAGTTTGTACAAAAAAGCAGGCTATGTCACTTCCACCCGCCA | Tb927.11.16600_rev | GGGGACCACTTTGTACAAGAAAGCTGGGTTTATCCTTCCTTTAGGGGATGTGGG |
| 2_known | C5 | Tb927.11.10820_for | GGGGACAAGTTTGTACAAAAAAGCAGGCTATGAACTACCCGATGTATGTTGGC | Tb927.11.10820_rev | GGGGACCACTTTGTACAAGAAAGCTGGGTTTATCTCTTTGGTGTCTCCTCAGGC |
| 2_known | C6 | Tb927.10.8400_for | GGGGACAAGTTTGTACAAAAAAGCAGGCTATGCCGCCGACCCCATTA | Tb927.10.8400_rev | GGGGACCACTTTGTACAAGAAAGCTGGGTTTAGTCACTTAACTCCTGTGTAACCTTGT |
| 2_known | C7 | Tb927.4.3640_for | GGGGACAAGTTTGTACAAAAAAGCAGGCTATGACCCTAGTACAGACACTGCTG | Tb927.4.3640_rev | GGGGACCACTTTGTACAAGAAAGCTGGGTTTAGCGAGCTGCAATGATAGCAA |
| 2_known | C8 | Tb927.4.3620_for | GGGGACAAGTTTGTACAAAAAAGCAGGCTATGACCCTAGTACAGACACTGCTG | Tb927.4.3620_rev | GGGGACCACTTTGTACAAGAAAGCTGGGTTCAGCGAGCAGGTATAATGAGAAAGG |
| 2_known | C9 | Tb927.11.14870_for | GGGGACAAGTTTGTACAAAAAAGCAGGCTATGCCGAAGGAGCCCATTCT | Tb927.11.14870_rev | GGGGACCACTTTGTACAAGAAAGCTGGGTCTACAGGTTCACAATACCGTTTAACTTATG |
| 2_known | C10 | Tb927.6.2670_for | GGGGACAAGTTTGTACAAAAAAGCAGGCTATGCCAAACGAAACCGCGG | Tb927.6.2670_rev | GGGGACCACTTTGTACAAGAAAGCTGGGTTCACTCGCAGGAGGTGGG |
| 2_known | C11 | Tb927.5.1470_for | GGGGACAAGTTTGTACAAAAAAGCAGGCTATGGTGCGACTCGCCTTATCT | Tb927.5.1470_rev | GGGGACCACTTTGTACAAGAAAGCTGGGTTCAGTATTTAAACACCTGCGACTCC |
| 2_known | C13 | Tb11.02.5410_for | GGGGACAAGTTTGTACAAAAAAGCAGGCTATGAATGCGGACGAAAAAACGC | Tb11.02.5410_rev | GGGGACCACTTTGTACAAGAAAGCTGGGTCTACGGGTTGCGCAGGTG |
| 2_known | C14 | Tb927.1.1660_for | GGGGACAAGTTTGTACAAAAAAGCAGGCTATGCACAATAGTGTTCCAGCATCC | Tb927.1.1660_rev | GGGGACCACTTTGTACAAGAAAGCTGGGTTCAAAATATAAGAAAATCAGGCCGTCCA |
| 2_known | C15 | Tb927.8.1920_for | GGGGACAAGTTTGTACAAAAAAGCAGGCTATGCCTGGATTGCTAATATCCCGG | Tb927.8.1920_rev | GGGGACCACTTTGTACAAGAAAGCTGGGTTCATCTCCGTCGGTGAATGAAAAAA |
| 2_known | C16 | Tb927.10.4310_for | GGGGACAAGTTTGTACAAAAAAGCAGGCTATGTCCCGGGCACCTCCA | Tb927.10.4310_rev | GGGGACCACTTTGTACAAGAAAGCTGGGTCTAATGTCCAATAGTTTGGGTGTTCAGA |
| 2_known | C17 | Tb927.10.340_for | GGGGACAAGTTTGTACAAAAAAGCAGGCTATGTCGGTACGGGAACGAGC | Tb927.10.340_rev | GGGGACCACTTTGTACAAGAAAGCTGGGTTTATAGCTTCAGGAAGAATTCACACACC |
| 2_known | C18 | Tb927.6.1080_for | GGGGACAAGTTTGTACAAAAAAGCAGGCTATGGAAGTTGTAGTGAAGAGCATCG | Tb927.6.1080_rev | GGGGACCACTTTGTACAAGAAAGCTGGGTTTACGGACACGTATTATAGAGGAAGTCC |
| 2_known | C19 | Tb927.10.12990_for | GGGGACAAGTTTGTACAAAAAAGCAGGCTATGAGTGATCACCTGTCGGAAATATGC | Tb927.10.12990_rev | GGGGACCACTTTGTACAAGAAAGCTGGGTTTAAACCTCCCAGAATTCTTTTCCCTC |
| 2_known | C20 | Tb927.6.1640_for | GGGGACAAGTTTGTACAAAAAAGCAGGCTATGAAGAATCCATCACTTTCATCACTTTGC | Tb927.6.1640_rev | GGGGACCACTTTGTACAAGAAAGCTGGGTTTATTTCCCCTTCTTTCGGAGCAC |
| 2_known | C21 | Tb927.6.3120_for | GGGGACAAGTTTGTACAAAAAAGCAGGCTATGCAAAAAAATCAGACTGCATATTATCGT | Tb927.6.3120_rev | GGGGACCACTTTGTACAAGAAAGCTGGGTTTATGATCTTCTTGTTTTTGCGGTCAC |
| 2_known | C22 | Tb927.8.870_for | GGGGACAAGTTTGTACAAAAAAGCAGGCTATGAGAGGTACGAAACGCATTGG | Tb927.8.870_rev | GGGGACCACTTTGTACAAGAAAGCTGGGTCTAGACGGTTTGTACCGCATTTTGC |
| 2_known | C23 | Tb927.9.15510_for | GGGGACAAGTTTGTACAAAAAAGCAGGCTATGAGCATCACTTTTAATAATTTATGGCTA | Tb927.9.15510_rev | GGGGACCACTTTGTACAAGAAAGCTGGGTTTACAGTACAGAACCGAGAAACAATCC |
| 2_known | C24 | Tb927.9.14160_for | GGGGACAAGTTTGTACAAAAAAGCAGGCTATGTTCCGCAGGTCCTGCA | Tb927.9.14160_rev | GGGGACCACTTTGTACAAGAAAGCTGGGTCTATAGTTTTCCCAAATATATCGTCTTATC |
| 2_known | D1 | Tb927.10.13970_for | GGGGACAAGTTTGTACAAAAAAGCAGGCTATGGTGCAACGGACACTGTTT | Tb927.10.13970_rev | GGGGACCACTTTGTACAAGAAAGCTGGGTCTACTTTCCCTTACTGAGGCATACGG |
| 2_known | D2 | Tb927.3.3970_for | GGGGACAAGTTTGTACAAAAAAGCAGGCTATGGTTTCCTTCACTTGTAGCGTC | Tb927.3.3970_rev | GGGGACCACTTTGTACAAGAAAGCTGGGTTTATCCCGCCTCTTCATCTCGC |
| 2_known | D3 | Tb927.8.3400_for | GGGGACAAGTTTGTACAAAAAAGCAGGCTATGCAACAGAACATTCACTCTTTCACA | Tb927.8.3400_rev | GGGGACCACTTTGTACAAGAAAGCTGGGTTCAAACAAGTGAAGCCGCCTC |
| 2_known | D4 | Tb927.9.7770_for | GGGGACAAGTTTGTACAAAAAAGCAGGCTATGCCTGGACCCGGTCTTC | Tb927.9.7770_rev | GGGGACCACTTTGTACAAGAAAGCTGGGTTCAAGAATAGTTTTGGTTGAGATGGCG |
| 2_known | D5 | Tb927.7.1600_for | GGGGACAAGTTTGTACAAAAAAGCAGGCTATGCCTGAGGCGGTTGTAATTG | Tb927.7.1600_rev | GGGGACCACTTTGTACAAGAAAGCTGGGTTCACAAGGCACGAACGCTTC |
| 2_known | D6 | Tb927.3.1840_for | GGGGACAAGTTTGTACAAAAAAGCAGGCTATGAAGGTTACCGTGATATCAGGCT | Tb927.3.1840_rev | GGGGACCACTTTGTACAAGAAAGCTGGGTCTACAGAAGGAATGGTAACATGGCC |
| 2_known | D7 | Tb927.10.4680_for | GGGGACAAGTTTGTACAAAAAAGCAGGCTATGGCTATGGGAAGAGTTTCCGT | Tb927.10.4680_rev | GGGGACCACTTTGTACAAGAAAGCTGGGTCTAACGCTTTCCAAGTGGGTGC |
| 2_known | D8 | Tb927.3.2410_for | GGGGACAAGTTTGTACAAAAAAGCAGGCTATGCAACCGGCCACTGCA | Tb927.3.2410_rev | GGGGACCACTTTGTACAAGAAAGCTGGGTCTACGACGCACCGCTGTC |
| 2_known | D9 | Tb927.10.13310_for | GGGGACAAGTTTGTACAAAAAAGCAGGCTATGGATTCCGTCAAAAGTTTCAAGATGA | Tb927.10.13310_rev | GGGGACCACTTTGTACAAGAAAGCTGGGTCTAAATAACATGTCGATACATTTCGTACGT |
| 2_known | D10 | Tb927.10.2250_for | GGGGACAAGTTTGTACAAAAAAGCAGGCTATGTCGATAGATGTGGTTCCTGCT | Tb927.10.2250_rev | GGGGACCACTTTGTACAAGAAAGCTGGGTTCACAACCCTCCAAATTCCTCATTG |
| 2_known | D11 | Tb927.6.1280_for | GGGGACAAGTTTGTACAAAAAAGCAGGCTATGGAAGTGCCCCATGATGACT | Tb927.6.1280_rev | GGGGACCACTTTGTACAAGAAAGCTGGGTCTACTGGCATTTCCCCGGC |
| 2_known | D12 | Tb927.10.7500_for | GGGGACAAGTTTGTACAAAAAAGCAGGCTATGCGAGGTGGGTTTGGACG | Tb927.10.7500_rev | GGGGACCACTTTGTACAAGAAAGCTGGGTTTATTGTTGTACTGCTTTCTTCATAGAGCG |
| 2_known | D13 | Tb927.6.1570_for | GGGGACAAGTTTGTACAAAAAAGCAGGCTATGTCGCTGCGAGTTGGC | Tb927.6.1570_rev | GGGGACCACTTTGTACAAGAAAGCTGGGTTCATGGCTTGGTCTCCTTAATCTTGC |
| 2_known | D14 | Tb927.10.4460_for | GGGGACAAGTTTGTACAAAAAAGCAGGCTATGGCGCTTCAGCAAACGG | Tb927.10.4460_rev | GGGGACCACTTTGTACAAGAAAGCTGGGTTCACGTCCACAACGACGTG |
| 2_known | D15 | Tb927.1.4830_for | GGGGACAAGTTTGTACAAAAAAGCAGGCTATGTCGGAACCCGTTGATCAGT | Tb927.1.4830_rev | GGGGACCACTTTGTACAAGAAAGCTGGGTTCATGAAGAACCGAACCCTCCG |
| 2_known | D16 | Tb927.1.1980_for | GGGGACAAGTTTGTACAAAAAAGCAGGCTATGGACTCCTTTGGAAACCAGAAAC | Tb927.1.1980_rev | GGGGACCACTTTGTACAAGAAAGCTGGGTCTACTGAAAGGGCCGTATCGC |
| 2_known | D17 | Tb927.6.2740_for | GGGGACAAGTTTGTACAAAAAAGCAGGCTATGTCAGAGAAGACGGTTTTATCGATTC | Tb927.6.2740_rev | GGGGACCACTTTGTACAAGAAAGCTGGGTTCATGAAATTGGTTTAACGTCAACAACTG |
| 2_known | D18 | Tb927.10.16160_for | GGGGACAAGTTTGTACAAAAAAGCAGGCTATGAGCGACAGTCAAATGGACTG | Tb927.10.16160_rev | GGGGACCACTTTGTACAAGAAAGCTGGGTTTAACTTCTCAGACTACAATGCTCAGAAAC |
| 2_known | D19 | Tb927.3.2310_for | GGGGACAAGTTTGTACAAAAAAGCAGGCTATGAGTTACGAGATACAACCCATATTGAAG | Tb927.3.2310_rev | GGGGACCACTTTGTACAAGAAAGCTGGGTTCAACTGAAAATACAGCTCTCATACGTC |
| 2_known | D20 | Tb927.10.8150_for | GGGGACAAGTTTGTACAAAAAAGCAGGCTATGTCCGATACTACTAGAAGAGTGAAGC | Tb927.10.8150_rev | GGGGACCACTTTGTACAAGAAAGCTGGGTTCAGTTTTCCACCCTCGCCT |
| 2_known | D21 | Tb927.1.1580_for | GGGGACAAGTTTGTACAAAAAAGCAGGCTATGTCACTTCGGCGCCAC | Tb927.1.1580_rev | GGGGACCACTTTGTACAAGAAAGCTGGGTTCAACCGCGTGTGTCCTTAAATC |
| 2_known | D22 | Tb927.3.2230_for | GGGGACAAGTTTGTACAAAAAAGCAGGCTATGCTCTCAAGGACAGTTCGCC | Tb927.3.2230_rev | GGGGACCACTTTGTACAAGAAAGCTGGGTTTAGATTGCCCTCTTTTTCATTAACTCAGC |
| 2_known | D23 | Tb927.7.4930_for | GGGGACAAGTTTGTACAAAAAAGCAGGCTATGGGAAAGAAAAGGTTTTACGCGG | Tb927.7.4930_rev | GGGGACCACTTTGTACAAGAAAGCTGGGTTTAGCCAAAGGGCGATTCACAG |
| 2_known | D24 | Tb927.10.1070_for | GGGGACAAGTTTGTACAAAAAAGCAGGCTATGGGGAGTCGTTACGAGCG | Tb927.10.1070_rev | GGGGACCACTTTGTACAAGAAAGCTGGGTTTAGAACTCGACAGAAAAGTATGGGTG |
| 2_known | E1 | Tb927.7.4160_for | GGGGACAAGTTTGTACAAAAAAGCAGGCTATGTTGATGAACTTCGGGGGC | Tb927.7.4160_rev | GGGGACCACTTTGTACAAGAAAGCTGGGTTTACTCTTTTTTCTTTATTTCCGCTGCTAC |
| 2_known | E2 | Tb927.7.4670_for | GGGGACAAGTTTGTACAAAAAAGCAGGCTATGAGTGTCCCGACGTGCA | Tb927.7.4670_rev | GGGGACCACTTTGTACAAGAAAGCTGGGTTCAATAAATTGTACGCTTACGGGTGT |
| 2_known | E3 | Tb927.3.1540_for | GGGGACAAGTTTGTACAAAAAAGCAGGCTATGAGTGATGTCGGCATCTGC | Tb927.3.1540_rev | GGGGACCACTTTGTACAAGAAAGCTGGGTTTATGTTGACCTTCTAGCAACTGTTGAA |
| 2_known | E4 | Tb927.2.4870_for | GGGGACAAGTTTGTACAAAAAAGCAGGCTATGGTGAGACGGGGTGCA | Tb927.2.4870_rev | GGGGACCACTTTGTACAAGAAAGCTGGGTTCACGCAACACCCAAATCACG |
| 2_known | E5 | Tb927.10.4980_for | GGGGACAAGTTTGTACAAAAAAGCAGGCTATGAGCATCACCATAAAGTTGACCAA | Tb927.10.4980_rev | GGGGACCACTTTGTACAAGAAAGCTGGGTTTACCCATTTAAAAGCCTTTCGATCGC |
| 2_known | E6 | Tb927.10.12540_for | GGGGACAAGTTTGTACAAAAAAGCAGGCTATGAGGCGTTTTTCTAACTGCTTGC | Tb927.10.12540_rev | GGGGACCACTTTGTACAAGAAAGCTGGGTCTATTTTCCTAGCGTTACTGTTCCTCTATC |
| 2_known | E7 | Tb927.10.14280_for | GGGGACAAGTTTGTACAAAAAAGCAGGCTATGGACTCGTTTGTGGCAGGT | Tb927.10.14280_rev | GGGGACCACTTTGTACAAGAAAGCTGGGTTTATTTTTCATGGAAACGATAACAACTACA |
| 2_known | E8 | Tb927.11.16970_for | GGGGACAAGTTTGTACAAAAAAGCAGGCTATGGACACGATAGACCAGTTCCTG | Tb927.11.16970_rev | GGGGACCACTTTGTACAAGAAAGCTGGGTTTACAAGAAATAATCCGGCACAGTAGG |
| 2_known | E9 | Tb927.11.8090_for | GGGGACAAGTTTGTACAAAAAAGCAGGCTATGTCTGTAGATGTCGATGCCATTATC | Tb927.11.8090_rev | GGGGACCACTTTGTACAAGAAAGCTGGGTCTACTGAAAGTACTTGGGCTTCTTTACG |
| 2_known | E10 | Tb927.10.14750_for | GGGGACAAGTTTGTACAAAAAAGCAGGCTATGCGCGGTGGTAGAGGG | Tb927.10.14750_rev | GGGGACCACTTTGTACAAGAAAGCTGGGTCTATTGCGGGGGGGATTTGTAGT |
| 2_known | E11 | Tb927.10.12840_for | GGGGACAAGTTTGTACAAAAAAGCAGGCTATGTCGAAAGAGACAAAGGCGC | Tb927.10.12840_rev | GGGGACCACTTTGTACAAGAAAGCTGGGTCTACTGCTTTGCCTGAAGCTTGG |
| 2_known | E12 | Tb927.11.15810_for | GGGGACAAGTTTGTACAAAAAAGCAGGCTATGATGCGCCGTTGCTTTCC | Tb927.11.15810_rev | GGGGACCACTTTGTACAAGAAAGCTGGGTCTATCTCCTGGCAATATTACTACCTGCA |
| 2_known | E13 | Tb927.11.10340_for | GGGGACAAGTTTGTACAAAAAAGCAGGCTATGACGGTGTTGTCACGGGT | Tb927.11.10340_rev | GGGGACCACTTTGTACAAGAAAGCTGGGTTTACGGAACGTTATGAGTTGTGACC |
| 2_known | E14 | Tb927.9.7070_for | GGGGACAAGTTTGTACAAAAAAGCAGGCTATGAGTGAAACAAACCCAGTAGTGAG | Tb927.9.7070_rev | GGGGACCACTTTGTACAAGAAAGCTGGGTTTACTCCTTGGGAAAATAACCGCG |
| 2_known | E15 | Tb927.10.14160_for | GGGGACAAGTTTGTACAAAAAAGCAGGCTATGCAGAGCCAACCAGACAATG | Tb927.10.14160_rev | GGGGACCACTTTGTACAAGAAAGCTGGGTTTAGTGTGGCACAAAATATTTGTACAGAAA |
| 2_known | E16 | Tb927.9.13990_for | GGGGACAAGTTTGTACAAAAAAGCAGGCTATGCAAGGAGGAAATATGTTCGACG | Tb927.9.13990_rev | GGGGACCACTTTGTACAAGAAAGCTGGGTTCATGAAGTGGATTTATTGTCAAGACGG |
| 2_known | E17 | Tb927.9.13030_for | GGGGACAAGTTTGTACAAAAAAGCAGGCTATGGCGACCCGTGACCGT | Tb927.9.13030_rev | GGGGACCACTTTGTACAAGAAAGCTGGGTTTAAGACAGCATCTTTTGAAACAGAAGAAT |
| 2_known | E18 | Tb927.3.4820_for | GGGGACAAGTTTGTACAAAAAAGCAGGCTATGGCGATCGTCGTGACGA | Tb927.3.4820_rev | GGGGACCACTTTGTACAAGAAAGCTGGGTTTAGGTAGCTTTTTCCGTGGAGGC |
| 2_known | E19 | Tb927.10.3030_for | GGGGACAAGTTTGTACAAAAAAGCAGGCTATGTTCCATCCCCAATTGGTGGC | Tb927.10.3030_rev | GGGGACCACTTTGTACAAGAAAGCTGGGTTCAGAAGACAACCCCATTAATCATCG |
| 2_known | E20 | Tb927.10.180_for | GGGGACAAGTTTGTACAAAAAAGCAGGCTATGTCAGGTAAGCTTCGTCTTTACAAA | Tb927.10.180_rev | GGGGACCACTTTGTACAAGAAAGCTGGGTCTACTTGGTTACTGCCCCTTCCC |
| 2_known | E22 | Tb927.11.7360_for | GGGGACAAGTTTGTACAAAAAAGCAGGCTATGTCTAGTCTACCTTCAGATTCAGCG | Tb927.11.7360_rev | GGGGACCACTTTGTACAAGAAAGCTGGGTCTATACCATGTGAACATCGGATGCG |
| 2_known | E23 | Tb927.7.4180_for | GGGGACAAGTTTGTACAAAAAAGCAGGCTATGTTTTTTACTCCTCCACAACTTCAGA | Tb927.7.4180_rev | GGGGACCACTTTGTACAAGAAAGCTGGGTTTATTTAACTTTCTTCTCGACAGAACCGT |
| 2_known | E24 | Tb927.1.1060_for | GGGGACAAGTTTGTACAAAAAAGCAGGCTATGTCTGTTCAGTGCCAGGTTG | Tb927.1.1060_rev | GGGGACCACTTTGTACAAGAAAGCTGGGTTTACTGATCGTGCAGATTGTATATGGG |
| 2_known | F1 | Tb927.7.820_for | GGGGACAAGTTTGTACAAAAAAGCAGGCTATGGCAACCTTGAACGCACC | Tb927.7.820_rev | GGGGACCACTTTGTACAAGAAAGCTGGGTTCACATGCCTGGTGTCGGG |
| 2_known | F2 | Tb927.10.6690_for | GGGGACAAGTTTGTACAAAAAAGCAGGCTATGTCCACAGCGAAGAGTTTTCC | Tb927.10.6690_rev | GGGGACCACTTTGTACAAGAAAGCTGGGTCTACGCTTTTAAGTTAAGTGTCGACAC |
| 2_known | F3 | Tb927.11.3750_for | GGGGACAAGTTTGTACAAAAAAGCAGGCTATGATGCTCGCAGTTATCGTAGC | Tb927.11.3750_rev | GGGGACCACTTTGTACAAGAAAGCTGGGTTTAAAAGGTAAATATATTGTCTTGTGAGTA |
| 2_known | F4 | Tb927.10.7790_for | GGGGACAAGTTTGTACAAAAAAGCAGGCTATGTATCCTGGAAAGGAGAGCCC | Tb927.10.7790_rev | GGGGACCACTTTGTACAAGAAAGCTGGGTTCAGTTTTTTGCTGCGTTGAATCTG |
| 2_known | F5 | Tb927.11.11940_for | GGGGACAAGTTTGTACAAAAAAGCAGGCTATGAGCGACTTAAACGCCATGAT | Tb927.11.11940_rev | GGGGACCACTTTGTACAAGAAAGCTGGGTTTATAGGAAATACTCCGGTGGCGT |
| 2_known | F6 | Tb927.8.2780_for | GGGGACAAGTTTGTACAAAAAAGCAGGCTATGGGAGACTCGATATCACCTTCG | Tb927.8.2780_rev | GGGGACCACTTTGTACAAGAAAGCTGGGTTCACTCCATTCGAACCGGAGG |
| 2_known | F7 | Tb927.8.3140_for | GGGGACAAGTTTGTACAAAAAAGCAGGCTATGGCTGACCGCCTTGCT | Tb927.8.3140_rev | GGGGACCACTTTGTACAAGAAAGCTGGGTTTACTTCGTGCCGCCCAAC |
| 2_known | F8 | Tb927.7.2750_for | GGGGACAAGTTTGTACAAAAAAGCAGGCTATGGGAATAGTAGGAGAGTATGAAATATTA | Tb927.7.2750_rev | GGGGACCACTTTGTACAAGAAAGCTGGGTTTAGCTGTTTGCAACTTGTTTCTCCT |
| 2_known | F9 | Tb927.6.3730_for | GGGGACAAGTTTGTACAAAAAAGCAGGCTATGCGCTCGTCGTTTTTTTTGC | Tb927.6.3730_rev | GGGGACCACTTTGTACAAGAAAGCTGGGTCTAACCACCAGCCACAATGCTG |
| 2_known | F10 | Tb927.9.11040_for | GGGGACAAGTTTGTACAAAAAAGCAGGCTATGGATGTAATATGCTCCGCTCTTG | Tb927.9.11040_rev | GGGGACCACTTTGTACAAGAAAGCTGGGTTTACGCCATATTACCTTTAAGATAGCTGTA |
| 2_known | F11 | Tb927.3.2400_for | GGGGACAAGTTTGTACAAAAAAGCAGGCTATGCTCAGCTTTCTTTCCCCCA | Tb927.3.2400_rev | GGGGACCACTTTGTACAAGAAAGCTGGGTTCACACCTCCGCCTCTTCAA |
| 2_known | F12 | Tb927.10.14830_for | GGGGACAAGTTTGTACAAAAAAGCAGGCTATGACGGATAAAAAGCGGGAACC | Tb927.10.14830_rev | GGGGACCACTTTGTACAAGAAAGCTGGGTCTAATTCGATCTGCGCCACTCC |
| 2_known | F13 | Tb927.11.9990_for | GGGGACAAGTTTGTACAAAAAAGCAGGCTATGTCACTACCAAGAAGAATGTGCATG | Tb927.11.9990_rev | GGGGACCACTTTGTACAAGAAAGCTGGGTCTACGGTACAAGGGTTCCAGTCT |
| 2_known | F14 | Tb927.10.14680_for | GGGGACAAGTTTGTACAAAAAAGCAGGCTATGGTTAAGAGGGAGAAACGTCAGC | Tb927.10.14680_rev | GGGGACCACTTTGTACAAGAAAGCTGGGTTCAATCGCGGAATACTGAGTCCA |
| 2_known | F15 | Tb927.7.2730_for | GGGGACAAGTTTGTACAAAAAAGCAGGCTATGATAGAAGAAGTTGAATCTGTGTACAAG | Tb927.7.2730_rev | GGGGACCACTTTGTACAAGAAAGCTGGGTCTACGAAGCGTTCTTTTTCCTGGA |
| 2_known | F16 | Tb927.9.11280_for | GGGGACAAGTTTGTACAAAAAAGCAGGCTATGCTTCGGCGATCACTCTTAGC | Tb927.9.11280_rev | GGGGACCACTTTGTACAAGAAAGCTGGGTCTAAGGAGACTTCTTAAAGCTCTTGAAGTC |
| 2_known | F17 | Tb927.10.9300_for | GGGGACAAGTTTGTACAAAAAAGCAGGCTATGGAATCCACAGTCACCGGT | Tb927.10.9300_rev | GGGGACCACTTTGTACAAGAAAGCTGGGTTTAACAAATGTGTCGGCTCACCA |
| 2_known | F20 | Tb927.11.14460_for | GGGGACAAGTTTGTACAAAAAAGCAGGCTATGTCACATACTGCTGAGGATGCC | Tb927.11.14460_rev | GGGGACCACTTTGTACAAGAAAGCTGGGTTTAGTCTTTCTTTGATTGGAAGTCCCC |
| 2_known | F21 | Tb927.11.1350_for | GGGGACAAGTTTGTACAAAAAAGCAGGCTATGTGGAGGCCTTATACTTTGTTTGTT | Tb927.11.1350_rev | GGGGACCACTTTGTACAAGAAAGCTGGGTTCAGTGATGTTTAAACCACTTCCGC |
| 2_known | F22 | Tb927.8.3330_for | GGGGACAAGTTTGTACAAAAAAGCAGGCTATGTCTCGCACCATAAAGAAGGAAAA | Tb927.8.3330_rev | GGGGACCACTTTGTACAAGAAAGCTGGGTCTACATGCAAGTGGTGAAAAGCTCA |
| 2_known | F23 | Tb927.10.16170_for | GGGGACAAGTTTGTACAAAAAAGCAGGCTATGAGGTTCATCAACCAAGCAGC | Tb927.10.16170_rev | GGGGACCACTTTGTACAAGAAAGCTGGGTTCAGTCCACCGTCCAAATATTAACTG |
| 2_known | F24 | Tb927.10.3620_for | GGGGACAAGTTTGTACAAAAAAGCAGGCTATGATGTTTCACACATTCACTGCGC | Tb927.10.3620_rev | GGGGACCACTTTGTACAAGAAAGCTGGGTTTATGGGGCCGCGGCACT |
| 2_known | G1 | Tb927.7.2530_for | GGGGACAAGTTTGTACAAAAAAGCAGGCTATGAACAATAAAAATGTGTTAGTTTCTGGC | Tb927.7.2530_rev | GGGGACCACTTTGTACAAGAAAGCTGGGTTTACGTCTGCTCGTTTTCAGGAAC |
| 2_known | G2 | Tb927.9.15110_for | GGGGACAAGTTTGTACAAAAAAGCAGGCTATGACATTCGTTAAGGTTGTGAAGAACA | Tb927.9.15110_rev | GGGGACCACTTTGTACAAGAAAGCTGGGTTTACTTCGCACGGTCACGGA |
| 2_known | G3 | Tb927.7.2360_for | GGGGACAAGTTTGTACAAAAAAGCAGGCTATGCCAAAAAAAGGGAAAAAGAATGCG | Tb927.7.2360_rev | GGGGACCACTTTGTACAAGAAAGCTGGGTTCAGATAGGAGAAGTAGAGGTAGAAGGA |
| 2_known | G4 | Tb927.11.16980_for | GGGGACAAGTTTGTACAAAAAAGCAGGCTATGGGGTCAGATGTGTTTGAGCT | Tb927.11.16980_rev | GGGGACCACTTTGTACAAGAAAGCTGGGTCTATCTACTTCGGTCATAGTTTCCCAGG |
| 2_known | G6 | Tb927.11.15550_for | GGGGACAAGTTTGTACAAAAAAGCAGGCTATGAAATTCCTTACTTTTACTGCTGCCT | Tb927.11.15550_rev | GGGGACCACTTTGTACAAGAAAGCTGGGTTTAGAAACGATAGACGTGATCTTTTGTATA |
| 2_known | G7 | Tb927.11.5140_for | GGGGACAAGTTTGTACAAAAAAGCAGGCTATGCCCAGTTGGTGTCTTATTGAAA | Tb927.11.5140_rev | GGGGACCACTTTGTACAAGAAAGCTGGGTCTACTGCTTAGCCCCTTGACCA |
| 2_known | G8 | Tb927.5.1200_for | GGGGACAAGTTTGTACAAAAAAGCAGGCTATGGCAGTCATAGTAAAGACTGGTAAGT | Tb927.5.1200_rev | GGGGACCACTTTGTACAAGAAAGCTGGGTTTAGACTAACCGAATTAGCGGGCA |
| 2_known | G9 | Tb927.11.15820_for | GGGGACAAGTTTGTACAAAAAAGCAGGCTATGCGTCGCGTGGCTTCA | Tb927.11.15820_rev | GGGGACCACTTTGTACAAGAAAGCTGGGTTTACCAGAACTTCATCTCGTAGTAGTCTT |
| 2_known | G10 | Tb927.11.8220_for | GGGGACAAGTTTGTACAAAAAAGCAGGCTATGAGGTCAACTGAGGTCGGG | Tb927.11.8220_rev | GGGGACCACTTTGTACAAGAAAGCTGGGTTTAATTCTCTTTCCCTGCAGTTGGC |
| 2_known | G11 | Tb927.7.7040_for | GGGGACAAGTTTGTACAAAAAAGCAGGCTATGATGTACACGAGTCCCCACG | Tb927.7.7040_rev | GGGGACCACTTTGTACAAGAAAGCTGGGTTTACGGAGCGAATATGGGATATTTTGC |
| 2_known | G12 | Tb927.10.6080_for | GGGGACAAGTTTGTACAAAAAAGCAGGCTATGTTGGCAGATTTTGAAAGTGTTCTC | Tb927.10.6080_rev | GGGGACCACTTTGTACAAGAAAGCTGGGTCTATGCACTCTGAGAAGAATAACGGTC |
| 2_known | G13 | Tb927.4.4360_for | GGGGACAAGTTTGTACAAAAAAGCAGGCTATGGGTTGCTGTTCCTGCTG | Tb927.4.4360_rev | GGGGACCACTTTGTACAAGAAAGCTGGGTCTACTGGCAATGTGCATTAAGAAATTTTAA |

| | | | | | |
|---|---|---|---|---|---|
| 2_known | G14 | Tb927.7.6960_for | GGGGACAAGTTTGTACAAAAAAGCAGGCTATGAACGTACAGAAGGCCATTCAG | Tb927.7.6960_rev | GGGGACCACTTTGTACAAGAAAGCTGGGTTTATCCCCAAAGTGCTTTCTTCACG |
| 2_known | G15 | Tb927.10.11870_for | GGGGACAAGTTTGTACAAAAAAGCAGGCTATGCTGCGCCACACATCAC | Tb927.10.11870_rev | GGGGACCACTTTGTACAAGAAAGCTGGGTTTATTTCTGCAGTTGATGCGTCTGC |
| 2_known | G16 | Tb927.11.270_for | GGGGACAAGTTTGTACAAAAAAGCAGGCTATGCATGATGGAAACTTTCTGAAGGA | Tb927.11.270_rev | GGGGACCACTTTGTACAAGAAAGCTGGGTTTATCCATTCAAACCGCTGCACA |
| 2_known | G17 | Tb927.5.1550_for | GGGGACAAGTTTGTACAAAAAAGCAGGCTATGGCACTCCCGACATCGC | Tb927.5.1550_rev | GGGGACCACTTTGTACAAGAAAGCTGGGTTTATGCGGGAGCGCAAGAAAC |
| 2_known | G18 | Tb927.5.1210_for | GGGGACAAGTTTGTACAAAAAAGCAGGCTATGTTATTTGACTGCACTTTGCTGTC | Tb927.5.1210_rev | GGGGACCACTTTGTACAAGAAAGCTGGGTTTACTGTTTCGCCTTGCGCT |
| 2_known | G19 | Tb927.3.5630_for | GGGGACAAGTTTGTACAAAAAAGCAGGCTATGGCTGCAACACCTTACAGGT | Tb927.3.5630_rev | GGGGACCACTTTGTACAAGAAAGCTGGGTCTACGCACTACCTGCAATGTACG |
| 2_known | G20 | Tb927.10.4990_for | GGGGACAAGTTTGTACAAAAAAGCAGGCTATGACAATGCTTGGGGCGTT | Tb927.10.4990_rev | GGGGACCACTTTGTACAAGAAAGCTGGGTCTAAAACATGGCATCACTAAACCATGG |
| 2_known | G21 | Tb927.9.9940_for | GGGGACAAGTTTGTACAAAAAAGCAGGCTATGGCGTTCTCACGAAAGGAGC | Tb927.9.9940_rev | GGGGACCACTTTGTACAAGAAAGCTGGGTTCACATGAGGCATGACTCATACGT |
| 2_known | G22 | Tb927.11.7370_for | GGGGACAAGTTTGTACAAAAAAGCAGGCTATGCCCCAATGTGGCACG | Tb927.11.7370_rev | GGGGACCACTTTGTACAAGAAAGCTGGGTCTAGCTCAGCAGAAACACTTCACG |
| 2_known | G23 | Tb927.2.2970_for | GGGGACAAGTTTGTACAAAAAAGCAGGCTATGTCATCCGAACACGCACC | Tb927.2.2970_rev | GGGGACCACTTTGTACAAGAAAGCTGGGTTTAATGCCGAACCACCCCTCT |
| 2_known | G24 | Tb927.7.3500_for | GGGGACAAGTTTGTACAAAAAAGCAGGCTATGTTCACAAAGGGGTCCATGTC | Tb927.7.3500_rev | GGGGACCACTTTGTACAAGAAAGCTGGGTTCACTTTGGGATGTATTTCTCATTACCC |
| 2_known | H1 | Tb927.10.14170_for | GGGGACAAGTTTGTACAAAAAAGCAGGCTATGCAGAGCCAACCAGACAATG | Tb927.10.14170_rev | GGGGACCACTTTGTACAAGAAAGCTGGGTTTAGTGTGGAAGAAAATATTTGTACAGAAA |
| 2_known | H2 | Tb927.10.1690_for | GGGGACAAGTTTGTACAAAAAAGCAGGCTATGAACTTGGCAAATTCTCATTGTATCG | Tb927.10.1690_rev | GGGGACCACTTTGTACAAGAAAGCTGGGTCTACTCCATCGCTAAACTTTCAATGAGT |
| 2_known | H3 | Tb927.11.5360_for | GGGGACAAGTTTGTACAAAAAAGCAGGCTATGGACCGTATTCCATATTTGGTTTCAT | Tb927.11.5360_rev | GGGGACCACTTTGTACAAGAAAGCTGGGTTTAATCTATCGTGTTGCTATGCCAACA |
| 2_known | H4 | Tb927.5.2380_for | GGGGACAAGTTTGTACAAAAAAGCAGGCTATGTTGACTACGACTGACACTATTGTTAAG | Tb927.5.2380_rev | GGGGACCACTTTGTACAAGAAAGCTGGGTTTAAGCCAGAGTCAGTTCCCGT |
| 2_known | H5 | Tb927.10.4500_for | GGGGACAAGTTTGTACAAAAAAGCAGGCTATGGAGAGCCTACGGACTGC | Tb927.10.4500_rev | GGGGACCACTTTGTACAAGAAAGCTGGGTTCAGCTGGCTTTCCGGAAGC |
| 2_known | H6 | Tb927.8.1810_for | GGGGACAAGTTTGTACAAAAAAGCAGGCTATGGCTGGTGAGGGTGCA | Tb927.8.1810_rev | GGGGACCACTTTGTACAAGAAAGCTGGGTTCACACTACAGCAGCATCGTGC |
| 2_known | H7 | Tb927.11.10020_for | GGGGACAAGTTTGTACAAAAAAGCAGGCTATGCAACTCTTATCGTGGTCTAGCG | Tb927.11.10020_rev | GGGGACCACTTTGTACAAGAAAGCTGGGTTCATTTCTGGAGAGAGCGATGACT |
| 2_known | H8 | Tb927.11.7140_for | GGGGACAAGTTTGTACAAAAAAGCAGGCTATGGCCGCAACAATTCTCAAGG | Tb927.11.7140_rev | GGGGACCACTTTGTACAAGAAAGCTGGGTCTACGAAGGTGTTTCTGGTTCCATC |
| 2_known | H9 | Tb927.5.3120_for | GGGGACAAGTTTGTACAAAAAAGCAGGCTATGGCTGAACAACCGGAAGTTT | Tb927.5.3120_rev | GGGGACCACTTTGTACAAGAAAGCTGGGTCTAAAGTGTCATGGCTTGGCGT |
| 2_known | H10 | Tb927.5.3830_for | GGGGACAAGTTTGTACAAAAAAGCAGGCTATGAGTTTGAAAGTTAATATTCTCGGACAT | Tb927.5.3830_rev | GGGGACCACTTTGTACAAGAAAGCTGGGTTCAATCCATCGTCTTCACGCG |
| 2_known | H11 | Tb927.4.2110_for | GGGGACAAGTTTGTACAAAAAAGCAGGCTATGCATCAACACGGCAAGACA | Tb927.4.2110_rev | GGGGACCACTTTGTACAAGAAAGCTGGGTCTACATATTGTGTAGAGGGTCGCCA |
| 2_known | H12 | Tb927.8.8020_for | GGGGACAAGTTTGTACAAAAAAGCAGGCTATGGGATGTTTTTGCTGCTGCT | Tb927.8.8020_rev | GGGGACCACTTTGTACAAGAAAGCTGGGTCTATTTACAGTGTGCATTAATAAACTTTAA |
| 2_known | H13 | Tb927.10.6860_for | GGGGACAAGTTTGTACAAAAAAGCAGGCTATGGGTGTGGGAGAAATGAGTGA | Tb927.10.6860_rev | GGGGACCACTTTGTACAAGAAAGCTGGGTTCAGCATTCATTTAGTTGAATCCGTCG |
| 2_known | H14 | Tb927.4.410_for | GGGGACAAGTTTGTACAAAAAAGCAGGCTATGCACCAAACCCAAGCGTATT | Tb927.4.410_rev | GGGGACCACTTTGTACAAGAAAGCTGGGTTCACGCACTGGCATCACCG |
| 2_known | H15 | Tb927.11.13440_for | GGGGACAAGTTTGTACAAAAAAGCAGGCTATGATTGATTGGGCACTCACGTT | Tb927.11.13440_rev | GGGGACCACTTTGTACAAGAAAGCTGGGTCTACCTCCATTCCTTCTGCCACC |
| 2_known | H16 | Tb927.6.1650_for | GGGGACAAGTTTGTACAAAAAAGCAGGCTATGAAGAATCCATCACTTTCATCACTTTGC | Tb927.6.1650_rev | GGGGACCACTTTGTACAAGAAAGCTGGGTTCACTGGTCTTCAGCAAACAGATG |
| 2_known | H17 | Tb927.10.12760_for | GGGGACAAGTTTGTACAAAAAAGCAGGCTATGTCATCGCAGGACAATCGC | Tb927.10.12760_rev | GGGGACCACTTTGTACAAGAAAGCTGGGTTTATGACCTGCAGAGAACAGGATAGT |
| 2_known | H18 | Tb927.9.10310_for | GGGGACAAGTTTGTACAAAAAAGCAGGCTATGAGCGCAAAGAACAAAACTTGG | Tb927.9.10310_rev | GGGGACCACTTTGTACAAGAAAGCTGGGTTTACTTCTTTCCGGAACCACCACT |
| 2_known | H19 | Tb927.9.2580_for | GGGGACAAGTTTGTACAAAAAAGCAGGCTATGGATCAGGACCTCTCTGTTTGT | Tb927.9.2580_rev | GGGGACCACTTTGTACAAGAAAGCTGGGTCTAGTGTTGGATTCTCTGATTGTTAAATTG |
| 2_known | H20 | Tb927.11.11360_for | GGGGACAAGTTTGTACAAAAAAGCAGGCTATGGCTGTCGCTTACGAAGGA | Tb927.11.11360_rev | GGGGACCACTTTGTACAAGAAAGCTGGGTTCACGCGTTCTCCGATACACC |
| 2_known | H21 | Tb927.10.2560_for | GGGGACAAGTTTGTACAAAAAAGCAGGCTATGCGTACCAGTGCAACTCTCC | Tb927.10.2560_rev | GGGGACCACTTTGTACAAGAAAGCTGGGTTCACTGTCTGGAACCGAAGGC |
| 2_known | H23 | Tb927.9.3440_for | GGGGACAAGTTTGTACAAAAAAGCAGGCTATGGCAATCGTCGGGGACT | Tb927.9.3440_rev | GGGGACCACTTTGTACAAGAAAGCTGGGTTCACTCTGTAAGGTCAATCACACCG |
| 2_known | H24 | Tb927.11.8790_for | GGGGACAAGTTTGTACAAAAAAGCAGGCTATGGACCGCTTGCTTCAGTTC | Tb927.11.8790_rev | GGGGACCACTTTGTACAAGAAAGCTGGGTTTACAATAACGACCCTAGCAAACCAG |
| 2_known | I1 | Tb927.9.3880_for | GGGGACAAGTTTGTACAAAAAAGCAGGCTATGGCTCCAGTAGTGTGGCC | Tb927.9.3880_rev | GGGGACCACTTTGTACAAGAAAGCTGGGTTCAAAACAGGTTGTGAACGCGG |
| 2_known | I2 | Tb927.8.990_for | GGGGACAAGTTTGTACAAAAAAGCAGGCTATGAGCACGGGTGATCCAACT | Tb927.8.990_rev | GGGGACCACTTTGTACAAGAAAGCTGGGTCTATGAATTGTGACGCATCACGGA |
| 2_known | I3 | Tb927.2.6210_for | GGGGACAAGTTTGTACAAAAAAGCAGGCTATGACACGCGCCTCGCTT | Tb927.2.6210_rev | GGGGACCACTTTGTACAAGAAAGCTGGGTCTACATTTTTGAGTAATCTTTCCCATATGC |
| 2_known | I4 | Tb927.2.6180_for | GGGGACAAGTTTGTACAAAAAAGCAGGCTATGACACGCGCCTCGCTT | Tb927.2.6180_rev | GGGGACCACTTTGTACAAGAAAGCTGGGTCTACATTTTAGGTTTTGCAAGCGAAAAC |
| 2_known | I6 | Tb927.10.13860_for | GGGGACAAGTTTGTACAAAAAAGCAGGCTATGTTGCCCATGTTACTGTGGC | Tb927.10.13860_rev | GGGGACCACTTTGTACAAGAAAGCTGGGTCTAGAACAAATCGTAACGTAACTCTACTGG |
| 2_known | I7 | Tb927.6.2540_for | GGGGACAAGTTTGTACAAAAAAGCAGGCTATGATTCCGTCACAACTGAAGAGC | Tb927.6.2540_rev | GGGGACCACTTTGTACAAGAAAGCTGGGTTCAAGTATGGATAGTACTACCATCTGTATC |
| 2_known | I9 | Tb927.2.5050_for | GGGGACAAGTTTGTACAAAAAAGCAGGCTATGGACTTATTTGGAGCCTATGCTTC | Tb927.2.5050_rev | GGGGACCACTTTGTACAAGAAAGCTGGGTTCATAATTTTACATTACTCGAATGATCGTC |
| 2_known | I11 | Tb927.11.13990_for | GGGGACAAGTTTGTACAAAAAAGCAGGCTATGTTAGGGTCCCTACCGAGTGT | Tb927.11.13990_rev | GGGGACCACTTTGTACAAGAAAGCTGGGTTCACTTCCGTGCGACATCGT |
| 2_known | I12 | Tb927.11.15130_for | GGGGACAAGTTTGTACAAAAAAGCAGGCTATGGCCGATCTATACGATGTTCTCG | Tb927.11.15130_rev | GGGGACCACTTTGTACAAGAAAGCTGGGTTTACTTGGCATCTAGAATACGGCTCA |
| 2_known | I13 | Tb927.5.300b_for | GGGGACAAGTTTGTACAAAAAAGCAGGCTATGGCTCACGGCTCGATTCC | Tb927.5.300b_rev | GGGGACCACTTTGTACAAGAAAGCTGGGTTCACATCTTTGTTTTGCGATGGGC |
| 2_known | I14 | Tb927.10.10830_for | GGGGACAAGTTTGTACAAAAAAGCAGGCTATGAAGCGCACACCTGTTAGG | Tb927.10.10830_rev | GGGGACCACTTTGTACAAGAAAGCTGGGTTTACACCTTCTGACTGGCATCACC |
| 2_known | I15 | Tb927.7.7230_for | GGGGACAAGTTTGTACAAAAAAGCAGGCTATGAACGTGAGCCGCTGC | Tb927.7.7230_rev | GGGGACCACTTTGTACAAGAAAGCTGGGTTCAGAAGCGATTCCATCGGACG |
| 2_known | I16 | Tb927.4.2350_for | GGGGACAAGTTTGTACAAAAAAGCAGGCTATGGCAAAAGCCCTTAGTGTCAG | Tb927.4.2350_rev | GGGGACCACTTTGTACAAGAAAGCTGGGTTTACGGTTTTCCGTCTCCACTGG |
| 2_known | I18 | Tb927.7.7500_for | GGGGACAAGTTTGTACAAAAAAGCAGGCTATGAATCACACTTCGATTCCAGTTATTGA | Tb927.7.7500_rev | GGGGACCACTTTGTACAAGAAAGCTGGGTTCACATATTTGCCTTGCGATGGG |
| 2_known | I19 | Tb927.10.6850_for | GGGGACAAGTTTGTACAAAAAAGCAGGCTATGAATCGTACGGGTGGGTCG | Tb927.10.6850_rev | GGGGACCACTTTGTACAAGAAAGCTGGGTTTACACCTTCTTGATACCCAGTGTCG |
| 2_known | I20 | Tb927.6.1520_for | GGGGACAAGTTTGTACAAAAAAGCAGGCTATGTCTGACGAAAAAATTAACGTGCAT | Tb927.6.1520_rev | GGGGACCACTTTGTACAAGAAAGCTGGGTTTAGTAGGGAACAAAGTATTTGTAGAGGAA |
| 2_known | I21 | Tb927.9.10570_for | GGGGACAAGTTTGTACAAAAAAGCAGGCTATGGCCCTTCTCGAGCATGT | Tb927.9.10570_rev | GGGGACCACTTTGTACAAGAAAGCTGGGTTTATTTCTGAGCGGCACCCTCT |
| 2_known | I22 | Tb927.9.7740_for | GGGGACAAGTTTGTACAAAAAAGCAGGCTATGACACTTCCGGGGCAACC | Tb927.9.7740_rev | GGGGACCACTTTGTACAAGAAAGCTGGGTTCACTTCGCACCTCGCCG |
| 2_known | I23 | Tb927.3.2890_for | GGGGACAAGTTTGTACAAAAAAGCAGGCTATGTCGAAGCAAAAGAATGTGGAGA | Tb927.3.2890_rev | GGGGACCACTTTGTACAAGAAAGCTGGGTCTAATCGGTCCACGTCATGCC |
| 2_known | I24 | Tb927.9.14180_for | GGGGACAAGTTTGTACAAAAAAGCAGGCTATGGATGACACTCGCATCAATTGG | Tb927.9.14180_rev | GGGGACCACTTTGTACAAGAAAGCTGGGTTCAATCAGTTAAAACACAAAGTGGAGAGG |
| 2_known | J1 | Tb927.11.7420_for | GGGGACAAGTTTGTACAAAAAAGCAGGCTATGGGAGAGGAGGTAGTTAGGTTAGA | Tb927.11.7420_rev | GGGGACCACTTTGTACAAGAAAGCTGGGTTCATGTGCTTAGCCTTCCACATC |
| 2_known | J2 | Tb927.11.1910_for | GGGGACAAGTTTGTACAAAAAAGCAGGCTATGCTTCCCGGGTTTCCCT | Tb927.11.1910_rev | GGGGACCACTTTGTACAAGAAAGCTGGGTTTATTCTACATGAACAATAATTCCGGTATT |
| 2_known | J3 | Tb927.9.9400_for | GGGGACAAGTTTGTACAAAAAAGCAGGCTATGGCTGTCCCACCAGTGG | Tb927.9.9400_rev | GGGGACCACTTTGTACAAGAAAGCTGGGTCTATGACTTTCCCTCGAAATTCATCATTTC |
| 2_known | J4 | Tb927.10.14690_for | GGGGACAAGTTTGTACAAAAAAGCAGGCTATGCCTAATTACTTACCCAACAGGGA | Tb927.10.14690_rev | GGGGACCACTTTGTACAAGAAAGCTGGGTTCATATGACGTCAAACTGCATGAGAA |
| 2_known | J5 | Tb927.3.1240_for | GGGGACAAGTTTGTACAAAAAAGCAGGCTATGCCTGCGACTACCTCAATTCC | Tb927.3.1240_rev | GGGGACCACTTTGTACAAGAAAGCTGGGTTCACAAAAAGTAGTCAGGTGTCTTTTTTG |
| 2_known | J6 | Tb927.10.15710_for | GGGGACAAGTTTGTACAAAAAAGCAGGCTATGTCTTCGGCGGCGGAA | Tb927.10.15710_rev | GGGGACCACTTTGTACAAGAAAGCTGGGTTCACGGCTGCGCGTCGCT |
| 2_known | J7 | Tb927.4.460_for | GGGGACAAGTTTGTACAAAAAAGCAGGCTATGGAAGACCCCGTGCGC | Tb927.4.460_rev | GGGGACCACTTTGTACAAGAAAGCTGGGTTCATTCGTTAAGGAACTCACACCTG |
| 2_known | J8 | Tb927.10.15410_for | GGGGACAAGTTTGTACAAAAAAGCAGGCTATGGTCAGCGTTGCCGTAATT | Tb927.10.15410_rev | GGGGACCACTTTGTACAAGAAAGCTGGGTTTACAACTTTGAGCGCGCGA |
| 2_known | J9 | Tb927.11.2060_for | GGGGACAAGTTTGTACAAAAAAGCAGGCTATGCCATCCGTTTCCCAGGA | Tb927.11.2060_rev | GGGGACCACTTTGTACAAGAAAGCTGGGTTTAGAAGAGAGCACCCATGCCA |
| 2_known | J10 | Tb927.10.12940_for | GGGGACAAGTTTGTACAAAAAAGCAGGCTATGGGGGGGAGCTCGTCG | Tb927.10.12940_rev | GGGGACCACTTTGTACAAGAAAGCTGGGTTCATACCTGGGAGCTGAGAGGC |
| 2_known | J11 | Tb927.11.16400_for | GGGGACAAGTTTGTACAAAAAAGCAGGCTATGTTGTTCCGCACCGTCAA | Tb927.11.16400_rev | GGGGACCACTTTGTACAAGAAAGCTGGGTCTAACGCCTCCTTACGGGTGG |
| 2_known | J12 | Tb927.11.14900_for | GGGGACAAGTTTGTACAAAAAAGCAGGCTATGGCGGATCCATTTTACGATGC | Tb927.11.14900_rev | GGGGACCACTTTGTACAAGAAAGCTGGGTCTATGCAGATGCCGTAAATTCTTCAATAG |
| 2_known | J13 | Tb927.11.7540_for | GGGGACAAGTTTGTACAAAAAAGCAGGCTATGTTTCGTTACACCGCTGCA | Tb927.11.7540_rev | GGGGACCACTTTGTACAAGAAAGCTGGGTTCACTTCCTTGACTTCGTAACCAGT |
| 2_known | J14 | Tb927.11.2050_for | GGGGACAAGTTTGTACAAAAAAGCAGGCTATGCCATCCGTTTCCCAGGA | Tb927.11.2050_rev | GGGGACCACTTTGTACAAGAAAGCTGGGTCTAGAAGAGAGCACCCATGCCA |
| 2_known | J15 | Tb927.10.14360_for | GGGGACAAGTTTGTACAAAAAAGCAGGCTATGAGGCTGACTCTTGACACCA | Tb927.10.14360_rev | GGGGACCACTTTGTACAAGAAAGCTGGGTTTATGATGTGCGAGTCTTCTTTGTCC |
| 2_known | J16 | Tb927.11.1180_for | GGGGACAAGTTTGTACAAAAAAGCAGGCTATGTCTGATACATTAAGTAGCAAACTAGAC | Tb927.11.1180_rev | GGGGACCACTTTGTACAAGAAAGCTGGGTCTAGTCACCTTGCAGAAAAGGATGT |
| 2_known | J17 | Tb927.6.4470_for | GGGGACAAGTTTGTACAAAAAAGCAGGCTATGTCGGTCACGCGGATTAAC | Tb927.6.4470_rev | GGGGACCACTTTGTACAAGAAAGCTGGGTTCAGGCACTGCGTGCGTA |
| 2_known | J18 | Tb927.11.1310_for | GGGGACAAGTTTGTACAAAAAAGCAGGCTATGCGCTTTGTTCCACCTTTCC | Tb927.11.1310_rev | GGGGACCACTTTGTACAAGAAAGCTGGGTTTAGAAGCGATAAACTTCATCATTGGAGT |
| 2_known | J19 | Tb927.11.11150_for | GGGGACAAGTTTGTACAAAAAAGCAGGCTATGGAAAGGATGTCTCTCGCGG | Tb927.11.11150_rev | GGGGACCACTTTGTACAAGAAAGCTGGGTTTAAATTCGCTGTAATATTACCTCGGTCTC |
| 2_known | J20 | Tb927.9.15590_for | GGGGACAAGTTTGTACAAAAAAGCAGGCTATGAGCATCACTTTTCATAATTTATGGCTA | Tb927.9.15590_rev | GGGGACCACTTTGTACAAGAAAGCTGGGTTCATAGAAGTGTATAGAAAGTGAATATGGT |
| 2_known | J21 | Tb927.4.3560_for | GGGGACAAGTTTGTACAAAAAAGCAGGCTATGTCGACTATTGTGCGGAAACG | Tb927.4.3560_rev | GGGGACCACTTTGTACAAGAAAGCTGGGTTTAATCGTATTGTGGTCTAAATATCTCAAA |
| 2_known | J22 | Tb927.10.5070_for | GGGGACAAGTTTGTACAAAAAAGCAGGCTATGGCGGAACCTCGGTCTT | Tb927.10.5070_rev | GGGGACCACTTTGTACAAGAAAGCTGGGTCTAAGGGACTTCCCGCGACA |
| 2_known | J23 | Tb927.3.3020_for | GGGGACAAGTTTGTACAAAAAAGCAGGCTATGGACGCTACATGTGTAATTGATTATGG | Tb927.3.3020_rev | GGGGACCACTTTGTACAAGAAAGCTGGGTTCAAATAATCTTGGGCCGTACGC |
| 2_known | J24 | Tb927.8.7450_for | GGGGACAAGTTTGTACAAAAAAGCAGGCTATGCAGCCCGCGGGGGGG | Tb927.8.7450_rev | GGGGACCACTTTGTACAAGAAAGCTGGGTTCAGTAAGTGACGCGGGACA |
| 2_known | K1 | Tb927.10.12240_for | GGGGACAAGTTTGTACAAAAAAGCAGGCTATGTTCTACATTGACGATTTATTTTGGACA | Tb927.10.12240_rev | GGGGACCACTTTGTACAAGAAAGCTGGGTTCAGGACTGCTTCTTCCCGC |
| 2_known | K2 | Tb927.9.4310_for | GGGGACAAGTTTGTACAAAAAAGCAGGCTATGCTTCCATGTCCGTCGTTC | Tb927.9.4310_rev | GGGGACCACTTTGTACAAGAAAGCTGGGTCTACAAACCCTTGTAATACGTGAATATTTC |
| 2_known | K3 | Tb927.3.2960_for | GGGGACAAGTTTGTACAAAAAAGCAGGCTATGGCGAAAACAGTGATCCTCG | Tb927.3.2960_rev | GGGGACCACTTTGTACAAGAAAGCTGGGTTCAGCAAACGCGTGTGGAA |
| 2_known | K4 | Tb927.5.3820_for | GGGGACAAGTTTGTACAAAAAAGCAGGCTATGGCGGAGCTGCAACCT | Tb927.5.3820_rev | GGGGACCACTTTGTACAAGAAAGCTGGGTTTAAGCGAGAACACTGTAAAGTATGGC |
| 2_known | K5 | Tb927.11.8120_for | GGGGACAAGTTTGTACAAAAAAGCAGGCTATGTCTAATGTTGCTTTTGACTCGGG | Tb927.11.8120_rev | GGGGACCACTTTGTACAAGAAAGCTGGGTTTAAGGTAGCTTACACGAACGTTCC |
| 2_known | K6 | Tb927.9.14600_for | GGGGACAAGTTTGTACAAAAAAGCAGGCTATGTATTCTGCTACGCCGTTCATG | Tb927.9.14600_rev | GGGGACCACTTTGTACAAGAAAGCTGGGTTTACATTTTCGATTTGTAAGAATAGAGATC |
| 2_known | K7 | Tb927.4.4970_for | GGGGACAAGTTTGTACAAAAAAGCAGGCTATGCAGTCAGCAGGAGGAGATT | Tb927.4.4970_rev | GGGGACCACTTTGTACAAGAAAGCTGGGTTCAGTAAGTGACACGGGACAGC |
| 2_known | K8 | Tb927.9.8740_for | GGGGACAAGTTTGTACAAAAAAGCAGGCTATGTACAGCCAACCTTTTACTCAAAACA | Tb927.9.8740_rev | GGGGACCACTTTGTACAAGAAAGCTGGGTTTAACTCCACTGCTGTGTAATGCC |
| 2_known | K9 | Tb927.6.1690_for | GGGGACAAGTTTGTACAAAAAAGCAGGCTATGGAGCGTCTGAAAAATTTCGTACA | Tb927.6.1690_rev | GGGGACCACTTTGTACAAGAAAGCTGGGTTCATGGTTCAGTTGGCCACTCC |
| 2_known | K10 | Tb927.10.14200_for | GGGGACAAGTTTGTACAAAAAAGCAGGCTATGGTTGTAGAGCGCGACCG | Tb927.10.14200_rev | GGGGACCACTTTGTACAAGAAAGCTGGGTCTAGCGCACAACGAGAAACCC |
| 2_known | K11 | Tb927.7.2070_for | GGGGACAAGTTTGTACAAAAAAGCAGGCTATGTTTGCCTTTTCGGACCATATGG | Tb927.7.2070_rev | GGGGACCACTTTGTACAAGAAAGCTGGGTTCACTGCGCAGTACACCCA |
| 2_known | K12 | Tb927.11.11250_for | GGGGACAAGTTTGTACAAAAAAGCAGGCTATGTCTAACACATGTAAGCGTGTGG | Tb927.11.11250_rev | GGGGACCACTTTGTACAAGAAAGCTGGGTCTAGGCTGTGGAGACCTCTCTAC |
| 2_known | K13 | Tb927.8.900_for | GGGGACAAGTTTGTACAAAAAAGCAGGCTATGGATTCCAGAGACGGGAGTG | Tb927.8.900_rev | GGGGACCACTTTGTACAAGAAAGCTGGGTCTAATCTCGGCGTTTGCGGT |
| 2_known | K15 | Tb927.9.9390_for | GGGGACAAGTTTGTACAAAAAAGCAGGCTATGGCTGTCCCACCAGTGG | Tb927.9.9390_rev | GGGGACCACTTTGTACAAGAAAGCTGGGTTCAATGATTTCTCTCATCTACGACTTTCC |
| 2_known | K16 | Tb927.3.5150_for | GGGGACAAGTTTGTACAAAAAAGCAGGCTATGGTTTCTTGCAAGGCAATACCG | Tb927.3.5150_rev | GGGGACCACTTTGTACAAGAAAGCTGGGTTTACTCGCGAAGGTGTCGTTG |
| 2_known | K17 | Tb927.2.2160_for | GGGGACAAGTTTGTACAAAAAAGCAGGCTATGACATCGGAGGAAGCCGT | Tb927.2.2160_rev | GGGGACCACTTTGTACAAGAAAGCTGGGTTTAAATGTATTGCTCCAAATGCTCATACAA |
| 2_known | K18 | Tb927.10.7090_for | GGGGACAAGTTTGTACAAAAAAGCAGGCTATGTTTCGTAACCACGCATCGA | Tb927.10.7090_rev | GGGGACCACTTTGTACAAGAAAGCTGGGTTTACACGTGTTTGTTTACATTACTCGC |
| 2_known | K19 | Tb927.9.8260_for | GGGGACAAGTTTGTACAAAAAAGCAGGCTATGTTACGGTACTCGCCGATGC | Tb927.9.8260_rev | GGGGACCACTTTGTACAAGAAAGCTGGGTTTACAATGCTGCTGTAAGTTTGTCAAG |
| 2_known | K20 | Tb927.4.4070_for | GGGGACAAGTTTGTACAAAAAAGCAGGCTATGCACGTGGCTGTTAAGGAC | Tb927.4.4070_rev | GGGGACCACTTTGTACAAGAAAGCTGGGTCTATAGCTTACTTCCGCCGGGC |
| 2_known | K21 | Tb927.5.790_for | GGGGACAAGTTTGTACAAAAAAGCAGGCTATGGGTGGCGACGGGCGC | Tb927.5.790_rev | GGGGACCACTTTGTACAAGAAAGCTGGGTTCATTCGGTGTCATCACTCTCCT |
| 2_known | K22 | Tb927.9.12120_for | GGGGACAAGTTTGTACAAAAAAGCAGGCTATGGATGCAACACTTCACGATAAAAGC | Tb927.9.12120_rev | GGGGACCACTTTGTACAAGAAAGCTGGGTTCAGTCTGCCACCAAGGGAA |
| 2_known | K23 | Tb927.6.850_for | GGGGACAAGTTTGTACAAAAAAGCAGGCTATGAACAATAGCAACCAGGGAACAC | Tb927.6.850_rev | GGGGACCACTTTGTACAAGAAAGCTGGGTTCAGTCCCCTCCCACCACA |
| 2_known | K24 | Tb927.10.13400_for | GGGGACAAGTTTGTACAAAAAAGCAGGCTATGACCACCGAAAGCGGGA | Tb927.10.13400_rev | GGGGACCACTTTGTACAAGAAAGCTGGGTTCATTTAGTTATATCAGACAACACCTGCC |
| 2_known | L1 | Tb927.8.2020_for | GGGGACAAGTTTGTACAAAAAAGCAGGCTATGTTCTCCAAGTTCCTGATGAACG | Tb927.8.2020_rev | GGGGACCACTTTGTACAAGAAAGCTGGGTTCAATGATCGGGGTACGGATTTTC |
| 2_known | L2 | Tb927.10.6880_for | GGGGACAAGTTTGTACAAAAAAGCAGGCTATGGTAATCAGAGTCGGCATCAATG | Tb927.10.6880_rev | GGGGACCACTTTGTACAAGAAAGCTGGGTTCACTTTGTGATGTGCGCAATGA |
| 2_known | L3 | Tb927.8.570_for | GGGGACAAGTTTGTACAAAAAAGCAGGCTATGCCTGAGGCTTGCTTTCTC | Tb927.8.570_rev | GGGGACCACTTTGTACAAGAAAGCTGGGTTTAGTCCTCCTTCTCTTCGTTCTGATC |
| 2_known | L4 | Tb927.11.1560_for | GGGGACAAGTTTGTACAAAAAAGCAGGCTATGCCGAGCAGTTGGTTATTGAA | Tb927.11.1560_rev | GGGGACCACTTTGTACAAGAAAGCTGGGTCTACAATTTCTCCACCCACTTCTCATAAC |
| 2_known | L5 | Tb927.10.10140_for | GGGGACAAGTTTGTACAAAAAAGCAGGCTATGTACCGGTCCCGGCCC | Tb927.10.10140_rev | GGGGACCACTTTGTACAAGAAAGCTGGGTTTACGAAGCTTTCACTGAGGAAGC |
| 2_known | L6 | Tb927.11.1050_for | GGGGACAAGTTTGTACAAAAAAGCAGGCTATGGGAAAAGGTGCGCGC | Tb927.11.1050_rev | GGGGACCACTTTGTACAAGAAAGCTGGGTTTAGAGGTACCACTCCAGCTTGTC |
| 2_known | L7 | Tb927.7.3940_for | GGGGACAAGTTTGTACAAAAAAGCAGGCTATGGATCACGATCAACTATACGACTCT | Tb927.7.3940_rev | GGGGACCACTTTGTACAAGAAAGCTGGGTCTAAACGGCCCAGAAATGGCG |
| 2_known | L8 | Tb927.4.690_for | GGGGACAAGTTTGTACAAAAAAGCAGGCTATGTTTGGAGCGCCAAACACA | Tb927.4.690_rev | GGGGACCACTTTGTACAAGAAAGCTGGGTTCACGCACAGGTACGCTTCC |
| 2_known | L9 | Tb927.3.2340_for | GGGGACAAGTTTGTACAAAAAAGCAGGCTATGGCTTGGATAACTGATGCTGC | Tb927.3.2340_rev | GGGGACCACTTTGTACAAGAAAGCTGGGTTCACTGTGCCCATGGGGTG |
| 2_known | L10 | Tb927.11.16070_for | GGGGACAAGTTTGTACAAAAAAGCAGGCTATGCCATCCTATATTTTCATCCTGCATT | Tb927.11.16070_rev | GGGGACCACTTTGTACAAGAAAGCTGGGTTCACTTAGAAGCGGCGCAGT |
| 2_known | L11 | Tb927.8.480_for | GGGGACAAGTTTGTACAAAAAAGCAGGCTATGGCAGATGATGGTAAGACAGGG | Tb927.8.480_rev | GGGGACCACTTTGTACAAGAAAGCTGGGTTCACTCAGCGACATCAGCGA |
| 2_known | L12 | Tb927.10.13930_for | GGGGACAAGTTTGTACAAAAAAGCAGGCTATGGCCACTGGAGGTCGTT | Tb927.10.13930_rev | GGGGACCACTTTGTACAAGAAAGCTGGGTTCAAACAGTATCATTGTCGTTTGGCG |
| 2_known | L13 | Tb927.11.1150_for | GGGGACAAGTTTGTACAAAAAAGCAGGCTATGCAAGGGCAAAACCAAGCA | Tb927.11.1150_rev | GGGGACCACTTTGTACAAGAAAGCTGGGTTCAGCTATTCAACTGTATGGCAACTAC |
| 2_known | L14 | Tb927.6.2790_for | GGGGACAAGTTTGTACAAAAAAGCAGGCTATGTTTTGTTTTTCTCGTTTCGCTCT | Tb927.6.2790_rev | GGGGACCACTTTGTACAAGAAAGCTGGGTTCACAAGGATGGAAGTCCATGAAGG |
| 2_known | L15 | Tb927.11.1490_for | GGGGACAAGTTTGTACAAAAAAGCAGGCTATGCCGGCAGTTTTGTGCG | Tb927.11.1490_rev | GGGGACCACTTTGTACAAGAAAGCTGGGTCTACAGACAGTGTTTCAGAATAGTTGANNN |
| 2_known | L16 | Tb927.8.5600_for | GGGGACAAGTTTGTACAAAAAAGCAGGCTATGAATCAACTGGAGAGCCTCAAGA | Tb927.8.5600_rev | GGGGACCACTTTGTACAAGAAAGCTGGGTTTACAAAAGTGTAGCGATGAGTGCT |
| 2_known | L17 | Tb927.2.5800_for | GGGGACAAGTTTGTACAAAAAAGCAGGCTATGATGCGACAAAATCTGACAAGTGA | Tb927.2.5800_rev | GGGGACCACTTTGTACAAGAAAGCTGGGTCTATAGTTTGCTGCACCCGCG |
| 2_known | L18 | Tb927.5.800_for | GGGGACAAGTTTGTACAAAAAAGCAGGCTATGAGCGTAGAGCTTCGTGTGG | Tb927.5.800_rev | GGGGACCACTTTGTACAAGAAAGCTGGGTTTAGACGGGATGTTCATCTTCCTTTTC |
| 2_known | L19 | Tb927.11.15650_for | GGGGACAAGTTTGTACAAAAAAGCAGGCTATGCCACCATCACCAGCAGT | Tb927.11.15650_rev | GGGGACCACTTTGTACAAGAAAGCTGGGTCTAGACAATAGTGAGTAGCGCCGT |
| 2_known | L20 | Tb927.11.1020_for | GGGGACAAGTTTGTACAAAAAAGCAGGCTATGAGTGCTGCCGGCGTT | Tb927.11.1020_rev | GGGGACCACTTTGTACAAGAAAGCTGGGTCTATATTTTGCATTCCTTCACCTCAGCA |
| 2_known | L21 | Tb927.10.15600_for | GGGGACAAGTTTGTACAAAAAAGCAGGCTATGAACAATGGCGATAATTCCTCCG | Tb927.10.15600_rev | GGGGACCACTTTGTACAAGAAAGCTGGGTTTACAGTGTGCCTTCACCGTTAC |
| 2_known | L22 | Tb927.8.810_for | GGGGACAAGTTTGTACAAAAAAGCAGGCTATGTCATCTGTTTCCGATTGTATATTTGGC | Tb927.8.810_rev | GGGGACCACTTTGTACAAGAAAGCTGGGTTTACGGAAGGAGAAAGCAAACCTC |
| 2_known | L23 | Tb927.9.11100_for | GGGGACAAGTTTGTACAAAAAAGCAGGCTATGACGACAACTCCCACTGGT | Tb927.9.11100_rev | GGGGACCACTTTGTACAAGAAAGCTGGGTCTAAAAACCACGGAATGCAACCTG |
| 2_known | L24 | Tb927.8.1310_for | GGGGACAAGTTTGTACAAAAAAGCAGGCTATGGTTGGTGGCGATGGTGA | Tb927.8.1310_rev | GGGGACCACTTTGTACAAGAAAGCTGGGTTCAGGCAGCCGGTAAAAACC |
| 2_known | M1 | Tb927.11.6090_for | GGGGACAAGTTTGTACAAAAAAGCAGGCTATGTCGTCAGTTACAGCCAAATATTTACC | Tb927.11.6090_rev | GGGGACCACTTTGTACAAGAAAGCTGGGTTTACCCATCTGGGTGACACGG |
| 2_known | M2 | Tb927.10.690_for | GGGGACAAGTTTGTACAAAAAAGCAGGCTATGACACGCACATCACGGAG | Tb927.10.690_rev | GGGGACCACTTTGTACAAGAAAGCTGGGTTTAAACGGCATGCACCACGT |
| 2_known | M3 | Tb927.11.1780_for | GGGGACAAGTTTGTACAAAAAAGCAGGCTATGCATGTGGTAAAGTTGTTGTTGAG | Tb927.11.1780_rev | GGGGACCACTTTGTACAAGAAAGCTGGGTTCAAGGCCAATCCCTGTAAGGG |
| 2_known | M4 | Tb927.11.2130_for | GGGGACAAGTTTGTACAAAAAAGCAGGCTATGCAAACGACCGGTGAGC | Tb927.11.2130_rev | GGGGACCACTTTGTACAAGAAAGCTGGGTTTACGGTAGGCCGTCTTCGAA |
| 2_known | M5 | Tb927.11.12610_for | GGGGACAAGTTTGTACAAAAAAGCAGGCTATGATGACAAACTTGAATGTGCGGA | Tb927.11.12610_rev | GGGGACCACTTTGTACAAGAAAGCTGGGTTCACCTGGGTGCACATAGTAGC |

| | | | | | |
|---|---|---|---|---|---|
| 2_known | M6 | Tb927.9.11260_for | GGGGACAAGTTTGTACAAAAAAGCAGGCTATGACAGCTCTAGAGCGGTACG | Tb927.9.11260_rev | GGGGACCACTTTGTACAAGAAAGCTGGGTCTAAGCGTTCTGCAGAAGCGA |
| 2_known | M7 | Tb927.10.9560_for | GGGGACAAGTTTGTACAAAAAAGCAGGCTATGCAGAGGAAAATTGTAGCGAAGA | Tb927.10.9560_rev | GGGGACCACTTTGTACAAGAAAGCTGGGTTCAAATCTCAACGATGACCTTTCCC |
| 2_known | M8 | Tb927.3.1400_for | GGGGACAAGTTTGTACAAAAAAGCAGGCTATGACCCACAATGTGATGTGTGATG | Tb927.3.1400_rev | GGGGACCACTTTGTACAAGAAAGCTGGGTCTAAGATCCCGTAGACCCATTGATTATATT |
| 2_known | M9 | Tb927.4.2970_for | GGGGACAAGTTTGTACAAAAAAGCAGGCTATGACAACATATGGTCACCGTCCT | Tb927.4.2970_rev | GGGGACCACTTTGTACAAGAAAGCTGGGTCTATGCCACAATTCGCCGCT |
| 2_known | M10 | Tb927.8.2480_for | GGGGACAAGTTTGTACAAAAAAGCAGGCTATGATGATTTCCCAAAGTCATTCGCC | Tb927.8.2480_rev | GGGGACCACTTTGTACAAGAAAGCTGGGTTCAGTACACGAATGGTATCACCGC |
| 2_known | M11 | Tb927.10.5660_for | GGGGACAAGTTTGTACAAAAAAGCAGGCTATGCGAGGTCTAATAAAATCATTTAGCGG | Tb927.10.5660_rev | GGGGACCACTTTGTACAAGAAAGCTGGGTTCAGATCTCAAATGTACGGTCGACA |
| 2_known | M12 | Tb927.10.2370_for | GGGGACAAGTTTGTACAAAAAAGCAGGCTATGCCACTTTCCTCCGAGAACA | Tb927.10.2370_rev | GGGGACCACTTTGTACAAGAAAGCTGGGTTTATTCACGTGACCGCTTGTGTC |
| 2_known | M13 | Tb927.6.4430_for | GGGGACAAGTTTGTACAAAAAAGCAGGCTATGTGTGTCCTTCCAAGGAAGGT | Tb927.6.4430_rev | GGGGACCACTTTGTACAAGAAAGCTGGGTCTAAGTCGATACATAAACAAGATTATGCCG |
| 2_known | M14 | Tb927.7.7410_for | GGGGACAAGTTTGTACAAAAAAGCAGGCTATGTCGAGGCTGTCTTCGAAGG | Tb927.7.7410_rev | GGGGACCACTTTGTACAAGAAAGCTGGGTTTACTTCTTGACAACAACAATACCCTTTGT |
| 2_known | M15 | Tb927.5.3520_for | GGGGACAAGTTTGTACAAAAAAGCAGGCTATGCACGGAGTGTATCCTATTTTAGTTG | Tb927.5.3520_rev | GGGGACCACTTTGTACAAGAAAGCTGGGTTCACAACTGTGCCAGCACC |
| 2_known | M16 | Tb927.4.5300_for | GGGGACAAGTTTGTACAAAAAAGCAGGCTATGTTCGTTCCGTCTAATGTTGCT | Tb927.4.5300_rev | GGGGACCACTTTGTACAAGAAAGCTGGGTTCAGGCCCTCTTACGCCAA |
| 2_known | M17 | Tb927.10.8540_for | GGGGACAAGTTTGTACAAAAAAGCAGGCTATGGGTATTGACTACTACAAGGTGCT | Tb927.10.8540_rev | GGGGACCACTTTGTACAAGAAAGCTGGGTCTAAAATTCACATTTTTCAACCAATCGCTT |
| 2_known | M18 | Tb927.8.6340_for | GGGGACAAGTTTGTACAAAAAAGCAGGCTATGAAGAAGGCTTTGATCTCGGATTT | Tb927.8.6340_rev | GGGGACCACTTTGTACAAGAAAGCTGGGTTCAAACTGCTGCGCCTTCC |
| 2_known | M19 | Tb927.8.4410_for | GGGGACAAGTTTGTACAAAAAAGCAGGCTATGCTGTTGCTGTTAGAGAGGGA | Tb927.8.4410_rev | GGGGACCACTTTGTACAAGAAAGCTGGGTTCAGGTTATTGTAATCTTCGTTTCTTCAGC |
| 2_known | M20 | Tb927.10.12830_for | GGGGACAAGTTTGTACAAAAAAGCAGGCTATGAGCGCAGTGGAGCTTG | Tb927.10.12830_rev | GGGGACCACTTTGTACAAGAAAGCTGGGTTTAACTTCTGAATAACTCAATATCAGCAAC |
| 2_known | M21 | Tb927.9.11030_for | GGGGACAAGTTTGTACAAAAAAGCAGGCTATGCCTGTAGCCACGTCGA | Tb927.9.11030_rev | GGGGACCACTTTGTACAAGAAAGCTGGGTCTAAAACCCACGGAACTCCAATTGT |
| 2_known | M22 | Tb927.2.4830_for | GGGGACAAGTTTGTACAAAAAAGCAGGCTATGTATGGCCGCTGTCTGCA | Tb927.2.4830_rev | GGGGACCACTTTGTACAAGAAAGCTGGGTTCAAGAAGCGCTACTGTGCTTG |
| 2_known | M23 | Tb927.11.12810_for | GGGGACAAGTTTGTACAAAAAAGCAGGCTATGCCACCCAAATCTCACAAGC | Tb927.11.12810_rev | GGGGACCACTTTGTACAAGAAAGCTGGGTCTAGAAATCTGCGTCCAGAGAGAAAAC |
| 2_known | M24 | Tb927.10.13010_for | GGGGACAAGTTTGTACAAAAAAGCAGGCTATGAAGTCGGATGGGTGCCT | Tb927.10.13010_rev | GGGGACCACTTTGTACAAGAAAGCTGGGTTCAGATCCTCGTGTATTCACCATCG |
| 2_known | N2 | Tb927.10.15370_for | GGGGACAAGTTTGTACAAAAAAGCAGGCTATGGTCAAAGTGTACAATTTGCAACG | Tb927.10.15370_rev | GGGGACCACTTTGTACAAGAAAGCTGGGTCTATTCCGTAAACGCAGTGGCT |
| 2_known | N3 | Tb927.3.5500_for | GGGGACAAGTTTGTACAAAAAAGCAGGCTATGAAGGTGACGATTGTAGACTGC | Tb927.3.5500_rev | GGGGACCACTTTGTACAAGAAAGCTGGGTTCAATTCATCACGAACTTGAGATCGT |
| 2_known | N4 | Tb927.9.9410_for | GGGGACAAGTTTGTACAAAAAAGCAGGCTATGGCTGTCCCACCAGTGG | Tb927.9.9410_rev | GGGGACCACTTTGTACAAGAAAGCTGGGTTCATACCTCGTTAGTTGATACCTCACA |
| 2_known | N5 | Tb927.10.11160_for | GGGGACAAGTTTGTACAAAAAAGCAGGCTATGCGGGTCGGTTCCTGG | Tb927.10.11160_rev | GGGGACCACTTTGTACAAGAAAGCTGGGTTTAGTCCTTGCCTTGCCGTTG |
| 2_known | N9 | Tb927.6.3760_for | GGGGACAAGTTTGTACAAAAAAGCAGGCTATGAAGCTAGAGAGATCGTCCTGC | Tb927.6.3760_rev | GGGGACCACTTTGTACAAGAAAGCTGGGTTCAAATAGGAATGGCTAGTATGTGTGAAAC |
| 2_known | N10 | Tb927.8.2740_for | GGGGACAAGTTTGTACAAAAAAGCAGGCTATGTTCCGAAGGCTCACACAGT | Tb927.8.2740_rev | GGGGACCACTTTGTACAAGAAAGCTGGGTTCACTGGACAAAGAGTCCAGCC |
| 2_known | N11 | Tb927.7.4860_for | GGGGACAAGTTTGTACAAAAAAGCAGGCTATGACTGAGCATATCTGTGCTTCG | Tb927.7.4860_rev | GGGGACCACTTTGTACAAGAAAGCTGGGTCTATTGTACGGTTTCCCTGGCTAC |
| 2_known | N12 | Tb927.10.8920_for | GGGGACAAGTTTGTACAAAAAAGCAGGCTATGACATCCAGTTTTCTCGCACAT | Tb927.10.8920_rev | GGGGACCACTTTGTACAAGAAAGCTGGGTCTAGTTGCGTCGGAAAATACTATAAGCA |
| 2_known | N13 | Tb927.11.14360_for | GGGGACAAGTTTGTACAAAAAAGCAGGCTATGGCACAAAGTACTGCTCGTC | Tb927.11.14360_rev | GGGGACCACTTTGTACAAGAAAGCTGGGTCTAAAAAAAATTCTGCTGCATTAACTTCTC |
| 2_known | N14 | Tb927.9.15930_for | GGGGACAAGTTTGTACAAAAAAGCAGGCTATGCCAGCACCCGCCTCT | Tb927.9.15930_rev | GGGGACCACTTTGTACAAGAAAGCTGGGTTTATTGGCAGCAACAATTATCATTTCTCTG |
| 2_known | N15 | Tb927.3.4990_for | GGGGACAAGTTTGTACAAAAAAGCAGGCTATGAAAACTATGCGTGCGGTTACG | Tb927.3.4990_rev | GGGGACCACTTTGTACAAGAAAGCTGGGTTCACTCACCTGTAGAGTCGCAC |
| 2_known | N16 | Tb927.4.1170_for | GGGGACAAGTTTGTACAAAAAAGCAGGCTATGGAAGCCGCCTTTGATGC | Tb927.4.1170_rev | GGGGACCACTTTGTACAAGAAAGCTGGGTTTAATCCCTACGTAGAGAAAATTCCATGG |
| 2_known | N17 | Tb927.2.5180_for | GGGGACAAGTTTGTACAAAAAAGCAGGCTATGCCTGGGGTTAACGTGATGA | Tb927.2.5180_rev | GGGGACCACTTTGTACAAGAAAGCTGGGTTTACGAAAAGAAACTCTCGCCGG |
| 2_known | N18 | Tb927.4.1640_for | GGGGACAAGTTTGTACAAAAAAGCAGGCTATGACGGCGCGTGACGGG | Tb927.4.1640_rev | GGGGACCACTTTGTACAAGAAAGCTGGGTTTAAATAGTTTCTTCCGACCTTGACCTC |
| 2_known | N19 | Tb927.10.8750_for | GGGGACAAGTTTGTACAAAAAAGCAGGCTATGGTGGGGTCGAGGACCA | Tb927.10.8750_rev | GGGGACCACTTTGTACAAGAAAGCTGGGTTTAGAGCTGTTGACAAAACTTCCTCTC |
| 2_known | N20 | Tb927.8.2230_for | GGGGACAAGTTTGTACAAAAAAGCAGGCTATGAAATTCAAAGCAACAGTACGAGATCG | Tb927.8.2230_rev | GGGGACCACTTTGTACAAGAAAGCTGGGTTCAATTTGTTACAGCCGGTATGCA |
| 2_known | N21 | Tb927.7.3730_for | GGGGACAAGTTTGTACAAAAAAGCAGGCTATGGAAACGCACCAGGATCCG | Tb927.7.3730_rev | GGGGACCACTTTGTACAAGAAAGCTGGGTTTAGTTTGGGCTCATGAAGTTGGTG |
| 2_known | N22 | Tb927.6.560_for | GGGGACAAGTTTGTACAAAAAAGCAGGCTATGCATCTCATGCGTGCCTG | Tb927.6.560_rev | GGGGACCACTTTGTACAAGAAAGCTGGGTCTACGCCGTGTTGGGTGC |
| 2_known | N23 | Tb927.11.9550_for | GGGGACAAGTTTGTACAAAAAAGCAGGCTATGACACAACAGACAGTACCTTGGG | Tb927.11.9550_rev | GGGGACCACTTTGTACAAGAAAGCTGGGTTTACGACTTCGTTGCGACAATGA |
| 2_known | N24 | Tb927.10.14080_for | GGGGACAAGTTTGTACAAAAAAGCAGGCTATGGCCGATCGAGGTGACG | Tb927.10.14080_rev | GGGGACCACTTTGTACAAGAAAGCTGGGTCTAGGCCACACTCCGGCA |
| 2_known | O1 | Tb927.8.4430_for | GGGGACAAGTTTGTACAAAAAAGCAGGCTATGGCTGCATCCGCTAATGG | Tb927.8.4430_rev | GGGGACCACTTTGTACAAGAAAGCTGGGTCTAAGCGCTGACAAGCGCT |
| 2_known | O3 | Tb927.11.5860_for | GGGGACAAGTTTGTACAAAAAAGCAGGCTATGTGTGGGAAGGATTCATCACATG | Tb927.11.5860_rev | GGGGACCACTTTGTACAAGAAAGCTGGGTCTAAAAACCGGCCATAAGCTCTTCA |
| 2_known | O4 | Tb927.1.870_for | GGGGACAAGTTTGTACAAAAAAGCAGGCTATGTCAGGTGTACCTTTTCCTTCACG | Tb927.1.870_rev | GGGGACCACTTTGTACAAGAAAGCTGGGTCTAGTTCCCATCCTCCCTCACAC |
| 2_known | O5 | Tb927.8.1150_for | GGGGACAAGTTTGTACAAAAAAGCAGGCTATGTCCTCGGTTCTTACGCGG | Tb927.8.1150_rev | GGGGACCACTTTGTACAAGAAAGCTGGGTTCAGATGTTCGTGGAGTCTCGA |
| 2_known | O6 | Tb927.10.6140_for | GGGGACAAGTTTGTACAAAAAAGCAGGCTATGCGCAATTGTCAGCCGG | Tb927.10.6140_rev | GGGGACCACTTTGTACAAGAAAGCTGGGTTCACTGAAGCATGCCTGCG |
| 2_known | O7 | Tb927.10.380_for | GGGGACAAGTTTGTACAAAAAAGCAGGCTATGCTCCGCCGTAGTGATTTACT | Tb927.10.380_rev | GGGGACCACTTTGTACAAGAAAGCTGGGTCTAATATGCAGTTCCACAAACCTCTGC |
| 2_known | O8 | Tb927.11.9610_for | GGGGACAAGTTTGTACAAAAAAGCAGGCTATGAACATTCAAGGCATGGCTCTG | Tb927.11.9610_rev | GGGGACCACTTTGTACAAGAAAGCTGGGTTTATATGCCTTCGGGTGTCCAGA |
| 2_known | O9 | Tb927.9.6090_for | GGGGACAAGTTTGTACAAAAAAGCAGGCTATGGTGAGGACGACACGCT | Tb927.9.6090_rev | GGGGACCACTTTGTACAAGAAAGCTGGGTCTAAAGTCTTGAAGGAGTGTGTCGC |
| 2_known | O11 | Tb927.6.1610_for | GGGGACAAGTTTGTACAAAAAAGCAGGCTATGCCGAAGGAACAACGCAG | Tb927.6.1610_rev | GGGGACCACTTTGTACAAGAAAGCTGGGTCTACACAAAATGGATATCATTCTTTGTGAA |
| 2_known | O12 | Tb927.8.3910_for | GGGGACAAGTTTGTACAAAAAAGCAGGCTATGGTGGCCGTGAACCGA | Tb927.8.3910_rev | GGGGACCACTTTGTACAAGAAAGCTGGGTTTAAATATGTTTTGCGACGGCATTGTATAT |
| 2_known | O13 | Tb927.9.9520_for | GGGGACAAGTTTGTACAAAAAAGCAGGCTATGACACAGGTCATCACGGAGA | Tb927.9.9520_rev | GGGGACCACTTTGTACAAGAAAGCTGGGTTCACCACAACGGCACGGA |
| 2_known | O14 | Tb927.10.1940_for | GGGGACAAGTTTGTACAAAAAAGCAGGCTATGGTAGACTGCACGGTCGA | Tb927.10.1940_rev | GGGGACCACTTTGTACAAGAAAGCTGGGTTTACGGCATCCTCAGAAAGGGG |
| 2_known | O15 | Tb927.11.4060_for | GGGGACAAGTTTGTACAAAAAAGCAGGCTATGACTTCCGCAGGTCGTCC | Tb927.11.4060_rev | GGGGACCACTTTGTACAAGAAAGCTGGGTCTACCGGAGTGGGTTCGTCT |
| 2_known | O16 | Tb927.6.2480_for | GGGGACAAGTTTGTACAAAAAAGCAGGCTATGCGACGCGTGACTTCC | Tb927.6.2480_rev | GGGGACCACTTTGTACAAGAAAGCTGGGTTTAGTATGGTCTGTGATATGAGTACAACTT |
| 2_known | O17 | Tb927.9.5710_for | GGGGACAAGTTTGTACAAAAAAGCAGGCTATGTCGACTACCACAGATTTTGTTGG | Tb927.9.5710_rev | GGGGACCACTTTGTACAAGAAAGCTGGGTTCATGTTTCCTCCCTCTTCACGC |
| 2_known | O18 | Tb927.6.2360_for | GGGGACAAGTTTGTACAAAAAAGCAGGCTATGTCATCCGCTCCTCTGAGG | Tb927.6.2360_rev | GGGGACCACTTTGTACAAGAAAGCTGGGTTCAAGGAGAGAAACTGGGCTTCT |
| 2_known | O20 | Tb927.7.7360_for | GGGGACAAGTTTGTACAAAAAAGCAGGCTATGCAGGTGCAGGTGCAGG | Tb927.7.7360_rev | GGGGACCACTTTGTACAAGAAAGCTGGGTTTAAGCTCCGTTGAACCGCATC |
| 2_known | O22 | Tb927.1.4690_for | GGGGACAAGTTTGTACAAAAAAGCAGGCTATGACGGTGGACGCAAATGC | Tb927.1.4690_rev | GGGGACCACTTTGTACAAGAAAGCTGGGTCTACCGCAGCCGAAAATCCTG |
| 2_known | O23 | Tb927.9.800_for | GGGGACAAGTTTGTACAAAAAAGCAGGCTATGTACTTTTGTCTCTATACAGCTTGTTTG | Tb927.9.800_rev | GGGGACCACTTTGTACAAGAAAGCTGGGTTTAACAGCTAGCTTCATGCCAGAATG |
| 2_known | O24 | Tb927.11.14350_for | GGGGACAAGTTTGTACAAAAAAGCAGGCTATGACAACCGCCCTACGAGA | Tb927.11.14350_rev | GGGGACCACTTTGTACAAGAAAGCTGGGTTCAGACGCGGCGGGAGAA |
| 2_known | P1 | Tb927.10.15380_for | GGGGACAAGTTTGTACAAAAAAGCAGGCTATGACAAAGGGCAAGGGCCG | Tb927.10.15380_rev | GGGGACCACTTTGTACAAGAAAGCTGGGTTTAACGAGCGCGTCGCTTAG |
| 2_known | P2 | Tb927.4.5030_for | GGGGACAAGTTTGTACAAAAAAGCAGGCTATGAACTGTGACGAAATCATAAAGAAACTC | Tb927.4.5030_rev | GGGGACCACTTTGTACAAGAAAGCTGGGTCTAAAAGAAGTACCTCAACAGCGTGC |
| 2_known | P3 | Tb927.8.7390_for | GGGGACAAGTTTGTACAAAAAAGCAGGCTATGAATTGTCGTGAGATCATAAGGAAGC | Tb927.8.7390_rev | GGGGACCACTTTGTACAAGAAAGCTGGGTCTAAAAGAAGTACCTCAGCAGCGTAC |
| 2_known | P4 | Tb927.6.940_for | GGGGACAAGTTTGTACAAAAAAGCAGGCTATGTGCTCCTTAATTACACAACTCTGTG | Tb927.6.940_rev | GGGGACCACTTTGTACAAGAAAGCTGGGTCTATTGGATAGATCTGTCAACAGAAAACAT |
| 2_known | P5 | Tb927.6.3890_for | GGGGACAAGTTTGTACAAAAAAGCAGGCTATGTTGCCAGACCCACCACT | Tb927.6.3890_rev | GGGGACCACTTTGTACAAGAAAGCTGGGTTTACACTGAGCATATGAAACTTCCTAATTC |
| 2_known | P6 | Tb927.6.3480_for | GGGGACAAGTTTGTACAAAAAAGCAGGCTATGTGGTACAGTAATTTGTTCGTAGCA | Tb927.6.3480_rev | GGGGACCACTTTGTACAAGAAAGCTGGGTTTACCAGGGTCCATATACAAGTGTTCG |
| 2_known | P7 | Tb927.9.9380_for | GGGGACAAGTTTGTACAAAAAAGCAGGCTATGGCTGTCCCACCAGTGG | Tb927.9.9380_rev | GGGGACCACTTTGTACAAGAAAGCTGGGTTCACACATACGCCCCACATTTAAA |
| 2_known | P8 | Tb927.10.4040_for | GGGGACAAGTTTGTACAAAAAAGCAGGCTATGGGAGTTATATTTCTTGTGTGTGCG | Tb927.10.4040_rev | GGGGACCACTTTGTACAAGAAAGCTGGGTTTACATGACTTTTTTACCCCTCTCTGATTC |
| 2_known | P9 | Tb927.6.3000_for | GGGGACAAGTTTGTACAAAAAAGCAGGCTATGCTGGCTTCTCAACTGAAAACC | Tb927.6.3000_rev | GGGGACCACTTTGTACAAGAAAGCTGGGTTTAATTTTCTTCCTTTCCGATTACCTCAGC |
| 2_known | P10 | Tb927.11.16290_for | GGGGACAAGTTTGTACAAAAAAGCAGGCTATGATTGGTTTCGTGTACAATATAGTGTGT | Tb927.11.16290_rev | GGGGACCACTTTGTACAAGAAAGCTGGGTCTAGGTAACCGGTTGGGATGCT |
| 2_known | P11 | Tb927.3.1790_for | GGGGACAAGTTTGTACAAAAAAGCAGGCTATGCGCCGTTTTGCTACTACTT | Tb927.3.1790_rev | GGGGACCACTTTGTACAAGAAAGCTGGGTTTAGCTCAGAATGCGACGTGC |
| 2_known | P12 | Tb927.11.20730_for | GGGGACAAGTTTGTACAAAAAAGCAGGCTATGGCCTTTTCTGCACTAGCC | Tb927.11.20730_rev | GGGGACCACTTTGTACAAGAAAGCTGGGTTCAGTTCAGCAAATTCATCGTATTAGAAAC |
| 2_known | P13 | Tb927.4.1720_for | GGGGACAAGTTTGTACAAAAAAGCAGGCTATGGCGCGTTCACTGCAG | Tb927.4.1720_rev | GGGGACCACTTTGTACAAGAAAGCTGGGTCTATTCATCTACATCCTCCGGTCTAAAGA |
| 2_known | P14 | Tb927.1.3320_for | GGGGACAAGTTTGTACAAAAAAGCAGGCTATGTCGACATCAAAATCTGAAAAAGGTG | Tb927.1.3320_rev | GGGGACCACTTTGTACAAGAAAGCTGGGTTTAACGAGGGTACGAGAGAATTTCCT |
| 2_known | P15 | Tb927.9.3760_for | GGGGACAAGTTTGTACAAAAAAGCAGGCTATGTTTCCGTTTTACTCATGCGACC | Tb927.9.3760_rev | GGGGACCACTTTGTACAAGAAAGCTGGGTCTATCTTGCTTTCAGCTGTGTAGGATT |
| 2_known | P16 | Tb927.9.14430_for | GGGGACAAGTTTGTACAAAAAAGCAGGCTATGTCCGTCTCTTCGCAACCT | Tb927.9.14430_rev | GGGGACCACTTTGTACAAGAAAGCTGGGTCTACTCCGTGACAACATTCCCTCT |
| 2_known | P17 | Tb927.9.9620_for | GGGGACAAGTTTGTACAAAAAAGCAGGCTATGCAGCACGTGGGAACG | Tb927.9.9620_rev | GGGGACCACTTTGTACAAGAAAGCTGGGTCTACTCACGTGCGTCAACAATCC |
| 2_known | P18 | Tb927.10.7990_for | GGGGACAAGTTTGTACAAAAAAGCAGGCTATGCTGTGGGTCGACAAGTACC | Tb927.10.7990_rev | GGGGACCACTTTGTACAAGAAAGCTGGGTTCAGTTTCGCTTGAGAAGTTGCA |
| 2_known | P19 | Tb927.10.7780_for | GGGGACAAGTTTGTACAAAAAAGCAGGCTATGGTGTCGTTCAGCATCGATG | Tb927.10.7780_rev | GGGGACCACTTTGTACAAGAAAGCTGGGTTTACAAGGCTAACTTTTGCTTTCGACG |
| 2_known | P20 | Tb927.9.8720_for | GGGGACAAGTTTGTACAAAAAAGCAGGCTATGCCTGACTACAGCCGCG | Tb927.9.8720_rev | GGGGACCACTTTGTACAAGAAAGCTGGGTTTAAAGCTTGCTCCTCTCCCTTTG |
| 2_known | P21 | Tb927.10.9760_for | GGGGACAAGTTTGTACAAAAAAGCAGGCTATGCTGCTCACAAAACTTGCGC | Tb927.10.9760_rev | GGGGACCACTTTGTACAAGAAAGCTGGGTTTATTGCACCTCGTGCATTTCGT |
| 2_known | P22 | Tb927.7.1110_for | GGGGACAAGTTTGTACAAAAAAGCAGGCTATGGGCGACGACGGTTATTCG | Tb927.7.1110_rev | GGGGACCACTTTGTACAAGAAAGCTGGGTTTACAACAAATTGTGCTTTTTCCGAATCT |
| 2_known | P23 | Tb927.6.600_for | GGGGACAAGTTTGTACAAAAAAGCAGGCTATGATGCAGTATGGTGGCGGG | Tb927.6.600_rev | GGGGACCACTTTGTACAAGAAAGCTGGGTTCAGCTATGACCCTTTACCGCT |
| 2_known | P24 | Tb927.8.6540_for | GGGGACAAGTTTGTACAAAAAAGCAGGCTATGCTTAAAACCGTGCTACTACTTGA | Tb927.8.6540_rev | GGGGACCACTTTGTACAAGAAAGCTGGGTTCACGAGGTGGCAACGCA |
| 3_hypothetical | A1 | Tb927.7.4270_for | GGGGACAAGTTTGTACAAAAAAGCAGGCTATGGTGATGAAGGCTTTCCACCT | Tb927.7.4270_rev | GGGGACCACTTTGTACAAGAAAGCTGGGTTTATTTGTTTGTTTCCTCAGTCCTATAGTG |
| 3_hypothetical | A2 | Tb927.10.11750_for | GGGGACAAGTTTGTACAAAAAAGCAGGCTATGCCAGTTGAGCAGTTGGAAG | Tb927.10.11750_rev | GGGGACCACTTTGTACAAGAAAGCTGGGTCTAGTTAGGTTTCCACAGCTTTAACGC |
| 3_hypothetical | A3 | Tb927.11.2410_for | GGGGACAAGTTTGTACAAAAAAGCAGGCTATGCCCTGCTGCAGCAAC | Tb927.11.2410_rev | GGGGACCACTTTGTACAAGAAAGCTGGGTTTACGTCGAACTCGGAACAACTTT |
| 3_hypothetical | A4 | Tb927.3.4250_for | GGGGACAAGTTTGTACAAAAAAGCAGGCTATGTTGCAACTAACGGTTGAGGAC | Tb927.3.4250_rev | GGGGACCACTTTGTACAAGAAAGCTGGGTTCACGTTGATGTATCATACGGAGGA |
| 3_hypothetical | A5 | Tb927.5.140_for | GGGGACAAGTTTGTACAAAAAAGCAGGCTATGCCCATTATGTGCCATCGG | Tb927.5.140_rev | GGGGACCACTTTGTACAAGAAAGCTGGGTTTACACGTTATTCGTTGGCAGCT |
| 3_hypothetical | A6 | Tb927.11.16180_for | GGGGACAAGTTTGTACAAAAAAGCAGGCTATGACTCAGCAGCAACCCCC | Tb927.11.16180_rev | GGGGACCACTTTGTACAAGAAAGCTGGGTTTAAGCTATTTTTTCACCAAGTTGCTTGG |
| 3_hypothetical | A7 | Tb927.4.970_for | GGGGACAAGTTTGTACAAAAAAGCAGGCTATGACAAGCCGCCGTGTC | Tb927.4.970_rev | GGGGACCACTTTGTACAAGAAAGCTGGGTTTAGAGGGGATTTAGTCCTCGAATCC |
| 3_hypothetical | A8 | Tb927.9.4920_for | GGGGACAAGTTTGTACAAAAAAGCAGGCTATGGCCGCGGATGATGAGT | Tb927.9.4920_rev | GGGGACCACTTTGTACAAGAAAGCTGGGTCTAAATATTTCTATTCACCAGACGAATGCA |
| 3_hypothetical | A9 | Tb927.11.1950_for | GGGGACAAGTTTGTACAAAAAAGCAGGCTATGAAGCGTACACGCAATGCT | Tb927.11.1950_rev | GGGGACCACTTTGTACAAGAAAGCTGGGTCTATTCTAGTGCCTCAAGAACGTGTT |
| 3_hypothetical | A10 | Tb927.3.5670_for | GGGGACAAGTTTGTACAAAAAAGCAGGCTATGGCTAAGGTTGATACGCTGC | Tb927.3.5670_rev | GGGGACCACTTTGTACAAGAAAGCTGGGTCTAGACAACAACAAAAAAAAGTATTGCATC |
| 3_hypothetical | A11 | Tb927.6.1470_for | GGGGACAAGTTTGTACAAAAAAGCAGGCTATGCGCCGAGACGACTTCG | Tb927.6.1470_rev | GGGGACCACTTTGTACAAGAAAGCTGGGTTCAAGCCGCCCCTCGGAA |
| 3_hypothetical | A12 | Tb927.11.14830_for | GGGGACAAGTTTGTACAAAAAAGCAGGCTATGAGTTTGCTTTTGGATGACGACG | Tb927.11.14830_rev | GGGGACCACTTTGTACAAGAAAGCTGGGTTTACATCTTCCTTACACATGCCATTTTGG |
| 3_hypothetical | A13 | Tb927.5.289b_for | GGGGACAAGTTTGTACAAAAAAGCAGGCTATGCCTCTCGTTCCAACTGCT | Tb927.5.289b_rev | GGGGACCACTTTGTACAAGAAAGCTGGGTTTATCTCGTGGCACCAGAGCA |
| 3_hypothetical | A14 | Tb927.11.4240_for | GGGGACAAGTTTGTACAAAAAAGCAGGCTATGCATCACAATGAGTTCAATAGTGTAGA | Tb927.11.4240_rev | GGGGACCACTTTGTACAAGAAAGCTGGGTTCAAGGGGCAATGCCCATACT |
| 3_hypothetical | A15 | Tb927.4.1890_for | GGGGACAAGTTTGTACAAAAAAGCAGGCTATGTCTGACCTTGACCGTGATAGT | Tb927.4.1890_rev | GGGGACCACTTTGTACAAGAAAGCTGGGTCTAATAATCCTCCTCTTCCTCTTCATTCTC |
| 3_hypothetical | A16 | Tb927.8.6570_for | GGGGACAAGTTTGTACAAAAAAGCAGGCTATGGGAAGTTTTGTGGTGCGT | Tb927.8.6570_rev | GGGGACCACTTTGTACAAGAAAGCTGGGTTTAGCTGGAAATGATGGCAGTAATGG |
| 3_hypothetical | A17 | Tb927.3.2580_for | GGGGACAAGTTTGTACAAAAAAGCAGGCTATGACGAGACCTGTGTGTGTCT | Tb927.3.2580_rev | GGGGACCACTTTGTACAAGAAAGCTGGGTTCAGAGAAATAGGAGCAACACTGGA |
| 3_hypothetical | A18 | Tb927.2.4410_for | GGGGACAAGTTTGTACAAAAAAGCAGGCTATGCTTGAGCCTTCGCAGC | Tb927.2.4410_rev | GGGGACCACTTTGTACAAGAAAGCTGGGTCTATACACGGCCAACGGCTTG |
| 3_hypothetical | A19 | Tb927.10.4880_for | GGGGACAAGTTTGTACAAAAAAGCAGGCTATGTTACGCAAATCTTCGGTTTCGT | Tb927.10.4880_rev | GGGGACCACTTTGTACAAGAAAGCTGGGTCTACTCCGCGTTCAGGTACACC |
| 3_hypothetical | A20 | Tb927.11.4090_for | GGGGACAAGTTTGTACAAAAAAGCAGGCTATGAGGAGAGTGCTATTATTCCTGTGT | Tb927.11.4090_rev | GGGGACCACTTTGTACAAGAAAGCTGGGTTCAAAGTAACATGCTGTATAAAAGAGGAAT |
| 3_hypothetical | A21 | Tb927.4.4200_for | GGGGACAAGTTTGTACAAAAAAGCAGGCTATGCGATCCAACTGCGCAC | Tb927.4.4200_rev | GGGGACCACTTTGTACAAGAAAGCTGGGTTCAGTGTCTCCATAGAACAGGGAG |
| 3_hypothetical | A22 | Tb927.8.5890_for | GGGGACAAGTTTGTACAAAAAAGCAGGCTATGACCACAGCCGGTCAGT | Tb927.8.5890_rev | GGGGACCACTTTGTACAAGAAAGCTGGGTTTAACGGTGGTGCTTTTCTTTCTCT |
| 3_hypothetical | A23 | Tb927.3.4830_for | GGGGACAAGTTTGTACAAAAAAGCAGGCTATGGAGATTCCCGGTGTATTTTCAGA | Tb927.3.4830_rev | GGGGACCACTTTGTACAAGAAAGCTGGGTTCAGCACGCCGAAGCCAT |
| 3_hypothetical | A24 | Tb927.8.860_for | GGGGACAAGTTTGTACAAAAAAGCAGGCTATGTATTTTTTTTTCTTTCAGGTAATCGAT | Tb927.8.860_rev | GGGGACCACTTTGTACAAGAAAGCTGGGTTCAAAACACCTCTTGAAGGGTTCC |
| 3_hypothetical | B1 | Tb927.11.4070_for | GGGGACAAGTTTGTACAAAAAAGCAGGCTATGCTGTCGGAAATGTTGTTGCT | Tb927.11.4070_rev | GGGGACCACTTTGTACAAGAAAGCTGGGTTCAAAGTAACATGCTGTATAAAAGAGGAAT |
| 3_hypothetical | B2 | Tb927.8.7720_for | GGGGACAAGTTTGTACAAAAAAGCAGGCTATGGCGACAAATAAGGGAATAACGT | Tb927.8.7720_rev | GGGGACCACTTTGTACAAGAAAGCTGGGTTTATCGGGAAGCGACCAGACT |
| 3_hypothetical | B3 | Tb927.9.7020_for | GGGGACAAGTTTGTACAAAAAAGCAGGCTATGGCACTCACGCTGATACCG | Tb927.9.7020_rev | GGGGACCACTTTGTACAAGAAAGCTGGGTTCAAAATATTATTTCCCCGTCATCATCGT |
| 3_hypothetical | B4 | Tb927.7.3600_for | GGGGACAAGTTTGTACAAAAAAGCAGGCTATGTACCGCAGTGGTACGACG | Tb927.7.3600_rev | GGGGACCACTTTGTACAAGAAAGCTGGGTCTAGGCCACTGCTCGGGT |
| 3_hypothetical | B5 | Tb927.10.12200_for | GGGGACAAGTTTGTACAAAAAAGCAGGCTATGACAGCACTGGTGGTCCA | Tb927.10.12200_rev | GGGGACCACTTTGTACAAGAAAGCTGGGTTCAACTCAGTAGCATCACCCATGC |
| 3_hypothetical | B6 | Tb927.8.4220_for | GGGGACAAGTTTGTACAAAAAAGCAGGCTATGGGCAAGAAGGCTGAGCA | Tb927.8.4220_rev | GGGGACCACTTTGTACAAGAAAGCTGGGTTCAAGAAAGATCATACGCGGCTTC |
| 3_hypothetical | B7 | Tb927.11.5540_for | GGGGACAAGTTTGTACAAAAAAGCAGGCTATGTTGCAGTGTCTCACATATGCG | Tb927.11.5540_rev | GGGGACCACTTTGTACAAGAAAGCTGGGTTCAAATGTACTTTTTGGTCGTGAAAAAGT |
| 3_hypothetical | B8 | Tb927.5.3690_for | GGGGACAAGTTTGTACAAAAAAGCAGGCTATGTTCGTAAGGTTGCCGACC | Tb927.5.3690_rev | GGGGACCACTTTGTACAAGAAAGCTGGGTTTATGGCAGCCTGTTCACCTTATTTT |
| 3_hypothetical | B9 | Tb927.9.10160_for | GGGGACAAGTTTGTACAAAAAAGCAGGCTATGTTGCATTCCTCAAGAATTGTCCT | Tb927.9.10160_rev | GGGGACCACTTTGTACAAGAAAGCTGGGTCTTCACCAGATATGGGTCTTTCCACA |
| 3_hypothetical | B10 | Tb927.9.9720_for | GGGGACAAGTTTGTACAAAAAAGCAGGCTATGGCAGAGCCGGTGTTGG | Tb927.9.9720_rev | GGGGACCACTTTGTACAAGAAAGCTGGGTTTACAGTGCAAATATCATATCAGCATCGG |
| 3_hypothetical | B11 | Tb927.11.4370_for | GGGGACAAGTTTGTACAAAAAAGCAGGCTATGAATGTTCCCGTAAAGCTTAGCG | Tb927.11.4370_rev | GGGGACCACTTTGTACAAGAAAGCTGGGTTTAACGGTGCAAACGCACTGC |
| 3_hypothetical | B12 | Tb927.10.11150_for | GGGGACAAGTTTGTACAAAAAAGCAGGCTATGGAAGAGCGTGATGAAATAACGG | Tb927.10.11150_rev | GGGGACCACTTTGTACAAGAAAGCTGGGTTCAACGAATCAGCCTATACATCTGC |
| 3_hypothetical | B13 | Tb927.10.3540_for | GGGGACAAGTTTGTACAAAAAAGCAGGCTATGAAGCTTCTAAAGTTCGTCAACTGT | Tb927.10.3540_rev | GGGGACCACTTTGTACAAGAAAGCTGGGTTCACGTTTCCTTTTTAATGCTTCGC |
| 3_hypothetical | B14 | Tb927.10.670_for | GGGGACAAGTTTGTACAAAAAAGCAGGCTATGTCGTATCGATTTAAGGGTCGC | Tb927.10.670_rev | GGGGACCACTTTGTACAAGAAAGCTGGGTTCACATGTCCCGCAGCCG |
| 3_hypothetical | B15 | Tb927.11.1770_for | GGGGACAAGTTTGTACAAAAAAGCAGGCTATGAGCGGTTGTGGCAGGA | Tb927.11.1770_rev | GGGGACCACTTTGTACAAGAAAGCTGGGTCTACTCAATACCAGTTTCCTCATCACTTC |
| 3_hypothetical | B16 | Tb927.9.11660_for | GGGGACAAGTTTGTACAAAAAAGCAGGCTATGACGGAGTATTTGAAGGATATTGGGC | Tb927.9.11660_rev | GGGGACCACTTTGTACAAGAAAGCTGGGTTTAATATAATGTTTCGGGTCTTTTGTCCTC |
| 3_hypothetical | B17 | Tb927.7.4880_for | GGGGACAAGTTTGTACAAAAAAGCAGGCTATGAGTCTCCTCTGCAACATTTCCA | Tb927.7.4880_rev | GGGGACCACTTTGTACAAGAAAGCTGGGTTCAAATTTTCGCACCAGTTTTAGATCC |
| 3_hypothetical | B18 | Tb927.10.5710_for | GGGGACAAGTTTGTACAAAAAAGCAGGCTATGGACAAGCAGGAAATGTTGATTCT | Tb927.10.5710_rev | GGGGACCACTTTGTACAAGAAAGCTGGGTTTATAGCAAGAAAAATAACGCTGAAAGCAT |
| 3_hypothetical | B19 | Tb927.7.6560_for | GGGGACAAGTTTGTACAAAAAAGCAGGCTATGGCCATGAGGCTTGCG | Tb927.7.6560_rev | GGGGACCACTTTGTACAAGAAAGCTGGGTTCACTGCGAGGTTACATTTTCACC |
| 3_hypothetical | B20 | Tb927.5.297b_for | GGGGACAAGTTTGTACAAAAAAGCAGGCTATGGGCCGCCACCATCAA | Tb927.5.297b_rev | GGGGACCACTTTGTACAAGAAAGCTGGGTCTACGAGTTCCCTACTCTCGGTT |
| 3_hypothetical | B21 | Tb927.11.12020_for | GGGGACAAGTTTGTACAAAAAAGCAGGCTATGGTGGACGAGCTTCCACC | Tb927.11.12020_rev | GGGGACCACTTTGTACAAGAAAGCTGGGTTCATCGAGCTAATTCAAAGTATTTTTCAAT |
| 3_hypothetical | B22 | Tb927.8.7330_for | GGGGACAAGTTTGTACAAAAAAGCAGGCTATGATGGGGAAAGTGCTGTTATTTCTG | Tb927.8.7330_rev | GGGGACCACTTTGTACAAGAAAGCTGGGTTCAAAGTAACGTGCTGTATAAAAAAGGAAT |
| 3_hypothetical | B23 | Tb927.1.1790_for | GGGGACAAGTTTGTACAAAAAAGCAGGCTATGGGAATTTATTCCGACACTGATGATG | Tb927.1.1790_rev | GGGGACCACTTTGTACAAGAAAGCTGGGTTCATTCAAATTTCCCACCTCTTAAACCA |

| | | | | | |
|---|---|---|---|---|---|
| 3_hypothetical | B24 | Tb927.4.4790_for | GGGGACAAGTTTGTACAAAAAAGCAGGCTATGAGTAATGCCCCAACACTGC | Tb927.4.4790_rev | GGGGACCACTTTGTACAAGAAAGCTGGGTTTACCGTGGAGAGAAAATGAGGGG |
| 3_hypothetical | C1 | Tb927.6.2500_for | GGGGACAAGTTTGTACAAAAAAGCAGGCTATGAGTTGGTTAACTCCTTCAGGTTCC | Tb927.6.2500_rev | GGGGACCACTTTGTACAAGAAAGCTGGGTTTACCACAGCCCTTTGCGAGC |
| 3_hypothetical | C2 | Tb927.5.1160_for | GGGGACAAGTTTGTACAAAAAAGCAGGCTATGTTCTCTATGTCGCCATGCG | Tb927.5.1160_rev | GGGGACCACTTTGTACAAGAAAGCTGGGTTCAAAGATCGCCACGGCCG |
| 3_hypothetical | C3 | Tb927.7.6880_for | GGGGACAAGTTTGTACAAAAAAGCAGGCTATGAAATACTGCGAAGTGCAACCG | Tb927.7.6880_rev | GGGGACCACTTTGTACAAGAAAGCTGGGTCTATCTTCTGCATATTTTCGCCCTCA |
| 3_hypothetical | C4 | Tb927.2.5190_for | GGGGACAAGTTTGTACAAAAAAGCAGGCTATGGCGTCCAAGAAAGGAGGG | Tb927.2.5190_rev | GGGGACCACTTTGTACAAGAAAGCTGGGTTCAACTACTCACCGTCGCTGG |
| 3_hypothetical | C5 | Tb927.9.15850_for | GGGGACAAGTTTGTACAAAAAAGCAGGCTATGGACGGAAACTCGGAGGAT | Tb927.9.15850_rev | GGGGACCACTTTGTACAAGAAAGCTGGGTTCACGATTTTGTGGAACCGAGAAT |
| 3_hypothetical | C6 | Tb927.10.7210_for | GGGGACAAGTTTGTACAAAAAAGCAGGCTATGTCCCGCGCCACTTATAGC | Tb927.10.7210_rev | GGGGACCACTTTGTACAAGAAAGCTGGGTTTATCCTTCATCAGAATCATCCATATCATT |
| 3_hypothetical | C7 | Tb927.10.12410_for | GGGGACAAGTTTGTACAAAAAAGCAGGCTATGGAGGTTAAGATTGGAAAGGATGTTG | Tb927.10.12410_rev | GGGGACCACTTTGTACAAGAAAGCTGGGTTTATGACTCAAGCAGTAATACAGACGC |
| 3_hypothetical | C8 | Tb927.3.510_for | GGGGACAAGTTTGTACAAAAAAGCAGGCTATGAAATACTGCGAAGTACAACCGC | Tb927.3.510_rev | GGGGACCACTTTGTACAAGAAAGCTGGGTCTATCTTCTGCATATTTTCGCCCTCA |
| 3_hypothetical | C9 | Tb927.8.7180_for | GGGGACAAGTTTGTACAAAAAAGCAGGCTATGCCCACGCCACAGCAG | Tb927.8.7180_rev | GGGGACCACTTTGTACAAGAAAGCTGGGTTCATAGCTCACCCTCGGAATTGA |
| 3_hypothetical | C10 | Tb927.10.11330_for | GGGGACAAGTTTGTACAAAAAAGCAGGCTATGTCCAAAAAGGAAGCAATATCGACA | Tb927.10.11330_rev | GGGGACCACTTTGTACAAGAAAGCTGGGTCTATTCATTGGCTAGGACGTCGAC |
| 3_hypothetical | C11 | Tb927.3.2260_for | GGGGACAAGTTTGTACAAAAAAGCAGGCTATGAGGCGCGGCGCCCTT | Tb927.3.2260_rev | GGGGACCACTTTGTACAAGAAAGCTGGGTTCATGAACTCTGCTTCATCCCACT |
| 3_hypothetical | C12 | Tb927.11.9630_for | GGGGACAAGTTTGTACAAAAAAGCAGGCTATGTTTGGCCGGTTATTTGGTTTTAAG | Tb927.11.9630_rev | GGGGACCACTTTGTACAAGAAAGCTGGGTTCACGACAGTAGTTCCCTCTCTAGT |
| 3_hypothetical | C13 | Tb927.6.1410_for | GGGGACAAGTTTGTACAAAAAAGCAGGCTATGCGATGCGTGAGATCCG | Tb927.6.1410_rev | GGGGACCACTTTGTACAAGAAAGCTGGGTTCATCCCTTGCCGATGGACT |
| 3_hypothetical | C14 | Tb927.9.2600_for | GGGGACAAGTTTGTACAAAAAAGCAGGCTATGAGCACCGCTCAGTCGT | Tb927.9.2600_rev | GGGGACCACTTTGTACAAGAAAGCTGGGTTCAAGTGAAGGCGCTCGC |
| 3_hypothetical | C15 | Tb927.4.340_for | GGGGACAAGTTTGTACAAAAAAGCAGGCTATGCGTTTGACAAGTAGAGGGAGT | Tb927.4.340_rev | GGGGACCACTTTGTACAAGAAAGCTGGGTTCACATTTCACCTAACCTCCCTTCT |
| 3_hypothetical | C16 | Tb927.4.3720_for | GGGGACAAGTTTGTACAAAAAAGCAGGCTATGCCTAGTAGGCGAGTCACG | Tb927.4.3720_rev | GGGGACCACTTTGTACAAGAAAGCTGGGTCTATTCCTCATCAATAGAGTCATCATCCTC |
| 3_hypothetical | C17 | Tb927.7.5840_for | GGGGACAAGTTTGTACAAAAAAGCAGGCTATGAGCAGTTCCAATGATATCGCC | Tb927.7.5840_rev | GGGGACCACTTTGTACAAGAAAGCTGGGTTCAATGGAAGAATCCAATGATTTGCCT |
| 3_hypothetical | C18 | Tb927.7.4430_for | GGGGACAAGTTTGTACAAAAAAGCAGGCTATGCGTTCCGTTATTCGTTCCTG | Tb927.7.4430_rev | GGGGACCACTTTGTACAAGAAAGCTGGGTTCAATCATTCCATGGACTTTCTAATGCT |
| 3_hypothetical | C19 | Tb927.2.2270_for | GGGGACAAGTTTGTACAAAAAAGCAGGCTATGTCACATGTGAAGGCATTTGACC | Tb927.2.2270_rev | GGGGACCACTTTGTACAAGAAAGCTGGGTTTACCCAATAATAAAAAAGAGTTCATAAAG |
| 3_hypothetical | C20 | Tb927.10.7770_for | GGGGACAAGTTTGTACAAAAAAGCAGGCTATGCCATATAAGTCGGGTGTCGC | Tb927.10.7770_rev | GGGGACCACTTTGTACAAGAAAGCTGGGTCTAATCAGAAGCGGAACTGTAATCACT |
| 3_hypothetical | C22 | Tb927.10.15110_for | GGGGACAAGTTTGTACAAAAAAGCAGGCTATGGTGGTCATACATAGTGGACTTGT | Tb927.10.15110_rev | GGGGACCACTTTGTACAAGAAAGCTGGGTTTACATAATATGTGAGGATACACCACACAC |
| 3_hypothetical | C23 | Tb927.10.15960_for | GGGGACAAGTTTGTACAAAAAAGCAGGCTATGTCTACGTGGGAGCAGGC | Tb927.10.15960_rev | GGGGACCACTTTGTACAAGAAAGCTGGGTTCAACACGAGTTGAGGAGTTGC |
| 3_hypothetical | C24 | Tb927.9.3620_for | GGGGACAAGTTTGTACAAAAAAGCAGGCTATGACACCTGCTGTAGTTTCGATG | Tb927.9.3620_rev | GGGGACCACTTTGTACAAGAAAGCTGGGTTCATCCTCCCAAAGACCGGTC |
| 3_hypothetical | D1 | Tb927.4.900_for | GGGGACAAGTTTGTACAAAAAAGCAGGCTATGCGTACAAGTGCGGACATT | Tb927.4.900_rev | GGGGACCACTTTGTACAAGAAAGCTGGGTTTAACTGTGTACTAAATCAGGACTCCGC |
| 3_hypothetical | D2 | Tb927.10.15090_for | GGGGACAAGTTTGTACAAAAAAGCAGGCTATGGTGGTCATACATAGTGGACTTGT | Tb927.10.15090_rev | GGGGACCACTTTGTACAAGAAAGCTGGGTTTACATAACATGTGAGGATACACCACAC |
| 3_hypothetical | D3 | Tb927.8.4190_for | GGGGACAAGTTTGTACAAAAAAGCAGGCTATGAGCGGCCATAGGCATTTG | Tb927.8.4190_rev | GGGGACCACTTTGTACAAGAAAGCTGGGTTTACTCCCCAGTAAAAATGTTATATGTTTG |
| 3_hypothetical | D4 | Tb927.4.4810_for | GGGGACAAGTTTGTACAAAAAAGCAGGCTATGGAGCTTACGCAAGCAATAAACC | Tb927.4.4810_rev | GGGGACCACTTTGTACAAGAAAGCTGGGTCTAATATGTGGTGGAGGGAACATCG |
| 3_hypothetical | D5 | Tb927.10.5700_for | GGGGACAAGTTTGTACAAAAAAGCAGGCTATGGACAAGCAGGAAATGTTGATTCT | Tb927.10.5700_rev | GGGGACCACTTTGTACAAGAAAGCTGGGTTTATAGCAAGAAAAATAACGTTGAAAGCAT |
| 3_hypothetical | D6 | Tb927.3.1730_for | GGGGACAAGTTTGTACAAAAAAGCAGGCTATGCTTCGCTCCCTCTTTGGA | Tb927.3.1730_rev | GGGGACCACTTTGTACAAGAAAGCTGGGTTCACGTCGCCCGTCCGCC |
| 3_hypothetical | D7 | Tb927.4.2830_for | GGGGACAAGTTTGTACAAAAAAGCAGGCTATGCGACCAGGTAGTCAGTTCA | Tb927.4.2830_rev | GGGGACCACTTTGTACAAGAAAGCTGGGTCTACACACGTTTCCCCTCCGA |
| 3_hypothetical | D8 | Tb927.7.4230_for | GGGGACAAGTTTGTACAAAAAAGCAGGCTATGGTGATGAAGGCTTTCCCCC | Tb927.7.4230_rev | GGGGACCACTTTGTACAAGAAAGCTGGGTCTACACGCGTGTTCTCGACC |
| 3_hypothetical | D9 | Tb927.4.5230_for | GGGGACAAGTTTGTACAAAAAAGCAGGCTATGCCGATACCCCCGAATAACG | Tb927.4.5230_rev | GGGGACCACTTTGTACAAGAAAGCTGGGTCTACACGTCATCTTCCGAGAATATGG |
| 3_hypothetical | D10 | Tb927.11.6930_for | GGGGACAAGTTTGTACAAAAAAGCAGGCTATGGAAAGAGACTTAAATGACAGTAGGGC | Tb927.11.6930_rev | GGGGACCACTTTGTACAAGAAAGCTGGGTTCATCGCCTAACCACAGACGG |
| 3_hypothetical | D11 | Tb927.8.2140_for | GGGGACAAGTTTGTACAAAAAAGCAGGCTATGGCTTGTGGTGCGATTGT | Tb927.8.2140_rev | GGGGACCACTTTGTACAAGAAAGCTGGGTTTAGAACGCGTTGTGACGCA |
| 3_hypothetical | D12 | Tb927.3.600_for | GGGGACAAGTTTGTACAAAAAAGCAGGCTATGTCAAGGTTTGGGGCTTGG | Tb927.3.600_rev | GGGGACCACTTTGTACAAGAAAGCTGGGTCTACAGGGAGAATAACAATACTGAATATAT |
| 3_hypothetical | D13 | Tb927.9.15870_for | GGGGACAAGTTTGTACAAAAAAGCAGGCTATGACAAGCCTCACCGAAGAGC | Tb927.9.15870_rev | GGGGACCACTTTGTACAAGAAAGCTGGGTCTACTCTGCATTGTAACCCGATTCC |
| 3_hypothetical | D14 | Tb927.10.9270_for | GGGGACAAGTTTGTACAAAAAAGCAGGCTATGAAGAGTTTCCGCGTAGTTTGC | Tb927.10.9270_rev | GGGGACCACTTTGTACAAGAAAGCTGGGTTTAAGCAATCTTCTGCTTTATAAAGAATCG |
| 3_hypothetical | D15 | Tb927.4.480_for | GGGGACAAGTTTGTACAAAAAAGCAGGCTATGGTAACGGGGGATATGTCGAC | Tb927.4.480_rev | GGGGACCACTTTGTACAAGAAAGCTGGGTTCATGGTGTTTCACAGGTCTCCC |
| 3_hypothetical | D16 | Tb927.10.5570_for | GGGGACAAGTTTGTACAAAAAAGCAGGCTATGGAGTCTCCGCCCTCAAAT | Tb927.10.5570_rev | GGGGACCACTTTGTACAAGAAAGCTGGGTTCAAAGGCGCATACCGCAC |
| 3_hypothetical | D17 | Tb927.4.3090_for | GGGGACAAGTTTGTACAAAAAAGCAGGCTATGCACCCGTTCTACACTATACGT | Tb927.4.3090_rev | GGGGACCACTTTGTACAAGAAAGCTGGGTTCATCGTGCCTGATCCACAAATAC |
| 3_hypothetical | D18 | Tb927.7.7510_for | GGGGACAAGTTTGTACAAAAAAGCAGGCTATGGAAATGGAGCAGGCTCCA | Tb927.7.7510_rev | GGGGACCACTTTGTACAAGAAAGCTGGGTCTAGGAGGAAGGAGTTTGAAGAGTGT |
| 3_hypothetical | D19 | Tb927.10.5650_for | GGGGACAAGTTTGTACAAAAAAGCAGGCTATGTACCCACTCCTCGTGTGC | Tb927.10.5650_rev | GGGGACCACTTTGTACAAGAAAGCTGGGTCTAGCGGCCAGGCGAAAC |
| 3_hypothetical | D20 | Tb927.9.7000_for | GGGGACAAGTTTGTACAAAAAAGCAGGCTATGGAACCTAAGAGCGTCGAGG | Tb927.9.7000_rev | GGGGACCACTTTGTACAAGAAAGCTGGGTTTATTCCCTGACCGTGTAAACAAACA |
| 3_hypothetical | D21 | Tb927.7.2270_for | GGGGACAAGTTTGTACAAAAAAGCAGGCTATGATTGAACGCACCGATGTTGT | Tb927.7.2270_rev | GGGGACCACTTTGTACAAGAAAGCTGGGTCTAAACAAATGGATGTAGCATTAGTTGAAA |
| 3_hypothetical | D22 | Tb927.7.1260_for | GGGGACAAGTTTGTACAAAAAAGCAGGCTATGATGGAAAGTATGAACCTTGAACAACC | Tb927.7.1260_rev | GGGGACCACTTTGTACAAGAAAGCTGGGTTCAGAAGAAAATCAACGATATTGCTACAAG |
| 3_hypothetical | D23 | Tb927.9.12440_for | GGGGACAAGTTTGTACAAAAAAGCAGGCTATGTACAACTCTGGCGGGGAG | Tb927.9.12440_rev | GGGGACCACTTTGTACAAGAAAGCTGGGTTCACGATGATAATGGTCGCATATAATTTTC |
| 3_hypothetical | D24 | Tb927.11.1590_for | GGGGACAAGTTTGTACAAAAAAGCAGGCTATGAAGCGCGGGCGGTCG | Tb927.11.1590_rev | GGGGACCACTTTGTACAAGAAAGCTGGGTCTACTTTACTCGCGCCAACACC |
| 3_hypothetical | E1 | Tb927.7.1280_for | GGGGACAAGTTTGTACAAAAAAGCAGGCTATGATGGAAAGTATGAACCTTGAACAACC | Tb927.7.1280_rev | GGGGACCACTTTGTACAAGAAAGCTGGGTTCAAAAGAAAATCACCAGTATAGCCACA |
| 3_hypothetical | E2 | Tb927.9.9690_for | GGGGACAAGTTTGTACAAAAAAGCAGGCTATGGGGTGCCAGATTTCTGTGG | Tb927.9.9690_rev | GGGGACCACTTTGTACAAGAAAGCTGGGTTCATTCATTACCTTCACCCTCATCGC |
| 3_hypothetical | E3 | Tb927.4.1600_for | GGGGACAAGTTTGTACAAAAAAGCAGGCTATGAAAGAAAATGGTGTTTTAGGCGG | Tb927.4.1600_rev | GGGGACCACTTTGTACAAGAAAGCTGGGTTTAGTGCATTTGGCTCCACCCC |
| 3_hypothetical | E4 | Tb927.11.16750_for | GGGGACAAGTTTGTACAAAAAAGCAGGCTATGTCTTCAGGTGACAGCAGCG | Tb927.11.16750_rev | GGGGACCACTTTGTACAAGAAAGCTGGGTTCACGATGACTGACCTGGCA |
| 3_hypothetical | E5 | Tb927.8.3470_for | GGGGACAAGTTTGTACAAAAAAGCAGGCTATGTCATCTGATCTTTTGCGAGAGTAC | Tb927.8.3470_rev | GGGGACCACTTTGTACAAGAAAGCTGGGTTCACTTCCAAAATACAATGATTAGAAAGGT |
| 3_hypothetical | E6 | Tb927.9.6470_for | GGGGACAAGTTTGTACAAAAAAGCAGGCTATGACACTGGAGGGCGGC | Tb927.9.6470_rev | GGGGACCACTTTGTACAAGAAAGCTGGGTTTAGGCCCCTTCGCCAATCG |
| 3_hypothetical | E7 | Tb927.8.2330_for | GGGGACAAGTTTGTACAAAAAAGCAGGCTATGAGCTCTGCTGAGCGGC | Tb927.8.2330_rev | GGGGACCACTTTGTACAAGAAAGCTGGGTCTAATCAAACTCACCATCTGCGTACC |
| 3_hypothetical | E8 | Tb927.11.5080_for | GGGGACAAGTTTGTACAAAAAAGCAGGCTATGCAGCCGCATGGTGGA | Tb927.11.5080_rev | GGGGACCACTTTGTACAAGAAAGCTGGGTTCAACCGCGCCTCACATCC |
| 3_hypothetical | E9 | Tb927.5.730_for | GGGGACAAGTTTGTACAAAAAAGCAGGCTATGTTCAGTGCAGGAGCTAGTGA | Tb927.5.730_rev | GGGGACCACTTTGTACAAGAAAGCTGGGTTTACGATTTGAGAAACTGGTAAGAGCG |
| 3_hypothetical | E10 | Tb927.9.7670_for | GGGGACAAGTTTGTACAAAAAAGCAGGCTATGGCGAAGTCCACCCGC | Tb927.9.7670_rev | GGGGACCACTTTGTACAAGAAAGCTGGGTTTACCTCCACTGCACCTTCTTCT |
| 3_hypothetical | E11 | Tb927.1.30_for | GGGGACAAGTTTGTACAAAAAAGCAGGCTATGAGGACCAACAAAAAAACACAGAGG | Tb927.1.30_rev | GGGGACCACTTTGTACAAGAAAGCTGGGTCTACACATATCGTCCCCCTATTACCA |
| 3_hypothetical | E12 | Tb927.11.5230_for | GGGGACAAGTTTGTACAAAAAAGCAGGCTATGAGTGCTTCTGGAAGTGGAGT | Tb927.11.5230_rev | GGGGACCACTTTGTACAAGAAAGCTGGGTTCAGTAGACATGCTCTGAAGGCA |
| 3_hypothetical | E13 | Tb927.7.6570_for | GGGGACAAGTTTGTACAAAAAAGCAGGCTATGAATCTAGCGAAGAAAATCATCGTGT | Tb927.7.6570_rev | GGGGACCACTTTGTACAAGAAAGCTGGGTTCAATTAGCAGATTCTGTGCCCG |
| 3_hypothetical | E14 | Tb927.11.2350_for | GGGGACAAGTTTGTACAAAAAAGCAGGCTATGTATGCTTCGCATAAAAAGGATATTAAC | Tb927.11.2350_rev | GGGGACCACTTTGTACAAGAAAGCTGGGTTCACTTTTTCATATCCAGCAGTTCTTGC |
| 3_hypothetical | E15 | Tb927.11.13470_for | GGGGACAAGTTTGTACAAAAAAGCAGGCTATGGAACAAGGTCAACCCGC | Tb927.11.13470_rev | GGGGACCACTTTGTACAAGAAAGCTGGGTTCACCGTACTGCCTTTCTGTCT |
| 3_hypothetical | E16 | Tb927.6.4270_for | GGGGACAAGTTTGTACAAAAAAGCAGGCTATGAAGCGGTCTTTTTTGTTTCTGAC | Tb927.6.4270_rev | GGGGACCACTTTGTACAAGAAAGCTGGGTTCAATGAACTGCAGCACTACTATTTCC |
| 3_hypothetical | E17 | Tb927.8.3570_for | GGGGACAAGTTTGTACAAAAAAGCAGGCTATGACCACGAATCTTGTCCTTCCA | Tb927.8.3570_rev | GGGGACCACTTTGTACAAGAAAGCTGGGTTTACGGTATAAACACACGAGTTGTTTTTC |
| 3_hypothetical | E18 | Tb927.6.4250_for | GGGGACAAGTTTGTACAAAAAAGCAGGCTATGTGGTTTGCTGAAGTGCTTGT | Tb927.6.4250_rev | GGGGACCACTTTGTACAAGAAAGCTGGGTCTACACTTTTTGTGGCACACCGA |
| 3_hypothetical | E19 | Tb927.7.1800_for | GGGGACAAGTTTGTACAAAAAAGCAGGCTATGTCGTCTTTAATGTCGGCAAATAATAGT | Tb927.7.1800_rev | GGGGACCACTTTGTACAAGAAAGCTGGGTTTATTTTTTTCGCGTAGTAATGCCCATG |
| 3_hypothetical | E20 | Tb927.8.6400_for | GGGGACAAGTTTGTACAAAAAAGCAGGCTATGCGCCACTCCGTGAAGT | Tb927.8.6400_rev | GGGGACCACTTTGTACAAGAAAGCTGGGTTCAATCACTCTTCTTCCCCTGTAGG |
| 3_hypothetical | E21 | Tb927.6.3060_for | GGGGACAAGTTTGTACAAAAAAGCAGGCTATGAAAGGCTTGGAAGAGTGGC | Tb927.6.3060_rev | GGGGACCACTTTGTACAAGAAAGCTGGGTTCAATACCGCGGCATCGGT |
| 3_hypothetical | E22 | Tb927.3.5200_for | GGGGACAAGTTTGTACAAAAAAGCAGGCTATGGCATGTGATGATTCGACGAA | Tb927.3.5200_rev | GGGGACCACTTTGTACAAGAAAGCTGGGTTTATCGACACCACCCTCCTGTG |
| 3_hypothetical | E23 | Tb927.10.8960_for | GGGGACAAGTTTGTACAAAAAAGCAGGCTATGTTGCGTTTTACGGTTGCG | Tb927.10.8960_rev | GGGGACCACTTTGTACAAGAAAGCTGGGTTTATTTTTTGCCTGCTGCCTTGTTC |
| 3_hypothetical | E24 | Tb927.11.6650_for | GGGGACAAGTTTGTACAAAAAAGCAGGCTATGGTTAAGCCTGCCGTGTTTC | Tb927.11.6650_rev | GGGGACCACTTTGTACAAGAAAGCTGGGTTCAAAGACAACCAGGCGGTTTT |
| 3_hypothetical | F1 | Tb927.2.4530_for | GGGGACAAGTTTGTACAAAAAAGCAGGCTATGCAATCTCCGAAGTTTGGTCG | Tb927.2.4530_rev | GGGGACCACTTTGTACAAGAAAGCTGGGTCTAGGAATGACCACCCGACACA |
| 3_hypothetical | F2 | Tb927.8.2830_for | GGGGACAAGTTTGTACAAAAAAGCAGGCTATGGAGGCTGTGGTCGCA | Tb927.8.2830_rev | GGGGACCACTTTGTACAAGAAAGCTGGGTCTAGTATTCGGGGCTGGTCCTC |
| 3_hypothetical | F3 | Tb927.10.9120_for | GGGGACAAGTTTGTACAAAAAAGCAGGCTATGTCCCGGTACAGTACAGGC | Tb927.10.9120_rev | GGGGACCACTTTGTACAAGAAAGCTGGGTTTAGCTGCAATAGCCGGCAATG |
| 3_hypothetical | F4 | Tb927.7.4720_for | GGGGACAAGTTTGTACAAAAAAGCAGGCTATGACTGTGCGTGGCGCC | Tb927.7.4720_rev | GGGGACCACTTTGTACAAGAAAGCTGGGTTTACTCGTTTGCCCGCTCCT |
| 3_hypothetical | F5 | Tb927.10.580_for | GGGGACAAGTTTGTACAAAAAAGCAGGCTATGCTTCTACAGTGTGGTTCTTGTAG | Tb927.10.580_rev | GGGGACCACTTTGTACAAGAAAGCTGGGTTCATTTCTGCAGCGACACGC |
| 3_hypothetical | F6 | Tb927.11.1920_for | GGGGACAAGTTTGTACAAAAAAGCAGGCTATGATGATCTTCATTATGTCAATAGAGCTG | Tb927.11.1920_rev | GGGGACCACTTTGTACAAGAAAGCTGGGTTTAAAGCGCAGCGCACCA |
| 3_hypothetical | F7 | Tb927.8.4790_for | GGGGACAAGTTTGTACAAAAAAGCAGGCTATGGCAGAGAACAGTCCTACATCT | Tb927.8.4790_rev | GGGGACCACTTTGTACAAGAAAGCTGGGTCTACAACCGAATTTGCATCAAGTCG |
| 3_hypothetical | F8 | Tb927.10.3070_for | GGGGACAAGTTTGTACAAAAAAGCAGGCTATGGACGCCGCAGATGCC | Tb927.10.3070_rev | GGGGACCACTTTGTACAAGAAAGCTGGGTTTACTGTGAAGCAAACCTTCTCCTC |
| 3_hypothetical | F9 | Tb927.7.5130_for | GGGGACAAGTTTGTACAAAAAAGCAGGCTATGAGCAACCACGGGGGC | Tb927.7.5130_rev | GGGGACCACTTTGTACAAGAAAGCTGGGTTCACAAGGGACACTTTGAGACCC |
| 3_hypothetical | F10 | Tb927.11.11620_for | GGGGACAAGTTTGTACAAAAAAGCAGGCTATGACCATTCTCCACAATCCCGG | Tb927.11.11620_rev | GGGGACCACTTTGTACAAGAAAGCTGGGTCTACAACTTTTTCCCTGCCTGCA |
| 3_hypothetical | F11 | Tb927.10.8410_for | GGGGACAAGTTTGTACAAAAAAGCAGGCTATGGACCGGAGCGAGACATT | Tb927.10.8410_rev | GGGGACCACTTTGTACAAGAAAGCTGGGTTTAGTCTGCTTTAATGCTATCGACTCG |
| 3_hypothetical | F12 | Tb927.8.2700_for | GGGGACAAGTTTGTACAAAAAAGCAGGCTATGATCCGACTGCACCGCA | Tb927.8.2700_rev | GGGGACCACTTTGTACAAGAAAGCTGGGTTCACCGTTCGGTGGCAAGG |
| 3_hypothetical | F13 | Tb927.8.6040_for | GGGGACAAGTTTGTACAAAAAAGCAGGCTATGCACCCGCGATTGTCG | Tb927.8.6040_rev | GGGGACCACTTTGTACAAGAAAGCTGGGTTCAACCACGAGGTAAAGTGGGG |
| 3_hypothetical | F14 | Tb927.7.350_for | GGGGACAAGTTTGTACAAAAAAGCAGGCTATGTCCTTGGCCTCAGTTCCT | Tb927.7.350_rev | GGGGACCACTTTGTACAAGAAAGCTGGGTCTATTGCATGCAAGCACAGGC |
| 3_hypothetical | F15 | Tb927.10.15200_for | GGGGACAAGTTTGTACAAAAAAGCAGGCTATGGCAGGCGGATCTACCTC | Tb927.10.15200_rev | GGGGACCACTTTGTACAAGAAAGCTGGGTCTAATCGGAATCGTACATGTCCCTCT |
| 3_hypothetical | F16 | Tb927.10.4390_for | GGGGACAAGTTTGTACAAAAAAGCAGGCTATGAATAGCATAAAAGTGTTGCTCGTTTT | Tb927.10.4390_rev | GGGGACCACTTTGTACAAGAAAGCTGGGTTTACGCTAAAAGAAAAAAAGGGATGAGAAA |
| 3_hypothetical | F17 | Tb927.5.2630_for | GGGGACAAGTTTGTACAAAAAAGCAGGCTATGGCCTTTTGGGAAAAGATATGGTG | Tb927.5.2630_rev | GGGGACCACTTTGTACAAGAAAGCTGGGTTTAGTGGAGGGCCTCCGC |
| 3_hypothetical | F18 | Tb927.7.3860_for | GGGGACAAGTTTGTACAAAAAAGCAGGCTATGCAACAAAAATATCTGGGCAGAGA | Tb927.7.3860_rev | GGGGACCACTTTGTACAAGAAAGCTGGGTTTAGTTGTTCTTTTTAACGACAGCCTTG |
| 3_hypothetical | F19 | Tb927.11.7800_for | GGGGACAAGTTTGTACAAAAAAGCAGGCTATGGCGACGGAACAACACC | Tb927.11.7800_rev | GGGGACCACTTTGTACAAGAAAGCTGGGTTCATACACGACGCAGTCCCC |
| 3_hypothetical | F20 | Tb927.10.6550_for | GGGGACAAGTTTGTACAAAAAAGCAGGCTATGCCGCAGCTGCGGGTG | Tb927.10.6550_rev | GGGGACCACTTTGTACAAGAAAGCTGGGTTTAAACTGAATGCCTTATAACAGCCCG |
| 3_hypothetical | F21 | Tb927.11.4250_for | GGGGACAAGTTTGTACAAAAAAGCAGGCTATGCAACAACCGATTGGCTTAACT | Tb927.11.4250_rev | GGGGACCACTTTGTACAAGAAAGCTGGGTCTATGTCAAGTAATTGTTCACATGTAGATT |
| 3_hypothetical | F22 | Tb927.11.6850_for | GGGGACAAGTTTGTACAAAAAAGCAGGCTATGCCAGCACCAGTTTACCG | Tb927.11.6850_rev | GGGGACCACTTTGTACAAGAAAGCTGGGTCTATGTTCCAAGTGCCTTTTTTGCA |
| 3_hypothetical | F23 | Tb927.8.1320_for | GGGGACAAGTTTGTACAAAAAAGCAGGCTATGACGTCCAAAGTTGAGGAGC | Tb927.8.1320_rev | GGGGACCACTTTGTACAAGAAAGCTGGGTTCATTTTTCAGCGTTTTTTTTCGCCT |
| 3_hypothetical | F24 | Tb927.11.3760_for | GGGGACAAGTTTGTACAAAAAAGCAGGCTATGGAAACGAAAGAGAGTTGCAGG | Tb927.11.3760_rev | GGGGACCACTTTGTACAAGAAAGCTGGGTTCAGTCATCATCATTGGCATGTAAACC |
| 3_hypothetical | G1 | Tb927.6.2490_for | GGGGACAAGTTTGTACAAAAAAGCAGGCTATGCGGCGGCAAATTCTTGG | Tb927.6.2490_rev | GGGGACCACTTTGTACAAGAAAGCTGGGTTTACGACAACTGTGGGAACGC |
| 3_hypothetical | G2 | Tb927.10.8660_for | GGGGACAAGTTTGTACAAAAAAGCAGGCTATGGATAGGGCCTCGGAACTTATG | Tb927.10.8660_rev | GGGGACCACTTTGTACAAGAAAGCTGGGTTTACGACCGTCGGAAAGCCA |
| 3_hypothetical | G3 | Tb927.11.7700_for | GGGGACAAGTTTGTACAAAAAAGCAGGCTATGTCCCGCGCCACCACA | Tb927.11.7700_rev | GGGGACCACTTTGTACAAGAAAGCTGGGTTTAACGAAGTGTAGGTAAGGGACCA |
| 3_hypothetical | G4 | Tb927.11.3520_for | GGGGACAAGTTTGTACAAAAAAGCAGGCTATGAAGGGGAATAAAGGTGCGAAC | Tb927.11.3520_rev | GGGGACCACTTTGTACAAGAAAGCTGGGTTTAGATGAGCCCAAACCACTCACC |
| 3_hypothetical | G5 | Tb927.5.720_for | GGGGACAAGTTTGTACAAAAAAGCAGGCTATGGCGTCTGTGTCTAGGCG | Tb927.5.720_rev | GGGGACCACTTTGTACAAGAAAGCTGGGTTCAATCAATCTCAAAAAGCCCCTCC |
| 3_hypothetical | G6 | Tb927.10.9090_for | GGGGACAAGTTTGTACAAAAAAGCAGGCTATGAGCTGGAGGGTTTCCTGC | Tb927.10.9090_rev | GGGGACCACTTTGTACAAGAAAGCTGGGTTCACACTTTTGATTTAACGTACTCAACAAA |
| 3_hypothetical | G7 | Tb927.11.2250_for | GGGGACAAGTTTGTACAAAAAAGCAGGCTATGTACAATAACGACTCGGCGG | Tb927.11.2250_rev | GGGGACCACTTTGTACAAGAAAGCTGGGTTTAGGAAGGCCCATCTGCCTT |
| 3_hypothetical | G8 | Tb927.11.13150_for | GGGGACAAGTTTGTACAAAAAAGCAGGCTATGTTTGACGATGCATCCCGTT | Tb927.11.13150_rev | GGGGACCACTTTGTACAAGAAAGCTGGGTTTAGTTGTACACCCGTTGGCG |
| 3_hypothetical | G9 | Tb927.6.4040_for | GGGGACAAGTTTGTACAAAAAAGCAGGCTATGTTACGGAACACAGATGTTCGAC | Tb927.6.4040_rev | GGGGACCACTTTGTACAAGAAAGCTGGGTTCAATTAGTCGAGACCTCCACATGA |
| 3_hypothetical | G10 | Tb927.5.4090_for | GGGGACAAGTTTGTACAAAAAAGCAGGCTATGGAAGAGGGCCATGCCA | Tb927.5.4090_rev | GGGGACCACTTTGTACAAGAAAGCTGGGTTTACAAATGAATCACGGCGTCGA |
| 3_hypothetical | G11 | Tb927.8.3880_for | GGGGACAAGTTTGTACAAAAAAGCAGGCTATGTGCGCTGGGCGGGAT | Tb927.8.3880_rev | GGGGACCACTTTGTACAAGAAAGCTGGGTCTATTTCATAATTCTGCACCACGGATATCC |
| 3_hypothetical | G12 | Tb927.11.12370_for | GGGGACAAGTTTGTACAAAAAAGCAGGCTATGTCAGCCGCTGGCATTG | Tb927.11.12370_rev | GGGGACCACTTTGTACAAGAAAGCTGGGTCTAACGTAGCTCGGATATGGGACG |
| 3_hypothetical | G13 | Tb927.7.1290_for | GGGGACAAGTTTGTACAAAAAAGCAGGCTATGTTTGGACGTGGTTGCATGC | Tb927.7.1290_rev | GGGGACCACTTTGTACAAGAAAGCTGGGTTCACTTGTGCTTCCTCTCGTGT |
| 3_hypothetical | G14 | Tb927.11.10440_for | GGGGACAAGTTTGTACAAAAAAGCAGGCTATGAGACAAGCGTCTGCGGG | Tb927.11.10440_rev | GGGGACCACTTTGTACAAGAAAGCTGGGTTCAGTCATCCCAGATATCGGGGT |
| 3_hypothetical | G15 | Tb927.11.11630_for | GGGGACAAGTTTGTACAAAAAAGCAGGCTATGTTGCGTGGCACTCGTG | Tb927.11.11630_rev | GGGGACCACTTTGTACAAGAAAGCTGGGTTTATTCCGAAAGAAACGCCTTGATATTTAT |
| 3_hypothetical | G16 | Tb927.9.1790_for | GGGGACAAGTTTGTACAAAAAAGCAGGCTATGGGTCGGGGGTTCGTG | Tb927.9.1790_rev | GGGGACCACTTTGTACAAGAAAGCTGGGTTCACTTGCTTTGTGGAAACGAAATATC |
| 3_hypothetical | G17 | Tb927.10.4760_for | GGGGACAAGTTTGTACAAAAAAGCAGGCTATGACGCAGAACATTCTTTTGGACC | Tb927.10.4760_rev | GGGGACCACTTTGTACAAGAAAGCTGGGTCTATTTACCTTCAAGGAGTAAACGGGG |
| 3_hypothetical | G18 | Tb927.8.2510_for | GGGGACAAGTTTGTACAAAAAAGCAGGCTATGAGCATTTCATGTTCCCTGGC | Tb927.8.2510_rev | GGGGACCACTTTGTACAAGAAAGCTGGGTCTACCAAAAAAATATGAGGAAGTCGCG |
| 3_hypothetical | G19 | Tb927.10.1750_for | GGGGACAAGTTTGTACAAAAAAGCAGGCTATGCGCGCGGTGCGGGCC | Tb927.10.1750_rev | GGGGACCACTTTGTACAAGAAAGCTGGGTTCACTTCGTCCGCTTTGATAAGTAG |
| 3_hypothetical | G20 | Tb927.10.7670_for | GGGGACAAGTTTGTACAAAAAAGCAGGCTATGGCGGGAAAAGGAAATGCC | Tb927.10.7670_rev | GGGGACCACTTTGTACAAGAAAGCTGGGTTTAATTGGCCCTTCGCGTCTTC |
| 3_hypothetical | G21 | Tb927.4.1910_for | GGGGACAAGTTTGTACAAAAAAGCAGGCTATGTCTTCCTTTAACATGCAAGCGT | Tb927.4.1910_rev | GGGGACCACTTTGTACAAGAAAGCTGGGTTTACGAAGTCTGTCCCTCTTCATTTACT |
| 3_hypothetical | G22 | Tb927.4.2620_for | GGGGACAAGTTTGTACAAAAAAGCAGGCTATGCGCGGCGTGGTGTAC | Tb927.4.2620_rev | GGGGACCACTTTGTACAAGAAAGCTGGGTCTAAGTGGCCGTGGGATAAATGTT |
| 3_hypothetical | G23 | Tb927.1.690_for | GGGGACAAGTTTGTACAAAAAAGCAGGCTATGTACCCTTCTTTTGTAAGTGAGTTGTC | Tb927.1.690_rev | GGGGACCACTTTGTACAAGAAAGCTGGGTTCATTTCACAAGAGCAGATGAAACAATAGA |
| 3_hypothetical | G24 | Tb927.11.5830_for | GGGGACAAGTTTGTACAAAAAAGCAGGCTATGGACGGTGTTGACGACTCC | Tb927.11.5830_rev | GGGGACCACTTTGTACAAGAAAGCTGGGTTCAGTTGGAACCCGCTCCTG |
| 3_hypothetical | H1 | Tb927.11.4930_for | GGGGACAAGTTTGTACAAAAAAGCAGGCTATGGTCGAAAGAACAGCGGATC | Tb927.11.4930_rev | GGGGACCACTTTGTACAAGAAAGCTGGGTTCATGCATCATCATCATCATCATCGT |
| 3_hypothetical | H2 | Tb927.11.6490_for | GGGGACAAGTTTGTACAAAAAAGCAGGCTATGTCTGCTAATGCTTGGCGG | Tb927.11.6490_rev | GGGGACCACTTTGTACAAGAAAGCTGGGTCTAACCCTTCGTGTCGCCCT |
| 3_hypothetical | H3 | Tb927.8.4760_for | GGGGACAAGTTTGTACAAAAAAGCAGGCTATGAAGGACAAAAATAAGTGCACTTTTAGG | Tb927.8.4760_rev | GGGGACCACTTTGTACAAGAAAGCTGGGTCTACCTTTGTCCTCCTTTTAAATACCATTC |
| 3_hypothetical | H4 | Tb927.10.1410_for | GGGGACAAGTTTGTACAAAAAAGCAGGCTATGTTTCGGCGATGTGGTTTGC | Tb927.10.1410_rev | GGGGACCACTTTGTACAAGAAAGCTGGGTTCATACGCTTTGGTGTCCTCTTGT |
| 3_hypothetical | H5 | Tb927.10.8010_for | GGGGACAAGTTTGTACAAAAAAGCAGGCTATGGCAGCCGAGCAGCCA | Tb927.10.8010_rev | GGGGACCACTTTGTACAAGAAAGCTGGGTCTACAGGTTCGTCGTCCATGGT |
| 3_hypothetical | H6 | Tb927.3.880_for | GGGGACAAGTTTGTACAAAAAAGCAGGCTATGTTAGGGTTGACAATCAGTCGGT | Tb927.3.880_rev | GGGGACCACTTTGTACAAGAAAGCTGGGTTCATGATAAAGAAACCAACGATTGAGGA |
| 3_hypothetical | H7 | Tb927.6.2750_for | GGGGACAAGTTTGTACAAAAAAGCAGGCTATGGGCGAGACTTTAGGCAGC | Tb927.6.2750_rev | GGGGACCACTTTGTACAAGAAAGCTGGGTTCAACGTCGTTTCCCTGCAC |
| 3_hypothetical | H8 | Tb927.9.1660_for | GGGGACAAGTTTGTACAAAAAAGCAGGCTATGACACTTGATCTCGTTGGTAAACTT | Tb927.9.1660_rev | GGGGACCACTTTGTACAAGAAAGCTGGGTTCACTCAGATCGGGGGCG |
| 3_hypothetical | H9 | Tb927.10.1430_for | GGGGACAAGTTTGTACAAAAAAGCAGGCTATGGTGGAATATAAGAGCGCCGC | Tb927.10.1430_rev | GGGGACCACTTTGTACAAGAAAGCTGGGTTCAGGGCGCCTTCTTCTCG |
| 3_hypothetical | H10 | Tb927.1.1390_for | GGGGACAAGTTTGTACAAAAAAGCAGGCTATGCTGGTTCGAACATGGATGTT | Tb927.1.1390_rev | GGGGACCACTTTGTACAAGAAAGCTGGGTCTATGCCACTTGCTGCCCATC |

| | | | | | |
|---|---|---|---|---|---|
| 3_hypothetical | H11 | Tb927.11.770_for | GGGGACAAGTTTGTACAAAAAAGCAGGCTATGGCGTCAGAGAAGTCACTACA | Tb927.11.770_rev | GGGGACCACTTTGTACAAGAAAGCTGGGTCTACAGGCTTAGCCACGGGT |
| 3_hypothetical | H12 | Tb927.10.5000_for | GGGGACAAGTTTGTACAAAAAAGCAGGCTATGACAGCACTATCGAATGAAGAAATGT | Tb927.10.5000_rev | GGGGACCACTTTGTACAAGAAAGCTGGGTTCACATACTCGCATCTTGATGCAA |
| 3_hypothetical | H13 | Tb927.7.3800_for | GGGGACAAGTTTGTACAAAAAAGCAGGCTATGAAGGCGACTGAAGAAGAAGC | Tb927.7.3800_rev | GGGGACCACTTTGTACAAGAAAGCTGGGTCTACAACTCCCACCACAGGGA |
| 3_hypothetical | H14 | Tb927.10.4380_for | GGGGACAAGTTTGTACAAAAAAGCAGGCTATGAATAACCTGAAAGTGTTGCTTGTGT | Tb927.10.4380_rev | GGGGACCACTTTGTACAAGAAAGCTGGGTTTACGCTAAAAGAAAAAAAGGGATGAGAAA |
| 3_hypothetical | H15 | Tb927.9.7100_for | GGGGACAAGTTTGTACAAAAAAGCAGGCTATGGCATCCAACGGTTTGGC | Tb927.9.7100_rev | GGGGACCACTTTGTACAAGAAAGCTGGGTCTATGCACGGAATTCACCTACGAA |
| 3_hypothetical | H16 | Tb927.7.1490_for | GGGGACAAGTTTGTACAAAAAAGCAGGCTATGAGACCGTGTGCGCCT | Tb927.7.1490_rev | GGGGACCACTTTGTACAAGAAAGCTGGGTTTACAGCTCCTCTTTGATCTCCTTCG |
| 3_hypothetical | H17 | Tb927.6.4310_for | GGGGACAAGTTTGTACAAAAAAGCAGGCTATGATGAATCACTCACCCCATGAAC | Tb927.6.4310_rev | GGGGACCACTTTGTACAAGAAAGCTGGGTCTACTGAGGCGGAAGGCTTAGG |
| 3_hypothetical | H18 | Tb927.5.292b_for | GGGGACAAGTTTGTACAAAAAAGCAGGCTATGCTACCTGTGATTTTCGTGTACG | Tb927.5.292b_rev | GGGGACCACTTTGTACAAGAAAGCTGGGTCTAAACCATTGACAAATATCCCAAGACTAA |
| 3_hypothetical | H19 | Tb927.1.2970_for | GGGGACAAGTTTGTACAAAAAAGCAGGCTATGAAACATGTTTCTGAAATGTTCATCCC | Tb927.1.2970_rev | GGGGACCACTTTGTACAAGAAAGCTGGGTTCAAATACGGTTGGGCTTCTGTTC |
| 3_hypothetical | H20 | Tb927.6.3680_for | GGGGACAAGTTTGTACAAAAAAGCAGGCTATGGAGCATCTACCAACTATCGACT | Tb927.6.3680_rev | GGGGACCACTTTGTACAAGAAAGCTGGGTTTATTCTTGCTTTTTATTTGCGTCTTCCC |
| 3_hypothetical | H21 | Tb927.11.870_for | GGGGACAAGTTTGTACAAAAAAGCAGGCTATGGTGGTTAGTCTGAGGAGGTTT | Tb927.11.870_rev | GGGGACCACTTTGTACAAGAAAGCTGGGTTCATTGAGTGATTCGTTCCTGGGG |
| 3_hypothetical | H22 | Tb927.10.6120_for | GGGGACAAGTTTGTACAAAAAAGCAGGCTATGGCCTCTGTGGAACCTGAT | Tb927.10.6120_rev | GGGGACCACTTTGTACAAGAAAGCTGGGTCTACAACTCGAACTTCTCGTCACC |
| 3_hypothetical | H23 | Tb927.9.11640_for | GGGGACAAGTTTGTACAAAAAAGCAGGCTATGAGTTCCCTTCCTCCGGAC | Tb927.9.11640_rev | GGGGACCACTTTGTACAAGAAAGCTGGGTTCATGGTGACATAAAAAAGGACGGTAA |
| 3_hypothetical | H24 | Tb927.3.1160_for | GGGGACAAGTTTGTACAAAAAAGCAGGCTATGTTTGCCCCATTCTTTCGTTCA | Tb927.3.1160_rev | GGGGACCACTTTGTACAAGAAAGCTGGGTTTAATCCCCTGTGTAACGCGGC |
| 3_hypothetical | I1 | Tb927.4.5440_for | GGGGACAAGTTTGTACAAAAAAGCAGGCTATGTCACATGTGTGCGATGTCTT | Tb927.4.5440_rev | GGGGACCACTTTGTACAAGAAAGCTGGGTTCACTGTCGTGGCGTGCG |
| 3_hypothetical | I2 | Tb927.8.7280_for | GGGGACAAGTTTGTACAAAAAAGCAGGCTATGATTGTTCCATCGTTGAACGGC | Tb927.8.7280_rev | GGGGACCACTTTGTACAAGAAAGCTGGGTCTACTGAGGCTTACACCGCAGA |
| 3_hypothetical | I3 | Tb927.4.1710_for | GGGGACAAGTTTGTACAAAAAAGCAGGCTATGGCTATGCGAGACGTCATTG | Tb927.4.1710_rev | GGGGACCACTTTGTACAAGAAAGCTGGGTTCATCGCGGACGGGGCTT |
| 3_hypothetical | I4 | Tb927.11.7470_for | GGGGACAAGTTTGTACAAAAAAGCAGGCTATGTCATTTGTACCCGCTTCACAA | Tb927.11.7470_rev | GGGGACCACTTTGTACAAGAAAGCTGGGTCTACCACTCACGCGTTGGTAAG |
| 3_hypothetical | I5 | Tb927.9.13120_for | GGGGACAAGTTTGTACAAAAAAGCAGGCTATGACAGACTTGGACGCGACC | Tb927.9.13120_rev | GGGGACCACTTTGTACAAGAAAGCTGGGTTTACTCTGAAAGTGAGACATTTGCCTG |
| 3_hypothetical | I6 | Tb927.9.15240_for | GGGGACAAGTTTGTACAAAAAAGCAGGCTATGCTTGCTGCTACCCGAAGT | Tb927.9.15240_rev | GGGGACCACTTTGTACAAGAAAGCTGGGTCTACTCTTTCCTATATCCATTGCAGCC |
| 3_hypothetical | I7 | Tb927.8.6280_for | GGGGACAAGTTTGTACAAAAAAGCAGGCTATGAGCGGAAAACATCTTTTGCATC | Tb927.8.6280_rev | GGGGACCACTTTGTACAAGAAAGCTGGGTTCACATCTCTCCGCACAGAGTG |
| 3_hypothetical | I8 | Tb927.4.2840_for | GGGGACAAGTTTGTACAAAAAAGCAGGCTATGCGTTGCCATGTGATGGG | Tb927.4.2840_rev | GGGGACCACTTTGTACAAGAAAGCTGGGTCTAATCTTCGTAGCACTCACGCAG |
| 3_hypothetical | I9 | Tb927.11.4260_for | GGGGACAAGTTTGTACAAAAAAGCAGGCTATGAACCCACTTTTTGCCGGC | Tb927.11.4260_rev | GGGGACCACTTTGTACAAGAAAGCTGGGTCTACACCTTTCCGTAGCGAGC |
| 3_hypothetical | I10 | Tb927.10.11120_for | GGGGACAAGTTTGTACAAAAAAGCAGGCTATGCAGGCCCAACACTATCATCT | Tb927.10.11120_rev | GGGGACCACTTTGTACAAGAAAGCTGGGTTCACGATAAAGGTGGAAGACCATTTTC |
| 3_hypothetical | I11 | Tb927.6.690_for | GGGGACAAGTTTGTACAAAAAAGCAGGCTATGAACTGTATCGAGTCAAAGTGTGC | Tb927.6.690_rev | GGGGACCACTTTGTACAAGAAAGCTGGGTTTAACAGAAGAGATACGACTCCTTTGGA |
| 3_hypothetical | I12 | Tb927.10.2080_for | GGGGACAAGTTTGTACAAAAAAGCAGGCTATGCTGTGGATTCAAATATGTACATTTGTA | Tb927.10.2080_rev | GGGGACCACTTTGTACAAGAAAGCTGGGTCTATTGCTTATGATTTTTACTCTTACTTC |
| 3_hypothetical | I13 | Tb927.7.630_for | GGGGACAAGTTTGTACAAAAAAGCAGGCTATGCGCGCAGGCTGCGTT | Tb927.7.630_rev | GGGGACCACTTTGTACAAGAAAGCTGGGTCTACACTTGATACTTAGATTTGGGGCG |
| 3_hypothetical | I14 | Tb927.10.9450_for | GGGGACAAGTTTGTACAAAAAAGCAGGCTATGAGCACCTTCGCGGTACT | Tb927.10.9450_rev | GGGGACCACTTTGTACAAGAAAGCTGGGTTTATTTTTGCGTCTCCCCTTCTTTTGA |
| 3_hypothetical | I15 | Tb927.10.12030_for | GGGGACAAGTTTGTACAAAAAAGCAGGCTATGTTAACTATCCGTGTCGGTAACCT | Tb927.10.12030_rev | GGGGACCACTTTGTACAAGAAAGCTGGGTCTATGAAGAAATGAGCTCCGTCGC |
| 3_hypothetical | I16 | Tb927.6.2120_for | GGGGACAAGTTTGTACAAAAAAGCAGGCTATGACAGGTGCGGATAGGTTTAACA | Tb927.6.2120_rev | GGGGACCACTTTGTACAAGAAAGCTGGGTTCATCGCTTCCTTCTCAGCGT |
| 3_hypothetical | I17 | Tb927.9.4290_for | GGGGACAAGTTTGTACAAAAAAGCAGGCTATGGCACACTGTTGCTTTCTCA | Tb927.9.4290_rev | GGGGACCACTTTGTACAAGAAAGCTGGGTTTATTTATCGAACCCCAACTCTCTAATGAG |
| 3_hypothetical | I18 | Tb927.4.700_for | GGGGACAAGTTTGTACAAAAAAGCAGGCTATGATGAGGCGGCAAACTTATGTT | Tb927.4.700_rev | GGGGACCACTTTGTACAAGAAAGCTGGGTCTACGTAAACGGGCTAACGGC |
| 3_hypothetical | I19 | Tb927.11.5180_for | GGGGACAAGTTTGTACAAAAAAGCAGGCTATGCAAAGGCACAGTTCGCA | Tb927.11.5180_rev | GGGGACCACTTTGTACAAGAAAGCTGGGTCTATAGATCACTAGCAGCTTCTTCAGC |
| 3_hypothetical | I21 | Tb927.10.7750_for | GGGGACAAGTTTGTACAAAAAAGCAGGCTATGAACAAGGAAAAGGTGGATCGC | Tb927.10.7750_rev | GGGGACCACTTTGTACAAGAAAGCTGGGTTCATTTCCCCTTTTGGAAGAACGATC |
| 3_hypothetical | I22 | Tb927.9.15000_for | GGGGACAAGTTTGTACAAAAAAGCAGGCTATGAAGGGCGTTGGCAGTAATAATT | Tb927.9.15000_rev | GGGGACCACTTTGTACAAGAAAGCTGGGTTCATTTCTTCACTTGCTTCAGTAATCCG |
| 3_hypothetical | I23 | Tb927.3.1080_for | GGGGACAAGTTTGTACAAAAAAGCAGGCTATGGCACAGGAGGGATTGAAAGA | Tb927.3.1080_rev | GGGGACCACTTTGTACAAGAAAGCTGGGTTCATTCTACCGTTGTAGTTTTCTGCTT |
| 3_hypothetical | I24 | Tb927.8.2040_for | GGGGACAAGTTTGTACAAAAAAGCAGGCTATGGCGGCTCTCCACGAG | Tb927.8.2040_rev | GGGGACCACTTTGTACAAGAAAGCTGGGTCTACTGAGGTCGGTTATAGTGGTGG |
| 3_hypothetical | J1 | Tb927.2.4850_for | GGGGACAAGTTTGTACAAAAAAGCAGGCTATGCTCGGCTCAGGAAAAAGC | Tb927.2.4850_rev | GGGGACCACTTTGTACAAGAAAGCTGGGTTTATTCGCATTCCTCCTGCTTACAG |
| 3_hypothetical | J2 | Tb927.10.15510_for | GGGGACAAGTTTGTACAAAAAAGCAGGCTATGAACTTGGGTCAAGTGCCCC | Tb927.10.15510_rev | GGGGACCACTTTGTACAAGAAAGCTGGGTTTAAACGCTGCTTGAACGCCA |
| 3_hypothetical | J3 | Tb927.1.3310_for | GGGGACAAGTTTGTACAAAAAAGCAGGCTATGGAAGAAGTACTCGATAACAAAGAGAC | Tb927.1.3310_rev | GGGGACCACTTTGTACAAGAAAGCTGGGTTTACGCCATTGGTGTGTTTTCCT |
| 3_hypothetical | J4 | Tb927.11.13480_for | GGGGACAAGTTTGTACAAAAAAGCAGGCTATGGACGAGAACACCACATCCT | Tb927.11.13480_rev | GGGGACCACTTTGTACAAGAAAGCTGGGTTCAAGGGGTCTCGTCATCGC |
| 3_hypothetical | J5 | Tb927.3.2480_for | GGGGACAAGTTTGTACAAAAAAGCAGGCTATGCTCCGGTGTATACAATGTAATGC | Tb927.3.2480_rev | GGGGACCACTTTGTACAAGAAAGCTGGGTTCACACCAGAAGTTGTGGACTCC |
| 3_hypothetical | J6 | Tb927.5.1620_for | GGGGACAAGTTTGTACAAAAAAGCAGGCTATGGAAAATAACGATACGACTCCATCC | Tb927.5.1620_rev | GGGGACCACTTTGTACAAGAAAGCTGGGTTTAGTCCACATGTAGATGAGGATCCAC |
| 3_hypothetical | J7 | Tb927.10.1240_for | GGGGACAAGTTTGTACAAAAAAGCAGGCTATGCGCATGTCGTTAACAACACG | Tb927.10.1240_rev | GGGGACCACTTTGTACAAGAAAGCTGGGTTTATTTCCGCAACCTCTTGGACTC |
| 3_hypothetical | J8 | Tb927.11.14770_for | GGGGACAAGTTTGTACAAAAAAGCAGGCTATGAGTACTAAATCCTCAATTTCACCTAGT | Tb927.11.14770_rev | GGGGACCACTTTGTACAAGAAAGCTGGGTTTACATAACACCACACGTTTGCGC |
| 3_hypothetical | J9 | Tb927.11.14010_for | GGGGACAAGTTTGTACAAAAAAGCAGGCTATGCTGCTGAAAAAGTTGCCGG | Tb927.11.14010_rev | GGGGACCACTTTGTACAAGAAAGCTGGGTTTAAACGGCACTCTTACGGGGG |
| 3_hypothetical | J10 | Tb927.9.3390_for | GGGGACAAGTTTGTACAAAAAAGCAGGCTATGCCGACCGGTGCCGCC | Tb927.9.3390_rev | GGGGACCACTTTGTACAAGAAAGCTGGGTCTATATGGGAACGAAGAGCTCCTTTG |
| 3_hypothetical | J11 | Tb927.4.920_for | GGGGACAAGTTTGTACAAAAAAGCAGGCTATGACCACAAGCGACGAATTACC | Tb927.4.920_rev | GGGGACCACTTTGTACAAGAAAGCTGGGTTCACTGCTCAAAGCGTACAGAATAG |
| 3_hypothetical | J12 | Tb927.8.6050_for | GGGGACAAGTTTGTACAAAAAAGCAGGCTATGCCTCCTAAGAGACCACCCC | Tb927.8.6050_rev | GGGGACCACTTTGTACAAGAAAGCTGGGTTCATTTCTTCGCCGGAAGGTTG |
| 3_hypothetical | J13 | Tb927.9.10520_for | GGGGACAAGTTTGTACAAAAAAGCAGGCTATGAAACGAACAGCTTTTCACTCCA | Tb927.9.10520_rev | GGGGACCACTTTGTACAAGAAAGCTGGGTTTAAATAAGTCGCTTTGACTTGAGATGCT |
| 3_hypothetical | J14 | Tb927.9.9080_for | GGGGACAAGTTTGTACAAAAAAGCAGGCTATGTCGCTACTCAACACTACTCTTCA | Tb927.9.9080_rev | GGGGACCACTTTGTACAAGAAAGCTGGGTTTAAGCATGAAAAATGTAGTGCATACCTCT |
| 3_hypothetical | J15 | Tb927.2.4650_for | GGGGACAAGTTTGTACAAAAAAGCAGGCTATGGCGATTCGGGCAAAGATG | Tb927.2.4650_rev | GGGGACCACTTTGTACAAGAAAGCTGGGTTCAACCACCCCATAAGGAAGATTCC |
| 3_hypothetical | J16 | Tb927.11.12590_for | GGGGACAAGTTTGTACAAAAAAGCAGGCTATGGCCCAAAGCTTTGATGACTG | Tb927.11.12590_rev | GGGGACCACTTTGTACAAGAAAGCTGGGTTTAAGTTGTGCCAAGCCGGTG |
| 3_hypothetical | J17 | Tb927.10.300_for | GGGGACAAGTTTGTACAAAAAAGCAGGCTATGGAGCACGTAGTAGCCAGG | Tb927.10.300_rev | GGGGACCACTTTGTACAAGAAAGCTGGGTTTAAGCTTTGTCTTTGGTATCGGAAGAA |
| 3_hypothetical | J18 | Tb927.6.3470_for | GGGGACAAGTTTGTACAAAAAAGCAGGCTATGTTTCTCAACAATAACATGGTTGACG | Tb927.6.3470_rev | GGGGACCACTTTGTACAAGAAAGCTGGGTCTAAACAGCGAGCCTCTGCG |
| 3_hypothetical | J19 | Tb927.5.1590_for | GGGGACAAGTTTGTACAAAAAAGCAGGCTATGGTGATAAAGAAGGATGTTCCTTTCAA | Tb927.5.1590_rev | GGGGACCACTTTGTACAAGAAAGCTGGGTTTAACTCTCATTGTCTCCTCCTGACA |
| 3_hypothetical | J20 | Tb927.5.3100_for | GGGGACAAGTTTGTACAAAAAAGCAGGCTATGATTAACAAAGAAAACAGTGGGCG | Tb927.5.3100_rev | GGGGACCACTTTGTACAAGAAAGCTGGGTTTATTCCCCCCTAATCTCAAAATCCGG |
| 3_hypothetical | J21 | Tb927.5.2990_for | GGGGACAAGTTTGTACAAAAAAGCAGGCTATGTCAAGGTCGCGAAATATCGTT | Tb927.5.2990_rev | GGGGACCACTTTGTACAAGAAAGCTGGGTTCATGCCCTCTGTCGCGC |
| 3_hypothetical | J22 | Tb927.9.8190_for | GGGGACAAGTTTGTACAAAAAAGCAGGCTATGTCACATCTCTTGCGCTCTCG | Tb927.9.8190_rev | GGGGACCACTTTGTACAAGAAAGCTGGGTCTACAGGCAATACTTCCGCTCAG |
| 3_hypothetical | J23 | Tb927.6.2820_for | GGGGACAAGTTTGTACAAAAAAGCAGGCTATGACTCACAGGACGGCGG | Tb927.6.2820_rev | GGGGACCACTTTGTACAAGAAAGCTGGGTTCATGCTAACTGCCCAGTTAGAAGG |
| 3_hypothetical | J24 | Tb927.7.6890_for | GGGGACAAGTTTGTACAAAAAAGCAGGCTATGTCCAAGTACGCTAACCCGG | Tb927.7.6890_rev | GGGGACCACTTTGTACAAGAAAGCTGGGTTCAGTCAAACCTTGCGGCATC |
| 3_hypothetical | K1 | Tb927.10.8030_for | GGGGACAAGTTTGTACAAAAAAGCAGGCTATGTTCCGCCGACTTTCTTCAAG | Tb927.10.8030_rev | GGGGACCACTTTGTACAAGAAAGCTGGGTTCAAACGCCATATTTCGCCTCC |
| 3_hypothetical | K2 | Tb927.3.4530_for | GGGGACAAGTTTGTACAAAAAAGCAGGCTATGTGTCGCGTCGTTGATGC | Tb927.3.4530_rev | GGGGACCACTTTGTACAAGAAAGCTGGGTTCAACCAGATATAACGACGGTAACGT |
| 3_hypothetical | K3 | Tb927.10.8630_for | GGGGACAAGTTTGTACAAAAAAGCAGGCTATGATGAATTCACATAACGCGATCGT | Tb927.10.8630_rev | GGGGACCACTTTGTACAAGAAAGCTGGGTCTACATCACACTGCAATTGAGGCC |
| 3_hypothetical | K4 | Tb927.5.480_for | GGGGACAAGTTTGTACAAAAAAGCAGGCTATGCATGCTGCTGGACAACG | Tb927.5.480_rev | GGGGACCACTTTGTACAAGAAAGCTGGGTTCACCTCGTAATACTCCACATGAGT |
| 3_hypothetical | K5 | Tb927.5.3680_for | GGGGACAAGTTTGTACAAAAAAGCAGGCTATGGCGCCGCCGATATCG | Tb927.5.3680_rev | GGGGACCACTTTGTACAAGAAAGCTGGGTTCAGTTGAGAACGTTTTGTGTAGCC |
| 3_hypothetical | K6 | Tb927.6.3700_for | GGGGACAAGTTTGTACAAAAAAGCAGGCTATGTTGGAGGGAAACGTGTTGG | Tb927.6.3700_rev | GGGGACCACTTTGTACAAGAAAGCTGGGTTCAGGGGAAAAGTTTCAAATTCCCAG |
| 3_hypothetical | K7 | Tb927.9.3490_for | GGGGACAAGTTTGTACAAAAAAGCAGGCTATGGGAAGTTGTTGTGCGAAGAG | Tb927.9.3490_rev | GGGGACCACTTTGTACAAGAAAGCTGGGTTCATTTACCACGGTTCCGAAGC |
| 3_hypothetical | K8 | Tb927.7.3870_for | GGGGACAAGTTTGTACAAAAAAGCAGGCTATGCTCATTAATTTTTTAGTAATCTTGATG | Tb927.7.3870_rev | GGGGACCACTTTGTACAAGAAAGCTGGGTTCAAAGCTCATCAAAGTCTACTTTCGC |
| 3_hypothetical | K9 | Tb927.1.890_for | GGGGACAAGTTTGTACAAAAAAGCAGGCTATGAATCGGACAGTTGCAATCTGT | Tb927.1.890_rev | GGGGACCACTTTGTACAAGAAAGCTGGGTCTACTTCCTTTCGCGTGTACGC |
| 3_hypothetical | K10 | Tb927.8.5190_for | GGGGACAAGTTTGTACAAAAAAGCAGGCTATGCCTGTCTGTGTTCTAGTGCC | Tb927.8.5190_rev | GGGGACCACTTTGTACAAGAAAGCTGGGTTTAGCCATTAACCGCAATACGTTCC |
| 3_hypothetical | K11 | Tb927.7.5810_for | GGGGACAAGTTTGTACAAAAAAGCAGGCTATGTTGGCGCTTCGTGCTG | Tb927.7.5810_rev | GGGGACCACTTTGTACAAGAAAGCTGGGTTTATGCTCCGTGCCGCAGG |
| 3_hypothetical | K12 | Tb927.11.6290_for | GGGGACAAGTTTGTACAAAAAAGCAGGCTATGGTCGGCCCCCGCATC | Tb927.11.6290_rev | GGGGACCACTTTGTACAAGAAAGCTGGGTTCACACTATGAACTTCATACACCTAGTTGC |
| 3_hypothetical | K13 | Tb927.9.9370_for | GGGGACAAGTTTGTACAAAAAAGCAGGCTATGGTCGCAGCAGCTGCT | Tb927.9.9370_rev | GGGGACCACTTTGTACAAGAAAGCTGGGTTCATGACCGAATTCGCGCC |
| 3_hypothetical | K14 | Tb927.8.1000_for | GGGGACAAGTTTGTACAAAAAAGCAGGCTATGAGGCAACGACAGTTGCAG | Tb927.8.1000_rev | GGGGACCACTTTGTACAAGAAAGCTGGGTCTACCGTCTCCCACGGTTTGG |
| 3_hypothetical | K15 | Tb927.11.4360_for | GGGGACAAGTTTGTACAAAAAAGCAGGCTATGCCGAGCGGAGCAAAG | Tb927.11.4360_rev | GGGGACCACTTTGTACAAGAAAGCTGGGTTTACTGACCACTATCCTCTTCTTCGC |
| 3_hypothetical | K16 | Tb927.8.5280_for | GGGGACAAGTTTGTACAAAAAAGCAGGCTATGTTACGGTGCGCACGC | Tb927.8.5280_rev | GGGGACCACTTTGTACAAGAAAGCTGGGTCTACTTTCCATCATAATGCACCGACC |
| 3_hypothetical | K17 | Tb927.10.12390_for | GGGGACAAGTTTGTACAAAAAAGCAGGCTATGGAAGAACGTCAGCGGC | Tb927.10.12390_rev | GGGGACCACTTTGTACAAGAAAGCTGGGTTTATACTAACGGCATAGCATCGTCCG |
| 3_hypothetical | K18 | Tb927.4.4920_for | GGGGACAAGTTTGTACAAAAAAGCAGGCTATGGATTCTTCATCAAGGAGTTTGAACC | Tb927.4.4920_rev | GGGGACCACTTTGTACAAGAAAGCTGGGTTCACCTAAAAGAACTAAACGGTTCAGC |
| 3_hypothetical | K19 | Tb927.11.2020_for | GGGGACAAGTTTGTACAAAAAAGCAGGCTATGACTAGTGAGTGCGTTACCGT | Tb927.11.2020_rev | GGGGACCACTTTGTACAAGAAAGCTGGGTCTACTGAAGATTACTGCGGTGGC |
| 3_hypothetical | K20 | Tb927.3.3710_for | GGGGACAAGTTTGTACAAAAAAGCAGGCTATGAAGCGGGCGGTTGCA | Tb927.3.3710_rev | GGGGACCACTTTGTACAAGAAAGCTGGGTCTACTGTTGCACTGGAGCCG |
| 3_hypothetical | K21 | Tb927.4.910_for | GGGGACAAGTTTGTACAAAAAAGCAGGCTATGTCAGAGGTGGATCAACAGCA | Tb927.4.910_rev | GGGGACCACTTTGTACAAGAAAGCTGGGTCTAATGGGGTGAAGCGCTGAC |
| 3_hypothetical | K22 | Tb927.6.1440_for | GGGGACAAGTTTGTACAAAAAAGCAGGCTATGAAGAGGTCCTTGATCGCGC | Tb927.6.1440_rev | GGGGACCACTTTGTACAAGAAAGCTGGGTTCAAGACGTCGACGCTGTTG |
| 3_hypothetical | K23 | Tb927.4.4040_for | GGGGACAAGTTTGTACAAAAAAGCAGGCTATGGATAATACGGAAGAAACGAGCAG | Tb927.4.4040_rev | GGGGACCACTTTGTACAAGAAAGCTGGGTTCAAACCCATCCGATTGCCCT |
| 3_hypothetical | K24 | Tb927.5.3200_for | GGGGACAAGTTTGTACAAAAAAGCAGGCTATGGCGACGAGGCCAAAC | Tb927.5.3200_rev | GGGGACCACTTTGTACAAGAAAGCTGGGTTCACTGTGCTGTCGGTGGG |
| 3_hypothetical | L1 | Tb927.11.4650_for | GGGGACAAGTTTGTACAAAAAAGCAGGCTATGCGCCTACATACGGTCCG | Tb927.11.4650_rev | GGGGACCACTTTGTACAAGAAAGCTGGGTCTACTCGTCCTTGATGGTGTTCTTTG |
| 3_hypothetical | L2 | Tb927.11.12760_for | GGGGACAAGTTTGTACAAAAAAGCAGGCTATGACAATTTTGTCTTTGCACGGC | Tb927.11.12760_rev | GGGGACCACTTTGTACAAGAAAGCTGGGTTCATTCCCTTACAGTCGGTTTAGTGG |
| 3_hypothetical | L3 | Tb927.4.4600_for | GGGGACAAGTTTGTACAAAAAAGCAGGCTATGAGCCGCAATGGCGAG | Tb927.4.4600_rev | GGGGACCACTTTGTACAAGAAAGCTGGGTCTAGGGGGGCTTTAATTGTTCCTTTC |
| 3_hypothetical | L4 | Tb927.10.10630_for | GGGGACAAGTTTGTACAAAAAAGCAGGCTATGCGCCCTAGGGATCTGTG | Tb927.10.10630_rev | GGGGACCACTTTGTACAAGAAAGCTGGGTCTATTCGTCATCACCACCGCTG |
| 3_hypothetical | L5 | Tb927.10.12080_for | GGGGACAAGTTTGTACAAAAAAGCAGGCTATGTCTGTTTCCCCCAAGCGG | Tb927.10.12080_rev | GGGGACCACTTTGTACAAGAAAGCTGGGTCTACCATCCGGCCATCGATGG |
| 3_hypothetical | L6 | Tb927.11.10970_for | GGGGACAAGTTTGTACAAAAAAGCAGGCTATGGAAAATGATGTGGATGAAGAGGTT | Tb927.11.10970_rev | GGGGACCACTTTGTACAAGAAAGCTGGGTTTACGGCATTGTCAAGAGAATATTGTGC |
| 3_hypothetical | L7 | Tb927.3.2080_for | GGGGACAAGTTTGTACAAAAAAGCAGGCTATGTCACTCTTCGCGAAGAAGAAG | Tb927.3.2080_rev | GGGGACCACTTTGTACAAGAAAGCTGGGTCTACAAACCGAGTTCAGTGGGATC |
| 3_hypothetical | L8 | Tb927.8.3390_for | GGGGACAAGTTTGTACAAAAAAGCAGGCTATGGTGAAAATGCCGTCGACC | Tb927.8.3390_rev | GGGGACCACTTTGTACAAGAAAGCTGGGTTTAACCGAGGTGCCCATCACC |
| 3_hypothetical | L9 | Tb927.11.13760_for | GGGGACAAGTTTGTACAAAAAAGCAGGCTATGATTTCACGGAGGAGCTTTGTAC | Tb927.11.13760_rev | GGGGACCACTTTGTACAAGAAAGCTGGGTTCAGCTTTGCTTTGCCTTTACGT |
| 3_hypothetical | L10 | Tb927.11.7600_for | GGGGACAAGTTTGTACAAAAAAGCAGGCTATGCTTGCCAACTACGACTATGAAA | Tb927.11.7600_rev | GGGGACCACTTTGTACAAGAAAGCTGGGTTTAGCCTCTATAAGAGCCCCGC |
| 3_hypothetical | L11 | Tb927.8.2400_for | GGGGACAAGTTTGTACAAAAAAGCAGGCTATGAATATCGTCACAAGCTACTTCCTG | Tb927.8.2400_rev | GGGGACCACTTTGTACAAGAAAGCTGGGTTTAATGCCGTTCCAACCTGTCG |
| 3_hypothetical | L12 | Tb927.6.4070_for | GGGGACAAGTTTGTACAAAAAAGCAGGCTATGACTGTATCAGGAATGACTGGGA | Tb927.6.4070_rev | GGGGACCACTTTGTACAAGAAAGCTGGGTTCAGTGTGCCGTTGCTATAGGG |
| 3_hypothetical | L13 | Tb927.10.1250_for | GGGGACAAGTTTGTACAAAAAAGCAGGCTATGACAAGTGTGGAAACCCGC | Tb927.10.1250_rev | GGGGACCACTTTGTACAAGAAAGCTGGGTTCAGTCCACCAACTGCCCC |
| 3_hypothetical | L14 | Tb927.10.13820_for | GGGGACAAGTTTGTACAAAAAAGCAGGCTATGTTCAACCGTAACTTTATTCTCCGA | Tb927.10.13820_rev | GGGGACCACTTTGTACAAGAAAGCTGGGTTTAGCAAGCCAACCTCTTCCTCC |
| 3_hypothetical | L15 | Tb927.7.6210_for | GGGGACAAGTTTGTACAAAAAAGCAGGCTATGGACACGTCGGGCACC | Tb927.7.6210_rev | GGGGACCACTTTGTACAAGAAAGCTGGGTCTACAAGTCGTAAAAGCGGCACA |
| 3_hypothetical | L16 | Tb927.11.8430_for | GGGGACAAGTTTGTACAAAAAAGCAGGCTATGTTTTGTGGCTCTAGGATGTTCC | Tb927.11.8430_rev | GGGGACCACTTTGTACAAGAAAGCTGGGTTTAAAGGTGATACTGCTCCACTTCATC |
| 3_hypothetical | L17 | Tb927.10.700_for | GGGGACAAGTTTGTACAAAAAAGCAGGCTATGCTACCGTGGCTATTATTTATCAACG | Tb927.10.700_rev | GGGGACCACTTTGTACAAGAAAGCTGGGTTCATGACACGACTTTACTAAATCCTTCTAT |
| 3_hypothetical | L18 | Tb927.8.1930_for | GGGGACAAGTTTGTACAAAAAAGCAGGCTATGCAAGACTACTTGCGGGAAATC | Tb927.8.1930_rev | GGGGACCACTTTGTACAAGAAAGCTGGGTCTAATTCTTGAGTGCATCAAGGCGC |
| 3_hypothetical | L19 | Tb927.8.8280_for | GGGGACAAGTTTGTACAAAAAAGCAGGCTATGGAATATGAGGGGTGTGCATCG | Tb927.8.8280_rev | GGGGACCACTTTGTACAAGAAAGCTGGGTTTAATTCCCCTTTTCGAAATTAAGTTCGC |
| 3_hypothetical | L20 | Tb927.11.3770_for | GGGGACAAGTTTGTACAAAAAAGCAGGCTATGATTCCTGCCCTTAATGGTGTTG | Tb927.11.3770_rev | GGGGACCACTTTGTACAAGAAAGCTGGGTCTACTGTTCTGTAACCTCAGTATCGC |
| 3_hypothetical | L21 | Tb927.11.10800_for | GGGGACAAGTTTGTACAAAAAAGCAGGCTATGGAGCCAAGTAAAACCAGCAC | Tb927.11.10800_rev | GGGGACCACTTTGTACAAGAAAGCTGGGTTCAAGAAGGTGATAGCGTAAACTGTTT |
| 3_hypothetical | L22 | Tb927.9.9430_for | GGGGACAAGTTTGTACAAAAAAGCAGGCTATGTCGGCAGCAGTTCCTGT | Tb927.9.9430_rev | GGGGACCACTTTGTACAAGAAAGCTGGGTCTAACGAGTAATCCTTTTTGCCAGCA |
| 3_hypothetical | L23 | Tb927.11.10700_for | GGGGACAAGTTTGTACAAAAAAGCAGGCTATGTATGCGACTCGTGCTAAGGT | Tb927.11.10700_rev | GGGGACCACTTTGTACAAGAAAGCTGGGTCTAACCCTGTTGTTTATTGTTCCAATCTAC |
| 3_hypothetical | L24 | Tb927.11.5530_for | GGGGACAAGTTTGTACAAAAAAGCAGGCTATGAAGCGACTATTCCCTTCTGCA | Tb927.11.5530_rev | GGGGACCACTTTGTACAAGAAAGCTGGGTTTACTCACAACGTTCCGCCCA |
| 3_hypothetical | M1 | Tb927.5.3410_for | GGGGACAAGTTTGTACAAAAAAGCAGGCTATGCGTCGCACGGTGGTG | Tb927.5.3410_rev | GGGGACCACTTTGTACAAGAAAGCTGGGTTTAGAAGAGCTTGATTTTTCTAGCCTTGC |
| 3_hypothetical | M2 | Tb927.8.3890_for | GGGGACAAGTTTGTACAAAAAAGCAGGCTATGTGCGCTGGGCGGGAT | Tb927.8.3890_rev | GGGGACCACTTTGTACAAGAAAGCTGGGTCTATTCCATTGTTTGCTCTCTTGGATATCC |
| 3_hypothetical | M3 | Tb927.5.3790_for | GGGGACAAGTTTGTACAAAAAAGCAGGCTATGAAAGTAGAATCACCTCATTCGAACTAT | Tb927.5.3790_rev | GGGGACCACTTTGTACAAGAAAGCTGGGTTCACGACTCTGCATGAGGATATGC |
| 3_hypothetical | M4 | Tb927.7.4260_for | GGGGACAAGTTTGTACAAAAAAGCAGGCTATGACAGTGAAGGTTTCATTTTCTCTTACA | Tb927.7.4260_rev | GGGGACCACTTTGTACAAGAAAGCTGGGTTTACTTCGGATTGTTGTCAGCACTATT |
| 3_hypothetical | M5 | Tb927.9.7800_for | GGGGACAAGTTTGTACAAAAAAGCAGGCTATGGGTAATCAAAAAGCTGTAGCTGG | Tb927.9.7800_rev | GGGGACCACTTTGTACAAGAAAGCTGGGTTCACGCTTTCGCAAGAAAATTCG |
| 3_hypothetical | M6 | Tb927.11.720_for | GGGGACAAGTTTGTACAAAAAAGCAGGCTATGGAACTGATGGAGGAGCGC | Tb927.11.720_rev | GGGGACCACTTTGTACAAGAAAGCTGGGTTTACCTGAATATAATAAGATTCCTTTCACT |
| 3_hypothetical | M7 | Tb927.10.1820_for | GGGGACAAGTTTGTACAAAAAAGCAGGCTATGATGAAGAGAACAATATTACAAAGATGC | Tb927.10.1820_rev | GGGGACCACTTTGTACAAGAAAGCTGGGTTCACCTGCCTCCCATCATAGTTT |
| 3_hypothetical | M8 | Tb927.7.1540_for | GGGGACAAGTTTGTACAAAAAAGCAGGCTATGCACGAGAAAGGATATGCCG | Tb927.7.1540_rev | GGGGACCACTTTGTACAAGAAAGCTGGGTCTATCTGCTAGCAAACAACCTGCG |
| 3_hypothetical | M9 | Tb927.5.3530_for | GGGGACAAGTTTGTACAAAAAAGCAGGCTATGGCCGCTGTCCCTTCA | Tb927.5.3530_rev | GGGGACCACTTTGTACAAGAAAGCTGGGTTTATCCGGCCCAATTATGGCAC |
| 3_hypothetical | M10 | Tb927.5.1990_for | GGGGACAAGTTTGTACAAAAAAGCAGGCTATGACGCAATCGCAATCGCA | Tb927.5.1990_rev | GGGGACCACTTTGTACAAGAAAGCTGGGTTTAGCGACTGAAGCGGGCG |
| 3_hypothetical | M11 | Tb927.8.4480_for | GGGGACAAGTTTGTACAAAAAAGCAGGCTATGACGGCGAGCCGTTATCA | Tb927.8.4480_rev | GGGGACCACTTTGTACAAGAAAGCTGGGTTTACACCACAGTAAAGAGCACATCC |
| 3_hypothetical | M12 | Tb927.10.1770_for | GGGGACAAGTTTGTACAAAAAAGCAGGCTATGAAAACGGAAAATTTTCCAAGAGGAC | Tb927.10.1770_rev | GGGGACCACTTTGTACAAGAAAGCTGGGTCTATGCTCCGGTGTTCTGTTTACC |
| 3_hypothetical | M13 | Tb927.8.6920_for | GGGGACAAGTTTGTACAAAAAAGCAGGCTATGACAAAAGGTGGGTCGAGTGT | Tb927.8.6920_rev | GGGGACCACTTTGTACAAGAAAGCTGGGTTTAATACTTCAACGGCTGATCAACAGG |
| 3_hypothetical | M14 | Tb927.3.3660_for | GGGGACAAGTTTGTACAAAAAAGCAGGCTATGCTGACTTTACGCAAGCAAAC | Tb927.3.3660_rev | GGGGACCACTTTGTACAAGAAAGCTGGGTCTACCAGTTTCCCGCCTCATGC |
| 3_hypothetical | M15 | Tb927.10.15820_for | GGGGACAAGTTTGTACAAAAAAGCAGGCTATGACAGCAGTACCTGTGGCA | Tb927.10.15820_rev | GGGGACCACTTTGTACAAGAAAGCTGGGTTCACTTCGGTACACCAATACCACA |
| 3_hypothetical | M16 | Tb927.10.8250_for | GGGGACAAGTTTGTACAAAAAAGCAGGCTATGCAGGGGAAGGTTGTCAGT | Tb927.10.8250_rev | GGGGACCACTTTGTACAAGAAAGCTGGGTTCATGGCTTTCGCTGTAACACG |
| 3_hypothetical | M17 | Tb927.7.6490_for | GGGGACAAGTTTGTACAAAAAAGCAGGCTATGAGCTCCACGAGGGCTG | Tb927.7.6490_rev | GGGGACCACTTTGTACAAGAAAGCTGGGTTTATTCGTCGTCGCCGGC |
| 3_hypothetical | M18 | Tb927.11.13860_for | GGGGACAAGTTTGTACAAAAAAGCAGGCTATGCATAACAGCAGGTCCACCA | Tb927.11.13860_rev | GGGGACCACTTTGTACAAGAAAGCTGGGTTTAAATGAAGAAAGATGATACGGCATCCC |
| 3_hypothetical | M19 | Tb927.8.7250_for | GGGGACAAGTTTGTACAAAAAAGCAGGCTATGGAGTTCGTTCCCCTCGAG | Tb927.8.7250_rev | GGGGACCACTTTGTACAAGAAAGCTGGGTTCACGGGCGATAATAATAATGTACATTCTT |
| 3_hypothetical | M20 | Tb927.4.5130_for | GGGGACAAGTTTGTACAAAAAAGCAGGCTATGCATGTCACGGGGTGC | Tb927.4.5130_rev | GGGGACCACTTTGTACAAGAAAGCTGGGTTCAAAAACAATTGCATGTCGCTTTCT |
| 3_hypothetical | M21 | Tb927.10.8790_for | GGGGACAAGTTTGTACAAAAAAGCAGGCTATGCTGCGCTTCCGGGTT | Tb927.10.8790_rev | GGGGACCACTTTGTACAAGAAAGCTGGGTTTACCGCAAACGCAACTTTTTAGC |

| | | | | | |
|---|---|---|---|---|---|
| 3_hypothetical | M22 | Tb927.4.5140_for | GGGGACAAGTTTGTACAAAAAAGCAGGCTATGGAGTTCGTTCCCCTCGAG | Tb927.4.5140_rev | GGGGACCACTTTGTACAAGAAAGCTGGGTTTACGCACGGCAATAATAATGTACATTC |
| 3_hypothetical | M23 | Tb927.4.2810_for | GGGGACAAGTTTGTACAAAAAAGCAGGCTATGCATCGTGTGATCACCATTTTATCT | Tb927.4.2810_rev | GGGGACCACTTTGTACAAGAAAGCTGGGTTCATTCGCTCATTAACCAGAATTCCA |
| 3_hypothetical | M24 | Tb927.11.13910_for | GGGGACAAGTTTGTACAAAAAAGCAGGCTATGTCCGTTGTTGGTGTCTCTTC | Tb927.11.13910_rev | GGGGACCACTTTGTACAAGAAAGCTGGGTTCACATTTCACACCGCCGG |
| 3_hypothetical | N1 | Tb927.10.12350_for | GGGGACAAGTTTGTACAAAAAAGCAGGCTATGCGCGTGGCGCTAACT | Tb927.10.12350_rev | GGGGACCACTTTGTACAAGAAAGCTGGGTTTATTTTCACTACTCTGTTGGAAAAACTT |
| 3_hypothetical | N2 | Tb927.10.9870_for | GGGGACAAGTTTGTACAAAAAAGCAGGCTATGGTTCTTATTGAGTGCAAGCGT | Tb927.10.9870_rev | GGGGACCACTTTGTACAAGAAAGCTGGGTTTACCCAGGTTTGTACTTAACTCCATCG |
| 3_hypothetical | N3 | Tb927.11.14390_for | GGGGACAAGTTTGTACAAAAAAGCAGGCTATGGCCACACTTCAAGTGCG | Tb927.11.14390_rev | GGGGACCACTTTGTACAAGAAAGCTGGGTTTACTTTACCACTTCAACGAGTGGAG |
| 3_hypothetical | N4 | Tb927.5.2590_for | GGGGACAAGTTTGTACAAAAAAGCAGGCTATGAAACGAGAGCGAGGTGGA | Tb927.5.2590_rev | GGGGACCACTTTGTACAAGAAAGCTGGGTTTAGAGACTGCTCCTCCCAACG |
| 3_hypothetical | N5 | Tb927.7.2190_for | GGGGACAAGTTTGTACAAAAAAGCAGGCTATGGCTTGCCGTCGTTCC | Tb927.7.2190_rev | GGGGACCACTTTGTACAAGAAAGCTGGGTTCATTTCTTCTTTTTCGGTGATTGGGA |
| 3_hypothetical | N6 | Tb927.9.10630_for | GGGGACAAGTTTGTACAAAAAAGCAGGCTATGTCGTGGCGTACGCGT | Tb927.9.10630_rev | GGGGACCACTTTGTACAAGAAAGCTGGGTTTACAGAATTCCTAACGCAGAACACC |
| 3_hypothetical | N7 | Tb927.6.1620_for | GGGGACAAGTTTGTACAAAAAAGCAGGCTATGGAATTCTTCCGGCGGACG | Tb927.6.1620_rev | GGGGACCACTTTGTACAAGAAAGCTGGGTTCACACGTGAGACACCCCAC |
| 3_hypothetical | N8 | Tb927.11.11320_for | GGGGACAAGTTTGTACAAAAAAGCAGGCTATGACGAGGAAAGTGGCCGG | Tb927.11.11320_rev | GGGGACCACTTTGTACAAGAAAGCTGGGTTCAACTGCGAGGGATTGTGATG |
| 3_hypothetical | N9 | Tb927.10.15830_for | GGGGACAAGTTTGTACAAAAAAGCAGGCTATGCGACGCTGCGCGGCA | Tb927.10.15830_rev | GGGGACCACTTTGTACAAGAAAGCTGGGTTCAGAAGTACCGTTTGGGCCG |
| 3_hypothetical | N10 | Tb927.11.2810_for | GGGGACAAGTTTGTACAAAAAAGCAGGCTATGGTGCTCCGCTGTGCT | Tb927.11.2810_rev | GGGGACCACTTTGTACAAGAAAGCTGGGTTTACCGTGAAGGGTAATGTTGCAA |
| 3_hypothetical | N11 | Tb927.9.3560_for | GGGGACAAGTTTGTACAAAAAAGCAGGCTATGGAGAGGAGTAATTCCCGGAGG | Tb927.9.3560_rev | GGGGACCACTTTGTACAAGAAAGCTGGGTCTACCGTCGAGCAGCATATGC |
| 3_hypothetical | N12 | Tb927.6.2140_for | GGGGACAAGTTTGTACAAAAAAGCAGGCTATGTTCCGCTGGTCGTGGT | Tb927.6.2140_rev | GGGGACCACTTTGTACAAGAAAGCTGGGTTCAGGTTAAGGACGCAGAAACAC |
| 3_hypothetical | N13 | Tb927.5.2730_for | GGGGACAAGTTTGTACAAAAAAGCAGGCTATGAGCAGCGATTCAAAAGCCA | Tb927.5.2730_rev | GGGGACCACTTTGTACAAGAAAGCTGGGTTTACGGCAACGTGTCGTCC |
| 3_hypothetical | N14 | Tb927.3.3040_for | GGGGACAAGTTTGTACAAAAAAGCAGGCTATGGAAACTACAACACACGTTGTCA | Tb927.3.3040_rev | GGGGACCACTTTGTACAAGAAAGCTGGGTTCATGTCGGTGGATCGACGT |
| 3_hypothetical | N15 | Tb927.8.6380_for | GGGGACAAGTTTGTACAAAAAAGCAGGCTATGATAGTGCGGTTGCTTTTCTTTTC | Tb927.8.6380_rev | GGGGACCACTTTGTACAAGAAAGCTGGGTCTACTTCATCTTTTTCCTCTTCTCCGC |
| 3_hypothetical | N16 | Tb927.7.6600_for | GGGGACAAGTTTGTACAAAAAAGCAGGCTATGGTTTTCGCAAGGGGCG | Tb927.7.6600_rev | GGGGACCACTTTGTACAAGAAAGCTGGGTCTAATCCACCTCTGCCTGGGC |
| 3_hypothetical | N17 | Tb927.9.10890_for | GGGGACAAGTTTGTACAAAAAAGCAGGCTATGGAAACATTCCGTGAGCGC | Tb927.9.10890_rev | GGGGACCACTTTGTACAAGAAAGCTGGGTTCAGTCGTCACTGTCCAACTGA |
| 3_hypothetical | N18 | Tb927.11.13890_for | GGGGACAAGTTTGTACAAAAAAGCAGGCTATGCTTAGGCGTGTCCCGA | Tb927.11.13890_rev | GGGGACCACTTTGTACAAGAAAGCTGGGTCTACGATAAGTGCACAGCGCC |
| 3_hypothetical | N19 | Tb927.8.8120_for | GGGGACAAGTTTGTACAAAAAAGCAGGCTATGCTATTCATCAGAATATTAAACCCATTA | Tb927.8.8120_rev | GGGGACCACTTTGTACAAGAAAGCTGGGTTCAGTCCTTCTTCACGTCACCT |
| 3_hypothetical | N20 | Tb927.10.12270_for | GGGGACAAGTTTGTACAAAAAAGCAGGCTATGAGGCGAGTGGTAGTGGATT | Tb927.10.12270_rev | GGGGACCACTTTGTACAAGAAAGCTGGGTTCAGACTGCCCGCCTGGC |
| 3_hypothetical | N21 | Tb927.11.15270_for | GGGGACAAGTTTGTACAAAAAAGCAGGCTATGGATCCGCTTCTGTTAGGCA | Tb927.11.15270_rev | GGGGACCACTTTGTACAAGAAAGCTGGGTTCAGGTACTCAACTCACATAACCGC |
| 3_hypothetical | N22 | Tb927.10.7030_for | GGGGACAAGTTTGTACAAAAAAGCAGGCTATGGATCCATCGAAGCGCATATG | Tb927.10.7030_rev | GGGGACCACTTTGTACAAGAAAGCTGGGTTTACTTCATTTTGGCGCGTAACGC |
| 3_hypothetical | N23 | Tb927.11.6020_for | GGGGACAAGTTTGTACAAAAAAGCAGGCTATGTTTGATGTGTCACCGCGT | Tb927.11.6020_rev | GGGGACCACTTTGTACAAGAAAGCTGGGTTTAGTACTTCCCATATCTCCTGTAGGTCG |
| 3_hypothetical | N24 | Tb927.11.10140_for | GGGGACAAGTTTGTACAAAAAAGCAGGCTATGGTGTCGTACGCCTCTGC | Tb927.11.10140_rev | GGGGACCACTTTGTACAAGAAAGCTGGGTCTACAATTTGGAGGGCTGCGT |
| 3_hypothetical | O1 | Tb927.10.5750_for | GGGGACAAGTTTGTACAAAAAAGCAGGCTATGACGGTGTACCCCTCACTATTT | Tb927.10.5750_rev | GGGGACCACTTTGTACAAGAAAGCTGGGTTCATTCAGTAAACAAGTAATTCCGTACATT |
| 3_hypothetical | O2 | Tb927.3.2670_for | GGGGACAAGTTTGTACAAAAAAGCAGGCTATGCAGAGACTCTTTTTTAGATCGACAT | Tb927.3.2670_rev | GGGGACCACTTTGTACAAGAAAGCTGGGTTTATTTCTGCACTGTGGGCGTC |
| 3_hypothetical | O3 | Tb927.11.1280_for | GGGGACAAGTTTGTACAAAAAAGCAGGCTATGAGGCCAACAGGAGAGTCTG | Tb927.11.1280_rev | GGGGACCACTTTGTACAAGAAAGCTGGGTTTATCTGCCCGTCCGGGC |
| 3_hypothetical | O4 | Tb927.10.8200_for | GGGGACAAGTTTGTACAAAAAAGCAGGCTATGGAGACCCCAAGAAGCAGAG | Tb927.10.8200_rev | GGGGACCACTTTGTACAAGAAAGCTGGGTCTACTCTTTCTTTCTTTTCTTCGTCTGACG |
| 3_hypothetical | O5 | Tb927.8.3740_for | GGGGACAAGTTTGTACAAAAAAGCAGGCTATGAATGTGCTAACATCTCACACTCAT | Tb927.8.3740_rev | GGGGACCACTTTGTACAAGAAAGCTGGGTTTACGGGGATGTGCAAGAAGAAGG |
| 3_hypothetical | O6 | Tb927.4.4700_for | GGGGACAAGTTTGTACAAAAAAGCAGGCTATGGCTCAGACGAGCAACAATG | Tb927.4.4700_rev | GGGGACCACTTTGTACAAGAAAGCTGGGTTCATACCCGCCCGGATGTC |
| 3_hypothetical | O7 | Tb927.4.1750_for | GGGGACAAGTTTGTACAAAAAAGCAGGCTATGTTCCGCGGGTCAGGG | Tb927.4.1750_rev | GGGGACCACTTTGTACAAGAAAGCTGGGTTCACCGCGATCCACTCGC |
| 3_hypothetical | O8 | Tb927.11.6250_for | GGGGACAAGTTTGTACAAAAAAGCAGGCTATGTCGAAGCAACTAACGTTTATATCAGC | Tb927.11.6250_rev | GGGGACCACTTTGTACAAGAAAGCTGGGTCTAAAGTGCAGAAACCACCGCC |
| 3_hypothetical | O9 | Tb927.2.1730_for | GGGGACAAGTTTGTACAAAAAAGCAGGCTATGAACACGGAGGACTCCTCC | Tb927.2.1730_rev | GGGGACCACTTTGTACAAGAAAGCTGGGTTCAGATAGCACAAAAATACAAATACGCAG |
| 3_hypothetical | O10 | Tb927.5.3640_for | GGGGACAAGTTTGTACAAAAAAGCAGGCTATGTTTAGCGAATCTCTCTGTATTCTCTCA | Tb927.5.3640_rev | GGGGACCACTTTGTACAAGAAAGCTGGGTTTAGAACCAGTGACCCCACATGA |
| 3_hypothetical | O11 | Tb927.11.13270_for | GGGGACAAGTTTGTACAAAAAAGCAGGCTATGAACCCCGGAAATGATTTGGATG | Tb927.11.13270_rev | GGGGACCACTTTGTACAAGAAAGCTGGGTTTAAATTTTGAAGGATGGTCTTACTCCCC |
| 3_hypothetical | O12 | Tb927.1.3040_for | GGGGACAAGTTTGTACAAAAAAGCAGGCTATGCGCACGTTCAGTCGTAC | Tb927.1.3040_rev | GGGGACCACTTTGTACAAGAAAGCTGGGTTCACGTGCCTTTGGGCGC |
| 3_hypothetical | O13 | Tb927.11.7830_for | GGGGACAAGTTTGTACAAAAAAGCAGGCTATGCGGGCACTTAGAGGGG | Tb927.11.7830_rev | GGGGACCACTTTGTACAAGAAAGCTGGGTCTACCGGCTCCTGCCATATGA |
| 3_hypothetical | O14 | Tb927.5.1050_for | GGGGACAAGTTTGTACAAAAAAGCAGGCTATGGCCAGTAGCACGGATTCA | Tb927.5.1050_rev | GGGGACCACTTTGTACAAGAAAGCTGGGTTCATTTTCCCCGATGTCTGTCCG |
| 3_hypothetical | O15 | Tb927.9.8490_for | GGGGACAAGTTTGTACAAAAAAGCAGGCTATGAGCACCCACACCAGCT | Tb927.9.8490_rev | GGGGACCACTTTGTACAAGAAAGCTGGGTTTATCCTCGGCGCCTCCG |
| 3_hypothetical | O16 | Tb927.1.1610_for | GGGGACAAGTTTGTACAAAAAAGCAGGCTATGCAGCGCGATGTGTACG | Tb927.1.1610_rev | GGGGACCACTTTGTACAAGAAAGCTGGGTTCAGTGAACGAATTGGTACTCGTC |
| 3_hypothetical | O17 | Tb927.7.4870_for | GGGGACAAGTTTGTACAAAAAAGCAGGCTATGGACGTTCTTATTGCAAAAGCAGT | Tb927.7.4870_rev | GGGGACCACTTTGTACAAGAAAGCTGGGTTCACATATTCATGGTGTTGGGTCCC |
| 3_hypothetical | O18 | Tb927.2.4090_for | GGGGACAAGTTTGTACAAAAAAGCAGGCTATGCTTTTAATTGCCGTCACAATCATT | Tb927.2.4090_rev | GGGGACCACTTTGTACAAGAAAGCTGGGTTTAACTCGTCGCGAACTCACG |
| 3_hypothetical | O19 | Tb927.8.5590_for | GGGGACAAGTTTGTACAAAAAAGCAGGCTATGGAAACGAGAAAAGTTACAGAGTGC | Tb927.8.5590_rev | GGGGACCACTTTGTACAAGAAAGCTGGGTTCACCTCGTAACCACACTTGATTG |
| 3_hypothetical | O20 | Tb927.11.14690_for | GGGGACAAGTTTGTACAAAAAAGCAGGCTATGGCCACTGAAGGAGAAGTGG | Tb927.11.14690_rev | GGGGACCACTTTGTACAAGAAAGCTGGGTTCAGGTAAGCAGCACCATCAGT |
| 3_hypothetical | O21 | Tb927.10.15230_for | GGGGACAAGTTTGTACAAAAAAGCAGGCTATGTCAGATTGGAAGGCCGCT | Tb927.10.15230_rev | GGGGACCACTTTGTACAAGAAAGCTGGGTCTATGGAATGGCCTCAACACTGTC |
| 3_hypothetical | O22 | Tb927.11.2770_for | GGGGACAAGTTTGTACAAAAAAGCAGGCTATGTTTTCTCGGCGCCGTAAG | Tb927.11.2770_rev | GGGGACCACTTTGTACAAGAAAGCTGGGTTCAGTCACCATAAAAAGTTGCTATAACTCT |
| 3_hypothetical | O23 | Tb927.7.5640_for | GGGGACAAGTTTGTACAAAAAAGCAGGCTATGAGCGAAGACCCAAACGGC | Tb927.7.5640_rev | GGGGACCACTTTGTACAAGAAAGCTGGGTTCACTCGATGAAAGGCGTATCGG |
| 3_hypothetical | O24 | Tb927.5.150_for | GGGGACAAGTTTGTACAAAAAAGCAGGCTATGCTTAAAATTGCAGCAGTCATTTTTTTG | Tb927.5.150_rev | GGGGACCACTTTGTACAAGAAAGCTGGGTTCAGGAGAACACAACACTTAATCCTGC |
| 3_hypothetical | P1 | Tb927.11.2800_for | GGGGACAAGTTTGTACAAAAAAGCAGGCTATGAGCCCCGTTAAAGCTTTCTG | Tb927.11.2800_rev | GGGGACCACTTTGTACAAGAAAGCTGGGTTCACGACTCCGCAGCATTCC |
| 3_hypothetical | P2 | Tb927.8.510_for | GGGGACAAGTTTGTACAAAAAAGCAGGCTATGAAAACCATAGTACTGGCGCTC | Tb927.8.510_rev | GGGGACCACTTTGTACAAGAAAGCTGGGTTTAAAAATAACATAAAATGCTTAGTAAAAA |
| 3_hypothetical | P3 | Tb927.7.2090_for | GGGGACAAGTTTGTACAAAAAAGCAGGCTATGGACAACGTTGTGAAGAATTTAATGC | Tb927.7.2090_rev | GGGGACCACTTTGTACAAGAAAGCTGGGTCTACTGCATTGCTGGAACTAAAACG |
| 3_hypothetical | P4 | Tb927.10.5540_for | GGGGACAAGTTTGTACAAAAAAGCAGGCTATGCTTAGCCGCTGTTTTGTCG | Tb927.10.5540_rev | GGGGACCACTTTGTACAAGAAAGCTGGGTTCACGAATCCAGTTCAAGTCGC |
| 3_hypothetical | P5 | Tb927.7.3540_for | GGGGACAAGTTTGTACAAAAAAGCAGGCTATGGCATGTGGTGACTTTGGC | Tb927.7.3540_rev | GGGGACCACTTTGTACAAGAAAGCTGGGTTCACACACTTTCCTCGTCAGACT |
| 3_hypothetical | P6 | Tb927.6.3240_for | GGGGACAAGTTTGTACAAAAAAGCAGGCTATGATGGGAGGAATCTCTATTTGTGTTG | Tb927.6.3240_rev | GGGGACCACTTTGTACAAGAAAGCTGGGTTCATATTCGTTTAATGGGGGTACTAATAGT |
| 3_hypothetical | P7 | Tb927.3.5600_for | GGGGACAAGTTTGTACAAAAAAGCAGGCTATGAAGGGTCTGCGGTTATTTTTTACA | Tb927.3.5600_rev | GGGGACCACTTTGTACAAGAAAGCTGGGTCTACACTCCTCTGCCTCCTGC |
| 3_hypothetical | P8 | Tb927.3.5430_for | GGGGACAAGTTTGTACAAAAAAGCAGGCTATGCCAAAGGTTCTCATCCCTATGT | Tb927.3.5430_rev | GGGGACCACTTTGTACAAGAAAGCTGGGTTCAGCCCCTTCGTCTTCTCG |
| 3_hypothetical | P9 | Tb927.11.13950_for | GGGGACAAGTTTGTACAAAAAAGCAGGCTATGCATTCGTCACCAGCGAC | Tb927.11.13950_rev | GGGGACCACTTTGTACAAGAAAGCTGGGTTCACCGAACCGACTGGGAC |
| 3_hypothetical | P10 | Tb927.1.5000_for | GGGGACAAGTTTGTACAAAAAAGCAGGCTATGACAACTGAAGAGGGAAAAGTAGAGA | Tb927.1.5000_rev | GGGGACCACTTTGTACAAGAAAGCTGGGTTTACCGCAGCCGAGGAACA |
| 3_hypothetical | P11 | Tb927.9.12680_for | GGGGACAAGTTTGTACAAAAAAGCAGGCTATGAGTAGTATCTTCACCTATGGTGCG | Tb927.9.12680_rev | GGGGACCACTTTGTACAAGAAAGCTGGGTTTAGTTTCTCCACTCATCGTGGTCG |
| 3_hypothetical | P12 | Tb927.4.5060_for | GGGGACAAGTTTGTACAAAAAAGCAGGCTATGGGTTCTGCTGCGGGA | Tb927.4.5060_rev | GGGGACCACTTTGTACAAGAAAGCTGGGTTCACGGTGTTGGTGTACAACG |
| 3_hypothetical | P13 | Tb927.4.2930_for | GGGGACAAGTTTGTACAAAAAAGCAGGCTATGCGCCGCTCTTCGATG | Tb927.4.2930_rev | GGGGACCACTTTGTACAAGAAAGCTGGGTTCACTTGTCGGGTCCCTTTCC |
| 3_hypothetical | P14 | Tb927.11.4920_for | GGGGACAAGTTTGTACAAAAAAGCAGGCTATGTCTACAATCTACAATGCAGTGGATG | Tb927.11.4920_rev | GGGGACCACTTTGTACAAGAAAGCTGGGTCTACTTGCTCGATGTTTCCCCAAA |
| 3_hypothetical | P15 | Tb927.11.4760_for | GGGGACAAGTTTGTACAAAAAAGCAGGCTATGTTTATCAGTGGGCTGTTCGTG | Tb927.11.4760_rev | GGGGACCACTTTGTACAAGAAAGCTGGGTTTACAAAACACGCGTAATGGTGGC |
| 3_hypothetical | P16 | Tb927.8.1570_for | GGGGACAAGTTTGTACAAAAAAGCAGGCTATGCCCGGGCAGGTTTCT | Tb927.8.1570_rev | GGGGACCACTTTGTACAAGAAAGCTGGGTTCAGGCGGAGACTGTTTGGT |
| 3_hypothetical | P17 | Tb927.5.4020_for | GGGGACAAGTTTGTACAAAAAAGCAGGCTATGATGATTTCCCGCGCTTTGT | Tb927.5.4020_rev | GGGGACCACTTTGTACAAGAAAGCTGGGTTTAGAAAGAGAGGGCCGTGGC |
| 3_hypothetical | P18 | Tb927.11.11390_for | GGGGACAAGTTTGTACAAAAAAGCAGGCTATGCAAACGTCGCCGGTC | Tb927.11.11390_rev | GGGGACCACTTTGTACAAGAAAGCTGGGTTCATGTGTGGCCCTTTCTTTCC |
| 3_hypothetical | P19 | Tb927.8.4530_for | GGGGACAAGTTTGTACAAAAAAGCAGGCTATGATGAATGACAAAATGCTTGGGGG | Tb927.8.4530_rev | GGGGACCACTTTGTACAAGAAAGCTGGGTCTATGGAGTATGCGCAAGAATCCTC |
| 3_hypothetical | P20 | Tb927.10.3290_for | GGGGACAAGTTTGTACAAAAAAGCAGGCTATGCGCAGCCTCCGCTCA | Tb927.10.3290_rev | GGGGACCACTTTGTACAAGAAAGCTGGGTTTAACGGCCTCTGGGGTAGC |
| 3_hypothetical | P21 | Tb927.8.6550_for | GGGGACAAGTTTGTACAAAAAAGCAGGCTATGTGGCGGCGGACGCCA | Tb927.8.6550_rev | GGGGACCACTTTGTACAAGAAAGCTGGGTTTAGCGGAAGTTGGGGTGGG |
| 3_hypothetical | P22 | Tb927.5.3570_for | GGGGACAAGTTTGTACAAAAAAGCAGGCTATGAATTCGTGTGATAACGTCGGC | Tb927.5.3570_rev | GGGGACCACTTTGTACAAGAAAGCTGGGTTCAACTTCTGTCAGACACCATAACCG |
| 3_hypothetical | P23 | Tb927.7.6700_for | GGGGACAAGTTTGTACAAAAAAGCAGGCTATGGAAGACATTCCACAAGATCCTGA | Tb927.7.6700_rev | GGGGACCACTTTGTACAAGAAAGCTGGGTTTACGTGATCTCGCAGTCACGC |
| 3_hypothetical | P24 | Tb927.11.5790_for | GGGGACAAGTTTGTACAAAAAAGCAGGCTATGGACTACCGTGGAAAGCACG | Tb927.11.5790_rev | GGGGACCACTTTGTACAAGAAAGCTGGGTTCAGGCGCTCACGTCTCTC |
| 3_known | A1 | Tb927.5.4430_for | GGGGACAAGTTTGTACAAAAAAGCAGGCTATGTCGCACGTTCCCTTTCG | Tb927.5.4430_rev | GGGGACCACTTTGTACAAGAAAGCTGGGTCTATCCCTCTTCACATTGGCTCGT |
| 3_known | A2 | Tb927.8.5950_for | GGGGACAAGTTTGTACAAAAAAGCAGGCTATGCCACCCAAACGACCCA | Tb927.8.5950_rev | GGGGACCACTTTGTACAAGAAAGCTGGGTTCAGGATACTGTGGTGAACCATCTT |
| 3_known | A3 | Tb927.7.1690_for | GGGGACAAGTTTGTACAAAAAAGCAGGCTATGACAGAACCGAAGTTAGCAACC | Tb927.7.1690_rev | GGGGACCACTTTGTACAAGAAAGCTGGGTTTAACCCTCAACGACTTTTTCCCTAC |
| 3_known | A4 | Tb927.6.4440_for | GGGGACAAGTTTGTACAAAAAAGCAGGCTATGCCAACGCCAACTGAGG | Tb927.6.4440_rev | GGGGACCACTTTGTACAAGAAAGCTGGGTTCAGCTGCGCTGCTTCTG |
| 3_known | A5 | Tb927.3.4910_for | GGGGACAAGTTTGTACAAAAAAGCAGGCTATGCCGAGTCCATCTGAAACAATAC | Tb927.3.4910_rev | GGGGACCACTTTGTACAAGAAAGCTGGGTCTAGTCATTCTTTTTTGTTCCCTCACATAC |
| 3_known | A6 | Tb927.10.13780_for | GGGGACAAGTTTGTACAAAAAAGCAGGCTATGTCGCTCAACCTTACCGATGC | Tb927.10.13780_rev | GGGGACCACTTTGTACAAGAAAGCTGGGTTTACTTCTTCAGCAGATACTCCCGC |
| 3_known | A7 | Tb927.6.1550_for | GGGGACAAGTTTGTACAAAAAAGCAGGCTATGGACATTGATCTTTCATATGGTGACC | Tb927.6.1550_rev | GGGGACCACTTTGTACAAGAAAGCTGGGTTCAGCGCGCTGCGACCCG |
| 3_known | A8 | Tb927.10.2440_for | GGGGACAAGTTTGTACAAAAAAGCAGGCTATGGGAGGCTGCGTCAGT | Tb927.10.2440_rev | GGGGACCACTTTGTACAAGAAAGCTGGGTCTACAGGAGCGTCCGTTCCT |
| 3_known | A9 | Tb927.1.4100_for | GGGGACAAGTTTGTACAAAAAAGCAGGCTATGTTTGCTCGCCGCTCAC | Tb927.1.4100_rev | GGGGACCACTTTGTACAAGAAAGCTGGGTCTAAATCTTGTTTGAGAGCTTCTTCCATTG |
| 3_known | A10 | Tb927.8.2650_for | GGGGACAAGTTTGTACAAAAAAGCAGGCTATGCGCCACTTCAGATTAAACCG | Tb927.8.2650_rev | GGGGACCACTTTGTACAAGAAAGCTGGGTTTAAAGATAGCCTTCATTAAAACCCACTGA |
| 3_known | A11 | Tb927.11.9470_for | GGGGACAAGTTTGTACAAAAAAGCAGGCTATGAGTCGTAATAGGGATGAGGTGC | Tb927.11.9470_rev | GGGGACCACTTTGTACAAGAAAGCTGGGTTTAGTGCTTCTTACCAAAAGGATTTAGAGA |
| 3_known | A12 | Tb927.8.3530_for | GGGGACAAGTTTGTACAAAAAAGCAGGCTATGGTCTCCGGTGTGACATATCTG | Tb927.8.3530_rev | GGGGACCACTTTGTACAAGAAAGCTGGGTTTACATTTTGGAAGTGCGCGGG |
| 3_known | A13 | Tb927.11.1600_for | GGGGACAAGTTTGTACAAAAAAGCAGGCTATGAGGGGATCTTCCGTTCCTTG | Tb927.11.1600_rev | GGGGACCACTTTGTACAAGAAAGCTGGGTTCAGATGTGAAACACCTCAACACATG |
| 3_known | A14 | Tb927.9.9780_for | GGGGACAAGTTTGTACAAAAAAGCAGGCTATGGGCAAGTTCTGCTGGC | Tb927.9.9780_rev | GGGGACCACTTTGTACAAGAAAGCTGGGTTCACGAAAACATCTTCTTCAAACCGG |
| 3_known | A15 | Tb927.10.10700_for | GGGGACAAGTTTGTACAAAAAAGCAGGCTATGAGCTATATCGAAACCGGATGGG | Tb927.10.10700_rev | GGGGACCACTTTGTACAAGAAAGCTGGGTTTATCTTAAATTCCGTCGCGGCC |
| 3_known | A16 | Tb927.11.3320_for | GGGGACAAGTTTGTACAAAAAAGCAGGCTATGCACGAACAAATCCAGAAGCT | Tb927.11.3320_rev | GGGGACCACTTTGTACAAGAAAGCTGGGTCTACAACACTTTACGCATTTCCTCAGC |
| 3_known | A17 | Tb927.9.9820_for | GGGGACAAGTTTGTACAAAAAAGCAGGCTATGGAGGCGGAAACGAGCA | Tb927.9.9820_rev | GGGGACCACTTTGTACAAGAAAGCTGGGTCTACACATTATTCTTCTTTGCATCCGC |
| 3_known | A18 | Tb927.9.6290_for | GGGGACAAGTTTGTACAAAAAAGCAGGCTATGGCTACCCGCGACGTT | Tb927.9.6290_rev | GGGGACCACTTTGTACAAGAAAGCTGGGTCTACTTCGACTTCTCCAGTTTGATGAG |
| 3_known | A19 | Tb927.9.9000_for | GGGGACAAGTTTGTACAAAAAAGCAGGCTATGTCCGAGTCAAGGAATGCAGA | Tb927.9.9000_rev | GGGGACCACTTTGTACAAGAAAGCTGGGTCTAGAGTGGTGCGCGCAAG |
| 3_known | A20 | Tb927.3.790_for | GGGGACAAGTTTGTACAAAAAAGCAGGCTATGGCCGAGCGGCGGCAG | Tb927.3.790_rev | GGGGACCACTTTGTACAAGAAAGCTGGGTCTAGCGCTTCCATAACCGCAG |
| 3_known | A21 | Tb927.11.3030_for | GGGGACAAGTTTGTACAAAAAAGCAGGCTATGCAGGGGCAAGCCGAC | Tb927.11.3030_rev | GGGGACCACTTTGTACAAGAAAGCTGGGTCTATGTGCGGAACAGAGAGGATAAC |
| 3_known | A22 | Tb927.9.3360_for | GGGGACAAGTTTGTACAAAAAAGCAGGCTATGAACCGAGCGGCTCGC | Tb927.9.3360_rev | GGGGACCACTTTGTACAAGAAAGCTGGGTTCAGCCGCTCCCAGCCAC |
| 3_known | A23 | Tb927.11.18280_for | GGGGACAAGTTTGTACAAAAAAGCAGGCTATGGTTTCACTAACAGCACTGTTTTTC | Tb927.11.18280_rev | GGGGACCACTTTGTACAAGAAAGCTGGGTTCAACTTCTGCGCTGTCGAATG |
| 3_known | A24 | Tb927.9.10080_for | GGGGACAAGTTTGTACAAAAAAGCAGGCTATGGCTTCGCTCACACGAC | Tb927.9.10080_rev | GGGGACCACTTTGTACAAGAAAGCTGGGTCTACTTGCCACTCTCCGACAATC |
| 3_known | B1 | Tb927.6.4650_for | GGGGACAAGTTTGTACAAAAAAGCAGGCTATGAGGATAATTCTCAAGAGCGTGC | Tb927.6.4650_rev | GGGGACCACTTTGTACAAGAAAGCTGGGTTCAGTCGGTAATCCCGAGAAAGT |
| 3_known | B2 | Tb927.7.3350_for | GGGGACAAGTTTGTACAAAAAAGCAGGCTATGTGCAGCTGTTGCTGTGAG | Tb927.7.3350_rev | GGGGACCACTTTGTACAAGAAAGCTGGGTTCAAGCAATAGGTTCGTTATCATGTGT |
| 3_known | B3 | Tb927.9.12300_for | GGGGACAAGTTTGTACAAAAAAGCAGGCTATGACGGGAGAAGGCACACT | Tb927.9.12300_rev | GGGGACCACTTTGTACAAGAAAGCTGGGTTCAGCACAGGCCACCTGC |
| 3_known | B4 | Tb927.1.1140_for | GGGGACAAGTTTGTACAAAAAAGCAGGCTATGGCGACTATTTACCACGAGGG | Tb927.1.1140_rev | GGGGACCACTTTGTACAAGAAAGCTGGGTCTAGCGCGGTGCATAACCAC |
| 3_known | B5 | Tb927.3.1030_for | GGGGACAAGTTTGTACAAAAAAGCAGGCTATGCCGGCAAAAAAGAAAGGAGG | Tb927.3.1030_rev | GGGGACCACTTTGTACAAGAAAGCTGGGTTCATCTCCGCAGTTCCTCCC |
| 3_known | B6 | Tb927.6.930_for | GGGGACAAGTTTGTACAAAAAAGCAGGCTATGGCCGTGGACCCAAGG | Tb927.6.930_rev | GGGGACCACTTTGTACAAGAAAGCTGGGTCTATTGGATAGATCTGTCAACAGAAAACAT |
| 3_known | B7 | Tb927.3.4340_for | GGGGACAAGTTTGTACAAAAAAGCAGGCTATGGAGGCGGAGTTGGGTC | Tb927.3.4340_rev | GGGGACCACTTTGTACAAGAAAGCTGGGTTTATAGGGATTTGTTTGTGTAGACAGCC |
| 3_known | B8 | Tb927.11.10010_for | GGGGACAAGTTTGTACAAAAAAGCAGGCTATGTTGGAGCGGGAGCGG | Tb927.11.10010_rev | GGGGACCACTTTGTACAAGAAAGCTGGGTTTAATGAGATGTTTCACCCGCCTC |
| 3_known | B9 | Tb927.7.4570_for | GGGGACAAGTTTGTACAAAAAAGCAGGCTATGGTTCATCGAAAACTTATCATCGACA | Tb927.7.4570_rev | GGGGACCACTTTGTACAAGAAAGCTGGGTTTACGCGACATATTCTACGATATCCCG |
| 3_known | B10 | Tb927.10.2550_for | GGGGACAAGTTTGTACAAAAAAGCAGGCTATGGTAATTTTCTTTGGTCGAGCACA | Tb927.10.2550_rev | GGGGACCACTTTGTACAAGAAAGCTGGGTTCATGAAACTGCATCTACCCCTGT |
| 3_known | B11 | Tb927.2.6000_for | GGGGACAAGTTTGTACAAAAAAGCAGGCTATGTTTGGTGGTGTAAAGTGGTCAC | Tb927.2.6000_rev | GGGGACCACTTTGTACAAGAAAGCTGGGTTTATGACCTTGCGGTTTGGTTGG |
| 3_known | B12 | Tb927.6.4670_for | GGGGACAAGTTTGTACAAAAAAGCAGGCTATGATATACTCTGGTGAGATAGAAAATGGC | Tb927.6.4670_rev | GGGGACCACTTTGTACAAGAAAGCTGGGTTTAATAATCCGGATTGCTGAGATCACAG |
| 3_known | B13 | Tb927.8.7530_for | GGGGACAAGTTTGTACAAAAAAGCAGGCTATGCGACGCAGTGTTGTGAC | Tb927.8.7530_rev | GGGGACCACTTTGTACAAGAAAGCTGGGTTTACTTTTTCGCTGCCACTCCC |
| 3_known | B14 | Tb927.6.4300_for | GGGGACAAGTTTGTACAAAAAAGCAGGCTATGACTATTAAAGTTGGCATCAATGGTTTC | Tb927.6.4300_rev | GGGGACCACTTTGTACAAGAAAGCTGGGTTTATAGCTTTGCTGCACGGTCC |
| 3_known | B15 | Tb927.10.13900_for | GGGGACAAGTTTGTACAAAAAAGCAGGCTATGCCTTCCGTTAAATGGCTTTCG | Tb927.10.13900_rev | GGGGACCACTTTGTACAAGAAAGCTGGGTCTAATGATGTTCCTCTCCTTTCGAAGTT |
| 3_known | B16 | Tb927.9.5890_for | GGGGACAAGTTTGTACAAAAAAGCAGGCTATGCACCGTGCTAATATTATATTACGCA | Tb927.9.5890_rev | GGGGACCACTTTGTACAAGAAAGCTGGGTTCACAATTCCCGTGTCAGGAGG |
| 3_known | B17 | Tb927.6.2720_for | GGGGACAAGTTTGTACAAAAAAGCAGGCTATGCCGCACTCCATGGACA | Tb927.6.2720_rev | GGGGACCACTTTGTACAAGAAAGCTGGGTTTACAAAAGTTTCCATTGATTATTGAGAAC |
| 3_known | B18 | Tb927.10.6950_for | GGGGACAAGTTTGTACAAAAAAGCAGGCTATGTCGGCCGGATCTCGT | Tb927.10.6950_rev | GGGGACCACTTTGTACAAGAAAGCTGGGTTTAGCACGACAGCTCTTCCCC |
| 3_known | B21 | Tb927.11.11210_for | GGGGACAAGTTTGTACAAAAAAGCAGGCTATGAAGACTAAGTTCTATACATATAAGAAC | Tb927.11.11210_rev | GGGGACCACTTTGTACAAGAAAGCTGGGTTCACCTTCTGTTTCTGCGCTC |
| 3_known | B22 | Tb927.9.14400_for | GGGGACAAGTTTGTACAAAAAAGCAGGCTATGGAACTGACACTGACTAGACGG | Tb927.9.14400_rev | GGGGACCACTTTGTACAAGAAAGCTGGGTTTAAGCAAAACCAGTGGACTGGTT |
| 3_known | B23 | Tb927.11.10230_for | GGGGACAAGTTTGTACAAAAAAGCAGGCTATGAGTCTTCGCGAGGACTGG | Tb927.11.10230_rev | GGGGACCACTTTGTACAAGAAAGCTGGGTCTACGGCTGGTCCGGCGG |
| 3_known | B24 | Tb927.3.3610_for | GGGGACAAGTTTGTACAAAAAAGCAGGCTATGCCTTCATTTGTTCTCCACCC | Tb927.3.3610_rev | GGGGACCACTTTGTACAAGAAAGCTGGGTCTACCTGACTGGGGGCACG |
| 3_known | C2 | Tb927.11.14260_for | GGGGACAAGTTTGTACAAAAAAGCAGGCTATGTATCAAGGACGTAGTCAACAGC | Tb927.11.14260_rev | GGGGACCACTTTGTACAAGAAAGCTGGGTCTAGTCAAGGTCCTGGACAATTGAC |
| 3_known | C3 | Tb927.11.11660_for | GGGGACAAGTTTGTACAAAAAAGCAGGCTATGCTAGCACCTAGAGGCTCTG | Tb927.11.11660_rev | GGGGACCACTTTGTACAAGAAAGCTGGGTTTAGATTGTACCAGTGTTCACACCAC |
| 3_known | C4 | Tb927.11.16480_for | GGGGACAAGTTTGTACAAAAAAGCAGGCTATGCGTTGCACGCTTCCG | Tb927.11.16480_rev | GGGGACCACTTTGTACAAGAAAGCTGGGTCTAGTACGGCTCTGTGGGATGC |
| 3_known | C5 | Tb927.3.2150_for | GGGGACAAGTTTGTACAAAAAAGCAGGCTATGGGAGGACAAAGCAGTGTAACC | Tb927.3.2150_rev | GGGGACCACTTTGTACAAGAAAGCTGGGTTTACCATGTGCTAGTAGCGTTAAGTTC |
| 3_known | C6 | Tb927.10.15490_for | GGGGACAAGTTTGTACAAAAAAGCAGGCTATGTCAGCACGTCGACTGGC | Tb927.10.15490_rev | GGGGACCACTTTGTACAAGAAAGCTGGGTTCAGCTGGTTTCGGCGGT |
| 3_known | C7 | Tb927.9.14410_for | GGGGACAAGTTTGTACAAAAAAGCAGGCTATGCTAACGTTCGAGGGAAGCA | Tb927.9.14410_rev | GGGGACCACTTTGTACAAGAAAGCTGGGTCTATCCACTCTTCTTCCAGACATTCAC |
| 3_known | C8 | Tb11.02.5400_for | GGGGACAAGTTTGTACAAAAAAGCAGGCTATGAAGGACCAACGAACCACTATTC | Tb11.02.5400_rev | GGGGACCACTTTGTACAAGAAAGCTGGGTTCACTTCCCAATTAAACCGAGTTTGG |
| 3_known | C9 | Tb927.9.10370_for | GGGGACAAGTTTGTACAAAAAAGCAGGCTATGCAAGGCGCCAATTTAAAAGTG | Tb927.9.10370_rev | GGGGACCACTTTGTACAAGAAAGCTGGGTCTACTGCTCCGCGGCAAC |
| 3_known | C10 | Tb927.9.12670_for | GGGGACAAGTTTGTACAAAAAAGCAGGCTATGGTAGCAACTATTACCTTCAAGTTGC | Tb927.9.12670_rev | GGGGACCACTTTGTACAAGAAAGCTGGGTTCAGTCCCGCGCACGCTT |

| | | | | | |
|---|---|---|---|---|---|
| 3_known | C11 | Tb927.6.670_for | GGGGACAAGTTTGTACAAAAAAGCAGGCTATGCAGTACCAACGAGCAGC | Tb927.6.670_rev | GGGGACCACTTTGTACAAGAAAGCTGGGTTCACGTTTCTGTTTTTATCTTCTTCACAC |
| 3_known | C12 | Tb927.9.9320_for | GGGGACAAGTTTGTACAAAAAAGCAGGCTATGCAAAAGTACGCAATATTAGGAAAGAAA | Tb927.9.9320_rev | GGGGACCACTTTGTACAAGAAAGCTGGGTTCACCCTTGAAGTTTGGGCAGA |
| 3_known | C13 | Tb927.9.6350_for | GGGGACAAGTTTGTACAAAAAAGCAGGCTATGGCTGAGTATGACATTAATTTGGAGG | Tb927.9.6350_rev | GGGGACCACTTTGTACAAGAAAGCTGGGTCTACAGCCAAAATCTTCCATAGCTCA |
| 3_known | C14 | Tb927.5.810_for | GGGGACAAGTTTGTACAAAAAAGCAGGCTATGAGCACTGCAACATCTGCA | Tb927.5.810_rev | GGGGACCACTTTGTACAAGAAAGCTGGGTTCACAAGGAAAGAAACATATGCAGACC |
| 3_known | C15 | Tb927.3.3380_for | GGGGACAAGTTTGTACAAAAAAGCAGGCTATGGTCGCAGAGTGCGCT | Tb927.3.3380_rev | GGGGACCACTTTGTACAAGAAAGCTGGGTCTATGGTATAAGTCTCTGCCACTGCT |
| 3_known | C16 | Tb927.11.15280_for | GGGGACAAGTTTGTACAAAAAAGCAGGCTATGGAGGAGGTGGTGGTACCG | Tb927.11.15280_rev | GGGGACCACTTTGTACAAGAAAGCTGGGTTCAAGGCACCGTAAGGTCCTC |
| 3_known | C17 | Tb927.9.4940_for | GGGGACAAGTTTGTACAAAAAAGCAGGCTATGAGGCGGAGCAGGGTAC | Tb927.9.4940_rev | GGGGACCACTTTGTACAAGAAAGCTGGGTCTACAGTGCAAGCACAGAGAGAG |
| 3_known | C18 | Tb927.10.16030_for | GGGGACAAGTTTGTACAAAAAAGCAGGCTATGGACATACCACAAGAACTGGCT | Tb927.10.16030_rev | GGGGACCACTTTGTACAAGAAAGCTGGGTCTAGTCACCCTTTGGACCGGG |
| 3_known | C19 | Tb927.10.6870_for | GGGGACAAGTTTGTACAAAAAAGCAGGCTATGATATTTTTACTGTTCATAAGCTTTATT | Tb927.10.6870_rev | GGGGACCACTTTGTACAAGAAAGCTGGGTTCATTTCTCCGAACTACTCCTGCT |
| 3_known | C20 | Tb927.10.240_for | GGGGACAAGTTTGTACAAAAAAGCAGGCTATGTCTTTGCTGCTGTCGGG | Tb927.10.240_rev | GGGGACCACTTTGTACAAGAAAGCTGGGTTCAAGCTGCCTCGCCGCC |
| 3_known | C21 | Tb927.8.750_for | GGGGACAAGTTTGTACAAAAAAGCAGGCTATGGAGGGTTTCTACGGTGTTGA | Tb927.8.750_rev | GGGGACCACTTTGTACAAGAAAGCTGGGTTTAGTAACGACCCCTCCCAGAGT |
| 3_known | C22 | Tb927.6.4630_for | GGGGACAAGTTTGTACAAAAAAGCAGGCTATGCCGAACGCTGATGCTC | Tb927.6.4630_rev | GGGGACCACTTTGTACAAGAAAGCTGGGTCTACACAGGGGAAAAAGTCCCATATC |
| 3_known | C23 | Tb927.6.3540_for | GGGGACAAGTTTGTACAAAAAAGCAGGCTATGTCGCTGCCAGCTCCT | Tb927.6.3540_rev | GGGGACCACTTTGTACAAGAAAGCTGGGTCTAATTCACTCTACCCTCACCATCATAAC |
| 3_known | C24 | Tb927.10.15180_for | GGGGACAAGTTTGTACAAAAAAGCAGGCTATGCCGCCGAAGCGAAAA | Tb927.10.15180_rev | GGGGACCACTTTGTACAAGAAAGCTGGGTTTACTGGTGTTTGCATTGTTGCTG |
| 3_known | D1 | Tb927.11.16000_for | GGGGACAAGTTTGTACAAAAAAGCAGGCTATGAGTGCCAATAAACACGGACAT | Tb927.11.16000_rev | GGGGACCACTTTGTACAAGAAAGCTGGGTCTATACAGTTTCCGTATTATCGGGATCCC |
| 3_known | D2 | Tb927.9.8210_for | GGGGACAAGTTTGTACAAAAAAGCAGGCTATGCGTATACATCATGCATTCTTTAGGC | Tb927.9.8210_rev | GGGGACCACTTTGTACAAGAAAGCTGGGTCTACTGAGGGGCGTTATCCTTCA |
| 3_known | D3 | Tb927.11.9430_for | GGGGACAAGTTTGTACAAAAAAGCAGGCTATGGAGGTTTTGTGTAATGGTAGACAAT | Tb927.11.9430_rev | GGGGACCACTTTGTACAAGAAAGCTGGGTCTACTTGGCAGGCGTAGCTCC |
| 3_known | D4 | Tb927.10.13540_for | GGGGACAAGTTTGTACAAAAAAGCAGGCTATGAATTTTTCACCATTGGCGAATCC | Tb927.10.13540_rev | GGGGACCACTTTGTACAAGAAAGCTGGGTCTACTCGCCCGATCCGCT |
| 3_known | D5 | Tb927.2.4590_for | GGGGACAAGTTTGTACAAAAAAGCAGGCTATGACTTTCCTCGCCAAGGACC | Tb927.2.4590_rev | GGGGACCACTTTGTACAAGAAAGCTGGGTTTAAGAAGTGGTACCAACAATGCGT |
| 3_known | D6 | Tb927.2.4610_for | GGGGACAAGTTTGTACAAAAAAGCAGGCTATGACTTTCCTCGCCAAAAAAATTTTTGT | Tb927.2.4610_rev | GGGGACCACTTTGTACAAGAAAGCTGGGTTTAAGAAGTGGTACCAACAATGCGT |
| 3_known | D7 | Tb927.7.3360_for | GGGGACAAGTTTGTACAAAAAAGCAGGCTATGGTAATGCAAATGTTTATGCAGGTATAT | Tb927.7.3360_rev | GGGGACCACTTTGTACAAGAAAGCTGGGTTCACTTCTGGCGCTTGTAAGTC |
| 3_known | D8 | Tb927.8.4440_for | GGGGACAAGTTTGTACAAAAAAGCAGGCTATGTCGGATGTACCGTGGGC | Tb927.8.4440_rev | GGGGACCACTTTGTACAAGAAAGCTGGGTTCAACCACTTTGGTGTGCCC |
| 3_known | D9 | Tb927.2.4240_for | GGGGACAAGTTTGTACAAAAAAGCAGGCTATGCACACTTCGTCATTGCAACT | Tb927.2.4240_rev | GGGGACCACTTTGTACAAGAAAGCTGGGTTTATCGCGAGGCTCCATGCT |
| 3_known | D10 | Tb927.7.300_for | GGGGACAAGTTTGTACAAAAAAGCAGGCTATGGTGTGGAGTGGGCATAAATACA | Tb927.7.300_rev | GGGGACCACTTTGTACAAGAAAGCTGGGTTCATGTGCACGAGGCGTG |
| 3_known | D11 | Tb927.10.6800_for | GGGGACAAGTTTGTACAAAAAAGCAGGCTATGTCCAATTTGCAGAAGATAAATGACATC | Tb927.10.6800_rev | GGGGACCACTTTGTACAAGAAAGCTGGGTCTAAACCTTCTTCACAACCTGCACC |
| 3_known | D12 | Tb927.5.3960_for | GGGGACAAGTTTGTACAAAAAAGCAGGCTATGGAGTCCGGGGGGTTTG | Tb927.5.3960_rev | GGGGACCACTTTGTACAAGAAAGCTGGGTTTATTTTAACTCGAGCTCAATGGTGTAGAA |
| 3_known | D13 | Tb927.8.5690_for | GGGGACAAGTTTGTACAAAAAAGCAGGCTATGCACCGCGGACCGCCG | Tb927.8.5690_rev | GGGGACCACTTTGTACAAGAAAGCTGGGTCTACAGCAAATTCATTGCTGCAAGT |
| 3_known | D14 | Tb927.8.7440_for | GGGGACAAGTTTGTACAAAAAAGCAGGCTATGAGTGAGCGGAAAGGCAAC | Tb927.8.7440_rev | GGGGACCACTTTGTACAAGAAAGCTGGGTTCAGTCCGCAGGCTTGTCG |
| 3_known | D15 | Tb927.10.4330_for | GGGGACAAGTTTGTACAAAAAAGCAGGCTATGAGGCGGTGGGCCAGT | Tb927.10.4330_rev | GGGGACCACTTTGTACAAGAAAGCTGGGTCTAGAACGCGATCACTTCCTTCAC |
| 3_known | D16 | Tb927.11.11110_for | GGGGACAAGTTTGTACAAAAAAGCAGGCTATGCACCGGGGGAGCCAA | Tb927.11.11110_rev | GGGGACCACTTTGTACAAGAAAGCTGGGTTCACTCTAGGGACGAAATCGGG |
| 3_known | D17 | Tb927.1.1120_for | GGGGACAAGTTTGTACAAAAAAGCAGGCTATGTCTCGGGGTGGAATATTTGGT | Tb927.1.1120_rev | GGGGACCACTTTGTACAAGAAAGCTGGGTTCACCGCTTTTTGTACAGTGACG |
| 3_known | D18 | Tb927.11.5010_for | GGGGACAAGTTTGTACAAAAAAGCAGGCTATGTCGGATCTCCGTCGAGTT | Tb927.11.5010_rev | GGGGACCACTTTGTACAAGAAAGCTGGGTTTACACCGTCCAGTGCTGCA |
| 3_known | D19 | Tb927.9.2670_for | GGGGACAAGTTTGTACAAAAAAGCAGGCTATGGGTTCCAGTATGTCTTGTCTGG | Tb927.9.2670_rev | GGGGACCACTTTGTACAAGAAAGCTGGGTCTACTCGGGGGAGAAAAAGCTCT |
| 3_known | D20 | Tb927.8.2050_for | GGGGACAAGTTTGTACAAAAAAGCAGGCTATGAGGGCCGTCATTCTTGTTG | Tb927.8.2050_rev | GGGGACCACTTTGTACAAGAAAGCTGGGTTTACATGACCACTTCTGGTTCGGG |
| 3_known | D21 | Tb927.11.14610_for | GGGGACAAGTTTGTACAAAAAAGCAGGCTATGAGGTTCATGCTAAGGTTAGTTATGTTG | Tb927.11.14610_rev | GGGGACCACTTTGTACAAGAAAGCTGGGTTTACCACAATACCACAGAGAATACAAAAAA |
| 3_known | D22 | Tb927.6.4460_for | GGGGACAAGTTTGTACAAAAAAGCAGGCTATGTCCTCTTGCAAGGACTCTCT | Tb927.6.4460_rev | GGGGACCACTTTGTACAAGAAAGCTGGGTTCATTCCTTCGCTCCAGAAGCT |
| 3_known | D23 | Tb927.9.6230_for | GGGGACAAGTTTGTACAAAAAAGCAGGCTATGGCTACCCGCGACGTT | Tb927.9.6230_rev | GGGGACCACTTTGTACAAGAAAGCTGGGTCTACAAGTTCGAACCAACAAATGGTTT |
| 3_known | D24 | Tb927.9.10970_for | GGGGACAAGTTTGTACAAAAAAGCAGGCTATGAGCGCTGAGCGGGTT | Tb927.9.10970_rev | GGGGACCACTTTGTACAAGAAAGCTGGGTCTAAATTAGCTGTTCTCTATGGCGAAGC |
| 3_known | E1 | Tb927.7.270_for | GGGGACAAGTTTGTACAAAAAAGCAGGCTATGTCCTCTATCGGTGGCTTCAA | Tb927.7.270_rev | GGGGACCACTTTGTACAAGAAAGCTGGGTTCAAATATCCCTATCGGGGTTATGTCA |
| 3_known | E3 | Tb927.10.7940_for | GGGGACAAGTTTGTACAAAAAAGCAGGCTATGCCTGCCGTTGCAGAC | Tb927.10.7940_rev | GGGGACCACTTTGTACAAGAAAGCTGGGTCTATGACTTGACCTCCTCCATTGC |
| 3_known | E4 | Tb927.10.10230_for | GGGGACAAGTTTGTACAAAAAAGCAGGCTATGCGAAGAACAGTGGCATTTTTAGT | Tb927.10.10230_rev | GGGGACCACTTTGTACAAGAAAGCTGGGTTTATACCCCACAATCCTTCAACTCTTCC |
| 3_known | E5 | Tb927.11.14880_for | GGGGACAAGTTTGTACAAAAAAGCAGGCTATGTCTGCTGACCCCACATCT | Tb927.11.14880_rev | GGGGACCACTTTGTACAAGAAAGCTGGGTCTACTCCTGAGCCTCCGCAT |
| 3_known | E6 | Tb927.8.3550_for | GGGGACAAGTTTGTACAAAAAAGCAGGCTATGCACAAAAGTAATCAAGAGTTGTGC | Tb927.8.3550_rev | GGGGACCACTTTGTACAAGAAAGCTGGGTCTACGCGCCTTTCGGTATTTGG |
| 3_known | E7 | Tb927.5.1310_for | GGGGACAAGTTTGTACAAAAAAGCAGGCTATGATGCGCCGCGTACTC | Tb927.5.1310_rev | GGGGACCACTTTGTACAAGAAAGCTGGGTCTATAATGCAACAGCAGTGAAGTGC |
| 3_known | E8 | Tb927.11.1010_for | GGGGACAAGTTTGTACAAAAAAGCAGGCTATGCGCCGCTTTGTTTCCC | Tb927.11.1010_rev | GGGGACCACTTTGTACAAGAAAGCTGGGTTCAAAAAAGGCGCATCCGCT |
| 3_known | E9 | Tb927.10.12290_for | GGGGACAAGTTTGTACAAAAAAGCAGGCTATGGTTGGACAAATTTTGAGTAGGAGG | Tb927.10.12290_rev | GGGGACCACTTTGTACAAGAAAGCTGGGTTCACCGCTTGCCGCATGT |
| 3_known | E10 | Tb927.4.980_for | GGGGACAAGTTTGTACAAAAAAGCAGGCTATGGCAGTAGTGGAGATTGGATGG | Tb927.4.980_rev | GGGGACCACTTTGTACAAGAAAGCTGGGTTCAGTAAAATAACTTGCGGTACAACACC |
| 3_known | E11 | Tb927.5.3910_for | GGGGACAAGTTTGTACAAAAAAGCAGGCTATGCGTAGGAACAATCGCACAC | Tb927.5.3910_rev | GGGGACCACTTTGTACAAGAAAGCTGGGTTCAAACTTCTTCATCCAGTTTATGTTTCTT |
| 3_known | E12 | Tb927.10.5620_for | GGGGACAAGTTTGTACAAAAAAGCAGGCTATGTCCAAGCGTGTTGAAGTTCT | Tb927.10.5620_rev | GGGGACCACTTTGTACAAGAAAGCTGGGTTTAGTATGTGTTGCCAGCAACGT |
| 3_known | E13 | Tb927.11.3220_for | GGGGACAAGTTTGTACAAAAAAGCAGGCTATGTCGTCTGAAAAAGTCGCTGG | Tb927.11.3220_rev | GGGGACCACTTTGTACAAGAAAGCTGGGTTTATTTTCCACCAAGAGGGGTGGT |
| 3_known | E14 | Tb927.7.6470_for | GGGGACAAGTTTGTACAAAAAAGCAGGCTATGAGCGCCAACACTCAGC | Tb927.7.6470_rev | GGGGACCACTTTGTACAAGAAAGCTGGGTTTAAGTGCGCCACACTACGTTG |
| 3_known | E15 | Tb927.2.2430_for | GGGGACAAGTTTGTACAAAAAAGCAGGCTATGTCCCGCAGCGGTGGT | Tb927.2.2430_rev | GGGGACCACTTTGTACAAGAAAGCTGGGTCTAACCTCTGCGCCTCGGG |
| 3_known | E16 | Tb927.1.3010_for | GGGGACAAGTTTGTACAAAAAAGCAGGCTATGCGGACGTTCCTGGCA | Tb927.1.3010_rev | GGGGACCACTTTGTACAAGAAAGCTGGGTTTACAGCGGAACAAACTGTTTCTGA |
| 3_known | E17 | Tb927.3.4860_for | GGGGACAAGTTTGTACAAAAAAGCAGGCTATGAAAAGGAAGGGGATTGCGTT | Tb927.3.4860_rev | GGGGACCACTTTGTACAAGAAAGCTGGGTCTACAGAACACCTAGCATTTCCAATTTTAT |
| 3_known | E18 | Tb927.7.7080_for | GGGGACAAGTTTGTACAAAAAAGCAGGCTATGCACAGTTCTCTCATAATACTGCG | Tb927.7.7080_rev | GGGGACCACTTTGTACAAGAAAGCTGGGTTCACGCAGCCACGAAGTCC |
| 3_known | E19 | Tb927.10.3230_for | GGGGACAAGTTTGTACAAAAAAGCAGGCTATGAACAGTAATGCTCCGCTTATCTTT | Tb927.10.3230_rev | GGGGACCACTTTGTACAAGAAAGCTGGGTTCAATCCTCATCATCGCTAACTAGTAGG |
| 3_known | E20 | Tb927.11.8190_for | GGGGACAAGTTTGTACAAAAAAGCAGGCTATGAACACTCGCACCAAAAATAAGAAAC | Tb927.11.8190_rev | GGGGACCACTTTGTACAAGAAAGCTGGGTCTAGTCCCTAACGTCTCCCACAC |
| 3_known | E21 | Tb927.9.15380_for | GGGGACAAGTTTGTACAAAAAAGCAGGCTATGTCCCGGCGGGTGTTT | Tb927.9.15380_rev | GGGGACCACTTTGTACAAGAAAGCTGGGTTTACATTCGTGGGGCCAAGTCT |
| 3_known | E22 | Tb927.10.11800_for | GGGGACAAGTTTGTACAAAAAAGCAGGCTATGGGGTTGCTTAAGGTGGAGA | Tb927.10.11800_rev | GGGGACCACTTTGTACAAGAAAGCTGGGTCTAGAGCTCTACACCGAGGTCAG |
| 3_known | E23 | Tb927.10.10210_for | GGGGACAAGTTTGTACAAAAAAGCAGGCTATGACAACCATGGGGAAGACAATAAAA | Tb927.10.10210_rev | GGGGACCACTTTGTACAAGAAAGCTGGGTTTACACCATCGCAGCGACAAG |
| 3_known | E24 | Tb927.3.2290_for | GGGGACAAGTTTGTACAAAAAAGCAGGCTATGTCCGGGCCGCCTAAG | Tb927.3.2290_rev | GGGGACCACTTTGTACAAGAAAGCTGGGTTTACACGCAGGTACTGATATCTTTCTCC |
| 3_known | F1 | Tb927.4.1300_for | GGGGACAAGTTTGTACAAAAAAGCAGGCTATGTCGACTGCTCTGTTCGTTC | Tb927.4.1300_rev | GGGGACCACTTTGTACAAGAAAGCTGGGTTCAGCAACCGCCGTCCTT |
| 3_known | F2 | Tb927.9.9020_for | GGGGACAAGTTTGTACAAAAAAGCAGGCTATGGGCTTACTTGAACGCCG | Tb927.9.9020_rev | GGGGACCACTTTGTACAAGAAAGCTGGGTTTATTCGTTAGACGTTTTAACCACTATTTG |
| 3_known | F3 | Tb927.1.990_for | GGGGACAAGTTTGTACAAAAAAGCAGGCTATGAGTGGCGAAAACATTAAAAATTTGC | Tb927.1.990_rev | GGGGACCACTTTGTACAAGAAAGCTGGGTCTAGTCGGAGGCAAAGAGACGG |
| 3_known | F4 | Tb927.7.5920_for | GGGGACAAGTTTGTACAAAAAAGCAGGCTATGCACGGTGACGGTTTGG | Tb927.7.5920_rev | GGGGACCACTTTGTACAAGAAAGCTGGGTTTACGTGTCTATGCGAGGATGAGC |
| 3_known | F5 | Tb927.10.14180_for | GGGGACAAGTTTGTACAAAAAAGCAGGCTATGCTTCATTCAACTAGATACCCGTCG | Tb927.10.14180_rev | GGGGACCACTTTGTACAAGAAAGCTGGGTTCAGTGAACCCGCTGATTTTGG |
| 3_known | F6 | Tb927.6.750_for | GGGGACAAGTTTGTACAAAAAAGCAGGCTATGGTGGGGAAATACTTGTTAATTTCGC | Tb927.6.750_rev | GGGGACCACTTTGTACAAGAAAGCTGGGTTCACAAGTCGTCTAGAACATCGTTC |
| 3_known | F7 | Tb927.11.9670_for | GGGGACAAGTTTGTACAAAAAAGCAGGCTATGTCAACTTCTCTACGGCACAC | Tb927.11.9670_rev | GGGGACCACTTTGTACAAGAAAGCTGGGTTCATCCCCTATAATAGCGCGGG |
| 3_known | F8 | Tb927.10.160_for | GGGGACAAGTTTGTACAAAAAAGCAGGCTATGGCTTTAGTTCGTAGGATAGGATCA | Tb927.10.160_rev | GGGGACCACTTTGTACAAGAAAGCTGGGTTCAAACACTAGAAAGCCTTCCTTGTT |
| 3_known | F9 | Tb927.11.3150_for | GGGGACAAGTTTGTACAAAAAAGCAGGCTATGACTGAGTACAAGAAACTTTGCGC | Tb927.11.3150_rev | GGGGACCACTTTGTACAAGAAAGCTGGGTCTAAAACTTGCAAGCTTCCAAAGAGTG |
| 3_known | F10 | Tb927.8.7150_for | GGGGACAAGTTTGTACAAAAAAGCAGGCTATGACACAAAGGAAAGTGTACCCG | Tb927.8.7150_rev | GGGGACCACTTTGTACAAGAAAGCTGGGTTTAGCACAGAACCATAGCGCG |
| 3_known | F11 | Tb927.9.8850_for | GGGGACAAGTTTGTACAAAAAAGCAGGCTATGTCGGACGAGGAACAAACTG | Tb927.9.8850_rev | GGGGACCACTTTGTACAAGAAAGCTGGGTTCAAAAGCATTTGCTGTGTACGATG |
| 3_known | F12 | Tb927.9.8880_for | GGGGACAAGTTTGTACAAAAAAGCAGGCTATGTCGGACGAGGAACAAACTG | Tb927.9.8880_rev | GGGGACCACTTTGTACAAGAAAGCTGGGTTTAAAAGCATTTGCTGTGTACGATGC |
| 3_known | F13 | Tb927.7.1300_for | GGGGACAAGTTTGTACAAAAAAGCAGGCTATGACCCCGTTTTCATGTCGC | Tb927.7.1300_rev | GGGGACCACTTTGTACAAGAAAGCTGGGTTTACTCACCTTTGATTGCCGAAAGAA |
| 3_known | F14 | Tb927.2.3370_for | GGGGACAAGTTTGTACAAAAAAGCAGGCTATGGTGTCGAAGGGTTTACAGATGG | Tb927.2.3370_rev | GGGGACCACTTTGTACAAGAAAGCTGGGTTCAGCATTTCCCACGAGCG |
| 3_known | F15 | Tb927.10.9430_for | GGGGACAAGTTTGTACAAAAAAGCAGGCTATGTCTTCTTCAACGTGTGCCTT | Tb927.10.9430_rev | GGGGACCACTTTGTACAAGAAAGCTGGGTTTATGGAGTGGCACCGGTTGC |
| 3_known | F16 | Tb927.3.3960_for | GGGGACAAGTTTGTACAAAAAAGCAGGCTATGATGCCGTGCAAGTCATTTCC | Tb927.3.3960_rev | GGGGACCACTTTGTACAAGAAAGCTGGGTTTAATCCACTGGTGAAGGCGGC |
| 3_known | F17 | Tb927.10.13720_for | GGGGACAAGTTTGTACAAAAAAGCAGGCTATGGCGCATGTTGGACAGC | Tb927.10.13720_rev | GGGGACCACTTTGTACAAGAAAGCTGGGTTCAACATTTATTTTCAACCTCTTCATCAAC |
| 3_known | F18 | Tb927.11.15760_for | GGGGACAAGTTTGTACAAAAAAGCAGGCTATGAGGAGCAGCGGTGATCT | Tb927.11.15760_rev | GGGGACCACTTTGTACAAGAAAGCTGGGTCTACGAGTTTTGAACAACCTGCGC |
| 3_known | F19 | Tb927.3.4920_for | GGGGACAAGTTTGTACAAAAAAGCAGGCTATGTTTCCGGACCTGATTCCCT | Tb927.3.4920_rev | GGGGACCACTTTGTACAAGAAAGCTGGGTCTAATTTTTTGCAATCACCTCTGAAGGC |
| 3_known | F20 | Tb927.9.4390_for | GGGGACAAGTTTGTACAAAAAAGCAGGCTATGAAAATTACCTGTACGGATGAAAGTGG | Tb927.9.4390_rev | GGGGACCACTTTGTACAAGAAAGCTGGGTTCACTCCTCCATCAGAAGTGACAT |
| 3_known | F21 | Tb927.10.5270_for | GGGGACAAGTTTGTACAAAAAAGCAGGCTATGACCGGGTTGAAGATTGACATC | Tb927.10.5270_rev | GGGGACCACTTTGTACAAGAAAGCTGGGTTCACTGCACCAACTTATTTAGTTCCTC |
| 3_known | F22 | Tb927.10.2910_for | GGGGACAAGTTTGTACAAAAAAGCAGGCTATGGATGCGATCCCACCTGA | Tb927.10.2910_rev | GGGGACCACTTTGTACAAGAAAGCTGGGTCTACCTCCTGTATCGCCCTTCA |
| 3_known | F23 | Tb927.11.12880_for | GGGGACAAGTTTGTACAAAAAAGCAGGCTATGGAGGGTGGTGCCAAGT | Tb927.11.12880_rev | GGGGACCACTTTGTACAAGAAAGCTGGGTCTAACGGAGTTCGTAACGGCC |
| 3_known | F24 | Tb927.10.12700_for | GGGGACAAGTTTGTACAAAAAAGCAGGCTATGCTTAAGTGTGTCAGCCGAG | Tb927.10.12700_rev | GGGGACCACTTTGTACAAGAAAGCTGGGTTCACGGCTTGGCATAAACTGTT |
| 3_known | G1 | Tb927.10.2390_for | GGGGACAAGTTTGTACAAAAAAGCAGGCTATGACTGATGTCTCGCTTCGACC | Tb927.10.2390_rev | GGGGACCACTTTGTACAAGAAAGCTGGGTTCATCCCTTTCCGGCGGT |
| 3_known | G2 | Tb927.6.4220_for | GGGGACAAGTTTGTACAAAAAAGCAGGCTATGGTAACAGCAAATGGCTCCTC | Tb927.6.4220_rev | GGGGACCACTTTGTACAAGAAAGCTGGGTTTACTGATAGTGCTGAATCTCTTTCCATAT |
| 3_known | G3 | Tb927.1.780_for | GGGGACAAGTTTGTACAAAAAAGCAGGCTATGCATTCGCTACGGAGCAAC | Tb927.1.780_rev | GGGGACCACTTTGTACAAGAAAGCTGGGTTCAAAGGTGAAGCATCTTCACGG |
| 3_known | G4 | Tb927.10.10440_for | GGGGACAAGTTTGTACAAAAAAGCAGGCTATGTTAGGTCGATTTGATTTTTTAGGGAAA | Tb927.10.10440_rev | GGGGACCACTTTGTACAAGAAAGCTGGGTTCACTTATTATCTCCTGGGGGCC |
| 3_known | G5 | Tb927.3.3920_for | GGGGACAAGTTTGTACAAAAAAGCAGGCTATGGTTATTTGGTTTTCCGCTGGT | Tb927.3.3920_rev | GGGGACCACTTTGTACAAGAAAGCTGGGTTCACACGCGAGGACGTTTTC |
| 3_known | G6 | Tb927.10.14040_for | GGGGACAAGTTTGTACAAAAAAGCAGGCTATGTCGGAGCACTTTCTCCTTTTG | Tb927.10.14040_rev | GGGGACCACTTTGTACAAGAAAGCTGGGTCTAACACCTTTGCAAATTGGTCATCTTAA |
| 3_known | G7 | Tb927.7.4970_for | GGGGACAAGTTTGTACAAAAAAGCAGGCTATGACAAACTCTTCAGTGCAGCA | Tb927.7.4970_rev | GGGGACCACTTTGTACAAGAAAGCTGGGTCTACACCTTAGAAGTACTTGCCATCCA |
| 3_known | G8 | Tb927.2.3480_for | GGGGACAAGTTTGTACAAAAAAGCAGGCTATGATTCCATCATTTGCTCCAGGG | Tb927.2.3480_rev | GGGGACCACTTTGTACAAGAAAGCTGGGTTCACTGTTCAGCACCATCCACC |
| 3_known | G9 | Tb927.10.14530_for | GGGGACAAGTTTGTACAAAAAAGCAGGCTATGGCCAATGATGACCCCAAGG | Tb927.10.14530_rev | GGGGACCACTTTGTACAAGAAAGCTGGGTTTACTTCCCTTCTGAAGTTCCACCG |
| 3_known | G10 | Tb927.11.6710_for | GGGGACAAGTTTGTACAAAAAAGCAGGCTATGGCAAAGCGCTCGAAGTC | Tb927.11.6710_rev | GGGGACCACTTTGTACAAGAAAGCTGGGTTTAATGCCTATCTGCAAGGAAGCG |
| 3_known | G11 | Tb927.4.820_for | GGGGACAAGTTTGTACAAAAAAGCAGGCTATGAGCAGGAAGTGTAGGAAGAGAC | Tb927.4.820_rev | GGGGACCACTTTGTACAAGAAAGCTGGGTTTAACTTCCAGATGCGGGGAAAGG |
| 3_known | G12 | Tb927.6.1800_for | GGGGACAAGTTTGTACAAAAAAGCAGGCTATGGGGGTTATGTTACCGAAACCG | Tb927.6.1800_rev | GGGGACCACTTTGTACAAGAAAGCTGGGTTTATTTGTTGCTTTGCTCTGCTAGCG |
| 3_known | G13 | Tb927.10.14070_for | GGGGACAAGTTTGTACAAAAAAGCAGGCTATGTCCGACCAGTGTGCCA | Tb927.10.14070_rev | GGGGACCACTTTGTACAAGAAAGCTGGGTTCATACCCGAGGTGGAATGAGTG |
| 3_known | G14 | Tb927.4.5240_for | GGGGACAAGTTTGTACAAAAAAGCAGGCTATGTCACCGCTGAAGATGCG | Tb927.4.5240_rev | GGGGACCACTTTGTACAAGAAAGCTGGGTTCAATGCTGAAAACAAGGTAGCGT |
| 3_known | G15 | Tb927.10.13560_for | GGGGACAAGTTTGTACAAAAAAGCAGGCTATGTCCGACCAGTGTGTCACC | Tb927.10.13560_rev | GGGGACCACTTTGTACAAGAAAGCTGGGTTCATACCCGAGGTGGAATGAGTG |
| 3_known | G16 | Tb927.11.940_for | GGGGACAAGTTTGTACAAAAAAGCAGGCTATGAGACGGGCTGTGGTACTC | Tb927.11.940_rev | GGGGACCACTTTGTACAAGAAAGCTGGGTTTACCGCCCTCCCAGTGC |
| 3_known | G17 | Tb927.10.12850_for | GGGGACAAGTTTGTACAAAAAAGCAGGCTATGTACTGTCACGCTGGCGT | Tb927.10.12850_rev | GGGGACCACTTTGTACAAGAAAGCTGGGTTCACTCGTTATTCTCCATATTGGAGGC |
| 3_known | G18 | Tb927.10.2770_for | GGGGACAAGTTTGTACAAAAAAGCAGGCTATGACGTCTTCAATGATCCCTGTGG | Tb927.10.2770_rev | GGGGACCACTTTGTACAAGAAAGCTGGGTTCACGCTGCGACACACTTG |
| 3_known | G19 | Tb927.10.14790_for | GGGGACAAGTTTGTACAAAAAAGCAGGCTATGTCTCGGCCAGAAGGTGAG | Tb927.10.14790_rev | GGGGACCACTTTGTACAAGAAAGCTGGGTCTACTCCTTGCGCTTCTTCTCTAGC |
| 3_known | G20 | Tb927.7.2480_for | GGGGACAAGTTTGTACAAAAAAGCAGGCTATGGTTACCTCTACTGGTCTTCTCTC | Tb927.7.2480_rev | GGGGACCACTTTGTACAAGAAAGCTGGGTTCACTCAACGCGATATATCAAATGCA |
| 3_known | G21 | Tb927.5.550_for | GGGGACAAGTTTGTACAAAAAAGCAGGCTATGGGGCGAGGAATTCTGAACT | Tb927.5.550_rev | GGGGACCACTTTGTACAAGAAAGCTGGGTTTAAAATATGCTCACGTACTCATCTACGC |
| 3_known | G22 | Tb927.9.3280_for | GGGGACAAGTTTGTACAAAAAAGCAGGCTATGACGGCAGTGGTGAATGAGT | Tb927.9.3280_rev | GGGGACCACTTTGTACAAGAAAGCTGGGTTTACAAATTGTTCCACACTGACAAAAAACT |
| 3_known | G23 | Tb927.3.1430_for | GGGGACAAGTTTGTACAAAAAAGCAGGCTATGCGCTGCACAATGTCTTCG | Tb927.3.1430_rev | GGGGACCACTTTGTACAAGAAAGCTGGGTTCAGTCGAGTGCCCTTTCAATAGC |
| 3_known | G24 | Tb927.11.11680_for | GGGGACAAGTTTGTACAAAAAAGCAGGCTATGCTCCGTCGTTTGGCC | Tb927.11.11680_rev | GGGGACCACTTTGTACAAGAAAGCTGGGTTTAGGCGAGGTCGAGCACA |
| 3_known | H1 | Tb927.11.2910_for | GGGGACAAGTTTGTACAAAAAAGCAGGCTATGGCAGTTGTACGCAGGTG | Tb927.11.2910_rev | GGGGACCACTTTGTACAAGAAAGCTGGGTTCACATGAAAGGCGCTCCAGT |
| 3_known | H3 | Tb927.3.5340_for | GGGGACAAGTTTGTACAAAAAAGCAGGCTATGCATACTCTGTCCTCATGTGATTTT | Tb927.3.5340_rev | GGGGACCACTTTGTACAAGAAAGCTGGGTTTAATCCAACTCGTCAGGGTCATGA |
| 3_known | H4 | Tb927.11.14140_for | GGGGACAAGTTTGTACAAAAAAGCAGGCTATGAAACGGTCGGTGTCGAAG | Tb927.11.14140_rev | GGGGACCACTTTGTACAAGAAAGCTGGGTTCACACCTCTCTGACATTTCTGTACA |
| 3_known | H5 | Tb927.5.2760_for | GGGGACAAGTTTGTACAAAAAAGCAGGCTATGTTTGCTAGGGTTAAAAGGGTAAGG | Tb927.5.2760_rev | GGGGACCACTTTGTACAAGAAAGCTGGGTTCACCACCTTCCACCCAAATGC |
| 3_known | H6 | Tb927.4.4250_for | GGGGACAAGTTTGTACAAAAAAGCAGGCTATGGCCCCCGCGCTTCGG | Tb927.4.4250_rev | GGGGACCACTTTGTACAAGAAAGCTGGGTCTACAAACAACTCATTTCCCGGAGA |
| 3_known | H8 | Tb927.9.4910_for | GGGGACAAGTTTGTACAAAAAAGCAGGCTATGTCAGCGTCCCGGATGG | Tb927.9.4910_rev | GGGGACCACTTTGTACAAGAAAGCTGGGTTCAATCCCCCCTAATCTTGCCA |
| 3_known | H9 | Tb927.4.400_for | GGGGACAAGTTTGTACAAAAAAGCAGGCTATGGAAAGTATGCCGTCTGAGGA | Tb927.4.400_rev | GGGGACCACTTTGTACAAGAAAGCTGGGTTTAGGCATAAAGCCTTGCCACC |
| 3_known | H10 | Tb927.11.16740_for | GGGGACAAGTTTGTACAAAAAAGCAGGCTATGCGGGGTTTGGCGGCA | Tb927.11.16740_rev | GGGGACCACTTTGTACAAGAAAGCTGGGTCTACAGATGCTCCGCCAACTC |
| 3_known | H11 | Tb927.9.14420_for | GGGGACAAGTTTGTACAAAAAAGCAGGCTATGAGCGCGCTCGTTCGA | Tb927.9.14420_rev | GGGGACCACTTTGTACAAGAAAGCTGGGTCTACAAGAACCTCCACCAAGGAGT |
| 3_known | H12 | Tb927.10.13190_for | GGGGACAAGTTTGTACAAAAAAGCAGGCTATGAGACCTCAAAAAGAAAAAAACACCG | Tb927.10.13190_rev | GGGGACCACTTTGTACAAGAAAGCTGGGTCTAAAGCTTCGTGACGTCTTTTCTTG |
| 3_known | H13 | Tb927.4.1660_for | GGGGACAAGTTTGTACAAAAAAGCAGGCTATGGGGGTTGAAGTAGCAGACT | Tb927.4.1660_rev | GGGGACCACTTTGTACAAGAAAGCTGGGTTTAGTCATCACCGACCAAAATGCTC |
| 3_known | H14 | Tb927.4.4290_for | GGGGACAAGTTTGTACAAAAAAGCAGGCTATGTGTAGGCAATTTAACAGTCGCG | Tb927.4.4290_rev | GGGGACCACTTTGTACAAGAAAGCTGGGTCTACAAACAACTCATTTCCCGGAGA |
| 3_known | H15 | Tb927.8.4640_for | GGGGACAAGTTTGTACAAAAAAGCAGGCTATGGAGCCAGCAGCTGAGA | Tb927.8.4640_rev | GGGGACCACTTTGTACAAGAAAGCTGGGTCTACATACAGAGATCCCAATACTTGAAATA |
| 3_known | H16 | Tb927.9.2450_for | GGGGACAAGTTTGTACAAAAAAGCAGGCTATGATACGCGGCTCACACCT | Tb927.9.2450_rev | GGGGACCACTTTGTACAAGAAAGCTGGGTTTAGATTTCACTGGGTTTCTGGACG |
| 3_known | H17 | Tb927.5.2960_for | GGGGACAAGTTTGTACAAAAAAGCAGGCTATGAGTAAACCTACCGTTTCGGAGG | Tb927.5.2960_rev | GGGGACCACTTTGTACAAGAAAGCTGGGTTTATGGAGAGTCGACGGATCCC |
| 3_known | H18 | Tb927.11.10990_for | GGGGACAAGTTTGTACAAAAAAGCAGGCTATGAGCAGTGCGCACAACA | Tb927.11.10990_rev | GGGGACCACTTTGTACAAGAAAGCTGGGTTCAAGGCTCAACAATACGGGATATG |
| 3_known | H19 | Tb927.10.5690_for | GGGGACAAGTTTGTACAAAAAAGCAGGCTATGCTCAAGTTTGTTCTTGTGTTAATACTG | Tb927.10.5690_rev | GGGGACCACTTTGTACAAGAAAGCTGGGTTTACGTACACTCCTTCATTTCCTCCG |
| 3_known | H20 | Tb927.10.4160_for | GGGGACAAGTTTGTACAAAAAAGCAGGCTATGTCTTTTTTCAGGGGCTTAAACTTCA | Tb927.10.4160_rev | GGGGACCACTTTGTACAAGAAAGCTGGGTTCACCGTTTTTTCAGTCGTGGG |
| 3_known | H21 | Tb927.10.7800_for | GGGGACAAGTTTGTACAAAAAAGCAGGCTATGATAAAGGAGGGGGGAAATGAGG | Tb927.10.7800_rev | GGGGACCACTTTGTACAAGAAAGCTGGGTTCACAATAGGGGACTTTCACATAGCT |
| 3_known | H22 | Tb927.9.6040_for | GGGGACAAGTTTGTACAAAAAAGCAGGCTATGAGTCAGGATAGGGGCAAGC | Tb927.9.6040_rev | GGGGACCACTTTGTACAAGAAAGCTGGGTCTACATGGCTCTCATTTTGGCCA |
| 3_known | H23 | Tb927.11.9030_for | GGGGACAAGTTTGTACAAAAAAGCAGGCTATGGCTAACGTTAATAACGAGACCAC | Tb927.11.9030_rev | GGGGACCACTTTGTACAAGAAAGCTGGGTTTAGATCCATTTTCCAATAAAGGCCATTAA |

| | | | | | |
|---|---|---|---|---|---|
| 3_known | H24 | Tb927.10.14800_for | GGGGACAAGTTTGTACAAAAAAGCAGGCTATGAATCGGTATACGATATTGGGTCAGC | Tb927.10.14800_rev | GGGGACCACTTTGTACAAGAAAGCTGGGTTTAAAAGTTGAACGCTTCATCGTCAAG |
| 3_known | I1 | Tb927.11.6970_for | GGGGACAAGTTTGTACAAAAAAGCAGGCTATGGATCCCAGCAGTGGTAACG | Tb927.11.6970_rev | GGGGACCACTTTGTACAAGAAAGCTGGGTTCATTGGTAGGCCGTTAAATGTGTC |
| 3_known | I2 | Tb927.11.8310_for | GGGGACAAGTTTGTACAAAAAAGCAGGCTATGTCGACGCTAGCTTCTGCT | Tb927.11.8310_rev | GGGGACCACTTTGTACAAGAAAGCTGGGTTCAAAACGCCCTCCTATGAAAATGA |
| 3_known | I3 | Tb927.3.5660_for | GGGGACAAGTTTGTACAAAAAAGCAGGCTATGGCAATCAAATCACGAGGAGTT | Tb927.3.5660_rev | GGGGACCACTTTGTACAAGAAAGCTGGGTTTATGCCGTCCATTCCGCATC |
| 3_known | I4 | Tb927.10.1680_for | GGGGACAAGTTTGTACAAAAAAGCAGGCTATGAATGAGGATGGTGATTGTCGG | Tb927.10.1680_rev | GGGGACCACTTTGTACAAGAAAGCTGGGTTCAAACAGTCATTATATTTGGACAGTGCT |
| 3_known | I5 | Tb927.3.5580_for | GGGGACAAGTTTGTACAAAAAAGCAGGCTATGACTGAAGCCGTTGTAACACC | Tb927.3.5580_rev | GGGGACCACTTTGTACAAGAAAGCTGGGTTCACTGGGTTGCCGCCTT |
| 3_known | I6 | Tb927.10.3560_for | GGGGACAAGTTTGTACAAAAAAGCAGGCTATGTCACCGAAGAAAAACTCGGC | Tb927.10.3560_rev | GGGGACCACTTTGTACAAGAAAGCTGGGTTCAATACCTTTGGTAGTTGTACGTGC |
| 3_known | I7 | Tb927.8.7510_for | GGGGACAAGTTTGTACAAAAAAGCAGGCTATGGCTAGCGTGTCATCAGGA | Tb927.8.7510_rev | GGGGACCACTTTGTACAAGAAAGCTGGGTTTAGTCGTCACGTGCATCAGGC |
| 3_known | I8 | Tb927.7.5490_for | GGGGACAAGTTTGTACAAAAAAGCAGGCTATGCCCCCAAAGCAGCAC | Tb927.7.5490_rev | GGGGACCACTTTGTACAAGAAAGCTGGGTTCAGGTTGTTTTGCCCTCGC |
| 3_known | I9 | Tb927.8.7170_for | GGGGACAAGTTTGTACAAAAAAGCAGGCTATGCCTCAGGTGGATCTCGTT | Tb927.8.7170_rev | GGGGACCACTTTGTACAAGAAAGCTGGGTCTACTGCCATCCCAAACGCC |
| 3_known | I10 | Tb927.8.5630_for | GGGGACAAGTTTGTACAAAAAAGCAGGCTATGGCGCGCGTTGAACTG | Tb927.8.5630_rev | GGGGACCACTTTGTACAAGAAAGCTGGGTTCACATTACTTTCCCCACAAGCTTC |
| 3_known | I11 | Tb927.8.8100_for | GGGGACAAGTTTGTACAAAAAAGCAGGCTATGTCCCTGTCGGAGCCC | Tb927.8.8100_rev | GGGGACCACTTTGTACAAGAAAGCTGGGTCTACGAGCAGTCGAAAGTTTTCACA |
| 3_known | I12 | Tb927.8.8090_for | GGGGACAAGTTTGTACAAAAAAGCAGGCTATGTCCCTGTCGGAGCCC | Tb927.8.8090_rev | GGGGACCACTTTGTACAAGAAAGCTGGGTCTACCGGCAACTAAAACCCTTCAT |
| 3_known | I13 | Tb927.9.12720_for | GGGGACAAGTTTGTACAAAAAAGCAGGCTATGTTCAGGACTATAGGAATCGTGCG | Tb927.9.12720_rev | GGGGACCACTTTGTACAAGAAAGCTGGGTTCAATGTAAAGAATTTTCCCCCAAACGT |
| 3_known | I14 | Tb927.10.6460_for | GGGGACAAGTTTGTACAAAAAAGCAGGCTATGCTGCCTATACTAAAGGCCGG | Tb927.10.6460_rev | GGGGACCACTTTGTACAAGAAAGCTGGGTTCATGCTCCCTCACACGAAATG |
| 3_known | I15 | Tb927.10.14720_for | GGGGACAAGTTTGTACAAAAAAGCAGGCTATGTACGGTGGTTATGGTGCAGG | Tb927.10.14720_rev | GGGGACCACTTTGTACAAGAAAGCTGGGTCTAGAGTTTTGTCTCCCTCTCCAGAA |
| 3_known | I16 | Tb927.11.8990_for | GGGGACAAGTTTGTACAAAAAAGCAGGCTATGCTTGGAGCGAACCTTGG | Tb927.11.8990_rev | GGGGACCACTTTGTACAAGAAAGCTGGGTCTAGTGCCACTTGGACAGCAG |
| 3_known | I17 | Tb927.11.1960_for | GGGGACAAGTTTGTACAAAAAAGCAGGCTATGAACCACCCTGATGAACTTGAAG | Tb927.11.1960_rev | GGGGACCACTTTGTACAAGAAAGCTGGGTTTACATCACTACCTTGGCACAGCA |
| 3_known | I18 | Tb927.11.4910_for | GGGGACAAGTTTGTACAAAAAAGCAGGCTATGGACAACGGTAAGCGCATT | Tb927.11.4910_rev | GGGGACCACTTTGTACAAGAAAGCTGGGTTCACTTCGACCTCCGCGT |
| 3_known | I19 | Tb927.8.1520_for | GGGGACAAGTTTGTACAAAAAAGCAGGCTATGTCGCTCCCCGCTGTG | Tb927.8.1520_rev | GGGGACCACTTTGTACAAGAAAGCTGGGTCTACACCTCTCCATCATTGTTTGATGC |
| 3_known | I20 | Tb927.11.11950_for | GGGGACAAGTTTGTACAAAAAAGCAGGCTATGATTCGCCTTCGTTATCGTGC | Tb927.11.11950_rev | GGGGACCACTTTGTACAAGAAAGCTGGGTTTACACAAGTACCAACCCTCGAGA |
| 3_known | I21 | Tb927.2.5250_for | GGGGACAAGTTTGTACAAAAAAGCAGGCTATGAGTGAGTTCCCGTTGGTGT | Tb927.2.5250_rev | GGGGACCACTTTGTACAAGAAAGCTGGGTTCACGTGCTCCCGATAGGC |
| 3_known | I22 | Tb927.3.830_for | GGGGACAAGTTTGTACAAAAAAGCAGGCTATGGGTGTTCTTGGCCTTTCG | Tb927.3.830_rev | GGGGACCACTTTGTACAAGAAAGCTGGGTCTACTTTTTGACAGCCTTTTTGTGTCC |
| 3_known | I23 | Tb927.10.9410_for | GGGGACAAGTTTGTACAAAAAAGCAGGCTATGTCAGAGAAGGAAGGAAAAGTAATACCC | Tb927.10.9410_rev | GGGGACCACTTTGTACAAGAAAGCTGGGTTTAAATAATCTTATCATTTACCTTCGCTCC |
| 3_known | I24 | Tb927.6.4120_for | GGGGACAAGTTTGTACAAAAAAGCAGGCTATGTCTGGCTTCCCGCCT | Tb927.6.4120_rev | GGGGACCACTTTGTACAAGAAAGCTGGGTCTACGATTTCCACATAGAAGAGACCCA |
| 3_known | J1 | Tb927.8.620_for | GGGGACAAGTTTGTACAAAAAAGCAGGCTATGAAGCGTGTTACTTCACATATTTCGC | Tb927.8.620_rev | GGGGACCACTTTGTACAAGAAAGCTGGGTTCACACCTTCAACACTGACCCA |
| 3_known | J2 | Tb927.11.10450_for | GGGGACAAGTTTGTACAAAAAAGCAGGCTATGCACGCGACAGTCACTG | Tb927.11.10450_rev | GGGGACCACTTTGTACAAGAAAGCTGGGTTCAGTAGACGTGGTCGTACAGG |
| 3_known | J3 | Tb927.6.2040_for | GGGGACAAGTTTGTACAAAAAAGCAGGCTATGGGATGTTTGCCCAGTGTG | Tb927.6.2040_rev | GGGGACCACTTTGTACAAGAAAGCTGGGTTTACAAGACGATTACCCGCTCTTCC |
| 3_known | J4 | Tb927.2.5060_for | GGGGACAAGTTTGTACAAAAAAGCAGGCTATGCCACCTAAGAAGAAGGATGACA | Tb927.2.5060_rev | GGGGACCACTTTGTACAAGAAAGCTGGGTTTACTTCTTACCTCCTTTAGCTGCATTAAA |
| 3_known | J5 | Tb927.10.5680_for | GGGGACAAGTTTGTACAAAAAAGCAGGCTATGAAGGTGAATTGTTTTCTACCTTTGTTG | Tb927.10.5680_rev | GGGGACCACTTTGTACAAGAAAGCTGGGTTCAAACAAAAACAGGAACGAAAACTAAGAA |
| 3_known | J6 | Tb927.11.10110_for | GGGGACAAGTTTGTACAAAAAAGCAGGCTATGGTACAACAACAGCCCGTTG | Tb927.11.10110_rev | GGGGACCACTTTGTACAAGAAAGCTGGGTCTAAATCCTCTCGTGAACGACGG |
| 3_known | J7 | Tb927.8.1910_for | GGGGACAAGTTTGTACAAAAAAGCAGGCTATGAACTGCAGAGAGTGGTTGGC | Tb927.8.1910_rev | GGGGACCACTTTGTACAAGAAAGCTGGGTCTACAAATGCGGTTTTCTTCCAATATCG |
| 3_known | J8 | Tb927.10.3080_for | GGGGACAAGTTTGTACAAAAAAGCAGGCTATGGGGATAGTCCGAGTACAGAGG | Tb927.10.3080_rev | GGGGACCACTTTGTACAAGAAAGCTGGGTTTATTCCGGCTCCTGTAAACTGC |
| 3_known | J9 | Tb927.11.9010_for | GGGGACAAGTTTGTACAAAAAAGCAGGCTATGGCTAACGTTAATAACGAGACCAC | Tb927.11.9010_rev | GGGGACCACTTTGTACAAGAAAGCTGGGTTTAGATCCACTTTCCGATAAAGGCCA |
| 3_known | J10 | Tb927.11.9000_for | GGGGACAAGTTTGTACAAAAAAGCAGGCTATGGCTAACGTTAATAACGAGACCAC | Tb927.11.9000_rev | GGGGACCACTTTGTACAAGAAAGCTGGGTTTAGACCCATTTTCCTAACACGGC |
| 3_known | J11 | Tb927.7.4980_for | GGGGACAAGTTTGTACAAAAAAGCAGGCTATGCCTGGTTGCCTTTCGG | Tb927.7.4980_rev | GGGGACCACTTTGTACAAGAAAGCTGGGTTCAGTAAACCACGATCAGCACAAC |
| 3_known | J12 | Tb927.8.5530_for | GGGGACAAGTTTGTACAAAAAAGCAGGCTATGATAGATGATGTGGTAACCGATGATG | Tb927.8.5530_rev | GGGGACCACTTTGTACAAGAAAGCTGGGTTCAAAGTGGATCGACACCGTATTG |
| 3_known | J13 | Tb927.8.7410_for | GGGGACAAGTTTGTACAAAAAAGCAGGCTATGCTAATGTGCATGCGCCC | Tb927.8.7410_rev | GGGGACCACTTTGTACAAGAAAGCTGGGTCTACAAGTCTGACTTATCCTCCTTCTTCT |
| 3_known | J14 | Tb927.10.15850_for | GGGGACAAGTTTGTACAAAAAAGCAGGCTATGATAGAGAGTAACTTGCTCTCGCA | Tb927.10.15850_rev | GGGGACCACTTTGTACAAGAAAGCTGGGTTCAGCACTCATAAATGCGTCGAA |
| 3_known | J15 | Tb927.9.4950_for | GGGGACAAGTTTGTACAAAAAAGCAGGCTATGAACAAGTGCAAGCAAGCCG | Tb927.9.4950_rev | GGGGACCACTTTGTACAAGAAAGCTGGGTTCACGGGGCAGATCCCTCG |
| 3_known | J17 | Tb927.11.15640_for | GGGGACAAGTTTGTACAAAAAAGCAGGCTATGCGCAAGCAATTATTTTTCACCC | Tb927.11.15640_rev | GGGGACCACTTTGTACAAGAAAGCTGGGTTCACGATGCATCTTCCCCGT |
| 3_known | J18 | Tb927.10.15270_for | GGGGACAAGTTTGTACAAAAAAGCAGGCTATGTCGTGCGAAGGTTGCG | Tb927.10.15270_rev | GGGGACCACTTTGTACAAGAAAGCTGGGTTCAAGCAGCGGTAGCCGA |
| 3_known | J19 | Tb927.11.2730_for | GGGGACAAGTTTGTACAAAAAAGCAGGCTATGCGCGTTTTGGTTTGTGG | Tb927.11.2730_rev | GGGGACCACTTTGTACAAGAAAGCTGGGTTTACAACTTAGTGGTCCTTCCCCC |
| 3_known | J20 | Tb927.8.5510_for | GGGGACAAGTTTGTACAAAAAAGCAGGCTATGCCACCGAAAAAACTATCCAAGC | Tb927.8.5510_rev | GGGGACCACTTTGTACAAGAAAGCTGGGTTTACTTACGGAGCCACATCTGCA |
| 3_known | J21 | Tb927.8.7140_for | GGGGACAAGTTTGTACAAAAAAGCAGGCTATGAGGACAAGGCGTCGTAGG | Tb927.8.7140_rev | GGGGACCACTTTGTACAAGAAAGCTGGGTCTAAGGGTGCCGACGGAAAG |
| 3_known | J22 | Tb927.6.4850_for | GGGGACAAGTTTGTACAAAAAAGCAGGCTATGTCCGTGCGCCAGATATTC | Tb927.6.4850_rev | GGGGACCACTTTGTACAAGAAAGCTGGGTCTACTGCACGTCACTAAGACCCT |
| 3_known | K2 | Tb927.11.9350_for | GGGGACAAGTTTGTACAAAAAAGCAGGCTATGCAGATGTTGGCCCCTGG | Tb927.11.9350_rev | GGGGACCACTTTGTACAAGAAAGCTGGGTCTAGCCCCTCAAGCAGTTAGTGA |
| 3_known | K3 | Tb927.1.2580_for | GGGGACAAGTTTGTACAAAAAAGCAGGCTATGCAGTTTGGCCCTGTTGTC | Tb927.1.2580_rev | GGGGACCACTTTGTACAAGAAAGCTGGGTTTACCCACTACTCCCCGCAAG |
| 3_known | K5 | Tb927.8.7730_for | GGGGACAAGTTTGTACAAAAAAGCAGGCTATGGCACGGCACCCTTTAGG | Tb927.8.7730_rev | GGGGACCACTTTGTACAAGAAAGCTGGGTCTACGACGTCTTGGAGTGCTTG |
| 3_known | K10 | Tb927.11.7160_for | GGGGACAAGTTTGTACAAAAAAGCAGGCTATGGAGCAGCAGGGAGATAGTG | Tb927.11.7160_rev | GGGGACCACTTTGTACAAGAAAGCTGGGTCTAACGAGTGCGTTTCCGTTCC |
| 3_known | K11 | Tb927.11.16030_for | GGGGACAAGTTTGTACAAAAAAGCAGGCTATGCCGCCTCAGCAACCG | Tb927.11.16030_rev | GGGGACCACTTTGTACAAGAAAGCTGGGTTCACACTTCTGCAAGACGAGATAATTT |
| 3_known | K12 | Tb927.10.3520_for | GGGGACAAGTTTGTACAAAAAAGCAGGCTATGCAGGAACGTCAGGAGGAG | Tb927.10.3520_rev | GGGGACCACTTTGTACAAGAAAGCTGGGTCTATTGCTCTGAGTATTGATGTGGAGC |
| 3_known | K13 | Tb927.4.5110_for | GGGGACAAGTTTGTACAAAAAAGCAGGCTATGCGCGCATCAACGCCA | Tb927.4.5110_rev | GGGGACCACTTTGTACAAGAAAGCTGGGTTCACACGGCAGGCACCAC |
| 3_known | K14 | Tb927.4.5260_for | GGGGACAAGTTTGTACAAAAAAGCAGGCTATGTCACCGCTGAAGATGCG | Tb927.4.5260_rev | GGGGACCACTTTGTACAAGAAAGCTGGGTTCAGGCCCTCTTACGCCAA |
| 3_known | K15 | Tb927.4.4740_for | GGGGACAAGTTTGTACAAAAAAGCAGGCTATGGCCCGGCAACCTTTG | Tb927.4.4740_rev | GGGGACCACTTTGTACAAGAAAGCTGGGTCTACTCCCTATGAGGTTTTGAATTCTTTGG |
| 3_known | K16 | Tb927.3.1630_for | GGGGACAAGTTTGTACAAAAAAGCAGGCTATGGCTGAGGCAACCGAAAG | Tb927.3.1630_rev | GGGGACCACTTTGTACAAGAAAGCTGGGTTTATTGTCGCCTGAAGGAATGACTC |
| 3_known | K17 | Tb927.4.4910_for | GGGGACAAGTTTGTACAAAAAAGCAGGCTATGCGTCGCAACGTCCTG | Tb927.4.4910_rev | GGGGACCACTTTGTACAAGAAAGCTGGGTTCACTTCCTTACTGCCTTACCCTT |
| 3_known | K18 | Tb927.2.5160_for | GGGGACAAGTTTGTACAAAAAAGCAGGCTATGGTGAAAGAAACAAAATACTACGACGC | Tb927.2.5160_rev | GGGGACCACTTTGTACAAGAAAGCTGGGTCTATTGCTGCGTACACGTTGCA |
| 3_known | K19 | Tb927.4.5290_for | GGGGACAAGTTTGTACAAAAAAGCAGGCTATGTCACCGCTGAAGATGCG | Tb927.4.5290_rev | GGGGACCACTTTGTACAAGAAAGCTGGGTTCAGGCCCTCTTACGCCAA |
| 3_known | K20 | Tb927.4.5250_for | GGGGACAAGTTTGTACAAAAAAGCAGGCTATGTCACCGCTGAAGATGCG | Tb927.4.5250_rev | GGGGACCACTTTGTACAAGAAAGCTGGGTTCAGGCCCTCTTACGCCAA |
| 3_known | K21 | Tb927.4.5270_for | GGGGACAAGTTTGTACAAAAAAGCAGGCTATGTCACCGCTGAAGATGCG | Tb927.4.5270_rev | GGGGACCACTTTGTACAAGAAAGCTGGGTTCAGGCCCTCTTACGCCAA |
| 3_known | K22 | Tb927.10.15800_for | GGGGACAAGTTTGTACAAAAAAGCAGGCTATGAGCGAAAAGCTCCCCGT | Tb927.10.15800_rev | GGGGACCACTTTGTACAAGAAAGCTGGGTCTAAACCCCAAATCGGGCCTG |
| 3_known | K23 | Tb927.4.5280_for | GGGGACAAGTTTGTACAAAAAAGCAGGCTATGTCACCGCTGAAGATGCG | Tb927.4.5280_rev | GGGGACCACTTTGTACAAGAAAGCTGGGTTCAGGCCCTCTTACGCCAA |
| 3_known | K24 | Tb927.5.630_for | GGGGACAAGTTTGTACAAAAAAGCAGGCTATGTCGCGTACTATTGTGCTTGG | Tb927.5.630_rev | GGGGACCACTTTGTACAAGAAAGCTGGGTTTAAAACGGGTCGATGTTTGTCGA |
| 3_known | L1 | Tb927.10.13120_for | GGGGACAAGTTTGTACAAAAAAGCAGGCTATGAGTATCACCAACATCGCTGC | Tb927.10.13120_rev | GGGGACCACTTTGTACAAGAAAGCTGGGTTCATATTTGTTTCCAGAGTTCTCGGTC |
| 3_known | L2 | Tb927.11.8770_for | GGGGACAAGTTTGTACAAAAAAGCAGGCTATGACAGCAACCGCAAGGAG | Tb927.11.8770_rev | GGGGACCACTTTGTACAAGAAAGCTGGGTTTAGAACTGTTCACCGACGTTTGC |
| 3_known | L3 | Tb927.5.2370_for | GGGGACAAGTTTGTACAAAAAAGCAGGCTATGTCTCAAGCTGATAATGCTGAGC | Tb927.5.2370_rev | GGGGACCACTTTGTACAAGAAAGCTGGGTTCAAAGTTTGGCCGCAACGG |
| 3_known | L4 | Tb927.1.3000_for | GGGGACAAGTTTGTACAAAAAAGCAGGCTATGACGTTTATCAGCTCCCTAATACAGG | Tb927.1.3000_rev | GGGGACCACTTTGTACAAGAAAGCTGGGTTTACGTTCCTGGACTAGGAGCG |
| 3_known | L5 | Tb927.10.10220_for | GGGGACAAGTTTGTACAAAAAAGCAGGCTATGATAAGGGTCGTATTCACAATAGCG | Tb927.10.10220_rev | GGGGACCACTTTGTACAAGAAAGCTGGGTTTAGTCGTCCTTTCCCCAATTTCCG |
| 3_known | L6 | Tb927.3.5320_for | GGGGACAAGTTTGTACAAAAAAGCAGGCTATGGATGAAGATGTGGTGCAAATCT | Tb927.3.5320_rev | GGGGACCACTTTGTACAAGAAAGCTGGGTTCAATGGATGTGAAGAACCACTGTATC |
| 3_known | L7 | Tb927.10.3660_for | GGGGACAAGTTTGTACAAAAAAGCAGGCTATGTCCAGGCCCTTTAAGGACTT | Tb927.10.3660_rev | GGGGACCACTTTGTACAAGAAAGCTGGGTCTACTTGTTACGCACGTGTCGG |
| 3_known | L8 | Tb927.11.14600_for | GGGGACAAGTTTGTACAAAAAAGCAGGCTATGCTTGCAACAGTAGTTTTGTTTGT | Tb927.11.14600_rev | GGGGACCACTTTGTACAAGAAAGCTGGGTCTAACCCTCCGATGTCCAACTTATACG |
| 3_known | L9 | Tb927.6.1090_for | GGGGACAAGTTTGTACAAAAAAGCAGGCTATGACTTCACTTTTAGCGCAAACAAC | Tb927.6.1090_rev | GGGGACCACTTTGTACAAGAAAGCTGGGTCTACGAATAAAAATCATATTGTTCATCACC |
| 3_known | L10 | Tb927.5.590_for | GGGGACAAGTTTGTACAAAAAAGCAGGCTATGTCACAGGAGGGTGGGAAAT | Tb927.5.590_rev | GGGGACCACTTTGTACAAGAAAGCTGGGTTCAAGCTTTTCGCTTACGATCAGC |
| 3_known | L11 | Tb927.10.6240_for | GGGGACAAGTTTGTACAAAAAAGCAGGCTATGAACCTACGTGGTGATGGGA | Tb927.10.6240_rev | GGGGACCACTTTGTACAAGAAAGCTGGGTCTACATGATTACACACGAACCTGTGC |
| 3_known | L12 | Tb927.6.440_for | GGGGACAAGTTTGTACAAAAAAGCAGGCTATGGAGAAACCGTCTTGCAGGG | Tb927.6.440_rev | GGGGACCACTTTGTACAAGAAAGCTGGGTCTACACCACCACCTGGAGCA |
| 3_known | L13 | Tb927.9.4680_for | GGGGACAAGTTTGTACAAAAAAGCAGGCTATGGCCCAACAAGGAAAGGTCG | Tb927.9.4680_rev | GGGGACCACTTTGTACAAGAAAGCTGGGTTTACTCACCAAGGTAGGCAGCA |
| 3_known | L14 | Tb927.7.4950_for | GGGGACAAGTTTGTACAAAAAAGCAGGCTATGGGTAAACTAAACCCCGCCC | Tb927.7.4950_rev | GGGGACCACTTTGTACAAGAAAGCTGGGTTTACCTTGTCATCCTTGCCTGTATGT |
| 3_known | L15 | Tb927.8.6060_for | GGGGACAAGTTTGTACAAAAAAGCAGGCTATGTTCCGCACTGTTCCTCGT | Tb927.8.6060_rev | GGGGACCACTTTGTACAAGAAAGCTGGGTCTATACGTTAAGTTCTTTCTTGATTTTTGT |
| 3_known | L16 | Tb927.9.6170_for | GGGGACAAGTTTGTACAAAAAAGCAGGCTATGGGCTTCGGATCATCAAAACC | Tb927.9.6170_rev | GGGGACCACTTTGTACAAGAAAGCTGGGTTTATTCATTTTCATTCGTGGCTACACCG |
| 3_known | L17 | Tb927.11.13090_for | GGGGACAAGTTTGTACAAAAAAGCAGGCTATGTCTCTCACACTCTGGTTTCAGC | Tb927.11.13090_rev | GGGGACCACTTTGTACAAGAAAGCTGGGTCTATTTAAAGACGCGGCCCTCAA |
| 3_known | L18 | Tb927.8.1710_for | GGGGACAAGTTTGTACAAAAAAGCAGGCTATGTCGCAGGTAGCTGGTAATCC | Tb927.8.1710_rev | GGGGACCACTTTGTACAAGAAAGCTGGGTCTACTGTTGCTCAGGCCATGG |
| 3_known | L19 | Tb927.10.6530_for | GGGGACAAGTTTGTACAAAAAAGCAGGCTATGGAGTATCGGTGGCTGAATAGG | Tb927.10.6530_rev | GGGGACCACTTTGTACAAGAAAGCTGGGTCTACTTTCCCCTTTTCACAGCTGT |
| 3_known | L21 | Tb927.10.10240_for | GGGGACAAGTTTGTACAAAAAAGCAGGCTATGGTCTCTTTGATTATCCTTTTTCGCT | Tb927.10.10240_rev | GGGGACCACTTTGTACAAGAAAGCTGGGTTTAAAACAACAAAAATAACAACGCGAGGG |
| 3_known | L22 | Tb927.10.15720_for | GGGGACAAGTTTGTACAAAAAAGCAGGCTATGGAGGAGCGGACGCTG | Tb927.10.15720_rev | GGGGACCACTTTGTACAAGAAAGCTGGGTCTAGTTGGGAGTTTGCTGCGT |
| 3_known | L23 | Tb927.7.2660_for | GGGGACAAGTTTGTACAAAAAAGCAGGCTATGCAGCTCGTAGCCTCGC | Tb927.7.2660_rev | GGGGACCACTTTGTACAAGAAAGCTGGGTTTAATTCCTGCCCCTCCCTTCC |
| 3_known | L24 | Tb927.7.1710_for | GGGGACAAGTTTGTACAAAAAAGCAGGCTATGCGCTTGTTGAAAAAGTCGG | Tb927.7.1710_rev | GGGGACCACTTTGTACAAGAAAGCTGGGTTTACACAACTACACTTGCCGCAG |
| 3_known | M1 | Tb927.3.2440_for | GGGGACAAGTTTGTACAAAAAAGCAGGCTATGCTCAATAGACTATTTGGCCGG | Tb927.3.2440_rev | GGGGACCACTTTGTACAAGAAAGCTGGGTCTAACTACCACCACCCAGATGATTATCA |
| 3_known | M2 | Tb927.1.1330_for | GGGGACAAGTTTGTACAAAAAAGCAGGCTATGAATGTAGCCAAACGAGAGTTTATTAGA | Tb927.1.1330_rev | GGGGACCACTTTGTACAAGAAAGCTGGGTTCATTGGGGTCGGAACGTTTTAAC |
| 3_known | M3 | Tb927.10.3990_for | GGGGACAAGTTTGTACAAAAAAGCAGGCTATGGTAACCGATGATGATTGGAGGG | Tb927.10.3990_rev | GGGGACCACTTTGTACAAGAAAGCTGGGTTTACGCGGTGTACAGCTCAGG |
| 3_known | M4 | Tb927.9.3350_for | GGGGACAAGTTTGTACAAAAAAGCAGGCTATGTTGCATGCCGTTCGCC | Tb927.9.3350_rev | GGGGACCACTTTGTACAAGAAAGCTGGGTTCAGCCCGGTATCAGCGG |
| 3_known | M5 | Tb927.7.220_for | GGGGACAAGTTTGTACAAAAAAGCAGGCTATGAAAAGCAAGTCCAGCAAGCA | Tb927.7.220_rev | GGGGACCACTTTGTACAAGAAAGCTGGGTCTATTCTGTGAGAGACCTATTAAGAGTCGA |
| 3_known | M6 | Tb927.8.5460_for | GGGGACAAGTTTGTACAAAAAAGCAGGCTATGGGTTGCTCTGCATCGAAG | Tb927.8.5460_rev | GGGGACCACTTTGTACAAGAAAGCTGGGTTCAATTACCTTCATTTGCTCCGTTCT |
| 3_known | M7 | Tb927.11.6690_for | GGGGACAAGTTTGTACAAAAAAGCAGGCTATGGAACGAGACGATGGTGCG | Tb927.11.6690_rev | GGGGACCACTTTGTACAAGAAAGCTGGGTCTAAGCATTCGCTCCTGATAGAAAGG |
| 3_known | M8 | Tb927.5.2780_for | GGGGACAAGTTTGTACAAAAAAGCAGGCTATGCTCAGACGAACCTTCTTCCG | Tb927.5.2780_rev | GGGGACCACTTTGTACAAGAAAGCTGGGTCTATGGGTCCCGATTCTCCGG |
| 3_known | M9 | Tb927.10.9740_for | GGGGACAAGTTTGTACAAAAAAGCAGGCTATGGCAACCAAGATGGTGACAC | Tb927.10.9740_rev | GGGGACCACTTTGTACAAGAAAGCTGGGTCTATTTCCACATCTTCTTGAGTGAGACG |
| 3_known | M10 | Tb927.11.760_for | GGGGACAAGTTTGTACAAAAAAGCAGGCTATGGGTGTTTCGCTACCAAAGC | Tb927.11.760_rev | GGGGACCACTTTGTACAAGAAAGCTGGGTTCATTCTTCCGACTGCCCCG |
| 3_known | M11 | Tb927.2.3080_for | GGGGACAAGTTTGTACAAAAAAGCAGGCTATGTTGCCTAAGCAACAGATGGG | Tb927.2.3080_rev | GGGGACCACTTTGTACAAGAAAGCTGGGTTTAACTGCGATAAATGCAGATCCCC |
| 3_known | M12 | Tb927.1.1720_for | GGGGACAAGTTTGTACAAAAAAGCAGGCTATGCAAGCTACACTGGGGTTTG | Tb927.1.1720_rev | GGGGACCACTTTGTACAAGAAAGCTGGGTTTAGTATCGCCTTCGCAGAACCT |
| 3_known | M13 | Tb927.11.14780_for | GGGGACAAGTTTGTACAAAAAAGCAGGCTATGTCGAAGTTAATTAAATTGGATTGTGGC | Tb927.11.14780_rev | GGGGACCACTTTGTACAAGAAAGCTGGGTTCACATATGCGATGCCGAATGG |
| 3_known | M14 | Tb927.8.3770_for | GGGGACAAGTTTGTACAAAAAAGCAGGCTATGGGTTCTAAAGGCGAGGAGC | Tb927.8.3770_rev | GGGGACCACTTTGTACAAGAAAGCTGGGTTTAGGGCATAGAGGTGGGAAGATC |
| 3_known | M15 | Tb927.9.4930_for | GGGGACAAGTTTGTACAAAAAAGCAGGCTATGACTGTGTCTGAATTCGGCAG | Tb927.9.4930_rev | GGGGACCACTTTGTACAAGAAAGCTGGGTCTAGTGCGAAGTGGGCACC |
| 3_known | M16 | Tb927.8.1820_for | GGGGACAAGTTTGTACAAAAAAGCAGGCTATGGCATCTCAGGTTGTGCG | Tb927.8.1820_rev | GGGGACCACTTTGTACAAGAAAGCTGGGTCTACAAGTCACTAATCAAAATGGCAACG |
| 3_known | M17 | Tb927.6.3620_for | GGGGACAAGTTTGTACAAAAAAGCAGGCTATGGGTTTTGAATTTAAGTTGCACCAG | Tb927.6.3620_rev | GGGGACCACTTTGTACAAGAAAGCTGGGTCTAGTGAATCACCTTGTCGATACAGC |
| 3_known | M18 | Tb927.10.15010_for | GGGGACAAGTTTGTACAAAAAAGCAGGCTATGTTTCAGCGATGGTCGTTTGC | Tb927.10.15010_rev | GGGGACCACTTTGTACAAGAAAGCTGGGTCTACTCGGAATCCTTTGGTTCATTCC |
| 3_known | M19 | Tb927.8.1850_for | GGGGACAAGTTTGTACAAAAAAGCAGGCTATGCTCTCAATATGGCGAGCG | Tb927.8.1850_rev | GGGGACCACTTTGTACAAGAAAGCTGGGTCTAACCTACTGAGCCCCTGAGTG |
| 3_known | M20 | Tb927.9.12320_for | GGGGACAAGTTTGTACAAAAAAGCAGGCTATGTCTAAGCAGCAGCATTTGGT | Tb927.9.12320_rev | GGGGACCACTTTGTACAAGAAAGCTGGGTCTAATGGCACATGGCATCCAGC |
| 3_known | M21 | Tb927.7.930_for | GGGGACAAGTTTGTACAAAAAAGCAGGCTATGTCTCGTCGGCGTCCT | Tb927.7.930_rev | GGGGACCACTTTGTACAAGAAAGCTGGGTTCACCCTATTTCAGCATTGGGATGC |
| 3_known | M22 | Tb927.11.4120_for | GGGGACAAGTTTGTACAAAAAAGCAGGCTATGCTAATGAACACCCCGCTAAAG | Tb927.11.4120_rev | GGGGACCACTTTGTACAAGAAAGCTGGGTCTACCACAGCTTCACAGCCAC |
| 3_known | M23 | Tb927.10.5080_for | GGGGACAAGTTTGTACAAAAAAGCAGGCTATGTTGGGCAACAAAGTCGTGT | Tb927.10.5080_rev | GGGGACCACTTTGTACAAGAAAGCTGGGTTCAAGCCTGCTTTTCTTTGGTAATGC |
| 3_known | M24 | Tb927.10.8940_for | GGGGACAAGTTTGTACAAAAAAGCAGGCTATGTCTGCAACCGTCGCAC | Tb927.10.8940_rev | GGGGACCACTTTGTACAAGAAAGCTGGGTTTAACACAGTGAGATTGAACTTCCCC |
| 3_known | N1 | Tb927.6.1780_for | GGGGACAAGTTTGTACAAAAAAGCAGGCTATGTTGACGCATGTGTCCAGTC | Tb927.6.1780_rev | GGGGACCACTTTGTACAAGAAAGCTGGGTTTACCGTCTCGAAGCACTGAAATC |
| 3_known | N2 | Tb927.9.5630_for | GGGGACAAGTTTGTACAAAAAAGCAGGCTATGTTTCAAAAAGCACTTCGTCGAC | Tb927.9.5630_rev | GGGGACCACTTTGTACAAGAAAGCTGGGTCTACACACCTTTAGAGAGTGATTGAGTAAC |
| 3_known | N3 | Tb927.11.1540_for | GGGGACAAGTTTGTACAAAAAAGCAGGCTATGCGCTCGTATCGTTTAGCG | Tb927.11.1540_rev | GGGGACCACTTTGTACAAGAAAGCTGGGTTTAAAGACTGTTGATCAAATCTTTCGTAAT |
| 3_known | N4 | Tb927.10.13290_for | GGGGACAAGTTTGTACAAAAAAGCAGGCTATGGGAGTAGTAGACTTACTCACTTCTACG | Tb927.10.13290_rev | GGGGACCACTTTGTACAAGAAAGCTGGGTTCACTCCACTTTCCTTACGGAGC |
| 3_known | N5 | Tb927.11.900_for | GGGGACAAGTTTGTACAAAAAAGCAGGCTATGAACAAGATTAGTGCTACCGGTG | Tb927.11.900_rev | GGGGACCACTTTGTACAAGAAAGCTGGGTCTACACTTTGCTTTTCTGCATTGCC |
| 3_known | N6 | Tb927.4.2450_for | GGGGACAAGTTTGTACAAAAAAGCAGGCTATGACGATGAAGCGCCCTTC | Tb927.4.2450_rev | GGGGACCACTTTGTACAAGAAAGCTGGGTTTACTTCGGCTGCTCCTTCGT |
| 3_known | N7 | Tb927.9.15490_for | GGGGACAAGTTTGTACAAAAAAGCAGGCTATGCCACCAACTTTACCAAAGGATT | Tb927.9.15490_rev | GGGGACCACTTTGTACAAGAAAGCTGGGTCTAAAGATCATCAAGGAATTCCTTTACTTT |
| 3_known | N8 | Tb927.10.12040_for | GGGGACAAGTTTGTACAAAAAAGCAGGCTATGAACCGTAGTTCGAGGAACCT | Tb927.10.12040_rev | GGGGACCACTTTGTACAAGAAAGCTGGGTTCACTCAACGGAACCCACATATTTG |
| 3_known | N9 | Tb927.11.11590_for | GGGGACAAGTTTGTACAAAAAAGCAGGCTATGACTGAAATGCTGAGCTGTTTGG | Tb927.11.11590_rev | GGGGACCACTTTGTACAAGAAAGCTGGGTTCAGCGCCGACCGTGGTC |
| 3_known | N10 | Tb927.10.11310_for | GGGGACAAGTTTGTACAAAAAAGCAGGCTATGAGCGAAGAGACACAACCCC | Tb927.10.11310_rev | GGGGACCACTTTGTACAAGAAAGCTGGGTTTACATATAATCCGTTTCTTCCCACTCATC |
| 3_known | N11 | Tb927.11.2990_for | GGGGACAAGTTTGTACAAAAAAGCAGGCTATGTTACGTGTTGAAAACTTGCGG | Tb927.11.2990_rev | GGGGACCACTTTGTACAAGAAAGCTGGGTCTATTTTTTCTTTGTGGCAAGGCGG |
| 3_known | N12 | Tb927.11.7060_for | GGGGACAAGTTTGTACAAAAAAGCAGGCTATGAACAACACACACGGTGTGG | Tb927.11.7060_rev | GGGGACCACTTTGTACAAGAAAGCTGGGTTCACAGTTCTCCCCTCTTGTTCA |
| 3_known | N13 | Tb927.6.1460_for | GGGGACAAGTTTGTACAAAAAAGCAGGCTATGAACATACGTACAACTGTGACGC | Tb927.6.1460_rev | GGGGACCACTTTGTACAAGAAAGCTGGGTCTATGAATTTCTTGTCGCAAGGCAC |
| 3_known | N14 | Tb927.6.1960_for | GGGGACAAGTTTGTACAAAAAAGCAGGCTATGGCTCTGTTCCTCTGGCTT | Tb927.6.1960_rev | GGGGACCACTTTGTACAAGAAAGCTGGGTTCACAAACTGAAGAGTGGGGTGA |
| 3_known | N16 | Tb927.11.7320_for | GGGGACAAGTTTGTACAAAAAAGCAGGCTATGAGCCGGTTGTTGTCTCCA | Tb927.11.7320_rev | GGGGACCACTTTGTACAAGAAAGCTGGGTTTACATATGATGCTTTTGATTGTTTAGTTC |
| 3_known | N17 | Tb927.8.3860_for | GGGGACAAGTTTGTACAAAAAAGCAGGCTATGGAGCATGTGGATGCTCGG | Tb927.8.3860_rev | GGGGACCACTTTGTACAAGAAAGCTGGGTTCACAGCGCTGCAGTCAC |
| 3_known | N18 | Tb927.9.5350_for | GGGGACAAGTTTGTACAAAAAAGCAGGCTATGGCCTACCCGGTAGTTGTAAT | Tb927.9.5350_rev | GGGGACCACTTTGTACAAGAAAGCTGGGTCTACTCAAAAACAGAGTGGAACATGTTG |
| 3_known | N19 | Tb927.8.2980_for | GGGGACAAGTTTGTACAAAAAAGCAGGCTATGAGGAGTGCTCAGTTGCCA | Tb927.8.2980_rev | GGGGACCACTTTGTACAAGAAAGCTGGGTTCAACCGCAAGTTGCGCA |
| 3_known | N20 | Tb927.9.2050_for | GGGGACAAGTTTGTACAAAAAAGCAGGCTATGCAACCGCAACCAAACGG | Tb927.9.2050_rev | GGGGACCACTTTGTACAAGAAAGCTGGGTCTAAAATATTGTTAGTGTGGCGAAGACCG |

| | | | | | |
|---|---|---|---|---|---|
| 3_known | N21 | Tb927.6.1230_for | GGGGACAAGTTTGTACAAAAAAGCAGGCTATGCCATCTGTATCGACAGAAGCA | Tb927.6.1230_rev | GGGGACCACTTTGTACAAGAAAGCTGGGTCTATTTTTGCAGCAAGTGTTTTTTTACTAG |
| 3_known | N22 | Tb927.10.2290_for | GGGGACAAGTTTGTACAAAAAAGCAGGCTATGGTCAAGGAGACAGAGTACTACG | Tb927.10.2290_rev | GGGGACCACTTTGTACAAGAAAGCTGGGTCTACTGTTGAGTACACTGATGTGCTG |
| 3_known | N23 | Tb927.11.1410_for | GGGGACAAGTTTGTACAAAAAAGCAGGCTATGTCACATCGAATAAGTTCAACTAACAAC | Tb927.11.1410_rev | GGGGACCACTTTGTACAAGAAAGCTGGGTTTATGTTTTTCGGCTCCAAGAACTCG |
| 3_known | N24 | Tb927.1.3030_for | GGGGACAAGTTTGTACAAAAAAGCAGGCTATGTTGCGTCGCCTCGGT | Tb927.1.3030_rev | GGGGACCACTTTGTACAAGAAAGCTGGGTTCATTCGCTAAAGTCAGGAGACTGA |
| 3_known | O1 | Tb927.10.660_for | GGGGACAAGTTTGTACAAAAAAGCAGGCTATGCTCCGAGTCTCGTGCC | Tb927.10.660_rev | GGGGACCACTTTGTACAAGAAAGCTGGGTTCACTTGCCGTAGAATCCCCTG |
| 3_known | O2 | Tb927.6.4000_for | GGGGACAAGTTTGTACAAAAAAGCAGGCTATGAGTGGAGCAGCTGAAGGT | Tb927.6.4000_rev | GGGGACCACTTTGTACAAGAAAGCTGGGTTTAGGAGGAAGCACCACCGTT |
| 3_known | O3 | Tb927.11.5090_for | GGGGACAAGTTTGTACAAAAAAGCAGGCTATGCTCAATCCCACAACACAGC | Tb927.11.5090_rev | GGGGACCACTTTGTACAAGAAAGCTGGGTTCATTTAGTAACGTTGTGAATAGCTTGGG |
| 3_known | O4 | Tb927.11.16810_for | GGGGACAAGTTTGTACAAAAAAGCAGGCTATGAGTATGCAAGGCGGTGC | Tb927.11.16810_rev | GGGGACCACTTTGTACAAGAAAGCTGGGTTTATTTGTCACCCTTGTCCCGTCT |
| 3_known | O5 | Tb927.3.2900_for | GGGGACAAGTTTGTACAAAAAAGCAGGCTATGGCAGCTTACGGTATAGTGGA | Tb927.3.2900_rev | GGGGACCACTTTGTACAAGAAAGCTGGGTTCAGTCCGTGTCTTCGCCT |
| 3_known | O6 | Tb927.10.8770_for | GGGGACAAGTTTGTACAAAAAAGCAGGCTATGTCGGATGGGGGGAATGTT | Tb927.10.8770_rev | GGGGACCACTTTGTACAAGAAAGCTGGGTTCACACCTGCCCTTCACGT |
| 3_known | O7 | Tb927.10.7410_for | GGGGACAAGTTTGTACAAAAAAGCAGGCTATGTTTACTCGCATTGGTCGTGT | Tb927.10.7410_rev | GGGGACCACTTTGTACAAGAAAGCTGGGTTCACGCGGCGAGTTTGCA |
| 3_known | O8 | Tb927.9.13380_for | GGGGACAAGTTTGTACAAAAAAGCAGGCTATGTCTGAAAATGTCTTTGAGTTCCGT | Tb927.9.13380_rev | GGGGACCACTTTGTACAAGAAAGCTGGGTCTAATCGTCATCAATCAAATGGTCACTG |
| 3_known | O9 | Tb927.7.5880_for | GGGGACAAGTTTGTACAAAAAAGCAGGCTATGCAGCCTGTGCTGGTTG | Tb927.7.5880_rev | GGGGACCACTTTGTACAAGAAAGCTGGGTTTATTTGCCGTAGCGTTTGTCACG |
| 3_known | O10 | Tb927.4.3110_for | GGGGACAAGTTTGTACAAAAAAGCAGGCTATGAGTGGCGTGGAACCCG | Tb927.4.3110_rev | GGGGACCACTTTGTACAAGAAAGCTGGGTTTATTCCACCTGAAACCTTCCCTCC |
| 3_known | O11 | Tb927.10.9860_for | GGGGACAAGTTTGTACAAAAAAGCAGGCTATGCGTCGATGGCTGCGT | Tb927.10.9860_rev | GGGGACCACTTTGTACAAGAAAGCTGGGTTTAGTGGGAGGAAGAGGCGG |
| 3_known | O12 | Tb927.1.1000_for | GGGGACAAGTTTGTACAAAAAAGCAGGCTATGCGTCGCTTATTGTGTCTCTC | Tb927.1.1000_rev | GGGGACCACTTTGTACAAGAAAGCTGGGTTCAAATTTGGATAAAATAATAGACACTGGT |
| 3_known | O13 | Tb927.1.710_for | GGGGACAAGTTTGTACAAAAAAGCAGGCTATGTCACTAAAAGAGAGGAAGAGCATTAAT | Tb927.1.710_rev | GGGGACCACTTTGTACAAGAAAGCTGGGTTTACTCCTTTTCGTCCAATACTGTAACG |
| 3_known | O14 | Tb927.7.6270_for | GGGGACAAGTTTGTACAAAAAAGCAGGCTATGCCAGCAGAGACCATTGAGC | Tb927.7.6270_rev | GGGGACCACTTTGTACAAGAAAGCTGGGTTCACCGTTTCCGCAGCTTG |
| 3_known | O15 | Tb927.10.7100_for | GGGGACAAGTTTGTACAAAAAAGCAGGCTATGAGTTCGGTAAAGAGTAAAGGTTCAAG | Tb927.10.7100_rev | GGGGACCACTTTGTACAAGAAAGCTGGGTTCACAACCGTTTGTCTTCTATGCC |
| 3_known | O16 | Tb927.7.2310_for | GGGGACAAGTTTGTACAAAAAAGCAGGCTATGATAGACGAAACATCGCTACGAG | Tb927.7.2310_rev | GGGGACCACTTTGTACAAGAAAGCTGGGTCTATCTCTCCTCTGCAGGATCTCG |
| 3_known | O17 | Tb927.9.5440_for | GGGGACAAGTTTGTACAAAAAAGCAGGCTATGCGTATCCCGGAAGGCA | Tb927.9.5440_rev | GGGGACCACTTTGTACAAGAAAGCTGGGTTCACAAACCTCGCCGCAG |
| 3_known | O18 | Tb927.9.12450_for | GGGGACAAGTTTGTACAAAAAAGCAGGCTATGCCGGAGGTCGGCACT | Tb927.9.12450_rev | GGGGACCACTTTGTACAAGAAAGCTGGGTTCAATTAGTTAGAGCGAAAAGTTTATTCCT |
| 3_known | O19 | Tb927.11.6620_for | GGGGACAAGTTTGTACAAAAAAGCAGGCTATGCAGGAAGGGAAATCGATAAAACC | Tb927.11.6620_rev | GGGGACCACTTTGTACAAGAAAGCTGGGTCTAGAAGAACTTGTCGTAGTATTCATCATC |
| 3_known | O20 | Tb927.9.5730_for | GGGGACAAGTTTGTACAAAAAAGCAGGCTATGCCCCGTGAGAACTTCTCA | Tb927.9.5730_rev | GGGGACCACTTTGTACAAGAAAGCTGGGTTTACTGCTGCTGGCACTCCG |
| 3_known | O21 | Tb927.11.8260_for | GGGGACAAGTTTGTACAAAAAAGCAGGCTATGAACACCATTTTCCGCCTTTTTC | Tb927.11.8260_rev | GGGGACCACTTTGTACAAGAAAGCTGGGTTTAGAGGGAGGATGTCCCGTACA |
| 3_known | O22 | Tb927.6.3690_for | GGGGACAAGTTTGTACAAAAAAGCAGGCTATGTCTTCTAATTGTCGCACTGAAGAA | Tb927.6.3690_rev | GGGGACCACTTTGTACAAGAAAGCTGGGTTCATGGGGCTTGTAAAGGTGGC |
| 3_known | O23 | Tb927.3.2110_for | GGGGACAAGTTTGTACAAAAAAGCAGGCTATGAACCACGATGCCATGTCATT | Tb927.3.2110_rev | GGGGACCACTTTGTACAAGAAAGCTGGGTTCAGTAATGCGGAGAGTTTTGTCTC |
| 3_known | O24 | Tb927.11.13730_for | GGGGACAAGTTTGTACAAAAAAGCAGGCTATGGACATTGTCGTGAACGATGAC | Tb927.11.13730_rev | GGGGACCACTTTGTACAAGAAAGCTGGGTTTATGATTTTTGACTTTTCAACTCCCGGA |
| 3_known | P1 | Tb927.11.5710_for | GGGGACAAGTTTGTACAAAAAAGCAGGCTATGTTTTTTGTGACGTCAATAGTTCGC | Tb927.11.5710_rev | GGGGACCACTTTGTACAAGAAAGCTGGGTTTATGTGTAGCACCCTGATCTATCGT |
| 3_known | P2 | Tb927.10.5490_for | GGGGACAAGTTTGTACAAAAAAGCAGGCTATGACATCCACCGCGACGG | Tb927.10.5490_rev | GGGGACCACTTTGTACAAGAAAGCTGGGTCTATGCACGAACTCGCTTACGC |
| 3_known | P3 | Tb927.10.16100_for | GGGGACAAGTTTGTACAAAAAAGCAGGCTATGAGCTCACCAGAGAGCAGTG | Tb927.10.16100_rev | GGGGACCACTTTGTACAAGAAAGCTGGGTCTACGAAAACATCTTCTTATATGCCTCACG |
| 3_known | P4 | Tb927.5.1660_for | GGGGACAAGTTTGTACAAAAAAGCAGGCTATGCCACCCAAAAATAGGAATCGG | Tb927.5.1660_rev | GGGGACCACTTTGTACAAGAAAGCTGGGTCTACTTAAGATCCACAATCATTATTGTCAT |
| 3_known | P5 | Tb927.10.11220_for | GGGGACAAGTTTGTACAAAAAAGCAGGCTATGTGCATCGAACAGCTCGTG | Tb927.10.11220_rev | GGGGACCACTTTGTACAAGAAAGCTGGGTCTACACTGTAGGATTTGGTTGGGGG |
| 3_known | P6 | Tb927.7.2400_for | GGGGACAAGTTTGTACAAAAAAGCAGGCTATGTACAAGTACGTTCTTCAGGAAAAATCA | Tb927.7.2400_rev | GGGGACCACTTTGTACAAGAAAGCTGGGTTCACTTTACAATAGCATTTGTTATTCCCGG |
| 3_known | P7 | Tb927.4.1870_for | GGGGACAAGTTTGTACAAAAAAGCAGGCTATGCAAGAAGAGGAAGGGAATTTGG | Tb927.4.1870_rev | GGGGACCACTTTGTACAAGAAAGCTGGGTTCAACTGTCACCTGAAAAGTAATGTGC |
| 3_known | P8 | Tb927.9.14040_for | GGGGACAAGTTTGTACAAAAAAGCAGGCTATGTTGCATGTCGAGTCATCTGAG | Tb927.9.14040_rev | GGGGACCACTTTGTACAAGAAAGCTGGGTTCACAAAAGTTTTACTGCAATAAGTCGC |
| 3_known | P9 | Tb927.4.2020_for | GGGGACAAGTTTGTACAAAAAAGCAGGCTATGATAACCGGCCTTTTCTTCCTTAATAA | Tb927.4.2020_rev | GGGGACCACTTTGTACAAGAAAGCTGGGTTCAAATTGTCCGCACGGCA |
| 3_known | P10 | Tb927.1.1020_for | GGGGACAAGTTTGTACAAAAAAGCAGGCTATGGGTTGTGCAGGGAGTCA | Tb927.1.1020_rev | GGGGACCACTTTGTACAAGAAAGCTGGGTCTATCCCACAACACTCGCAGTATC |
| 3_known | P11 | Tb927.1.820_for | GGGGACAAGTTTGTACAAAAAAGCAGGCTATGATGAGCAATACCACCAGTAGCG | Tb927.1.820_rev | GGGGACCACTTTGTACAAGAAAGCTGGGTTCAAACCACACGTGTGCACG |
| 3_known | P12 | Tb927.3.3930_for | GGGGACAAGTTTGTACAAAAAAGCAGGCTATGTCCCAGGAAGGGGTTCATC | Tb927.3.3930_rev | GGGGACCACTTTGTACAAGAAAGCTGGGTCTACTTTTCATCCAAGTATTGGCGGG |
| 3_known | P13 | Tb927.10.5980_for | GGGGACAAGTTTGTACAAAAAAGCAGGCTATGAGCACATCCAATATAGCGCC | Tb927.10.5980_rev | GGGGACCACTTTGTACAAGAAAGCTGGGTCTAAGTCATGTATTTGCTTCCAGACTTCA |
| 3_known | P14 | Tb927.11.16720_for | GGGGACAAGTTTGTACAAAAAAGCAGGCTATGAATCCCACGGCACTTCGT | Tb927.11.16720_rev | GGGGACCACTTTGTACAAGAAAGCTGGGTTCAAAAGTCAGGTACTTCACTAGGCA |
| 3_known | P15 | Tb927.9.5320_for | GGGGACAAGTTTGTACAAAAAAGCAGGCTATGGTTTTTGCCTTATTGGAGTTGC | Tb927.9.5320_rev | GGGGACCACTTTGTACAAGAAAGCTGGGTTTACCTGAACCCACCGCCG |
| 3_known | P16 | Tb927.10.170_for | GGGGACAAGTTTGTACAAAAAAGCAGGCTATGGTCAGTAAGAAGCTTGGTGAGG | Tb927.10.170_rev | GGGGACCACTTTGTACAAGAAAGCTGGGTTTAGGTGTCATCTGCTTCTGACCT |
| 3_known | P17 | Tb927.10.7020_for | GGGGACAAGTTTGTACAAAAAAGCAGGCTATGAGGAAGTACGCCATATTTTCCTTAG | Tb927.10.7020_rev | GGGGACCACTTTGTACAAGAAAGCTGGGTCTAACGAATGTCCGTGTCATGGT |
| 3_known | P18 | Tb927.9.8950_for | GGGGACAAGTTTGTACAAAAAAGCAGGCTATGGTTGCGTGGGGTGGG | Tb927.9.8950_rev | GGGGACCACTTTGTACAAGAAAGCTGGGTTCATTTCCGTTTCTTATTTTCCTCCTTAAT |
| 3_known | P19 | Tb927.6.1870_for | GGGGACAAGTTTGTACAAAAAAGCAGGCTATGCAAAATTTACGTGCTGATGCG | Tb927.6.1870_rev | GGGGACCACTTTGTACAAGAAAGCTGGGTTCACCAAAGCCGACGATTGC |
| 3_known | P20 | Tb927.7.6830_for | GGGGACAAGTTTGTACAAAAAAGCAGGCTATGGAGGAACTCCACCAACAAATG | Tb927.7.6830_rev | GGGGACCACTTTGTACAAGAAAGCTGGGTTTAGTACACCATCACGAGTTGCCA |
| 3_known | P21 | Tb927.8.4450_for | GGGGACAAGTTTGTACAAAAAAGCAGGCTATGGGGATCAAGGCGAAGAAGG | Tb927.8.4450_rev | GGGGACCACTTTGTACAAGAAAGCTGGGTTTACGACGGGTCGCGCTT |
| 3_known | P22 | Tb927.5.610_for | GGGGACAAGTTTGTACAAAAAAGCAGGCTATGTCGCGTACTATTGTGCTTGG | Tb927.5.610_rev | GGGGACCACTTTGTACAAGAAAGCTGGGTCTAAAGAATATTACCTTCTAAGACGTCAGA |
| 3_known | P23 | Tb927.3.4670_for | GGGGACAAGTTTGTACAAAAAAGCAGGCTATGACAGACGCTGATCGTTCG | Tb927.3.4670_rev | GGGGACCACTTTGTACAAGAAAGCTGGGTCTACGAATCTTTTGTGTCTTTCTCAGCT |
| 3_known | P24 | Tb927.6.2970_for | GGGGACAAGTTTGTACAAAAAAGCAGGCTATGTTGAGTGGCAACACCGAAG | Tb927.6.2970_rev | GGGGACCACTTTGTACAAGAAAGCTGGGTTTAAGTTGGGTTACCACACCGATG |
| 4_hypothetical | A1 | Tb927.1.530_for | GGGGACAAGTTTGTACAAAAAAGCAGGCTATGCGTAGTCGGTACATGCGC | Tb927.1.530_rev | GGGGACCACTTTGTACAAGAAAGCTGGGTTCATAAATTAAGTGGAATATATTCCTTTAC |
| 4_hypothetical | A2 | Tb927.7.2040_for | GGGGACAAGTTTGTACAAAAAAGCAGGCTATGTGGCGCCAGATCCTTCA | Tb927.7.2040_rev | GGGGACCACTTTGTACAAGAAAGCTGGGTTTAATCGAACATATGGAAGCGAGAAGC |
| 4_hypothetical | A3 | Tb927.8.7230_for | GGGGACAAGTTTGTACAAAAAAGCAGGCTATGGAGCTTTCGTTCCCCTTTCT | Tb927.8.7230_rev | GGGGACCACTTTGTACAAGAAAGCTGGGTTTAAACAACCGCATTGTACCCCTG |
| 4_hypothetical | A4 | Tb927.11.4880_for | GGGGACAAGTTTGTACAAAAAAGCAGGCTATGTCGGGTGAGAGTGTCTTCA | Tb927.11.4880_rev | GGGGACCACTTTGTACAAGAAAGCTGGGTCTAATTTCGATCATCGTAAACAAATACATG |
| 4_hypothetical | A5 | Tb927.10.8110_for | GGGGACAAGTTTGTACAAAAAAGCAGGCTATGAGACGGATACAGTGTTGTATTCCG | Tb927.10.8110_rev | GGGGACCACTTTGTACAAGAAAGCTGGGTTCAATTGGTTTTAGCAACAGCTAACG |
| 4_hypothetical | A6 | Tb927.3.5360_for | GGGGACAAGTTTGTACAAAAAAGCAGGCTATGGAGAGGGTAACATACGCTCG | Tb927.3.5360_rev | GGGGACCACTTTGTACAAGAAAGCTGGGTTTAAGCTCGGACAATCAAGTACTTCC |
| 4_hypothetical | A7 | Tb927.4.710_for | GGGGACAAGTTTGTACAAAAAAGCAGGCTATGACTATATTCTACGAGTACCCTCGC | Tb927.4.710_rev | GGGGACCACTTTGTACAAGAAAGCTGGGTTCATCCAAAGGGAGGAAACCCC |
| 4_hypothetical | A8 | Tb927.4.5160_for | GGGGACAAGTTTGTACAAAAAAGCAGGCTATGGAACTTTCGTTCGCCTTTCT | Tb927.4.5160_rev | GGGGACCACTTTGTACAAGAAAGCTGGGTTTAGGCAAGTGCATCATACCCATG |
| 4_hypothetical | A9 | Tb927.11.1970_for | GGGGACAAGTTTGTACAAAAAAGCAGGCTATGGTCGATCGAATTGAAGCAACA | Tb927.11.1970_rev | GGGGACCACTTTGTACAAGAAAGCTGGGTTCACATCTTTCCTCGCCCAAAC |
| 4_hypothetical | A10 | Tb927.9.5000_for | GGGGACAAGTTTGTACAAAAAAGCAGGCTATGCTGATTAGTGCTGCTGATGTG | Tb927.9.5000_rev | GGGGACCACTTTGTACAAGAAAGCTGGGTTCACTCGCCTTGACTTCTACCTT |
| 4_hypothetical | A11 | Tb927.11.13450_for | GGGGACAAGTTTGTACAAAAAAGCAGGCTATGATGAGTAGTGCAACGTTAAGTGAT | Tb927.11.13450_rev | GGGGACCACTTTGTACAAGAAAGCTGGGTTTACCTGGTAACGGTAAAAGGCCT |
| 4_hypothetical | A12 | Tb927.7.4620_for | GGGGACAAGTTTGTACAAAAAAGCAGGCTATGAGGTACCGACTGCGTTTAGA | Tb927.7.4620_rev | GGGGACCACTTTGTACAAGAAAGCTGGGTTCAGAACCATTTCCTGGATCGGT |
| 4_hypothetical | A13 | Tb927.5.1970_for | GGGGACAAGTTTGTACAAAAAAGCAGGCTATGGAGTCATATCCCAACAGCCC | Tb927.5.1970_rev | GGGGACCACTTTGTACAAGAAAGCTGGGTTTAAAAATAGTAAAACTGCCCAATCCCATG |
| 4_hypothetical | A14 | Tb927.10.4290_for | GGGGACAAGTTTGTACAAAAAAGCAGGCTATGCAGAAGCTTGCTGATCGC | Tb927.10.4290_rev | GGGGACCACTTTGTACAAGAAAGCTGGGTTCACTGTTCCACCTCCAGGAG |
| 4_hypothetical | A15 | Tb927.4.1030_for | GGGGACAAGTTTGTACAAAAAAGCAGGCTATGCAGAGCCCGAGTGCG | Tb927.4.1030_rev | GGGGACCACTTTGTACAAGAAAGCTGGGTTCAGTTAATCCGATGCTTCTTCCG |
| 4_hypothetical | A16 | Tb927.10.2330_for | GGGGACAAGTTTGTACAAAAAAGCAGGCTATGGAAAACAAGGACGTTGCAACC | Tb927.10.2330_rev | GGGGACCACTTTGTACAAGAAAGCTGGGTTTACCTGCCACTGTTTTCATTTCCG |
| 4_hypothetical | A17 | Tb927.10.16180_for | GGGGACAAGTTTGTACAAAAAAGCAGGCTATGCTCTTCCCGCACCTTAGT | Tb927.10.16180_rev | GGGGACCACTTTGTACAAGAAAGCTGGGTTTACACAGCAAATGGAAGAATTTTCAAATC |
| 4_hypothetical | A18 | Tb927.6.3660_for | GGGGACAAGTTTGTACAAAAAAGCAGGCTATGGAGTCGTCTGTTGCTTCAC | Tb927.6.3660_rev | GGGGACCACTTTGTACAAGAAAGCTGGGTTCAAGAATTGCTGCTACACGACAC |
| 4_hypothetical | A19 | Tb927.7.3380_for | GGGGACAAGTTTGTACAAAAAAGCAGGCTATGAACCATGATGATGCGATGTGG | Tb927.7.3380_rev | GGGGACCACTTTGTACAAGAAAGCTGGGTTCACCAATCCAAATCAGCCGTG |
| 4_hypothetical | A20 | Tb927.11.4040_for | GGGGACAAGTTTGTACAAAAAAGCAGGCTATGCTTCGAGCTTCTGTGTTGC | Tb927.11.4040_rev | GGGGACCACTTTGTACAAGAAAGCTGGGTCTACTTACCTTTCCCCGTTTTGCA |
| 4_hypothetical | A21 | Tb927.9.2120_for | GGGGACAAGTTTGTACAAAAAAGCAGGCTATGCCCCGTGGTGAACGA | Tb927.9.2120_rev | GGGGACCACTTTGTACAAGAAAGCTGGGTCTAGCGATGTCTTTCCAAGAACATGC |
| 4_hypothetical | A22 | Tb927.2.2490_for | GGGGACAAGTTTGTACAAAAAAGCAGGCTATGTCAAGTCGCTACTTCGCAG | Tb927.2.2490_rev | GGGGACCACTTTGTACAAGAAAGCTGGGTTTAATTGGTATTACGTGCTGCGGA |
| 4_hypothetical | A23 | Tb927.8.1580_for | GGGGACAAGTTTGTACAAAAAAGCAGGCTATGTTACGCTGCGCTACCAG | Tb927.8.1580_rev | GGGGACCACTTTGTACAAGAAAGCTGGGTTCACTGTCCATACGCAATCTGGT |
| 4_hypothetical | A24 | Tb927.10.2630_for | GGGGACAAGTTTGTACAAAAAAGCAGGCTATGAGCAGTTTTTGGTCCAAGAAGT | Tb927.10.2630_rev | GGGGACCACTTTGTACAAGAAAGCTGGGTTCAACCACGTTTAAGCTGAAGTAAGC |
| 4_hypothetical | B1 | Tb927.11.3110_for | GGGGACAAGTTTGTACAAAAAAGCAGGCTATGTTTGTACTACCGGCTATGAAGGA | Tb927.11.3110_rev | GGGGACCACTTTGTACAAGAAAGCTGGGTTCACTGAATGCAACGAAGTTTTCG |
| 4_hypothetical | B2 | Tb927.7.4320_for | GGGGACAAGTTTGTACAAAAAAGCAGGCTATGCTGCAACGTATTCCACCAT | Tb927.7.4320_rev | GGGGACCACTTTGTACAAGAAAGCTGGGTTCACTTCACCGTCAGGAACTGT |
| 4_hypothetical | B3 | Tb927.5.2640_for | GGGGACAAGTTTGTACAAAAAAGCAGGCTATGTCTTACAGATTTTTCGCGTCGA | Tb927.5.2640_rev | GGGGACCACTTTGTACAAGAAAGCTGGGTTTAGGAGATACTCACCGGCTCTTTC |
| 4_hypothetical | B4 | Tb927.11.8800_for | GGGGACAAGTTTGTACAAAAAAGCAGGCTATGATAAATCAAACAGGCTTCGCAAGC | Tb927.11.8800_rev | GGGGACCACTTTGTACAAGAAAGCTGGGTCTACAGGTTCATTAACCCACGATACAC |
| 4_hypothetical | B5 | Tb927.11.4590_for | GGGGACAAGTTTGTACAAAAAAGCAGGCTATGCCATCAGTGTTGGTCTTCCT | Tb927.11.4590_rev | GGGGACCACTTTGTACAAGAAAGCTGGGTTCACGATACTTCAGCGAGCAGC |
| 4_hypothetical | B6 | Tb927.11.8280_for | GGGGACAAGTTTGTACAAAAAAGCAGGCTATGGATGATGTTGGTGACGAAGAGG | Tb927.11.8280_rev | GGGGACCACTTTGTACAAGAAAGCTGGGTCTACACACTCTCGTCGTCACCC |
| 4_hypothetical | B7 | Tb927.11.4270_for | GGGGACAAGTTTGTACAAAAAAGCAGGCTATGTTCGGTTTAGATCGTTTATCCATCTT | Tb927.11.4270_rev | GGGGACCACTTTGTACAAGAAAGCTGGGTTCAAGAATCCAACTTCGCCCCC |
| 4_hypothetical | B8 | Tb927.4.4690_for | GGGGACAAGTTTGTACAAAAAAGCAGGCTATGATTGGAGGGGCGGTGT | Tb927.4.4690_rev | GGGGACCACTTTGTACAAGAAAGCTGGGTTTAGGGATGGATAATCGGCGGAA |
| 4_hypothetical | B9 | Tb927.7.2760_for | GGGGACAAGTTTGTACAAAAAAGCAGGCTATGCCAAGACCTGCCCCA | Tb927.7.2760_rev | GGGGACCACTTTGTACAAGAAAGCTGGGTCTAAATGCCCTCAGCAGTAATAGCG |
| 4_hypothetical | B10 | Tb927.7.6180_for | GGGGACAAGTTTGTACAAAAAAGCAGGCTATGAACGCTGCCAGTTATAGTGG | Tb927.7.6180_rev | GGGGACCACTTTGTACAAGAAAGCTGGGTCTACTCCTGTTGTGTACCCTTTCGG |
| 4_hypothetical | B11 | Tb927.7.7490_for | GGGGACAAGTTTGTACAAAAAAGCAGGCTATGAGTTCCATCCCCGCCA | Tb927.7.7490_rev | GGGGACCACTTTGTACAAGAAAGCTGGGTCTATGATTGTCCCATCCCGCG |
| 4_hypothetical | B12 | Tb927.3.4640_for | GGGGACAAGTTTGTACAAAAAAGCAGGCTATGGACAGCTTTCGCCTCACG | Tb927.3.4640_rev | GGGGACCACTTTGTACAAGAAAGCTGGGTTCACACACCGCTACGACAAGC |
| 4_hypothetical | B13 | Tb927.8.5040_for | GGGGACAAGTTTGTACAAAAAAGCAGGCTATGGATTACCGACAGAAAGCTGC | Tb927.8.5040_rev | GGGGACCACTTTGTACAAGAAAGCTGGGTCTAACACGCCTCGTTATCCTCG |
| 4_hypothetical | B14 | Tb927.9.13770_for | GGGGACAAGTTTGTACAAAAAAGCAGGCTATGGCTTCTGCAGACTCCAAAGC | Tb927.9.13770_rev | GGGGACCACTTTGTACAAGAAAGCTGGGTTCAGTTGTCACTCTCTCCCATTTGC |
| 4_hypothetical | B15 | Tb927.2.4980_for | GGGGACAAGTTTGTACAAAAAAGCAGGCTATGCCCCCCTCGCACACG | Tb927.2.4980_rev | GGGGACCACTTTGTACAAGAAAGCTGGGTTTAACGATTGCCGGTGGCTG |
| 4_hypothetical | B16 | Tb927.10.7590_for | GGGGACAAGTTTGTACAAAAAAGCAGGCTATGGAGCGTTGTCTCTCTCTCG | Tb927.10.7590_rev | GGGGACCACTTTGTACAAGAAAGCTGGGTTCACTTGACCCACGGTGGG |
| 4_hypothetical | B17 | Tb927.7.2260_for | GGGGACAAGTTTGTACAAAAAAGCAGGCTATGAGTGGCGGTGGCCCG | Tb927.7.2260_rev | GGGGACCACTTTGTACAAGAAAGCTGGGTTTAAAGGGTTCTCACAATCACGGTAC |
| 4_hypothetical | B18 | Tb927.11.10780_for | GGGGACAAGTTTGTACAAAAAAGCAGGCTATGGGTCCCAGTGAGTACGATG | Tb927.11.10780_rev | GGGGACCACTTTGTACAAGAAAGCTGGGTTCACCTTCCTGTGAAAACAATACCAA |
| 4_hypothetical | B19 | Tb927.8.5270_for | GGGGACAAGTTTGTACAAAAAAGCAGGCTATGGACGCCACCAATTCGTG | Tb927.8.5270_rev | GGGGACCACTTTGTACAAGAAAGCTGGGTTCATAAATCATGTGAAAAACTACAACAGCC |
| 4_hypothetical | B20 | Tb927.6.3530_for | GGGGACAAGTTTGTACAAAAAAGCAGGCTATGGTTCTCACTACTGTTGTGGGT | Tb927.6.3530_rev | GGGGACCACTTTGTACAAGAAAGCTGGGTTCAATATCCAAAGCCCGCGC |
| 4_hypothetical | B21 | Tb927.9.15040_for | GGGGACAAGTTTGTACAAAAAAGCAGGCTATGAAGCGCGCGCGCAAA | Tb927.9.15040_rev | GGGGACCACTTTGTACAAGAAAGCTGGGTTCACGAAGTTCTGTGGTTGACCA |
| 4_hypothetical | B22 | Tb927.8.8340_for | GGGGACAAGTTTGTACAAAAAAGCAGGCTATGACTTCCTCTTCTTTTCTAGAGGAACT | Tb927.8.8340_rev | GGGGACCACTTTGTACAAGAAAGCTGGGTTTATTCATTAGCAGAAAGGCACTCCAA |
| 4_hypothetical | B23 | Tb927.6.1450_for | GGGGACAAGTTTGTACAAAAAAGCAGGCTATGGCACCATACGTGCGC | Tb927.6.1450_rev | GGGGACCACTTTGTACAAGAAAGCTGGGTCTAAGTGAGGTTCTTACTTCGCTCG |
| 4_hypothetical | B24 | Tb927.11.550_for | GGGGACAAGTTTGTACAAAAAAGCAGGCTATGGCTGCGACACGTGCT | Tb927.11.550_rev | GGGGACCACTTTGTACAAGAAAGCTGGGTTTAGTTGTAGCGGCCGTTGTAAC |
| 4_hypothetical | C1 | Tb927.7.3890_for | GGGGACAAGTTTGTACAAAAAAGCAGGCTATGGAGGGTATGCTTCTGGATGC | Tb927.7.3890_rev | GGGGACCACTTTGTACAAGAAAGCTGGGTTCACAGTATTCCACAAATTATCGACCG |
| 4_hypothetical | C2 | Tb927.9.8520_for | GGGGACAAGTTTGTACAAAAAAGCAGGCTATGAGCACCGCTGCGACT | Tb927.9.8520_rev | GGGGACCACTTTGTACAAGAAAGCTGGGTTTACTCATCATCAAGAAATTTGGATGTTTT |
| 4_hypothetical | C3 | Tb927.3.1150_for | GGGGACAAGTTTGTACAAAAAAGCAGGCTATGGGCAAATACGCTGTTGTCG | Tb927.3.1150_rev | GGGGACCACTTTGTACAAGAAAGCTGGGTTTACCACATCACATCCTCACCGG |
| 4_hypothetical | C4 | Tb927.11.13410_for | GGGGACAAGTTTGTACAAAAAAGCAGGCTATGTGGACACCTCGTTCCGA | Tb927.11.13410_rev | GGGGACCACTTTGTACAAGAAAGCTGGGTTCACACACGTGAAATTATTTGTTGCAC |
| 4_hypothetical | C5 | Tb927.2.2500_for | GGGGACAAGTTTGTACAAAAAAGCAGGCTATGATTCCCACATGCATAGCAATCC | Tb927.2.2500_rev | GGGGACCACTTTGTACAAGAAAGCTGGGTTTAAGTGTATAACGTGCGGCGAA |
| 4_hypothetical | C6 | Tb927.10.5430_for | GGGGACAAGTTTGTACAAAAAAGCAGGCTATGGCACAGTCTTTGGTGTGG | Tb927.10.5430_rev | GGGGACCACTTTGTACAAGAAAGCTGGGTCTATTTTTTTCGCTTACCACCCTTCCC |
| 4_hypothetical | C7 | Tb927.3.800_for | GGGGACAAGTTTGTACAAAAAAGCAGGCTATGTCCGAATCAGAAAAAATTGATGTAAGT | Tb927.3.800_rev | GGGGACCACTTTGTACAAGAAAGCTGGGTTCACACTTCGATGCTAGATCGCA |
| 4_hypothetical | C8 | Tb927.9.7880_for | GGGGACAAGTTTGTACAAAAAAGCAGGCTATGAAAGAGTATGGGACACTTATGTTTTTT | Tb927.9.7880_rev | GGGGACCACTTTGTACAAGAAAGCTGGGTTTACATGTTTTCATTATATTTTGAATTACT |
| 4_hypothetical | C9 | Tb927.11.13370_for | GGGGACAAGTTTGTACAAAAAAGCAGGCTATGCTGCACACGGACCCA | Tb927.11.13370_rev | GGGGACCACTTTGTACAAGAAAGCTGGGTTCAGTACCGACAATTGTGGTCTGT |
| 4_hypothetical | C10 | Tb927.11.11470_for | GGGGACAAGTTTGTACAAAAAAGCAGGCTATGCGTCGTTTCTGCACTGC | Tb927.11.11470_rev | GGGGACCACTTTGTACAAGAAAGCTGGGTTCAATGGGCACCGGCAGG |
| 4_hypothetical | C11 | Tb927.11.5380_for | GGGGACAAGTTTGTACAAAAAAGCAGGCTATGAGATGCAGCCGTGTCACA | Tb927.11.5380_rev | GGGGACCACTTTGTACAAGAAAGCTGGGTTCAGTAGGGATACCGGTGAGAAAC |
| 4_hypothetical | C12 | Tb927.11.9660_for | GGGGACAAGTTTGTACAAAAAAGCAGGCTATGGGAAACGCGCCCTCA | Tb927.11.9660_rev | GGGGACCACTTTGTACAAGAAAGCTGGGTTCAACAAAAAGACTTCACTGACGACT |
| 4_hypothetical | C13 | Tb927.11.8180_for | GGGGACAAGTTTGTACAAAAAAGCAGGCTATGTCACATTTGGGGGGTCACA | Tb927.11.8180_rev | GGGGACCACTTTGTACAAGAAAGCTGGGTTCATGCAAGCGACGCATGA |
| 4_hypothetical | C14 | Tb927.10.8840_for | GGGGACAAGTTTGTACAAAAAAGCAGGCTATGGGCATGCATGAAGGGGC | Tb927.10.8840_rev | GGGGACCACTTTGTACAAGAAAGCTGGGTTCACACAAATTCACGGGGGTG |
| 4_hypothetical | C15 | Tb927.10.1370_for | GGGGACAAGTTTGTACAAAAAAGCAGGCTATGTCCGCCTCCTGCTTCA | Tb927.10.1370_rev | GGGGACCACTTTGTACAAGAAAGCTGGGTTCACCAAGTGCCTAGAGGCAG |
| 4_hypothetical | C16 | Tb927.11.10940_for | GGGGACAAGTTTGTACAAAAAAGCAGGCTATGTCAACCACCTGCACCAAG | Tb927.11.10940_rev | GGGGACCACTTTGTACAAGAAAGCTGGGTCTACTCATACCTCACTGCAGGATAAGT |
| 4_hypothetical | C17 | Tb927.11.15500_for | GGGGACAAGTTTGTACAAAAAAGCAGGCTATGCGTCCGACCTTTCCGG | Tb927.11.15500_rev | GGGGACCACTTTGTACAAGAAAGCTGGGTTTAAGCCGGGTACATATTGTCCTTGA |
| 4_hypothetical | C19 | Tb927.9.2560_for | GGGGACAAGTTTGTACAAAAAAGCAGGCTATGGGGCTTTATGGTGCATATGAC | Tb927.9.2560_rev | GGGGACCACTTTGTACAAGAAAGCTGGGTTTATTCTTTCTTCTTCCTTTTACTGACGGA |
| 4_hypothetical | C20 | Tb927.4.2870_for | GGGGACAAGTTTGTACAAAAAAGCAGGCTATGAACCGCCAGCAACGC | Tb927.4.2870_rev | GGGGACCACTTTGTACAAGAAAGCTGGGTCTATTTTTTGCGTGAGAGGCGC |
| 4_hypothetical | C21 | Tb927.11.15090_for | GGGGACAAGTTTGTACAAAAAAGCAGGCTATGCTTCGTTTCCGTGTCGAA | Tb927.11.15090_rev | GGGGACCACTTTGTACAAGAAAGCTGGGTTTACCCAATTGTTGAATGTGAAGGGT |
| 4_hypothetical | C22 | Tb927.10.7560_for | GGGGACAAGTTTGTACAAAAAAGCAGGCTATGAGTGCCCCTACTGAAAGGG | Tb927.10.7560_rev | GGGGACCACTTTGTACAAGAAAGCTGGGTTCACTGTTCACCCCTCTTACTCTTC |
| 4_hypothetical | C23 | Tb927.11.6310_for | GGGGACAAGTTTGTACAAAAAAGCAGGCTATGCGCACTGGTTTTCGTAGG | Tb927.11.6310_rev | GGGGACCACTTTGTACAAGAAAGCTGGGTTCAAAAAAAGCCATTATCGTCACCGC |
| 4_hypothetical | C24 | Tb927.10.3640_for | GGGGACAAGTTTGTACAAAAAAGCAGGCTATGGAGGGCTTTCGATATTTTACACAG | Tb927.10.3640_rev | GGGGACCACTTTGTACAAGAAAGCTGGGTTCACACAAATACAGAGGCGATACG |
| 4_hypothetical | D1 | Tb927.6.4680_for | GGGGACAAGTTTGTACAAAAAAGCAGGCTATGGAGTCTTTTAAGGGGTTGGCG | Tb927.6.4680_rev | GGGGACCACTTTGTACAAGAAAGCTGGGTTCAAGCTGCCACATGTGTTGT |
| 4_hypothetical | D2 | Tb927.8.2460_for | GGGGACAAGTTTGTACAAAAAAGCAGGCTATGAAAAAGCGAGTGGCGGT | Tb927.8.2460_rev | GGGGACCACTTTGTACAAGAAAGCTGGGTTTAGTTTTCCGCTTCTGCATCCTC |
| 4_hypothetical | D3 | Tb927.3.4520_for | GGGGACAAGTTTGTACAAAAAAGCAGGCTATGCAAATCGCAACCGTGCC | Tb927.3.4520_rev | GGGGACCACTTTGTACAAGAAAGCTGGGTTCAAATAATGGCGGTGAGAGTTGTAG |
| 4_hypothetical | D4 | Tb927.10.8760_for | GGGGACAAGTTTGTACAAAAAAGCAGGCTATGCACATTCATGAAACGATCCCT | Tb927.10.8760_rev | GGGGACCACTTTGTACAAGAAAGCTGGGTTTAACATCGGAGTGCGAAGTAGC |
| 4_hypothetical | D5 | Tb927.9.7940_for | GGGGACAAGTTTGTACAAAAAAGCAGGCTATGCGAGGTGACAATATGAAAGAGTATG | Tb927.9.7940_rev | GGGGACCACTTTGTACAAGAAAGCTGGGTTTACATGTTTTCATTATATTTTGAATGATT |
| 4_hypothetical | D6 | Tb927.6.350_for | GGGGACAAGTTTGTACAAAAAAGCAGGCTATGACGATTCAAGCCGTATCGC | Tb927.6.350_rev | GGGGACCACTTTGTACAAGAAAGCTGGGTTTACATAATCGAGCATCCCGCATTT |
| 4_hypothetical | D7 | Tb927.10.15440_for | GGGGACAAGTTTGTACAAAAAAGCAGGCTATGAAGAATGCACTGGAGCAGC | Tb927.10.15440_rev | GGGGACCACTTTGTACAAGAAAGCTGGGTTTACAATGCCCCCTGTGTACTGG |

| | | | | | |
|---|---|---|---|---|---|
| 4_hypothetical | D8 | Tb927.1.2320_for | GGGGACAAGTTTGTACAAAAAAGCAGGCTATGCGTTGCTGCAGCACG | Tb927.1.2320_rev | GGGGACCACTTTGTACAAGAAAGCTGGGTTCACATATAGAACGTTTTGATGTAATCCCA |
| 4_hypothetical | D10 | Tb927.5.1380_for | GGGGACAAGTTTGTACAAAAAAGCAGGCTATGGCGATACTTCTGACACTCGT | Tb927.5.1380_rev | GGGGACCACTTTGTACAAGAAAGCTGGGTTCAGCCACTAAGAAAAAGTAACAAGCA |
| 4_hypothetical | D11 | Tb927.9.7860_for | GGGGACAAGTTTGTACAAAAAAGCAGGCTATGAAAGGGCGTGGGAAAGC | Tb927.9.7860_rev | GGGGACCACTTTGTACAAGAAAGCTGGGTTTACATGTTTTCATTATATTTTGAATTATT |
| 4_hypothetical | D12 | Tb927.11.9100_for | GGGGACAAGTTTGTACAAAAAAGCAGGCTATGGTCTTTTCAATTCCCTCATATCTTTGC | Tb927.11.9100_rev | GGGGACCACTTTGTACAAGAAAGCTGGGTCTCACAGGCGCCTAAACACGA |
| 4_hypothetical | D13 | Tb927.10.710_for | GGGGACAAGTTTGTACAAAAAAGCAGGCTATGGCTCATTCAAATGTTGTAGTGGC | Tb927.10.710_rev | GGGGACCACTTTGTACAAGAAAGCTGGGTTCAAAGGTCCCCACGGCTT |
| 4_hypothetical | D14 | Tb927.11.11040_for | GGGGACAAGTTTGTACAAAAAAGCAGGCTATGGCAACGTCGGAATCTGAC | Tb927.11.11040_rev | GGGGACCACTTTGTACAAGAAAGCTGGGTTTACACACGTCCCACACTGTTTG |
| 4_hypothetical | D15 | Tb927.8.6850_for | GGGGACAAGTTTGTACAAAAAAGCAGGCTATGACTTCATTTATGCTGGCAACTAACT | Tb927.8.6850_rev | GGGGACCACTTTGTACAAGAAAGCTGGGTCTATCGGAACATCCTTCCAACACAG |
| 4_hypothetical | D17 | Tb927.10.4530_for | GGGGACAAGTTTGTACAAAAAAGCAGGCTATGATGAATGCAAGCTGGAGGG | Tb927.10.4530_rev | GGGGACCACTTTGTACAAGAAAGCTGGGTCTACCGCTTCCCATCCTCCA |
| 4_hypothetical | D18 | Tb927.11.15590_for | GGGGACAAGTTTGTACAAAAAAGCAGGCTATGGGTGACCGTGTGGGC | Tb927.11.15590_rev | GGGGACCACTTTGTACAAGAAAGCTGGGTCTAAGGCCTCTGCCCAGCT |
| 4_hypothetical | D19 | Tb927.6.2190_for | GGGGACAAGTTTGTACAAAAAAGCAGGCTATGCTCCGTGTTGTGTCAAGG | Tb927.6.2190_rev | GGGGACCACTTTGTACAAGAAAGCTGGGTCTACTCGGCGACACACTCAGT |
| 4_hypothetical | D20 | Tb927.11.15510_for | GGGGACAAGTTTGTACAAAAAAGCAGGCTATGGTCAAGCGGTCCCCC | Tb927.11.15510_rev | GGGGACCACTTTGTACAAGAAAGCTGGGTTCAACTCAGCTCGCGTTGC |
| 4_hypothetical | D21 | Tb927.6.3980_for | GGGGACAAGTTTGTACAAAAAAGCAGGCTATGATTATTGACCACGAGCGTCG | Tb927.6.3980_rev | GGGGACCACTTTGTACAAGAAAGCTGGGTCTACCGCGTCTGACCGCG |
| 4_hypothetical | D22 | Tb927.10.4230_for | GGGGACAAGTTTGTACAAAAAAGCAGGCTATGGACAGCTGTATCGTTGACGG | Tb927.10.4230_rev | GGGGACCACTTTGTACAAGAAAGCTGGGTTCAGATAAATCGACGATGTGGCAC |
| 4_hypothetical | D23 | Tb927.10.10160_for | GGGGACAAGTTTGTACAAAAAAGCAGGCTATGGCAATGCTTTCACAGGCT | Tb927.10.10160_rev | GGGGACCACTTTGTACAAGAAAGCTGGGTTTACCCTTTCCCTTCCTCCTCCT |
| 4_hypothetical | D24 | Tb927.9.10070_for | GGGGACAAGTTTGTACAAAAAAGCAGGCTATGGCGAGCATTGGCCCA | Tb927.9.10070_rev | GGGGACCACTTTGTACAAGAAAGCTGGGTTCAGTTCTTGAAAAATACAACTGGTTCTTC |
| 4_hypothetical | E1 | Tb927.10.8870_for | GGGGACAAGTTTGTACAAAAAAGCAGGCTATGTCATCGCTTGCCGAGAGA | Tb927.10.8870_rev | GGGGACCACTTTGTACAAGAAAGCTGGGTCTATGTTGCCGCATCCGTACTC |
| 4_hypothetical | E2 | Tb927.6.3020_for | GGGGACAAGTTTGTACAAAAAAGCAGGCTATGCAACCAAGAAAAACGGTTAAAAAAGT | Tb927.6.3020_rev | GGGGACCACTTTGTACAAGAAAGCTGGGTTCAAGCTCGTTTCTGGCGTATC |
| 4_hypothetical | E3 | Tb927.7.1890_for | GGGGACAAGTTTGTACAAAAAAGCAGGCTATGAATGTTTCTGAGTTTCTAAAAGTTTGG | Tb927.7.1890_rev | GGGGACCACTTTGTACAAGAAAGCTGGGTCTATGACGTAAACCTCTTTCCAAAAAAATC |
| 4_hypothetical | E4 | Tb927.1.4400_for | GGGGACAAGTTTGTACAAAAAAGCAGGCTATGCCCATCACTGGACCGT | Tb927.1.4400_rev | GGGGACCACTTTGTACAAGAAAGCTGGGTTCATTGTTGCACAAACCGGGC |
| 4_hypothetical | E5 | Tb927.10.3680_for | GGGGACAAGTTTGTACAAAAAAGCAGGCTATGGGAGGCAATAAAGTCTCCGT | Tb927.10.3680_rev | GGGGACCACTTTGTACAAGAAAGCTGGGTTTACATAAAAGAGGATTCGTCGTCGC |
| 4_hypothetical | E6 | Tb927.3.1870_for | GGGGACAAGTTTGTACAAAAAAGCAGGCTATGGTGTCTCTAAACCGAAAACAGAAG | Tb927.3.1870_rev | GGGGACCACTTTGTACAAGAAAGCTGGGTTCACAAGTCACCGAAGAAACTCG |
| 4_hypothetical | E7 | Tb927.9.7200_for | GGGGACAAGTTTGTACAAAAAAGCAGGCTATGCTCGCGACCCAAACC | Tb927.9.7200_rev | GGGGACCACTTTGTACAAGAAAGCTGGGTTCAGTGTGTGTGTACGTCGGC |
| 4_hypothetical | E8 | Tb927.10.12050_for | GGGGACAAGTTTGTACAAAAAAGCAGGCTATGGTACTCCCACGCAAGTTGG | Tb927.10.12050_rev | GGGGACCACTTTGTACAAGAAAGCTGGGTCTACATCTTCCTAAAGGGGTGATAGGC |
| 4_hypothetical | E9 | Tb927.10.14250_for | GGGGACAAGTTTGTACAAAAAAGCAGGCTATGGTAGTCTCTCAGATAGGGGTGC | Tb927.10.14250_rev | GGGGACCACTTTGTACAAGAAAGCTGGGTCTACTGAACAAAGTCTGGCATCTCTG |
| 4_hypothetical | E10 | Tb927.11.4560_for | GGGGACAAGTTTGTACAAAAAAGCAGGCTATGATGACGTACCGGCCGT | Tb927.11.4560_rev | GGGGACCACTTTGTACAAGAAAGCTGGGTTTACGGGCCACGGCACAA |
| 4_hypothetical | E11 | Tb927.9.7960_for | GGGGACAAGTTTGTACAAAAAAGCAGGCTATGCGAGGTGACAATATGAAAGAGTATG | Tb927.9.7960_rev | GGGGACCACTTTGTACAAGAAAGCTGGGTTTATTTCCATAACTGTTTACATGCTTCCGT |
| 4_hypothetical | E12 | Tb927.8.670_for | GGGGACAAGTTTGTACAAAAAAGCAGGCTATGTCTCTACTAGTGGCGGGTG | Tb927.8.670_rev | GGGGACCACTTTGTACAAGAAAGCTGGGTTTACGGGGTGGAGAGAGCAG |
| 4_hypothetical | E13 | Tb927.8.3990_for | GGGGACAAGTTTGTACAAAAAAGCAGGCTATGCCTCCCAAGCAACATAAGGA | Tb927.8.3990_rev | GGGGACCACTTTGTACAAGAAAGCTGGGTCTACACTGCGGTTAGCGCAA |
| 4_hypothetical | E14 | Tb927.10.680_for | GGGGACAAGTTTGTACAAAAAAGCAGGCTATGCGTCGTTCATTAGTATTTCCTCC | Tb927.10.680_rev | GGGGACCACTTTGTACAAGAAAGCTGGGTCTATTTTGCAGCAACGCGATGG |
| 4_hypothetical | E15 | Tb927.11.1730_for | GGGGACAAGTTTGTACAAAAAAGCAGGCTATGTGCGAATTCCAGTTTCATGACG | Tb927.11.1730_rev | GGGGACCACTTTGTACAAGAAAGCTGGGTTCAATCGCTCTCATCACTTTCACAA |
| 4_hypothetical | E16 | Tb927.7.5100_for | GGGGACAAGTTTGTACAAAAAAGCAGGCTATGGCCTTTTCACTTCCAAAAAAATATTAC | Tb927.7.5100_rev | GGGGACCACTTTGTACAAGAAAGCTGGGTCTATTCTGGAACCGGAGGCTCC |
| 4_hypothetical | E17 | Tb927.11.12210_for | GGGGACAAGTTTGTACAAAAAAGCAGGCTATGTTCAGTGAACCTGTGTATCTGAC | Tb927.11.12210_rev | GGGGACCACTTTGTACAAGAAAGCTGGGTCTACACCACTGGGGGAGGG |
| 4_hypothetical | E18 | Tb927.9.6320_for | GGGGACAAGTTTGTACAAAAAAGCAGGCTATGAGGTCGACATTACCCTCTGC | Tb927.9.6320_rev | GGGGACCACTTTGTACAAGAAAGCTGGGTTTACGGCGGGCAGTAATGC |
| 4_hypothetical | E19 | Tb927.11.12070_for | GGGGACAAGTTTGTACAAAAAAGCAGGCTATGAACGCTAGGAAGCCACTCG | Tb927.11.12070_rev | GGGGACCACTTTGTACAAGAAAGCTGGGTCTACAACTGAAGACTTTCTTTAAGGTCTTT |
| 4_hypothetical | E20 | Tb927.11.3790_for | GGGGACAAGTTTGTACAAAAAAGCAGGCTATGAAAGCCAGGCGGGCT | Tb927.11.3790_rev | GGGGACCACTTTGTACAAGAAAGCTGGGTTTACAAAAGCACCGAGGCGG |
| 4_hypothetical | E21 | Tb927.11.6390_for | GGGGACAAGTTTGTACAAAAAAGCAGGCTATGCCATCAGATACCACCAAAGAGG | Tb927.11.6390_rev | GGGGACCACTTTGTACAAGAAAGCTGGGTTCAACTGCTGCCGCTCTCT |
| 4_hypothetical | E22 | Tb927.9.6760_for | GGGGACAAGTTTGTACAAAAAAGCAGGCTATGCACAAGCAAGGGTTGACA | Tb927.9.6760_rev | GGGGACCACTTTGTACAAGAAAGCTGGGTTCAGTGGATTCCCACAGGAATC |
| 4_hypothetical | E23 | Tb927.5.3700_for | GGGGACAAGTTTGTACAAAAAAGCAGGCTATGCCCGCAACTATTGTGTCC | Tb927.5.3700_rev | GGGGACCACTTTGTACAAGAAAGCTGGGTTTAAGCATTAGCAGTTTGATCAGCAAC |
| 4_hypothetical | E24 | Tb927.7.4440_for | GGGGACAAGTTTGTACAAAAAAGCAGGCTATGAAGTCGGGTAAGCTATTGATATTTGG | Tb927.7.4440_rev | GGGGACCACTTTGTACAAGAAAGCTGGGTTCATTTATTACTTTGATTGTTATCACCACA |
| 4_hypothetical | F1 | Tb927.5.293b_for | GGGGACAAGTTTGTACAAAAAAGCAGGCTATGGGAGGCGATGAATCGTGC | Tb927.5.293b_rev | GGGGACCACTTTGTACAAGAAAGCTGGGTTCAAATGACACAAAGAAACTTAAATTCTGC |
| 4_hypothetical | F2 | Tb927.11.16080_for | GGGGACAAGTTTGTACAAAAAAGCAGGCTATGTCTAAACCCCCGGAACAGT | Tb927.11.16080_rev | GGGGACCACTTTGTACAAGAAAGCTGGGTCTACATGATGCTTTCACTGCGGC |
| 4_hypothetical | F3 | Tb927.4.1360_for | GGGGACAAGTTTGTACAAAAAAGCAGGCTATGGCGAAATCTCCTATTGTGGC | Tb927.4.1360_rev | GGGGACCACTTTGTACAAGAAAGCTGGGTTTACATTTTAGAACTGGGAAGCACAAGG |
| 4_hypothetical | F4 | Tb927.9.13780_for | GGGGACAAGTTTGTACAAAAAAGCAGGCTATGCTGTGCCGTACCTTTTTACG | Tb927.9.13780_rev | GGGGACCACTTTGTACAAGAAAGCTGGGTCTACGGATAAGGTGCCAGGAAGC |
| 4_hypothetical | F5 | Tb927.9.15020_for | GGGGACAAGTTTGTACAAAAAAGCAGGCTATGTTCGCTACGTTGGGGTCG | Tb927.9.15020_rev | GGGGACCACTTTGTACAAGAAAGCTGGGTTCATGATCGCCTCTTCGAACTCC |
| 4_hypothetical | F6 | Tb927.10.2190_for | GGGGACAAGTTTGTACAAAAAAGCAGGCTATGTTCCACAACAGTTACCAGTCG | Tb927.10.2190_rev | GGGGACCACTTTGTACAAGAAAGCTGGGTCTAAGCGCCCTCTACTGGTGA |
| 4_hypothetical | F7 | Tb927.10.2580_for | GGGGACAAGTTTGTACAAAAAAGCAGGCTATGAATAGGAGCACCTCCCTTGAA | Tb927.10.2580_rev | GGGGACCACTTTGTACAAGAAAGCTGGGTTCAACGTCTTCGATTCTTCCTTTTACC |
| 4_hypothetical | F8 | Tb927.7.2620_for | GGGGACAAGTTTGTACAAAAAAGCAGGCTATGGAAGTGTACCGTCGTATCACT | Tb927.7.2620_rev | GGGGACCACTTTGTACAAGAAAGCTGGGTCTCACCAGCGGTTAATCGCCT |
| 4_hypothetical | F9 | Tb927.6.1190_for | GGGGACAAGTTTGTACAAAAAAGCAGGCTATGGCCGCTACACCTCAGAC | Tb927.6.1190_rev | GGGGACCACTTTGTACAAGAAAGCTGGGTCTACAATGTATATGCCTCAAATGTCTCCA |
| 4_hypothetical | F10 | Tb927.10.2730_for | GGGGACAAGTTTGTACAAAAAAGCAGGCTATGTGGGACGTCGTAAGCCG | Tb927.10.2730_rev | GGGGACCACTTTGTACAAGAAAGCTGGGTTTAAGAGGCCTCGGGTTGGG |
| 4_hypothetical | F11 | Tb927.11.9960_for | GGGGACAAGTTTGTACAAAAAAGCAGGCTATGGTCTTACTGAGTGTTAAGGGTACG | Tb927.11.9960_rev | GGGGACCACTTTGTACAAGAAAGCTGGGTTCAATTCACCTCATGTTCCTCTCGC |
| 4_hypothetical | F12 | Tb927.6.2160_for | GGGGACAAGTTTGTACAAAAAAGCAGGCTATGACCATGAGACGCTTCGGA | Tb927.6.2160_rev | GGGGACCACTTTGTACAAGAAAGCTGGGTTCACGGTGGCAACAGGGT |
| 4_hypothetical | F13 | Tb927.10.1660_for | GGGGACAAGTTTGTACAAAAAAGCAGGCTATGTTTCGTCAGGAACTACGCC | Tb927.10.1660_rev | GGGGACCACTTTGTACAAGAAAGCTGGGTCTACTGGAACGGTACAGATCCATGT |
| 4_hypothetical | F14 | Tb927.10.13750_for | GGGGACAAGTTTGTACAAAAAAGCAGGCTATGCCGGTGCCGTGGCGT | Tb927.10.13750_rev | GGGGACCACTTTGTACAAGAAAGCTGGGTCTAATCGAAGTGGGCAGTTACCC |
| 4_hypothetical | F15 | Tb927.10.6840_for | GGGGACAAGTTTGTACAAAAAAGCAGGCTATGGAGCCACTGGAAAAGCTACA | Tb927.10.6840_rev | GGGGACCACTTTGTACAAGAAAGCTGGGTCTACTCGCCCGGCATTTCC |
| 4_hypothetical | F16 | Tb927.5.2580_for | GGGGACAAGTTTGTACAAAAAAGCAGGCTATGGCAGTAATGAGGTGTATCTTGC | Tb927.5.2580_rev | GGGGACCACTTTGTACAAGAAAGCTGGGTTTAGCGCTTACGGAAGTCATCAAC |
| 4_hypothetical | F17 | Tb927.1.4500_for | GGGGACAAGTTTGTACAAAAAAGCAGGCTATGAAGGGTGCCGTGCCA | Tb927.1.4500_rev | GGGGACCACTTTGTACAAGAAAGCTGGGTTTACGTGTTGCAATCCACTTTCCC |
| 4_hypothetical | F18 | Tb927.8.3460_for | GGGGACAAGTTTGTACAAAAAAGCAGGCTATGGACGGACGGAACGGC | Tb927.8.3460_rev | GGGGACCACTTTGTACAAGAAAGCTGGGTTCAAACAACCTCTTGCGTGCA |
| 4_hypothetical | F19 | Tb927.11.7390_for | GGGGACAAGTTTGTACAAAAAAGCAGGCTATGGCCACAGCTAAAGAACTTGG | Tb927.11.7390_rev | GGGGACCACTTTGTACAAGAAAGCTGGGTCTACTTTTCATCACTCGGCGGTG |
| 4_hypothetical | F20 | Tb927.3.1480_for | GGGGACAAGTTTGTACAAAAAAGCAGGCTATGACGAGTGCTTCAAAAAAATCAACG | Tb927.3.1480_rev | GGGGACCACTTTGTACAAGAAAGCTGGGTTCATTCGTTCTGAACAATCATTGGTAATAC |
| 4_hypothetical | F21 | Tb927.9.7110_for | GGGGACAAGTTTGTACAAAAAAGCAGGCTATGGAGGGAAGCACTGCGC | Tb927.9.7110_rev | GGGGACCACTTTGTACAAGAAAGCTGGGTTTACTGAGTGTACTGAACACCCGG |
| 4_hypothetical | F22 | Tb927.11.9600_for | GGGGACAAGTTTGTACAAAAAAGCAGGCTATGTGGCGGCGTGGAGTT | Tb927.11.9600_rev | GGGGACCACTTTGTACAAGAAAGCTGGGTCTATATGTAATGCGCCTCCAACCAC |
| 4_hypothetical | F23 | Tb927.3.640_for | GGGGACAAGTTTGTACAAAAAAGCAGGCTATGCTAACGATGATGAGGGGT | Tb927.3.640_rev | GGGGACCACTTTGTACAAGAAAGCTGGGTTCACCTCGCTGCACTTGACT |
| 4_hypothetical | F24 | Tb927.10.12550_for | GGGGACAAGTTTGTACAAAAAAGCAGGCTATGGGGCAGGCGTTGACG | Tb927.10.12550_rev | GGGGACCACTTTGTACAAGAAAGCTGGGTTCAGCCCTTTGTATAATCAACTGTGAC |
| 4_hypothetical | G1 | Tb927.3.1980_for | GGGGACAAGTTTGTACAAAAAAGCAGGCTATGTCTACGGGTGCCGAGTT | Tb927.3.1980_rev | GGGGACCACTTTGTACAAGAAAGCTGGGTTCACGGAAAAACAGACGAAGCC |
| 4_hypothetical | G2 | Tb927.8.6130_for | GGGGACAAGTTTGTACAAAAAAGCAGGCTATGGACACAACTACCTCGTCTGA | Tb927.8.6130_rev | GGGGACCACTTTGTACAAGAAAGCTGGGTTCACTCACCATCATCCCGTGG |
| 4_hypothetical | G3 | Tb927.7.4710_for | GGGGACAAGTTTGTACAAAAAAGCAGGCTATGCGTCCGGTAGCACGT | Tb927.7.4710_rev | GGGGACCACTTTGTACAAGAAAGCTGGGTCTCAACTCTCAAATGTGCTGTCAGC |
| 4_hypothetical | G4 | Tb927.11.6380_for | GGGGACAAGTTTGTACAAAAAAGCAGGCTATGCATAGGAGTGGCACTTCCG | Tb927.11.6380_rev | GGGGACCACTTTGTACAAGAAAGCTGGGTCTACACAGAAAGCCACTGGACTC |
| 4_hypothetical | G5 | Tb927.10.10620_for | GGGGACAAGTTTGTACAAAAAAGCAGGCTATGCAGGAGCGAGAGAGTTTGC | Tb927.10.10620_rev | GGGGACCACTTTGTACAAGAAAGCTGGGTTCAACGCCTCTTTTTCGCACA |
| 4_hypothetical | G6 | Tb927.9.7060_for | GGGGACAAGTTTGTACAAAAAAGCAGGCTATGATACTCCCTTCTGCCCCTCC | Tb927.9.7060_rev | GGGGACCACTTTGTACAAGAAAGCTGGGTTCACTGCAGGTTGCGCAAC |
| 4_hypothetical | G7 | Tb927.11.16420_for | GGGGACAAGTTTGTACAAAAAAGCAGGCTATGGTGATGAGGGCGCATCT | Tb927.11.16420_rev | GGGGACCACTTTGTACAAGAAAGCTGGGTTTACAGAAGTCGCTTCTGAAGTTTCTC |
| 4_hypothetical | G8 | Tb927.7.4290_for | GGGGACAAGTTTGTACAAAAAAGCAGGCTATGGCGACGGACGAACGA | Tb927.7.4290_rev | GGGGACCACTTTGTACAAGAAAGCTGGGTTCAGCATATTTTGGCTTGTGAGAAGT |
| 4_hypothetical | G9 | Tb927.6.2950_for | GGGGACAAGTTTGTACAAAAAAGCAGGCTATGGATGGCAAGGAAGATGGTTTAAA | Tb927.6.2950_rev | GGGGACCACTTTGTACAAGAAAGCTGGGTCTAACAAAAGGGCCAGCGGT |
| 4_hypothetical | G10 | Tb927.10.920_for | GGGGACAAGTTTGTACAAAAAAGCAGGCTATGCCGCAGGATAAACGATTAATTATAACT | Tb927.10.920_rev | GGGGACCACTTTGTACAAGAAAGCTGGGTTTACTTCTGTATGTGAACCTCCATGAAGC |
| 4_hypothetical | G11 | Tb927.6.3330_for | GGGGACAAGTTTGTACAAAAAAGCAGGCTATGGGGGGGAAAACAAATGTAAAGG | Tb927.6.3330_rev | GGGGACCACTTTGTACAAGAAAGCTGGGTTTATAGTGGCAATCCCGTAATAATATCATA |
| 4_hypothetical | G12 | Tb927.7.470_for | GGGGACAAGTTTGTACAAAAAAGCAGGCTATGACCCGATATTCACTTGCCCT | Tb927.7.470_rev | GGGGACCACTTTGTACAAGAAAGCTGGGTTCATAGTTGCGTTGGTTTTTGCATT |
| 4_hypothetical | G13 | Tb927.9.5410_for | GGGGACAAGTTTGTACAAAAAAGCAGGCTATGATTTACAGCGATAACAACAATCCACG | Tb927.9.5410_rev | GGGGACCACTTTGTACAAGAAAGCTGGGTTTACCTCCCGTGCAGTGTATTCG |
| 4_hypothetical | G14 | Tb927.7.2280_for | GGGGACAAGTTTGTACAAAAAAGCAGGCTATGGGGCAGAAGGGGAGAATG | Tb927.7.2280_rev | GGGGACCACTTTGTACAAGAAAGCTGGGTTTAGTCCACATCCCGCACACT |
| 4_hypothetical | G15 | Tb927.11.2540_for | GGGGACAAGTTTGTACAAAAAAGCAGGCTATGTCACTGGCTGATATTTTTGCATTTG | Tb927.11.2540_rev | GGGGACCACTTTGTACAAGAAAGCTGGGTCTACAGTTCAAAGGCAATGTCATGGT |
| 4_hypothetical | G16 | Tb927.10.11010_for | GGGGACAAGTTTGTACAAAAAAGCAGGCTATGAAGGAGCGGAGAAGTGTCT | Tb927.10.11010_rev | GGGGACCACTTTGTACAAGAAAGCTGGGTTTATCTTCCGCGCGACTTTTTCA |
| 4_hypothetical | G17 | Tb927.11.9570_for | GGGGACAAGTTTGTACAAAAAAGCAGGCTATGGACGGTCTTTACTTCACCGC | Tb927.11.9570_rev | GGGGACCACTTTGTACAAGAAAGCTGGGTTTATCCCCCTGAGGGCAACTG |
| 4_hypothetical | G18 | Tb927.10.7840_for | GGGGACAAGTTTGTACAAAAAAGCAGGCTATGGATTACAGTGACTTCGATTTTCCG | Tb927.10.7840_rev | GGGGACCACTTTGTACAAGAAAGCTGGGTTTACCAATCAATAATGAAATCGTCGGTGT |
| 4_hypothetical | G19 | Tb927.3.2250_for | GGGGACAAGTTTGTACAAAAAAGCAGGCTATGACCGTGAAGAGGGAGACC | Tb927.3.2250_rev | GGGGACCACTTTGTACAAGAAAGCTGGGTTCAAAGACATGCGGACGACAAA |
| 4_hypothetical | G20 | Tb927.11.3460_for | GGGGACAAGTTTGTACAAAAAAGCAGGCTATGCTGTCAGGTGCAGAGGA | Tb927.11.3460_rev | GGGGACCACTTTGTACAAGAAAGCTGGGTTTAACGATGCGCCCTCGC |
| 4_hypothetical | G21 | Tb927.10.6390_for | GGGGACAAGTTTGTACAAAAAAGCAGGCTATGGCACTCCACCTCTGCC | Tb927.10.6390_rev | GGGGACCACTTTGTACAAGAAAGCTGGGTTCAACATTCCACAGGTTCACAGC |
| 4_hypothetical | G22 | Tb927.7.5710_for | GGGGACAAGTTTGTACAAAAAAGCAGGCTATGTTTACCTCAACTGCTGCGTAC | Tb927.7.5710_rev | GGGGACCACTTTGTACAAGAAAGCTGGGTTTACAACAATAACAGCGACAAAATAACAAT |
| 4_hypothetical | G23 | Tb927.9.4810_for | GGGGACAAGTTTGTACAAAAAAGCAGGCTATGAGACACGTGGGCTTTGC | Tb927.9.4810_rev | GGGGACCACTTTGTACAAGAAAGCTGGGTCTAAACGTCCTCCCGCGTG |
| 4_hypothetical | G24 | Tb927.4.2750_for | GGGGACAAGTTTGTACAAAAAAGCAGGCTATGAGCACAACATCGTCACGTAC | Tb927.4.2750_rev | GGGGACCACTTTGTACAAGAAAGCTGGGTCTACTTTTTTTGGGCTGATCCTTCCG |
| 4_hypothetical | H1 | Tb927.11.6100_for | GGGGACAAGTTTGTACAAAAAAGCAGGCTATGAACTCTTTGATGTTGCAAAAGGC | Tb927.11.6100_rev | GGGGACCACTTTGTACAAGAAAGCTGGGTTCACTCACAACCGGTGTCGG |
| 4_hypothetical | H2 | Tb927.11.4550_for | GGGGACAAGTTTGTACAAAAAAGCAGGCTATGTCGAATGCACTTGAGAGCAT | Tb927.11.4550_rev | GGGGACCACTTTGTACAAGAAAGCTGGGTCTCACGCGTCATCATCTCTGGA |
| 4_hypothetical | H3 | Tb927.11.8450_for | GGGGACAAGTTTGTACAAAAAAGCAGGCTATGCATGGTGGCGCAAACT | Tb927.11.8450_rev | GGGGACCACTTTGTACAAGAAAGCTGGGTCTATGCCCTAAGTGCGAAAACTACA |
| 4_hypothetical | H4 | Tb927.2.2940_for | GGGGACAAGTTTGTACAAAAAAGCAGGCTATGACGGTCGCAACAGCC | Tb927.2.2940_rev | GGGGACCACTTTGTACAAGAAAGCTGGGTTCATAGCTGCACGGCAACATC |
| 4_hypothetical | H5 | Tb927.11.640_for | GGGGACAAGTTTGTACAAAAAAGCAGGCTATGCGAGCCAGTGCTATGACG | Tb927.11.640_rev | GGGGACCACTTTGTACAAGAAAGCTGGGTCTAAATACCATAGTTGAACAGGTGGAGG |
| 4_hypothetical | H6 | Tb927.8.4850_for | GGGGACAAGTTTGTACAAAAAAGCAGGCTATGCTGCAGTGGGAGTGGC | Tb927.8.4850_rev | GGGGACCACTTTGTACAAGAAAGCTGGGTTCATATGTTGAAGATGATACCCTCTTCTAT |
| 4_hypothetical | H7 | Tb927.7.1620_for | GGGGACAAGTTTGTACAAAAAAGCAGGCTATGTCACTTAAAGCGAAGATCGACG | Tb927.7.1620_rev | GGGGACCACTTTGTACAAGAAAGCTGGGTTTACCTGCAATAGTTTATGTTTACACGAGA |
| 4_hypothetical | H8 | Tb927.3.2860_for | GGGGACAAGTTTGTACAAAAAAGCAGGCTATGTTTCGCGGAGTTTTGGCT | Tb927.3.2860_rev | GGGGACCACTTTGTACAAGAAAGCTGGGTTTACACCATTTTGTAACTATATAACCACGG |
| 4_hypothetical | H9 | Tb927.10.9150_for | GGGGACAAGTTTGTACAAAAAAGCAGGCTATGGGGAAAGCACCAGTCAAAG | Tb927.10.9150_rev | GGGGACCACTTTGTACAAGAAAGCTGGGTCTACAAATCAGGGTTCACCTCCG |
| 4_hypothetical | H10 | Tb927.7.6990_for | GGGGACAAGTTTGTACAAAAAAGCAGGCTATGTATCGATACGCGTCGAGTGC | Tb927.7.6990_rev | GGGGACCACTTTGTACAAGAAAGCTGGGTTTACTTTGCTGTTACCAATGCCGC |
| 4_hypothetical | H11 | Tb927.4.860_for | GGGGACAAGTTTGTACAAAAAAGCAGGCTATGGGCCACACCGAACACC | Tb927.4.860_rev | GGGGACCACTTTGTACAAGAAAGCTGGGTTCACAAGAAGAACTCGTCGACAATG |
| 4_hypothetical | H12 | Tb927.10.10180_for | GGGGACAAGTTTGTACAAAAAAGCAGGCTATGGGCGCTGCGGATTCG | Tb927.10.10180_rev | GGGGACCACTTTGTACAAGAAAGCTGGGTTTACGCGGACCTCAGCGA |
| 4_hypothetical | H13 | Tb927.8.4150_for | GGGGACAAGTTTGTACAAAAAAGCAGGCTATGCTGCGGTTGCGGTGT | Tb927.8.4150_rev | GGGGACCACTTTGTACAAGAAAGCTGGGTCTACACGGCACCTGTGGTCA |
| 4_hypothetical | H14 | Tb927.10.4890_for | GGGGACAAGTTTGTACAAAAAAGCAGGCTATGCCGCCTCTCTTTCGCA | Tb927.10.4890_rev | GGGGACCACTTTGTACAAGAAAGCTGGGTCTACGCCCTGCCAAATGCC |
| 4_hypothetical | H15 | Tb927.11.7695_for | GGGGACAAGTTTGTACAAAAAAGCAGGCTATGAGCATTAGTTCCAGTGATGCG | Tb927.11.7695_rev | GGGGACCACTTTGTACAAGAAAGCTGGGTCTAATCAAAATGAAGTCCAAGTCGCTG |
| 4_hypothetical | H16 | Tb927.10.10720_for | GGGGACAAGTTTGTACAAAAAAGCAGGCTATGGCCGGAGGAAAACTGCT | Tb927.10.10720_rev | GGGGACCACTTTGTACAAGAAAGCTGGGTTTAGTACTCGTACTGAAAATATAACGGACT |
| 4_hypothetical | H17 | Tb927.11.15220_for | GGGGACAAGTTTGTACAAAAAAGCAGGCTATGCCGAGGAAAGCGCTTTT | Tb927.11.15220_rev | GGGGACCACTTTGTACAAGAAAGCTGGGTTCACAGCTTGTCCCCGTTGG |
| 4_hypothetical | H18 | Tb927.11.3200_for | GGGGACAAGTTTGTACAAAAAAGCAGGCTATGACCTTTTTCTACGCTGCAGC | Tb927.11.3200_rev | GGGGACCACTTTGTACAAGAAAGCTGGGTTCAAGTAATTTTTCTGTAGATAAAAAACGG |
| 4_hypothetical | H19 | Tb927.10.9810_for | GGGGACAAGTTTGTACAAAAAAGCAGGCTATGGGCAGTAGTAGCAGCAGT | Tb927.10.9810_rev | GGGGACCACTTTGTACAAGAAAGCTGGGTCTACGCGAAAGGAACATTCCAACT |
| 4_hypothetical | H20 | Tb927.4.630_for | GGGGACAAGTTTGTACAAAAAAGCAGGCTATGCAGTTCATTTACTCCTTATTTACGCC | Tb927.4.630_rev | GGGGACCACTTTGTACAAGAAAGCTGGGTCTACTGTTCCTTTAACATATTCTCCACACA |
| 4_hypothetical | H21 | Tb927.1.610_for | GGGGACAAGTTTGTACAAAAAAGCAGGCTATGTGTGATCCCCTCGTTGAGA | Tb927.1.610_rev | GGGGACCACTTTGTACAAGAAAGCTGGGTTCACTTTCCACAACCGTGCTTT |
| 4_hypothetical | H22 | Tb927.3.2940_for | GGGGACAAGTTTGTACAAAAAAGCAGGCTATGCCTCGTACACTTTTTAAACTTTCCA | Tb927.3.2940_rev | GGGGACCACTTTGTACAAGAAAGCTGGGTTCATTTTTCAATGGCTGCGGAGA |
| 4_hypothetical | H23 | Tb927.11.13310_for | GGGGACAAGTTTGTACAAAAAAGCAGGCTATGCCACCTCTGAATAGCCGC | Tb927.11.13310_rev | GGGGACCACTTTGTACAAGAAAGCTGGGTCTCATCTCTTCCCCCTGCCCT |
| 4_hypothetical | H24 | Tb927.11.5820_for | GGGGACAAGTTTGTACAAAAAAGCAGGCTATGCTTCGCCGGCTCAAC | Tb927.11.5820_rev | GGGGACCACTTTGTACAAGAAAGCTGGGTTTAACGGTCACCAGAACCCAGA |
| 4_hypothetical | I1 | Tb927.5.1440_for | GGGGACAAGTTTGTACAAAAAAGCAGGCTATGAACATTGTGGCGAAGAGGG | Tb927.5.1440_rev | GGGGACCACTTTGTACAAGAAAGCTGGGTCTACCTAACAACTTGTTCATTATGAGTTGA |
| 4_hypothetical | I2 | Tb927.4.2550_for | GGGGACAAGTTTGTACAAAAAAGCAGGCTATGTCACTGTGGCCGTTTCG | Tb927.4.2550_rev | GGGGACCACTTTGTACAAGAAAGCTGGGTCTATAAATCTCTGTATTTTTTTGCCCGATT |
| 4_hypothetical | I3 | Tb927.7.420_for | GGGGACAAGTTTGTACAAAAAAGCAGGCTATGACTGCCCGTTTTTTGTGCT | Tb927.7.420_rev | GGGGACCACTTTGTACAAGAAAGCTGGGTTTAAAAATGTGCGGCAGCACTG |
| 4_hypothetical | I5 | Tb927.8.7030_for | GGGGACAAGTTTGTACAAAAAAGCAGGCTATGTATGTCGATTCCGATGAGGAGG | Tb927.8.7030_rev | GGGGACCACTTTGTACAAGAAAGCTGGGTTTAGTCGCTTCCCACTATTTGCAC |
| 4_hypothetical | I6 | Tb927.3.730_for | GGGGACAAGTTTGTACAAAAAAGCAGGCTATGGAAGAGGAAGAGGAAACACGT | Tb927.3.730_rev | GGGGACCACTTTGTACAAGAAAGCTGGGTTTAAACTTGAAAAGAGTAGATGCCCAGC |
| 4_hypothetical | I7 | Tb927.8.600_for | GGGGACAAGTTTGTACAAAAAAGCAGGCTATGAGTTCCAGCTCTTCTTCAACTTC | Tb927.8.600_rev | GGGGACCACTTTGTACAAGAAAGCTGGGTTTAGTTCATCTCACTTTCGCCACG |
| 4_hypothetical | I8 | Tb927.11.7212_for | GGGGACAAGTTTGTACAAAAAAGCAGGCTATGAAACGGTGTAGGCTTGCG | Tb927.11.7212_rev | GGGGACCACTTTGTACAAGAAAGCTGGGTTTAATAACCCTGCTGAATACTTTCCGC |
| 4_hypothetical | I9 | Tb927.10.12970_for | GGGGACAAGTTTGTACAAAAAAGCAGGCTATGCAAGTTGGCGTTTGTGACG | Tb927.10.12970_rev | GGGGACCACTTTGTACAAGAAAGCTGGGTCTCACATACACCAACAGCCACTCT |
| 4_hypothetical | I10 | Tb927.2.3970_for | GGGGACAAGTTTGTACAAAAAAGCAGGCTATGAACGAAAGCAACGGCCG | Tb927.2.3970_rev | GGGGACCACTTTGTACAAGAAAGCTGGGTTTACAGTAACTCATCACAATCCAACTGC |
| 4_hypothetical | I11 | Tb927.10.360_for | GGGGACAAGTTTGTACAAAAAAGCAGGCTATGTTTAGTCAGAAGCCGCCAC | Tb927.10.360_rev | GGGGACCACTTTGTACAAGAAAGCTGGGTTCACGAAGACACCTTAAGCTTAGAGC |
| 4_hypothetical | I12 | Tb927.5.4010_for | GGGGACAAGTTTGTACAAAAAAGCAGGCTATGGAGGATATTTGTCGTGTTTTTAGTCTTG | Tb927.5.4010_rev | GGGGACCACTTTGTACAAGAAAGCTGGGTCAGAATATACTGAGGAGCAAAAAAGAAAC |
| 4_hypothetical | I13 | Tb927.5.3490_for | GGGGACAAGTTTGTACAAAAAAGCAGGCTATGACTGATGTGGCAGAGGGAC | Tb927.5.3490_rev | GGGGACCACTTTGTACAAGAAAGCTGGGTTCATCTATCTATATGATTGGTGTAGTACTC |
| 4_hypothetical | I14 | Tb927.11.8080_for | GGGGACAAGTTTGTACAAAAAAGCAGGCTATGAGAGAGAGAAAACCCATTAGGAAGA | Tb927.11.8080_rev | GGGGACCACTTTGTACAAGAAAGCTGGGTCTACATATCCTCGTCTTTAACACCTGC |
| 4_hypothetical | I15 | Tb927.10.6450_for | GGGGACAAGTTTGTACAAAAAAGCAGGCTATGGATCACCCTCCCGCAG | Tb927.10.6450_rev | GGGGACCACTTTGTACAAGAAAGCTGGGTCTAAATGAATGTCGTAGCGGACATTACG |
| 4_hypothetical | I16 | Tb927.8.6590_for | GGGGACAAGTTTGTACAAAAAAGCAGGCTATGGATGTTAGCGCTGGTGTTG | Tb927.8.6590_rev | GGGGACCACTTTGTACAAGAAAGCTGGGTTTATGACACACAAGAGCCCCGG |
| 4_hypothetical | I17 | Tb927.8.2970_for | GGGGACAAGTTTGTACAAAAAAGCAGGCTATGGAGTTGCATATTTTCGACTTCGA | Tb927.8.2970_rev | GGGGACCACTTTGTACAAGAAAGCTGGGTTTAGCTTCTCTTCCGCAAGCCC |
| 4_hypothetical | I18 | Tb927.10.4670_for | GGGGACAAGTTTGTACAAAAAAGCAGGCTATGTTGCACATACCACGCATGG | Tb927.10.4670_rev | GGGGACCACTTTGTACAAGAAAGCTGGGTCATGTGTCCTCAGTGTTGGGA |
| 4_hypothetical | I19 | Tb927.9.6420_for | GGGGACAAGTTTGTACAAAAAAGCAGGCTATGGTAGGCTCCAGCTTTTCCT | Tb927.9.6420_rev | GGGGACCACTTTGTACAAGAAAGCTGGGTTTACGTTACTTCCTGTGATGTGAACC |
| 4_hypothetical | I20 | Tb927.8.4680_for | GGGGACAAGTTTGTACAAAAAAGCAGGCTATGACTGATCACGAGGGTTCTATAAGG | Tb927.8.4680_rev | GGGGACCACTTTGTACAAGAAAGCTGGGTTTATTCTGTGTGCGTCTGCAGC |

| | | | | | |
|---|---|---|---|---|---|
| 4_hypothetical | I21 | Tb927.8.1970_for | GGGGACAAGTTTGTACAAAAAAGCAGGCTATGATCGGCGAGTCAGAAGCA | Tb927.8.1970_rev | GGGGACCACTTTGTACAAGAAAGCTGGGTCTACAGCGGTAAAACTTGGACACA |
| 4_hypothetical | I22 | Tb927.8.5910_for | GGGGACAAGTTTGTACAAAAAAGCAGGCTATGCTTCCCGTGTCCTGCG | Tb927.8.5910_rev | GGGGACCACTTTGTACAAGAAAGCTGGGTTCAGGCGGTTTCCTCTGACTC |
| 4_hypothetical | I23 | Tb927.5.3890_for | GGGGACAAGTTTGTACAAAAAAGCAGGCTATGATGACTGGGGGAGTCGTG | Tb927.5.3890_rev | GGGGACCACTTTGTACAAGAAAGCTGGGTCTACGAGGCGAGTAGTGAGAAACT |
| 4_hypothetical | I24 | Tb927.3.3060_for | GGGGACAAGTTTGTACAAAAAAGCAGGCTATGGATTTTGATTTTGTTCGTATGCTACAA | Tb927.3.3060_rev | GGGGACCACTTTGTACAAGAAAGCTGGGTCTAAAACGGATCCGCTGAGGTTATG |
| 4_hypothetical | J1 | Tb927.8.4930_for | GGGGACAAGTTTGTACAAAAAAGCAGGCTATGTTTCAGAGGGCTAGATTTCTCCT | Tb927.8.4930_rev | GGGGACCACTTTGTACAAGAAAGCTGGGTTCACAAAACCAAAAGATCTCTGTGAATGC |
| 4_hypothetical | J2 | Tb927.8.2420_for | GGGGACAAGTTTGTACAAAAAAGCAGGCTATGTACGGCCCGCACCAC | Tb927.8.2420_rev | GGGGACCACTTTGTACAAGAAAGCTGGGTTCACATAGAGAAAATGTCATCTTCGATGTA |
| 4_hypothetical | J3 | Tb927.3.890_for | GGGGACAAGTTTGTACAAAAAAGCAGGCTATGAGCGATGATGGAGAGGAAAATAATAG | Tb927.3.890_rev | GGGGACCACTTTGTACAAGAAAGCTGGGTTCAATAGTCGGTGACCCCTGTT |
| 4_hypothetical | J4 | Tb927.3.4590_for | GGGGACAAGTTTGTACAAAAAAGCAGGCTATGGGAGGGGCTGTCGTC | Tb927.3.4590_rev | GGGGACCACTTTGTACAAGAAAGCTGGGTTCAAAACTTTGGAAAGCGTGTGGA |
| 4_hypothetical | J5 | Tb927.9.10790_for | GGGGACAAGTTTGTACAAAAAAGCAGGCTATGTCCACGCTGGCATCATC | Tb927.9.10790_rev | GGGGACCACTTTGTACAAGAAAGCTGGGTTCAGTAACCCAAAAGCCTCGGG |
| 4_hypothetical | J6 | Tb927.7.3700_for | GGGGACAAGTTTGTACAAAAAAGCAGGCTATGACTGAGCAAACCGAAGCTG | Tb927.7.3700_rev | GGGGACCACTTTGTACAAGAAAGCTGGGTCTAAGCAGGGCTTCCATTTTCTGA |
| 4_hypothetical | J7 | Tb927.11.3950_for | GGGGACAAGTTTGTACAAAAAAGCAGGCTATGTTAAAGCATCGAGACAACATATCTTAT | Tb927.11.3950_rev | GGGGACCACTTTGTACAAGAAAGCTGGGTTTAAAAAGAATCACTGCTCTCTACCGC |
| 4_hypothetical | J8 | Tb927.9.7850_for | GGGGACAAGTTTGTACAAAAAAGCAGGCTATGCTATTTGGAGTTTTGCTACTGATGT | Tb927.9.7850_rev | GGGGACCACTTTGTACAAGAAAGCTGGGTTTAAAGAAAAAAGGAAAAAAATGAAATCAC |
| 4_hypothetical | J9 | Tb927.10.9320_for | GGGGACAAGTTTGTACAAAAAAGCAGGCTATGAAATCATTGGAAGATTTGAACGGATG | Tb927.10.9320_rev | GGGGACCACTTTGTACAAGAAAGCTGGGTTTAAAGTTCCCCAATGAACTCGCC |
| 4_hypothetical | J10 | Tb927.3.2590_for | GGGGACAAGTTTGTACAAAAAAGCAGGCTATGTTGACATTTACAGCGTTGTTTCTAG | Tb927.3.2590_rev | GGGGACCACTTTGTACAAGAAAGCTGGGTTTATCCGGTGATAAAGCATACAATTGCG |
| 4_hypothetical | J11 | Tb927.4.3370_for | GGGGACAAGTTTGTACAAAAAAGCAGGCTATGTCAAGCAACACGTGCATAAAC | Tb927.4.3370_rev | GGGGACCACTTTGTACAAGAAAGCTGGGTTCAGCCCTTCAATTCGTCGC |
| 4_hypothetical | J12 | Tb927.5.980_for | GGGGACAAGTTTGTACAAAAAAGCAGGCTATGAAGCGTTTGTTGGACAGCG | Tb927.5.980_rev | GGGGACCACTTTGTACAAGAAAGCTGGGTCTAAACACGCAACATGTCCTCCA |
| 4_hypothetical | J13 | Tb927.5.2610_for | GGGGACAAGTTTGTACAAAAAAGCAGGCTATGAACAGTGACCCCAATGCCT | Tb927.5.2610_rev | GGGGACCACTTTGTACAAGAAAGCTGGGTTTATTTGGTGTGATGTGATGCATGATTG |
| 4_hypothetical | J14 | Tb927.1.3170_for | GGGGACAAGTTTGTACAAAAAAGCAGGCTATGAAGGTTTATTTCACATACGAGGGC | Tb927.1.3170_rev | GGGGACCACTTTGTACAAGAAAGCTGGGTTTACAGCAGGTGCGGCCC |
| 4_hypothetical | J15 | Tb927.1.3150_for | GGGGACAAGTTTGTACAAAAAAGCAGGCTATGCATGTTGTGAGTGAAATTCAGC | Tb927.1.3150_rev | GGGGACCACTTTGTACAAGAAAGCTGGGTTCATGTTCCCCACTGCTGTGC |
| 4_hypothetical | J16 | Tb927.8.7810_for | GGGGACAAGTTTGTACAAAAAAGCAGGCTATGCTCAGTTCCTTGGATCACACG | Tb927.8.7810_rev | GGGGACCACTTTGTACAAGAAAGCTGGGTTCAAGCAACAAACGAACCAACAAC |
| 4_hypothetical | J17 | Tb927.4.1880_for | GGGGACAAGTTTGTACAAAAAAGCAGGCTATGCTGCGACAGCATAGTCCT | Tb927.4.1880_rev | GGGGACCACTTTGTACAAGAAAGCTGGGTCTAGTCATCCCTTCGACTACCAATAGG |
| 4_hypothetical | J18 | Tb927.3.2740_for | GGGGACAAGTTTGTACAAAAAAGCAGGCTATGGTGCCGCTACCGGCC | Tb927.3.2740_rev | GGGGACCACTTTGTACAAGAAAGCTGGGTTTAACGATCTAAGAACCATTTTTCCCAGA |
| 4_hypothetical | J19 | Tb927.1.1300_for | GGGGACAAGTTTGTACAAAAAAGCAGGCTATGACAGAGATTGTGTCGGATAATTACTTT | Tb927.1.1300_rev | GGGGACCACTTTGTACAAGAAAGCTGGGTCTACATGGTGCGTGAAAGTGATATTGG |
| 4_hypothetical | J20 | Tb927.7.5670_for | GGGGACAAGTTTGTACAAAAAAGCAGGCTATGGCTACTAACGTTGCTTCCG | Tb927.7.5670_rev | GGGGACCACTTTGTACAAGAAAGCTGGGTTCACTGCAATCGATTTTTTTTCTTGTTTTT |
| 4_hypothetical | J21 | Tb927.2.5830_for | GGGGACAAGTTTGTACAAAAAAGCAGGCTATGACGACCGCTTCTGTGAGT | Tb927.2.5830_rev | GGGGACCACTTTGTACAAGAAAGCTGGGTTCACTTCCCAATCACTGAGTCCC |
| 4_hypothetical | J22 | Tb927.10.3250_for | GGGGACAAGTTTGTACAAAAAAGCAGGCTATGCGATGTAGTTGCGCTTTTCT | Tb927.10.3250_rev | GGGGACCACTTTGTACAAGAAAGCTGGGTTCAATCTGGAGTCATTCCCGGG |
| 4_hypothetical | J23 | Tb927.9.3600_for | GGGGACAAGTTTGTACAAAAAAGCAGGCTATGAGCAACAAGGCATTCCTAACA | Tb927.9.3600_rev | GGGGACCACTTTGTACAAGAAAGCTGGGTCTACAGGAGGATCCAGTCGTTACA |
| 4_hypothetical | J24 | Tb927.9.1420_for | GGGGACAAGTTTGTACAAAAAAGCAGGCTATGACTCAGCGGTGGATTAAAAGG | Tb927.9.1420_rev | GGGGACCACTTTGTACAAGAAAGCTGGGTTTATGGTGGGGCATGAGACTGA |
| 4_hypothetical | K1 | Tb927.11.9480_for | GGGGACAAGTTTGTACAAAAAAGCAGGCTATGGAAGAAAGGTTTCTTAGAACGCG | Tb927.11.9480_rev | GGGGACCACTTTGTACAAGAAAGCTGGGTTCATACCGCAGCAGCATCCT |
| 4_hypothetical | K2 | Tb927.2.2830_for | GGGGACAAGTTTGTACAAAAAAGCAGGCTATGTTTACCTCGAGCGGTTGTAGC | Tb927.2.2830_rev | GGGGACCACTTTGTACAAGAAAGCTGGGTTTATATAATTTCATACACAATGATCTCCAT |
| 4_hypothetical | K3 | Tb927.8.6120_for | GGGGACAAGTTTGTACAAAAAAGCAGGCTATGTCGTGGTTCGGTTGGGA | Tb927.8.6120_rev | GGGGACCACTTTGTACAAGAAAGCTGGGTTCAGCGGCTGCTGCTTTC |
| 4_hypothetical | K4 | Tb927.3.2100_for | GGGGACAAGTTTGTACAAAAAAGCAGGCTATGACGACTATTACCTCACAGCTGC | Tb927.3.2100_rev | GGGGACCACTTTGTACAAGAAAGCTGGGTTTACAACCCACCAGTAAGTAATATCACAGC |
| 4_hypothetical | K6 | Tb927.7.2990_for | GGGGACAAGTTTGTACAAAAAAGCAGGCTATGTCAAACCCACCTAAGCACTACA | Tb927.7.2990_rev | GGGGACCACTTTGTACAAGAAAGCTGGGTTTACTTCTTTCTGCCACCACCCT |
| 4_hypothetical | K8 | Tb927.6.2560_for | GGGGACAAGTTTGTACAAAAAAGCAGGCTATGCCACGCCATTTACGCTC | Tb927.6.2560_rev | GGGGACCACTTTGTACAAGAAAGCTGGGTTCAGTTGACCTTCTTGCCCTTGG |
| 4_hypothetical | K9 | Tb927.10.4220_for | GGGGACAAGTTTGTACAAAAAAGCAGGCTATGTCGTACCGGCGAAAGGT | Tb927.10.4220_rev | GGGGACCACTTTGTACAAGAAAGCTGGGTCTACGTAAACACTGAAGTACGTTTCGA |
| 4_hypothetical | K10 | Tb927.9.7870_for | GGGGACAAGTTTGTACAAAAAAGCAGGCTATGCGAATACTGTTTGGATTATTGTTGC | Tb927.9.7870_rev | GGGGACCACTTTGTACAAGAAAGCTGGGTTCAAATTAAAAAGCATAAAAATGAAATTAG |
| 4_hypothetical | K11 | Tb927.11.2460_for | GGGGACAAGTTTGTACAAAAAAGCAGGCTATGTTATCGATGCTGGCGGAAG | Tb927.11.2460_rev | GGGGACCACTTTGTACAAGAAAGCTGGGTTCAACTTTGAAACTTCCCTGGTGAAG |
| 4_hypothetical | K12 | Tb927.7.4660_for | GGGGACAAGTTTGTACAAAAAAGCAGGCTATGCGAACACGATTTAGAAACAAGC | Tb927.7.4660_rev | GGGGACCACTTTGTACAAGAAAGCTGGGTTTACTTCCATGGAACACGCCG |
| 4_hypothetical | K13 | Tb927.9.11350_for | GGGGACAAGTTTGTACAAAAAAGCAGGCTATGAAGGTGCGGGTGGCC | Tb927.9.11350_rev | GGGGACCACTTTGTACAAGAAAGCTGGGTTCACGAATCTACCACGTGATCGC |
| 4_hypothetical | K14 | Tb927.8.2490_for | GGGGACAAGTTTGTACAAAAAAGCAGGCTATGCCAGTGTTTGAGAACAAGCA | Tb927.8.2490_rev | GGGGACCACTTTGTACAAGAAAGCTGGGTTCACAAAACGCGTTGCATTCC |
| 4_hypothetical | K15 | Tb06.3A7.960_for | GGGGACAAGTTTGTACAAAAAAGCAGGCTATGGCGACATTCACTTGCGG | Tb06.3A7.960_rev | GGGGACCACTTTGTACAAGAAAGCTGGGTCTATGGGTGCTGGTCAATTAACACA |
| 4_hypothetical | K16 | Tb927.8.2450_for | GGGGACAAGTTTGTACAAAAAAGCAGGCTATGGGACAACACAGCGCCA | Tb927.8.2450_rev | GGGGACCACTTTGTACAAGAAAGCTGGGTTTATTGGAGGGATGCCGTGGG |
| 4_hypothetical | K17 | Tb927.2.1600_for | GGGGACAAGTTTGTACAAAAAAGCAGGCTATGGCATCACCTTCTGATAGCAGA | Tb927.2.1600_rev | GGGGACCACTTTGTACAAGAAAGCTGGGTCTAAATTTGCACAGCGACAAGAATCG |
| 4_hypothetical | K18 | Tb927.11.10460_for | GGGGACAAGTTTGTACAAAAAAGCAGGCTATGTCTGCCGGTAGTGCCG | Tb927.11.10460_rev | GGGGACCACTTTGTACAAGAAAGCTGGGTTTACGTCTGTCTTAGCGTCCGC |
| 4_hypothetical | K19 | Tb927.6.3410_for | GGGGACAAGTTTGTACAAAAAAGCAGGCTATGACACTAAAGAACAAGAGACCGGG | Tb927.6.3410_rev | GGGGACCACTTTGTACAAGAAAGCTGGGTTTACCCGTGGGGAGGGGT |
| 4_hypothetical | K20 | Tb927.7.4540_for | GGGGACAAGTTTGTACAAAAAAGCAGGCTATGTCCATCCACCAAACAAAAGTCT | Tb927.7.4540_rev | GGGGACCACTTTGTACAAGAAAGCTGGGTCTAGGATGACTCCATAACGACGGA |
| 4_hypothetical | K21 | Tb927.9.13340_for | GGGGACAAGTTTGTACAAAAAAGCAGGCTATGTCAAAACTGCGTGACTTATGGT | Tb927.9.13340_rev | GGGGACCACTTTGTACAAGAAAGCTGGGTTTAAAAGGATGTTTCGGGAATCACACC |
| 4_hypothetical | K22 | Tb927.9.7950_for | GGGGACAAGTTTGTACAAAAAAGCAGGCTATGCGAACGATAGTTGGAGTTTTGT | Tb927.9.7950_rev | GGGGACCACTTTGTACAAGAAAGCTGGGTTTAAAGAAAAAAGCATAAAAATGAAATTAC |
| 4_hypothetical | K23 | Tb927.10.14590_for | GGGGACAAGTTTGTACAAAAAAGCAGGCTATGCGGCGGATAGCTTGTG | Tb927.10.14590_rev | GGGGACCACTTTGTACAAGAAAGCTGGGTTTACTTTCCCGCCTTACAACCTGC |
| 4_hypothetical | K24 | Tb927.5.1510_for | GGGGACAAGTTTGTACAAAAAAGCAGGCTATGCGACAATGTGTAGTGCGG | Tb927.5.1510_rev | GGGGACCACTTTGTACAAGAAAGCTGGGTTCAGCTCGTTCCACTGTATGTCG |
| 4_hypothetical | L1 | Tb927.8.7190_for | GGGGACAAGTTTGTACAAAAAAGCAGGCTATGCCCCCACACAGGAAGC | Tb927.8.7190_rev | GGGGACCACTTTGTACAAGAAAGCTGGGTCTAGTGGGCAGCATCAGAATTCTG |
| 4_hypothetical | L2 | Tb927.11.13120_for | GGGGACAAGTTTGTACAAAAAAGCAGGCTATGCGTTTGGAAGAGGTGGATG | Tb927.11.13120_rev | GGGGACCACTTTGTACAAGAAAGCTGGGTTCACAAAAACAGTTGTGAAAACGAGAC |
| 4_hypothetical | L3 | Tb927.4.2270_for | GGGGACAAGTTTGTACAAAAAAGCAGGCTATGGACGAATACGGTGATTTTGACA | Tb927.4.2270_rev | GGGGACCACTTTGTACAAGAAAGCTGGGTTTATATGCAGTCGGCCGCATTG |
| 4_hypothetical | L4 | Tb927.11.3400_for | GGGGACAAGTTTGTACAAAAAAGCAGGCTATGGAGGGAAGAGGGGGAGA | Tb927.11.3400_rev | GGGGACCACTTTGTACAAGAAAGCTGGGTCTAATATGTATAGCCGTTGGCGCG |
| 4_hypothetical | L5 | Tb927.5.1130_for | GGGGACAAGTTTGTACAAAAAAGCAGGCTATGTGCGGAAAGGACAGCG | Tb927.5.1130_rev | GGGGACCACTTTGTACAAGAAAGCTGGGTTTAGTGGTATGCCGGACCTTCC |
| 4_hypothetical | L6 | Tb927.9.9420_for | GGGGACAAGTTTGTACAAAAAAGCAGGCTATGTCGCCTAGGTCGTTGTATGT | Tb927.9.9420_rev | GGGGACCACTTTGTACAAGAAAGCTGGGTTTACTTGCTCAACTCAGAACACCTG |
| 4_hypothetical | L7 | Tb927.9.7900_for | GGGGACAAGTTTGTACAAAAAAGCAGGCTATGCAAATACTATTTGGAGTGTTGTTGC | Tb927.9.7900_rev | GGGGACCACTTTGTACAAGAAAGCTGGGTTCAAATTAAAAAGCATAAAAATGAAATTAG |
| 4_hypothetical | L8 | Tb927.2.2220_for | GGGGACAAGTTTGTACAAAAAAGCAGGCTATGCATTCACATCTACAAAACACGTTTAG | Tb927.2.2220_rev | GGGGACCACTTTGTACAAGAAAGCTGGGTTTAGTTTCGCTGCCGAGAGAGC |
| 4_hypothetical | L9 | Tb927.11.3990_for | GGGGACAAGTTTGTACAAAAAAGCAGGCTATGTTGGATGGGAGGGAATTACGA | Tb927.11.3990_rev | GGGGACCACTTTGTACAAGAAAGCTGGGTTCATTTCTTGGCCTCTGTCTTAATCTTT |
| 4_hypothetical | L10 | Tb927.11.14810_for | GGGGACAAGTTTGTACAAAAAAGCAGGCTATGTTCGCTGAGTTAGACTTCGATG | Tb927.11.14810_rev | GGGGACCACTTTGTACAAGAAAGCTGGGTTCACTTATGATGAATGATGCGTCTCAC |
| 4_hypothetical | L11 | Tb927.3.5370_for | GGGGACAAGTTTGTACAAAAAAGCAGGCTATGCCATTCTGGAAGTATTTCGTAATTACC | Tb927.3.5370_rev | GGGGACCACTTTGTACAAGAAAGCTGGGTTTATAGCGCATCCTTTATCTTGTTAAAGAT |
| 4_hypothetical | L13 | Tb927.11.13830_for | GGGGACAAGTTTGTACAAAAAAGCAGGCTATGTGGCGCTTTCGAGCG | Tb927.11.13830_rev | GGGGACCACTTTGTACAAGAAAGCTGGGTTTAATGGCTCAAAATCCCTTGTCGA |
| 4_hypothetical | L14 | Tb927.4.1690_for | GGGGACAAGTTTGTACAAAAAAGCAGGCTATGTCGGACGTGGATGATTTAATTGA | Tb927.4.1690_rev | GGGGACCACTTTGTACAAGAAAGCTGGGTTTAATGCTTTGGCACGTGTCCC |
| 4_hypothetical | L15 | Tb927.6.1530_for | GGGGACAAGTTTGTACAAAAAAGCAGGCTATGCGGCCCGTGATTGTG | Tb927.6.1530_rev | GGGGACCACTTTGTACAAGAAAGCTGGGTTTAAAAAAGGAAACCAGAACACCACGT |
| 4_hypothetical | L16 | Tb927.8.500_for | GGGGACAAGTTTGTACAAAAAAGCAGGCTATGTGTAAACAGATAGGCGCTATTTTAATA | Tb927.8.500_rev | GGGGACCACTTTGTACAAGAAAGCTGGGTTTAGAAGAAAAGAAAAATAACAATTATCAA |
| 4_hypothetical | L17 | Tb927.10.11050_for | GGGGACAAGTTTGTACAAAAAAGCAGGCTATGAAGATGGGGGCTTCAGCG | Tb927.10.11050_rev | GGGGACCACTTTGTACAAGAAAGCTGGGTTTACTTCGGCACTGGGATACGA |
| 4_hypothetical | L18 | Tb927.8.3190_for | GGGGACAAGTTTGTACAAAAAAGCAGGCTATGCCTCCGCCGCCATTC | Tb927.8.3190_rev | GGGGACCACTTTGTACAAGAAAGCTGGGTTCACTGCATCGAAAATGAAAACCCC |
| 4_hypothetical | L19 | Tb927.10.6820_for | GGGGACAAGTTTGTACAAAAAAGCAGGCTATGGGTTCGACGTCGTCCG | Tb927.10.6820_rev | GGGGACCACTTTGTACAAGAAAGCTGGGTTTAAAATTGTTTCACTAGTTCAACCCAATT |
| 4_hypothetical | L20 | Tb927.11.5030_for | GGGGACAAGTTTGTACAAAAAAGCAGGCTATGGGGCCCCGCATATATCC | Tb927.11.5030_rev | GGGGACCACTTTGTACAAGAAAGCTGGGTTTATATCTTGAAGGTCGCAAATTTATCACC |
| 4_hypothetical | L21 | Tb927.9.14100_for | GGGGACAAGTTTGTACAAAAAAGCAGGCTATGATACCAGCCGCTATGCTCG | Tb927.9.14100_rev | GGGGACCACTTTGTACAAGAAAGCTGGGTTTACGCGGAATTCAACTCCTCCC |
| 4_hypothetical | L22 | Tb927.8.6600_for | GGGGACAAGTTTGTACAAAAAAGCAGGCTATGAATGAAGGTTCTAGCAGAGCG | Tb927.8.6600_rev | GGGGACCACTTTGTACAAGAAAGCTGGGTTCAAGAACAGCCTTGTCTACACGT |
| 4_hypothetical | L23 | Tb927.10.15760_for | GGGGACAAGTTTGTACAAAAAAGCAGGCTATGAAATCAGAGGGGACGCCG | Tb927.10.15760_rev | GGGGACCACTTTGTACAAGAAAGCTGGGTTTACGATTGCTCAGCAGCGG |
| 4_hypothetical | L24 | Tb927.10.5190_for | GGGGACAAGTTTGTACAAAAAAGCAGGCTATGGGCTCAATGGTAATCGGAGG | Tb927.10.5190_rev | GGGGACCACTTTGTACAAGAAAGCTGGGTTCAGTTAGTGGACTCCTCCAATCTAG |
| 4_hypothetical | M1 | Tb927.4.1570_for | GGGGACAAGTTTGTACAAAAAAGCAGGCTATGCCTTTAGACTCACCAGTGGG | Tb927.4.1570_rev | GGGGACCACTTTGTACAAGAAAGCTGGGTTCACAGTTCCAAATCCTTGGTTAAGAG |
| 4_hypothetical | M2 | Tb927.9.15260_for | GGGGACAAGTTTGTACAAAAAAGCAGGCTATGCCGGACGTTGCCGAA | Tb927.9.15260_rev | GGGGACCACTTTGTACAAGAAAGCTGGGTTCACAGGCGACGCATCATTAC |
| 4_hypothetical | M3 | Tb927.8.7070_for | GGGGACAAGTTTGTACAAAAAAGCAGGCTATGTGGGAAGCCGCTTGC | Tb927.8.7070_rev | GGGGACCACTTTGTACAAGAAAGCTGGGTTCAGTTCACACTCTGCGACCG |
| 4_hypothetical | M4 | Tb927.11.12060_for | GGGGACAAGTTTGTACAAAAAAGCAGGCTATGAAGAACACTAATAACCTCCGCG | Tb927.11.12060_rev | GGGGACCACTTTGTACAAGAAAGCTGGGTTTAGCGGCCATACGTCCAAAC |
| 4_hypothetical | M5 | Tb927.6.370_for | GGGGACAAGTTTGTACAAAAAAGCAGGCTATGGGCACTGAGATAAGCATGACG | Tb927.6.370_rev | GGGGACCACTTTGTACAAGAAAGCTGGGTTCAGCACACAATTGCGAGGG |
| 4_hypothetical | M6 | Tb927.7.7140_for | GGGGACAAGTTTGTACAAAAAAGCAGGCTATGTCTTTAGCAGATGAAGTCCCCC | Tb927.7.7140_rev | GGGGACCACTTTGTACAAGAAAGCTGGGTTTAACGTTGAGCGAGCAGGTC |
| 4_hypothetical | M8 | Tb927.9.11750_for | GGGGACAAGTTTGTACAAAAAAGCAGGCTATGCCTGATAAGATTGAAAATAATACCCGC | Tb927.9.11750_rev | GGGGACCACTTTGTACAAGAAAGCTGGGTTCACTTTTGGGGCATGTTCCTG |
| 4_hypothetical | M9 | Tb927.11.220_for | GGGGACAAGTTTGTACAAAAAAGCAGGCTATGATTGAAGCGGACGTCATACG | Tb927.11.220_rev | GGGGACCACTTTGTACAAGAAAGCTGGGTTCACATCAACGACCTGGTGGG |
| 4_hypothetical | M10 | Tb927.10.11890_for | GGGGACAAGTTTGTACAAAAAAGCAGGCTATGGTGGTGGGACACCGG | Tb927.10.11890_rev | GGGGACCACTTTGTACAAGAAAGCTGGGTTTACTCCGACAAGCCCTTTGGC |
| 4_hypothetical | M11 | Tb927.11.10920_for | GGGGACAAGTTTGTACAAAAAAGCAGGCTATGTGGCTTTTCTTTGGAACTTTGTG | Tb927.11.10920_rev | GGGGACCACTTTGTACAAGAAAGCTGGGTTCAGTCTTTCCGGTGACTGGAC |
| 4_hypothetical | M12 | Tb927.4.600_for | GGGGACAAGTTTGTACAAAAAAGCAGGCTATGAATGTAATCAATCGCATGCTTTCG | Tb927.4.600_rev | GGGGACCACTTTGTACAAGAAAGCTGGGTCTATTCCTCCCCACCTGTTGTCT |
| 4_hypothetical | M13 | Tb927.7.1020_for | GGGGACAAGTTTGTACAAAAAAGCAGGCTATGTCCTCCGGCGAACTGC | Tb927.7.1020_rev | GGGGACCACTTTGTACAAGAAAGCTGGGTCTAGTCGGAGTCGCTGTGAGT |
| 4_hypothetical | M14 | Tb927.5.3950_for | GGGGACAAGTTTGTACAAAAAAGCAGGCTATGTACAGCTATAACAACATTATTGAAAAA | Tb927.5.3950_rev | GGGGACCACTTTGTACAAGAAAGCTGGGTTCACAGTTGGTATAACTCTGCTACTCC |
| 4_hypothetical | M15 | Tb927.11.1810_for | GGGGACAAGTTTGTACAAAAAAGCAGGCTATGATCGCCGATAAGTGTGGC | Tb927.11.1810_rev | GGGGACCACTTTGTACAAGAAAGCTGGGTTTACCGTTTAAAACAGCAAAATACCCTACT |
| 4_hypothetical | M16 | Tb927.4.2130_for | GGGGACAAGTTTGTACAAAAAAGCAGGCTATGGGTAGTGACGTTGATTTGAGGA | Tb927.4.2130_rev | GGGGACCACTTTGTACAAGAAAGCTGGGTTCACCGAATCACGCGTGC |
| 4_hypothetical | M17 | Tb927.1.2210_for | GGGGACAAGTTTGTACAAAAAAGCAGGCTATGGGTAGTCCTGCTGCGC | Tb927.1.2210_rev | GGGGACCACTTTGTACAAGAAAGCTGGGTTCATTCGTGTTTGTTTGGCTTTTCG |
| 4_hypothetical | M18 | Tb927.11.7970_for | GGGGACAAGTTTGTACAAAAAAGCAGGCTATGCTTGCTGGATTGTTTTATAAACAGATT | Tb927.11.7970_rev | GGGGACCACTTTGTACAAGAAAGCTGGGTTCAGTTTTCCGATATTTGTTCAATGTGC |
| 4_hypothetical | M19 | Tb927.10.9110_for | GGGGACAAGTTTGTACAAAAAAGCAGGCTATGTGGCAACGCCTCAGGT | Tb927.10.9110_rev | GGGGACCACTTTGTACAAGAAAGCTGGGTCTAAGTTTGATGGGAAGTCCACGAA |
| 4_hypothetical | M20 | Tb927.3.5210_for | GGGGACAAGTTTGTACAAAAAAGCAGGCTATGTTGAGGCTTATCAACAGAGGAAAC | Tb927.3.5210_rev | GGGGACCACTTTGTACAAGAAAGCTGGGTCTACGAATCTGCGTCCTTTACATCTT |
| 4_hypothetical | M21 | Tb927.11.15210_for | GGGGACAAGTTTGTACAAAAAAGCAGGCTATGTATGCACGACTGAGTAACACTTC | Tb927.11.15210_rev | GGGGACCACTTTGTACAAGAAAGCTGGGTCTACAAGTCGAAATTCATGAACACTGAATC |
| 4_hypothetical | M22 | Tb927.1.2640_for | GGGGACAAGTTTGTACAAAAAAGCAGGCTATGAAGTCAGAGAAAGCAGAGAAGGC | Tb927.1.2640_rev | GGGGACCACTTTGTACAAGAAAGCTGGGTTTAATACAACACTTCCGTACGACTGG |
| 4_hypothetical | M23 | Tb927.1.4700_for | GGGGACAAGTTTGTACAAAAAAGCAGGCTATGCATACCGATGACAAGCAGG | Tb927.1.4700_rev | GGGGACCACTTTGTACAAGAAAGCTGGGTTCACACGGCATCAGCGGA |
| 4_hypothetical | M24 | Tb927.4.4610_for | GGGGACAAGTTTGTACAAAAAAGCAGGCTATGTTTACTGTATCTTGTAATGTTGCCTTC | Tb927.4.4610_rev | GGGGACCACTTTGTACAAGAAAGCTGGGTTTACCTATGAAGACCCTGCATCGT |
| 4_hypothetical | N1 | Tb927.2.4150_for | GGGGACAAGTTTGTACAAAAAAGCAGGCTATGAAGTCTGTGGCCTTTCATTTTCT | Tb927.2.4150_rev | GGGGACCACTTTGTACAAGAAAGCTGGGTTTATGATATATGTGACAAATGTTCTTCTGT |
| 4_hypothetical | N2 | Tb927.7.4690_for | GGGGACAAGTTTGTACAAAAAAGCAGGCTATGATCTTGATTTATTACGTATATCTTAGC | Tb927.7.4690_rev | GGGGACCACTTTGTACAAGAAAGCTGGGTTCAAGAGTCCGCGAAAGCTTG |
| 4_hypothetical | N3 | Tb927.10.6960_for | GGGGACAAGTTTGTACAAAAAAGCAGGCTATGGTAACACCCGAGCAAGCT | Tb927.10.6960_rev | GGGGACCACTTTGTACAAGAAAGCTGGGTTCAAATGTTCATTGCACTAGCCACA |
| 4_hypothetical | N4 | Tb927.10.6920_for | GGGGACAAGTTTGTACAAAAAAGCAGGCTATGGTAACACCCGAGCAAGCT | Tb927.10.6920_rev | GGGGACCACTTTGTACAAGAAAGCTGGGTTCAAATGTTCATTGCACTGCCCA |
| 4_hypothetical | N5 | Tb927.7.5700_for | GGGGACAAGTTTGTACAAAAAAGCAGGCTATGTCATCAACACTCCTTGCAACA | Tb927.7.5700_rev | GGGGACCACTTTGTACAAGAAAGCTGGGTTCATCGACGCCCCTCCCT |
| 4_hypothetical | N6 | Tb927.7.3970_for | GGGGACAAGTTTGTACAAAAAAGCAGGCTATGACGTTGTCGTCCGGGA | Tb927.7.3970_rev | GGGGACCACTTTGTACAAGAAAGCTGGGTTTAAAAAACATAGGGCGGTGGCA |
| 4_hypothetical | N7 | Tb927.10.6570_for | GGGGACAAGTTTGTACAAAAAAGCAGGCTATGCGAGCTGCGAAGGAAC | Tb927.10.6570_rev | GGGGACCACTTTGTACAAGAAAGCTGGGTTCACCACATCGTACTCTCATACTTATAAGA |
| 4_hypothetical | N8 | Tb927.4.5220_for | GGGGACAAGTTTGTACAAAAAAGCAGGCTATGTTTCCCAATGCACACTACCC | Tb927.4.5220_rev | GGGGACCACTTTGTACAAGAAAGCTGGGTTTACCCTGTGTTGATCGCCGC |
| 4_hypothetical | N9 | Tb927.3.3890_for | GGGGACAAGTTTGTACAAAAAAGCAGGCTATGTGGGTGAGGACATTCCTGC | Tb927.3.3890_rev | GGGGACCACTTTGTACAAGAAAGCTGGGTTCACGGTGCCGCATGGCA |
| 4_hypothetical | N10 | Tb927.7.3920_for | GGGGACAAGTTTGTACAAAAAAGCAGGCTATGACCGAGGGGCGTTTTGA | Tb927.7.3920_rev | GGGGACCACTTTGTACAAGAAAGCTGGGTTTATTTGAGCCTTGCGGGCC |
| 4_hypothetical | N11 | Tb927.4.2360_for | GGGGACAAGTTTGTACAAAAAAGCAGGCTATGTTCCTGCGGCGTCTGA | Tb927.4.2360_rev | GGGGACCACTTTGTACAAGAAAGCTGGGTCTACACTTTATCTCCCTCCTCTTTTTCTG |
| 4_hypothetical | N12 | Tb927.8.2990_for | GGGGACAAGTTTGTACAAAAAAGCAGGCTATGTACCCGCTGAGTGCCA | Tb927.8.2990_rev | GGGGACCACTTTGTACAAGAAAGCTGGGTTCACGAGCACTTAACAAACTCAGTG |
| 4_hypothetical | N13 | Tb927.7.1680_for | GGGGACAAGTTTGTACAAAAAAGCAGGCTATGGAAGAGGAGGTAAACACAACGT | Tb927.7.1680_rev | GGGGACCACTTTGTACAAGAAAGCTGGGTTCACATGAACGGTAAAATACCTTTCTCTAT |
| 4_hypothetical | N14 | Tb927.3.4160_for | GGGGACAAGTTTGTACAAAAAAGCAGGCTATGAAGCTGATAGCGAAACTCGG | Tb927.3.4160_rev | GGGGACCACTTTGTACAAGAAAGCTGGGTTTATAACTGCGGCGGGTTCTCG |
| 4_hypothetical | N15 | Tb927.11.11960_for | GGGGACAAGTTTGTACAAAAAAGCAGGCTATGATGATATGGCTGCAACGCA | Tb927.11.11960_rev | GGGGACCACTTTGTACAAGAAAGCTGGGTTTAATAAGCGGGGGGGCCC |
| 4_hypothetical | N16 | Tb927.7.1450_for | GGGGACAAGTTTGTACAAAAAAGCAGGCTATGTTCGGAGAGCTGGTCTGC | Tb927.7.1450_rev | GGGGACCACTTTGTACAAGAAAGCTGGGTTCATAGAAAGTTGCCGATGCAATCA |
| 4_hypothetical | N17 | Tb927.11.3940_for | GGGGACAAGTTTGTACAAAAAAGCAGGCTATGCCGCTCCCCACTGAC | Tb927.11.3940_rev | GGGGACCACTTTGTACAAGAAAGCTGGGTCTACAGTTCAAATCTTTCCGCTGCC |
| 4_hypothetical | N18 | Tb927.11.5900_for | GGGGACAAGTTTGTACAAAAAAGCAGGCTATGGAAACCAGTGGCCCTAAGA | Tb927.11.5900_rev | GGGGACCACTTTGTACAAGAAAGCTGGGTTTATTTGTAGGAGTGACTCTTCTTAGCAGC |
| 4_hypothetical | N19 | Tb927.5.2550_for | GGGGACAAGTTTGTACAAAAAAGCAGGCTATGCTTCGCCGGTTAGCATG | Tb927.5.2550_rev | GGGGACCACTTTGTACAAGAAAGCTGGGTTTACACATACGCCGCCCACA |
| 4_hypothetical | N20 | Tb927.7.6260_for | GGGGACAAGTTTGTACAAAAAAGCAGGCTATGAAAGGGCCAACCGACAAC | Tb927.7.6260_rev | GGGGACCACTTTGTACAAGAAAGCTGGGTTTAATCATCACGATGTAGCATCGCC |
| 4_hypothetical | N21 | Tb927.3.1970_for | GGGGACAAGTTTGTACAAAAAAGCAGGCTATGAGTGCCGAGGGACTGAG | Tb927.3.1970_rev | GGGGACCACTTTGTACAAGAAAGCTGGGTTCATCTTCTCCTCCCCTCAAGTTTG |
| 4_hypothetical | N22 | Tb927.4.3800_for | GGGGACAAGTTTGTACAAAAAAGCAGGCTATGGAGAAACGGGTGATACCAACA | Tb927.4.3800_rev | GGGGACCACTTTGTACAAGAAAGCTGGGTTCATGACAGAAGAAATAGTATTTGCGTCT |
| 4_hypothetical | N23 | Tb927.9.11820_for | GGGGACAAGTTTGTACAAAAAAGCAGGCTATGGGTAGCAAGGGCGACG | Tb927.9.11820_rev | GGGGACCACTTTGTACAAGAAAGCTGGGTTTACCGACTCTTCTGTTCACTGCT |
| 4_hypothetical | N24 | Tb927.5.4360_for | GGGGACAAGTTTGTACAAAAAAGCAGGCTATGTTTTCATATTCGTTCAAGGCGC | Tb927.5.4360_rev | GGGGACCACTTTGTACAAGAAAGCTGGGTTCACCCCTCCGCTGGAAATG |
| 4_hypothetical | O1 | Tb927.3.3260_for | GGGGACAAGTTTGTACAAAAAAGCAGGCTATGAGCTTTCACGATGACAATCATTTCG | Tb927.3.3260_rev | GGGGACCACTTTGTACAAGAAAGCTGGGTTCACGCAGAACCTGCGAGA |
| 4_hypothetical | O2 | Tb927.6.3960_for | GGGGACAAGTTTGTACAAAAAAGCAGGCTATGACGAGACATGAAAATGTGGAGC | Tb927.6.3960_rev | GGGGACCACTTTGTACAAGAAAGCTGGGTTTATGTCAATCTTGATTCCTTCGCTGA |
| 4_hypothetical | O3 | Tb927.10.5520_for | GGGGACAAGTTTGTACAAAAAAGCAGGCTATGACATACTCACGCCGCGC | Tb927.10.5520_rev | GGGGACCACTTTGTACAAGAAAGCTGGGTTCATGGCGCAGATTTATCACCG |
| 4_hypothetical | O4 | Tb927.11.13850_for | GGGGACAAGTTTGTACAAAAAAGCAGGCTATGTACTCATGTAAGACGGCCCA | Tb927.11.13850_rev | GGGGACCACTTTGTACAAGAAAGCTGGGTTTATTCGGTTGGCTTACTTAGCAACT |
| 4_hypothetical | O5 | Tb927.6.4700_for | GGGGACAAGTTTGTACAAAAAAGCAGGCTATGAGATCGTCTCCCCGTTTAGC | Tb927.6.4700_rev | GGGGACCACTTTGTACAAGAAAGCTGGGTCTATGCAAGCCCCGATCGC |
| 4_hypothetical | O6 | Tb927.3.4870_for | GGGGACAAGTTTGTACAAAAAAGCAGGCTATGGGAGTAGTCAAAGTGTGGTTGC | Tb927.3.4870_rev | GGGGACCACTTTGTACAAGAAAGCTGGGTCTACCATTGATCTTTAGGCGAAGGTATG |
| 4_hypothetical | O7 | Tb927.6.3250_for | GGGGACAAGTTTGTACAAAAAAGCAGGCTATGTCGGATTTAGAGAAACTACGTGC | Tb927.6.3250_rev | GGGGACCACTTTGTACAAGAAAGCTGGGTTCATGCCCCGCCCTTTGC |
| 4_hypothetical | O8 | Tb927.6.4260_for | GGGGACAAGTTTGTACAAAAAAGCAGGCTATGTTCGTTTTTTTCGTAGCCCTCG | Tb927.6.4260_rev | GGGGACCACTTTGTACAAGAAAGCTGGGTCTATGGAGTCGAGTTTGTTTCATCTCC |
| 4_hypothetical | O9 | Tb927.10.610_for | GGGGACAAGTTTGTACAAAAAAGCAGGCTATGGTAGTGAATACACGCGTGTC | Tb927.10.610_rev | GGGGACCACTTTGTACAAGAAAGCTGGGTCTACGTGGCCTTTGGTGCAG |
| 4_hypothetical | O10 | Tb927.10.3580_for | GGGGACAAGTTTGTACAAAAAAGCAGGCTATGTCGTTTCGCTATACGAACAACT | Tb927.10.3580_rev | GGGGACCACTTTGTACAAGAAAGCTGGGTCTACTGCGTAAACCGGAGTTGC |

| | | | | | |
|---|---|---|---|---|---|
| 4_hypothetical | O11 | Tb927.6.1990_for | GGGGACAAGTTTGTACAAAAAAGCAGGCTATGTTGAGAGATGGTGACCGTTATACC | Tb927.6.1990_rev | GGGGACCACTTTGTACAAGAAAGCTGGGTTTAAGCAAAGTCAACCTTGACTTGCG |
| 4_hypothetical | O12 | Tb927.11.10130_for | GGGGACAAGTTTGTACAAAAAAGCAGGCTATGACAGACATGGAACTTGTGGAAG | Tb927.11.10130_rev | GGGGACCACTTTGTACAAGAAAGCTGGGTTCAACGGTAAAAATGTCCGCCA |
| 4_hypothetical | O13 | Tb927.10.9260_for | GGGGACAAGTTTGTACAAAAAAGCAGGCTATGACTTCGCTCAACACAGAGC | Tb927.10.9260_rev | GGGGACCACTTTGTACAAGAAAGCTGGGTTCACCACCACTTCGACAGCG |
| 4_hypothetical | O14 | Tb927.4.950_for | GGGGACAAGTTTGTACAAAAAAGCAGGCTATGCGGGGGCAACGGAGC | Tb927.4.950_rev | GGGGACCACTTTGTACAAGAAAGCTGGGTTCAAAAACGGTACCTCAGTAACCTCC |
| 4_hypothetical | O15 | Tb927.10.7120_for | GGGGACAAGTTTGTACAAAAAAGCAGGCTATGCCTTTAAGGACCGATTACGC | Tb927.10.7120_rev | GGGGACCACTTTGTACAAGAAAGCTGGGTCTAGTAGTCATATACGATTCGATGACAAAA |
| 4_hypothetical | O16 | Tb927.10.11960_for | GGGGACAAGTTTGTACAAAAAAGCAGGCTATGAAGTACCTGGAGAATACCACTACTATT | Tb927.10.11960_rev | GGGGACCACTTTGTACAAGAAAGCTGGGTTTAACTGTGTTGCCCATTTACTTGAGC |
| 4_hypothetical | O17 | Tb927.6.4510_for | GGGGACAAGTTTGTACAAAAAAGCAGGCTATGAACACATCAATTCGTCTGCTTTCG | Tb927.6.4510_rev | GGGGACCACTTTGTACAAGAAAGCTGGGTTCACAACTTGAGTGCTTCCAACC |
| 4_hypothetical | O18 | Tb927.11.9690_for | GGGGACAAGTTTGTACAAAAAAGCAGGCTATGCTAGACAAAGTTGTAGGCGATC | Tb927.11.9690_rev | GGGGACCACTTTGTACAAGAAAGCTGGGTTCACCGGACAGTAAGAGGAAAGT |
| 4_hypothetical | O19 | Tb927.9.11560_for | GGGGACAAGTTTGTACAAAAAAGCAGGCTATGTGGCTTTTCTCACGAAGTAATGA | Tb927.9.11560_rev | GGGGACCACTTTGTACAAGAAAGCTGGGTTCACGAGTACCTTCCACGCA |
| 4_hypothetical | O20 | Tb927.3.1810_for | GGGGACAAGTTTGTACAAAAAAGCAGGCTATGCGAACAACCACGCATGG | Tb927.3.1810_rev | GGGGACCACTTTGTACAAGAAAGCTGGGTTCACTGGGTGTGCTGTGGC |
| 4_hypothetical | O21 | Tb927.5.2040_for | GGGGACAAGTTTGTACAAAAAAGCAGGCTATGGCGGCACTACGCTATTTC | Tb927.5.2040_rev | GGGGACCACTTTGTACAAGAAAGCTGGGTTTAAATGCGGTTAAGTAAACAACGGTTG |
| 4_hypothetical | O22 | Tb927.5.1460_for | GGGGACAAGTTTGTACAAAAAAGCAGGCTATGCCCGGTACCGACGAA | Tb927.5.1460_rev | GGGGACCACTTTGTACAAGAAAGCTGGGTTTACGATGCTCCATTCTTTACAAAGAAATT |
| 4_hypothetical | O23 | Tb927.10.10400_for | GGGGACAAGTTTGTACAAAAAAGCAGGCTATGATGCAGAAATCAGTCACCTTCTTAC | Tb927.10.10400_rev | GGGGACCACTTTGTACAAGAAAGCTGGGTTTACTTGAAGACGCCACGAGGC |
| 4_hypothetical | O24 | Tb927.2.4750_for | GGGGACAAGTTTGTACAAAAAAGCAGGCTATGAAGGGTGGTGGGGGC | Tb927.2.4750_rev | GGGGACCACTTTGTACAAGAAAGCTGGGTCTACTCAATTATACCACACAGAAACGACA |
| 4_hypothetical | P2 | Tb927.8.660_for | GGGGACAAGTTTGTACAAAAAAGCAGGCTATGTCCGGACATCGTGCCT | Tb927.8.660_rev | GGGGACCACTTTGTACAAGAAAGCTGGGTTTACTTACAGCGGAGAAGCCGT |
| 4_hypothetical | P3 | Tb927.9.10680_for | GGGGACAAGTTTGTACAAAAAAGCAGGCTATGCCACCTAAGGTCCCCAAC | Tb927.9.10680_rev | GGGGACCACTTTGTACAAGAAAGCTGGGTTCAGTGCGCGTCGACGTT |
| 4_hypothetical | P4 | Tb927.11.1890_for | GGGGACAAGTTTGTACAAAAAAGCAGGCTATGAACAACAGACATTCGGGATCC | Tb927.11.1890_rev | GGGGACCACTTTGTACAAGAAAGCTGGGTTTAGTTCTCGTTATCGTATGTCTCGATTAT |
| 4_hypothetical | P5 | Tb927.7.5360_for | GGGGACAAGTTTGTACAAAAAAGCAGGCTATGAGTTCCCTCTTGAAGGTTCATCA | Tb927.7.5360_rev | GGGGACCACTTTGTACAAGAAAGCTGGGTTCAAGTCACGCACCGCGC |
| 4_hypothetical | P6 | Tb927.10.9390_for | GGGGACAAGTTTGTACAAAAAAGCAGGCTATGTTTTTTCTAACCCGCTTCAAACTG | Tb927.10.9390_rev | GGGGACCACTTTGTACAAGAAAGCTGGGTCTACGGCTTTGTAAAGTACACAGCA |
| 4_hypothetical | P7 | Tb927.7.340_for | GGGGACAAGTTTGTACAAAAAAGCAGGCTATGGGGAATGTGAGGGCGC | Tb927.7.340_rev | GGGGACCACTTTGTACAAGAAAGCTGGGTCTAGGAAGTTTGCAGTGTCTCCAAT |
| 4_hypothetical | P8 | Tb927.11.1750_for | GGGGACAAGTTTGTACAAAAAAGCAGGCTATGTCTCTGCAAGCTGTTGGC | Tb927.11.1750_rev | GGGGACCACTTTGTACAAGAAAGCTGGGTTTAGAAGGTTGGGGGCGGC |
| 4_hypothetical | P9 | Tb927.4.4770_for | GGGGACAAGTTTGTACAAAAAAGCAGGCTATGCCGATTTCCATTCTTTACGTGA | Tb927.4.4770_rev | GGGGACCACTTTGTACAAGAAAGCTGGGTTTACCCCGCCGCGTATTCT |
| 4_hypothetical | P10 | Tb927.10.4210_for | GGGGACAAGTTTGTACAAAAAAGCAGGCTATGGAAAGCAGATGCCTGAGTAGC | Tb927.10.4210_rev | GGGGACCACTTTGTACAAGAAAGCTGGGTTTACTCTTCTAGGGACATGGTCAGAAG |
| 4_hypothetical | P11 | Tb927.7.3310_for | GGGGACAAGTTTGTACAAAAAAGCAGGCTATGTTTTACACGGGCTGCAGC | Tb927.7.3310_rev | GGGGACCACTTTGTACAAGAAAGCTGGGTCTACAGAGAATCCTCTACAATATTAAGACA |
| 4_hypothetical | P12 | Tb927.11.14160_for | GGGGACAAGTTTGTACAAAAAAGCAGGCTATGCCGCTTGGTCCCATTG | Tb927.11.14160_rev | GGGGACCACTTTGTACAAGAAAGCTGGGTTCATTTGACAGGTTTACGAATGCTGC |
| 4_hypothetical | P13 | Tb927.7.1250_for | GGGGACAAGTTTGTACAAAAAAGCAGGCTATGCAACGCCGCTTGCAG | Tb927.7.1250_rev | GGGGACCACTTTGTACAAGAAAGCTGGGTTTATGATGTTCTCACACATGACGAAAATTT |
| 4_hypothetical | P15 | Tb927.4.2940_for | GGGGACAAGTTTGTACAAAAAAGCAGGCTATGAAAAAATGTTGGGTATTCGACTGC | Tb927.4.2940_rev | GGGGACCACTTTGTACAAGAAAGCTGGGTTCACAAAAGGCTAACCTCATCGC |
| 4_hypothetical | P16 | Tb927.11.2080_for | GGGGACAAGTTTGTACAAAAAAGCAGGCTATGAATGATTGGATTACTAAGGCTTTGGA | Tb927.11.2080_rev | GGGGACCACTTTGTACAAGAAAGCTGGGTTCACGGGGGCGGGTGCTC |
| 4_hypothetical | P17 | Tb927.8.6230_for | GGGGACAAGTTTGTACAAAAAAGCAGGCTATGGTCCGCTGGGGTGGT | Tb927.8.6230_rev | GGGGACCACTTTGTACAAGAAAGCTGGGTCTACGGATGTCCACCGCACA |
| 4_hypothetical | P18 | Tb927.11.10950_for | GGGGACAAGTTTGTACAAAAAAGCAGGCTATGTTTCGTTTCAACCAAACCCTCC | Tb927.11.10950_rev | GGGGACCACTTTGTACAAGAAAGCTGGGTTCACGTCTTTTTGTAATTCCCAGTGA |
| 4_hypothetical | P19 | Tb927.4.4090_for | GGGGACAAGTTTGTACAAAAAAGCAGGCTATGACTCCAGCTGCTTCTCGC | Tb927.4.4090_rev | GGGGACCACTTTGTACAAGAAAGCTGGGTTTACCGTTCGTATACGTACGATTTTGG |
| 4_hypothetical | P21 | Tb927.7.4360_for | GGGGACAAGTTTGTACAAAAAAGCAGGCTATGGGTTTCTTTAAAAAGCGTTATGGTG | Tb927.7.4360_rev | GGGGACCACTTTGTACAAGAAAGCTGGGTCTAGGTTGCCAGTCGGTGTTG |
| 4_hypothetical | P22 | Tb927.4.4280_for | GGGGACAAGTTTGTACAAAAAAGCAGGCTATGGAAACGGAGAGCAAGACAAC | Tb927.4.4280_rev | GGGGACCACTTTGTACAAGAAAGCTGGGTTCATTGTCCTGCAGCGGC |
| 4_hypothetical | P23 | Tb927.11.5420_for | GGGGACAAGTTTGTACAAAAAAGCAGGCTATGACCCAAGCTGTAGAAATGGC | Tb927.11.5420_rev | GGGGACCACTTTGTACAAGAAAGCTGGGTTCAATTCTTTACTTGTGCTTGACCCT |
| 4_known | A1 | Tb927.10.11590_for | GGGGACAAGTTTGTACAAAAAAGCAGGCTATGAAGCCAATTATAAAGTCGATCCTTGA | Tb927.10.11590_rev | GGGGACCACTTTGTACAAGAAAGCTGGGTTCATTGTACCTGTGTTGGGTGGT |
| 4_known | A2 | Tb927.11.14910_for | GGGGACAAGTTTGTACAAAAAAGCAGGCTATGAAAGGCGCGATGCCTC | Tb927.11.14910_rev | GGGGACCACTTTGTACAAGAAAGCTGGGTTCAACTCTCCTCCCCATCTTCTG |
| 4_known | A3 | Tb927.10.2890_for | GGGGACAAGTTTGTACAAAAAAGCAGGCTATGACGATCCAGAAAGTTCACGG | Tb927.10.2890_rev | GGGGACCACTTTGTACAAGAAAGCTGGGTTTACGACCAACCGGGAAAGCC |
| 4_known | A4 | Tb927.9.13280_for | GGGGACAAGTTTGTACAAAAAAGCAGGCTATGTCGCTCACCGCTGAGC | Tb927.9.13280_rev | GGGGACCACTTTGTACAAGAAAGCTGGGTCTAGTGTCCCTTCTCCGGGTTC |
| 4_known | A5 | Tb927.10.11970_for | GGGGACAAGTTTGTACAAAAAAGCAGGCTATGCAACAGTCAATTCGGCTTGC | Tb927.10.11970_rev | GGGGACCACTTTGTACAAGAAAGCTGGGTCTAATGGGCAGGAAGCAGGTACT |
| 4_known | A6 | Tb927.11.7890_for | GGGGACAAGTTTGTACAAAAAAGCAGGCTATGTCAATGCACAGCGTAGAGG | Tb927.11.7890_rev | GGGGACCACTTTGTACAAGAAAGCTGGGTTCAGCTGGATCTCCCGTCCC |
| 4_known | A7 | Tb927.6.3300_for | GGGGACAAGTTTGTACAAAAAAGCAGGCTATGGAACTGCAGTCGCTTATAGACA | Tb927.6.3300_rev | GGGGACCACTTTGTACAAGAAAGCTGGGTCTAAGCTACTTTGATCGACTTCGCT |
| 4_known | A8 | Tb927.9.8160_for | GGGGACAAGTTTGTACAAAAAAGCAGGCTATGCTGGGTCGAAAATGCAGTG | Tb927.9.8160_rev | GGGGACCACTTTGTACAAGAAAGCTGGGTTCACACCACTTCATCAGCCCC |
| 4_known | A9 | Tb927.1.1310_for | GGGGACAAGTTTGTACAAAAAAGCAGGCTATGTCACTTCTAGTTGCTCTCGTACG | Tb927.1.1310_rev | GGGGACCACTTTGTACAAGAAAGCTGGGTTCACAATGCACCGCGCCC |
| 4_known | A10 | Tb927.7.4730_for | GGGGACAAGTTTGTACAAAAAAGCAGGCTATGCTTCGTAGGGGTGAACGC | Tb927.7.4730_rev | GGGGACCACTTTGTACAAGAAAGCTGGGTTCACTCACCGACTGCCCC |
| 4_known | A11 | Tb927.9.1540_for | GGGGACAAGTTTGTACAAAAAAGCAGGCTATGTCGAGTGGTGCTTCACAC | Tb927.9.1540_rev | GGGGACCACTTTGTACAAGAAAGCTGGGTTCAAGTAACTATATGATGCATCGCACAA |
| 4_known | A12 | Tb927.8.6930_for | GGGGACAAGTTTGTACAAAAAAGCAGGCTATGGCCGAACCGTTCTCGA | Tb927.8.6930_rev | GGGGACCACTTTGTACAAGAAAGCTGGGTTTAGGCGACACCTATACCACGT |
| 4_known | A13 | Tb927.4.2700_for | GGGGACAAGTTTGTACAAAAAAGCAGGCTATGTTTCGAACGTTACCCTTGAGG | Tb927.4.2700_rev | GGGGACCACTTTGTACAAGAAAGCTGGGTTCAGCAGGAAGATGCTTCAACG |
| 4_known | A15 | Tb927.11.2090_for | GGGGACAAGTTTGTACAAAAAAGCAGGCTATGGAGGTGGCTGTGGGG | Tb927.11.2090_rev | GGGGACCACTTTGTACAAGAAAGCTGGGTTTATGAAGATGCACTAAACTCCTTCCTC |
| 4_known | A16 | Tb927.9.8250_for | GGGGACAAGTTTGTACAAAAAAGCAGGCTATGAAAAGTCTTCACTCGCCCCC | Tb927.9.8250_rev | GGGGACCACTTTGTACAAGAAAGCTGGGTTCAAACGGAATTCCTGCGCC |
| 4_known | A17 | Tb927.8.6000_for | GGGGACAAGTTTGTACAAAAAAGCAGGCTATGAGCGAAGCAATGGAAAGCA | Tb927.8.6000_rev | GGGGACCACTTTGTACAAGAAAGCTGGGTTCACTCATTCAAGTAGCCCACTTGT |
| 4_known | A18 | Tb927.10.8900_for | GGGGACAAGTTTGTACAAAAAAGCAGGCTATGAACGCGCCTCACAGAAC | Tb927.10.8900_rev | GGGGACCACTTTGTACAAGAAAGCTGGGTCTACTCCCACTTCGTGTTGGAGT |
| 4_known | A19 | Tb927.10.14950_for | GGGGACAAGTTTGTACAAAAAAGCAGGCTATGTACGGTAACCAGTCTAGTGGTG | Tb927.10.14950_rev | GGGGACCACTTTGTACAAGAAAGCTGGGTTTAAGTACTTCCCTCAGAAGGTTCCG |
| 4_known | A20 | Tb927.2.2450_for | GGGGACAAGTTTGTACAAAAAAGCAGGCTATGGGTCGTGCTTCAAAGGAC | Tb927.2.2450_rev | GGGGACCACTTTGTACAAGAAAGCTGGGTTTATCCGACACGCTGCCGT |
| 4_known | A21 | Tb927.7.3180_for | GGGGACAAGTTTGTACAAAAAAGCAGGCTATGGCTTCCGTGTTCTATATTCTTGAC | Tb927.7.3180_rev | GGGGACCACTTTGTACAAGAAAGCTGGGTTTACGCCGTCCGAATCTGGT |
| 4_known | A22 | Tb927.11.3880_for | GGGGACAAGTTTGTACAAAAAAGCAGGCTATGGTTTGCAGCACGGAGC | Tb927.11.3880_rev | GGGGACCACTTTGTACAAGAAAGCTGGGTCTACTGGCACTTGCAGTGCA |
| 4_known | A23 | Tb927.8.6810_for | GGGGACAAGTTTGTACAAAAAAGCAGGCTATGAGTTGCACACCTGCAACG | Tb927.8.6810_rev | GGGGACCACTTTGTACAAGAAAGCTGGGTTTATTCCGTTTCCCCCAGTATCCA |
| 4_known | A24 | Tb927.1.1270_for | GGGGACAAGTTTGTACAAAAAAGCAGGCTATGAAGGACATTAGTTGCCCTACGG | Tb927.1.1270_rev | GGGGACCACTTTGTACAAGAAAGCTGGGTTCACACTAAACTGCGTATCATAGCAAT |
| 4_known | B1 | Tb927.8.3970_for | GGGGACAAGTTTGTACAAAAAAGCAGGCTATGGGCGCAGGGACGTCG | Tb927.8.3970_rev | GGGGACCACTTTGTACAAGAAAGCTGGGTTTATCGCGCGGTGGCATC |
| 4_known | B2 | Tb927.11.16020_for | GGGGACAAGTTTGTACAAAAAAGCAGGCTATGGCAGCAGCGTTACTTAAGAG | Tb927.11.16020_rev | GGGGACCACTTTGTACAAGAAAGCTGGGTTTAGTTTGCCTCATTGTTGGAACTCT |
| 4_known | B3 | Tb927.6.4540_for | GGGGACAAGTTTGTACAAAAAAGCAGGCTATGCTTCGTAGGACTCTCTCATGT | Tb927.6.4540_rev | GGGGACCACTTTGTACAAGAAAGCTGGGTTTACTTCCTACGATTCAACCGCATG |
| 4_known | B4 | Tb927.10.13870_for | GGGGACAAGTTTGTACAAAAAAGCAGGCTATGCCGGCGGTAGGGGAG | Tb927.10.13870_rev | GGGGACCACTTTGTACAAGAAAGCTGGGTCTACGTCAGAAGGCGCTCAC |
| 4_known | B5 | Tb927.5.2280_for | GGGGACAAGTTTGTACAAAAAAGCAGGCTATGGCACTCACTGCACGTG | Tb927.5.2280_rev | GGGGACCACTTTGTACAAGAAAGCTGGGTTTAGGGTATTAAATAGTAGCTCAAGTTACC |
| 4_known | B6 | Tb927.10.4870_for | GGGGACAAGTTTGTACAAAAAAGCAGGCTATGTCCATTTTCAATGGGAAAACAATTCTT | Tb927.10.4870_rev | GGGGACCACTTTGTACAAGAAAGCTGGGTTTACCGACCCATCTCCGCC |
| 4_known | B7 | Tb927.1.740_for | GGGGACAAGTTTGTACAAAAAAGCAGGCTATGTTTGTGCGCCGGTTGG | Tb927.1.740_rev | GGGGACCACTTTGTACAAGAAAGCTGGGTTTATCCCTTTGAGGTAAAGGCCTTCC |
| 4_known | B8 | Tb927.11.350_for | GGGGACAAGTTTGTACAAAAAAGCAGGCTATGGAGGAGAAAGAAAGTGATTCTTCG | Tb927.11.350_rev | GGGGACCACTTTGTACAAGAAAGCTGGGTTTAAAACTGCCGACGAGGGTTC |
| 4_known | B9 | Tb927.11.3620_for | GGGGACAAGTTTGTACAAAAAAGCAGGCTATGGCTCTTGGATTTTCCTCTGC | Tb927.11.3620_rev | GGGGACCACTTTGTACAAGAAAGCTGGGTTCATGGTAGGAAGACAACCACAATCA |
| 4_known | B11 | Tb927.3.2060_for | GGGGACAAGTTTGTACAAAAAAGCAGGCTATGGAGAACATTCGTAGGACGAATCG | Tb927.3.2060_rev | GGGGACCACTTTGTACAAGAAAGCTGGGTCTACTCTGGGTCTTTACTCGACCG |
| 4_known | B12 | Tb927.6.1490_for | GGGGACAAGTTTGTACAAAAAAGCAGGCTATGTGTTCCAACCCCGGGG | Tb927.6.1490_rev | GGGGACCACTTTGTACAAGAAAGCTGGGTTTACATGGAGCAAATGTGGCAACG |
| 4_known | B13 | Tb927.10.540_for | GGGGACAAGTTTGTACAAAAAAGCAGGCTATGAGTAGTGGTCTCGCTGACTT | Tb927.10.540_rev | GGGGACCACTTTGTACAAGAAAGCTGGGTCTATTGGTTCATATACTGGCTCTGGTTC |
| 4_known | B14 | Tb927.11.14110_for | GGGGACAAGTTTGTACAAAAAAGCAGGCTATGGATGTGTGGGATGCCTCT | Tb927.11.14110_rev | GGGGACCACTTTGTACAAGAAAGCTGGGTTCAGATAGAGAATTCCCTGCGGAG |
| 4_known | B15 | Tb927.11.7216_for | GGGGACAAGTTTGTACAAAAAAGCAGGCTATGAAACGAAGGAGGGACAGTAACG | Tb927.11.7216_rev | GGGGACCACTTTGTACAAGAAAGCTGGGTCTATATCGAAAACAGTGGGCATATCCC |
| 4_known | B16 | Tb927.11.1670_for | GGGGACAAGTTTGTACAAAAAAGCAGGCTATGTTTAGTGGTGTTCGCGTACTT | Tb927.11.1670_rev | GGGGACCACTTTGTACAAGAAAGCTGGGTCTACCGCCACTCCACGTCT |
| 4_known | B17 | Tb927.8.2940_for | GGGGACAAGTTTGTACAAAAAAGCAGGCTATGAAAGGCGCTCAGTTTTTGAAGA | Tb927.8.2940_rev | GGGGACCACTTTGTACAAGAAAGCTGGGTTCAAAAAAAAAAACAGCTCAACGCAG |
| 4_known | B18 | Tb927.8.3690_for | GGGGACAAGTTTGTACAAAAAAGCAGGCTATGTTCCGCCGTGTAGCCT | Tb927.8.3690_rev | GGGGACCACTTTGTACAAGAAAGCTGGGTTCACCCCTTTACTGAACGCGC |
| 4_known | B19 | Tb927.10.4940_for | GGGGACAAGTTTGTACAAAAAAGCAGGCTATGTCAGATGTGCCTAAATCTCAAGTTG | Tb927.10.4940_rev | GGGGACCACTTTGTACAAGAAAGCTGGGTTTAATCATCATCTGAAAAATTCTGCAGCTC |
| 4_known | B20 | Tb927.11.9590_for | GGGGACAAGTTTGTACAAAAAAGCAGGCTATGACTGATTACAAGGTACGTGATATCAGC | Tb927.11.9590_rev | GGGGACCACTTTGTACAAGAAAGCTGGGTTTAGTAACGGTAGTGGTCAGGTTTGA |
| 4_known | B21 | Tb927.7.2500_for | GGGGACAAGTTTGTACAAAAAAGCAGGCTATGGCAAGCGAGAAGGGAGG | Tb927.7.2500_rev | GGGGACCACTTTGTACAAGAAAGCTGGGTTTAGTTGTACACCATGTATTTGGCTGT |
| 4_known | B22 | Tb927.11.3740_for | GGGGACAAGTTTGTACAAAAAAGCAGGCTATGGGGCAAGCAGGTGGG | Tb927.11.3740_rev | GGGGACCACTTTGTACAAGAAAGCTGGGTCTAAAGGTAAAACTTGGAGAAGCGACC |
| 4_known | B23 | Tb927.3.3990_for | GGGGACAAGTTTGTACAAAAAAGCAGGCTATGACTCCTGTATTTGGTACCGCT | Tb927.3.3990_rev | GGGGACCACTTTGTACAAGAAAGCTGGGTTCACCTCATCAAAGACAATCTCTTCG |
| 4_known | B24 | Tb927.8.2540_for | GGGGACAAGTTTGTACAAAAAAGCAGGCTATGCTCCGTCGTACGTCCTC | Tb927.8.2540_rev | GGGGACCACTTTGTACAAGAAAGCTGGGTTTAGATATTCGTTAGCACCATAGACACTGC |
| 4_known | C1 | Tb927.6.3110_for | GGGGACAAGTTTGTACAAAAAAGCAGGCTATGGATTCACCAACTGAGTTTTCTTGT | Tb927.6.3110_rev | GGGGACCACTTTGTACAAGAAAGCTGGGTTCAAAGTGTGGCCATGTTCATGT |
| 4_known | C2 | Tb927.5.4330_for | GGGGACAAGTTTGTACAAAAAAGCAGGCTATGAGATCGTTTATGTACATGTGCCG | Tb927.5.4330_rev | GGGGACCACTTTGTACAAGAAAGCTGGGTTCACGCCACCTCGCCCGC |
| 4_known | C3 | Tb927.11.3630_for | GGGGACAAGTTTGTACAAAAAAGCAGGCTATGGCTCTTGGATTTTCCTCTGC | Tb927.11.3630_rev | GGGGACCACTTTGTACAAGAAAGCTGGGTTCATGGTAGGAAGATAACCACAATCATCG |
| 4_known | C4 | Tb927.3.2320_for | GGGGACAAGTTTGTACAAAAAAGCAGGCTATGGAGGATTATTATTATTACGAAGACGAC | Tb927.3.2320_rev | GGGGACCACTTTGTACAAGAAAGCTGGGTTCAAAGGTTACGTTCTATGTCCTTGTATG |
| 4_known | C5 | Tb927.1.700_for | GGGGACAAGTTTGTACAAAAAAGCAGGCTATGACCCTTAACGAGAAGAAGAGCA | Tb927.1.700_rev | GGGGACCACTTTGTACAAGAAAGCTGGGTTTAAAGAGAGCTCCACCGGTTAGA |
| 4_known | C6 | Tb927.8.7110_for | GGGGACAAGTTTGTACAAAAAAGCAGGCTATGACCGCGCACAAACGT | Tb927.8.7110_rev | GGGGACCACTTTGTACAAGAAAGCTGGGTTCAATGCGATATGGCTTTCTGAAGG |
| 4_known | C7 | Tb927.4.5310_for | GGGGACAAGTTTGTACAAAAAAGCAGGCTATGACGGAACATAAGCGTCCCG | Tb927.4.5310_rev | GGGGACCACTTTGTACAAGAAAGCTGGGTTCAACGAGGCACGGCATTC |
| 4_known | C8 | Tb927.11.5040_for | GGGGACAAGTTTGTACAAAAAAGCAGGCTATGAAAGTGGACAGCCTTCTGTCT | Tb927.11.5040_rev | GGGGACCACTTTGTACAAGAAAGCTGGGTCTATCCAGGTGTGGGCTTATTATGATATTC |
| 4_known | C9 | Tb927.1.1920_for | GGGGACAAGTTTGTACAAAAAAGCAGGCTATGAAGGGATGTACGCTCCAGAA | Tb927.1.1920_rev | GGGGACCACTTTGTACAAGAAAGCTGGGTCTAACGCTTTTCACGCGATCCA |
| 4_known | C10 | Tb927.2.4710_for | GGGGACAAGTTTGTACAAAAAAGCAGGCTATGCAACAATATACCCTTCGGGTG | Tb927.2.4710_rev | GGGGACCACTTTGTACAAGAAAGCTGGGTTCAGTCCCTTACGCGGTCG |
| 4_known | C11 | Tb927.11.16560_for | GGGGACAAGTTTGTACAAAAAAGCAGGCTATGTTTCCCTTGAGGAGTGTGAGT | Tb927.11.16560_rev | GGGGACCACTTTGTACAAGAAAGCTGGGTTCAACCGCGACTATAAATTATCAGCG |
| 4_known | C12 | Tb927.8.4890_for | GGGGACAAGTTTGTACAAAAAAGCAGGCTATGTTGAAGATGCGATTATTGATAGTAGTT | Tb927.8.4890_rev | GGGGACCACTTTGTACAAGAAAGCTGGGTTCAACTCATAGCCGTGTAATTGAATTTTTC |
| 4_known | C13 | Tb927.10.5140_for | GGGGACAAGTTTGTACAAAAAAGCAGGCTATGTCATCAGAAATAGAGCCACATATCCT | Tb927.10.5140_rev | GGGGACCACTTTGTACAAGAAAGCTGGGTTCATTTGTGCAACACACGAGAGG |
| 4_known | C14 | Tb927.11.14480_for | GGGGACAAGTTTGTACAAAAAAGCAGGCTATGGCTAGTGTTTTACCATCCGC | Tb927.11.14480_rev | GGGGACCACTTTGTACAAGAAAGCTGGGTTCATGTCTTGCGCTGGTACAC |
| 4_known | C15 | Tb927.1.2330_for | GGGGACAAGTTTGTACAAAAAAGCAGGCTATGCGCGAAATCGTCTGCG | Tb927.1.2330_rev | GGGGACCACTTTGTACAAGAAAGCTGGGTCTAGTATTGCTCCTCCTCGTCGA |
| 4_known | C19 | Tb927.11.11770_for | GGGGACAAGTTTGTACAAAAAAGCAGGCTATGAATCCAGAAGCGGAGGAATTTG | Tb927.11.11770_rev | GGGGACCACTTTGTACAAGAAAGCTGGGTCTAAAGCGTGAAGTCGGGTGTT |
| 4_known | C20 | Tb927.7.1930_for | GGGGACAAGTTTGTACAAAAAAGCAGGCTATGGTTACCCTACGGATGGCC | Tb927.7.1930_rev | GGGGACCACTTTGTACAAGAAAGCTGGGTCTAACGCCGCACCTTCCC |
| 4_known | C21 | Tb927.10.14730_for | GGGGACAAGTTTGTACAAAAAAGCAGGCTATGCAGCAGGCGGACTATTATG | Tb927.10.14730_rev | GGGGACCACTTTGTACAAGAAAGCTGGGTTTATAGTTTCCACTCATCCAAGTTAATTGG |
| 4_known | C22 | Tb927.9.11250_for | GGGGACAAGTTTGTACAAAAAAGCAGGCTATGGTAAAGGTGAACGTTATCAGCC | Tb927.9.11250_rev | GGGGACCACTTTGTACAAGAAAGCTGGGTCTACTTCAGATACTGATAAACCTTGCGC |
| 4_known | C23 | Tb927.3.3280_for | GGGGACAAGTTTGTACAAAAAAGCAGGCTATGTCCGTTGATTTCACGTCCAA | Tb927.3.3280_rev | GGGGACCACTTTGTACAAGAAAGCTGGGTTTATCCCTCCTGCCCAAACTCTT |
| 4_known | C24 | Tb927.10.10170_for | GGGGACAAGTTTGTACAAAAAAGCAGGCTATGAAATGTGAGCTGGCATCCC | Tb927.10.10170_rev | GGGGACCACTTTGTACAAGAAAGCTGGGTTTACTCCTTCGTGCGCCACA |
| 4_known | D1 | Tb927.3.1070_for | GGGGACAAGTTTGTACAAAAAAGCAGGCTATGAGGAATGCCCCATTGCTCT | Tb927.3.1070_rev | GGGGACCACTTTGTACAAGAAAGCTGGGTTTACGTAACACAGTTGAAGTCGCTC |
| 4_known | D2 | Tb927.11.4780_for | GGGGACAAGTTTGTACAAAAAAGCAGGCTATGTTTCGGTGTTATGGACGAAAACT | Tb927.11.4780_rev | GGGGACCACTTTGTACAAGAAAGCTGGGTCTAAAACGAACCCTCTCTATCCCG |
| 4_known | D3 | Tb927.5.3620_for | GGGGACAAGTTTGTACAAAAAAGCAGGCTATGCAGTCGTCAAGTTTTGCTGG | Tb927.5.3620_rev | GGGGACCACTTTGTACAAGAAAGCTGGGTTCAGGATTCATTAGCCAACGACTCC |
| 4_known | D4 | Tb927.9.7190_for | GGGGACAAGTTTGTACAAAAAAGCAGGCTATGCGACGCTCTGCCACC | Tb927.9.7190_rev | GGGGACCACTTTGTACAAGAAAGCTGGGTTCAATGTGTGCTCTCTGCCGC |
| 4_known | D5 | Tb927.3.4680_for | GGGGACAAGTTTGTACAAAAAAGCAGGCTATGGAGGAAACGTACGATGCAATT | Tb927.3.4680_rev | GGGGACCACTTTGTACAAGAAAGCTGGGTCTACCTATCCTCGTTGGAAGCAGT |
| 4_known | D6 | Tb927.7.2550_for | GGGGACAAGTTTGTACAAAAAAGCAGGCTATGACGCAGGACACCGCA | Tb927.7.2550_rev | GGGGACCACTTTGTACAAGAAAGCTGGGTCTATGCGTAATAGTTCAAGCTAGATTTTTT |
| 4_known | D7 | Tb927.9.13430_for | GGGGACAAGTTTGTACAAAAAAGCAGGCTATGCGTGCGGAGATTCCCG | Tb927.9.13430_rev | GGGGACCACTTTGTACAAGAAAGCTGGGTTCAAAAAAAAGTTGGTTCAACCTGAGG |
| 4_known | D8 | Tb927.10.14240_for | GGGGACAAGTTTGTACAAAAAAGCAGGCTATGACTGACAAAGCATTTACGGAACA | Tb927.10.14240_rev | GGGGACCACTTTGTACAAGAAAGCTGGGTTTAAACCATCACAAGACCAACTTCATTAGC |
| 4_known | D9 | Tb927.10.4770_for | GGGGACAAGTTTGTACAAAAAAGCAGGCTATGGGTTCATGCCAGGCGG | Tb927.10.4770_rev | GGGGACCACTTTGTACAAGAAAGCTGGGTCTATTTCTCTGGTGGAGGTATTTTGCC |
| 4_known | D10 | Tb927.3.910_for | GGGGACAAGTTTGTACAAAAAAGCAGGCTATGCCACGGGAAATCATCACAC | Tb927.3.910_rev | GGGGACCACTTTGTACAAGAAAGCTGGGTTCAAAAGTTTCGTATATAATCGGAGCTCTC |
| 4_known | D11 | Tb927.6.3130_for | GGGGACAAGTTTGTACAAAAAAGCAGGCTATGTCCAAGGAGGAGGCGG | Tb927.6.3130_rev | GGGGACCACTTTGTACAAGAAAGCTGGGTTCAGCGGGGGAGCTCAAC |
| 4_known | D12 | Tb927.9.12700_for | GGGGACAAGTTTGTACAAAAAAGCAGGCTATGGTAACGTGGGCGCTGA | Tb927.9.12700_rev | GGGGACCACTTTGTACAAGAAAGCTGGGTCTAACACGTTGAACACACTTCGGT |
| 4_known | D13 | Tb927.6.2830_for | GGGGACAAGTTTGTACAAAAAAGCAGGCTATGCTCCAACGCGGTTTGG | Tb927.6.2830_rev | GGGGACCACTTTGTACAAGAAAGCTGGGTCTACTGCGCGAGATGTGCAA |
| 4_known | D14 | Tb927.9.9970_for | GGGGACAAGTTTGTACAAAAAAGCAGGCTATGCCTCCTAAACAATTGGTGGAC | Tb927.9.9970_rev | GGGGACCACTTTGTACAAGAAAGCTGGGTTTAAGCAGTGTAGTACTTTCCTTGAGGG |
| 4_known | D15 | Tb927.7.1430_for | GGGGACAAGTTTGTACAAAAAAGCAGGCTATGCTCGCGGACCGCATA | Tb927.7.1430_rev | GGGGACCACTTTGTACAAGAAAGCTGGGTTCATGCTCGAGAGGATGATGTGC |
| 4_known | D16 | Tb927.10.2110_for | GGGGACAAGTTTGTACAAAAAAGCAGGCTATGGGAAAGGAAAAGGTGCACATG | Tb927.10.2110_rev | GGGGACCACTTTGTACAAGAAAGCTGGGTTTATTTCTTCGAAGCCTTCACCGC |
| 4_known | D21 | Tb927.11.4220_for | GGGGACAAGTTTGTACAAAAAAGCAGGCTATGAGCGTCGGCCTCTACTG | Tb927.11.4220_rev | GGGGACCACTTTGTACAAGAAAGCTGGGTTCAAATAACACCCACACTACCATCCT |
| 4_known | E2 | Tb927.6.960_for | GGGGACAAGTTTGTACAAAAAAGCAGGCTATGCCTCGAACAGAAATGGTGC | Tb927.6.960_rev | GGGGACCACTTTGTACAAGAAAGCTGGGTTCACTGGTGTGGGACCAGC |
| 4_known | E5 | Tb927.4.4850_for | GGGGACAAGTTTGTACAAAAAAGCAGGCTATGGAAGTCAGGAACGAACCAATC | Tb927.4.4850_rev | GGGGACCACTTTGTACAAGAAAGCTGGGTTTATATGTTAATAGCTCCCCAGATGGAAGC |
| 4_known | E9 | Tb927.9.9150_for | GGGGACAAGTTTGTACAAAAAAGCAGGCTATGCGCCGTGGACTCACT | Tb927.9.9150_rev | GGGGACCACTTTGTACAAGAAAGCTGGGTTCACCGCCCTGGAATAATGCG |
| 4_known | E10 | Tb927.9.12730_for | GGGGACAAGTTTGTACAAAAAAGCAGGCTATGTTGCGGTTCACGTCCG | Tb927.9.12730_rev | GGGGACCACTTTGTACAAGAAAGCTGGGTCTACTGATGAACGCACTCGGC |
| 4_known | E11 | Tb927.10.7570_for | GGGGACAAGTTTGTACAAAAAAGCAGGCTATGTTCCGCCGTCCCGTC | Tb927.10.7570_rev | GGGGACCACTTTGTACAAGAAAGCTGGGTCTACAGCAATAGCGAGAGTGGGT |
| 4_known | E12 | Tb927.9.12910_for | GGGGACAAGTTTGTACAAAAAAGCAGGCTATGTGTAGCATTGAGGGCCCG | Tb927.9.12910_rev | GGGGACCACTTTGTACAAGAAAGCTGGGTCTAAAACTCACCGAACTGTTGCAC |
| 4_known | E13 | Tb927.1.2400_for | GGGGACAAGTTTGTACAAAAAAGCAGGCTATGCGTGAGGCTATCTGCATCC | Tb927.1.2400_rev | GGGGACCACTTTGTACAAGAAAGCTGGGTCTAGTACTCCTCCACATCCTCCTCA |
| 4_known | E17 | Tb927.4.4870_for | GGGGACAAGTTTGTACAAAAAAGCAGGCTATGGAAGTCAGGAACGAACCAATC | Tb927.4.4870_rev | GGGGACCACTTTGTACAAGAAAGCTGGGTTCAATTTCCCACCACTACGCCA |
| 4_known | E18 | Tb927.3.3410_for | GGGGACAAGTTTGTACAAAAAAGCAGGCTATGTTGCCACTTGACTCACCG | Tb927.3.3410_rev | GGGGACCACTTTGTACAAGAAAGCTGGGTTTAGGAGGTCACCATGTCCCTG |
| 4_known | E19 | Tb927.11.2300_for | GGGGACAAGTTTGTACAAAAAAGCAGGCTATGGCCGACCACGAGTTGT | Tb927.11.2300_rev | GGGGACCACTTTGTACAAGAAAGCTGGGTTTACATAAAGTCGTCGTCGAAGTCAAA |
| 4_known | E20 | Tb927.10.10800_for | GGGGACAAGTTTGTACAAAAAAGCAGGCTATGTGTTGTGGTCGGCGTATC | Tb927.10.10800_rev | GGGGACCACTTTGTACAAGAAAGCTGGGTTCAAACAGGAGTAGGACTTGTCAACG |
| 4_known | E21 | Tb927.11.12650_for | GGGGACAAGTTTGTACAAAAAAGCAGGCTATGGAATCGTTCGGCCCTCT | Tb927.11.12650_rev | GGGGACCACTTTGTACAAGAAAGCTGGGTTCAACTGTAGGGAAACATCACCTCC |

| | | | | | |
|---|---|---|---|---|---|
| 4_known | E22 | Tb927.2.2120_for | GGGGACAAGTTTGTACAAAAAAGCAGGCTATGGCTGATCTCCTTGAGCTTCG | Tb927.2.2120_rev | GGGGACCACTTTGTACAAGAAAGCTGGGTTTAATTACTGCTGCCGCTGCTC |
| 4_known | E23 | Tb927.11.4540_for | GGGGACAAGTTTGTACAAAAAAGCAGGCTATGTACCAGCCACTCCTTGAGG | Tb927.11.4540_rev | GGGGACCACTTTGTACAAGAAAGCTGGGTTTATGCATGATACGCAGGAACAAATGA |
| 4_known | E24 | Tb927.10.6500_for | GGGGACAAGTTTGTACAAAAAAGCAGGCTATGGTTGTGAATTCTGATGGGGTG | Tb927.10.6500_rev | GGGGACCACTTTGTACAAGAAAGCTGGGTCTAATTGATAATCGCATAAATCGTCGCC |
| 4_known | F1 | Tb927.4.3930_for | GGGGACAAGTTTGTACAAAAAAGCAGGCTATGACGGAAGGACTAGGCGAA | Tb927.4.3930_rev | GGGGACCACTTTGTACAAGAAAGCTGGGTTCACACAATCACGTCATAAACCGC |
| 4_known | F2 | Tb927.10.6350_for | GGGGACAAGTTTGTACAAAAAAGCAGGCTATGCCACCACGGACCGCT | Tb927.10.6350_rev | GGGGACCACTTTGTACAAGAAAGCTGGGTCTACTCAAAGTTGCTCCGTGGC |
| 4_known | F3 | Tb927.6.540_for | GGGGACAAGTTTGTACAAAAAAGCAGGCTATGATGCACGAATTAGTGTTTATCATTGG | Tb927.6.540_rev | GGGGACCACTTTGTACAAGAAAGCTGGGTCTAAGATATACAAATAAAAGATGAAAACAA |
| 4_known | F4 | Tb927.8.7050_for | GGGGACAAGTTTGTACAAAAAAGCAGGCTATGCCCATTTCGCAAGTGTTCA | Tb927.8.7050_rev | GGGGACCACTTTGTACAAGAAAGCTGGGTTCAATGTGTTCGGAATACGTAGGAATC |
| 4_known | F5 | Tb927.11.11440_for | GGGGACAAGTTTGTACAAAAAAGCAGGCTATGCCAGCGATGAAAAATAATTGTGC | Tb927.11.11440_rev | GGGGACCACTTTGTACAAGAAAGCTGGGTCTATCCGGCCAATCTAGGTCGT |
| 4_known | F6 | Tb927.2.1780_for | GGGGACAAGTTTGTACAAAAAAGCAGGCTATGCGTCGTCACCGCGTG | Tb927.2.1780_rev | GGGGACCACTTTGTACAAGAAAGCTGGGTTCAGGATGGGCTTCCCACAA |
| 4_known | F7 | Tb927.10.880_for | GGGGACAAGTTTGTACAAAAAAGCAGGCTATGCACGACGGAGATGGCA | Tb927.10.880_rev | GGGGACCACTTTGTACAAGAAAGCTGGGTCTAAGTAGTATCAACGGCCATTTGATTTCT |
| 4_known | F8 | Tb927.11.3140_for | GGGGACAAGTTTGTACAAAAAAGCAGGCTATGGATGTCTCTCGCAGGTTGG | Tb927.11.3140_rev | GGGGACCACTTTGTACAAGAAAGCTGGGTCTAGCACCCGCATCTGTCGG |
| 4_known | F9 | Tb927.11.9160_for | GGGGACAAGTTTGTACAAAAAAGCAGGCTATGGAGGATTTCGTTGTTGTGCC | Tb927.11.9160_rev | GGGGACCACTTTGTACAAGAAAGCTGGGTCTATGATGACACCATCATTCTCCTCG |
| 4_known | F10 | Tb927.4.4000_for | GGGGACAAGTTTGTACAAAAAAGCAGGCTATGAGCAACTCCAAGCAACAACA | Tb927.4.4000_rev | GGGGACCACTTTGTACAAGAAAGCTGGGTTCATCCCCTGACGGTATCCAAAA |
| 4_known | F11 | Tb927.9.1400_for | GGGGACAAGTTTGTACAAAAAAGCAGGCTATGCGTGTTGTCATCGTTGGT | Tb927.9.1400_rev | GGGGACCACTTTGTACAAGAAAGCTGGGTTTAGTTCCTGTACGCCGTTTCCA |
| 4_known | F14 | Tb927.8.7120_for | GGGGACAAGTTTGTACAAAAAAGCAGGCTATGGATTACGTTAGCGAATTGTACGC | Tb927.8.7120_rev | GGGGACCACTTTGTACAAGAAAGCTGGGTTCACCTGCCCAAATAACCGAAGA |
| 4_known | F15 | Tb927.6.2000_for | GGGGACAAGTTTGTACAAAAAAGCAGGCTATGAAGGAAGCAAAAAAGAAAAAGTTACTG | Tb927.6.2000_rev | GGGGACCACTTTGTACAAGAAAGCTGGGTTCAAAATTTTGTTGGTGCACGTGG |
| 4_known | F16 | Tb927.11.8460_for | GGGGACAAGTTTGTACAAAAAAGCAGGCTATGGAATCAAAACACCCCGAAGTG | Tb927.11.8460_rev | GGGGACCACTTTGTACAAGAAAGCTGGGTCAACCCTTTGGAAATACCTTCATGAC |
| 4_known | F17 | Tb927.6.500_for | GGGGACAAGTTTGTACAAAAAAGCAGGCTATGAGGCACGAATTAGTGTTTACCAT | Tb927.6.500_rev | GGGGACCACTTTGTACAAGAAAGCTGGGTCTAAGATATACAAATGAAAGATGAAAACAA |
| 4_known | F18 | Tb927.1.1030_for | GGGGACAAGTTTGTACAAAAAAGCAGGCTATGAGTCGTTTTGCCGCAAGC | Tb927.1.1030_rev | GGGGACCACTTTGTACAAGAAAGCTGGGTTTACCAACCTGAGGCCTCAGTC |
| 4_known | F19 | Tb927.6.470_for | GGGGACAAGTTTGTACAAAAAAGCAGGCTATGAGGCACGAATTAGTGTTTACCAT | Tb927.6.470_rev | GGGGACCACTTTGTACAAGAAAGCTGGGTCTAAGATATACAAATAAAAGATGAAAACAA |
| 4_known | F20 | Tb927.11.16590_for | GGGGACAAGTTTGTACAAAAAAGCAGGCTATGAGCACATCAGCGGGGA | Tb927.11.16590_rev | GGGGACCACTTTGTACAAGAAAGCTGGGTCTACACCCACCTCCTCCCATTT |
| 4_known | F21 | Tb927.7.540_for | GGGGACAAGTTTGTACAAAAAAGCAGGCTATGTCGTCCGCAGATCCCA | Tb927.7.540_rev | GGGGACCACTTTGTACAAGAAAGCTGGGTTTACAGAAAACCAACATACCGCAATGG |
| 4_known | F22 | Tb927.8.4700_for | GGGGACAAGTTTGTACAAAAAAGCAGGCTATGCTGAGCCCTACAGAACCA | Tb927.8.4700_rev | GGGGACCACTTTGTACAAGAAAGCTGGGTTTACACAGCCACATTGTAGATTGAGG |
| 4_known | F24 | Tb927.3.4900_for | GGGGACAAGTTTGTACAAAAAAGCAGGCTATGAGGTCTATTTACTTTGTCATAGGCG | Tb927.3.4900_rev | GGGGACCACTTTGTACAAGAAAGCTGGGTTTACGACCGCTTAACTCTCTTAGAAGG |
| 4_known | G1 | Tb927.8.4730_for | GGGGACAAGTTTGTACAAAAAAGCAGGCTATGCTGAGCCCTACAGAAGCA | Tb927.8.4730_rev | GGGGACCACTTTGTACAAGAAAGCTGGGTTTACACAGCCACATTGTAGATTGAGG |
| 4_known | G3 | Tb927.11.15530_for | GGGGACAAGTTTGTACAAAAAAGCAGGCTATGCCGCCACGGAGCCGC | Tb927.11.15530_rev | GGGGACCACTTTGTACAAGAAAGCTGGGTCTAATAGACGTATGGGATCAAGCGATAAC |
| 4_known | G4 | Tb927.5.3810_for | GGGGACAAGTTTGTACAAAAAAGCAGGCTATGTCCTCTACTTTCTTTGATATGCTTAAC | Tb927.5.3810_rev | GGGGACCACTTTGTACAAGAAAGCTGGGTTTACAGCTTGCTAGTGGATTCCAAC |
| 4_known | G5 | Tb927.6.2250_for | GGGGACAAGTTTGTACAAAAAAGCAGGCTATGACCATCGATTACTCAGGTTTTCTTG | Tb927.6.2250_rev | GGGGACCACTTTGTACAAGAAAGCTGGGTCTATGTAGGTCTTGATCTCCCATTAAACG |
| 4_known | G7 | Tb927.10.5960_for | GGGGACAAGTTTGTACAAAAAAGCAGGCTATGGGAACTTCGTCCACCGA | Tb927.10.5960_rev | GGGGACCACTTTGTACAAGAAAGCTGGGTTCACTCTTCATTATTATCATCATGATCCCC |
| 4_known | G8 | Tb927.4.1270_for | GGGGACAAGTTTGTACAAAAAAGCAGGCTATGTCTGGAATCAAAATTGAGGAGGTT | Tb927.4.1270_rev | GGGGACCACTTTGTACAAGAAAGCTGGGTCTATTGGTAGACGTAATCCTCTGCGT |
| 4_known | G9 | Tb927.4.1400_for | GGGGACAAGTTTGTACAAAAAAGCAGGCTATGGAAGATTCCTTCAACCTGAAGAAAG | Tb927.4.1400_rev | GGGGACCACTTTGTACAAGAAAGCTGGGTTCATTTTCCTCCTGTGCGCC |
| 4_known | G10 | Tb927.8.8220_for | GGGGACAAGTTTGTACAAAAAAGCAGGCTATGCGAGCCACCAAAAAGCA | Tb927.8.8220_rev | GGGGACCACTTTGTACAAGAAAGCTGGGTCTACGGCACCGTGTTGTATATTGT |
| 4_known | G11 | Tb927.9.8410_for | GGGGACAAGTTTGTACAAAAAAGCAGGCTATGAGGCAACGGAGTCCTTTGT | Tb927.9.8410_rev | GGGGACCACTTTGTACAAGAAAGCTGGGTCTAGGCGGCAAACCAAGATTTGT |
| 4_known | G13 | Tb927.8.8260_for | GGGGACAAGTTTGTACAAAAAAGCAGGCTATGACGAATCAATCCAGCGGGG | Tb927.8.8260_rev | GGGGACCACTTTGTACAAGAAAGCTGGGTCTACACAGCATAATATACTGCCGATCC |
| 4_known | G14 | Tb927.10.210_for | GGGGACAAGTTTGTACAAAAAAGCAGGCTATGTTGCAGGCGGGAACAAA | Tb927.10.210_rev | GGGGACCACTTTGTACAAGAAAGCTGGGTCTAAGCACCCAACGTCGACC |
| 4_known | G15 | Tb927.8.8250_for | GGGGACAAGTTTGTACAAAAAAGCAGGCTATGACGAATCAATCCAGCGGGG | Tb927.8.8250_rev | GGGGACCACTTTGTACAAGAAAGCTGGGTCTACGGCACCGTGTTGTATATTGT |
| 4_known | G16 | Tb927.8.8230_for | GGGGACAAGTTTGTACAAAAAAGCAGGCTATGACGAATCAATCCAGCGGGG | Tb927.8.8230_rev | GGGGACCACTTTGTACAAGAAAGCTGGGTTCAAGACAGTGCACCGGCT |
| 4_known | G17 | Tb927.9.15050_for | GGGGACAAGTTTGTACAAAAAAGCAGGCTATGCCACCAAAGAAGGAAGCAG | Tb927.9.15050_rev | GGGGACCACTTTGTACAAGAAAGCTGGGTCTACAGGGTGCTGCTTGATCCC |
| 4_known | G18 | Tb927.10.2750_for | GGGGACAAGTTTGTACAAAAAAGCAGGCTATGGCTGAGTTGGCCAAGAGT | Tb927.10.2750_rev | GGGGACCACTTTGTACAAGAAAGCTGGGTTCACGAGCGGATATTCTCCTTTCC |
| 4_known | G19 | Tb927.7.2670_for | GGGGACAAGTTTGTACAAAAAAGCAGGCTATGAGCCATCCACAATCTCATCGC | Tb927.7.2670_rev | GGGGACCACTTTGTACAAGAAAGCTGGGTTCAATCGCTGATTCCTCCCGT |
| 4_known | G20 | Tb927.2.6150_for | GGGGACAAGTTTGTACAAAAAAGCAGGCTATGGCAATGCTTGGTTTCGAGT | Tb927.2.6150_rev | GGGGACCACTTTGTACAAGAAAGCTGGGTTTATTTAGGAAGTCCCTCCTTAACAGCC |
| 4_known | G21 | Tb927.3.590_for | GGGGACAAGTTTGTACAAAAAAGCAGGCTATGATGCTCGGGTTCGAATCG | Tb927.3.590_rev | GGGGACCACTTTGTACAAGAAAGCTGGGTCTATTGAGGAAGTCCCTCCTTGACG |
| 4_known | G22 | Tb927.2.6220_for | GGGGACAAGTTTGTACAAAAAAGCAGGCTATGGCAATGCTTGGTTTCGAGT | Tb927.2.6220_rev | GGGGACCACTTTGTACAAGAAAGCTGGGTCTACTCTGTTTTAGGAAGAGCCTCCT |
| 4_known | G23 | Tb927.9.7470_for | GGGGACAAGTTTGTACAAAAAAGCAGGCTATGCTCGGGTTTGAGTCGTTTT | Tb927.9.7470_rev | GGGGACCACTTTGTACAAGAAAGCTGGGTTTATGATTCTGGAAGAGCCGTCATAAC |
| 4_known | G24 | Tb927.6.3230_for | GGGGACAAGTTTGTACAAAAAAGCAGGCTATGGTACGCAGCATTCGCTAC | Tb927.6.3230_rev | GGGGACCACTTTGTACAAGAAAGCTGGGTTCACTGATAACAGAAGGTAGTGATAACCT |
| 4_known | H1 | Tb927.11.15860_for | GGGGACAAGTTTGTACAAAAAAGCAGGCTATGTCACCCAGGGCAACAGA | Tb927.11.15860_rev | GGGGACCACTTTGTACAAGAAAGCTGGGTCTACTTCTTAACCTCGTCGTAAATGGTT |
| 4_known | H4 | Tb927.10.14010_for | GGGGACAAGTTTGTACAAAAAAGCAGGCTATGACCTACCAAGCGAAGGAAGT | Tb927.10.14010_rev | GGGGACCACTTTGTACAAGAAAGCTGGGTCTAAAAGTCGTCACTGTCATCGTCG |
| 4_known | H5 | Tb927.9.2010_for | GGGGACAAGTTTGTACAAAAAAGCAGGCTATGCGCAATAACGCAGCGG | Tb927.9.2010_rev | GGGGACCACTTTGTACAAGAAAGCTGGGTTTAAAAACACTCGGCTATGGCACG |
| 4_known | H6 | Tb927.11.9130_for | GGGGACAAGTTTGTACAAAAAAGCAGGCTATGCAGCAGCCATCACAACA | Tb927.11.9130_rev | GGGGACCACTTTGTACAAGAAAGCTGGGTTTACAAGTAGGCATTAATGCATTTAATCAT |
| 4_known | H7 | Tb927.2.6240_for | GGGGACAAGTTTGTACAAAAAAGCAGGCTATGGCAATGCTTGGTTTCGAGT | Tb927.2.6240_rev | GGGGACCACTTTGTACAAGAAAGCTGGGTTTATTCAGGAAGTCCCTCCTTAACAGC |
| 4_known | H8 | Tb927.6.220_for | GGGGACAAGTTTGTACAAAAAAGCAGGCTATGCTCGGTTTTTATTCAGTACCTGAA | Tb927.6.220_rev | GGGGACCACTTTGTACAAGAAAGCTGGGTTTAATGCGGAAGAGCTTTTATAACCGC |
| 4_known | H9 | Tb927.10.1420_for | GGGGACAAGTTTGTACAAAAAAGCAGGCTATGCCACCTAAAGGCGGGG | Tb927.10.1420_rev | GGGGACCACTTTGTACAAGAAAGCTGGGTTTAGTAGTCGTCGCCCTTAGAAAGG |
| 4_known | H10 | Tb927.9.14070_for | GGGGACAAGTTTGTACAAAAAAGCAGGCTATGGGTGTTGGGTTGGGTTTC | Tb927.9.14070_rev | GGGGACCACTTTGTACAAGAAAGCTGGGTCTACGCGGAAAGGTCTTCCTTTG |
| 4_known | H11 | Tb927.5.286b_for | GGGGACAAGTTTGTACAAAAAAGCAGGCTATGCTCGGGTTTGACTCAGCT | Tb927.5.286b_rev | GGGGACCACTTTGTACAAGAAAGCTGGGTCTACTTGGGAAGCCCCTCATTGA |
| 4_known | H12 | Tb927.10.1040_for | GGGGACAAGTTTGTACAAAAAAGCAGGCTATGATGCTGTGCCACACGTC | Tb927.10.1040_rev | GGGGACCACTTTGTACAAGAAAGCTGGGTTTACGAGAGTGGTTCGTTCCTCA |
| 4_known | H13 | Tb927.2.6200_for | GGGGACAAGTTTGTACAAAAAAGCAGGCTATGGCAATGCTTGGTTTCGAGT | Tb927.2.6200_rev | GGGGACCACTTTGTACAAGAAAGCTGGGTTTATTCAGGAAGTCCCTCTTTAACAGCC |
| 4_known | H14 | Tb927.7.5770_for | GGGGACAAGTTTGTACAAAAAAGCAGGCTATGTCCACGCGGAAAGAGC | Tb927.7.5770_rev | GGGGACCACTTTGTACAAGAAAGCTGGGTTTACCGTTTGAGGTTGAATTTGTAGTCT |
| 4_known | H15 | Tb927.8.5390_for | GGGGACAAGTTTGTACAAAAAAGCAGGCTATGTCAACTGTCAACAGACTGCAA | Tb927.8.5390_rev | GGGGACCACTTTGTACAAGAAAGCTGGGTTTACGGATATACACTGTGTTCTCGAAGT |
| 4_known | H17 | Tb927.9.1500_for | GGGGACAAGTTTGTACAAAAAAGCAGGCTATGGATGGCGTGTGCGAAC | Tb927.9.1500_rev | GGGGACCACTTTGTACAAGAAAGCTGGGTTCACTTGACAATCTTCTGCGATACC |
| 4_known | H18 | Tb927.11.1390_for | GGGGACAAGTTTGTACAAAAAAGCAGGCTATGTCAAACGAAGTTGAATCTTCTGCA | Tb927.11.1390_rev | GGGGACCACTTTGTACAAGAAAGCTGGGTTTATCTTAGATCCTTATCTGTATTGATAAC |
| 4_known | H19 | Tb927.11.12410_for | GGGGACAAGTTTGTACAAAAAAGCAGGCTATGGATAGCATAAGCTCTGGTCTAACC | Tb927.11.12410_rev | GGGGACCACTTTGTACAAGAAAGCTGGGTTTACATGACAGGTGTTGGGGGG |
| 4_known | H20 | Tb927.5.2100_for | GGGGACAAGTTTGTACAAAAAAGCAGGCTATGCGTTTTGTTTGTCTTTTCTCACG | Tb927.5.2100_rev | GGGGACCACTTTGTACAAGAAAGCTGGGTTCATTTCTTTTCTGATTCCCTCTCCAAC |
| 4_known | H21 | Tb927.4.3790_for | GGGGACAAGTTTGTACAAAAAAGCAGGCTATGGTTCCTGTTAAAGTGAAATGGGG | Tb927.4.3790_rev | GGGGACCACTTTGTACAAGAAAGCTGGGTCTACAGCGGAATGACCGACTG |
| 4_known | H22 | Tb927.7.4560_for | GGGGACAAGTTTGTACAAAAAAGCAGGCTATGGTAATGTTTGAGGTGCGGC | Tb927.7.4560_rev | GGGGACCACTTTGTACAAGAAAGCTGGGTTTATACTTTCTCAGCTCTACGCAAGTCA |
| 4_known | H23 | Tb927.2.6280_for | GGGGACAAGTTTGTACAAAAAAGCAGGCTATGGCAATGCTTGGTTTCGAGT | Tb927.2.6280_rev | GGGGACCACTTTGTACAAGAAAGCTGGGTCTACTTTTTTTCCTTCGGAAGTCCCT |
| 4_known | H24 | Tb927.10.4600_for | GGGGACAAGTTTGTACAAAAAAGCAGGCTATGCAAACAGAAGCCCTGCAA | Tb927.10.4600_rev | GGGGACCACTTTGTACAAGAAAGCTGGGTTCAAAGATCATTTAGGTGTGGCGC |
| 4_known | I2 | Tb927.8.7650_for | GGGGACAAGTTTGTACAAAAAAGCAGGCTATGACCAGCATCAAGACCAAGTCG | Tb927.8.7650_rev | GGGGACCACTTTGTACAAGAAAGCTGGGTCTAACCAGTGAAAGTTCCCCAGATAGC |
| 4_known | I3 | Tb927.10.6830_for | GGGGACAAGTTTGTACAAAAAAGCAGGCTATGGAAAAGAAAGGATCGGGAAGGG | Tb927.10.6830_rev | GGGGACCACTTTGTACAAGAAAGCTGGGTTCATTTGCGTAGCTTTGCCGT |
| 4_known | I4 | Tb927.3.3360_for | GGGGACAAGTTTGTACAAAAAAGCAGGCTATGGACGCCGGTAAACTTTTTCA | Tb927.3.3360_rev | GGGGACCACTTTGTACAAGAAAGCTGGGTTTATCCCTTGGTGAACCTCTTAGGTG |
| 4_known | I5 | Tb927.10.1050_for | GGGGACAAGTTTGTACAAAAAAGCAGGCTATGCGGCTCATTTCGTATCCG | Tb927.10.1050_rev | GGGGACCACTTTGTACAAGAAAGCTGGGTTTACGAGAGTGGTTCGTTCCTCA |
| 4_known | I6 | Tb927.8.7640_for | GGGGACAAGTTTGTACAAAAAAGCAGGCTATGACCAGCATCAAGACCAAGTCG | Tb927.8.7640_rev | GGGGACCACTTTGTACAAGAAAGCTGGGTTCATCCCACAGTAACTGCCCA |
| 4_known | I7 | Tb927.8.7620_for | GGGGACAAGTTTGTACAAAAAAGCAGGCTATGACCAGCATCAATGCCCAAC | Tb927.8.7620_rev | GGGGACCACTTTGTACAAGAAAGCTGGGTTCATCCCACAGTAACTGCCCA |
| 4_known | I8 | Tb927.8.7600_for | GGGGACAAGTTTGTACAAAAAAGCAGGCTATGACCAGCATCAATGCCCAAC | Tb927.8.7600_rev | GGGGACCACTTTGTACAAGAAAGCTGGGTTTAACCAGTCGCAGTTCCCCA |
| 4_known | I9 | Tb927.4.4960_for | GGGGACAAGTTTGTACAAAAAAGCAGGCTATGGTGGAGAAACCCCGTGA | Tb927.4.4960_rev | GGGGACCACTTTGTACAAGAAAGCTGGGTTTAAACTGTTCCACAAGTTGAGTGACA |
| 4_known | I10 | Tb927.9.4620_for | GGGGACAAGTTTGTACAAAAAAGCAGGCTATGTCATGCAATCCGCTGCG | Tb927.9.4620_rev | GGGGACCACTTTGTACAAGAAAGCTGGGTTCACGAGATGTGTATATCAGCCCC |
| 4_known | I11 | Tb927.10.3690_for | GGGGACAAGTTTGTACAAAAAAGCAGGCTATGTCCCGTCGATTCCTCAACC | Tb927.10.3690_rev | GGGGACCACTTTGTACAAGAAAGCTGGGTTTAGGTCCCTGAGCCATGCG |
| 4_known | I13 | Tb927.8.7610_for | GGGGACAAGTTTGTACAAAAAAGCAGGCTATGATCAGCATCAATGCCCAACC | Tb927.8.7610_rev | GGGGACCACTTTGTACAAGAAAGCTGGGTTCATCCCACAGTAACTGCCCA |
| 4_known | I14 | Tb927.11.4970_for | GGGGACAAGTTTGTACAAAAAAGCAGGCTATGCGAAAAGCCTTGGTGGAA | Tb927.11.4970_rev | GGGGACCACTTTGTACAAGAAAGCTGGGTTTAAGAAGTATAGAAAATATCCCTTGGTCT |
| 4_known | I15 | Tb927.2.3580_for | GGGGACAAGTTTGTACAAAAAAGCAGGCTATGCTTCAAGAAAGAGTATTTCATATTAAC | Tb927.2.3580_rev | GGGGACCACTTTGTACAAGAAAGCTGGGTCTAATTCTCTGACCAATTATGTTTGCACTT |
| 4_known | I16 | Tb927.8.7740_for | GGGGACAAGTTTGTACAAAAAAGCAGGCTATGACATCCGGAGCTGCTGC | Tb927.8.7740_rev | GGGGACCACTTTGTACAAGAAAGCTGGGTTTAATAGTACTTCTTAATTGTGTAATAAAT |
| 4_known | I17 | Tb927.10.730_for | GGGGACAAGTTTGTACAAAAAAGCAGGCTATGTCGTCCACGCCCGAG | Tb927.10.730_rev | GGGGACCACTTTGTACAAGAAAGCTGGGTTTAAATGCACTCCCACCGTTGAA |
| 4_known | I18 | Tb927.11.11270_for | GGGGACAAGTTTGTACAAAAAAGCAGGCTATGATGTCCACTGAATCCCTTGAGT | Tb927.11.11270_rev | GGGGACCACTTTGTACAAGAAAGCTGGGTTCACCGAAAACCCTCCAGCG |
| 4_known | I19 | Tb927.10.70_for | GGGGACAAGTTTGTACAAAAAAGCAGGCTATGGCTGATGGCCGCGCC | Tb927.10.70_rev | GGGGACCACTTTGTACAAGAAAGCTGGGTTTAACGCACTTCCCTTTTCCAAAATC |
| 4_known | I20 | Tb927.4.1680_for | GGGGACAAGTTTGTACAAAAAAGCAGGCTATGCCTCCCAAGCGAGCG | Tb927.4.1680_rev | GGGGACCACTTTGTACAAGAAAGCTGGGTTCAGTAAACTGGTTGCCAGTTCG |
| 4_known | I21 | Tb927.11.9070_for | GGGGACAAGTTTGTACAAAAAAGCAGGCTATGACGGGGTTCCGTCTCC | Tb927.11.9070_rev | GGGGACCACTTTGTACAAGAAAGCTGGGTTTACAGTATGAAATCATCTGCCTTCTCC |
| 4_known | I22 | Tb927.7.1590_for | GGGGACAAGTTTGTACAAAAAAGCAGGCTATGAGTTACAATACCAGTAGCAGCAGC | Tb927.7.1590_rev | GGGGACCACTTTGTACAAGAAAGCTGGGTTCAAAGACCGTACAGAAGGTCTGG |
| 4_known | I23 | Tb927.2.4110_for | GGGGACAAGTTTGTACAAAAAAGCAGGCTATGCTGCGTCATTGTATTCCACG | Tb927.2.4110_rev | GGGGACCACTTTGTACAAGAAAGCTGGGTTCAGTTTAGTTTCTCATAAGTTTTGCGAAC |
| 4_known | I24 | Tb927.9.4360_for | GGGGACAAGTTTGTACAAAAAAGCAGGCTATGCAACTCCAAAGGTTGGGTG | Tb927.9.4360_rev | GGGGACCACTTTGTACAAGAAAGCTGGGTTCATTCGCCCTTTGTGGGGG |
| 4_known | J1 | Tb927.9.9200_for | GGGGACAAGTTTGTACAAAAAAGCAGGCTATGTTACTCACCGCACCGTTATTAG | Tb927.9.9200_rev | GGGGACCACTTTGTACAAGAAAGCTGGGTCTATGACGGAAGATGCCGGC |
| 4_known | J2 | Tb927.1.1750_for | GGGGACAAGTTTGTACAAAAAAGCAGGCTATGGAGCTCAATCTTACACAACTTCC | Tb927.1.1750_rev | GGGGACCACTTTGTACAAGAAAGCTGGGTTTAATACACCGAGTCTGCAGATGAGG |
| 4_known | J3 | Tb927.10.13360_for | GGGGACAAGTTTGTACAAAAAAGCAGGCTATGTTTCGCGCTGCCTTTCG | Tb927.10.13360_rev | GGGGACCACTTTGTACAAGAAAGCTGGGTTTACTTCTTTTTCTTGCTCTTCCCCG |
| 4_known | J4 | Tb927.8.8300_for | GGGGACAAGTTTGTACAAAAAAGCAGGCTATGGACAGATCTGGCAGTCAATCC | Tb927.8.8300_rev | GGGGACCACTTTGTACAAGAAAGCTGGGTTCAACCCACAGCAGCGCC |
| 4_known | J5 | Tb927.5.2720_for | GGGGACAAGTTTGTACAAAAAAGCAGGCTATGGCTGAAACACCAAAGGAGATTC | Tb927.5.2720_rev | GGGGACCACTTTGTACAAGAAAGCTGGGTCTAGTTGACATTCATCTGATGGAAGCC |
| 4_known | J6 | Tb927.4.4020_for | GGGGACAAGTTTGTACAAAAAAGCAGGCTATGTGTATTGCCAGAGAAAATACAAGCA | Tb927.4.4020_rev | GGGGACCACTTTGTACAAGAAAGCTGGGTTTAACCAGTTATAGTGCCCCATATACTTGC |
| 4_known | J7 | Tb927.9.5170_for | GGGGACAAGTTTGTACAAAAAAGCAGGCTATGGTCGTGGCATCGTGTCC | Tb927.9.5170_rev | GGGGACCACTTTGTACAAGAAAGCTGGGTTTAATCAAGAACAAGCCCATTTGCATC |
| 4_known | J8 | Tb927.10.2010_for | GGGGACAAGTTTGTACAAAAAAGCAGGCTATGTCTAGACGCCTAAACAATATCCTCG | Tb927.10.2010_rev | GGGGACCACTTTGTACAAGAAAGCTGGGTTTACTTGTCGTTCACCACCATTGC |
| 4_known | J9 | Tb927.4.1310_for | GGGGACAAGTTTGTACAAAAAAGCAGGCTATGTTCGGTGGTGGCGAAGA | Tb927.4.1310_rev | GGGGACCACTTTGTACAAGAAAGCTGGGTTCAAAACTCTTTTGTCATAGACACGAGA |
| 4_known | J10 | Tb927.8.8290_for | GGGGACAAGTTTGTACAAAAAAGCAGGCTATGAGTAATGTCCGTGGAAATATAACCC | Tb927.8.8290_rev | GGGGACCACTTTGTACAAGAAAGCTGGGTTTAGCCAACAATTGTGCCCCAA |
| 4_known | J11 | Tb927.8.7460_for | GGGGACAAGTTTGTACAAAAAAGCAGGCTATGGAAGAGGCAAGTAGCGAGC | Tb927.8.7460_rev | GGGGACCACTTTGTACAAGAAAGCTGGGTTTAGGCGATAGTTTGTGCCGAGG |
| 4_known | J12 | Tb927.11.5850_for | GGGGACAAGTTTGTACAAAAAAGCAGGCTATGCCTGTGCTTCCGGAGT | Tb927.11.5850_rev | GGGGACCACTTTGTACAAGAAAGCTGGGTTCACAAGGGGCGACCACG |
| 4_known | J13 | Tb927.10.2020_for | GGGGACAAGTTTGTACAAAAAAGCAGGCTATGTCTAGACGCCTAAACAATATCCTCG | Tb927.10.2020_rev | GGGGACCACTTTGTACAAGAAAGCTGGGTTCACTTCCCGTCAGCAACGA |
| 4_known | J14 | Tb927.10.15940_for | GGGGACAAGTTTGTACAAAAAAGCAGGCTATGCGGCGACGGCTCCAT | Tb927.10.15940_rev | GGGGACCACTTTGTACAAGAAAGCTGGGTTCAACAACAACCCACTGAAGGATG |
| 4_known | J15 | Tb927.10.15480_for | GGGGACAAGTTTGTACAAAAAAGCAGGCTATGGCAATCTGGGCATGTATTGC | Tb927.10.15480_rev | GGGGACCACTTTGTACAAGAAAGCTGGGTTTATTCCTCCAAGACGCACAAATAAACC |
| 4_known | J16 | Tb927.9.7550_for | GGGGACAAGTTTGTACAAAAAAGCAGGCTATGGAGAAGGGTTCACCATCCG | Tb927.9.7550_rev | GGGGACCACTTTGTACAAGAAAGCTGGGTTTATTTAGCACTGAAGCGAGGAACG |
| 4_known | J17 | Tb927.11.950_for | GGGGACAAGTTTGTACAAAAAAGCAGGCTATGCAGGAAGGCCAACTGTTTC | Tb927.11.950_rev | GGGGACCACTTTGTACAAGAAAGCTGGGTTTAGTACCCCTTGTCGCTTGCC |
| 4_known | J18 | Tb927.3.3290_for | GGGGACAAGTTTGTACAAAAAAGCAGGCTATGTTCACGACAATGGAGAGCG | Tb927.3.3290_rev | GGGGACCACTTTGTACAAGAAAGCTGGGTTCACTTCTTCTTCAGAAGTAAACTTAACAT |
| 4_known | J19 | Tb927.10.13670_for | GGGGACAAGTTTGTACAAAAAAGCAGGCTATGGCCGGGGTAGAGGAAGC | Tb927.10.13670_rev | GGGGACCACTTTGTACAAGAAAGCTGGGTTCAAAACAGGCCAAATCCTGTGG |
| 4_known | J20 | Tb927.10.7430_for | GGGGACAAGTTTGTACAAAAAAGCAGGCTATGCTTCAACCCTGCCTGC | Tb927.10.7430_rev | GGGGACCACTTTGTACAAGAAAGCTGGGTCACTGCCACTTCTCACTGGT |
| 4_known | J21 | Tb927.11.6320_for | GGGGACAAGTTTGTACAAAAAAGCAGGCTATGCAGGGGACTGGATCCG | Tb927.11.6320_rev | GGGGACCACTTTGTACAAGAAAGCTGGGTTTATTTACGCCGGTGAGAACGAAC |
| 4_known | J22 | Tb927.4.4730_for | GGGGACAAGTTTGTACAAAAAAGCAGGCTATGACATTTGAAGCTGCTGCACA | Tb927.4.4730_rev | GGGGACCACTTTGTACAAGAAAGCTGGGTTTAAGAATACCTCTTAATTGTGTAGTAAAC |
| 4_known | J23 | Tb927.4.4840_for | GGGGACAAGTTTGTACAAAAAAGCAGGCTATGGCGTCCGCAGTGGTC | Tb927.4.4840_rev | GGGGACCACTTTGTACAAGAAAGCTGGGTTTACCAGCCCACAGCCACT |
| 4_known | J24 | Tb927.8.5730_for | GGGGACAAGTTTGTACAAAAAAGCAGGCTATGAGCGACGAGGGGGTTTGA | Tb927.8.5730_rev | GGGGACCACTTTGTACAAGAAAGCTGGGTCTACAGGCGGTAGACACTCTTCA |
| 4_known | K1 | Tb927.8.2240_for | GGGGACAAGTTTGTACAAAAAAGCAGGCTATGCGGCGGGCTTCGCAT | Tb927.8.2240_rev | GGGGACCACTTTGTACAAGAAAGCTGGGTCTACGCCATAATGTTGCGCACC |
| 4_known | K2 | Tb927.9.15795_for | GGGGACAAGTTTGTACAAAAAAGCAGGCTATGCGGAGAAGTATGTACTGGGT | Tb927.9.15795_rev | GGGGACCACTTTGTACAAGAAAGCTGGGTTTATACCAAAAGCAAAATTAATCCCAGTGG |
| 4_known | K3 | Tb927.7.2570_for | GGGGACAAGTTTGTACAAAAAAGCAGGCTATGCTTCGCTTATTGCGGCG | Tb927.7.2570_rev | GGGGACCACTTTGTACAAGAAAGCTGGGTTTACAACTTCGCCTCACAGCC |
| 4_known | K5 | Tb927.4.4860_for | GGGGACAAGTTTGTACAAAAAAGCAGGCTATGGCGTCCGCAGTGGTC | Tb927.4.4860_rev | GGGGACCACTTTGTACAAGAAAGCTGGGTTTACCAACCCACAGCCACTCC |
| 4_known | K6 | Tb927.8.3130_for | GGGGACAAGTTTGTACAAAAAAGCAGGCTATGAAAAGAATCGTTGTACCCCATCG | Tb927.8.3130_rev | GGGGACCACTTTGTACAAGAAAGCTGGGTTCATTTGTCATCCAGCCCCACG |
| 4_known | K7 | Tb927.4.2000_for | GGGGACAAGTTTGTACAAAAAAGCAGGCTATGGCTGAGACAAATATACGTGCAG | Tb927.4.2000_rev | GGGGACCACTTTGTACAAGAAAGCTGGGTTCACTGCTCTTCCTCATTCTGACA |
| 4_known | K8 | Tb927.8.6410_for | GGGGACAAGTTTGTACAAAAAAGCAGGCTATGCATCTTGTGAGGAAGGGGA | Tb927.8.6410_rev | GGGGACCACTTTGTACAAGAAAGCTGGGTTCAGTTCGCACTGCTAACCTTAC |
| 4_known | K9 | Tb927.10.12740_for | GGGGACAAGTTTGTACAAAAAAGCAGGCTATGACGATGGATATTATCGAGGGGG | Tb927.10.12740_rev | GGGGACCACTTTGTACAAGAAAGCTGGGTTTACCTCGCGTATGGCCAATTTAAA |
| 4_known | K10 | Tb927.10.12260_for | GGGGACAAGTTTGTACAAAAAAGCAGGCTATGTCGGTGGATTGGAATACCGT | Tb927.10.12260_rev | GGGGACCACTTTGTACAAGAAAGCTGGGTCTAATCCTTTCGCAACTTTGGCG |
| 4_known | K11 | Tb927.8.4390_for | GGGGACAAGTTTGTACAAAAAAGCAGGCTATGGGGAAACGATCGATTCGGA | Tb927.8.4390_rev | GGGGACCACTTTGTACAAGAAAGCTGGGTTCAGTCCTCAGGACTGTAGTTTTCC |
| 4_known | K12 | Tb927.10.10650_for | GGGGACAAGTTTGTACAAAAAAGCAGGCTATGTCGGCCAACAATTTCTCCG | Tb927.10.10650_rev | GGGGACCACTTTGTACAAGAAAGCTGGGTTCACCATTTCAAACTATGCACCCC |
| 4_known | K13 | Tb927.5.1520_for | GGGGACAAGTTTGTACAAAAAAGCAGGCTATGATGCGGCGAGTAACTTCTTC | Tb927.5.1520_rev | GGGGACCACTTTGTACAAGAAAGCTGGGTCTAGAGGATGAATTTCTTGATGTCCACC |
| 4_known | K14 | Tb927.6.400_for | GGGGACAAGTTTGTACAAAAAAGCAGGCTATGCAGGTTGATTGGGGTGC | Tb927.6.400_rev | GGGGACCACTTTGTACAAGAAAGCTGGGTTCACTGCTTATGTATCTTTGGCGTT |
| 4_known | K15 | Tb927.6.1900_for | GGGGACAAGTTTGTACAAAAAAGCAGGCTATGCCGGGACGCGAAAAG | Tb927.6.1900_rev | GGGGACCACTTTGTACAAGAAAGCTGGGTTCAACTGTTCCCTCCTTTCGCC |
| 4_known | K16 | Tb927.1.4650_for | GGGGACAAGTTTGTACAAAAAAGCAGGCTATGTCCTTGGAGGAGGGAAGC | Tb927.1.4650_rev | GGGGACCACTTTGTACAAGAAAGCTGGGTCTACATCTCAGCGTAATCCACTTCATG |
| 4_known | K17 | Tb927.6.1580_for | GGGGACAAGTTTGTACAAAAAAGCAGGCTATGAAGCGCGCAAGGAGC | Tb927.6.1580_rev | GGGGACCACTTTGTACAAGAAAGCTGGGTTCATGAGAGCAGGAAAAACTCCCT |
| 4_known | K18 | Tb927.11.13250_for | GGGGACAAGTTTGTACAAAAAAGCAGGCTATGAGCGAGAATGACATTGCCG | Tb927.11.13250_rev | GGGGACCACTTTGTACAAGAAAGCTGGGTTTAATTCAGTTTCACTGGCACTCCAC |
| 4_known | K19 | Tb927.4.1350_for | GGGGACAAGTTTGTACAAAAAAGCAGGCTATGATGGAAAGCGTGAAGCAGTT | Tb927.4.1350_rev | GGGGACCACTTTGTACAAGAAAGCTGGGTCTACCCCGTGTCGTTCCGC |

| | | | | | |
|---|---|---|---|---|---|
| 4_known | K20 | Tb927.3.5230_for | GGGGACAAGTTTGTACAAAAAAGCAGGCTATGGCTAGCGTGAATGCTCCC | Tb927.3.5230_rev | GGGGACCACTTTGTACAAGAAAGCTGGGTTCATTCGTCGTTGCATATACCTTCG |
| 4_known | K21 | Tb927.11.16730_for | GGGGACAAGTTTGTACAAAAAAGCAGGCTATGTTCCGTCGCTGCTTTCC | Tb927.11.16730_rev | GGGGACCACTTTGTACAAGAAAGCTGGGTTTAGAAGTTGATTGTTTTGGCAAAGCA |
| 4_known | K22 | Tb927.3.1110_for | GGGGACAAGTTTGTACAAAAAAGCAGGCTATGTCTTTTAGAGAGCTTCGTTCCTTTG | Tb927.3.1110_rev | GGGGACCACTTTGTACAAGAAAGCTGGGTCTAGAGGTCGCTGCTGCTTTC |
| 4_known | K23 | Tb927.10.4730_for | GGGGACAAGTTTGTACAAAAAAGCAGGCTATGTTGGGTGTGAGGGCGT | Tb927.10.4730_rev | GGGGACCACTTTGTACAAGAAAGCTGGGTCTATGAGTGGCTATCTTTCATTTTGCG |
| 4_known | K24 | Tb927.9.4520_for | GGGGACAAGTTTGTACAAAAAAGCAGGCTATGCTTCGTCCATCCTTTTGCA | Tb927.9.4520_rev | GGGGACCACTTTGTACAAGAAAGCTGGGTTCAAAGTATGTGCTTTACTTGTGTGAGG |
| 4_known | L1 | Tb927.6.2060_for | GGGGACAAGTTTGTACAAAAAAGCAGGCTATGGCAAAGACAGTAGATAACGCC | Tb927.6.2060_rev | GGGGACCACTTTGTACAAGAAAGCTGGGTCTACACGATTTTATCAACTGGAAGAACG |
| 4_known | L2 | Tb927.10.4520_for | GGGGACAAGTTTGTACAAAAAAGCAGGCTATGACTCAGTCAACGATCCCTTTGC | Tb927.10.4520_rev | GGGGACCACTTTGTACAAGAAAGCTGGGTTCACCCGACGGAGGTGCT |
| 4_known | L3 | Tb927.3.4380_for | GGGGACAAGTTTGTACAAAAAAGCAGGCTATGACCTTTTCTAGCGAGAGTAGTGT | Tb927.3.4380_rev | GGGGACCACTTTGTACAAGAAAGCTGGGTTCACATGGAAAAAGCGGATGACC |
| 4_known | L4 | Tb927.11.7170_for | GGGGACAAGTTTGTACAAAAAAGCAGGCTATGGTGCTTGATATACAGCTGTTTCG | Tb927.11.7170_rev | GGGGACCACTTTGTACAAGAAAGCTGGGTTCACTCCCCCTTGTCGGGT |
| 4_known | L5 | Tb927.9.12110_for | GGGGACAAGTTTGTACAAAAAAGCAGGCTATGTCAATGGATGTCGGTGTTGTC | Tb927.9.12110_rev | GGGGACCACTTTGTACAAGAAAGCTGGGTTTATTGCAACTCAGGCCATTGGAA |
| 4_known | L6 | Tb927.10.14150_for | GGGGACAAGTTTGTACAAAAAAGCAGGCTATGAGCAAGACAGAGACGGCA | Tb927.10.14150_rev | GGGGACCACTTTGTACAAGAAAGCTGGGTTTACGCAAAATCCTCCCCCTCT |
| 4_known | L7 | Tb927.6.3780_for | GGGGACAAGTTTGTACAAAAAAGCAGGCTATGAACGTTAATTCGTGTGGCAATATC | Tb927.6.3780_rev | GGGGACCACTTTGTACAAGAAAGCTGGGTTTAATCCACTTCCTCCATACGCAGC |
| 4_known | L8 | Tb927.8.3580_for | GGGGACAAGTTTGTACAAAAAAGCAGGCTATGAATCTTGCTGCTCTCGCC | Tb927.8.3580_rev | GGGGACCACTTTGTACAAGAAAGCTGGGTTCATTGCAAGCTGTGTAACATCGG |
| 4_known | L9 | Tb927.1.1150_for | GGGGACAAGTTTGTACAAAAAAGCAGGCTATGGCAATTGTTGTCGTGCAGG | Tb927.1.1150_rev | GGGGACCACTTTGTACAAGAAAGCTGGGTTCAATCCTCCTCCGCATCAGC |
| 4_known | L10 | Tb927.1.1230_for | GGGGACAAGTTTGTACAAAAAAGCAGGCTATGCGAGGCATTACATTGGCG | Tb927.1.1230_rev | GGGGACCACTTTGTACAAGAAAGCTGGGTTTACGTACCCGGCAGCCA |
| 4_known | L11 | Tb927.4.1620_for | GGGGACAAGTTTGTACAAAAAAGCAGGCTATGCCGTTCAATAGAGAAATACTTGCC | Tb927.4.1620_rev | GGGGACCACTTTGTACAAGAAAGCTGGGTTTACACAATAATGGACGAGACAAAACTGA |
| 4_known | L12 | Tb927.7.940_for | GGGGACAAGTTTGTACAAAAAAGCAGGCTATGACCGATATGTTGTTTGCTGTTTTATTG | Tb927.7.940_rev | GGGGACCACTTTGTACAAGAAAGCTGGGTTTACTCGCATATAGCGGGTGTTTG |
| 4_known | L13 | Tb927.8.8310_for | GGGGACAAGTTTGTACAAAAAAGCAGGCTATGAAGAAAGAATCCGTTCCTTCCAAA | Tb927.8.8310_rev | GGGGACCACTTTGTACAAGAAAGCTGGGTTTATTGTGCTTCAATTACCACACTGGA |
| 4_known | L14 | Tb927.11.6210_for | GGGGACAAGTTTGTACAAAAAAGCAGGCTATGTTGCTTGAAGTTGCCATTTTCTT | Tb927.11.6210_rev | GGGGACCACTTTGTACAAGAAAGCTGGGTCTAAGCAGCTGCCGCCTTC |
| 4_known | L15 | Tb927.11.6230_for | GGGGACAAGTTTGTACAAAAAAGCAGGCTATGACCGACATCCTCATGAAGATGA | Tb927.11.6230_rev | GGGGACCACTTTGTACAAGAAAGCTGGGTTCACGCAACCTTCCGCTTG |
| 4_known | L16 | Tb927.4.2480_for | GGGGACAAGTTTGTACAAAAAAGCAGGCTATGCGCGATTGGCATGCG | Tb927.4.2480_rev | GGGGACCACTTTGTACAAGAAAGCTGGGTTCATGCTGTTGCTTCCTTGTTCC |
| 4_known | L17 | Tb927.10.12400_for | GGGGACAAGTTTGTACAAAAAAGCAGGCTATGACAGCCAAGCCTGCATC | Tb927.10.12400_rev | GGGGACCACTTTGTACAAGAAAGCTGGGTTCAGTTAGTGGAAAGCAGGGCC |
| 4_known | L18 | Tb927.9.9870_for | GGGGACAAGTTTGTACAAAAAAGCAGGCTATGCCCAACATACCGTTAGTGGC | Tb927.9.9870_rev | GGGGACCACTTTGTACAAGAAAGCTGGGTTCATCTCCTCTGCGTAGCACA |
| 4_known | L19 | Tb927.10.1550_for | GGGGACAAGTTTGTACAAAAAAGCAGGCTATGACCGACGTAGACCAGTGG | Tb927.10.1550_rev | GGGGACCACTTTGTACAAGAAAGCTGGGTCTATGAAGCAGTCGCACCCG |
| 4_known | L20 | Tb927.7.610_for | GGGGACAAGTTTGTACAAAAAAGCAGGCTATGCGGTTTAGTCGCCCATG | Tb927.7.610_rev | GGGGACCACTTTGTACAAGAAAGCTGGGTTTAAACCCAATCACTACCAGTCTTTGTG |
| 4_known | L21 | Tb927.9.11810_for | GGGGACAAGTTTGTACAAAAAAGCAGGCTATGGTGACCATGGAAGGAACCC | Tb927.9.11810_rev | GGGGACCACTTTGTACAAGAAAGCTGGGTCTATGTGAGTTCGTCCTCATCCACC |
| 4_known | L22 | Tb927.11.16860_for | GGGGACAAGTTTGTACAAAAAAGCAGGCTATGAGTAACCCTTTTGAGAAAGTTGCG | Tb927.11.16860_rev | GGGGACCACTTTGTACAAGAAAGCTGGGTCTACTTACCACCATCTTGCGACAAC |
| 4_known | L23 | Tb927.9.15980_for | GGGGACAAGTTTGTACAAAAAAGCAGGCTATGCTTGGCTTCGGTTCTGTG | Tb927.9.15980_rev | GGGGACCACTTTGTACAAGAAAGCTGGGTTTACTGTGACTCATCTTTCGGGAGA |
| 4_known | L24 | Tb927.11.5350_for | GGGGACAAGTTTGTACAAAAAAGCAGGCTATGAAGTGGCGCGTGAAGATATT | Tb927.11.5350_rev | GGGGACCACTTTGTACAAGAAAGCTGGGTCTAAATGGGCGCACGGGC |
| 4_known | M1 | Tb927.8.3750_for | GGGGACAAGTTTGTACAAAAAAGCAGGCTATGTCAAAAACACTTTATCTCTTATATGAG | Tb927.8.3750_rev | GGGGACCACTTTGTACAAGAAAGCTGGGTCTATGCCGATTCTTGTGGCACT |
| 4_known | M2 | Tb927.10.13430_for | GGGGACAAGTTTGTACAAAAAAGCAGGCTATGTGCATGCGTGCTCGT | Tb927.10.13430_rev | GGGGACCACTTTGTACAAGAAAGCTGGGTCTACGCTATGTTGTACTTTGTGAGAAGT |
| 4_known | M3 | Tb927.11.12420_for | GGGGACAAGTTTGTACAAAAAAGCAGGCTATGGCTTCATACATAGCGTCAAACG | Tb927.11.12420_rev | GGGGACCACTTTGTACAAGAAAGCTGGGTTTACATGACAGGTGTTGGGGGG |
| 4_known | M4 | Tb927.10.13130_for | GGGGACAAGTTTGTACAAAAAAGCAGGCTATGCCGCTAAACCCTCCTTCA | Tb927.10.13130_rev | GGGGACCACTTTGTACAAGAAAGCTGGGTCTACTCGACTACCACAACCTCATTG |
| 4_known | M5 | Tb927.4.3770_for | GGGGACAAGTTTGTACAAAAAAGCAGGCTATGTCATACACAACAGAAAGTGCCTT | Tb927.4.3770_rev | GGGGACCACTTTGTACAAGAAAGCTGGGTCTATTTACAGCCGGAAACCAAAGCT |
| 4_known | M6 | Tb927.10.13620_for | GGGGACAAGTTTGTACAAAAAAGCAGGCTATGTTGCGCCGTGTAGGAGA | Tb927.10.13620_rev | GGGGACCACTTTGTACAAGAAAGCTGGGTCTAATGAATAGCCACAGGTCCGGG |
| 4_known | M7 | Tb927.10.15250_for | GGGGACAAGTTTGTACAAAAAAGCAGGCTATGGCAAGTTCTACAGCAACCG | Tb927.10.15250_rev | GGGGACCACTTTGTACAAGAAAGCTGGGTTTACATCTCTGGGTTCTCCTCGC |
| 4_known | M8 | Tb927.1.1080_for | GGGGACAAGTTTGTACAAAAAAGCAGGCTATGAGTCAACTCAGCGAGTTGC | Tb927.1.1080_rev | GGGGACCACTTTGTACAAGAAAGCTGGGTTCAAGCATCTTTCGGAAAAGAGCG |
| 4_known | M9 | Tb927.11.2640_for | GGGGACAAGTTTGTACAAAAAAGCAGGCTATGGGTGGACAGTCTAAGAAGCA | Tb927.11.2640_rev | GGGGACCACTTTGTACAAGAAAGCTGGGTTTACTGCACACCACTTTTCATCTTACT |
| 4_known | M10 | Tb927.8.1270_for | GGGGACAAGTTTGTACAAAAAAGCAGGCTATGGCCGTCACATCAAAGGC | Tb927.8.1270_rev | GGGGACCACTTTGTACAAGAAAGCTGGGTCTACTCCTCAGCAGACCCCTTT |
| 4_known | M11 | Tb927.11.8400_for | GGGGACAAGTTTGTACAAAAAAGCAGGCTATGATTCGCTTGAGCGAAGTTGT | Tb927.11.8400_rev | GGGGACCACTTTGTACAAGAAAGCTGGGTTCAGGGCGGCTGCCCACC |
| 4_known | M12 | Tb927.3.3270_for | GGGGACAAGTTTGTACAAAAAAGCAGGCTATGGCCGTTGAATCTCGCAG | Tb927.3.3270_rev | GGGGACCACTTTGTACAAGAAAGCTGGGTTTAGAGCTTGGCCAGTTCCTCG |
| 4_known | M13 | Tb927.8.7680_for | GGGGACAAGTTTGTACAAAAAAGCAGGCTATGCATCCTGACGGAACCACT | Tb927.8.7680_rev | GGGGACCACTTTGTACAAGAAAGCTGGGTTTACCACTCCACTGCGACACT |
| 4_known | M14 | Tb927.10.1140_for | GGGGACAAGTTTGTACAAAAAAGCAGGCTATGCCCCGTGAGGTTGTTACA | Tb927.10.1140_rev | GGGGACCACTTTGTACAAGAAAGCTGGGTTCAAAAGTAAACAAGGTCACGCATGC |
| 4_known | M15 | Tb927.11.5400_for | GGGGACAAGTTTGTACAAAAAAGCAGGCTATGGTGTTAGCAGAACTCGGACA | Tb927.11.5400_rev | GGGGACCACTTTGTACAAGAAAGCTGGGTTCACTTTATGCTCTGAAGTTGCTTCA |
| 4_known | M16 | Tb927.2.3880_for | GGGGACAAGTTTGTACAAAAAAGCAGGCTATGGAACAGTCTCCTGAGCAGC | Tb927.2.3880_rev | GGGGACCACTTTGTACAAGAAAGCTGGGTTTAACAGAACATCTGTGGTACCATTGG |
| 4_known | M17 | Tb927.7.1550_for | GGGGACAAGTTTGTACAAAAAAGCAGGCTATGTTGATGCATACCGCGCC | Tb927.7.1550_rev | GGGGACCACTTTGTACAAGAAAGCTGGGTTCATTCACGGTCTCTGTCACGC |
| 4_known | M18 | Tb927.11.8170_for | GGGGACAAGTTTGTACAAAAAAGCAGGCTATGGAAGACTACGAAACCGTGGC | Tb927.11.8170_rev | GGGGACCACTTTGTACAAGAAAGCTGGGTTCACCGCTTCGAGCCATTG |
| 4_known | M20 | Tb927.7.690_for | GGGGACAAGTTTGTACAAAAAAGCAGGCTATGCAAAAGAAGAACTGCAGGTGG | Tb927.7.690_rev | GGGGACCACTTTGTACAAGAAAGCTGGGTTTACACATTCTGTTGTAACCAATCACGA |
| 4_known | M21 | Tb927.7.1440_for | GGGGACAAGTTTGTACAAAAAAGCAGGCTATGAGCGCAAGCGACGGA | Tb927.7.1440_rev | GGGGACCACTTTGTACAAGAAAGCTGGGTTTACACCCCAATGGGATTTCCTACG |
| 4_known | M22 | Tb927.8.5450_for | GGGGACAAGTTTGTACAAAAAAGCAGGCTATGAGAGAGCCGATACAAACTTCAAC | Tb927.8.5450_rev | GGGGACCACTTTGTACAAGAAAGCTGGGTTCAGAGTTCAGCAATGACGCTG |
| 4_known | M23 | Tb927.9.5260_for | GGGGACAAGTTTGTACAAAAAAGCAGGCTATGCTTTCCCTATATGCGTACATTTTCC | Tb927.9.5260_rev | GGGGACCACTTTGTACAAGAAAGCTGGGTCTAGGAGGCATCAATGGATTTCAGAC |
| 4_known | M24 | Tb927.10.4640_for | GGGGACAAGTTTGTACAAAAAAGCAGGCTATGGCGTCGAACCCTGATTCA | Tb927.10.4640_rev | GGGGACCACTTTGTACAAGAAAGCTGGGTTTAGTTAGTATTGTGGGCATCCGGT |
| 4_known | N1 | Tb927.9.1570_for | GGGGACAAGTTTGTACAAAAAAGCAGGCTATGCGCGGGGAGCATGAA | Tb927.9.1570_rev | GGGGACCACTTTGTACAAGAAAGCTGGGTTTAGTCCCAGACAACAGACTTCAAGC |
| 4_known | N2 | Tb927.2.5150_for | GGGGACAAGTTTGTACAAAAAAGCAGGCTATGGATTACGCCGCCGCA | Tb927.2.5150_rev | GGGGACCACTTTGTACAAGAAAGCTGGGTCTATATAACCCCAAGTTGCCGCC |
| 4_known | N3 | Tb927.10.7510_for | GGGGACAAGTTTGTACAAAAAAGCAGGCTATGGCGCCTTATGCTGTGG | Tb927.10.7510_rev | GGGGACCACTTTGTACAAGAAAGCTGGGTTCAAATAACAGCTAAAAAGTGGTCGGT |
| 4_known | N4 | Tb927.11.10930_for | GGGGACAAGTTTGTACAAAAAAGCAGGCTATGGCATGTGTGCATGTCCTG | Tb927.11.10930_rev | GGGGACCACTTTGTACAAGAAAGCTGGGTTCAAGCCGCCCCGTATGC |
| 4_known | N5 | Tb927.9.13490_for | GGGGACAAGTTTGTACAAAAAAGCAGGCTATGTCGTCAAATTGTATTCGCAACAC | Tb927.9.13490_rev | GGGGACCACTTTGTACAAGAAAGCTGGGTTCAAGGGGCATTTTTTTGGTTAACG |
| 4_known | N6 | Tb927.5.1060_for | GGGGACAAGTTTGTACAAAAAAGCAGGCTATGTCTGTCAGTTTTTCCACTCTGATT | Tb927.5.1060_rev | GGGGACCACTTTGTACAAGAAAGCTGGGTTTAATACCAGTACTTGTAGGACCAGTGT |
| 4_known | N7 | Tb927.10.5930_for | GGGGACAAGTTTGTACAAAAAAGCAGGCTATGTCGTCAACGGATGCAATGA | Tb927.10.5930_rev | GGGGACCACTTTGTACAAGAAAGCTGGGTTTACAATCCAAGAACCTCCTGTATTTTCTG |
| 4_known | N8 | Tb927.3.1700_for | GGGGACAAGTTTGTACAAAAAAGCAGGCTATGCTTCAAAAGATCAGCGCTGA | Tb927.3.1700_rev | GGGGACCACTTTGTACAAGAAAGCTGGGTCTATCGTATAACTAACCCTTTTGAATCCGC |
| 4_known | N9 | Tb927.9.10010_for | GGGGACAAGTTTGTACAAAAAAGCAGGCTATGGTGGAAAGTGGAATTGGTGG | Tb927.9.10010_rev | GGGGACCACTTTGTACAAGAAAGCTGGGTCTAGAGCCGGCGCAGGCT |
| 4_known | N10 | Tb927.11.9360_for | GGGGACAAGTTTGTACAAAAAAGCAGGCTATGTTAAGGCGTACAATTTTCCTGGG | Tb927.11.9360_rev | GGGGACCACTTTGTACAAGAAAGCTGGGTCTAAACGGCGATGCTGGAGAAC |
| 4_known | N11 | Tb927.4.1020_for | GGGGACAAGTTTGTACAAAAAAGCAGGCTATGGTGGGATTTAAGACTCTTTCATCCA | Tb927.4.1020_rev | GGGGACCACTTTGTACAAGAAAGCTGGGTTCAAACAAAAACACTCTTCACCGC |
| 4_known | N12 | Tb927.7.6930_for | GGGGACAAGTTTGTACAAAAAAGCAGGCTATGTTTTGGCGCTTCCCTGC | Tb927.7.6930_rev | GGGGACCACTTTGTACAAGAAAGCTGGGTTCACGCTCCGGTACTTAATGACA |
| 4_known | N13 | Tb927.8.1010_for | GGGGACAAGTTTGTACAAAAAAGCAGGCTATGAGTTATGGCAAGTGGGCTG | Tb927.8.1010_rev | GGGGACCACTTTGTACAAGAAAGCTGGGTTTACAACACCCGGTTCTCAGCT |
| 4_known | N14 | Tb927.9.4850_for | GGGGACAAGTTTGTACAAAAAAGCAGGCTATGCTTGAAGGAAGTGATGCGAG | Tb927.9.4850_rev | GGGGACCACTTTGTACAAGAAAGCTGGGTTCATCGAAGCAGATTCACTCCCC |
| 4_known | N15 | Tb927.5.2080_for | GGGGACAAGTTTGTACAAAAAAGCAGGCTATGTCCTTCAATGAATCGGCATCC | Tb927.5.2080_rev | GGGGACCACTTTGTACAAGAAAGCTGGGTTTAAAGTTTGGCAACACCGTGACT |
| 4_known | N16 | Tb927.10.9440_for | GGGGACAAGTTTGTACAAAAAAGCAGGCTATGATATGCCGCACATCTTTTCTGC | Tb927.10.9440_rev | GGGGACCACTTTGTACAAGAAAGCTGGGTCTACTCGTTCTGTTTCTTTGTCGCG |
| 4_known | N17 | Tb927.1.4300_for | GGGGACAAGTTTGTACAAAAAAGCAGGCTATGGCTCATATATCCGTTACTGACACC | Tb927.1.4300_rev | GGGGACCACTTTGTACAAGAAAGCTGGGTTTAACGACGACTCGTAGCCGC |
| 4_known | N18 | Tb927.10.8070_for | GGGGACAAGTTTGTACAAAAAAGCAGGCTATGAGAGCTTCGTTGTGTACCCG | Tb927.10.8070_rev | GGGGACCACTTTGTACAAGAAAGCTGGGTCTACTCCAGAAGGGCAGCGA |
| 4_known | N19 | Tb927.2.3800_for | GGGGACAAGTTTGTACAAAAAAGCAGGCTATGCTGCGCGCGCGCCTG | Tb927.2.3800_rev | GGGGACCACTTTGTACAAGAAAGCTGGGTCTAGTATGCCGAAACGGCAGTC |
| 4_known | N20 | Tb927.9.10340_for | GGGGACAAGTTTGTACAAAAAAGCAGGCTATGCACATGTCTAATGTCACCGAC | Tb927.9.10340_rev | GGGGACCACTTTGTACAAGAAAGCTGGGTTCATTTGAGAGACTCATAAGCTTGCAT |
| 4_known | N21 | Tb927.10.10390_for | GGGGACAAGTTTGTACAAAAAAGCAGGCTATGTCCAAGGCCTTCGATTTGG | Tb927.10.10390_rev | GGGGACCACTTTGTACAAGAAAGCTGGGTTCACAGGTTAGAGTCCGGAAGC |
| 4_known | N22 | Tb927.11.4800_for | GGGGACAAGTTTGTACAAAAAAGCAGGCTATGAACTCTAAAGACACGAATGAGTTGAA | Tb927.11.4800_rev | GGGGACCACTTTGTACAAGAAAGCTGGGTCTACTGGCCTCCCTTCTGGC |
| 4_known | N23 | Tb927.1.1700_for | GGGGACAAGTTTGTACAAAAAAGCAGGCTATGTCTACACGTAGCGGTAAGCA | Tb927.1.1700_rev | GGGGACCACTTTGTACAAGAAAGCTGGGTTTATTGGAATAAAGACTTGAACAACACCTC |
| 4_known | N24 | Tb927.11.2690_for | GGGGACAAGTTTGTACAAAAAAGCAGGCTATGCTCCGCCGAACAAATTTTTTC | Tb927.11.2690_rev | GGGGACCACTTTGTACAAGAAAGCTGGGTCTACGTGAGGGGCGCCAA |
| 4_known | O1 | Tb927.11.13880_for | GGGGACAAGTTTGTACAAAAAAGCAGGCTATGTCGCGCGATCCGCCA | Tb927.11.13880_rev | GGGGACCACTTTGTACAAGAAAGCTGGGTTCAGAGCACATCTGCAGACTCT |
| 4_known | O2 | Tb927.10.5950_for | GGGGACAAGTTTGTACAAAAAAGCAGGCTATGTCGTCAACGGATGCAATGA | Tb927.10.5950_rev | GGGGACCACTTTGTACAAGAAAGCTGGGTTCACTCAGCATCACCAATGCC |
| 4_known | O4 | Tb927.5.3760_for | GGGGACAAGTTTGTACAAAAAAGCAGGCTATGCACAGAGCCGCGTATG | Tb927.5.3760_rev | GGGGACCACTTTGTACAAGAAAGCTGGGTCTAAATGTAGTCCCGCCTAAATAATCCC |
| 4_known | O5 | Tb927.11.15960_for | GGGGACAAGTTTGTACAAAAAAGCAGGCTATGTCTGACAAAGGCAAGTGTGC | Tb927.11.15960_rev | GGGGACCACTTTGTACAAGAAAGCTGGGTTCAGCAGACAGGCGGGGG |
| 4_known | O6 | Tb927.11.11690_for | GGGGACAAGTTTGTACAAAAAAGCAGGCTATGGGCGGACATAATGAAGAAGTG | Tb927.11.11690_rev | GGGGACCACTTTGTACAAGAAAGCTGGGTTCAACGGTTGGCGGGTTG |
| 4_known | O7 | Tb927.4.3980_for | GGGGACAAGTTTGTACAAAAAAGCAGGCTATGTTTGTTTTATGTACAGTTGAAAACATG | Tb927.4.3980_rev | GGGGACCACTTTGTACAAGAAAGCTGGGTCTAAAGCCGTTGGGACGTCG |
| 4_known | O8 | Tb927.11.11750_for | GGGGACAAGTTTGTACAAAAAAGCAGGCTATGATGCGTCCACTCTCACTTGT | Tb927.11.11750_rev | GGGGACCACTTTGTACAAGAAAGCTGGGTTTAGACAGCCATCTTTTGTTTAAACATCAC |
| 4_known | O9 | Tb927.11.15950_for | GGGGACAAGTTTGTACAAAAAAGCAGGCTATGTCTGACAAAGGCAAGTGTGC | Tb927.11.15950_rev | GGGGACCACTTTGTACAAGAAAGCTGGGTTCAGCACTGAGTTTCCGGCG |
| 4_known | O10 | Tb927.11.14120_for | GGGGACAAGTTTGTACAAAAAAGCAGGCTATGAGTACCATGGAGAACACCATTCT | Tb927.11.14120_rev | GGGGACCACTTTGTACAAGAAAGCTGGGTCTAAAAACGCATTAGCTTTGAGTTCCTG |
| 4_known | O11 | Tb927.11.12230_for | GGGGACAAGTTTGTACAAAAAAGCAGGCTATGATTCGTTTCAGTTGGGTTCGC | Tb927.11.12230_rev | GGGGACCACTTTGTACAAGAAAGCTGGGTTCAGAGAATGTACTTTGCTAAGTCAATGT |
| 4_known | O12 | Tb927.5.450_for | GGGGACAAGTTTGTACAAAAAAGCAGGCTATGCTCCGGCGTGTGGGT | Tb927.5.450_rev | GGGGACCACTTTGTACAAGAAAGCTGGGTTTAGTTCCAATTCCGCACGATCTTT |
| 4_known | O13 | Tb927.10.8230_for | GGGGACAAGTTTGTACAAAAAAGCAGGCTATGCGCGCTATTTTTCTTGTTGC | Tb927.10.8230_rev | GGGGACCACTTTGTACAAGAAAGCTGGGTCTACAGATCTTGTTTATCAACATTGCTCTC |
| 4_known | O14 | Tb927.5.1750_for | GGGGACAAGTTTGTACAAAAAAGCAGGCTATGTACGCAAAGGGTACGCG | Tb927.5.1750_rev | GGGGACCACTTTGTACAAGAAAGCTGGGTTTAGCTGCGAGACACCGCT |
| 4_known | O15 | Tb927.8.6420_for | GGGGACAAGTTTGTACAAAAAAGCAGGCTATGCCATTCCAGCTAAGCCGT | Tb927.8.6420_rev | GGGGACCACTTTGTACAAGAAAGCTGGGTTTATCTGTCCCATTGCATATATGGTTGC |
| 4_known | O16 | Tb927.8.6110_for | GGGGACAAGTTTGTACAAAAAAGCAGGCTATGCGCCGAACTGTGCGT | Tb927.8.6110_rev | GGGGACCACTTTGTACAAGAAAGCTGGGTCTACTTCACGAACCGGTAATATTCGAT |
| 4_known | O17 | Tb927.10.4900_for | GGGGACAAGTTTGTACAAAAAAGCAGGCTATGGAGCCTGATGATGGTGTTGC | Tb927.10.4900_rev | GGGGACCACTTTGTACAAGAAAGCTGGGTTCACCTATAGTGAAATGTTTGCTTTTGC |
| 4_known | O18 | Tb927.8.3100_for | GGGGACAAGTTTGTACAAAAAAGCAGGCTATGGACGTATCGCGGTTTCG | Tb927.8.3100_rev | GGGGACCACTTTGTACAAGAAAGCTGGGTTTACGCTTGACTACCAGAGGCA |
| 4_known | O19 | Tb927.7.5740_for | GGGGACAAGTTTGTACAAAAAAGCAGGCTATGCTACGGTCCTTTAGAAGTAATGC | Tb927.7.5740_rev | GGGGACCACTTTGTACAAGAAAGCTGGGTTCAATTCATGAGATCAGTATCGCGTAAG |
| 4_known | O20 | Tb927.11.4610_for | GGGGACAAGTTTGTACAAAAAAGCAGGCTATGTCTGAAAAGGGAACATCGTTAAACC | Tb927.11.4610_rev | GGGGACCACTTTGTACAAGAAAGCTGGGTCTACTTCCTCCCCTCTGCCC |
| 4_known | O21 | Tb927.11.2660_for | GGGGACAAGTTTGTACAAAAAAGCAGGCTATGGGACAGGGGAAAAGCGA | Tb927.11.2660_rev | GGGGACCACTTTGTACAAGAAAGCTGGGTTTAGCGAGATGGTGTGGCTGG |
| 4_known | O22 | Tb927.1.3130_for | GGGGACAAGTTTGTACAAAAAAGCAGGCTATGGCCATCGCTCCCGCC | Tb927.1.3130_rev | GGGGACCACTTTGTACAAGAAAGCTGGGTTCAGCCATGTGCGCCAAAC |
| 4_known | O23 | Tb927.10.14140_for | GGGGACAAGTTTGTACAAAAAAGCAGGCTATGTCGCAGCTAGAGCACAAC | Tb927.10.14140_rev | GGGGACCACTTTGTACAAGAAAGCTGGGTTTAAGGAAGATAGATGAGGCGAGTTTGG |
| 4_known | O24 | Tb927.3.3080_for | GGGGACAAGTTTGTACAAAAAAGCAGGCTATGAAGCCGCGGGATATTTCC | Tb927.3.3080_rev | GGGGACCACTTTGTACAAGAAAGCTGGGTTCAACACAACCCATAATTTGTAACCTTGT |
| 4_known | P1 | Tb927.10.7520_for | GGGGACAAGTTTGTACAAAAAAGCAGGCTATGACTGCGTGGCACCGC | Tb927.10.7520_rev | GGGGACCACTTTGTACAAGAAAGCTGGGTTTAAGACCCTTGCGAGTCATACAAATT |
| 4_known | P2 | Tb927.10.5210_for | GGGGACAAGTTTGTACAAAAAAGCAGGCTATGAACCCGTCCTATTTGAATCAGATATTG | Tb927.10.5210_rev | GGGGACCACTTTGTACAAGAAAGCTGGGTTTAATCCGGAGTAATGAAATTGGACAGTAC |
| 4_known | P3 | Tb927.9.14220_for | GGGGACAAGTTTGTACAAAAAAGCAGGCTATGGACGCAGCTCTCGCT | Tb927.9.14220_rev | GGGGACCACTTTGTACAAGAAAGCTGGGTTCACTTGCGGCCGGGTTT |
| 4_known | P4 | Tb927.11.11740_for | GGGGACAAGTTTGTACAAAAAAGCAGGCTATGATGCGTCCACTCTCACTTGT | Tb927.11.11740_rev | GGGGACCACTTTGTACAAGAAAGCTGGGTTTACCTTACGGTGGGATCTTTCGG |
| 4_known | P5 | Tb927.2.3910_for | GGGGACAAGTTTGTACAAAAAAGCAGGCTATGGCATTTGGAACGTTCCCC | Tb927.2.3910_rev | GGGGACCACTTTGTACAAGAAAGCTGGGTTCAAAACTGTGTGAAGAGTAGCGC |
| 4_known | P6 | Tb927.10.7140_for | GGGGACAAGTTTGTACAAAAAAGCAGGCTATGCCTTACGCCTGCCTCC | Tb927.10.7140_rev | GGGGACCACTTTGTACAAGAAAGCTGGGTCTAGCCCTTCGCCCTCGT |
| 4_known | P7 | Tb927.8.3430_for | GGGGACAAGTTTGTACAAAAAAGCAGGCTATGGATTTCCTTCAGTTTTGCTCATCG | Tb927.8.3430_rev | GGGGACCACTTTGTACAAGAAAGCTGGGTCTAGAGTGGTCCCACTTGATGAAAC |
| 4_known | P8 | Tb927.7.2420_for | GGGGACAAGTTTGTACAAAAAAGCAGGCTATGAGTGAGCGGATTTTGCCG | Tb927.7.2420_rev | GGGGACCACTTTGTACAAGAAAGCTGGGTTTATATCGCATTCTCCGGCGC |
| 4_known | P9 | Tb927.10.3700_for | GGGGACAAGTTTGTACAAAAAAGCAGGCTATGTCTCAGTATGACCATTTATTTGATACC | Tb927.10.3700_rev | GGGGACCACTTTGTACAAGAAAGCTGGGTTCAGGTACCACAGCCGCA |
| 4_known | P10 | Tb927.11.2500_for | GGGGACAAGTTTGTACAAAAAAGCAGGCTATGAAGGCATACAAAGAGCTCGAG | Tb927.11.2500_rev | GGGGACCACTTTGTACAAGAAAGCTGGGTTCAGTTGGCATCGTCACGGT |
| 4_known | P11 | Tb927.9.11790_for | GGGGACAAGTTTGTACAAAAAAGCAGGCTATGATTGGCACTATGGAAGACGATG | Tb927.9.11790_rev | GGGGACCACTTTGTACAAGAAAGCTGGGTCTAATCCTTACTTCCAAGACTCTCAACAG |
| 4_known | P12 | Tb927.3.3190_for | GGGGACAAGTTTGTACAAAAAAGCAGGCTATGGCGCAGGTTGCAGATG | Tb927.3.3190_rev | GGGGACCACTTTGTACAAGAAAGCTGGGTCTACATCTTCTTCCCCTCGTACACG |
| 4_known | P13 | Tb927.10.6190_for | GGGGACAAGTTTGTACAAAAAAGCAGGCTATGCGAAAGACGACTCGTCTACT | Tb927.10.6190_rev | GGGGACCACTTTGTACAAGAAAGCTGGGTCTACACACGTGACTCCAGAACATATTT |
| 4_known | P14 | Tb927.1.3950_for | GGGGACAAGTTTGTACAAAAAAGCAGGCTATGAAAGGTAACCAGGAATTTTTTCAGAAG | Tb927.1.3950_rev | GGGGACCACTTTGTACAAGAAAGCTGGGTTTATGCATACTTGGCCCAAATGCC |
| 4_known | P15 | Tb927.11.14100_for | GGGGACAAGTTTGTACAAAAAAGCAGGCTATGCAAACCTGTCTTTTGAGTAACATTC | Tb927.11.14100_rev | GGGGACCACTTTGTACAAGAAAGCTGGGTTCACTCATTCTGTTCGTCCTGTGG |
| 4_known | P16 | Tb927.11.670_for | GGGGACAAGTTTGTACAAAAAAGCAGGCTATGTCATTTCCGACTTCTCTCCACG | Tb927.11.670_rev | GGGGACCACTTTGTACAAGAAAGCTGGGTCTACTGACCTAACCGGCGACC |
| 4_known | P17 | Tb927.3.4390_for | GGGGACAAGTTTGTACAAAAAAGCAGGCTATGTTTGCTAAAACCCGGCTCA | Tb927.3.4390_rev | GGGGACCACTTTGTACAAGAAAGCTGGGTCTAATATGCTCTGCCTCGGTCAAG |
| 4_known | P18 | Tb927.11.2330_for | GGGGACAAGTTTGTACAAAAAAGCAGGCTATGAATGTCCTTGAAGATCTTGAAGCTC | Tb927.11.2330_rev | GGGGACCACTTTGTACAAGAAAGCTGGGTTTAGTAAAAGGGGAAGTTTCTGTCTACTTC |
| 4_known | P19 | Tb927.8.4090_for | GGGGACAAGTTTGTACAAAAAAGCAGGCTATGCATCGCATCACCGTACG | Tb927.8.4090_rev | GGGGACCACTTTGTACAAGAAAGCTGGGTTTAACCGGTGTCGTTGGTCGA |
| 4_known | P20 | Tb927.6.4370_for | GGGGACAAGTTTGTACAAAAAAGCAGGCTATGGCCTTTGTGTTACCCCAGA | Tb927.6.4370_rev | GGGGACCACTTTGTACAAGAAAGCTGGGTCTAACCGTCTTCCTCACCATCATCA |
| 4_known | P22 | Tb927.10.12810_for | GGGGACAAGTTTGTACAAAAAAGCAGGCTATGGTATCCCACTACCTGAGTGAC | Tb927.10.12810_rev | GGGGACCACTTTGTACAAGAAAGCTGGGTCTACGACCCCTCCATAACAGTTGA |
| 4_known | P23 | Tb927.11.7240_for | GGGGACAAGTTTGTACAAAAAAGCAGGCTATGCAAGCCGATGCAAAGGG | Tb927.11.7240_rev | GGGGACCACTTTGTACAAGAAAGCTGGGTCTACCTCGTCATGTCGGAGACG |
| 4_known | P24 | Tb927.11.16410_for | GGGGACAAGTTTGTACAAAAAAGCAGGCTATGGAATCCGTCTCATGGAAATTGTATG | Tb927.11.16410_rev | GGGGACCACTTTGTACAAGAAAGCTGGGTTCACTTCTTGCGCTTCTTCTTATCC |
| 5_hypothetical | A1 | Tb927.10.4070_for | GGGGACAAGTTTGTACAAAAAAGCAGGCTATGGACGTGAGAGATCAGTTGATGC | Tb927.10.4070_rev | GGGGACCACTTTGTACAAGAAAGCTGGGTTTACCCGCCAGCAACCGA |
| 5_hypothetical | A3 | Tb927.3.5690_for | GGGGACAAGTTTGTACAAAAAAGCAGGCTATGGAGAACAAATATTTAGTGTGGTGCA | Tb927.3.5690_rev | GGGGACCACTTTGTACAAGAAAGCTGGGTTTAAAGCAACAATAACGAAAGTGTTAAAAG |
| 5_hypothetical | A4 | Tb927.8.1190_for | GGGGACAAGTTTGTACAAAAAAGCAGGCTATGTTTCACGGTTTTCCGCAGG | Tb927.8.1190_rev | GGGGACCACTTTGTACAAGAAAGCTGGGTCTACAGGTCACGTAGGAGTGATGA |
| 5_hypothetical | A5 | Tb927.10.13710_for | GGGGACAAGTTTGTACAAAAAAGCAGGCTATGGCGCGTATAGGTGAAGGA | Tb927.10.13710_rev | GGGGACCACTTTGTACAAGAAAGCTGGGTTCACAGATATTCCAACGAGTAACCGA |
| 5_hypothetical | A6 | Tb927.7.4740_for | GGGGACAAGTTTGTACAAAAAAGCAGGCTATGAGTGATGGCGTTCCTCCA | Tb927.7.4740_rev | GGGGACCACTTTGTACAAGAAAGCTGGGTTTATATGGGGGCGAGAAACCGA |
| 5_hypothetical | A7 | Tb927.1.4680_for | GGGGACAAGTTTGTACAAAAAAGCAGGCTATGTCAACAGAGGAACTCGTGC | Tb927.1.4680_rev | GGGGACCACTTTGTACAAGAAAGCTGGGTTTATCTGTGATTATTCGCATTAGCTTTGAT |
| 5_hypothetical | A8 | Tb927.4.1230_for | GGGGACAAGTTTGTACAAAAAAGCAGGCTATGGGAAGAAAGACTGGTGCGC | Tb927.4.1230_rev | GGGGACCACTTTGTACAAGAAAGCTGGGTTCAAGACAATGCAGTTAACAGAAGCA |
| 5_hypothetical | A9 | Tb927.11.3540_for | GGGGACAAGTTTGTACAAAAAAGCAGGCTATGAGCGTGCCTCGTTTGC | Tb927.11.3540_rev | GGGGACCACTTTGTACAAGAAAGCTGGGTTCAGTCGAGGGCAAATGCAAG |

| | | | | | |
|---|---|---|---|---|---|
| 5_hypothetical | A10 | Tb927.11.610_for | GGGGACAAGTTTGTACAAAAAAGCAGGCTATGAGTTTCGACTACGCCGTTC | Tb927.11.610_rev | GGGGACCACTTTGTACAAGAAAGCTGGGTCTACGCCAGTTTGAAGTTGTGTCG |
| 5_hypothetical | A12 | Tb927.11.5760_for | GGGGACAAGTTTGTACAAAAAAGCAGGCTATGTTCCCCCCGGTGTTCA | Tb927.11.5760_rev | GGGGACCACTTTGTACAAGAAAGCTGGGTTTATCGCTTTCGGCGTGGAG |
| 5_hypothetical | A13 | Tb927.10.7000_for | GGGGACAAGTTTGTACAAAAAAGCAGGCTATGAAGACAGTATACATTGTAGCGATGTG | Tb927.10.7000_rev | GGGGACCACTTTGTACAAGAAAGCTGGGTTCATTTCTCAACAACATCTACTGTGAAAAC |
| 5_hypothetical | A14 | Tb927.6.2990_for | GGGGACAAGTTTGTACAAAAAAGCAGGCTATGGCACTCATGTGGCGGT | Tb927.6.2990_rev | GGGGACCACTTTGTACAAGAAAGCTGGGTTCAGCATTGCGTAACGTAACGC |
| 5_hypothetical | A15 | Tb927.3.2120_for | GGGGACAAGTTTGTACAAAAAAGCAGGCTATGAGCGCTATTTATAGTCTTCTATTCGAC | Tb927.3.2120_rev | GGGGACCACTTTGTACAAGAAAGCTGGGTTCATCCAATAAACGTTTTCTCGCCT |
| 5_hypothetical | A16 | Tb927.11.4730_for | GGGGACAAGTTTGTACAAAAAAGCAGGCTATGGCAGGATTGCAGAGTGC | Tb927.11.4730_rev | GGGGACCACTTTGTACAAGAAAGCTGGGTTTACGTCAGCCCCGCTACG |
| 5_hypothetical | A17 | Tb927.10.11430_for | GGGGACAAGTTTGTACAAAAAAGCAGGCTATGTCATTGTTTGATGATGATGATGATGTG | Tb927.10.11430_rev | GGGGACCACTTTGTACAAGAAAGCTGGGTTTACACAAAACGGTCCATAGATCGTC |
| 5_hypothetical | A18 | Tb927.9.6730_for | GGGGACAAGTTTGTACAAAAAAGCAGGCTATGGATGATGAGACATTCGCGTG | Tb927.9.6730_rev | GGGGACCACTTTGTACAAGAAAGCTGGGTTTACGGGCGACCCCACTC |
| 5_hypothetical | A19 | Tb927.4.380_for | GGGGACAAGTTTGTACAAAAAAGCAGGCTATGTCGGAGGAACATGCGAATG | Tb927.4.380_rev | GGGGACCACTTTGTACAAGAAAGCTGGGTTCACTGCTTTACCGTAGCGAAAAG |
| 5_hypothetical | A20 | Tb927.6.1930_for | GGGGACAAGTTTGTACAAAAAAGCAGGCTATGAGCACTCGCAGTGGTGT | Tb927.6.1930_rev | GGGGACCACTTTGTACAAGAAAGCTGGGTTCAGCCACTTTTATCACTCGGC |
| 5_hypothetical | A21 | Tb927.4.2250_for | GGGGACAAGTTTGTACAAAAAAGCAGGCTATGATATTACGGCACTCTCGCCG | Tb927.4.2250_rev | GGGGACCACTTTGTACAAGAAAGCTGGGTTCAAAACCATCGTCCGATGCG |
| 5_hypothetical | A22 | Tb927.10.10820_for | GGGGACAAGTTTGTACAAAAAAGCAGGCTATGCCAAGTTCGGTGGCCC | Tb927.10.10820_rev | GGGGACCACTTTGTACAAGAAAGCTGGGTTCACTGCCCCGATTGAGGG |
| 5_hypothetical | A23 | Tb927.10.9730_for | GGGGACAAGTTTGTACAAAAAAGCAGGCTATGTGGCGTCCAGCTGAAGT | Tb927.10.9730_rev | GGGGACCACTTTGTACAAGAAAGCTGGGTCTACATTCGCACTGAATCGGTTGT |
| 5_hypothetical | A24 | Tb927.11.10570_for | GGGGACAAGTTTGTACAAAAAAGCAGGCTATGCTTCGTCTGTCGTCATGGA | Tb927.11.10570_rev | GGGGACCACTTTGTACAAGAAAGCTGGGTCTAGTTACCCGCTGCTGCCG |
| 5_hypothetical | B1 | Tb927.8.5640_for | GGGGACAAGTTTGTACAAAAAAGCAGGCTATGCGTCGCTTCCCACCC | Tb927.8.5640_rev | GGGGACCACTTTGTACAAGAAAGCTGGGTTTATTCAGGTGATCGTTGGAGGCA |
| 5_hypothetical | B2 | Tb927.8.6080_for | GGGGACAAGTTTGTACAAAAAAGCAGGCTATGAAGAGTGTATTTGTAGCATCTAGTATG | Tb927.8.6080_rev | GGGGACCACTTTGTACAAGAAAGCTGGGTTCATTTGCTGGGAAGTTTTGCTGC |
| 5_hypothetical | B3 | Tb927.5.3060_for | GGGGACAAGTTTGTACAAAAAAGCAGGCTATGGCGTCGGTTGTCAGC | Tb927.5.3060_rev | GGGGACCACTTTGTACAAGAAAGCTGGGTTCACACACTCAACCCCGTGT |
| 5_hypothetical | B4 | Tb927.10.11820_for | GGGGACAAGTTTGTACAAAAAAGCAGGCTATGACGGATCCTGTGCGCC | Tb927.10.11820_rev | GGGGACCACTTTGTACAAGAAAGCTGGGTTCAAATACGGTAAAACCCCGGAATAC |
| 5_hypothetical | B5 | Tb927.3.810_for | GGGGACAAGTTTGTACAAAAAAGCAGGCTATGTTGAATGCTGGTGGAGCC | Tb927.3.810_rev | GGGGACCACTTTGTACAAGAAAGCTGGGTTCAAGAACCGTAGCTGAGAAACAGC |
| 5_hypothetical | B6 | Tb927.11.13260.2_for | GGGGACAAGTTTGTACAAAAAAGCAGGCTATGCACCCTTATTTATTTATTTGTAAAGAA | Tb927.11.13260.2_rev | GGGGACCACTTTGTACAAGAAAGCTGGGTTCACATCTTCAAGAGACGAAACATCTG |
| 5_hypothetical | B7 | Tb927.4.1260_for | GGGGACAAGTTTGTACAAAAAAGCAGGCTATGAAGTTTGTGTGGTTAGACACGAA | Tb927.4.1260_rev | GGGGACCACTTTGTACAAGAAAGCTGGGTTTATATTCCTGTGGGCACACGC |
| 5_hypothetical | B8 | Tb927.5.3080_for | GGGGACAAGTTTGTACAAAAAAGCAGGCTATGTGGGGACCAAACCAGGAG | Tb927.5.3080_rev | GGGGACCACTTTGTACAAGAAAGCTGGGTTCACTGCACTTCGCAATCCTC |
| 5_hypothetical | B9 | Tb927.9.2130_for | GGGGACAAGTTTGTACAAAAAAGCAGGCTATGATGGGGGAAAAACAAGCACA | Tb927.9.2130_rev | GGGGACCACTTTGTACAAGAAAGCTGGGTTTACTTCTTTTCATCTCCCCCACTTAGT |
| 5_hypothetical | B10 | Tb927.7.850_for | GGGGACAAGTTTGTACAAAAAAGCAGGCTATGATGACATTGCGTTCCTCAACG | Tb927.7.850_rev | GGGGACCACTTTGTACAAGAAAGCTGGGTTCATAAACCCATACTCGCCAAGTTTG |
| 5_hypothetical | B11 | Tb927.11.13420_for | GGGGACAAGTTTGTACAAAAAAGCAGGCTATGGGCGGAGGCTGTTGC | Tb927.11.13420_rev | GGGGACCACTTTGTACAAGAAAGCTGGGTTTATTCTGACCGCTGTTGTCTCTGT |
| 5_hypothetical | B12 | Tb927.3.750_for | GGGGACAAGTTTGTACAAAAAAGCAGGCTATGCGGCTGCGGGGTCAA | Tb927.3.750_rev | GGGGACCACTTTGTACAAGAAAGCTGGGTTCATACGCCACGCTGCTTG |
| 5_hypothetical | B13 | Tb927.7.640_for | GGGGACAAGTTTGTACAAAAAAGCAGGCTATGGGTTGGTGGAGGAGAGGC | Tb927.7.640_rev | GGGGACCACTTTGTACAAGAAAGCTGGGTCTAAAGTAGAAATAGGAGCACAGCCAA |
| 5_hypothetical | B14 | Tb927.11.11050_for | GGGGACAAGTTTGTACAAAAAAGCAGGCTATGACGATTGTTGCGGACTGTG | Tb927.11.11050_rev | GGGGACCACTTTGTACAAGAAAGCTGGGTTCATGGCGGCGCTTTGGA |
| 5_hypothetical | B15 | Tb927.10.510_for | GGGGACAAGTTTGTACAAAAAAGCAGGCTATGCCTCTCATGTCTGCAGATGG | Tb927.10.510_rev | GGGGACCACTTTGTACAAGAAAGCTGGGTTCACTCTTCATCTTCTAAAAACTCTTCTTC |
| 5_hypothetical | B16 | Tb927.8.4570_for | GGGGACAAGTTTGTACAAAAAAGCAGGCTATGGGGCAAAAATCTTCAACGGC | Tb927.8.4570_rev | GGGGACCACTTTGTACAAGAAAGCTGGGTCTAAGCGGTCATTAGGCGTTCAA |
| 5_hypothetical | B17 | Tb927.10.11230_for | GGGGACAAGTTTGTACAAAAAAGCAGGCTATGGCAGGGACTGGTGCC | Tb927.10.11230_rev | GGGGACCACTTTGTACAAGAAAGCTGGGTTCACAACTTCTTTACTTCACCCACAG |
| 5_hypothetical | B18 | Tb927.2.3400_for | GGGGACAAGTTTGTACAAAAAAGCAGGCTATGAGTGCAACACCACAATTAGGAG | Tb927.2.3400_rev | GGGGACCACTTTGTACAAGAAAGCTGGGTTTACTCACCCTTCTTGCGCTCG |
| 5_hypothetical | B19 | Tb927.7.4840_for | GGGGACAAGTTTGTACAAAAAAGCAGGCTATGCCTCCGCGTCGCAAG | Tb927.7.4840_rev | GGGGACCACTTTGTACAAGAAAGCTGGGTTCACTGTGGTTGCACAAACGG |
| 5_hypothetical | B20 | Tb927.3.2460_for | GGGGACAAGTTTGTACAAAAAAGCAGGCTATGAAAGAAGTTAAAGGGGAAAATGAGTTT | Tb927.3.2460_rev | GGGGACCACTTTGTACAAGAAAGCTGGGTTCAACAATGCTTTTTGACACGAAGAG |
| 5_hypothetical | B21 | Tb927.3.2920_for | GGGGACAAGTTTGTACAAAAAAGCAGGCTATGAATCAAAAGCGTCGTTGCAC | Tb927.3.2920_rev | GGGGACCACTTTGTACAAGAAAGCTGGGTTTAGTTAACAACCCTAATAATGTCATTGGG |
| 5_hypothetical | B22 | Tb927.9.9710_for | GGGGACAAGTTTGTACAAAAAAGCAGGCTATGTTTTATCTGATGGTGGTTGCACT | Tb927.9.9710_rev | GGGGACCACTTTGTACAAGAAAGCTGGGTTCAGAAACCCTTGTACAACACTGATAC |
| 5_hypothetical | B23 | Tb927.11.12520_for | GGGGACAAGTTTGTACAAAAAAGCAGGCTATGACTGCTGGTGGAGGCA | Tb927.11.12520_rev | GGGGACCACTTTGTACAAGAAAGCTGGGTTCACTTGAACATGTGCTGATGCT |
| 5_hypothetical | B24 | Tb927.4.1180_for | GGGGACAAGTTTGTACAAAAAAGCAGGCTATGGCAAAGAAGGAAGTTGTCTGG | Tb927.4.1180_rev | GGGGACCACTTTGTACAAGAAAGCTGGGTCTACACTTCGTCCTCAACGACAC |
| 5_hypothetical | C1 | Tb927.5.960_for | GGGGACAAGTTTGTACAAAAAAGCAGGCTATGCGACCTCTGGACGTGT | Tb927.5.960_rev | GGGGACCACTTTGTACAAGAAAGCTGGGTTCAAACGCTTGATGCCATCGA |
| 5_hypothetical | C2 | Tb927.7.1850_for | GGGGACAAGTTTGTACAAAAAAGCAGGCTATGGGTTGCCAACGTATGGGG | Tb927.7.1850_rev | GGGGACCACTTTGTACAAGAAAGCTGGGTTTAGCGAAAATCTGGAACCCGC |
| 5_hypothetical | C4 | Tb927.9.10870_for | GGGGACAAGTTTGTACAAAAAAGCAGGCTATGAGTATCTTGGTGAACCGTCGT | Tb927.9.10870_rev | GGGGACCACTTTGTACAAGAAAGCTGGGTTCAACGACGCCTTGAAGAAGAC |
| 5_hypothetical | C5 | Tb927.11.15630_for | GGGGACAAGTTTGTACAAAAAAGCAGGCTATGCACTCGAGAAGGAAAGGGA | Tb927.11.15630_rev | GGGGACCACTTTGTACAAGAAAGCTGGGTTTATGAACTCGTGCTATCATCGTCATTT |
| 5_hypothetical | C6 | Tb927.3.3250_for | GGGGACAAGTTTGTACAAAAAAGCAGGCTATGCCGCACAGGAAACTACTG | Tb927.3.3250_rev | GGGGACCACTTTGTACAAGAAAGCTGGGTTCACCTCCCAAGCGGGCT |
| 5_hypothetical | C7 | Tb927.5.1350_for | GGGGACAAGTTTGTACAAAAAAGCAGGCTATGCTGTGGGCATTCAGTAGAGG | Tb927.5.1350_rev | GGGGACCACTTTGTACAAGAAAGCTGGGTTCATACCGCAATCTGTGAAGAGTTG |
| 5_hypothetical | C8 | Tb927.10.8690_for | GGGGACAAGTTTGTACAAAAAAGCAGGCTATGCCGCCACAGAGGCAA | Tb927.10.8690_rev | GGGGACCACTTTGTACAAGAAAGCTGGGTTTATTGTCCCTCTACCTCCACATTTTGC |
| 5_hypothetical | C9 | Tb927.9.13350_for | GGGGACAAGTTTGTACAAAAAAGCAGGCTATGGCAGCGAAGCGTAGC | Tb927.9.13350_rev | GGGGACCACTTTGTACAAGAAAGCTGGGTCTAGAACTTTTTAAGTGAACGCCGTTG |
| 5_hypothetical | C10 | Tb927.10.14500_for | GGGGACAAGTTTGTACAAAAAAGCAGGCTATGTCTGTCAAGTTGCCACACG | Tb927.10.14500_rev | GGGGACCACTTTGTACAAGAAAGCTGGGTCTAATTAGTTGTGGTTGGTTGATTCTTCAG |
| 5_hypothetical | C11 | Tb927.10.14310_for | GGGGACAAGTTTGTACAAAAAAGCAGGCTATGGTCATTTTCGATGATGTGGAAAAC | Tb927.10.14310_rev | GGGGACCACTTTGTACAAGAAAGCTGGGTTCAGTTTTTTGCATGGTTGTCGGG |
| 5_hypothetical | C12 | Tb927.7.6820_for | GGGGACAAGTTTGTACAAAAAAGCAGGCTATGGGCGGTGATGGCAACG | Tb927.7.6820_rev | GGGGACCACTTTGTACAAGAAAGCTGGGTTCAAGAGTGTATATCCACAGCTCCC |
| 5_hypothetical | C13 | Tb927.6.3070_for | GGGGACAAGTTTGTACAAAAAAGCAGGCTATGGACTCCATGGAAGAGTTTTATAACAAC | Tb927.6.3070_rev | GGGGACCACTTTGTACAAGAAAGCTGGGTTTAACGCTCAAGTGCTGACAAGG |
| 5_hypothetical | C14 | Tb927.10.11500_for | GGGGACAAGTTTGTACAAAAAAGCAGGCTATGTGGCGCCGGGGGATT | Tb927.10.11500_rev | GGGGACCACTTTGTACAAGAAAGCTGGGTTTATGTAAGTTTACTCAGCGCCTCAC |
| 5_hypothetical | C15 | Tb927.10.8650_for | GGGGACAAGTTTGTACAAAAAAGCAGGCTATGTCGGACCGCCTTAGAGG | Tb927.10.8650_rev | GGGGACCACTTTGTACAAGAAAGCTGGGTTCACTTAGACTGCTCCTGTGCG |
| 5_hypothetical | C16 | Tb927.9.1360_for | GGGGACAAGTTTGTACAAAAAAGCAGGCTATGAACTCAGCCGATTCTTCAGGT | Tb927.9.1360_rev | GGGGACCACTTTGTACAAGAAAGCTGGGTTTACGGTGTGGGAACCGAGT |
| 5_hypothetical | C17 | Tb927.8.7370_for | GGGGACAAGTTTGTACAAAAAAGCAGGCTATGGGTTCCGGGGTAGGAAGA | Tb927.8.7370_rev | GGGGACCACTTTGTACAAGAAAGCTGGGTTTACGGGTAGTAACCCACAGAGG |
| 5_hypothetical | C18 | Tb927.9.13290_for | GGGGACAAGTTTGTACAAAAAAGCAGGCTATGCCCGGGATTGATTACTCAAAAT | Tb927.9.13290_rev | GGGGACCACTTTGTACAAGAAAGCTGGGTTCACGTACTTCCCCCACGC |
| 5_hypothetical | C19 | Tb927.3.1680_for | GGGGACAAGTTTGTACAAAAAAGCAGGCTATGGAAGGTCTTGTGGTTCCCC | Tb927.3.1680_rev | GGGGACCACTTTGTACAAGAAAGCTGGGTTTAACGGGTATTATTTCCTCTGTTGGG |
| 5_hypothetical | C20 | Tb927.10.2200_for | GGGGACAAGTTTGTACAAAAAAGCAGGCTATGAGCCGTGTGCCGCAT | Tb927.10.2200_rev | GGGGACCACTTTGTACAAGAAAGCTGGGTTCATCCTCGCATAACGCTGTG |
| 5_hypothetical | C21 | Tb927.10.2810_for | GGGGACAAGTTTGTACAAAAAAGCAGGCTATGTGTAACGACGATGCTCGG | Tb927.10.2810_rev | GGGGACCACTTTGTACAAGAAAGCTGGGTCTACATGGCATCGGGGCATTC |
| 5_hypothetical | C22 | Tb927.11.8110_for | GGGGACAAGTTTGTACAAAAAAGCAGGCTATGGAGCCAAGGCGTTGGC | Tb927.11.8110_rev | GGGGACCACTTTGTACAAGAAAGCTGGGTTCAAACCAACATGAGCGACTTGT |
| 5_hypothetical | C23 | Tb927.9.6720_for | GGGGACAAGTTTGTACAAAAAAGCAGGCTATGCTGGCTGATGGATTTCGC | Tb927.9.6720_rev | GGGGACCACTTTGTACAAGAAAGCTGGGTTTACCTTTTGCATTCGACTCGTAAAATG |
| 5_hypothetical | C24 | Tb927.10.11000_for | GGGGACAAGTTTGTACAAAAAAGCAGGCTATGAACCCCGAAGCGCCG | Tb927.10.11000_rev | GGGGACCACTTTGTACAAGAAAGCTGGGTTCACCTCCTACGTGGCGG |
| 5_hypothetical | D1 | Tb927.8.3820_for | GGGGACAAGTTTGTACAAAAAAGCAGGCTATGAATCGATCTTCTGATGCTGAGTG | Tb927.8.3820_rev | GGGGACCACTTTGTACAAGAAAGCTGGGTTTACTTTCCCCTGTTCCTTTGATAATTAGA |
| 5_hypothetical | D2 | Tb927.9.13970_for | GGGGACAAGTTTGTACAAAAAAGCAGGCTATGCACTGCCGCCTGCTC | Tb927.9.13970_rev | GGGGACCACTTTGTACAAGAAAGCTGGGTTCATTGTCTAGGAATAAGGACATTACCAAA |
| 5_hypothetical | D3 | Tb927.7.5510_for | GGGGACAAGTTTGTACAAAAAAGCAGGCTATGATTTTGAGGGGAAGGGTTGGA | Tb927.7.5510_rev | GGGGACCACTTTGTACAAGAAAGCTGGGTCTCACTTTTCCCTGTGAAGAAAAGGAAT |
| 5_hypothetical | D4 | Tb927.10.6480_for | GGGGACAAGTTTGTACAAAAAAGCAGGCTATGGTGGATAAACTAACCCCACCA | Tb927.10.6480_rev | GGGGACCACTTTGTACAAGAAAGCTGGGTTCAAGTGAGCCGTTGCCG |
| 5_hypothetical | D5 | Tb927.8.6480_for | GGGGACAAGTTTGTACAAAAAAGCAGGCTATGGGGTTCAAGTGTCTCCTCTC | Tb927.8.6480_rev | GGGGACCACTTTGTACAAGAAAGCTGGGTTCATGATTTGGTGAGGACTTTGCG |
| 5_hypothetical | D6 | Tb927.11.13680_for | GGGGACAAGTTTGTACAAAAAAGCAGGCTATGAGTAGGAAAAATGCGCCTCC | Tb927.11.13680_rev | GGGGACCACTTTGTACAAGAAAGCTGGGTTCAACCCTTTACCCGCTTCTTCT |
| 5_hypothetical | D7 | Tb927.11.10350_for | GGGGACAAGTTTGTACAAAAAAGCAGGCTATGGACGTGGGGTATTACGTTCC | Tb927.11.10350_rev | GGGGACCACTTTGTACAAGAAAGCTGGGTCTATAGAGAAGCGGAGGCTGAGAC |
| 5_hypothetical | D8 | Tb927.9.6960_for | GGGGACAAGTTTGTACAAAAAAGCAGGCTATGAACTCCCCTCACGACTCG | Tb927.9.6960_rev | GGGGACCACTTTGTACAAGAAAGCTGGGTCTATCTCGTGTGACGCGGGT |
| 5_hypothetical | D9 | Tb927.10.1800_for | GGGGACAAGTTTGTACAAAAAAGCAGGCTATGTCCTTCTCCAGTGACGAGAA | Tb927.10.1800_rev | GGGGACCACTTTGTACAAGAAAGCTGGGTTTAAGATAAACAAAAGCTGCCATTTATGAA |
| 5_hypothetical | D10 | Tb927.11.9800_for | GGGGACAAGTTTGTACAAAAAAGCAGGCTATGTTAAACATATTGTCGTACTCAAAGTTA | Tb927.11.9800_rev | GGGGACCACTTTGTACAAGAAAGCTGGGTTCAAGAACGGTCATCACCATCGT |
| 5_hypothetical | D11 | Tb927.4.2770_for | GGGGACAAGTTTGTACAAAAAAGCAGGCTATGGCGGTGACACCAATAGAAAAA | Tb927.4.2770_rev | GGGGACCACTTTGTACAAGAAAGCTGGGTTCAGGAGGCAGTGGCTTCC |
| 5_hypothetical | D13 | Tb927.10.9980_for | GGGGACAAGTTTGTACAAAAAAGCAGGCTATGAGCTCAAGGAACTGCAGC | Tb927.10.9980_rev | GGGGACCACTTTGTACAAGAAAGCTGGGTTCACCACATGGAAACATGGTAGGA |
| 5_hypothetical | D14 | Tb927.8.5480_for | GGGGACAAGTTTGTACAAAAAAGCAGGCTATGCGCAACGGTGTGTCG | Tb927.8.5480_rev | GGGGACCACTTTGTACAAGAAAGCTGGGTTTACAAAACATGTACATGAGGCCCAA |
| 5_hypothetical | D15 | Tb927.11.11710_for | GGGGACAAGTTTGTACAAAAAAGCAGGCTATGACAGTCGCAAAGAAGTGTATGG | Tb927.11.11710_rev | GGGGACCACTTTGTACAAGAAAGCTGGGTCTACTCCTGGAAAGTTGCCAGC |
| 5_hypothetical | D16 | Tb927.7.2160_for | GGGGACAAGTTTGTACAAAAAAGCAGGCTATGCAAGCATCTGAATCAGGAGATC | Tb927.7.2160_rev | GGGGACCACTTTGTACAAGAAAGCTGGGTTCACGCTGCAACCTTCTGC |
| 5_hypothetical | D17 | Tb927.10.4820_for | GGGGACAAGTTTGTACAAAAAAGCAGGCTATGGTGACGGAGGCCCTG | Tb927.10.4820_rev | GGGGACCACTTTGTACAAGAAAGCTGGGTTTACCGCTTGAGGAATTGCTGAAT |
| 5_hypothetical | D18 | Tb927.10.9640_for | GGGGACAAGTTTGTACAAAAAAGCAGGCTATGAGTGGAAACTCACAAACTGCTAG | Tb927.10.9640_rev | GGGGACCACTTTGTACAAGAAAGCTGGGTTTACTTCGGCTTTGTCCCAATTTTTG |
| 5_hypothetical | D19 | Tb927.11.14400_for | GGGGACAAGTTTGTACAAAAAAGCAGGCTATGAGTTACGAAAATTTCGCTGCAC | Tb927.11.14400_rev | GGGGACCACTTTGTACAAGAAAGCTGGGTTTAGACGAGCACAGAACCAACAATT |
| 5_hypothetical | D20 | Tb927.10.14220_for | GGGGACAAGTTTGTACAAAAAAGCAGGCTATGACGGAAGCAGTAGCGGC | Tb927.10.14220_rev | GGGGACCACTTTGTACAAGAAAGCTGGGTTTAGTCGTCAAGTTGCGTATGCTC |
| 5_hypothetical | D21 | Tb927.5.2390_for | GGGGACAAGTTTGTACAAAAAAGCAGGCTATGCGATCACCGAACGGATCT | Tb927.5.2390_rev | GGGGACCACTTTGTACAAGAAAGCTGGGTTCATAACATGAACTTGCCAAATTCACG |
| 5_hypothetical | D22 | Tb927.9.2780_for | GGGGACAAGTTTGTACAAAAAAGCAGGCTATGGATAACGAAGGCTGCAGC | Tb927.9.2780_rev | GGGGACCACTTTGTACAAGAAAGCTGGGTCTAGCAGGGTGCCAATATACCGA |
| 5_hypothetical | D23 | Tb927.10.1230_for | GGGGACAAGTTTGTACAAAAAAGCAGGCTATGGCGGAAGAAAGGTGCCT | Tb927.10.1230_rev | GGGGACCACTTTGTACAAGAAAGCTGGGTTTACACAGAAAGAGTGGTTGCCC |
| 5_hypothetical | D24 | Tb927.10.13840_for | GGGGACAAGTTTGTACAAAAAAGCAGGCTATGAGTCTTTTGTCTAATGCGGGTAAC | Tb927.10.13840_rev | GGGGACCACTTTGTACAAGAAAGCTGGGTTCAAACACGTGCTTTGCCACA |
| 5_hypothetical | E1 | Tb927.11.15410_for | GGGGACAAGTTTGTACAAAAAAGCAGGCTATGACAAGTATTACAGCGTATATGCGT | Tb927.11.15410_rev | GGGGACCACTTTGTACAAGAAAGCTGGGTTTATGCACCACTCCGTCCGA |
| 5_hypothetical | E2 | Tb927.7.310_for | GGGGACAAGTTTGTACAAAAAAGCAGGCTATGCGGTGGCCATCTGCC | Tb927.7.310_rev | GGGGACCACTTTGTACAAGAAAGCTGGGTTTATTTTTTTGTCACATGCGTCCGC |
| 5_hypothetical | E3 | Tb927.10.14700_for | GGGGACAAGTTTGTACAAAAAAGCAGGCTATGAACGCGAGAGGTAAAGGACG | Tb927.10.14700_rev | GGGGACCACTTTGTACAAGAAAGCTGGGTTTACCATACACCCTCACCGTTGC |
| 5_hypothetical | E4 | Tb927.11.13210_for | GGGGACAAGTTTGTACAAAAAAGCAGGCTATGCCCCTGGGAAGTACTAATCAG | Tb927.11.13210_rev | GGGGACCACTTTGTACAAGAAAGCTGGGTTCAACAACTGAGCCGACTGTTG |
| 5_hypothetical | E5 | Tb927.9.12470_for | GGGGACAAGTTTGTACAAAAAAGCAGGCTATGTTAAATATTTGCCAAAACTTGTCTTCC | Tb927.9.12470_rev | GGGGACCACTTTGTACAAGAAAGCTGGGTTCATGAAAGAAGAAAAATAATTTTTTCCAA |
| 5_hypothetical | E6 | Tb927.11.9490_for | GGGGACAAGTTTGTACAAAAAAGCAGGCTATGAGCGATGATGAATGCCCG | Tb927.11.9490_rev | GGGGACCACTTTGTACAAGAAAGCTGGGTCTACGCACAACCCCGGAGG |
| 5_hypothetical | E7 | Tb927.10.13170_for | GGGGACAAGTTTGTACAAAAAAGCAGGCTATGCCACGCCACCTTTCG | Tb927.10.13170_rev | GGGGACCACTTTGTACAAGAAAGCTGGGTTCAGCCGTGGGTGTGGTG |
| 5_hypothetical | E8 | Tb927.9.12750_for | GGGGACAAGTTTGTACAAAAAAGCAGGCTATGCTTCGCCGCACGCTG | Tb927.9.12750_rev | GGGGACCACTTTGTACAAGAAAGCTGGGTTTATGAGGTAATTTTCTTTGTTTCTTCCAC |
| 5_hypothetical | E9 | Tb927.9.10470_for | GGGGACAAGTTTGTACAAAAAAGCAGGCTATGTTATGTCGTACGTTTGTCAGACG | Tb927.9.10470_rev | GGGGACCACTTTGTACAAGAAAGCTGGGTTCACTTGTTCAGTGCACGTCC |
| 5_hypothetical | E10 | Tb927.9.9180_for | GGGGACAAGTTTGTACAAAAAAGCAGGCTATGGACAATAGGCCGGCGC | Tb927.9.9180_rev | GGGGACCACTTTGTACAAGAAAGCTGGGTTCAAGCCGTTGGAGGTTGATCT |
| 5_hypothetical | E11 | Tb927.4.3150_for | GGGGACAAGTTTGTACAAAAAAGCAGGCTATGTGGAACGCATTTAGCGGG | Tb927.4.3150_rev | GGGGACCACTTTGTACAAGAAAGCTGGGTCTACTTCGCATGTGGCATTAGAGG |
| 5_hypothetical | E12 | Tb927.9.10880_for | GGGGACAAGTTTGTACAAAAAAGCAGGCTATGGATGTCAAGACCCTCTTAGCT | Tb927.9.10880_rev | GGGGACCACTTTGTACAAGAAAGCTGGGTTTAAGCGATGTCATCATCACCCTCA |
| 5_hypothetical | E13 | Tb927.10.9370_for | GGGGACAAGTTTGTACAAAAAAGCAGGCTATGTCGCGCTTTCCTCACTC | Tb927.10.9370_rev | GGGGACCACTTTGTACAAGAAAGCTGGGTCTAAATTGGAATACAGGGAATGGGATTAAC |
| 5_hypothetical | E14 | Tb927.5.820_for | GGGGACAAGTTTGTACAAAAAAGCAGGCTATGCCGAGGAATTCTCAAACATTTCG | Tb927.5.820_rev | GGGGACCACTTTGTACAAGAAAGCTGGGTTCAGCCGACCGCCGGTGC |
| 5_hypothetical | E15 | Tb927.6.1840_for | GGGGACAAGTTTGTACAAAAAAGCAGGCTATGAGCCGTTTCAAACGCTGC | Tb927.6.1840_rev | GGGGACCACTTTGTACAAGAAAGCTGGGTTCAGCAGGATGCCAGCAGC |
| 5_hypothetical | E16 | Tb927.5.2480_for | GGGGACAAGTTTGTACAAAAAAGCAGGCTATGGCACAAACCTCTTTCACACC | Tb927.5.2480_rev | GGGGACCACTTTGTACAAGAAAGCTGGGTTCAAAGCACAAATACCATAATACCGTAAAC |
| 5_hypothetical | E17 | Tb927.9.11360_for | GGGGACAAGTTTGTACAAAAAAGCAGGCTATGGCGTCATTGGAAGATAAGGTTG | Tb927.9.11360_rev | GGGGACCACTTTGTACAAGAAAGCTGGGTTCACGATAACGACAACCTCACCC |
| 5_hypothetical | E18 | Tb927.7.770_for | GGGGACAAGTTTGTACAAAAAAGCAGGCTATGCTAATGCCTCAGTTTGAGTTTTTG | Tb927.7.770_rev | GGGGACCACTTTGTACAAGAAAGCTGGGTCTATAATCTCGGTTTCTTCTTTGGATGCA |
| 5_hypothetical | E19 | Tb927.8.5830_for | GGGGACAAGTTTGTACAAAAAAGCAGGCTATGGAGTTCGTTGCAGAGGCA | Tb927.8.5830_rev | GGGGACCACTTTGTACAAGAAAGCTGGGTTCACTCCGCTTGACCCCC |
| 5_hypothetical | E20 | Tb927.10.10430_for | GGGGACAAGTTTGTACAAAAAAGCAGGCTATGCTCGACATTACAGCCATCCC | Tb927.10.10430_rev | GGGGACCACTTTGTACAAGAAAGCTGGGTTTAACCGCTCTTCCCCAGGG |
| 5_hypothetical | E21 | Tb927.6.3610_for | GGGGACAAGTTTGTACAAAAAAGCAGGCTATGACGCTGCTTCACATGTTGG | Tb927.6.3610_rev | GGGGACCACTTTGTACAAGAAAGCTGGGTTCATTCGAAGTGGTCGTGCTCA |
| 5_hypothetical | E22 | Tb927.3.3440_for | GGGGACAAGTTTGTACAAAAAAGCAGGCTATGAACAATTCACTTAAGAAGGGGTCC | Tb927.3.3440_rev | GGGGACCACTTTGTACAAGAAAGCTGGGTCTATTCCACTTTGCATCTTTTTGAAACCT |
| 5_hypothetical | E23 | Tb927.6.1350_for | GGGGACAAGTTTGTACAAAAAAGCAGGCTATGTGGCGATCATTTATGCAAGTGG | Tb927.6.1350_rev | GGGGACCACTTTGTACAAGAAAGCTGGGTTCAGATAAAACAGAGGGATACTGATAGTAA |
| 5_hypothetical | E24 | Tb927.10.820_for | GGGGACAAGTTTGTACAAAAAAGCAGGCTATGGAGTCCCCTGTGGAAATGT | Tb927.10.820_rev | GGGGACCACTTTGTACAAGAAAGCTGGGTTTAATCTTCGTCTTCCACAATGCTATCTT |
| 5_hypothetical | F1 | Tb927.8.6610_for | GGGGACAAGTTTGTACAAAAAAGCAGGCTATGGATTCACTGCAGTCGGCG | Tb927.8.6610_rev | GGGGACCACTTTGTACAAGAAAGCTGGGTCTACACCCCGTCGACCGC |
| 5_hypothetical | F2 | Tb927.9.8990_for | GGGGACAAGTTTGTACAAAAAAGCAGGCTATGGCACAAGGTGGTGTGGT | Tb927.9.8990_rev | GGGGACCACTTTGTACAAGAAAGCTGGGTTTACATTTCCTGCCGCATTTTTTTTGT |
| 5_hypothetical | F3 | Tb927.10.1200_for | GGGGACAAGTTTGTACAAAAAAGCAGGCTATGCGAGAGGTATTGCTTGGTG | Tb927.10.1200_rev | GGGGACCACTTTGTACAAGAAAGCTGGGTCTACGTCCTTCCTGTCATAAAGGTG |
| 5_hypothetical | F4 | Tb927.8.4600_for | GGGGACAAGTTTGTACAAAAAAGCAGGCTATGTGTACAGGAGGTTCAACAAAGTT | Tb927.8.4600_rev | GGGGACCACTTTGTACAAGAAAGCTGGGTCTAATCGTTGCCTTTGGAGAGGC |
| 5_hypothetical | F5 | Tb927.11.11760_for | GGGGACAAGTTTGTACAAAAAAGCAGGCTATGGATTTTGGCACGGATGTAGC | Tb927.11.11760_rev | GGGGACCACTTTGTACAAGAAAGCTGGGTTCAGATGTCCATCACAACTCCATCA |
| 5_hypothetical | F6 | Tb927.7.2810_for | GGGGACAAGTTTGTACAAAAAAGCAGGCTATGCGCGACTCGGGTGTG | Tb927.7.2810_rev | GGGGACCACTTTGTACAAGAAAGCTGGGTTTAATGCTTGGTGGTGGTGTCG |
| 5_hypothetical | F7 | Tb927.10.5500_for | GGGGACAAGTTTGTACAAAAAAGCAGGCTATGCGGTCGGTAGGTGGTG | Tb927.10.5500_rev | GGGGACCACTTTGTACAAGAAAGCTGGGTCTAAACTGCATTCATTTCTGCTTCCG |
| 5_hypothetical | F8 | Tb927.9.15650_for | GGGGACAAGTTTGTACAAAAAAGCAGGCTATGCACTCATGTTGGATTGGTTCG | Tb927.9.15650_rev | GGGGACCACTTTGTACAAGAAAGCTGGGTTTATATGGTAAACGCAATTACGAAAAAAGC |
| 5_hypothetical | F9 | Tb927.10.5580_for | GGGGACAAGTTTGTACAAAAAAGCAGGCTATGGTCCTTTATGTAATGGGCCCC | Tb927.10.5580_rev | GGGGACCACTTTGTACAAGAAAGCTGGGTCTAGAGTATTTCAAGAAAATGTCTCTCCTT |
| 5_hypothetical | F10 | Tb927.9.7520_for | GGGGACAAGTTTGTACAAAAAAGCAGGCTATGAGACAAGGACGCGTGC | Tb927.9.7520_rev | GGGGACCACTTTGTACAAGAAAGCTGGGTTTACAACGACTCCCGAAGCTCT |
| 5_hypothetical | F11 | Tb927.3.4440_for | GGGGACAAGTTTGTACAAAAAAGCAGGCTATGATAGAGAAATGGAAGGATAACCCACT | Tb927.3.4440_rev | GGGGACCACTTTGTACAAGAAAGCTGGGTTTACCGTTTGGCCCTTCCTCT |
| 5_hypothetical | F12 | Tb927.11.4680_for | GGGGACAAGTTTGTACAAAAAAGCAGGCTATGAGGCCAGATTACATGGGTTTCA | Tb927.11.4680_rev | GGGGACCACTTTGTACAAGAAAGCTGGGTTTACGACTTCTCCCCTGAGGC |
| 5_hypothetical | F13 | Tb927.9.8700_for | GGGGACAAGTTTGTACAAAAAAGCAGGCTATGGAGGGGTCATTTAAGGACTTCG | Tb927.9.8700_rev | GGGGACCACTTTGTACAAGAAAGCTGGGTTCAATCATCTTTATCAATAGAAAGGGCGG |
| 5_hypothetical | F14 | Tb927.6.1390_for | GGGGACAAGTTTGTACAAAAAAGCAGGCTATGTGGAGAGTAATTATGAAAGTGGTCATG | Tb927.6.1390_rev | GGGGACCACTTTGTACAAGAAAGCTGGGTTCAGATAAAACAGAGGGATACTGACAGT |
| 5_hypothetical | F15 | Tb927.6.1370_for | GGGGACAAGTTTGTACAAAAAAGCAGGCTATGTGGAGAGTAATTATGAAAGTGGTCATG | Tb927.6.1370_rev | GGGGACCACTTTGTACAAGAAAGCTGGGTTCAGATAAAACAGAGGGATACTGATAGTAA |
| 5_hypothetical | F16 | Tb927.4.2860_for | GGGGACAAGTTTGTACAAAAAAGCAGGCTATGCGGCGTGGCGTCGGT | Tb927.4.2860_rev | GGGGACCACTTTGTACAAGAAAGCTGGGTTCAGTTACCAGAGGCGACACC |
| 5_hypothetical | F17 | Tb927.3.5060_for | GGGGACAAGTTTGTACAAAAAAGCAGGCTATGAGCGACTTCCCCAAACTGA | Tb927.3.5060_rev | GGGGACCACTTTGTACAAGAAAGCTGGGTCTATTTGGAGGCTTTAGGGCGTC |
| 5_hypothetical | F18 | Tb927.11.15300_for | GGGGACAAGTTTGTACAAAAAAGCAGGCTATGTGGACCGTTGTGCAGC | Tb927.11.15300_rev | GGGGACCACTTTGTACAAGAAAGCTGGGTCTACACCATATCAAGTCCCACACATG |
| 5_hypothetical | F19 | Tb927.4.1960_for | GGGGACAAGTTTGTACAAAAAAGCAGGCTATGGATGGCTTCATTAGCCTTGAAA | Tb927.4.1960_rev | GGGGACCACTTTGTACAAGAAAGCTGGGTTTAAGCCTCCTCCCCTGAGGC |
| 5_hypothetical | F20 | Tb927.10.12690_for | GGGGACAAGTTTGTACAAAAAAGCAGGCTATGTTGCGGTGGCGCATT | Tb927.10.12690_rev | GGGGACCACTTTGTACAAGAAAGCTGGGTTTATTTTTTTTTGCTTGGTGGTGGTTCA |
| 5_hypothetical | F21 | Tb927.5.3000_for | GGGGACAAGTTTGTACAAAAAAGCAGGCTATGTTGCGGTGCAGCTGG | Tb927.5.3000_rev | GGGGACCACTTTGTACAAGAAAGCTGGGTCTATGTGGACACACCCCCCG |
| 5_hypothetical | F22 | Tb927.7.3390_for | GGGGACAAGTTTGTACAAAAAAGCAGGCTATGGCATCGTTAGGCGAATTCTG | Tb927.7.3390_rev | GGGGACCACTTTGTACAAGAAAGCTGGGTTCACGAAGGCACGTCCGT |

| | | | | | |
|---|---|---|---|---|---|
| 5_hypothetical | F23 | Tb927.11.9340_for | GGGGACAAGTTTGTACAAAAAAGCAGGCTATGGAGTGGGTTCCGTGCA | Tb927.11.9340_rev | GGGGACCACTTTGTACAAGAAAGCTGGGTTTACCGTTGCAAAGTAACACCCG |
| 5_hypothetical | F24 | Tb927.10.6600_for | GGGGACAAGTTTGTACAAAAAAGCAGGCTATGTTACTTAGTGATGAGCACCTGGC | Tb927.10.6600_rev | GGGGACCACTTTGTACAAGAAAGCTGGGTCTAGGCATCCATTAAACCCACATGT |
| 5_hypothetical | G1 | Tb927.8.5310_for | GGGGACAAGTTTGTACAAAAAAGCAGGCTATGCGATTTGTTTCTCGACGACT | Tb927.8.5310_rev | GGGGACCACTTTGTACAAGAAAGCTGGGTTTAGTTGTGCGGCAGATGCAC |
| 5_hypothetical | G2 | Tb927.11.11010_for | GGGGACAAGTTTGTACAAAAAAGCAGGCTATGCCTTTACCGCCTGTTGAG | Tb927.11.11010_rev | GGGGACCACTTTGTACAAGAAAGCTGGGTTCACACACCATCTAAGTGGCTCA |
| 5_hypothetical | G3 | Tb927.3.2200_for | GGGGACAAGTTTGTACAAAAAAGCAGGCTATGGCCAATGCAGCGTTAGAG | Tb927.3.2200_rev | GGGGACCACTTTGTACAAGAAAGCTGGGTTCATGAAGTGCGAGAGCGC |
| 5_hypothetical | G4 | Tb927.5.1670_for | GGGGACAAGTTTGTACAAAAAAGCAGGCTATGTTAACTTCCACTGCTTATGGCA | Tb927.5.1670_rev | GGGGACCACTTTGTACAAGAAAGCTGGGTTCAACCCTCCTTTCCAATAAATTGCA |
| 5_hypothetical | G5 | Tb927.1.3400_for | GGGGACAAGTTTGTACAAAAAAGCAGGCTATGCGACATGTCGATGTGTTACA | Tb927.1.3400_rev | GGGGACCACTTTGTACAAGAAAGCTGGGTCTAAACACCTGCAATACGCTGCG |
| 5_hypothetical | G6 | Tb927.10.3330_for | GGGGACAAGTTTGTACAAAAAAGCAGGCTATGCTTCTTATAGTGTTCGTAGTCGGA | Tb927.10.3330_rev | GGGGACCACTTTGTACAAGAAAGCTGGGTTCACAACATGATTATCCTGTTACACTCTC |
| 5_hypothetical | G7 | Tb927.7.5720_for | GGGGACAAGTTTGTACAAAAAAGCAGGCTATGGTCCGCGCCCTACGC | Tb927.7.5720_rev | GGGGACCACTTTGTACAAGAAAGCTGGGTTCACTCTCTGTCCGCAGTAACC |
| 5_hypothetical | G8 | Tb927.11.11870_for | GGGGACAAGTTTGTACAAAAAAGCAGGCTATGGTAGCGAAGCTGCAGC | Tb927.11.11870_rev | GGGGACCACTTTGTACAAGAAAGCTGGGTTCAGTGGTGCGGTGGATTCC |
| 5_hypothetical | G9 | Tb927.10.9310_for | GGGGACAAGTTTGTACAAAAAAGCAGGCTATGATTTCATCGAGATTTCGAGGGG | Tb927.10.9310_rev | GGGGACCACTTTGTACAAGAAAGCTGGGTCTAGCAGTTGAGCGCCATTTTATCA |
| 5_hypothetical | G10 | Tb927.11.12740.1_for | GGGGACAAGTTTGTACAAAAAAGCAGGCTATGAGACGCACCTGCTGTGT | Tb927.11.12740.1_rev | GGGGACCACTTTGTACAAGAAAGCTGGGTTCACACACGACACAAACCCG |
| 5_hypothetical | G11 | Tb927.6.3420_for | GGGGACAAGTTTGTACAAAAAAGCAGGCTATGCGGCGAATTTTTAGCAAATCA | Tb927.6.3420_rev | GGGGACCACTTTGTACAAGAAAGCTGGGTTCAAACTTGCGTCAGCAGGAC |
| 5_hypothetical | G12 | Tb927.10.7380_for | GGGGACAAGTTTGTACAAAAAAGCAGGCTATGCTTGTTCCCGGGTTGTC | Tb927.10.7380_rev | GGGGACCACTTTGTACAAGAAAGCTGGGTTTACTTCCCTACAGCTGCCTGC |
| 5_hypothetical | G13 | Tb927.6.1630_for | GGGGACAAGTTTGTACAAAAAAGCAGGCTATGCAAACGTCCACTGGCAC | Tb927.6.1630_rev | GGGGACCACTTTGTACAAGAAAGCTGGGTTCAACCCCGCTTAAACTCGTAGG |
| 5_hypothetical | G14 | Tb927.11.8830_for | GGGGACAAGTTTGTACAAAAAAGCAGGCTATGACCTTACTCTCCCATGTAGTGG | Tb927.11.8830_rev | GGGGACCACTTTGTACAAGAAAGCTGGGTTTAAGGAGCGGTAATGCGAGC |
| 5_hypothetical | G15 | Tb927.3.3840_for | GGGGACAAGTTTGTACAAAAAAGCAGGCTATGGCCAAAATTAATGCAACTTATGGTT | Tb927.3.3840_rev | GGGGACCACTTTGTACAAGAAAGCTGGGTTCAATCGTCAAGCAAGGAAGCC |
| 5_hypothetical | G16 | Tb927.9.4760_for | GGGGACAAGTTTGTACAAAAAAGCAGGCTATGCTCTCCGGTGTGTATGGT | Tb927.9.4760_rev | GGGGACCACTTTGTACAAGAAAGCTGGGTTCATTGGAGGCTGCGGCT |
| 5_hypothetical | G17 | Tb927.8.3340_for | GGGGACAAGTTTGTACAAAAAAGCAGGCTATGGAGGGACACGGAATTGATGT | Tb927.8.3340_rev | GGGGACCACTTTGTACAAGAAAGCTGGGTTCACTCGGAACCCACGCA |
| 5_hypothetical | G18 | Tb927.7.1350_for | GGGGACAAGTTTGTACAAAAAAGCAGGCTATGTTTCGCTATCGTTGCCGC | Tb927.7.1350_rev | GGGGACCACTTTGTACAAGAAAGCTGGGTCTATAAGATGACAGACTCCGACGCA |
| 5_hypothetical | G19 | Tb927.4.990_for | GGGGACAAGTTTGTACAAAAAAGCAGGCTATGGACACCGCTGCTGAAATG | Tb927.4.990_rev | GGGGACCACTTTGTACAAGAAAGCTGGGTTCAAACAAACGATTTTCGAACGGC |
| 5_hypothetical | G20 | Tb927.3.3200_for | GGGGACAAGTTTGTACAAAAAAGCAGGCTATGACTCACTTGAGCGCGTTT | Tb927.3.3200_rev | GGGGACCACTTTGTACAAGAAAGCTGGGTTTAATAAAAACGGCGCAAACATGATTGA |
| 5_hypothetical | G21 | Tb927.10.12480_for | GGGGACAAGTTTGTACAAAAAAGCAGGCTATGCTCGATGCGGCCCCC | Tb927.10.12480_rev | GGGGACCACTTTGTACAAGAAAGCTGGGTCTAAGCCTTTGGTAGAAGAGGGTACA |
| 5_hypothetical | G22 | Tb927.6.840_for | GGGGACAAGTTTGTACAAAAAAGCAGGCTATGGTTCTCACATTGGTTGGGTC | Tb927.6.840_rev | GGGGACCACTTTGTACAAGAAAGCTGGGTCTACATGATGTTTATCTTCAGCGCCG |
| 5_hypothetical | G23 | Tb927.11.16360_for | GGGGACAAGTTTGTACAAAAAAGCAGGCTATGAAAAATTATACACCGGTAGAAATCGCA | Tb927.11.16360_rev | GGGGACCACTTTGTACAAGAAAGCTGGGTTTACCGTGGAGGAAAAGACCACC |
| 5_hypothetical | G24 | Tb927.7.3640_for | GGGGACAAGTTTGTACAAAAAAGCAGGCTATGTACGACCCCGCGGTAC | Tb927.7.3640_rev | GGGGACCACTTTGTACAAGAAAGCTGGGTTTAGGTTCCACATAAAGGCTCACCA |
| 5_hypothetical | H1 | Tb927.6.1310_for | GGGGACAAGTTTGTACAAAAAAGCAGGCTATGTGGCAAGCAATTGTACAAGTGG | Tb927.6.1310_rev | GGGGACCACTTTGTACAAGAAAGCTGGGTTCAGATAAAACAGAGGGATACTGATAGTAA |
| 5_hypothetical | H2 | Tb927.11.12500_for | GGGGACAAGTTTGTACAAAAAAGCAGGCTATGGAGAAGTACAGGAAGTTTGGTGA | Tb927.11.12500_rev | GGGGACCACTTTGTACAAGAAAGCTGGGTTTACTGAGCCATGAAAACCTCAACG |
| 5_hypothetical | H3 | Tb927.5.2120_for | GGGGACAAGTTTGTACAAAAAAGCAGGCTATGGAGAATCAGGCACCCCC | Tb927.5.2120_rev | GGGGACCACTTTGTACAAGAAAGCTGGGTCTACTTCATCATGGTATTTGTTCCAGCG |
| 5_hypothetical | H4 | Tb927.2.5220_for | GGGGACAAGTTTGTACAAAAAAGCAGGCTATGCGGTTGAGTGTTTCGGG | Tb927.2.5220_rev | GGGGACCACTTTGTACAAGAAAGCTGGGTTCAATGTGGGAGATTCTTATCCGAAGC |
| 5_hypothetical | H5 | Tb927.11.12690_for | GGGGACAAGTTTGTACAAAAAAGCAGGCTATGCAGGTTAGCGACGGCA | Tb927.11.12690_rev | GGGGACCACTTTGTACAAGAAAGCTGGGTTTATAAAGGCTTCACCGGTCGGT |
| 5_hypothetical | H6 | Tb927.2.4920_for | GGGGACAAGTTTGTACAAAAAAGCAGGCTATGAGCATGGACACCCGTATGG | Tb927.2.4920_rev | GGGGACCACTTTGTACAAGAAAGCTGGGTTTAGCCATAATATTGCAGGGCTAAGGG |
| 5_hypothetical | H7 | Tb927.8.1140_for | GGGGACAAGTTTGTACAAAAAAGCAGGCTATGTTTCTTCGCGTTCGCGG | Tb927.8.1140_rev | GGGGACCACTTTGTACAAGAAAGCTGGGTTTAGTGCCCACCACAGTGTCC |
| 5_hypothetical | H8 | Tb927.9.9660_for | GGGGACAAGTTTGTACAAAAAAGCAGGCTATGCTGAAGGAATGGCTTCGAG | Tb927.9.9660_rev | GGGGACCACTTTGTACAAGAAAGCTGGGTTTAGGCAGTGAATACCACACCATGT |
| 5_hypothetical | H9 | Tb927.5.1240_for | GGGGACAAGTTTGTACAAAAAAGCAGGCTATGCCACTAAATGCTCAACAATTGTTATT | Tb927.5.1240_rev | GGGGACCACTTTGTACAAGAAAGCTGGGTTCAAAGCAACTCCAACGGAATTCG |
| 5_hypothetical | H10 | Tb927.9.8500_for | GGGGACAAGTTTGTACAAAAAAGCAGGCTATGGTCCGAAATGAACATATACTTCTACAA | Tb927.9.8500_rev | GGGGACCACTTTGTACAAGAAAGCTGGGTCTACTCCACAGGTTGCAAAGACG |
| 5_hypothetical | H11 | Tb927.10.760_for | GGGGACAAGTTTGTACAAAAAAGCAGGCTATGCAGAAATGTTTAGTCATACGTCATGG | Tb927.10.760_rev | GGGGACCACTTTGTACAAGAAAGCTGGGTTCATATGGTGGATGGGGCATCTG |
| 5_hypothetical | H12 | Tb927.6.4420_for | GGGGACAAGTTTGTACAAAAAAGCAGGCTATGGATGATGAACGGACAACGG | Tb927.6.4420_rev | GGGGACCACTTTGTACAAGAAAGCTGGGTTCAGTCCGCCTGCCCAGA |
| 5_hypothetical | H13 | Tb927.6.680_for | GGGGACAAGTTTGTACAAAAAAGCAGGCTATGTGGCAGCTGTCAAGCTG | Tb927.6.680_rev | GGGGACCACTTTGTACAAGAAAGCTGGGTTCACTCATCGTTCCCGATTTCGT |
| 5_hypothetical | H14 | Tb927.3.5700_for | GGGGACAAGTTTGTACAAAAAAGCAGGCTATGAGGAAACAACATGTCATATACTACGTA | Tb927.3.5700_rev | GGGGACCACTTTGTACAAGAAAGCTGGGTCTAAAGCAACAATAAAGAAAGTGTTAAAAA |
| 5_hypothetical | H15 | Tb927.11.9230_for | GGGGACAAGTTTGTACAAAAAAGCAGGCTATGCTTCTGCAATTCAACTGGGA | Tb927.11.9230_rev | GGGGACCACTTTGTACAAGAAAGCTGGGTCTAAAGGGACAGTGGCACCAC |
| 5_hypothetical | H16 | Tb927.6.3440_for | GGGGACAAGTTTGTACAAAAAAGCAGGCTATGAGTAAGAGTGCAATTTACGGGTT | Tb927.6.3440_rev | GGGGACCACTTTGTACAAGAAAGCTGGGTCTACGCAAAGCTATGCGTCTGT |
| 5_hypothetical | H17 | Tb927.10.16020_for | GGGGACAAGTTTGTACAAAAAAGCAGGCTATGCTTCAACCAGAGTGTCAACC | Tb927.10.16020_rev | GGGGACCACTTTGTACAAGAAAGCTGGGTCTAAGCTATCGGACACGTCAGAATTAA |
| 5_hypothetical | H20 | Tb927.9.2060_for | GGGGACAAGTTTGTACAAAAAAGCAGGCTATGTTTTCGCTTGTCCGGGG | Tb927.9.2060_rev | GGGGACCACTTTGTACAAGAAAGCTGGGTTTATTTACCATCATCTGGTAGTGGCTGC |
| 5_hypothetical | H21 | Tb927.1.2730_for | GGGGACAAGTTTGTACAAAAAAGCAGGCTATGTTTAGATTGTCATTTGCCGCATTC | Tb927.1.2730_rev | GGGGACCACTTTGTACAAGAAAGCTGGGTCTATAGAGGCAGAAACCCAAAGCG |
| 5_hypothetical | H23 | Tb927.7.3200_for | GGGGACAAGTTTGTACAAAAAAGCAGGCTATGAAAAGCACGGTAACGATGGC | Tb927.7.3200_rev | GGGGACCACTTTGTACAAGAAAGCTGGGTCTAGCGCGGCAATTCCTTTCC |
| 5_hypothetical | H24 | Tb927.10.2500_for | GGGGACAAGTTTGTACAAAAAAGCAGGCTATGAAGCCCTTTCCCGCCT | Tb927.10.2500_rev | GGGGACCACTTTGTACAAGAAAGCTGGGTTCATAAGTGTCCATGCGGCG |
| 5_hypothetical | I1 | Tb927.7.4200_for | GGGGACAAGTTTGTACAAAAAAGCAGGCTATGTGGTTTACATCGCTTGCCC | Tb927.7.4200_rev | GGGGACCACTTTGTACAAGAAAGCTGGGTTTATGCACCACGCCGACCA |
| 5_hypothetical | I2 | Tb927.8.1380_for | GGGGACAAGTTTGTACAAAAAAGCAGGCTATGGAAATGGCGGCGCCA | Tb927.8.1380_rev | GGGGACCACTTTGTACAAGAAAGCTGGGTTTATTCAGGTCTACGTCCACCAAGA |
| 5_hypothetical | I3 | Tb927.11.16900_for | GGGGACAAGTTTGTACAAAAAAGCAGGCTATGTCCATCGCAGCGTCTAGT | Tb927.11.16900_rev | GGGGACCACTTTGTACAAGAAAGCTGGGTCTAATTCATTGGTGATAACGACGTTAAACG |
| 5_hypothetical | I4 | Tb927.10.8100_for | GGGGACAAGTTTGTACAAAAAAGCAGGCTATGATATGCAAGGTTTTGGAGCGT | Tb927.10.8100_rev | GGGGACCACTTTGTACAAGAAAGCTGGGTCTACGACGCAGACCACGTG |
| 5_hypothetical | I5 | Tb927.11.3830_for | GGGGACAAGTTTGTACAAAAAAGCAGGCTATGGGGCACTCGAAGGAAACT | Tb927.11.3830_rev | GGGGACCACTTTGTACAAGAAAGCTGGGTCTATATGGTGTTCTCTTCCGCAGAAAG |
| 5_hypothetical | I6 | Tb927.10.2940_for | GGGGACAAGTTTGTACAAAAAAGCAGGCTATGTCAACCGGAGCATTTACTGAG | Tb927.10.2940_rev | GGGGACCACTTTGTACAAGAAAGCTGGGTTTATCGCAAGTCTTCCTCACCGT |
| 5_hypothetical | I7 | Tb927.11.13600_for | GGGGACAAGTTTGTACAAAAAAGCAGGCTATGCGCAGTTGGGCTGCA | Tb927.11.13600_rev | GGGGACCACTTTGTACAAGAAAGCTGGGTTTAGCGCTGCCGCCGTCT |
| 5_hypothetical | I8 | Tb927.11.700_for | GGGGACAAGTTTGTACAAAAAAGCAGGCTATGGGGGATAAAGTTGAATATGGCAC | Tb927.11.700_rev | GGGGACCACTTTGTACAAGAAAGCTGGGTTCACTTAGCACCCTTAAACGGC |
| 5_hypothetical | I9 | Tb927.9.9630_for | GGGGACAAGTTTGTACAAAAAAGCAGGCTATGACAACCGCAACGGAATTGA | Tb927.9.9630_rev | GGGGACCACTTTGTACAAGAAAGCTGGGTTCAAAACGGTTGCTGAGAGAACTG |
| 5_hypothetical | I10 | Tb927.7.1330_for | GGGGACAAGTTTGTACAAAAAAGCAGGCTATGTTTTGTGACGCTGAAGATGGG | Tb927.7.1330_rev | GGGGACCACTTTGTACAAGAAAGCTGGGTTCAGCACCGCAAAAGCACC |
| 5_hypothetical | I11 | Tb927.3.840_for | GGGGACAAGTTTGTACAAAAAAGCAGGCTATGAATGTTTCTGTGGTGCTTCCA | Tb927.3.840_rev | GGGGACCACTTTGTACAAGAAAGCTGGGTTTAAGGGGTTTGCAACAACCAAGA |
| 5_hypothetical | I12 | Tb927.8.580_for | GGGGACAAGTTTGTACAAAAAAGCAGGCTATGAGTTCACACTCTTCTCTTGGCG | Tb927.8.580_rev | GGGGACCACTTTGTACAAGAAAGCTGGGTCTATGACGGATGCCTCGTTTTTGT |
| 5_hypothetical | I13 | Tb927.3.3910_for | GGGGACAAGTTTGTACAAAAAAGCAGGCTATGACCTCAGTGCCGGGTG | Tb927.3.3910_rev | GGGGACCACTTTGTACAAGAAAGCTGGGTTCACGACATCCACTCCTTGAGG |
| 5_hypothetical | I14 | Tb927.11.14420_for | GGGGACAAGTTTGTACAAAAAAGCAGGCTATGGGTGACTGCGACAGGT | Tb927.11.14420_rev | GGGGACCACTTTGTACAAGAAAGCTGGGTTTACTCGAAAAACAGGGTGGGACC |
| 5_hypothetical | I15 | Tb927.11.3050_for | GGGGACAAGTTTGTACAAAAAAGCAGGCTATGGAGTGCAAGACTAACTTGGC | Tb927.11.3050_rev | GGGGACCACTTTGTACAAGAAAGCTGGGTTCATCTAAATACCATACCAATATCGGACGA |
| 5_hypothetical | I16 | Tb927.7.510_for | GGGGACAAGTTTGTACAAAAAAGCAGGCTATGGGAAGCGGATGTTCAACC | Tb927.7.510_rev | GGGGACCACTTTGTACAAGAAAGCTGGGTCTACCCGCCACTGTTCAAAGC |
| 5_hypothetical | I17 | Tb927.11.2760_for | GGGGACAAGTTTGTACAAAAAAGCAGGCTATGGAAACGGAGGGGGTCG | Tb927.11.2760_rev | GGGGACCACTTTGTACAAGAAAGCTGGGTTCACTTACTTAGGTTCCGCACAAAC |
| 5_hypothetical | I18 | Tb927.3.900_for | GGGGACAAGTTTGTACAAAAAAGCAGGCTATGAAAACCTTACACCTGTTGCTTGG | Tb927.3.900_rev | GGGGACCACTTTGTACAAGAAAGCTGGGTTCACTTAAAATAAACAAAGCTATTCCCTTG |
| 5_hypothetical | I19 | Tb927.11.12670_for | GGGGACAAGTTTGTACAAAAAAGCAGGCTATGTTCGACTTGATGGCAGACG | Tb927.11.12670_rev | GGGGACCACTTTGTACAAGAAAGCTGGGTCTACGCATCGGGGTGTTTGC |
| 5_hypothetical | I20 | Tb927.11.12030_for | GGGGACAAGTTTGTACAAAAAAGCAGGCTATGGCTTGTGTAACCGGTGAC | Tb927.11.12030_rev | GGGGACCACTTTGTACAAGAAAGCTGGGTTTACTGCTGCCAGTTGATATCATCG |
| 5_hypothetical | I21 | Tb927.9.11220_for | GGGGACAAGTTTGTACAAAAAAGCAGGCTATGGCGTCGCGTTTGGCT | Tb927.9.11220_rev | GGGGACCACTTTGTACAAGAAAGCTGGGTTTACAGAATCTCCACATCCCCAACG |
| 5_hypothetical | I23 | Tb927.9.13540_for | GGGGACAAGTTTGTACAAAAAAGCAGGCTATGCGCCGCTTAGCTACCG | Tb927.9.13540_rev | GGGGACCACTTTGTACAAGAAAGCTGGGTCTAAAGAATCTCTGACAAAAAACGATTCCT |
| 5_hypothetical | I24 | Tb927.10.1600_for | GGGGACAAGTTTGTACAAAAAAGCAGGCTATGCGGCATACAACACAACATTACA | Tb927.10.1600_rev | GGGGACCACTTTGTACAAGAAAGCTGGGTTTACTTCAGAACTTGCCTGACGGA |
| 5_hypothetical | J1 | Tb927.11.15770_for | GGGGACAAGTTTGTACAAAAAAGCAGGCTATGCAGGTGCATCACAGTGTG | Tb927.11.15770_rev | GGGGACCACTTTGTACAAGAAAGCTGGGTTCAGCTGGCTCCCTTAATATACGG |
| 5_hypothetical | J2 | Tb927.5.3310_for | GGGGACAAGTTTGTACAAAAAAGCAGGCTATGACGGATCCACAGGAGGATAAC | Tb927.5.3310_rev | GGGGACCACTTTGTACAAGAAAGCTGGGTTCATGCAAGTGACAGGAAACCC |
| 5_hypothetical | J3 | Tb927.2.4800_for | GGGGACAAGTTTGTACAAAAAAGCAGGCTATGTTACGTCCGTCGTCCCT | Tb927.2.4800_rev | GGGGACCACTTTGTACAAGAAAGCTGGGTCTATTCATTATTCAGCAACCGATTCGC |
| 5_hypothetical | J4 | Tb927.11.13770_for | GGGGACAAGTTTGTACAAAAAAGCAGGCTATGTCCACAAGTGGCATAGTCCG | Tb927.11.13770_rev | GGGGACCACTTTGTACAAGAAAGCTGGGTCTAAAGAACCCATTGCTGGATGATAACA |
| 5_hypothetical | J5 | Tb927.8.2110_for | GGGGACAAGTTTGTACAAAAAAGCAGGCTATGCAGGGCGCGACAGTC | Tb927.8.2110_rev | GGGGACCACTTTGTACAAGAAAGCTGGGTTTAGGCCAACAACTCAGCAAGTG |
| 5_hypothetical | J6 | Tb927.10.1730_for | GGGGACAAGTTTGTACAAAAAAGCAGGCTATGTGGCGTTGCTCTACTCTACT | Tb927.10.1730_rev | GGGGACCACTTTGTACAAGAAAGCTGGGTTCAACTTTTCCCCACAGTTTTTTCCC |
| 5_hypothetical | J7 | Tb927.11.16300_for | GGGGACAAGTTTGTACAAAAAAGCAGGCTATGGAGGTGGTTGGGCGT | Tb927.11.16300_rev | GGGGACCACTTTGTACAAGAAAGCTGGGTTTATGCAACATTGCACGTCGTCA |
| 5_hypothetical | J8 | Tb927.10.4510_for | GGGGACAAGTTTGTACAAAAAAGCAGGCTATGACTGACCGAAATGGGGAACA | Tb927.10.4510_rev | GGGGACCACTTTGTACAAGAAAGCTGGGTTCACGGATAACTGGATAGCGCT |
| 5_hypothetical | J9 | Tb927.3.5190_for | GGGGACAAGTTTGTACAAAAAAGCAGGCTATGTTCGTTCGGAAATTTCAGGGA | Tb927.3.5190_rev | GGGGACCACTTTGTACAAGAAAGCTGGGTTTAGATACGAAAATTTGCTTTAGATGTGTT |
| 5_hypothetical | J10 | Tb927.3.3810_for | GGGGACAAGTTTGTACAAAAAAGCAGGCTATGTCTTCTGGTGTCCCTGTTTCA | Tb927.3.3810_rev | GGGGACCACTTTGTACAAGAAAGCTGGGTTCATTGCGCCACATGCACA |
| 5_hypothetical | J11 | Tb927.4.1610_for | GGGGACAAGTTTGTACAAAAAAGCAGGCTATGATTCCTCCGGCAAAATTAAATGATTTC | Tb927.4.1610_rev | GGGGACCACTTTGTACAAGAAAGCTGGGTTCAACCGGATTCCACGGAAAGA |
| 5_hypothetical | J12 | Tb927.8.2350_for | GGGGACAAGTTTGTACAAAAAAGCAGGCTATGGAGGCGGGCACATCC | Tb927.8.2350_rev | GGGGACCACTTTGTACAAGAAAGCTGGGTTTAGCTGCTCAACCTAAGCCAGA |
| 5_hypothetical | J13 | Tb927.3.4400_for | GGGGACAAGTTTGTACAAAAAAGCAGGCTATGGACTACACTGACATATATTTGCGG | Tb927.3.4400_rev | GGGGACCACTTTGTACAAGAAAGCTGGGTTCATACGTCCTTCACTGCCCC |
| 5_hypothetical | J14 | Tb927.10.1860_for | GGGGACAAGTTTGTACAAAAAAGCAGGCTATGACAGATAATCAAGGTAGCTATATCACT | Tb927.10.1860_rev | GGGGACCACTTTGTACAAGAAAGCTGGGTTCAGAGATCGATCAGCTTCCTTCC |
| 5_hypothetical | J15 | Tb927.11.1300_for | GGGGACAAGTTTGTACAAAAAAGCAGGCTATGAATGTTTTTGATTCCAACAGCATGC | Tb927.11.1300_rev | GGGGACCACTTTGTACAAGAAAGCTGGGTTCAAGAGCTATCTTTGTTCACAATGAAGT |
| 5_hypothetical | J16 | Tb927.10.4720_for | GGGGACAAGTTTGTACAAAAAAGCAGGCTATGAGCACATCAGCAACAAGCA | Tb927.10.4720_rev | GGGGACCACTTTGTACAAGAAAGCTGGGTTTAACAGAGCTTCAAGTACTCCAATGTG |
| 5_hypothetical | J17 | Tb927.7.5030_for | GGGGACAAGTTTGTACAAAAAAGCAGGCTATGATTCTATCCCTCGACATGAAAAATGC | Tb927.7.5030_rev | GGGGACCACTTTGTACAAGAAAGCTGGGTTCATTCCCGTGCGATCGCA |
| 5_hypothetical | J18 | Tb927.10.11670_for | GGGGACAAGTTTGTACAAAAAAGCAGGCTATGCTGCCTCGTTACACCCC | Tb927.10.11670_rev | GGGGACCACTTTGTACAAGAAAGCTGGGTTTAATTACAGTATCTGTCGTAGAGTCCCTC |
| 5_hypothetical | J19 | Tb927.7.1080_for | GGGGACAAGTTTGTACAAAAAAGCAGGCTATGATGCAACGTTGCGGCA | Tb927.7.1080_rev | GGGGACCACTTTGTACAAGAAAGCTGGGTTTAATCCATCTTCCTCCCGTATTTGTCT |
| 5_hypothetical | J20 | Tb927.10.9290_for | GGGGACAAGTTTGTACAAAAAAGCAGGCTATGACAGTTTGTCTTGTTCTACTAACTGG | Tb927.10.9290_rev | GGGGACCACTTTGTACAAGAAAGCTGGGTTCAAAGATCTACTAAGGCATGCTCGA |
| 5_hypothetical | J21 | Tb927.5.2000_for | GGGGACAAGTTTGTACAAAAAAGCAGGCTATGCGCCCCACCGGAGTA | Tb927.5.2000_rev | GGGGACCACTTTGTACAAGAAAGCTGGGTTCACCCCCTTCTTCCGTTCAT |
| 5_hypothetical | J22 | Tb927.11.13100_for | GGGGACAAGTTTGTACAAAAAAGCAGGCTATGCAGCAGAAGAAGTGCTACAC | Tb927.11.13100_rev | GGGGACCACTTTGTACAAGAAAGCTGGGTTCATGTTGCCGTTGAAACAGATTGC |
| 5_hypothetical | J23 | Tb927.5.4290_for | GGGGACAAGTTTGTACAAAAAAGCAGGCTATGACATCGAAGACCCCTTCGC | Tb927.5.4290_rev | GGGGACCACTTTGTACAAGAAAGCTGGGTTCACATGTATTCTGCCTCAGGTGT |
| 5_hypothetical | J24 | Tb927.10.13990_for | GGGGACAAGTTTGTACAAAAAAGCAGGCTATGCCACCTCTTACTCTTGAACAGG | Tb927.10.13990_rev | GGGGACCACTTTGTACAAGAAAGCTGGGTCTAAAAGCGCCAGCGGCG |
| 5_hypothetical | K1 | Tb927.9.3930_for | GGGGACAAGTTTGTACAAAAAAGCAGGCTATGGTCGTCAAACGCCGTG | Tb927.9.3930_rev | GGGGACCACTTTGTACAAGAAAGCTGGGTTCAAGGCGACAAGAACCCGG |
| 5_hypothetical | K2 | Tb927.3.3500_for | GGGGACAAGTTTGTACAAAAAAGCAGGCTATGTCGAGCTGTGAGCGACT | Tb927.3.3500_rev | GGGGACCACTTTGTACAAGAAAGCTGGGTTTAGTTCATGAGTGAGAAGCCGTTG |
| 5_hypothetical | K3 | Tb927.6.4160_for | GGGGACAAGTTTGTACAAAAAAGCAGGCTATGAGTACTGTGTTTCCCCTTCTTCG | Tb927.6.4160_rev | GGGGACCACTTTGTACAAGAAAGCTGGGTCTACTTCAAAAGGCAGTGCGACG |
| 5_hypothetical | K4 | Tb927.9.9090_for | GGGGACAAGTTTGTACAAAAAAGCAGGCTATGGGAAGCCAGATTCCGCA | Tb927.9.9090_rev | GGGGACCACTTTGTACAAGAAAGCTGGGTTTACACGTGTAGTAGCTGCGC |
| 5_hypothetical | K5 | Tb927.8.1350_for | GGGGACAAGTTTGTACAAAAAAGCAGGCTATGCAGAAACTGTGGCACGAC | Tb927.8.1350_rev | GGGGACCACTTTGTACAAGAAAGCTGGGTTTAATCACGACCGTTGGGTCTACT |
| 5_hypothetical | K6 | Tb927.11.3100_for | GGGGACAAGTTTGTACAAAAAAGCAGGCTATGACCGCACTGTCAAATGGG | Tb927.11.3100_rev | GGGGACCACTTTGTACAAGAAAGCTGGGTCTAGCGGGAGGAAAACCGC |
| 5_hypothetical | K8 | Tb927.9.6920_for | GGGGACAAGTTTGTACAAAAAAGCAGGCTATGACACCAACAGTAACCCCTCT | Tb927.9.6920_rev | GGGGACCACTTTGTACAAGAAAGCTGGGTCTAGCGCTTAGCGTTCCACTG |
| 5_hypothetical | K9 | Tb927.11.10600_for | GGGGACAAGTTTGTACAAAAAAGCAGGCTATGCTTTGTCACAATCGACTGCT | Tb927.11.10600_rev | GGGGACCACTTTGTACAAGAAAGCTGGGTTTAGCGAACGAATTGAGGAGGAGG |
| 5_hypothetical | K10 | Tb927.7.870_for | GGGGACAAGTTTGTACAAAAAAGCAGGCTATGCTCCTTCCGTTTCCACGG | Tb927.7.870_rev | GGGGACCACTTTGTACAAGAAAGCTGGGTTCATTCCAAGGCACGGCAGG |
| 5_hypothetical | K11 | Tb927.1.3550_for | GGGGACAAGTTTGTACAAAAAAGCAGGCTATGGGTGTTGGTTTTCAGAGTAAAAAAC | Tb927.1.3550_rev | GGGGACCACTTTGTACAAGAAAGCTGGGTCTAACAATGCGCGTCAAGCCC |
| 5_hypothetical | K13 | Tb927.7.5550_for | GGGGACAAGTTTGTACAAAAAAGCAGGCTATGGTCGGTAAGTGTCCACGA | Tb927.7.5550_rev | GGGGACCACTTTGTACAAGAAAGCTGGGTTCACTGGGCACAGAAGTGCC |
| 5_hypothetical | K14 | Tb927.11.7520_for | GGGGACAAGTTTGTACAAAAAAGCAGGCTATGAACGGAGAGGAAGGGAAGG | Tb927.11.7520_rev | GGGGACCACTTTGTACAAGAAAGCTGGGTTCAGTTGGGGGGACGGAGG |
| 5_hypothetical | K15 | Tb927.4.3100_for | GGGGACAAGTTTGTACAAAAAAGCAGGCTATGCCGTTATGTGTTCCTCTTCTTC | Tb927.4.3100_rev | GGGGACCACTTTGTACAAGAAAGCTGGGTTTAAATTGCAATGGAAACCCGCTTAC |
| 5_hypothetical | K16 | Tb927.8.3720_for | GGGGACAAGTTTGTACAAAAAAGCAGGCTATGAAAGCATGCTGTGGCGG | Tb927.8.3720_rev | GGGGACCACTTTGTACAAGAAAGCTGGGTTCAGATAGTCAGTGGGTTTCGCC |
| 5_hypothetical | K17 | Tb927.11.6610_for | GGGGACAAGTTTGTACAAAAAAGCAGGCTATGAGTGAACAGAATAGGGATGCGG | Tb927.11.6610_rev | GGGGACCACTTTGTACAAGAAAGCTGGGTTTAATCCGCCGAGCGGTGT |
| 5_hypothetical | K18 | Tb927.10.550_for | GGGGACAAGTTTGTACAAAAAAGCAGGCTATGAACGTTTACGGCAAGGACC | Tb927.10.550_rev | GGGGACCACTTTGTACAAGAAAGCTGGGTCTATCTATGCGTCGAGTCTCCAATAATATC |
| 5_hypothetical | K19 | Tb927.3.2830_for | GGGGACAAGTTTGTACAAAAAAGCAGGCTATGGGGAAATCTAAAAAGATGCCGG | Tb927.3.2830_rev | GGGGACCACTTTGTACAAGAAAGCTGGGTTTAGTCCATATCAGCTCCGTAGCC |
| 5_hypothetical | K20 | Tb927.6.3940_for | GGGGACAAGTTTGTACAAAAAAGCAGGCTATGCCGGGACACGGAAAGG | Tb927.6.3940_rev | GGGGACCACTTTGTACAAGAAAGCTGGGTTTAAACGTTTCCAGCTGGCGC |
| 5_hypothetical | K21 | Tb927.10.9240_for | GGGGACAAGTTTGTACAAAAAAGCAGGCTATGGATGACAAACAGGCGTTCTT | Tb927.10.9240_rev | GGGGACCACTTTGTACAAGAAAGCTGGGTCTAGTTAAGCAGTTCATTTATGAGCGAAC |
| 5_hypothetical | K22 | Tb927.2.4760_for | GGGGACAAGTTTGTACAAAAAAGCAGGCTATGAGCATGGACACCCGTATGG | Tb927.2.4760_rev | GGGGACCACTTTGTACAAGAAAGCTGGGTCTAATACGATTGACCGGTCATCATTTGG |
| 5_hypothetical | K23 | Tb927.4.4340_for | GGGGACAAGTTTGTACAAAAAAGCAGGCTATGGTGCAACGAGAAGTATTTACTTTTG | Tb927.4.4340_rev | GGGGACCACTTTGTACAAGAAAGCTGGGTTCAAAAACTGTGATACCACACAACAAC |
| 5_hypothetical | L1 | Tb927.5.4140_for | GGGGACAAGTTTGTACAAAAAAGCAGGCTATGCTACCCTTTGTTGCCCTGC | Tb927.5.4140_rev | GGGGACCACTTTGTACAAGAAAGCTGGGTTCATATTTCAATTCGCCCAAGGAGC |
| 5_hypothetical | L2 | Tb927.10.7600_for | GGGGACAAGTTTGTACAAAAAAGCAGGCTATGCCGGACGGAGCTATTGG | Tb927.10.7600_rev | GGGGACCACTTTGTACAAGAAAGCTGGGTTTAACGAGTAGAACGCGAGCCA |
| 5_hypothetical | L3 | Tb927.8.2800_for | GGGGACAAGTTTGTACAAAAAAGCAGGCTATGGGTTGTCTCCCTCCCG | Tb927.8.2800_rev | GGGGACCACTTTGTACAAGAAAGCTGGGTTCACGAACTCAAAACTGAGCGAA |
| 5_hypothetical | L4 | Tb927.10.7970_for | GGGGACAAGTTTGTACAAAAAAGCAGGCTATGGCCATACATGTCGCTCGT | Tb927.10.7970_rev | GGGGACCACTTTGTACAAGAAAGCTGGGTCTACTGATTACGCACGATAGCTTTGA |
| 5_hypothetical | L5 | Tb927.9.7260_for | GGGGACAAGTTTGTACAAAAAAGCAGGCTATGCGATCCGGTCGCAAAC | Tb927.9.7260_rev | GGGGACCACTTTGTACAAGAAAGCTGGGTTTACGCCTCCTCTTCGGATGG |
| 5_hypothetical | L6 | Tb927.7.1360_for | GGGGACAAGTTTGTACAAAAAAGCAGGCTATGCCTGAGCAGTTGGGCG | Tb927.7.1360_rev | GGGGACCACTTTGTACAAGAAAGCTGGGTCTACATGCCGCCATCCTCTTG |
| 5_hypothetical | L7 | Tb927.10.8640_for | GGGGACAAGTTTGTACAAAAAAGCAGGCTATGTCGACCGAGGTTGTTCACA | Tb927.10.8640_rev | GGGGACCACTTTGTACAAGAAAGCTGGGTCTATCCACCTTTCTTTCGGGAACC |
| 5_hypothetical | L8 | Tb927.8.2120_for | GGGGACAAGTTTGTACAAAAAAGCAGGCTATGTGGGAAGAAACGTACGTACG | Tb927.8.2120_rev | GGGGACCACTTTGTACAAGAAAGCTGGGTCTACTCCCTTGCCACCATAGGAC |
| 5_hypothetical | L9 | Tb927.11.9910_for | GGGGACAAGTTTGTACAAAAAAGCAGGCTATGTCTACCGGTGAGAGCGC | Tb927.11.9910_rev | GGGGACCACTTTGTACAAGAAAGCTGGGTTTAAGTTGAGGCGAACATCTTGGC |
| 5_hypothetical | L10 | Tb927.7.1870_for | GGGGACAAGTTTGTACAAAAAAGCAGGCTATGAGTAGAGATGCGACACCCTAC | Tb927.7.1870_rev | GGGGACCACTTTGTACAAGAAAGCTGGGTTTACTCGGGGAACTTGGGAAGC |
| 5_hypothetical | L11 | Tb927.9.1650_for | GGGGACAAGTTTGTACAAAAAAGCAGGCTATGTGGGGATTGACAAGGAGATGC | Tb927.9.1650_rev | GGGGACCACTTTGTACAAGAAAGCTGGGTTCAACGAACAAAAGCAATATCACAATGAA |
| 5_hypothetical | L13 | Tb927.5.3130_for | GGGGACAAGTTTGTACAAAAAAGCAGGCTATGAAAAATTTTGTGAATGCAAACCCCA | Tb927.5.3130_rev | GGGGACCACTTTGTACAAGAAAGCTGGGTTTACGTGGGACCAAATAAGGGAAAG |
| 5_hypothetical | L14 | Tb927.2.5290_for | GGGGACAAGTTTGTACAAAAAAGCAGGCTATGAGCATGGACACCCGTATGG | Tb927.2.5290_rev | GGGGACCACTTTGTACAAGAAAGCTGGGTTTATTCACTGGTTCTCATTTGGTTATAATA |
| 5_hypothetical | L15 | Tb927.9.2320_for | GGGGACAAGTTTGTACAAAAAAGCAGGCTATGCTGGCGTTGTGCAAATGT | Tb927.9.2320_rev | GGGGACCACTTTGTACAAGAAAGCTGGGTTCAGCGGCGCCGCCACAC |
| 5_hypothetical | L16 | Tb927.9.7090_for | GGGGACAAGTTTGTACAAAAAAGCAGGCTATGTACTCTCCAAGTGAAGAAGCAGG | Tb927.9.7090_rev | GGGGACCACTTTGTACAAGAAAGCTGGGTCTATCGTGCGAAACGGCAGT |

| | | | | | |
|---|---|---|---|---|---|
| 5_hypothetical | L17 | Tb927.8.3280_for | GGGGACAAGTTTGTACAAAAAAGCAGGCTATGAAGGGAGGCGTGGGAAG | Tb927.8.3280_rev | GGGGACCACTTTGTACAAGAAAGCTGGGTTCATAAACGTCCACTAAGGTGCTGC |
| 5_hypothetical | L18 | Tb927.3.4260_for | GGGGACAAGTTTGTACAAAAAAGCAGGCTATGTTGCGTCGCTGTTTTACCC | Tb927.3.4260_rev | GGGGACCACTTTGTACAAGAAAGCTGGGTCTACCGCCTGCTATTGCGC |
| 5_hypothetical | L19 | Tb927.10.4060_for | GGGGACAAGTTTGTACAAAAAAGCAGGCTATGCAACAAGTTGGTGAACCATTTC | Tb927.10.4060_rev | GGGGACCACTTTGTACAAGAAAGCTGGGTCTAGGCCGCCTTCGGTCG |
| 5_hypothetical | L20 | Tb927.5.2050_for | GGGGACAAGTTTGTACAAAAAAGCAGGCTATGCTCCAATGGGTCGTGAAAG | Tb927.5.2050_rev | GGGGACCACTTTGTACAAGAAAGCTGGGTCTATCGAGTCCGCTCTCGCT |
| 5_hypothetical | L21 | Tb927.11.16820_for | GGGGACAAGTTTGTACAAAAAAGCAGGCTATGACATTTTTCGGGTTAATGCGGT | Tb927.11.16820_rev | GGGGACCACTTTGTACAAGAAAGCTGGGTTCATCCGCCAGGCCCTGC |
| 5_hypothetical | L22 | Tb927.8.3640_for | GGGGACAAGTTTGTACAAAAAAGCAGGCTATGATGGCTGCTCGTCGTGG | Tb927.8.3640_rev | GGGGACCACTTTGTACAAGAAAGCTGGGTTTAAACTATTCTCCGGTCACGCTTC |
| 5_hypothetical | L23 | Tb927.2.2860_for | GGGGACAAGTTTGTACAAAAAAGCAGGCTATGGTTCATTTTACTGTTTATTACGGCAAC | Tb927.2.2860_rev | GGGGACCACTTTGTACAAGAAAGCTGGGTCTAATTCTTTTTTGACCGCTTCCGTT |
| 5_hypothetical | L24 | Tb927.8.2690_for | GGGGACAAGTTTGTACAAAAAAGCAGGCTATGGACCCCTCTCAAGAGGGA | Tb927.8.2690_rev | GGGGACCACTTTGTACAAGAAAGCTGGGTTCACTCTTCCTCTTCACAAAGAGGA |
| 5_hypothetical | M1 | Tb927.10.12220_for | GGGGACAAGTTTGTACAAAAAAGCAGGCTATGAGAAAGGGAATATTTTCTCACATCATC | Tb927.10.12220_rev | GGGGACCACTTTGTACAAGAAAGCTGGGTTCAAGGGATCATAATCTGCCGTAGG |
| 5_hypothetical | M2 | Tb927.3.3030_for | GGGGACAAGTTTGTACAAAAAAGCAGGCTATGTCGACAGAAGATGTGTTTGTTGA | Tb927.3.3030_rev | GGGGACCACTTTGTACAAGAAAGCTGGGTTTAATGGGGAGATGTACGCCGG |
| 5_hypothetical | M5 | Tb927.1.4140_for | GGGGACAAGTTTGTACAAAAAAGCAGGCTATGCCAGCTGGTGGTGACA | Tb927.1.4140_rev | GGGGACCACTTTGTACAAGAAAGCTGGGTTTATGTCACGACATTCCACAAATCCT |
| 5_hypothetical | M6 | Tb927.7.4910_for | GGGGACAAGTTTGTACAAAAAAGCAGGCTATGTTCTCCACTACCACCGGTT | Tb927.7.4910_rev | GGGGACCACTTTGTACAAGAAAGCTGGGTTCAGTCCACCAGCACCTGG |
| 5_hypothetical | M9 | Tb927.8.6020_for | GGGGACAAGTTTGTACAAAAAAGCAGGCTATGGGTATGGGTCTCACCGG | Tb927.8.6020_rev | GGGGACCACTTTGTACAAGAAAGCTGGGTTTAGTTCTCGCGCACGGC |
| 5_hypothetical | M10 | Tb927.3.970_for | GGGGACAAGTTTGTACAAAAAAGCAGGCTATGTCGGCCATTAAACGCTCC | Tb927.3.970_rev | GGGGACCACTTTGTACAAGAAAGCTGGGTCTAGTAAAAGTCCTCCACCCCAAATTC |
| 5_hypothetical | M11 | Tb927.5.2930_for | GGGGACAAGTTTGTACAAAAAAGCAGGCTATGCGCCGTGTATCTTCCCC | Tb927.5.2930_rev | GGGGACCACTTTGTACAAGAAAGCTGGGTTTAGTGATGGGCCTTTTTCTTTTGTTCG |
| 5_hypothetical | M12 | Tb927.11.6940_for | GGGGACAAGTTTGTACAAAAAAGCAGGCTATGAGTGGCGCTCCCACG | Tb927.11.6940_rev | GGGGACCACTTTGTACAAGAAAGCTGGGTTCACTCCCAAGGGATAAGTCGTT |
| 5_hypothetical | M13 | Tb927.10.12160_for | GGGGACAAGTTTGTACAAAAAAGCAGGCTATGCGTGCAACTGCGGAC | Tb927.10.12160_rev | GGGGACCACTTTGTACAAGAAAGCTGGGTTCATTTTGAGTTAGAATGCGGGGAA |
| 5_hypothetical | M14 | Tb927.8.6620_for | GGGGACAAGTTTGTACAAAAAAGCAGGCTATGGCATTAGCACACGCCC | Tb927.8.6620_rev | GGGGACCACTTTGTACAAGAAAGCTGGGTCTAGGCACCAACAAGCACATTTACA |
| 5_hypothetical | M15 | Tb927.10.11410_for | GGGGACAAGTTTGTACAAAAAAGCAGGCTATGTTGCAGGGGGAAATGGC | Tb927.10.11410_rev | GGGGACCACTTTGTACAAGAAAGCTGGGTTCACTGCGCCGCGTTCGA |
| 5_hypothetical | M16 | Tb927.9.4250_for | GGGGACAAGTTTGTACAAAAAAGCAGGCTATGCTTTCGACGGTTTGTGGT | Tb927.9.4250_rev | GGGGACCACTTTGTACAAGAAAGCTGGGTTTATGATTTCAGTTTTACGCGTATGCC |
| 5_hypothetical | M17 | Tb927.11.5670_for | GGGGACAAGTTTGTACAAAAAAGCAGGCTATGAACAAATCCGCAAAAAAGAAGGG | Tb927.11.5670_rev | GGGGACCACTTTGTACAAGAAAGCTGGGTTTACCTTCGGCAACGCCG |
| 5_hypothetical | M18 | Tb927.11.13490_for | GGGGACAAGTTTGTACAAAAAAGCAGGCTATGACGGAGTATGGCCCAAGG | Tb927.11.13490_rev | GGGGACCACTTTGTACAAGAAAGCTGGGTCTAGAACAAGCCGCTTCTCCTAAG |
| 5_hypothetical | M19 | Tb927.9.13630_for | GGGGACAAGTTTGTACAAAAAAGCAGGCTATGCGGTCTATACCTTACGGGAG | Tb927.9.13630_rev | GGGGACCACTTTGTACAAGAAAGCTGGGTTTACTTCAGGTGTGGTAAGCGC |
| 5_hypothetical | M20 | Tb927.8.6530_for | GGGGACAAGTTTGTACAAAAAAGCAGGCTATGAACAGGGGGTCGGTGG | Tb927.8.6530_rev | GGGGACCACTTTGTACAAGAAAGCTGGGTTTACCTCTCCCTCCTGCATCG |
| 5_hypothetical | M21 | Tb927.11.14750_for | GGGGACAAGTTTGTACAAAAAAGCAGGCTATGTTCCACGATCTGACTAACGTG | Tb927.11.14750_rev | GGGGACCACTTTGTACAAGAAAGCTGGGTTTATTTGTACGTTGGAGGGCGG |
| 5_hypothetical | M22 | Tb927.10.3630_for | GGGGACAAGTTTGTACAAAAAAGCAGGCTATGGCGACACCGCGGACG | Tb927.10.3630_rev | GGGGACCACTTTGTACAAGAAAGCTGGGTTCACACCGATGTGGAGAGTTCC |
| 5_hypothetical | M23 | Tb927.5.420_for | GGGGACAAGTTTGTACAAAAAAGCAGGCTATGGGCGCAAGATTGCGT | Tb927.5.420_rev | GGGGACCACTTTGTACAAGAAAGCTGGGTCTAACAATCCTCTTCTTCCTCAGCG |
| 5_hypothetical | M24 | Tb927.8.6560_for | GGGGACAAGTTTGTACAAAAAAGCAGGCTATGAACACAGTGAAAGTTCTGTTACGG | Tb927.8.6560_rev | GGGGACCACTTTGTACAAGAAAGCTGGGTTCAACGCAAGAGGAGGGGC |
| 5_hypothetical | N1 | Tb927.10.430_for | GGGGACAAGTTTGTACAAAAAAGCAGGCTATGCGGGTAGCATCTTTTAGGGC | Tb927.10.430_rev | GGGGACCACTTTGTACAAGAAAGCTGGGTCTAATCTGTTGATGCGTCGTATGGG |
| 5_hypothetical | N2 | Tb927.7.3290_for | GGGGACAAGTTTGTACAAAAAAGCAGGCTATGCTGCTCTCTGTCCCCG | Tb927.7.3290_rev | GGGGACCACTTTGTACAAGAAAGCTGGGTTCAACACCCATCTCCACCGC |
| 5_hypothetical | N3 | Tb927.4.5150_for | GGGGACAAGTTTGTACAAAAAAGCAGGCTATGTTTCGGAAGACGTTGTTGTTTC | Tb927.4.5150_rev | GGGGACCACTTTGTACAAGAAAGCTGGGTTCATTTCTTTTTTACTTTCTTTAGTTTCGC |
| 5_hypothetical | N4 | Tb927.5.3980_for | GGGGACAAGTTTGTACAAAAAAGCAGGCTATGCTCTCTGTGAGTCCGCC | Tb927.5.3980_rev | GGGGACCACTTTGTACAAGAAAGCTGGGTCTACAAACGACCTGCCGCTC |
| 5_hypothetical | N5 | Tb927.10.8240_for | GGGGACAAGTTTGTACAAAAAAGCAGGCTATGGACGCTAAGCAAAGGGAATC | Tb927.10.8240_rev | GGGGACCACTTTGTACAAGAAAGCTGGGTCTACTTAGGTCCCAATGTTGAGGCC |
| 5_hypothetical | N6 | Tb927.8.7240_for | GGGGACAAGTTTGTACAAAAAAGCAGGCTATGCTCCGGCGGACGTTG | Tb927.8.7240_rev | GGGGACCACTTTGTACAAGAAAGCTGGGTTCATTTCTTTTTTACTTTCTTTAGTTTCGC |
| 5_hypothetical | N7 | Tb927.11.3690_for | GGGGACAAGTTTGTACAAAAAAGCAGGCTATGAAAACTTTAGCCAAGAAAACACTGC | Tb927.11.3690_rev | GGGGACCACTTTGTACAAGAAAGCTGGGTTTACAAATACTCCGTCAAAAACGGCA |
| 5_hypothetical | N8 | Tb927.11.2940_for | GGGGACAAGTTTGTACAAAAAAGCAGGCTATGGGCGATGACTGCGAGA | Tb927.11.2940_rev | GGGGACCACTTTGTACAAGAAAGCTGGGTTTAGTCGAGTTCATCACCGCTAGT |
| 5_hypothetical | N9 | Tb927.10.10710_for | GGGGACAAGTTTGTACAAAAAAGCAGGCTATGAACGAGGTTATGCGTGTGC | Tb927.10.10710_rev | GGGGACCACTTTGTACAAGAAAGCTGGGTCTAATAACCGTCATCCGTATCATTCCAA |
| 5_hypothetical | N10 | Tb927.11.14720_for | GGGGACAAGTTTGTACAAAAAAGCAGGCTATGATTGAGACACCTGATGCGTC | Tb927.11.14720_rev | GGGGACCACTTTGTACAAGAAAGCTGGGTTCACGAAGACGCCGCTTTC |
| 5_hypothetical | N11 | Tb927.6.3260_for | GGGGACAAGTTTGTACAAAAAAGCAGGCTATGGACGATGACCTCATCCTCAA | Tb927.6.3260_rev | GGGGACCACTTTGTACAAGAAAGCTGGGTTCAAGGTCGTAATAGCGGAGTATCTAA |
| 5_hypothetical | N12 | Tb927.11.15160_for | GGGGACAAGTTTGTACAAAAAAGCAGGCTATGTTTTATGGTACCAACGACGTCG | Tb927.11.15160_rev | GGGGACCACTTTGTACAAGAAAGCTGGGTTTACTCCTCCTGTTCTTCAAGTTCATATAC |
| 5_hypothetical | N13 | Tb927.7.2140_for | GGGGACAAGTTTGTACAAAAAAGCAGGCTATGTCCTTCTCAATGAACACAGTGAC | Tb927.7.2140_rev | GGGGACCACTTTGTACAAGAAAGCTGGGTTCACTGCCCTAAAGAAGTTGCG |
| 5_hypothetical | N14 | Tb927.9.3030_for | GGGGACAAGTTTGTACAAAAAAGCAGGCTATGTCGGTTTTTCCCCCTGGC | Tb927.9.3030_rev | GGGGACCACTTTGTACAAGAAAGCTGGGTTTAGCGCTTTTTACCTATTGGTGGT |
| 5_hypothetical | N15 | Tb927.6.2910_for | GGGGACAAGTTTGTACAAAAAAGCAGGCTATGGTCTCGCCACCAGCT | Tb927.6.2910_rev | GGGGACCACTTTGTACAAGAAAGCTGGGTCTAGGGTTTGTCTTCTATTATTAACGCAAA |
| 5_hypothetical | N16 | Tb927.3.2640_for | GGGGACAAGTTTGTACAAAAAAGCAGGCTATGTCCATCACCACAGCCGG | Tb927.3.2640_rev | GGGGACCACTTTGTACAAGAAAGCTGGGTCTACAAAACCTCTTCGTCCCCCA |
| 5_hypothetical | N17 | Tb927.11.5610_for | GGGGACAAGTTTGTACAAAAAAGCAGGCTATGAAAGGCGATGATTCTGTTCATGG | Tb927.11.5610_rev | GGGGACCACTTTGTACAAGAAAGCTGGGTCTAAAGCAGTCCTTGTGAAAGTACCG |
| 5_hypothetical | N18 | Tb927.2.4420_for | GGGGACAAGTTTGTACAAAAAAGCAGGCTATGCTTCCACCCATTTCCGTG | Tb927.2.4420_rev | GGGGACCACTTTGTACAAGAAAGCTGGGTTCATGTGGTTAGTGGTAGTGAAGGA |
| 5_hypothetical | N19 | Tb927.5.430_for | GGGGACAAGTTTGTACAAAAAAGCAGGCTATGGGCGCAAGATTGCGT | Tb927.5.430_rev | GGGGACCACTTTGTACAAGAAAGCTGGGTCTATATCATGAGTCGAGACCCTGACC |
| 5_hypothetical | N20 | Tb927.3.5130_for | GGGGACAAGTTTGTACAAAAAAGCAGGCTATGTCGCCCCCACAACAAGC | Tb927.3.5130_rev | GGGGACCACTTTGTACAAGAAAGCTGGGTTTAGAGCAGCGTCATCACTTTCC |
| 5_hypothetical | N21 | Tb927.3.660_for | GGGGACAAGTTTGTACAAAAAAGCAGGCTATGAATGAAGCGGTGGTAAGCG | Tb927.3.660_rev | GGGGACCACTTTGTACAAGAAAGCTGGGTTTACTCGTCAGGTTTGTGGTTGTAC |
| 5_hypothetical | N22 | Tb927.7.2230_for | GGGGACAAGTTTGTACAAAAAAGCAGGCTATGACCACGCAAGATGAAAAGAACC | Tb927.7.2230_rev | GGGGACCACTTTGTACAAGAAAGCTGGGTTCAAACAGCCTTCCAGCAGC |
| 5_hypothetical | N23 | Tb927.7.6800_for | GGGGACAAGTTTGTACAAAAAAGCAGGCTATGCGGCTGGCGCGGACC | Tb927.7.6800_rev | GGGGACCACTTTGTACAAGAAAGCTGGGTTCACGCCTTTGATTTCTTTTTCTCCT |
| 5_hypothetical | N24 | Tb927.8.5820_for | GGGGACAAGTTTGTACAAAAAAGCAGGCTATGGAGCATGAGAGGGGGAGT | Tb927.8.5820_rev | GGGGACCACTTTGTACAAGAAAGCTGGGTCTAAGAGAGCGTCCAATCACCTACA |
| 5_hypothetical | O1 | Tb927.10.2850_for | GGGGACAAGTTTGTACAAAAAAGCAGGCTATGGAGCGCGTTACCCTCA | Tb927.10.2850_rev | GGGGACCACTTTGTACAAGAAAGCTGGGTTTACTGGGTCGTAGAAGGAATGCC |
| 5_hypothetical | O2 | Tb927.3.3830_for | GGGGACAAGTTTGTACAAAAAAGCAGGCTATGGGCATTATTAAACATGTGGCAGT | Tb927.3.3830_rev | GGGGACCACTTTGTACAAGAAAGCTGGGTTTAACGGGAAAGCCGTCGC |
| 5_hypothetical | O3 | Tb927.3.1710_for | GGGGACAAGTTTGTACAAAAAAGCAGGCTATGTACGCTCGTTTCCGGACT | Tb927.3.1710_rev | GGGGACCACTTTGTACAAGAAAGCTGGGTTCAAACATGCTTCACACTTTCCTCA |
| 5_hypothetical | O4 | Tb927.10.4580_for | GGGGACAAGTTTGTACAAAAAAGCAGGCTATGGTTGGGAAGGTTGAAGTAATCCC | Tb927.10.4580_rev | GGGGACCACTTTGTACAAGAAAGCTGGGTTTATGTGACCCACTGTCGCACG |
| 5_hypothetical | O5 | Tb927.9.9170_for | GGGGACAAGTTTGTACAAAAAAGCAGGCTATGGCGTTTTGGTCAAAGGCG | Tb927.9.9170_rev | GGGGACCACTTTGTACAAGAAAGCTGGGTTCAAGTTGGGCTCAGCGGC |
| 5_hypothetical | O6 | Tb927.11.8880_for | GGGGACAAGTTTGTACAAAAAAGCAGGCTATGGCGAGGATAATTGACGTATCCG | Tb927.11.8880_rev | GGGGACCACTTTGTACAAGAAAGCTGGGTCTCATCCGAAGCGAATGTTAGCTATC |
| 5_hypothetical | O7 | Tb927.9.11510_for | GGGGACAAGTTTGTACAAAAAAGCAGGCTATGAGACGCCAACCGTTTCC | Tb927.9.11510_rev | GGGGACCACTTTGTACAAGAAAGCTGGGTTCAACTGGTTGCTGCTGTGAC |
| 5_hypothetical | O8 | Tb927.4.680_for | GGGGACAAGTTTGTACAAAAAAGCAGGCTATGCGGCGCATTCCAATTGC | Tb927.4.680_rev | GGGGACCACTTTGTACAAGAAAGCTGGGTCTACTTTAAATCGGAAACCAATGGCG |
| 5_hypothetical | O9 | Tb927.4.1900_for | GGGGACAAGTTTGTACAAAAAAGCAGGCTATGCCGCCAACACGTCCC | Tb927.4.1900_rev | GGGGACCACTTTGTACAAGAAAGCTGGGTCTAAAAACGGGGTCGCCGC |
| 5_hypothetical | O10 | Tb927.11.1640_for | GGGGACAAGTTTGTACAAAAAAGCAGGCTATGAGTGGACAGCGCAACTCT | Tb927.11.1640_rev | GGGGACCACTTTGTACAAGAAAGCTGGGTCTACGACAACCTGCGGTATAATACAA |
| 5_hypothetical | O11 | Tb927.4.330_for | GGGGACAAGTTTGTACAAAAAAGCAGGCTATGCCACGAACGTTCTATAGGAGG | Tb927.4.330_rev | GGGGACCACTTTGTACAAGAAAGCTGGGTCTAGAACTTACCCTCACTGTTTAGATCCC |
| 5_hypothetical | O12 | Tb927.9.1380_for | GGGGACAAGTTTGTACAAAAAAGCAGGCTATGTTATTCCGGAGGCTCAAGCG | Tb927.9.1380_rev | GGGGACCACTTTGTACAAGAAAGCTGGGTTCATCGAGGAATGTCACCCCTC |
| 5_hypothetical | O13 | Tb927.4.3540_for | GGGGACAAGTTTGTACAAAAAAGCAGGCTATGCACTCACGTGGTGTCTACA | Tb927.4.3540_rev | GGGGACCACTTTGTACAAGAAAGCTGGGTTCACTTCTGGAGCCCCTTTAGC |
| 5_hypothetical | O14 | Tb927.3.1560_for | GGGGACAAGTTTGTACAAAAAAGCAGGCTATGAAAAAGAATTGTAGCTTAATGTTGAGC | Tb927.3.1560_rev | GGGGACCACTTTGTACAAGAAAGCTGGGTCTATGAAAAGAACCCCCGAAGACAA |
| 5_hypothetical | O15 | Tb927.5.1960_for | GGGGACAAGTTTGTACAAAAAAGCAGGCTATGATCAATGAAGCAGCTGCTGAA | Tb927.5.1960_rev | GGGGACCACTTTGTACAAGAAAGCTGGGTTCATAGTTTTAATGTTCTCAACGCACGC |
| 5_hypothetical | O16 | Tb927.4.1780_for | GGGGACAAGTTTGTACAAAAAAGCAGGCTATGCATAAGCTGACTCAGCTTCTG | Tb927.4.1780_rev | GGGGACCACTTTGTACAAGAAAGCTGGGTTTACAATACAATTGGTCTGAAGACCCG |
| 5_hypothetical | O17 | Tb927.7.5530_for | GGGGACAAGTTTGTACAAAAAAGCAGGCTATGAATCAGAAGATTCTCCTAGCGGA | Tb927.7.5530_rev | GGGGACCACTTTGTACAAGAAAGCTGGGTTCATGTTTCGTCGCCCTCGT |
| 5_hypothetical | O18 | Tb927.10.13850_for | GGGGACAAGTTTGTACAAAAAAGCAGGCTATGCGAAAGTTTTTACGGCCTCA | Tb927.10.13850_rev | GGGGACCACTTTGTACAAGAAAGCTGGGTTCAAATAACCTCTTCTAGAAACGCAGAC |
| 5_hypothetical | O19 | Tb927.9.5330_for | GGGGACAAGTTTGTACAAAAAAGCAGGCTATGCAGCGCTGTAAGCTTCTG | Tb927.9.5330_rev | GGGGACCACTTTGTACAAGAAAGCTGGGTTCACGCTCCATCACTTGAGTAATCC |
| 5_hypothetical | O20 | Tb927.3.3530_for | GGGGACAAGTTTGTACAAAAAAGCAGGCTATGTTGAGGAGGATGGAAGATTGCT | Tb927.3.3530_rev | GGGGACCACTTTGTACAAGAAAGCTGGGTTTATATCGCAGGCCAGAGACCA |
| 5_hypothetical | O21 | Tb927.1.1820_for | GGGGACAAGTTTGTACAAAAAAGCAGGCTATGGATAATAGCACGGAACTTGGGC | Tb927.1.1820_rev | GGGGACCACTTTGTACAAGAAAGCTGGGTTTAGCACCCGTAATAATCCAATAAACCTT |
| 5_hypothetical | O22 | Tb927.7.3810_for | GGGGACAAGTTTGTACAAAAAAGCAGGCTATGTTCCGACTCGCGGGC | Tb927.7.3810_rev | GGGGACCACTTTGTACAAGAAAGCTGGGTTCACTCAAGTAGATTGAATTCATCGTCG |
| 5_hypothetical | O23 | Tb927.5.3770_for | GGGGACAAGTTTGTACAAAAAAGCAGGCTATGAGACGGTTTGCACCACAC | Tb927.5.3770_rev | GGGGACCACTTTGTACAAGAAAGCTGGGTCTAAACTTGAATCACAGAAACCCGGT |
| 5_hypothetical | O24 | Tb927.9.9140_for | GGGGACAAGTTTGTACAAAAAAGCAGGCTATGTGTGAGGTTGACACTACCAAAC | Tb927.9.9140_rev | GGGGACCACTTTGTACAAGAAAGCTGGGTTTACTCGATTTTATCTCGGGAGTGTTTC |
| 5_hypothetical | P1 | Tb927.11.9060_for | GGGGACAAGTTTGTACAAAAAAGCAGGCTATGGCATCGGCAGTGGTGG | Tb927.11.9060_rev | GGGGACCACTTTGTACAAGAAAGCTGGGTCTCACTCACAAAGAAAACATGACCCC |
| 5_hypothetical | P2 | Tb927.9.13020_for | GGGGACAAGTTTGTACAAAAAAGCAGGCTATGAAGTTGGTCCGTCATATAGAAACC | Tb927.9.13020_rev | GGGGACCACTTTGTACAAGAAAGCTGGGTTTATTCACTCCGCAACACGAGAG |
| 5_hypothetical | P4 | Tb927.10.2320_for | GGGGACAAGTTTGTACAAAAAAGCAGGCTATGATGGCGCTTGAGCTGC | Tb927.10.2320_rev | GGGGACCACTTTGTACAAGAAAGCTGGGTTTACGACATGGAGAAAATGCTGAGAA |
| 5_hypothetical | P5 | Tb927.8.2730_for | GGGGACAAGTTTGTACAAAAAAGCAGGCTATGCTTGGTGCTGTTCCTCCT | Tb927.8.2730_rev | GGGGACCACTTTGTACAAGAAAGCTGGGTTCATCGCGACCGCCAAAAC |
| 5_hypothetical | P6 | Tb927.10.1740_for | GGGGACAAGTTTGTACAAAAAAGCAGGCTATGGATACTTTCCAGGAGAAACTGGA | Tb927.10.1740_rev | GGGGACCACTTTGTACAAGAAAGCTGGGTCTAACAGTCTCGATCTCGCTTCCT |
| 5_hypothetical | P7 | Tb927.5.4160_for | GGGGACAAGTTTGTACAAAAAAGCAGGCTATGAATTCTTCTCTTGTTTTTGATAAGGAT | Tb927.5.4160_rev | GGGGACCACTTTGTACAAGAAAGCTGGGTTCAAAGGAAACGGCCGATGAG |
| 5_hypothetical | P8 | Tb927.11.480_for | GGGGACAAGTTTGTACAAAAAAGCAGGCTATGCTGTCTCGTACTTTTTCCCG | Tb927.11.480_rev | GGGGACCACTTTGTACAAGAAAGCTGGGTTTACTTGCAGGACATTCCACCTGG |
| 5_hypothetical | P10 | Tb927.10.3530_for | GGGGACAAGTTTGTACAAAAAAGCAGGCTATGGATCCTTTGGACGATTGGGA | Tb927.10.3530_rev | GGGGACCACTTTGTACAAGAAAGCTGGGTTCAGCGGGATGAAGGGGTAC |
| 5_hypothetical | P11 | Tb927.4.2280_for | GGGGACAAGTTTGTACAAAAAAGCAGGCTATGTTGCCTCATTCGGTGCC | Tb927.4.2280_rev | GGGGACCACTTTGTACAAGAAAGCTGGGTTTAGACGGATCGTGTATCTTCACCG |
| 5_hypothetical | P12 | Tb927.10.7730_for | GGGGACAAGTTTGTACAAAAAAGCAGGCTATGTTGAGCGTACAGGGAGCA | Tb927.10.7730_rev | GGGGACCACTTTGTACAAGAAAGCTGGGTCTATAGGTCTTTCAAAAGTGTGGCCA |
| 5_hypothetical | P13 | Tb927.3.4730_for | GGGGACAAGTTTGTACAAAAAAGCAGGCTATGGGCCGCATCACTACCG | Tb927.3.4730_rev | GGGGACCACTTTGTACAAGAAAGCTGGGTTCACCCGGTTCCATCCTCATT |
| 5_hypothetical | P15 | Tb927.9.2250_for | GGGGACAAGTTTGTACAAAAAAGCAGGCTATGAACCATGGCAGTCGGGG | Tb927.9.2250_rev | GGGGACCACTTTGTACAAGAAAGCTGGGTTTATCGCTCCCTCACGTCTTCC |
| 5_hypothetical | P16 | Tb927.10.10690_for | GGGGACAAGTTTGTACAAAAAAGCAGGCTATGCAGTTTGTGTCTGATCTGGG | Tb927.10.10690_rev | GGGGACCACTTTGTACAAGAAAGCTGGGTCTAGTCCGAGTCGTCCCACTC |
| 5_hypothetical | P17 | Tb927.11.13820_for | GGGGACAAGTTTGTACAAAAAAGCAGGCTATGGATTTAGGGGAAGATTCGACACT | Tb927.11.13820_rev | GGGGACCACTTTGTACAAGAAAGCTGGGTTTAAATAGTACTGAACTGCTCCTTAGCCA |
| 5_hypothetical | P18 | Tb927.11.15870_for | GGGGACAAGTTTGTACAAAAAAGCAGGCTATGTCTGGGGAGATTCGATATGCT | Tb927.11.15870_rev | GGGGACCACTTTGTACAAGAAAGCTGGGTTTACGAAGGGCTCATAAATAACTTTACTAC |
| 5_hypothetical | P19 | Tb927.11.10380_for | GGGGACAAGTTTGTACAAAAAAGCAGGCTATGACTGAGCTATTAAGTGCAGGGG | Tb927.11.10380_rev | GGGGACCACTTTGTACAAGAAAGCTGGGTTCAGGGTTCTTCGTGCAAAAGC |
| 5_hypothetical | P20 | Tb927.11.6070_for | GGGGACAAGTTTGTACAAAAAAGCAGGCTATGTGGTCTGTCATTCGTAAACTTCAG | Tb927.11.6070_rev | GGGGACCACTTTGTACAAGAAAGCTGGGTTTAACACTTTCGTTCCAGCTGGC |
| 5_hypothetical | P21 | Tb927.11.5980_for | GGGGACAAGTTTGTACAAAAAAGCAGGCTATGTCTATTGCATGCGCACTTGA | Tb927.11.5980_rev | GGGGACCACTTTGTACAAGAAAGCTGGGTTTAGATGCGAGTTGTTCCGTTGG |
| 5_hypothetical | P22 | Tb927.4.2440_for | GGGGACAAGTTTGTACAAAAAAGCAGGCTATGTTTTCACAAACGCGCTGTTC | Tb927.4.2440_rev | GGGGACCACTTTGTACAAGAAAGCTGGGTTTACGAATCCGCTGGCACAC |
| 5_hypothetical | P23 | Tb927.6.5010_for | GGGGACAAGTTTGTACAAAAAAGCAGGCTATGGAACATTCGCGTCGGAAG | Tb927.6.5010_rev | GGGGACCACTTTGTACAAGAAAGCTGGGTTTAAACACTCTTCCCTCCTGGAACA |
| 5_hypothetical | P24 | Tb927.10.990_for | GGGGACAAGTTTGTACAAAAAAGCAGGCTATGGCAGCAGTGATGAAATTAGAAAAGT | Tb927.10.990_rev | GGGGACCACTTTGTACAAGAAAGCTGGGTTCATTGAGTCCATAAACACAGCGAG |
| 5_known | A1 | Tb927.11.2550_for | GGGGACAAGTTTGTACAAAAAAGCAGGCTATGTCCGTGGAGCAATGCTCC | Tb927.11.2550_rev | GGGGACCACTTTGTACAAGAAAGCTGGGTTCAAAGAGTGGGTCGGAAAACATC |
| 5_known | A2 | Tb927.11.6680_for | GGGGACAAGTTTGTACAAAAAAGCAGGCTATGGCTGGGGATGGATTCGT | Tb927.11.6680_rev | GGGGACCACTTTGTACAAGAAAGCTGGGTCTACTGTTTTTCATCCTCGCTCAGAG |
| 5_known | A3 | Tb927.7.3760_for | GGGGACAAGTTTGTACAAAAAAGCAGGCTATGGCCGGTAAGCTCAACAGT | Tb927.7.3760_rev | GGGGACCACTTTGTACAAGAAAGCTGGGTTTATCGCCAAAAGATTATGTAGCGCT |
| 5_known | A4 | Tb927.10.8360_for | GGGGACAAGTTTGTACAAAAAAGCAGGCTATGGACGTCGACAAATATAACTGTTCC | Tb927.10.8360_rev | GGGGACCACTTTGTACAAGAAAGCTGGGTCTACCCACTAAGACGGTAAACACTTAGT |
| 5_known | A5 | Tb927.1.720_for | GGGGACAAGTTTGTACAAAAAAGCAGGCTATGTCCACTGCACCGAATGC | Tb927.1.720_rev | GGGGACCACTTTGTACAAGAAAGCTGGGTTTACTCCTTATCATCCAATACTGCAACAC |
| 5_known | A6 | Tb927.2.5240_for | GGGGACAAGTTTGTACAAAAAAGCAGGCTATGTTTTGTTGCATTTCTAACTGTGTGC | Tb927.2.5240_rev | GGGGACCACTTTGTACAAGAAAGCTGGGTTCATGTTGATCTGTACTCCTTCACCA |
| 5_known | A7 | Tb927.10.2000_for | GGGGACAAGTTTGTACAAAAAAGCAGGCTATGACTGCTGGATATCCGGTTCT | Tb927.10.2000_rev | GGGGACCACTTTGTACAAGAAAGCTGGGTTTACAGTAAATTTTGAAGCGCCGAGA |
| 5_known | A8 | Tb927.11.4170_for | GGGGACAAGTTTGTACAAAAAAGCAGGCTATGATGGGGGACGATGATCGT | Tb927.11.4170_rev | GGGGACCACTTTGTACAAGAAAGCTGGGTTCAACTATTACGCTCTCCATAATTTAGTCG |
| 5_known | A9 | Tb927.4.2230_for | GGGGACAAGTTTGTACAAAAAAGCAGGCTATGATTTACGAACTTGCGCTACTTATATCC | Tb927.4.2230_rev | GGGGACCACTTTGTACAAGAAAGCTGGGTTCAGTCCGCCTTTTTTCCTCCC |
| 5_known | A10 | Tb927.7.4590_for | GGGGACAAGTTTGTACAAAAAAGCAGGCTATGTTTCAACTACGGTCGTGTGTT | Tb927.7.4590_rev | GGGGACCACTTTGTACAAGAAAGCTGGGTTTATCCTCCTTCATCGTACTGGATACTAG |
| 5_known | A11 | Tb927.11.16550_for | GGGGACAAGTTTGTACAAAAAAGCAGGCTATGAGTGAAGCGAAGGACTCTTTG | Tb927.11.16550_rev | GGGGACCACTTTGTACAAGAAAGCTGGGTTTACACATATGTAAGCAACAGCGGC |
| 5_known | A12 | Tb927.3.3540_for | GGGGACAAGTTTGTACAAAAAAGCAGGCTATGATGTCAACTGCCCCAACG | Tb927.3.3540_rev | GGGGACCACTTTGTACAAGAAAGCTGGGTTTATGAGCGACCGGTTTTGTTCA |
| 5_known | A13 | Tb927.9.12570_for | GGGGACAAGTTTGTACAAAAAAGCAGGCTATGAAGTACGTCGGATCCATTGACC | Tb927.9.12570_rev | GGGGACCACTTTGTACAAGAAAGCTGGGTCTACAACTTTGCCCACTTCGTCC |
| 5_known | A16 | Tb927.8.630_for | GGGGACAAGTTTGTACAAAAAAGCAGGCTATGCGCGCTGCCAGACTG | Tb927.8.630_rev | GGGGACCACTTTGTACAAGAAAGCTGGGTTTAATAACTTATGGGGTAGAAGACGTTTTT |
| 5_known | A19 | Tb927.10.16120_for | GGGGACAAGTTTGTACAAAAAAGCAGGCTATGGAAAACACCAACCTACGCAC | Tb927.10.16120_rev | GGGGACCACTTTGTACAAGAAAGCTGGGTTTAGAGCTTCGAGGCAAAGAGTTTC |
| 5_known | A20 | Tb927.10.350_for | GGGGACAAGTTTGTACAAAAAAGCAGGCTATGATTTCCGTCTTGTCGCCC | Tb927.10.350_rev | GGGGACCACTTTGTACAAGAAAGCTGGGTTCAAATAGACGTGTCTTTGATGCCA |
| 5_known | A21 | Tb927.11.13970_for | GGGGACAAGTTTGTACAAAAAAGCAGGCTATGGATTCAAAGGGACGCGGT | Tb927.11.13970_rev | GGGGACCACTTTGTACAAGAAAGCTGGGTTTACAACGTTGCCGATGTCGT |
| 5_known | A22 | Tb927.10.6290_for | GGGGACAAGTTTGTACAAAAAAGCAGGCTATGGTTAAGGAGAAGTTGGCGGG | Tb927.10.6290_rev | GGGGACCACTTTGTACAAGAAAGCTGGGTTCATGACTTCATCTGCTTCCGC |
| 5_known | A23 | Tb927.6.2450_for | GGGGACAAGTTTGTACAAAAAAGCAGGCTATGCCCCGTAGTGGTGATGC | Tb927.6.2450_rev | GGGGACCACTTTGTACAAGAAAGCTGGGTTTACTTCATGAAATCTTCATGAACATTATT |
| 5_known | A24 | Tb927.1.2670_for | GGGGACAAGTTTGTACAAAAAAGCAGGCTATGCCCAACAGGCAGATCATTC | Tb927.1.2670_rev | GGGGACCACTTTGTACAAGAAAGCTGGGTTTAATGCTGCTGCACTTGTTGGA |
| 5_known | B1 | Tb927.2.2440_for | GGGGACAAGTTTGTACAAAAAAGCAGGCTATGGAAAAGGATTTGGACTCACTTTGG | Tb927.2.2440_rev | GGGGACCACTTTGTACAAGAAAGCTGGGTTTATTCCTTCTTCGGCTCACTTTTCG |
| 5_known | B2 | Tb927.7.600_for | GGGGACAAGTTTGTACAAAAAAGCAGGCTATGTGGCGGATTTCTAATTTATGGCG | Tb927.7.600_rev | GGGGACCACTTTGTACAAGAAAGCTGGGTTTAGAGCCAGTCCTTTTCGGACT |
| 5_known | B3 | Tb927.11.4740_for | GGGGACAAGTTTGTACAAAAAAGCAGGCTATGGCGAAAAAGGGTAAAAAGAATTCG | Tb927.11.4740_rev | GGGGACCACTTTGTACAAGAAAGCTGGGTTCAGTGTGAAGGCAACATATCCCA |
| 5_known | B4 | Tb927.10.5150_for | GGGGACAAGTTTGTACAAAAAAGCAGGCTATGTATCGGCCAACTAACAGATGC | Tb927.10.5150_rev | GGGGACCACTTTGTACAAGAAAGCTGGGTTTAGATGGTTTCACGAGGGTGTGG |
| 5_known | B5 | Tb927.7.5820_for | GGGGACAAGTTTGTACAAAAAAGCAGGCTATGAATGGTCGTACACGGGTATTG | Tb927.7.5820_rev | GGGGACCACTTTGTACAAGAAAGCTGGGTTTATGCAAGGAGAAATGACTCCCGG |
| 5_known | B6 | Tb927.10.9900_for | GGGGACAAGTTTGTACAAAAAAGCAGGCTATGACAGTAAACGCAATTACAGGTTCT | Tb927.10.9900_rev | GGGGACCACTTTGTACAAGAAAGCTGGGTTCACACGCTCGTTAATTCATCGT |
| 5_known | B8 | Tb927.11.9510_for | GGGGACAAGTTTGTACAAAAAAGCAGGCTATGTCTGAAGCCATTGAGGGCG | Tb927.11.9510_rev | GGGGACCACTTTGTACAAGAAAGCTGGGTCTACTGAAAGACACTATGGGTGGGA |
| 5_known | B9 | Tb927.1.580_for | GGGGACAAGTTTGTACAAAAAAGCAGGCTATGGTGCATCCATACTTATGGGTCA | Tb927.1.580_rev | GGGGACCACTTTGTACAAGAAAGCTGGGTTTACTGCGAGGTGAGACTCGG |
| 5_known | B10 | Tb927.7.4830_for | GGGGACAAGTTTGTACAAAAAAGCAGGCTATGGAACACATAGGTGCTTTTCTACG | Tb927.7.4830_rev | GGGGACCACTTTGTACAAGAAAGCTGGGTCTATGAGGCTTGTGGCCCGA |
| 5_known | B11 | Tb927.5.3010_for | GGGGACAAGTTTGTACAAAAAAGCAGGCTATGCGGAGCGCGTTGCGT | Tb927.5.3010_rev | GGGGACCACTTTGTACAAGAAAGCTGGGTTTATGCTTGCGGTGTAGACCCA |
| 5_known | B12 | Tb927.5.4560_for | GGGGACAAGTTTGTACAAAAAAGCAGGCTATGGAAACCGCCGTGCCT | Tb927.5.4560_rev | GGGGACCACTTTGTACAAGAAAGCTGGGTCTAAAGTTCTACCCTCCCGTGGA |
| 5_known | B13 | Tb927.11.1980_for | GGGGACAAGTTTGTACAAAAAAGCAGGCTATGAGCAGCGCGGTAGATTCC | Tb927.11.1980_rev | GGGGACCACTTTGTACAAGAAAGCTGGGTTCACAGGGTCTTGGGCTTCA |
| 5_known | B14 | Tb927.10.14910_for | GGGGACAAGTTTGTACAAAAAAGCAGGCTATGAGTTCGGGTAATAGGGACCTTT | Tb927.10.14910_rev | GGGGACCACTTTGTACAAGAAAGCTGGGTCTACCGTGGAGCATAGAACTGGT |

| | | | | | |
|---|---|---|---|---|---|
| 5_known | B15 | Tb927.5.2850_for | GGGGACAAGTTTGTACAAAAAAGCAGGCTATGACTGAGAGTGCATACCTCAAATG | Tb927.5.2850_rev | GGGGACCACTTTGTACAAGAAAGCTGGGTTTAGTTGCCTTCGTCCCCATCA |
| 5_known | B16 | Tb927.11.3980_for | GGGGACAAGTTTGTACAAAAAAGCAGGCTATGTACCGACGCGCCGTT | Tb927.11.3980_rev | GGGGACCACTTTGTACAAGAAAGCTGGGTTTACGCTCCCAACTGACGGA |
| 5_known | B17 | Tb927.8.1420_for | GGGGACAAGTTTGTACAAAAAAGCAGGCTATGAGACGTTTCATTTCTGTTTCTAAGCA | Tb927.8.1420_rev | GGGGACCACTTTGTACAAGAAAGCTGGGTCTACTGTAAAGCATCTTCGCAAATATCG |
| 5_known | B18 | Tb927.3.1380_for | GGGGACAAGTTTGTACAAAAAAGCAGGCTATGCTGACTCGTTTCCGAAGTG | Tb927.3.1380_rev | GGGGACCACTTTGTACAAGAAAGCTGGGTTTAGCTACTGGCTTGAGCAACAC |
| 5_known | B19 | Tb927.8.7570_for | GGGGACAAGTTTGTACAAAAAAGCAGGCTATGACATACAATGAGGCGGCG | Tb927.8.7570_rev | GGGGACCACTTTGTACAAGAAAGCTGGGTTCACATCAAGAAGTCGATGCCAAG |
| 5_known | B20 | Tb927.10.10380_for | GGGGACAAGTTTGTACAAAAAAGCAGGCTATGAGCGAGTTCGAAGCACTG | Tb927.10.10380_rev | GGGGACCACTTTGTACAAGAAAGCTGGGTTTATTCCCCTGCCATGCTGGT |
| 5_known | B21 | Tb927.5.3750_for | GGGGACAAGTTTGTACAAAAAAGCAGGCTATGCCAAGTTCGTCGTTGTCG | Tb927.5.3750_rev | GGGGACCACTTTGTACAAGAAAGCTGGGTTTATTGGCGCACCCTCTTGC |
| 5_known | B22 | Tb927.10.7400_for | GGGGACAAGTTTGTACAAAAAAGCAGGCTATGACACCGCTTCAAGTATTGTTCA | Tb927.10.7400_rev | GGGGACCACTTTGTACAAGAAAGCTGGGTTCAGCTAGAGTCCCTCTGCCG |
| 5_known | B23 | Tb927.5.2650_for | GGGGACAAGTTTGTACAAAAAAGCAGGCTATGGGACAGGAGACGATGTCC | Tb927.5.2650_rev | GGGGACCACTTTGTACAAGAAAGCTGGGTCTACAGATGACTAATAGTGCTGTTAAGTAC |
| 5_known | B24 | Tb927.3.5460_for | GGGGACAAGTTTGTACAAAAAAGCAGGCTATGGAGGGCCGCAAGGGA | Tb927.3.5460_rev | GGGGACCACTTTGTACAAGAAAGCTGGGTTCAGGAGGAATGCGGTAGCA |
| 5_known | C1 | Tb927.11.11200_for | GGGGACAAGTTTGTACAAAAAAGCAGGCTATGGAACTTGAGGTGAGTGAGGAG | Tb927.11.11200_rev | GGGGACCACTTTGTACAAGAAAGCTGGGTTCACGAATGTTTCTCCCTCGTTTC |
| 5_known | C2 | Tb927.11.6590_for | GGGGACAAGTTTGTACAAAAAAGCAGGCTATGCCTACTTTACCCAAGGCCG | Tb927.11.6590_rev | GGGGACCACTTTGTACAAGAAAGCTGGGTCTACAATTTTCGGAAGTACTCGGTAAGT |
| 5_known | C3 | Tb927.11.4940_for | GGGGACAAGTTTGTACAAAAAAGCAGGCTATGTCCGTCGATGGGGCG | Tb927.11.4940_rev | GGGGACCACTTTGTACAAGAAAGCTGGGTTCAAAGGGCAGCAAATGCATAATCA |
| 5_known | C4 | Tb927.5.2430_for | GGGGACAAGTTTGTACAAAAAAGCAGGCTATGCAGACACTTTCAGCCTCTTTT | Tb927.5.2430_rev | GGGGACCACTTTGTACAAGAAAGCTGGGTTTACTGGTGTCGCCGACTCA |
| 5_known | C5 | Tb927.6.2810_for | GGGGACAAGTTTGTACAAAAAAGCAGGCTATGGCACTGTCATTTACCGATCC | Tb927.6.2810_rev | GGGGACCACTTTGTACAAGAAAGCTGGGTTCACACGATTACGTTCACGTGC |
| 5_known | C6 | Tb927.5.3860_for | GGGGACAAGTTTGTACAAAAAAGCAGGCTATGTTCCCTAATACATCGCTGCG | Tb927.5.3860_rev | GGGGACCACTTTGTACAAGAAAGCTGGGTTCAGCACTCTCCTACTTGCAAAAG |
| 5_known | C7 | Tb927.6.1680_for | GGGGACAAGTTTGTACAAAAAAGCAGGCTATGCGGCGCTGGTTGGTG | Tb927.6.1680_rev | GGGGACCACTTTGTACAAGAAAGCTGGGTTTACGACGTCACCTCACTTACCT |
| 5_known | C8 | Tb927.11.13130_for | GGGGACAAGTTTGTACAAAAAAGCAGGCTATGAAGGAGAGAAAGGGCTCTTGG | Tb927.11.13130_rev | GGGGACCACTTTGTACAAGAAAGCTGGGTCTATGGGACGGGGCCATCA |
| 5_known | C9 | Tb927.2.4210_for | GGGGACAAGTTTGTACAAAAAAGCAGGCTATGGCTCCTATAATTCACAAGAATCTCAC | Tb927.2.4210_rev | GGGGACCACTTTGTACAAGAAAGCTGGGTTTAAAGACGCGAAGCCTCAACAT |
| 5_known | C10 | Tb927.10.470_for | GGGGACAAGTTTGTACAAAAAAGCAGGCTATGCGGTGCAACAAAGCATTAC | Tb927.10.470_rev | GGGGACCACTTTGTACAAGAAAGCTGGGTTCAATAAATTATACGCGCCTTCGCA |
| 5_known | C11 | Tb927.10.1710_for | GGGGACAAGTTTGTACAAAAAAGCAGGCTATGCGCTGCATCGCCCAC | Tb927.10.1710_rev | GGGGACCACTTTGTACAAGAAAGCTGGGTTCAGTCAATGATGTGTACATCCTTCTCT |
| 5_known | C12 | Tb927.11.8720_for | GGGGACAAGTTTGTACAAAAAAGCAGGCTATGGGTCGTAGGAGCAACTCTG | Tb927.11.8720_rev | GGGGACCACTTTGTACAAGAAAGCTGGGTTCATACAACCAAAGAAAGGGCACTG |
| 5_known | C13 | Tb927.6.1850_for | GGGGACAAGTTTGTACAAAAAAGCAGGCTATGTTCTCTCGATCATTTGTAGTCTTTGC | Tb927.6.1850_rev | GGGGACCACTTTGTACAAGAAAGCTGGGTCTAGTAACGGTCCCGCCTCTT |
| 5_known | C14 | Tb927.8.8070_for | GGGGACAAGTTTGTACAAAAAAGCAGGCTATGCGGACCAGAACAACAATGAC | Tb927.8.8070_rev | GGGGACCACTTTGTACAAGAAAGCTGGGTTTAGCAAATGCGTTTCATCACAGC |
| 5_known | C15 | Tb927.9.8780_for | GGGGACAAGTTTGTACAAAAAAGCAGGCTATGTTGCTACCAAAGAAGGTTAGATTGA | Tb927.9.8780_rev | GGGGACCACTTTGTACAAGAAAGCTGGGTTTAGGAGGAACCTCCCTGTCTTAAC |
| 5_known | C16 | Tb927.10.14090_for | GGGGACAAGTTTGTACAAAAAAGCAGGCTATGATGGCTTCCGAAGATAAAACGC | Tb927.10.14090_rev | GGGGACCACTTTGTACAAGAAAGCTGGGTCTACAAAGTGTCTTTGCTTCTTTCCG |
| 5_known | C17 | Tb927.10.13580_for | GGGGACAAGTTTGTACAAAAAAGCAGGCTATGATGGCTTCCGAAGATAAAACGC | Tb927.10.13580_rev | GGGGACCACTTTGTACAAGAAAGCTGGGTCTACGAATTAGCCTTTTCAGCCTCC |
| 5_known | C18 | Tb927.10.8440_for | GGGGACAAGTTTGTACAAAAAAGCAGGCTATGACTGAGCGTCGTGATAACG | Tb927.10.8440_rev | GGGGACCACTTTGTACAAGAAAGCTGGGTTTAGTTCCGCGGAGATGCTTCA |
| 5_known | C20 | Tb927.10.8480_for | GGGGACAAGTTTGTACAAAAAAGCAGGCTATGACTGAGCGTCGTGATAACG | Tb927.10.8480_rev | GGGGACCACTTTGTACAAGAAAGCTGGGTTTAGTTACGCGGAGATGCTTCAAC |
| 5_known | C21 | Tb927.7.5480_for | GGGGACAAGTTTGTACAAAAAAGCAGGCTATGCTCAGTCTTACGCGTATCCT | Tb927.7.5480_rev | GGGGACCACTTTGTACAAGAAAGCTGGGTCTACACCGCCATCTCCATAGAAATTAC |
| 5_known | C22 | Tb927.1.4600_for | GGGGACAAGTTTGTACAAAAAAGCAGGCTATGTTTTTGAAGGAGGAAACAGTGA | Tb927.1.4600_rev | GGGGACCACTTTGTACAAGAAAGCTGGGTCTAAGATGCCGCTAGTACGCAC |
| 5_known | C24 | Tb927.11.7960_for | GGGGACAAGTTTGTACAAAAAAGCAGGCTATGAGAAAGTTTTCAGCTTTTCGGTCA | Tb927.11.7960_rev | GGGGACCACTTTGTACAAGAAAGCTGGGTCTACTGATGATTGTCGCATTTTATCCAGC |
| 5_known | D6 | Tb927.7.6450_for | GGGGACAAGTTTGTACAAAAAAGCAGGCTATGAGTGACAAAGATGATGGTGGTAC | Tb927.7.6450_rev | GGGGACCACTTTGTACAAGAAAGCTGGGTTCAACCCTCAACATCCATATCCGC |
| 5_known | D7 | Tb927.7.3630_for | GGGGACAAGTTTGTACAAAAAAGCAGGCTATGTACATTACAGGGGAGAACTCTGC | Tb927.7.3630_rev | GGGGACCACTTTGTACAAGAAAGCTGGGTTTAGTCGAAGTGAAAATAAAACTGCTGAAA |
| 5_known | D8 | Tb927.10.7110_for | GGGGACAAGTTTGTACAAAAAAGCAGGCTATGCCAGCCGTCCGTACG | Tb927.10.7110_rev | GGGGACCACTTTGTACAAGAAAGCTGGGTTCAACTTCCCACGCCGCG |
| 5_known | D9 | Tb927.10.8490_for | GGGGACAAGTTTGTACAAAAAAGCAGGCTATGACTGAGCGTCGTGATAACG | Tb927.10.8490_rev | GGGGACCACTTTGTACAAGAAAGCTGGGTTTAGTTCCGCGGCGTGGG |
| 5_known | D12 | Tb927.11.1900_for | GGGGACAAGTTTGTACAAAAAAGCAGGCTATGCTTTTCGCTAACCAAGCTCA | Tb927.11.1900_rev | GGGGACCACTTTGTACAAGAAAGCTGGGTTCATTGAGTCCGTTTGCGTGG |
| 5_known | D15 | Tb927.10.7680_for | GGGGACAAGTTTGTACAAAAAAGCAGGCTATGATCAACAACGGTTTAGAGGCC | Tb927.10.7680_rev | GGGGACCACTTTGTACAAGAAAGCTGGGTTTAGTATTTAGTGTAAGTTAACCCTGGCG |
| 5_known | D16 | Tb927.10.13160_for | GGGGACAAGTTTGTACAAAAAAGCAGGCTATGTCGCCAACTCGTGCG | Tb927.10.13160_rev | GGGGACCACTTTGTACAAGAAAGCTGGGTTCAGGTCCATGGCAAGCCA |
| 5_known | D17 | Tb927.11.6550_for | GGGGACAAGTTTGTACAAAAAAGCAGGCTATGTTGTCCCGAGCCCCA | Tb927.11.6550_rev | GGGGACCACTTTGTACAAGAAAGCTGGGTTTAGCCGTTCTCACTGCTCACG |
| 5_known | D18 | Tb927.11.16060_for | GGGGACAAGTTTGTACAAAAAAGCAGGCTATGGCGATGTTGCGCGGT | Tb927.11.16060_rev | GGGGACCACTTTGTACAAGAAAGCTGGGTTTACTTCCACCGCTCCTTCGC |
| 5_known | D19 | Tb927.9.12810_for | GGGGACAAGTTTGTACAAAAAAGCAGGCTATGACTTCTCACAACATAGTGATGAGGC | Tb927.9.12810_rev | GGGGACCACTTTGTACAAGAAAGCTGGGTCTAAATTAACACCAGATGTGTGCCCA |
| 5_known | D20 | Tb927.5.520_for | GGGGACAAGTTTGTACAAAAAAGCAGGCTATGTTACGTAGTTTGGTGCACTCG | Tb927.5.520_rev | GGGGACCACTTTGTACAAGAAAGCTGGGTCTACCTAAATCCGCTACTATTATTATTATT |
| 5_known | D21 | Tb927.8.5250_for | GGGGACAAGTTTGTACAAAAAAGCAGGCTATGACCGTTATTGTTGCTGGGATAG | Tb927.8.5250_rev | GGGGACCACTTTGTACAAGAAAGCTGGGTTTACTGTACGTTGTATCCTTCCGCA |
| 5_known | D22 | Tb927.10.9420_for | GGGGACAAGTTTGTACAAAAAAGCAGGCTATGAGGCGGTGTGTAGCTTATGT | Tb927.10.9420_rev | GGGGACCACTTTGTACAAGAAAGCTGGGTTCATCGACGACATTTTCTGTAAATAGCG |
| 5_known | D23 | Tb927.6.2640_for | GGGGACAAGTTTGTACAAAAAAGCAGGCTATGTTCAGTGGTCAGAATGAGAAACC | Tb927.6.2640_rev | GGGGACCACTTTGTACAAGAAAGCTGGGTTCAAAGATCGAACTGGCCTTGTG |
| 5_known | D24 | Tb927.1.3220_for | GGGGACAAGTTTGTACAAAAAAGCAGGCTATGGTGCTGTCCGCCGCA | Tb927.1.3220_rev | GGGGACCACTTTGTACAAGAAAGCTGGGTTCAAACTGTGAAAACTGAAGATCTTATGCT |
| 5_known | E1 | Tb927.3.5540_for | GGGGACAAGTTTGTACAAAAAAGCAGGCTATGCCCATTCCGTGTAATTTCAAGG | Tb927.3.5540_rev | GGGGACCACTTTGTACAAGAAAGCTGGGTTCAGAGAGAGAAACCAAACAGAGATATGA |
| 5_known | E2 | Tb927.7.3110_for | GGGGACAAGTTTGTACAAAAAAGCAGGCTATGGCGAAGCGCCGTGAA | Tb927.7.3110_rev | GGGGACCACTTTGTACAAGAAAGCTGGGTCTACTTGGCATCCACAGTGTCATC |
| 5_known | E3 | Tb927.9.15030_for | GGGGACAAGTTTGTACAAAAAAGCAGGCTATGGATTACCGTAATGATATCCAACAGTTA | Tb927.9.15030_rev | GGGGACCACTTTGTACAAGAAAGCTGGGTTCATACTTGGTCGTCGACACTGC |
| 5_known | E4 | Tb927.6.3510_for | GGGGACAAGTTTGTACAAAAAAGCAGGCTATGAAGAGAAACAGTGCGCGAG | Tb927.6.3510_rev | GGGGACCACTTTGTACAAGAAAGCTGGGTCTATTTGCACCGCCTGCGG |
| 5_known | E5 | Tb927.10.1060_for | GGGGACAAGTTTGTACAAAAAAGCAGGCTATGTCTACCAAGAAGGCAAATGAAAACA | Tb927.10.1060_rev | GGGGACCACTTTGTACAAGAAAGCTGGGTTTACGCCCGTGTCATGATGATG |
| 5_known | E6 | Tb927.10.8190_for | GGGGACAAGTTTGTACAAAAAAGCAGGCTATGTCGTTCATGCAAACTGGTCC | Tb927.10.8190_rev | GGGGACCACTTTGTACAAGAAAGCTGGGTTTAATCATCATCTCTTGCCTGATCCG |
| 5_known | E7 | Tb927.7.290_for | GGGGACAAGTTTGTACAAAAAAGCAGGCTATGCCAAATGCTATTGGCCTGC | Tb927.7.290_rev | GGGGACCACTTTGTACAAGAAAGCTGGGTTTATTTATTAACCCATACGTTCATTGACTT |
| 5_known | E8 | Tb927.10.5440_for | GGGGACAAGTTTGTACAAAAAAGCAGGCTATGGAGCGCGGAGGGCGC | Tb927.10.5440_rev | GGGGACCACTTTGTACAAGAAAGCTGGGTCTAAACTAATTCGGCCACTAGTAGTGC |
| 5_known | E9 | Tb927.10.9600_for | GGGGACAAGTTTGTACAAAAAAGCAGGCTATGCTAGGTGACCGCTACGC | Tb927.10.9600_rev | GGGGACCACTTTGTACAAGAAAGCTGGGTTCAGGCTTTTGAGGAGGCGT |
| 5_known | E10 | Tb927.11.14250_for | GGGGACAAGTTTGTACAAAAAAGCAGGCTATGTCTCTGGCCTTTGATGAGTACG | Tb927.11.14250_rev | GGGGACCACTTTGTACAAGAAAGCTGGGTCTATTCCTCCTCTTCGGGCTTTGT |
| 5_known | E11 | Tb927.6.3630_for | GGGGACAAGTTTGTACAAAAAAGCAGGCTATGTCGCTGTCATGCTTTTTGGA | Tb927.6.3630_rev | GGGGACCACTTTGTACAAGAAAGCTGGGTCTAATGTTCGGTGCTGTAGTACTTGT |
| 5_known | E12 | Tb927.9.9960_for | GGGGACAAGTTTGTACAAAAAAGCAGGCTATGCCAGTTATCGATTCGATGACAC | Tb927.9.9960_rev | GGGGACCACTTTGTACAAGAAAGCTGGGTTCACGAAGAAAGTGGTATTTCATTTAATAC |
| 5_known | E13 | Tb927.11.15560_for | GGGGACAAGTTTGTACAAAAAAGCAGGCTATGAAGGGCTTTGCTGTGCG | Tb927.11.15560_rev | GGGGACCACTTTGTACAAGAAAGCTGGGTCTAAGCGATCGTTGACTTCCCTAGC |
| 5_known | E14 | Tb927.6.4660_for | GGGGACAAGTTTGTACAAAAAAGCAGGCTATGGTCTCATGGTCCCGTCC | Tb927.6.4660_rev | GGGGACCACTTTGTACAAGAAAGCTGGGTTCAATTATGGACATAACGCTCAGTCAA |
| 5_known | E15 | Tb927.1.2740_for | GGGGACAAGTTTGTACAAAAAAGCAGGCTATGCAAATGATGGGACGGGC | Tb927.1.2740_rev | GGGGACCACTTTGTACAAGAAAGCTGGGTTTATGATAAGGATTTGTAATACCGCTGCC |
| 5_known | E16 | Tb927.8.6500_for | GGGGACAAGTTTGTACAAAAAAGCAGGCTATGAACTTCGAGTTGAGCATCAACA | Tb927.8.6500_rev | GGGGACCACTTTGTACAAGAAAGCTGGGTTCAGCGCAAGGGGAGCTC |
| 5_known | E17 | Tb927.11.5480_for | GGGGACAAGTTTGTACAAAAAAGCAGGCTATGATGAACATATCACCTCAATTTAAACGG | Tb927.11.5480_rev | GGGGACCACTTTGTACAAGAAAGCTGGGTTCACCGTGGTAGTAGATCATTTGTCC |
| 5_known | E18 | Tb927.11.2440_for | GGGGACAAGTTTGTACAAAAAAGCAGGCTATGACTAACAAATACGCCCCTGGA | Tb927.11.2440_rev | GGGGACCACTTTGTACAAGAAAGCTGGGTCTACGACTGGCCGTTCGC |
| 5_known | E19 | Tb927.6.4090_for | GGGGACAAGTTTGTACAAAAAAGCAGGCTATGCTTTATTCAACACGCCGTCT | Tb927.6.4090_rev | GGGGACCACTTTGTACAAGAAAGCTGGGTTCAACCACTATCCCGTTGACGC |
| 5_known | E20 | Tb927.10.1260_for | GGGGACAAGTTTGTACAAAAAAGCAGGCTATGTATCGTTTCTGGCCGAGC | Tb927.10.1260_rev | GGGGACCACTTTGTACAAGAAAGCTGGGTTCATGGCGCCAACCGCCT |
| 5_known | E21 | Tb927.11.6960_for | GGGGACAAGTTTGTACAAAAAAGCAGGCTATGTCATGGATTGGCATTATTAACCGC | Tb927.11.6960_rev | GGGGACCACTTTGTACAAGAAAGCTGGGTTCAATTCCACAAGACCAACCGC |
| 5_known | E22 | Tb927.6.3160_for | GGGGACAAGTTTGTACAAAAAAGCAGGCTATGTTTGGTGGAGTTTTAGAAAAGATTCG | Tb927.6.3160_rev | GGGGACCACTTTGTACAAGAAAGCTGGGTTCAACCTGTTTTCCTCGACATCTG |
| 5_known | E23 | Tb927.7.800_for | GGGGACAAGTTTGTACAAAAAAGCAGGCTATGCGACGTCGGGTAGTTTTATG | Tb927.7.800_rev | GGGGACCACTTTGTACAAGAAAGCTGGGTCTAAAGCGATGCTTCGTCAGGT |
| 5_known | E24 | Tb927.6.3050_for | GGGGACAAGTTTGTACAAAAAAGCAGGCTATGCCAGCAGGTGTTCCGG | Tb927.6.3050_rev | GGGGACCACTTTGTACAAGAAAGCTGGGTTCATTCATTGCTCCTCGGACCT |
| 5_known | F1 | Tb927.7.1920_for | GGGGACAAGTTTGTACAAAAAAGCAGGCTATGTTACAAGCGGCTGAGCAG | Tb927.7.1920_rev | GGGGACCACTTTGTACAAGAAAGCTGGGTTCATCTCTCAATATATAATCGTCGATTAAA |
| 5_known | F2 | Tb927.11.6760_for | GGGGACAAGTTTGTACAAAAAAGCAGGCTATGAAGATCGACCTCTCTCACCG | Tb927.11.6760_rev | GGGGACCACTTTGTACAAGAAAGCTGGGTTCAGTTTTCGAAATCCGCGTCG |
| 5_known | F3 | Tb927.11.3240_for | GGGGACAAGTTTGTACAAAAAAGCAGGCTATGATGGCATCTAACCTAGCATACATAAAT | Tb927.11.3240_rev | GGGGACCACTTTGTACAAGAAAGCTGGGTTTACTGCGACGGGGGGGC |
| 5_known | F4 | Tb927.11.9560_for | GGGGACAAGTTTGTACAAAAAAGCAGGCTATGACATGCAAACGGACATTTGATG | Tb927.11.9560_rev | GGGGACCACTTTGTACAAGAAAGCTGGGTCTACCCAACGGGGTCTTGTAAAAA |
| 5_known | F5 | Tb927.11.2520_for | GGGGACAAGTTTGTACAAAAAAGCAGGCTATGAGTGACAGGGACGTGTGC | Tb927.11.2520_rev | GGGGACCACTTTGTACAAGAAAGCTGGGTTTACATGTTCGATGATTCGGAGACC |
| 5_known | F6 | Tb927.11.4200_for | GGGGACAAGTTTGTACAAAAAAGCAGGCTATGTCGGCCATCGTGCAC | Tb927.11.4200_rev | GGGGACCACTTTGTACAAGAAAGCTGGGTCTAGTCACGACGATCTCTCGAGC |
| 5_known | F7 | Tb927.10.10130_for | GGGGACAAGTTTGTACAAAAAAGCAGGCTATGCTCAACGTTCTCTCGAGC | Tb927.10.10130_rev | GGGGACCACTTTGTACAAGAAAGCTGGGTTCACATAACTCCGGGAGCAGC |
| 5_known | F8 | Tb927.3.5640_for | GGGGACAAGTTTGTACAAAAAAGCAGGCTATGGTGGAAATCACTGAACTCGC | Tb927.3.5640_rev | GGGGACCACTTTGTACAAGAAAGCTGGGTTCACTCAAGTGCAGGAAATGGTG |
| 5_known | F9 | Tb927.7.6220_for | GGGGACAAGTTTGTACAAAAAAGCAGGCTATGGACTTCCATTCGTTGCAGAG | Tb927.7.6220_rev | GGGGACCACTTTGTACAAGAAAGCTGGGTTCATCTCTCGTTGCGGCGC |
| 5_known | F10 | Tb927.9.11050_for | GGGGACAAGTTTGTACAAAAAAGCAGGCTATGAGAACTACAATTCGATACACTCGC | Tb927.9.11050_rev | GGGGACCACTTTGTACAAGAAAGCTGGGTCTAGCGGCGCTGGTTGGG |
| 5_known | F11 | Tb927.11.2670_for | GGGGACAAGTTTGTACAAAAAAGCAGGCTATGGTGAACCGGAAGGCAAAG | Tb927.11.2670_rev | GGGGACCACTTTGTACAAGAAAGCTGGGTTTAGGGCTTTGATTGCATCGTCAT |
| 5_known | F12 | Tb927.8.7380_for | GGGGACAAGTTTGTACAAAAAAGCAGGCTATGTTACGGCACACGCGTT | Tb927.8.7380_rev | GGGGACCACTTTGTACAAGAAAGCTGGGTCTATTGTTTGCCACCGCCTTGA |
| 5_known | F13 | Tb927.8.4010_for | GGGGACAAGTTTGTACAAAAAAGCAGGCTATGGGTGGCAGGACTGAATCA | Tb927.8.4010_rev | GGGGACCACTTTGTACAAGAAAGCTGGGTTCACTCGAACGCCGGCAA |
| 5_known | F14 | Tb927.11.8870_for | GGGGACAAGTTTGTACAAAAAAGCAGGCTATGAGGGCCCTGCGTTGT | Tb927.11.8870_rev | GGGGACCACTTTGTACAAGAAAGCTGGGTCTACCAACCACTGTCACGAGGT |
| 5_known | F15 | Tb927.7.740_for | GGGGACAAGTTTGTACAAAAAAGCAGGCTATGTCAAGTGGACTAACAGCCAGG | Tb927.7.740_rev | GGGGACCACTTTGTACAAGAAAGCTGGGTTTAACCACCTTCACCTCGTTTCTG |
| 5_known | F16 | Tb927.6.2150_for | GGGGACAAGTTTGTACAAAAAAGCAGGCTATGGAGCTGTCAGCTGTACTTGA | Tb927.6.2150_rev | GGGGACCACTTTGTACAAGAAAGCTGGGTTCACGCGGAGAAGAACAGACT |
| 5_known | F17 | Tb927.5.470_for | GGGGACAAGTTTGTACAAAAAAGCAGGCTATGCTTCCAAGTTTCACCCGTAAA | Tb927.5.470_rev | GGGGACCACTTTGTACAAGAAAGCTGGGTTCACATAATCTCAACAAGTTTTGTCTTCCG |
| 5_known | F18 | Tb927.11.7820_for | GGGGACAAGTTTGTACAAAAAAGCAGGCTATGTATACAACAATGCCTAACTCCCCA | Tb927.11.7820_rev | GGGGACCACTTTGTACAAGAAAGCTGGGTTTAAGAGAGTAAGCACAGCGGAACT |
| 5_known | F19 | Tb927.1.1840_for | GGGGACAAGTTTGTACAAAAAAGCAGGCTATGCGGCCCGCTCCCACG | Tb927.1.1840_rev | GGGGACCACTTTGTACAAGAAAGCTGGGTCTAAGAGCGTTTGCAACTCATTGC |
| 5_known | F20 | Tb927.3.3690_for | GGGGACAAGTTTGTACAAAAAAGCAGGCTATGGCTAGTCTCACAGATGATGACA | Tb927.3.3690_rev | GGGGACCACTTTGTACAAGAAAGCTGGGTCTAATCATATTCTTCTTCATCTTCCCCAGC |
| 5_known | F21 | Tb927.11.7070_for | GGGGACAAGTTTGTACAAAAAAGCAGGCTATGAAAACCTTTGGGGTGCGAT | Tb927.11.7070_rev | GGGGACCACTTTGTACAAGAAAGCTGGGTTCAATATGCACATTTAGCACCTGAAACC |
| 5_known | F22 | Tb927.8.4540_for | GGGGACAAGTTTGTACAAAAAAGCAGGCTATGGAGGGGTCTCATAGCAGC | Tb927.8.4540_rev | GGGGACCACTTTGTACAAGAAAGCTGGGTCTATTTCCCAACTCGTTTCTTCGGC |
| 5_known | F23 | Tb927.9.11900_for | GGGGACAAGTTTGTACAAAAAAGCAGGCTATGTCGAAATCGCTGCAGCG | Tb927.9.11900_rev | GGGGACCACTTTGTACAAGAAAGCTGGGTCTACAAGAAACGTTCCTCGGATGG |
| 5_known | F24 | Tb927.11.3420_for | GGGGACAAGTTTGTACAAAAAAGCAGGCTATGGTGCATAACACCTCCACTGC | Tb927.11.3420_rev | GGGGACCACTTTGTACAAGAAAGCTGGGTCTACTGCGGTGCATTGGCA |
| 5_known | G1 | Tb927.5.2940_for | GGGGACAAGTTTGTACAAAAAAGCAGGCTATGGACGCAACGGAGTTAAAGAAT | Tb927.5.2940_rev | GGGGACCACTTTGTACAAGAAAGCTGGGTCTACTGGCCGAAACGGATAATCC |
| 5_known | G2 | Tb927.10.7050_for | GGGGACAAGTTTGTACAAAAAAGCAGGCTATGGCAACATCCATTCAGCAAAATC | Tb927.10.7050_rev | GGGGACCACTTTGTACAAGAAAGCTGGGTTTAATCGTTGAGGTTATCCATAAGTTTGGA |
| 5_known | G3 | Tb927.10.7930_for | GGGGACAAGTTTGTACAAAAAAGCAGGCTATGGCACTCACGCTTGCTG | Tb927.10.7930_rev | GGGGACCACTTTGTACAAGAAAGCTGGGTCTACGCTACCTCAATGAGCGAC |
| 5_known | G4 | Tb927.4.2290_for | GGGGACAAGTTTGTACAAAAAAGCAGGCTATGATTGAGGGAAGCAGAAACTCAAG | Tb927.4.2290_rev | GGGGACCACTTTGTACAAGAAAGCTGGGTCTAAGATGATTGCCATAAGCGTAAATGC |
| 5_known | G5 | Tb927.11.16760_for | GGGGACAAGTTTGTACAAAAAAGCAGGCTATGCAAACCAAGCAGAAGGCC | Tb927.11.16760_rev | GGGGACCACTTTGTACAAGAAAGCTGGGTTTAACGGCGTCCCTCCTCC |
| 5_known | G6 | Tb927.6.5030_for | GGGGACAAGTTTGTACAAAAAAGCAGGCTATGTCCAAAGGTTACACGGTATATTCC | Tb927.6.5030_rev | GGGGACCACTTTGTACAAGAAAGCTGGGTCTAACAACTGTGGACCCACAGGG |
| 5_known | G7 | Tb927.2.5750_for | GGGGACAAGTTTGTACAAAAAAGCAGGCTATGGAGGAAACAAAGTTATGCCCATT | Tb927.2.5750_rev | GGGGACCACTTTGTACAAGAAAGCTGGGTTCACGCATCCCTGTTGGTTCG |
| 5_known | G8 | Tb927.11.9900_for | GGGGACAAGTTTGTACAAAAAAGCAGGCTATGTTGCGCCGTTCTTTCTTGC | Tb927.11.9900_rev | GGGGACCACTTTGTACAAGAAAGCTGGGTTCACTTCCGCCCGAATCCA |
| 5_known | G9 | Tb927.8.1640_for | GGGGACAAGTTTGTACAAAAAAGCAGGCTATGATGCTACCGGCGTGTGT | Tb927.8.1640_rev | GGGGACCACTTTGTACAAGAAAGCTGGGTTTAACAGGCAGCATAAACTATAGCGAA |
| 5_known | G10 | Tb927.8.2960_for | GGGGACAAGTTTGTACAAAAAAGCAGGCTATGGGCGGGAACCGACGG | Tb927.8.2960_rev | GGGGACCACTTTGTACAAGAAAGCTGGGTTCACCATGGGCTTTCATCGC |
| 5_known | G11 | Tb927.11.8940_for | GGGGACAAGTTTGTACAAAAAAGCAGGCTATGTCTACGGGAAAGATAATTGGTGATC | Tb927.11.8940_rev | GGGGACCACTTTGTACAAGAAAGCTGGGTCTAGTTCTTCTCCAGCCAACGGA |
| 5_known | G14 | Tb927.11.1930_for | GGGGACAAGTTTGTACAAAAAAGCAGGCTATGTCCTTTTTCCAGTTGGATTGGG | Tb927.11.1930_rev | GGGGACCACTTTGTACAAGAAAGCTGGGTCTAAGCAGAACGTACGTAACACAGA |
| 5_known | G16 | Tb927.11.8350_for | GGGGACAAGTTTGTACAAAAAAGCAGGCTATGAGCACATCTAATGACGTGCTG | Tb927.11.8350_rev | GGGGACCACTTTGTACAAGAAAGCTGGGTCTAAACATTTACCTCGAACTCCTTCTTCC |
| 5_known | G17 | Tb927.10.1620_for | GGGGACAAGTTTGTACAAAAAAGCAGGCTATGCAGGAACTTATCGCTGTCCG | Tb927.10.1620_rev | GGGGACCACTTTGTACAAGAAAGCTGGGTTCAACTCCTGCTGGATGGTATGG |
| 5_known | G18 | Tb927.10.8720_for | GGGGACAAGTTTGTACAAAAAAGCAGGCTATGACGGATGATGTCTTGAGACAAC | Tb927.10.8720_rev | GGGGACCACTTTGTACAAGAAAGCTGGGTTCATGCGCACCATGAAAGCC |
| 5_known | G19 | Tb927.11.5050_for | GGGGACAAGTTTGTACAAAAAAGCAGGCTATGTTACGACGGTCGTCCCT | Tb927.11.5050_rev | GGGGACCACTTTGTACAAGAAAGCTGGGTCTACCTGAGCTGTTGGAAGAAGTCA |
| 5_known | G20 | Tb927.9.10770_for | GGGGACAAGTTTGTACAAAAAAGCAGGCTATGGCTGCATTTGCTGCTGC | Tb927.9.10770_rev | GGGGACCACTTTGTACAAGAAAGCTGGGTCTACATGCCAATGTGACGGTTGA |
| 5_known | G21 | Tb927.7.4000_for | GGGGACAAGTTTGTACAAAAAAGCAGGCTATGGTGTTAAAATTGTTGCTGGAGC | Tb927.7.4000_rev | GGGGACCACTTTGTACAAGAAAGCTGGGTTTACGGTACAACCGCTAAGGAATCA |
| 5_known | G22 | Tb927.8.3150_for | GGGGACAAGTTTGTACAAAAAAGCAGGCTATGCAGCAGCAACAGCCC | Tb927.8.3150_rev | GGGGACCACTTTGTACAAGAAAGCTGGGTTTAGGCAGCTTCAGCCTCACC |
| 5_known | G23 | Tb927.7.210_for | GGGGACAAGTTTGTACAAAAAAGCAGGCTATGTTTCGCCTCCTGTGCCG | Tb927.7.210_rev | GGGGACCACTTTGTACAAGAAAGCTGGGTTCACATCCAAAAGACGCGACG |
| 5_known | G24 | Tb927.11.13810_for | GGGGACAAGTTTGTACAAAAAAGCAGGCTATGGATATATTTAGTGCTCTTTCCACGG | Tb927.11.13810_rev | GGGGACCACTTTGTACAAGAAAGCTGGGTCTACTGCACCTCTTCCACGC |
| 5_known | H1 | Tb927.4.260_for | GGGGACAAGTTTGTACAAAAAAGCAGGCTATGAGGAGGCCACGGGATG | Tb927.4.260_rev | GGGGACCACTTTGTACAAGAAAGCTGGGTTTAATTCATCATCATATTTCTGATCATCAG |
| 5_known | H2 | Tb927.9.12790_for | GGGGACAAGTTTGTACAAAAAAGCAGGCTATGATGATATATCCTCGCATGAGAGTGG | Tb927.9.12790_rev | GGGGACCACTTTGTACAAGAAAGCTGGGTCTAAGCACAACTATCTTCACCCCAC |
| 5_known | H5 | Tb927.11.790_for | GGGGACAAGTTTGTACAAAAAAGCAGGCTATGCCCTTGAATTCATCTCACCG | Tb927.11.790_rev | GGGGACCACTTTGTACAAGAAAGCTGGGTTTACGCCACGAGCGATAGTTTC |
| 5_known | H6 | Tb927.10.4050_for | GGGGACAAGTTTGTACAAAAAAGCAGGCTATGCCTACTTACGTGGAGGCG | Tb927.10.4050_rev | GGGGACCACTTTGTACAAGAAAGCTGGGTTTACTCCTCGCCCTGCGG |
| 5_known | H7 | Tb927.4.280_for | GGGGACAAGTTTGTACAAAAAAGCAGGCTATGAGGAGGCCACGGGATG | Tb927.4.280_rev | GGGGACCACTTTGTACAAGAAAGCTGGGTTTAATTCATCATCATATTTCTGATCATAAG |
| 5_known | H8 | Tb927.8.1830_for | GGGGACAAGTTTGTACAAAAAAGCAGGCTATGACGTCCCAGGCGGTG | Tb927.8.1830_rev | GGGGACCACTTTGTACAAGAAAGCTGGGTCTAAATAATCCAACCACTGGCTACCAC |
| 5_known | H9 | Tb927.10.2490_for | GGGGACAAGTTTGTACAAAAAAGCAGGCTATGGACGGTGATCTTTCCCAACT | Tb927.10.2490_rev | GGGGACCACTTTGTACAAGAAAGCTGGGTTTACAAATGATGAAGCTTCCGCTGC |
| 5_known | H10 | Tb927.10.4780_for | GGGGACAAGTTTGTACAAAAAAGCAGGCTATGAGTCGACTCTATATATTTTACAAACTG | Tb927.10.4780_rev | GGGGACCACTTTGTACAAGAAAGCTGGGTTCAGTAGCCCCCCTGGTCC |
| 5_known | H11 | Tb927.10.310_for | GGGGACAAGTTTGTACAAAAAAGCAGGCTATGTGGACTGTGCAGGAGGAC | Tb927.10.310_rev | GGGGACCACTTTGTACAAGAAAGCTGGGTTCAGCCGGAGAGCGGTTG |
| 5_known | H12 | Tb927.6.1880_for | GGGGACAAGTTTGTACAAAAAAGCAGGCTATGTCGGACCCCGCTCAA | Tb927.6.1880_rev | GGGGACCACTTTGTACAAGAAAGCTGGGTCTACGGTGTCACACGCTGC |
| 5_known | H13 | Tb927.11.6780_for | GGGGACAAGTTTGTACAAAAAAGCAGGCTATGGAAGAACGCGTTTCTTGTGT | Tb927.11.6780_rev | GGGGACCACTTTGTACAAGAAAGCTGGGTCTAACAAAGTGTTGCGGGGTCG |
| 5_known | H14 | Tb927.11.4900_for | GGGGACAAGTTTGTACAAAAAAGCAGGCTATGCGAATCAAGGTAGACGTTGAG | Tb927.11.4900_rev | GGGGACCACTTTGTACAAGAAAGCTGGGTTTACGCAGCATACGCTTCAGTC |
| 5_known | H15 | Tb927.6.2850_for | GGGGACAAGTTTGTACAAAAAAGCAGGCTATGTCCCACTTTCAGCCTTCGC | Tb927.6.2850_rev | GGGGACCACTTTGTACAAGAAAGCTGGGTTTAGTCGCTGCAAAAGTCCGC |
| 5_known | H16 | Tb927.9.12360_for | GGGGACAAGTTTGTACAAAAAAGCAGGCTATGGAGGGCACTGGCGGT | Tb927.9.12360_rev | GGGGACCACTTTGTACAAGAAAGCTGGGTTTATGCTGACTGATGCGGTGGG |

| | | | | | |
|---|---|---|---|---|---|
| 5_known | H17 | Tb927.4.4590_for | GGGGACAAGTTTGTACAAAAAAGCAGGCTATGGTGCCGTCTGAACCCC | Tb927.4.4590_rev | GGGGACCACTTTGTACAAGAAAGCTGGGTTCAAGGCCCCGGCTTCGA |
| 5_known | H18 | Tb927.9.4300_for | GGGGACAAGTTTGTACAAAAAAGCAGGCTATGCCTGCGAAGCGACGG | Tb927.9.4300_rev | GGGGACCACTTTGTACAAGAAAGCTGGGTTCACACAGTTGACTGACTTTCCG |
| 5_known | H19 | Tb927.10.3210_for | GGGGACAAGTTTGTACAAAAAAGCAGGCTATGCTTCGCCGTACGTTGC | Tb927.10.3210_rev | GGGGACCACTTTGTACAAGAAAGCTGGGTTCACAGAGTCACGTAGTCGGG |
| 5_known | H20 | Tb927.11.7750_for | GGGGACAAGTTTGTACAAAAAAGCAGGCTATGACAGTGATTATGTTCCCTCGTTATATC | Tb927.11.7750_rev | GGGGACCACTTTGTACAAGAAAGCTGGGTTTATAATAGGGAAAAGCAAGAAACGAAGAG |
| 5_known | H21 | Tb927.10.6400_for | GGGGACAAGTTTGTACAAAAAAGCAGGCTATGTTCCGCTGTGTCGTCCG | Tb927.10.6400_rev | GGGGACCACTTTGTACAAGAAAGCTGGGTTCAATACATGCCGTCCATACCTCC |
| 5_known | H23 | Tb927.10.9850_for | GGGGACAAGTTTGTACAAAAAAGCAGGCTATGCATTCAGTCCACGAATCGC | Tb927.10.9850_rev | GGGGACCACTTTGTACAAGAAAGCTGGGTTTACAAAGCAGGAAGGACGTACTGT |
| 5_known | H24 | Tb927.10.15570_for | GGGGACAAGTTTGTACAAAAAAGCAGGCTATGAATGGAAGGTATGCTCGGCA | Tb927.10.15570_rev | GGGGACCACTTTGTACAAGAAAGCTGGGTCTATATTTCACGAGAAGTTACGGCCC |
| 5_known | I1 | Tb927.4.4220_for | GGGGACAAGTTTGTACAAAAAAGCAGGCTATGCGCCCCGAGTTCCCG | Tb927.4.4220_rev | GGGGACCACTTTGTACAAGAAAGCTGGGTTCAACAACCCGAGCGTTTGG |
| 5_known | I2 | Tb927.11.11500_for | GGGGACAAGTTTGTACAAAAAAGCAGGCTATGAACGATCTGTCCGCAGTCG | Tb927.11.11500_rev | GGGGACCACTTTGTACAAGAAAGCTGGGTTTAGGGCCACGCAAACTTCTTAC |
| 5_known | I3 | Tb927.10.15970_for | GGGGACAAGTTTGTACAAAAAAGCAGGCTATGTGTACGTGCACCTGCTG | Tb927.10.15970_rev | GGGGACCACTTTGTACAAGAAAGCTGGGTCTAGGTGGATCGTCCGTTGGT |
| 5_known | I4 | Tb927.11.5440_for | GGGGACAAGTTTGTACAAAAAAGCAGGCTATGACGCATCAATCAACTGATGTGT | Tb927.11.5440_rev | GGGGACCACTTTGTACAAGAAAGCTGGGTTTACTCGTCAGGGAGAATGCCTG |
| 5_known | I5 | Tb927.11.7710_for | GGGGACAAGTTTGTACAAAAAAGCAGGCTATGACAGTGATTATGTTCCCTCGTTATATC | Tb927.11.7710_rev | GGGGACCACTTTGTACAAGAAAGCTGGGTTTATAATAGGGGAAAACAAGAAACGAAGAG |
| 5_known | I6 | Tb927.8.1240_for | GGGGACAAGTTTGTACAAAAAAGCAGGCTATGCTTCGTCGAACAAGTTGTTGT | Tb927.8.1240_rev | GGGGACCACTTTGTACAAGAAAGCTGGGTTCACATTTGTGAGTTATAGTTGGGACC |
| 5_known | I7 | Tb927.10.1540_for | GGGGACAAGTTTGTACAAAAAAGCAGGCTATGCCTCCCGCTGACAGG | Tb927.10.1540_rev | GGGGACCACTTTGTACAAGAAAGCTGGGTCTACGCTTTTTGCCGACGTTTC |
| 5_known | I8 | Tb927.11.6270_for | GGGGACAAGTTTGTACAAAAAAGCAGGCTATGCGCAGCCTACTGGAAGG | Tb927.11.6270_rev | GGGGACCACTTTGTACAAGAAAGCTGGGTTCAAAAAAACAGTGTCACAGCGGC |
| 5_known | I9 | Tb927.10.5510_for | GGGGACAAGTTTGTACAAAAAAGCAGGCTATGAGGGCTGCCGAGGGT | Tb927.10.5510_rev | GGGGACCACTTTGTACAAGAAAGCTGGGTTTATTTGTTGCTTGGAATTGTTGCAACG |
| 5_known | I10 | Tb927.3.4500_for | GGGGACAAGTTTGTACAAAAAAGCAGGCTATGAGTCTCTGCGAAAACTGTGAG | Tb927.3.4500_rev | GGGGACCACTTTGTACAAGAAAGCTGGGTCTAGACGAGTTTAGCATAGAAGTCATTACC |
| 5_known | I11 | Tb927.11.2470_for | GGGGACAAGTTTGTACAAAAAAGCAGGCTATGCCTGCAACCCTCAAGC | Tb927.11.2470_rev | GGGGACCACTTTGTACAAGAAAGCTGGGTTTATGCTGTCCCAAAGAGTGAGGC |
| 5_known | I12 | Tb927.10.5970_for | GGGGACAAGTTTGTACAAAAAAGCAGGCTATGTCGACCACTTCTTTCGGGT | Tb927.10.5970_rev | GGGGACCACTTTGTACAAGAAAGCTGGGTTCACCTACTTCTCTTGCGCCC |
| 5_known | I13 | Tb927.11.15600_for | GGGGACAAGTTTGTACAAAAAAGCAGGCTATGGTTGTGGAGCCTCCTGC | Tb927.11.15600_rev | GGGGACCACTTTGTACAAGAAAGCTGGGTTCACTGTCCCTCACTTAGACTCCC |
| 5_known | I14 | Tb927.3.1130_for | GGGGACAAGTTTGTACAAAAAAGCAGGCTATGTCTGAGGCTCTGATTGACGA | Tb927.3.1130_rev | GGGGACCACTTTGTACAAGAAAGCTGGGTTTAACGTGGGTTGCGCAGC |
| 5_known | I15 | Tb927.11.5300_for | GGGGACAAGTTTGTACAAAAAAGCAGGCTATGAATAATAGCCGCATTTGCGTTG | Tb927.11.5300_rev | GGGGACCACTTTGTACAAGAAAGCTGGGTTTAATTGCGTCTCTTCATGGAAGAGTT |
| 5_known | I16 | Tb927.5.1580_for | GGGGACAAGTTTGTACAAAAAAGCAGGCTATGGCGAGAAACCGGAAACATG | Tb927.5.1580_rev | GGGGACCACTTTGTACAAGAAAGCTGGGTTCACAGAGGGTACACGGGC |
| 5_known | I17 | Tb927.9.9290_for | GGGGACAAGTTTGTACAAAAAAGCAGGCTATGACAATCGCTGCACAGGG | Tb927.9.9290_rev | GGGGACCACTTTGTACAAGAAAGCTGGGTTTAAGCGCTTGAGGCGTGTAC |
| 5_known | I18 | Tb927.3.3000_for | GGGGACAAGTTTGTACAAAAAAGCAGGCTATGCTGTCTACAAGTGGGAGGG | Tb927.3.3000_rev | GGGGACCACTTTGTACAAGAAAGCTGGGTTCACTTCTTCTTCAATTTGCTCTCCTT |
| 5_known | I19 | Tb927.3.2820_for | GGGGACAAGTTTGTACAAAAAAGCAGGCTATGTTAAATGTTCGGGGTTTGTTAAGC | Tb927.3.2820_rev | GGGGACCACTTTGTACAAGAAAGCTGGGTTTAGCTCCCCAGCCGACC |
| 5_known | I20 | Tb927.11.3870_for | GGGGACAAGTTTGTACAAAAAAGCAGGCTATGAGTAAAAAGGGTAACATTCGCCTT | Tb927.11.3870_rev | GGGGACCACTTTGTACAAGAAAGCTGGGTTTATGACGACGCTCCGGAGC |
| 5_known | I21 | Tb927.10.6630_for | GGGGACAAGTTTGTACAAAAAAGCAGGCTATGGAGGAGACGTACAGTCCCT | Tb927.10.6630_rev | GGGGACCACTTTGTACAAGAAAGCTGGGTCTATCTCGCGTATTTCTTGCGGG |
| 5_known | I22 | Tb927.9.11270_for | GGGGACAAGTTTGTACAAAAAAGCAGGCTATGATTCAGCCACAATTAATTTTACTCCG | Tb927.9.11270_rev | GGGGACCACTTTGTACAAGAAAGCTGGGTTCAGCGGCCGGCACGACC |
| 5_known | I23 | Tb927.10.11100_for | GGGGACAAGTTTGTACAAAAAAGCAGGCTATGTACGTACCGAGTCGTGTTGT | Tb927.10.11100_rev | GGGGACCACTTTGTACAAGAAAGCTGGGTCTAAATCACCTCTGCTGCATGTTTCT |
| 5_known | I24 | Tb927.11.16570_for | GGGGACAAGTTTGTACAAAAAAGCAGGCTATGAGATGCTGTGTCCGAGCC | Tb927.11.16570_rev | GGGGACCACTTTGTACAAGAAAGCTGGGTCTACGATCCAAAGGAACGTCGC |
| 5_known | J1 | Tb927.10.13510_for | GGGGACAAGTTTGTACAAAAAAGCAGGCTATGAGCACTGCTAGCAACACC | Tb927.10.13510_rev | GGGGACCACTTTGTACAAGAAAGCTGGGTCTAGTACTGCGGCTGGTTTTGC |
| 5_known | J2 | Tb927.1.730_for | GGGGACAAGTTTGTACAAAAAAGCAGGCTATGTGGCGGCTGCTTCGT | Tb927.1.730_rev | GGGGACCACTTTGTACAAGAAAGCTGGGTTCACTTCAGTTTGGAGTGTTTACCAC |
| 5_known | J3 | Tb927.10.10870_for | GGGGACAAGTTTGTACAAAAAAGCAGGCTATGCACAATGTCCCGTTACTCG | Tb927.10.10870_rev | GGGGACCACTTTGTACAAGAAAGCTGGGTTTAACGAGGTGATGAAAAGTGCCTC |
| 5_known | J6 | Tb927.11.5450_for | GGGGACAAGTTTGTACAAAAAAGCAGGCTATGTTGGGTCGTTCGTTTAAACTTTG | Tb927.11.5450_rev | GGGGACCACTTTGTACAAGAAAGCTGGGTTTACTCCAGCTTCTTCGACAGGT |
| 5_known | J7 | Tb927.11.8620_for | GGGGACAAGTTTGTACAAAAAAGCAGGCTATGTGTGGGGGAGAAGAGAATGG | Tb927.11.8620_rev | GGGGACCACTTTGTACAAGAAAGCTGGGTCTAAATTATGGTAACTTCCCTCTCAACAGT |
| 5_known | J11 | Tb927.4.300_for | GGGGACAAGTTTGTACAAAAAAGCAGGCTATGTCAAGAGCCAATAGCCCCG | Tb927.4.300_rev | GGGGACCACTTTGTACAAGAAAGCTGGGTTTAATTCATCATCATATTTCTGATCATCAG |
| 5_known | J12 | Tb927.11.8490_for | GGGGACAAGTTTGTACAAAAAAGCAGGCTATGTGTGGGGGAGAAGAGAATGG | Tb927.11.8490_rev | GGGGACCACTTTGTACAAGAAAGCTGGGTTCACTCAATTATGGTAACTTCCCTCTCA |
| 5_known | J13 | Tb927.10.4430_for | GGGGACAAGTTTGTACAAAAAAGCAGGCTATGTCGTCGGATGAGGAAACCA | Tb927.10.4430_rev | GGGGACCACTTTGTACAAGAAAGCTGGGTCTATGTTCCTTTCTTGCTTTCCGGA |
| 5_known | J15 | Tb927.11.10640_for | GGGGACAAGTTTGTACAAAAAAGCAGGCTATGTTTCTCATCAGCTGGAATGTGG | Tb927.11.10640_rev | GGGGACCACTTTGTACAAGAAAGCTGGGTTTATTCACTTCTTGGCCGCTTTTGG |
| 5_known | J16 | Tb927.5.3050_for | GGGGACAAGTTTGTACAAAAAAGCAGGCTATGGCGGCTAAAGGAAAGTCAAC | Tb927.5.3050_rev | GGGGACCACTTTGTACAAGAAAGCTGGGTCTATTTATACTCAAAAGAAACACGAACTAG |
| 5_known | J17 | Tb927.11.15010_for | GGGGACAAGTTTGTACAAAAAAGCAGGCTATGGGAAACGGTGTTGTGAGAGT | Tb927.11.15010_rev | GGGGACCACTTTGTACAAGAAAGCTGGGTCTACCCCATGTCAAGAGTAAACATGAG |
| 5_known | J18 | Tb927.1.1380_for | GGGGACAAGTTTGTACAAAAAAGCAGGCTATGTCAGAGCACTTTAAAAATGTCAAGTC | Tb927.1.1380_rev | GGGGACCACTTTGTACAAGAAAGCTGGGTTTAATTCTCTGTGGCGGCAAAACG |
| 5_known | J19 | Tb927.11.6750_for | GGGGACAAGTTTGTACAAAAAAGCAGGCTATGTTCCGTCTTGTAAGACCCCG | Tb927.11.6750_rev | GGGGACCACTTTGTACAAGAAAGCTGGGTTCACACGAACCACCCCTTGC |
| 5_known | J20 | Tb927.8.5140_for | GGGGACAAGTTTGTACAAAAAAGCAGGCTATGTCGTTATCTAACGTTGTTGCAAATAAC | Tb927.8.5140_rev | GGGGACCACTTTGTACAAGAAAGCTGGGTTCACGCCCCTTCCCCTTC |
| 5_known | J21 | Tb927.11.3070_for | GGGGACAAGTTTGTACAAAAAAGCAGGCTATGCGGAAAGCCGCTAAACG | Tb927.11.3070_rev | GGGGACCACTTTGTACAAGAAAGCTGGGTTCACTTCATCAGTGACCGCCA |
| 5_known | J22 | Tb927.7.1640_for | GGGGACAAGTTTGTACAAAAAAGCAGGCTATGAGACGCTGCATCCCTGC | Tb927.7.1640_rev | GGGGACCACTTTGTACAAGAAAGCTGGGTTTAGATGTAGTCAGGGTTGGAATCCTT |
| 5_known | J23 | Tb927.10.5780_for | GGGGACAAGTTTGTACAAAAAAGCAGGCTATGACAAGTCGTGAGGGTATTCACC | Tb927.10.5780_rev | GGGGACCACTTTGTACAAGAAAGCTGGGTTTACTTTGATAGCATTGGGGCGAAA |
| 5_known | J24 | Tb927.9.5210_for | GGGGACAAGTTTGTACAAAAAAGCAGGCTATGACAACAACGGTTGATTCCGC | Tb927.9.5210_rev | GGGGACCACTTTGTACAAGAAAGCTGGGTTTACTGTTTCTTTCCCTTCACCCCC |
| 5_known | K1 | Tb927.10.3110_for | GGGGACAAGTTTGTACAAAAAAGCAGGCTATGCAAGCGATAGGAACTTCGC | Tb927.10.3110_rev | GGGGACCACTTTGTACAAGAAAGCTGGGTCTACAAAGAATCACGCGGACGG |
| 5_known | K2 | Tb927.7.3790_for | GGGGACAAGTTTGTACAAAAAAGCAGGCTATGGATCATATTGATCTTGCTGAAGAGC | Tb927.7.3790_rev | GGGGACCACTTTGTACAAGAAAGCTGGGTTTAGTCCAATAGCACGTCAATAGCC |
| 5_known | K3 | Tb927.10.16150_for | GGGGACAAGTTTGTACAAAAAAGCAGGCTATGGATCCCCTGTCGCGTG | Tb927.10.16150_rev | GGGGACCACTTTGTACAAGAAAGCTGGGTTCAGGCTGTCACATCACTGAACG |
| 5_known | K4 | Tb927.11.6220_for | GGGGACAAGTTTGTACAAAAAAGCAGGCTATGCCAACGCGGGAGGGT | Tb927.11.6220_rev | GGGGACCACTTTGTACAAGAAAGCTGGGTCTAAAGTCGCTCTAGTTCCTTTGCC |
| 5_known | K5 | Tb927.3.3400_for | GGGGACAAGTTTGTACAAAAAAGCAGGCTATGCATGACTCTGTGGGAACGG | Tb927.3.3400_rev | GGGGACCACTTTGTACAAGAAAGCTGGGTTTAAACACATTCAGATAGGATAAACCCGTC |
| 5_known | K6 | Tb927.9.2490_for | GGGGACAAGTTTGTACAAAAAAGCAGGCTATGACGGGAATGAACAGATCGTG | Tb927.9.2490_rev | GGGGACCACTTTGTACAAGAAAGCTGGGTCTATCCATCAACGTCCTCCCCT |
| 5_known | K7 | Tb927.2.1180_for | GGGGACAAGTTTGTACAAAAAAGCAGGCTATGAATCAACAGGTTCCTATTGAAGGAAG | Tb927.2.1180_rev | GGGGACCACTTTGTACAAGAAAGCTGGGTTTAACTCATCATCATATTTCGAATCATCAG |
| 5_known | K8 | Tb927.4.3840_for | GGGGACAAGTTTGTACAAAAAAGCAGGCTATGTCCAAGCCGGATCGAAAGG | Tb927.4.3840_rev | GGGGACCACTTTGTACAAGAAAGCTGGGTTCATCCACTTTTCCCTTTACTCTTTGG |
| 5_known | K9 | Tb927.3.5250_for | GGGGACAAGTTTGTACAAAAAAGCAGGCTATGGGGCGTGGAAAGGGG | Tb927.3.5250_rev | GGGGACCACTTTGTACAAGAAAGCTGGGTCTATTCTGTGGTTAGCAGCGCC |
| 5_known | K10 | Tb927.8.5500_for | GGGGACAAGTTTGTACAAAAAAGCAGGCTATGGCTCATGGAGCTTCTCCG | Tb927.8.5500_rev | GGGGACCACTTTGTACAAGAAAGCTGGGTCTATGTATGATGATTCTGGATGACTTTAGC |
| 5_known | K11 | Tb927.5.4480_for | GGGGACAAGTTTGTACAAAAAAGCAGGCTATGGCTCCGAAGAGGGGAAG | Tb927.5.4480_rev | GGGGACCACTTTGTACAAGAAAGCTGGGTCTACATCTCATATGGTATAAAGTGAACCTC |
| 5_known | K12 | Tb927.11.3270_for | GGGGACAAGTTTGTACAAAAAAGCAGGCTATGGAGGCGGCCATTATTGGA | Tb927.11.3270_rev | GGGGACCACTTTGTACAAGAAAGCTGGGTTCACAAACCAATTGGCTTCCTTTTCC |
| 5_known | K13 | Tb927.9.12900_for | GGGGACAAGTTTGTACAAAAAAGCAGGCTATGTATCGACACGGTCTGGAATCA | Tb927.9.12900_rev | GGGGACCACTTTGTACAAGAAAGCTGGGTTCACTTTCGCTCATGTAGCTTGC |
| 5_known | K14 | Tb927.10.10990_for | GGGGACAAGTTTGTACAAAAAAGCAGGCTATGAGACATTGGGCGCAGGC | Tb927.10.10990_rev | GGGGACCACTTTGTACAAGAAAGCTGGGTCTACCGCAACTCGTCTTCAAGG |
| 5_known | K15 | Tb927.10.12380_for | GGGGACAAGTTTGTACAAAAAAGCAGGCTATGGAAGATGCGAAATACCGTACG | Tb927.10.12380_rev | GGGGACCACTTTGTACAAGAAAGCTGGGTTCAGTCTACGATGATGCCGATAAGC |
| 5_known | K16 | Tb927.5.3710_for | GGGGACAAGTTTGTACAAAAAAGCAGGCTATGGCTGCGGAGATAACTGTTCT | Tb927.5.3710_rev | GGGGACCACTTTGTACAAGAAAGCTGGGTTCATTTACAACCATTACCTTTATGCATTTC |
| 5_known | K17 | Tb927.11.10750_for | GGGGACAAGTTTGTACAAAAAAGCAGGCTATGTCACGCTACATCCCACCA | Tb927.11.10750_rev | GGGGACCACTTTGTACAAGAAAGCTGGGTTCACGCGTACTCCAGTGCG |
| 5_known | K18 | Tb927.4.1140_for | GGGGACAAGTTTGTACAAAAAAGCAGGCTATGTCGAATGCTTTGTTTTGTCTTCAC | Tb927.4.1140_rev | GGGGACCACTTTGTACAAGAAAGCTGGGTCTAGAGTATACCATTCTGGAGAGTTTGGA |
| 5_known | K19 | Tb927.4.5040_for | GGGGACAAGTTTGTACAAAAAAGCAGGCTATGATTCGGCTGTGCCGGG | Tb927.4.5040_rev | GGGGACCACTTTGTACAAGAAAGCTGGGTCTAGCGTCGCTCAATATCCTCG |
| 5_known | K20 | Tb927.11.9890_for | GGGGACAAGTTTGTACAAAAAAGCAGGCTATGATCGACACGCTTTCGATAGTTAG | Tb927.11.9890_rev | GGGGACCACTTTGTACAAGAAAGCTGGGTCTACGCCATTAAAGCGCTGACA |
| 5_known | K21 | Tb927.3.690_for | GGGGACAAGTTTGTACAAAAAAGCAGGCTATGGAGAACTACCAACTTCTCAATCAGT | Tb927.3.690_rev | GGGGACCACTTTGTACAAGAAAGCTGGGTCTAACTGTTGGAACCTTGGTTGTGC |
| 5_known | K22 | Tb927.7.6900_for | GGGGACAAGTTTGTACAAAAAAGCAGGCTATGTTCTACTCAACCTACGTGCTGA | Tb927.7.6900_rev | GGGGACCACTTTGTACAAGAAAGCTGGGTTCACACAGTGAGTTGCACCTCC |
| 5_known | K23 | Tb927.7.7420_for | GGGGACAAGTTTGTACAAAAAAGCAGGCTATGCGTCGCTTTGGTTCCAA | Tb927.7.7420_rev | GGGGACCACTTTGTACAAGAAAGCTGGGTCTACACTGCCCGCTTACAAGAG |
| 5_known | K24 | Tb927.7.250_for | GGGGACAAGTTTGTACAAAAAAGCAGGCTATGCAGAAACAGCGTATCACTTTCG | Tb927.7.250_rev | GGGGACCACTTTGTACAAGAAAGCTGGGTTTACACCCTTGCTTCCATGATTACAC |
| 5_known | L1 | Tb927.3.680_for | GGGGACAAGTTTGTACAAAAAAGCAGGCTATGGCTGTAATAACGAATACCTCAAGTC | Tb927.3.680_rev | GGGGACCACTTTGTACAAGAAAGCTGGGTTTAATAACGAGGTCGCGGATGGC |
| 5_known | L3 | Tb927.8.1600_for | GGGGACAAGTTTGTACAAAAAAGCAGGCTATGTCGGCTGTGGAGGAGC | Tb927.8.1600_rev | GGGGACCACTTTGTACAAGAAAGCTGGGTTCAAAGAAGTGGCACCCCTTGT |
| 5_known | L4 | Tb927.1.1470_for | GGGGACAAGTTTGTACAAAAAAGCAGGCTATGGGTGGTATTCCTTCACGCG | Tb927.1.1470_rev | GGGGACCACTTTGTACAAGAAAGCTGGGTCTACTGACCCCAGAAGCCC |
| 5_known | L5 | Tb927.7.5970_for | GGGGACAAGTTTGTACAAAAAAGCAGGCTATGAGCGCACCCGTCGAC | Tb927.7.5970_rev | GGGGACCACTTTGTACAAGAAAGCTGGGTCTATTCTGTGTGGGACAGTGACTCA |
| 5_known | L6 | Tb927.7.460_for | GGGGACAAGTTTGTACAAAAAAGCAGGCTATGTCTTTTGTACCACCTCCAGCA | Tb927.7.460_rev | GGGGACCACTTTGTACAAGAAAGCTGGGTTCAGCACATGAAAGTCTTTGCGC |
| 5_known | L7 | Tb927.11.4480_for | GGGGACAAGTTTGTACAAAAAAGCAGGCTATGCCAGTAAATCCTGAACCACAG | Tb927.11.4480_rev | GGGGACCACTTTGTACAAGAAAGCTGGGTTTACATTTCTTCTTCCTCTTCCTCAGCT |
| 5_known | L8 | Tb927.9.8400_for | GGGGACAAGTTTGTACAAAAAAGCAGGCTATGATTGGAAATGGGCGAGGATTG | Tb927.9.8400_rev | GGGGACCACTTTGTACAAGAAAGCTGGGTTTAGCAACGCCTTGACTGACC |
| 5_known | L10 | Tb927.11.14580_for | GGGGACAAGTTTGTACAAAAAAGCAGGCTATGTCAGCTACTGATGTCGCATCT | Tb927.11.14580_rev | GGGGACCACTTTGTACAAGAAAGCTGGGTTCACACATTTCTGTAAAAGGGAACTTTCT |
| 5_known | L11 | Tb927.8.2850_for | GGGGACAAGTTTGTACAAAAAAGCAGGCTATGCAGGTGACACGGGACA | Tb927.8.2850_rev | GGGGACCACTTTGTACAAGAAAGCTGGGTTCACCTTCTTAGTGCTTTGCAAACC |
| 5_known | L12 | Tb927.5.2900_for | GGGGACAAGTTTGTACAAAAAAGCAGGCTATGTCTGTGAGGGAACGGAATAAAAAT | Tb927.5.2900_rev | GGGGACCACTTTGTACAAGAAAGCTGGGTTCACTTGATGATATTGGACTCCGCA |
| 5_known | L13 | Tb927.3.1340_for | GGGGACAAGTTTGTACAAAAAAGCAGGCTATGCCTCCCAGACGTTCCTT | Tb927.3.1340_rev | GGGGACCACTTTGTACAAGAAAGCTGGGTTTACTCCTTTTCTTCCACCTGGGA |
| 5_known | L15 | Tb927.11.12150_for | GGGGACAAGTTTGTACAAAAAAGCAGGCTATGGCGTTTCTCGTACAAGTAGC | Tb927.11.12150_rev | GGGGACCACTTTGTACAAGAAAGCTGGGTTTAATCTCGCGGATAGGACCTCC |
| 5_known | L16 | Tb927.11.5650_for | GGGGACAAGTTTGTACAAAAAAGCAGGCTATGATGACCAGCAGTCCGAGT | Tb927.11.5650_rev | GGGGACCACTTTGTACAAGAAAGCTGGGTCTAATAGTCATCGGAACTGTAGCTCTCA |
| 5_known | L17 | Tb927.10.8210_for | GGGGACAAGTTTGTACAAAAAAGCAGGCTATGTACCGCAGGTTCTTTCGC | Tb927.10.8210_rev | GGGGACCACTTTGTACAAGAAAGCTGGGTCTATGTGATCACAGCTAGAGTCCCT |
| 5_known | L18 | Tb927.10.15650_for | GGGGACAAGTTTGTACAAAAAAGCAGGCTATGTTTGTTACCTCGCAACTAACACA | Tb927.10.15650_rev | GGGGACCACTTTGTACAAGAAAGCTGGGTTTATATGTCACGGGCGATGTTCAC |
| 5_known | L19 | Tb927.8.2910_for | GGGGACAAGTTTGTACAAAAAAGCAGGCTATGCCGTGGGTTCGTGCT | Tb927.8.2910_rev | GGGGACCACTTTGTACAAGAAAGCTGGGTTTATCGAAACACAGACTTCGACAGC |
| 5_known | L20 | Tb927.10.13960_for | GGGGACAAGTTTGTACAAAAAAGCAGGCTATGGCGCAGGTCGGCGAG | Tb927.10.13960_rev | GGGGACCACTTTGTACAAGAAAGCTGGGTTCAGCACCAGTACCTCACCAC |
| 5_known | L21 | Tb927.10.5560_for | GGGGACAAGTTTGTACAAAAAAGCAGGCTATGAAGCGAGCACCTTGTTTGT | Tb927.10.5560_rev | GGGGACCACTTTGTACAAGAAAGCTGGGTTCACTTCGTCACCTTGACCCA |
| 5_known | L22 | Tb927.3.4310_for | GGGGACAAGTTTGTACAAAAAAGCAGGCTATGGCCGCAGTTGACGATG | Tb927.3.4310_rev | GGGGACCACTTTGTACAAGAAAGCTGGGTCTATTCGAGGCGTGCCGG |
| 5_known | M1 | Tb927.10.6780_for | GGGGACAAGTTTGTACAAAAAAGCAGGCTATGTCGACCGCGCCTCCC | Tb927.10.6780_rev | GGGGACCACTTTGTACAAGAAAGCTGGGTTCATGCGTGCGCTGCTAG |
| 5_known | M2 | Tb927.1.1240_for | GGGGACAAGTTTGTACAAAAAAGCAGGCTATGGAAGGCAGTGCAACTGC | Tb927.1.1240_rev | GGGGACCACTTTGTACAAGAAAGCTGGGTTTACGCCTGTTTCAACCGGC |
| 5_known | M5 | Tb927.5.1020_for | GGGGACAAGTTTGTACAAAAAAGCAGGCTATGCAACGAACGAAAAGTTTGCTT | Tb927.5.1020_rev | GGGGACCACTTTGTACAAGAAAGCTGGGTTTAGTTCCTCTTCTTTTCAACTGCAGAA |
| 5_known | M6 | Tb927.9.15810_for | GGGGACAAGTTTGTACAAAAAAGCAGGCTATGCCAAGGGCAAACAGACC | Tb927.9.15810_rev | GGGGACCACTTTGTACAAGAAAGCTGGGTTTAATTCATCATCATATTTCTAATCATCAC |
| 5_known | M7 | Tb927.7.5950_for | GGGGACAAGTTTGTACAAAAAAGCAGGCTATGAGCGCACCCGTCGAC | Tb927.7.5950_rev | GGGGACCACTTTGTACAAGAAAGCTGGGTTTAAATTACACCTATCGGTTCCGTAGAAAC |
| 5_known | M8 | Tb927.8.4110_for | GGGGACAAGTTTGTACAAAAAAGCAGGCTATGGGTGGCAGGACTGAATCA | Tb927.8.4110_rev | GGGGACCACTTTGTACAAGAAAGCTGGGTCTACTCCAGCGCCGGCAG |
| 5_known | M10 | Tb927.9.12960_for | GGGGACAAGTTTGTACAAAAAAGCAGGCTATGCAACAAAAATCGGACGCCG | Tb927.9.12960_rev | GGGGACCACTTTGTACAAGAAAGCTGGGTTCACCCCTTAATACCTGTGAAAATTCC |
| 5_known | M11 | Tb927.10.9720_for | GGGGACAAGTTTGTACAAAAAAGCAGGCTATGTATCACTACGAGTACGATGACGA | Tb927.10.9720_rev | GGGGACCACTTTGTACAAGAAAGCTGGGTTCAACCGACAGTGGCGGC |
| 5_known | M12 | Tb927.10.2410_for | GGGGACAAGTTTGTACAAAAAAGCAGGCTATGACCCAACTGTTAGGAACCGC | Tb927.10.2410_rev | GGGGACCACTTTGTACAAGAAAGCTGGGTTCACATTTGACGAGGGTTCACG |
| 5_known | M13 | Tb927.3.3390_for | GGGGACAAGTTTGTACAAAAAAGCAGGCTATGCATGACTCTGTGGGAACGG | Tb927.3.3390_rev | GGGGACCACTTTGTACAAGAAAGCTGGGTTCACTCCATGCCGAAGGCG |
| 5_known | M14 | Tb927.8.1650_for | GGGGACAAGTTTGTACAAAAAAGCAGGCTATGAACTGCGCGGCGGAC | Tb927.8.1650_rev | GGGGACCACTTTGTACAAGAAAGCTGGGTTTAGTTGGGAAGTTCCGCCGT |
| 5_known | M15 | Tb927.4.2190_for | GGGGACAAGTTTGTACAAAAAAGCAGGCTATGGAAACTAAAGAGAAATCACCAATGTTG | Tb927.4.2190_rev | GGGGACCACTTTGTACAAGAAAGCTGGGTCTACACATTTGCAAATACCGATTGCA |
| 5_known | M16 | Tb927.8.1670_for | GGGGACAAGTTTGTACAAAAAAGCAGGCTATGGGGATGGATCACTACACCG | Tb927.8.1670_rev | GGGGACCACTTTGTACAAGAAAGCTGGGTTCACTGCACAGCAGGTGCG |
| 5_known | M17 | Tb927.11.13500_for | GGGGACAAGTTTGTACAAAAAAGCAGGCTATGATCGAGGTGCAACTTAACAGTT | Tb927.11.13500_rev | GGGGACCACTTTGTACAAGAAAGCTGGGTCTACGCACGACCGACGGG |
| 5_known | M18 | Tb927.3.3940_for | GGGGACAAGTTTGTACAAAAAAGCAGGCTATGGAAGTTCCTTTTGAAGAAGGGAC | Tb927.3.3940_rev | GGGGACCACTTTGTACAAGAAAGCTGGGTTTACAAAAACGGCACAAATGAGCC |
| 5_known | M19 | Tb927.10.13740_for | GGGGACAAGTTTGTACAAAAAAGCAGGCTATGGAAAAGGCAAATGAACTGATGC | Tb927.10.13740_rev | GGGGACCACTTTGTACAAGAAAGCTGGGTTTAGTGGCGCAGAAGCTTAAGTTC |
| 5_known | M20 | Tb927.11.15040_for | GGGGACAAGTTTGTACAAAAAAGCAGGCTATGCTGAGGCGCTCCTTCG | Tb927.11.15040_rev | GGGGACCACTTTGTACAAGAAAGCTGGGTCTAAATACCCTTCATTTGAAACTTCATCGG |
| 5_known | M21 | Tb927.9.6710_for | GGGGACAAGTTTGTACAAAAAAGCAGGCTATGTGGCGGCGCTCTCGG | Tb927.9.6710_rev | GGGGACCACTTTGTACAAGAAAGCTGGGTCTACGTATATACCCTGCCAATGTGATG |
| 5_known | M22 | Tb927.6.4210_for | GGGGACAAGTTTGTACAAAAAAGCAGGCTATGGAAGGTGTATCTGCCTTACTGG | Tb927.6.4210_rev | GGGGACCACTTTGTACAAGAAAGCTGGGTTCAAGAATCCTTCTTGCGTGGAGC |
| 5_known | M23 | Tb927.10.5320_for | GGGGACAAGTTTGTACAAAAAAGCAGGCTATGCGGAACTCGCTCATTCAG | Tb927.10.5320_rev | GGGGACCACTTTGTACAAGAAAGCTGGGTTCAGCCGGCCGATTGTGC |
| 5_known | M24 | Tb927.7.5930_for | GGGGACAAGTTTGTACAAAAAAGCAGGCTATGAGCGCACCCGTCGAC | Tb927.7.5930_rev | GGGGACCACTTTGTACAAGAAAGCTGGGTTCATTGCGGAGCAGCCTCA |
| 5_known | N1 | Tb927.10.1720_for | GGGGACAAGTTTGTACAAAAAAGCAGGCTATGAACATCTACAGTTGGGAGAGCT | Tb927.10.1720_rev | GGGGACCACTTTGTACAAGAAAGCTGGGTTTAATTTTGTGGTGTGACTTCCTTCAATG |
| 5_known | N2 | Tb927.7.3580_for | GGGGACAAGTTTGTACAAAAAAGCAGGCTATGGATAGGTTTACTAAGGTCCGTAATATT | Tb927.7.3580_rev | GGGGACCACTTTGTACAAGAAAGCTGGGTTCAAGTAGCATTCCGCCTTGC |
| 5_known | N3 | Tb927.8.2920_for | GGGGACAAGTTTGTACAAAAAAGCAGGCTATGAAAGGCGCTCAGTTTTTGAAGA | Tb927.8.2920_rev | GGGGACCACTTTGTACAAGAAAGCTGGGTTTATCGAAACACAGACTTCGACAGC |
| 5_known | N4 | Tb927.9.10670_for | GGGGACAAGTTTGTACAAAAAAGCAGGCTATGCGTCGGGAGCAGTTAGG | Tb927.9.10670_rev | GGGGACCACTTTGTACAAGAAAGCTGGGTCTATGAATTGTCTTCTATGAATCCAGCGT |
| 5_known | N6 | Tb927.3.2280_for | GGGGACAAGTTTGTACAAAAAAGCAGGCTATGAACCCTTTATGCGAAATCATTAAGC | Tb927.3.2280_rev | GGGGACCACTTTGTACAAGAAAGCTGGGTCTACTCTTCCAAAGGATAGGGCAGT |
| 5_known | N7 | Tb927.11.11670_for | GGGGACAAGTTTGTACAAAAAAGCAGGCTATGGACTTCAAACGACAGCTGG | Tb927.11.11670_rev | GGGGACCACTTTGTACAAGAAAGCTGGGTCTACTCGCCGCTTCTTTTTGAGC |
| 5_known | N8 | Tb927.7.5980_for | GGGGACAAGTTTGTACAAAAAAGCAGGCTATGAGCGCACCCGTCGAC | Tb927.7.5980_rev | GGGGACCACTTTGTACAAGAAAGCTGGGTTCACTTGAGTTGATGCGCAGG |
| 5_known | N9 | Tb927.9.3800_for | GGGGACAAGTTTGTACAAAAAAGCAGGCTATGACCTGCGTAAAAGAGTTGTTGC | Tb927.9.3800_rev | GGGGACCACTTTGTACAAGAAAGCTGGGTCTATGTGCTAGCACCAGCCAC |
| 5_known | N10 | Tb927.10.5590_for | GGGGACAAGTTTGTACAAAAAAGCAGGCTATGTCCGGTCAGGGAAATTTGC | Tb927.10.5590_rev | GGGGACCACTTTGTACAAGAAAGCTGGGTTCAAACCTTCCTTTCCGGCAAAC |
| 5_known | N12 | Tb927.7.5960_for | GGGGACAAGTTTGTACAAAAAAGCAGGCTATGAGCGCACCCGTCGAC | Tb927.7.5960_rev | GGGGACCACTTTGTACAAGAAAGCTGGGTTCACTTGAGTTGATGCACAGGTTC |
| 5_known | N13 | Tb927.11.7450_for | GGGGACAAGTTTGTACAAAAAAGCAGGCTATGAGTAGGGATCGTGGTGGTATAG | Tb927.11.7450_rev | GGGGACCACTTTGTACAAGAAAGCTGGGTCTAAAAACGCCGCCTGCCT |
| 5_known | N14 | Tb927.10.4300_for | GGGGACAAGTTTGTACAAAAAAGCAGGCTATGGCCAGTCAAGGAGAAACGG | Tb927.10.4300_rev | GGGGACCACTTTGTACAAGAAAGCTGGGTTCACGGCAGCGAATTGAAACC |
| 5_known | N15 | Tb927.7.5380_for | GGGGACAAGTTTGTACAAAAAAGCAGGCTATGAGCTTATTTTCTTTTACGCCAACTG | Tb927.7.5380_rev | GGGGACCACTTTGTACAAGAAAGCTGGGTTTACGTAAGTGCAGCAAATGTCCC |
| 5_known | N16 | Tb927.11.360_for | GGGGACAAGTTTGTACAAAAAAGCAGGCTATGAAGACTATTGCCCTCATCTCCG | Tb927.11.360_rev | GGGGACCACTTTGTACAAGAAAGCTGGGTTCAAGTACCTTCCCGCTCATCTG |
| 5_known | N17 | Tb927.8.4990_for | GGGGACAAGTTTGTACAAAAAAGCAGGCTATGAGCGGAAAGGAAGTTGAAGG | Tb927.8.4990_rev | GGGGACCACTTTGTACAAGAAAGCTGGGTCTACTGAGTGATCTGCGGCATC |
| 5_known | N21 | Tb927.10.2230_for | GGGGACAAGTTTGTACAAAAAAGCAGGCTATGCTCTTGCGGCGGAGC | Tb927.10.2230_rev | GGGGACCACTTTGTACAAGAAAGCTGGGTTCACAGCTGTGATGGATGGAGC |
| 5_known | N23 | Tb927.11.4980_for | GGGGACAAGTTTGTACAAAAAAGCAGGCTATGAGTGGGTCACCGAGTGAC | Tb927.11.4980_rev | GGGGACCACTTTGTACAAGAAAGCTGGGTTTACAACTTGCGCCGCTTCA |

| | | | | | |
|---|---|---|---|---|---|
| 5_known | N24 | Tb927.7.7450_for | GGGGACAAGTTTGTACAAAAAAGCAGGCTATGGGGAAGCCTGGTAAAAAGGC | Tb927.7.7450_rev | GGGGACCACTTTGTACAAGAAAGCTGGGTTCAGTCCTGAGCTGCAAGAAGG |
| 5_known | O1 | Tb927.11.3650_for | GGGGACAAGTTTGTACAAAAAAGCAGGCTATGGCCGCGGCTCCAAGT | Tb927.11.3650_rev | GGGGACCACTTTGTACAAGAAAGCTGGGTTTATTGCTTGAGGTATATAATGTCTGAACC |
| 5_known | O2 | Tb927.10.15530_for | GGGGACAAGTTTGTACAAAAAAGCAGGCTATGGCCAAACCGAAGTACCTAAAAA | Tb927.10.15530_rev | GGGGACCACTTTGTACAAGAAAGCTGGGTTCACTTCCGGTTGGCGTAGT |
| 5_known | O3 | Tb927.8.980_for | GGGGACAAGTTTGTACAAAAAAGCAGGCTATGGTGCTGCAGGCTGCG | Tb927.8.980_rev | GGGGACCACTTTGTACAAGAAAGCTGGGTCTACGCTCCACCGCAGAAAC |
| 5_known | O4 | Tb927.11.14300_for | GGGGACAAGTTTGTACAAAAAAGCAGGCTATGACGGATGTGACGTACAGTTACT | Tb927.11.14300_rev | GGGGACCACTTTGTACAAGAAAGCTGGGTTCAATCCGAATCAACATCTATTCCCCC |
| 5_known | O5 | Tb927.4.3420_for | GGGGACAAGTTTGTACAAAAAAGCAGGCTATGTTGGCGCCTAAAGGGGG | Tb927.4.3420_rev | GGGGACCACTTTGTACAAGAAAGCTGGGTCTAAAATTTTGCACTATCTCCCCTATCCC |
| 5_known | O6 | Tb927.8.3800_for | GGGGACAAGTTTGTACAAAAAAGCAGGCTATGTTGCACTCAGACCACCCC | Tb927.8.3800_rev | GGGGACCACTTTGTACAAGAAAGCTGGGTTCACACACAGTGCTCCTCTAAGC |
| 5_known | O7 | Tb927.11.7380_for | GGGGACAAGTTTGTACAAAAAAGCAGGCTATGTCTCGTATTCCTGAGAAGATTCTCT | Tb927.11.7380_rev | GGGGACCACTTTGTACAAGAAAGCTGGGTTTATTGGCCGTGAAATGATTCCAGC |
| 5_known | O8 | Tb927.2.4020_for | GGGGACAAGTTTGTACAAAAAAGCAGGCTATGAGTAATGAAAAATATGACAGGCAGCT | Tb927.2.4020_rev | GGGGACCACTTTGTACAAGAAAGCTGGGTTTATTGAGCCTCTGCAACCCAGA |
| 5_known | O9 | Tb927.10.6260_for | GGGGACAAGTTTGTACAAAAAAGCAGGCTATGGGTGTGGATGACGTAGGC | Tb927.10.6260_rev | GGGGACCACTTTGTACAAGAAAGCTGGGTCTACCACTCACCACTGATGTTCTCC |
| 5_known | O10 | Tb927.11.11530_for | GGGGACAAGTTTGTACAAAAAAGCAGGCTATGGCGTCGTTAGCAGCAAA | Tb927.11.11530_rev | GGGGACCACTTTGTACAAGAAAGCTGGGTTCAGCTCTGGTGCTGATTGTGC |
| 5_known | O11 | Tb927.3.2470_for | GGGGACAAGTTTGTACAAAAAAGCAGGCTATGGGTAAAACTAACACAAAAAAGCAGG | Tb927.3.2470_rev | GGGGACCACTTTGTACAAGAAAGCTGGGTTTATTTCTTGGGGAGAACCTTCTTTTTCT |
| 5_known | O12 | Tb927.3.1920_for | GGGGACAAGTTTGTACAAAAAAGCAGGCTATGACAAATAATAAAAAGAGTCAACAGGAG | Tb927.3.1920_rev | GGGGACCACTTTGTACAAGAAAGCTGGGTTCAACGCAGCTCATTCTCGAG |
| 5_known | O13 | Tb927.7.1370_for | GGGGACAAGTTTGTACAAAAAAGCAGGCTATGAGAAGTAGTGCTAATTTTTTTTTAGCG | Tb927.7.1370_rev | GGGGACCACTTTGTACAAGAAAGCTGGGTTCAAAAAATAACTTCTTCCTCCGTAGCC |
| 5_known | O14 | Tb927.7.5940_for | GGGGACAAGTTTGTACAAAAAAGCAGGCTATGAGCGCACCCGTCGAC | Tb927.7.5940_rev | GGGGACCACTTTGTACAAGAAAGCTGGGTTTACTTGCTTGAATCAGCAGCTTTCT |
| 5_known | O15 | Tb927.10.10350_for | GGGGACAAGTTTGTACAAAAAAGCAGGCTATGAGTACACAAGAATCTCATGGTTCTG | Tb927.10.10350_rev | GGGGACCACTTTGTACAAGAAAGCTGGGTTCAAAACGTAAGAAATGGATGTTTCAAAAG |
| 5_known | O16 | Tb927.8.6660_for | GGGGACAAGTTTGTACAAAAAAGCAGGCTATGACGACGATGCAGAGGGT | Tb927.8.6660_rev | GGGGACCACTTTGTACAAGAAAGCTGGGTCTAAGGTGCGAAGGCTGCC |
| 5_known | O17 | Tb927.11.3470_for | GGGGACAAGTTTGTACAAAAAAGCAGGCTATGGGCAAGCAGGGAAATGTTG | Tb927.11.3470_rev | GGGGACCACTTTGTACAAGAAAGCTGGGTTTAATCAAGCTTCACCGAAGTGTATATCTT |
| 5_known | O18 | Tb927.11.7630_for | GGGGACAAGTTTGTACAAAAAAGCAGGCTATGACAGTGATTATGTTCCCTCGTTATG | Tb927.11.7630_rev | GGGGACCACTTTGTACAAGAAAGCTGGGTTTATAATAGGGAAAAGCAAGAAACGAAGAG |
| 5_known | O19 | Tb927.11.7640_for | GGGGACAAGTTTGTACAAAAAAGCAGGCTATGACGCTGATTATGTTCCCCCG | Tb927.11.7640_rev | GGGGACCACTTTGTACAAGAAAGCTGGGTTCATAATAGGGGAAAGCAAGAAACGAAG |
| 5_known | O20 | Tb927.6.4590_for | GGGGACAAGTTTGTACAAAAAAGCAGGCTATGCTCCGGTTAGCTAACAAGTTTAC | Tb927.6.4590_rev | GGGGACCACTTTGTACAAGAAAGCTGGGTCTAGACAGCTTGTTCTTTGTTCTGACT |
| 5_known | O21 | Tb927.11.11780_for | GGGGACAAGTTTGTACAAAAAAGCAGGCTATGTTTCGTCGTAGCCTTTCCC | Tb927.11.11780_rev | GGGGACCACTTTGTACAAGAAAGCTGGGTTCACCTAAGATCCCATGTCGACTC |
| 5_known | O22 | Tb927.1.3830_for | GGGGACAAGTTTGTACAAAAAAGCAGGCTATGAGCAATTACTTGGATGATTTGCG | Tb927.1.3830_rev | GGGGACCACTTTGTACAAGAAAGCTGGGTTCAAAGGTGCGACAATTCATTAAACATATT |
| 5_known | O23 | Tb927.5.4070_for | GGGGACAAGTTTGTACAAAAAAGCAGGCTATGCTCTGGTGTAGGGCAACG | Tb927.5.4070_rev | GGGGACCACTTTGTACAAGAAAGCTGGGTCTAAGCGTCCTGCGACGG |
| 5_known | O24 | Tb927.11.6990_for | GGGGACAAGTTTGTACAAAAAAGCAGGCTATGAACACCTTCAAGGGTGATGGG | Tb927.11.6990_rev | GGGGACCACTTTGTACAAGAAAGCTGGGTTCATGCAATCGTGGGGTTCCA |
| 5_known | P1 | Tb927.11.470_for | GGGGACAAGTTTGTACAAAAAAGCAGGCTATGTCTAGTTGTTCGCATCCTACCT | Tb927.11.470_rev | GGGGACCACTTTGTACAAGAAAGCTGGGTCTAAGCGACCCATTCATCCTCCG |
| 5_known | P2 | Tb927.10.15910_for | GGGGACAAGTTTGTACAAAAAAGCAGGCTATGTCAGACGACGAGCACAAAC | Tb927.10.15910_rev | GGGGACCACTTTGTACAAGAAAGCTGGGTTCAACGAACTATACCGTACCTTTCCT |
| 5_known | P3 | Tb927.10.960_for | GGGGACAAGTTTGTACAAAAAAGCAGGCTATGGAGTTAATTGATCCCGTAGTCCC | Tb927.10.960_rev | GGGGACCACTTTGTACAAGAAAGCTGGGTTTACTTTCTCTTCAGTAGTTGCATTTCATC |
| 5_known | P4 | Tb927.8.6580_for | GGGGACAAGTTTGTACAAAAAAGCAGGCTATGCTACGGCGTTCTTTTAACCG | Tb927.8.6580_rev | GGGGACCACTTTGTACAAGAAAGCTGGGTCTAGTAGACGCGCTTCGCTG |
| 5_known | P5 | Tb927.4.1950_for | GGGGACAAGTTTGTACAAAAAAGCAGGCTATGGAGGAAAATGAAGGGATTGGAAC | Tb927.4.1950_rev | GGGGACCACTTTGTACAAGAAAGCTGGGTCTACACAGACCATGTATCCACCACA |
| 5_known | P6 | Tb927.9.14550_for | GGGGACAAGTTTGTACAAAAAAGCAGGCTATGGTGTGGAAGGGTCAAGCC | Tb927.9.14550_rev | GGGGACCACTTTGTACAAGAAAGCTGGGTTTATTTAGCTCTCCACCAATCAAATAATGA |
| 5_known | P7 | Tb927.10.13210_for | GGGGACAAGTTTGTACAAAAAAGCAGGCTATGTTTAGCTACGATGATCTTGTGTGG | Tb927.10.13210_rev | GGGGACCACTTTGTACAAGAAAGCTGGGTTCAGAGTAAGTTCGCTAACAAGGGC |
| 5_known | P8 | Tb927.4.1080_for | GGGGACAAGTTTGTACAAAAAAGCAGGCTATGTCGAGCGACCGCAAC | Tb927.4.1080_rev | GGGGACCACTTTGTACAAGAAAGCTGGGTCTACTGCACGAGGGAAGCGA |
| 5_known | P9 | Tb927.11.6630_for | GGGGACAAGTTTGTACAAAAAAGCAGGCTATGAAGTCGTTTTGCCGGTTGG | Tb927.11.6630_rev | GGGGACCACTTTGTACAAGAAAGCTGGGTTCACATACGAAAAACCCCAAACTTTGT |
| 5_known | P10 | Tb927.3.4490_for | GGGGACAAGTTTGTACAAAAAAGCAGGCTATGAATAAGAGCGCGGTGCG | Tb927.3.4490_rev | GGGGACCACTTTGTACAAGAAAGCTGGGTTCAAACATAGTCCCGGTTCATAACG |
| 5_known | P11 | Tb927.6.140_for | GGGGACAAGTTTGTACAAAAAAGCAGGCTATGGACAACGAAACTCAACTGCT | Tb927.6.140_rev | GGGGACCACTTTGTACAAGAAAGCTGGGTTTACGTTCGGCCCGCAGC |
| 5_known | P12 | Tb927.7.7280_for | GGGGACAAGTTTGTACAAAAAAGCAGGCTATGCCAAACAGTGATCACAGCTAC | Tb927.7.7280_rev | GGGGACCACTTTGTACAAGAAAGCTGGGTCTACGTTGTTTCAATCGTCTGCG |
| 5_known | P13 | Tb927.6.1500_for | GGGGACAAGTTTGTACAAAAAAGCAGGCTATGGATAAGAGAATGATTACTGATGCATTT | Tb927.6.1500_rev | GGGGACCACTTTGTACAAGAAAGCTGGGTTCAGAGGTGAGCCTGAAGAGC |
| 5_known | P14 | Tb927.10.4480_for | GGGGACAAGTTTGTACAAAAAAGCAGGCTATGGAGACGAATGGACCGAAGA | Tb927.10.4480_rev | GGGGACCACTTTGTACAAGAAAGCTGGGTCTATGTAAAAGCAACAGGGTGAGGC |
| 5_known | P15 | Tb927.8.6740_for | GGGGACAAGTTTGTACAAAAAAGCAGGCTATGCCGAACAGCCATAACGAAA | Tb927.8.6740_rev | GGGGACCACTTTGTACAAGAAAGCTGGGTCTAGCAGTAACTCCTCGTCAATATAAATTT |
| 5_known | P16 | Tb927.1.670_for | GGGGACAAGTTTGTACAAAAAAGCAGGCTATGATGCAAGTCAAACTGAAGCATTTT | Tb927.1.670_rev | GGGGACCACTTTGTACAAGAAAGCTGGGTTCATGCGGCAGCTCGACA |
| 5_known | P17 | Tb927.8.1720_for | GGGGACAAGTTTGTACAAAAAAGCAGGCTATGCTCTTTCTTCTCGCTTACGC | Tb927.8.1720_rev | GGGGACCACTTTGTACAAGAAAGCTGGGTCTACAAAAGGTCCTGCCCCATTT |
| 5_known | P18 | Tb927.11.7620_for | GGGGACAAGTTTGTACAAAAAAGCAGGCTATGACAGTGATTATGTTCCCTCGTTATG | Tb927.11.7620_rev | GGGGACCACTTTGTACAAGAAAGCTGGGTTTATAGAGAAAAAAAAGAAACGAAGAGTAG |
| 5_known | P19 | Tb927.7.3210_for | GGGGACAAGTTTGTACAAAAAAGCAGGCTATGGCGACCTCTCGCAAGC | Tb927.7.3210_rev | GGGGACCACTTTGTACAAGAAAGCTGGGTTTAATCAGAGTCCGCAACAGTTTTAATGG |
| 5_known | P20 | Tb927.3.2090_for | GGGGACAAGTTTGTACAAAAAAGCAGGCTATGTCGACCAACAGTGCCAAG | Tb927.3.2090_rev | GGGGACCACTTTGTACAAGAAAGCTGGGTTCAACAACCGCCCTCAAGC |
| 5_known | P21 | Tb927.8.1870_for | GGGGACAAGTTTGTACAAAAAAGCAGGCTATGCGGCGTTCTTTCTTTGTTG | Tb927.8.1870_rev | GGGGACCACTTTGTACAAGAAAGCTGGGTCTAATCCTCCACCTCAATGACGGA |
| 5_known | P22 | Tb927.1.1130_for | GGGGACAAGTTTGTACAAAAAAGCAGGCTATGGGTCGCTATACGCGCC | Tb927.1.1130_rev | GGGGACCACTTTGTACAAGAAAGCTGGGTTCATGTGAGAGCCGTGGCG |
| 5_known | P23 | Tb927.11.10760_for | GGGGACAAGTTTGTACAAAAAAGCAGGCTATGCAAGCCGACACATTTGCT | Tb927.11.10760_rev | GGGGACCACTTTGTACAAGAAAGCTGGGTTTACATGGTAACACACTGGCATCC |
| 5_known | P24 | Tb927.10.10810_for | GGGGACAAGTTTGTACAAAAAAGCAGGCTATGCATGTGGATGTGCCTCAC | Tb927.10.10810_rev | GGGGACCACTTTGTACAAGAAAGCTGGGTTCATGCAACGCGTTGCTCC |
| 6_hypothetical | A1 | Tb927.10.14740_for | GGGGACAAGTTTGTACAAAAAAGCAGGCTATGATACCTCGAGCAGCGCA | Tb927.10.14740_rev | GGGGACCACTTTGTACAAGAAAGCTGGGTCTATTCACCGTTCCGCCGTC |
| 6_hypothetical | A3 | Tb927.10.7900_for | GGGGACAAGTTTGTACAAAAAAGCAGGCTATGCTCGGTAACTCACAAGGCG | Tb927.10.7900_rev | GGGGACCACTTTGTACAAGAAAGCTGGGTCTACAAGTCAAACAGCATTTCCTCCT |
| 6_hypothetical | A4 | Tb927.11.2580_for | GGGGACAAGTTTGTACAAAAAAGCAGGCTATGACGTCGTTCAAGGACAATATTTCA | Tb927.11.2580_rev | GGGGACCACTTTGTACAAGAAAGCTGGGTTCATACACGCGTTAAGACGAGGC |
| 6_hypothetical | A5 | Tb927.9.5020_for | GGGGACAAGTTTGTACAAAAAAGCAGGCTATGAGGCGGTGCTGTTGTG | Tb927.9.5020_rev | GGGGACCACTTTGTACAAGAAAGCTGGGTTCATTCTTTCCCTTTTTTCTTGCTTGTAG |
| 6_hypothetical | A6 | Tb927.8.6960_for | GGGGACAAGTTTGTACAAAAAAGCAGGCTATGTACGCTGTGAAACCGTGG | Tb927.8.6960_rev | GGGGACCACTTTGTACAAGAAAGCTGGGTTTACTTTGGCAATTGAGAGAACGTGC |
| 6_hypothetical | A7 | Tb927.11.4840_for | GGGGACAAGTTTGTACAAAAAAGCAGGCTATGATTGTTAGGGCGAAGTCCCT | Tb927.11.4840_rev | GGGGACCACTTTGTACAAGAAAGCTGGGTCTACTGCTGGCGGCTGGA |
| 6_hypothetical | A8 | Tb927.6.1540_for | GGGGACAAGTTTGTACAAAAAAGCAGGCTATGGCAGCCAGTAGAAGTGATGA | Tb927.6.1540_rev | GGGGACCACTTTGTACAAGAAAGCTGGGTCTAAAAGCTGTAGTTAAGGAAGTAAAACTT |
| 6_hypothetical | A9 | Tb927.8.3850_for | GGGGACAAGTTTGTACAAAAAAGCAGGCTATGATAGTCAGAACGCCTCTTGCG | Tb927.8.3850_rev | GGGGACCACTTTGTACAAGAAAGCTGGGTTCAAGCGGCTACCGCGCC |
| 6_hypothetical | A10 | Tb927.9.14200_for | GGGGACAAGTTTGTACAAAAAAGCAGGCTATGTTCCGGCGGTTTATTCCATG | Tb927.9.14200_rev | GGGGACCACTTTGTACAAGAAAGCTGGGTCTATTCCCTGGGTTTTTGGACGTC |
| 6_hypothetical | A11 | Tb927.10.16110_for | GGGGACAAGTTTGTACAAAAAAGCAGGCTATGTCTTACTTGACAGAGGAGGAGG | Tb927.10.16110_rev | GGGGACCACTTTGTACAAGAAAGCTGGGTTCATTGTACGAGCGACAACTTGG |
| 6_hypothetical | A12 | Tb927.5.4150_for | GGGGACAAGTTTGTACAAAAAAGCAGGCTATGATAACACTAAAGACTGCTGTAAACACC | Tb927.5.4150_rev | GGGGACCACTTTGTACAAGAAAGCTGGGTCTAAAAAGTTTCCTTGGTTTCTTATCCAC |
| 6_hypothetical | A13 | Tb927.2.4670_for | GGGGACAAGTTTGTACAAAAAAGCAGGCTATGGAGTCAAACAAGAGAGTAACTGGG | Tb927.2.4670_rev | GGGGACCACTTTGTACAAGAAAGCTGGGTCTACACGTCAGCGATGGCTTC |
| 6_hypothetical | A14 | Tb927.11.6800_for | GGGGACAAGTTTGTACAAAAAAGCAGGCTATGGCCGTTGCCACTTCG | Tb927.11.6800_rev | GGGGACCACTTTGTACAAGAAAGCTGGGTTCATGATCCGCGTGAAGTTGAAC |
| 6_hypothetical | A15 | Tb927.7.3060_for | GGGGACAAGTTTGTACAAAAAAGCAGGCTATGTTTTGTTTCTTCCAAAACCGCTG | Tb927.7.3060_rev | GGGGACCACTTTGTACAAGAAAGCTGGGTCTAAAGGGCGGCTTCAAGTCC |
| 6_hypothetical | A16 | Tb927.9.12000_for | GGGGACAAGTTTGTACAAAAAAGCAGGCTATGTCTTTCGGTAGGTTATTGAAGCAG | Tb927.9.12000_rev | GGGGACCACTTTGTACAAGAAAGCTGGGTTTATTTTGTTAGCAATCGCTCCACGA |
| 6_hypothetical | A17 | Tb927.6.2440_for | GGGGACAAGTTTGTACAAAAAAGCAGGCTATGCGCAAGAAGAGCGTGC | Tb927.6.2440_rev | GGGGACCACTTTGTACAAGAAAGCTGGGTCTACGCCGACGCATTCAACA |
| 6_hypothetical | A18 | Tb927.3.2700_for | GGGGACAAGTTTGTACAAAAAAGCAGGCTATGATCCTTTCAGAAGGCGTTCC | Tb927.3.2700_rev | GGGGACCACTTTGTACAAGAAAGCTGGGTCTAGGCGAATGCCTCCAAAATACC |
| 6_hypothetical | A19 | Tb927.3.4460_for | GGGGACAAGTTTGTACAAAAAAGCAGGCTATGATAGCTGAACAACAGAACCAGG | Tb927.3.4460_rev | GGGGACCACTTTGTACAAGAAAGCTGGGTTTACCAAAAGGGAACACTTTCATATCCC |
| 6_hypothetical | A20 | Tb927.2.2230_for | GGGGACAAGTTTGTACAAAAAAGCAGGCTATGCGGCGCGGTGGTGCG | Tb927.2.2230_rev | GGGGACCACTTTGTACAAGAAAGCTGGGTTTATTTACGTCGTCGGTTGTTGTGC |
| 6_hypothetical | A21 | Tb927.7.6720_for | GGGGACAAGTTTGTACAAAAAAGCAGGCTATGGGGTGTGGCGATTCGC | Tb927.7.6720_rev | GGGGACCACTTTGTACAAGAAAGCTGGGTTTAGCGACTAATGTGAACCACATGG |
| 6_hypothetical | A22 | Tb927.10.12020_for | GGGGACAAGTTTGTACAAAAAAGCAGGCTATGTCTGTGTGTCCTCCACCA | Tb927.10.12020_rev | GGGGACCACTTTGTACAAGAAAGCTGGGTTCAGGGTAAAAAGATGTTTCCCACG |
| 6_hypothetical | A23 | Tb927.10.11380_for | GGGGACAAGTTTGTACAAAAAAGCAGGCTATGGATTTGTCCGGCCTAGTGC | Tb927.10.11380_rev | GGGGACCACTTTGTACAAGAAAGCTGGGTCTACCTTAACATCTGAACATCCCACTC |
| 6_hypothetical | A24 | Tb927.4.2240_for | GGGGACAAGTTTGTACAAAAAAGCAGGCTATGCAGACCGGCCCATTCG | Tb927.4.2240_rev | GGGGACCACTTTGTACAAGAAAGCTGGGTTTAAAGAAGAGCCTCCGCTTCATAGG |
| 6_hypothetical | B1 | Tb927.2.5130_for | GGGGACAAGTTTGTACAAAAAAGCAGGCTATGGTGGTTACTTTGGTGGAACC | Tb927.2.5130_rev | GGGGACCACTTTGTACAAGAAAGCTGGGTTTACCATTCACGGAATCTCTTGGAAAC |
| 6_hypothetical | B2 | Tb927.3.1140_for | GGGGACAAGTTTGTACAAAAAAGCAGGCTATGTCTGGTGAGGCCGATTCT | Tb927.3.1140_rev | GGGGACCACTTTGTACAAGAAAGCTGGGTTCACGCAGAGTTATTTCCGTCCA |
| 6_hypothetical | B3 | Tb927.10.15420_for | GGGGACAAGTTTGTACAAAAAAGCAGGCTATGCCATCTCCAATTGTGCTCAC | Tb927.10.15420_rev | GGGGACCACTTTGTACAAGAAAGCTGGGTCTAACTCTCCTTTTTCTGGAAGGGAGG |
| 6_hypothetical | B4 | Tb927.3.3370_for | GGGGACAAGTTTGTACAAAAAAGCAGGCTATGTCTGAGGTGGTACCTACTCGT | Tb927.3.3370_rev | GGGGACCACTTTGTACAAGAAAGCTGGGTTCAGCTTCGCTTTGAGTCATCC |
| 6_hypothetical | B5 | Tb927.8.3810_for | GGGGACAAGTTTGTACAAAAAAGCAGGCTATGAGAGCAAGTATGAATTGCTCCG | Tb927.8.3810_rev | GGGGACCACTTTGTACAAGAAAGCTGGGTCTACGGGTCTCCGAATGTGCG |
| 6_hypothetical | B6 | Tb927.11.3160_for | GGGGACAAGTTTGTACAAAAAAGCAGGCTATGTCCCGCCTGTTCCTCG | Tb927.11.3160_rev | GGGGACCACTTTGTACAAGAAAGCTGGGTTTAAAAGTCGGGGGAACAACCAG |
| 6_hypothetical | B7 | Tb927.11.520_for | GGGGACAAGTTTGTACAAAAAAGCAGGCTATGAGCAATGGTGGTGTTACTGC | Tb927.11.520_rev | GGGGACCACTTTGTACAAGAAAGCTGGGTTCATTGGAGGCCACTCCTCAA |
| 6_hypothetical | B8 | Tb927.6.1920_for | GGGGACAAGTTTGTACAAAAAAGCAGGCTATGTCCTACCGGAGGCCTACC | Tb927.6.1920_rev | GGGGACCACTTTGTACAAGAAAGCTGGGTTCAGTCCGCCCGACTTGT |
| 6_hypothetical | B9 | Tb927.10.4750_for | GGGGACAAGTTTGTACAAAAAAGCAGGCTATGGCGGTGCGAGCTCTG | Tb927.10.4750_rev | GGGGACCACTTTGTACAAGAAAGCTGGGTTTATTCCCATTCATTAGATGGATCGTAACG |
| 6_hypothetical | B10 | Tb927.7.5120_for | GGGGACAAGTTTGTACAAAAAAGCAGGCTATGCAGAGATGGAGCCAGAGAC | Tb927.7.5120_rev | GGGGACCACTTTGTACAAGAAAGCTGGGTTTAGTGATTCCTATACACCTCCACAGC |
| 6_hypothetical | B11 | Tb927.6.3860_for | GGGGACAAGTTTGTACAAAAAAGCAGGCTATGGATGTGTGTGGTTTGTCGC | Tb927.6.3860_rev | GGGGACCACTTTGTACAAGAAAGCTGGGTCTAACAGCGAACAACAGCATTGG |
| 6_hypothetical | B12 | Tb927.10.14660_for | GGGGACAAGTTTGTACAAAAAAGCAGGCTATGGAGGGCCTGTCGCCC | Tb927.10.14660_rev | GGGGACCACTTTGTACAAGAAAGCTGGGTTCATCTGTCTTCAGTTCTCTGCTTGT |
| 6_hypothetical | B13 | Tb927.5.850_for | GGGGACAAGTTTGTACAAAAAAGCAGGCTATGAGCGCCCAGGTGCCC | Tb927.5.850_rev | GGGGACCACTTTGTACAAGAAAGCTGGGTTCAGTCAATGGAAATCGTGTGAAGC |
| 6_hypothetical | B14 | Tb927.7.3170_for | GGGGACAAGTTTGTACAAAAAAGCAGGCTATGATGAGCAGCGGGAACCA | Tb927.7.3170_rev | GGGGACCACTTTGTACAAGAAAGCTGGGTTCACGACAGAGGAAGTGCCA |
| 6_hypothetical | B15 | Tb927.11.7213_for | GGGGACAAGTTTGTACAAAAAAGCAGGCTATGATAGGCCTCCGGAAACGC | Tb927.11.7213_rev | GGGGACCACTTTGTACAAGAAAGCTGGGTTTACACAAGTCGTGTCAGACGC |
| 6_hypothetical | B16 | Tb927.7.2520_for | GGGGACAAGTTTGTACAAAAAAGCAGGCTATGGAGGAGAGTACTGAAAGATCACTT | Tb927.7.2520_rev | GGGGACCACTTTGTACAAGAAAGCTGGGTTCACGCCACTGCTTCCTCT |
| 6_hypothetical | B17 | Tb927.11.9840_for | GGGGACAAGTTTGTACAAAAAAGCAGGCTATGCTAGATTTTCTCGACACGATTAACA | Tb927.11.9840_rev | GGGGACCACTTTGTACAAGAAAGCTGGGTTCACTGACAGTCGGAATGGCT |
| 6_hypothetical | B18 | Tb927.8.6140_for | GGGGACAAGTTTGTACAAAAAAGCAGGCTATGGCTCTATGTAACAGTCCTTCACG | Tb927.8.6140_rev | GGGGACCACTTTGTACAAGAAAGCTGGGTTCATTCATGCTGAGAGTTGCTGTC |
| 6_hypothetical | B19 | Tb927.3.1450_for | GGGGACAAGTTTGTACAAAAAAGCAGGCTATGCCGTGTGCCACTAAAGG | Tb927.3.1450_rev | GGGGACCACTTTGTACAAGAAAGCTGGGTTTATCCACCCTTTTCCCTGATGAGG |
| 6_hypothetical | B20 | Tb927.10.4860_for | GGGGACAAGTTTGTACAAAAAAGCAGGCTATGAGCGGGATGCAGGCA | Tb927.10.4860_rev | GGGGACCACTTTGTACAAGAAAGCTGGGTTTATATGAGGCCGACAGTCTTCAAATATTG |
| 6_hypothetical | B21 | Tb927.7.2610_for | GGGGACAAGTTTGTACAAAAAAGCAGGCTATGGGTAGCGCTTCTCAGGT | Tb927.7.2610_rev | GGGGACCACTTTGTACAAGAAAGCTGGGTTTATCTCAAATCCATCTCTTCACCTGTG |
| 6_hypothetical | B22 | Tb927.11.12580_for | GGGGACAAGTTTGTACAAAAAAGCAGGCTATGGGTGCCAACAACATAAGGG | Tb927.11.12580_rev | GGGGACCACTTTGTACAAGAAAGCTGGGTCTAATCCGACTTATCGCTGCCTTC |
| 6_hypothetical | B24 | Tb927.5.410_for | GGGGACAAGTTTGTACAAAAAAGCAGGCTATGCATATGTATTTCATGCGTGTGAAG | Tb927.5.410_rev | GGGGACCACTTTGTACAAGAAAGCTGGGTTCACTCCCTCAGGTAACTTGGTG |
| 6_hypothetical | C1 | Tb927.5.309b_for | GGGGACAAGTTTGTACAAAAAAGCAGGCTATGGCACGTGTCACCTTTTGC | Tb927.5.309b_rev | GGGGACCACTTTGTACAAGAAAGCTGGGTCTACAGTTTGGGCTGCGCA |
| 6_hypothetical | C2 | Tb927.10.9950_for | GGGGACAAGTTTGTACAAAAAAGCAGGCTATGACGGAGCGTACCGACA | Tb927.10.9950_rev | GGGGACCACTTTGTACAAGAAAGCTGGGTTTAGCGAATGATCCAGTTGGCTAAC |
| 6_hypothetical | C3 | Tb927.8.6780_for | GGGGACAAGTTTGTACAAAAAAGCAGGCTATGTTGAGAGGGCGTGCCA | Tb927.8.6780_rev | GGGGACCACTTTGTACAAGAAAGCTGGGTCTAAAGATTAGATGACAACAGCGGCA |
| 6_hypothetical | C4 | Tb927.8.1900_for | GGGGACAAGTTTGTACAAAAAAGCAGGCTATGCACGAAGAGGACCTTATGAAGT | Tb927.8.1900_rev | GGGGACCACTTTGTACAAGAAAGCTGGGTCTACTTTTTAATCCCTGGAGCACAAGG |
| 6_hypothetical | C5 | Tb927.11.12440_for | GGGGACAAGTTTGTACAAAAAAGCAGGCTATGAGAGTAGAGGAGCAACATCAGT | Tb927.11.12440_rev | GGGGACCACTTTGTACAAGAAAGCTGGGTTCAAATTTCTTTGGGCTCAAACAGTTC |
| 6_hypothetical | C6 | Tb927.6.5000_for | GGGGACAAGTTTGTACAAAAAAGCAGGCTATGGAACAACAAACCATATCATTGCAC | Tb927.6.5000_rev | GGGGACCACTTTGTACAAGAAAGCTGGGTCTCACAAAAATTTGTTGATATCGCCCAC |
| 6_hypothetical | C7 | Tb927.9.12710_for | GGGGACAAGTTTGTACAAAAAAGCAGGCTATGCTCGGTGCAGGCACG | Tb927.9.12710_rev | GGGGACCACTTTGTACAAGAAAGCTGGGTTTACGTTACTCTCAGCATCTCCACG |
| 6_hypothetical | C8 | Tb927.11.12940_for | GGGGACAAGTTTGTACAAAAAAGCAGGCTATGTCTCGGGGGCATATGGC | Tb927.11.12940_rev | GGGGACCACTTTGTACAAGAAAGCTGGGTTCACACAGCCACGAGTGCC |
| 6_hypothetical | C9 | Tb927.3.4700_for | GGGGACAAGTTTGTACAAAAAAGCAGGCTATGTTGAGACAACTCCTCCACCA | Tb927.3.4700_rev | GGGGACCACTTTGTACAAGAAAGCTGGGTTCAGGACTTCCCTTTTCCGTGG |
| 6_hypothetical | C10 | Tb927.10.14450_for | GGGGACAAGTTTGTACAAAAAAGCAGGCTATGAGTACGGAGCCGCACA | Tb927.10.14450_rev | GGGGACCACTTTGTACAAGAAAGCTGGGTTCATCGAAACTTTTGTGACACCAATG |
| 6_hypothetical | C11 | Tb927.3.2500_for | GGGGACAAGTTTGTACAAAAAAGCAGGCTATGAGGAGTTACATCGTGTTTGTTGC | Tb927.3.2500_rev | GGGGACCACTTTGTACAAGAAAGCTGGGTTTAAAAAACTAGAAAAAATAACTTAAGTGG |
| 6_hypothetical | C12 | Tb927.11.3930_for | GGGGACAAGTTTGTACAAAAAAGCAGGCTATGGCGGGAACTTCTTTACCGT | Tb927.11.3930_rev | GGGGACCACTTTGTACAAGAAAGCTGGGTTTACGGCTCAAACGATGCTGC |
| 6_hypothetical | C13 | Tb927.7.620_for | GGGGACAAGTTTGTACAAAAAAGCAGGCTATGAGTGGTTCGGCCGGT | Tb927.7.620_rev | GGGGACCACTTTGTACAAGAAAGCTGGGTTCATCCTCCCATTGACGAAGTTGC |
| 6_hypothetical | C14 | Tb927.11.6450_for | GGGGACAAGTTTGTACAAAAAAGCAGGCTATGGGTTCGAGTGATGGTACTTCT | Tb927.11.6450_rev | GGGGACCACTTTGTACAAGAAAGCTGGGTTTACCAATCGTCCACAGCAATACC |
| 6_hypothetical | C15 | Tb927.10.520_for | GGGGACAAGTTTGTACAAAAAAGCAGGCTATGCAGGGCAGTTGGTCGG | Tb927.10.520_rev | GGGGACCACTTTGTACAAGAAAGCTGGGTCTAAGCTGTGTGTCGGCCC |
| 6_hypothetical | C16 | Tb927.10.6710_for | GGGGACAAGTTTGTACAAAAAAGCAGGCTATGGAAGCTGTTGTGTATGAGATCG | Tb927.10.6710_rev | GGGGACCACTTTGTACAAGAAAGCTGGGTTTAAGACCCGCAGTCTTCATACCA |
| 6_hypothetical | C17 | Tb927.6.2080_for | GGGGACAAGTTTGTACAAAAAAGCAGGCTATGTTTCGGTCGACCCGCC | Tb927.6.2080_rev | GGGGACCACTTTGTACAAGAAAGCTGGGTTCACGGGTCACCCGTGTAAC |
| 6_hypothetical | C18 | Tb927.11.8230_for | GGGGACAAGTTTGTACAAAAAAGCAGGCTATGGAGTTCTCTGGATCTACATTATCCG | Tb927.11.8230_rev | GGGGACCACTTTGTACAAGAAAGCTGGGTTTACTTCAACTGATCGGAAAACATTTCG |
| 6_hypothetical | C19 | Tb927.3.1100_for | GGGGACAAGTTTGTACAAAAAAGCAGGCTATGAGACGCGCTTGTGTACATT | Tb927.3.1100_rev | GGGGACCACTTTGTACAAGAAAGCTGGGTCTAAATGATGAACTCCTCTCCACGATC |
| 6_hypothetical | C20 | Tb927.3.1170_for | GGGGACAAGTTTGTACAAAAAAGCAGGCTATGTCTACCGATGGCAGTGAAGT | Tb927.3.1170_rev | GGGGACCACTTTGTACAAGAAAGCTGGGTTCATGCGTAGTTTCCTCTTGCAG |
| 6_hypothetical | C21 | Tb927.11.9050_for | GGGGACAAGTTTGTACAAAAAAGCAGGCTATGCCTAAGCGCCACCGT | Tb927.11.9050_rev | GGGGACCACTTTGTACAAGAAAGCTGGGTTCAGTAATGCAGCACACTGGC |
| 6_hypothetical | C22 | Tb927.6.4170_for | GGGGACAAGTTTGTACAAAAAAGCAGGCTATGCATCAGACGTTCGCATCG | Tb927.6.4170_rev | GGGGACCACTTTGTACAAGAAAGCTGGGTTCACGGCTCTCCTTCAGCA |
| 6_hypothetical | C23 | Tb927.4.3900_for | GGGGACAAGTTTGTACAAAAAAGCAGGCTATGGCGAGTAACTCCTCGGC | Tb927.4.3900_rev | GGGGACCACTTTGTACAAGAAAGCTGGGTTTATTTGGGCTTTTCACGTGATATTCTTTC |
| 6_hypothetical | C24 | Tb927.11.15790_for | GGGGACAAGTTTGTACAAAAAAGCAGGCTATGAAGGGCAAGGCAGTTAAAGAAC | Tb927.11.15790_rev | GGGGACCACTTTGTACAAGAAAGCTGGGTTCATTGTGTTACACTTGAGGCTGC |
| 6_hypothetical | D1 | Tb927.4.1280_for | GGGGACAAGTTTGTACAAAAAAGCAGGCTATGAGGCGCTTCATGTCCCT | Tb927.4.1280_rev | GGGGACCACTTTGTACAAGAAAGCTGGGTTTACCAACCGTCTTTAGCACGAG |
| 6_hypothetical | D2 | Tb927.8.5700_for | GGGGACAAGTTTGTACAAAAAAGCAGGCTATGATGTACCGGTGGAGCCC | Tb927.8.5700_rev | GGGGACCACTTTGTACAAGAAAGCTGGGTTCACGCTGCGCCTATTAATACC |
| 6_hypothetical | D3 | Tb927.11.1290_for | GGGGACAAGTTTGTACAAAAAAGCAGGCTATGAAATTGTCACAACGGTTCCTATCA | Tb927.11.1290_rev | GGGGACCACTTTGTACAAGAAAGCTGGGTCTAACGTGTTTGACGCACCAAAAG |
| 6_hypothetical | D4 | Tb927.6.4320_for | GGGGACAAGTTTGTACAAAAAAGCAGGCTATGGCGGACGTCGAGGCC | Tb927.6.4320_rev | GGGGACCACTTTGTACAAGAAAGCTGGGTTCAAGTGGACAAATAGTCCATAGCGT |
| 6_hypothetical | D5 | Tb927.1.4220_for | GGGGACAAGTTTGTACAAAAAAGCAGGCTATGCGAGCCGTCAAGAGGA | Tb927.1.4220_rev | GGGGACCACTTTGTACAAGAAAGCTGGGTCTATGTTTCTGCAGTCCCGCAT |
| 6_hypothetical | D6 | Tb927.9.9730_for | GGGGACAAGTTTGTACAAAAAAGCAGGCTATGCCCGTAACGAAGCCAAG | Tb927.9.9730_rev | GGGGACCACTTTGTACAAGAAAGCTGGGTTTAAGCCGCTGCACGCTT |
| 6_hypothetical | D7 | Tb927.11.10320_for | GGGGACAAGTTTGTACAAAAAAGCAGGCTATGATACATCTCCCAACATCAACTACAC | Tb927.11.10320_rev | GGGGACCACTTTGTACAAGAAAGCTGGGTCTAAACACCAGGTAACACACCCG |
| 6_hypothetical | D8 | Tb927.11.1840_for | GGGGACAAGTTTGTACAAAAAAGCAGGCTATGGAAACAGCAGAGTTAGCAAGAG | Tb927.11.1840_rev | GGGGACCACTTTGTACAAGAAAGCTGGGTCTACTGGCTTAGCTTTGAGCGG |
| 6_hypothetical | D9 | Tb927.9.2260_for | GGGGACAAGTTTGTACAAAAAAGCAGGCTATGGAAACAACACCAACGTTTTCTG | Tb927.9.2260_rev | GGGGACCACTTTGTACAAGAAAGCTGGGTTCATTCCCTACGTACGGGACTC |
| 6_hypothetical | D10 | Tb927.3.2240_for | GGGGACAAGTTTGTACAAAAAAGCAGGCTATGTTACTTAGGTTTGTCACTTTTGCCT | Tb927.3.2240_rev | GGGGACCACTTTGTACAAGAAAGCTGGGTTTAAACCCCAATAGAGATCTTAACTGGAGG |
| 6_hypothetical | D11 | Tb927.11.9870_for | GGGGACAAGTTTGTACAAAAAAGCAGGCTATGGAGTCATCGCTCGGTGC | Tb927.11.9870_rev | GGGGACCACTTTGTACAAGAAAGCTGGGTTCAAGACAGGGGTATGTCCTCG |

| | | | | | |
|---|---|---|---|---|---|
| 6_hypothetical | D12 | Tb927.11.6340_for | GGGGACAAGTTTGTACAAAAAAGCAGGCTATGTCATCCCTCTCTGGAGAGC | Tb927.11.6340_rev | GGGGACCACTTTGTACAAGAAAGCTGGGTTTACTCTTCTCGGTACCCGAACC |
| 6_hypothetical | D13 | Tb927.9.3100_for | GGGGACAAGTTTGTACAAAAAAGCAGGCTATGAGACGTACGATTGTCCAACTTTG | Tb927.9.3100_rev | GGGGACCACTTTGTACAAGAAAGCTGGGTTCACCGCTGGCTGGATGC |
| 6_hypothetical | D14 | Tb927.11.710_for | GGGGACAAGTTTGTACAAAAAAGCAGGCTATGTTTGGTCGGGGATTTGGTG | Tb927.11.710_rev | GGGGACCACTTTGTACAAGAAAGCTGGGTCTACACACCAATTACATTATCAATATGACT |
| 6_hypothetical | D15 | Tb927.5.660_for | GGGGACAAGTTTGTACAAAAAAGCAGGCTATGTTGGGGATGAGGAGTTACATCG | Tb927.5.660_rev | GGGGACCACTTTGTACAAGAAAGCTGGGTTTAAAAAACTAGAAAAAATAACTTAAGTGG |
| 6_hypothetical | D16 | Tb927.4.3760_for | GGGGACAAGTTTGTACAAAAAAGCAGGCTATGGTAAGAAACATGGGTGGAAATGC | Tb927.4.3760_rev | GGGGACCACTTTGTACAAGAAAGCTGGGTTCATCCTTCACTTTGAATTTCGTTTCTTTT |
| 6_hypothetical | D17 | Tb927.8.3320_for | GGGGACAAGTTTGTACAAAAAAGCAGGCTATGTGGAGGTTTGGTGCCTCC | Tb927.8.3320_rev | GGGGACCACTTTGTACAAGAAAGCTGGGTCTACTCTTCAATTTCATTCATACGCTTCTG |
| 6_hypothetical | D18 | Tb927.7.1480_for | GGGGACAAGTTTGTACAAAAAAGCAGGCTATGGTTGTAAATGAGGCTCCGTG | Tb927.7.1480_rev | GGGGACCACTTTGTACAAGAAAGCTGGGTTCACAGTCCCTCCACAAGAAGC |
| 6_hypothetical | D19 | Tb927.10.12320_for | GGGGACAAGTTTGTACAAAAAAGCAGGCTATGTTGAGTAGTGGAGAGCGCA | Tb927.10.12320_rev | GGGGACCACTTTGTACAAGAAAGCTGGGTTTAGTCCCCAAAGAGGGCTTGT |
| 6_hypothetical | D20 | Tb927.10.6340_for | GGGGACAAGTTTGTACAAAAAAGCAGGCTATGATGAGTCGAAGGCCGCC | Tb927.10.6340_rev | GGGGACCACTTTGTACAAGAAAGCTGGGTCTAAAAGCGCTTGGGCCGT |
| 6_hypothetical | D21 | Tb927.1.2770_for | GGGGACAAGTTTGTACAAAAAAGCAGGCTATGTCGGTCAGTTCCCCGC | Tb927.1.2770_rev | GGGGACCACTTTGTACAAGAAAGCTGGGTTCACATCAAGTGAAGCGCTCG |
| 6_hypothetical | D22 | Tb927.8.4140_for | GGGGACAAGTTTGTACAAAAAAGCAGGCTATGATTACAGAAGAACTCTTTCGATCCTTC | Tb927.8.4140_rev | GGGGACCACTTTGTACAAGAAAGCTGGGTCTAAACGTGCCGGTCGTCTAAA |
| 6_hypothetical | D23 | Tb927.11.6560_for | GGGGACAAGTTTGTACAAAAAAGCAGGCTATGCCCACTGCAGTTAGGCA | Tb927.11.6560_rev | GGGGACCACTTTGTACAAGAAAGCTGGGTCTAACTGGAGAACCACATTCGTCTC |
| 6_hypothetical | D24 | Tb927.7.1760_for | GGGGACAAGTTTGTACAAAAAAGCAGGCTATGGAGCATCTAATCGCTGCG | Tb927.7.1760_rev | GGGGACCACTTTGTACAAGAAAGCTGGGTCTAATCAAGCCATATATCAAGTGGTTCTTG |
| 6_hypothetical | E1 | Tb927.5.2030_for | GGGGACAAGTTTGTACAAAAAAGCAGGCTATGTTTCGCTGCACTGTGAGG | Tb927.5.2030_rev | GGGGACCACTTTGTACAAGAAAGCTGGGTTCAACCGCCGTCGTGGCC |
| 6_hypothetical | E2 | Tb927.8.7270_for | GGGGACAAGTTTGTACAAAAAAGCAGGCTATGAAGGGCCTCAGCCGC | Tb927.8.7270_rev | GGGGACCACTTTGTACAAGAAAGCTGGGTTCAAAACTGCTTCCCTCTGTTTATCTT |
| 6_hypothetical | E3 | Tb927.2.2910_for | GGGGACAAGTTTGTACAAAAAAGCAGGCTATGGTCTCCACGCGAAGAATAGC | Tb927.2.2910_rev | GGGGACCACTTTGTACAAGAAAGCTGGGTTCAAACATCATTAATATTCCATAATGAAAC |
| 6_hypothetical | E4 | Tb927.6.2240_for | GGGGACAAGTTTGTACAAAAAAGCAGGCTATGTCTGGGCTATGGAGTTTTAGGG | Tb927.6.2240_rev | GGGGACCACTTTGTACAAGAAAGCTGGGTTTATGAATGTATGGGACCAAATCTTCCTG |
| 6_hypothetical | E5 | Tb927.9.12950_for | GGGGACAAGTTTGTACAAAAAAGCAGGCTATGGAAACGAAAGGTTCTTCTGTTCA | Tb927.9.12950_rev | GGGGACCACTTTGTACAAGAAAGCTGGGTTTAATCGTCAACACTAAGTCTTTCCCAC |
| 6_hypothetical | E6 | Tb927.4.5100_for | GGGGACAAGTTTGTACAAAAAAGCAGGCTATGAAAGGCCTCAGCCGCC | Tb927.4.5100_rev | GGGGACCACTTTGTACAAGAAAGCTGGGTTCAAAACTGCTTCCCTCTATTCATCTTTAT |
| 6_hypothetical | E7 | Tb927.9.9030_for | GGGGACAAGTTTGTACAAAAAAGCAGGCTATGAGTGAAGACGGCTCCGA | Tb927.9.9030_rev | GGGGACCACTTTGTACAAGAAAGCTGGGTTCAAGCCTTCACTCCGTCTTTTC |
| 6_hypothetical | E8 | Tb927.4.370_for | GGGGACAAGTTTGTACAAAAAAGCAGGCTATGGTGATAGATGGTGACTTAGTTGCC | Tb927.4.370_rev | GGGGACCACTTTGTACAAGAAAGCTGGGTTCACAAATGAATACGAATGGATGGTAACA |
| 6_hypothetical | E9 | Tb927.10.800_for | GGGGACAAGTTTGTACAAAAAAGCAGGCTATGCATCCGGAAATTGAAGCACT | Tb927.10.800_rev | GGGGACCACTTTGTACAAGAAAGCTGGGTTTAATCAATCTCGAATATATCTGGGTCATT |
| 6_hypothetical | E10 | Tb927.3.3820_for | GGGGACAAGTTTGTACAAAAAAGCAGGCTATGTACCAAAGCAGAACAATGGTATCTT | Tb927.3.3820_rev | GGGGACCACTTTGTACAAGAAAGCTGGGTTTATCCATCAAGTTTCCAACTCCATTCC |
| 6_hypothetical | E11 | Tb927.11.16500_for | GGGGACAAGTTTGTACAAAAAAGCAGGCTATGTGCCTTCCGATCATCCGC | Tb927.11.16500_rev | GGGGACCACTTTGTACAAGAAAGCTGGGTTTAGGGGTTTACGGTACCTTCCAG |
| 6_hypothetical | E12 | Tb927.2.5000_for | GGGGACAAGTTTGTACAAAAAAGCAGGCTATGGCAACAAGAGGTGTGGC | Tb927.2.5000_rev | GGGGACCACTTTGTACAAGAAAGCTGGGTTCACTTGTGTTGAAGCCTGAGAGC |
| 6_hypothetical | E13 | Tb927.9.1950_for | GGGGACAAGTTTGTACAAAAAAGCAGGCTATGTATCGTAGGCTATGTGCGAGA | Tb927.9.1950_rev | GGGGACCACTTTGTACAAGAAAGCTGGGTCTAGTGTAGACCGTACCCACTCTC |
| 6_hypothetical | E14 | Tb927.8.1460_for | GGGGACAAGTTTGTACAAAAAAGCAGGCTATGGCATCCCAAAATGCAGCA | Tb927.8.1460_rev | GGGGACCACTTTGTACAAGAAAGCTGGGTCTAACTGGAGCGCGAACCC |
| 6_hypothetical | E15 | Tb927.6.4560_for | GGGGACAAGTTTGTACAAAAAAGCAGGCTATGCGAAAGTTTTGTCATTTCATGGC | Tb927.6.4560_rev | GGGGACCACTTTGTACAAGAAAGCTGGGTTTACAAAGTGAGCCATTCCTCCTCG |
| 6_hypothetical | E16 | Tb927.7.4510_for | GGGGACAAGTTTGTACAAAAAAGCAGGCTATGTACGAGTATGATGACCAGCAGG | Tb927.7.4510_rev | GGGGACCACTTTGTACAAGAAAGCTGGGTTTAACTTGCCAGAGACTCTTTTGCTT |
| 6_hypothetical | E17 | Tb927.5.2770_for | GGGGACAAGTTTGTACAAAAAAGCAGGCTATGACGGGGGCGGACGAG | Tb927.5.2770_rev | GGGGACCACTTTGTACAAGAAAGCTGGGTTTACCACTGGCGTCCATTTCG |
| 6_hypothetical | E18 | Tb927.10.10000_for | GGGGACAAGTTTGTACAAAAAAGCAGGCTATGCAACGCTGCACACCA | Tb927.10.10000_rev | GGGGACCACTTTGTACAAGAAAGCTGGGTTCAATAACTGCCGAGTACACGC |
| 6_hypothetical | E19 | Tb927.5.310_for | GGGGACAAGTTTGTACAAAAAAGCAGGCTATGGCACGTGTCACCTTTTGC | Tb927.5.310_rev | GGGGACCACTTTGTACAAGAAAGCTGGGTTTACATTTTCACGGAATGGTATGGATCC |
| 6_hypothetical | E20 | Tb927.4.2120_for | GGGGACAAGTTTGTACAAAAAAGCAGGCTATGGAATCACCAACGTGCGG | Tb927.4.2120_rev | GGGGACCACTTTGTACAAGAAAGCTGGGTCTACGAACCCCCTCCCCG |
| 6_hypothetical | E21 | Tb927.11.5660_for | GGGGACAAGTTTGTACAAAAAAGCAGGCTATGATGACGCAAAACGTGAAACAG | Tb927.11.5660_rev | GGGGACCACTTTGTACAAGAAAGCTGGGTTCACATCTTGCAGCCTTTTTGAAAC |
| 6_hypothetical | E22 | Tb927.3.4180_for | GGGGACAAGTTTGTACAAAAAAGCAGGCTATGCCGGCAGAGGGTCCA | Tb927.3.4180_rev | GGGGACCACTTTGTACAAGAAAGCTGGGTTTAAAGGCTTGCATTTCCATTGCG |
| 6_hypothetical | E23 | Tb927.11.10690_for | GGGGACAAGTTTGTACAAAAAAGCAGGCTATGAAGAGGCAAAGAACTGCATCC | Tb927.11.10690_rev | GGGGACCACTTTGTACAAGAAAGCTGGGTTCAGCTGCATTCAGCGAATCC |
| 6_hypothetical | E24 | Tb927.5.4280_for | GGGGACAAGTTTGTACAAAAAAGCAGGCTATGCCCCCCAAAAAGGGTAAAAA | Tb927.5.4280_rev | GGGGACCACTTTGTACAAGAAAGCTGGGTTTAGTATGCACGATACACAATATCCATGTC |
| 6_hypothetical | F1 | Tb927.3.1280_for | GGGGACAAGTTTGTACAAAAAAGCAGGCTATGGCTGATGCTTCTGCACTTG | Tb927.3.1280_rev | GGGGACCACTTTGTACAAGAAAGCTGGGTCTAACTTTCGATGAGCAACTGCAGT |
| 6_hypothetical | F2 | Tb927.7.2770_for | GGGGACAAGTTTGTACAAAAAAGCAGGCTATGTTTAGGAAACGTTACAGACCCC | Tb927.7.2770_rev | GGGGACCACTTTGTACAAGAAAGCTGGGTTCACACATGAGCAAGCCACTG |
| 6_hypothetical | F3 | Tb927.10.8130_for | GGGGACAAGTTTGTACAAAAAAGCAGGCTATGAGTGGTTGGTTTGGTTCGC | Tb927.10.8130_rev | GGGGACCACTTTGTACAAGAAAGCTGGGTCTATGGTTGCGTCCCAGCAG |
| 6_hypothetical | F4 | Tb927.11.16490_for | GGGGACAAGTTTGTACAAAAAAGCAGGCTATGCCAGCAAAACGCAAAGGC | Tb927.11.16490_rev | GGGGACCACTTTGTACAAGAAAGCTGGGTTCAGACATTGAAAATAGCAGCACCA |
| 6_hypothetical | F5 | Tb927.8.3960_for | GGGGACAAGTTTGTACAAAAAAGCAGGCTATGCGCTTTTTTGTTGTGGCAG | Tb927.8.3960_rev | GGGGACCACTTTGTACAAGAAAGCTGGGTCTATTCACGCTCCCTTTTGTCTCG |
| 6_hypothetical | F6 | Tb927.11.2560_for | GGGGACAAGTTTGTACAAAAAAGCAGGCTATGACAGATGAAAAAAAAGAGGAATGGATT | Tb927.11.2560_rev | GGGGACCACTTTGTACAAGAAAGCTGGGTTCAACAAAAAAAACTTTTTTGCAACTCCA |
| 6_hypothetical | F7 | Tb11.02.5430_for | GGGGACAAGTTTGTACAAAAAAGCAGGCTATGACACCACCAACGACTCGA | Tb11.02.5430_rev | GGGGACCACTTTGTACAAGAAAGCTGGGTTTACTCAGCACCCGCAGCG |
| 6_hypothetical | F8 | Tb927.10.11830_for | GGGGACAAGTTTGTACAAAAAAGCAGGCTATGCCTGCGCTCAGTTGG | Tb927.10.11830_rev | GGGGACCACTTTGTACAAGAAAGCTGGGTCTACACCAATCGTGGTTGCTGA |
| 6_hypothetical | F9 | Tb927.9.2816_for | GGGGACAAGTTTGTACAAAAAAGCAGGCTATGGAAATCCGCATGGTGCC | Tb927.9.2816_rev | GGGGACCACTTTGTACAAGAAAGCTGGGTTCACTCATCAGCTGCTGGAGT |
| 6_hypothetical | F10 | Tb927.10.11560_for | GGGGACAAGTTTGTACAAAAAAGCAGGCTATGGTTGGCGAAGATTCTAATTTAGGG | Tb927.10.11560_rev | GGGGACCACTTTGTACAAGAAAGCTGGGTTTAGAGACCAAGCCGCGACA |
| 6_hypothetical | F11 | Tb927.10.9920_for | GGGGACAAGTTTGTACAAAAAAGCAGGCTATGTCCTCACGAGCATGGGT | Tb927.10.9920_rev | GGGGACCACTTTGTACAAGAAAGCTGGGTTTAGAGCGCATAGTTTGTCACTCG |
| 6_hypothetical | F12 | Tb927.5.2540_for | GGGGACAAGTTTGTACAAAAAAGCAGGCTATGTCTGATTCTTCTCTGGTGTTCCT | Tb927.5.2540_rev | GGGGACCACTTTGTACAAGAAAGCTGGGTTCACTCCTCATCCATAATGTCCAGC |
| 6_hypothetical | F13 | Tb927.1.2700_for | GGGGACAAGTTTGTACAAAAAAGCAGGCTATGGAACAGTGCTGCGGGG | Tb927.1.2700_rev | GGGGACCACTTTGTACAAGAAAGCTGGGTTTACCACGGACACGACACCA |
| 6_hypothetical | F14 | Tb927.3.1300_for | GGGGACAAGTTTGTACAAAAAAGCAGGCTATGTGGTTCACAAACATTACGAAGGA | Tb927.3.1300_rev | GGGGACCACTTTGTACAAGAAAGCTGGGTCTACTTTACATTTGTCCAACCATCATTGTC |
| 6_hypothetical | F15 | Tb927.9.2802_for | GGGGACAAGTTTGTACAAAAAAGCAGGCTATGGAAATCCGCATGGTGCC | Tb927.9.2802_rev | GGGGACCACTTTGTACAAGAAAGCTGGGTTCACTCATCAGCTGCTGGAGT |
| 6_hypothetical | F16 | Tb927.2.4680_for | GGGGACAAGTTTGTACAAAAAAGCAGGCTATGCCCTCGGGATTCTTCCAG | Tb927.2.4680_rev | GGGGACCACTTTGTACAAGAAAGCTGGGTTCACCTCCTCTCTGTTATTTGTAGGTAA |
| 6_hypothetical | F17 | Tb927.11.10300_for | GGGGACAAGTTTGTACAAAAAAGCAGGCTATGAGCCACCCACAGCAGC | Tb927.11.10300_rev | GGGGACCACTTTGTACAAGAAAGCTGGGTTCATGATTCCGCCCTGAAGCG |
| 6_hypothetical | F18 | Tb927.2.4380_for | GGGGACAAGTTTGTACAAAAAAGCAGGCTATGAAGAGATTTGGAATTTGGACCCG | Tb927.2.4380_rev | GGGGACCACTTTGTACAAGAAAGCTGGGTTCAGTTTTGATCAATGCCAGCGC |
| 6_hypothetical | F19 | Tb927.5.3630_for | GGGGACAAGTTTGTACAAAAAAGCAGGCTATGGAAAAAGTAACGCCAACTTCCC | Tb927.5.3630_rev | GGGGACCACTTTGTACAAGAAAGCTGGGTTCAATATTTCTGGAGGGTACTCCACG |
| 6_hypothetical | F20 | Tb927.11.16040_for | GGGGACAAGTTTGTACAAAAAAGCAGGCTATGGACTCGGACGAAGGGG | Tb927.11.16040_rev | GGGGACCACTTTGTACAAGAAAGCTGGGTTCATAAACTGCCTACACCACCCA |
| 6_hypothetical | F21 | Tb927.4.840_for | GGGGACAAGTTTGTACAAAAAAGCAGGCTATGAGATGTCTTTTCGCTTCAGCA | Tb927.4.840_rev | GGGGACCACTTTGTACAAGAAAGCTGGGTTCACCACGGCGCACTCAT |
| 6_hypothetical | F22 | Tb927.7.7330_for | GGGGACAAGTTTGTACAAAAAAGCAGGCTATGTCTTCAGGCGGAAATGTTGG | Tb927.7.7330_rev | GGGGACCACTTTGTACAAGAAAGCTGGGTTCAAATATGCCGATAGTCGTAGTCCT |
| 6_hypothetical | F23 | Tb927.10.2050_for | GGGGACAAGTTTGTACAAAAAAGCAGGCTATGTTCCGAGGTAGTGGGGC | Tb927.10.2050_rev | GGGGACCACTTTGTACAAGAAAGCTGGGTTCAACAAAGTGAGACAACGAAAGATATGG |
| 6_hypothetical | F24 | Tb927.8.4800_for | GGGGACAAGTTTGTACAAAAAAGCAGGCTATGACAACGCTAGGTGTAGGAGAC | Tb927.8.4800_rev | GGGGACCACTTTGTACAAGAAAGCTGGGTTCACGTCGTAGCGGTGTCG |
| 6_hypothetical | G1 | Tb927.9.12740_for | GGGGACAAGTTTGTACAAAAAAGCAGGCTATGGGTTCAAAGTATGACCGAGTTAAG | Tb927.9.12740_rev | GGGGACCACTTTGTACAAGAAAGCTGGGTTCACGTATCGTTAGTAGGCGACA |
| 6_hypothetical | G2 | Tb927.3.5170_for | GGGGACAAGTTTGTACAAAAAAGCAGGCTATGGGGAGGGAGGTTTCTTGG | Tb927.3.5170_rev | GGGGACCACTTTGTACAAGAAAGCTGGGTCTATAGACCGAGTAACCCTCTCACAA |
| 6_hypothetical | G3 | Tb927.3.4690_for | GGGGACAAGTTTGTACAAAAAAGCAGGCTATGAAAATTGCGGTTAACCCGTACA | Tb927.3.4690_rev | GGGGACCACTTTGTACAAGAAAGCTGGGTCTAATGTTCCCCTGATTCCTTGTAGTG |
| 6_hypothetical | G4 | Tb927.10.9200_for | GGGGACAAGTTTGTACAAAAAAGCAGGCTATGACGCACTATGTTGATAGTGTATTGTAC | Tb927.10.9200_rev | GGGGACCACTTTGTACAAGAAAGCTGGGTTTAGGATCCGTCCGTTTCCCG |
| 6_hypothetical | G5 | Tb927.10.6130_for | GGGGACAAGTTTGTACAAAAAAGCAGGCTATGAACAACGGTGGTCCTTACAATG | Tb927.10.6130_rev | GGGGACCACTTTGTACAAGAAAGCTGGGTTTACCGGTACGAGTAACTACGTAACC |
| 6_hypothetical | G6 | Tb927.6.4610_for | GGGGACAAGTTTGTACAAAAAAGCAGGCTATGAACTCCGGCACTTGCATTG | Tb927.6.4610_rev | GGGGACCACTTTGTACAAGAAAGCTGGGTTTACTTCTTCATAGTGCTCGCACC |
| 6_hypothetical | G7 | Tb927.6.1810_for | GGGGACAAGTTTGTACAAAAAAGCAGGCTATGAATACGATACGTTACCTAGTGTCGG | Tb927.6.1810_rev | GGGGACCACTTTGTACAAGAAAGCTGGGTTCAATCTTTTCTCTCTAACAGCAAATCATT |
| 6_hypothetical | G8 | Tb927.7.4750_for | GGGGACAAGTTTGTACAAAAAAGCAGGCTATGTACCATTGTAAAACCAGTGATACCAT | Tb927.7.4750_rev | GGGGACCACTTTGTACAAGAAAGCTGGGTTTATGCTCGCTTGCAGGGTTT |
| 6_hypothetical | G9 | Tb927.1.3070_for | GGGGACAAGTTTGTACAAAAAAGCAGGCTATGTCTGCGTGGGCCTCT | Tb927.1.3070_rev | GGGGACCACTTTGTACAAGAAAGCTGGGTTCATGGTGCGGAAGATGTGTG |
| 6_hypothetical | G10 | Tb927.8.2720_for | GGGGACAAGTTTGTACAAAAAAGCAGGCTATGAAAACGGTAATCAGGGACCCG | Tb927.8.2720_rev | GGGGACCACTTTGTACAAGAAAGCTGGGTTTACGCATTCAGTATTTCGGCAGC |
| 6_hypothetical | G11 | Tb927.11.14570_for | GGGGACAAGTTTGTACAAAAAAGCAGGCTATGAGTTATAGCTGGCGCGGC | Tb927.11.14570_rev | GGGGACCACTTTGTACAAGAAAGCTGGGTCTACTCGTCAATGATATTTGCAAAGCC |
| 6_hypothetical | G12 | Tb927.11.9260_for | GGGGACAAGTTTGTACAAAAAAGCAGGCTATGGGATGTCAACAAAGTGGAGTG | Tb927.11.9260_rev | GGGGACCACTTTGTACAAGAAAGCTGGGTTCACGCCTGCGCCTTATGA |
| 6_hypothetical | G13 | Tb927.8.6520_for | GGGGACAAGTTTGTACAAAAAAGCAGGCTATGGAAGAAAAGATGACGAAGATTGACA | Tb927.8.6520_rev | GGGGACCACTTTGTACAAGAAAGCTGGGTTTACCACCACACAAGCCTCTGA |
| 6_hypothetical | G14 | Tb927.11.12380_for | GGGGACAAGTTTGTACAAAAAAGCAGGCTATGTCCACCAACTACGCACTGG | Tb927.11.12380_rev | GGGGACCACTTTGTACAAGAAAGCTGGGTTTAATTCTTATTCTCTGCCATTTTTTCCTG |
| 6_hypothetical | G15 | Tb927.8.5420_for | GGGGACAAGTTTGTACAAAAAAGCAGGCTATGGCAATACTTAGCCGTGTCATTG | Tb927.8.5420_rev | GGGGACCACTTTGTACAAGAAAGCTGGGTTTACAACTCCGAGAGTGATGATCGA |
| 6_hypothetical | G16 | Tb927.11.7920_for | GGGGACAAGTTTGTACAAAAAAGCAGGCTATGATACGGCAATGTGTTAAAACCATCC | Tb927.11.7920_rev | GGGGACCACTTTGTACAAGAAAGCTGGGTTCACGGGATAGTATCAGGATGTTCG |
| 6_hypothetical | G17 | Tb927.9.2100_for | GGGGACAAGTTTGTACAAAAAAGCAGGCTATGGAAACCCCCGATAACGTTG | Tb927.9.2100_rev | GGGGACCACTTTGTACAAGAAAGCTGGGTTTACCGGGCATCCTCTTCCC |
| 6_hypothetical | G18 | Tb927.11.15580_for | GGGGACAAGTTTGTACAAAAAAGCAGGCTATGATACGGAAGACCTTGGATGTGG | Tb927.11.15580_rev | GGGGACCACTTTGTACAAGAAAGCTGGGTTTAAGCATTAGACCTCCGTGATAGGT |
| 6_hypothetical | G19 | Tb927.9.4420_for | GGGGACAAGTTTGTACAAAAAAGCAGGCTATGTCCAAGTCTAGTCGTAAAATCACTTG | Tb927.9.4420_rev | GGGGACCACTTTGTACAAGAAAGCTGGGTTTAAATGCGGGCTCCGTCATC |
| 6_hypothetical | G20 | Tb927.11.1240_for | GGGGACAAGTTTGTACAAAAAAGCAGGCTATGAGTCACACAATGACAAATGCTTCT | Tb927.11.1240_rev | GGGGACCACTTTGTACAAGAAAGCTGGGTCTAAAACAAAAGTTGGCTGACAAGGC |
| 6_hypothetical | G21 | Tb927.9.13040_for | GGGGACAAGTTTGTACAAAAAAGCAGGCTATGAACTTCGTTCCTCTTTATAGATCAATT | Tb927.9.13040_rev | GGGGACCACTTTGTACAAGAAAGCTGGGTCTATTCGTCGTTAAACAAAAGAATGTGCC |
| 6_hypothetical | G22 | Tb927.7.2120_for | GGGGACAAGTTTGTACAAAAAAGCAGGCTATGCATCAAACGTCTGCAACTGT | Tb927.7.2120_rev | GGGGACCACTTTGTACAAGAAAGCTGGGTTCACACCACGCCATCAACCT |
| 6_hypothetical | G23 | Tb927.11.6530_for | GGGGACAAGTTTGTACAAAAAAGCAGGCTATGTTCGGTGGGTCTAAATCGGA | Tb927.11.6530_rev | GGGGACCACTTTGTACAAGAAAGCTGGGTTCATTGATGTGCGCGGTGATT |
| 6_hypothetical | G24 | Tb927.11.10090_for | GGGGACAAGTTTGTACAAAAAAGCAGGCTATGTCATCACCCATGTGTTCAGTC | Tb927.11.10090_rev | GGGGACCACTTTGTACAAGAAAGCTGGGTCTAAGATAGGTACCCCATTTCTACGGC |
| 6_hypothetical | H1 | Tb927.11.5470_for | GGGGACAAGTTTGTACAAAAAAGCAGGCTATGTCTTTTTTTGAAGATAGCGCAGAC | Tb927.11.5470_rev | GGGGACCACTTTGTACAAGAAAGCTGGGTCTACCGCTGAGTAACCATCATGC |
| 6_hypothetical | H2 | Tb927.10.15890_for | GGGGACAAGTTTGTACAAAAAAGCAGGCTATGCGGTCATCTTCCGCCG | Tb927.10.15890_rev | GGGGACCACTTTGTACAAGAAAGCTGGGTCTATGACTTCGCACTTCTGGCAA |
| 6_hypothetical | H3 | Tb927.8.6320_for | GGGGACAAGTTTGTACAAAAAAGCAGGCTATGATGTCATCCTCAGCTTTTACGG | Tb927.8.6320_rev | GGGGACCACTTTGTACAAGAAAGCTGGGTTTATTTGTGGCCATCAGAGTCGATAAC |
| 6_hypothetical | H4 | Tb927.4.3350_for | GGGGACAAGTTTGTACAAAAAAGCAGGCTATGCTTGGTGACGACCGC | Tb927.4.3350_rev | GGGGACCACTTTGTACAAGAAAGCTGGGTCTACGCAGCTTCCCTCGTATCG |
| 6_hypothetical | H5 | Tb927.11.4890_for | GGGGACAAGTTTGTACAAAAAAGCAGGCTATGAACTTTCCGGATAGTAGGGTACTT | Tb927.11.4890_rev | GGGGACCACTTTGTACAAGAAAGCTGGGTTCACGGTGCACGATCAGAGG |
| 6_hypothetical | H6 | Tb927.11.8030_for | GGGGACAAGTTTGTACAAAAAAGCAGGCTATGGAAGAAGGAACGTATGCGGG | Tb927.11.8030_rev | GGGGACCACTTTGTACAAGAAAGCTGGGTTTATTTGATAGCATTACGTGCGTGAAGT |
| 6_hypothetical | H7 | Tb927.9.12530_for | GGGGACAAGTTTGTACAAAAAAGCAGGCTATGTTTAAGTGCCGCGTGGC | Tb927.9.12530_rev | GGGGACCACTTTGTACAAGAAAGCTGGGTTCATAACTTCATGTGTTCAAGCGCG |
| 6_hypothetical | H8 | Tb927.5.3780_for | GGGGACAAGTTTGTACAAAAAAGCAGGCTATGAGTGTATCTCGGGCACAAGT | Tb927.5.3780_rev | GGGGACCACTTTGTACAAGAAAGCTGGGTTTAATTCCACCCCAAAAGTAATCTTACCTC |
| 6_hypothetical | H9 | Tb927.9.15230_for | GGGGACAAGTTTGTACAAAAAAGCAGGCTATGGAGTGCTCTGTGCACTCG | Tb927.9.15230_rev | GGGGACCACTTTGTACAAGAAAGCTGGGTTCAGAAGAACACCGCACGGG |
| 6_hypothetical | H10 | Tb927.10.9180_for | GGGGACAAGTTTGTACAAAAAAGCAGGCTATGTCAGCGTCAAGTGATTTGGAT | Tb927.10.9180_rev | GGGGACCACTTTGTACAAGAAAGCTGGGTTTACATCAGGCACGGTGCAGC |
| 6_hypothetical | H11 | Tb927.4.320_for | GGGGACAAGTTTGTACAAAAAAGCAGGCTATGAGGAAGTCGGCATTGAAGTTG | Tb927.4.320_rev | GGGGACCACTTTGTACAAGAAAGCTGGGTTCAAGGAAGCCCGTTGCG |
| 6_hypothetical | H12 | Tb927.9.12100_for | GGGGACAAGTTTGTACAAAAAAGCAGGCTATGTCTGTTTCTAGTCAAATAGGCATAAGG | Tb927.9.12100_rev | GGGGACCACTTTGTACAAGAAAGCTGGGTTTAATCTCTTGTTGCGTCCGTACG |
| 6_hypothetical | H13 | Tb927.11.6150_for | GGGGACAAGTTTGTACAAAAAAGCAGGCTATGTCGGCCCACCGCTTT | Tb927.11.6150_rev | GGGGACCACTTTGTACAAGAAAGCTGGGTTTAGTCTTTCACCTTTCTCCCCCTT |
| 6_hypothetical | H14 | Tb927.10.8990_for | GGGGACAAGTTTGTACAAAAAAGCAGGCTATGCCAATAAAGGGTTCGAGGAAG | Tb927.10.8990_rev | GGGGACCACTTTGTACAAGAAAGCTGGGTTCACTGGGGTTGCTCCCG |
| 6_hypothetical | H15 | Tb927.6.4800_for | GGGGACAAGTTTGTACAAAAAAGCAGGCTATGACTAGTATCGAGATGCAGGTGG | Tb927.6.4800_rev | GGGGACCACTTTGTACAAGAAAGCTGGGTTTATATAAATAGCCTTGTCGAGACCTGTGG |
| 6_hypothetical | H16 | Tb927.2.3000_for | GGGGACAAGTTTGTACAAAAAAGCAGGCTATGAATTACACTGCGGCCCG | Tb927.2.3000_rev | GGGGACCACTTTGTACAAGAAAGCTGGGTCTATACATCCAACTGAGTGTCCTGAGA |
| 6_hypothetical | H17 | Tb927.8.6770_for | GGGGACAAGTTTGTACAAAAAAGCAGGCTATGCTTACGTCCCTTTCGTGGG | Tb927.8.6770_rev | GGGGACCACTTTGTACAAGAAAGCTGGGTTTAGCAAAACACTCGCTGCACA |
| 6_hypothetical | H18 | Tb927.11.7780_for | GGGGACAAGTTTGTACAAAAAAGCAGGCTATGGAGTCGCCGATCAACCC | Tb927.11.7780_rev | GGGGACCACTTTGTACAAGAAAGCTGGGTTTACATGCCGCCACCCATCA |
| 6_hypothetical | H19 | Tb927.11.10850_for | GGGGACAAGTTTGTACAAAAAAGCAGGCTATGCTAAACCTGGAGTCGCTTAAC | Tb927.11.10850_rev | GGGGACCACTTTGTACAAGAAAGCTGGGTTCATAGCCCACGTCGTTCCC |
| 6_hypothetical | H20 | Tb927.8.4210_for | GGGGACAAGTTTGTACAAAAAAGCAGGCTATGCGGCTTCGTTCCTTCTTG | Tb927.8.4210_rev | GGGGACCACTTTGTACAAGAAAGCTGGGTTCAACTTTTCCATATACCACACGTATTATC |
| 6_hypothetical | H21 | Tb927.3.950_for | GGGGACAAGTTTGTACAAAAAAGCAGGCTATGCGTCGTACTTTTTCGCTTTTC | Tb927.3.950_rev | GGGGACCACTTTGTACAAGAAAGCTGGGTTTAAGGTGCTGCTGGCATTAGATC |
| 6_hypothetical | H22 | Tb927.5.3020_for | GGGGACAAGTTTGTACAAAAAAGCAGGCTATGCTGAACAACAAGAAAGAAAACACG | Tb927.5.3020_rev | GGGGACCACTTTGTACAAGAAAGCTGGGTCTACTCGAGTTCCTTTGTTGTCGAA |
| 6_hypothetical | H23 | Tb927.11.11580_for | GGGGACAAGTTTGTACAAAAAAGCAGGCTATGGACTTTGACACGTCAACTACG | Tb927.11.11580_rev | GGGGACCACTTTGTACAAGAAAGCTGGGTTCATGAACGAATAAGCTCCAGTAGGT |
| 6_hypothetical | H24 | Tb927.10.14490_for | GGGGACAAGTTTGTACAAAAAAGCAGGCTATGCTCCGTGAGCAGTGCA | Tb927.10.14490_rev | GGGGACCACTTTGTACAAGAAAGCTGGGTCTAATTAGCTGCTGCGGGAAGG |
| 6_hypothetical | I1 | Tb927.2.5970_for | GGGGACAAGTTTGTACAAAAAAGCAGGCTATGTCGCTTTCATGTGGACCAC | Tb927.2.5970_rev | GGGGACCACTTTGTACAAGAAAGCTGGGTTCAAACATGATCTGCACGGTCAC |
| 6_hypothetical | I2 | Tb927.8.4490_for | GGGGACAAGTTTGTACAAAAAAGCAGGCTATGCAAGATAATGTTGCGGGCT | Tb927.8.4490_rev | GGGGACCACTTTGTACAAGAAAGCTGGGTTTACTTTGGGGCGTCGCG |
| 6_hypothetical | I3 | Tb927.11.13350_for | GGGGACAAGTTTGTACAAAAAAGCAGGCTATGGCGGCACACGATGAGG | Tb927.11.13350_rev | GGGGACCACTTTGTACAAGAAAGCTGGGTCTAACGATTCTTAACTTTGGACTCCAGTAT |
| 6_hypothetical | I4 | Tb927.11.8390_for | GGGGACAAGTTTGTACAAAAAAGCAGGCTATGGTGTTCATCCTCTTGGCATG | Tb927.11.8390_rev | GGGGACCACTTTGTACAAGAAAGCTGGGTTCAGTCATGAATGTCGTACCCATATTTG |
| 6_hypothetical | I5 | Tb927.7.390_for | GGGGACAAGTTTGTACAAAAAAGCAGGCTATGCATGATTTGGTCGCTGTTGT | Tb927.7.390_rev | GGGGACCACTTTGTACAAGAAAGCTGGGTTTATTCTTTCGACTGTCCCTGTTCAG |
| 6_hypothetical | I6 | Tb927.8.2030_for | GGGGACAAGTTTGTACAAAAAAGCAGGCTATGGCCGATTCAAATGCTGTCG | Tb927.8.2030_rev | GGGGACCACTTTGTACAAGAAAGCTGGGTCTACTTACCCAACTTACTTACAAGTGCA |
| 6_hypothetical | I7 | Tb927.5.620_for | GGGGACAAGTTTGTACAAAAAAGCAGGCTATGTCATACAACGCCCTTTGCA | Tb927.5.620_rev | GGGGACCACTTTGTACAAGAAAGCTGGGTTCAGATTACCTTGTCTCCCGGTAG |
| 6_hypothetical | I8 | Tb927.7.3610_for | GGGGACAAGTTTGTACAAAAAAGCAGGCTATGTCCTCTAGGAGCGGCG | Tb927.7.3610_rev | GGGGACCACTTTGTACAAGAAAGCTGGGTTCACGGGACGTTGGTACTAATCA |
| 6_hypothetical | I9 | Tb927.8.6290_for | GGGGACAAGTTTGTACAAAAAAGCAGGCTATGGAAGACGCGGTAGAGGC | Tb927.8.6290_rev | GGGGACCACTTTGTACAAGAAAGCTGGGTTTAGCAATCTTTTGAAACAACACTTGACTC |
| 6_hypothetical | I10 | Tb927.4.1450_for | GGGGACAAGTTTGTACAAAAAAGCAGGCTATGATGATAGGTGTCCCGCCTG | Tb927.4.1450_rev | GGGGACCACTTTGTACAAGAAAGCTGGGTTCACATAAGTTTATTAAAGCAAGTACTTTC |
| 6_hypothetical | I11 | Tb927.4.1740_for | GGGGACAAGTTTGTACAAAAAAGCAGGCTATGCCGTTGTTGTATGCCTCAA | Tb927.4.1740_rev | GGGGACCACTTTGTACAAGAAAGCTGGGTTTAGGAGAAGTCCCATCGTTTCTTTAAC |
| 6_hypothetical | I12 | Tb927.6.4380_for | GGGGACAAGTTTGTACAAAAAAGCAGGCTATGGAGTGTGAATTCATGTGTGACG | Tb927.6.4380_rev | GGGGACCACTTTGTACAAGAAAGCTGGGTTCAATCAACCTTCCCCTTTTCGG |
| 6_hypothetical | I13 | Tb927.9.12980_for | GGGGACAAGTTTGTACAAAAAAGCAGGCTATGTCCGTCGTTGAAATGGAGAAG | Tb927.9.12980_rev | GGGGACCACTTTGTACAAGAAAGCTGGGTTTATTCATAAAGTTGAATGATACGCCTTTG |
| 6_hypothetical | I14 | Tb927.9.13580_for | GGGGACAAGTTTGTACAAAAAAGCAGGCTATGACCAAACATGGTGGCAGC | Tb927.9.13580_rev | GGGGACCACTTTGTACAAGAAAGCTGGGTTTAATAGAAGGGTTTGAACCGCGC |
| 6_hypothetical | I15 | Tb927.9.2814_for | GGGGACAAGTTTGTACAAAAAAGCAGGCTATGGAAATCCGCATGGTGCC | Tb927.9.2814_rev | GGGGACCACTTTGTACAAGAAAGCTGGGTTCACTCATCAGCTGCTGGAGT |
| 6_hypothetical | I17 | Tb927.5.3470_for | GGGGACAAGTTTGTACAAAAAAGCAGGCTATGCAAATGAGAGTAGAGTCTCCACA | Tb927.5.3470_rev | GGGGACCACTTTGTACAAGAAAGCTGGGTTTACTGGTCTTCCGAAAGGGACC |
| 6_hypothetical | I18 | Tb927.8.2890_for | GGGGACAAGTTTGTACAAAAAAGCAGGCTATGATGTCGAGTTTATTAGCGGACAC | Tb927.8.2890_rev | GGGGACCACTTTGTACAAGAAAGCTGGGTCTACTGCAAGGTTAGTGCTGACAA |
| 6_hypothetical | I19 | Tb927.10.1300_for | GGGGACAAGTTTGTACAAAAAAGCAGGCTATGCCAGCTGTCTGTCAAAGTC | Tb927.10.1300_rev | GGGGACCACTTTGTACAAGAAAGCTGGGTTCAAGAGAAGTCGACGGGAGG |
| 6_hypothetical | I20 | Tb927.10.15000_for | GGGGACAAGTTTGTACAAAAAAGCAGGCTATGATAGGCGGTCAGCACAAGA | Tb927.10.15000_rev | GGGGACCACTTTGTACAAGAAAGCTGGGTTCATCGGCGTTTCGGGATGA |
| 6_hypothetical | I21 | Tb927.11.2170_for | GGGGACAAGTTTGTACAAAAAAGCAGGCTATGATGGAATTAATCGGCTGCCC | Tb927.11.2170_rev | GGGGACCACTTTGTACAAGAAAGCTGGGTTTACCACACGCCATAGAACAAATCC |
| 6_hypothetical | I22 | Tb927.4.4750_for | GGGGACAAGTTTGTACAAAAAAGCAGGCTATGCAAGCATCGAGGTCAAAGG | Tb927.4.4750_rev | GGGGACCACTTTGTACAAGAAAGCTGGGTTTACGATCTTGCTGCTGTTTTCCT |

| | | | | | |
|---|---|---|---|---|---|
| 6_hypothetical | I23 | Tb927.11.7030_for | GGGGACAAGTTTGTACAAAAAAGCAGGCTATGGATGCACTCTGGCCAGT | Tb927.11.7030_rev | GGGGACCACTTTGTACAAGAAAGCTGGGTCTAGCAGTGAACAAATGTTGGCAC |
| 6_hypothetical | I24 | Tb927.9.10050_for | GGGGACAAGTTTGTACAAAAAAGCAGGCTATGCGGAAATTGGTTCGAAGCT | Tb927.9.10050_rev | GGGGACCACTTTGTACAAGAAAGCTGGGTTCACGAGCGGACCCTTGT |
| 6_hypothetical | J1 | Tb927.3.4780_for | GGGGACAAGTTTGTACAAAAAAGCAGGCTATGCGCACCCGACTACTTGT | Tb927.3.4780_rev | GGGGACCACTTTGTACAAGAAAGCTGGGTTCAGAGTGGCGGCATGGT |
| 6_hypothetical | J2 | Tb927.5.3550_for | GGGGACAAGTTTGTACAAAAAAGCAGGCTATGTCCAGGAGGTGGTCTTGC | Tb927.5.3550_rev | GGGGACCACTTTGTACAAGAAAGCTGGGTCTAAAGGTATGCTTGCAGCAACTCG |
| 6_hypothetical | J4 | Tb927.11.9090_for | GGGGACAAGTTTGTACAAAAAAGCAGGCTATGTTGCTACCTTGCACACCG | Tb927.11.9090_rev | GGGGACCACTTTGTACAAGAAAGCTGGGTCTAACAAATCGACCTCAGTCGAATCA |
| 6_hypothetical | J5 | Tb927.7.4410_for | GGGGACAAGTTTGTACAAAAAAGCAGGCTATGATAGAGTTCGCCAGCCCG | Tb927.7.4410_rev | GGGGACCACTTTGTACAAGAAAGCTGGGTCTACATGTTCTCTCTTAAAAAATTCCCTGA |
| 6_hypothetical | J6 | Tb927.6.3930_for | GGGGACAAGTTTGTACAAAAAAGCAGGCTATGTCCAGAAACGCTTTTTCGTCT | Tb927.6.3930_rev | GGGGACCACTTTGTACAAGAAAGCTGGGTCTATTCATTGTACTCTTTCTGCCTGACG |
| 6_hypothetical | J7 | Tb927.4.3820_for | GGGGACAAGTTTGTACAAAAAAGCAGGCTATGAATGATTTACGTCAACGGGCC | Tb927.4.3820_rev | GGGGACCACTTTGTACAAGAAAGCTGGGTTCAATACCCTTCATCATCGGGCG |
| 6_hypothetical | J8 | Tb927.1.4230_for | GGGGACAAGTTTGTACAAAAAAGCAGGCTATGCTTAAGGCCTGTTTGCCG | Tb927.1.4230_rev | GGGGACCACTTTGTACAAGAAAGCTGGGTTTATTTGGGAGGTTCCTTGGCAC |
| 6_hypothetical | J9 | Tb927.5.2750_for | GGGGACAAGTTTGTACAAAAAAGCAGGCTATGACTTCCTTAAAGGAGAAGGTACTGT | Tb927.5.2750_rev | GGGGACCACTTTGTACAAGAAAGCTGGGTTCAATACAAACAAATACCTTCAGTTGGTTC |
| 6_hypothetical | J10 | Tb927.11.14500_for | GGGGACAAGTTTGTACAAAAAAGCAGGCTATGTCAAGCGGTTGCGCA | Tb927.11.14500_rev | GGGGACCACTTTGTACAAGAAAGCTGGGTTTAACCAATCTGCTTTATCCACGAGATC |
| 6_hypothetical | J11 | Tb927.7.6250_for | GGGGACAAGTTTGTACAAAAAAGCAGGCTATGAGTGGGACAAGTTCTTCACTTG | Tb927.7.6250_rev | GGGGACCACTTTGTACAAGAAAGCTGGGTTCATACAGCGATCGAACACGAAAC |
| 6_hypothetical | J12 | Tb927.10.11510_for | GGGGACAAGTTTGTACAAAAAAGCAGGCTATGTCTGTTGCGTCTTCGACC | Tb927.10.11510_rev | GGGGACCACTTTGTACAAGAAAGCTGGGTCTAAAGGAGAAATGCCATGCCATTATCG |
| 6_hypothetical | J13 | Tb927.4.580_for | GGGGACAAGTTTGTACAAAAAAGCAGGCTATGGTAAATGGTGTGGTGCCCT | Tb927.4.580_rev | GGGGACCACTTTGTACAAGAAAGCTGGGTCTAATGCGTCATGCGACGTGA |
| 6_hypothetical | J14 | Tb927.3.3520_for | GGGGACAAGTTTGTACAAAAAAGCAGGCTATGGTCTACCCCTTTTCCAAGCA | Tb927.3.3520_rev | GGGGACCACTTTGTACAAGAAAGCTGGGTTTAACCCCCGGAACTTTCAACCT |
| 6_hypothetical | J15 | Tb927.3.1530_for | GGGGACAAGTTTGTACAAAAAAGCAGGCTATGCCGTGTGCCACTAAAGG | Tb927.3.1530_rev | GGGGACCACTTTGTACAAGAAAGCTGGGTTCACTCATATGGAGCAATCGGATCA |
| 6_hypothetical | J16 | Tb927.11.4390_for | GGGGACAAGTTTGTACAAAAAAGCAGGCTATGACGTCCGTGCAGTATGC | Tb927.11.4390_rev | GGGGACCACTTTGTACAAGAAAGCTGGGTCTAGGTGTATTGTAACACGGTAGCAG |
| 6_hypothetical | J17 | Tb927.9.2812_for | GGGGACAAGTTTGTACAAAAAAGCAGGCTATGGAAATCCGCATGGTGCC | Tb927.9.2812_rev | GGGGACCACTTTGTACAAGAAAGCTGGGTTCACTCATCAGCTGCTGGAGT |
| 6_hypothetical | J18 | Tb927.11.2720_for | GGGGACAAGTTTGTACAAAAAAGCAGGCTATGAGGGGAATGTGGGCGA | Tb927.11.2720_rev | GGGGACCACTTTGTACAAGAAAGCTGGGTTCAACGGCAGTAGAGGTACCC |
| 6_hypothetical | J19 | Tb927.10.11240_for | GGGGACAAGTTTGTACAAAAAAGCAGGCTATGAGTGCTGAGGATGCACTTG | Tb927.10.11240_rev | GGGGACCACTTTGTACAAGAAAGCTGGGTTTACTTATCCTCCCTTTTGGCCAGT |
| 6_hypothetical | J20 | Tb927.9.3080_for | GGGGACAAGTTTGTACAAAAAAGCAGGCTATGCCACAGCCGTCAGCG | Tb927.9.3080_rev | GGGGACCACTTTGTACAAGAAAGCTGGGTTCAAACGATTTTGTCAATCGGCG |
| 6_hypothetical | J21 | Tb927.8.2670_for | GGGGACAAGTTTGTACAAAAAAGCAGGCTATGCTCAGAAGAGCGGTACCC | Tb927.8.2670_rev | GGGGACCACTTTGTACAAGAAAGCTGGGTTCACAACGAAGAGGGAAGCTCG |
| 6_hypothetical | J22 | Tb927.9.2860_for | GGGGACAAGTTTGTACAAAAAAGCAGGCTATGGAAATCCGCATGGTGCC | Tb927.9.2860_rev | GGGGACCACTTTGTACAAGAAAGCTGGGTTCACTCATCAGCTGCTGGAGT |
| 6_hypothetical | J23 | Tb927.6.1170_for | GGGGACAAGTTTGTACAAAAAAGCAGGCTATGCACTTTATTGGCATTGTTGGC | Tb927.6.1170_rev | GGGGACCACTTTGTACAAGAAAGCTGGGTTCAATTCTCCAATCCCGCTGC |
| 6_hypothetical | J24 | Tb927.2.3160_for | GGGGACAAGTTTGTACAAAAAAGCAGGCTATGTCCAGTCGGTGCTCCTC | Tb927.2.3160_rev | GGGGACCACTTTGTACAAGAAAGCTGGGTTCACCTCGCAATCCAAGAAGGA |
| 6_hypothetical | K1 | Tb927.10.14020_for | GGGGACAAGTTTGTACAAAAAAGCAGGCTATGGCAGAAGCGGACTGTTCC | Tb927.10.14020_rev | GGGGACCACTTTGTACAAGAAAGCTGGGTTCATTCATACTGTTCAGTGGGTGTTAAG |
| 6_hypothetical | K2 | Tb927.7.5620_for | GGGGACAAGTTTGTACAAAAAAGCAGGCTATGGTGAGGCTTCGCTCGG | Tb927.7.5620_rev | GGGGACCACTTTGTACAAGAAAGCTGGGTTTACTCCCTTCGAATACATTTTGGGC |
| 6_hypothetical | K3 | Tb927.7.3080_for | GGGGACAAGTTTGTACAAAAAAGCAGGCTATGGCAGAGGGAGTACCGGC | Tb927.7.3080_rev | GGGGACCACTTTGTACAAGAAAGCTGGGTCTACCGGCCTTCACCCTCG |
| 6_hypothetical | K4 | Tb927.4.500_for | GGGGACAAGTTTGTACAAAAAAGCAGGCTATGAAGAGGTATTTTCTCATTGATGAATCT | Tb927.4.500_rev | GGGGACCACTTTGTACAAGAAAGCTGGGTTTAAGCTGGAACTTTACCGCGC |
| 6_hypothetical | K5 | Tb927.11.9760_for | GGGGACAAGTTTGTACAAAAAAGCAGGCTATGAGCTCACCGTACCAACGA | Tb927.11.9760_rev | GGGGACCACTTTGTACAAGAAAGCTGGGTTTATATCGCCACCGCTTCCTCT |
| 6_hypothetical | K6 | Tb927.11.12920_for | GGGGACAAGTTTGTACAAAAAAGCAGGCTATGCCCCATTCGGGATACCC | Tb927.11.12920_rev | GGGGACCACTTTGTACAAGAAAGCTGGGTCTACAAGTCGTTCATGCTATGCGG |
| 6_hypothetical | K7 | Tb927.4.1590.1_for | GGGGACAAGTTTGTACAAAAAAGCAGGCTATGCTGTCACCCTTGTTACTATGGA | Tb927.4.1590.1_rev | GGGGACCACTTTGTACAAGAAAGCTGGGTCTACTTGCCGTACTCTGTGGAGA |
| 6_hypothetical | K8 | Tb927.7.2800_for | GGGGACAAGTTTGTACAAAAAAGCAGGCTATGAGTTTTGATGGCCTTCGGG | Tb927.7.2800_rev | GGGGACCACTTTGTACAAGAAAGCTGGGTTTATCCCGCTGTTTCCTTGTTCG |
| 6_hypothetical | K9 | Tb927.2.4340_for | GGGGACAAGTTTGTACAAAAAAGCAGGCTATGCAAAATCCTACGATTTATCGAAATTCC | Tb927.2.4340_rev | GGGGACCACTTTGTACAAGAAAGCTGGGTTCATCTTGAAAGCAGGGATCGC |
| 6_hypothetical | K10 | Tb927.8.6820_for | GGGGACAAGTTTGTACAAAAAAGCAGGCTATGTGGAAATGTTACCCTCTTCGC | Tb927.8.6820_rev | GGGGACCACTTTGTACAAGAAAGCTGGGTCTAAACGACCGATCCTTGTGGC |
| 6_hypothetical | K11 | Tb927.11.5870_for | GGGGACAAGTTTGTACAAAAAAGCAGGCTATGAGCACATCCCTCACGTCG | Tb927.11.5870_rev | GGGGACCACTTTGTACAAGAAAGCTGGGTTTACGATTTCTTCCCAAACAGAGAAAGA |
| 6_hypothetical | K12 | Tb927.11.10890_for | GGGGACAAGTTTGTACAAAAAAGCAGGCTATGGATACGAGCTCGCCGA | Tb927.11.10890_rev | GGGGACCACTTTGTACAAGAAAGCTGGGTTCATGGGGGCGCTATGATAGC |
| 6_hypothetical | K13 | Tb927.9.4440_for | GGGGACAAGTTTGTACAAAAAAGCAGGCTATGTCACTCAGGTCCACAATTCAGA | Tb927.9.4440_rev | GGGGACCACTTTGTACAAGAAAGCTGGGTTCAGCAAAAGGTGCTTTCCTCAA |
| 6_hypothetical | K14 | Tb927.10.13800_for | GGGGACAAGTTTGTACAAAAAAGCAGGCTATGTCTGAGGCTAGCGACCG | Tb927.10.13800_rev | GGGGACCACTTTGTACAAGAAAGCTGGGTCTACTGAGAACAATCGGCGTTGG |
| 6_hypothetical | K15 | Tb927.11.10880_for | GGGGACAAGTTTGTACAAAAAAGCAGGCTATGCTTCGCCGTACACTTTCG | Tb927.11.10880_rev | GGGGACCACTTTGTACAAGAAAGCTGGGTCTACCAGCGGAAGTCGTTAAATATATTTGT |
| 6_hypothetical | K16 | Tb927.4.740_for | GGGGACAAGTTTGTACAAAAAAGCAGGCTATGTTGTTGCTCAGGGCAACG | Tb927.4.740_rev | GGGGACCACTTTGTACAAGAAAGCTGGGTTCAAAAAATAACTTCGAGCGTTATGCC |
| 6_hypothetical | K17 | Tb927.10.5200_for | GGGGACAAGTTTGTACAAAAAAGCAGGCTATGCAACACATTAACCATAGATTCAGTTAC | Tb927.10.5200_rev | GGGGACCACTTTGTACAAGAAAGCTGGGTCTACGTCACGCATTCTACATCGG |
| 6_hypothetical | K18 | Tb927.7.2460_for | GGGGACAAGTTTGTACAAAAAAGCAGGCTATGTCCACACACCGTGAGAGT | Tb927.7.2460_rev | GGGGACCACTTTGTACAAGAAAGCTGGGTTCATAGCTGAACACCACAAAGCAC |
| 6_hypothetical | K19 | Tb927.5.2860_for | GGGGACAAGTTTGTACAAAAAAGCAGGCTATGAAGGAGTTTCGACGTCGAATG | Tb927.5.2860_rev | GGGGACCACTTTGTACAAGAAAGCTGGGTTTACTTGCGTATCTGGCACTCAGC |
| 6_hypothetical | K20 | Tb927.11.10860_for | GGGGACAAGTTTGTACAAAAAAGCAGGCTATGTCGTGGGCTGCACTTG | Tb927.11.10860_rev | GGGGACCACTTTGTACAAGAAAGCTGGGTTCACTTCCGTCGCGCCGC |
| 6_hypothetical | K21 | Tb927.3.870_for | GGGGACAAGTTTGTACAAAAAAGCAGGCTATGCAAACTCCGCTACGTCC | Tb927.3.870_rev | GGGGACCACTTTGTACAAGAAAGCTGGGTTTACTCTGTGAATAATATCTGTTGGGCAC |
| 6_hypothetical | K22 | Tb927.5.4400_for | GGGGACAAGTTTGTACAAAAAAGCAGGCTATGGGACGGAGGAAGGAGAAGA | Tb927.5.4400_rev | GGGGACCACTTTGTACAAGAAAGCTGGGTCTAAGTGCTTGCAGCCTTCTTTCC |
| 6_hypothetical | K23 | Tb927.10.5810_for | GGGGACAAGTTTGTACAAAAAAGCAGGCTATGGACGAGAGAGGCATCTTAGAAT | Tb927.10.5810_rev | GGGGACCACTTTGTACAAGAAAGCTGGGTTTATTTCTTTGTTTCATCAAGAATGTCGTT |
| 6_hypothetical | K24 | Tb927.3.3120_for | GGGGACAAGTTTGTACAAAAAAGCAGGCTATGGCGGAGGCCGCCCGC | Tb927.3.3120_rev | GGGGACCACTTTGTACAAGAAAGCTGGGTTTACACTGCAGCGTCAAACTCC |
| 6_hypothetical | L1 | Tb927.10.9590_for | GGGGACAAGTTTGTACAAAAAAGCAGGCTATGTCGTCGTCGCGGGCC | Tb927.10.9590_rev | GGGGACCACTTTGTACAAGAAAGCTGGGTTTATAGCGTGTCACGCGAAAGAG |
| 6_hypothetical | L2 | Tb927.11.14060_for | GGGGACAAGTTTGTACAAAAAAGCAGGCTATGGAACTTGATGAAGAGACGAATGC | Tb927.11.14060_rev | GGGGACCACTTTGTACAAGAAAGCTGGGTCTACCGCAAAAGAAGTTCCCACG |
| 6_hypothetical | L3 | Tb927.7.6370_for | GGGGACAAGTTTGTACAAAAAAGCAGGCTATGGCTGCCCGGGCTTTA | Tb927.7.6370_rev | GGGGACCACTTTGTACAAGAAAGCTGGGTTTAGTGTTGCCCCTGACACTCC |
| 6_hypothetical | L4 | Tb927.10.480_for | GGGGACAAGTTTGTACAAAAAAGCAGGCTATGGTTGATGTTAATGGAACGACAATTTT | Tb927.10.480_rev | GGGGACCACTTTGTACAAGAAAGCTGGGTTTACTCTTCCATTTCAGCCACCATTC |
| 6_hypothetical | L5 | Tb927.10.1850_for | GGGGACAAGTTTGTACAAAAAAGCAGGCTATGGGTCAGTGGCTTTGCTTTC | Tb927.10.1850_rev | GGGGACCACTTTGTACAAGAAAGCTGGGTTTAGCCTGGGCTGGCCCC |
| 6_hypothetical | L6 | Tb927.10.11290_for | GGGGACAAGTTTGTACAAAAAAGCAGGCTATGCAACGGTATGGTTCCCTGA | Tb927.10.11290_rev | GGGGACCACTTTGTACAAGAAAGCTGGGTCTAGCTGACAAATAACAGTGCAGTCT |
| 6_hypothetical | L7 | Tb927.10.12230_for | GGGGACAAGTTTGTACAAAAAAGCAGGCTATGGACTCGGTTTGCCCGG | Tb927.10.12230_rev | GGGGACCACTTTGTACAAGAAAGCTGGGTCTACATTTCACTTTTTGGAAAGTAGAAGCT |
| 6_hypothetical | L8 | Tb927.8.2570_for | GGGGACAAGTTTGTACAAAAAAGCAGGCTATGGCAAGGTTTCAATCTTCTGTGG | Tb927.8.2570_rev | GGGGACCACTTTGTACAAGAAAGCTGGGTCTACGATTTGCACTTTTCGGTGTC |
| 6_hypothetical | L9 | Tb927.2.2880_for | GGGGACAAGTTTGTACAAAAAAGCAGGCTATGGCGGCGATGAGCGAG | Tb927.2.2880_rev | GGGGACCACTTTGTACAAGAAAGCTGGGTCTACCGCGCCTTTCGCCG |
| 6_hypothetical | L10 | Tb927.4.3000_for | GGGGACAAGTTTGTACAAAAAAGCAGGCTATGGACAGCCAAGCACGGG | Tb927.4.3000_rev | GGGGACCACTTTGTACAAGAAAGCTGGGTCTACGAAGAGCCGCTTTGCA |
| 6_hypothetical | L11 | Tb927.8.1960_for | GGGGACAAGTTTGTACAAAAAAGCAGGCTATGGTGGTGGAGTACCTTGGTC | Tb927.8.1960_rev | GGGGACCACTTTGTACAAGAAAGCTGGGTTTACTTTAGTTCATCCCATCTCGTAACCA |
| 6_hypothetical | L13 | Tb927.3.2720_for | GGGGACAAGTTTGTACAAAAAAGCAGGCTATGCAAAAGTCAGTTCCTTCATATGTTG | Tb927.3.2720_rev | GGGGACCACTTTGTACAAGAAAGCTGGGTTCATGAAAGGAAATAAACCTTCATTTCCTT |
| 6_hypothetical | L14 | Tb927.11.8330_for | GGGGACAAGTTTGTACAAAAAAGCAGGCTATGCGTGTAGGAGGATATATGGGTAC | Tb927.11.8330_rev | GGGGACCACTTTGTACAAGAAAGCTGGGTTTACAGAGTGGATTCCAGGATTCGG |
| 6_hypothetical | L17 | Tb927.11.8320_for | GGGGACAAGTTTGTACAAAAAAGCAGGCTATGATGATGCGAGTGGTAGGTAAGG | Tb927.11.8320_rev | GGGGACCACTTTGTACAAGAAAGCTGGGTTCAAAGGGTTCGAATCACTATAGGAGA |
| 6_hypothetical | L18 | Tb927.11.3580_for | GGGGACAAGTTTGTACAAAAAAGCAGGCTATGACACACAACAGGAGCGGG | Tb927.11.3580_rev | GGGGACCACTTTGTACAAGAAAGCTGGGTCTAGCCCCCCTCCTTTTTTTGC |
| 6_hypothetical | L19 | Tb927.10.6300_for | GGGGACAAGTTTGTACAAAAAAGCAGGCTATGTTTCGGTTGACTGCTGCTT | Tb927.10.6300_rev | GGGGACCACTTTGTACAAGAAAGCTGGGTTCAGTGTAACTGGTAGTTACGTTTTATGTG |
| 6_hypothetical | L20 | Tb927.10.330_for | GGGGACAAGTTTGTACAAAAAAGCAGGCTATGGAAAAGGCCCGTCAATGG | Tb927.10.330_rev | GGGGACCACTTTGTACAAGAAAGCTGGGTTTAATGTGACATTGTTTCTACACCCACT |
| 6_hypothetical | L21 | Tb927.6.3100_for | GGGGACAAGTTTGTACAAAAAAGCAGGCTATGCCCCGAGGATCAAGCA | Tb927.6.3100_rev | GGGGACCACTTTGTACAAGAAAGCTGGGTCTATTCGTGTTGAAAGTGTTGATTCGC |
| 6_hypothetical | L22 | Tb927.10.6100_for | GGGGACAAGTTTGTACAAAAAAGCAGGCTATGCAGCCGGTAAGGACGT | Tb927.10.6100_rev | GGGGACCACTTTGTACAAGAAAGCTGGGTCTAGTACATAATGGTGCGGCATAAGT |
| 6_hypothetical | L23 | Tb927.11.2010_for | GGGGACAAGTTTGTACAAAAAAGCAGGCTATGGGTCTTTTAACGAGGTATTATGAGAAA | Tb927.11.2010_rev | GGGGACCACTTTGTACAAGAAAGCTGGGTTCAGTTTCCGGCCAGCCC |
| 6_hypothetical | L24 | Tb927.4.3340_for | GGGGACAAGTTTGTACAAAAAAGCAGGCTATGAACTTGCCTTTCCCTACTCACA | Tb927.4.3340_rev | GGGGACCACTTTGTACAAGAAAGCTGGGTTTAGCGTGCACATCCCTCTTTG |
| 6_hypothetical | M1 | Tb927.11.920_for | GGGGACAAGTTTGTACAAAAAAGCAGGCTATGCGTACAGGGCGCCTG | Tb927.11.920_rev | GGGGACCACTTTGTACAAGAAAGCTGGGTCTACCATACGCTGGGGGCG |
| 6_hypothetical | M2 | Tb927.8.8320_for | GGGGACAAGTTTGTACAAAAAAGCAGGCTATGAATGTACTTTCTGACATTGCTCACT | Tb927.8.8320_rev | GGGGACCACTTTGTACAAGAAAGCTGGGTTCACTTGAGGGATATCATGCACGC |
| 6_hypothetical | M3 | Tb927.6.4520_for | GGGGACAAGTTTGTACAAAAAAGCAGGCTATGGACGCTGGAGATGCCC | Tb927.6.4520_rev | GGGGACCACTTTGTACAAGAAAGCTGGGTCTATTTCGTAATCCCAAATTTCTTCTTTTC |
| 6_hypothetical | M4 | Tb927.7.3400_for | GGGGACAAGTTTGTACAAAAAAGCAGGCTATGGCTTTGGGATCAGGAATCGG | Tb927.7.3400_rev | GGGGACCACTTTGTACAAGAAAGCTGGGTTCATGTTGGTGCATCCGCC |
| 6_hypothetical | M5 | Tb927.7.180_for | GGGGACAAGTTTGTACAAAAAAGCAGGCTATGTATGCTATTAAAATTTTTGTTATTTCG | Tb927.7.180_rev | GGGGACCACTTTGTACAAGAAAGCTGGGTTCATTCCACTGGAAATAGTTGCTTCAG |
| 6_hypothetical | M6 | Tb927.11.5580_for | GGGGACAAGTTTGTACAAAAAAGCAGGCTATGGTAAGAAAGGAGGAAGAAAAAGTTTCA | Tb927.11.5580_rev | GGGGACCACTTTGTACAAGAAAGCTGGGTCTATTTTCTCCAGCGAGTCCATATACC |
| 6_hypothetical | M7 | Tb927.1.3410_for | GGGGACAAGTTTGTACAAAAAAGCAGGCTATGAAGCGCAGGGTATTGCG | Tb927.1.3410_rev | GGGGACCACTTTGTACAAGAAAGCTGGGTTCACTCTTTTCGCTCTTTCTTCCTC |
| 6_hypothetical | M8 | Tb927.11.4750_for | GGGGACAAGTTTGTACAAAAAAGCAGGCTATGAATCGCATACTGTTTGCACTTG | Tb927.11.4750_rev | GGGGACCACTTTGTACAAGAAAGCTGGGTTTAGACGACAAAGGTAAGAAACACAAACA |
| 6_hypothetical | M9 | Tb927.10.13100_for | GGGGACAAGTTTGTACAAAAAAGCAGGCTATGAAAGACGATGAGTTGGAAAGAAAAATT | Tb927.10.13100_rev | GGGGACCACTTTGTACAAGAAAGCTGGGTCTAGTGAGCCAATAGTGACATCTGTTT |
| 6_hypothetical | M10 | Tb927.3.1940_for | GGGGACAAGTTTGTACAAAAAAGCAGGCTATGGAGCGGACACGGATTAACT | Tb927.3.1940_rev | GGGGACCACTTTGTACAAGAAAGCTGGGTCTACAGCTTTATTCCGGTCTTCTTGT |
| 6_hypothetical | M11 | Tb927.9.11540_for | GGGGACAAGTTTGTACAAAAAAGCAGGCTATGGCTGACGTTAGCGCCG | Tb927.9.11540_rev | GGGGACCACTTTGTACAAGAAAGCTGGGTTTATCGCATTGGGCGGGTTC |
| 6_hypothetical | M12 | Tb927.9.13110_for | GGGGACAAGTTTGTACAAAAAAGCAGGCTATGGAGAATTCGTGTCCAGGGG | Tb927.9.13110_rev | GGGGACCACTTTGTACAAGAAAGCTGGGTTTATTTCCCCTTCTTTGCTTCGAAAACC |
| 6_hypothetical | M13 | Tb927.4.3690_for | GGGGACAAGTTTGTACAAAAAAGCAGGCTATGCTGCGGAGGACTGTAACA | Tb927.4.3690_rev | GGGGACCACTTTGTACAAGAAAGCTGGGTTTACCGCCTAATCATCTCCTCCTTC |
| 6_hypothetical | M14 | Tb927.6.3600_for | GGGGACAAGTTTGTACAAAAAAGCAGGCTATGTGGTCCAAGGTGTTTACACC | Tb927.6.3600_rev | GGGGACCACTTTGTACAAGAAAGCTGGGTCTACGATGCTAGTTCTAGCAAGAAAGC |
| 6_hypothetical | M15 | Tb927.6.2630_for | GGGGACAAGTTTGTACAAAAAAGCAGGCTATGTGGTTCAGCCGCTTTTCG | Tb927.6.2630_rev | GGGGACCACTTTGTACAAGAAAGCTGGGTTTATTTTTTAAAGGAAGTACGGTCGCGC |
| 6_hypothetical | M16 | Tb927.11.13980_for | GGGGACAAGTTTGTACAAAAAAGCAGGCTATGACGGAAATGACGAAGGATGTTT | Tb927.11.13980_rev | GGGGACCACTTTGTACAAGAAAGCTGGGTTTACTCTATACGTTTCTCCCACGAATAAAA |
| 6_hypothetical | M17 | Tb927.10.12880_for | GGGGACAAGTTTGTACAAAAAAGCAGGCTATGCCGACTCAGAGCCCC | Tb927.10.12880_rev | GGGGACCACTTTGTACAAGAAAGCTGGGTCTAGGGCGGAAAAAGTTTCTGGC |
| 6_hypothetical | M18 | Tb927.9.2811_for | GGGGACAAGTTTGTACAAAAAAGCAGGCTATGGAAATCCGCATGGTGCC | Tb927.9.2811_rev | GGGGACCACTTTGTACAAGAAAGCTGGGTTCACTCATCAGCTGCTGGAGT |
| 6_hypothetical | M19 | Tb927.9.11880_for | GGGGACAAGTTTGTACAAAAAAGCAGGCTATGATGCGCAGTTCTTCTTTTTGC | Tb927.9.11880_rev | GGGGACCACTTTGTACAAGAAAGCTGGGTTTAAAGATCCTCACCGTCAGATTTATCG |
| 6_hypothetical | M20 | Tb927.11.4630_for | GGGGACAAGTTTGTACAAAAAAGCAGGCTATGCAACGGCACCAGTTATATGTT | Tb927.11.4630_rev | GGGGACCACTTTGTACAAGAAAGCTGGGTTTACTCCACATCCAGCGCTCT |
| 6_hypothetical | M21 | Tb927.7.1200_for | GGGGACAAGTTTGTACAAAAAAGCAGGCTATGTCTGTCACTTGCTGCGC | Tb927.7.1200_rev | GGGGACCACTTTGTACAAGAAAGCTGGGTTCATTTCGATCCCTCTTTTGGAGC |
| 6_hypothetical | M22 | Tb927.10.7980_for | GGGGACAAGTTTGTACAAAAAAGCAGGCTATGGCAGCCCAAACACCAC | Tb927.10.7980_rev | GGGGACCACTTTGTACAAGAAAGCTGGGTTCATCGTGGAATGAGGTCGTAGA |
| 6_hypothetical | M23 | Tb927.11.6440_for | GGGGACAAGTTTGTACAAAAAAGCAGGCTATGCCTCCAAAAAACAATCCGAAAC | Tb927.11.6440_rev | GGGGACCACTTTGTACAAGAAAGCTGGGTCTACCACCTATTAGGGAGAGCGG |
| 6_hypothetical | M24 | Tb927.3.2270_for | GGGGACAAGTTTGTACAAAAAAGCAGGCTATGTCCGCCATAGAACAGGCT | Tb927.3.2270_rev | GGGGACCACTTTGTACAAGAAAGCTGGGTTTAACTTGTTACCCACCGGAAAGC |
| 6_hypothetical | N1 | Tb927.11.13170_for | GGGGACAAGTTTGTACAAAAAAGCAGGCTATGAACCGCAGAGTGCCGA | Tb927.11.13170_rev | GGGGACCACTTTGTACAAGAAAGCTGGGTTCATTGCACCGGCGGGTT |
| 6_hypothetical | N2 | Tb927.7.6340_for | GGGGACAAGTTTGTACAAAAAAGCAGGCTATGTTCCATTCCAAAACCGCGC | Tb927.7.6340_rev | GGGGACCACTTTGTACAAGAAAGCTGGGTTTAATTCGTCCGTTCATGATAAGAAACAGC |
| 6_hypothetical | N3 | Tb927.1.560_for | GGGGACAAGTTTGTACAAAAAAGCAGGCTATGTTGTTCGTCGCTTCCAGC | Tb927.1.560_rev | GGGGACCACTTTGTACAAGAAAGCTGGGTTCAACTTGCAGCCCTCAATTTCC |
| 6_hypothetical | N4 | Tb927.3.2390_for | GGGGACAAGTTTGTACAAAAAAGCAGGCTATGAAAATATTACGCGATGTTTACAGGC | Tb927.3.2390_rev | GGGGACCACTTTGTACAAGAAAGCTGGGTTCATTCCCCAACAAGGCTGTAATAATT |
| 6_hypothetical | N5 | Tb927.6.1830_for | GGGGACAAGTTTGTACAAAAAAGCAGGCTATGGAGTTTGAGACGGAGCGC | Tb927.6.1830_rev | GGGGACCACTTTGTACAAGAAAGCTGGGTTTATTGTTGCCTATTCATCTCAAAAACATC |
| 6_hypothetical | N6 | Tb927.11.4710_for | GGGGACAAGTTTGTACAAAAAAGCAGGCTATGCCACGTCAGACTACTTTGTC | Tb927.11.4710_rev | GGGGACCACTTTGTACAAGAAAGCTGGGTTCATGTCTTCAGAGCCGTGAATCT |
| 6_hypothetical | N7 | Tb927.8.3110_for | GGGGACAAGTTTGTACAAAAAAGCAGGCTATGCAGAAGCTCTTATCGCCAAG | Tb927.8.3110_rev | GGGGACCACTTTGTACAAGAAAGCTGGGTCTACCGCTTGGCAAAGCGA |
| 6_hypothetical | N8 | Tb927.7.7070_for | GGGGACAAGTTTGTACAAAAAAGCAGGCTATGCCCTCGAACAATAGCAGC | Tb927.7.7070_rev | GGGGACCACTTTGTACAAGAAAGCTGGGTTTAAGATCCGCGGACGGTGT |
| 6_hypothetical | N9 | Tb927.1.4470_for | GGGGACAAGTTTGTACAAAAAAGCAGGCTATGATAAAAAACGCGTCGCCAC | Tb927.1.4470_rev | GGGGACCACTTTGTACAAGAAAGCTGGGTCTATGCATTTGATGAGGCTGCAGT |
| 6_hypothetical | N10 | Tb927.11.4850_for | GGGGACAAGTTTGTACAAAAAAGCAGGCTATGCTTCGGCCGAGATCGG | Tb927.11.4850_rev | GGGGACCACTTTGTACAAGAAAGCTGGGTCTACCGAAGCCCAGTGTGTTTT |
| 6_hypothetical | N11 | Tb927.10.6560_for | GGGGACAAGTTTGTACAAAAAAGCAGGCTATGTCCGGGCCTGGGGTT | Tb927.10.6560_rev | GGGGACCACTTTGTACAAGAAAGCTGGGTCTAATCACTTATTATGAATGTCTCCTTTAC |
| 6_hypothetical | N12 | Tb927.8.5020_for | GGGGACAAGTTTGTACAAAAAAGCAGGCTATGGAAATGGATAGGTTGACACGC | Tb927.8.5020_rev | GGGGACCACTTTGTACAAGAAAGCTGGGTCTATTTGTTCTTCCGCTTCCTTTTTCG |
| 6_hypothetical | N13 | Tb927.10.5990_for | GGGGACAAGTTTGTACAAAAAAGCAGGCTATGTCATCTTCCCGTGTCACATCC | Tb927.10.5990_rev | GGGGACCACTTTGTACAAGAAAGCTGGGTCTACCGATGCTGGGAATATAAGGAGC |
| 6_hypothetical | N14 | Tb927.3.4420_for | GGGGACAAGTTTGTACAAAAAAGCAGGCTATGAGTATCTTTTACAACTTCCTCCCCG | Tb927.3.4420_rev | GGGGACCACTTTGTACAAGAAAGCTGGGTTCACATTTTGCTCGCATTTTCCAG |
| 6_hypothetical | N15 | Tb927.8.2750_for | GGGGACAAGTTTGTACAAAAAAGCAGGCTATGCGGTGTTCAGCGCGC | Tb927.8.2750_rev | GGGGACCACTTTGTACAAGAAAGCTGGGTCTACCCCAACTCATCACCCGC |
| 6_hypothetical | N16 | Tb927.4.3360_for | GGGGACAAGTTTGTACAAAAAAGCAGGCTATGAAGGTAGCTGCGGTTGGA | Tb927.4.3360_rev | GGGGACCACTTTGTACAAGAAAGCTGGGTTCACATCATAACCCCCCACTGAC |
| 6_hypothetical | N17 | Tb927.11.12980_for | GGGGACAAGTTTGTACAAAAAAGCAGGCTATGCTCCCCTCGCTCATCC | Tb927.11.12980_rev | GGGGACCACTTTGTACAAGAAAGCTGGGTTTACTTCTTACCTGCTCTCTTCCGC |
| 6_hypothetical | N18 | Tb927.8.6370_for | GGGGACAAGTTTGTACAAAAAAGCAGGCTATGAAAGAAAAAAAAGAGGTGAATTCAAGC | Tb927.8.6370_rev | GGGGACCACTTTGTACAAGAAAGCTGGGTTTATTTTATGAGTAACTCATCGGGGACG |
| 6_hypothetical | N19 | Tb927.6.4200_for | GGGGACAAGTTTGTACAAAAAAGCAGGCTATGCCGATATGCCCTGGACT | Tb927.6.4200_rev | GGGGACCACTTTGTACAAGAAAGCTGGGTTCATGCTGCAACCGTTTCTCTC |
| 6_hypothetical | N20 | Tb927.11.12970_for | GGGGACAAGTTTGTACAAAAAAGCAGGCTATGGGGGACGGAGAGCAGG | Tb927.11.12970_rev | GGGGACCACTTTGTACAAGAAAGCTGGGTTCATTTGTTTGTCACCGCAGCC |
| 6_hypothetical | N21 | Tb927.4.5070_for | GGGGACAAGTTTGTACAAAAAAGCAGGCTATGTCTACTCATGAAGTCGACGTACT | Tb927.4.5070_rev | GGGGACCACTTTGTACAAGAAAGCTGGGTTTACACATTCATTCCCCAAGGCAAA |
| 6_hypothetical | N22 | Tb927.11.8810_for | GGGGACAAGTTTGTACAAAAAAGCAGGCTATGATAAAACGAAGACAACACACAGATGT | Tb927.11.8810_rev | GGGGACCACTTTGTACAAGAAAGCTGGGTCTAACTAAGAGTACTTCCAGAGTCGATTTC |
| 6_hypothetical | N23 | Tb927.11.2220_for | GGGGACAAGTTTGTACAAAAAAGCAGGCTATGCAGCGGACGCTTCGC | Tb927.11.2220_rev | GGGGACCACTTTGTACAAGAAAGCTGGGTTCAGACTCGTCGCTCTGGG |
| 6_hypothetical | N24 | Tb927.6.2430_for | GGGGACAAGTTTGTACAAAAAAGCAGGCTATGGCTACAGAAGCGGAGGAA | Tb927.6.2430_rev | GGGGACCACTTTGTACAAGAAAGCTGGGTTCAAAACACTGTAAATGTGACGTACAAC |
| 6_hypothetical | O1 | Tb927.8.3610_for | GGGGACAAGTTTGTACAAAAAAGCAGGCTATGTCGTGGAGGGCTCTTTCG | Tb927.8.3610_rev | GGGGACCACTTTGTACAAGAAAGCTGGGTCTAAAGGTAAACCACGGCCTCG |
| 6_hypothetical | O2 | Tb927.10.8680_for | GGGGACAAGTTTGTACAAAAAAGCAGGCTATGACTTTTGAGCAATGCTCTGGA | Tb927.10.8680_rev | GGGGACCACTTTGTACAAGAAAGCTGGGTTTACGGTTGCCGCTGCAAC |
| 6_hypothetical | O3 | Tb927.11.11260_for | GGGGACAAGTTTGTACAAAAAAGCAGGCTATGGCGCTTTCATTAGAGGACAG | Tb927.11.11260_rev | GGGGACCACTTTGTACAAGAAAGCTGGGTCTAGCTGGTTGCAGTGAGATGTG |
| 6_hypothetical | O4 | Tb927.8.7960_for | GGGGACAAGTTTGTACAAAAAAGCAGGCTATGACAAATGAACCATCTGAAGCAGC | Tb927.8.7960_rev | GGGGACCACTTTGTACAAGAAAGCTGGGTTTACCCAACATCATCGCCATCATC |
| 6_hypothetical | O5 | Tb927.5.2190_for | GGGGACAAGTTTGTACAAAAAAGCAGGCTATGATGGATATTACTGCGTTGTGCG | Tb927.5.2190_rev | GGGGACCACTTTGTACAAGAAAGCTGGGTCTACTGGTGCACTGCGTCATAG |
| 6_hypothetical | O6 | Tb927.10.4920_for | GGGGACAAGTTTGTACAAAAAAGCAGGCTATGGCCTTTCCTCATCGTGAAC | Tb927.10.4920_rev | GGGGACCACTTTGTACAAGAAAGCTGGGTCTAGTTGATATCTGTTTTGTTCCGCTTCC |
| 6_hypothetical | O7 | Tb927.4.2570_for | GGGGACAAGTTTGTACAAAAAAGCAGGCTATGACTAACAATCATGTAGAAAATAATTGC | Tb927.4.2570_rev | GGGGACCACTTTGTACAAGAAAGCTGGGTCTATGAGAGGTCTGTTTTCTTGTGAGC |
| 6_hypothetical | O10 | Tb927.10.5220_for | GGGGACAAGTTTGTACAAAAAAGCAGGCTATGCGTAGGGGACCACAATGG | Tb927.10.5220_rev | GGGGACCACTTTGTACAAGAAAGCTGGGTCTAAAGTGTGGGTTGCACCACA |
| 6_hypothetical | O11 | Tb927.10.3180_for | GGGGACAAGTTTGTACAAAAAAGCAGGCTATGACAAAACAGCAGATGTGCCT | Tb927.10.3180_rev | GGGGACCACTTTGTACAAGAAAGCTGGGTTTATTGAAAATGAGACGACGCACCG |
| 6_hypothetical | O12 | Tb927.11.5620_for | GGGGACAAGTTTGTACAAAAAAGCAGGCTATGAACGTTTCACTTGGTACGGC | Tb927.11.5620_rev | GGGGACCACTTTGTACAAGAAAGCTGGGTCTATCGCATCACATTGAAGCTACCG |
| 6_hypothetical | O13 | Tb927.4.3070_for | GGGGACAAGTTTGTACAAAAAAGCAGGCTATGCTGCGCCATACATGCA | Tb927.4.3070_rev | GGGGACCACTTTGTACAAGAAAGCTGGGTTTATTTTGTGTGCGAGTACAAATCCCC |
| 6_hypothetical | O14 | Tb927.7.7340_for | GGGGACAAGTTTGTACAAAAAAGCAGGCTATGGCACTGGTGGCGTCC | Tb927.7.7340_rev | GGGGACCACTTTGTACAAGAAAGCTGGGTTCAGATCAACGCCATGCGTTC |

| | | | | | |
|---|---|---|---|---|---|
| 6_hypothetical | O15 | Tb927.2.2920_for | GGGGACAAGTTTGTACAAAAAAGCAGGCTATGGCTGCACATATATCGGAACAC | Tb927.2.2920_rev | GGGGACCACTTTGTACAAGAAAGCTGGGTTCATGCACGTTCATCATTCCTTCTAC |
| 6_hypothetical | O16 | Tb927.11.1230_for | GGGGACAAGTTTGTACAAAAAAGCAGGCTATGGTGACCACACAACCTAACCA | Tb927.11.1230_rev | GGGGACCACTTTGTACAAGAAAGCTGGGTTTATGAAACCTTGTTAGAACCTACGGTG |
| 6_hypothetical | O17 | Tb927.10.9910_for | GGGGACAAGTTTGTACAAAAAAGCAGGCTATGCAGCATCAACTTAGAGAAGAGC | Tb927.10.9910_rev | GGGGACCACTTTGTACAAGAAAGCTGGGTTTAGTTCTCCACTGCCTCCATCC |
| 6_hypothetical | O18 | Tb927.10.15460_for | GGGGACAAGTTTGTACAAAAAAGCAGGCTATGAGTTCATGGGAATCGTGTGAGG | Tb927.10.15460_rev | GGGGACCACTTTGTACAAGAAAGCTGGGTTTAAAAGACAAATATATTCCAATGGAACGT |
| 6_hypothetical | O19 | Tb927.7.3460_for | GGGGACAAGTTTGTACAAAAAAGCAGGCTATGCTTTTCCGGAGTGTTTCATGT | Tb927.7.3460_rev | GGGGACCACTTTGTACAAGAAAGCTGGGTTCATAACCTCTCCATCGATCTCGATG |
| 6_hypothetical | O20 | Tb927.10.14030_for | GGGGACAAGTTTGTACAAAAAAGCAGGCTATGCTCAGCCTCACTTTTCTTCTTT | Tb927.10.14030_rev | GGGGACCACTTTGTACAAGAAAGCTGGGTCTACCAGAAAAACGCTTGTTTTTCTAAATC |
| 6_hypothetical | O21 | Tb927.6.3830_for | GGGGACAAGTTTGTACAAAAAAGCAGGCTATGCCACTGGAAAACAATAGCAACG | Tb927.6.3830_rev | GGGGACCACTTTGTACAAGAAAGCTGGGTTTACCGCACCTCAAATATTCCCCA |
| 6_hypothetical | O22 | Tb927.4.1550_for | GGGGACAAGTTTGTACAAAAAAGCAGGCTATGGAGCTGCGTTGCTTTTTAGC | Tb927.4.1550_rev | GGGGACCACTTTGTACAAGAAAGCTGGGTTCAGTGGCGGTAGTACACAACC |
| 6_hypothetical | O23 | Tb927.5.920_for | GGGGACAAGTTTGTACAAAAAAGCAGGCTATGCTTCGGTTTCTCCACCGC | Tb927.5.920_rev | GGGGACCACTTTGTACAAGAAAGCTGGGTCTAACGTGAACGATTGTTGTCGTCC |
| 6_hypothetical | O24 | Tb927.11.1190_for | GGGGACAAGTTTGTACAAAAAAGCAGGCTATGAGCACCTCAATTGAGGGAGA | Tb927.11.1190_rev | GGGGACCACTTTGTACAAGAAAGCTGGGTTCAGTCAACCTTCGGATCCCC |
| 6_hypothetical | P1 | Tb927.11.1760_for | GGGGACAAGTTTGTACAAAAAAGCAGGCTATGCTGGAACTGGACGGACG | Tb927.11.1760_rev | GGGGACCACTTTGTACAAGAAAGCTGGGTTCACCCCCTCACGCAGTG |
| 6_hypothetical | P2 | Tb927.4.2490_for | GGGGACAAGTTTGTACAAAAAAGCAGGCTATGAGCAACAATGTGGAGTTGTTGC | Tb927.4.2490_rev | GGGGACCACTTTGTACAAGAAAGCTGGGTTCAACCTTCACGCGCAAGTG |
| 6_hypothetical | P3 | Tb927.4.3440_for | GGGGACAAGTTTGTACAAAAAAGCAGGCTATGCACGCTCATGCAGCTG | Tb927.4.3440_rev | GGGGACCACTTTGTACAAGAAAGCTGGGTTCAGTCGCTAACGAACTCCAGT |
| 6_hypothetical | P4 | Tb927.3.4150_for | GGGGACAAGTTTGTACAAAAAAGCAGGCTATGAGCGACGTCTGCGCG | Tb927.3.4150_rev | GGGGACCACTTTGTACAAGAAAGCTGGGTTCAGCCCAGCAAATAACGCTT |
| 6_hypothetical | P5 | Tb927.11.3860_for | GGGGACAAGTTTGTACAAAAAAGCAGGCTATGAGGTGGCCACGCGGG | Tb927.11.3860_rev | GGGGACCACTTTGTACAAGAAAGCTGGGTTTATTTGCTTGAAGCCGTCGGG |
| 6_hypothetical | P6 | Tb927.11.9880_for | GGGGACAAGTTTGTACAAAAAAGCAGGCTATGCGCTACGAGAGTGTCTTACT | Tb927.11.9880_rev | GGGGACCACTTTGTACAAGAAAGCTGGGTTTAGTCAAATCCATTGAAGCCTGTTAAGA |
| 6_hypothetical | P7 | Tb927.10.12150_for | GGGGACAAGTTTGTACAAAAAAGCAGGCTATGAGGGATGTACTAGCGAGCTT | Tb927.10.12150_rev | GGGGACCACTTTGTACAAGAAAGCTGGGTTTACGAGCGAAATAATCCCCGC |
| 6_hypothetical | P8 | Tb927.10.15310_for | GGGGACAAGTTTGTACAAAAAAGCAGGCTATGTATCCAAAAAATAAGCCCCTGCC | Tb927.10.15310_rev | GGGGACCACTTTGTACAAGAAAGCTGGGTTTACAAAGGGGTTGTAGTCTCTTCCT |
| 6_hypothetical | P9 | Tb927.5.2620_for | GGGGACAAGTTTGTACAAAAAAGCAGGCTATGCACGTACTTGGCAGAGATCC | Tb927.5.2620_rev | GGGGACCACTTTGTACAAGAAAGCTGGGTTTACAGCTGTTCAAGAGAAACAGCA |
| 6_hypothetical | P10 | Tb927.9.2815_for | GGGGACAAGTTTGTACAAAAAAGCAGGCTATGGAAATCCGCATGGTGCC | Tb927.9.2815_rev | GGGGACCACTTTGTACAAGAAAGCTGGGTTCACTCATCAACTGCTGGAGTTTC |
| 6_hypothetical | P11 | Tb927.2.5890_for | GGGGACAAGTTTGTACAAAAAAGCAGGCTATGTGGCGGATCACGCGG | Tb927.2.5890_rev | GGGGACCACTTTGTACAAGAAAGCTGGGTTCATTGACCACACGGCAGGC |
| 6_hypothetical | P12 | Tb927.11.16940_for | GGGGACAAGTTTGTACAAAAAAGCAGGCTATGCCAACACGCTCCTCAGT | Tb927.11.16940_rev | GGGGACCACTTTGTACAAGAAAGCTGGGTCTATCTCCCGGAACCACTGAGAC |
| 6_hypothetical | P13 | Tb927.10.7220_for | GGGGACAAGTTTGTACAAAAAAGCAGGCTATGCTCAGTGACAAGGAGGTGA | Tb927.10.7220_rev | GGGGACCACTTTGTACAAGAAAGCTGGGTTCACCTGTGAAGAAAGTCGATAATAAGC |
| 6_hypothetical | P14 | Tb927.9.6450_for | GGGGACAAGTTTGTACAAAAAAGCAGGCTATGGTGTCCCGGGTATTCAGTG | Tb927.9.6450_rev | GGGGACCACTTTGTACAAGAAAGCTGGGTTCACTCTCCGATAGAGTACATGAACG |
| 6_hypothetical | P15 | Tb927.3.3980_for | GGGGACAAGTTTGTACAAAAAAGCAGGCTATGAAGTGCTGGTCTCCGCA | Tb927.3.3980_rev | GGGGACCACTTTGTACAAGAAAGCTGGGTTTACCCAATGGTGGTGTATGGACT |
| 6_hypothetical | P16 | Tb927.10.11660_for | GGGGACAAGTTTGTACAAAAAAGCAGGCTATGCTGCCGGCTCCGGTA | Tb927.10.11660_rev | GGGGACCACTTTGTACAAGAAAGCTGGGTCTAGTACAACTCATCAAAGTCATCCAATGC |
| 6_hypothetical | P17 | Tb927.7.2640_for | GGGGACAAGTTTGTACAAAAAAGCAGGCTATGAGCACCACTGCACTGTCT | Tb927.7.2640_rev | GGGGACCACTTTGTACAAGAAAGCTGGGTCTATTTGTCGGCCTCCCGC |
| 6_hypothetical | P18 | Tb927.10.3400_for | GGGGACAAGTTTGTACAAAAAAGCAGGCTATGGAGGCATCCGTGACAACA | Tb927.10.3400_rev | GGGGACCACTTTGTACAAGAAAGCTGGGTTCACCAACGGAGCGCAAG |
| 6_hypothetical | P19 | Tb927.10.11840_for | GGGGACAAGTTTGTACAAAAAAGCAGGCTATGTACACTACGACAGCCAAACAG | Tb927.10.11840_rev | GGGGACCACTTTGTACAAGAAAGCTGGGTTTAAACGGTATTAGTTGTGTTAACCGGA |
| 6_hypothetical | P20 | Tb927.4.3850_for | GGGGACAAGTTTGTACAAAAAAGCAGGCTATGCAAGAAGAAGCTATTTCCTCCG | Tb927.4.3850_rev | GGGGACCACTTTGTACAAGAAAGCTGGGTTTAATTAAGAAGAGAGTCCCCTTTGAAACA |
| 6_hypothetical | P21 | Tb927.11.14230_for | GGGGACAAGTTTGTACAAAAAAGCAGGCTATGAATGATAGGGAGTTGCGGGC | Tb927.11.14230_rev | GGGGACCACTTTGTACAAGAAAGCTGGGTCTACGACATAATGGCATGAAGCTGT |
| 6_hypothetical | P22 | Tb927.11.5110_for | GGGGACAAGTTTGTACAAAAAAGCAGGCTATGGCACTGAATGTCGTGAGC | Tb927.11.5110_rev | GGGGACCACTTTGTACAAGAAAGCTGGGTTCAAAAGTAACCGAGCATGTTCACC |
| 6_hypothetical | P23 | Tb927.7.1530_for | GGGGACAAGTTTGTACAAAAAAGCAGGCTATGGAGACTCACAACACCGAGC | Tb927.7.1530_rev | GGGGACCACTTTGTACAAGAAAGCTGGGTTTAAGTACCATCACATAAGCTGGGACA |
| 6_hypothetical | P24 | Tb927.6.5070_for | GGGGACAAGTTTGTACAAAAAAGCAGGCTATGCGAGCAAACCCAATTGAGC | Tb927.6.5070_rev | GGGGACCACTTTGTACAAGAAAGCTGGGTCTAATACGCCCTCCACAGTGGA |
| 6_known | A1 | Tb927.11.7950_for | GGGGACAAGTTTGTACAAAAAAGCAGGCTATGGCCCTTGAAAAGATCGACTG | Tb927.11.7950_rev | GGGGACCACTTTGTACAAGAAAGCTGGGTTCATTCACCGGAACAGGAGATGT |
| 6_known | A2 | Tb927.9.10710_for | GGGGACAAGTTTGTACAAAAAAGCAGGCTATGAATCGCCACAGGAAGTTTTACA | Tb927.9.10710_rev | GGGGACCACTTTGTACAAGAAAGCTGGGTTCAAAGCATGATCGACTCCACCA |
| 6_known | A3 | Tb927.10.10770_for | GGGGACAAGTTTGTACAAAAAAGCAGGCTATGCCGACGGAGACGTTATCA | Tb927.10.10770_rev | GGGGACCACTTTGTACAAGAAAGCTGGGTTCAGTGCACTGTGTTAACGTACG |
| 6_known | A4 | Tb927.3.2330_for | GGGGACAAGTTTGTACAAAAAAGCAGGCTATGAAGTCTCGACAACCGACGA | Tb927.3.2330_rev | GGGGACCACTTTGTACAAGAAAGCTGGGTCTAACAAATGCTTTGGTTACCCTCGG |
| 6_known | A5 | Tb927.10.14550_for | GGGGACAAGTTTGTACAAAAAAGCAGGCTATGCACGGCATGAATTTCGGC | Tb927.10.14550_rev | GGGGACCACTTTGTACAAGAAAGCTGGGTTTACATGGTGGGGCCTCCTC |
| 6_known | A6 | Tb927.7.1510_for | GGGGACAAGTTTGTACAAAAAAGCAGGCTATGTCTAAGCGTAAGAATTTTAACACGC | Tb927.7.1510_rev | GGGGACCACTTTGTACAAGAAAGCTGGGTTCACTCGCTTCCGCCGCC |
| 6_known | A7 | Tb927.11.4130_for | GGGGACAAGTTTGTACAAAAAAGCAGGCTATGCACATCAACGTCTCCGG | Tb927.11.4130_rev | GGGGACCACTTTGTACAAGAAAGCTGGGTTCACCTCCTCTCCAGTTGCTTC |
| 6_known | A8 | Tb927.11.7760_for | GGGGACAAGTTTGTACAAAAAAGCAGGCTATGGCAGTGATTATGTTCCCTCGT | Tb927.11.7760_rev | GGGGACCACTTTGTACAAGAAAGCTGGGTTTATAATAGGGAAAAGCAAGAAACGAAGAG |
| 6_known | A9 | Tb927.10.13000_for | GGGGACAAGTTTGTACAAAAAAGCAGGCTATGTTGGAAGCGCTCCGAAAG | Tb927.10.13000_rev | GGGGACCACTTTGTACAAGAAAGCTGGGTCTAGGTTCCCTGTCCAGTGTTACC |
| 6_known | A10 | Tb927.2.1820_for | GGGGACAAGTTTGTACAAAAAAGCAGGCTATGGAGTTTAGTACGGAACCTCAGG | Tb927.2.1820_rev | GGGGACCACTTTGTACAAGAAAGCTGGGTCTACATCAGTTCCGGTTTCCCCT |
| 6_known | A11 | Tb927.3.3900_for | GGGGACAAGTTTGTACAAAAAAGCAGGCTATGAAGCGCATCCCCGTAAAG | Tb927.3.3900_rev | GGGGACCACTTTGTACAAGAAAGCTGGGTCTATTTTAGTTTAGTCCCTGCCTTAAGCA |
| 6_known | A12 | Tb927.10.16040_for | GGGGACAAGTTTGTACAAAAAAGCAGGCTATGAAGCATGACTTTCCAAAGCGT | Tb927.10.16040_rev | GGGGACCACTTTGTACAAGAAAGCTGGGTTCAGCTGAAGAGGCCAACCG |
| 6_known | A13 | Tb927.3.1360_for | GGGGACAAGTTTGTACAAAAAAGCAGGCTATGTGCTCTGTTGCACCGTT | Tb927.3.1360_rev | GGGGACCACTTTGTACAAGAAAGCTGGGTTTACTTAAGGATACGACTTAAAAGTGCACG |
| 6_known | A14 | Tb927.8.4820_for | GGGGACAAGTTTGTACAAAAAAGCAGGCTATGCACGTCTACACCATCGACC | Tb927.8.4820_rev | GGGGACCACTTTGTACAAGAAAGCTGGGTTCAAGTCAAGGTCCCCTTCCC |
| 6_known | A15 | Tb927.3.1570_for | GGGGACAAGTTTGTACAAAAAAGCAGGCTATGAGTGGCCATTGGGAGGG | Tb927.3.1570_rev | GGGGACCACTTTGTACAAGAAAGCTGGGTTCACTCGTACACAATCTTTACGTAGTCA |
| 6_known | A16 | Tb927.11.11720_for | GGGGACAAGTTTGTACAAAAAAGCAGGCTATGAATAGGTGCAATCCTATCTCTCTCA | Tb927.11.11720_rev | GGGGACCACTTTGTACAAGAAAGCTGGGTCTACCTCAGCTTCAGCTCCCG |
| 6_known | A17 | Tb927.3.3670_for | GGGGACAAGTTTGTACAAAAAAGCAGGCTATGCGAGGAAGAGGGAGAGGT | Tb927.3.3670_rev | GGGGACCACTTTGTACAAGAAAGCTGGGTCTAAGAAGATTTCTTCCCCTTCCCCT |
| 6_known | A18 | Tb927.4.4310_for | GGGGACAAGTTTGTACAAAAAAGCAGGCTATGCACCGGAAGACCATCGT | Tb927.4.4310_rev | GGGGACCACTTTGTACAAGAAAGCTGGGTTTAGTACCTCCTACACGGCCTTTT |
| 6_known | A19 | Tb927.9.13510_for | GGGGACAAGTTTGTACAAAAAAGCAGGCTATGAAGCTCATTACACGGAACAATTTTG | Tb927.9.13510_rev | GGGGACCACTTTGTACAAGAAAGCTGGGTCTACTTCTCAGTCAAATGTTTGATCCGA |
| 6_known | A20 | Tb927.11.2360_for | GGGGACAAGTTTGTACAAAAAAGCAGGCTATGCCAACCCTCGCTGTTG | Tb927.11.2360_rev | GGGGACCACTTTGTACAAGAAAGCTGGGTCTACTGGGCAAACTGAACATTCAGC |
| 6_known | A21 | Tb927.11.9640_for | GGGGACAAGTTTGTACAAAAAAGCAGGCTATGAGCACCGGGCTGCCC | Tb927.11.9640_rev | GGGGACCACTTTGTACAAGAAAGCTGGGTTCACGTTGCGGCTGCCTG |
| 6_known | A22 | Tb927.4.2380_for | GGGGACAAGTTTGTACAAAAAAGCAGGCTATGTCGTCACACGGCAAATTAGT | Tb927.4.2380_rev | GGGGACCACTTTGTACAAGAAAGCTGGGTCTAAGGAAACAATTGTCCCTGCTGC |
| 6_known | A23 | Tb927.7.5040_for | GGGGACAAGTTTGTACAAAAAAGCAGGCTATGGAGCAAACCAGCAAACCC | Tb927.7.5040_rev | GGGGACCACTTTGTACAAGAAAGCTGGGTCTACACAGCGGAGTACCGCG |
| 6_known | A24 | Tb927.10.5880_for | GGGGACAAGTTTGTACAAAAAAGCAGGCTATGCTCCAGAGCAAAGTTGTGG | Tb927.10.5880_rev | GGGGACCACTTTGTACAAGAAAGCTGGGTCTAGCTTTCAGTTGGCGCTTGC |
| 6_known | B1 | Tb927.4.1990_for | GGGGACAAGTTTGTACAAAAAAGCAGGCTATGCGCCGTTTGTCAGTTACT | Tb927.4.1990_rev | GGGGACCACTTTGTACAAGAAAGCTGGGTTCACAACCTCCGCAACAGC |
| 6_known | B2 | Tb927.2.4370_for | GGGGACAAGTTTGTACAAAAAAGCAGGCTATGACGAAGTCGGCACTTGC | Tb927.2.4370_rev | GGGGACCACTTTGTACAAGAAAGCTGGGTCTACATTTGAATACGTACGGGACCAA |
| 6_known | B3 | Tb927.9.1780_for | GGGGACAAGTTTGTACAAAAAAGCAGGCTATGCCACCGCAGCGGAAG | Tb927.9.1780_rev | GGGGACCACTTTGTACAAGAAAGCTGGGTTTATGATATTCCTTCACCAAGAACAGTGAG |
| 6_known | B4 | Tb927.10.3490_for | GGGGACAAGTTTGTACAAAAAAGCAGGCTATGCGTATGATAGACAGTGAAGATGTG | Tb927.10.3490_rev | GGGGACCACTTTGTACAAGAAAGCTGGGTCTACGCTAAGTGCAAATGGCGG |
| 6_known | B5 | Tb927.8.1940_for | GGGGACAAGTTTGTACAAAAAAGCAGGCTATGGCGATGCTTCTCAACAATCTATATG | Tb927.8.1940_rev | GGGGACCACTTTGTACAAGAAAGCTGGGTTTAGTCGGCCTTTCCATAACGATATATCC |
| 6_known | B6 | Tb927.5.1140_for | GGGGACAAGTTTGTACAAAAAAGCAGGCTATGGCATTACGACCGTTCCCG | Tb927.5.1140_rev | GGGGACCACTTTGTACAAGAAAGCTGGGTTCAGGAAAGAAGCCCCTTCTCTG |
| 6_known | B7 | Tb927.6.1770_for | GGGGACAAGTTTGTACAAAAAAGCAGGCTATGGCAAAACCACTCAGCGG | Tb927.6.1770_rev | GGGGACCACTTTGTACAAGAAAGCTGGGTTTACGCTTTCAGTTTCTCAAGATGGT |
| 6_known | B8 | Tb927.11.1820_for | GGGGACAAGTTTGTACAAAAAAGCAGGCTATGTCCTCCGCAGTGGGAG | Tb927.11.1820_rev | GGGGACCACTTTGTACAAGAAAGCTGGGTTCACGTAGCCGACATCAAATAAAGATC |
| 6_known | B9 | Tb927.11.16880_for | GGGGACAAGTTTGTACAAAAAAGCAGGCTATGCTCTTTGAACCTTCGGGGG | Tb927.11.16880_rev | GGGGACCACTTTGTACAAGAAAGCTGGGTTCATTTCTTACCGCGCGGC |
| 6_known | B10 | Tb927.11.7560_for | GGGGACAAGTTTGTACAAAAAAGCAGGCTATGGCAGCGAAGGAGTACGA | Tb927.11.7560_rev | GGGGACCACTTTGTACAAGAAAGCTGGGTTTACAGCGTCTTAAACTCCAAGTCAC |
| 6_known | B11 | Tb927.11.14590_for | GGGGACAAGTTTGTACAAAAAAGCAGGCTATGTCACTAAACCCTAACGCTCCT | Tb927.11.14590_rev | GGGGACCACTTTGTACAAGAAAGCTGGGTTTACCCAGAAGCTTCATTCTGCAC |
| 6_known | B12 | Tb927.11.8380_for | GGGGACAAGTTTGTACAAAAAAGCAGGCTATGCTATTAGTGGTGTCGGCCA | Tb927.11.8380_rev | GGGGACCACTTTGTACAAGAAAGCTGGGTTCACGCCGTCCAGACATCC |
| 6_known | B13 | Tb927.8.2550_for | GGGGACAAGTTTGTACAAAAAAGCAGGCTATGCTCCGCATCACCTTACCA | Tb927.8.2550_rev | GGGGACCACTTTGTACAAGAAAGCTGGGTTTACGCTCCTTGTGGAGGGATG |
| 6_known | B14 | Tb927.10.3140_for | GGGGACAAGTTTGTACAAAAAAGCAGGCTATGTACACCGTGGCTCCTGC | Tb927.10.3140_rev | GGGGACCACTTTGTACAAGAAAGCTGGGTTCATTCAAGACACGCGCTCTC |
| 6_known | B15 | Tb927.10.12640_for | GGGGACAAGTTTGTACAAAAAAGCAGGCTATGAACAATGACCTTGAGCGATCAA | Tb927.10.12640_rev | GGGGACCACTTTGTACAAGAAAGCTGGGTTCAACCCTGTGGTTTCCTCAGC |
| 6_known | B16 | Tb11.02.5420_for | GGGGACAAGTTTGTACAAAAAAGCAGGCTATGATTTTCCTCGTGATATCCACCG | Tb11.02.5420_rev | GGGGACCACTTTGTACAAGAAAGCTGGGTTCACGCCGTCCAGACATCC |
| 6_known | B17 | Tb927.10.1630_for | GGGGACAAGTTTGTACAAAAAAGCAGGCTATGCCACCTAAAAACAGGCAAGAT | Tb927.10.1630_rev | GGGGACCACTTTGTACAAGAAAGCTGGGTCTACTCGTTGCTTTCATCATCTTCGT |
| 6_known | B18 | Tb927.3.4560_for | GGGGACAAGTTTGTACAAAAAAGCAGGCTATGCCGGAAGGTGAACAGGC | Tb927.3.4560_rev | GGGGACCACTTTGTACAAGAAAGCTGGGTTCACGCCAATTTCTGGACAAGAG |
| 6_known | B19 | Tb927.5.4420_for | GGGGACAAGTTTGTACAAAAAAGCAGGCTATGTCTTCCAGAGCTGAGGAAGG | Tb927.5.4420_rev | GGGGACCACTTTGTACAAGAAAGCTGGGTCTAATAACGCCTTTGAGTCGACCG |
| 6_known | B20 | Tb927.8.8140_for | GGGGACAAGTTTGTACAAAAAAGCAGGCTATGCGTGTGGAGGACCCG | Tb927.8.8140_rev | GGGGACCACTTTGTACAAGAAAGCTGGGTTCAGCAACCTGAACGTTTAGACC |
| 6_known | B21 | Tb927.6.3570_for | GGGGACAAGTTTGTACAAAAAAGCAGGCTATGATAAAACTGTGCCGATCGCT | Tb927.6.3570_rev | GGGGACCACTTTGTACAAGAAAGCTGGGTTCAAGCCTCCAATGCCCACA |
| 6_known | B22 | Tb927.11.16090_for | GGGGACAAGTTTGTACAAAAAAGCAGGCTATGCCTCCGAGCCAGCTG | Tb927.11.16090_rev | GGGGACCACTTTGTACAAGAAAGCTGGGTTTACTTTCTGCGGAGTATAGGCGG |
| 6_known | B23 | Tb927.6.1510_for | GGGGACAAGTTTGTACAAAAAAGCAGGCTATGCGCCGTTGGGGAGCG | Tb927.6.1510_rev | GGGGACCACTTTGTACAAGAAAGCTGGGTTCAGGAACGTGGTCTACCACAC |
| 6_known | B24 | Tb927.4.1090_for | GGGGACAAGTTTGTACAAAAAAGCAGGCTATGGGCTACAAGTTTCACTTTTACCG | Tb927.4.1090_rev | GGGGACCACTTTGTACAAGAAAGCTGGGTTTAGTCAGCTTTTATGGAGTTGTAAATAGT |
| 6_known | C1 | Tb927.6.1760_for | GGGGACAAGTTTGTACAAAAAAGCAGGCTATGGCTTTCCGTACCTCAACTATTTTC | Tb927.6.1760_rev | GGGGACCACTTTGTACAAGAAAGCTGGGTTTACGACGAGCCGCTCCC |
| 6_known | C2 | Tb927.5.4380_for | GGGGACAAGTTTGTACAAAAAAGCAGGCTATGGATCCACCTAATGTACCTCCC | Tb927.5.4380_rev | GGGGACCACTTTGTACAAGAAAGCTGGGTTTAAATGCGCTTAAGACGCGGT |
| 6_known | C3 | Tb927.11.12480_for | GGGGACAAGTTTGTACAAAAAAGCAGGCTATGTCATCGTCAGAAGCGGTGA | Tb927.11.12480_rev | GGGGACCACTTTGTACAAGAAAGCTGGGTTCACCGCGGCCCGTTTTC |
| 6_known | C4 | Tb927.10.6540_for | GGGGACAAGTTTGTACAAAAAAGCAGGCTATGGGATTCTTCTCGTTTAAGAGCAAC | Tb927.10.6540_rev | GGGGACCACTTTGTACAAGAAAGCTGGGTTTAACTACAAAATTCCTTTGGTGGGACC |
| 6_known | C5 | Tb927.7.3650_for | GGGGACAAGTTTGTACAAAAAAGCAGGCTATGTCCAATTCGAGTGGCAAAACA | Tb927.7.3650_rev | GGGGACCACTTTGTACAAGAAAGCTGGGTTCAAATATTGGGGAAAGAAACACCCA |
| 6_known | C6 | Tb927.4.880_for | GGGGACAAGTTTGTACAAAAAAGCAGGCTATGAGTCACCATATCCAACTGCCT | Tb927.4.880_rev | GGGGACCACTTTGTACAAGAAAGCTGGGTTCATGTAGCCGCTGAAAAAGTCAT |
| 6_known | C7 | Tb927.7.5650_for | GGGGACAAGTTTGTACAAAAAAGCAGGCTATGGCTGAAAAAGTACAGGTGGTG | Tb927.7.5650_rev | GGGGACCACTTTGTACAAGAAAGCTGGGTTTATACTTGACACCTCCCGTCGAT |
| 6_known | C8 | Tb927.7.2580_for | GGGGACAAGTTTGTACAAAAAAGCAGGCTATGCGTCACGGAGCATCCC | Tb927.7.2580_rev | GGGGACCACTTTGTACAAGAAAGCTGGGTCTATGCATCTATCGGCGAAGCG |
| 6_known | C9 | Tb927.4.2560_for | GGGGACAAGTTTGTACAAAAAAGCAGGCTATGACCTTCAATGGAAGGTTTCGTG | Tb927.4.2560_rev | GGGGACCACTTTGTACAAGAAAGCTGGGTCTAGTCTATCTCGGAGGAGAGATCGG |
| 6_known | C10 | Tb927.6.2030_for | GGGGACAAGTTTGTACAAAAAAGCAGGCTATGTCCCTCTTGGAAAATTGCCAC | Tb927.6.2030_rev | GGGGACCACTTTGTACAAGAAAGCTGGGTTCAGCGGCTGACAATGGGC |
| 6_known | C11 | Tb927.11.13220_for | GGGGACAAGTTTGTACAAAAAAGCAGGCTATGACTTACAAACATATGGCAAACCACC | Tb927.11.13220_rev | GGGGACCACTTTGTACAAGAAAGCTGGGTTCACTGGCCTTCCATTAAAAGACAAT |
| 6_known | C12 | Tb927.7.710_for | GGGGACAAGTTTGTACAAAAAAGCAGGCTATGCCTGCTCCGGCGATC | Tb927.7.710_rev | GGGGACCACTTTGTACAAGAAAGCTGGGTTTAGTCAACATCATCGGGGACTGG |
| 6_known | C13 | Tb927.10.14770_for | GGGGACAAGTTTGTACAAAAAAGCAGGCTATGGAATCCACTGACACAGAAGTGA | Tb927.10.14770_rev | GGGGACCACTTTGTACAAGAAAGCTGGGTCTACACCTTCTTGACTTGGGACG |
| 6_known | C14 | Tb927.10.9080_for | GGGGACAAGTTTGTACAAAAAAGCAGGCTATGAGTTTCGCCCCTACGCG | Tb927.10.9080_rev | GGGGACCACTTTGTACAAGAAAGCTGGGTTTAAGTACTACCGCTCTCCTGCAC |
| 6_known | C15 | Tb927.11.3130_for | GGGGACAAGTTTGTACAAAAAAGCAGGCTATGAGTTCTGTACTTGACGGGCT | Tb927.11.3130_rev | GGGGACCACTTTGTACAAGAAAGCTGGGTTCAAATGGTTACCCTGGGCTGC |
| 6_known | C16 | Tb927.4.5200_for | GGGGACAAGTTTGTACAAAAAAGCAGGCTATGAGTGTGCTGGGATCAGGC | Tb927.4.5200_rev | GGGGACCACTTTGTACAAGAAAGCTGGGTTCAGGAGGAAGCGACGCG |
| 6_known | C17 | Tb927.9.1970_for | GGGGACAAGTTTGTACAAAAAAGCAGGCTATGATGAGCAACCATGCAAAGCT | Tb927.9.1970_rev | GGGGACCACTTTGTACAAGAAAGCTGGGTTTATTCACTCAGTACCTTAACAAACTCATT |
| 6_known | C18 | Tb927.11.14380_for | GGGGACAAGTTTGTACAAAAAAGCAGGCTATGTTTGCCTGTCGTCGGC | Tb927.11.14380_rev | GGGGACCACTTTGTACAAGAAAGCTGGGTTTACCACCGTCTTTTGCGTCG |
| 6_known | C19 | Tb927.11.1030_for | GGGGACAAGTTTGTACAAAAAAGCAGGCTATGAGCGGATATGGCGAATCTATGC | Tb927.11.1030_rev | GGGGACCACTTTGTACAAGAAAGCTGGGTTCACCGCGTTCTCCCCCC |
| 6_known | C20 | Tb927.7.7180_for | GGGGACAAGTTTGTACAAAAAAGCAGGCTATGGCGAAGCGAAGCCGC | Tb927.7.7180_rev | GGGGACCACTTTGTACAAGAAAGCTGGGTTTACCAACCGTGACGATTCTTAGC |
| 6_known | C21 | Tb927.3.4190_for | GGGGACAAGTTTGTACAAAAAAGCAGGCTATGACTCGCTTCTCGTGTGTTG | Tb927.3.4190_rev | GGGGACCACTTTGTACAAGAAAGCTGGGTTCAGTCCACCTTGATGCTTCCA |
| 6_known | C22 | Tb927.7.7110_for | GGGGACAAGTTTGTACAAAAAAGCAGGCTATGGTGAAGCGACTCGGCT | Tb927.7.7110_rev | GGGGACCACTTTGTACAAGAAAGCTGGGTTTACCAACCGTGACGATTCTTAGC |
| 6_known | C23 | Tb927.8.3680_for | GGGGACAAGTTTGTACAAAAAAGCAGGCTATGGGAGACAATCCACTGTCTGT | Tb927.8.3680_rev | GGGGACCACTTTGTACAAGAAAGCTGGGTTTAATCACGACTTATAGCGAAACGTAACA |
| 6_known | C24 | Tb927.10.7360_for | GGGGACAAGTTTGTACAAAAAAGCAGGCTATGGAGGGCAATGGCAGCT | Tb927.10.7360_rev | GGGGACCACTTTGTACAAGAAAGCTGGGTTCACTGAAGATACTTCGCACGC |
| 6_known | D1 | Tb927.9.6310_for | GGGGACAAGTTTGTACAAAAAAGCAGGCTATGCCGGATTCACCTGAAACTG | Tb927.9.6310_rev | GGGGACCACTTTGTACAAGAAAGCTGGGTTCACTTCCCCTCTTCCGTTTCA |
| 6_known | D2 | Tb927.11.190_for | GGGGACAAGTTTGTACAAAAAAGCAGGCTATGGCGATGCGACGTTTGT | Tb927.11.190_rev | GGGGACCACTTTGTACAAGAAAGCTGGGTTCATTTCCTTTTGGTAGCGTTGCG |
| 6_known | D3 | Tb927.4.5180_for | GGGGACAAGTTTGTACAAAAAAGCAGGCTATGAAAAACATTGAGGTTAACCCCCA | Tb927.4.5180_rev | GGGGACCACTTTGTACAAGAAAGCTGGGTTCAGGAGAGATTGCTCAACTTCTCC |
| 6_known | D4 | Tb927.8.5570_for | GGGGACAAGTTTGTACAAAAAAGCAGGCTATGATCGCCGACGCGGTG | Tb927.8.5570_rev | GGGGACCACTTTGTACAAGAAAGCTGGGTTCATCTAACTATATAAATTGACAACCCGTA |
| 6_known | D5 | Tb927.9.1670_for | GGGGACAAGTTTGTACAAAAAAGCAGGCTATGAGCACACGTCTGTGCG | Tb927.9.1670_rev | GGGGACCACTTTGTACAAGAAAGCTGGGTCTATCTCACGTCATATGACGACCCT |
| 6_known | D6 | Tb927.8.7220_for | GGGGACAAGTTTGTACAAAAAAGCAGGCTATGAAAAACAACGACGCTAACCATATG | Tb927.8.7220_rev | GGGGACCACTTTGTACAAGAAAGCTGGGTTCAGGGGATGTTACTTAACTTCTCCTC |
| 6_known | D7 | Tb927.11.15470_for | GGGGACAAGTTTGTACAAAAAAGCAGGCTATGCGGCAGTGCATATCGAG | Tb927.11.15470_rev | GGGGACCACTTTGTACAAGAAAGCTGGGTTCAACAGGATGCCAGGTTGC |
| 6_known | D8 | Tb927.8.7520_for | GGGGACAAGTTTGTACAAAAAAGCAGGCTATGCACCGTTGGCGTCCA | Tb927.8.7520_rev | GGGGACCACTTTGTACAAGAAAGCTGGGTTCACGTACAGTACAGCGCAGC |
| 6_known | D9 | Tb927.1.1540_for | GGGGACAAGTTTGTACAAAAAAGCAGGCTATGACAGAAAGAGAGCTTGTCACG | Tb927.1.1540_rev | GGGGACCACTTTGTACAAGAAAGCTGGGTTCAGTTCATGCTAATATCATCACCGTTATC |
| 6_known | D10 | Tb927.9.6880_for | GGGGACAAGTTTGTACAAAAAAGCAGGCTATGGAAGGAAAAAATGAGAGAGACCAATAC | Tb927.9.6880_rev | GGGGACCACTTTGTACAAGAAAGCTGGGTCTACTTTCTTCCACGGTCCATACCC |
| 6_known | D11 | Tb927.1.2820_for | GGGGACAAGTTTGTACAAAAAAGCAGGCTATGGGTCGACAGGAAGATGGTT | Tb927.1.2820_rev | GGGGACCACTTTGTACAAGAAAGCTGGGTTCAGGACCTCTTGGGCGC |
| 6_known | D13 | Tb927.11.8730_for | GGGGACAAGTTTGTACAAAAAAGCAGGCTATGCGCAAGGGGGGAAATG | Tb927.11.8730_rev | GGGGACCACTTTGTACAAGAAAGCTGGGTCTAGTAACGCACTGCGCTCG |
| 6_known | D15 | Tb927.4.670_for | GGGGACAAGTTTGTACAAAAAAGCAGGCTATGGATTTTGTGGAAGTGCTTCCTG | Tb927.4.670_rev | GGGGACCACTTTGTACAAGAAAGCTGGGTTCAACGAGGTGAGACACTGCG |
| 6_known | D16 | Tb927.7.1610_for | GGGGACAAGTTTGTACAAAAAAGCAGGCTATGCGCCCAGCTCCGATG | Tb927.7.1610_rev | GGGGACCACTTTGTACAAGAAAGCTGGGTTCAAAGTTTGTTGAAGTCCTCTGTGC |
| 6_known | D17 | Tb927.11.7460_for | GGGGACAAGTTTGTACAAAAAAGCAGGCTATGTCGAGGATGTGGCTGACC | Tb927.11.7460_rev | GGGGACCACTTTGTACAAGAAAGCTGGGTTTACAGATCGTCCATGGGTTGTGG |
| 6_known | D19 | Tb927.5.3170_for | GGGGACAAGTTTGTACAAAAAAGCAGGCTATGCGCACAGGAAAGGGTCG | Tb927.5.3170_rev | GGGGACCACTTTGTACAAGAAAGCTGGGTTTACTTTTCAAAGAGCGTGGTGAGAG |
| 6_known | D20 | Tb927.10.11940_for | GGGGACAAGTTTGTACAAAAAAGCAGGCTATGACGCTATGGAGTGCGACG | Tb927.10.11940_rev | GGGGACCACTTTGTACAAGAAAGCTGGGTTTACCGTATGGAATAAAACTTGTTTTCCTT |
| 6_known | D21 | Tb927.10.14980_for | GGGGACAAGTTTGTACAAAAAAGCAGGCTATGACGGATGCCCCAAGGG | Tb927.10.14980_rev | GGGGACCACTTTGTACAAGAAAGCTGGGTTCAAAGCTCCTCAATTTCATAGACGAAA |
| 6_known | D22 | Tb927.5.1100_for | GGGGACAAGTTTGTACAAAAAAGCAGGCTATGGACTGCGGCGCCGGT | Tb927.5.1100_rev | GGGGACCACTTTGTACAAGAAAGCTGGGTTCACAGTAACATAGACTGCAGGCC |
| 6_known | D23 | Tb927.4.1820_for | GGGGACAAGTTTGTACAAAAAAGCAGGCTATGACAGAAGTTAATGATGTTGCCTCTG | Tb927.4.1820_rev | GGGGACCACTTTGTACAAGAAAGCTGGGTCTACTGCTGAAGCTGACTGTGG |
| 6_known | D24 | Tb927.10.5250_for | GGGGACAAGTTTGTACAAAAAAGCAGGCTATGTCTGGCACTAATCACCTCCG | Tb927.10.5250_rev | GGGGACCACTTTGTACAAGAAAGCTGGGTTCATCTCTGTTTCTGCGTGGC |
| 6_known | E1 | Tb927.11.3120_for | GGGGACAAGTTTGTACAAAAAAGCAGGCTATGTCGACAATCTACAATTTCAAAACGG | Tb927.11.3120_rev | GGGGACCACTTTGTACAAGAAAGCTGGGTTCAGCGTCTGTCGCGTCT |
| 6_known | E2 | Tb927.11.5000_for | GGGGACAAGTTTGTACAAAAAAGCAGGCTATGGATACCAATGCACGCCG | Tb927.11.5000_rev | GGGGACCACTTTGTACAAGAAAGCTGGGTTCACGCCTGCCCCAGGAG |
| 6_known | E3 | Tb927.6.3800_for | GGGGACAAGTTTGTACAAAAAAGCAGGCTATGTTGGCCCGTCGTGTATGC | Tb927.6.3800_rev | GGGGACCACTTTGTACAAGAAAGCTGGGTTTACTCCTTCTTTTCACCAGATGCC |

| | | | | | |
|---|---|---|---|---|---|
| 6_known | E4 | Tb927.9.9580_for | GGGGACAAGTTTGTACAAAAAAGCAGGCTATGGCCGAACCCGTGCTT | Tb927.9.9580_rev | GGGGACCACTTTGTACAAGAAAGCTGGGTTCACGCTGTCTTCTTTTCCTGC |
| 6_known | E5 | Tb927.11.16790_for | GGGGACAAGTTTGTACAAAAAAGCAGGCTATGGATGCGTACGAAACACTTGG | Tb927.11.16790_rev | GGGGACCACTTTGTACAAGAAAGCTGGGTTCATCGTTTTGTTGCACCATCTCG |
| 6_known | E6 | Tb927.11.960_for | GGGGACAAGTTTGTACAAAAAAGCAGGCTATGATTAGGGTGCCCTCCCG | Tb927.11.960_rev | GGGGACCACTTTGTACAAGAAAGCTGGGTTCATACAGCTCCCGAGTCGG |
| 6_known | E7 | Tb927.10.7620_for | GGGGACAAGTTTGTACAAAAAAGCAGGCTATGCACCGGCGCTGTCCC | Tb927.10.7620_rev | GGGGACCACTTTGTACAAGAAAGCTGGGTTTACGTTATGGAAACGGGGCG |
| 6_known | E9 | Tb927.6.3750_for | GGGGACAAGTTTGTACAAAAAAGCAGGCTATGTTGGCCCGCCGTGTA | Tb927.6.3750_rev | GGGGACCACTTTGTACAAGAAAGCTGGGTTTACTCCTTCTTTTCACCAGATGCC |
| 6_known | E10 | Tb927.11.11480_for | GGGGACAAGTTTGTACAAAAAAGCAGGCTATGCCTGACAGCGAGGGT | Tb927.11.11480_rev | GGGGACCACTTTGTACAAGAAAGCTGGGTCTACGAAACGGGCACGAACTC |
| 6_known | E11 | Tb927.1.1960_for | GGGGACAAGTTTGTACAAAAAAGCAGGCTATGATAGTACTTCCCACTTTTGATTTCCC | Tb927.1.1960_rev | GGGGACCACTTTGTACAAGAAAGCTGGGTCTACTTGCGCTTATTCATAGCACAGA |
| 6_known | E12 | Tb927.7.2100_for | GGGGACAAGTTTGTACAAAAAAGCAGGCTATGTCAGTTCAACAGGAGCCGG | Tb927.7.2100_rev | GGGGACCACTTTGTACAAGAAAGCTGGGTTTATTCCCATTCCGTCGTCCCC |
| 6_known | E13 | Tb927.3.1610_for | GGGGACAAGTTTGTACAAAAAAGCAGGCTATGCAACCAACGCGAAGATGC | Tb927.3.1610_rev | GGGGACCACTTTGTACAAGAAAGCTGGGTCTAAACAGGTGAAGTAAAAAAAGGATGACT |
| 6_known | E14 | Tb927.8.1690_for | GGGGACAAGTTTGTACAAAAAAGCAGGCTATGAATGAATATCATATCATAGAAAAGATG | Tb927.8.1690_rev | GGGGACCACTTTGTACAAGAAAGCTGGGTTCATGCTGGTCCCGCCAC |
| 6_known | E16 | Tb927.7.5560_for | GGGGACAAGTTTGTACAAAAAAGCAGGCTATGTGCGGGATATTCGGTTACATTC | Tb927.7.5560_rev | GGGGACCACTTTGTACAAGAAAGCTGGGTTCATTGAACCGTTACACTCTTCGC |
| 6_known | E17 | Tb927.3.4720_for | GGGGACAAGTTTGTACAAAAAAGCAGGCTATGGAGCGACTCATCTCGGTC | Tb927.3.4720_rev | GGGGACCACTTTGTACAAGAAAGCTGGGTCTAAATAAGAGCAAACTCTCGAACACTGT |
| 6_known | E19 | Tb927.5.1760_for | GGGGACAAGTTTGTACAAAAAAGCAGGCTATGCGGAGTTATGTATTTGGCCAA | Tb927.5.1760_rev | GGGGACCACTTTGTACAAGAAAGCTGGGTCTAGAATGGGCACACGTATGAAGC |
| 6_known | E20 | Tb927.9.11070_for | GGGGACAAGTTTGTACAAAAAAGCAGGCTATGGTGATCCGACCTGCAACC | Tb927.9.11070_rev | GGGGACCACTTTGTACAAGAAAGCTGGGTTTACAACACAATGGGATGATCCGC |
| 6_known | E21 | Tb927.5.1830_for | GGGGACAAGTTTGTACAAAAAAGCAGGCTATGATGTGTACCGCAACCGTTATG | Tb927.5.1830_rev | GGGGACCACTTTGTACAAGAAAGCTGGGTCTAGTTCTGAGGATCTATAAGACCTTCAGC |
| 6_known | E22 | Tb927.11.11330_for | GGGGACAAGTTTGTACAAAAAAGCAGGCTATGACATACGAAGGCGCCATTG | Tb927.11.11330_rev | GGGGACCACTTTGTACAAGAAAGCTGGGTTTAGTCAACCTCCTCGACTTTAGGC |
| 6_known | E23 | Tb927.11.1660_for | GGGGACAAGTTTGTACAAAAAAGCAGGCTATGTCTGACCAGATGCGGCG | Tb927.11.1660_rev | GGGGACCACTTTGTACAAGAAAGCTGGGTTCAGCAAGGGCCCTTCGC |
| 6_known | E24 | Tb927.10.2210_for | GGGGACAAGTTTGTACAAAAAAGCAGGCTATGGGGGGAAGTTATTCCACGG | Tb927.10.2210_rev | GGGGACCACTTTGTACAAGAAAGCTGGGTTTACGCAACTCGCTCGTACAG |
| 6_known | F1 | Tb927.1.1420_for | GGGGACAAGTTTGTACAAAAAAGCAGGCTATGGGTGCCTTTCCGTCTCA | Tb927.1.1420_rev | GGGGACCACTTTGTACAAGAAAGCTGGGTCTACTGTTGAGCTACTTGCCCCG |
| 6_known | F2 | Tb927.1.1500_for | GGGGACAAGTTTGTACAAAAAAGCAGGCTATGGGAGGTGCGCCTTCT | Tb927.1.1500_rev | GGGGACCACTTTGTACAAGAAAGCTGGGTCTACGACAGCGAGCTTGGAATG |
| 6_known | F3 | Tb927.11.4490_for | GGGGACAAGTTTGTACAAAAAAGCAGGCTATGAGGCGTTTGCTCGGC | Tb927.11.4490_rev | GGGGACCACTTTGTACAAGAAAGCTGGGTTCAAAACAGATCATCATAGTCGTCCAC |
| 6_known | F4 | Tb927.11.540_for | GGGGACAAGTTTGTACAAAAAAGCAGGCTATGGTCCGCCACGCTCAT | Tb927.11.540_rev | GGGGACCACTTTGTACAAGAAAGCTGGGTTCACACCTCCGCCCCTGC |
| 6_known | F5 | Tb927.10.9050_for | GGGGACAAGTTTGTACAAAAAAGCAGGCTATGACTGTAGTCGAGTCTCCTTTCA | Tb927.10.9050_rev | GGGGACCACTTTGTACAAGAAAGCTGGGTTCATCCAGCGGATGGCTCA |
| 6_known | F6 | Tb927.8.6650_for | GGGGACAAGTTTGTACAAAAAAGCAGGCTATGACAGACCACCAAGTCATCTGC | Tb927.8.6650_rev | GGGGACCACTTTGTACAAGAAAGCTGGGTTCAACCTACACGCATGGTGGG |
| 6_known | F7 | Tb927.8.3310_for | GGGGACAAGTTTGTACAAAAAAGCAGGCTATGTCGCTAAAGCGCCCTCG | Tb927.8.3310_rev | GGGGACCACTTTGTACAAGAAAGCTGGGTTCATTTACCGAACATAAATCCCTTCAGC |
| 6_known | F8 | Tb927.4.2310_for | GGGGACAAGTTTGTACAAAAAAGCAGGCTATGTCGGAGGAAATAATTACAAAGCTTCG | Tb927.4.2310_rev | GGGGACCACTTTGTACAAGAAAGCTGGGTTTAGGGTTGGCATCGACCCA |
| 6_known | F9 | Tb927.1.2600_for | GGGGACAAGTTTGTACAAAAAAGCAGGCTATGGAAGTACGCGATGTGAATACAC | Tb927.1.2600_rev | GGGGACCACTTTGTACAAGAAAGCTGGGTCTAACATTCTCCGTCATCACAATCAACC |
| 6_known | F10 | Tb927.7.2250_for | GGGGACAAGTTTGTACAAAAAAGCAGGCTATGCGCCTTTCCTCTGTTTGC | Tb927.7.2250_rev | GGGGACCACTTTGTACAAGAAAGCTGGGTCTAATCCTCGAGCATATCGTAGCAATC |
| 6_known | F11 | Tb927.3.1590_for | GGGGACAAGTTTGTACAAAAAAGCAGGCTATGTACCATCGTGGCTACGGC | Tb927.3.1590_rev | GGGGACCACTTTGTACAAGAAAGCTGGGTTCATCTTTTCGCAGTAACGGTCC |
| 6_known | F12 | Tb927.10.7700_for | GGGGACAAGTTTGTACAAAAAAGCAGGCTATGCGCTGCTTTCGAAAAAAGG | Tb927.10.7700_rev | GGGGACCACTTTGTACAAGAAAGCTGGGTTTATAGTTTGTGTCGCGCCGC |
| 6_known | F13 | Tb927.8.6170_for | GGGGACAAGTTTGTACAAAAAAGCAGGCTATGTCGTTTAATGATAACCTGGCCAC | Tb927.8.6170_rev | GGGGACCACTTTGTACAAGAAAGCTGGGTTCACAAATGGGACCGCTTCG |
| 6_known | F14 | Tb927.11.9110_for | GGGGACAAGTTTGTACAAAAAAGCAGGCTATGCAAAGATTGGCAGGGAAGC | Tb927.11.9110_rev | GGGGACCACTTTGTACAAGAAAGCTGGGTTTAGAAGTTACCGACACGCTTCG |
| 6_known | F15 | Tb927.1.4780_for | GGGGACAAGTTTGTACAAAAAAGCAGGCTATGGCGGCCTTCTTCGGT | Tb927.1.4780_rev | GGGGACCACTTTGTACAAGAAAGCTGGGTTCATGACGCGCTGAAAGAGG |
| 6_known | F16 | Tb927.6.1940_for | GGGGACAAGTTTGTACAAAAAAGCAGGCTATGATTGTACCTCCTCCCTGGGG | Tb927.6.1940_rev | GGGGACCACTTTGTACAAGAAAGCTGGGTTCAAGCAACTGGCGCCGC |
| 6_known | F17 | Tb927.5.3940_for | GGGGACAAGTTTGTACAAAAAAGCAGGCTATGTCGTCTTCAGACATCGCG | Tb927.5.3940_rev | GGGGACCACTTTGTACAAGAAAGCTGGGTTCAGTTAAGCAGGAGGGCGT |
| 6_known | F18 | Tb927.8.3060_for | GGGGACAAGTTTGTACAAAAAAGCAGGCTATGCTCAAGAGAGTTATTGCGAGG | Tb927.8.3060_rev | GGGGACCACTTTGTACAAGAAAGCTGGGTTCAATTGCCAGACCTTTTCTTTCCT |
| 6_known | F19 | Tb927.6.160_for | GGGGACAAGTTTGTACAAAAAAGCAGGCTATGAATCAGAACGGTGGTAGTGTGG | Tb927.6.160_rev | GGGGACCACTTTGTACAAGAAAGCTGGGTCTAATTGTTAACTCCGTTGTTATTGTTACC |
| 6_known | F20 | Tb927.3.3590_for | GGGGACAAGTTTGTACAAAAAAGCAGGCTATGGTAACTGCGATGGCAAACA | Tb927.3.3590_rev | GGGGACCACTTTGTACAAGAAAGCTGGGTCTATGCAGCAGCCTTCCGTAG |
| 6_known | F21 | Tb927.6.4710_for | GGGGACAAGTTTGTACAAAAAAGCAGGCTATGACGCACGTTGGTGCTG | Tb927.6.4710_rev | GGGGACCACTTTGTACAAGAAAGCTGGGTCTACATCCCAACACGGCGC |
| 6_known | F22 | Tb927.8.2520_for | GGGGACAAGTTTGTACAAAAAAGCAGGCTATGCGTGACACAGGTCGC | Tb927.8.2520_rev | GGGGACCACTTTGTACAAGAAAGCTGGGTTCACCCCACTGTTTGCTTCCT |
| 6_known | F23 | Tb927.11.5340_for | GGGGACAAGTTTGTACAAAAAAGCAGGCTATGAGGGTGATGAGACACCTCTC | Tb927.11.5340_rev | GGGGACCACTTTGTACAAGAAAGCTGGGTTCACTCAAACAGCACCTTGCG |
| 6_known | F24 | Tb927.6.1140_for | GGGGACAAGTTTGTACAAAAAAGCAGGCTATGTGGCTCTTTCTCGGGGC | Tb927.6.1140_rev | GGGGACCACTTTGTACAAGAAAGCTGGGTTTACTTTGGGTTTTGTTTGCTCAGC |
| 6_known | G1 | Tb927.2.240_for | GGGGACAAGTTTGTACAAAAAAGCAGGCTATGAATCAGAACGGCGGTAGTGT | Tb927.2.240_rev | GGGGACCACTTTGTACAAGAAAGCTGGGTTTACCTTCGGCCCACAGCA |
| 6_known | G3 | Tb927.9.5490_for | GGGGACAAGTTTGTACAAAAAAGCAGGCTATGAGTGGATCGTTGAACGTGG | Tb927.9.5490_rev | GGGGACCACTTTGTACAAGAAAGCTGGGTCTACTCCAGAAGTCCTCGTCGG |
| 6_known | G4 | Tb927.10.9820_for | GGGGACAAGTTTGTACAAAAAAGCAGGCTATGTTGCGGCGTGTCACC | Tb927.10.9820_rev | GGGGACCACTTTGTACAAGAAAGCTGGGTTCACACCCATATGTCGATTTCATCG |
| 6_known | G5 | Tb927.1.420_for | GGGGACAAGTTTGTACAAAAAAGCAGGCTATGAATCAGAATGGCGGTAGTGTG | Tb927.1.420_rev | GGGGACCACTTTGTACAAGAAAGCTGGGTTTACCTTCGGCCCACAGCA |
| 6_known | G6 | Tb927.11.7410_for | GGGGACAAGTTTGTACAAAAAAGCAGGCTATGCTTCACCATAAATTAGATACAGTGAGC | Tb927.11.7410_rev | GGGGACCACTTTGTACAAGAAAGCTGGGTTTAGAAGGGAAGCATCGATCGGT |
| 6_known | G7 | Tb927.4.250_for | GGGGACAAGTTTGTACAAAAAAGCAGGCTATGAATCAGAACGGCGGTAGTGT | Tb927.4.250_rev | GGGGACCACTTTGTACAAGAAAGCTGGGTTTACCTTCGGCCCGCAGC |
| 6_known | G8 | Tb927.4.3390_for | GGGGACAAGTTTGTACAAAAAAGCAGGCTATGTGGCGGCGGACATGG | Tb927.4.3390_rev | GGGGACCACTTTGTACAAGAAAGCTGGGTCTATTGCCAGCAGCGAGCC |
| 6_known | G10 | Tb927.7.4390_for | GGGGACAAGTTTGTACAAAAAAGCAGGCTATGCTCACCTTACGCGTATGTG | Tb927.7.4390_rev | GGGGACCACTTTGTACAAGAAAGCTGGGTTCAGCAAGAGCCCTCACTGG |
| 6_known | G11 | Tb927.11.11290_for | GGGGACAAGTTTGTACAAAAAAGCAGGCTATGACATACGAAGGCGCCATTG | Tb927.11.11290_rev | GGGGACCACTTTGTACAAGAAAGCTGGGTTTACTCTGTGTTTGCGTTGATAGCC |
| 6_known | G12 | Tb927.10.15520_for | GGGGACAAGTTTGTACAAAAAAGCAGGCTATGAGTGATGTGAGTAAGTTCGTGAGG | Tb927.10.15520_rev | GGGGACCACTTTGTACAAGAAAGCTGGGTCTACGACGACGGGTCTGCT |
| 6_known | G13 | Tb927.3.3340_for | GGGGACAAGTTTGTACAAAAAAGCAGGCTATGGGCTGTGGAATCGCAAC | Tb927.3.3340_rev | GGGGACCACTTTGTACAAGAAAGCTGGGTCTACTCACACCATGCAGTGCG |
| 6_known | G14 | Tb927.10.11360_for | GGGGACAAGTTTGTACAAAAAAGCAGGCTATGTCCTCCAAATTAGGGACCGC | Tb927.10.11360_rev | GGGGACCACTTTGTACAAGAAAGCTGGGTTTACAGGCGGAGGCTGGG |
| 6_known | G15 | Tb927.10.14270_for | GGGGACAAGTTTGTACAAAAAAGCAGGCTATGCGACGTAGTGGATTCGTCG | Tb927.10.14270_rev | GGGGACCACTTTGTACAAGAAAGCTGGGTCTACTCGGGTGGTGAAGGTACG |
| 6_known | G16 | Tb927.11.14730_for | GGGGACAAGTTTGTACAAAAAAGCAGGCTATGCAGGGAGAACCGCGT | Tb927.11.14730_rev | GGGGACCACTTTGTACAAGAAAGCTGGGTCTAGCTGGTATCAGTGACTATTGGACG |
| 6_known | G17 | Tb927.8.590_for | GGGGACAAGTTTGTACAAAAAAGCAGGCTATGACACGGGTGTCCACGT | Tb927.8.590_rev | GGGGACCACTTTGTACAAGAAAGCTGGGTCTAAAGCTTCGACTGGAGTGGAAG |
| 6_known | G18 | Tb927.8.6970_for | GGGGACAAGTTTGTACAAAAAAGCAGGCTATGCTCCGATACAACGTGTTCTAC | Tb927.8.6970_rev | GGGGACCACTTTGTACAAGAAAGCTGGGTTTAAACTGCAGCAGAGGCAATATGG |
| 6_known | G19 | Tb927.4.200_for | GGGGACAAGTTTGTACAAAAAAGCAGGCTATGAATCAGAACGGCGGTAGTGT | Tb927.4.200_rev | GGGGACCACTTTGTACAAGAAAGCTGGGTCTATGCATTGTTACCACCTCTATTGTTAAC |
| 6_known | G20 | Tb927.8.5770_for | GGGGACAAGTTTGTACAAAAAAGCAGGCTATGCTACACGCTAGTGGTAATGGC | Tb927.8.5770_rev | GGGGACCACTTTGTACAAGAAAGCTGGGTTCACTGGCCTCGGGGCCG |
| 6_known | G21 | Tb927.1.1370_for | GGGGACAAGTTTGTACAAAAAAGCAGGCTATGCAGTATCGTGAGTATCGCGT | Tb927.1.1370_rev | GGGGACCACTTTGTACAAGAAAGCTGGGTTCACTCTTCCGCCTCACCATC |
| 6_known | G22 | Tb927.10.12370_for | GGGGACAAGTTTGTACAAAAAAGCAGGCTATGGGTCTTCTAACCACTGGCG | Tb927.10.12370_rev | GGGGACCACTTTGTACAAGAAAGCTGGGTTCACCCTTCGCGTTGTCTTTTAC |
| 6_known | G23 | Tb927.10.1210_for | GGGGACAAGTTTGTACAAAAAAGCAGGCTATGGACGACGACGTGGGG | Tb927.10.1210_rev | GGGGACCACTTTGTACAAGAAAGCTGGGTTCACACTGACCCCAGCTCTTC |
| 6_known | G24 | Tb927.6.2690_for | GGGGACAAGTTTGTACAAAAAAGCAGGCTATGTCGAAAACAATTTGCAAATGTTGTG | Tb927.6.2690_rev | GGGGACCACTTTGTACAAGAAAGCTGGGTTTAATGGTAAAGCTGGTCGTAGAAGAGC |
| 6_known | H1 | Tb927.9.15470_for | GGGGACAAGTTTGTACAAAAAAGCAGGCTATGGCGGATATGAAGAAGGTAACAGT | Tb927.9.15470_rev | GGGGACCACTTTGTACAAGAAAGCTGGGTTCATGCGGTTCCGACATGTTC |
| 6_known | H2 | Tb927.11.11410_for | GGGGACAAGTTTGTACAAAAAAGCAGGCTATGAGAATGCAAATACTCATTTTACTGTTA | Tb927.11.11410_rev | GGGGACCACTTTGTACAAGAAAGCTGGGTTCAAAAAGTGACAACAAAGATCATAGCAG |
| 6_known | H3 | Tb927.7.580_for | GGGGACAAGTTTGTACAAAAAAGCAGGCTATGGCACCATTTAGCGGGAAAC | Tb927.7.580_rev | GGGGACCACTTTGTACAAGAAAGCTGGGTTTACACTGAACTGGCCAATGGC |
| 6_known | H4 | Tb927.1.2110_for | GGGGACAAGTTTGTACAAAAAAGCAGGCTATGAGCTCGCGTAACGGC | Tb927.1.2110_rev | GGGGACCACTTTGTACAAGAAAGCTGGGTTTAGGACAGCTCAATAGAGTTACCAAGA |
| 6_known | H5 | Tb927.4.4050_for | GGGGACAAGTTTGTACAAAAAAGCAGGCTATGACAGTGTTTGAGCTATGCAAGG | Tb927.4.4050_rev | GGGGACCACTTTGTACAAGAAAGCTGGGTTCACTGTTTCCCGTCAGATACATCA |
| 6_known | H6 | Tb927.11.6740_for | GGGGACAAGTTTGTACAAAAAAGCAGGCTATGGGCAAGAAGGGAACGCT | Tb927.11.6740_rev | GGGGACCACTTTGTACAAGAAAGCTGGGTTCATGTGGATTCACTTATCTGCACAA |
| 6_known | H7 | Tb927.5.1650_for | GGGGACAAGTTTGTACAAAAAAGCAGGCTATGAGCGGCAGTGCAACTAC | Tb927.5.1650_rev | GGGGACCACTTTGTACAAGAAAGCTGGGTTCATCGCTTCATCGGTTCTAAAAAGG |
| 6_known | H8 | Tb927.11.1070_for | GGGGACAAGTTTGTACAAAAAAGCAGGCTATGACAACGCATTGGGTTCGT | Tb927.11.1070_rev | GGGGACCACTTTGTACAAGAAAGCTGGGTTTAGAAACCCCTGTGCAAATCAACT |
| 6_known | H9 | Tb927.10.10880_for | GGGGACAAGTTTGTACAAAAAAGCAGGCTATGTCTTCGCTTAGTACTGCTGGA | Tb927.10.10880_rev | GGGGACCACTTTGTACAAGAAAGCTGGGTTTAACCATCGTTAGCCTTCTCCTCC |
| 6_known | H10 | Tb927.11.2710_for | GGGGACAAGTTTGTACAAAAAAGCAGGCTATGGCTGATACACTTTCACCGTG | Tb927.11.2710_rev | GGGGACCACTTTGTACAAGAAAGCTGGGTTCAGCGCCCGCCAGCGAG |
| 6_known | H11 | Tb927.10.7240_for | GGGGACAAGTTTGTACAAAAAAGCAGGCTATGGAAGAAGTCAATGCGTATTCACC | Tb927.10.7240_rev | GGGGACCACTTTGTACAAGAAAGCTGGGTCTAACGAGGAAATCGGAGCGC |
| 6_known | H12 | Tb927.1.70_for | GGGGACAAGTTTGTACAAAAAAGCAGGCTATGAATCAGAATGGTGGTTGTGTGG | Tb927.1.70_rev | GGGGACCACTTTGTACAAGAAAGCTGGGTTCACCTTCGCTTTGCTTTCATCG |
| 6_known | H13 | Tb927.7.190_for | GGGGACAAGTTTGTACAAAAAAGCAGGCTATGACGTCGAGACTGTTTGCG | Tb927.7.190_rev | GGGGACCACTTTGTACAAGAAAGCTGGGTTTATACACCAAGGCTTTTGAAAAAAGCG |
| 6_known | H14 | Tb927.2.2190_for | GGGGACAAGTTTGTACAAAAAAGCAGGCTATGGGCAAAGAAACGCGTGA | Tb927.2.2190_rev | GGGGACCACTTTGTACAAGAAAGCTGGGTTTACTTGCACATATGGTCAAAAGAATTGG |
| 6_known | H15 | Tb927.11.9920_for | GGGGACAAGTTTGTACAAAAAAGCAGGCTATGCAGATCTTCGTGAAAACTCTTACC | Tb927.11.9920_rev | GGGGACCACTTTGTACAAGAAAGCTGGGTCTACCAACCACCTCGAAGACGA |
| 6_known | H16 | Tb927.7.1970_for | GGGGACAAGTTTGTACAAAAAAGCAGGCTATGATTCAAAACAACGGCAATATGAGAAA | Tb927.7.1970_rev | GGGGACCACTTTGTACAAGAAAGCTGGGTTCATTTCCGTGTGAATGCTGGTG |
| 6_known | H17 | Tb927.4.1920_for | GGGGACAAGTTTGTACAAAAAAGCAGGCTATGAAGCGAGTGCCGTGC | Tb927.4.1920_rev | GGGGACCACTTTGTACAAGAAAGCTGGGTTTAGCCAGTGCCAGCAGCG |
| 6_known | H18 | Tb927.9.12880_for | GGGGACAAGTTTGTACAAAAAAGCAGGCTATGGATGACGGAACTGGACACC | Tb927.9.12880_rev | GGGGACCACTTTGTACAAGAAAGCTGGGTTCATATGGGACCCATAGCCCTTG |
| 6_known | H19 | Tb927.7.3780_for | GGGGACAAGTTTGTACAAAAAAGCAGGCTATGTCACGAACAGACAAGTCGTTC | Tb927.7.3780_rev | GGGGACCACTTTGTACAAGAAAGCTGGGTTCAAAAACCGAGTCCCAGCG |
| 6_known | H21 | Tb927.7.2030_for | GGGGACAAGTTTGTACAAAAAAGCAGGCTATGATTCAAAACAACGGCAATATGGG | Tb927.7.2030_rev | GGGGACCACTTTGTACAAGAAAGCTGGGTTCATTTCCGTGTGAATGCTGGTG |
| 6_known | H22 | Tb927.10.7080_for | GGGGACAAGTTTGTACAAAAAAGCAGGCTATGAGGCGCTTCGCGAGG | Tb927.10.7080_rev | GGGGACCACTTTGTACAAGAAAGCTGGGTTCAAAGAGGCCTAAAGTTCTTGTTCAG |
| 6_known | H23 | Tb927.7.6970_for | GGGGACAAGTTTGTACAAAAAAGCAGGCTATGACCAAAAAGAGTGGTCATGAGG | Tb927.7.6970_rev | GGGGACCACTTTGTACAAGAAAGCTGGGTCTATGAAGACGTTTTAGGCGCCA |
| 6_known | H24 | Tb927.10.6090_for | GGGGACAAGTTTGTACAAAAAAGCAGGCTATGCGCCGTGTCAGTCTGC | Tb927.10.6090_rev | GGGGACCACTTTGTACAAGAAAGCTGGGTTTAAACAGCGGCCGTGATGTC |
| 6_known | I1 | Tb927.3.5490_for | GGGGACAAGTTTGTACAAAAAAGCAGGCTATGACATCCCGACCCTTGACG | Tb927.3.5490_rev | GGGGACCACTTTGTACAAGAAAGCTGGGTTTACTCACGCATATGCAAATATAAATGCTT |
| 6_known | I2 | Tb927.10.9130_for | GGGGACAAGTTTGTACAAAAAAGCAGGCTATGTTGAATGAACGGCGGCC | Tb927.10.9130_rev | GGGGACCACTTTGTACAAGAAAGCTGGGTTTACAACATCATACGTGAGAAAGACGAT |
| 6_known | I3 | Tb927.10.6230_for | GGGGACAAGTTTGTACAAAAAAGCAGGCTATGTTAACTGTAACGGAGGATAAGATCTCC | Tb927.10.6230_rev | GGGGACCACTTTGTACAAGAAAGCTGGGTTTAGGCAGCGTAAATACCGTTTTTGA |
| 6_known | I4 | Tb927.1.1550_for | GGGGACAAGTTTGTACAAAAAAGCAGGCTATGACAGACGGCGTTACATTACATG | Tb927.1.1550_rev | GGGGACCACTTTGTACAAGAAAGCTGGGTTTAAGAAGCCCACGATGGGGG |
| 6_known | I5 | Tb927.7.1950_for | GGGGACAAGTTTGTACAAAAAAGCAGGCTATGATTCAAAACAACGGCAATATGAGAAA | Tb927.7.1950_rev | GGGGACCACTTTGTACAAGAAAGCTGGGTCTACTCATGACCTCGTTTCCGTG |
| 6_known | I6 | Tb927.11.4660_for | GGGGACAAGTTTGTACAAAAAAGCAGGCTATGACCTCACTCTGTCCAATTACGA | Tb927.11.4660_rev | GGGGACCACTTTGTACAAGAAAGCTGGGTTCACACGCTTTCAAGTTCATGAAGC |
| 6_known | I7 | Tb927.9.13250_for | GGGGACAAGTTTGTACAAAAAAGCAGGCTATGTTTCGTCGGTTCTGCACC | Tb927.9.13250_rev | GGGGACCACTTTGTACAAGAAAGCTGGGTTCAGTACACGTCCTTCAAGTATCTACC |
| 6_known | I8 | Tb927.7.4090_for | GGGGACAAGTTTGTACAAAAAAGCAGGCTATGCAACAACAACAGCAGCAGC | Tb927.7.4090_rev | GGGGACCACTTTGTACAAGAAAGCTGGGTCTAAGAGGAACCAACGCCCAAAT |
| 6_known | I11 | Tb927.11.7010_for | GGGGACAAGTTTGTACAAAAAAGCAGGCTATGACTCAGGAGGAATGCACGG | Tb927.11.7010_rev | GGGGACCACTTTGTACAAGAAAGCTGGGTTCACTCCTCGTCTCCTTTACAAGG |
| 6_known | I12 | Tb927.9.3650_for | GGGGACAAGTTTGTACAAAAAAGCAGGCTATGGCGAAGTGGGAATTAAAGCTG | Tb927.9.3650_rev | GGGGACCACTTTGTACAAGAAAGCTGGGTCTAAATCCCGTTTTGCTCGAGACG |
| 6_known | I13 | Tb927.7.2000_for | GGGGACAAGTTTGTACAAAAAAGCAGGCTATGATTCAAAACAACGGCAATATGGG | Tb927.7.2000_rev | GGGGACCACTTTGTACAAGAAAGCTGGGTCTACTCATGACCTCGTTTCCGTG |
| 6_known | I14 | Tb927.4.1700_for | GGGGACAAGTTTGTACAAAAAAGCAGGCTATGAACAGAACGCTACTCGGTTTAG | Tb927.4.1700_rev | GGGGACCACTTTGTACAAGAAAGCTGGGTTCACATGCAATGGCATGTACAGG |
| 6_known | I15 | Tb927.11.1990_for | GGGGACAAGTTTGTACAAAAAAGCAGGCTATGGCGAAATATATCGCAGGGC | Tb927.11.1990_rev | GGGGACCACTTTGTACAAGAAAGCTGGGTTCAAATCTTCTCGGCGACGTC |
| 6_known | I16 | Tb927.7.1000_for | GGGGACAAGTTTGTACAAAAAAGCAGGCTATGACACGTTGCCGGCTC | Tb927.7.1000_rev | GGGGACCACTTTGTACAAGAAAGCTGGGTTCAAGCTTCTCCACCGTTGC |
| 6_known | I17 | Tb927.7.1980_for | GGGGACAAGTTTGTACAAAAAAGCAGGCTATGATTCAAAACAACGGCAATATGGG | Tb927.7.1980_rev | GGGGACCACTTTGTACAAGAAAGCTGGGTCTACTCATGACCTCGTTTCCGTG |
| 6_known | I18 | Tb927.1.1730_for | GGGGACAAGTTTGTACAAAAAAGCAGGCTATGTTGCGCTACACCAGCTG | Tb927.1.1730_rev | GGGGACCACTTTGTACAAGAAAGCTGGGTTTAAGCTGGAGCTGCTACTTTATCTTTAG |
| 6_known | I19 | Tb927.10.8040_for | GGGGACAAGTTTGTACAAAAAAGCAGGCTATGGAAGCGGTTCTTCGCAAG | Tb927.10.8040_rev | GGGGACCACTTTGTACAAGAAAGCTGGGTTCATGAAAACAAATCATCCAGAGTTGACG |
| 6_known | I20 | Tb927.4.1330_for | GGGGACAAGTTTGTACAAAAAAGCAGGCTATGGGTAAGGCACAGAAGCCG | Tb927.4.1330_rev | GGGGACCACTTTGTACAAGAAAGCTGGGTTTATTTTGCTATGATGTTGACAAGCGG |
| 6_known | I21 | Tb927.10.2700_for | GGGGACAAGTTTGTACAAAAAAGCAGGCTATGCTTGCCAAGAAGAGGGC | Tb927.10.2700_rev | GGGGACCACTTTGTACAAGAAAGCTGGGTTTACTTATCCTCTTCGCCAGCAGT |
| 6_known | I22 | Tb927.10.13680_for | GGGGACAAGTTTGTACAAAAAAGCAGGCTATGCCGGCGTTGCAATGG | Tb927.10.13680_rev | GGGGACCACTTTGTACAAGAAAGCTGGGTCTACACAACGCCCATGCTACC |
| 6_known | I23 | Tb927.5.2570_for | GGGGACAAGTTTGTACAAAAAAGCAGGCTATGGAGGACGATGGCTTGATGT | Tb927.5.2570_rev | GGGGACCACTTTGTACAAGAAAGCTGGGTCTAATTTAGCGGGCGCCGG |
| 6_known | I24 | Tb927.7.3620_for | GGGGACAAGTTTGTACAAAAAAGCAGGCTATGACAACCAACACCGCCG | Tb927.7.3620_rev | GGGGACCACTTTGTACAAGAAAGCTGGGTTTACTTACTCTTAGAAAGCGACTTCTCCG |
| 6_known | J1 | Tb927.11.10560_for | GGGGACAAGTTTGTACAAAAAAGCAGGCTATGCTGTTCAAACCTCGTGGTG | Tb927.11.10560_rev | GGGGACCACTTTGTACAAGAAAGCTGGGTTCACGCATATATAACTGAGAGTTCTTCCA |
| 6_known | J2 | Tb927.11.11860_for | GGGGACAAGTTTGTACAAAAAAGCAGGCTATGAGAGAAAAGGAAGCACATAAGGG | Tb927.11.11860_rev | GGGGACCACTTTGTACAAGAAAGCTGGGTTCAAAGTTCCGGGCTGATACGA |
| 6_known | J3 | Tb927.11.6160_for | GGGGACAAGTTTGTACAAAAAAGCAGGCTATGTCAGGTTGGGCACAACCA | Tb927.11.6160_rev | GGGGACCACTTTGTACAAGAAAGCTGGGTTTAAGCTTTACCTGCGTTCGCC |
| 6_known | J4 | Tb927.10.3100_for | GGGGACAAGTTTGTACAAAAAAGCAGGCTATGGAGGGAAAGAGCGTAGTAGG | Tb927.10.3100_rev | GGGGACCACTTTGTACAAGAAAGCTGGGTTCAATTTTTGATATCCGCAAGCCGG |
| 6_known | J5 | Tb927.10.8020_for | GGGGACAAGTTTGTACAAAAAAGCAGGCTATGCGCCTCGCTTACCCC | Tb927.10.8020_rev | GGGGACCACTTTGTACAAGAAAGCTGGGTTCAGTCTGTCCACTGGGCG |
| 6_known | J6 | Tb927.9.4190_for | GGGGACAAGTTTGTACAAAAAAGCAGGCTATGGGAATGTACGTCACTAAGCTTCT | Tb927.9.4190_rev | GGGGACCACTTTGTACAAGAAAGCTGGGTTTACTCATACTTGAAAAGTTGTTCAATGAC |
| 6_known | J7 | Tb927.6.4780_for | GGGGACAAGTTTGTACAAAAAAGCAGGCTATGGTAAGTGGACCTACAGCTACTG | Tb927.6.4780_rev | GGGGACCACTTTGTACAAGAAAGCTGGGTTTACTCCATATCGCCCACACCC |
| 6_known | J8 | Tb927.11.11490_for | GGGGACAAGTTTGTACAAAAAAGCAGGCTATGAGGAATGCGATGGCGGA | Tb927.11.11490_rev | GGGGACCACTTTGTACAAGAAAGCTGGGTTTACTGGCGGTAGTGATACCAATGG |
| 6_known | J9 | Tb927.10.3770_for | GGGGACAAGTTTGTACAAAAAAGCAGGCTATGATGAACACCGCCACTTTTTCT | Tb927.10.3770_rev | GGGGACCACTTTGTACAAGAAAGCTGGGTTCACCTTCCATCGCCTTTATTATCAC |
| 6_known | J10 | Tb927.9.10960_for | GGGGACAAGTTTGTACAAAAAAGCAGGCTATGAGCGGTGTCAGCAGCA | Tb927.9.10960_rev | GGGGACCACTTTGTACAAGAAAGCTGGGTTCATTTCACTGCCCGGCG |
| 6_known | J11 | Tb927.8.1020_for | GGGGACAAGTTTGTACAAAAAAGCAGGCTATGACGGAGGCTCCTAAGCC | Tb927.8.1020_rev | GGGGACCACTTTGTACAAGAAAGCTGGGTTCATTCCACCGGTGCAGCT |
| 6_known | J12 | Tb927.9.4210_for | GGGGACAAGTTTGTACAAAAAAGCAGGCTATGGGGGGATGTGTGATTTCCG | Tb927.9.4210_rev | GGGGACCACTTTGTACAAGAAAGCTGGGTTCAGCACTCCTCCACGAAAAGT |
| 6_known | J14 | Tb927.2.5280_for | GGGGACAAGTTTGTACAAAAAAGCAGGCTATGCGCGTTGTATACCAGCG | Tb927.2.5280_rev | GGGGACCACTTTGTACAAGAAAGCTGGGTTTACATAAATAGGGAATATAAGTGCAGAAC |
| 6_known | J15 | Tb927.10.3260_for | GGGGACAAGTTTGTACAAAAAAGCAGGCTATGGGTGGCTGCGTTGCA | Tb927.10.3260_rev | GGGGACCACTTTGTACAAGAAAGCTGGGTTCAAACATCATTGCGGAATATCTCTTCG |
| 6_known | J16 | Tb927.4.1630_for | GGGGACAAGTTTGTACAAAAAAGCAGGCTATGTCTGGCTCTCTTCCACAAACA | Tb927.4.1630_rev | GGGGACCACTTTGTACAAGAAAGCTGGGTTCACGATTTGTTTTTCTTTCCGAGC |
| 6_known | J17 | Tb927.11.16110_for | GGGGACAAGTTTGTACAAAAAAGCAGGCTATGCGACTTCGTGATGTTTTCATTG | Tb927.11.16110_rev | GGGGACCACTTTGTACAAGAAAGCTGGGTTCAGTCCGTTGCACGGCC |
| 6_known | J18 | Tb927.3.3160_for | GGGGACAAGTTTGTACAAAAAAGCAGGCTATGGAGTGGATGTATGGCCCAA | Tb927.3.3160_rev | GGGGACCACTTTGTACAAGAAAGCTGGGTTTACTTCAGGTCTTGTAACAGGTGC |
| 6_known | J19 | Tb927.10.15610_for | GGGGACAAGTTTGTACAAAAAAGCAGGCTATGGACCCCTCGAGAGCG | Tb927.10.15610_rev | GGGGACCACTTTGTACAAGAAAGCTGGGTTTACGGTTTGAGGCATAACCGC |
| 6_known | J20 | Tb927.10.10890_for | GGGGACAAGTTTGTACAAAAAAGCAGGCTATGACGGAGACATTCGCATTCC | Tb927.10.10890_rev | GGGGACCACTTTGTACAAGAAAGCTGGGTTCAGTCCACTTCCTCCATACCCG |
| 6_known | K6 | Tb927.11.14960_for | GGGGACAAGTTTGTACAAAAAAGCAGGCTATGCCAAAAATGCGTTTAGACTTCTTAAA | Tb927.11.14960_rev | GGGGACCACTTTGTACAAGAAAGCTGGGTTCATTCGGCCGTTTTGAAAGGC |
| 6_known | K7 | Tb927.10.4910_for | GGGGACAAGTTTGTACAAAAAAGCAGGCTATGCTGGCCAAGGACGTTG | Tb927.10.4910_rev | GGGGACCACTTTGTACAAGAAAGCTGGGTTTACCTTGTAAAGAAAATGTCGTGCAAAA |

| | | | | | |
|---|---|---|---|---|---|
| 6_known | K8 | Tb927.11.16210_for | GGGGACAAGTTTGTACAAAAAAGCAGGCTATGGGTAGTTATGAATACCCAGACTACG | Tb927.11.16210_rev | GGGGACCACTTTGTACAAGAAAGCTGGGTTTACCTTTTCGCCATGAACTCTTTGC |
| 6_known | K9 | Tb927.4.2220_for | GGGGACAAGTTTGTACAAAAAAGCAGGCTATGGAGACGACATGCAAGGAGG | Tb927.4.2220_rev | GGGGACCACTTTGTACAAGAAAGCTGGGTTCACCAAAATCCACTACTGCCTTG |
| 6_known | K10 | Tb927.3.580_for | GGGGACAAGTTTGTACAAAAAAGCAGGCTATGGTGAAGATTTCCTGGGTGACG | Tb927.3.580_rev | GGGGACCACTTTGTACAAGAAAGCTGGGTCTACATATATTCATCATCCATCCTTTTAAC |
| 6_known | K11 | Tb927.7.2680_for | GGGGACAAGTTTGTACAAAAAAGCAGGCTATGTTCCTTCCCGAACAATCGAAC | Tb927.7.2680_rev | GGGGACCACTTTGTACAAGAAAGCTGGGTTCAAAAGGGTACGTGTAGGGGAA |
| 6_known | K12 | Tb927.9.4200_for | GGGGACAAGTTTGTACAAAAAAGCAGGCTATGAACACCGCTTCTGCGC | Tb927.9.4200_rev | GGGGACCACTTTGTACAAGAAAGCTGGGTTCAGCACTCCTCCACGAAAAGT |
| 6_known | K13 | Tb927.10.7490_for | GGGGACAAGTTTGTACAAAAAAGCAGGCTATGTTGGACGCTGCGCTG | Tb927.10.7490_rev | GGGGACCACTTTGTACAAGAAAGCTGGGTTCAAGCTTCGTAGACGTAGATGTCG |
| 6_known | K14 | Tb927.10.8930_for | GGGGACAAGTTTGTACAAAAAAGCAGGCTATGGACTGTGTGTATAACTCAGTGGG | Tb927.10.8930_rev | GGGGACCACTTTGTACAAGAAAGCTGGGTTTACTGGCGAGCTAATGCAAGC |
| 6_known | K15 | Tb927.10.5670_for | GGGGACAAGTTTGTACAAAAAAGCAGGCTATGGCTACAACCTTGCCTTTACAG | Tb927.10.5670_rev | GGGGACCACTTTGTACAAGAAAGCTGGGTTTAATTCAGTTTTTCCAAAGCGGTAGTAAG |
| 6_known | K16 | Tb927.10.1950_for | GGGGACAAGTTTGTACAAAAAAGCAGGCTATGCGCCGAGTTTTTAGGAAAAGT | Tb927.10.1950_rev | GGGGACCACTTTGTACAAGAAAGCTGGGTTCAAACGTGGACAGTTATTCCCTTAC |
| 6_known | K17 | Tb927.10.10410_for | GGGGACAAGTTTGTACAAAAAAGCAGGCTATGTCGATGAATGGCGTTCCAC | Tb927.10.10410_rev | GGGGACCACTTTGTACAAGAAAGCTGGGTCTAAACACCGCGCAGTTGGT |
| 6_known | K18 | Tb927.11.8950_for | GGGGACAAGTTTGTACAAAAAAGCAGGCTATGCATTCGAAAGCATCCTCTGTT | Tb927.11.8950_rev | GGGGACCACTTTGTACAAGAAAGCTGGGTTCAATATATGGATTGGACGGAACCTTTAC |
| 6_known | K19 | Tb927.11.5970_for | GGGGACAAGTTTGTACAAAAAAGCAGGCTATGGGAGGGTGTACGTCACG | Tb927.11.5970_rev | GGGGACCACTTTGTACAAGAAAGCTGGGTCTACTGATAAGATACTTGAACCATTAAAAA |
| 6_known | K20 | Tb927.11.12850_for | GGGGACAAGTTTGTACAAAAAAGCAGGCTATGCAAACTGAACGTGGTCCAA | Tb927.11.12850_rev | GGGGACCACTTTGTACAAGAAAGCTGGGTCTACTTCCGCAGCAGCTGC |
| 6_known | K21 | Tb927.2.5030_for | GGGGACAAGTTTGTACAAAAAAGCAGGCTATGTCCGACGAACTAGGTGTAAAGG | Tb927.2.5030_rev | GGGGACCACTTTGTACAAGAAAGCTGGGTTTACTCTATGTCCATCTCGTACCACTC |
| 6_known | K22 | Tb927.1.3230_for | GGGGACAAGTTTGTACAAAAAAGCAGGCTATGGCATACAGTGGTGCAGGG | Tb927.1.3230_rev | GGGGACCACTTTGTACAAGAAAGCTGGGTCTACAGTGGAAGGACCTTTGTAAAACA |
| 6_known | K23 | Tb927.10.12890_for | GGGGACAAGTTTGTACAAAAAAGCAGGCTATGTCAGCAAGTAATTGTCGAGGC | Tb927.10.12890_rev | GGGGACCACTTTGTACAAGAAAGCTGGGTTCAATAGCTTCTCCCAAATAAAACCCAC |
| 6_known | K24 | Tb927.7.7240_for | GGGGACAAGTTTGTACAAAAAAGCAGGCTATGGGTATTGGACAGAGCCGT | Tb927.7.7240_rev | GGGGACCACTTTGTACAAGAAAGCTGGGTTTACTCCTTGAGGCCCCGG |
| 6_known | L1 | Tb927.7.2470_for | GGGGACAAGTTTGTACAAAAAAGCAGGCTATGGATGTTGGTAGTAAAGTCTTCGTG | Tb927.7.2470_rev | GGGGACCACTTTGTACAAGAAAGCTGGGTTCACGTTGTGTTGCCTTCCAA |
| 6_known | L2 | Tb927.11.15490_for | GGGGACAAGTTTGTACAAAAAAGCAGGCTATGTGGAACTTTTTAATAGACATAAAGAAC | Tb927.11.15490_rev | GGGGACCACTTTGTACAAGAAAGCTGGGTCTACCAGTCATCCCACTCCTCC |
| 6_known | L3 | Tb927.11.16930_for | GGGGACAAGTTTGTACAAAAAAGCAGGCTATGCGGCGCGCTGCGTTC | Tb927.11.16930_rev | GGGGACCACTTTGTACAAGAAAGCTGGGTCTACTCCTCAGCACGATGTGC |
| 6_known | L4 | Tb927.11.3280_for | GGGGACAAGTTTGTACAAAAAAGCAGGCTATGGAGCGACAGCTTCGCG | Tb927.11.3280_rev | GGGGACCACTTTGTACAAGAAAGCTGGGTTTACCGGCGTTGCCGAGA |
| 6_known | L5 | Tb927.1.1010_for | GGGGACAAGTTTGTACAAAAAAGCAGGCTATGGACGGCAGTGAGTGCT | Tb927.1.1010_rev | GGGGACCACTTTGTACAAGAAAGCTGGGTCTATCCTTCCAGAAGAAGCTTTACCG |
| 6_known | L6 | Tb927.4.3300_for | GGGGACAAGTTTGTACAAAAAAGCAGGCTATGTTTAGGATACCCCGATCAGCC | Tb927.4.3300_rev | GGGGACCACTTTGTACAAGAAAGCTGGGTTTAGCTCTCCAGATAAGTGCGGAA |
| 6_known | L7 | Tb927.5.1150_for | GGGGACAAGTTTGTACAAAAAAGCAGGCTATGGGGTGCGGAGAGAAAAATC | Tb927.5.1150_rev | GGGGACCACTTTGTACAAGAAAGCTGGGTCTAAGACGAATCATCATCATCATCATCATC |
| 6_known | L8 | Tb927.6.4970_for | GGGGACAAGTTTGTACAAAAAAGCAGGCTATGCCCCGACCCGACAGA | Tb927.6.4970_rev | GGGGACCACTTTGTACAAGAAAGCTGGGTTTAGTCATTAGTGCAGCTGCTGTC |
| 6_known | L9 | Tb927.10.3170_for | GGGGACAAGTTTGTACAAAAAAGCAGGCTATGGATCTTGCACAGGCAAAGAC | Tb927.10.3170_rev | GGGGACCACTTTGTACAAGAAAGCTGGGTTCAGCGCATCTGCTTCATAACAA |
| 6_known | L10 | Tb927.7.5080_for | GGGGACAAGTTTGTACAAAAAAGCAGGCTATGCTTCGGCACGCGTTG | Tb927.7.5080_rev | GGGGACCACTTTGTACAAGAAAGCTGGGTTCACATGTTCGCGGGGAACT |
| 6_known | L11 | Tb927.10.7690_for | GGGGACAAGTTTGTACAAAAAAGCAGGCTATGACGGACCGGACCATTTATGA | Tb927.10.7690_rev | GGGGACCACTTTGTACAAGAAAGCTGGGTTTAAAACATAATGAGCCGTTTTATTTCTTT |
| 6_known | L12 | Tb927.7.7170_for | GGGGACAAGTTTGTACAAAAAAGCAGGCTATGAATCCACACGCCTGTGGT | Tb927.7.7170_rev | GGGGACCACTTTGTACAAGAAAGCTGGGTTTATCGGTAGTCCCTGTAGTGATCAG |
| 6_known | L13 | Tb927.11.3890_for | GGGGACAAGTTTGTACAAAAAAGCAGGCTATGCGCTCCGGTTTGTTCG | Tb927.11.3890_rev | GGGGACCACTTTGTACAAGAAAGCTGGGTTCATGGAGCATCGACAGCGG |
| 6_known | L14 | Tb927.1.1560_for | GGGGACAAGTTTGTACAAAAAAGCAGGCTATGGGGAAAAGTTTTGTCGTACGG | Tb927.1.1560_rev | GGGGACCACTTTGTACAAGAAAGCTGGGTCTACTCTGACCTTCCCGCAATGG |
| 6_known | L15 | Tb927.11.15460_for | GGGGACAAGTTTGTACAAAAAAGCAGGCTATGGGCATCAAGGGTCGTAGG | Tb927.11.15460_rev | GGGGACCACTTTGTACAAGAAAGCTGGGTTTAAGACTTCGGAAAAGCCATTCCAA |
| 6_known | L16 | Tb927.9.6560_for | GGGGACAAGTTTGTACAAAAAAGCAGGCTATGGGCAGTGATAGTAAATCTAGTAAGTCC | Tb927.9.6560_rev | GGGGACCACTTTGTACAAGAAAGCTGGGTTTACTTGAAGAGACTGGCGAAGGG |
| 6_known | L17 | Tb927.7.5540_for | GGGGACAAGTTTGTACAAAAAAGCAGGCTATGCCGCTAGGGATTGTTGC | Tb927.7.5540_rev | GGGGACCACTTTGTACAAGAAAGCTGGGTTCAAATCTTGTAACCGATTTCATACCCC |
| 6_known | L18 | Tb927.5.2060_for | GGGGACAAGTTTGTACAAAAAAGCAGGCTATGACGCGCCTACACTTCAAG | Tb927.5.2060_rev | GGGGACCACTTTGTACAAGAAAGCTGGGTTTAGCTGACCTGTTTGTTGCGG |
| 6_known | L19 | Tb927.11.7530_for | GGGGACAAGTTTGTACAAAAAAGCAGGCTATGTTTACTCTTGTCGGGCGC | Tb927.11.7530_rev | GGGGACCACTTTGTACAAGAAAGCTGGGTTTAAACAAGGTTAAACATGGATCTGAGCG |
| 6_known | L20 | Tb927.2.5020_for | GGGGACAAGTTTGTACAAAAAAGCAGGCTATGATGTGCAAGGGTCTATTTTCAAAATTG | Tb927.2.5020_rev | GGGGACCACTTTGTACAAGAAAGCTGGGTTCATTTCTCTTGTGCAAAGGACTGT |
| 6_known | L21 | Tb927.11.11160_for | GGGGACAAGTTTGTACAAAAAAGCAGGCTATGAAGTTCGGAAAGCGGCTT | Tb927.11.11160_rev | GGGGACCACTTTGTACAAGAAAGCTGGGTCTACGGTTTGCCTTCAAACACAAC |
| 6_known | L22 | Tb927.3.4570_for | GGGGACAAGTTTGTACAAAAAAGCAGGCTATGATGGTAACCAACACATGCATAATAATG | Tb927.3.4570_rev | GGGGACCACTTTGTACAAGAAAGCTGGGTTTACGCAACGGCCGTCTTTG |
| 6_known | L23 | Tb927.10.5310_for | GGGGACAAGTTTGTACAAAAAAGCAGGCTATGAGTCAGCGGTTCGGACC | Tb927.10.5310_rev | GGGGACCACTTTGTACAAGAAAGCTGGGTTCAAGTGCATACGGTGAACTTACC |
| 6_known | L24 | Tb927.10.11450_for | GGGGACAAGTTTGTACAAAAAAGCAGGCTATGAATATTTTTTCCTTTGGGAGGGAAAGC | Tb927.10.11450_rev | GGGGACCACTTTGTACAAGAAAGCTGGGTTCAGGAACACGACACTTCCCG |
| 6_known | M1 | Tb927.3.4840_for | GGGGACAAGTTTGTACAAAAAAGCAGGCTATGCCGCCTCCCTTTTCTATAACT | Tb927.3.4840_rev | GGGGACCACTTTGTACAAGAAAGCTGGGTCTAATTAGAAACCCTTTTTCTCTTGTAAAA |
| 6_known | M2 | Tb927.11.16140_for | GGGGACAAGTTTGTACAAAAAAGCAGGCTATGGCGGACGGGCCGACG | Tb927.11.16140_rev | GGGGACCACTTTGTACAAGAAAGCTGGGTTCACTCCAGTGAAAAGTGAACAGTG |
| 6_known | M3 | Tb927.10.2520_for | GGGGACAAGTTTGTACAAAAAAGCAGGCTATGGGTCCGAAAACTCAGGAGG | Tb927.10.2520_rev | GGGGACCACTTTGTACAAGAAAGCTGGGTCTCACCACTGTTGGACGATATCTTCA |
| 6_known | M4 | Tb927.1.2120_for | GGGGACAAGTTTGTACAAAAAAGCAGGCTATGGGGTGTGGTGCGTCT | Tb927.1.2120_rev | GGGGACCACTTTGTACAAGAAAGCTGGGTTTACTTTTTTTTATTTCCACATACCAGTC |
| 6_known | M5 | Tb927.6.640_for | GGGGACAAGTTTGTACAAAAAAGCAGGCTATGACTTCAAGCACTTTACTGGAAAAGT | Tb927.6.640_rev | GGGGACCACTTTGTACAAGAAAGCTGGGTTTAGGCATTTTGGGGACTCACGG |
| 6_known | M6 | Tb927.6.4770_for | GGGGACAAGTTTGTACAAAAAAGCAGGCTATGTACCCCCGACACGATGAT | Tb927.6.4770_rev | GGGGACCACTTTGTACAAGAAAGCTGGGTTCAATAATATGTTTCTCGCACTCCCG |
| 6_known | M7 | Tb927.1.5030_for | GGGGACAAGTTTGTACAAAAAAGCAGGCTATGGGTGGTGTGGTCGGC | Tb927.1.5030_rev | GGGGACCACTTTGTACAAGAAAGCTGGGTTTACATTTGTTTAACTACAATACCGCGTTT |
| 6_known | M8 | Tb927.6.2900_for | GGGGACAAGTTTGTACAAAAAAGCAGGCTATGGAAGGGGAACTTGCCTTCG | Tb927.6.2900_rev | GGGGACCACTTTGTACAAGAAAGCTGGGTTTAGCGGCTGCGTTTATTTCCC |
| 6_known | M9 | Tb927.9.12510_for | GGGGACAAGTTTGTACAAAAAAGCAGGCTATGAGTTTATCCCGCACAGAAAACA | Tb927.9.12510_rev | GGGGACCACTTTGTACAAGAAAGCTGGGTTTAGAAGCCTCCGTCATCCACA |
| 6_known | M10 | Tb927.11.2230_for | GGGGACAAGTTTGTACAAAAAAGCAGGCTATGGCAACATCAGAAAGTTCAGGTG | Tb927.11.2230_rev | GGGGACCACTTTGTACAAGAAAGCTGGGTTTACAAATTTGGTGAGGCATCAGCA |
| 6_known | M11 | Tb927.1.1620_for | GGGGACAAGTTTGTACAAAAAAGCAGGCTATGCTAAGTCCCTGGAATGATTGTG | Tb927.1.1620_rev | GGGGACCACTTTGTACAAGAAAGCTGGGTTCAGCCACGTTCTCTCTTTGTTTG |
| 6_known | M12 | Tb927.8.8050_for | GGGGACAAGTTTGTACAAAAAAGCAGGCTATGCATCGAAAAAGCATCGTGTCG | Tb927.8.8050_rev | GGGGACCACTTTGTACAAGAAAGCTGGGTTCAGAACCGCTTCGGTGTCC |
| 6_known | M13 | Tb927.10.8290_for | GGGGACAAGTTTGTACAAAAAAGCAGGCTATGGCGGATTTCTTTAACGTAAGTGA | Tb927.10.8290_rev | GGGGACCACTTTGTACAAGAAAGCTGGGTTTACTGCCCTCCCCTAGCAC |
| 6_known | M14 | Tb927.5.760_for | GGGGACAAGTTTGTACAAAAAAGCAGGCTATGAGTGGAAGCGCTGATATTCTCG | Tb927.5.760_rev | GGGGACCACTTTGTACAAGAAAGCTGGGTCTAACTGCTGCTGCGCTTTGC |
| 6_known | M15 | Tb927.10.6680_for | GGGGACAAGTTTGTACAAAAAAGCAGGCTATGCAGCCACTACAGGCAAC | Tb927.10.6680_rev | GGGGACCACTTTGTACAAGAAAGCTGGGTTTAAGGAATGCGCTCCGCC |
| 6_known | M16 | Tb927.10.3900_for | GGGGACAAGTTTGTACAAAAAAGCAGGCTATGACCTCAACGACCGAATTCAC | Tb927.10.3900_rev | GGGGACCACTTTGTACAAGAAAGCTGGGTTTATATAAGTACGCTTTCGCCCCTGG |
| 6_known | M17 | Tb927.10.8270_for | GGGGACAAGTTTGTACAAAAAAGCAGGCTATGGCGGATTTCTTTAACGTAAGTGA | Tb927.10.8270_rev | GGGGACCACTTTGTACAAGAAAGCTGGGTTTACTGCCCTCCTCTAGCACG |
| 6_known | M18 | Tb927.6.1160_for | GGGGACAAGTTTGTACAAAAAAGCAGGCTATGTCTCGTATTGCGTCCGTTG | Tb927.6.1160_rev | GGGGACCACTTTGTACAAGAAAGCTGGGTTTAGTCTATGGTTTTGTATGCCGTAATGG |
| 6_known | M19 | Tb927.8.1060_for | GGGGACAAGTTTGTACAAAAAAGCAGGCTATGTTGACCTTAGTCCGGCGC | Tb927.8.1060_rev | GGGGACCACTTTGTACAAGAAAGCTGGGTTTAGGATATGTTGAGGAGTCGAAGAACA |
| 6_known | M20 | Tb927.1.1340_for | GGGGACAAGTTTGTACAAAAAAGCAGGCTATGGCATGCAGTACCAAAAGGG | Tb927.1.1340_rev | GGGGACCACTTTGTACAAGAAAGCTGGGTCTACTCGCCCAGCTGCTGA |
| 6_known | M21 | Tb927.4.2390_for | GGGGACAAGTTTGTACAAAAAAGCAGGCTATGGAAGGTACAAATAGCGGTAATGAAG | Tb927.4.2390_rev | GGGGACCACTTTGTACAAGAAAGCTGGGTTTATCCCTTTTGTGTTGTGCCCC |
| 6_known | M22 | Tb927.8.5970_for | GGGGACAAGTTTGTACAAAAAAGCAGGCTATGCTTGAGGAAGACCTTGAGGC | Tb927.8.5970_rev | GGGGACCACTTTGTACAAGAAAGCTGGGTTCAGGCGGCATACCTGTACA |
| 6_known | M23 | Tb927.9.7210_for | GGGGACAAGTTTGTACAAAAAAGCAGGCTATGACCCCTCGGCGCGTC | Tb927.9.7210_rev | GGGGACCACTTTGTACAAGAAAGCTGGGTTCACGAGTCCAGAGATGGAAGAAG |
| 6_known | M24 | Tb927.1.4180_for | GGGGACAAGTTTGTACAAAAAAGCAGGCTATGGAGACATCGATCGGCTTAACG | Tb927.1.4180_rev | GGGGACCACTTTGTACAAGAAAGCTGGGTCTATTTCTTCTTGCTTCCCTTGCCT |
| 6_known | N1 | Tb927.10.7740_for | GGGGACAAGTTTGTACAAAAAAGCAGGCTATGGACCAATACGGAATACCACAGC | Tb927.10.7740_rev | GGGGACCACTTTGTACAAGAAAGCTGGGTCTACTGGGAAACTGCGAGCC |
| 6_known | N2 | Tb927.9.2220_for | GGGGACAAGTTTGTACAAAAAAGCAGGCTATGGCAGATATCCTTTTAAATGCCGC | Tb927.9.2220_rev | GGGGACCACTTTGTACAAGAAAGCTGGGTTCACGCCTTCGTGTATAAGGCC |
| 6_known | N3 | Tb927.3.5030_for | GGGGACAAGTTTGTACAAAAAAGCAGGCTATGGCGCACTATGATGAGTGGC | Tb927.3.5030_rev | GGGGACCACTTTGTACAAGAAAGCTGGGTCTAAATGTCGCCTTGCAATTGAGATATC |
| 6_known | N4 | Tb927.8.3830_for | GGGGACAAGTTTGTACAAAAAAGCAGGCTATGCGACACTGGACATTTCACTG | Tb927.8.3830_rev | GGGGACCACTTTGTACAAGAAAGCTGGGTTTATTTACGGGTCGCTGGTCGC |
| 6_known | N5 | Tb927.6.2020_for | GGGGACAAGTTTGTACAAAAAAGCAGGCTATGAGTCTTCAGACAAACTGCTACTCC | Tb927.6.2020_rev | GGGGACCACTTTGTACAAGAAAGCTGGGTCTCACTTTTCGCCAGTACGAATGAG |
| 6_known | N6 | Tb927.8.1090_for | GGGGACAAGTTTGTACAAAAAAGCAGGCTATGAAAGATCGGTCGGCCGC | Tb927.8.1090_rev | GGGGACCACTTTGTACAAGAAAGCTGGGTTTAACGTCGGGTCAATGACCGA |
| 6_known | N7 | Tb927.9.11760_for | GGGGACAAGTTTGTACAAAAAAGCAGGCTATGGGTGTCCATGGACTGTGG | Tb927.9.11760_rev | GGGGACCACTTTGTACAAGAAAGCTGGGTTCACGGATCACCCCTGGC |
| 6_known | N8 | Tb927.10.13030_for | GGGGACAAGTTTGTACAAAAAAGCAGGCTATGGAAATAATTAGTGTGGCTGAGGAG | Tb927.10.13030_rev | GGGGACCACTTTGTACAAGAAAGCTGGGTTTACTCCCATTCTAGAGGAACAACACA |
| 6_known | N9 | Tb927.10.5010_for | GGGGACAAGTTTGTACAAAAAAGCAGGCTATGAGGAGGTTTTGTGATGTCGTC | Tb927.10.5010_rev | GGGGACCACTTTGTACAAGAAAGCTGGGTTTACTGTCCACTCAACCTGTACAGC |
| 6_known | N10 | Tb927.7.4020_for | GGGGACAAGTTTGTACAAAAAAGCAGGCTATGTATACCAGTGTTAGAAAGCCTCCG | Tb927.7.4020_rev | GGGGACCACTTTGTACAAGAAAGCTGGGTTTACCACAGAGAGCGACTCCG |
| 6_known | N11 | Tb927.3.2020_for | GGGGACAAGTTTGTACAAAAAAGCAGGCTATGAGCAACATTAAGGTGGCAGTT | Tb927.3.2020_rev | GGGGACCACTTTGTACAAGAAAGCTGGGTCTATTTTTCACTGGGCGCTGAGG |
| 6_known | N12 | Tb927.7.6940_for | GGGGACAAGTTTGTACAAAAAAGCAGGCTATGAAAAGCAACTGGAGAGACAAGAA | Tb927.7.6940_rev | GGGGACCACTTTGTACAAGAAAGCTGGGTTCAACTTGCATCCATCCATCCCT |
| 6_known | N13 | Tb927.11.2530_for | GGGGACAAGTTTGTACAAAAAAGCAGGCTATGCGGCGGTGTTCCTTTG | Tb927.11.2530_rev | GGGGACCACTTTGTACAAGAAAGCTGGGTTCACAAACTTACCTTCCCTACAGAAGG |
| 6_known | N14 | Tb927.8.2870_for | GGGGACAAGTTTGTACAAAAAAGCAGGCTATGGCGGATCACAGTGTCAAAG | Tb927.8.2870_rev | GGGGACCACTTTGTACAAGAAAGCTGGGTTCAAAGAAGGGCGAGAGTCTCG |
| 6_known | N15 | Tb927.8.7340_for | GGGGACAAGTTTGTACAAAAAAGCAGGCTATGGCATCCTACATGTTGTTGCTAG | Tb927.8.7340_rev | GGGGACCACTTTGTACAAGAAAGCTGGGTCTATGCTGACAGTAACTGGAGCAG |
| 6_known | N17 | Tb927.3.4940_for | GGGGACAAGTTTGTACAAAAAAGCAGGCTATGACCTCACTCGTCCTAAATCACT | Tb927.3.4940_rev | GGGGACCACTTTGTACAAGAAAGCTGGGTTTATTTCTTTCGGTAGACGGACAGC |
| 6_known | N18 | Tb927.8.4500_for | GGGGACAAGTTTGTACAAAAAAGCAGGCTATGGAGCACCGCGCCGCA | Tb927.8.4500_rev | GGGGACCACTTTGTACAAGAAAGCTGGGTTTATTTAGATTTCTCAAACGCAGCAAAGG |
| 6_known | N19 | Tb927.8.2630_for | GGGGACAAGTTTGTACAAAAAAGCAGGCTATGCGGACAAATGAAGAAAGAGATGA | Tb927.8.2630_rev | GGGGACCACTTTGTACAAGAAAGCTGGGTCTACTTTCTCTTCACCATCGCCTTC |
| 6_known | N20 | Tb927.11.14270_for | GGGGACAAGTTTGTACAAAAAAGCAGGCTATGGTCTCTCTCCGGATCAGGA | Tb927.11.14270_rev | GGGGACCACTTTGTACAAGAAAGCTGGGTTCAACACAAGTGGGGAAATTTTTCATC |
| 6_known | N21 | Tb927.10.15260_for | GGGGACAAGTTTGTACAAAAAAGCAGGCTATGAAGAAATCCACAGTACTTGATGCG | Tb927.10.15260_rev | GGGGACCACTTTGTACAAGAAAGCTGGGTCTAAAATACAACAGGACTAACATCCTCTTC |
| 6_known | N22 | Tb927.11.2040_for | GGGGACAAGTTTGTACAAAAAAGCAGGCTATGAAGCTGAGGGCTGACTCC | Tb927.11.2040_rev | GGGGACCACTTTGTACAAGAAAGCTGGGTTTATTTGATCAACCACTTGATCACCGA |
| 6_known | N23 | Tb927.11.11800_for | GGGGACAAGTTTGTACAAAAAAGCAGGCTATGCGGCGCTTAGTGCGT | Tb927.11.11800_rev | GGGGACCACTTTGTACAAGAAAGCTGGGTTCAGTCCGTCAAACCAACAAGTAAC |
| 6_known | N24 | Tb927.3.3640_for | GGGGACAAGTTTGTACAAAAAAGCAGGCTATGATTAGCTCCCATGATATTGAATTAGTT | Tb927.3.3640_rev | GGGGACCACTTTGTACAAGAAAGCTGGGTTTAACAACGAAAAATACGCCGAAGC |
| 6_known | O1 | Tb927.9.14240_for | GGGGACAAGTTTGTACAAAAAAGCAGGCTATGAGGTGCACGGCATCCG | Tb927.9.14240_rev | GGGGACCACTTTGTACAAGAAAGCTGGGTCTATAGAAGCCTCATCAGCTTCTCACG |
| 6_known | O2 | Tb927.2.4390_for | GGGGACAAGTTTGTACAAAAAAGCAGGCTATGGCCGAGAGGGCATCC | Tb927.2.4390_rev | GGGGACCACTTTGTACAAGAAAGCTGGGTCTAATAGTTATCTGGCATGCCAGAACTAC |
| 6_known | O3 | Tb927.5.1560_for | GGGGACAAGTTTGTACAAAAAAGCAGGCTATGCCACCGACCCCGTTG | Tb927.5.1560_rev | GGGGACCACTTTGTACAAGAAAGCTGGGTTTAATGCCTCGACATGCGCC |
| 6_known | O4 | Tb927.11.2650_for | GGGGACAAGTTTGTACAAAAAAGCAGGCTATGATGCGTCGTGTTTGTCAGA | Tb927.11.2650_rev | GGGGACCACTTTGTACAAGAAAGCTGGGTCTACTTATCGGCAGTGGGCTCC |
| 6_known | O5 | Tb927.11.12930_for | GGGGACAAGTTTGTACAAAAAAGCAGGCTATGGGACGACGCAACATAGTTTT | Tb927.11.12930_rev | GGGGACCACTTTGTACAAGAAAGCTGGGTTTATTTAGGTGGCCCTCGAGGTG |
| 6_known | O6 | Tb927.11.15850_for | GGGGACAAGTTTGTACAAAAAAGCAGGCTATGCGGCACACAATACCTTTTTTC | Tb927.11.15850_rev | GGGGACCACTTTGTACAAGAAAGCTGGGTTCAGTTATTGTCATCAAGTATATCCTTCGC |
| 6_known | O7 | Tb927.10.11180_for | GGGGACAAGTTTGTACAAAAAAGCAGGCTATGGAGCGCTTGAAGTATTTGATGC | Tb927.10.11180_rev | GGGGACCACTTTGTACAAGAAAGCTGGGTCTACTGATTGTCCACATCATCATCCC |
| 6_known | O8 | Tb927.9.10640_for | GGGGACAAGTTTGTACAAAAAAGCAGGCTATGTCTGTAGGGGATTTTTTTGGAAATATC | Tb927.9.10640_rev | GGGGACCACTTTGTACAAGAAAGCTGGGTTCATTTCAGCAAAGGGTAGTTTGTGT |
| 6_known | O9 | Tb927.4.4230_for | GGGGACAAGTTTGTACAAAAAAGCAGGCTATGGGAAGGCGGATGCGG | Tb927.4.4230_rev | GGGGACCACTTTGTACAAGAAAGCTGGGTTTAGCGATCCACGCCAAACG |
| 6_known | O10 | Tb927.6.2550_for | GGGGACAAGTTTGTACAAAAAAGCAGGCTATGAGCGCACCACCACCG | Tb927.6.2550_rev | GGGGACCACTTTGTACAAGAAAGCTGGGTTTACCGACGCCTCTTCTTTGTG |
| 6_known | O11 | Tb927.10.11730_for | GGGGACAAGTTTGTACAAAAAAGCAGGCTATGGATAGCTCGCTTTGCTTCG | Tb927.10.11730_rev | GGGGACCACTTTGTACAAGAAAGCTGGGTTCACGGTTCCCCACTCGTG |
| 6_known | O12 | Tb927.7.4400_for | GGGGACAAGTTTGTACAAAAAAGCAGGCTATGGGGGAAGAGGAGAATTTACGT | Tb927.7.4400_rev | GGGGACCACTTTGTACAAGAAAGCTGGGTTTATGTGAGCATGTCAAGTACAGTTTTCA |
| 6_known | O13 | Tb927.2.4720_for | GGGGACAAGTTTGTACAAAAAAGCAGGCTATGACTGTTTTATCGTTTTTTAGTTTTTTA | Tb927.2.4720_rev | GGGGACCACTTTGTACAAGAAAGCTGGGTTCAACATCGTGGACCAAGGGC |
| 6_known | O14 | Tb927.3.2950_for | GGGGACAAGTTTGTACAAAAAAGCAGGCTATGGTTGTAGGTGCTGTGAAAGAG | Tb927.3.2950_rev | GGGGACCACTTTGTACAAGAAAGCTGGGTTTATCGTCTTGAAGATCTCTGAGGAGAT |
| 6_known | O15 | Tb927.9.2470_for | GGGGACAAGTTTGTACAAAAAAGCAGGCTATGACGGGAATGAACAGATCGTG | Tb927.9.2470_rev | GGGGACCACTTTGTACAAGAAAGCTGGGTTTAAGATGCCTGTTGCACAGATGC |
| 6_known | O16 | Tb927.1.760_for | GGGGACAAGTTTGTACAAAAAAGCAGGCTATGTTCGCCCCACCGGAG | Tb927.1.760_rev | GGGGACCACTTTGTACAAGAAAGCTGGGTCTACTGTCCAATACGATCTTCAGCG |
| 6_known | O17 | Tb927.2.4330_for | GGGGACAAGTTTGTACAAAAAAGCAGGCTATGACGCAATCAGCTCAATCAACT | Tb927.2.4330_rev | GGGGACCACTTTGTACAAGAAAGCTGGGTTCAATCATCTTCATCACAGGGTGC |
| 6_known | O18 | Tb927.2.2470_for | GGGGACAAGTTTGTACAAAAAAGCAGGCTATGAGACGCCTCACCCGC | Tb927.2.2470_rev | GGGGACCACTTTGTACAAGAAAGCTGGGTTTACCCTACAACTTTCACATAGCCGA |
| 6_known | O19 | Tb927.11.8010_for | GGGGACAAGTTTGTACAAAAAAGCAGGCTATGTTGTTTTGTCGGCATTTATTGTTCA | Tb927.11.8010_rev | GGGGACCACTTTGTACAAGAAAGCTGGGTTTATCTTGAACTTTTACTGCCACTGGC |
| 6_known | O20 | Tb927.10.460_for | GGGGACAAGTTTGTACAAAAAAGCAGGCTATGGAGAATTACACCCAACTTCGTG | Tb927.10.460_rev | GGGGACCACTTTGTACAAGAAAGCTGGGTCTACCGCAGCATTTGACGATTAAATG |
| 6_known | O21 | Tb927.11.5270_for | GGGGACAAGTTTGTACAAAAAAGCAGGCTATGGATATCGTTAATAACAATGGTTCTTTT | Tb927.11.5270_rev | GGGGACCACTTTGTACAAGAAAGCTGGGTTTACGAGAATTTTCCGTGGGCG |
| 6_known | O22 | Tb927.11.5220_for | GGGGACAAGTTTGTACAAAAAAGCAGGCTATGCAGTTGCATTTATCTTCAATCAGC | Tb927.11.5220_rev | GGGGACCACTTTGTACAAGAAAGCTGGGTTCACAGCTCCGGAAGCGG |
| 6_known | O23 | Tb927.10.2640_for | GGGGACAAGTTTGTACAAAAAAGCAGGCTATGAACTTTGGCCACACCATGG | Tb927.10.2640_rev | GGGGACCACTTTGTACAAGAAAGCTGGGTCTAGTCACCCGGAAGTACCAACA |
| 6_known | O24 | Tb927.7.6310_for | GGGGACAAGTTTGTACAAAAAAGCAGGCTATGCACGCAACCGCTGAG | Tb927.7.6310_rev | GGGGACCACTTTGTACAAGAAAGCTGGGTCTAAATATCACGGTTTTGTATGAGCTCGC |
| 6_known | P1 | Tb927.8.8350_for | GGGGACAAGTTTGTACAAAAAAGCAGGCTATGCAGAAAAGTATACAAGATGAAGTGCG | Tb927.8.8350_rev | GGGGACCACTTTGTACAAGAAAGCTGGGTCTAGTTTTGTTGCTGTTGTATTTGACGT |
| 6_known | P2 | Tb927.10.2960_for | GGGGACAAGTTTGTACAAAAAAGCAGGCTATGAGGTTATGGAATAGATTCTCTCGTCT | Tb927.10.2960_rev | GGGGACCACTTTGTACAAGAAAGCTGGGTTCACAATGCAGTGGCACCC |
| 6_known | P3 | Tb927.9.11930_for | GGGGACAAGTTTGTACAAAAAAGCAGGCTATGGGGCAGCAGAAAAAAGATGC | Tb927.9.11930_rev | GGGGACCACTTTGTACAAGAAAGCTGGGTTCAGTGAGAGGAAAGATGTCCATGT |
| 6_known | P4 | Tb927.11.12310_for | GGGGACAAGTTTGTACAAAAAAGCAGGCTATGGGTATGGCAACACGTTCG | Tb927.11.12310_rev | GGGGACCACTTTGTACAAGAAAGCTGGGTTTACGCAGCGCTTGGATAAGG |
| 6_known | P5 | Tb927.4.2720_for | GGGGACAAGTTTGTACAAAAAAGCAGGCTATGGAAAGACTGTACCGTAGCCA | Tb927.4.2720_rev | GGGGACCACTTTGTACAAGAAAGCTGGGTTTATTTATTACTCTTCATACCATAAAGTGC |
| 6_known | P6 | Tb927.10.9230_for | GGGGACAAGTTTGTACAAAAAAGCAGGCTATGGAAACCTCATGTGCCTCTGC | Tb927.10.9230_rev | GGGGACCACTTTGTACAAGAAAGCTGGGTTCAAAACTGATCCATGATTTTGTGAAAATG |
| 6_known | P7 | Tb927.7.1240_for | GGGGACAAGTTTGTACAAAAAAGCAGGCTATGAATGCCAACGGGGTACCC | Tb927.7.1240_rev | GGGGACCACTTTGTACAAGAAAGCTGGGTCTACTTCCCAGGCAAACTCAGGC |
| 6_known | P8 | Tb927.11.5570_for | GGGGACAAGTTTGTACAAAAAAGCAGGCTATGGAATCGGATGCGGGTGT | Tb927.11.5570_rev | GGGGACCACTTTGTACAAGAAAGCTGGGTTTAACGTAACCGATGAATTTGGGTGC |
| 6_known | P9 | Tb927.4.1340_for | GGGGACAAGTTTGTACAAAAAAGCAGGCTATGTTTTCACACGCGGCGT | Tb927.4.1340_rev | GGGGACCACTTTGTACAAGAAAGCTGGGTCTATGCGCCCTCGTCAATGG |
| 6_known | P10 | Tb927.7.6850_for | GGGGACAAGTTTGTACAAAAAAGCAGGCTATGGAGGAACTCCACCAACAAATG | Tb927.7.6850_rev | GGGGACCACTTTGTACAAGAAAGCTGGGTTCAAATCGCCAACACATACATTAAGATCA |
| 6_known | P11 | Tb927.3.3580_for | GGGGACAAGTTTGTACAAAAAAGCAGGCTATGATCCAATCGGGTATGTTCTTCG | Tb927.3.3580_rev | GGGGACCACTTTGTACAAGAAAGCTGGGTTTACAAGTCTCCAGCATTGGCTCC |
| 6_known | P12 | Tb927.9.9600_for | GGGGACAAGTTTGTACAAAAAAGCAGGCTATGAATGGTGTTGCACCAATAACGG | Tb927.9.9600_rev | GGGGACCACTTTGTACAAGAAAGCTGGGTTCATGCATTAAACGACAAACTACGC |
| 6_known | P13 | Tb927.4.1840_for | GGGGACAAGTTTGTACAAAAAAGCAGGCTATGATGGTAGAAGCAAAGGTGCAG | Tb927.4.1840_rev | GGGGACCACTTTGTACAAGAAAGCTGGGTTCAAGAGAATGAAGTACGATTGGCC |
| 6_known | P14 | Tb927.10.1500_for | GGGGACAAGTTTGTACAAAAAAGCAGGCTATGGCTCTAAAGCTGCTTTCAGAAA | Tb927.10.1500_rev | GGGGACCACTTTGTACAAGAAAGCTGGGTTTATGTACTCTTTGTATTCTCTGTTGAGCG |
| 6_known | P15 | Tb927.7.680_for | GGGGACAAGTTTGTACAAAAAAGCAGGCTATGCGGCGGCATGTTTCT | Tb927.7.680_rev | GGGGACCACTTTGTACAAGAAAGCTGGGTTTAAAAAGAACCCGGTTTACTTGTAGAGG |
| 6_known | P16 | Tb927.2.4510_for | GGGGACAAGTTTGTACAAAAAAGCAGGCTATGGCTGAGTTGTTAAGAAAGAAAATTGAG | Tb927.2.4510_rev | GGGGACCACTTTGTACAAGAAAGCTGGGTTTAGCGGCGCCGCACTTC |
| 6_known | P17 | Tb927.5.440_for | GGGGACAAGTTTGTACAAAAAAGCAGGCTATGAGAGTATGGATGAAAGATGACTTCCC | Tb927.5.440_rev | GGGGACCACTTTGTACAAGAAAGCTGGGTTCATTTCCTTACATCCAGTGTGTCATC |
| 6_known | P18 | Tb927.11.11920_for | GGGGACAAGTTTGTACAAAAAAGCAGGCTATGTTGCGGCTCGTGCTG | Tb927.11.11920_rev | GGGGACCACTTTGTACAAGAAAGCTGGGTCTACCGTTTCTCAGGGCGTTG |

| | | | | | |
|---|---|---|---|---|---|
| 6_known | P19 | Tb927.7.3880_for | GGGGACAAGTTTGTACAAAAAAGCAGGCTATGGAACCCGCTGATGGGT | Tb927.7.3880_rev | GGGGACCACTTTGTACAAGAAAGCTGGGTTCAATCAACCTGCGCCAAAAAGG |
| 6_known | P20 | Tb927.8.1410_for | GGGGACAAGTTTGTACAAAAAAGCAGGCTATGGCACGTAATAAGGCTCACAAG | Tb927.8.1410_rev | GGGGACCACTTTGTACAAGAAAGCTGGGTCTACTCTTCGGCAACAACTTTCGG |
| 6_known | P21 | Tb927.11.16270_for | GGGGACAAGTTTGTACAAAAAAGCAGGCTATGCAGTGCCTGTTCTTGGAG | Tb927.11.16270_rev | GGGGACCACTTTGTACAAGAAAGCTGGGTCTAACTCTCATGATAGACAACGTCATCG |
| 6_known | P22 | Tb927.11.12830_for | GGGGACAAGTTTGTACAAAAAAGCAGGCTATGTGGAGCAAAGCGGAGC | Tb927.11.12830_rev | GGGGACCACTTTGTACAAGAAAGCTGGGTTCACCCAATTGAACTCTTACGATGC |
| 6_known | P23 | Tb927.8.1550_for | GGGGACAAGTTTGTACAAAAAAGCAGGCTATGTCGGCCACTTATCCAGAGC | Tb927.8.1550_rev | GGGGACCACTTTGTACAAGAAAGCTGGGTTTATGCGCACTCCTCTACAGCA |
| 6_known | P24 | Tb927.7.990_for | GGGGACAAGTTTGTACAAAAAAGCAGGCTATGTCTGACAGTGGTGACGAATACG | Tb927.7.990_rev | GGGGACCACTTTGTACAAGAAAGCTGGGTCTACGACTCAACCAAATGTTCCCTT |
| 7_hypothetical | A1 | Tb927.5.3930_for | GGGGACAAGTTTGTACAAAAAAGCAGGCTATGACGGGAAAAAATCACTCGGTC | Tb927.5.3930_rev | GGGGACCACTTTGTACAAGAAAGCTGGGTTTACTTCGTGTTAATAAACGCTAGTGTAAT |
| 7_hypothetical | A2 | Tb927.6.2860_for | GGGGACAAGTTTGTACAAAAAAGCAGGCTATGCAACAGCGTTCCATCAGT | Tb927.6.2860_rev | GGGGACCACTTTGTACAAGAAAGCTGGGTTCAGTTAAACCTATAAACCATAGCTTTGCC |
| 7_hypothetical | A3 | Tb927.7.4610_for | GGGGACAAGTTTGTACAAAAAAGCAGGCTATGGAAGCTGCTGCCGAGC | Tb927.7.4610_rev | GGGGACCACTTTGTACAAGAAAGCTGGGTTCACGTTTCGTCGTTGCTTTGC |
| 7_hypothetical | A4 | Tb927.10.5870_for | GGGGACAAGTTTGTACAAAAAAGCAGGCTATGATTTTTCAAGACACGAGCTCCT | Tb927.10.5870_rev | GGGGACCACTTTGTACAAGAAAGCTGGGTTCAACGACGATGACGAAGCATG |
| 7_hypothetical | A5 | Tb927.7.2960_for | GGGGACAAGTTTGTACAAAAAAGCAGGCTATGGCGGGCATCAGGTTTG | Tb927.7.2960_rev | GGGGACCACTTTGTACAAGAAAGCTGGGTTCACCTGAGCTGTTGCATGC |
| 7_hypothetical | A6 | Tb927.6.3810_for | GGGGACAAGTTTGTACAAAAAAGCAGGCTATGTTTGGTCCTGTAGACGATGC | Tb927.6.3810_rev | GGGGACCACTTTGTACAAGAAAGCTGGGTTCACAACGCTCGAATAAGGAAGATTT |
| 7_hypothetical | A7 | Tb927.10.13230_for | GGGGACAAGTTTGTACAAAAAAGCAGGCTATGATGAGCTCTCGCTCCGT | Tb927.10.13230_rev | GGGGACCACTTTGTACAAGAAAGCTGGGTTCAGAGAGAAATTACGTCGACTGTTTG |
| 7_hypothetical | A8 | Tb927.10.13640_for | GGGGACAAGTTTGTACAAAAAAGCAGGCTATGGCCGAAACCTTTCAGTTGC | Tb927.10.13640_rev | GGGGACCACTTTGTACAAGAAAGCTGGGTTTAACCATCCTCGTGACCCAGC |
| 7_hypothetical | A9 | Tb927.9.2310_for | GGGGACAAGTTTGTACAAAAAAGCAGGCTATGGGTGATTTGTTAGAACTGACACC | Tb927.9.2310_rev | GGGGACCACTTTGTACAAGAAAGCTGGGTTCACCAATACTTATGTGACTCTTTCATTCT |
| 7_hypothetical | A10 | Tb927.11.3670_for | GGGGACAAGTTTGTACAAAAAAGCAGGCTATGCTCGTATGTTGTCGAAGCAG | Tb927.11.3670_rev | GGGGACCACTTTGTACAAGAAAGCTGGGTCTATGCAGTTCGCGATACAGCA |
| 7_hypothetical | A11 | Tb927.4.3670_for | GGGGACAAGTTTGTACAAAAAAGCAGGCTATGGAGAAAGAATTCACGAAGGCAC | Tb927.4.3670_rev | GGGGACCACTTTGTACAAGAAAGCTGGGTTTACACAAACAGCAGCGCTCC |
| 7_hypothetical | A12 | Tb927.10.8060_for | GGGGACAAGTTTGTACAAAAAAGCAGGCTATGGTTCTCCCCTCACCACATT | Tb927.10.8060_rev | GGGGACCACTTTGTACAAGAAAGCTGGGTTTATTTCTCACTTCCATTGACCTGACG |
| 7_hypothetical | A13 | Tb927.11.10770_for | GGGGACAAGTTTGTACAAAAAAGCAGGCTATGGTGCTCCTCTCTGTTCTGG | Tb927.11.10770_rev | GGGGACCACTTTGTACAAGAAAGCTGGGTCTACTTTGGTGCCTGTCGCG |
| 7_hypothetical | A14 | Tb927.7.4810_for | GGGGACAAGTTTGTACAAAAAAGCAGGCTATGAGAAATGGAATTATCTGTCGATACTTG | Tb927.7.4810_rev | GGGGACCACTTTGTACAAGAAAGCTGGGTTTATGGAGTATTGTAGAAGGGAAGTGAGT |
| 7_hypothetical | A15 | Tb927.9.10840_for | GGGGACAAGTTTGTACAAAAAAGCAGGCTATGCACCGGTGCGCTCTG | Tb927.9.10840_rev | GGGGACCACTTTGTACAAGAAAGCTGGGTTCATGTGAAGGACACGGGTCT |
| 7_hypothetical | A16 | Tb927.8.5370_for | GGGGACAAGTTTGTACAAAAAAGCAGGCTATGTCCCTCTCACCACAGTGG | Tb927.8.5370_rev | GGGGACCACTTTGTACAAGAAAGCTGGGTTCAGGCACTCGGCGCCAA |
| 7_hypothetical | A17 | Tb927.9.5880_for | GGGGACAAGTTTGTACAAAAAAGCAGGCTATGTATGGCATTAGTGAAGATGGTGG | Tb927.9.5880_rev | GGGGACCACTTTGTACAAGAAAGCTGGGTCTACAGACGGCGCTTCTTTTTCT |
| 7_hypothetical | A18 | Tb927.10.7040_for | GGGGACAAGTTTGTACAAAAAAGCAGGCTATGGTGGGTTGCAGAAGGTTG | Tb927.10.7040_rev | GGGGACCACTTTGTACAAGAAAGCTGGGTTCATCTTCTCCCCGATGTCTCAAC |
| 7_hypothetical | A19 | Tb927.10.9960_for | GGGGACAAGTTTGTACAAAAAAGCAGGCTATGACGGCGGGACTCGTT | Tb927.10.9960_rev | GGGGACCACTTTGTACAAGAAAGCTGGGTTCACGAGGACGACGCTGC |
| 7_hypothetical | A20 | Tb927.5.2920_for | GGGGACAAGTTTGTACAAAAAAGCAGGCTATGTCCAGAAGTGTTCCGTGGC | Tb927.5.2920_rev | GGGGACCACTTTGTACAAGAAAGCTGGGTTCAGACAGTACCATAAAAGAATTGTTCGT |
| 7_hypothetical | A21 | Tb927.8.970_for | GGGGACAAGTTTGTACAAAAAAGCAGGCTATGACGCGATTCACCTGTTTCA | Tb927.8.970_rev | GGGGACCACTTTGTACAAGAAAGCTGGGTCTACAAAAGACCAGTTCCTAGCAAACA |
| 7_hypothetical | A22 | Tb927.4.1110_for | GGGGACAAGTTTGTACAAAAAAGCAGGCTATGCTGGATGCCCTTCCACT | Tb927.4.1110_rev | GGGGACCACTTTGTACAAGAAAGCTGGGTCTACGAGGCTGAGGAGCATGC |
| 7_hypothetical | A23 | Tb927.7.4960_for | GGGGACAAGTTTGTACAAAAAAGCAGGCTATGGGTGAAAATGACAACGCTCA | Tb927.7.4960_rev | GGGGACCACTTTGTACAAGAAAGCTGGGTTCACACCGGATGATGTGTAGCT |
| 7_hypothetical | A24 | Tb927.11.6000_for | GGGGACAAGTTTGTACAAAAAAGCAGGCTATGCGCCGCGTCTCTTTC | Tb927.11.6000_rev | GGGGACCACTTTGTACAAGAAAGCTGGGTCTAACTGAGCATTGACCACGATACTT |
| 7_hypothetical | B1 | Tb927.7.2130_for | GGGGACAAGTTTGTACAAAAAAGCAGGCTATGGCGCGGGTAACGCTA | Tb927.7.2130_rev | GGGGACCACTTTGTACAAGAAAGCTGGGTTTAAGAGAGGTTGCCATACATTACTGGA |
| 7_hypothetical | B2 | Tb927.10.14190_for | GGGGACAAGTTTGTACAAAAAAGCAGGCTATGATTCATCCGAAGCTTATGCCTTATC | Tb927.10.14190_rev | GGGGACCACTTTGTACAAGAAAGCTGGGTTCAACAATGCCGTCGCTCG |
| 7_hypothetical | B3 | Tb927.9.9700_for | GGGGACAAGTTTGTACAAAAAAGCAGGCTATGAGACTCCGGCGCATCG | Tb927.9.9700_rev | GGGGACCACTTTGTACAAGAAAGCTGGGTCTAGTGATGGTTGTTCCGGTAAAAACG |
| 7_hypothetical | B4 | Tb927.11.3720_for | GGGGACAAGTTTGTACAAAAAAGCAGGCTATGATCCGTTGGGACCATGGG | Tb927.11.3720_rev | GGGGACCACTTTGTACAAGAAAGCTGGGTCTAATGATCGCCGCCACGG |
| 7_hypothetical | B5 | Tb927.11.10360_for | GGGGACAAGTTTGTACAAAAAAGCAGGCTATGGCCACGACTGGGTATACTC | Tb927.11.10360_rev | GGGGACCACTTTGTACAAGAAAGCTGGGTTCATTCCTCCCACAGCTTTCTCA |
| 7_hypothetical | B6 | Tb927.4.1050_for | GGGGACAAGTTTGTACAAAAAAGCAGGCTATGCGCTTCCTCGGTGCA | Tb927.4.1050_rev | GGGGACCACTTTGTACAAGAAAGCTGGGTCTAACAATGAAACTCAGGTGCGCT |
| 7_hypothetical | B7 | Tb927.11.2610_for | GGGGACAAGTTTGTACAAAAAAGCAGGCTATGGCAGCCGCTGTTACAATT | Tb927.11.2610_rev | GGGGACCACTTTGTACAAGAAAGCTGGGTCTACGACGTGCGTTCGACTT |
| 7_hypothetical | B8 | Tb927.11.3840_for | GGGGACAAGTTTGTACAAAAAAGCAGGCTATGGCCCGTCGGCAACGT | Tb927.11.3840_rev | GGGGACCACTTTGTACAAGAAAGCTGGGTTCAGGAGAAAAGTGGAAATTGGTGAC |
| 7_hypothetical | B9 | Tb927.4.640_for | GGGGACAAGTTTGTACAAAAAAGCAGGCTATGGACGCTGTTGTAAATAGTGAAGT | Tb927.4.640_rev | GGGGACCACTTTGTACAAGAAAGCTGGGTTCAAAACCTTCTCTTGACCCACCC |
| 7_hypothetical | B10 | Tb927.9.13300_for | GGGGACAAGTTTGTACAAAAAAGCAGGCTATGCCGCCGGCAGGTGGA | Tb927.9.13300_rev | GGGGACCACTTTGTACAAGAAAGCTGGGTTTAGTAGTCAACCTTCACATAAGCCGA |
| 7_hypothetical | B11 | Tb927.5.1690_for | GGGGACAAGTTTGTACAAAAAAGCAGGCTATGGAGAGCTACAAGGATGTGATTCT | Tb927.5.1690_rev | GGGGACCACTTTGTACAAGAAAGCTGGGTTCATATGGCAGGGCAACGATACA |
| 7_hypothetical | B12 | Tb927.7.590_for | GGGGACAAGTTTGTACAAAAAAGCAGGCTATGTTGTCTGCTCCACGACGC | Tb927.7.590_rev | GGGGACCACTTTGTACAAGAAAGCTGGGTTCAGTACAGAAGAGAGGAGAGGCG |
| 7_hypothetical | B13 | Tb927.11.170_for | GGGGACAAGTTTGTACAAAAAAGCAGGCTATGAGATGGACTGCTGTGTTATGTAAAG | Tb927.11.170_rev | GGGGACCACTTTGTACAAGAAAGCTGGGTCTATCGCACTCCATAACTTCGAAGG |
| 7_hypothetical | B14 | Tb927.6.880_for | GGGGACAAGTTTGTACAAAAAAGCAGGCTATGGTGTACATTCTTGCCCATGATG | Tb927.6.880_rev | GGGGACCACTTTGTACAAGAAAGCTGGGTCTACGCATTTTTCTTAGCATTTAGCGC |
| 7_hypothetical | B15 | Tb927.9.1560_for | GGGGACAAGTTTGTACAAAAAAGCAGGCTATGGACGTCGTGGTTCTTGC | Tb927.9.1560_rev | GGGGACCACTTTGTACAAGAAAGCTGGGTCTAATCGTCATCGTCTGAAGAAGTTTCT |
| 7_hypothetical | B16 | Tb927.11.1940_for | GGGGACAAGTTTGTACAAAAAAGCAGGCTATGGGCTCTGAATGCTCAAGGC | Tb927.11.1940_rev | GGGGACCACTTTGTACAAGAAAGCTGGGTTTACTCTTGTAGAAGCGATACTAGTCTTTT |
| 7_hypothetical | B17 | Tb927.11.6060_for | GGGGACAAGTTTGTACAAAAAAGCAGGCTATGGAAGGGGGGTTAAGAAAGCG | Tb927.11.6060_rev | GGGGACCACTTTGTACAAGAAAGCTGGGTTTAGCCCATCAAAAAGGTGGGAAAC |
| 7_hypothetical | B18 | Tb927.10.15770_for | GGGGACAAGTTTGTACAAAAAAGCAGGCTATGATCCAGTCGTCTAAAGAGCGA | Tb927.10.15770_rev | GGGGACCACTTTGTACAAGAAAGCTGGGTCTACACGGCGTCGTGACC |
| 7_hypothetical | B19 | Tb927.11.6480_for | GGGGACAAGTTTGTACAAAAAAGCAGGCTATGGAGATTGCTATAGTTGCTGCAC | Tb927.11.6480_rev | GGGGACCACTTTGTACAAGAAAGCTGGGTTCACCTGCCAGCCGCACG |
| 7_hypothetical | B20 | Tb927.6.2870_for | GGGGACAAGTTTGTACAAAAAAGCAGGCTATGTATCGTTTTTCAATCAGTGGTAGTGC | Tb927.6.2870_rev | GGGGACCACTTTGTACAAGAAAGCTGGGTTTATACACAGCGCTGTTGTAACTCC |
| 7_hypothetical | B21 | Tb927.11.8710_for | GGGGACAAGTTTGTACAAAAAAGCAGGCTATGTGGCGTCACAGCTGTATTC | Tb927.11.8710_rev | GGGGACCACTTTGTACAAGAAAGCTGGGTTTACGGTTCAATGGTGGCGTG |
| 7_hypothetical | B22 | Tb927.9.1710_for | GGGGACAAGTTTGTACAAAAAAGCAGGCTATGACGGGGTACAACTTGAAAACTATAG | Tb927.9.1710_rev | GGGGACCACTTTGTACAAGAAAGCTGGGTCTAAGAAGTTACGTCAGCGCAGT |
| 7_hypothetical | B23 | Tb927.8.910_for | GGGGACAAGTTTGTACAAAAAAGCAGGCTATGACTGAATTCAATTACGATACGATGTTT | Tb927.8.910_rev | GGGGACCACTTTGTACAAGAAAGCTGGGTTTAATTGAAGCGACATCCCATGGG |
| 7_hypothetical | B24 | Tb927.11.4860_for | GGGGACAAGTTTGTACAAAAAAGCAGGCTATGGCCGTGGCCTACGTT | Tb927.11.4860_rev | GGGGACCACTTTGTACAAGAAAGCTGGGTTTACTGCGGTGGGAAGAAGTGT |
| 7_hypothetical | C1 | Tb927.10.14520_for | GGGGACAAGTTTGTACAAAAAAGCAGGCTATGAGCGCGGACACTATTTGTC | Tb927.10.14520_rev | GGGGACCACTTTGTACAAGAAAGCTGGGTTTACAAATGGTGAGCGCTTGTGC |
| 7_hypothetical | C2 | Tb927.7.660_for | GGGGACAAGTTTGTACAAAAAAGCAGGCTATGCCAAATGAAAAGGCGCAGA | Tb927.7.660_rev | GGGGACCACTTTGTACAAGAAAGCTGGGTTTACAATAGAAGACCAAACTCGCGC |
| 7_hypothetical | C3 | Tb927.8.4250_for | GGGGACAAGTTTGTACAAAAAAGCAGGCTATGTGGTTCAACCACCGCATC | Tb927.8.4250_rev | GGGGACCACTTTGTACAAGAAAGCTGGGTTTACAAAACGAACTTGTACGGCTTTC |
| 7_hypothetical | C4 | Tb927.5.2470_for | GGGGACAAGTTTGTACAAAAAAGCAGGCTATGGTAAACAATGAAGGGTATTCGCA | Tb927.5.2470_rev | GGGGACCACTTTGTACAAGAAAGCTGGGTTTATGAGGGAGTGATATCTGCAGGTG |
| 7_hypothetical | C5 | Tb927.5.1780_for | GGGGACAAGTTTGTACAAAAAAGCAGGCTATGTCATCGGTGGGAATCGAGT | Tb927.5.1780_rev | GGGGACCACTTTGTACAAGAAAGCTGGGTCTAACGAGGAAGCGATGCGAG |
| 7_hypothetical | C6 | Tb927.10.7250_for | GGGGACAAGTTTGTACAAAAAAGCAGGCTATGCCGGCCACGTTTAGC | Tb927.10.7250_rev | GGGGACCACTTTGTACAAGAAAGCTGGGTTTACCCACTGGGTGGGGC |
| 7_hypothetical | C7 | Tb927.10.5900_for | GGGGACAAGTTTGTACAAAAAAGCAGGCTATGTCTTTTTACCGTACAAAAAAGGTTATA | Tb927.10.5900_rev | GGGGACCACTTTGTACAAGAAAGCTGGGTTCACATAGCTACAGGGGCCTC |
| 7_hypothetical | C8 | Tb927.3.2160_for | GGGGACAAGTTTGTACAAAAAAGCAGGCTATGACGGGAGAGCCTGTTCG | Tb927.3.2160_rev | GGGGACCACTTTGTACAAGAAAGCTGGGTTCAGGGTCCCAAATTATCGGGG |
| 7_hypothetical | C9 | Tb927.4.1430_for | GGGGACAAGTTTGTACAAAAAAGCAGGCTATGACCTTGCCTGAATTGGCG | Tb927.4.1430_rev | GGGGACCACTTTGTACAAGAAAGCTGGGTTCATCCTTGGCTAGCTGGGTAAG |
| 7_hypothetical | C10 | Tb927.1.4800_for | GGGGACAAGTTTGTACAAAAAAGCAGGCTATGCGCTCGCGCGAGGTG | Tb927.1.4800_rev | GGGGACCACTTTGTACAAGAAAGCTGGGTTTATGCACACCGCCTTCCCC |
| 7_hypothetical | C11 | Tb927.10.9680_for | GGGGACAAGTTTGTACAAAAAAGCAGGCTATGAGTGTTTGCCTTGGCACG | Tb927.10.9680_rev | GGGGACCACTTTGTACAAGAAAGCTGGGTTTAACACATTCCTTTCACGTACTCCTG |
| 7_hypothetical | C12 | Tb927.4.3140_for | GGGGACAAGTTTGTACAAAAAAGCAGGCTATGAGCGCGCGCATGCAG | Tb927.4.3140_rev | GGGGACCACTTTGTACAAGAAAGCTGGGTTCACTCCTCTCCTACCCCTTCC |
| 7_hypothetical | C13 | Tb927.9.10400_for | GGGGACAAGTTTGTACAAAAAAGCAGGCTATGTCTGTGTTACGTCTTCCAAAAGAT | Tb927.9.10400_rev | GGGGACCACTTTGTACAAGAAAGCTGGGTTTAATCGCTGTCATCGAACCTTGTG |
| 7_hypothetical | C14 | Tb927.9.6650_for | GGGGACAAGTTTGTACAAAAAAGCAGGCTATGAAGACTGTAATTTCCACTGCTTTTGC | Tb927.9.6650_rev | GGGGACCACTTTGTACAAGAAAGCTGGGTTTACCCCTTGGAAGTCAACTCCTT |
| 7_hypothetical | C15 | Tb927.9.14290_for | GGGGACAAGTTTGTACAAAAAAGCAGGCTATGACCGCCTCAGCGAGG | Tb927.9.14290_rev | GGGGACCACTTTGTACAAGAAAGCTGGGTCTAAAACATTGCACAGCATGGCG |
| 7_hypothetical | C16 | Tb927.6.1130_for | GGGGACAAGTTTGTACAAAAAAGCAGGCTATGGGGCGGAGGTCCGAT | Tb927.6.1130_rev | GGGGACCACTTTGTACAAGAAAGCTGGGTCTAAGCGGGGGTGATTGTTTCTG |
| 7_hypothetical | C17 | Tb927.10.950_for | GGGGACAAGTTTGTACAAAAAAGCAGGCTATGCTGCGTGGCGTGACA | Tb927.10.950_rev | GGGGACCACTTTGTACAAGAAAGCTGGGTCTAACTGGACAGCATTAACGGAACC |
| 7_hypothetical | C18 | Tb927.6.1200_for | GGGGACAAGTTTGTACAAAAAAGCAGGCTATGCGGCGATGCGCTACA | Tb927.6.1200_rev | GGGGACCACTTTGTACAAGAAAGCTGGGTTTACTTCTCACTTAATAATTCCAACCAGTT |
| 7_hypothetical | C19 | Tb927.10.10200_for | GGGGACAAGTTTGTACAAAAAAGCAGGCTATGGCCATCACGGTGCAGC | Tb927.10.10200_rev | GGGGACCACTTTGTACAAGAAAGCTGGGTCTACTTCGGCCGTTCACCTCC |
| 7_hypothetical | C20 | Tb927.6.4940_for | GGGGACAAGTTTGTACAAAAAAGCAGGCTATGAGGCCACGATGTGGCT | Tb927.6.4940_rev | GGGGACCACTTTGTACAAGAAAGCTGGGTTTACACAACACCGTCCACGC |
| 7_hypothetical | C21 | Tb927.3.5260_for | GGGGACAAGTTTGTACAAAAAAGCAGGCTATGTTAAGTTTTTTGATTGACTGCAGCG | Tb927.3.5260_rev | GGGGACCACTTTGTACAAGAAAGCTGGGTTTATGTCACATGAGCCTTCCCCC |
| 7_hypothetical | C22 | Tb927.7.5140_for | GGGGACAAGTTTGTACAAAAAAGCAGGCTATGGGCTCCACGGAGGATG | Tb927.7.5140_rev | GGGGACCACTTTGTACAAGAAAGCTGGGTTTATACAATCCCTCCAGTGCTGTTCT |
| 7_hypothetical | C23 | Tb927.10.4260_for | GGGGACAAGTTTGTACAAAAAAGCAGGCTATGGATTGGGTATTTTTTCTCATGGCG | Tb927.10.4260_rev | GGGGACCACTTTGTACAAGAAAGCTGGGTTTACCACCGACTGACTCTAGCG |
| 7_hypothetical | C24 | Tb927.10.5470_for | GGGGACAAGTTTGTACAAAAAAGCAGGCTATGTTCCTTGCAGATGGTAGACGA | Tb927.10.5470_rev | GGGGACCACTTTGTACAAGAAAGCTGGGTTTATGGTCTTTTGTCCTCCCACAAC |
| 7_hypothetical | D1 | Tb927.11.13200_for | GGGGACAAGTTTGTACAAAAAAGCAGGCTATGGCCCGTGGCGGTGTG | Tb927.11.13200_rev | GGGGACCACTTTGTACAAGAAAGCTGGGTTCACTCATCATCAGAGCTTTGGGG |
| 7_hypothetical | D2 | Tb927.9.6370_for | GGGGACAAGTTTGTACAAAAAAGCAGGCTATGAGCAGCACAGACAATGATATCG | Tb927.9.6370_rev | GGGGACCACTTTGTACAAGAAAGCTGGGTTTACCGCAAAACCTTGCCGC |
| 7_hypothetical | D3 | Tb927.2.1850_for | GGGGACAAGTTTGTACAAAAAAGCAGGCTATGGCAGAGGGGGGTTTTAATATGC | Tb927.2.1850_rev | GGGGACCACTTTGTACAAGAAAGCTGGGTTCATGATGGGAGGAGACTCACAAC |
| 7_hypothetical | D4 | Tb927.8.950_for | GGGGACAAGTTTGTACAAAAAAGCAGGCTATGACACGATTCACCTGTTTCACA | Tb927.8.950_rev | GGGGACCACTTTGTACAAGAAAGCTGGGTCTACAAAAGACCAACTAATGTGAAAGTAAG |
| 7_hypothetical | D5 | Tb927.8.930_for | GGGGACAAGTTTGTACAAAAAAGCAGGCTATGACGCGATTCACCTGTTTCA | Tb927.8.930_rev | GGGGACCACTTTGTACAAGAAAGCTGGGTCTACAAAAGACCAACTAATGTGAAAGTAAG |
| 7_hypothetical | D6 | Tb927.8.2840_for | GGGGACAAGTTTGTACAAAAAAGCAGGCTATGGACATCCTGCGGAGGAC | Tb927.8.2840_rev | GGGGACCACTTTGTACAAGAAAGCTGGGTCTACGTGGCCAACCTCGTCG |
| 7_hypothetical | D7 | Tb927.4.610_for | GGGGACAAGTTTGTACAAAAAAGCAGGCTATGTCCGAAAACCCCGCTCA | Tb927.4.610_rev | GGGGACCACTTTGTACAAGAAAGCTGGGTTCACTGAGAACCCTCCTCCGG |
| 7_hypothetical | D8 | Tb927.8.5610_for | GGGGACAAGTTTGTACAAAAAAGCAGGCTATGCATGCGTATCATTTCCAGCA | Tb927.8.5610_rev | GGGGACCACTTTGTACAAGAAAGCTGGGTTCAAGAAAGGAGTACTAAGATAGGGTACG |
| 7_hypothetical | D9 | Tb927.11.4830_for | GGGGACAAGTTTGTACAAAAAAGCAGGCTATGTTCACTGTATCTTCATCGATGCG | Tb927.11.4830_rev | GGGGACCACTTTGTACAAGAAAGCTGGGTTCAAGCTGTATTTACACTTTGCTGTACT |
| 7_hypothetical | D10 | Tb927.4.3030_for | GGGGACAAGTTTGTACAAAAAAGCAGGCTATGACAGATGGAAACCCTGGCG | Tb927.4.3030_rev | GGGGACCACTTTGTACAAGAAAGCTGGGTTCAGCGCGTCACCGCTTC |
| 7_hypothetical | D11 | Tb927.9.7170_for | GGGGACAAGTTTGTACAAAAAAGCAGGCTATGCAGCGTAATTTAACGAAATTGAGC | Tb927.9.7170_rev | GGGGACCACTTTGTACAAGAAAGCTGGGTCTAAATGTCAGTAACGCCCACGT |
| 7_hypothetical | D12 | Tb927.10.9060_for | GGGGACAAGTTTGTACAAAAAAGCAGGCTATGTCCTCCACGACACTAGTGG | Tb927.10.9060_rev | GGGGACCACTTTGTACAAGAAAGCTGGGTTCACGCAGCAACGTCCTCG |
| 7_hypothetical | D13 | Tb927.10.11210_for | GGGGACAAGTTTGTACAAAAAAGCAGGCTATGATCTCGTGTGTTGCTGGTC | Tb927.10.11210_rev | GGGGACCACTTTGTACAAGAAAGCTGGGTTCAAGTGCGTGTTTTTTTCATGGC |
| 7_hypothetical | D14 | Tb927.10.4850_for | GGGGACAAGTTTGTACAAAAAAGCAGGCTATGATCCGGCGTTATTCACTAGC | Tb927.10.4850_rev | GGGGACCACTTTGTACAAGAAAGCTGGGTTCACTGTAAACCGTATGCCTTCTTTAC |
| 7_hypothetical | D15 | Tb927.11.1570_for | GGGGACAAGTTTGTACAAAAAAGCAGGCTATGAATAATGGAAGTGCTATGGAGGAGG | Tb927.11.1570_rev | GGGGACCACTTTGTACAAGAAAGCTGGGTTCAGGTCTTTCCGAAAGTGCG |
| 7_hypothetical | D16 | Tb927.5.1290_for | GGGGACAAGTTTGTACAAAAAAGCAGGCTATGTCGTGCGACACTGTCAC | Tb927.5.1290_rev | GGGGACCACTTTGTACAAGAAAGCTGGGTTCACCTGCGGTGAGAGTGAA |
| 7_hypothetical | D17 | Tb927.8.5940_for | GGGGACAAGTTTGTACAAAAAAGCAGGCTATGGACAAAGCAAGGCCGTTT | Tb927.8.5940_rev | GGGGACCACTTTGTACAAGAAAGCTGGGTTCAGCTACCTCCGTGAGGGG |
| 7_hypothetical | D18 | Tb927.7.3710_for | GGGGACAAGTTTGTACAAAAAAGCAGGCTATGCACGACGGTGCGGGG | Tb927.7.3710_rev | GGGGACCACTTTGTACAAGAAAGCTGGGTTTACGAAGGTACATCCAAAAACAACCT |
| 7_hypothetical | D19 | Tb927.11.14800_for | GGGGACAAGTTTGTACAAAAAAGCAGGCTATGCCCTCAGAATCTTTGGCCG | Tb927.11.14800_rev | GGGGACCACTTTGTACAAGAAAGCTGGGTCTATCGCTGACGCAGATTGCA |
| 7_hypothetical | D20 | Tb927.11.6820_for | GGGGACAAGTTTGTACAAAAAAGCAGGCTATGACTTTTGTTGTTATGTGTGACGGC | Tb927.11.6820_rev | GGGGACCACTTTGTACAAGAAAGCTGGGTTCATAGAATTGACGTCTCCTTACTCGG |
| 7_hypothetical | D21 | Tb927.11.2750_for | GGGGACAAGTTTGTACAAAAAAGCAGGCTATGTTTCCCTTTGATGAAAGGTCACC | Tb927.11.2750_rev | GGGGACCACTTTGTACAAGAAAGCTGGGTTCACATTACAGTACATCGAGGGCA |
| 7_hypothetical | D22 | Tb927.11.10260_for | GGGGACAAGTTTGTACAAAAAAGCAGGCTATGTGTGACGAGTTCTTTGGAGAC | Tb927.11.10260_rev | GGGGACCACTTTGTACAAGAAAGCTGGGTTTATAAATCGCGGCATGTAACTCTAATCG |
| 7_hypothetical | D23 | Tb927.11.9250_for | GGGGACAAGTTTGTACAAAAAAGCAGGCTATGCTGCCTCGCACAGCA | Tb927.11.9250_rev | GGGGACCACTTTGTACAAGAAAGCTGGGTCTACGGTTTTGATAGAGCTCCATTAGC |
| 7_hypothetical | D24 | Tb927.10.2180_for | GGGGACAAGTTTGTACAAAAAAGCAGGCTATGTCTTGCGGTGGCGCA | Tb927.10.2180_rev | GGGGACCACTTTGTACAAGAAAGCTGGGTTCATTCGACTACCAACGCGC |
| 7_hypothetical | E1 | Tb927.3.1830_for | GGGGACAAGTTTGTACAAAAAAGCAGGCTATGGAGCAGCCGAGCGCT | Tb927.3.1830_rev | GGGGACCACTTTGTACAAGAAAGCTGGGTTTACTCCAATTGTCGCACATGAACG |
| 7_hypothetical | E2 | Tb927.3.2360_for | GGGGACAAGTTTGTACAAAAAAGCAGGCTATGATTCACTTCGTCTTACTATGCTATCTC | Tb927.3.2360_rev | GGGGACCACTTTGTACAAGAAAGCTGGGTCTAGGTTGAACAAAAAAAATAAGCGGTAAG |
| 7_hypothetical | E3 | Tb927.11.4320_for | GGGGACAAGTTTGTACAAAAAAGCAGGCTATGGAGAAAATTGTTCGCACTGTATACA | Tb927.11.4320_rev | GGGGACCACTTTGTACAAGAAAGCTGGGTCTAAGCATCTGGTGGTGCATGC |
| 7_hypothetical | E4 | Tb927.9.11140_for | GGGGACAAGTTTGTACAAAAAAGCAGGCTATGAGTTATTTGGTCGAAGAATTGCCT | Tb927.9.11140_rev | GGGGACCACTTTGTACAAGAAAGCTGGGTCTACATATTATCGGACTGGTTGATTGTGC |
| 7_hypothetical | E5 | Tb927.5.2210_for | GGGGACAAGTTTGTACAAAAAAGCAGGCTATGGCATCAGACAATGAGTCCACT | Tb927.5.2210_rev | GGGGACCACTTTGTACAAGAAAGCTGGGTTCATCGTCTTCGCCGGGA |
| 7_hypothetical | E6 | Tb927.9.8670_for | GGGGACAAGTTTGTACAAAAAAGCAGGCTATGGTTGTACGTACTTTAACTGGCAA | Tb927.9.8670_rev | GGGGACCACTTTGTACAAGAAAGCTGGGTTCAAACCCGGTCCCAGATAACT |
| 7_hypothetical | E8 | Tb927.10.9020_for | GGGGACAAGTTTGTACAAAAAAGCAGGCTATGATGGGTGATGCTGGTAGTTTCT | Tb927.10.9020_rev | GGGGACCACTTTGTACAAGAAAGCTGGGTCTACGAGTCTGAAAGATTTCCTTCTTGT |
| 7_hypothetical | E10 | Tb927.11.15050_for | GGGGACAAGTTTGTACAAAAAAGCAGGCTATGATAGGAGAAGCTGTTCGTCTGT | Tb927.11.15050_rev | GGGGACCACTTTGTACAAGAAAGCTGGGTCTAAACTGCGGTGACGGAACC |
| 7_hypothetical | E11 | Tb927.8.1980_for | GGGGACAAGTTTGTACAAAAAAGCAGGCTATGCAAAATAAGTCTTGGGCCCTG | Tb927.8.1980_rev | GGGGACCACTTTGTACAAGAAAGCTGGGTTTATACGTCATTTACTATTAACTCCATTAT |
| 7_hypothetical | E12 | Tb927.8.1450_for | GGGGACAAGTTTGTACAAAAAAGCAGGCTATGATCCGCTGGACTGTAAATCTCG | Tb927.8.1450_rev | GGGGACCACTTTGTACAAGAAAGCTGGGTTCACTCCGAACCGTCAGAGTG |
| 7_hypothetical | E13 | Tb927.8.2760_for | GGGGACAAGTTTGTACAAAAAAGCAGGCTATGACGAGCAAGCATGTGGTG | Tb927.8.2760_rev | GGGGACCACTTTGTACAAGAAAGCTGGGTTTAAACAATGGCGCTGAAGTCAGT |
| 7_hypothetical | E14 | Tb927.10.410_for | GGGGACAAGTTTGTACAAAAAAGCAGGCTATGACTTCCGTGGGGTTCCC | Tb927.10.410_rev | GGGGACCACTTTGTACAAGAAAGCTGGGTCTACATTCGCACAGAGTGACTGC |
| 7_hypothetical | E15 | Tb927.5.2270_for | GGGGACAAGTTTGTACAAAAAAGCAGGCTATGAACACCGCGTCAACTGTT | Tb927.5.2270_rev | GGGGACCACTTTGTACAAGAAAGCTGGGTCTACCGAACACTGACAATCTTTGATAATTT |
| 7_hypothetical | E16 | Tb927.11.8930_for | GGGGACAAGTTTGTACAAAAAAGCAGGCTATGGATTTCCGCCTCCCCC | Tb927.11.8930_rev | GGGGACCACTTTGTACAAGAAAGCTGGGTTTAAAGCTCCTGTGACCGCTGT |
| 7_hypothetical | E17 | Tb927.6.2840_for | GGGGACAAGTTTGTACAAAAAAGCAGGCTATGGTCAAACTTGACGTCTCACTC | Tb927.6.2840_rev | GGGGACCACTTTGTACAAGAAAGCTGGGTTTAGTCAAAGAGTCCTTCATTTTCCGC |
| 7_hypothetical | E18 | Tb927.8.5760_for | GGGGACAAGTTTGTACAAAAAAGCAGGCTATGAACACACCACTGCATGCC | Tb927.8.5760_rev | GGGGACCACTTTGTACAAGAAAGCTGGGTTCACCCCCCACCGTACCG |
| 7_hypothetical | E19 | Tb927.3.1390_for | GGGGACAAGTTTGTACAAAAAAGCAGGCTATGTTGAACTGGTGGCTCATAGTACT | Tb927.3.1390_rev | GGGGACCACTTTGTACAAGAAAGCTGGGTTTATGTTGAATAGTTTGGCTTCTTGGGT |
| 7_hypothetical | E20 | Tb927.3.1930_for | GGGGACAAGTTTGTACAAAAAAGCAGGCTATGCTTTTTGACCTTGCACCGC | Tb927.3.1930_rev | GGGGACCACTTTGTACAAGAAAGCTGGGTTCACGCCCATGTCTTAATGGC |
| 7_hypothetical | E21 | Tb927.5.2740_for | GGGGACAAGTTTGTACAAAAAAGCAGGCTATGGATGAACTATCGGATATTTATGACTCC | Tb927.5.2740_rev | GGGGACCACTTTGTACAAGAAAGCTGGGTTTATTTCGGATATGCCCCATCTGC |
| 7_hypothetical | E22 | Tb927.10.3910_for | GGGGACAAGTTTGTACAAAAAAGCAGGCTATGAAGCGATCAGATGACGTCAGT | Tb927.10.3910_rev | GGGGACCACTTTGTACAAGAAAGCTGGGTTCACAATAAGTGACTGCTGAAATTCTTTCG |
| 7_hypothetical | E23 | Tb927.9.6360_for | GGGGACAAGTTTGTACAAAAAAGCAGGCTATGAGCAGCACAGACAATGATATCG | Tb927.9.6360_rev | GGGGACCACTTTGTACAAGAAAGCTGGGTTCAAGTTCGTGATTTCGCTTTTGC |
| 7_hypothetical | E24 | Tb927.6.2570_for | GGGGACAAGTTTGTACAAAAAAGCAGGCTATGCTCAGTGGAACTACTTTTGGAG | Tb927.6.2570_rev | GGGGACCACTTTGTACAAGAAAGCTGGGTTCAAATGACCGACCGCCAAC |
| 7_hypothetical | F2 | Tb927.5.4100_for | GGGGACAAGTTTGTACAAAAAAGCAGGCTATGCAGAGGCCCCTGACG | Tb927.5.4100_rev | GGGGACCACTTTGTACAAGAAAGCTGGGTTTAAAGGCTCAGTTTGAAAAAGCGAAC |
| 7_hypothetical | F3 | Tb927.5.2870_for | GGGGACAAGTTTGTACAAAAAAGCAGGCTATGCCGCACAAGGGGTGC | Tb927.5.2870_rev | GGGGACCACTTTGTACAAGAAAGCTGGGTTTAAACCCCCCAAACGTGTTCC |
| 7_hypothetical | F4 | Tb927.10.3740_for | GGGGACAAGTTTGTACAAAAAAGCAGGCTATGCCGCGGTTACATATTTGGAT | Tb927.10.3740_rev | GGGGACCACTTTGTACAAGAAAGCTGGGTTCACAACACGTTTGTAGGGGTATG |
| 7_hypothetical | F5 | Tb927.11.14090_for | GGGGACAAGTTTGTACAAAAAAGCAGGCTATGCAAGGCGCATACGGAG | Tb927.11.14090_rev | GGGGACCACTTTGTACAAGAAAGCTGGGTTTATGCTGAACCATTTTCCCCAGC |
| 7_hypothetical | F6 | Tb927.11.7650_for | GGGGACAAGTTTGTACAAAAAAGCAGGCTATGGAGCGCGAGCAACTG | Tb927.11.7650_rev | GGGGACCACTTTGTACAAGAAAGCTGGGTTCACGTGCTAGTCACTATAAGGGAC |
| 7_hypothetical | F7 | Tb927.6.3170_for | GGGGACAAGTTTGTACAAAAAAGCAGGCTATGACGTCCAGTCGTGCGG | Tb927.6.3170_rev | GGGGACCACTTTGTACAAGAAAGCTGGGTTTACTCCTTCTTCACCGATATAGTCGAG |

| | | | | | |
|---|---|---|---|---|---|
| 7_hypothetical | F8 | Tb927.10.1180_for | GGGGACAAGTTTGTACAAAAAAGCAGGCTATGCCCCTGACCTGTCTTGG | Tb927.10.1180_rev | GGGGACCACTTTGTACAAGAAAGCTGGGTTTAAAAACAAAAGGATCCGTCATCCTGG |
| 7_hypothetical | F9 | Tb927.4.3660_for | GGGGACAAGTTTGTACAAAAAAGCAGGCTATGAGATGTTTCGGGGCCTGC | Tb927.4.3660_rev | GGGGACCACTTTGTACAAGAAAGCTGGGTTCACGGAGTGGTGACGACG |
| 7_hypothetical | F10 | Tb927.3.1990_for | GGGGACAAGTTTGTACAAAAAAGCAGGCTATGGCTAGTGTTGGGCCTGT | Tb927.3.1990_rev | GGGGACCACTTTGTACAAGAAAGCTGGGTTTACAAGATGCCATTACTCATCCACTC |
| 7_hypothetical | F11 | Tb927.10.2310_for | GGGGACAAGTTTGTACAAAAAAGCAGGCTATGAACCGCAACGCGGCG | Tb927.10.2310_rev | GGGGACCACTTTGTACAAGAAAGCTGGGTTTAGTCCTCCATATCAATCTCTGGGATTT |
| 7_hypothetical | F12 | Tb927.4.1940_for | GGGGACAAGTTTGTACAAAAAAGCAGGCTATGGCGGACAACTCGTCTGC | Tb927.4.1940_rev | GGGGACCACTTTGTACAAGAAAGCTGGGTTCAGGTCTTCACGGGTACCCA |
| 7_hypothetical | F13 | Tb927.10.15470_for | GGGGACAAGTTTGTACAAAAAAGCAGGCTATGCGGGTCGATGTAGAAGCC | Tb927.10.15470_rev | GGGGACCACTTTGTACAAGAAAGCTGGGTTCATTGGACCATGTATACTCTCATATCCTT |
| 7_hypothetical | F14 | Tb927.10.10310_for | GGGGACAAGTTTGTACAAAAAAGCAGGCTATGCTGCGATCCTGTCGAAC | Tb927.10.10310_rev | GGGGACCACTTTGTACAAGAAAGCTGGGTCTACACACAGCGGCCAGAAC |
| 7_hypothetical | F15 | Tb927.8.1530_for | GGGGACAAGTTTGTACAAAAAAGCAGGCTATGCTGTTGAAGCTTGGCGC | Tb927.8.1530_rev | GGGGACCACTTTGTACAAGAAAGCTGGGTTTAATCTGCGGCGGCACAC |
| 7_hypothetical | F16 | Tb927.10.15450_for | GGGGACAAGTTTGTACAAAAAAGCAGGCTATGTACGTGGGTCCTTGGGAG | Tb927.10.15450_rev | GGGGACCACTTTGTACAAGAAAGCTGGGTTCAAACAATGCTATCAGGGTCCAAC |
| 7_hypothetical | F17 | Tb927.9.9300_for | GGGGACAAGTTTGTACAAAAAAGCAGGCTATGGCAACGGCACACTGTAC | Tb927.9.9300_rev | GGGGACCACTTTGTACAAGAAAGCTGGGTCTATCTCGTTCCTGGCCTAGTCTG |
| 7_hypothetical | F18 | Tb927.3.670_for | GGGGACAAGTTTGTACAAAAAAGCAGGCTATGCTATCAACGCCCTGTCCT | Tb927.3.670_rev | GGGGACCACTTTGTACAAGAAAGCTGGGTTTAGGTGATGGGAACACCACAACT |
| 7_hypothetical | F19 | Tb927.7.3510_for | GGGGACAAGTTTGTACAAAAAAGCAGGCTATGTGGCATTCGTCCCTCCG | Tb927.7.3510_rev | GGGGACCACTTTGTACAAGAAAGCTGGGTTCAGGAGCCATAACGCATGGA |
| 7_hypothetical | F20 | Tb927.5.1230_for | GGGGACAAGTTTGTACAAAAAAGCAGGCTATGGCTCACGTTGCAATCCG | Tb927.5.1230_rev | GGGGACCACTTTGTACAAGAAAGCTGGGTCTAGATTGGTGGGAATCGTTGCC |
| 7_hypothetical | F21 | Tb927.3.3460_for | GGGGACAAGTTTGTACAAAAAAGCAGGCTATGTTTTGCGTAAGTGCCGCA | Tb927.3.3460_rev | GGGGACCACTTTGTACAAGAAAGCTGGGTCTAAGCATCAGCAGAAGTAATCCTTTTAAG |
| 7_hypothetical | F22 | Tb927.10.11070_for | GGGGACAAGTTTGTACAAAAAAGCAGGCTATGGGCTCTCCGCCGCCT | Tb927.10.11070_rev | GGGGACCACTTTGTACAAGAAAGCTGGGTCTATTCGCTTGCATCGCATTTTTCC |
| 7_hypothetical | F23 | Tb927.8.4580_for | GGGGACAAGTTTGTACAAAAAAGCAGGCTATGATTGGTATGATGGCGCGTAG | Tb927.8.4580_rev | GGGGACCACTTTGTACAAGAAAGCTGGGTCTATCGCCGGGCCACCTT |
| 7_hypothetical | F24 | Tb927.9.2700_for | GGGGACAAGTTTGTACAAAAAAGCAGGCTATGCTTCGGTGGACCGGA | Tb927.9.2700_rev | GGGGACCACTTTGTACAAGAAAGCTGGGTCTACCGTGACCGCTGACGA |
| 7_hypothetical | G1 | Tb927.10.6620_for | GGGGACAAGTTTGTACAAAAAAGCAGGCTATGCCTCACTGCCCGCGG | Tb927.10.6620_rev | GGGGACCACTTTGTACAAGAAAGCTGGGTTTAAGGGCATGCCGAAGCAG |
| 7_hypothetical | G2 | Tb927.4.4670_for | GGGGACAAGTTTGTACAAAAAAGCAGGCTATGACAACTTCCCCATCCACGC | Tb927.4.4670_rev | GGGGACCACTTTGTACAAGAAAGCTGGGTTCAGCTGAAGGCAGCGGA |
| 7_hypothetical | G3 | Tb927.11.5720_for | GGGGACAAGTTTGTACAAAAAAGCAGGCTATGATGAATCGTAACGAGAACTCTGG | Tb927.11.5720_rev | GGGGACCACTTTGTACAAGAAAGCTGGGTTTAGACATTTCCCGTGCCTCGC |
| 7_hypothetical | G4 | Tb927.11.9650_for | GGGGACAAGTTTGTACAAAAAAGCAGGCTATGCCCTCAGGTACAGACACTC | Tb927.11.9650_rev | GGGGACCACTTTGTACAAGAAAGCTGGGTTCAAGAAAAGTTCTTCCTCAACTGCC |
| 7_hypothetical | G5 | Tb927.10.2610_for | GGGGACAAGTTTGTACAAAAAAGCAGGCTATGACGGAGTTGGTTCCCGA | Tb927.10.2610_rev | GGGGACCACTTTGTACAAGAAAGCTGGGTTTACGCGCATCGAACACTGC |
| 7_hypothetical | G6 | Tb927.10.11690_for | GGGGACAAGTTTGTACAAAAAAGCAGGCTATGCAGGCCACTTCTAAATCTACTGG | Tb927.10.11690_rev | GGGGACCACTTTGTACAAGAAAGCTGGGTCTAGCGCTTCCACGACGTC |
| 7_hypothetical | G7 | Tb927.10.11910_for | GGGGACAAGTTTGTACAAAAAAGCAGGCTATGGAAGGGGCCGCTGGA | Tb927.10.11910_rev | GGGGACCACTTTGTACAAGAAAGCTGGGTCTAGCACCTCTCGCTTTCACCA |
| 7_hypothetical | G8 | Tb927.10.15930_for | GGGGACAAGTTTGTACAAAAAAGCAGGCTATGTTATGGAGCACACCGCCA | Tb927.10.15930_rev | GGGGACCACTTTGTACAAGAAAGCTGGGTTCAAGAACCCGACGCACGT |
| 7_hypothetical | G9 | Tb927.9.8330_for | GGGGACAAGTTTGTACAAAAAAGCAGGCTATGTCTCCTTTTGAGCAACAGCG | Tb927.9.8330_rev | GGGGACCACTTTGTACAAGAAAGCTGGGTTCATCTCTTCCGAGCCCCG |
| 7_hypothetical | G10 | Tb927.3.5110_for | GGGGACAAGTTTGTACAAAAAAGCAGGCTATGTCGATACAAGGGGACCCAA | Tb927.3.5110_rev | GGGGACCACTTTGTACAAGAAAGCTGGGTTTACCTCACACGCGCCTTTG |
| 7_hypothetical | G11 | Tb927.8.6300_for | GGGGACAAGTTTGTACAAAAAAGCAGGCTATGATAGCTGCAACTCAGGTGC | Tb927.8.6300_rev | GGGGACCACTTTGTACAAGAAAGCTGGGTTCATCCACCGAGTATCGGCG |
| 7_hypothetical | G12 | Tb927.7.2450_for | GGGGACAAGTTTGTACAAAAAAGCAGGCTATGAGTGGTGATGACGATGACGT | Tb927.7.2450_rev | GGGGACCACTTTGTACAAGAAAGCTGGGTTTAATTTCGTTGACTTACATTTGTAGCGG |
| 7_hypothetical | G13 | Tb927.10.11190_for | GGGGACAAGTTTGTACAAAAAAGCAGGCTATGCGTCACGCTGTCACAAA | Tb927.10.11190_rev | GGGGACCACTTTGTACAAGAAAGCTGGGTTTACCCCTTAACGCACAACAACTG |
| 7_hypothetical | G14 | Tb927.4.2800_for | GGGGACAAGTTTGTACAAAAAAGCAGGCTATGACCGTGAGCGCAAACG | Tb927.4.2800_rev | GGGGACCACTTTGTACAAGAAAGCTGGGTTTAATTCGATGTCTTCGTTCCTTTACTCTT |
| 7_hypothetical | G15 | Tb927.2.6100_for | GGGGACAAGTTTGTACAAAAAAGCAGGCTATGCTGCGGTCCTTGTTGTTAC | Tb927.2.6100_rev | GGGGACCACTTTGTACAAGAAAGCTGGGTTTAAAGTCGAAGAAGTCCGGTATTTGTT |
| 7_hypothetical | G16 | Tb927.10.6360_for | GGGGACAAGTTTGTACAAAAAAGCAGGCTATGAGCAGTGAGACTGTTGCG | Tb927.10.6360_rev | GGGGACCACTTTGTACAAGAAAGCTGGGTTTAAACCGGCTCCCCTCTGTG |
| 7_hypothetical | G17 | Tb927.4.3970_for | GGGGACAAGTTTGTACAAAAAAGCAGGCTATGTCGGCAGGTTCGGAACC | Tb927.4.3970_rev | GGGGACCACTTTGTACAAGAAAGCTGGGTTTAATTGCTAAAGAACTCAATGATATTTTG |
| 7_hypothetical | G18 | Tb927.11.1060_for | GGGGACAAGTTTGTACAAAAAAGCAGGCTATGTATTCTTCAGACAGGGTTCGGA | Tb927.11.1060_rev | GGGGACCACTTTGTACAAGAAAGCTGGGTCTACGATCGGCCCACATCGT |
| 7_hypothetical | G19 | Tb927.3.5480_for | GGGGACAAGTTTGTACAAAAAAGCAGGCTATGTTTAGTGCTAAGAGAAAGGATGTGG | Tb927.3.5480_rev | GGGGACCACTTTGTACAAGAAAGCTGGGTTTAAACAACTGGATCGGACCCGT |
| 7_hypothetical | G20 | Tb927.7.6690_for | GGGGACAAGTTTGTACAAAAAAGCAGGCTATGAGCCTCTCCAGAAGCCC | Tb927.7.6690_rev | GGGGACCACTTTGTACAAGAAAGCTGGGTTCATGCGGAGCGAAGTGTCC |
| 7_hypothetical | G21 | Tb927.9.10690_for | GGGGACAAGTTTGTACAAAAAAGCAGGCTATGAACCTCTTTGGGCTTCGTTC | Tb927.9.10690_rev | GGGGACCACTTTGTACAAGAAAGCTGGGTTCATAGACTGAGGTCACCGTCG |
| 7_hypothetical | G22 | Tb927.10.3670_for | GGGGACAAGTTTGTACAAAAAAGCAGGCTATGAATGAAGATGATATTCTCTTTGCTCAA | Tb927.10.3670_rev | GGGGACCACTTTGTACAAGAAAGCTGGGTTCAGAACGGAAGCTGCAAAGAC |
| 7_hypothetical | G23 | Tb927.10.3920_for | GGGGACAAGTTTGTACAAAAAAGCAGGCTATGGTCGGTGGGTTCACTGA | Tb927.10.3920_rev | GGGGACCACTTTGTACAAGAAAGCTGGGTCTACCTTCCGACTAACCACGTCA |
| 7_hypothetical | G24 | Tb927.8.2950_for | GGGGACAAGTTTGTACAAAAAAGCAGGCTATGTTGTCTCGTTCATCTCAGGATCC | Tb927.8.2950_rev | GGGGACCACTTTGTACAAGAAAGCTGGGTCTACATGTCCGTGAGATCCGTTTC |
| 7_hypothetical | H1 | Tb927.9.9950_for | GGGGACAAGTTTGTACAAAAAAGCAGGCTATGGAGGAAGTCTCCGGTTCC | Tb927.9.9950_rev | GGGGACCACTTTGTACAAGAAAGCTGGGTTTACACACAACCAAAGTCACTTTTGTTT |
| 7_hypothetical | H2 | Tb927.10.6720_for | GGGGACAAGTTTGTACAAAAAAGCAGGCTATGATGCAAGGATTTCCCTCCTTAGG | Tb927.10.6720_rev | GGGGACCACTTTGTACAAGAAAGCTGGGTTTACACACGTGGAGGTGCTGC |
| 7_hypothetical | H3 | Tb927.10.8880_for | GGGGACAAGTTTGTACAAAAAAGCAGGCTATGTACTCGGATAGCGAGAGCAT | Tb927.10.8880_rev | GGGGACCACTTTGTACAAGAAAGCTGGGTTTAAAGATACCTCTGCACCACGC |
| 7_hypothetical | H4 | Tb927.7.5240_for | GGGGACAAGTTTGTACAAAAAAGCAGGCTATGGCCAACTTTAATAAAGTCTCTAGGGC | Tb927.7.5240_rev | GGGGACCACTTTGTACAAGAAAGCTGGGTTTAGCGACTATTTCGCCCCTCC |
| 7_hypothetical | H5 | Tb927.10.6730_for | GGGGACAAGTTTGTACAAAAAAGCAGGCTATGAACAAGCAATACGGAACAGAAACA | Tb927.10.6730_rev | GGGGACCACTTTGTACAAGAAAGCTGGGTTTACACACGTGGAGGTGCTGC |
| 7_hypothetical | H6 | Tb927.11.10200_for | GGGGACAAGTTTGTACAAAAAAGCAGGCTATGAAGTCCAAGGATGTGGAGGAT | Tb927.11.10200_rev | GGGGACCACTTTGTACAAGAAAGCTGGGTCTAGAAAACATTTGTAGCGTAACCTGTG |
| 7_hypothetical | H7 | Tb927.10.6740_for | GGGGACAAGTTTGTACAAAAAAGCAGGCTATGAACAAGCAACACGGAACAGA | Tb927.10.6740_rev | GGGGACCACTTTGTACAAGAAAGCTGGGTTTACACACGTGGAGGTGCTGC |
| 7_hypothetical | H8 | Tb927.10.11640_for | GGGGACAAGTTTGTACAAAAAAGCAGGCTATGCATGACCAACGAAAGGTACG | Tb927.10.11640_rev | GGGGACCACTTTGTACAAGAAAGCTGGGTTTACGCTGTGGTAACCTCAGAAAGG |
| 7_hypothetical | H9 | Tb927.6.1740_for | GGGGACAAGTTTGTACAAAAAAGCAGGCTATGAAACGGTACTACTTACCTGCGG | Tb927.6.1740_rev | GGGGACCACTTTGTACAAGAAAGCTGGGTTCAGCCCAAGTGAGTTGCGC |
| 7_hypothetical | H10 | Tb927.3.3510_for | GGGGACAAGTTTGTACAAAAAAGCAGGCTATGCTGCCCTCAGCGCAA | Tb927.3.3510_rev | GGGGACCACTTTGTACAAGAAAGCTGGGTTCACATCAGCATGTAAACAAGAACTGC |
| 7_hypothetical | H11 | Tb927.6.2470_for | GGGGACAAGTTTGTACAAAAAAGCAGGCTATGTTTCGTCTAGACGGCTGCC | Tb927.6.2470_rev | GGGGACCACTTTGTACAAGAAAGCTGGGTCTAGCACGCTGCTCGCAT |
| 7_hypothetical | H12 | Tb927.4.730_for | GGGGACAAGTTTGTACAAAAAAGCAGGCTATGGCGACGGTGGCTAATGA | Tb927.4.730_rev | GGGGACCACTTTGTACAAGAAAGCTGGGTTCAGCGGGGATCGAAGGTG |
| 7_hypothetical | H13 | Tb927.6.1100_for | GGGGACAAGTTTGTACAAAAAAGCAGGCTATGTGCGCGCCTTCTTGTG | Tb927.6.1100_rev | GGGGACCACTTTGTACAAGAAAGCTGGGTCTACTCGTGCGCCTCTTCATAGT |
| 7_hypothetical | H14 | Tb927.3.4470_for | GGGGACAAGTTTGTACAAAAAAGCAGGCTATGTTGGCACTATCACAGGGGT | Tb927.3.4470_rev | GGGGACCACTTTGTACAAGAAAGCTGGGTCTACCGCTGCTCGCGATTT |
| 7_hypothetical | H15 | Tb927.5.2460_for | GGGGACAAGTTTGTACAAAAAAGCAGGCTATGCCACTCCTTCTCCGTTTGT | Tb927.5.2460_rev | GGGGACCACTTTGTACAAGAAAGCTGGGTTCACAATTCGTCAAGTTCCTGTAAAAGG |
| 7_hypothetical | H16 | Tb927.6.3180_for | GGGGACAAGTTTGTACAAAAAAGCAGGCTATGGTCTCCAACGACCCCC | Tb927.6.3180_rev | GGGGACCACTTTGTACAAGAAAGCTGGGTTCACAGGCACGAACGTCGC |
| 7_hypothetical | H17 | Tb927.11.12720_for | GGGGACAAGTTTGTACAAAAAAGCAGGCTATGTGTGGTGATTCCTTTGATACTTCTC | Tb927.11.12720_rev | GGGGACCACTTTGTACAAGAAAGCTGGGTTTACATCCGTGGGGAGGAGC |
| 7_hypothetical | H18 | Tb927.3.1020_for | GGGGACAAGTTTGTACAAAAAAGCAGGCTATGTTTTCGGGTTCTGAAGGCAG | Tb927.3.1020_rev | GGGGACCACTTTGTACAAGAAAGCTGGGTTCACTTAGTTTCGTATTGCTCGGTG |
| 7_hypothetical | H19 | Tb927.11.11000_for | GGGGACAAGTTTGTACAAAAAAGCAGGCTATGACTCTTGAACACAATGATGAGTTCG | Tb927.11.11000_rev | GGGGACCACTTTGTACAAGAAAGCTGGGTTCAGTTAGAGAAGTCACGCTCATGC |
| 7_hypothetical | H20 | Tb927.7.6620_for | GGGGACAAGTTTGTACAAAAAAGCAGGCTATGACGGAAGCGGCTCCA | Tb927.7.6620_rev | GGGGACCACTTTGTACAAGAAAGCTGGGTTTATTTCCCGCCCTCTCCATTCC |
| 7_hypothetical | H21 | Tb927.5.560_for | GGGGACAAGTTTGTACAAAAAAGCAGGCTATGAGGAAATATCATCTATCCCCATTGGT | Tb927.5.560_rev | GGGGACCACTTTGTACAAGAAAGCTGGGTTTACCATTTATTCCCCGCACCCG |
| 7_hypothetical | H22 | Tb927.7.950_for | GGGGACAAGTTTGTACAAAAAAGCAGGCTATGAATCCGAAGCTGTTCTCGAC | Tb927.7.950_rev | GGGGACCACTTTGTACAAGAAAGCTGGGTCTACCCCCTCAACTTCCCCAA |
| 7_hypothetical | H23 | Tb927.7.1060_for | GGGGACAAGTTTGTACAAAAAAGCAGGCTATGTTTCGCGTTGTGTCGTTTG | Tb927.7.1060_rev | GGGGACCACTTTGTACAAGAAAGCTGGGTCTACGATGAGTCTGGGTTGCTGT |
| 7_hypothetical | H24 | Tb927.9.12380_for | GGGGACAAGTTTGTACAAAAAAGCAGGCTATGTATGCTGATGAAGAGGAGAGTCG | Tb927.9.12380_rev | GGGGACCACTTTGTACAAGAAAGCTGGGTTTAACCCGCCAAAGCTACCGA |
| 7_hypothetical | I1 | Tb927.8.7970_for | GGGGACAAGTTTGTACAAAAAAGCAGGCTATGGAAGACGCAACGAAAGAAACA | Tb927.8.7970_rev | GGGGACCACTTTGTACAAGAAAGCTGGGTTCATTCAATCATCTCCCCGGATGC |
| 7_hypothetical | I2 | Tb927.9.7040_for | GGGGACAAGTTTGTACAAAAAAGCAGGCTATGCACCGTACCAGTTTACGC | Tb927.9.7040_rev | GGGGACCACTTTGTACAAGAAAGCTGGGTTTAGACACGCCGAGATCGGC |
| 7_hypothetical | I3 | Tb927.11.1850_for | GGGGACAAGTTTGTACAAAAAAGCAGGCTATGGGCGCTGCGCGGAGT | Tb927.11.1850_rev | GGGGACCACTTTGTACAAGAAAGCTGGGTCTAGCCTGATTTAGGTGGATCCTCA |
| 7_hypothetical | I4 | Tb927.10.1700_for | GGGGACAAGTTTGTACAAAAAAGCAGGCTATGTCGGTAGTGGCGTCCTT | Tb927.10.1700_rev | GGGGACCACTTTGTACAAGAAAGCTGGGTTTAATGCTGCGGGCTCTTTCC |
| 7_hypothetical | I5 | Tb927.9.2940_for | GGGGACAAGTTTGTACAAAAAAGCAGGCTATGATCGACAAATCGCCCCG | Tb927.9.2940_rev | GGGGACCACTTTGTACAAGAAAGCTGGGTTCAACGATGTTGGGGAATGAGGT |
| 7_hypothetical | I6 | Tb927.11.2150_for | GGGGACAAGTTTGTACAAAAAAGCAGGCTATGCAGGACCTCAACTGGGC | Tb927.11.2150_rev | GGGGACCACTTTGTACAAGAAAGCTGGGTTTACTGTGGTCTGGGTGTCGT |
| 7_hypothetical | I7 | Tb927.9.4660_for | GGGGACAAGTTTGTACAAAAAAGCAGGCTATGATGTCAAAAGCGTCGTATATGATGC | Tb927.9.4660_rev | GGGGACCACTTTGTACAAGAAAGCTGGGTCTAAGATAGTGGAGGCATACCCCAC |
| 7_hypothetical | I8 | Tb927.10.5640_for | GGGGACAAGTTTGTACAAAAAAGCAGGCTATGGAAGACGATGCTGATGCCT | Tb927.10.5640_rev | GGGGACCACTTTGTACAAGAAAGCTGGGTTTACAGCGGAACCAAACGATTACC |
| 7_hypothetical | I9 | Tb927.10.2820_for | GGGGACAAGTTTGTACAAAAAAGCAGGCTATGCCAGCCACCAGGGAA | Tb927.10.2820_rev | GGGGACCACTTTGTACAAGAAAGCTGGGTTTACAGCTTCAGGAGACGTGTTTC |
| 7_hypothetical | I10 | Tb927.5.3660_for | GGGGACAAGTTTGTACAAAAAAGCAGGCTATGAACGCAGCAGTTGTGACA | Tb927.5.3660_rev | GGGGACCACTTTGTACAAGAAAGCTGGGTTCAAAACAAATTTTTCAGTCGAAGGATGA |
| 7_hypothetical | I11 | Tb927.4.1460_for | GGGGACAAGTTTGTACAAAAAAGCAGGCTATGCTTCGGTGTCGAGTAGCG | Tb927.4.1460_rev | GGGGACCACTTTGTACAAGAAAGCTGGGTCTACGCTCTCCCGTCTCTTGT |
| 7_hypothetical | I12 | Tb927.11.7250_for | GGGGACAAGTTTGTACAAAAAAGCAGGCTATGCGCCGTTTCTACTGCATT | Tb927.11.7250_rev | GGGGACCACTTTGTACAAGAAAGCTGGGTTTATTTGAGCCCCTCCAGGACA |
| 7_hypothetical | I13 | Tb927.3.4540_for | GGGGACAAGTTTGTACAAAAAAGCAGGCTATGAGTGATGATGCGTCTCAAAACT | Tb927.3.4540_rev | GGGGACCACTTTGTACAAGAAAGCTGGGTTTATGTCAGTGGTTTCTTCCATACTTTACT |
| 7_hypothetical | I14 | Tb927.9.1510_for | GGGGACAAGTTTGTACAAAAAAGCAGGCTATGCCGGCCAAACCCACT | Tb927.9.1510_rev | GGGGACCACTTTGTACAAGAAAGCTGGGTTTAGTAGTTGTTTCCATTTTCCTCTTTTGT |
| 7_hypothetical | I15 | Tb927.7.3070_for | GGGGACAAGTTTGTACAAAAAAGCAGGCTATGCTAAGGACAAATGGTAAGATTCTCAA | Tb927.7.3070_rev | GGGGACCACTTTGTACAAGAAAGCTGGGTTCACTTACTACTTACAGGCTCCTTGC |
| 7_hypothetical | I16 | Tb927.10.3950_for | GGGGACAAGTTTGTACAAAAAAGCAGGCTATGGCACTTTCTTTTCCCTCTTCATC | Tb927.10.3950_rev | GGGGACCACTTTGTACAAGAAAGCTGGGTTTAAAAAGCCATGAAAAACTCGTGGTC |
| 7_hypothetical | I17 | Tb927.9.8240_for | GGGGACAAGTTTGTACAAAAAAGCAGGCTATGTCCGGGACGCCGACA | Tb927.9.8240_rev | GGGGACCACTTTGTACAAGAAAGCTGGGTCTACGTCCACACTAATAAATCACGCA |
| 7_hypothetical | I18 | Tb927.11.11310_for | GGGGACAAGTTTGTACAAAAAAGCAGGCTATGAACTCTGAAGAATGCACGAAGC | Tb927.11.11310_rev | GGGGACCACTTTGTACAAGAAAGCTGGGTTTAAAATCGTGTATGCGCATCGTG |
| 7_hypothetical | I19 | Tb927.6.1250_for | GGGGACAAGTTTGTACAAAAAAGCAGGCTATGCGCTCCAAGACCCTAAAGA | Tb927.6.1250_rev | GGGGACCACTTTGTACAAGAAAGCTGGGTCTACATTACCCCCACGGCACC |
| 7_hypothetical | I20 | Tb927.7.5340_for | GGGGACAAGTTTGTACAAAAAAGCAGGCTATGGGAGGAGGTTCATCCGTTG | Tb927.7.5340_rev | GGGGACCACTTTGTACAAGAAAGCTGGGTTTAGTTCAATTGGTCATCAAACATCATGAA |
| 7_hypothetical | I21 | Tb927.10.3610_for | GGGGACAAGTTTGTACAAAAAAGCAGGCTATGGCGGTTGGTGCAGGC | Tb927.10.3610_rev | GGGGACCACTTTGTACAAGAAAGCTGGGTTTAGTTCGTTTCGTTAAGGAGGCG |
| 7_hypothetical | I22 | Tb927.11.10840_for | GGGGACAAGTTTGTACAAAAAAGCAGGCTATGAATCCGCCGACTACTCAGG | Tb927.11.10840_rev | GGGGACCACTTTGTACAAGAAAGCTGGGTTCAATCTCTTCCTCTTTTCTTATTCTCTTC |
| 7_hypothetical | I23 | Tb927.3.4950_for | GGGGACAAGTTTGTACAAAAAAGCAGGCTATGCGGCGTGCATTGTACC | Tb927.3.4950_rev | GGGGACCACTTTGTACAAGAAAGCTGGGTTTATGGAGGTCCAACAGGGCG |
| 7_hypothetical | I24 | Tb927.7.550_for | GGGGACAAGTTTGTACAAAAAAGCAGGCTATGGGTGGCGGTGGAAGC | Tb927.7.550_rev | GGGGACCACTTTGTACAAGAAAGCTGGGTTTACATTCTGTCAATGTAATCATAATTCCA |
| 7_hypothetical | J1 | Tb927.10.6180_for | GGGGACAAGTTTGTACAAAAAAGCAGGCTATGAACCCGTTTGTTTTTTCCCTTTAC | Tb927.10.6180_rev | GGGGACCACTTTGTACAAGAAAGCTGGGTTCAGTTGTTCGTTCCGTATCCAC |
| 7_hypothetical | J2 | Tb927.6.4230_for | GGGGACAAGTTTGTACAAAAAAGCAGGCTATGAGCCGCTGTAACGTTGC | Tb927.6.4230_rev | GGGGACCACTTTGTACAAGAAAGCTGGGTTCACGACCGGTACCTTTCTAAGG |
| 7_hypothetical | J3 | Tb927.11.14740_for | GGGGACAAGTTTGTACAAAAAAGCAGGCTATGGCCCAACCCAGGAAGAA | Tb927.11.14740_rev | GGGGACCACTTTGTACAAGAAAGCTGGGTTCAGCTAATAACCTCAACAGAAGAACATG |
| 7_hypothetical | J4 | Tb927.2.5010_for | GGGGACAAGTTTGTACAAAAAAGCAGGCTATGGACGCCGTACACGAGG | Tb927.2.5010_rev | GGGGACCACTTTGTACAAGAAAGCTGGGTTTATACTACATGTGCGCTCTGGTCA |
| 7_hypothetical | J5 | Tb927.5.2350_for | GGGGACAAGTTTGTACAAAAAAGCAGGCTATGGAAAGAGGGGGAAAAATTGCG | Tb927.5.2350_rev | GGGGACCACTTTGTACAAGAAAGCTGGGTCTAATCTTCAGAAGCCATATGTTTGATGGA |
| 7_hypothetical | J6 | Tb927.7.4100_for | GGGGACAAGTTTGTACAAAAAAGCAGGCTATGGCAACTGAGGTGAAGCCA | Tb927.7.4100_rev | GGGGACCACTTTGTACAAGAAAGCTGGGTCTATTGCCTTCTCTTCATGTCCAAAAGA |
| 7_hypothetical | J7 | Tb927.11.15980_for | GGGGACAAGTTTGTACAAAAAAGCAGGCTATGTCCCAACGTCACGCCG | Tb927.11.15980_rev | GGGGACCACTTTGTACAAGAAAGCTGGGTTCACATCACCGAACAGCATTTAGAC |
| 7_hypothetical | J8 | Tb927.6.3880_for | GGGGACAAGTTTGTACAAAAAAGCAGGCTATGACAATACACGAGTCGTGGC | Tb927.6.3880_rev | GGGGACCACTTTGTACAAGAAAGCTGGGTTCATTTTGAACCATATATCTCTTTTAATTT |
| 7_hypothetical | J9 | Tb927.8.1800_for | GGGGACAAGTTTGTACAAAAAAGCAGGCTATGCAAGCGCGGCTGCTG | Tb927.8.1800_rev | GGGGACCACTTTGTACAAGAAAGCTGGGTCTAACGCCGTAACTTCCCCAC |
| 7_hypothetical | J10 | Tb927.4.4900_for | GGGGACAAGTTTGTACAAAAAAGCAGGCTATGGCGCAAGTCACCCAAAG | Tb927.4.4900_rev | GGGGACCACTTTGTACAAGAAAGCTGGGTTTATACCCCGTAGCAGAAACCGC |
| 7_hypothetical | J11 | Tb927.11.12290_for | GGGGACAAGTTTGTACAAAAAAGCAGGCTATGATTGAACTCCTCCTTGAGGTTGC | Tb927.11.12290_rev | GGGGACCACTTTGTACAAGAAAGCTGGGTTTAGCAACAGAGAGGCAGAGATGC |
| 7_hypothetical | J13 | Tb927.7.3910_for | GGGGACAAGTTTGTACAAAAAAGCAGGCTATGTTTTTAAAGAAGACAATGTCGGGTG | Tb927.7.3910_rev | GGGGACCACTTTGTACAAGAAAGCTGGGTTTACTTAAACCCCCTTTCCCTCCTGC |
| 7_hypothetical | J14 | Tb927.9.7480_for | GGGGACAAGTTTGTACAAAAAAGCAGGCTATGAACCAACCGACAACCACAC | Tb927.9.7480_rev | GGGGACCACTTTGTACAAGAAAGCTGGGTCTACTTGGCACGTAGGTGCTG |
| 7_hypothetical | J15 | Tb927.4.4520_for | GGGGACAAGTTTGTACAAAAAAGCAGGCTATGCTCTTCCATCCCACTGTATGC | Tb927.4.4520_rev | GGGGACCACTTTGTACAAGAAAGCTGGGTTCAATTGTCATCAAAGCTACCAAATGGC |
| 7_hypothetical | J16 | Tb927.8.7820_for | GGGGACAAGTTTGTACAAAAAAGCAGGCTATGCTTTTTCACACGACACCATGC | Tb927.8.7820_rev | GGGGACCACTTTGTACAAGAAAGCTGGGTTCATTTCCCATCAAATCCACCAAAGA |
| 7_hypothetical | J17 | Tb927.11.13540_for | GGGGACAAGTTTGTACAAAAAAGCAGGCTATGTTTTCGAGGACTGAAGTTGATGC | Tb927.11.13540_rev | GGGGACCACTTTGTACAAGAAAGCTGGGTCTACTGTGTGAAGATACTAAACACCTCC |
| 7_hypothetical | J18 | Tb927.10.6750_for | GGGGACAAGTTTGTACAAAAAAGCAGGCTATGACGCAAAAGGGGTACGTC | Tb927.10.6750_rev | GGGGACCACTTTGTACAAGAAAGCTGGGTTTACACACGTGGAGGTGCTGC |
| 7_hypothetical | J19 | Tb927.10.11400_for | GGGGACAAGTTTGTACAAAAAAGCAGGCTATGAGTGACAGTGACACTGATTCGT | Tb927.10.11400_rev | GGGGACCACTTTGTACAAGAAAGCTGGGTCTACAATCTCAGCTCATCTCCGTCG |
| 7_hypothetical | J20 | Tb927.6.1910_for | GGGGACAAGTTTGTACAAAAAAGCAGGCTATGACGAAGGAGGAGGTTTTATCCT | Tb927.6.1910_rev | GGGGACCACTTTGTACAAGAAAGCTGGGTTTAGGGCGTCTTTTCCAAACGC |
| 7_hypothetical | J21 | Tb927.10.2280_for | GGGGACAAGTTTGTACAAAAAAGCAGGCTATGGAATCGCTTATAAAAAACATACAGAGT | Tb927.10.2280_rev | GGGGACCACTTTGTACAAGAAAGCTGGGTTCATATCCCTTTACTACCCCGTTTACC |
| 7_hypothetical | J22 | Tb927.11.4180_for | GGGGACAAGTTTGTACAAAAAAGCAGGCTATGATGAGCGGGAGTAGGGAATC | Tb927.11.4180_rev | GGGGACCACTTTGTACAAGAAAGCTGGGTTCAGCCAGTGTGTGCATCATTG |
| 7_hypothetical | J23 | Tb927.9.13320_for | GGGGACAAGTTTGTACAAAAAAGCAGGCTATGAGTGCCCCCCAACCC | Tb927.9.13320_rev | GGGGACCACTTTGTACAAGAAAGCTGGGTTCAACCGTCCAATGAAACAACAGG |
| 7_hypothetical | J24 | Tb927.7.4120_for | GGGGACAAGTTTGTACAAAAAAGCAGGCTATGGATCTGGATTCTTTCCTCGACA | Tb927.7.4120_rev | GGGGACCACTTTGTACAAGAAAGCTGGGTTTAGCTTGGTGGTCTAAAGGTGCT |
| 7_hypothetical | K1 | Tb927.11.14980_for | GGGGACAAGTTTGTACAAAAAAGCAGGCTATGGTAATGCGTCGATGTAGTTACAAC | Tb927.11.14980_rev | GGGGACCACTTTGTACAAGAAAGCTGGGTCTACCGTCTGTACGTTATGGGAACA |
| 7_hypothetical | K2 | Tb927.9.6270_for | GGGGACAAGTTTGTACAAAAAAGCAGGCTATGTTTTGTTGCACCTTCACTCAAGC | Tb927.9.6270_rev | GGGGACCACTTTGTACAAGAAAGCTGGGTTCATATTTGGTGCTTCTCACCGG |
| 7_hypothetical | K3 | Tb927.10.3720_for | GGGGACAAGTTTGTACAAAAAAGCAGGCTATGCGAGGGGATGTGCCA | Tb927.10.3720_rev | GGGGACCACTTTGTACAAGAAAGCTGGGTTCACCATCTATTCCGCTTACGGG |
| 7_hypothetical | K4 | Tb927.4.1540_for | GGGGACAAGTTTGTACAAAAAAGCAGGCTATGCTTCGTGTATTCTCCGGC | Tb927.4.1540_rev | GGGGACCACTTTGTACAAGAAAGCTGGGTTCAATTCTTTCCCCTCCTTCGCA |
| 7_hypothetical | K5 | Tb927.4.1120_for | GGGGACAAGTTTGTACAAAAAAGCAGGCTATGGAGAGGTGTTGCACGCT | Tb927.4.1120_rev | GGGGACCACTTTGTACAAGAAAGCTGGGTTCAATTGTAAACCTTTTGTGCGCG |
| 7_hypothetical | K6 | Tb927.10.7830_for | GGGGACAAGTTTGTACAAAAAAGCAGGCTATGCAGCTTTGTTATAGTTGTCGACA | Tb927.10.7830_rev | GGGGACCACTTTGTACAAGAAAGCTGGGTTCAAACCTCCTTTTCTGTAATGAAGATTGA |
| 7_hypothetical | K7 | Tb927.4.350_for | GGGGACAAGTTTGTACAAAAAAGCAGGCTATGAAAAGCGCAAATGTACCCACA | Tb927.4.350_rev | GGGGACCACTTTGTACAAGAAAGCTGGGTTTAATAACCTTTACGTCTTCGTTTACCCA |
| 7_hypothetical | K8 | Tb927.10.6700_for | GGGGACAAGTTTGTACAAAAAAGCAGGCTATGGCTGGTGCGGAACAAC | Tb927.10.6700_rev | GGGGACCACTTTGTACAAGAAAGCTGGGTCTACCGGTGATACAACGTCGC |
| 7_hypothetical | K9 | Tb927.5.1860_for | GGGGACAAGTTTGTACAAAAAAGCAGGCTATGCAATTCTCGACCTACATGACG | Tb927.5.1860_rev | GGGGACCACTTTGTACAAGAAAGCTGGGTCTATTCAGCTTCACAGAAAAGAGTACAAAC |
| 7_hypothetical | K10 | Tb927.8.7550_for | GGGGACAAGTTTGTACAAAAAAGCAGGCTATGCCCCAAACGGTTCAGAAG | Tb927.8.7550_rev | GGGGACCACTTTGTACAAGAAAGCTGGGTCTACAGCACAATCCACCGTCG |
| 7_hypothetical | K11 | Tb927.11.7660_for | GGGGACAAGTTTGTACAAAAAAGCAGGCTATGTTTGATGAGGACGCGACG | Tb927.11.7660_rev | GGGGACCACTTTGTACAAGAAAGCTGGGTCTATTTCCCCCCTTTACGCGATTT |
| 7_hypothetical | K12 | Tb927.8.6670_for | GGGGACAAGTTTGTACAAAAAAGCAGGCTATGGCTGCCGCCCCGGCG | Tb927.8.6670_rev | GGGGACCACTTTGTACAAGAAAGCTGGGTTCATTGTCCCATAAAGAGAGACATCATTG |
| 7_hypothetical | K13 | Tb927.4.530_for | GGGGACAAGTTTGTACAAAAAAGCAGGCTATGTCTCCTCTAGATTTACTGCAGCA | Tb927.4.530_rev | GGGGACCACTTTGTACAAGAAAGCTGGGTTCAGCGGTTGGTATCAGCACT |
| 7_hypothetical | K14 | Tb927.11.5240_for | GGGGACAAGTTTGTACAAAAAAGCAGGCTATGCAAGGGACCGCATGTG | Tb927.11.5240_rev | GGGGACCACTTTGTACAAGAAAGCTGGGTTTATTGAAAGGCAGGTAATGAGGGGT |
| 7_hypothetical | K15 | Tb927.8.3000_for | GGGGACAAGTTTGTACAAAAAAGCAGGCTATGTGCGCAATGGAATATAGCGA | Tb927.8.3000_rev | GGGGACCACTTTGTACAAGAAAGCTGGGTCTAACCCGACACATATTGTGGAGC |
| 7_hypothetical | K16 | Tb927.3.1220_for | GGGGACAAGTTTGTACAAAAAAGCAGGCTATGGACACCCGGTTTCACTGT | Tb927.3.1220_rev | GGGGACCACTTTGTACAAGAAAGCTGGGTTCAGTGGTTAAGCCACTCGTCG |
| 7_hypothetical | K17 | Tb927.9.12040_for | GGGGACAAGTTTGTACAAAAAAGCAGGCTATGTATCCTAGTCTTTTTAAGAAGCGACC | Tb927.9.12040_rev | GGGGACCACTTTGTACAAGAAAGCTGGGTTTACTGTTTCATTTTTGTTGTTACCCAAAC |
| 7_hypothetical | K18 | Tb927.4.4110_for | GGGGACAAGTTTGTACAAAAAAGCAGGCTATGGCAAACACGGCTCTGC | Tb927.4.4110_rev | GGGGACCACTTTGTACAAGAAAGCTGGGTTCACGTAGGGGGTTCCACATC |

| | | | | | |
|---|---|---|---|---|---|
| 7_hypothetical | K19 | Tb927.8.2130_for | GGGGACAAGTTTGTACAAAAAAGCAGGCTATGTGGACTGTTTTTGCTCCAGATC | Tb927.8.2130_rev | GGGGACCACTTTGTACAAGAAAGCTGGGTTCACTTTTGGACGGGTTGCAG |
| 7_hypothetical | K20 | Tb927.4.2340_for | GGGGACAAGTTTGTACAAAAAAGCAGGCTATGGACTTGATCTTCCCCGAAGA | Tb927.4.2340_rev | GGGGACCACTTTGTACAAGAAAGCTGGGTCTACAGAAGTCCAACCTCTTGAGAAGC |
| 7_hypothetical | K21 | Tb927.11.7300_for | GGGGACAAGTTTGTACAAAAAAGCAGGCTATGGAAAGACAATCTGATGAAGATATGCTC | Tb927.11.7300_rev | GGGGACCACTTTGTACAAGAAAGCTGGGTCTAGGGCTCACGATGCAAATATACC |
| 7_hypothetical | K22 | Tb927.11.15710_for | GGGGACAAGTTTGTACAAAAAAGCAGGCTATGTCCTTCAGGATTCCCAATAATTTACTG | Tb927.11.15710_rev | GGGGACCACTTTGTACAAGAAAGCTGGGTTCACGCACAGATAAACTGCCAC |
| 7_hypothetical | K23 | Tb927.7.4310_for | GGGGACAAGTTTGTACAAAAAAGCAGGCTATGCTCTGGAAGTTATTCACAAGTAGC | Tb927.7.4310_rev | GGGGACCACTTTGTACAAGAAAGCTGGGTTTAACAGTGAGAAAAGTCAACAATATCGGT |
| 7_hypothetical | K24 | Tb927.6.1590_for | GGGGACAAGTTTGTACAAAAAAGCAGGCTATGCTGCGTTTATCGTTGATTCGC | Tb927.6.1590_rev | GGGGACCACTTTGTACAAGAAAGCTGGGTTTATTGAGGTGGCGCCCTAAGT |
| 7_hypothetical | L1 | Tb927.3.5390_for | GGGGACAAGTTTGTACAAAAAAGCAGGCTATGACTTCTCCAAAACGGCCC | Tb927.3.5390_rev | GGGGACCACTTTGTACAAGAAAGCTGGGTTTACATGGGAGAACAGGAGCTGT |
| 7_hypothetical | L2 | Tb927.11.3710_for | GGGGACAAGTTTGTACAAAAAAGCAGGCTATGGTAGGGCAAACCCAATCGA | Tb927.11.3710_rev | GGGGACCACTTTGTACAAGAAAGCTGGGTTTACACGTGCAACTCCATTGCT |
| 7_hypothetical | L3 | Tb927.5.1500_for | GGGGACAAGTTTGTACAAAAAAGCAGGCTATGTCAGACGCCGGCTGC | Tb927.5.1500_rev | GGGGACCACTTTGTACAAGAAAGCTGGGTCTACGCACTGGTTGGCGAC |
| 7_hypothetical | L4 | Tb927.2.5080_for | GGGGACAAGTTTGTACAAAAAAGCAGGCTATGAATTTGAAAGAGTTGTTAACTTTTTTA | Tb927.2.5080_rev | GGGGACCACTTTGTACAAGAAAGCTGGGTCTACAATTTTAACTTAGAAATGAGCGAAAC |
| 7_hypothetical | L5 | Tb927.7.5060_for | GGGGACAAGTTTGTACAAAAAAGCAGGCTATGACCGGTGGGGAGTTGG | Tb927.7.5060_rev | GGGGACCACTTTGTACAAGAAAGCTGGGTCTAACTGCAACCACCACCGC |
| 7_hypothetical | L6 | Tb927.9.9270_for | GGGGACAAGTTTGTACAAAAAAGCAGGCTATGACCTCACCCATCGCAGT | Tb927.9.9270_rev | GGGGACCACTTTGTACAAGAAAGCTGGGTTTAATCAGTCTGCAAAACGTAATCAACAG |
| 7_hypothetical | L7 | Tb927.4.540_for | GGGGACAAGTTTGTACAAAAAAGCAGGCTATGTCTTTTCGTCACTTCAGCCG | Tb927.4.540_rev | GGGGACCACTTTGTACAAGAAAGCTGGGTTTAATCATCGTCCGGATCCACAAAAT |
| 7_hypothetical | L8 | Tb927.10.4020_for | GGGGACAAGTTTGTACAAAAAAGCAGGCTATGCCCTATTTTTTTAAACCACTGGGA | Tb927.10.4020_rev | GGGGACCACTTTGTACAAGAAAGCTGGGTTCACCATAGACGCCACCATATCG |
| 7_hypothetical | L9 | Tb927.7.5350_for | GGGGACAAGTTTGTACAAAAAAGCAGGCTATGTTGCGCGAAAGGAGTTGC | Tb927.7.5350_rev | GGGGACCACTTTGTACAAGAAAGCTGGGTTTACAGAAAGCATTTAAGTTCAAGGGAGC |
| 7_hypothetical | L10 | Tb927.10.1840_for | GGGGACAAGTTTGTACAAAAAAGCAGGCTATGGAGGACACGGCGGCT | Tb927.10.1840_rev | GGGGACCACTTTGTACAAGAAAGCTGGGTTTAAGTCATCATCGCAATATCTGTCAAGA |
| 7_hypothetical | L11 | Tb927.10.3970_for | GGGGACAAGTTTGTACAAAAAAGCAGGCTATGGTAGGTGCTTCTTCCAAGCG | Tb927.10.3970_rev | GGGGACCACTTTGTACAAGAAAGCTGGGTCTAAAATTCCTCATCAAAACTAAGTCCTTT |
| 7_hypothetical | L12 | Tb927.11.14220_for | GGGGACAAGTTTGTACAAAAAAGCAGGCTATGGACAGCATCACTGCAAACG | Tb927.11.14220_rev | GGGGACCACTTTGTACAAGAAAGCTGGGTCTATTGTTGCGGCTGCTGCA |
| 7_hypothetical | L13 | Tb927.8.6800_for | GGGGACAAGTTTGTACAAAAAAGCAGGCTATGTCGTGGCGACTATTCCTTGC | Tb927.8.6800_rev | GGGGACCACTTTGTACAAGAAAGCTGGGTTCAATCAGATATCTCCTCTGCCATCC |
| 7_hypothetical | L14 | Tb927.3.5380_for | GGGGACAAGTTTGTACAAAAAAGCAGGCTATGATAACGGACGTTACGAGCAC | Tb927.3.5380_rev | GGGGACCACTTTGTACAAGAAAGCTGGGTTTACCTCAACCCGTTTGCATGAC |
| 7_hypothetical | L15 | Tb927.10.15740_for | GGGGACAAGTTTGTACAAAAAAGCAGGCTATGTCTGGAGAAAACTTGAATAAATCGATG | Tb927.10.15740_rev | GGGGACCACTTTGTACAAGAAAGCTGGGTTCACGCCGTGGTCCATGC |
| 7_hypothetical | L16 | Tb927.9.13060_for | GGGGACAAGTTTGTACAAAAAAGCAGGCTATGCCTCCGAGAAGTGCTGG | Tb927.9.13060_rev | GGGGACCACTTTGTACAAGAAAGCTGGGTTCATTCGCCTTCTTGCGGTG |
| 7_hypothetical | L18 | Tb927.11.660_for | GGGGACAAGTTTGTACAAAAAAGCAGGCTATGAGCTTACAATGCGAAGGTGATG | Tb927.11.660_rev | GGGGACCACTTTGTACAAGAAAGCTGGGTCTACATGAGAGCACAGCTCTTCATC |
| 7_hypothetical | L19 | Tb927.3.3350_for | GGGGACAAGTTTGTACAAAAAAGCAGGCTATGGGAGATTGTTTTGACGAAGGC | Tb927.3.3350_rev | GGGGACCACTTTGTACAAGAAAGCTGGGTCTAGCGGCAGGCAGAACCA |
| 7_hypothetical | L20 | Tb927.8.5720_for | GGGGACAAGTTTGTACAAAAAAGCAGGCTATGGAAGAGGCAGCAACCATTG | Tb927.8.5720_rev | GGGGACCACTTTGTACAAGAAAGCTGGGTTCAGCCCTCGCGCGCTCG |
| 7_hypothetical | L21 | Tb927.10.3750_for | GGGGACAAGTTTGTACAAAAAAGCAGGCTATGTCCAGTACCGGTAAGTTTGTGG | Tb927.10.3750_rev | GGGGACCACTTTGTACAAGAAAGCTGGGTTTAAAGATTTCTGTTGATGATGGACCGC |
| 7_hypothetical | L22 | Tb927.11.15660_for | GGGGACAAGTTTGTACAAAAAAGCAGGCTATGATTCATGCGGTTGTTATGGCG | Tb927.11.15660_rev | GGGGACCACTTTGTACAAGAAAGCTGGGTCTAGGTCTCCGGCCTTTCAACC |
| 7_hypothetical | L23 | Tb927.11.13460_for | GGGGACAAGTTTGTACAAAAAAGCAGGCTATGCCGCCCAAGCGTTCC | Tb927.11.13460_rev | GGGGACCACTTTGTACAAGAAAGCTGGGTCTATTTCAATTTTGAAGCTGTGCGTTTG |
| 7_hypothetical | L24 | Tb927.11.560_for | GGGGACAAGTTTGTACAAAAAAGCAGGCTATGCCACCAAAGAAAGATGTGGC | Tb927.11.560_rev | GGGGACCACTTTGTACAAGAAAGCTGGGTCTAGTCAGATGACATGATCGAGCG |
| 7_hypothetical | M1 | Tb927.9.14130_for | GGGGACAAGTTTGTACAAAAAAGCAGGCTATGCAATGCAATATGAAAACTTGGATAATT | Tb927.9.14130_rev | GGGGACCACTTTGTACAAGAAAGCTGGGTTTAACTTGGTTCAACGTATGACCGC |
| 7_hypothetical | M2 | Tb927.5.780_for | GGGGACAAGTTTGTACAAAAAAGCAGGCTATGTACGTTTGTGAGAACGTTGATGC | Tb927.5.780_rev | GGGGACCACTTTGTACAAGAAAGCTGGGTTTAGGAGCGTCGCCTGACTT |
| 7_hypothetical | M3 | Tb927.10.9360_for | GGGGACAAGTTTGTACAAAAAAGCAGGCTATGGCCAAAAGTTTCCTCAAGTGG | Tb927.10.9360_rev | GGGGACCACTTTGTACAAGAAAGCTGGGTCTACCAGCGGAATTGTTCAGCA |
| 7_hypothetical | M4 | Tb927.10.5530_for | GGGGACAAGTTTGTACAAAAAAGCAGGCTATGAAAGCCCGGCGACGT | Tb927.10.5530_rev | GGGGACCACTTTGTACAAGAAAGCTGGGTCTACGTTGCCAGCAGGCG |
| 7_hypothetical | M5 | Tb927.5.2130_for | GGGGACAAGTTTGTACAAAAAAGCAGGCTATGTCGAGTCTGGTGCATCACT | Tb927.5.2130_rev | GGGGACCACTTTGTACAAGAAAGCTGGGTTCACACATCCCCTACGGCAT |
| 7_hypothetical | M6 | Tb927.8.4920_for | GGGGACAAGTTTGTACAAAAAAGCAGGCTATGTTCCAACTGACCTTGCGC | Tb927.8.4920_rev | GGGGACCACTTTGTACAAGAAAGCTGGGTTTAAGTGTGCGAGCACCGGG |
| 7_hypothetical | M7 | Tb927.4.4030_for | GGGGACAAGTTTGTACAAAAAAGCAGGCTATGGATGCCACAACTCAGGAAGG | Tb927.4.4030_rev | GGGGACCACTTTGTACAAGAAAGCTGGGTTTACCGAAAGGCGACAATACAAATGT |
| 7_hypothetical | M8 | Tb927.10.15040_for | GGGGACAAGTTTGTACAAAAAAGCAGGCTATGGGGGGAGGCGCTTGC | Tb927.10.15040_rev | GGGGACCACTTTGTACAAGAAAGCTGGGTCTACATCTCTTCCGCGGCG |
| 7_hypothetical | M9 | Tb927.10.13440_for | GGGGACAAGTTTGTACAAAAAAGCAGGCTATGGGTGTAGAAATCCCACTACCC | Tb927.10.13440_rev | GGGGACCACTTTGTACAAGAAAGCTGGGTTCATTTGTGGGTTTGCAAGCGC |
| 7_hypothetical | M10 | Tb927.6.2010_for | GGGGACAAGTTTGTACAAAAAAGCAGGCTATGCTCCGTCGCCTCGCC | Tb927.6.2010_rev | GGGGACCACTTTGTACAAGAAAGCTGGGTCTAGTACAATTGGGTGCCATGCA |
| 7_hypothetical | M11 | Tb927.11.8050_for | GGGGACAAGTTTGTACAAAAAAGCAGGCTATGATGAAGGGCAAACGCCTG | Tb927.11.8050_rev | GGGGACCACTTTGTACAAGAAAGCTGGGTCTACTGAAACCGAACGCTTTGCG |
| 7_hypothetical | M12 | Tb927.11.7670_for | GGGGACAAGTTTGTACAAAAAAGCAGGCTATGCCACCAGCGTACCGA | Tb927.11.7670_rev | GGGGACCACTTTGTACAAGAAAGCTGGGTTCAATTGTCAAGGTAGGACAAGAGC |
| 7_hypothetical | M13 | Tb927.2.5930_for | GGGGACAAGTTTGTACAAAAAAGCAGGCTATGCGGCGCTTTACACTTATAACG | Tb927.2.5930_rev | GGGGACCACTTTGTACAAGAAAGCTGGGTTCATTCCTGATCCGAGGAGTGATG |
| 7_hypothetical | M14 | Tb927.3.990_for | GGGGACAAGTTTGTACAAAAAAGCAGGCTATGGATCGATCACCTTCGCAAC | Tb927.3.990_rev | GGGGACCACTTTGTACAAGAAAGCTGGGTTCAAAGTTGACGCGCCGC |
| 7_hypothetical | M15 | Tb927.11.7110_for | GGGGACAAGTTTGTACAAAAAAGCAGGCTATGAGTGTGAAAGAGCAGCTTCG | Tb927.11.7110_rev | GGGGACCACTTTGTACAAGAAAGCTGGGTTTACAAGACAGCACTTGATATTGTCGA |
| 7_hypothetical | M16 | Tb927.11.1550_for | GGGGACAAGTTTGTACAAAAAAGCAGGCTATGGGCACCAAGGGGCTAC | Tb927.11.1550_rev | GGGGACCACTTTGTACAAGAAAGCTGGGTTCATTTTACACCTCCATCGCAGAC |
| 7_hypothetical | M17 | Tb927.9.12500_for | GGGGACAAGTTTGTACAAAAAAGCAGGCTATGCAAGCCGCTTGTACCAG | Tb927.9.12500_rev | GGGGACCACTTTGTACAAGAAAGCTGGGTTCATGCTGTTGGAACCTCCAAGT |
| 7_hypothetical | M18 | Tb927.8.5990_for | GGGGACAAGTTTGTACAAAAAAGCAGGCTATGCCAGTCCGAAAGAAGCAAC | Tb927.8.5990_rev | GGGGACCACTTTGTACAAGAAAGCTGGGTTTAGTGGACCCAAATTAGAACTTTCTTGT |
| 7_hypothetical | M19 | Tb927.9.14050_for | GGGGACAAGTTTGTACAAAAAAGCAGGCTATGAGTGTGCCATCCGCAATC | Tb927.9.14050_rev | GGGGACCACTTTGTACAAGAAAGCTGGGTTTACACTGGGCTTCTTAACTCGTCC |
| 7_hypothetical | M20 | Tb927.10.7270_for | GGGGACAAGTTTGTACAAAAAAGCAGGCTATGTTTTTCGTCTTTCCGTTTCCTACG | Tb927.10.7270_rev | GGGGACCACTTTGTACAAGAAAGCTGGGTCTATAGTGCGTCAGGAGAATCGATG |
| 7_hypothetical | M21 | Tb927.10.2510_for | GGGGACAAGTTTGTACAAAAAAGCAGGCTATGGACCTACCCATCGAACTGC | Tb927.10.2510_rev | GGGGACCACTTTGTACAAGAAAGCTGGGTTTATGGTCCAAACTCCACCCCA |
| 7_hypothetical | M22 | Tb927.11.13080_for | GGGGACAAGTTTGTACAAAAAAGCAGGCTATGGAAGGTTTTTGCTACACTGCT | Tb927.11.13080_rev | GGGGACCACTTTGTACAAGAAAGCTGGGTTCAGAACAGTAGGGCGGCTAG |
| 7_hypothetical | M23 | Tb927.11.12630_for | GGGGACAAGTTTGTACAAAAAAGCAGGCTATGACCGTTCCGGTTAGGCA | Tb927.11.12630_rev | GGGGACCACTTTGTACAAGAAAGCTGGGTTTATAACGCGGCACGCTTGC |
| 7_hypothetical | M24 | Tb927.8.1770_for | GGGGACAAGTTTGTACAAAAAAGCAGGCTATGCATAGTGTAGAGGTGATGGGG | Tb927.8.1770_rev | GGGGACCACTTTGTACAAGAAAGCTGGGTTCAGGACTGCTGTGTTGCGT |
| 7_hypothetical | N1 | Tb927.11.14410_for | GGGGACAAGTTTGTACAAAAAAGCAGGCTATGAACGGCGTAGAAATAAACGAAAGG | Tb927.11.14410_rev | GGGGACCACTTTGTACAAGAAAGCTGGGTCTACGATTTCTTTGAAATGTCAAATCTGCG |
| 7_hypothetical | N2 | Tb927.10.12470_for | GGGGACAAGTTTGTACAAAAAAGCAGGCTATGAGTTTACCGGAGCTTGATTCG | Tb927.10.12470_rev | GGGGACCACTTTGTACAAGAAAGCTGGGTTTACGGTTTCGTGTGCATGTTGA |
| 7_hypothetical | N3 | Tb927.4.4890_for | GGGGACAAGTTTGTACAAAAAAGCAGGCTATGGGCCACCAACTGTGTGA | Tb927.4.4890_rev | GGGGACCACTTTGTACAAGAAAGCTGGGTCTAACGAAAGACAAACTTGTCAGGTCT |
| 7_hypothetical | N4 | Tb927.4.4720_for | GGGGACAAGTTTGTACAAAAAAGCAGGCTATGGTCCTCGCAGAAGTCTGT | Tb927.4.4720_rev | GGGGACCACTTTGTACAAGAAAGCTGGGTTCACCGACCTGTGGCGGC |
| 7_hypothetical | N5 | Tb927.5.3450_for | GGGGACAAGTTTGTACAAAAAAGCAGGCTATGGAGTCGAATGTGTCCGATACT | Tb927.5.3450_rev | GGGGACCACTTTGTACAAGAAAGCTGGGTCTATCTCCGTTTCTTCTTCTGCACTAC |
| 7_hypothetical | N6 | Tb927.7.6240_for | GGGGACAAGTTTGTACAAAAAAGCAGGCTATGCATCCAACTTCGTTCCCG | Tb927.7.6240_rev | GGGGACCACTTTGTACAAGAAAGCTGGGTTCACTCCTCTTGCTCATCTATGATCC |
| 7_hypothetical | N7 | Tb927.7.4500_for | GGGGACAAGTTTGTACAAAAAAGCAGGCTATGATGCCTTCAGGTGACGGC | Tb927.7.4500_rev | GGGGACCACTTTGTACAAGAAAGCTGGGTTCACTCATCAGCGGTGTATGCC |
| 7_hypothetical | N8 | Tb927.11.2340_for | GGGGACAAGTTTGTACAAAAAAGCAGGCTATGAGCCGTCCGTATACCAAAGG | Tb927.11.2340_rev | GGGGACCACTTTGTACAAGAAAGCTGGGTTTAACCTCCAAGTAAGTTTGCCATGAC |
| 7_hypothetical | N9 | Tb927.11.5150_for | GGGGACAAGTTTGTACAAAAAAGCAGGCTATGGATAAGGCATTGTGTACCTCTCT | Tb927.11.5150_rev | GGGGACCACTTTGTACAAGAAAGCTGGGTTTATGTCACTGTTAATGGACAGGAAGAC |
| 7_hypothetical | N10 | Tb927.5.3420_for | GGGGACAAGTTTGTACAAAAAAGCAGGCTATGGCTTGGCGCACAATTCG | Tb927.5.3420_rev | GGGGACCACTTTGTACAAGAAAGCTGGGTCTAACCGCTAAGTTGCAGCTGC |
| 7_hypothetical | N11 | Tb927.2.6080_for | GGGGACAAGTTTGTACAAAAAAGCAGGCTATGCTAACGTTTCCCGGATTTGT | Tb927.2.6080_rev | GGGGACCACTTTGTACAAGAAAGCTGGGTCTACGCATATTCATCATTGGCTTCAGC |
| 7_hypothetical | N12 | Tb927.11.10500_for | GGGGACAAGTTTGTACAAAAAAGCAGGCTATGACTAGCAGGAAGCGGCC | Tb927.11.10500_rev | GGGGACCACTTTGTACAAGAAAGCTGGGTTTACAGGAGCTGCACTCGGC |
| 7_hypothetical | N13 | Tb927.9.10500_for | GGGGACAAGTTTGTACAAAAAAGCAGGCTATGTTCTCCGCCTCTTGGGG | Tb927.9.10500_rev | GGGGACCACTTTGTACAAGAAAGCTGGGTTTAACTGAAGCCATAATCTTCATGGAGATC |
| 7_hypothetical | N14 | Tb927.6.2930_for | GGGGACAAGTTTGTACAAAAAAGCAGGCTATGGACACTTACGCAGCGTTG | Tb927.6.2930_rev | GGGGACCACTTTGTACAAGAAAGCTGGGTCTAGAAGACAGATCGCCTACTTCCT |
| 7_hypothetical | N15 | Tb927.5.2150_for | GGGGACAAGTTTGTACAAAAAAGCAGGCTATGCTGGAGAGGCGTGCG | Tb927.5.2150_rev | GGGGACCACTTTGTACAAGAAAGCTGGGTCTACCTGGCCGCGAGGCG |
| 7_hypothetical | N16 | Tb927.11.4420_for | GGGGACAAGTTTGTACAAAAAAGCAGGCTATGGATCGTCCAACTAAGGAAGCA | Tb927.11.4420_rev | GGGGACCACTTTGTACAAGAAAGCTGGGTTCACCGAAAGTGGATCTCCTTTTTC |
| 7_hypothetical | N17 | Tb927.10.8730_for | GGGGACAAGTTTGTACAAAAAAGCAGGCTATGCTTCGCCGCATTTGGA | Tb927.10.8730_rev | GGGGACCACTTTGTACAAGAAAGCTGGGTCTACCATGCATTCGCATTTAAGTGAAC |
| 7_hypothetical | N18 | Tb927.8.7290_for | GGGGACAAGTTTGTACAAAAAAGCAGGCTATGCTCCGTCGGTTGCAC | Tb927.8.7290_rev | GGGGACCACTTTGTACAAGAAAGCTGGGTCTAACGCATGCTCCGCAGAG |
| 7_hypothetical | N19 | Tb927.11.6600_for | GGGGACAAGTTTGTACAAAAAAGCAGGCTATGGATCCCAGAAACCCAGCC | Tb927.11.6600_rev | GGGGACCACTTTGTACAAGAAAGCTGGGTTCAAGCACTTATGCCACACCTCC |
| 7_hypothetical | N20 | Tb927.6.4830_for | GGGGACAAGTTTGTACAAAAAAGCAGGCTATGCCGTACCCATTGGGGT | Tb927.6.4830_rev | GGGGACCACTTTGTACAAGAAAGCTGGGTCTACTCAAGCGTTTCGATGTTGTCA |
| 7_hypothetical | N21 | Tb927.11.3440_for | GGGGACAAGTTTGTACAAAAAAGCAGGCTATGGTGTATGAGTTTGTGCTTGAGC | Tb927.11.3440_rev | GGGGACCACTTTGTACAAGAAAGCTGGGTCTAAGACGGATGCATGTCCTGTC |
| 7_hypothetical | N22 | Tb927.2.5950_for | GGGGACAAGTTTGTACAAAAAAGCAGGCTATGAATCGCGACAAGGGTAAGTTC | Tb927.2.5950_rev | GGGGACCACTTTGTACAAGAAAGCTGGGTTCATTTGAACGTTTTGCGGTCG |
| 7_hypothetical | N23 | Tb927.1.4380_for | GGGGACAAGTTTGTACAAAAAAGCAGGCTATGTTGAGTCTGCTGCTTCTACTGC | Tb927.1.4380_rev | GGGGACCACTTTGTACAAGAAAGCTGGGTTCACCTATGTGGTCGACGTTCG |
| 7_hypothetical | N24 | Tb927.6.3350_for | GGGGACAAGTTTGTACAAAAAAGCAGGCTATGCTCTTGGGATGCCGTCG | Tb927.6.3350_rev | GGGGACCACTTTGTACAAGAAAGCTGGGTTTAATTAAAGTTATTCAGTTCGTCCAACAA |
| 7_hypothetical | O1 | Tb927.9.14120_for | GGGGACAAGTTTGTACAAAAAAGCAGGCTATGGAGCATTTTGCCCCATGTG | Tb927.9.14120_rev | GGGGACCACTTTGTACAAGAAAGCTGGGTCTACCGGTTCCTATGCTTACGTCG |
| 7_hypothetical | O2 | Tb927.3.2680_for | GGGGACAAGTTTGTACAAAAAAGCAGGCTATGGCGAGTCCGGAAGTCA | Tb927.3.2680_rev | GGGGACCACTTTGTACAAGAAAGCTGGGTTCAACTCCGAAGGGATGTGTCA |
| 7_hypothetical | O3 | Tb927.3.5080_for | GGGGACAAGTTTGTACAAAAAAGCAGGCTATGCCTGTAGTGTCGACTGTCG | Tb927.3.5080_rev | GGGGACCACTTTGTACAAGAAAGCTGGGTTCAATTACTTAAATCAGGAGTTGTGGCA |
| 7_hypothetical | O4 | Tb927.9.3670_for | GGGGACAAGTTTGTACAAAAAAGCAGGCTATGCTTCGCATCAGCACCC | Tb927.9.3670_rev | GGGGACCACTTTGTACAAGAAAGCTGGGTTCAATCCTTTATAGATATTTCCCCCACGT |
| 7_hypothetical | O5 | Tb927.5.4080_for | GGGGACAAGTTTGTACAAAAAAGCAGGCTATGTTCTGTTGCTTTTGGGGACTT | Tb927.5.4080_rev | GGGGACCACTTTGTACAAGAAAGCTGGGTTCACACATTCGTGTGGGGGC |
| 7_hypothetical | O6 | Tb927.8.5560_for | GGGGACAAGTTTGTACAAAAAAGCAGGCTATGGCTTTTGCAACCTCCCC | Tb927.8.5560_rev | GGGGACCACTTTGTACAAGAAAGCTGGGTTCACTTTTCCATCCACTTCAAATCAGT |
| 7_hypothetical | O7 | Tb927.7.4300_for | GGGGACAAGTTTGTACAAAAAAGCAGGCTATGCCGCAGATCTTTAGTGATGAATC | Tb927.7.4300_rev | GGGGACCACTTTGTACAAGAAAGCTGGGTTCACACATCTCTGCGATTGTGC |
| 7_hypothetical | O8 | Tb927.10.620_for | GGGGACAAGTTTGTACAAAAAAGCAGGCTATGGAGTTTCCGCAAGTCACATG | Tb927.10.620_rev | GGGGACCACTTTGTACAAGAAAGCTGGGTTTAGATAACAAAATGTGATTGCCACTCCG |
| 7_hypothetical | O9 | Tb927.7.790_for | GGGGACAAGTTTGTACAAAAAAGCAGGCTATGCTGGTCTCCTTGGCGC | Tb927.7.790_rev | GGGGACCACTTTGTACAAGAAAGCTGGGTCTATTCTTGCTGAAAGTCAGCCGC |
| 7_hypothetical | O10 | Tb927.4.1560_for | GGGGACAAGTTTGTACAAAAAAGCAGGCTATGAGTGACTTAAATTTTGAGCGACCT | Tb927.4.1560_rev | GGGGACCACTTTGTACAAGAAAGCTGGGTTCAAGGAACCAGCGCATCAGT |
| 7_hypothetical | O11 | Tb927.10.13520_for | GGGGACAAGTTTGTACAAAAAAGCAGGCTATGAATGTCAACTTGGACGATCTCTTT | Tb927.10.13520_rev | GGGGACCACTTTGTACAAGAAAGCTGGGTCTACTTACGAGCCTCGAAGCAAG |
| 7_hypothetical | O12 | Tb927.9.2760_for | GGGGACAAGTTTGTACAAAAAAGCAGGCTATGGACCATCGCAATACCCATGG | Tb927.9.2760_rev | GGGGACCACTTTGTACAAGAAAGCTGGGTTTACTCTGCAGCGTACAATACATCC |
| 7_hypothetical | O13 | Tb927.10.12670_for | GGGGACAAGTTTGTACAAAAAAGCAGGCTATGATGATAAGTGTAAGGAAAACGGTGAA | Tb927.10.12670_rev | GGGGACCACTTTGTACAAGAAAGCTGGGTCTATATGGACGGACGGACGACC |
| 7_hypothetical | O14 | Tb927.10.5790_for | GGGGACAAGTTTGTACAAAAAAGCAGGCTATGGCCACAATTCTTACAACTAGCA | Tb927.10.5790_rev | GGGGACCACTTTGTACAAGAAAGCTGGGTTCACAAAACTTTTGAGTGTGATATCACCG |
| 7_hypothetical | O15 | Tb927.11.12280_for | GGGGACAAGTTTGTACAAAAAAGCAGGCTATGAATGCCAAATCTTCTTGTCTCGT | Tb927.11.12280_rev | GGGGACCACTTTGTACAAGAAAGCTGGGTTCACTGCTTTACCCCACAACCT |
| 7_hypothetical | O16 | Tb927.9.8030_for | GGGGACAAGTTTGTACAAAAAAGCAGGCTATGAAGAGGAAAGGCATGACCAAC | Tb927.9.8030_rev | GGGGACCACTTTGTACAAGAAAGCTGGGTTCACTGTGTTGACGGGGGAT |
| 7_hypothetical | O17 | Tb927.9.1350_for | GGGGACAAGTTTGTACAAAAAAGCAGGCTATGGTATTCAGAGATGCCTCCCAA | Tb927.9.1350_rev | GGGGACCACTTTGTACAAGAAAGCTGGGTCTACGCACTGGGGTGCATG |
| 7_hypothetical | O18 | Tb11.02.5440_for | GGGGACAAGTTTGTACAAAAAAGCAGGCTATGGAGTGTGATAAACCATCAGGGA | Tb11.02.5440_rev | GGGGACCACTTTGTACAAGAAAGCTGGGTCTACGCTGCAAAGCGCATCC |
| 7_hypothetical | O19 | Tb927.10.9210_for | GGGGACAAGTTTGTACAAAAAAGCAGGCTATGCTTCGCACTAGCATTATTGTGC | Tb927.10.9210_rev | GGGGACCACTTTGTACAAGAAAGCTGGGTTCACTTACTTCTCGATACCTTTATCAAGTC |
| 7_hypothetical | O20 | Tb927.7.6790_for | GGGGACAAGTTTGTACAAAAAAGCAGGCTATGCTGGTGGAATGTTCGGAAAT | Tb927.7.6790_rev | GGGGACCACTTTGTACAAGAAAGCTGGGTTTATTGGGGTTGTAAAACATTACCACCT |
| 7_hypothetical | O21 | Tb927.5.3460_for | GGGGACAAGTTTGTACAAAAAAGCAGGCTATGAATAGCGTATCGTGCCAGATC | Tb927.5.3460_rev | GGGGACCACTTTGTACAAGAAAGCTGGGTTCACAAAACCTGCTCCACTGGC |
| 7_hypothetical | O22 | Tb927.7.2650_for | GGGGACAAGTTTGTACAAAAAAGCAGGCTATGAGCACCACTGCACTGTCT | Tb927.7.2650_rev | GGGGACCACTTTGTACAAGAAAGCTGGGTTCAGTAGTTGGCTTCACGAGCA |
| 7_hypothetical | O23 | Tb927.5.1530_for | GGGGACAAGTTTGTACAAAAAAGCAGGCTATGCAGTCCTGTGCTGTGATTG | Tb927.5.1530_rev | GGGGACCACTTTGTACAAGAAAGCTGGGTCTAAACGGTATTGCTGAAAAGGATGTG |
| 7_hypothetical | O24 | Tb927.7.4030_for | GGGGACAAGTTTGTACAAAAAAGCAGGCTATGATAATCTCTAAGGAGGAGCTGAGTG | Tb927.7.4030_rev | GGGGACCACTTTGTACAAGAAAGCTGGGTTTAAGACTTATTGGAAACAGAGTGCCC |
| 7_hypothetical | P1 | Tb927.5.3070_for | GGGGACAAGTTTGTACAAAAAAGCAGGCTATGGCTAAGAAAAAGAAGTCCAAACGG | Tb927.5.3070_rev | GGGGACCACTTTGTACAAGAAAGCTGGGTTTACCTCTTCCGCCCCTCAG |
| 7_hypothetical | P2 | Tb927.11.4190_for | GGGGACAAGTTTGTACAAAAAAGCAGGCTATGGGAAGAGGGTACAGTAACAGTC | Tb927.11.4190_rev | GGGGACCACTTTGTACAAGAAAGCTGGGTCTATCCTTCATCCTCCTCGTTACCG |
| 7_hypothetical | P3 | Tb927.7.200_for | GGGGACAAGTTTGTACAAAAAAGCAGGCTATGCCGGTAGGCTGGTCC | Tb927.7.200_rev | GGGGACCACTTTGTACAAGAAAGCTGGGTCTAAACATTGCGAATTAGTACGCATACG |
| 7_hypothetical | P4 | Tb927.4.4100_for | GGGGACAAGTTTGTACAAAAAAGCAGGCTATGTCAAGTGAGCAAGGTGGC | Tb927.4.4100_rev | GGGGACCACTTTGTACAAGAAAGCTGGGTTTAAGTACCCTGTTCTGAAGGAGTTATGC |
| 7_hypothetical | P5 | Tb927.7.4760_for | GGGGACAAGTTTGTACAAAAAAGCAGGCTATGCGACTCTCATATGCGGATCA | Tb927.7.4760_rev | GGGGACCACTTTGTACAAGAAAGCTGGGTCTACAAAACGTCCAACAGTCCTGA |
| 7_hypothetical | P6 | Tb927.10.3600_for | GGGGACAAGTTTGTACAAAAAAGCAGGCTATGAGCACCAACGGCGGC | Tb927.10.3600_rev | GGGGACCACTTTGTACAAGAAAGCTGGGTTCACTTCGCTAAGTGATACATCATTCG |
| 7_hypothetical | P7 | Tb927.1.4040_for | GGGGACAAGTTTGTACAAAAAAGCAGGCTATGTCTGATGATGCTGCGGAC | Tb927.1.4040_rev | GGGGACCACTTTGTACAAGAAAGCTGGGTTCAATCCACTAAAATGGTAACACGCC |
| 7_hypothetical | P8 | Tb927.11.14940_for | GGGGACAAGTTTGTACAAAAAAGCAGGCTATGTTTTTCAATCAACTCTTGTTACGTTTG | Tb927.11.14940_rev | GGGGACCACTTTGTACAAGAAAGCTGGGTCTACAGGGTCGTGATATTTGGATGC |
| 7_hypothetical | P9 | Tb927.10.7550_for | GGGGACAAGTTTGTACAAAAAAGCAGGCTATGCCGGCAAAGAAGGGTG | Tb927.10.7550_rev | GGGGACCACTTTGTACAAGAAAGCTGGGTCTATCCCTGCAGTGGCGCA |
| 7_hypothetical | P11 | Tb927.3.1270_for | GGGGACAAGTTTGTACAAAAAAGCAGGCTATGGTTGAACCGCACTTATTTTGTG | Tb927.3.1270_rev | GGGGACCACTTTGTACAAGAAAGCTGGGTTCAGAACTGAAGAAAATGCTCACAATGC |
| 7_hypothetical | P12 | Tb927.9.8050_for | GGGGACAAGTTTGTACAAAAAAGCAGGCTATGTCAAAAGACGGTGTATCACCC | Tb927.9.8050_rev | GGGGACCACTTTGTACAAGAAAGCTGGGTTTAACACACCAAAAGTTCGCAAATCAC |
| 7_hypothetical | P13 | Tb927.10.1340_for | GGGGACAAGTTTGTACAAAAAAGCAGGCTATGCGGCGATGGCTTAATATCG | Tb927.10.1340_rev | GGGGACCACTTTGTACAAGAAAGCTGGGTTCATTTCGCTGGTGCCACTTT |
| 7_hypothetical | P14 | Tb927.10.3300_for | GGGGACAAGTTTGTACAAAAAAGCAGGCTATGACAGAGGTAGCCGTTGATTGT | Tb927.10.3300_rev | GGGGACCACTTTGTACAAGAAAGCTGGGTTCACTCTTTTATCAACTGTGTTGCTCT |
| 7_hypothetical | P15 | Tb927.10.11860_for | GGGGACAAGTTTGTACAAAAAAGCAGGCTATGAGAAGCGCGCAGTTCC | Tb927.10.11860_rev | GGGGACCACTTTGTACAAGAAAGCTGGGTTTATTCAACCGGTGGCGGTCC |
| 7_hypothetical | P16 | Tb927.10.600_for | GGGGACAAGTTTGTACAAAAAAGCAGGCTATGTTGTACACTAGGCGGTTGATG | Tb927.10.600_rev | GGGGACCACTTTGTACAAGAAAGCTGGGTCTAATGAACACCATACGTCACGTTCT |
| 7_hypothetical | P17 | Tb927.7.970_for | GGGGACAAGTTTGTACAAAAAAGCAGGCTATGGATCTCGATGACGAGTTCATTACA | Tb927.7.970_rev | GGGGACCACTTTGTACAAGAAAGCTGGGTTTACACCTCAGGCCGTTTCTTTTT |
| 7_hypothetical | P18 | Tb927.8.8000_for | GGGGACAAGTTTGTACAAAAAAGCAGGCTATGAAAAGCAACCGCAAGAAGGA | Tb927.8.8000_rev | GGGGACCACTTTGTACAAGAAAGCTGGGTTCACATATTCACGTAGGAAACAGAAGG |
| 7_hypothetical | P19 | Tb927.11.15740_for | GGGGACAAGTTTGTACAAAAAAGCAGGCTATGAAAGGTGGTCCCTTGACTAGC | Tb927.11.15740_rev | GGGGACCACTTTGTACAAGAAAGCTGGGTCTACTGGGGCACACCGGT |
| 7_hypothetical | P20 | Tb927.4.2650_for | GGGGACAAGTTTGTACAAAAAAGCAGGCTATGATTCGGGATGCAGAGTTTTGG | Tb927.4.2650_rev | GGGGACCACTTTGTACAAGAAAGCTGGGTCTATAACACCACACAACACTTTCCACA |
| 7_hypothetical | P21 | Tb927.9.15270_for | GGGGACAAGTTTGTACAAAAAAGCAGGCTATGGTTCGGTTTCAGCCATGG | Tb927.9.15270_rev | GGGGACCACTTTGTACAAGAAAGCTGGGTTTAATGAGGTTCCCACAATCCTCTCA |
| 7_hypothetical | P22 | Tb927.7.490_for | GGGGACAAGTTTGTACAAAAAAGCAGGCTATGACAAAATTAGAAGCAAATGCAGCG | Tb927.7.490_rev | GGGGACCACTTTGTACAAGAAAGCTGGGTTTAATACGTGTGACCGTTGAGCAC |
| 7_hypothetical | P23 | Tb927.11.8000_for | GGGGACAAGTTTGTACAAAAAAGCAGGCTATGTTCGCTGATAGCCATTTAGGTG | Tb927.11.8000_rev | GGGGACCACTTTGTACAAGAAAGCTGGGTCTAAGCTACATCTTCTTCCGTTAATGCA |
| 7_hypothetical | P24 | Tb927.9.15450_for | GGGGACAAGTTTGTACAAAAAAGCAGGCTATGCCAGCAAAAAAGGCGACT | Tb927.9.15450_rev | GGGGACCACTTTGTACAAGAAAGCTGGGTTCAAAAAGGCGGTTCCGCTC |
| 7_known | A1 | Tb927.8.1780_for | GGGGACAAGTTTGTACAAAAAAGCAGGCTATGGACCATAAAACTGCTACGGTAGA | Tb927.8.1780_rev | GGGGACCACTTTGTACAAGAAAGCTGGGTTCAGCGCTTTTGAAGGTTTGAGC |
| 7_known | A2 | Tb927.11.4430_for | GGGGACAAGTTTGTACAAAAAAGCAGGCTATGGATTTAATCCCACACATAACGAGAAC | Tb927.11.4430_rev | GGGGACCACTTTGTACAAGAAAGCTGGGTTTATAATAAAAATGTTTCCTGGTGAAAGTC |
| 7_known | A3 | Tb927.9.2240_for | GGGGACAAGTTTGTACAAAAAAGCAGGCTATGGGGTGCAACACTTCCACC | Tb927.9.2240_rev | GGGGACCACTTTGTACAAGAAAGCTGGGTTCAATTGACAACTCCTGTGCATTCT |
| 7_known | A4 | Tb927.4.3410_for | GGGGACAAGTTTGTACAAAAAAGCAGGCTATGGTTTCTGAAGCGGGGAGG | Tb927.4.3410_rev | GGGGACCACTTTGTACAAGAAAGCTGGGTTTACCTGGGTCAGCTAATGCATATGA |
| 7_known | A5 | Tb927.10.5770_for | GGGGACAAGTTTGTACAAAAAAGCAGGCTATGTCAGCTAAGGCGCAACAA | Tb927.10.5770_rev | GGGGACCACTTTGTACAAGAAAGCTGGGTTCAACTATAGAGATCGTCATCTCCTGC |
| 7_known | A6 | Tb927.8.2610_for | GGGGACAAGTTTGTACAAAAAAGCAGGCTATGATGACGACCGCAACGC | Tb927.8.2610_rev | GGGGACCACTTTGTACAAGAAAGCTGGGTCTAGTGCACAGTAAAGTGCCGC |

| | | | | | |
|---|---|---|---|---|---|
| 7_known | A7 | Tb927.11.6790_for | GGGGACAAGTTTGTACAAAAAAGCAGGCTATGGTCAAGAAGCGTGAGCCC | Tb927.11.6790_rev | GGGGACCACTTTGTACAAGAAAGCTGGGTTTATTCCGTCCATGCTGATACAACAC |
| 7_known | A8 | Tb927.11.11340_for | GGGGACAAGTTTGTACAAAAAAGCAGGCTATGTCTGACGGGGTGCAGT | Tb927.11.11340_rev | GGGGACCACTTTGTACAAGAAAGCTGGGTTCAAAACTCTGTGTTGTCGCCG |
| 7_known | A9 | Tb927.10.14990_for | GGGGACAAGTTTGTACAAAAAAGCAGGCTATGCGGTTGAATATTACGATTCCCAA | Tb927.10.14990_rev | GGGGACCACTTTGTACAAGAAAGCTGGGTCTACTGGTAGGGCTTGGCACC |
| 7_known | A10 | Tb927.5.1300_for | GGGGACAAGTTTGTACAAAAAAGCAGGCTATGTCAAGTGAAGCTGCTAGTGGC | Tb927.5.1300_rev | GGGGACCACTTTGTACAAGAAAGCTGGGTTTAGTCCAGTTCCTTCAGCACTTCA |
| 7_known | A11 | Tb927.10.9770_for | GGGGACAAGTTTGTACAAAAAAGCAGGCTATGACGACCCACTTGTGGGC | Tb927.10.9770_rev | GGGGACCACTTTGTACAAGAAAGCTGGGTTCAACGCTTTCGCCTGTCC |
| 7_known | A12 | Tb927.10.640_for | GGGGACAAGTTTGTACAAAAAAGCAGGCTATGAAGCCATGTGTTTCCGCTG | Tb927.10.640_rev | GGGGACCACTTTGTACAAGAAAGCTGGGTTCACGTCAGCAGAGAAATGTGG |
| 7_known | A13 | Tb927.9.5680_for | GGGGACAAGTTTGTACAAAAAAGCAGGCTATGGTGAATGGTCTCCCACCG | Tb927.9.5680_rev | GGGGACCACTTTGTACAAGAAAGCTGGGTTCACAATATCACGGCTGGGGG |
| 7_known | A14 | Tb927.6.950_for | GGGGACAAGTTTGTACAAAAAAGCAGGCTATGAAGGAAAGTGACGGTTTGTTATTAGC | Tb927.6.950_rev | GGGGACCACTTTGTACAAGAAAGCTGGGTCTATGATACTTCGGTGCTTTTTAAGCTCT |
| 7_known | A15 | Tb927.7.3280_for | GGGGACAAGTTTGTACAAAAAAGCAGGCTATGCAGTCGGGACTTTGCCA | Tb927.7.3280_rev | GGGGACCACTTTGTACAAGAAAGCTGGGTTCATATGGTGGTCTTCACTTCGGA |
| 7_known | A16 | Tb927.8.6350_for | GGGGACAAGTTTGTACAAAAAAGCAGGCTATGACTGCTCAAACCCCCTCA | Tb927.8.6350_rev | GGGGACCACTTTGTACAAGAAAGCTGGGTTCAACTCCCTCGCCTTGCC |
| 7_known | A17 | Tb927.4.3910_for | GGGGACAAGTTTGTACAAAAAAGCAGGCTATGCAGGACGAACAGCATGAG | Tb927.4.3910_rev | GGGGACCACTTTGTACAAGAAAGCTGGGTTTAATTCGGTTGCTGCTGCACC |
| 7_known | A18 | Tb927.9.14330_for | GGGGACAAGTTTGTACAAAAAAGCAGGCTATGATTACCCTTTTGGACGAGGGA | Tb927.9.14330_rev | GGGGACCACTTTGTACAAGAAAGCTGGGTTCACGCACGGCGGGACTT |
| 7_known | A19 | Tb927.2.280_for | GGGGACAAGTTTGTACAAAAAAGCAGGCTATGTGGAATTCAATGCGGCGT | Tb927.2.280_rev | GGGGACCACTTTGTACAAGAAAGCTGGGTTCAGGCAGCGGGGCCACC |
| 7_known | A21 | Tb927.2.380_for | GGGGACAAGTTTGTACAAAAAAGCAGGCTATGTGGAATTCAATAAGGCGTTTCTATACT | Tb927.2.380_rev | GGGGACCACTTTGTACAAGAAAGCTGGGTTCAGGCAGCGGGGCCACC |
| 7_known | A22 | Tb927.3.4280_for | GGGGACAAGTTTGTACAAAAAAGCAGGCTATGGACGAAGGTTTGGATGATATTGC | Tb927.3.4280_rev | GGGGACCACTTTGTACAAGAAAGCTGGGTTCAAACACACTCACTGGTCATCAC |
| 7_known | A23 | Tb927.11.12240_for | GGGGACAAGTTTGTACAAAAAAGCAGGCTATGTTGTCATCTTCGTCGCCG | Tb927.11.12240_rev | GGGGACCACTTTGTACAAGAAAGCTGGGTTCATACGAGTGCGTCTTGTTCAC |
| 7_known | A24 | Tb927.9.6150_for | GGGGACAAGTTTGTACAAAAAAGCAGGCTATGAACTCCGGTAGCCCAAGT | Tb927.9.6150_rev | GGGGACCACTTTGTACAAGAAAGCTGGGTTCATGCCTTACGGAAATTTAAGTGGAG |
| 7_known | B1 | Tb927.4.3940_for | GGGGACAAGTTTGTACAAAAAAGCAGGCTATGACGAGCAGCGGTGGT | Tb927.4.3940_rev | GGGGACCACTTTGTACAAGAAAGCTGGGTTTACTCACCCAAATCGAACTCTTTCAG |
| 7_known | B2 | Tb927.11.12220_for | GGGGACAAGTTTGTACAAAAAAGCAGGCTATGCCCTTCAGCAAAGCATGG | Tb927.11.12220_rev | GGGGACCACTTTGTACAAGAAAGCTGGGTTCAGAAGGTGTCGCTTCCGG |
| 7_known | B3 | Tb927.6.2230_for | GGGGACAAGTTTGTACAAAAAAGCAGGCTATGTTGCGGCCATTCGTCG | Tb927.6.2230_rev | GGGGACCACTTTGTACAAGAAAGCTGGGTTTACCTAGTTTGGTGGGAAAGTACCT |
| 7_known | B4 | Tb927.6.3790_for | GGGGACAAGTTTGTACAAAAAAGCAGGCTATGGGCGCCAAAGCAAGG | Tb927.6.3790_rev | GGGGACCACTTTGTACAAGAAAGCTGGGTTTACTTCATCGACACACTTTGCCC |
| 7_known | B5 | Tb927.5.3430_for | GGGGACAAGTTTGTACAAAAAAGCAGGCTATGCACGTTAATGTCGGACATATTGT | Tb927.5.3430_rev | GGGGACCACTTTGTACAAGAAAGCTGGGTCTAATCAATTTCTACAACCTCGTCACTATC |
| 7_known | B6 | Tb927.9.3460_for | GGGGACAAGTTTGTACAAAAAAGCAGGCTATGGGAAGGAAGGATGACGCC | Tb927.9.3460_rev | GGGGACCACTTTGTACAAGAAAGCTGGGTTTACTTACGCCGCGGTGC |
| 7_known | B7 | Tb927.10.910_for | GGGGACAAGTTTGTACAAAAAAGCAGGCTATGGTAGTACTGCGGCCAAAGA | Tb927.10.910_rev | GGGGACCACTTTGTACAAGAAAGCTGGGTTCAATCATCGTCCACCATTAGATTGTT |
| 7_known | B8 | Tb927.11.2950_for | GGGGACAAGTTTGTACAAAAAAGCAGGCTATGCTGACAACATTTGAGCAGCA | Tb927.11.2950_rev | GGGGACCACTTTGTACAAGAAAGCTGGGTTCACCATTTCGGTTGTGATGTCAC |
| 7_known | B9 | Tb927.8.3220_for | GGGGACAAGTTTGTACAAAAAAGCAGGCTATGGGGATTAAAGGGTTATGGGGAG | Tb927.8.3220_rev | GGGGACCACTTTGTACAAGAAAGCTGGGTCTACCGTCGCGTATAAGATAACAGC |
| 7_known | B10 | Tb927.5.1090_for | GGGGACAAGTTTGTACAAAAAAGCAGGCTATGGCCACGGCTGTTGTTG | Tb927.5.1090_rev | GGGGACCACTTTGTACAAGAAAGCTGGGTTTAGAAGGACGGATCATATTTATCGGCC |
| 7_known | B11 | Tb927.11.1740_for | GGGGACAAGTTTGTACAAAAAAGCAGGCTATGGACTTACAACAGGGTGACGG | Tb927.11.1740_rev | GGGGACCACTTTGTACAAGAAAGCTGGGTCTATATTCCCGGGAGGTCAATTTCATC |
| 7_known | B12 | Tb927.11.11570_for | GGGGACAAGTTTGTACAAAAAAGCAGGCTATGGACATATCAGCCACGCGC | Tb927.11.11570_rev | GGGGACCACTTTGTACAAGAAAGCTGGGTTCAAAACATTCCCTCCTCTTCTTCTTC |
| 7_known | B13 | Tb927.5.890_for | GGGGACAAGTTTGTACAAAAAAGCAGGCTATGACGAAAGGTGGGAAAGTAGCT | Tb927.5.890_rev | GGGGACCACTTTGTACAAGAAAGCTGGGTTCATTCGTAATGGAACCGCTTCG |
| 7_known | B14 | Tb927.4.760_for | GGGGACAAGTTTGTACAAAAAAGCAGGCTATGAACAGCCAAAAGCTTCGTGA | Tb927.4.760_rev | GGGGACCACTTTGTACAAGAAAGCTGGGTCTAAACCTCTTGAGACACCTGCAG |
| 7_known | B15 | Tb927.10.6220_for | GGGGACAAGTTTGTACAAAAAAGCAGGCTATGGGGATTGCCGGTTTTTACC | Tb927.10.6220_rev | GGGGACCACTTTGTACAAGAAAGCTGGGTTCAGCGCCCAATGACTCTGT |
| 7_known | B16 | Tb927.8.5980_for | GGGGACAAGTTTGTACAAAAAAGCAGGCTATGAAGTTGTTCATTGAAGACGTGCC | Tb927.8.5980_rev | GGGGACCACTTTGTACAAGAAAGCTGGGTTCACTCTCGGGACCGCTTT |
| 7_known | B17 | Tb927.4.2730_for | GGGGACAAGTTTGTACAAAAAAGCAGGCTATGACCACTCTATCTTTAGATTCAACAGC | Tb927.4.2730_rev | GGGGACCACTTTGTACAAGAAAGCTGGGTTCATAACTCACCCCTCATACCTATTTCTT |
| 7_known | B18 | Tb927.2.4130_for | GGGGACAAGTTTGTACAAAAAAGCAGGCTATGCGTCGCTTGGAAACCAT | Tb927.2.4130_rev | GGGGACCACTTTGTACAAGAAAGCTGGGTTTACGGCCACGTGGAACGA |
| 7_known | B19 | Tb927.11.1110_for | GGGGACAAGTTTGTACAAAAAAGCAGGCTATGCAGCGGTCGTTGCGC | Tb927.11.1110_rev | GGGGACCACTTTGTACAAGAAAGCTGGGTTCATCTACGCTGCGGTGTCG |
| 7_known | B20 | Tb927.11.11430_for | GGGGACAAGTTTGTACAAAAAAGCAGGCTATGGAGCAGACCAACCATGACG | Tb927.11.11430_rev | GGGGACCACTTTGTACAAGAAAGCTGGGTTTATGTAACGTACGAAAACGTGTCTGT |
| 7_known | B21 | Tb927.2.3620_for | GGGGACAAGTTTGTACAAAAAAGCAGGCTATGGCTGACGGATCACATGACA | Tb927.2.3620_rev | GGGGACCACTTTGTACAAGAAAGCTGGGTCTACGTTTCTTTATGTTTCAACCTCTTCG |
| 7_known | B22 | Tb927.7.5450_for | GGGGACAAGTTTGTACAAAAAAGCAGGCTATGATATCTGGTTTTGCAGATTTCTTTAAC | Tb927.7.5450_rev | GGGGACCACTTTGTACAAGAAAGCTGGGTTCACACAGTGTCACCTTGTTGATAG |
| 7_known | B23 | Tb927.10.1900_for | GGGGACAAGTTTGTACAAAAAAGCAGGCTATGTTTCGGCAAACAGTTACATCACT | Tb927.10.1900_rev | GGGGACCACTTTGTACAAGAAAGCTGGGTTCACCCACCGCCCGCAAC |
| 7_known | B24 | Tb927.10.12980_for | GGGGACAAGTTTGTACAAAAAAGCAGGCTATGTGGAAACCGAAACAGCGC | Tb927.10.12980_rev | GGGGACCACTTTGTACAAGAAAGCTGGGTTCAAGGATCACGCACGTCTGG |
| 7_known | C1 | Tb927.10.13630_for | GGGGACAAGTTTGTACAAAAAAGCAGGCTATGATGCTATCGCTTGTGCTATCG | Tb927.10.13630_rev | GGGGACCACTTTGTACAAGAAAGCTGGGTCTATCTCTTCAGCACAATGGTCCAG |
| 7_known | C2 | Tb927.3.620_for | GGGGACAAGTTTGTACAAAAAAGCAGGCTATGTCACTAGCGGGTTCCGA | Tb927.3.620_rev | GGGGACCACTTTGTACAAGAAAGCTGGGTTTAGGCATCGAATTCAGTAAATTGCTGG |
| 7_known | C3 | Tb927.11.2270_for | GGGGACAAGTTTGTACAAAAAAGCAGGCTATGTCTACGGAAGCAGAGGATGAT | Tb927.11.2270_rev | GGGGACCACTTTGTACAAGAAAGCTGGGTTCAAATTGGTGAGGCCCGTG |
| 7_known | C4 | Tb927.7.7120_for | GGGGACAAGTTTGTACAAAAAAGCAGGCTATGAATGGGATTCATTTAAATTCGACTATT | Tb927.7.7120_rev | GGGGACCACTTTGTACAAGAAAGCTGGGTTCATAAGGATGTCTCGCAACTCTCG |
| 7_known | C5 | Tb927.10.12710_for | GGGGACAAGTTTGTACAAAAAAGCAGGCTATGTCAGTCTTCGGTGTGGACT | Tb927.10.12710_rev | GGGGACCACTTTGTACAAGAAAGCTGGGTCTAGTCAAGCTGATCATCCGACG |
| 7_known | C6 | Tb927.5.940_for | GGGGACAAGTTTGTACAAAAAAGCAGGCTATGTTTTCCGGTCCTGCTGAAG | Tb927.5.940_rev | GGGGACCACTTTGTACAAGAAAGCTGGGTTCAAAGAGTGACTATGTTTGATTTGTTATG |
| 7_known | C7 | Tb927.2.3030_for | GGGGACAAGTTTGTACAAAAAAGCAGGCTATGCGAAAGCTGGGTTCGG | Tb927.2.3030_rev | GGGGACCACTTTGTACAAGAAAGCTGGGTTCATTCTGTGAGTTGCGGACCC |
| 7_known | C8 | Tb927.10.15880_for | GGGGACAAGTTTGTACAAAAAAGCAGGCTATGTATTTAAAACCATATAACAATCCCTAC | Tb927.10.15880_rev | GGGGACCACTTTGTACAAGAAAGCTGGGTTTAGCAACGCCCCTCCTGT |
| 7_known | C9 | Tb927.2.3930_for | GGGGACAAGTTTGTACAAAAAAGCAGGCTATGCTTAGCAACAGGACCCTCA | Tb927.2.3930_rev | GGGGACCACTTTGTACAAGAAAGCTGGGTTCAAATGAACTGAACCCATTCATCTTCG |
| 7_known | C10 | Tb927.5.2790_for | GGGGACAAGTTTGTACAAAAAAGCAGGCTATGCTTTTGTTCCGCCCCAC | Tb927.5.2790_rev | GGGGACCACTTTGTACAAGAAAGCTGGGTTCATTTTGATCTTCGTTTTTTATCTGCTCC |
| 7_known | C11 | Tb927.9.1340_for | GGGGACAAGTTTGTACAAAAAAGCAGGCTATGAGCATCAGTGAGTCGGACA | Tb927.9.1340_rev | GGGGACCACTTTGTACAAGAAAGCTGGGTCTACTGTGGCTGCTTTACTTCCTC |
| 7_known | C12 | Tb927.5.4310_for | GGGGACAAGTTTGTACAAAAAAGCAGGCTATGCAATCGTTGGGGAGGAGC | Tb927.5.4310_rev | GGGGACCACTTTGTACAAGAAAGCTGGGTTTAGTAACCTTGAACAACCTGCCTG |
| 7_known | C13 | Tb927.3.1440_for | GGGGACAAGTTTGTACAAAAAAGCAGGCTATGGGTCGCCACCATGATGT | Tb927.3.1440_rev | GGGGACCACTTTGTACAAGAAAGCTGGGTCTACTTACTTCCGAAAGTTTTGCTCCA |
| 7_known | C14 | Tb927.3.2610_for | GGGGACAAGTTTGTACAAAAAAGCAGGCTATGCGCCTTTGTGGTGATGAA | Tb927.3.2610_rev | GGGGACCACTTTGTACAAGAAAGCTGGGTTTAGTTTGAGGTACCAGGAGCTAGAAG |
| 7_known | C15 | Tb927.3.1490_for | GGGGACAAGTTTGTACAAAAAAGCAGGCTATGGCAACACAGTTAAATAGTGATACGG | Tb927.3.1490_rev | GGGGACCACTTTGTACAAGAAAGCTGGGTTCAGATTGTTTTGACTCTAACACCCC |
| 7_known | C16 | Tb927.7.7480_for | GGGGACAAGTTTGTACAAAAAAGCAGGCTATGGAACGCCTGCAGGTGC | Tb927.7.7480_rev | GGGGACCACTTTGTACAAGAAAGCTGGGTTCAAATATTAGTAGGAACCACGTCCCG |
| 7_known | C17 | Tb927.5.640_for | GGGGACAAGTTTGTACAAAAAAGCAGGCTATGAAGCGCCTGCCTGTACG | Tb927.5.640_rev | GGGGACCACTTTGTACAAGAAAGCTGGGTTCAGATTATAGTAGAATCCCCGTCCCA |
| 7_known | C18 | Tb927.11.8980_for | GGGGACAAGTTTGTACAAAAAAGCAGGCTATGAAAGGGTTCCGGTGTGC | Tb927.11.8980_rev | GGGGACCACTTTGTACAAGAAAGCTGGGTTTAACTGTCGTATATACCTAAGTTCTTCAT |
| 7_known | C19 | Tb927.1.1690_for | GGGGACAAGTTTGTACAAAAAAGCAGGCTATGGGGCTACATTGGCCCC | Tb927.1.1690_rev | GGGGACCACTTTGTACAAGAAAGCTGGGTTCACGCACCAACCGAGATGC |
| 7_known | C20 | Tb927.11.230_for | GGGGACAAGTTTGTACAAAAAAGCAGGCTATGCTGCGCGCTTCGGCT | Tb927.11.230_rev | GGGGACCACTTTGTACAAGAAAGCTGGGTTCACAGAACTTGAGAAAAGTGCTTGT |
| 7_known | C21 | Tb927.11.12430_for | GGGGACAAGTTTGTACAAAAAAGCAGGCTATGCCGAGGCCACTCCGA | Tb927.11.12430_rev | GGGGACCACTTTGTACAAGAAAGCTGGGTTTAAGGTTGGAGGAATACAACCCGC |
| 7_known | C22 | Tb927.11.14790_for | GGGGACAAGTTTGTACAAAAAAGCAGGCTATGCTCCTCAACTCCACCAGG | Tb927.11.14790_rev | GGGGACCACTTTGTACAAGAAAGCTGGGTTCACTTGTGGTGTTTCCACGTC |
| 7_known | C23 | Tb927.10.14890_for | GGGGACAAGTTTGTACAAAAAAGCAGGCTATGTCTGTGGAACAACCCGAGC | Tb927.10.14890_rev | GGGGACCACTTTGTACAAGAAAGCTGGGTTTATGGATTCTTCTTGATCTGGCCAAG |
| 7_known | C24 | Tb927.7.3490_for | GGGGACAAGTTTGTACAAAAAAGCAGGCTATGTACGTGGAGAAGAGCCCG | Tb927.7.3490_rev | GGGGACCACTTTGTACAAGAAAGCTGGGTTTACTTGTATAATCCCCGCACGAAATT |
| 7_known | D1 | Tb927.9.7700_for | GGGGACAAGTTTGTACAAAAAAGCAGGCTATGTGGGTCTTTAGCGACCGA | Tb927.9.7700_rev | GGGGACCACTTTGTACAAGAAAGCTGGGTTTAGGCACTAGGCACAGTGGAG |
| 7_known | D2 | Tb927.5.910_for | GGGGACAAGTTTGTACAAAAAAGCAGGCTATGACGAAAGGTGGGAAAGTAGCT | Tb927.5.910_rev | GGGGACCACTTTGTACAAGAAAGCTGGGTCTAACCCATCTGACGCATATACGC |
| 7_known | D4 | Tb927.7.2970_for | GGGGACAAGTTTGTACAAAAAAGCAGGCTATGCCCGTCTCCCTTACTTCAG | Tb927.7.2970_rev | GGGGACCACTTTGTACAAGAAAGCTGGGTTTAATTCTCGGATTCGCTGGTTTCA |
| 7_known | D5 | Tb927.8.7980_for | GGGGACAAGTTTGTACAAAAAAGCAGGCTATGATTAAATTGTCACCACTGCTTCTAAAA | Tb927.8.7980_rev | GGGGACCACTTTGTACAAGAAAGCTGGGTTTACTTCACCAGAAGCCCTCCC |
| 7_known | D6 | Tb927.4.4380_for | GGGGACAAGTTTGTACAAAAAAGCAGGCTATGATTAAAATGTCACCGCTGCTTCT | Tb927.4.4380_rev | GGGGACCACTTTGTACAAGAAAGCTGGGTTTACTTCACCAGAAGCCCTCCC |
| 7_known | D7 | Tb927.10.7310_for | GGGGACAAGTTTGTACAAAAAAGCAGGCTATGTTCACATCGGTGCCTGTAGC | Tb927.10.7310_rev | GGGGACCACTTTGTACAAGAAAGCTGGGTCTAGTTTTTCTGACAGTTGTACGCCT |
| 7_known | D8 | Tb927.5.4270_for | GGGGACAAGTTTGTACAAAAAAGCAGGCTATGGATGAAAGGCAGATGGAGGA | Tb927.5.4270_rev | GGGGACCACTTTGTACAAGAAAGCTGGGTTTATTTACTTTTTTGTGGGGTGGGATTATT |
| 7_known | D9 | Tb927.5.295b_for | GGGGACAAGTTTGTACAAAAAAGCAGGCTATGTTTTCCAAACGCTGTTTTGCA | Tb927.5.295b_rev | GGGGACCACTTTGTACAAGAAAGCTGGGTCTATGCAATGTTACCACCCCTATTGT |
| 7_known | D10 | Tb927.2.100_for | GGGGACAAGTTTGTACAAAAAAGCAGGCTATGTCAAGGGCAAATAGTCCCGC | Tb927.2.100_rev | GGGGACCACTTTGTACAAGAAAGCTGGGTCTATGCATTGTTACCACCTCTATTGTTAAC |
| 7_known | D11 | Tb927.5.2970_for | GGGGACAAGTTTGTACAAAAAAGCAGGCTATGTTGACGCTCCGTGAACTAGA | Tb927.5.2970_rev | GGGGACCACTTTGTACAAGAAAGCTGGGTTTAAAAGACTGTGGGCGTCGG |
| 7_known | D12 | Tb927.2.3780_for | GGGGACAAGTTTGTACAAAAAAGCAGGCTATGCCGCCTAAAGGACCAAAG | Tb927.2.3780_rev | GGGGACCACTTTGTACAAGAAAGCTGGGTCTATCGAGGCGGCACCTTCA |
| 7_known | D13 | Tb927.1.180_for | GGGGACAAGTTTGTACAAAAAAGCAGGCTATGTCAAGAGCAAATAGTCCCGC | Tb927.1.180_rev | GGGGACCACTTTGTACAAGAAAGCTGGGTCTATGCATTGTTACCACCTCTATTGTTAAC |
| 7_known | D14 | Tb927.1.220_for | GGGGACAAGTTTGTACAAAAAAGCAGGCTATGTCAAGAGCAAATAGTCCCGC | Tb927.1.220_rev | GGGGACCACTTTGTACAAGAAAGCTGGGTCTATGCATTGTTACCACCTCTATTGTTAAC |
| 7_known | D15 | Tb927.11.4990_for | GGGGACAAGTTTGTACAAAAAAGCAGGCTATGGATTTAAACACAGTCGACGAGAG | Tb927.11.4990_rev | GGGGACCACTTTGTACAAGAAAGCTGGGTTCAATCAAGGTTAACCACAGGTTTACC |
| 7_known | D16 | Tb927.2.370_for | GGGGACAAGTTTGTACAAAAAAGCAGGCTATGTCAAGAGCAAATAGTCCCGC | Tb927.2.370_rev | GGGGACCACTTTGTACAAGAAAGCTGGGTCTATGCATTGTTACCACCTCTATTGTTAAC |
| 7_known | D17 | Tb927.10.15540_for | GGGGACAAGTTTGTACAAAAAAGCAGGCTATGCAGTCTGTGAGGTTCAGCG | Tb927.10.15540_rev | GGGGACCACTTTGTACAAGAAAGCTGGGTCTAATTAGCACTATTGTCGTTGCTGTCC |
| 7_known | D18 | Tb927.9.4500_for | GGGGACAAGTTTGTACAAAAAAGCAGGCTATGTCCATGCTCTGTTTGGCAC | Tb927.9.4500_rev | GGGGACCACTTTGTACAAGAAAGCTGGGTTCACTCACCGAGCTTTAGGTCC |
| 7_known | D19 | Tb927.11.12250_for | GGGGACAAGTTTGTACAAAAAAGCAGGCTATGAACAGGCATGCATTTGTGGA | Tb927.11.12250_rev | GGGGACCACTTTGTACAAGAAAGCTGGGTTCACGAAAGCCTACCGGCG |
| 7_known | D20 | Tb927.2.1380_for | GGGGACAAGTTTGTACAAAAAAGCAGGCTATGGGAATAACCCTAGCCAACAAAAA | Tb927.2.1380_rev | GGGGACCACTTTGTACAAGAAAGCTGGGTCTAACATTGAGAGCAGGGGACAGA |
| 7_known | D21 | Tb927.2.510_for | GGGGACAAGTTTGTACAAAAAAGCAGGCTATGTCAAGAGCAAATAGTCCCGC | Tb927.2.510_rev | GGGGACCACTTTGTACAAGAAAGCTGGGTTCACCTCCCCTGTGCCTTTT |
| 7_known | D22 | Tb927.11.7840_for | GGGGACAAGTTTGTACAAAAAAGCAGGCTATGTTGGAAACGGTGAAGCTTGT | Tb927.11.7840_rev | GGGGACCACTTTGTACAAGAAAGCTGGGTCTACGAACCACAATTCAGGCATCC |
| 7_known | D23 | Tb927.11.5070_for | GGGGACAAGTTTGTACAAAAAAGCAGGCTATGACCTCACGCTCTGATGCT | Tb927.11.5070_rev | GGGGACCACTTTGTACAAGAAAGCTGGGTTCATGCCAGTAACGCAGCC |
| 7_known | D24 | Tb927.5.4460_for | GGGGACAAGTTTGTACAAAAAAGCAGGCTATGAGTGATATCATACGAATTAAACGTCAT | Tb927.5.4460_rev | GGGGACCACTTTGTACAAGAAAGCTGGGTTTATGACTTTGTCTCGGTTGTTTGGC |
| 7_known | E1 | Tb927.10.3890_for | GGGGACAAGTTTGTACAAAAAAGCAGGCTATGCTGCAGTCCGATGTGTCT | Tb927.10.3890_rev | GGGGACCACTTTGTACAAGAAAGCTGGGTCTACATTGTAGCATTGCACAAAACACC |
| 7_known | E2 | Tb927.11.13640_for | GGGGACAAGTTTGTACAAAAAAGCAGGCTATGCGAAGGCAGGTGAAGAAG | Tb927.11.13640_rev | GGGGACCACTTTGTACAAGAAAGCTGGGTCTATTTCTCCCGTCTGCGGACG |
| 7_known | E3 | Tb927.7.1210_for | GGGGACAAGTTTGTACAAAAAAGCAGGCTATGCACGAACCTCGTGAAGTAGT | Tb927.7.1210_rev | GGGGACCACTTTGTACAAGAAAGCTGGGTCTAACGGGCACTTCGCCTG |
| 7_known | E4 | Tb927.8.7840_for | GGGGACAAGTTTGTACAAAAAAGCAGGCTATGACGACGAAGTCAGCCGT | Tb927.8.7840_rev | GGGGACCACTTTGTACAAGAAAGCTGGGTTCACACATTTTGGGGAATGGGAAG |
| 7_known | E5 | Tb927.6.5100_for | GGGGACAAGTTTGTACAAAAAAGCAGGCTATGTTTAACGTAAGAACGCCGATGT | Tb927.6.5100_rev | GGGGACCACTTTGTACAAGAAAGCTGGGTTTAAAAGCACTCCACAATATTTGCACAG |
| 7_known | E6 | Tb927.2.1210_for | GGGGACAAGTTTGTACAAAAAAGCAGGCTATGTCAAGGGCAAATAGTCCCGC | Tb927.2.1210_rev | GGGGACCACTTTGTACAAGAAAGCTGGGTTCACCCTCCTTGCGCTCC |
| 7_known | E7 | Tb927.11.9170_for | GGGGACAAGTTTGTACAAAAAAGCAGGCTATGGTCCAAACAGTACTTATGGTTGC | Tb927.11.9170_rev | GGGGACCACTTTGTACAAGAAAGCTGGGTCTAGTTTCGGTAGCCTCTTCCTCT |
| 7_known | E8 | Tb927.5.2140_for | GGGGACAAGTTTGTACAAAAAAGCAGGCTATGTTCAGTGAGCATGCTAGTGATG | Tb927.5.2140_rev | GGGGACCACTTTGTACAAGAAAGCTGGGTTCAAGACTCACCAGGCAAGAACA |
| 7_known | E9 | Tb927.4.2630_for | GGGGACAAGTTTGTACAAAAAAGCAGGCTATGAAAGGTGTAGGTACGTGGGA | Tb927.4.2630_rev | GGGGACCACTTTGTACAAGAAAGCTGGGTTCACCTTTTTCGTTTACCTTTCTGCT |
| 7_known | E10 | Tb927.11.12600_for | GGGGACAAGTTTGTACAAAAAAGCAGGCTATGGAGCACGAACAAAGGGTTT | Tb927.11.12600_rev | GGGGACCACTTTGTACAAGAAAGCTGGGTTCACTCTTGCTGCCCTTCACT |
| 7_known | E11 | Tb927.10.11760_for | GGGGACAAGTTTGTACAAAAAAGCAGGCTATGAGTTCAACCAAAGAGCCGC | Tb927.10.11760_rev | GGGGACCACTTTGTACAAGAAAGCTGGGTTCACTCGGCATCGAAGTGCT |
| 7_known | E12 | Tb927.7.6460_for | GGGGACAAGTTTGTACAAAAAAGCAGGCTATGACCCTCACAGGCAACATCA | Tb927.7.6460_rev | GGGGACCACTTTGTACAAGAAAGCTGGGTCTAACCAGTAAGAGGTACCTGATGTTTTT |
| 7_known | E13 | Tb927.2.560_for | GGGGACAAGTTTGTACAAAAAAGCAGGCTATGTCAAGAGCAAATAGTCCCGC | Tb927.2.560_rev | GGGGACCACTTTGTACAAGAAAGCTGGGTTCACCCTCCTTGCGCTCC |
| 7_known | E14 | Tb927.11.5550_for | GGGGACAAGTTTGTACAAAAAAGCAGGCTATGAAAAATCTTTCAATCTTCTCCGCAAG | Tb927.11.5550_rev | GGGGACCACTTTGTACAAGAAAGCTGGGTTTAAGGAACATCAAGTTTCTCGAGTGAA |
| 7_known | E15 | Tb927.6.3430_for | GGGGACAAGTTTGTACAAAAAAGCAGGCTATGGAGGATTACACTATTTACCGTCCA | Tb927.6.3430_rev | GGGGACCACTTTGTACAAGAAAGCTGGGTTCATACCTCACGGTTTTCAATCCAG |
| 7_known | E16 | Tb927.10.4570_for | GGGGACAAGTTTGTACAAAAAAGCAGGCTATGGTCAACTTCACCGTTGATGAG | Tb927.10.4570_rev | GGGGACCACTTTGTACAAGAAAGCTGGGTCTATAGCTTGTCCAGGAAGGTGTCG |
| 7_known | E17 | Tb927.10.1460_for | GGGGACAAGTTTGTACAAAAAAGCAGGCTATGAGATCTTGCTGTAGCTGTGGA | Tb927.10.1460_rev | GGGGACCACTTTGTACAAGAAAGCTGGGTCTACACTCTGACGAATAGGTTTCGG |
| 7_known | E19 | Tb927.11.10810_for | GGGGACAAGTTTGTACAAAAAAGCAGGCTATGAACGTCGCGTGGGGG | Tb927.11.10810_rev | GGGGACCACTTTGTACAAGAAAGCTGGGTTTATTTATGCGATGCAGGAATTCCGG |
| 7_known | E20 | Tb927.10.12660_for | GGGGACAAGTTTGTACAAAAAAGCAGGCTATGTCTGGTTGGGACGAACCC | Tb927.10.12660_rev | GGGGACCACTTTGTACAAGAAAGCTGGGTCTACAGCGTTGGCATGCAGG |
| 7_known | E21 | Tb927.11.10900_for | GGGGACAAGTTTGTACAAAAAAGCAGGCTATGGAAAACGGAAAGAATGACGAATTTAC | Tb927.11.10900_rev | GGGGACCACTTTGTACAAGAAAGCTGGGTTCACGACTTAATGCTCCTCGAAGC |
| 7_known | E22 | Tb927.11.12350_for | GGGGACAAGTTTGTACAAAAAAGCAGGCTATGAAGCGAATATTTAGGAGTTCGGTT | Tb927.11.12350_rev | GGGGACCACTTTGTACAAGAAAGCTGGGTCTACTCGTCGGAGGCACTTAGC |
| 7_known | E23 | Tb927.7.960_for | GGGGACAAGTTTGTACAAAAAAGCAGGCTATGACCAGAAATGCTGATGCACA | Tb927.7.960_rev | GGGGACCACTTTGTACAAGAAAGCTGGGTTCAGTTGTTGGTGGCATCATACAG |
| 7_known | E24 | Tb927.10.6940_for | GGGGACAAGTTTGTACAAAAAAGCAGGCTATGCCGACCTTGCCTTTGTC | Tb927.10.6940_rev | GGGGACCACTTTGTACAAGAAAGCTGGGTTTAAAGCACCTCCATAAGTGTCTTTGC |
| 7_known | F2 | Tb927.4.3950_for | GGGGACAAGTTTGTACAAAAAAGCAGGCTATGGGTTGTGGTGGATCAAAGGT | Tb927.4.3950_rev | GGGGACCACTTTGTACAAGAAAGCTGGGTTTATTCATCACTGTCAGTTTCATCACCC |
| 7_known | F3 | Tb927.2.6050_for | GGGGACAAGTTTGTACAAAAAAGCAGGCTATGTCGTCAAAAATGATGTCACCGC | Tb927.2.6050_rev | GGGGACCACTTTGTACAAGAAAGCTGGGTTTACTGCTCACCCCATTCGTCG |
| 7_known | F4 | Tb927.11.370_for | GGGGACAAGTTTGTACAAAAAAGCAGGCTATGACCGACGTTGACGGTGG | Tb927.11.370_rev | GGGGACCACTTTGTACAAGAAAGCTGGGTTTATTGTGGCGCTTTCGAAGTGG |
| 7_known | F5 | Tb927.9.7540_for | GGGGACAAGTTTGTACAAAAAAGCAGGCTATGATGCAGGCATCAGCAGATG | Tb927.9.7540_rev | GGGGACCACTTTGTACAAGAAAGCTGGGTTTAGGAAAAAGAGGAACCAACTAACTCTTC |
| 7_known | F6 | Tb927.1.120_for | GGGGACAAGTTTGTACAAAAAAGCAGGCTATGAATCAACAATTTCCTATTGAAGGAAGG | Tb927.1.120_rev | GGGGACCACTTTGTACAAGAAAGCTGGGTTCATCTCCTTCCACGTCGTCTC |
| 7_known | F7 | Tb927.8.4170_for | GGGGACAAGTTTGTACAAAAAAGCAGGCTATGTCTCGCGTACGTATTTTGAACC | Tb927.8.4170_rev | GGGGACCACTTTGTACAAGAAAGCTGGGTTTACGATTCGTCCAGCTTGGC |
| 7_known | F9 | Tb927.11.9270_for | GGGGACAAGTTTGTACAAAAAAGCAGGCTATGCTATCCAAGTTATTCCGGTATGACA | Tb927.11.9270_rev | GGGGACCACTTTGTACAAGAAAGCTGGGTTTAGCAGGAACCCGTCCGT |
| 7_known | F10 | Tb927.2.340_for | GGGGACAAGTTTGTACAAAAAAGCAGGCTATGAATCAACAGGTTCCTATTGAAGGAAG | Tb927.2.340_rev | GGGGACCACTTTGTACAAGAAAGCTGGGTTTACGCGTCTTCACCTCCTTCC |
| 7_known | F11 | Tb927.11.7770_for | GGGGACAAGTTTGTACAAAAAAGCAGGCTATGAGTGACGGGGATGAGGG | Tb927.11.7770_rev | GGGGACCACTTTGTACAAGAAAGCTGGGTCTACCAAGGTATTTCCCTAAAATCCTTGAA |
| 7_known | F13 | Tb927.11.2880_for | GGGGACAAGTTTGTACAAAAAAGCAGGCTATGAGTATTGAGGTTCATGTACGCATT | Tb927.11.2880_rev | GGGGACCACTTTGTACAAGAAAGCTGGGTTCAACGACGGCGGGAGCG |
| 7_known | F14 | Tb927.10.6310_for | GGGGACAAGTTTGTACAAAAAAGCAGGCTATGAATGATTATTTAGCGAATGAGCTGGT | Tb927.10.6310_rev | GGGGACCACTTTGTACAAGAAAGCTGGGTTTACTGCTGCACATGACCAGTC |
| 7_known | F15 | Tb927.10.1990_for | GGGGACAAGTTTGTACAAAAAAGCAGGCTATGGTGGACAAGGAGAATCAGGT | Tb927.10.1990_rev | GGGGACCACTTTGTACAAGAAAGCTGGGTTCATGGCAACGCGTCGTT |
| 7_known | F16 | Tb927.8.6880_for | GGGGACAAGTTTGTACAAAAAAGCAGGCTATGCGTCACACTCGTTTTTTTCAG | Tb927.8.6880_rev | GGGGACCACTTTGTACAAGAAAGCTGGGTTTATGTGGAAGGTGTTAGTAGAAGTTGTC |
| 7_known | F17 | Tb927.2.4060_for | GGGGACAAGTTTGTACAAAAAAGCAGGCTATGAAAAATGGCTCCTCGAGTGG | Tb927.2.4060_rev | GGGGACCACTTTGTACAAGAAAGCTGGGTTTATGTGGTCGCCTTTCGAACTAC |
| 7_known | F18 | Tb927.3.4220_for | GGGGACAAGTTTGTACAAAAAAGCAGGCTATGGACGCACCGCTGCAG | Tb927.3.4220_rev | GGGGACCACTTTGTACAAGAAAGCTGGGTCTACTCAAAATCCTCATCGTCGTCG |
| 7_known | F19 | Tb927.10.2900_for | GGGGACAAGTTTGTACAAAAAAGCAGGCTATGAGCTCCTTGACGGAATTGTTAAC | Tb927.10.2900_rev | GGGGACCACTTTGTACAAGAAAGCTGGGTTTACACCACCTTTGAAAGTTGTTGAAGT |
| 7_known | F20 | Tb927.10.12440_for | GGGGACAAGTTTGTACAAAAAAGCAGGCTATGGGACGAAGGACACATGTGG | Tb927.10.12440_rev | GGGGACCACTTTGTACAAGAAAGCTGGGTCTACATTATTACACAGTAATGGCAACCCT |
| 7_known | F21 | Tb927.8.2810_for | GGGGACAAGTTTGTACAAAAAAGCAGGCTATGGGTGTCCCATTACTTCTCACT | Tb927.8.2810_rev | GGGGACCACTTTGTACAAGAAAGCTGGGTTTAGGTCCCCAGGGTGGCA |
| 7_known | F22 | Tb927.11.11640_for | GGGGACAAGTTTGTACAAAAAAGCAGGCTATGGACCTCGTGCCAACAGG | Tb927.11.11640_rev | GGGGACCACTTTGTACAAGAAAGCTGGGTCTACTGCTTCGTAATATCTGGGTTATAGTT |

| | | | | | |
|---|---|---|---|---|---|
| 7_known | F23 | Tb927.2.5980_for | GGGGACAAGTTTGTACAAAAAAGCAGGCTATGGCACATTCAGATCGCCAAT | Tb927.2.5980_rev | GGGGACCACTTTGTACAAGAAAGCTGGGTTTATTCCCATTCGTCACTGGTAAGC |
| 7_known | F24 | Tb927.7.1030_for | GGGGACAAGTTTGTACAAAAAAGCAGGCTATGCACTGTTGTAAGGTTTTTTCGTTT | Tb927.7.1030_rev | GGGGACCACTTTGTACAAGAAAGCTGGGTCTAGTCCATCGCGAACAGTTTGG |
| 7_known | G1 | Tb927.3.4750_for | GGGGACAAGTTTGTACAAAAAAGCAGGCTATGCCAACTCTCCCCTCAGATG | Tb927.3.4750_rev | GGGGACCACTTTGTACAAGAAAGCTGGGTTCACCGCGGAAAGAACAGGT |
| 7_known | G3 | Tb927.11.3570_for | GGGGACAAGTTTGTACAAAAAAGCAGGCTATGTCCACGATCGAACGGGA | Tb927.11.3570_rev | GGGGACCACTTTGTACAAGAAAGCTGGGTCTATAACGCTGCAACCGAAAGAAAC |
| 7_known | G4 | Tb927.2.1860_for | GGGGACAAGTTTGTACAAAAAAGCAGGCTATGCTGCTGCCCACTCTCG | Tb927.2.1860_rev | GGGGACCACTTTGTACAAGAAAGCTGGGTTCACACCAGCTGTAAAGGCGG |
| 7_known | G5 | Tb927.11.1510_for | GGGGACAAGTTTGTACAAAAAAGCAGGCTATGGGAACTACAATGTCAGGAAACC | Tb927.11.1510_rev | GGGGACCACTTTGTACAAGAAAGCTGGGTCTAGCTCGCGAAGCTTAGGC |
| 7_known | G6 | Tb927.1.3300_for | GGGGACAAGTTTGTACAAAAAAGCAGGCTATGTCGTTGGGTACGCCATGT | Tb927.1.3300_rev | GGGGACCACTTTGTACAAGAAAGCTGGGTCTATGCACTATTTAACTTGTGTGCACTATA |
| 7_known | G7 | Tb927.10.2040_for | GGGGACAAGTTTGTACAAAAAAGCAGGCTATGCGGGCTAACGACATTGTG | Tb927.10.2040_rev | GGGGACCACTTTGTACAAGAAAGCTGGGTTCATCCCACATTTGTCTGTGCC |
| 7_known | G8 | Tb927.4.4210_for | GGGGACAAGTTTGTACAAAAAAGCAGGCTATGCGCCGCTACTTGCTC | Tb927.4.4210_rev | GGGGACCACTTTGTACAAGAAAGCTGGGTTCACACGCCCTTCTTACGGT |
| 7_known | G9 | Tb927.10.3500_for | GGGGACAAGTTTGTACAAAAAAGCAGGCTATGGGGCGTGATAGTCGCG | Tb927.10.3500_rev | GGGGACCACTTTGTACAAGAAAGCTGGGTTTAACCGTCAATACCTGCGATACCG |
| 7_known | G10 | Tb927.10.9780_for | GGGGACAAGTTTGTACAAAAAAGCAGGCTATGTCACTGAAGAAGAAGACGCTAAAG | Tb927.10.9780_rev | GGGGACCACTTTGTACAAGAAAGCTGGGTCTACTCCTTCTCTGTATTTTTTCGGGC |
| 7_known | G11 | Tb927.11.11900_for | GGGGACAAGTTTGTACAAAAAAGCAGGCTATGCAACACGACCGTGACC | Tb927.11.11900_rev | GGGGACCACTTTGTACAAGAAAGCTGGGTCTACGACAGCAGCGCACC |
| 7_known | G12 | Tb927.9.4820_for | GGGGACAAGTTTGTACAAAAAAGCAGGCTATGTCGAAATCACGCGCGG | Tb927.9.4820_rev | GGGGACCACTTTGTACAAGAAAGCTGGGTTCAACTAGCCGGTAGAAGGTAGAAA |
| 7_known | G13 | Tb927.11.6370_for | GGGGACAAGTTTGTACAAAAAAGCAGGCTATGCTGTTTGTGACGCCGAC | Tb927.11.6370_rev | GGGGACCACTTTGTACAAGAAAGCTGGGTTTAAACCTGAGGAGCCATCAGTGC |
| 7_known | G14 | Tb927.10.5280_for | GGGGACAAGTTTGTACAAAAAAGCAGGCTATGGACAGGGAAGCCATGCG | Tb927.10.5280_rev | GGGGACCACTTTGTACAAGAAAGCTGGGTTCAAACGTGTTTCCGCTTGTGC |
| 7_known | G15 | Tb927.10.7060_for | GGGGACAAGTTTGTACAAAAAAGCAGGCTATGATGTCCATCCCAAACCTCACG | Tb927.10.7060_rev | GGGGACCACTTTGTACAAGAAAGCTGGGTTCACTGCACAAACAACCGCT |
| 7_known | G16 | Tb927.8.1100_for | GGGGACAAGTTTGTACAAAAAAGCAGGCTATGAGTGAGGACAGCGCAGC | Tb927.8.1100_rev | GGGGACCACTTTGTACAAGAAAGCTGGGTTTAGCGGGTGACCAAGAAAGGA |
| 7_known | G17 | Tb927.1.1350_for | GGGGACAAGTTTGTACAAAAAAGCAGGCTATGCAAACATCGCGACCCTC | Tb927.1.1350_rev | GGGGACCACTTTGTACAAGAAAGCTGGGTTTACGCAAAGGCGTGTTCCA |
| 7_known | G18 | Tb927.5.1870_for | GGGGACAAGTTTGTACAAAAAAGCAGGCTATGCTTTCATGTTTGCGTCAGATCC | Tb927.5.1870_rev | GGGGACCACTTTGTACAAGAAAGCTGGGTTCACCGTTTGATGGTTGCCTC |
| 7_known | G19 | Tb927.8.6840_for | GGGGACAAGTTTGTACAAAAAAGCAGGCTATGAGGGGTATTGAGCGGCT | Tb927.8.6840_rev | GGGGACCACTTTGTACAAGAAAGCTGGGTTCAGCAACGCTCGAAGACCT |
| 7_known | G20 | Tb927.8.8330_for | GGGGACAAGTTTGTACAAAAAAGCAGGCTATGGGTTGTGGTGGATCAAAGC | Tb927.8.8330_rev | GGGGACCACTTTGTACAAGAAAGCTGGGTCTAGTCAGTGTCGCTATCATCCGA |
| 7_known | G21 | Tb927.11.1680_for | GGGGACAAGTTTGTACAAAAAAGCAGGCTATGATAGTCAAGACCACAGTTAGCAAAA | Tb927.11.1680_rev | GGGGACCACTTTGTACAAGAAAGCTGGGTTCAGTACATTTCCTCCCCGTGAA |
| 7_known | G22 | Tb927.3.2170_for | GGGGACAAGTTTGTACAAAAAAGCAGGCTATGAATCTCGAAACGACTGAGCAC | Tb927.3.2170_rev | GGGGACCACTTTGTACAAGAAAGCTGGGTCTACGCCCCCTTCAATGAAAATTTC |
| 7_known | G23 | Tb927.7.6290_for | GGGGACAAGTTTGTACAAAAAAGCAGGCTATGCGGTCACAGGTGCGT | Tb927.7.6290_rev | GGGGACCACTTTGTACAAGAAAGCTGGGTTCACCGCGTGCTGGGAAT |
| 7_known | G24 | Tb927.10.3340_for | GGGGACAAGTTTGTACAAAAAAGCAGGCTATGTCCTCACAGGACAGTGAAAGC | Tb927.10.3340_rev | GGGGACCACTTTGTACAAGAAAGCTGGGTTCACATGATAATGCAGCAGAAGTTAGG |
| 7_known | H1 | Tb927.7.1070_for | GGGGACAAGTTTGTACAAAAAAGCAGGCTATGGCATTGGCTCAGTCATGTC | Tb927.7.1070_rev | GGGGACCACTTTGTACAAGAAAGCTGGGTTTACGGTAACTTCAATGAAAACGTTATACT |
| 7_known | H2 | Tb927.11.8290_for | GGGGACAAGTTTGTACAAAAAAGCAGGCTATGGATGAAACACCGAATGCTGT | Tb927.11.8290_rev | GGGGACCACTTTGTACAAGAAAGCTGGGTTCAAACGAGACTTGTGGGTATTGC |
| 7_known | H3 | Tb927.4.4160_for | GGGGACAAGTTTGTACAAAAAAGCAGGCTATGCGATGTCTTAGTCTTCTCCCC | Tb927.4.4160_rev | GGGGACCACTTTGTACAAGAAAGCTGGGTCTAATAAAGCACCTTCGCTAACATTTCC |
| 7_known | H4 | Tb927.3.1770_for | GGGGACAAGTTTGTACAAAAAAGCAGGCTATGTACACAACCGCAAATAGGTCAT | Tb927.3.1770_rev | GGGGACCACTTTGTACAAGAAAGCTGGGTTCACACAGGGGTTGTGTTCGA |
| 7_known | H5 | Tb927.10.14000_for | GGGGACAAGTTTGTACAAAAAAGCAGGCTATGCTCAGCACGATGAAGTTACTG | Tb927.10.14000_rev | GGGGACCACTTTGTACAAGAAAGCTGGGTCTACAAATTACCCTTGATTTTCTCGCG |
| 7_known | H6 | Tb927.10.11440_for | GGGGACAAGTTTGTACAAAAAAGCAGGCTATGACATTGTCTGTCGACACCG | Tb927.10.11440_rev | GGGGACCACTTTGTACAAGAAAGCTGGGTTTATTTCCTGGGAAGACGGTGGT |
| 7_known | H7 | Tb927.7.6680_for | GGGGACAAGTTTGTACAAAAAAGCAGGCTATGCCTGTCGTGAAGGGCG | Tb927.7.6680_rev | GGGGACCACTTTGTACAAGAAAGCTGGGTTCACTCATCGTCTGGTGGCA |
| 7_known | H8 | Tb927.11.9310_for | GGGGACAAGTTTGTACAAAAAAGCAGGCTATGAGGGTGGGGTGGACAAC | Tb927.11.9310_rev | GGGGACCACTTTGTACAAGAAAGCTGGGTTCACCACGTCACGTTCCACG |
| 7_known | H9 | Tb927.7.4110_for | GGGGACAAGTTTGTACAAAAAAGCAGGCTATGTTTGATAAGGTAACAGCAATGAGCG | Tb927.7.4110_rev | GGGGACCACTTTGTACAAGAAAGCTGGGTTCACTTCACTGTAGGAGATACTGGATTAC |
| 7_known | H10 | Tb927.3.3220_for | GGGGACAAGTTTGTACAAAAAAGCAGGCTATGCAATATATATCGGAACCCGTTCAAC | Tb927.3.3220_rev | GGGGACCACTTTGTACAAGAAAGCTGGGTTCATATATTCCCATCATTAGCGGTGTCA |
| 7_known | H11 | Tb927.10.10850_for | GGGGACAAGTTTGTACAAAAAAGCAGGCTATGTCTGACTGGGAACGTGGA | Tb927.10.10850_rev | GGGGACCACTTTGTACAAGAAAGCTGGGTTTATAGATAATGCATTGTTGTCCGCAAGT |
| 7_known | H12 | Tb927.3.5420_for | GGGGACAAGTTTGTACAAAAAAGCAGGCTATGAACTTCGCCGTACCTCCA | Tb927.3.5420_rev | GGGGACCACTTTGTACAAGAAAGCTGGGTTCACTTAGCGATCAGTGAAGACCT |
| 7_known | H13 | Tb927.7.4940_for | GGGGACAAGTTTGTACAAAAAAGCAGGCTATGACGGCACTCTATCCGTGG | Tb927.7.4940_rev | GGGGACCACTTTGTACAAGAAAGCTGGGTCTATTTTCTCCCGTTTTCCGAGGC |
| 7_known | H14 | Tb927.7.3980_for | GGGGACAAGTTTGTACAAAAAAGCAGGCTATGACCATCACTGTGGATCTATTTGGT | Tb927.7.3980_rev | GGGGACCACTTTGTACAAGAAAGCTGGGTCTATGGCAGTAACTTCTCCAAGTTCTTG |
| 7_known | H15 | Tb927.2.4550_for | GGGGACAAGTTTGTACAAAAAAGCAGGCTATGGGTACCAAATCTAAAAAGAAGGCTAAG | Tb927.2.4550_rev | GGGGACCACTTTGTACAAGAAAGCTGGGTTCACTTACGCCTCTTCGCACG |
| 7_known | H16 | Tb927.8.8170_for | GGGGACAAGTTTGTACAAAAAAGCAGGCTATGCGACGGCTAAGCCTTCT | Tb927.8.8170_rev | GGGGACCACTTTGTACAAGAAAGCTGGGTCTAACAAAATACCTTCGCTAACATCTTCTG |
| 7_known | H17 | Tb927.4.2080_for | GGGGACAAGTTTGTACAAAAAAGCAGGCTATGCAGCAGGTTGATTTTGGTCA | Tb927.4.2080_rev | GGGGACCACTTTGTACAAGAAAGCTGGGTCTAATGGGTATTCATAGCCATTAGAACATC |
| 7_known | H18 | Tb927.7.5230_for | GGGGACAAGTTTGTACAAAAAAGCAGGCTATGATGCTCTATCAGAGAAGGGGC | Tb927.7.5230_rev | GGGGACCACTTTGTACAAGAAAGCTGGGTCTAATCCATCGATCGGGGAAAATGT |
| 7_known | H19 | Tb927.10.3570_for | GGGGACAAGTTTGTACAAAAAAGCAGGCTATGTTGCGCCGCAGTCGC | Tb927.10.3570_rev | GGGGACCACTTTGTACAAGAAAGCTGGGTCTAAACCACCTGAAACTCAACAAGGA |
| 7_known | H20 | Tb927.3.3070_for | GGGGACAAGTTTGTACAAAAAAGCAGGCTATGGTGTGCTACGACGGTTCG | Tb927.3.3070_rev | GGGGACCACTTTGTACAAGAAAGCTGGGTTCAAGGACCCACCGACGC |
| 7_known | H21 | Tb927.4.1250_for | GGGGACAAGTTTGTACAAAAAAGCAGGCTATGCAGCACAGCTCCTTTGAAG | Tb927.4.1250_rev | GGGGACCACTTTGTACAAGAAAGCTGGGTTCACCGTTGGGTCAATTTCCTTC |
| 7_known | H22 | Tb927.3.5520_for | GGGGACAAGTTTGTACAAAAAAGCAGGCTATGTCGGACAACAGAAAAGCAGT | Tb927.3.5520_rev | GGGGACCACTTTGTACAAGAAAGCTGGGTCTACTCCATATTTGGTTTCTCTTCCACAA |
| 7_known | H23 | Tb927.10.12500_for | GGGGACAAGTTTGTACAAAAAAGCAGGCTATGGGGGATACTGGACCAAAGG | Tb927.10.12500_rev | GGGGACCACTTTGTACAAGAAAGCTGGGTTTAAATAGGTTCCTTAGACGACTTAGCATT |
| 7_known | H24 | Tb927.11.2380_for | GGGGACAAGTTTGTACAAAAAAGCAGGCTATGTCTCTTAGCGCCTTACGGT | Tb927.11.2380_rev | GGGGACCACTTTGTACAAGAAAGCTGGGTCTATTTTGTTACTCGGGGCAACCG |
| 7_known | I1 | Tb927.10.13220_for | GGGGACAAGTTTGTACAAAAAAGCAGGCTATGCAGGCACATCGGTCGA | Tb927.10.13220_rev | GGGGACCACTTTGTACAAGAAAGCTGGGTCTATTCCAACGATAACCCTTCTTCAGATAC |
| 7_known | I2 | Tb927.6.3580_for | GGGGACAAGTTTGTACAAAAAAGCAGGCTATGCAAAATAGTTATGAAGAGATTCTTTTC | Tb927.6.3580_rev | GGGGACCACTTTGTACAAGAAAGCTGGGTCTACGTTTCTTCCTTTTCGTCCAAGT |
| 7_known | I3 | Tb927.11.6280_for | GGGGACAAGTTTGTACAAAAAAGCAGGCTATGGTGGCTAAAAAGTGGGTTTACTAC | Tb927.11.6280_rev | GGGGACCACTTTGTACAAGAAAGCTGGGTTTAAAGCTTGGCCGCGAATCC |
| 7_known | I4 | Tb927.10.13020_for | GGGGACAAGTTTGTACAAAAAAGCAGGCTATGTCACGTCCAAGAAGGTTTGTG | Tb927.10.13020_rev | GGGGACCACTTTGTACAAGAAAGCTGGGTTCATCCGAATAATTTGGCCAATCTTACG |
| 7_known | I5 | Tb927.11.12180_for | GGGGACAAGTTTGTACAAAAAAGCAGGCTATGGAAAATATTCGCGTGGTGGT | Tb927.11.12180_rev | GGGGACCACTTTGTACAAGAAAGCTGGGTTCATGAAGTTTGTCTAAAAGAATCACTAAT |
| 7_known | I6 | Tb927.11.9520_for | GGGGACAAGTTTGTACAAAAAAGCAGGCTATGACAATATGGCTGAATGTTGCGG | Tb927.11.9520_rev | GGGGACCACTTTGTACAAGAAAGCTGGGTTCAGTCCAACCACTGAAAAAATTGGC |
| 7_known | I7 | Tb927.8.5710_for | GGGGACAAGTTTGTACAAAAAAGCAGGCTATGCACATTATTGAAGTGAACCACGG | Tb927.8.5710_rev | GGGGACCACTTTGTACAAGAAAGCTGGGTTTAGACGACCAATGGTCGGACA |
| 7_known | I8 | Tb927.11.10650_for | GGGGACAAGTTTGTACAAAAAAGCAGGCTATGAACCGCGCTATCATTGCA | Tb927.11.10650_rev | GGGGACCACTTTGTACAAGAAAGCTGGGTTCACTCCGCGGGACCACT |
| 7_known | I9 | Tb927.11.16690_for | GGGGACAAGTTTGTACAAAAAAGCAGGCTATGTCCAACAAGTTGTTGCCATCG | Tb927.11.16690_rev | GGGGACCACTTTGTACAAGAAAGCTGGGTCTAGGCCCTCGGCACGGC |
| 7_known | I10 | Tb927.10.12510_for | GGGGACAAGTTTGTACAAAAAAGCAGGCTATGGGGGATACTGGACCAAAGG | Tb927.10.12510_rev | GGGGACCACTTTGTACAAGAAAGCTGGGTTTATAGCGTAGGTTCTTTAGGTGCACC |
| 7_known | I11 | Tb927.10.12310_for | GGGGACAAGTTTGTACAAAAAAGCAGGCTATGGTCCGCCTGACCGTG | Tb927.10.12310_rev | GGGGACCACTTTGTACAAGAAAGCTGGGTTCATTTCATGTTTTGTCGATGGGCC |
| 7_known | I12 | Tb927.1.4050_for | GGGGACAAGTTTGTACAAAAAAGCAGGCTATGGGTTGCTCGACCTCCAA | Tb927.1.4050_rev | GGGGACCACTTTGTACAAGAAAGCTGGGTTCAATCTCTCACAAAGGAGGAAAAGC |
| 7_known | I13 | Tb927.5.490_for | GGGGACAAGTTTGTACAAAAAAGCAGGCTATGTTATTTTTTTGTTGGTTTGCTTGTTCC | Tb927.5.490_rev | GGGGACCACTTTGTACAAGAAAGCTGGGTTTATTCCTTTTCATTTTGCGGTGCTG |
| 7_known | I14 | Tb927.9.5100_for | GGGGACAAGTTTGTACAAAAAAGCAGGCTATGACACACAACGGTGGTCG | Tb927.9.5100_rev | GGGGACCACTTTGTACAAGAAAGCTGGGTCTAAGACGAAGCCCCTGTACTCC |
| 7_known | I15 | Tb927.8.3560_for | GGGGACAAGTTTGTACAAAAAAGCAGGCTATGCTCTCCCGTCTAAGCGC | Tb927.8.3560_rev | GGGGACCACTTTGTACAAGAAAGCTGGGTTTATTCTTCCACTTCCCCTTCATACTCC |
| 7_known | I16 | Tb927.3.5650_for | GGGGACAAGTTTGTACAAAAAAGCAGGCTATGATAGAATATGCGTGTGGTTGGC | Tb927.3.5650_rev | GGGGACCACTTTGTACAAGAAAGCTGGGTCTACAAATGAAATTCGCTTCTCGAGATATT |
| 7_known | I17 | Tb927.11.6890_for | GGGGACAAGTTTGTACAAAAAAGCAGGCTATGTTGCGTCGGCTCCTCC | Tb927.11.6890_rev | GGGGACCACTTTGTACAAGAAAGCTGGGTCTATTCCAACGACTCACCAGAGGA |
| 7_known | I18 | Tb927.9.5040_for | GGGGACAAGTTTGTACAAAAAAGCAGGCTATGTTCATGAACAAGCCCTTTGGC | Tb927.9.5040_rev | GGGGACCACTTTGTACAAGAAAGCTGGGTTCAACGAGTACTGCTGTTGTTGC |
| 7_known | I19 | Tb927.10.2270_for | GGGGACAAGTTTGTACAAAAAAGCAGGCTATGTACGGCCATGTTGTGGTG | Tb927.10.2270_rev | GGGGACCACTTTGTACAAGAAAGCTGGGTCTACTCATTATCAGCGGCCTCGC |
| 7_known | I20 | Tb927.8.8180_for | GGGGACAAGTTTGTACAAAAAAGCAGGCTATGTACCGTCTTTATCGTCGGACT | Tb927.8.8180_rev | GGGGACCACTTTGTACAAGAAAGCTGGGTCTAAACTGCTGTGGCCAGTTTATTCT |
| 7_known | I22 | Tb927.9.5240_for | GGGGACAAGTTTGTACAAAAAAGCAGGCTATGAGCAAACGCCGCCGT | Tb927.9.5240_rev | GGGGACCACTTTGTACAAGAAAGCTGGGTCTAAGAATTGTTTTTGACGAAGCGCC |
| 7_known | I23 | Tb927.9.8540_for | GGGGACAAGTTTGTACAAAAAAGCAGGCTATGTATACCAGCATGTCCATCCAAGG | Tb927.9.8540_rev | GGGGACCACTTTGTACAAGAAAGCTGGGTTTAAGGACAAGAGTTGATCGCAGTG |
| 7_known | I24 | Tb927.3.5100_for | GGGGACAAGTTTGTACAAAAAAGCAGGCTATGATTCTTGGCCCTGGTGGG | Tb927.3.5100_rev | GGGGACCACTTTGTACAAGAAAGCTGGGTCTAGGTAGTCAGCGGAATTCGGT |
| 7_known | J1 | Tb927.8.4260_for | GGGGACAAGTTTGTACAAAAAAGCAGGCTATGAGTGAAATCAGACGACCACAGG | Tb927.8.4260_rev | GGGGACCACTTTGTACAAGAAAGCTGGGTCTAAGCGGTAAAGGAAGGAGTAACAG |
| 7_known | J2 | Tb927.10.13270_for | GGGGACAAGTTTGTACAAAAAAGCAGGCTATGGAGACAAACTTTCAGCTGGC | Tb927.10.13270_rev | GGGGACCACTTTGTACAAGAAAGCTGGGTTCATTCAGTAACTTCCGCGGC |
| 7_known | J3 | Tb927.10.3360_for | GGGGACAAGTTTGTACAAAAAAGCAGGCTATGCCGCGGTTGTTGTTACC | Tb927.10.3360_rev | GGGGACCACTTTGTACAAGAAAGCTGGGTCTATTCAACAAAATGCCACGATGTGG |
| 7_known | J4 | Tb927.10.7630_for | GGGGACAAGTTTGTACAAAAAAGCAGGCTATGAGTTTTGCGGGAGGTTATTCC | Tb927.10.7630_rev | GGGGACCACTTTGTACAAGAAAGCTGGGTTCAACAAATGAGTCCATATGAAGATATCAC |
| 7_known | J5 | Tb927.5.2300_for | GGGGACAAGTTTGTACAAAAAAGCAGGCTATGCAACGTGCCCAGATTCTC | Tb927.5.2300_rev | GGGGACCACTTTGTACAAGAAAGCTGGGTTTAACTTCTAATTGCACGACGGGG |
| 7_known | J6 | Tb927.1.4420_for | GGGGACAAGTTTGTACAAAAAAGCAGGCTATGACGACAACTCAAGGTGAAGTAATC | Tb927.1.4420_rev | GGGGACCACTTTGTACAAGAAAGCTGGGTTCACAATGTTACGGTCTCATCCGC |
| 7_known | J7 | Tb927.11.7218_for | GGGGACAAGTTTGTACAAAAAAGCAGGCTATGACATCTAGCGTAACCGGACC | Tb927.11.7218_rev | GGGGACCACTTTGTACAAGAAAGCTGGGTCTATTTTTTCTTCCCCTTGTTCTTTCCTTT |
| 7_known | J8 | Tb927.3.1290_for | GGGGACAAGTTTGTACAAAAAAGCAGGCTATGTCATCACGGACGCTGTATG | Tb927.3.1290_rev | GGGGACCACTTTGTACAAGAAAGCTGGGTCTACGAAATATATATATATTCATTTGTATT |
| 7_known | J9 | Tb927.11.7860_for | GGGGACAAGTTTGTACAAAAAAGCAGGCTATGCCTCCTCACCGCAAGC | Tb927.11.7860_rev | GGGGACCACTTTGTACAAGAAAGCTGGGTTTACACCAGGGAGTGCTCTATTTGA |
| 7_known | J10 | Tb927.5.3600_for | GGGGACAAGTTTGTACAAAAAAGCAGGCTATGCCTAAACGTGAAAGGGAAGAC | Tb927.5.3600_rev | GGGGACCACTTTGTACAAGAAAGCTGGGTCTAAAGCTGTGCCCCAACCG |
| 7_known | J11 | Tb927.11.860_for | GGGGACAAGTTTGTACAAAAAAGCAGGCTATGCACGCTGGCACTCCA | Tb927.11.860_rev | GGGGACCACTTTGTACAAGAAAGCTGGGTCTAGGCCCTTGTTGCACCTTC |
| 7_known | J12 | Tb927.11.460_for | GGGGACAAGTTTGTACAAAAAAGCAGGCTATGGAAGGAAAGAACTGCTTCCG | Tb927.11.460_rev | GGGGACCACTTTGTACAAGAAAGCTGGGTTCAAGTCTTCTCACGGCGCT |
| 7_known | J13 | Tb927.3.2140_for | GGGGACAAGTTTGTACAAAAAAGCAGGCTATGAAGTTTGAGAACCCCGTTGC | Tb927.3.2140_rev | GGGGACCACTTTGTACAAGAAAGCTGGGTCTAGTGAGATTCACTGAAACTGCCG |
| 7_known | J14 | Tb927.3.1230_for | GGGGACAAGTTTGTACAAAAAAGCAGGCTATGATTTTCGGGGAAGATATTGGACC | Tb927.3.1230_rev | GGGGACCACTTTGTACAAGAAAGCTGGGTTCACTGAGTGGTAATATTCAACGCATC |
| 7_known | J15 | Tb927.8.6210_for | GGGGACAAGTTTGTACAAAAAAGCAGGCTATGGCCACGAACGAGGACA | Tb927.8.6210_rev | GGGGACCACTTTGTACAAGAAAGCTGGGTTTAGGCCCGCCGCGCCTG |
| 7_known | J16 | Tb927.10.7440_for | GGGGACAAGTTTGTACAAAAAAGCAGGCTATGCAGGATGACATACCCGTGG | Tb927.10.7440_rev | GGGGACCACTTTGTACAAGAAAGCTGGGTCTACAGGTACAGCGACGACGC |
| 7_known | J17 | Tb927.10.11020_for | GGGGACAAGTTTGTACAAAAAAGCAGGCTATGAGTGACGACCGCGACC | Tb927.10.11020_rev | GGGGACCACTTTGTACAAGAAAGCTGGGTTCACACCTCCACCAATGACGA |
| 7_known | J18 | Tb927.11.7211_for | GGGGACAAGTTTGTACAAAAAAGCAGGCTATGGAGCAGCTACTTCGAAGGG | Tb927.11.7211_rev | GGGGACCACTTTGTACAAGAAAGCTGGGTCTAGACGGTGGGGCACGC |
| 7_known | J19 | Tb927.10.5450_for | GGGGACAAGTTTGTACAAAAAAGCAGGCTATGTCACAGAAAGAGGAGAATAGCGT | Tb927.10.5450_rev | GGGGACCACTTTGTACAAGAAAGCTGGGTTCACAACTCGAGCCTTAGCTTGT |
| 7_known | J20 | Tb927.4.2950_for | GGGGACAAGTTTGTACAAAAAAGCAGGCTATGTCGTTACATGGTCATCCGTACC | Tb927.4.2950_rev | GGGGACCACTTTGTACAAGAAAGCTGGGTTCAACTCGGCACCCGCAG |
| 7_known | J21 | Tb927.6.4740_for | GGGGACAAGTTTGTACAAAAAAGCAGGCTATGGATCCAATCAACCCGAATGATC | Tb927.6.4740_rev | GGGGACCACTTTGTACAAGAAAGCTGGGTTCACTGCAATTGCGCCATTACC |
| 7_known | J22 | Tb927.11.1260_for | GGGGACAAGTTTGTACAAAAAAGCAGGCTATGGAGGAGAACAACGCAACTACA | Tb927.11.1260_rev | GGGGACCACTTTGTACAAGAAAGCTGGGTCTAAATGATATCAATGAACTGTTCCCGTTT |
| 7_known | J23 | Tb927.3.3090_for | GGGGACAAGTTTGTACAAAAAAGCAGGCTATGTCGTCTGTATTTATTGACGGCG | Tb927.3.3090_rev | GGGGACCACTTTGTACAAGAAAGCTGGGTTCAGGAAACCCTTGGTCGCT |
| 7_known | J24 | Tb927.11.9330_for | GGGGACAAGTTTGTACAAAAAAGCAGGCTATGACAGAGGACGAGGAACCATATG | Tb927.11.9330_rev | GGGGACCACTTTGTACAAGAAAGCTGGGTTCACTGCGGAAACCCGTCC |
| 7_known | K1 | Tb927.6.4600_for | GGGGACAAGTTTGTACAAAAAAGCAGGCTATGACTAAGAAGAATGTTATTCGCTTTACT | Tb927.6.4600_rev | GGGGACCACTTTGTACAAGAAAGCTGGGTTTACTCGACAGTGACAGCGCA |
| 7_known | K2 | Tb927.6.700_for | GGGGACAAGTTTGTACAAAAAAGCAGGCTATGACAAAAGCGGTGACTGAATGG | Tb927.6.700_rev | GGGGACCACTTTGTACAAGAAAGCTGGGTTCAGTTCAAAAGCACTGCCATCA |
| 7_known | K3 | Tb927.7.7160_for | GGGGACAAGTTTGTACAAAAAAGCAGGCTATGGAAAGAGAAGCAACAAAACGGG | Tb927.7.7160_rev | GGGGACCACTTTGTACAAGAAAGCTGGGTCTAATGCGGCGAGGCGCA |
| 7_known | K4 | Tb927.11.5260_for | GGGGACAAGTTTGTACAAAAAAGCAGGCTATGAATTGCAACTGTGGCACACC | Tb927.11.5260_rev | GGGGACCACTTTGTACAAGAAAGCTGGGTTCACTGAGTCTCACCGGTTTCA |
| 7_known | K5 | Tb927.8.3660_for | GGGGACAAGTTTGTACAAAAAAGCAGGCTATGCAAGGACACGTACCACAGC | Tb927.8.3660_rev | GGGGACCACTTTGTACAAGAAAGCTGGGTCTACTTTTTCGACTCTGGCGTAACA |
| 7_known | K6 | Tb927.1.1100_for | GGGGACAAGTTTGTACAAAAAAGCAGGCTATGCCGAAGTCTTCCTCAGCC | Tb927.1.1100_rev | GGGGACCACTTTGTACAAGAAAGCTGGGTTTAGTCCAACACCCCATTACCATGA |
| 7_known | K7 | Tb927.7.1910_for | GGGGACAAGTTTGTACAAAAAAGCAGGCTATGTATCGCAGTGCGTTTCGC | Tb927.7.1910_rev | GGGGACCACTTTGTACAAGAAAGCTGGGTCTATTTGAATGAGTCCAACGACATACAGT |
| 7_known | K8 | Tb11.02.5380_for | GGGGACAAGTTTGTACAAAAAAGCAGGCTATGTTTACGAAAAAACGACTGGCTG | Tb11.02.5380_rev | GGGGACCACTTTGTACAAGAAAGCTGGGTTTACTGTTGCCCCTCCTCCAG |
| 7_known | K9 | Tb927.9.6430_for | GGGGACAAGTTTGTACAAAAAAGCAGGCTATGAAGTCATCTGATCATGACGAAGAGG | Tb927.9.6430_rev | GGGGACCACTTTGTACAAGAAAGCTGGGTTCAGGAACTTACTGAGGAAACGGG |
| 7_known | K10 | Tb927.3.1210_for | GGGGACAAGTTTGTACAAAAAAGCAGGCTATGTCGTACCCACACAACTTTGAC | Tb927.3.1210_rev | GGGGACCACTTTGTACAAGAAAGCTGGGTTCATAGCGCCAACTCGCCC |
| 7_known | K11 | Tb927.10.5180_for | GGGGACAAGTTTGTACAAAAAAGCAGGCTATGAGTGTAAGGCACCTGTTTGG | Tb927.10.5180_rev | GGGGACCACTTTGTACAAGAAAGCTGGGTTCAACGATTGCCGATCACCTG |
| 7_known | K12 | Tb927.11.15430_for | GGGGACAAGTTTGTACAAAAAAGCAGGCTATGAACTTTGGTTATGATGAAGAGGGG | Tb927.11.15430_rev | GGGGACCACTTTGTACAAGAAAGCTGGGTTCACATCTGGAACTCTGCGAGA |
| 7_known | K13 | Tb927.6.4480_for | GGGGACAAGTTTGTACAAAAAAGCAGGCTATGAAACAACTTGCTCCCCAGATG | Tb927.6.4480_rev | GGGGACCACTTTGTACAAGAAAGCTGGGTTTAGAGTAAACTCTTCATCTTTGTTAAACC |
| 7_known | K14 | Tb927.7.3950_for | GGGGACAAGTTTGTACAAAAAAGCAGGCTATGGTAAGTAAGTACCACCGCTTGC | Tb927.7.3950_rev | GGGGACCACTTTGTACAAGAAAGCTGGGTTCAACGAGGGCTCTTGAATGGA |
| 7_known | K15 | Tb927.11.9220_for | GGGGACAAGTTTGTACAAAAAAGCAGGCTATGGCCGTCGGACTCTCATC | Tb927.11.9220_rev | GGGGACCACTTTGTACAAGAAAGCTGGGTCTAGGGCCATGTAAACGGTTCTG |
| 7_known | K16 | Tb927.11.6810_for | GGGGACAAGTTTGTACAAAAAAGCAGGCTATGAGGAACATGATGGAGCCAGT | Tb927.11.6810_rev | GGGGACCACTTTGTACAAGAAAGCTGGGTCTACTCATCCAAATGGAATGAATCTTCTCT |
| 7_known | K17 | Tb927.5.3920_for | GGGGACAAGTTTGTACAAAAAAGCAGGCTATGATGTCACGGACCGTTGAGC | Tb927.5.3920_rev | GGGGACCACTTTGTACAAGAAAGCTGGGTTCAATCAGCCACACGACCCG |
| 7_known | K18 | Tb927.1.2570_for | GGGGACAAGTTTGTACAAAAAAGCAGGCTATGGAGGCCAAAGCTGAGGG | Tb927.1.2570_rev | GGGGACCACTTTGTACAAGAAAGCTGGGTTTATGACACTATACGTCGATTCAGTATATT |
| 7_known | K19 | Tb927.11.330_for | GGGGACAAGTTTGTACAAAAAAGCAGGCTATGTTCCCTGGGTCAAAAAGGTCG | Tb927.11.330_rev | GGGGACCACTTTGTACAAGAAAGCTGGGTTCAACTGGACTGGGATTCTCCTTC |
| 7_known | K20 | Tb927.11.750_for | GGGGACAAGTTTGTACAAAAAAGCAGGCTATGACAAGCAAACATAGTGCTGTATTAAC | Tb927.11.750_rev | GGGGACCACTTTGTACAAGAAAGCTGGGTCTACCTCGTGAACAGAAACTGAAGATC |
| 7_known | K21 | Tb927.10.11680_for | GGGGACAAGTTTGTACAAAAAAGCAGGCTATGAAGGGGCAGTCTTCACAGT | Tb927.10.11680_rev | GGGGACCACTTTGTACAAGAAAGCTGGGTTTAGAAAGTAGCATCGGGTTTCCAAG |
| 7_known | K22 | Tb927.5.2400_for | GGGGACAAGTTTGTACAAAAAAGCAGGCTATGCATCACGGTCGCAAAGT | Tb927.5.2400_rev | GGGGACCACTTTGTACAAGAAAGCTGGGTTCACTCGCTTCCCCGTGG |
| 7_known | K23 | Tb927.11.6400_for | GGGGACAAGTTTGTACAAAAAAGCAGGCTATGGGCGGAGAAGATAACGTCG | Tb927.11.6400_rev | GGGGACCACTTTGTACAAGAAAGCTGGGTTCAGGTGGAGACGTTGAGTCC |
| 7_known | K24 | Tb927.10.1380_for | GGGGACAAGTTTGTACAAAAAAGCAGGCTATGACAAGCCGAGCCCCT | Tb927.10.1380_rev | GGGGACCACTTTGTACAAGAAAGCTGGGTTTATGACATAGCCATCCATAGTTGTAGCC |
| 7_known | L1 | Tb927.6.920_for | GGGGACAAGTTTGTACAAAAAAGCAGGCTATGCATCGCTTTGTGTTGGTGC | Tb927.6.920_rev | GGGGACCACTTTGTACAAGAAAGCTGGGTTCATGAGGTATGGGGGAAGTATGAG |
| 7_known | L2 | Tb927.3.3180_for | GGGGACAAGTTTGTACAAAAAAGCAGGCTATGCGTGAGCGGCCGCGT | Tb927.3.3180_rev | GGGGACCACTTTGTACAAGAAAGCTGGGTTTAAGCATTCCCAGAAGAGGCACC |
| 7_known | L3 | Tb927.8.700_for | GGGGACAAGTTTGTACAAAAAAGCAGGCTATGTGGGATGGGCCGCTAC | Tb927.8.700_rev | GGGGACCACTTTGTACAAGAAAGCTGGGTTCAGTTCACCCTTCGGGCG |
| 7_known | L4 | Tb927.9.5900_for | GGGGACAAGTTTGTACAAAAAAGCAGGCTATGATGCGCCGTGCTTCTT | Tb927.9.5900_rev | GGGGACCACTTTGTACAAGAAAGCTGGGTTTAAGAGATGTGCTTGGCCTGC |
| 7_known | L5 | Tb927.10.6410_for | GGGGACAAGTTTGTACAAAAAAGCAGGCTATGCCTGAGGTTTCGCTTGAATG | Tb927.10.6410_rev | GGGGACCACTTTGTACAAGAAAGCTGGGTCTACCGATGCGAGAACTGTGTC |
| 7_known | L6 | Tb927.10.2570_for | GGGGACAAGTTTGTACAAAAAAGCAGGCTATGCTCACACTTGGGATGCATAC | Tb927.10.2570_rev | GGGGACCACTTTGTACAAGAAAGCTGGGTTTACACACCCGTGCAGCTCA |
| 7_known | L7 | Tb927.9.5460_for | GGGGACAAGTTTGTACAAAAAAGCAGGCTATGGACTCACCTGAAGCATGCC | Tb927.9.5460_rev | GGGGACCACTTTGTACAAGAAAGCTGGGTCTAAACCCGCCTCACAGCG |
| 7_known | L8 | Tb927.6.2460_for | GGGGACAAGTTTGTACAAAAAAGCAGGCTATGATTGAGGTAAGCAGTTTGGAGC | Tb927.6.2460_rev | GGGGACCACTTTGTACAAGAAAGCTGGGTTCATTCACGGTGCGCCTTC |
| 7_known | L9 | Tb927.11.17000_for | GGGGACAAGTTTGTACAAAAAAGCAGGCTATGAGTGCGTCTGCGATAAGGC | Tb927.11.17000_rev | GGGGACCACTTTGTACAAGAAAGCTGGGTCTATGCGCCGTCAGTCCG |
| 7_known | L10 | Tb927.4.2140_for | GGGGACAAGTTTGTACAAAAAAGCAGGCTATGGCAGATACCTTGCAACTTCTG | Tb927.4.2140_rev | GGGGACCACTTTGTACAAGAAAGCTGGGTTCACTGCGTCAGCGAAAAGTG |

| | | | | | |
|---|---|---|---|---|---|
| 7_known | L11 | Tb927.11.15990_for | GGGGACAAGTTTGTACAAAAAAGCAGGCTATGGCGGAGGTGCGGCGT | Tb927.11.15990_rev | GGGGACCACTTTGTACAAGAAAGCTGGGTTCATTTCCTCACCCTGTCATGATACT |
| 7_known | L12 | Tb927.2.3180_for | GGGGACAAGTTTGTACAAAAAAGCAGGCTATGTTCCAACGCCGGTTAGTG | Tb927.2.3180_rev | GGGGACCACTTTGTACAAGAAAGCTGGGTCTAAACATCCGCACCGGCA |
| 7_known | L13 | Tb927.10.5820_for | GGGGACAAGTTTGTACAAAAAAGCAGGCTATGTGGCGCCCACCAATCC | Tb927.10.5820_rev | GGGGACCACTTTGTACAAGAAAGCTGGGTCTATTTGTCGCATTTGGCAATGCC |
| 7_known | L14 | Tb927.9.2350_for | GGGGACAAGTTTGTACAAAAAAGCAGGCTATGGAGGAGTATCTGACTGTCAACAT | Tb927.9.2350_rev | GGGGACCACTTTGTACAAGAAAGCTGGGTTCACCACCTACGGTGGGC |
| 7_known | L15 | Tb927.10.2990_for | GGGGACAAGTTTGTACAAAAAAGCAGGCTATGTATGCAATGCCCGGTAATAGTG | Tb927.10.2990_rev | GGGGACCACTTTGTACAAGAAAGCTGGGTCTATACACTTACTTCGTATCGACCGC |
| 7_known | L16 | Tb927.11.5740_for | GGGGACAAGTTTGTACAAAAAAGCAGGCTATGCGTTTTTTTAGCGGGTGGT | Tb927.11.5740_rev | GGGGACCACTTTGTACAAGAAAGCTGGGTTCACTCTTTCTTGGAAGAGTCTTCAGG |
| 7_known | L17 | Tb927.11.9980_for | GGGGACAAGTTTGTACAAAAAAGCAGGCTATGATGCGAAGGCTCAGTCCT | Tb927.11.9980_rev | GGGGACCACTTTGTACAAGAAAGCTGGGTTTATGAAAATACGCATTCGCAAATGTACC |
| 7_known | L18 | Tb927.2.6130_for | GGGGACAAGTTTGTACAAAAAAGCAGGCTATGCACGCGAGAATAAATATACGTAACG | Tb927.2.6130_rev | GGGGACCACTTTGTACAAGAAAGCTGGGTTTATTTCACAAAGTGGGGCAATTGC |
| 7_known | L19 | Tb927.11.1450_for | GGGGACAAGTTTGTACAAAAAAGCAGGCTATGATGCGCCGTGCCTTT | Tb927.11.1450_rev | GGGGACCACTTTGTACAAGAAAGCTGGGTTTACGCCAGCGCCTGTCT |
| 7_known | L20 | Tb927.4.2500_for | GGGGACAAGTTTGTACAAAAAAGCAGGCTATGCCGGATGCCTATGTTTTTGC | Tb927.4.2500_rev | GGGGACCACTTTGTACAAGAAAGCTGGGTTCAGTTGGCTGACGGCAAAC |
| 7_known | L21 | Tb927.10.3800_for | GGGGACAAGTTTGTACAAAAAAGCAGGCTATGACCACAGAGCTTAACAGTTTTCG | Tb927.10.3800_rev | GGGGACCACTTTGTACAAGAAAGCTGGGTTCACTGATCAGAGCCTACACAAGA |
| 7_known | L22 | Tb927.11.15690_for | GGGGACAAGTTTGTACAAAAAAGCAGGCTATGAGCGCTGCGGCACCT | Tb927.11.15690_rev | GGGGACCACTTTGTACAAGAAAGCTGGGTTTACAATTTGATCATTTCCACGACTTGC |
| 7_known | L23 | Tb927.6.3310_for | GGGGACAAGTTTGTACAAAAAAGCAGGCTATGGTTGTGAGTCTTTCCGGTTATTC | Tb927.6.3310_rev | GGGGACCACTTTGTACAAGAAAGCTGGGTTTAAACACTCGACAACACTCTAACAGAA |
| 7_known | L24 | Tb927.8.1510_for | GGGGACAAGTTTGTACAAAAAAGCAGGCTATGTATGGGATGAACAGAGGAGGAG | Tb927.8.1510_rev | GGGGACCACTTTGTACAAGAAAGCTGGGTCTAATAAGAACCACCCGTCGGTTG |
| 7_known | M1 | Tb927.5.3400_for | GGGGACAAGTTTGTACAAAAAAGCAGGCTATGCTACCCGAGAACCTTCCG | Tb927.5.3400_rev | GGGGACCACTTTGTACAAGAAAGCTGGGTCTAGTCCTTCTTCTTTTCCTGGGCC |
| 7_known | M2 | Tb927.10.14460_for | GGGGACAAGTTTGTACAAAAAAGCAGGCTATGGAAGCGACTACAGATCCACC | Tb927.10.14460_rev | GGGGACCACTTTGTACAAGAAAGCTGGGTTCATATCCCACTGGGTTTGCTTCC |
| 7_known | M3 | Tb927.9.3120_for | GGGGACAAGTTTGTACAAAAAAGCAGGCTATGTTGCCTCACCCCATTTTAGC | Tb927.9.3120_rev | GGGGACCACTTTGTACAAGAAAGCTGGGTCTACGAATGGGGTTGTGTGGTG |
| 7_known | M4 | Tb927.8.4840_for | GGGGACAAGTTTGTACAAAAAAGCAGGCTATGGTAAATAGTTCCGCTTTGGTTAGTG | Tb927.8.4840_rev | GGGGACCACTTTGTACAAGAAAGCTGGGTTCAATAGGGTCCCAAAGTCGATGC |
| 7_known | M5 | Tb927.7.7300_for | GGGGACAAGTTTGTACAAAAAAGCAGGCTATGAACGCATGCAACAATAATACTGG | Tb927.7.7300_rev | GGGGACCACTTTGTACAAGAAAGCTGGGTTTAAGTTGTGATGGTGCCATGCG |
| 7_known | M6 | Tb927.3.2710_for | GGGGACAAGTTTGTACAAAAAAGCAGGCTATGAATCCACAAAATCACGTTTGTTGC | Tb927.3.2710_rev | GGGGACCACTTTGTACAAGAAAGCTGGGTTCATTCGCTCGGCTGCAGT |
| 7_known | M7 | Tb927.6.2980_for | GGGGACAAGTTTGTACAAAAAAGCAGGCTATGGCCAGCGAGTGCTCC | Tb927.6.2980_rev | GGGGACCACTTTGTACAAGAAAGCTGGGTTCACACCTCAGAAAAAAAGGAGAGC |
| 7_known | M8 | Tb927.6.2960_for | GGGGACAAGTTTGTACAAAAAAGCAGGCTATGTCTAAGCTTTACAAGGCTCAGAC | Tb927.6.2960_rev | GGGGACCACTTTGTACAAGAAAGCTGGGTCTACGACAATGGTATCGCGCTG |
| 7_known | M9 | Tb927.9.2140_for | GGGGACAAGTTTGTACAAAAAAGCAGGCTATGGCACGGGATAGCGGT | Tb927.9.2140_rev | GGGGACCACTTTGTACAAGAAAGCTGGGTTTACTCCGACATTATCTTTTCGTACAGAAA |
| 7_known | M10 | Tb927.2.1800_for | GGGGACAAGTTTGTACAAAAAAGCAGGCTATGACTGATTCACCTGATCTCTCTCG | Tb927.2.1800_rev | GGGGACCACTTTGTACAAGAAAGCTGGGTTCACCAATCGTCCACTACACCG |
| 7_known | M11 | Tb927.7.5600_for | GGGGACAAGTTTGTACAAAAAAGCAGGCTATGGGGTGTAACCTTTCCAGCT | Tb927.7.5600_rev | GGGGACCACTTTGTACAAGAAAGCTGGGTCTAATAAATACCCTTCTGACTATCTGCACA |
| 7_known | M12 | Tb927.8.7260_for | GGGGACAAGTTTGTACAAAAAAGCAGGCTATGACCGCTCGACGACGC | Tb927.8.7260_rev | GGGGACCACTTTGTACAAGAAAGCTGGGTTCAGTGCGGTTTTCTAAACACAAAC |
| 7_known | M13 | Tb927.5.4300_for | GGGGACAAGTTTGTACAAAAAAGCAGGCTATGTCAGCCTCCTCAGAGCATAG | Tb927.5.4300_rev | GGGGACCACTTTGTACAAGAAAGCTGGGTCTATGTGCTAGACTGCTCAGCAGC |
| 7_known | M14 | Tb927.9.9450_for | GGGGACAAGTTTGTACAAAAAAGCAGGCTATGTATTCTAGCGAAAAGGAGAAGGAAG | Tb927.9.9450_rev | GGGGACCACTTTGTACAAGAAAGCTGGGTTCACGAACGACCCGGATGC |
| 7_known | M15 | Tb927.5.2820_for | GGGGACAAGTTTGTACAAAAAAGCAGGCTATGAATGCCTTGGATGAGCTTAACG | Tb927.5.2820_rev | GGGGACCACTTTGTACAAGAAAGCTGGGTTCACTGTGCCGTAGCTCCA |
| 7_known | M16 | Tb927.8.1860_for | GGGGACAAGTTTGTACAAAAAAGCAGGCTATGATTCGACGTTGTGCTGTTCC | Tb927.8.1860_rev | GGGGACCACTTTGTACAAGAAAGCTGGGTCTACTCTTTCCGCTCCTCGGG |
| 7_known | M17 | Tb927.9.4560_for | GGGGACAAGTTTGTACAAAAAAGCAGGCTATGTATTTCCCGCGTCACCAAC | Tb927.9.4560_rev | GGGGACCACTTTGTACAAGAAAGCTGGGTTTAGAACAACCGAATACCCGCTAAATC |
| 7_known | M18 | Tb927.6.820_for | GGGGACAAGTTTGTACAAAAAAGCAGGCTATGACAACCATGGAGGAAGAAAACAT | Tb927.6.820_rev | GGGGACCACTTTGTACAAGAAAGCTGGGTTCATCCCTTCCTGCCGCG |
| 7_known | M19 | Tb927.11.14340_for | GGGGACAAGTTTGTACAAAAAAGCAGGCTATGGAAGATATTCTCGACTTCAGCAAG | Tb927.11.14340_rev | GGGGACCACTTTGTACAAGAAAGCTGGGTTCAGGGATAAAAAGAAGGTGTTGGTG |
| 7_known | M20 | Tb927.11.5430_for | GGGGACAAGTTTGTACAAAAAAGCAGGCTATGCCAGTTTTTCGTAATTCCACTGG | Tb927.11.5430_rev | GGGGACCACTTTGTACAAGAAAGCTGGGTCTAATCGTCACCAGAGGGATAATCATCA |
| 7_known | M21 | Tb927.2.4200_for | GGGGACAAGTTTGTACAAAAAAGCAGGCTATGTCACCCTCGAAGGGTGAA | Tb927.2.4200_rev | GGGGACCACTTTGTACAAGAAAGCTGGGTTTACTTTCTTTTTCCATGCGCTTCAG |
| 7_known | M22 | Tb927.9.15460_for | GGGGACAAGTTTGTACAAAAAAGCAGGCTATGACGGGCTCAGGTGGC | Tb927.9.15460_rev | GGGGACCACTTTGTACAAGAAAGCTGGGTTTACATCTTTCCTCCTTTCCCCCC |
| 7_known | M23 | Tb927.7.7260_for | GGGGACAAGTTTGTACAAAAAAGCAGGCTATGGGGAAGTCGGAGGGTATTC | Tb927.7.7260_rev | GGGGACCACTTTGTACAAGAAAGCTGGGTCTATTGAGGGGGGTCCAGCTG |
| 7_known | M24 | Tb927.2.4280_for | GGGGACAAGTTTGTACAAAAAAGCAGGCTATGGGTGCTGCCTTATGCAC | Tb927.2.4280_rev | GGGGACCACTTTGTACAAGAAAGCTGGGTTTAATCAAGTGCTAGCAACAGTTCATCC |
| 7_known | N1 | Tb927.11.5730_for | GGGGACAAGTTTGTACAAAAAAGCAGGCTATGCTTGACCGCCGGCGG | Tb927.11.5730_rev | GGGGACCACTTTGTACAAGAAAGCTGGGTTCAGATGTTGTTTCGCCGTCTTG |
| 7_known | N2 | Tb927.11.6900_for | GGGGACAAGTTTGTACAAAAAAGCAGGCTATGTCTTCAAAGACGGTAGTTTGGAG | Tb927.11.6900_rev | GGGGACCACTTTGTACAAGAAAGCTGGGTTTACGGTGTTGCTTCCTTCGC |
| 7_known | N3 | Tb927.7.2080_for | GGGGACAAGTTTGTACAAAAAAGCAGGCTATGGAATCGACGAGTTCACTTTCG | Tb927.7.2080_rev | GGGGACCACTTTGTACAAGAAAGCTGGGTTCAAATCTTGACGAACAAGAAGGTTCT |
| 7_known | N4 | Tb927.9.3400_for | GGGGACAAGTTTGTACAAAAAAGCAGGCTATGGCCACATCTCTTGTGCAG | Tb927.9.3400_rev | GGGGACCACTTTGTACAAGAAAGCTGGGTTTAGTTTCCAGAAGCTGATCCATAGAAGA |
| 7_known | N5 | Tb927.11.12540_for | GGGGACAAGTTTGTACAAAAAAGCAGGCTATGTCGGCGGTTGCAACC | Tb927.11.12540_rev | GGGGACCACTTTGTACAAGAAAGCTGGGTTCAGGCGATCCTCCGGTG |
| 7_known | N6 | Tb927.8.2640_for | GGGGACAAGTTTGTACAAAAAAGCAGGCTATGACTAGTGAAGAGCAAAGACGGC | Tb927.8.2640_rev | GGGGACCACTTTGTACAAGAAAGCTGGGTTTAAAAATCCTTGTACCTGTAATAAATTAG |
| 7_known | N7 | Tb927.11.10370_for | GGGGACAAGTTTGTACAAAAAAGCAGGCTATGCCCGCATACACTGGCC | Tb927.11.10370_rev | GGGGACCACTTTGTACAAGAAAGCTGGGTTCATAACCTCAAATCGAATGCAGCG |
| 7_known | N8 | Tb927.10.3550_for | GGGGACAAGTTTGTACAAAAAAGCAGGCTATGTATGCGGCTGGTCGCT | Tb927.10.3550_rev | GGGGACCACTTTGTACAAGAAAGCTGGGTTCACTTCCGCCGCTCGAT |
| 7_known | N9 | Tb927.11.6720_for | GGGGACAAGTTTGTACAAAAAAGCAGGCTATGTTGAACGCAAATGCCAGGT | Tb927.11.6720_rev | GGGGACCACTTTGTACAAGAAAGCTGGGTTTAATCCCTGTTAAGGTCAACGTTTTGT |
| 7_known | N10 | Tb927.8.3350_for | GGGGACAAGTTTGTACAAAAAAGCAGGCTATGCGCAAGACGGTGGTG | Tb927.8.3350_rev | GGGGACCACTTTGTACAAGAAAGCTGGGTTCAGTAGGTTGAAGATTGGAGGGTT |
| 7_known | N11 | Tb927.9.10920_for | GGGGACAAGTTTGTACAAAAAAGCAGGCTATGATTGGCACAGTAGAACAAATATTGTAC | Tb927.9.10920_rev | GGGGACCACTTTGTACAAGAAAGCTGGGTTTACACGCCACCGTAGAGCA |
| 7_known | N12 | Tb927.11.16310_for | GGGGACAAGTTTGTACAAAAAAGCAGGCTATGACCACGGAGCAAAAAGTCG | Tb927.11.16310_rev | GGGGACCACTTTGTACAAGAAAGCTGGGTCTACGTAACGTTGTTAAAAAGGCTACG |
| 7_known | N13 | Tb927.10.7280_for | GGGGACAAGTTTGTACAAAAAAGCAGGCTATGAGCGATGAGTTATGTGTGCTG | Tb927.10.7280_rev | GGGGACCACTTTGTACAAGAAAGCTGGGTTCACCTGCGACGCCTCTG |
| 7_known | N14 | Tb927.8.7020_for | GGGGACAAGTTTGTACAAAAAAGCAGGCTATGACGCGGGAGGAAGTCC | Tb927.8.7020_rev | GGGGACCACTTTGTACAAGAAAGCTGGGTCTAGAAGGTCCTGCAACGAATGAC |
| 7_known | N15 | Tb927.11.5690_for | GGGGACAAGTTTGTACAAAAAAGCAGGCTATGGATGGCGATGAATTATTCGTTTTAAC | Tb927.11.5690_rev | GGGGACCACTTTGTACAAGAAAGCTGGGTCTAGACAGTATCGTCATCAACGTATATGG |
| 7_known | N16 | Tb927.3.2040_for | GGGGACAAGTTTGTACAAAAAAGCAGGCTATGAACACTTCCGCGTGTTTTACA | Tb927.3.2040_rev | GGGGACCACTTTGTACAAGAAAGCTGGGTTCATGACTGCATCGCCTTCAC |
| 7_known | N17 | Tb927.11.3730_for | GGGGACAAGTTTGTACAAAAAAGCAGGCTATGTCGACTGTACGACGTGATAGA | Tb927.11.3730_rev | GGGGACCACTTTGTACAAGAAAGCTGGGTTCATGCCTTCTTGTCCGGAAAGG |
| 7_known | N18 | Tb927.11.9780_for | GGGGACAAGTTTGTACAAAAAAGCAGGCTATGGAACGGTTGATGGAGGGA | Tb927.11.9780_rev | GGGGACCACTTTGTACAAGAAAGCTGGGTCTATGAAACACTTTGTGAAGATGAGACAAA |
| 7_known | N19 | Tb927.7.4650_for | GGGGACAAGTTTGTACAAAAAAGCAGGCTATGCCTATGTTTATGGATGGGGCG | Tb927.7.4650_rev | GGGGACCACTTTGTACAAGAAAGCTGGGTTTATATCCACCGCTCGCTCTTTACA |
| 7_known | N20 | Tb927.8.6490_for | GGGGACAAGTTTGTACAAAAAAGCAGGCTATGGGTTGTGGTGCCTCCA | Tb927.8.6490_rev | GGGGACCACTTTGTACAAGAAAGCTGGGTTCACGCGTAGAGCCCCCG |
| 7_known | N21 | Tb927.4.1510_for | GGGGACAAGTTTGTACAAAAAAGCAGGCTATGCCAGAAGTAGAGAACGATAGCT | Tb927.4.1510_rev | GGGGACCACTTTGTACAAGAAAGCTGGGTTCACTTCATCCGCGCGTTG |
| 7_known | N22 | Tb927.8.1200_for | GGGGACAAGTTTGTACAAAAAAGCAGGCTATGCATCCACTTGAGTCACCGC | Tb927.8.1200_rev | GGGGACCACTTTGTACAAGAAAGCTGGGTTCATACCATCCTTAATGTGTCATAAAGTTG |
| 7_known | N23 | Tb927.1.1670_for | GGGGACAAGTTTGTACAAAAAAGCAGGCTATGTTGACAGGTGACATCTTGTTGC | Tb927.1.1670_rev | GGGGACCACTTTGTACAAGAAAGCTGGGTTCAGTAGGATGAACACGACTCGT |
| 7_known | N24 | Tb927.10.9570_for | GGGGACAAGTTTGTACAAAAAAGCAGGCTATGGAGAAAACACAACCGTTCAACG | Tb927.10.9570_rev | GGGGACCACTTTGTACAAGAAAGCTGGGTCTACTCTTTCACAAGAGCCCGAAC |
| 7_known | O1 | Tb927.10.14300_for | GGGGACAAGTTTGTACAAAAAAGCAGGCTATGAGTTCGATGGAGGAATTACACG | Tb927.10.14300_rev | GGGGACCACTTTGTACAAGAAAGCTGGGTTTACACATCCCCTGAATGCGTG |
| 7_known | O2 | Tb927.11.15100_for | GGGGACAAGTTTGTACAAAAAAGCAGGCTATGGCCGTTGTCCACAAGAAC | Tb927.11.15100_rev | GGGGACCACTTTGTACAAGAAAGCTGGGTTTACGCATCACCCTCACCAGT |
| 7_known | O3 | Tb927.6.2370_for | GGGGACAAGTTTGTACAAAAAAGCAGGCTATGCATGGAAATTTTGTATTTACCGGC | Tb927.6.2370_rev | GGGGACCACTTTGTACAAGAAAGCTGGGTTCAACTGAGACCGAAGTCATCCG |
| 7_known | O4 | Tb927.6.2880_for | GGGGACAAGTTTGTACAAAAAAGCAGGCTATGAACGTGCGTGTTGCTACA | Tb927.6.2880_rev | GGGGACCACTTTGTACAAGAAAGCTGGGTCTAACTCCGCCGAGTGATACCG |
| 7_known | O5 | Tb11.02.5390_for | GGGGACAAGTTTGTACAAAAAAGCAGGCTATGAATGGGAGGCTGTACTGTTTG | Tb11.02.5390_rev | GGGGACCACTTTGTACAAGAAAGCTGGGTTCAGCGTCTCAATAGAAGCTGAAGA |
| 7_known | O6 | Tb927.5.2090_for | GGGGACAAGTTTGTACAAAAAAGCAGGCTATGGGGAAGAAAACCAACAGCG | Tb927.5.2090_rev | GGGGACCACTTTGTACAAGAAAGCTGGGTTTAGAAATCAGCTCTTCGGACGTTC |
| 7_known | O7 | Tb927.11.4470_for | GGGGACAAGTTTGTACAAAAAAGCAGGCTATGGACAACTATACCATAGTCGAACTTATA | Tb927.11.4470_rev | GGGGACCACTTTGTACAAGAAAGCTGGGTTCACGAGGACGAAATCCTTTGC |
| 7_known | O8 | Tb927.11.13920_for | GGGGACAAGTTTGTACAAAAAAGCAGGCTATGTCTAAGAAACACCAAGGGGCG | Tb927.11.13920_rev | GGGGACCACTTTGTACAAGAAAGCTGGGTTCAGAGACGCCGCGTCGC |
| 7_known | O9 | Tb927.6.2770_for | GGGGACAAGTTTGTACAAAAAAGCAGGCTATGAGTATAGCGCGGTTGGAGC | Tb927.6.2770_rev | GGGGACCACTTTGTACAAGAAAGCTGGGTCTATGCGCTCTGCATTTCAACCT |
| 7_known | O10 | Tb927.4.3890_for | GGGGACAAGTTTGTACAAAAAAGCAGGCTATGGGCAAGAAGACCAGGGAT | Tb927.4.3890_rev | GGGGACCACTTTGTACAAGAAAGCTGGGTTCAGCCACGCTTTGGTAGGG |
| 7_known | O11 | Tb927.10.15020_for | GGGGACAAGTTTGTACAAAAAAGCAGGCTATGGACGCTGACGAGGGC | Tb927.10.15020_rev | GGGGACCACTTTGTACAAGAAAGCTGGGTCTAACGTTGTGAGTATACCTGTAAATATTT |
| 7_known | O12 | Tb927.10.15390_for | GGGGACAAGTTTGTACAAAAAAGCAGGCTATGTGTGACCGTCTAAATACAATGTCTATC | Tb927.10.15390_rev | GGGGACCACTTTGTACAAGAAAGCTGGGTTCAGCGTGGCCGTGAAGA |
| 7_known | O13 | Tb927.11.1100_for | GGGGACAAGTTTGTACAAAAAAGCAGGCTATGATGAATTGTAGGACTACGGAGCT | Tb927.11.1100_rev | GGGGACCACTTTGTACAAGAAAGCTGGGTTCAGTTTAGCAGCTCATGTGAATGC |
| 7_known | O14 | Tb927.6.1110_for | GGGGACAAGTTTGTACAAAAAAGCAGGCTATGGTTAGTGCGGGCAGTGA | Tb927.6.1110_rev | GGGGACCACTTTGTACAAGAAAGCTGGGTCTAATATGCAGTTTTAGCTGATCCACGC |
| 7_known | O15 | Tb927.11.11850_for | GGGGACAAGTTTGTACAAAAAAGCAGGCTATGGCCGATGAAGACGGTTCT | Tb927.11.11850_rev | GGGGACCACTTTGTACAAGAAAGCTGGGTTTACAGTAGCACCTCCAACTCGT |
| 7_known | O16 | Tb927.11.13060_for | GGGGACAAGTTTGTACAAAAAAGCAGGCTATGGTGAAACAGCGCATGACG | Tb927.11.13060_rev | GGGGACCACTTTGTACAAGAAAGCTGGGTCTATTTGCGAAGGCCAGTGCT |
| 7_known | O17 | Tb927.10.11620_for | GGGGACAAGTTTGTACAAAAAAGCAGGCTATGCGTGGGGAGAGCAGT | Tb927.10.11620_rev | GGGGACCACTTTGTACAAGAAAGCTGGGTTTAAACGTATGCAGGGGCAGTTT |
| 7_known | O18 | Tb927.8.1160_for | GGGGACAAGTTTGTACAAAAAAGCAGGCTATGATTTCCCCAAAAGACACTTCGAA | Tb927.8.1160_rev | GGGGACCACTTTGTACAAGAAAGCTGGGTTCATCTAATACGTTCCTCGCACCA |
| 7_known | O19 | Tb927.10.2220_for | GGGGACAAGTTTGTACAAAAAAGCAGGCTATGATGCGGAGGACTCTTCGT | Tb927.10.2220_rev | GGGGACCACTTTGTACAAGAAAGCTGGGTTTAAAAGAACCACAGTCTTTCATTCAAAGG |
| 7_known | O20 | Tb927.10.14610_for | GGGGACAAGTTTGTACAAAAAAGCAGGCTATGTCACGATCATGGTGTACATTTTCC | Tb927.10.14610_rev | GGGGACCACTTTGTACAAGAAAGCTGGGTCTACACCGTTGCGACTACCTG |
| 7_known | O21 | Tb927.8.1180_for | GGGGACAAGTTTGTACAAAAAAGCAGGCTATGATTTCCCCAAAAGACACTTCGAA | Tb927.8.1180_rev | GGGGACCACTTTGTACAAGAAAGCTGGGTTCATACCATCCTTAATGTGTCATAAAGTTG |
| 7_known | O22 | Tb927.2.690_for | GGGGACAAGTTTGTACAAAAAAGCAGGCTATGTCAGGAAACCAAGGAAGAAAGC | Tb927.2.690_rev | GGGGACCACTTTGTACAAGAAAGCTGGGTTTAACGGATATATACCCTCCTCTTTCTCAG |
| 7_known | O23 | Tb927.6.4390_for | GGGGACAAGTTTGTACAAAAAAGCAGGCTATGTCGGATGCCGATGTGAAAG | Tb927.6.4390_rev | GGGGACCACTTTGTACAAGAAAGCTGGGTTCAGTTGGAGCGCAGGCG |
| 7_known | O24 | Tb927.11.4640_for | GGGGACAAGTTTGTACAAAAAAGCAGGCTATGAATACGATCGATGCTGCTCTTG | Tb927.11.4640_rev | GGGGACCACTTTGTACAAGAAAGCTGGGTTTACAACACCTCCGCCTTCAGT |
| 7_known | P1 | Tb927.11.15330_for | GGGGACAAGTTTGTACAAAAAAGCAGGCTATGACCCAGTCGGCGGAA | Tb927.11.15330_rev | GGGGACCACTTTGTACAAGAAAGCTGGGTTCATTTTGTTTGATTTCGTGCATCCA |
| 7_known | P2 | Tb927.9.14650_for | GGGGACAAGTTTGTACAAAAAAGCAGGCTATGAGTAACAACGGTTACTGCAACA | Tb927.9.14650_rev | GGGGACCACTTTGTACAAGAAAGCTGGGTTCATGGATATCCTCCTGCGGG |
| 7_known | P3 | Tb927.5.1490_for | GGGGACAAGTTTGTACAAAAAAGCAGGCTATGAACTGCTCCGCGCCA | Tb927.5.1490_rev | GGGGACCACTTTGTACAAGAAAGCTGGGTTTAATTTCCCCCAAGGGACCGA |
| 7_known | P4 | Tb927.1.1050_for | GGGGACAAGTTTGTACAAAAAAGCAGGCTATGAATGTCACTAACCCCTCCACT | Tb927.1.1050_rev | GGGGACCACTTTGTACAAGAAAGCTGGGTTCACTCACGGAGAGGATCTGGC |
| 7_known | P5 | Tb927.7.4080_for | GGGGACAAGTTTGTACAAAAAAGCAGGCTATGGCAGATGAACTCTCCCAGT | Tb927.7.4080_rev | GGGGACCACTTTGTACAAGAAAGCTGGGTTTACGATCCCTCCACCCTGGG |
| 7_known | P6 | Tb927.5.3610_for | GGGGACAAGTTTGTACAAAAAAGCAGGCTATGACGGATGCAACTAAGATGGC | Tb927.5.3610_rev | GGGGACCACTTTGTACAAGAAAGCTGGGTCTACGACGTGGCCGTCATC |
| 7_known | P7 | Tb927.11.3350_for | GGGGACAAGTTTGTACAAAAAAGCAGGCTATGGCGGAAGACGCTGCA | Tb927.11.3350_rev | GGGGACCACTTTGTACAAGAAAGCTGGGTCTACCCATGAAGCGTATCCCCG |
| 7_known | P8 | Tb927.1.540_for | GGGGACAAGTTTGTACAAAAAAGCAGGCTATGACCCCTAGGGCCCGT | Tb927.1.540_rev | GGGGACCACTTTGTACAAGAAAGCTGGGTTCATTGCGATCCCATGAAATCAAGC |
| 7_known | P9 | Tb927.10.12490_for | GGGGACAAGTTTGTACAAAAAAGCAGGCTATGATGTCTTCTTCATCCTCAGCTCA | Tb927.10.12490_rev | GGGGACCACTTTGTACAAGAAAGCTGGGTTCAGAAGAAAAGAGCGGGCCT |
| 7_known | P10 | Tb927.5.930_for | GGGGACAAGTTTGTACAAAAAAGCAGGCTATGGTAGACGGGCGATCTTCTG | Tb927.5.930_rev | GGGGACCACTTTGTACAAGAAAGCTGGGTTCAAATTTTGGATGAGCCGCTCG |
| 7_known | P11 | Tb927.10.9190_for | GGGGACAAGTTTGTACAAAAAAGCAGGCTATGAATCGATGCTACTCCCTCCTT | Tb927.10.9190_rev | GGGGACCACTTTGTACAAGAAAGCTGGGTTTACGATTCACCAGCCGACGG |
| 7_known | P12 | Tb927.3.2790_for | GGGGACAAGTTTGTACAAAAAAGCAGGCTATGGTTCGCGACGATACGC | Tb927.3.2790_rev | GGGGACCACTTTGTACAAGAAAGCTGGGTTCAACGGCGCACACGCCA |
| 7_known | P13 | Tb927.1.2100_for | GGGGACAAGTTTGTACAAAAAAGCAGGCTATGTCGGAGTTCGAACTTATGAGTG | Tb927.1.2100_rev | GGGGACCACTTTGTACAAGAAAGCTGGGTTCAGCACACGGAGGAGTCC |
| 7_known | P14 | Tb927.10.7180_for | GGGGACAAGTTTGTACAAAAAAGCAGGCTATGTCCGCCAAATACTACATTCCTTT | Tb927.10.7180_rev | GGGGACCACTTTGTACAAGAAAGCTGGGTTTACGGTTTCGCTTCATCTGAGG |
| 7_known | P15 | Tb927.7.1190_for | GGGGACAAGTTTGTACAAAAAAGCAGGCTATGCGTCCATTAGTTGAGCGTT | Tb927.7.1190_rev | GGGGACCACTTTGTACAAGAAAGCTGGGTCTACTTCTCTCTATTCTGGTTGTTGCG |
| 7_known | P16 | Tb927.10.15300_for | GGGGACAAGTTTGTACAAAAAAGCAGGCTATGTTACTGGATTTCGTCCTAAAGTACG | Tb927.10.15300_rev | GGGGACCACTTTGTACAAGAAAGCTGGGTCTAACGTCCCTTTTGTAGCCACG |
| 7_known | P17 | Tb927.8.2380_for | GGGGACAAGTTTGTACAAAAAAGCAGGCTATGCACTCGGAATTGAAAGCCTATG | Tb927.8.2380_rev | GGGGACCACTTTGTACAAGAAAGCTGGGTTCAACGTGCGACAGTAGTGAATTG |
| 7_known | P18 | Tb927.1.3120_for | GGGGACAAGTTTGTACAAAAAAGCAGGCTATGAGGCGCACTCAGGTTCG | Tb927.1.3120_rev | GGGGACCACTTTGTACAAGAAAGCTGGGTTCACATCGCTCCACCCGG |
| 7_known | P19 | Tb927.2.1810_for | GGGGACAAGTTTGTACAAAAAAGCAGGCTATGGAGGCACCAGCAGGAC | Tb927.2.1810_rev | GGGGACCACTTTGTACAAGAAAGCTGGGTTTATTCCGGAAACTTCCGCTCCA |
| 7_known | P20 | Tb927.9.14470_for | GGGGACAAGTTTGTACAAAAAAGCAGGCTATGCTTCAGGTTCTAGGTATTGTGAAG | Tb927.9.14470_rev | GGGGACCACTTTGTACAAGAAAGCTGGGTTTAATTGTGGATAACAACCTTATCCTCTTT |
| 7_known | P21 | Tb927.8.5210_for | GGGGACAAGTTTGTACAAAAAAGCAGGCTATGCCCTTGTCTGGGGGC | Tb927.8.5210_rev | GGGGACCACTTTGTACAAGAAAGCTGGGTTCACACAACGTACTCGAGTAATTGTG |
| 7_known | P22 | Tb927.10.11080_for | GGGGACAAGTTTGTACAAAAAAGCAGGCTATGTATAATAATTCCCCAGGCGAGGG | Tb927.10.11080_rev | GGGGACCACTTTGTACAAGAAAGCTGGGTTCACCTCAGGCTGCGCAA |
| 7_known | P23 | Tb927.10.1270_for | GGGGACAAGTTTGTACAAAAAAGCAGGCTATGCGCAACAATTTCCCGATTCC | Tb927.10.1270_rev | GGGGACCACTTTGTACAAGAAAGCTGGGTTCACTCAGTAGAGGAACCGCATG |
| 7_known | P24 | Tb927.4.3380_for | GGGGACAAGTTTGTACAAAAAAGCAGGCTATGGCGCATCTGGACCTTAATAAAC | Tb927.4.3380_rev | GGGGACCACTTTGTACAAGAAAGCTGGGTTTACCGCTTTGACCGTTGCTC |
| 8_hypothetical | A1 | Tb927.11.400_for | GGGGACAAGTTTGTACAAAAAAGCAGGCTATGAATATTTATGAATGGCAGTCAATTGAC | Tb927.11.400_rev | GGGGACCACTTTGTACAAGAAAGCTGGGTCTACACATTAACACCCGTACTAATGGG |
| 8_hypothetical | A2 | Tb927.6.910_for | GGGGACAAGTTTGTACAAAAAAGCAGGCTATGGTACTTGGTGGTGTAGTTGGG | Tb927.6.910_rev | GGGGACCACTTTGTACAAGAAAGCTGGGTCTAGTCGACTGGTGAGGCCTC |
| 8_hypothetical | A3 | Tb927.7.2240_for | GGGGACAAGTTTGTACAAAAAAGCAGGCTATGCCGGTTAAGAAGAAGGTTGC | Tb927.7.2240_rev | GGGGACCACTTTGTACAAGAAAGCTGGGTTTAATACTTCCAACCACGACTTCTTCC |
| 8_hypothetical | A4 | Tb927.11.13560_for | GGGGACAAGTTTGTACAAAAAAGCAGGCTATGCGATGTTGTCGGACAATGC | Tb927.11.13560_rev | GGGGACCACTTTGTACAAGAAAGCTGGGTTTACGCGTTTTGGGGTGGAAA |
| 8_hypothetical | A5 | Tb927.11.12450.1_for | GGGGACAAGTTTGTACAAAAAAGCAGGCTATGGCTGAGATGACGGAGGC | Tb927.11.12450.1_rev | GGGGACCACTTTGTACAAGAAAGCTGGGTTTAGGGGGCCGCAAGTATTTTTC |
| 8_hypothetical | A6 | Tb927.11.6840_for | GGGGACAAGTTTGTACAAAAAAGCAGGCTATGGACACATCTGAGGAGGCAA | Tb927.11.6840_rev | GGGGACCACTTTGTACAAGAAAGCTGGGTTCACAACGAAAACGACCTAACGAC |
| 8_hypothetical | A7 | Tb927.8.6470_for | GGGGACAAGTTTGTACAAAAAAGCAGGCTATGCCTGGGTGTCTTGAGGT | Tb927.8.6470_rev | GGGGACCACTTTGTACAAGAAAGCTGGGTTTATGTTGGTTCACATTCAGTTACGCA |
| 8_hypothetical | A8 | Tb927.10.3350_for | GGGGACAAGTTTGTACAAAAAAGCAGGCTATGTGCCAAACTTTAACTACGGACG | Tb927.10.3350_rev | GGGGACCACTTTGTACAAGAAAGCTGGGTTCATCTGACTCTCTCGTGTGAGG |
| 8_hypothetical | A9 | Tb927.5.2340_for | GGGGACAAGTTTGTACAAAAAAGCAGGCTATGTTGGAGGGACTATGCGTGC | Tb927.5.2340_rev | GGGGACCACTTTGTACAAGAAAGCTGGGTTCACCTCATAAACTTTCTCCGCCA |
| 8_hypothetical | A10 | Tb927.11.1080_for | GGGGACAAGTTTGTACAAAAAAGCAGGCTATGACGAACAGAGAAAACACGTGG | Tb927.11.1080_rev | GGGGACCACTTTGTACAAGAAAGCTGGGTTTAGTCACCCTCAAAGGAAGATAATGACC |
| 8_hypothetical | A11 | Tb927.11.13070_for | GGGGACAAGTTTGTACAAAAAAGCAGGCTATGGATGAGCGGTCATTTGAGTTG | Tb927.11.13070_rev | GGGGACCACTTTGTACAAGAAAGCTGGGTTCATACCCCCTCGACCGGC |
| 8_hypothetical | A12 | Tb927.8.2590_for | GGGGACAAGTTTGTACAAAAAAGCAGGCTATGGCGGTGTCGGGATCG | Tb927.8.2590_rev | GGGGACCACTTTGTACAAGAAAGCTGGGTTTAATCTTCTTCCAGCAACCGGTTAAG |
| 8_hypothetical | A13 | Tb927.10.7010_for | GGGGACAAGTTTGTACAAAAAAGCAGGCTATGAACATTTCTGATATCGTGGGAGAGG | Tb927.10.7010_rev | GGGGACCACTTTGTACAAGAAAGCTGGGTTCACACGATCACTTGGGCTTCT |
| 8_hypothetical | A14 | Tb927.3.4710_for | GGGGACAAGTTTGTACAAAAAAGCAGGCTATGAGCATGCAACACACCCTC | Tb927.3.4710_rev | GGGGACCACTTTGTACAAGAAAGCTGGGTTTAGATCCTGGCGGTATCGCA |
| 8_hypothetical | A15 | Tb927.10.5830_for | GGGGACAAGTTTGTACAAAAAAGCAGGCTATGTATTTACATTCCATGAAGCGCTACA | Tb927.10.5830_rev | GGGGACCACTTTGTACAAGAAAGCTGGGTTCAAATTTTGACGGGCTTCCGC |
| 8_hypothetical | A16 | Tb927.10.8860_for | GGGGACAAGTTTGTACAAAAAAGCAGGCTATGACCACAATTTTAGAAAATGACAGAGC | Tb927.10.8860_rev | GGGGACCACTTTGTACAAGAAAGCTGGGTTTAGTCGTTCTGTGGGACGTACG |
| 8_hypothetical | A17 | Tb927.3.1580_for | GGGGACAAGTTTGTACAAAAAAGCAGGCTATGGAAACGGTCCCTGAGTCG | Tb927.3.1580_rev | GGGGACCACTTTGTACAAGAAAGCTGGGTTCACATGAAAGATGTAGCCAAGCTC |
| 8_hypothetical | A18 | Tb927.11.14700_for | GGGGACAAGTTTGTACAAAAAAGCAGGCTATGTATTTCAAGCCATCTCTGACGC | Tb927.11.14700_rev | GGGGACCACTTTGTACAAGAAAGCTGGGTTTACTTTTCGATAGTGTCCTTTTTTTCAGT |
| 8_hypothetical | A19 | Tb927.8.6700_for | GGGGACAAGTTTGTACAAAAAAGCAGGCTATGTCCTCCGTTACCACTGGC | Tb927.8.6700_rev | GGGGACCACTTTGTACAAGAAAGCTGGGTTTAAGAAGCCTCCGGCGAGG |
| 8_hypothetical | A20 | Tb927.10.9380_for | GGGGACAAGTTTGTACAAAAAAGCAGGCTATGCCAGTGGATGCGCCC | Tb927.10.9380_rev | GGGGACCACTTTGTACAAGAAAGCTGGGTCTATGACTTTGTTTCCCTTGGTGGC |

| | | | | | |
|---|---|---|---|---|---|
| 8_hypothetical | A21 | Tb927.4.490_for | GGGGACAAGTTTGTACAAAAAAGCAGGCTATGTCGAAACGCAAGCGGT | Tb927.4.490_rev | GGGGACCACTTTGTACAAGAAAGCTGGGTCTAACTATCCGACGATATTTCTATGCCG |
| 8_hypothetical | A22 | Tb927.6.3190_for | GGGGACAAGTTTGTACAAAAAAGCAGGCTATGGAAAGCACCATCATGTCTATCG | Tb927.6.3190_rev | GGGGACCACTTTGTACAAGAAAGCTGGGTTTATTCATTTTCTTCTTCGTCAGAATTATC |
| 8_hypothetical | A23 | Tb927.11.4050_for | GGGGACAAGTTTGTACAAAAAAGCAGGCTATGACCGTCTGCTTCAAATGCA | Tb927.11.4050_rev | GGGGACCACTTTGTACAAGAAAGCTGGGTTTAACTGTCTTTCTCACCATCATCTGTC |
| 8_hypothetical | A24 | Tb927.7.6300_for | GGGGACAAGTTTGTACAAAAAAGCAGGCTATGGGTTGCGGTTCGAGTGT | Tb927.7.6300_rev | GGGGACCACTTTGTACAAGAAAGCTGGGTCTACATGAGTTCTTGGATATATATATTCCC |
| 8_hypothetical | B1 | Tb927.10.6580_for | GGGGACAAGTTTGTACAAAAAAGCAGGCTATGAGCTTAAGACTTGGTCAGCCG | Tb927.10.6580_rev | GGGGACCACTTTGTACAAGAAAGCTGGGTCTACGCCCCGAGTGCTACA |
| 8_hypothetical | B2 | Tb927.11.3700_for | GGGGACAAGTTTGTACAAAAAAGCAGGCTATGACTGACAGCGAGTCGGA | Tb927.11.3700_rev | GGGGACCACTTTGTACAAGAAAGCTGGGTTCACTTCCCGCGTTCCACC |
| 8_hypothetical | B3 | Tb927.10.2470_for | GGGGACAAGTTTGTACAAAAAAGCAGGCTATGCCACAAAATATGGAAGTACGTGA | Tb927.10.2470_rev | GGGGACCACTTTGTACAAGAAAGCTGGGTCTAGTCCCCCTTTTCCCTCGTG |
| 8_hypothetical | B4 | Tb927.3.760_for | GGGGACAAGTTTGTACAAAAAAGCAGGCTATGTCAGTTTTAAGCGAATGGAGGC | Tb927.3.760_rev | GGGGACCACTTTGTACAAGAAAGCTGGGTTTAAGAGGCTCGGAGAATGACAGC |
| 8_hypothetical | B5 | Tb927.11.11380_for | GGGGACAAGTTTGTACAAAAAAGCAGGCTATGCTGCGCCGCGGTGCG | Tb927.11.11380_rev | GGGGACCACTTTGTACAAGAAAGCTGGGTTTAGCTCTTTTTTTTATCGTCGTCTTCATT |
| 8_hypothetical | B6 | Tb927.11.6540_for | GGGGACAAGTTTGTACAAAAAAGCAGGCTATGCAACTTCGACATGAATTGAAGTTTG | Tb927.11.6540_rev | GGGGACCACTTTGTACAAGAAAGCTGGGTTCAGTTAGTCACTGCAGGCTCA |
| 8_hypothetical | B7 | Tb927.11.2000_for | GGGGACAAGTTTGTACAAAAAAGCAGGCTATGGCGCAGAGCTGGTTCA | Tb927.11.2000_rev | GGGGACCACTTTGTACAAGAAAGCTGGGTTCAGAAGCTGAACGCAAAGCC |
| 8_hypothetical | B8 | Tb927.8.4940_for | GGGGACAAGTTTGTACAAAAAAGCAGGCTATGGGATCCTGCTGCGGAG | Tb927.8.4940_rev | GGGGACCACTTTGTACAAGAAAGCTGGGTTCAAATTCCCCCCGTCCTGC |
| 8_hypothetical | B9 | Tb927.3.3950_for | GGGGACAAGTTTGTACAAAAAAGCAGGCTATGCCTCCAAAACGCTCAAAAGG | Tb927.3.3950_rev | GGGGACCACTTTGTACAAGAAAGCTGGGTTTACAAGAAGGACAATGTCTTCCTCG |
| 8_hypothetical | B10 | Tb927.9.15480_for | GGGGACAAGTTTGTACAAAAAAGCAGGCTATGAAAGTTCCAATAGGCATCTTGCG | Tb927.9.15480_rev | GGGGACCACTTTGTACAAGAAAGCTGGGTTCAGACGAAGAGAACAGATGTGAGA |
| 8_hypothetical | B11 | Tb927.6.4190_for | GGGGACAAGTTTGTACAAAAAAGCAGGCTATGAAGTATATGGCAAAGGGGGCG | Tb927.6.4190_rev | GGGGACCACTTTGTACAAGAAAGCTGGGTTTATGACAAGTGGGGCTGAAGATGC |
| 8_hypothetical | B12 | Tb927.9.4050_for | GGGGACAAGTTTGTACAAAAAAGCAGGCTATGAGTGTCACCGGCTTCGG | Tb927.9.4050_rev | GGGGACCACTTTGTACAAGAAAGCTGGGTTCACTCCAAATCCCTCATCAAACTCT |
| 8_hypothetical | B13 | Tb927.7.1700_for | GGGGACAAGTTTGTACAAAAAAGCAGGCTATGAGGTACAGTGTTGTCAGAACGC | Tb927.7.1700_rev | GGGGACCACTTTGTACAAGAAAGCTGGGTTTACGGAAATAGTACAGTTCGGCG |
| 8_hypothetical | B14 | Tb927.11.9440_for | GGGGACAAGTTTGTACAAAAAAGCAGGCTATGAACGAATTAAAGAACCCGATGCG | Tb927.11.9440_rev | GGGGACCACTTTGTACAAGAAAGCTGGGTTTACATATTTAATCTAATGTCCTTGATGTC |
| 8_hypothetical | B15 | Tb927.11.9540_for | GGGGACAAGTTTGTACAAAAAAGCAGGCTATGAAAACGTACAGTGAGCTGCTG | Tb927.11.9540_rev | GGGGACCACTTTGTACAAGAAAGCTGGGTTCACTTTGACAACTCAACGAGCC |
| 8_hypothetical | B16 | Tb927.9.6660_for | GGGGACAAGTTTGTACAAAAAAGCAGGCTATGGCAGTTGATCGTATCGGAAAC | Tb927.9.6660_rev | GGGGACCACTTTGTACAAGAAAGCTGGGTTCAACTGGACTCAAACTCCTGCT |
| 8_hypothetical | B17 | Tb927.7.1400_for | GGGGACAAGTTTGTACAAAAAAGCAGGCTATGAACAATGCTAATGACACTTTAGATTAC | Tb927.7.1400_rev | GGGGACCACTTTGTACAAGAAAGCTGGGTTCACGCACCCTCCCGTCG |
| 8_hypothetical | B18 | Tb927.8.820_for | GGGGACAAGTTTGTACAAAAAAGCAGGCTATGTCATTCCACAAGTTATTTTCTTCAAAT | Tb927.8.820_rev | GGGGACCACTTTGTACAAGAAAGCTGGGTCTATTCACTCGTCGACCGCCA |
| 8_hypothetical | B19 | Tb927.11.11790_for | GGGGACAAGTTTGTACAAAAAAGCAGGCTATGGCGGCAACAGGTTCAC | Tb927.11.11790_rev | GGGGACCACTTTGTACAAGAAAGCTGGGTTTACGCGGGCTTGCTATCGC |
| 8_hypothetical | B20 | Tb927.9.3630_for | GGGGACAAGTTTGTACAAAAAAGCAGGCTATGGCCGCGTCGCAGGTA | Tb927.9.3630_rev | GGGGACCACTTTGTACAAGAAAGCTGGGTTCACTCATCGACCTCGTCTTCC |
| 8_hypothetical | B21 | Tb927.11.7214_for | GGGGACAAGTTTGTACAAAAAAGCAGGCTATGAAGAGATACCGTTCCTCTGTTGT | Tb927.11.7214_rev | GGGGACCACTTTGTACAAGAAAGCTGGGTTCAAAAACGATAACACGAAAGGCAAG |
| 8_hypothetical | B22 | Tb927.7.5400_for | GGGGACAAGTTTGTACAAAAAAGCAGGCTATGTATACAACAACAGACCTCATACAGAC | Tb927.7.5400_rev | GGGGACCACTTTGTACAAGAAAGCTGGGTCTACGTGTTAAACAAACCACCAGACG |
| 8_hypothetical | B23 | Tb927.11.12490_for | GGGGACAAGTTTGTACAAAAAAGCAGGCTATGAGCTCCAGGATGTCACAGAA | Tb927.11.12490_rev | GGGGACCACTTTGTACAAGAAAGCTGGGTCTAACGTGTGCGAACACGCT |
| 8_hypothetical | B24 | Tb927.10.15430_for | GGGGACAAGTTTGTACAAAAAAGCAGGCTATGAAGAAGAATAACTCCAAGGGTAACTCC | Tb927.10.15430_rev | GGGGACCACTTTGTACAAGAAAGCTGGGTTTAACCGAACAAACTTACAATGGTATTCCT |
| 8_hypothetical | C1 | Tb927.3.2010_for | GGGGACAAGTTTGTACAAAAAAGCAGGCTATGGATACCCTCGCATTCCGC | Tb927.3.2010_rev | GGGGACCACTTTGTACAAGAAAGCTGGGTTTACCGACTAGGCAAATCCGCG |
| 8_hypothetical | C2 | Tb927.11.8700_for | GGGGACAAGTTTGTACAAAAAAGCAGGCTATGGTTTCGTTTATTTACAGCTGCG | Tb927.11.8700_rev | GGGGACCACTTTGTACAAGAAAGCTGGGTTTATGTGACGATGCGAATTCCCG |
| 8_hypothetical | C3 | Tb927.11.12400_for | GGGGACAAGTTTGTACAAAAAAGCAGGCTATGATCCATGCCACCGGTGA | Tb927.11.12400_rev | GGGGACCACTTTGTACAAGAAAGCTGGGTTCACTTTCCGTACACCACAGTGC |
| 8_hypothetical | C4 | Tb927.10.10070_for | GGGGACAAGTTTGTACAAAAAAGCAGGCTATGGATCTTGACAACGTGGTTCTTG | Tb927.10.10070_rev | GGGGACCACTTTGTACAAGAAAGCTGGGTTCAGGAATTAAGATCGCCAAAGAAACC |
| 8_hypothetical | C5 | Tb927.10.14390_for | GGGGACAAGTTTGTACAAAAAAGCAGGCTATGTCGAGCTTGGCAGGAGT | Tb927.10.14390_rev | GGGGACCACTTTGTACAAGAAAGCTGGGTTTAATGATCTTTCTTGTGCTTTTTATGACG |
| 8_hypothetical | C6 | Tb927.11.14530_for | GGGGACAAGTTTGTACAAAAAAGCAGGCTATGCCGGTTACGACACACTTTT | Tb927.11.14530_rev | GGGGACCACTTTGTACAAGAAAGCTGGGTCTAATCAATTATGAAGACGACACCGGG |
| 8_hypothetical | C7 | Tb927.5.3480_for | GGGGACAAGTTTGTACAAAAAAGCAGGCTATGAGTTCTACAAAGGATGGATCTCAAATT | Tb927.5.3480_rev | GGGGACCACTTTGTACAAGAAAGCTGGGTCTAAGGTTTCTTAACCGCAAGTTGTACC |
| 8_hypothetical | C8 | Tb927.10.3320_for | GGGGACAAGTTTGTACAAAAAAGCAGGCTATGCGAGGGTGTGCAAAAGG | Tb927.10.3320_rev | GGGGACCACTTTGTACAAGAAAGCTGGGTTCAAGTACCTTTCGTGCTATTGTCG |
| 8_hypothetical | C9 | Tb927.3.3650_for | GGGGACAAGTTTGTACAAAAAAGCAGGCTATGGAGCCTCTGTGGGACTC | Tb927.3.3650_rev | GGGGACCACTTTGTACAAGAAAGCTGGGTCTAACGACCCTTTCTTGACCTCCC |
| 8_hypothetical | C10 | Tb927.11.12190_for | GGGGACAAGTTTGTACAAAAAAGCAGGCTATGTCAGGTTGCGACGGGA | Tb927.11.12190_rev | GGGGACCACTTTGTACAAGAAAGCTGGGTCTACTCTTTCCGCTTTTCTTGGTAGC |
| 8_hypothetical | C11 | Tb927.10.10330_for | GGGGACAAGTTTGTACAAAAAAGCAGGCTATGACAGCTCAGAGAGACCACC | Tb927.10.10330_rev | GGGGACCACTTTGTACAAGAAAGCTGGGTCTACTCAGGCTCTTCCATGGATATTCT |
| 8_hypothetical | C12 | Tb927.11.5880_for | GGGGACAAGTTTGTACAAAAAAGCAGGCTATGATGCACTTGTCGTTACAACTATGT | Tb927.11.5880_rev | GGGGACCACTTTGTACAAGAAAGCTGGGTTCACCCTGAGGGAAGTAAATGTTTTTT |
| 8_hypothetical | C13 | Tb927.11.8920_for | GGGGACAAGTTTGTACAAAAAAGCAGGCTATGGATGAGGACATGTGTGTTGC | Tb927.11.8920_rev | GGGGACCACTTTGTACAAGAAAGCTGGGTTTAAAAATTCTGTGACAGGAAATTTTCATA |
| 8_hypothetical | C14 | Tb927.1.3050_for | GGGGACAAGTTTGTACAAAAAAGCAGGCTATGGAGGGGACGGAGTGC | Tb927.1.3050_rev | GGGGACCACTTTGTACAAGAAAGCTGGGTTTATGAGCTATTACTTCTGGAAAGGACGG |
| 8_hypothetical | C15 | Tb927.8.7770_for | GGGGACAAGTTTGTACAAAAAAGCAGGCTATGCAGTTTTGTGGCGGGC | Tb927.8.7770_rev | GGGGACCACTTTGTACAAGAAAGCTGGGTTCACCAGTCATGCACAACATAATGT |
| 8_hypothetical | C16 | Tb927.11.7340_for | GGGGACAAGTTTGTACAAAAAAGCAGGCTATGCAAAGTTCCTCTGAGACGATG | Tb927.11.7340_rev | GGGGACCACTTTGTACAAGAAAGCTGGGTTCAAAACCGTGAACCCCTGC |
| 8_hypothetical | C17 | Tb927.10.3590_for | GGGGACAAGTTTGTACAAAAAAGCAGGCTATGTTCCGCCGGACAAGTCG | Tb927.10.3590_rev | GGGGACCACTTTGTACAAGAAAGCTGGGTCTATCCATGCGGCATGTTGTCC |
| 8_hypothetical | C18 | Tb927.11.13180_for | GGGGACAAGTTTGTACAAAAAAGCAGGCTATGCCTTCAAAGCGGAAAACTATCG | Tb927.11.13180_rev | GGGGACCACTTTGTACAAGAAAGCTGGGTTTAGATGAACTCGAGGGACTCCAC |
| 8_hypothetical | C19 | Tb927.11.9300_for | GGGGACAAGTTTGTACAAAAAAGCAGGCTATGGGAAACAGGTTTGAAGGGAGT | Tb927.11.9300_rev | GGGGACCACTTTGTACAAGAAAGCTGGGTCTAAGAAAGCAGTCTCGCACCG |
| 8_hypothetical | C20 | Tb927.4.1970_for | GGGGACAAGTTTGTACAAAAAAGCAGGCTATGGGTGGGCACCACGGA | Tb927.4.1970_rev | GGGGACCACTTTGTACAAGAAAGCTGGGTTCACAGTGAAATGTTGGAAAGCATTG |
| 8_hypothetical | C21 | Tb927.7.7150_for | GGGGACAAGTTTGTACAAAAAAGCAGGCTATGGAATTGTTGGGGAACGCAA | Tb927.7.7150_rev | GGGGACCACTTTGTACAAGAAAGCTGGGTTCATATGCTGTGCACAATCCTCC |
| 8_hypothetical | C22 | Tb927.3.4140_for | GGGGACAAGTTTGTACAAAAAAGCAGGCTATGGTACGGCGGCGGCCT | Tb927.3.4140_rev | GGGGACCACTTTGTACAAGAAAGCTGGGTTTACTGTCTGGTGAGTTCACAAACC |
| 8_hypothetical | C23 | Tb927.6.3640_for | GGGGACAAGTTTGTACAAAAAAGCAGGCTATGACGGAGGTTCCGGGG | Tb927.6.3640_rev | GGGGACCACTTTGTACAAGAAAGCTGGGTTCAGCACGACACAATCAAGGATTC |
| 8_hypothetical | C24 | Tb927.4.1650_for | GGGGACAAGTTTGTACAAAAAAGCAGGCTATGCCGGAGAATCCCCAACG | Tb927.4.1650_rev | GGGGACCACTTTGTACAAGAAAGCTGGGTTCACCGTGATGTGATTGCTGC |
| 8_hypothetical | D1 | Tb927.9.11480_for | GGGGACAAGTTTGTACAAAAAAGCAGGCTATGACAGGTGATGTTCGGCCG | Tb927.9.11480_rev | GGGGACCACTTTGTACAAGAAAGCTGGGTTTACGACTGACGCGCCACA |
| 8_hypothetical | D2 | Tb927.8.5300_for | GGGGACAAGTTTGTACAAAAAAGCAGGCTATGACCGCTGTTGGGGGC | Tb927.8.5300_rev | GGGGACCACTTTGTACAAGAAAGCTGGGTTTAGGACACTATTGCCGGCACC |
| 8_hypothetical | D3 | Tb927.7.3150_for | GGGGACAAGTTTGTACAAAAAAGCAGGCTATGGTGAAAACGGTGTCGTCG | Tb927.7.3150_rev | GGGGACCACTTTGTACAAGAAAGCTGGGTTCACATTACCGTACAACAGGAGGA |
| 8_hypothetical | D4 | Tb927.2.4560_for | GGGGACAAGTTTGTACAAAAAAGCAGGCTATGTTAACAGGTCGACAAAAGGTTGT | Tb927.2.4560_rev | GGGGACCACTTTGTACAAGAAAGCTGGGTTTACGCAAGCATGCTTGGCG |
| 8_hypothetical | D5 | Tb927.2.2990_for | GGGGACAAGTTTGTACAAAAAAGCAGGCTATGTCACGGCGTGCTGAGG | Tb927.2.2990_rev | GGGGACCACTTTGTACAAGAAAGCTGGGTTTAGTAAAAGTCCAAAACTAGCATCATGTC |
| 8_hypothetical | D6 | Tb927.11.1210_for | GGGGACAAGTTTGTACAAAAAAGCAGGCTATGAGCAACAAATACCTTTTGGATGAGA | Tb927.11.1210_rev | GGGGACCACTTTGTACAAGAAAGCTGGGTTCACGCCACCACAGAGCA |
| 8_hypothetical | D7 | Tb927.11.11350_for | GGGGACAAGTTTGTACAAAAAAGCAGGCTATGCAATCAACGTCTGCAGGT | Tb927.11.11350_rev | GGGGACCACTTTGTACAAGAAAGCTGGGTTTACTTTTCACCGTACCATTTAGCAAGT |
| 8_hypothetical | D8 | Tb927.3.4370_for | GGGGACAAGTTTGTACAAAAAAGCAGGCTATGCAGACGGTATGGAATGTTCC | Tb927.3.4370_rev | GGGGACCACTTTGTACAAGAAAGCTGGGTCTACGCACTCCCAAGAAAAATTAATGC |
| 8_hypothetical | D9 | Tb927.3.5040_for | GGGGACAAGTTTGTACAAAAAAGCAGGCTATGACACAACCTGCCCACCT | Tb927.3.5040_rev | GGGGACCACTTTGTACAAGAAAGCTGGGTTCACATGTACGCCACTTCATCTTC |
| 8_hypothetical | D10 | Tb927.8.7560_for | GGGGACAAGTTTGTACAAAAAAGCAGGCTATGATACCGAATGGCGGGGA | Tb927.8.7560_rev | GGGGACCACTTTGTACAAGAAAGCTGGGTCTAAAAGACTAGTTTGTCCGGTCTGC |
| 8_hypothetical | D11 | Tb927.6.3460_for | GGGGACAAGTTTGTACAAAAAAGCAGGCTATGGTACTAAATGTGATTCTTCTTGGTACG | Tb927.6.3460_rev | GGGGACCACTTTGTACAAGAAAGCTGGGTTCACAATGACGAATCCTGAGAAAGC |
| 8_hypothetical | D12 | Tb927.11.16250_for | GGGGACAAGTTTGTACAAAAAAGCAGGCTATGCAGCGGGTTCCGCCG | Tb927.11.16250_rev | GGGGACCACTTTGTACAAGAAAGCTGGGTCTAATTTTCCTCCGATTTGACAGCGG |
| 8_hypothetical | D13 | Tb927.11.16370_for | GGGGACAAGTTTGTACAAAAAAGCAGGCTATGTTAACGCCCGTGTTCGTT | Tb927.11.16370_rev | GGGGACCACTTTGTACAAGAAAGCTGGGTTCACGAAATTGGGAGGCCTAAGT |
| 8_hypothetical | D14 | Tb927.6.1220_for | GGGGACAAGTTTGTACAAAAAAGCAGGCTATGGAATGCGCGTCGCAG | Tb927.6.1220_rev | GGGGACCACTTTGTACAAGAAAGCTGGGTTTACCACAGTGTGGAGGTCCTG |
| 8_hypothetical | D15 | Tb927.5.3370_for | GGGGACAAGTTTGTACAAAAAAGCAGGCTATGGACACTACAGGGAATTCTCCAG | Tb927.5.3370_rev | GGGGACCACTTTGTACAAGAAAGCTGGGTTTACCTTTGTAATGATGGCGACTTCG |
| 8_hypothetical | D16 | Tb927.11.9210_for | GGGGACAAGTTTGTACAAAAAAGCAGGCTATGGATGTTTATCGACAAGCAAGTGA | Tb927.11.9210_rev | GGGGACCACTTTGTACAAGAAAGCTGGGTTCACCGCTTCTTCTTCGACCT |
| 8_hypothetical | D17 | Tb927.9.9280_for | GGGGACAAGTTTGTACAAAAAAGCAGGCTATGAGCAACAACCCATACCTTGC | Tb927.9.9280_rev | GGGGACCACTTTGTACAAGAAAGCTGGGTTCAGGGGCATCCCCCACT |
| 8_hypothetical | D18 | Tb927.3.5000_for | GGGGACAAGTTTGTACAAAAAAGCAGGCTATGTTCATTTATGTCCGCTGCTTACT | Tb927.3.5000_rev | GGGGACCACTTTGTACAAGAAAGCTGGGTCTAGAGCATACGTCGTCTCACCT |
| 8_hypothetical | D19 | Tb927.9.9220_for | GGGGACAAGTTTGTACAAAAAAGCAGGCTATGTCTTCAGATTGCGCTGGAC | Tb927.9.9220_rev | GGGGACCACTTTGTACAAGAAAGCTGGGTCTACCTATTTATGGCAGCTGTGCA |
| 8_hypothetical | D20 | Tb927.6.5080_for | GGGGACAAGTTTGTACAAAAAAGCAGGCTATGCATTGTAGGACAGCATCTGATG | Tb927.6.5080_rev | GGGGACCACTTTGTACAAGAAAGCTGGGTTTACCCATCATTAGTCTTTTGTTTAAGTTG |
| 8_hypothetical | D21 | Tb927.9.3130_for | GGGGACAAGTTTGTACAAAAAAGCAGGCTATGGATCTTGGTAGTGACGAGGATC | Tb927.9.3130_rev | GGGGACCACTTTGTACAAGAAAGCTGGGTCTATTGCTTTTCTGTTCCCCCCC |
| 8_hypothetical | D22 | Tb927.8.3940_for | GGGGACAAGTTTGTACAAAAAAGCAGGCTATGGTGAAGGTCGAACCCCC | Tb927.8.3940_rev | GGGGACCACTTTGTACAAGAAAGCTGGGTCTACTTCTGCACAAATCGGAGTATCC |
| 8_hypothetical | D23 | Tb927.11.4350_for | GGGGACAAGTTTGTACAAAAAAGCAGGCTATGCAGTCTAGAGTATCCCCGGG | Tb927.11.4350_rev | GGGGACCACTTTGTACAAGAAAGCTGGGTTCAGTTAGCTGTGACTGGCTCC |
| 8_hypothetical | D24 | Tb927.10.10740_for | GGGGACAAGTTTGTACAAAAAAGCAGGCTATGGGACTCAGGGACCTGGA | Tb927.10.10740_rev | GGGGACCACTTTGTACAAGAAAGCTGGGTTTACACAACAAGGTCGTTCGTTGA |
| 8_hypothetical | E1 | Tb927.8.3050_for | GGGGACAAGTTTGTACAAAAAAGCAGGCTATGACCACAGCAACCGTCAAG | Tb927.8.3050_rev | GGGGACCACTTTGTACAAGAAAGCTGGGTTCACTGTATGGCAAAGCCACG |
| 8_hypothetical | E2 | Tb927.11.14540_for | GGGGACAAGTTTGTACAAAAAAGCAGGCTATGGATACCGCGTACGCGA | Tb927.11.14540_rev | GGGGACCACTTTGTACAAGAAAGCTGGGTTTACAAGGGACAAAGTATTGTTTTTTTCTT |
| 8_hypothetical | E3 | Tb927.9.11970_for | GGGGACAAGTTTGTACAAAAAAGCAGGCTATGGCGATGCATCGGCTATG | Tb927.9.11970_rev | GGGGACCACTTTGTACAAGAAAGCTGGGTCTAGCAGAGTTTCATGGTTTTGTTTGG |
| 8_hypothetical | E4 | Tb927.11.10430_for | GGGGACAAGTTTGTACAAAAAAGCAGGCTATGCGAAACGATAAGTCGATGCC | Tb927.11.10430_rev | GGGGACCACTTTGTACAAGAAAGCTGGGTTCAGGTCTTGCGCTTATGCTTG |
| 8_hypothetical | E5 | Tb927.11.15140_for | GGGGACAAGTTTGTACAAAAAAGCAGGCTATGTTCCAACGCCTTGACGAG | Tb927.11.15140_rev | GGGGACCACTTTGTACAAGAAAGCTGGGTTCATTTTCGGGGTGTCACAAGATAAG |
| 8_hypothetical | E6 | Tb927.11.2030_for | GGGGACAAGTTTGTACAAAAAAGCAGGCTATGTCTTCATCTTTCTTGGTTCCTTTCA | Tb927.11.2030_rev | GGGGACCACTTTGTACAAGAAAGCTGGGTTCAACGGTTCAAGTAATATGGGTGATTC |
| 8_hypothetical | E7 | Tb927.11.1360_for | GGGGACAAGTTTGTACAAAAAAGCAGGCTATGACGATGTCGGACACAAGTG | Tb927.11.1360_rev | GGGGACCACTTTGTACAAGAAAGCTGGGTTTACAACCGTATTGGTGGGTTCCC |
| 8_hypothetical | E8 | Tb927.9.1880_for | GGGGACAAGTTTGTACAAAAAAGCAGGCTATGGCTAAACGCACAGTCACAG | Tb927.9.1880_rev | GGGGACCACTTTGTACAAGAAAGCTGGGTCTACAGATACGTAATACCTGGCGTTATG |
| 8_hypothetical | E9 | Tb927.7.760_for | GGGGACAAGTTTGTACAAAAAAGCAGGCTATGCCTCCGAAGAAAAGCGAG | Tb927.7.760_rev | GGGGACCACTTTGTACAAGAAAGCTGGGTTTAGTTGTCTTCACTCTCAAGGGGA |
| 8_hypothetical | E10 | Tb927.11.1040_for | GGGGACAAGTTTGTACAAAAAAGCAGGCTATGCTATTTTATCTGCATATCAATTTATTT | Tb927.11.1040_rev | GGGGACCACTTTGTACAAGAAAGCTGGGTTCACTCCTTCAAACGGTACTTTCTACC |
| 8_hypothetical | E11 | Tb927.7.5110_for | GGGGACAAGTTTGTACAAAAAAGCAGGCTATGGTGGTGGGTGGTGATCG | Tb927.7.5110_rev | GGGGACCACTTTGTACAAGAAAGCTGGGTTCACTTCACTATCTTGGGCATCAGC |
| 8_hypothetical | E12 | Tb927.10.10840_for | GGGGACAAGTTTGTACAAAAAAGCAGGCTATGTCCTTGTGGCTAACATTGTTTTCG | Tb927.10.10840_rev | GGGGACCACTTTGTACAAGAAAGCTGGGTTTACAAATGTATGATATTGTGGGGTAAAAG |
| 8_hypothetical | E13 | Tb927.4.2790_for | GGGGACAAGTTTGTACAAAAAAGCAGGCTATGATACACAGAAGTTTGCGTTCCA | Tb927.4.2790_rev | GGGGACCACTTTGTACAAGAAAGCTGGGTTCAGTCCACTTCACCGGTGG |
| 8_hypothetical | E14 | Tb927.11.1830_for | GGGGACAAGTTTGTACAAAAAAGCAGGCTATGTATAATGCCCTGAGGTCAGCA | Tb927.11.1830_rev | GGGGACCACTTTGTACAAGAAAGCTGGGTCTACGAAGCTTTCGCATCCCTAG |
| 8_hypothetical | E15 | Tb927.8.7470_for | GGGGACAAGTTTGTACAAAAAAGCAGGCTATGCAAAGCGGCAGTGAGAA | Tb927.8.7470_rev | GGGGACCACTTTGTACAAGAAAGCTGGGTCTACAGCCGCAACACATCCAT |
| 8_hypothetical | E16 | Tb927.6.1660_for | GGGGACAAGTTTGTACAAAAAAGCAGGCTATGTCGAGCAATCGCATGTTTGC | Tb927.6.1660_rev | GGGGACCACTTTGTACAAGAAAGCTGGGTTCACCAAATCCATTTCTCTTCAGAGAGG |
| 8_hypothetical | E17 | Tb927.11.1700_for | GGGGACAAGTTTGTACAAAAAAGCAGGCTATGTGGCGCCGCATCTCC | Tb927.11.1700_rev | GGGGACCACTTTGTACAAGAAAGCTGGGTTTAGTTATGTTCATCATCATCATCGTCGT |
| 8_hypothetical | E18 | Tb927.11.16670_for | GGGGACAAGTTTGTACAAAAAAGCAGGCTATGAGTACGGGCGGGGTG | Tb927.11.16670_rev | GGGGACCACTTTGTACAAGAAAGCTGGGTTCATCCAACCGTCTGCCCA |
| 8_hypothetical | E20 | Tb927.11.3010_for | GGGGACAAGTTTGTACAAAAAAGCAGGCTATGCGCTCCGTGTTGAACG | Tb927.11.3010_rev | GGGGACCACTTTGTACAAGAAAGCTGGGTCTAGCGTACACATAACCAAGGCTG |
| 8_hypothetical | E21 | Tb927.3.5400_for | GGGGACAAGTTTGTACAAAAAAGCAGGCTATGCCCACCCACAGAGATCC | Tb927.3.5400_rev | GGGGACCACTTTGTACAAGAAAGCTGGGTCTATTTCTTTTTCTTCGCCTTTCCATTTTT |
| 8_hypothetical | E22 | Tb927.4.2370_for | GGGGACAAGTTTGTACAAAAAAGCAGGCTATGGCGAAAGGTAATGAGGAGTGT | Tb927.4.2370_rev | GGGGACCACTTTGTACAAGAAAGCTGGGTTTATCGGAGTACGGCCGACTC |
| 8_hypothetical | E23 | Tb927.6.2390_for | GGGGACAAGTTTGTACAAAAAAGCAGGCTATGTCCTCTGTGTCAAGTGATCACA | Tb927.6.2390_rev | GGGGACCACTTTGTACAAGAAAGCTGGGTTCACACGCCTCGAACAGTCA |
| 8_hypothetical | E24 | Tb927.11.11610_for | GGGGACAAGTTTGTACAAAAAAGCAGGCTATGTTTAAACGCACCCGAGGTG | Tb927.11.11610_rev | GGGGACCACTTTGTACAAGAAAGCTGGGTCTACCGTCGATAATTGCTGTTGGT |
| 8_hypothetical | F1 | Tb927.8.6220_for | GGGGACAAGTTTGTACAAAAAAGCAGGCTATGAAACGGGGTAGTGATGGATGT | Tb927.8.6220_rev | GGGGACCACTTTGTACAAGAAAGCTGGGTTCACCGTCCGGCCGCCGT |
| 8_hypothetical | F2 | Tb927.10.14900_for | GGGGACAAGTTTGTACAAAAAAGCAGGCTATGGAGTACCCAGCGGAGTTC | Tb927.10.14900_rev | GGGGACCACTTTGTACAAGAAAGCTGGGTCTCACCCCTTGCGGCGAAT |
| 8_hypothetical | F3 | Tb927.11.8780_for | GGGGACAAGTTTGTACAAAAAAGCAGGCTATGGCGCTTTTATCACTGGACG | Tb927.11.8780_rev | GGGGACCACTTTGTACAAGAAAGCTGGGTTCATAACCGCAACACCAACCC |
| 8_hypothetical | F4 | Tb927.11.8060_for | GGGGACAAGTTTGTACAAAAAAGCAGGCTATGCAACCTATAGAACGTGCTGAG | Tb927.11.8060_rev | GGGGACCACTTTGTACAAGAAAGCTGGGTCTACAGCGTTTCCAGGCCAC |
| 8_hypothetical | F5 | Tb927.10.2720_for | GGGGACAAGTTTGTACAAAAAAGCAGGCTATGAGTTGCGCTGGATCCACA | Tb927.10.2720_rev | GGGGACCACTTTGTACAAGAAAGCTGGGTTTACGGCAACTCAATCTTACACGC |
| 8_hypothetical | F6 | Tb927.11.2570_for | GGGGACAAGTTTGTACAAAAAAGCAGGCTATGGTGGCGACAGGAACCT | Tb927.11.2570_rev | GGGGACCACTTTGTACAAGAAAGCTGGGTTTAGTAATCCTTCAAAACGTGTAGCATCA |
| 8_hypothetical | F7 | Tb927.5.1260_for | GGGGACAAGTTTGTACAAAAAAGCAGGCTATGGTGTCTTGTTTCGTGCGG | Tb927.5.1260_rev | GGGGACCACTTTGTACAAGAAAGCTGGGTTCAGTACAATTCTTGATACTCCAACTTGTG |
| 8_hypothetical | F8 | Tb927.11.10590_for | GGGGACAAGTTTGTACAAAAAAGCAGGCTATGACTTATCGTGTACTGGGCTCA | Tb927.11.10590_rev | GGGGACCACTTTGTACAAGAAAGCTGGGTTCAGACAACAATCGTCGAAGTGTT |
| 8_hypothetical | F9 | Tb927.11.15110_for | GGGGACAAGTTTGTACAAAAAAGCAGGCTATGATGCGTTTGGTTGCCTGC | Tb927.11.15110_rev | GGGGACCACTTTGTACAAGAAAGCTGGGTTTAATTAAGGGGGTTTAAATGTCGCATACA |
| 8_hypothetical | F10 | Tb927.10.10120_for | GGGGACAAGTTTGTACAAAAAAGCAGGCTATGCGTCGTTGCAAACTGTGG | Tb927.10.10120_rev | GGGGACCACTTTGTACAAGAAAGCTGGGTCTACATGGTGTCGCCCGAGA |
| 8_hypothetical | F11 | Tb927.7.4580_for | GGGGACAAGTTTGTACAAAAAAGCAGGCTATGCCGGACCCTCCGTTG | Tb927.7.4580_rev | GGGGACCACTTTGTACAAGAAAGCTGGGTCTAGTGCTGACGCCCTCTCA |
| 8_hypothetical | F12 | Tb927.11.5630_for | GGGGACAAGTTTGTACAAAAAAGCAGGCTATGACTGAGGTCTTCCGTGTGC | Tb927.11.5630_rev | GGGGACCACTTTGTACAAGAAAGCTGGGTCTACATAACTTTTGCGGCCAGCC |
| 8_hypothetical | F13 | Tb927.10.5400_for | GGGGACAAGTTTGTACAAAAAAGCAGGCTATGACACCCGGAGCAATGTACT | Tb927.10.5400_rev | GGGGACCACTTTGTACAAGAAAGCTGGGTCTACTGCACTTTTCTCTTATTGGCATTG |
| 8_hypothetical | F14 | Tb927.8.8110_for | GGGGACAAGTTTGTACAAAAAAGCAGGCTATGCGGACCAGAACAACAATGAC | Tb927.8.8110_rev | GGGGACCACTTTGTACAAGAAAGCTGGGTTCACTCTCGCAAATAGAAACTAAGCAC |
| 8_hypothetical | F15 | Tb927.10.2680_for | GGGGACAAGTTTGTACAAAAAAGCAGGCTATGTACCTTCCCAAACAACCGGT | Tb927.10.2680_rev | GGGGACCACTTTGTACAAGAAAGCTGGGTTTACAACATAGTTCCATCATCGCCAA |
| 8_hypothetical | F16 | Tb927.2.4970_for | GGGGACAAGTTTGTACAAAAAAGCAGGCTATGACGTGTTCGGTGGTTCCA | Tb927.2.4970_rev | GGGGACCACTTTGTACAAGAAAGCTGGGTTCACACTCCTTGATCATTTGGAGACG |
| 8_hypothetical | F17 | Tb927.5.3840_for | GGGGACAAGTTTGTACAAAAAAGCAGGCTATGCATGGAGCTTCAGTCGGA | Tb927.5.3840_rev | GGGGACCACTTTGTACAAGAAAGCTGGGTTTACAGTTCCCATATTTCTTCCTCTTCG |
| 8_hypothetical | F18 | Tb927.9.7580_for | GGGGACAAGTTTGTACAAAAAAGCAGGCTATGCCCTTTGATAATGTGACTGCTT | Tb927.9.7580_rev | GGGGACCACTTTGTACAAGAAAGCTGGGTTTACATAAGTATTCCAGATACATCGCTGGA |
| 8_hypothetical | F19 | Tb927.11.2370_for | GGGGACAAGTTTGTACAAAAAAGCAGGCTATGCCCAACCCATACAGGAAAAAC | Tb927.11.2370_rev | GGGGACCACTTTGTACAAGAAAGCTGGGTTCACTGCGGGACTGCCGC |
| 8_hypothetical | F20 | Tb927.2.2550_for | GGGGACAAGTTTGTACAAAAAAGCAGGCTATGTGTGGCCTTTTGCGCT | Tb927.2.2550_rev | GGGGACCACTTTGTACAAGAAAGCTGGGTTCACCAAAAAACGCGGAGCA |
| 8_hypothetical | F21 | Tb927.7.3770_for | GGGGACAAGTTTGTACAAAAAAGCAGGCTATGAGAAGGAACTCTTTTATTCGGACC | Tb927.7.3770_rev | GGGGACCACTTTGTACAAGAAAGCTGGGTCTAAAGGTAGCACTGAGAAGCTTGAA |
| 8_hypothetical | F22 | Tb927.7.5660_for | GGGGACAAGTTTGTACAAAAAAGCAGGCTATGCAAGCAAAGGGAGATGGG | Tb927.7.5660_rev | GGGGACCACTTTGTACAAGAAAGCTGGGTTTACTCTCCTCTAGCCGTTGCG |
| 8_hypothetical | F23 | Tb927.9.8060_for | GGGGACAAGTTTGTACAAAAAAGCAGGCTATGTACAAGGAGACCACTATGACACC | Tb927.9.8060_rev | GGGGACCACTTTGTACAAGAAAGCTGGGTTTACCAATCGTCATCATCGTCGGT |
| 8_hypothetical | F24 | Tb927.5.1250_for | GGGGACAAGTTTGTACAAAAAAGCAGGCTATGACCGAAAGCGTTGGAGC | Tb927.5.1250_rev | GGGGACCACTTTGTACAAGAAAGCTGGGTCTACGATGTGTTAATAGTGTTCCACCA |
| 8_hypothetical | G1 | Tb927.8.5130_for | GGGGACAAGTTTGTACAAAAAAGCAGGCTATGCAGCATCACTTCCACCTCA | Tb927.8.5130_rev | GGGGACCACTTTGTACAAGAAAGCTGGGTTCAGTAGGCTGAAACAACAAAAAAGAAC |
| 8_hypothetical | G2 | Tb927.5.3730_for | GGGGACAAGTTTGTACAAAAAAGCAGGCTATGGCGGCCCTTGTGGAA | Tb927.5.3730_rev | GGGGACCACTTTGTACAAGAAAGCTGGGTCTAATGGAGTCGGACATTCACTTCTTC |
| 8_hypothetical | G3 | Tb927.8.560_for | GGGGACAAGTTTGTACAAAAAAGCAGGCTATGGTACTGGCATCCATTCGTGT | Tb927.8.560_rev | GGGGACCACTTTGTACAAGAAAGCTGGGTCTACCGGCCTTTCTTCCGTG |
| 8_hypothetical | G4 | Tb927.4.1850_for | GGGGACAAGTTTGTACAAAAAAGCAGGCTATGGACTTTGATCTACTGTTGTTCCTAAAG | Tb927.4.1850_rev | GGGGACCACTTTGTACAAGAAAGCTGGGTTTACTTACGCCAACCCCAACTCC |
| 8_hypothetical | G5 | Tb927.11.6240_for | GGGGACAAGTTTGTACAAAAAAGCAGGCTATGCGTCCGAGGTTGTACCA | Tb927.11.6240_rev | GGGGACCACTTTGTACAAGAAAGCTGGGTTCAACAGCACCTTACTTCTTTTTTCCC |
| 8_hypothetical | G6 | Tb927.8.1470_for | GGGGACAAGTTTGTACAAAAAAGCAGGCTATGGAAAGGGGTATGTGCCTCG | Tb927.8.1470_rev | GGGGACCACTTTGTACAAGAAAGCTGGGTTCAAAGAATGGAAGCGACGCC |
| 8_hypothetical | G7 | Tb927.9.3580_for | GGGGACAAGTTTGTACAAAAAAGCAGGCTATGCATGTGGCCTCTCCCG | Tb927.9.3580_rev | GGGGACCACTTTGTACAAGAAAGCTGGGTCTAGTCAGAATCACTGTCACTGTCAC |

| | | | | | |
|---|---|---|---|---|---|
| 8_hypothetical | G8 | Tb927.6.610_for | GGGGACAAGTTTGTACAAAAAAGCAGGCTATGTTAATACGCCGCCTCCGC | Tb927.6.610_rev | GGGGACCACTTTGTACAAGAAAGCTGGGTTCAGAATGTGCCATCTGGATTGAC |
| 8_hypothetical | G9 | Tb927.11.9200_for | GGGGACAAGTTTGTACAAAAAAGCAGGCTATGGTGAAGTTGAAGGATGTTTCCC | Tb927.11.9200_rev | GGGGACCACTTTGTACAAGAAAGCTGGGTTTAATCGACATGAGTCCATAATTTAACAAA |
| 8_hypothetical | G10 | Tb927.3.5240_for | GGGGACAAGTTTGTACAAAAAAGCAGGCTATGCTTTCTCAGAATGTGGCCAA | Tb927.3.5240_rev | GGGGACCACTTTGTACAAGAAAGCTGGGTCTACCGTGACCCACCACCT |
| 8_hypothetical | G11 | Tb927.8.4670_for | GGGGACAAGTTTGTACAAAAAAGCAGGCTATGCCTCCTCGGAGGGCT | Tb927.8.4670_rev | GGGGACCACTTTGTACAAGAAAGCTGGGTTCAAAAGTTGTTGCTTAATGGGTTATCG |
| 8_hypothetical | G12 | Tb927.11.4000_for | GGGGACAAGTTTGTACAAAAAAGCAGGCTATGTTGGCGTCATGGGCCG | Tb927.11.4000_rev | GGGGACCACTTTGTACAAGAAAGCTGGGTCTAATATAGTTTCTTAGACAAAGTCCAGTG |
| 8_hypothetical | G13 | Tb927.10.4800_for | GGGGACAAGTTTGTACAAAAAAGCAGGCTATGGGTTCGTCGGGGTATGAC | Tb927.10.4800_rev | GGGGACCACTTTGTACAAGAAAGCTGGGTCTACGCAGAAGAACATTGACTTGTATCC |
| 8_hypothetical | G14 | Tb927.10.6250_for | GGGGACAAGTTTGTACAAAAAAGCAGGCTATGGCTCACAGTGGGGCC | Tb927.10.6250_rev | GGGGACCACTTTGTACAAGAAAGCTGGGTTCAGAAGAACACACTCCCCATACC |
| 8_hypothetical | G15 | Tb927.6.1980_for | GGGGACAAGTTTGTACAAAAAAGCAGGCTATGACTATTCCCGCCGTATACTCC | Tb927.6.1980_rev | GGGGACCACTTTGTACAAGAAAGCTGGGTTCACGACCGCCTCGCTCC |
| 8_hypothetical | G16 | Tb927.11.9820_for | GGGGACAAGTTTGTACAAAAAAGCAGGCTATGATCCGCGTGTATGTTGCTG | Tb927.11.9820_rev | GGGGACCACTTTGTACAAGAAAGCTGGGTTCATTTAAAGGGGTGAAAAACTGAATCGT |
| 8_hypothetical | G17 | Tb927.3.4080_for | GGGGACAAGTTTGTACAAAAAAGCAGGCTATGGTTTACCTTGTCGATGACCTTG | Tb927.3.4080_rev | GGGGACCACTTTGTACAAGAAAGCTGGGTTTAATTTTCTTTTTCTGAATCCTCTGTAAC |
| 8_hypothetical | G18 | Tb927.4.570_for | GGGGACAAGTTTGTACAAAAAAGCAGGCTATGTTCCAATTAACGACAGGATTCTGC | Tb927.4.570_rev | GGGGACCACTTTGTACAAGAAAGCTGGGTTTACAAACCTCCCTCACTCATTATTTGC |
| 8_hypothetical | G19 | Tb927.6.1560_for | GGGGACAAGTTTGTACAAAAAAGCAGGCTATGCCAGAGGAACTTAGGCAGT | Tb927.6.1560_rev | GGGGACCACTTTGTACAAGAAAGCTGGGTTCACACGGCTTGCTTTACTTTCT |
| 8_hypothetical | G20 | Tb927.11.7680_for | GGGGACAAGTTTGTACAAAAAAGCAGGCTATGCACAAGAGAGTCGGTTATACTGA | Tb927.11.7680_rev | GGGGACCACTTTGTACAAGAAAGCTGGGTTCAGAGATCATTCTGGGTTGATTGACC |
| 8_hypothetical | G21 | Tb927.8.3230_for | GGGGACAAGTTTGTACAAAAAAGCAGGCTATGAATCGAAAAAAAAGGTCCGTTGC | Tb927.8.3230_rev | GGGGACCACTTTGTACAAGAAAGCTGGGTTCAGCTTCCTCGTAACCTAAACGT |
| 8_hypothetical | G22 | Tb927.10.2600_for | GGGGACAAGTTTGTACAAAAAAGCAGGCTATGGTCGGTGAGGAGGTCG | Tb927.10.2600_rev | GGGGACCACTTTGTACAAGAAAGCTGGGTTCAAGTTTCTCGTGGAATTTTCTGACT |
| 8_hypothetical | G23 | Tb927.4.2910_for | GGGGACAAGTTTGTACAAAAAAGCAGGCTATGAGGCGTTGTGTTAGAGCAG | Tb927.4.2910_rev | GGGGACCACTTTGTACAAGAAAGCTGGGTTCACACTTCTCCTCCATCTGCA |
| 8_hypothetical | G24 | Tb927.10.8830_for | GGGGACAAGTTTGTACAAAAAAGCAGGCTATGGTATTCGACAAAGAGTTGAGGAATG | Tb927.10.8830_rev | GGGGACCACTTTGTACAAGAAAGCTGGGTCTAATTTCGTCGCACATTTTCAGAAGG |
| 8_hypothetical | H1 | Tb927.3.4100_for | GGGGACAAGTTTGTACAAAAAAGCAGGCTATGGTTTACCTTGTCGATGACCTTG | Tb927.3.4100_rev | GGGGACCACTTTGTACAAGAAAGCTGGGTTTATTGCTTCTTTCCTGAATGTCCACC |
| 8_hypothetical | H2 | Tb927.9.9010_for | GGGGACAAGTTTGTACAAAAAAGCAGGCTATGAAGACGAAGGACGGTAGTTGC | Tb927.9.9010_rev | GGGGACCACTTTGTACAAGAAAGCTGGGTCTAGGCGTCGCGGCGCGT |
| 8_hypothetical | H3 | Tb927.10.8850_for | GGGGACAAGTTTGTACAAAAAAGCAGGCTATGCCCGTCCATGCCCAT | Tb927.10.8850_rev | GGGGACCACTTTGTACAAGAAAGCTGGGTCTACTCATGAAGATCCGCCACAAA |
| 8_hypothetical | H4 | Tb927.9.10490_for | GGGGACAAGTTTGTACAAAAAAGCAGGCTATGACCGACACGCAAAAGGC | Tb927.9.10490_rev | GGGGACCACTTTGTACAAGAAAGCTGGGTCTAATCCAGAGCAAACAAATAGCGATTG |
| 8_hypothetical | H5 | Tb927.11.11700_for | GGGGACAAGTTTGTACAAAAAAGCAGGCTATGCAAAACCAGGCTCCAGTG | Tb927.11.11700_rev | GGGGACCACTTTGTACAAGAAAGCTGGGTTCAAACACGGATGACTGAATCTTTCTT |
| 8_hypothetical | H6 | Tb927.5.4340_for | GGGGACAAGTTTGTACAAAAAAGCAGGCTATGGTGACGCTGCGCTATTTAC | Tb927.5.4340_rev | GGGGACCACTTTGTACAAGAAAGCTGGGTCTATAACTCTGAGTGCAGTCGCAAC |
| 8_hypothetical | H7 | Tb927.5.680_for | GGGGACAAGTTTGTACAAAAAAGCAGGCTATGTGGCGAAGGCCACCG | Tb927.5.680_rev | GGGGACCACTTTGTACAAGAAAGCTGGGTCTACTGGCTGTAACGCATCCG |
| 8_hypothetical | H8 | Tb927.9.12850_for | GGGGACAAGTTTGTACAAAAAAGCAGGCTATGCACTGGCCGCACGGA | Tb927.9.12850_rev | GGGGACCACTTTGTACAAGAAAGCTGGGTTCACCTGTCGTAATTCTCCGGT |
| 8_hypothetical | H9 | Tb927.9.12830_for | GGGGACAAGTTTGTACAAAAAAGCAGGCTATGCAGAAAAATGCACCCAAGTTTT | Tb927.9.12830_rev | GGGGACCACTTTGTACAAGAAAGCTGGGTCTAACCAAAGGGGCCATGTATTTCC |
| 8_hypothetical | H10 | Tb927.4.2060_for | GGGGACAAGTTTGTACAAAAAAGCAGGCTATGGAGGCTTCTGTTGGAGTGG | Tb927.4.2060_rev | GGGGACCACTTTGTACAAGAAAGCTGGGTTTACGGAGCAGAAGGCAACCT |
| 8_hypothetical | H11 | Tb927.10.8800_for | GGGGACAAGTTTGTACAAAAAAGCAGGCTATGGCTTCGTTATCACCCCTGG | Tb927.10.8800_rev | GGGGACCACTTTGTACAAGAAAGCTGGGTTTAATCCGTGAGAAATGTGTTTACTTCAGG |
| 8_hypothetical | H12 | Tb927.5.3500_for | GGGGACAAGTTTGTACAAAAAAGCAGGCTATGGAGAGCGTGTCGCTTATCC | Tb927.5.3500_rev | GGGGACCACTTTGTACAAGAAAGCTGGGTCTAAACGCAGTCATGTAGGAAATAAACAC |
| 8_hypothetical | H13 | Tb927.8.1740_for | GGGGACAAGTTTGTACAAAAAAGCAGGCTATGCGCCGAGGTTTCTTTCTT | Tb927.8.1740_rev | GGGGACCACTTTGTACAAGAAAGCTGGGTCTAGTGTGTGGGAGTCTTCGCC |
| 8_hypothetical | H14 | Tb927.10.13090_for | GGGGACAAGTTTGTACAAAAAAGCAGGCTATGTTAAGGCGAGGCGTTTACAG | Tb927.10.13090_rev | GGGGACCACTTTGTACAAGAAAGCTGGGTTTACGCCTTCTTTGCCTTTCTGT |
| 8_hypothetical | H15 | Tb927.2.3730_for | GGGGACAAGTTTGTACAAAAAAGCAGGCTATGAGTTACACTGAAGGAGATGGGAA | Tb927.2.3730_rev | GGGGACCACTTTGTACAAGAAAGCTGGGTTCATCCGTCTGTTTTGACACTATTCG |
| 8_hypothetical | H16 | Tb927.10.2240_for | GGGGACAAGTTTGTACAAAAAAGCAGGCTATGTCCATTACACAAAGTGCACTCG | Tb927.10.2240_rev | GGGGACCACTTTGTACAAGAAAGCTGGGTCTAGGGATAGAACCGTGCCACC |
| 8_hypothetical | H17 | Tb927.7.3190_for | GGGGACAAGTTTGTACAAAAAAGCAGGCTATGCAGTGGAAAGAGGTATTAGATAGCG | Tb927.7.3190_rev | GGGGACCACTTTGTACAAGAAAGCTGGGTTCAATCATCTTCCAAAGAGTTTCCGTC |
| 8_hypothetical | H18 | Tb927.11.12170_for | GGGGACAAGTTTGTACAAAAAAGCAGGCTATGTCCGTGGCCTCTGAGG | Tb927.11.12170_rev | GGGGACCACTTTGTACAAGAAAGCTGGGTTCACGAACCCAACTGTGACGA |
| 8_hypothetical | H19 | Tb927.3.5220_for | GGGGACAAGTTTGTACAAAAAAGCAGGCTATGTCCTCCGCTAATTCCATCCA | Tb927.3.5220_rev | GGGGACCACTTTGTACAAGAAAGCTGGGTTTAAAGCACCGGCACTGCG |
| 8_hypothetical | H20 | Tb927.11.7870_for | GGGGACAAGTTTGTACAAAAAAGCAGGCTATGACCAAATGGAGGTGGAAAACAC | Tb927.11.7870_rev | GGGGACCACTTTGTACAAGAAAGCTGGGTCTAAACACCAATGTCACAGGCAGT |
| 8_hypothetical | H21 | Tb927.4.4370_for | GGGGACAAGTTTGTACAAAAAAGCAGGCTATGCTCATTTGCCCCACCG | Tb927.4.4370_rev | GGGGACCACTTTGTACAAGAAAGCTGGGTTCACAACTCCGCACCCTCA |
| 8_hypothetical | H22 | Tb927.9.11520_for | GGGGACAAGTTTGTACAAAAAAGCAGGCTATGGATGCTGATCTTAACGGTTTGC | Tb927.9.11520_rev | GGGGACCACTTTGTACAAGAAAGCTGGGTTCAATCGACCAACTCGCCACC |
| 8_hypothetical | H23 | Tb927.8.3520_for | GGGGACAAGTTTGTACAAAAAAGCAGGCTATGAACTTCTCTGGTGGAAACAAGG | Tb927.8.3520_rev | GGGGACCACTTTGTACAAGAAAGCTGGGTTTACTTCGATTCGGGTTGATTTTGAAATAC |
| 8_hypothetical | H24 | Tb927.10.9930_for | GGGGACAAGTTTGTACAAAAAAGCAGGCTATGCGTATTGTAATCGATGTCGATGC | Tb927.10.9930_rev | GGGGACCACTTTGTACAAGAAAGCTGGGTCTAATTGTCGTCGGCGATACCAC |
| 8_hypothetical | I1 | Tb927.3.1910_for | GGGGACAAGTTTGTACAAAAAAGCAGGCTATGTCCATGACTGAGAGCAGCA | Tb927.3.1910_rev | GGGGACCACTTTGTACAAGAAAGCTGGGTCTACTGACGCTGCTGGCG |
| 8_hypothetical | I2 | Tb927.11.16580_for | GGGGACAAGTTTGTACAAAAAAGCAGGCTATGCTCAACCCGGGTGATCA | Tb927.11.16580_rev | GGGGACCACTTTGTACAAGAAAGCTGGGTTCACAGCACCACGTACAAAAACC |
| 8_hypothetical | I3 | Tb927.9.12160_for | GGGGACAAGTTTGTACAAAAAAGCAGGCTATGCGCCGCTTTTGTGGG | Tb927.9.12160_rev | GGGGACCACTTTGTACAAGAAAGCTGGGTTCAGTTGCCCCCCCTCTCC |
| 8_hypothetical | I4 | Tb927.10.10300_for | GGGGACAAGTTTGTACAAAAAAGCAGGCTATGTCATGGATGAGCGCGC | Tb927.10.10300_rev | GGGGACCACTTTGTACAAGAAAGCTGGGTTTAAGCCGGAAGAAAGTGTATGGC |
| 8_hypothetical | I5 | Tb927.7.3340_for | GGGGACAAGTTTGTACAAAAAAGCAGGCTATGAGGTACTTCCCACTCGGTG | Tb927.7.3340_rev | GGGGACCACTTTGTACAAGAAAGCTGGGTTCACATCTCGGATGGATGGTGC |
| 8_hypothetical | I6 | Tb927.6.730_for | GGGGACAAGTTTGTACAAAAAAGCAGGCTATGAATGCTGCATCCCCTCCT | Tb927.6.730_rev | GGGGACCACTTTGTACAAGAAAGCTGGGTTCAAACCATTGGAAGCTCCAGGA |
| 8_hypothetical | I7 | Tb927.3.2210_for | GGGGACAAGTTTGTACAAAAAAGCAGGCTATGGACCACACTGCGTTCAC | Tb927.3.2210_rev | GGGGACCACTTTGTACAAGAAAGCTGGGTTCAAGCAGTACGGTTGTACACAC |
| 8_hypothetical | I8 | Tb927.8.6710_for | GGGGACAAGTTTGTACAAAAAAGCAGGCTATGTCCTCCGTTACCACTGGC | Tb927.8.6710_rev | GGGGACCACTTTGTACAAGAAAGCTGGGTTCATTCTCGAACGTATTCCTTATTCTCCC |
| 8_hypothetical | I9 | Tb927.8.6730_for | GGGGACAAGTTTGTACAAAAAAGCAGGCTATGTCCTCCGTTACCACTGGC | Tb927.8.6730_rev | GGGGACCACTTTGTACAAGAAAGCTGGGTTTATTTCCTTGCAGGGCTTTTGTCA |
| 8_hypothetical | I10 | Tb927.8.5080_for | GGGGACAAGTTTGTACAAAAAAGCAGGCTATGCTTACTCTGGCTATCATATGTGTCA | Tb927.8.5080_rev | GGGGACCACTTTGTACAAGAAAGCTGGGTTTACTCGAGAATAGCCCAGCCAG |
| 8_hypothetical | I11 | Tb927.3.4070_for | GGGGACAAGTTTGTACAAAAAAGCAGGCTATGGTTTACCTTGTCGATGACCTTG | Tb927.3.4070_rev | GGGGACCACTTTGTACAAGAAAGCTGGGTCTACTGCTGCGCTGCTCC |
| 8_hypothetical | I12 | Tb927.10.3810_for | GGGGACAAGTTTGTACAAAAAAGCAGGCTATGTGGAAGCAAGATATTGGGCAG | Tb927.10.3810_rev | GGGGACCACTTTGTACAAGAAAGCTGGGTTCAGAGTGTTTTTGCCCTGTATTTGT |
| 8_hypothetical | I13 | Tb927.3.1850_for | GGGGACAAGTTTGTACAAAAAAGCAGGCTATGAGGGGGAACTTAATTACCGGG | Tb927.3.1850_rev | GGGGACCACTTTGTACAAGAAAGCTGGGTCTAAGAACTCACATCAAGATTAAAGAACGG |
| 8_hypothetical | I14 | Tb927.11.5210_for | GGGGACAAGTTTGTACAAAAAAGCAGGCTATGGCCGAACTACGGGAAGG | Tb927.11.5210_rev | GGGGACCACTTTGTACAAGAAAGCTGGGTTTAGTCGGCATTGGCTGCAG |
| 8_hypothetical | I15 | Tb927.10.12600_for | GGGGACAAGTTTGTACAAAAAAGCAGGCTATGTCCGTATATGGGAACGGCG | Tb927.10.12600_rev | GGGGACCACTTTGTACAAGAAAGCTGGGTCTACGCCACATTATTCCAGCAACT |
| 8_hypothetical | I16 | Tb927.7.900_for | GGGGACAAGTTTGTACAAAAAAGCAGGCTATGTTTTTGTCGTTGGCACTACTGG | Tb927.7.900_rev | GGGGACCACTTTGTACAAGAAAGCTGGGTTCAACTGCTTCCACTGCTAGCA |
| 8_hypothetical | I17 | Tb927.3.4110_for | GGGGACAAGTTTGTACAAAAAAGCAGGCTATGACGTATATTATCGATGACCTTGCG | Tb927.3.4110_rev | GGGGACCACTTTGTACAAGAAAGCTGGGTCTACTGCTGCGCTGCTCC |
| 8_hypothetical | I19 | Tb927.5.830_for | GGGGACAAGTTTGTACAAAAAAGCAGGCTATGCTGTACAAAGACCTCCTGCA | Tb927.5.830_rev | GGGGACCACTTTGTACAAGAAAGCTGGGTCTACTCACCCCTCAATCGGACC |
| 8_hypothetical | I20 | Tb927.11.14550_for | GGGGACAAGTTTGTACAAAAAAGCAGGCTATGGAAAATTTGGTGGACGTTGGA | Tb927.11.14550_rev | GGGGACCACTTTGTACAAGAAAGCTGGGTCTATGCGTGGGGACACCTTAAAAT |
| 8_hypothetical | I21 | Tb927.6.580_for | GGGGACAAGTTTGTACAAAAAAGCAGGCTATGATAAAACTCGTGTCGGGTGTC | Tb927.6.580_rev | GGGGACCACTTTGTACAAGAAAGCTGGGTTCAAAGCACATCTTCACTCAGCAA |
| 8_hypothetical | I22 | Tb927.11.9680_for | GGGGACAAGTTTGTACAAAAAAGCAGGCTATGGTGACGCTGGGGAAGAA | Tb927.11.9680_rev | GGGGACCACTTTGTACAAGAAAGCTGGGTTTATTGAGAAAACCCATACTTGCTCATAAT |
| 8_hypothetical | I23 | Tb927.10.9070_for | GGGGACAAGTTTGTACAAAAAAGCAGGCTATGACCGCGAGGTTCACTGC | Tb927.10.9070_rev | GGGGACCACTTTGTACAAGAAAGCTGGGTCTATTGAGAGACAATGTAGTCAAGAAGGTT |
| 8_hypothetical | I24 | Tb927.7.3370_for | GGGGACAAGTTTGTACAAAAAAGCAGGCTATGCAGGGGGGATTTGGTCG | Tb927.7.3370_rev | GGGGACCACTTTGTACAAGAAAGCTGGGTTCAGCGGTCACCAACCATCA |
| 8_hypothetical | J1 | Tb927.2.4460_for | GGGGACAAGTTTGTACAAAAAAGCAGGCTATGGCGCTACAGCGACGC | Tb927.2.4460_rev | GGGGACCACTTTGTACAAGAAAGCTGGGTTTACATGATGGAGTAAAGTTCTTCAGCAG |
| 8_hypothetical | J2 | Tb927.3.4660_for | GGGGACAAGTTTGTACAAAAAAGCAGGCTATGTCAGTTCATCAACTGCAACTTGA | Tb927.3.4660_rev | GGGGACCACTTTGTACAAGAAAGCTGGGTCTAGTTGTTTGACTTCCATAACCGTATGC |
| 8_hypothetical | J3 | Tb927.10.6520_for | GGGGACAAGTTTGTACAAAAAAGCAGGCTATGGACGGTGGCTTCCCTT | Tb927.10.6520_rev | GGGGACCACTTTGTACAAGAAAGCTGGGTTTAGCGCAGAGGCATGGCC |
| 8_hypothetical | J4 | Tb927.11.10180_for | GGGGACAAGTTTGTACAAAAAAGCAGGCTATGCACCTACTCGGTGATATTGTAGC | Tb927.11.10180_rev | GGGGACCACTTTGTACAAGAAAGCTGGGTCTACAGTGTAACCGAGACATTTTCGT |
| 8_hypothetical | J5 | Tb927.4.2590_for | GGGGACAAGTTTGTACAAAAAAGCAGGCTATGAGGGCGGAGCCGAAT | Tb927.4.2590_rev | GGGGACCACTTTGTACAAGAAAGCTGGGTTCATAAGCTTTCCCTGCCGCG |
| 8_hypothetical | J6 | Tb927.10.11600_for | GGGGACAAGTTTGTACAAAAAAGCAGGCTATGGAGTATTCGTCAGATGCAGTGT | Tb927.10.11600_rev | GGGGACCACTTTGTACAAGAAAGCTGGGTCTATTGATAACGGTGATGGGTAAGAGC |
| 8_hypothetical | J7 | Tb927.10.13450_for | GGGGACAAGTTTGTACAAAAAAGCAGGCTATGCTCGACGAGAATCAACGAG | Tb927.10.13450_rev | GGGGACCACTTTGTACAAGAAAGCTGGGTTCACTGCTCGTCTTCACTGTCC |
| 8_hypothetical | J8 | Tb927.1.4370_for | GGGGACAAGTTTGTACAAAAAAGCAGGCTATGGATACTCTCTTTTATTTTCCTGTTGGA | Tb927.1.4370_rev | GGGGACCACTTTGTACAAGAAAGCTGGGTCTAATGATCACCCGCCTCCTGA |
| 8_hypothetical | J9 | Tb927.3.1010_for | GGGGACAAGTTTGTACAAAAAAGCAGGCTATGTTGGTCGAGGAACAGACTGG | Tb927.3.1010_rev | GGGGACCACTTTGTACAAGAAAGCTGGGTTTAAGCAGTCGATTCCTTCTCTGAAGC |
| 8_hypothetical | J10 | Tb927.7.2410_for | GGGGACAAGTTTGTACAAAAAAGCAGGCTATGAGGCAGTTAGGAAGTGCAGT | Tb927.7.2410_rev | GGGGACCACTTTGTACAAGAAAGCTGGGTTTATTTGACCCTCACCTCATTACCCA |
| 8_hypothetical | J11 | Tb927.9.1700_for | GGGGACAAGTTTGTACAAAAAAGCAGGCTATGGAACTACTTGAGACGGTGGA | Tb927.9.1700_rev | GGGGACCACTTTGTACAAGAAAGCTGGGTTTACAATTGGGTTTGTCCAATTGATTGG |
| 8_hypothetical | J12 | Tb927.10.5600_for | GGGGACAAGTTTGTACAAAAAAGCAGGCTATGCATCACACGGGGATGC | Tb927.10.5600_rev | GGGGACCACTTTGTACAAGAAAGCTGGGTCTAAGCGCCCCGCTGCAA |
| 8_hypothetical | J13 | Tb927.4.3470_for | GGGGACAAGTTTGTACAAAAAAGCAGGCTATGTCTCTTCGTGACGATATCTTGGC | Tb927.4.3470_rev | GGGGACCACTTTGTACAAGAAAGCTGGGTCTATGCACTCCCATGCATGCTC |
| 8_hypothetical | J14 | Tb927.8.1500_for | GGGGACAAGTTTGTACAAAAAAGCAGGCTATGGCAATGCAGTTATTTACCTTTGGA | Tb927.8.1500_rev | GGGGACCACTTTGTACAAGAAAGCTGGGTTCAGTCATCGCAGATTCCCTTGA |
| 8_hypothetical | J15 | Tb927.10.10020_for | GGGGACAAGTTTGTACAAAAAAGCAGGCTATGAATTCTGCTGCGAGTGAAGAG | Tb927.10.10020_rev | GGGGACCACTTTGTACAAGAAAGCTGGGTTTACGACCGGGCGTTCCG |
| 8_hypothetical | J16 | Tb927.9.10150_for | GGGGACAAGTTTGTACAAAAAAGCAGGCTATGGATGCGCCATCGCTG | Tb927.9.10150_rev | GGGGACCACTTTGTACAAGAAAGCTGGGTTCAGCACATACTTGGGGTACTATCC |
| 8_hypothetical | J17 | Tb927.9.11120_for | GGGGACAAGTTTGTACAAAAAAGCAGGCTATGCTTCGACGGGCTTACATC | Tb927.9.11120_rev | GGGGACCACTTTGTACAAGAAAGCTGGGTTCACAACTCGGGAATCGCCT |
| 8_hypothetical | J18 | Tb927.11.16050_for | GGGGACAAGTTTGTACAAAAAAGCAGGCTATGCCAGTCAAACCTGTATACGAGT | Tb927.11.16050_rev | GGGGACCACTTTGTACAAGAAAGCTGGGTTCAAGTGGCAATCGCTGGTG |
| 8_hypothetical | J19 | Tb927.9.2030_for | GGGGACAAGTTTGTACAAAAAAGCAGGCTATGATGGAAAATACCCCGCTATTGG | Tb927.9.2030_rev | GGGGACCACTTTGTACAAGAAAGCTGGGTTCACAGGGATAAAATGCCCTTGAGT |
| 8_hypothetical | J20 | Tb927.6.1890_for | GGGGACAAGTTTGTACAAAAAAGCAGGCTATGCGGCGGTGCTTATGG | Tb927.6.1890_rev | GGGGACCACTTTGTACAAGAAAGCTGGGTTTATCGACGACGATGGTGCGC |
| 8_hypothetical | J21 | Tb927.11.14990_for | GGGGACAAGTTTGTACAAAAAAGCAGGCTATGGGCGGCGGTACTAGAG | Tb927.11.14990_rev | GGGGACCACTTTGTACAAGAAAGCTGGGTTCAAAATAATTCGTCCAGTATTGAGTCCC |
| 8_hypothetical | J22 | Tb927.10.1330_for | GGGGACAAGTTTGTACAAAAAAGCAGGCTATGAACTTGTCCAATCGGGTTGTG | Tb927.10.1330_rev | GGGGACCACTTTGTACAAGAAAGCTGGGTCTAGTTTGTCGGTTCCGCCTG |
| 8_hypothetical | J23 | Tb927.10.810_for | GGGGACAAGTTTGTACAAAAAAGCAGGCTATGGGGGCGGAAAAAAAAATAAATCG | Tb927.10.810_rev | GGGGACCACTTTGTACAAGAAAGCTGGGTTTAATTATCACGCTTCTTTTTCGCATCAT |
| 8_hypothetical | J24 | Tb927.6.660_for | GGGGACAAGTTTGTACAAAAAAGCAGGCTATGGATCTCCATTCCCCCTCCG | Tb927.6.660_rev | GGGGACCACTTTGTACAAGAAAGCTGGGTTTAAGTGCGGACCCAACCTCT |
| 8_hypothetical | K1 | Tb927.6.4450_for | GGGGACAAGTTTGTACAAAAAAGCAGGCTATGATTCGGCGCCTCAATGTG | Tb927.6.4450_rev | GGGGACCACTTTGTACAAGAAAGCTGGGTCTAACCTGTAGAACCGCTTGTTGAT |
| 8_hypothetical | K2 | Tb927.4.1490_for | GGGGACAAGTTTGTACAAAAAAGCAGGCTATGTCAGACGGCGGTATCTCC | Tb927.4.1490_rev | GGGGACCACTTTGTACAAGAAAGCTGGGTTCACGCGTGTTTCCATCTGC |
| 8_hypothetical | K3 | Tb927.8.3020_for | GGGGACAAGTTTGTACAAAAAAGCAGGCTATGGAGACGAAGCGTAGTTTCGA | Tb927.8.3020_rev | GGGGACCACTTTGTACAAGAAAGCTGGGTTCAACCACGGTGGCGCCC |
| 8_hypothetical | K5 | Tb927.10.9330_for | GGGGACAAGTTTGTACAAAAAAGCAGGCTATGGAACACGAAATTCATAAGGGAATGG | Tb927.10.9330_rev | GGGGACCACTTTGTACAAGAAAGCTGGGTTTACATTCTTCCCGGTACATTTTGAGG |
| 8_hypothetical | K6 | Tb927.8.6640_for | GGGGACAAGTTTGTACAAAAAAGCAGGCTATGCCACTCGACGTCCATGA | Tb927.8.6640_rev | GGGGACCACTTTGTACAAGAAAGCTGGGTTCAAAGGGACGAGGCCCTC |
| 8_hypothetical | K7 | Tb927.9.8350_for | GGGGACAAGTTTGTACAAAAAAGCAGGCTATGGGGGCTGGAGCGTCG | Tb927.9.8350_rev | GGGGACCACTTTGTACAAGAAAGCTGGGTCTACTCGGGTTTCCCCGCT |
| 8_hypothetical | K8 | Tb927.6.2580_for | GGGGACAAGTTTGTACAAAAAAGCAGGCTATGCTGTTCTGGTGTCGTCATG | Tb927.6.2580_rev | GGGGACCACTTTGTACAAGAAAGCTGGGTTTATGCACCACTACCAGTATAGTAACATCC |
| 8_hypothetical | K9 | Tb927.11.6670_for | GGGGACAAGTTTGTACAAAAAAGCAGGCTATGACACAAAAAAATGTGCTGTGTCA | Tb927.11.6670_rev | GGGGACCACTTTGTACAAGAAAGCTGGGTCTACAGACCATCTGCCGTCATG |
| 8_hypothetical | K10 | Tb927.4.3400_for | GGGGACAAGTTTGTACAAAAAAGCAGGCTATGGATATTCGGGAGATTATTGATCTTGTG | Tb927.4.3400_rev | GGGGACCACTTTGTACAAGAAAGCTGGGTCTATCGGCGAGGGCGCTT |
| 8_hypothetical | K11 | Tb927.11.10490_for | GGGGACAAGTTTGTACAAAAAAGCAGGCTATGTTTAGTGAGTCTTTGTACTCGTTTCA | Tb927.11.10490_rev | GGGGACCACTTTGTACAAGAAAGCTGGGTTCAAAGTTTCCATTTGTAGTACTCAAGCT |
| 8_hypothetical | K12 | Tb927.11.14640_for | GGGGACAAGTTTGTACAAAAAAGCAGGCTATGCCGTTGGGCTTGGATG | Tb927.11.14640_rev | GGGGACCACTTTGTACAAGAAAGCTGGGTTTAGGGAGTCCAAAAAACATTACCGG |
| 8_hypothetical | K13 | Tb927.7.2320_for | GGGGACAAGTTTGTACAAAAAAGCAGGCTATGAGCATCGTCAATCAGCTCAC | Tb927.7.2320_rev | GGGGACCACTTTGTACAAGAAAGCTGGGTTCACTTGACACGACTGCAACC |
| 8_hypothetical | K14 | Tb927.8.6270_for | GGGGACAAGTTTGTACAAAAAAGCAGGCTATGCGGGGAATATCAGCTCTGT | Tb927.8.6270_rev | GGGGACCACTTTGTACAAGAAAGCTGGGTCTACGCATCCGTCATGACGG |
| 8_hypothetical | K15 | Tb927.7.1770_for | GGGGACAAGTTTGTACAAAAAAGCAGGCTATGGATCGTATATACAGCCAGCCG | Tb927.7.1770_rev | GGGGACCACTTTGTACAAGAAAGCTGGGTCTATGTTTCATCATAACAACGTGCTACAC |
| 8_hypothetical | K16 | Tb927.10.15330_for | GGGGACAAGTTTGTACAAAAAAGCAGGCTATGACACGCATGCGAGTGG | Tb927.10.15330_rev | GGGGACCACTTTGTACAAGAAAGCTGGGTTTACGCTGCAGCCAAGGAAG |
| 8_hypothetical | K17 | Tb927.10.980_for | GGGGACAAGTTTGTACAAAAAAGCAGGCTATGGCCGCCGTGGAGCCA | Tb927.10.980_rev | GGGGACCACTTTGTACAAGAAAGCTGGGTTCATCCGTCATCTGGATGCTGG |
| 8_hypothetical | K18 | Tb927.9.2650_for | GGGGACAAGTTTGTACAAAAAAGCAGGCTATGACTAGCCAGCTGAATGCG | Tb927.9.2650_rev | GGGGACCACTTTGTACAAGAAAGCTGGGTTCACTTAGTCTTTCTCCTCCGCC |
| 8_hypothetical | K19 | Tb927.9.13130_for | GGGGACAAGTTTGTACAAAAAAGCAGGCTATGACGAAAGATAAAGTTAACTTTAGGTAC | Tb927.9.13130_rev | GGGGACCACTTTGTACAAGAAAGCTGGGTTTACTTCAGAAGCCTGAATATCGCTC |
| 8_hypothetical | K20 | Tb927.11.16620_for | GGGGACAAGTTTGTACAAAAAAGCAGGCTATGCTCACGCGGAAGCTG | Tb927.11.16620_rev | GGGGACCACTTTGTACAAGAAAGCTGGGTTTAAATTTCCACCGCAGCATAACCA |
| 8_hypothetical | K21 | Tb927.8.5680_for | GGGGACAAGTTTGTACAAAAAAGCAGGCTATGTTACGGCATACAGTGAAATTGTGC | Tb927.8.5680_rev | GGGGACCACTTTGTACAAGAAAGCTGGGTCTAAACGACACGGCCCTTATTTCT |
| 8_hypothetical | K22 | Tb927.10.13890_for | GGGGACAAGTTTGTACAAAAAAGCAGGCTATGCGTCCACGAATGGGTTC | Tb927.10.13890_rev | GGGGACCACTTTGTACAAGAAAGCTGGGTCTAAGTGTGTTGCCTTATGGGTGAT |
| 8_hypothetical | K23 | Tb927.11.6980_for | GGGGACAAGTTTGTACAAAAAAGCAGGCTATGCTTCGTAGAGTCCTCTCCG | Tb927.11.6980_rev | GGGGACCACTTTGTACAAGAAAGCTGGGTTCACACTGAAGCGTGCGC |
| 8_hypothetical | K24 | Tb927.8.3480_for | GGGGACAAGTTTGTACAAAAAAGCAGGCTATGACAGAAGGGATGGACCGC | Tb927.8.3480_rev | GGGGACCACTTTGTACAAGAAAGCTGGGTTTATTTTTGTACACTGCCCCCAGC |
| 8_hypothetical | L1 | Tb927.3.4430_for | GGGGACAAGTTTGTACAAAAAAGCAGGCTATGCAACGGTTAGTTAGTGCACTC | Tb927.3.4430_rev | GGGGACCACTTTGTACAAGAAAGCTGGGTCTAGCGCTGAACACGTTCCC |
| 8_hypothetical | L2 | Tb927.7.5690_for | GGGGACAAGTTTGTACAAAAAAGCAGGCTATGCCGACAACGGCGGAT | Tb927.7.5690_rev | GGGGACCACTTTGTACAAGAAAGCTGGGTTCAGGAGGAAAACTGTATAACGCCT |
| 8_hypothetical | L3 | Tb927.5.3590_for | GGGGACAAGTTTGTACAAAAAAGCAGGCTATGCCACCGAACCGTTGGG | Tb927.5.3590_rev | GGGGACCACTTTGTACAAGAAAGCTGGGTTTAGTTCACCTCCATAAGTTGCTTCC |
| 8_hypothetical | L4 | Tb927.7.730_for | GGGGACAAGTTTGTACAAAAAAGCAGGCTATGACGGAAAATTCACTCTCCACG | Tb927.7.730_rev | GGGGACCACTTTGTACAAGAAAGCTGGGTTCAGTTTCTCGGTGGGATAAATATATCACC |
| 8_hypothetical | L5 | Tb927.7.3100_for | GGGGACAAGTTTGTACAAAAAAGCAGGCTATGACGGGGCCAACGCGT | Tb927.7.3100_rev | GGGGACCACTTTGTACAAGAAAGCTGGGTTTAGCTGCTACGCCACTTATGTTC |
| 8_hypothetical | L6 | Tb927.3.3230_for | GGGGACAAGTTTGTACAAAAAAGCAGGCTATGAGCCGAAATCACAACAGAAGG | Tb927.3.3230_rev | GGGGACCACTTTGTACAAGAAAGCTGGGTTCAACTCTCTTCCGCGCAAC |
| 8_hypothetical | L7 | Tb927.1.3840_for | GGGGACAAGTTTGTACAAAAAAGCAGGCTATGGCTCACGCCACCAGT | Tb927.1.3840_rev | GGGGACCACTTTGTACAAGAAAGCTGGGTCTATGGTGACGCTATACCTGCTGA |
| 8_hypothetical | L8 | Tb927.11.6570_for | GGGGACAAGTTTGTACAAAAAAGCAGGCTATGAGTGTCACTGAAGGAGATGAACG | Tb927.11.6570_rev | GGGGACCACTTTGTACAAGAAAGCTGGGTTCACTCAGTCCCCCTGCG |
| 8_hypothetical | L9 | Tb927.11.7580_for | GGGGACAAGTTTGTACAAAAAAGCAGGCTATGAGAAGGCCTCTGTCGTTTATTG | Tb927.11.7580_rev | GGGGACCACTTTGTACAAGAAAGCTGGGTTTAAATTGAAAAGGCAACAGCTTCCTC |
| 8_hypothetical | L10 | Tb927.11.16120_for | GGGGACAAGTTTGTACAAAAAAGCAGGCTATGCGTGGTGTATTTTACCTTACCG | Tb927.11.16120_rev | GGGGACCACTTTGTACAAGAAAGCTGGGTTTAGAGTGTGGTGCCCTGGG |
| 8_hypothetical | L11 | Tb927.5.1900_for | GGGGACAAGTTTGTACAAAAAAGCAGGCTATGCACGAAGAGGACCCCG | Tb927.5.1900_rev | GGGGACCACTTTGTACAAGAAAGCTGGGTTCATATGGTTTCATCCTTCACTCGC |
| 8_hypothetical | L12 | Tb927.5.1950_for | GGGGACAAGTTTGTACAAAAAAGCAGGCTATGGAGGAAGTGCAACCGCT | Tb927.5.1950_rev | GGGGACCACTTTGTACAAGAAAGCTGGGTTTAGCCGATTTCACTCATAGCAATATCTAA |
| 8_hypothetical | L13 | Tb927.11.5130_for | GGGGACAAGTTTGTACAAAAAAGCAGGCTATGGTGGAACTTACAGTGTACGAAGG | Tb927.11.5130_rev | GGGGACCACTTTGTACAAGAAAGCTGGGTTCACGACAATGACCACATTTCTTTTTG |
| 8_hypothetical | L14 | Tb927.10.6210_for | GGGGACAAGTTTGTACAAAAAAGCAGGCTATGTCGGTGTCGCCTGAACC | Tb927.10.6210_rev | GGGGACCACTTTGTACAAGAAAGCTGGGTTTAGGTACATGTCGTGGAGTGGC |
| 8_hypothetical | L15 | Tb927.11.7200_for | GGGGACAAGTTTGTACAAAAAAGCAGGCTATGGCCGCTGAGGATCAAGG | Tb927.11.7200_rev | GGGGACCACTTTGTACAAGAAAGCTGGGTTTACTTCAATGGGGGACACTGTGG |
| 8_hypothetical | L16 | Tb927.10.2030_for | GGGGACAAGTTTGTACAAAAAAGCAGGCTATGCGGAGAGCCTTGTGGC | Tb927.10.2030_rev | GGGGACCACTTTGTACAAGAAAGCTGGGTTCAAATGACGATGTCATTTCGGGC |
| 8_hypothetical | L17 | Tb927.10.6320_for | GGGGACAAGTTTGTACAAAAAAGCAGGCTATGGCAGCCATGCAGCGC | Tb927.10.6320_rev | GGGGACCACTTTGTACAAGAAAGCTGGGTCTACTCACCAGCGGGATCTGA |
| 8_hypothetical | L18 | Tb927.10.1120_for | GGGGACAAGTTTGTACAAAAAAGCAGGCTATGCTTTTTGCCCAGCGCA | Tb927.10.1120_rev | GGGGACCACTTTGTACAAGAAAGCTGGGTCTATGCATTGTTACCGGCGAGT |
| 8_hypothetical | L19 | Tb927.9.1410_for | GGGGACAAGTTTGTACAAAAAAGCAGGCTATGTCCAGCAATAGCAGTGACG | Tb927.9.1410_rev | GGGGACCACTTTGTACAAGAAAGCTGGGTTCACTCCTCCCATAATCCACCAC |

| | | | | | |
|---|---|---|---|---|---|
| 8_hypothetical | L20 | Tb927.11.780_for | GGGGACAAGTTTGTACAAAAAAGCAGGCTATGGGGTTTAGCGACGGAGC | Tb927.11.780_rev | GGGGACCACTTTGTACAAGAAAGCTGGGTCTATGATTTTAATGTGTGACGATACCGC |
| 8_hypothetical | L21 | Tb927.11.1200_for | GGGGACAAGTTTGTACAAAAAAGCAGGCTATGGGAGACAGTGTGATACCCC | Tb927.11.1200_rev | GGGGACCACTTTGTACAAGAAAGCTGGGTTCACGCCTCTGCTTCCCA |
| 8_hypothetical | L22 | Tb927.4.1410_for | GGGGACAAGTTTGTACAAAAAAGCAGGCTATGTTTTCCTTCCTTACTTCGGGAGG | Tb927.4.1410_rev | GGGGACCACTTTGTACAAGAAAGCTGGGTTCATTCGCGCCCCCGCTG |
| 8_hypothetical | L23 | Tb927.9.2821_for | GGGGACAAGTTTGTACAAAAAAGCAGGCTATGGAAATCCGCATGGTGCC | Tb927.9.2821_rev | GGGGACCACTTTGTACAAGAAAGCTGGGTTCACTCATCAGCTGCTGGAGT |
| 8_hypothetical | L24 | Tb927.8.6720_for | GGGGACAAGTTTGTACAAAAAAGCAGGCTATGTCCTCCGTTACCACTGGC | Tb927.8.6720_rev | GGGGACCACTTTGTACAAGAAAGCTGGGTTTACTGTTCTTCTTTCTTCCCATACCCT |
| 8_hypothetical | M1 | Tb927.11.16610_for | GGGGACAAGTTTGTACAAAAAAGCAGGCTATGAGCTCTCAGCACCGCG | Tb927.11.16610_rev | GGGGACCACTTTGTACAAGAAAGCTGGGTTTACGCATCCACTACGAGCTTCG |
| 8_hypothetical | M2 | Tb927.11.5990_for | GGGGACAAGTTTGTACAAAAAAGCAGGCTATGAAGGTAAGTTCAAGGGGCCC | Tb927.11.5990_rev | GGGGACCACTTTGTACAAGAAAGCTGGGTCTACAATAAGCCATAGAGGTCGTGAAAA |
| 8_hypothetical | M3 | Tb927.8.1560_for | GGGGACAAGTTTGTACAAAAAAGCAGGCTATGGAGTTTGAGCAGCTGGAGA | Tb927.8.1560_rev | GGGGACCACTTTGTACAAGAAAGCTGGGTCTATATTTTTTTCCTAACGTCAGAAACATT |
| 8_hypothetical | M4 | Tb927.4.4300_for | GGGGACAAGTTTGTACAAAAAAGCAGGCTATGAGGAAGAGGGATGCGCT | Tb927.4.4300_rev | GGGGACCACTTTGTACAAGAAAGCTGGGTTCAGCGTTGAGTTCTCAGAACTGC |
| 8_hypothetical | M5 | Tb927.10.3430_for | GGGGACAAGTTTGTACAAAAAAGCAGGCTATGGTAATGGCTTGCGGTGC | Tb927.10.3430_rev | GGGGACCACTTTGTACAAGAAAGCTGGGTCTATGAGTACTGGTAGCCTTTGGCC |
| 8_hypothetical | M6 | Tb927.8.5290_for | GGGGACAAGTTTGTACAAAAAAGCAGGCTATGTTCCTTTATCTTATCATCGATGGAGAC | Tb927.8.5290_rev | GGGGACCACTTTGTACAAGAAAGCTGGGTTCAAGGACGTGTCCAAGTTGTCT |
| 8_hypothetical | M7 | Tb927.8.4270_for | GGGGACAAGTTTGTACAAAAAAGCAGGCTATGGAGGGGCGGTTATATGACTT | Tb927.8.4270_rev | GGGGACCACTTTGTACAAGAAAGCTGGGTTTACGTGGAGGATCCCCAAGC |
| 8_hypothetical | M8 | Tb927.5.1600_for | GGGGACAAGTTTGTACAAAAAAGCAGGCTATGAGTGAACCATTGATGGCTAGGT | Tb927.5.1600_rev | GGGGACCACTTTGTACAAGAAAGCTGGGTCTATTCTTCATTTTCCTCACCATCGGA |
| 8_hypothetical | M9 | Tb927.11.14170_for | GGGGACAAGTTTGTACAAAAAAGCAGGCTATGCTTCTCGTAGCGCTGTTTATC | Tb927.11.14170_rev | GGGGACCACTTTGTACAAGAAAGCTGGGTCTAAATCGATAGAAGCTCGGCAGC |
| 8_hypothetical | M10 | Tb927.10.13240_for | GGGGACAAGTTTGTACAAAAAAGCAGGCTATGAAGGCGCCATCCCTCC | Tb927.10.13240_rev | GGGGACCACTTTGTACAAGAAAGCTGGGTTCACATCTTTGACGTGTAGTTGTGAT |
| 8_hypothetical | M11 | Tb927.3.4800_for | GGGGACAAGTTTGTACAAAAAAGCAGGCTATGTCTACGAAGGAGGAAGTTGCG | Tb927.3.4800_rev | GGGGACCACTTTGTACAAGAAAGCTGGGTCTACAAAGGCTTTTCGTTGTTTGTGT |
| 8_hypothetical | M12 | Tb927.3.4040_for | GGGGACAAGTTTGTACAAAAAAGCAGGCTATGCAGACGCCATTGACTGATG | Tb927.3.4040_rev | GGGGACCACTTTGTACAAGAAAGCTGGGTCTAAATGAAGAAGCATGGAATAGCTACCT |
| 8_hypothetical | M13 | Tb927.9.9160_for | GGGGACAAGTTTGTACAAAAAAGCAGGCTATGGCAAAGGACTCGGAGAAGA | Tb927.9.9160_rev | GGGGACCACTTTGTACAAGAAAGCTGGGTTCACAACACACCCAAGAAAGACAA |
| 8_hypothetical | M14 | Tb927.11.8760_for | GGGGACAAGTTTGTACAAAAAAGCAGGCTATGGAAGAGGTCAGTAACGTGGT | Tb927.11.8760_rev | GGGGACCACTTTGTACAAGAAAGCTGGGTTCACAACATTATATCATCTACGGCAGC |
| 8_hypothetical | M15 | Tb927.11.9370_for | GGGGACAAGTTTGTACAAAAAAGCAGGCTATGGAGAATGTGGTTATGGAAGCG | Tb927.11.9370_rev | GGGGACCACTTTGTACAAGAAAGCTGGGTTCACCATAAGTCAACTGACTCTCGT |
| 8_hypothetical | M16 | Tb927.11.12260_for | GGGGACAAGTTTGTACAAAAAAGCAGGCTATGCAGGAAGCACAGCAGAC | Tb927.11.12260_rev | GGGGACCACTTTGTACAAGAAAGCTGGGTTTAAGCTGTTCTCTCCGTGCTGA |
| 8_hypothetical | M17 | Tb927.3.3740_for | GGGGACAAGTTTGTACAAAAAAGCAGGCTATGAGAAACATCTGGGTTTCTGTTCC | Tb927.3.3740_rev | GGGGACCACTTTGTACAAGAAAGCTGGGTTTAACGAGGCTCAAAGAAATTACTCTTCA |
| 8_hypothetical | M18 | Tb927.11.2620_for | GGGGACAAGTTTGTACAAAAAAGCAGGCTATGCTTCAAAACTGTTGGTTACTTGC | Tb927.11.2620_rev | GGGGACCACTTTGTACAAGAAAGCTGGGTCTATAAGTGAGCCATACTCGGGTTCA |
| 8_hypothetical | M19 | Tb927.11.15320_for | GGGGACAAGTTTGTACAAAAAAGCAGGCTATGATCGCATCTTCTCCGGCC | Tb927.11.15320_rev | GGGGACCACTTTGTACAAGAAAGCTGGGTCTAGCTGTTTCCCTTTGGATTGAGG |
| 8_hypothetical | M20 | Tb927.11.12050_for | GGGGACAAGTTTGTACAAAAAAGCAGGCTATGGATATAGGCAACTTTGACTCACG | Tb927.11.12050_rev | GGGGACCACTTTGTACAAGAAAGCTGGGTCTACGCTTTTTTCTTCGCGAGGT |
| 8_hypothetical | M21 | Tb927.11.15250_for | GGGGACAAGTTTGTACAAAAAAGCAGGCTATGCACCTTCCACCGTTACCC | Tb927.11.15250_rev | GGGGACCACTTTGTACAAGAAAGCTGGGTTCATGACAAAGACACCCTCCCTT |
| 8_hypothetical | M22 | Tb927.10.8570_for | GGGGACAAGTTTGTACAAAAAAGCAGGCTATGCCACTTTTTAAACCATCGGAAATAC | Tb927.10.8570_rev | GGGGACCACTTTGTACAAGAAAGCTGGGTTCAATCTCGTATAACCTCAAACGTTCC |
| 8_hypothetical | M23 | Tb927.9.6010_for | GGGGACAAGTTTGTACAAAAAAGCAGGCTATGCTTCTAAAGTTCACGGCAACT | Tb927.9.6010_rev | GGGGACCACTTTGTACAAGAAAGCTGGGTTTAGTAACTCTGTGTTTGGGGGACA |
| 8_hypothetical | M24 | Tb927.8.5490_for | GGGGACAAGTTTGTACAAAAAAGCAGGCTATGAAAATCAAAAAGAAGGACTTGCGC | Tb927.8.5490_rev | GGGGACCACTTTGTACAAGAAAGCTGGGTCTATTCATCACCGTATAAATCCTCCACC |
| 8_hypothetical | N1 | Tb927.11.11460_for | GGGGACAAGTTTGTACAAAAAAGCAGGCTATGTCTAGTGACGCCACTGCT | Tb927.11.11460_rev | GGGGACCACTTTGTACAAGAAAGCTGGGTTCAGAACTGGAGAGGTTGTGAGG |
| 8_hypothetical | N2 | Tb927.7.6920_for | GGGGACAAGTTTGTACAAAAAAGCAGGCTATGCGAATGCCTGATACGAGGC | Tb927.7.6920_rev | GGGGACCACTTTGTACAAGAAAGCTGGGTTCATTGTTTCTCCTTCACCTCAGTATTTC |
| 8_hypothetical | N3 | Tb927.6.4750_for | GGGGACAAGTTTGTACAAAAAAGCAGGCTATGTTCGCCTACATCTGTGCCA | Tb927.6.4750_rev | GGGGACCACTTTGTACAAGAAAGCTGGGTTTATTCGCTGGTTGCATTGAGTGA |
| 8_hypothetical | N4 | Tb927.11.15620_for | GGGGACAAGTTTGTACAAAAAAGCAGGCTATGAAGCACTGTTCTGCTTGCG | Tb927.11.15620_rev | GGGGACCACTTTGTACAAGAAAGCTGGGTTTATACTGGAGAGTTGCGCGGA |
| 8_hypothetical | N5 | Tb927.3.3560_for | GGGGACAAGTTTGTACAAAAAAGCAGGCTATGGCGACAGTTAAGAAACTCTTGG | Tb927.3.3560_rev | GGGGACCACTTTGTACAAGAAAGCTGGGTCTAAAAGTCATAAATATCCACTTCATTCTC |
| 8_hypothetical | N6 | Tb927.7.5850_for | GGGGACAAGTTTGTACAAAAAAGCAGGCTATGTTGCGTACATCCCGCG | Tb927.7.5850_rev | GGGGACCACTTTGTACAAGAAAGCTGGGTTTATTCCGAATCGTCGAGCAAAAGC |
| 8_hypothetical | N7 | Tb927.6.3010_for | GGGGACAAGTTTGTACAAAAAAGCAGGCTATGACGCACGGTAGTAATAATGATTCTAC | Tb927.6.3010_rev | GGGGACCACTTTGTACAAGAAAGCTGGGTCTATTCTGCTCTTTCGTCATAATTAGCAGG |
| 8_hypothetical | N8 | Tb927.9.11960_for | GGGGACAAGTTTGTACAAAAAAGCAGGCTATGAAGCATATGTTGCCGATAAGCA | Tb927.9.11960_rev | GGGGACCACTTTGTACAAGAAAGCTGGGTCTATGATATATTGCTTTCTTGGAGTTGGGT |
| 8_hypothetical | N9 | Tb927.11.15340_for | GGGGACAAGTTTGTACAAAAAAGCAGGCTATGCCACGGGATCCGTTTAATG | Tb927.11.15340_rev | GGGGACCACTTTGTACAAGAAAGCTGGGTTCACCTTCGCATCGACCAGA |
| 8_hypothetical | N10 | Tb927.8.4200_for | GGGGACAAGTTTGTACAAAAAAGCAGGCTATGCTAGATCTATTTGCATCGTTGGTT | Tb927.8.4200_rev | GGGGACCACTTTGTACAAGAAAGCTGGGTTCACCCCTTATGTTGCGTTACCA |
| 8_hypothetical | N11 | Tb927.6.3900_for | GGGGACAAGTTTGTACAAAAAAGCAGGCTATGCAGGGTGTAAGGCGAGA | Tb927.6.3900_rev | GGGGACCACTTTGTACAAGAAAGCTGGGTCTACGAAGCACTCCTCGCAC |
| 8_hypothetical | N12 | Tb927.1.1850_for | GGGGACAAGTTTGTACAAAAAAGCAGGCTATGAATCGTGCTCGAAAGCCAG | Tb927.1.1850_rev | GGGGACCACTTTGTACAAGAAAGCTGGGTTTAAAAGTAATATGGGTTCACACTAGAGCA |
| 8_hypothetical | N13 | Tb927.8.5410_for | GGGGACAAGTTTGTACAAAAAAGCAGGCTATGGCTGCCATTTTCCCCAAA | Tb927.8.5410_rev | GGGGACCACTTTGTACAAGAAAGCTGGGTTCAATTAGGTTTGCTGAAAACAGTACTCA |
| 8_hypothetical | N14 | Tb927.11.2700_for | GGGGACAAGTTTGTACAAAAAAGCAGGCTATGACATGCAACGTTGGGACG | Tb927.11.2700_rev | GGGGACCACTTTGTACAAGAAAGCTGGGTTTACTGTTGAATTATTGCAACACGGC |
| 8_hypothetical | N15 | Tb927.5.1720_for | GGGGACAAGTTTGTACAAAAAAGCAGGCTATGAGGGGGTCTTTGCCTCTT | Tb927.5.1720_rev | GGGGACCACTTTGTACAAGAAAGCTGGGTTTATTCACATGTTCCCCTCGATACGG |
| 8_hypothetical | N16 | Tb927.11.9750_for | GGGGACAAGTTTGTACAAAAAAGCAGGCTATGCGCTGCTCTTTGGTTCA | Tb927.11.9750_rev | GGGGACCACTTTGTACAAGAAAGCTGGGTCTAAACCATGATGGGGTTCGTCC |
| 8_hypothetical | N17 | Tb927.11.4210_for | GGGGACAAGTTTGTACAAAAAAGCAGGCTATGTCCGTACACAGTGCCATCT | Tb927.11.4210_rev | GGGGACCACTTTGTACAAGAAAGCTGGGTTCACGCCACGGCACTGGC |
| 8_hypothetical | N18 | Tb927.7.4380_for | GGGGACAAGTTTGTACAAAAAAGCAGGCTATGTCGGGGGGCACCTCC | Tb927.7.4380_rev | GGGGACCACTTTGTACAAGAAAGCTGGGTTCAGTTAAGTTCATTTTCAACCCGTTTAAG |
| 8_hypothetical | N19 | Tb927.7.780_for | GGGGACAAGTTTGTACAAAAAAGCAGGCTATGGAGGCAACACGGCTCC | Tb927.7.780_rev | GGGGACCACTTTGTACAAGAAAGCTGGGTTTACTCTACAACCCAAGGTGGCG |
| 8_hypothetical | N20 | Tb927.5.1790_for | GGGGACAAGTTTGTACAAAAAAGCAGGCTATGTTCCGTCGTGCGATTCC | Tb927.5.1790_rev | GGGGACCACTTTGTACAAGAAAGCTGGGTTCATGCCTCCTGTAGCCAGAG |
| 8_hypothetical | N21 | Tb927.1.4720_for | GGGGACAAGTTTGTACAAAAAAGCAGGCTATGGAGGAGTTGCGTAAACGC | Tb927.1.4720_rev | GGGGACCACTTTGTACAAGAAAGCTGGGTCTAAAGGTAATAAACCTGCGCCAATTTC |
| 8_hypothetical | N22 | Tb927.4.2890_for | GGGGACAAGTTTGTACAAAAAAGCAGGCTATGAATTCTGCCTCACAGGATATCGC | Tb927.4.2890_rev | GGGGACCACTTTGTACAAGAAAGCTGGGTTCAGGTAACGAAGATGCGCTTTATT |
| 8_hypothetical | N23 | Tb927.10.770_for | GGGGACAAGTTTGTACAAAAAAGCAGGCTATGCATCGCCAGCGCCGC | Tb927.10.770_rev | GGGGACCACTTTGTACAAGAAAGCTGGGTCTATGAGGAGGCAGCGGCC |
| 8_hypothetical | N24 | Tb927.11.8860_for | GGGGACAAGTTTGTACAAAAAAGCAGGCTATGTCTGGACCGGTTTCCCA | Tb927.11.8860_rev | GGGGACCACTTTGTACAAGAAAGCTGGGTTCATGGAGACCTGTAGGCTGC |
| 8_hypothetical | O1 | Tb927.5.1920_for | GGGGACAAGTTTGTACAAAAAAGCAGGCTATGCACGCGGGAGGAACA | Tb927.5.1920_rev | GGGGACCACTTTGTACAAGAAAGCTGGGTTCACGCCGCATCGTTTCG |
| 8_hypothetical | O2 | Tb927.11.2100_for | GGGGACAAGTTTGTACAAAAAAGCAGGCTATGTTTCAAATTGATGTCACCGACG | Tb927.11.2100_rev | GGGGACCACTTTGTACAAGAAAGCTGGGTTTATGGCAGTAGAGGAGTTATACGTCG |
| 8_hypothetical | O3 | Tb927.9.13010_for | GGGGACAAGTTTGTACAAAAAAGCAGGCTATGTGGTGGAGTGTTCGCCT | Tb927.9.13010_rev | GGGGACCACTTTGTACAAGAAAGCTGGGTTTACTTTCCTGCGCCTAAGGGT |
| 8_hypothetical | O4 | Tb927.10.11630_for | GGGGACAAGTTTGTACAAAAAAGCAGGCTATGAGTGCCAAGTTACAACCCATC | Tb927.10.11630_rev | GGGGACCACTTTGTACAAGAAAGCTGGGTTTAAACTACGATCAAATCTCCCGCG |
| 8_hypothetical | O5 | Tb927.11.16650_for | GGGGACAAGTTTGTACAAAAAAGCAGGCTATGAGCGCCCGGGAAAGT | Tb927.11.16650_rev | GGGGACCACTTTGTACAAGAAAGCTGGGTTCATTCAACACCCGCAATCGC |
| 8_hypothetical | O6 | Tb927.5.2070_for | GGGGACAAGTTTGTACAAAAAAGCAGGCTATGAAATCTGTGTTTGATATTGCCAGAAG | Tb927.5.2070_rev | GGGGACCACTTTGTACAAGAAAGCTGGGTCTACACTACCGGCACCCCA |
| 8_hypothetical | O7 | Tb927.8.4510_for | GGGGACAAGTTTGTACAAAAAAGCAGGCTATGTTCAAAAAAACGAGAAAGGCACA | Tb927.8.4510_rev | GGGGACCACTTTGTACAAGAAAGCTGGGTTCATTGCTTAAAGAGAAACATAATGACGGA |
| 8_hypothetical | O8 | Tb927.7.3040_for | GGGGACAAGTTTGTACAAAAAAGCAGGCTATGATTGACCCATTTCGCTCTTCTT | Tb927.7.3040_rev | GGGGACCACTTTGTACAAGAAAGCTGGGTTCAGTCTCTCACGGAATACACACT |
| 8_hypothetical | O9 | Tb927.8.7480_for | GGGGACAAGTTTGTACAAAAAAGCAGGCTATGAAGTCTCGTGGTTTTACTCGAC | Tb927.8.7480_rev | GGGGACCACTTTGTACAAGAAAGCTGGGTCTAATATTGCACCACATGGCGCA |
| 8_hypothetical | O10 | Tb927.8.4000_for | GGGGACAAGTTTGTACAAAAAAGCAGGCTATGATGGGAAGAATTCGTGTTTGCC | Tb927.8.4000_rev | GGGGACCACTTTGTACAAGAAAGCTGGGTTCACGCTTTCTCCTCTTGCGA |
| 8_hypothetical | O11 | Tb927.10.3850_for | GGGGACAAGTTTGTACAAAAAAGCAGGCTATGACAGAAGTGAAGGGGCACC | Tb927.10.3850_rev | GGGGACCACTTTGTACAAGAAAGCTGGGTTCAGTCACTCATAGGAAATTGAGCGC |
| 8_hypothetical | O12 | Tb927.10.11480_for | GGGGACAAGTTTGTACAAAAAAGCAGGCTATGAGATGCGCACGTTTCTATCTT | Tb927.10.11480_rev | GGGGACCACTTTGTACAAGAAAGCTGGGTCTACAGCCTTGGCTTTTCCAACG |
| 8_hypothetical | O13 | Tb927.10.11810_for | GGGGACAAGTTTGTACAAAAAAGCAGGCTATGTCACAGGCAGGTGTGAGT | Tb927.10.11810_rev | GGGGACCACTTTGTACAAGAAAGCTGGGTTCATAACTTCGATACCACATACGCCG |
| 8_hypothetical | O14 | Tb927.10.9160_for | GGGGACAAGTTTGTACAAAAAAGCAGGCTATGACTGCGACTTCCGGCT | Tb927.10.9160_rev | GGGGACCACTTTGTACAAGAAAGCTGGGTCTACCTTCGACGCCGCTG |
| 8_hypothetical | O15 | Tb927.7.6320_for | GGGGACAAGTTTGTACAAAAAAGCAGGCTATGATGCGACGGACGGGT | Tb927.7.6320_rev | GGGGACCACTTTGTACAAGAAAGCTGGGTTTAAGTTGTTGGGAAATCATCACCACG |
| 8_hypothetical | O16 | Tb927.8.770_for | GGGGACAAGTTTGTACAAAAAAGCAGGCTATGAATGAGGGGAAGGGTACGTC | Tb927.8.770_rev | GGGGACCACTTTGTACAAGAAAGCTGGGTTCAGGTTTTCCACGTTATTGTGGAA |
| 8_hypothetical | O17 | Tb927.7.5070_for | GGGGACAAGTTTGTACAAAAAAGCAGGCTATGGCACAAGGACGAGTTATTGAG | Tb927.7.5070_rev | GGGGACCACTTTGTACAAGAAAGCTGGGTTTAGCACGTGAGTTCCTTGCG |
| 8_hypothetical | O18 | Tb927.11.13550_for | GGGGACAAGTTTGTACAAAAAAGCAGGCTATGACCCACCAAACCCATGCG | Tb927.11.13550_rev | GGGGACCACTTTGTACAAGAAAGCTGGGTTCACCAAAAACAAACGTACGCAGC |
| 8_hypothetical | O19 | Tb927.7.6280_for | GGGGACAAGTTTGTACAAAAAAGCAGGCTATGGACGTCACGCCTGTGG | Tb927.7.6280_rev | GGGGACCACTTTGTACAAGAAAGCTGGGTTTAATACAGGAAGGTCTTCTGTACCGT |
| 8_hypothetical | O20 | Tb927.11.4150_for | GGGGACAAGTTTGTACAAAAAAGCAGGCTATGCGCGTGCTAGCTGTG | Tb927.11.4150_rev | GGGGACCACTTTGTACAAGAAAGCTGGGTTCAAATCTCACCCGTTTCACCATC |
| 8_hypothetical | O21 | Tb927.10.450_for | GGGGACAAGTTTGTACAAAAAAGCAGGCTATGCCCGAGGCGTATATTGAGG | Tb927.10.450_rev | GGGGACCACTTTGTACAAGAAAGCTGGGTCTAACGGGTAAGGGCATCGGG |
| 8_hypothetical | O22 | Tb927.9.8180_for | GGGGACAAGTTTGTACAAAAAAGCAGGCTATGGGGAGTGATCTTTCCATTGTGT | Tb927.9.8180_rev | GGGGACCACTTTGTACAAGAAAGCTGGGTTTATTTTACACGCTTTGGTGGCTCC |
| 8_hypothetical | O23 | Tb927.11.2180_for | GGGGACAAGTTTGTACAAAAAAGCAGGCTATGCCGTCTGCTAGTTCCGG | Tb927.11.2180_rev | GGGGACCACTTTGTACAAGAAAGCTGGGTTCAAGTTTGATTCCTCAGCGCG |
| 8_hypothetical | O24 | Tb927.8.3930_for | GGGGACAAGTTTGTACAAAAAAGCAGGCTATGAACTTCTTATCCCAGATTTACACCG | Tb927.8.3930_rev | GGGGACCACTTTGTACAAGAAAGCTGGGTCTAATAGACGGCGGTGCCG |
| 8_hypothetical | P1 | Tb927.11.2890_for | GGGGACAAGTTTGTACAAAAAAGCAGGCTATGCTTTTTCGTGAAACGGCAAG | Tb927.11.2890_rev | GGGGACCACTTTGTACAAGAAAGCTGGGTTCACTGCAGTGGCGGAGG |
| 8_hypothetical | P2 | Tb927.11.6040_for | GGGGACAAGTTTGTACAAAAAAGCAGGCTATGCATTTGGAGCAAGCTGGAG | Tb927.11.6040_rev | GGGGACCACTTTGTACAAGAAAGCTGGGTTTAATAAATGAACCCACCCCTCTTTTTGA |
| 8_hypothetical | P3 | Tb927.10.9140_for | GGGGACAAGTTTGTACAAAAAAGCAGGCTATGGGTTCTTTTACTTCCTGTGACCA | Tb927.10.9140_rev | GGGGACCACTTTGTACAAGAAAGCTGGGTCTACGATGACGATGACGGAGTGG |
| 8_hypothetical | P4 | Tb927.11.14860_for | GGGGACAAGTTTGTACAAAAAAGCAGGCTATGTTTGCGGACCGCGAA | Tb927.11.14860_rev | GGGGACCACTTTGTACAAGAAAGCTGGGTTTAGTTTGCGCACTGTTGGCG |
| 8_hypothetical | P5 | Tb927.9.10820_for | GGGGACAAGTTTGTACAAAAAAGCAGGCTATGAGCGATAAGGTAAATGTCCATAAAAAT | Tb927.9.10820_rev | GGGGACCACTTTGTACAAGAAAGCTGGGTTTATATCCCGCTCATCGCCAAAC |
| 8_hypothetical | P6 | Tb927.10.930_for | GGGGACAAGTTTGTACAAAAAAGCAGGCTATGCAGAACGTTATCCTTGCCC | Tb927.10.930_rev | GGGGACCACTTTGTACAAGAAAGCTGGGTTTATTCCTGCGGCTGCTGC |
| 8_hypothetical | P7 | Tb927.3.5070_for | GGGGACAAGTTTGTACAAAAAAGCAGGCTATGTCCTCTCTCCCTTACCGC | Tb927.3.5070_rev | GGGGACCACTTTGTACAAGAAAGCTGGGTTCACTCAACATATTTCCCCCCCC |
| 8_hypothetical | P8 | Tb927.8.2600_for | GGGGACAAGTTTGTACAAAAAAGCAGGCTATGGGCATGAAGCTGCAGC | Tb927.8.2600_rev | GGGGACCACTTTGTACAAGAAAGCTGGGTCTACCCCATCAAACGCCTGAGG |
| 8_hypothetical | P9 | Tb927.11.4400_for | GGGGACAAGTTTGTACAAAAAAGCAGGCTATGCTTGGTTCATGGCTGAACG | Tb927.11.4400_rev | GGGGACCACTTTGTACAAGAAAGCTGGGTTCAACTCTGGGATTTCGTCTGTTG |
| 8_hypothetical | P10 | Tb927.11.2420_for | GGGGACAAGTTTGTACAAAAAAGCAGGCTATGATGAAAGACTTTGAAGCATTCTCGA | Tb927.11.2420_rev | GGGGACCACTTTGTACAAGAAAGCTGGGTTCACAGTGGTGTCATGTTCGTTG |
| 8_hypothetical | P11 | Tb927.11.10330_for | GGGGACAAGTTTGTACAAAAAAGCAGGCTATGTTCTTCTGGGGTTCGTACAAC | Tb927.11.10330_rev | GGGGACCACTTTGTACAAGAAAGCTGGGTTCATTTTCCGACGACATAGTTTCCAG |
| 8_hypothetical | P12 | Tb927.2.2410_for | GGGGACAAGTTTGTACAAAAAAGCAGGCTATGGTAGTGCACCGTCGCA | Tb927.2.2410_rev | GGGGACCACTTTGTACAAGAAAGCTGGGTTCACCTTCCCGCGAGAAGT |
| 8_hypothetical | P13 | Tb927.10.1930_for | GGGGACAAGTTTGTACAAAAAAGCAGGCTATGCGCCCTGCAGCCACT | Tb927.10.1930_rev | GGGGACCACTTTGTACAAGAAAGCTGGGTCTATCCCATCTCCTTACGCCCG |
| 8_hypothetical | P14 | Tb927.8.4310_for | GGGGACAAGTTTGTACAAAAAAGCAGGCTATGGAAAGGCGTATTGAAGAGGATG | Tb927.8.4310_rev | GGGGACCACTTTGTACAAGAAAGCTGGGTTTAATTGCGATAGCAACCGCGC |
| 8_hypothetical | P15 | Tb927.11.280_for | GGGGACAAGTTTGTACAAAAAAGCAGGCTATGAGTTTGACACTGGGAGAATTAAGTG | Tb927.11.280_rev | GGGGACCACTTTGTACAAGAAAGCTGGGTTCACTGAGTAGCAGAAATGTAGTTTCC |
| 8_hypothetical | P16 | Tb927.11.14890_for | GGGGACAAGTTTGTACAAAAAAGCAGGCTATGCAACAATCAACGCGATGGA | Tb927.11.14890_rev | GGGGACCACTTTGTACAAGAAAGCTGGGTTCACGCAGCGGCGGCGAT |
| 8_hypothetical | P17 | Tb927.5.1980_for | GGGGACAAGTTTGTACAAAAAAGCAGGCTATGATACCGAGGCATTATGTAGCAATG | Tb927.5.1980_rev | GGGGACCACTTTGTACAAGAAAGCTGGGTTTAACCCCTCAGTTCCTCGGG |
| 8_hypothetical | P18 | Tb927.7.3660_for | GGGGACAAGTTTGTACAAAAAAGCAGGCTATGTCGACACTGGAGGGCTG | Tb927.7.3660_rev | GGGGACCACTTTGTACAAGAAAGCTGGGTTCAATACGGGAAGATATCTGAAAAGACG |
| 8_hypothetical | P19 | Tb927.10.12430_for | GGGGACAAGTTTGTACAAAAAAGCAGGCTATGGCAAAGGAAAAAAAATCGTTCAAGA | Tb927.10.12430_rev | GGGGACCACTTTGTACAAGAAAGCTGGGTCTAGGAGTTGTGGTCGCTCCT |
| 8_hypothetical | P20 | Tb927.11.10960_for | GGGGACAAGTTTGTACAAAAAAGCAGGCTATGCGCAGGTGCCTTATTTTGC | Tb927.11.10960_rev | GGGGACCACTTTGTACAAGAAAGCTGGGTTTACGCAGTTGGCTGGCCG |
| 8_hypothetical | P21 | Tb927.11.5120_for | GGGGACAAGTTTGTACAAAAAAGCAGGCTATGCAGTTAGCTGAGGAACTAAAACAA | Tb927.11.5120_rev | GGGGACCACTTTGTACAAGAAAGCTGGGTTTACACCTTCTCGACTGCCTTTTTAAT |
| 8_hypothetical | P22 | Tb927.11.5410_for | GGGGACAAGTTTGTACAAAAAAGCAGGCTATGATCGAAAATGTTACCGCAGTGT | Tb927.11.5410_rev | GGGGACCACTTTGTACAAGAAAGCTGGGTCTATTCAGCCGGTATTGTAGTAGCAC |
| 8_hypothetical | P23 | Tb927.10.13690_for | GGGGACAAGTTTGTACAAAAAAGCAGGCTATGACATTTATTTCCTTCCATAGTTTGGCG | Tb927.10.13690_rev | GGGGACCACTTTGTACAAGAAAGCTGGGTCTACGCAACTAGCTCCGTATCCA |
| 8_hypothetical | P24 | Tb927.6.3520_for | GGGGACAAGTTTGTACAAAAAAGCAGGCTATGAACCAAAAAGAGGATCACGAGG | Tb927.6.3520_rev | GGGGACCACTTTGTACAAGAAAGCTGGGTTCAGTCCTCTGCACGGGG |
| 9_hypothetical | A1 | Tb927.10.13390_for | GGGGACAAGTTTGTACAAAAAAGCAGGCTATGACATTTATTTCCTTCCATAGTTTGGCG | Tb927.10.13390_rev | GGGGACCACTTTGTACAAGAAAGCTGGGTTTACGCAACTAGCTCCGTATCCA |
| 9_hypothetical | A2 | Tb927.10.1350_for | GGGGACAAGTTTGTACAAAAAAGCAGGCTATGGGAGCCGCGTGCACC | Tb927.10.1350_rev | GGGGACCACTTTGTACAAGAAAGCTGGGTTCACGAAACCCTAGTGACAAATCG |
| 9_hypothetical | A3 | Tb927.10.7880_for | GGGGACAAGTTTGTACAAAAAAGCAGGCTATGGAGACGGGAGGCGAC | Tb927.10.7880_rev | GGGGACCACTTTGTACAAGAAAGCTGGGTTCAGGTATGGGCACGGGTTT |
| 9_hypothetical | A4 | Tb927.7.7030_for | GGGGACAAGTTTGTACAAAAAAGCAGGCTATGCCCCACGAGCCGTCG | Tb927.7.7030_rev | GGGGACCACTTTGTACAAGAAAGCTGGGTCTACCGCCGCGTTGGCCG |
| 9_hypothetical | A5 | Tb927.8.3590_for | GGGGACAAGTTTGTACAAAAAAGCAGGCTATGAGTTCTGTGGGTGTGAGCG | Tb927.8.3590_rev | GGGGACCACTTTGTACAAGAAAGCTGGGTCTATGACGGGACAGGAGAGGC |
| 9_hypothetical | A6 | Tb927.11.10470_for | GGGGACAAGTTTGTACAAAAAAGCAGGCTATGAAGACTCATCGTGCCGC | Tb927.11.10470_rev | GGGGACCACTTTGTACAAGAAAGCTGGGTCTACTCAGTTAATGAACCTTCTGTGTAGTA |
| 9_hypothetical | A7 | Tb927.9.11830_for | GGGGACAAGTTTGTACAAAAAAGCAGGCTATGCTTACTTCATCCTTCATCAGCG | Tb927.9.11830_rev | GGGGACCACTTTGTACAAGAAAGCTGGGTCTAGTCTAGCGCTAAGCGAGCG |
| 9_hypothetical | A8 | Tb927.11.7180_for | GGGGACAAGTTTGTACAAAAAAGCAGGCTATGACAACGAAGGAAGCAGTGG | Tb927.11.7180_rev | GGGGACCACTTTGTACAAGAAAGCTGGGTTCACACCCCGTCATGAGAGG |
| 9_hypothetical | A9 | Tb927.8.2680_for | GGGGACAAGTTTGTACAAAAAAGCAGGCTATGCGGTGGATCCCTGCA | Tb927.8.2680_rev | GGGGACCACTTTGTACAAGAAAGCTGGGTTTACATGTCTCTCGGCCGCT |
| 9_hypothetical | A10 | Tb927.7.3670_for | GGGGACAAGTTTGTACAAAAAAGCAGGCTATGCAGGAAGACGTAAAACACCTT | Tb927.7.3670_rev | GGGGACCACTTTGTACAAGAAAGCTGGGTCTACGGTGAATTTGCCGAGCA |
| 9_hypothetical | A11 | Tb927.4.2430_for | GGGGACAAGTTTGTACAAAAAAGCAGGCTATGGCACGGCACCCCAAC | Tb927.4.2430_rev | GGGGACCACTTTGTACAAGAAAGCTGGGTCTATTTGTACGACAAATCGGCAATCAA |
| 9_hypothetical | A12 | Tb927.3.3170_for | GGGGACAAGTTTGTACAAAAAAGCAGGCTATGCATACAGCGGCATCGAAG | Tb927.3.3170_rev | GGGGACCACTTTGTACAAGAAAGCTGGGTCTAGACGGCGGCACGAGG |
| 9_hypothetical | A13 | Tb927.11.990_for | GGGGACAAGTTTGTACAAAAAAGCAGGCTATGTTCTTCACACATGGGCCG | Tb927.11.990_rev | GGGGACCACTTTGTACAAGAAAGCTGGGTTTATGTCCCGCATCCCTCTCTC |
| 9_hypothetical | A14 | Tb927.3.2870_for | GGGGACAAGTTTGTACAAAAAAGCAGGCTATGAATGGACGAATCACCGGC | Tb927.3.2870_rev | GGGGACCACTTTGTACAAGAAAGCTGGGTCTAAGGAATCTGCGTCTCAGCTTT |
| 9_hypothetical | A15 | Tb927.10.6900_for | GGGGACAAGTTTGTACAAAAAAGCAGGCTATGGGAGGTCCAAGTGAGGGG | Tb927.10.6900_rev | GGGGACCACTTTGTACAAGAAAGCTGGGTTTAGTCTGCCTGCGACCCTC |
| 9_hypothetical | A16 | Tb927.11.16990_for | GGGGACAAGTTTGTACAAAAAAGCAGGCTATGTGCATATGTACACGTCTCTTGTG | Tb927.11.16990_rev | GGGGACCACTTTGTACAAGAAAGCTGGGTTTACGCAGCCCAGAAACCGG |
| 9_hypothetical | A17 | Tb927.9.14170_for | GGGGACAAGTTTGTACAAAAAAGCAGGCTATGGATGGGCGGATTAAAGGCC | Tb927.9.14170_rev | GGGGACCACTTTGTACAAGAAAGCTGGGTTTACACAGCACTCACTGAGGGA |
| 9_hypothetical | A18 | Tb927.10.2460_for | GGGGACAAGTTTGTACAAAAAAGCAGGCTATGTTCCTCAGAAGTGATCAGCATCG | Tb927.10.2460_rev | GGGGACCACTTTGTACAAGAAAGCTGGGTCTAGAGGAAATGATATAACAGAACGATCGT |
| 9_hypothetical | A19 | Tb927.9.4450_for | GGGGACAAGTTTGTACAAAAAAGCAGGCTATGGAAAACGCAAGGAATCTGGAG | Tb927.9.4450_rev | GGGGACCACTTTGTACAAGAAAGCTGGGTTTAATTTTGATTACTAGTATCATTAGGACG |
| 9_hypothetical | A20 | Tb927.11.5310_for | GGGGACAAGTTTGTACAAAAAAGCAGGCTATGCCTCTCTGCTGTTGTAAAGATG | Tb927.11.5310_rev | GGGGACCACTTTGTACAAGAAAGCTGGGTTTAGAATGGCGAAGAGTAAGTTCTCATATG |
| 9_hypothetical | A21 | Tb927.10.630_for | GGGGACAAGTTTGTACAAAAAAGCAGGCTATGTCGTACAAATATAGCAGTAGAGAATAT | Tb927.10.630_rev | GGGGACCACTTTGTACAAGAAAGCTGGGTTCATTTTTTCTCTTCACGACTAAATAACAT |
| 9_hypothetical | A22 | Tb927.9.13140_for | GGGGACAAGTTTGTACAAAAAAGCAGGCTATGCACAAACCGGTGACGG | Tb927.9.13140_rev | GGGGACCACTTTGTACAAGAAAGCTGGGTCTATGCTCCCTCATTCTCATCCGG |
| 9_hypothetical | A23 | Tb927.11.2630_for | GGGGACAAGTTTGTACAAAAAAGCAGGCTATGTACCACGATGCTCCGCC | Tb927.11.2630_rev | GGGGACCACTTTGTACAAGAAAGCTGGGTTCATTGTTCCTTTTCTGTTGCATATCCA |
| 9_hypothetical | A24 | Tb927.9.8200_for | GGGGACAAGTTTGTACAAAAAAGCAGGCTATGCACCGAAAGCAGGCATG | Tb927.9.8200_rev | GGGGACCACTTTGTACAAGAAAGCTGGGTTCAACGCACCCACTTTGGAAG |
| 9_hypothetical | B1 | Tb927.11.14930_for | GGGGACAAGTTTGTACAAAAAAGCAGGCTATGGATAGGGAAAAACTAGCGTGGG | Tb927.11.14930_rev | GGGGACCACTTTGTACAAGAAAGCTGGGTTTACACATACACGTTAAACGGTGCT |
| 9_hypothetical | B2 | Tb927.11.2970_for | GGGGACAAGTTTGTACAAAAAAGCAGGCTATGAGACCGCGCCCTGTT | Tb927.11.2970_rev | GGGGACCACTTTGTACAAGAAAGCTGGGTTCATAAAAGACCACTGAGGTTAAGGAGT |
| 9_hypothetical | B3 | Tb927.11.6770_for | GGGGACAAGTTTGTACAAAAAAGCAGGCTATGCTGTGCCACTTCTTTGCA | Tb927.11.6770_rev | GGGGACCACTTTGTACAAGAAAGCTGGGTCTAGTCGTCTTCGTCTCCATCCT |
| 9_hypothetical | B4 | Tb927.9.14300_for | GGGGACAAGTTTGTACAAAAAAGCAGGCTATGGATGTGTGTGAGGGATTGAGG | Tb927.9.14300_rev | GGGGACCACTTTGTACAAGAAAGCTGGGTTCATAGAAGCTCACGGTGTCTGG |
| 9_hypothetical | B5 | Tb927.10.15190_for | GGGGACAAGTTTGTACAAAAAAGCAGGCTATGGAAGGTGCTGCAAGGACG | Tb927.10.15190_rev | GGGGACCACTTTGTACAAGAAAGCTGGGTTTACTGTAATTCGTGATGGGTTAAACCC |

| | | | | | |
|---|---|---|---|---|---|
| 9_hypothetical | B6 | Tb927.9.11690_for | GGGGACAAGTTTGTACAAAAAAGCAGGCTATGAGTGGTTCCCATTGGATGCT | Tb927.9.11690_rev | GGGGACCACTTTGTACAAGAAAGCTGGGTTTATGGAGAACGAACTGATGAGGTGA |
| 9_hypothetical | B7 | Tb927.11.13400_for | GGGGACAAGTTTGTACAAAAAAGCAGGCTATGAGTCAAAACCGGCAGCTG | Tb927.11.13400_rev | GGGGACCACTTTGTACAAGAAAGCTGGGTTCACAAGGCGGGAGCCTC |
| 9_hypothetical | B8 | Tb927.5.530_for | GGGGACAAGTTTGTACAAAAAAGCAGGCTATGAGGACGTTCTTTTCGCACG | Tb927.5.530_rev | GGGGACCACTTTGTACAAGAAAGCTGGGTTCAGACGCTAGTTTTGTTGTTGTTGC |
| 9_hypothetical | B9 | Tb927.10.10150_for | GGGGACAAGTTTGTACAAAAAAGCAGGCTATGACAGACGCCGAATCTGACG | Tb927.10.10150_rev | GGGGACCACTTTGTACAAGAAAGCTGGGTTCACCAGGGGCCCCGCGT |
| 9_hypothetical | B10 | Tb927.11.2780_for | GGGGACAAGTTTGTACAAAAAAGCAGGCTATGCTAGCAGGAAAGATTAGCGGA | Tb927.11.2780_rev | GGGGACCACTTTGTACAAGAAAGCTGGGTCTATTTTACTCGATCACTCCTCTTCTTCG |
| 9_hypothetical | B11 | Tb927.7.5310_for | GGGGACAAGTTTGTACAAAAAAGCAGGCTATGATTCACGGCAATGCTGCG | Tb927.7.5310_rev | GGGGACCACTTTGTACAAGAAAGCTGGGTTTATGTGTACTGCTGGTAGATATGAGCC |
| 9_hypothetical | B12 | Tb927.4.4140_for | GGGGACAAGTTTGTACAAAAAAGCAGGCTATGACTGCAGTTAAGGAGCCTCG | Tb927.4.4140_rev | GGGGACCACTTTGTACAAGAAAGCTGGGTTCATAAGTTCCACGTCAGCCCG |
| 9_hypothetical | B13 | Tb927.11.8300_for | GGGGACAAGTTTGTACAAAAAAGCAGGCTATGTCGTCAAAGAAGAAGGGCTCG | Tb927.11.8300_rev | GGGGACCACTTTGTACAAGAAAGCTGGGTCTATTTCAGCAGCGTTAACAACACC |
| 9_hypothetical | B15 | Tb927.7.2720_for | GGGGACAAGTTTGTACAAAAAAGCAGGCTATGTTTCACAAGAAATATGTCGTGAAAGG | Tb927.7.2720_rev | GGGGACCACTTTGTACAAGAAAGCTGGGTTCATGCACAGCGTTCAGCTATT |
| 9_hypothetical | B16 | Tb927.10.4320_for | GGGGACAAGTTTGTACAAAAAAGCAGGCTATGAAACCGCGAGACCTATGGG | Tb927.10.4320_rev | GGGGACCACTTTGTACAAGAAAGCTGGGTCTACGCCTGCATAGAGTAAAGTCTCT |
| 9_hypothetical | B17 | Tb927.5.1640_for | GGGGACAAGTTTGTACAAAAAAGCAGGCTATGCGCCCCAACTCGATGC | Tb927.5.1640_rev | GGGGACCACTTTGTACAAGAAAGCTGGGTTCAGTAGGTTGTGGACCGCC |
| 9_hypothetical | B18 | Tb927.3.710_for | GGGGACAAGTTTGTACAAAAAAGCAGGCTATGCTCTTATATTTTATTGTTGTTGCCGTA | Tb927.3.710_rev | GGGGACCACTTTGTACAAGAAAGCTGGGTCTACACGATGTCGGCGTTACG |
| 9_hypothetical | B19 | Tb927.7.700_for | GGGGACAAGTTTGTACAAAAAAGCAGGCTATGCAGGTATCGCTCCATAACAAG | Tb927.7.700_rev | GGGGACCACTTTGTACAAGAAAGCTGGGTTCATTTAGACGACACTCTCCGAAGC |
| 9_hypothetical | B20 | Tb927.7.5630_for | GGGGACAAGTTTGTACAAAAAAGCAGGCTATGCGGGCATCTGCAGCT | Tb927.7.5630_rev | GGGGACCACTTTGTACAAGAAAGCTGGGTTTACTCAGCCTTTGCACTAGGGA |
| 9_hypothetical | B21 | Tb927.9.6510_for | GGGGACAAGTTTGTACAAAAAAGCAGGCTATGTCAGTGAGCGGTGTGTTC | Tb927.9.6510_rev | GGGGACCACTTTGTACAAGAAAGCTGGGTCTAAAACTCCGGAGTCGCGC |
| 9_hypothetical | B22 | Tb927.7.5890_for | GGGGACAAGTTTGTACAAAAAAGCAGGCTATGAATGAGGATAAAAAAGCACCCTTGA | Tb927.7.5890_rev | GGGGACCACTTTGTACAAGAAAGCTGGGTTCACACTCTCCTTTCCATGTCCTT |
| 9_hypothetical | B23 | Tb927.11.3020_for | GGGGACAAGTTTGTACAAAAAAGCAGGCTATGCATGGGAGGAATAATGCGG | Tb927.11.3020_rev | GGGGACCACTTTGTACAAGAAAGCTGGGTCTAGAAGAGGGAACATATTCGCTTTTTCA |
| 9_hypothetical | B24 | Tb927.3.2800_for | GGGGACAAGTTTGTACAAAAAAGCAGGCTATGATTGGCAGGTTATTGGGGTTTAG | Tb927.3.2800_rev | GGGGACCACTTTGTACAAGAAAGCTGGGTCTATGAAAAGCAAACGACGTTCAGC |
| 9_hypothetical | C1 | Tb927.8.4400_for | GGGGACAAGTTTGTACAAAAAAGCAGGCTATGACTAGGTTGCCTTCAGCGG | Tb927.8.4400_rev | GGGGACCACTTTGTACAAGAAAGCTGGGTTCACCGCTGGCGTTCCTC |
| 9_hypothetical | C2 | Tb927.11.9280_for | GGGGACAAGTTTGTACAAAAAAGCAGGCTATGCTCTCCAGTGCGCTGA | Tb927.11.9280_rev | GGGGACCACTTTGTACAAGAAAGCTGGGTTCAACCTACGCGCATAACCTTATC |
| 9_hypothetical | C3 | Tb927.11.16680_for | GGGGACAAGTTTGTACAAAAAAGCAGGCTATGCTTCAGGCCTGTGGTGT | Tb927.11.16680_rev | GGGGACCACTTTGTACAAGAAAGCTGGGTTCACCACCAAGTGTCACAGCC |
| 9_hypothetical | C4 | Tb927.10.14560_for | GGGGACAAGTTTGTACAAAAAAGCAGGCTATGAGCGATTCAAAGAAAAAGGCCG | Tb927.10.14560_rev | GGGGACCACTTTGTACAAGAAAGCTGGGTTTATCTCCGTTTAACCTTCTTTCCGC |
| 9_hypothetical | C5 | Tb927.4.4940_for | GGGGACAAGTTTGTACAAAAAAGCAGGCTATGAAGTCCCGTGGTTTCACCC | Tb927.4.4940_rev | GGGGACCACTTTGTACAAGAAAGCTGGGTCTAGCATAGCGCAACGTGGC |
| 9_hypothetical | C6 | Tb927.8.1700_for | GGGGACAAGTTTGTACAAAAAAGCAGGCTATGCCGTTGAAGTTTGTTCACCA | Tb927.8.1700_rev | GGGGACCACTTTGTACAAGAAAGCTGGGTCTAGACGGCGTCAAACCCAAG |
| 9_hypothetical | C7 | Tb927.6.2400_for | GGGGACAAGTTTGTACAAAAAAGCAGGCTATGGAAGAACTGCAGTGCGC | Tb927.6.2400_rev | GGGGACCACTTTGTACAAGAAAGCTGGGTCTACCGGTGCCCTGCTTTC |
| 9_hypothetical | C8 | Tb927.7.2220_for | GGGGACAAGTTTGTACAAAAAAGCAGGCTATGCGCGGCTTCCGCTCA | Tb927.7.2220_rev | GGGGACCACTTTGTACAAGAAAGCTGGGTCTACTGTTCCCTCGACCCACC |
| 9_hypothetical | C9 | Tb927.10.2690_for | GGGGACAAGTTTGTACAAAAAAGCAGGCTATGATGAACTTAGGAGAAACTCTTGACAC | Tb927.10.2690_rev | GGGGACCACTTTGTACAAGAAAGCTGGGTTCACCACACCCTCGACTTGC |
| 9_hypothetical | C10 | Tb927.9.6410_for | GGGGACAAGTTTGTACAAAAAAGCAGGCTATGATTCTCCGGGGACGGTT | Tb927.9.6410_rev | GGGGACCACTTTGTACAAGAAAGCTGGGTTCAAGAACCATCAATTTCCGCGC |
| 9_hypothetical | C11 | Tb927.6.3720_for | GGGGACAAGTTTGTACAAAAAAGCAGGCTATGTCCCACACTGGGTGTCC | Tb927.6.3720_rev | GGGGACCACTTTGTACAAGAAAGCTGGGTTCACAAATGTGGAGCAACATCCC |
| 9_hypothetical | C12 | Tb927.7.2950_for | GGGGACAAGTTTGTACAAAAAAGCAGGCTATGTTTACATCTTTTTTAATAGTTTTAAAT | Tb927.7.2950_rev | GGGGACCACTTTGTACAAGAAAGCTGGGTTCACTGCACGTTGGCAGAAG |
| 9_hypothetical | C13 | Tb927.4.2760_for | GGGGACAAGTTTGTACAAAAAAGCAGGCTATGGAAGAGGGTGATCATATGGAGTT | Tb927.4.2760_rev | GGGGACCACTTTGTACAAGAAAGCTGGGTCTAAATGATGTGACTGACCACTGGC |
| 9_hypothetical | C14 | Tb927.10.3730_for | GGGGACAAGTTTGTACAAAAAAGCAGGCTATGCGTCCGTATTTCCCAGGT | Tb927.10.3730_rev | GGGGACCACTTTGTACAAGAAAGCTGGGTTCACCGTGCTGGTCGCGC |
| 9_hypothetical | C15 | Tb927.9.10190_for | GGGGACAAGTTTGTACAAAAAAGCAGGCTATGGAAGGAGACGTCATCCCTTC | Tb927.9.10190_rev | GGGGACCACTTTGTACAAGAAAGCTGGGTTTAGTCAGCCGCATCATACACCA |
| 9_hypothetical | C16 | Tb927.7.1560_for | GGGGACAAGTTTGTACAAAAAAGCAGGCTATGACTCTGGGCGCTGCT | Tb927.7.1560_rev | GGGGACCACTTTGTACAAGAAAGCTGGGTTCACTCCACCTGCTCATTACCC |
| 9_hypothetical | C17 | Tb927.4.3310_for | GGGGACAAGTTTGTACAAAAAAGCAGGCTATGAGTGGTGGTGTGTCCCC | Tb927.4.3310_rev | GGGGACCACTTTGTACAAGAAAGCTGGGTTCACAACGCCCTCTGGCA |
| 9_hypothetical | C18 | Tb927.3.1880_for | GGGGACAAGTTTGTACAAAAAAGCAGGCTATGGGACCTTTAGTTGTTGTATCGC | Tb927.3.1880_rev | GGGGACCACTTTGTACAAGAAAGCTGGGTTCAGTAGACACAAACACAGTTACATCC |
| 9_hypothetical | C19 | Tb927.3.4550_for | GGGGACAAGTTTGTACAAAAAAGCAGGCTATGTTAAACCGCTGTTACATACCACG | Tb927.3.4550_rev | GGGGACCACTTTGTACAAGAAAGCTGGGTTCAAGACGGTACAAAGAGTCGCT |
| 9_hypothetical | C20 | Tb927.9.2620_for | GGGGACAAGTTTGTACAAAAAAGCAGGCTATGTACAAGAAGGCCCTTCATTCCT | Tb927.9.2620_rev | GGGGACCACTTTGTACAAGAAAGCTGGGTTTAAAATCCAGGTTTATCCCCCTGGT |
| 9_hypothetical | C21 | Tb927.3.4120_for | GGGGACAAGTTTGTACAAAAAAGCAGGCTATGAGTAGCGAGGGGCCCC | Tb927.3.4120_rev | GGGGACCACTTTGTACAAGAAAGCTGGGTCTAGCGACGCAGTAAGAAGGC |
| 9_hypothetical | C22 | Tb927.11.8410_for | GGGGACAAGTTTGTACAAAAAAGCAGGCTATGTCACAACTGCCGTTAATTCATACG | Tb927.11.8410_rev | GGGGACCACTTTGTACAAGAAAGCTGGGTTCAAATAGGTGAAACGGGGCG |
| 9_hypothetical | C23 | Tb927.7.5470_for | GGGGACAAGTTTGTACAAAAAAGCAGGCTATGTCCCGCCCCTGTGGC | Tb927.7.5470_rev | GGGGACCACTTTGTACAAGAAAGCTGGGTCTACGTTGCATGCGCTAGTCT |
| 9_hypothetical | C24 | Tb927.8.4860_for | GGGGACAAGTTTGTACAAAAAAGCAGGCTATGCTGTCGCACGCCGTC | Tb927.8.4860_rev | GGGGACCACTTTGTACAAGAAAGCTGGGTTTATACCAATTGCTCCCCAGGCG |
| 9_hypothetical | D1 | Tb927.10.13790_for | GGGGACAAGTTTGTACAAAAAAGCAGGCTATGGCAACCATAGCGGAGTTG | Tb927.10.13790_rev | GGGGACCACTTTGTACAAGAAAGCTGGGTTTACGCGTTGCGAACACCTT |
| 9_hypothetical | D2 | Tb927.2.2360_for | GGGGACAAGTTTGTACAAAAAAGCAGGCTATGTTTAGTGGGAATCCATATATAACCCCT | Tb927.2.2360_rev | GGGGACCACTTTGTACAAGAAAGCTGGGTTTACGTGTATATTTGCATCGCTTGCT |
| 9_hypothetical | D3 | Tb927.9.12340_for | GGGGACAAGTTTGTACAAAAAAGCAGGCTATGGCAACCACGAACCCCC | Tb927.9.12340_rev | GGGGACCACTTTGTACAAGAAAGCTGGGTCTACGCCTCACTAGCCATACGA |
| 9_hypothetical | D4 | Tb927.10.14870_for | GGGGACAAGTTTGTACAAAAAAGCAGGCTATGCCTAAAGTAAAGAGTGGGGCT | Tb927.10.14870_rev | GGGGACCACTTTGTACAAGAAAGCTGGGTTCAACAAGATCGATCATCCTCTCCC |
| 9_hypothetical | D5 | Tb927.4.2850_for | GGGGACAAGTTTGTACAAAAAAGCAGGCTATGCAAAGCACGAAAGGAACGA | Tb927.4.2850_rev | GGGGACCACTTTGTACAAGAAAGCTGGGTTCAGCCTCTCCACACGGAG |
| 9_hypothetical | D6 | Tb927.11.15190_for | GGGGACAAGTTTGTACAAAAAAGCAGGCTATGTCTTTCCAGAAGTCTCCACTTCT | Tb927.11.15190_rev | GGGGACCACTTTGTACAAGAAAGCTGGGTCTAGGCACGGAAGAACTCGTCA |
| 9_hypothetical | D7 | Tb927.11.11450_for | GGGGACAAGTTTGTACAAAAAAGCAGGCTATGCAGGCATGGTACCAATTCG | Tb927.11.11450_rev | GGGGACCACTTTGTACAAGAAAGCTGGGTTCATTGGCCACGAAGTCGC |
| 9_hypothetical | D8 | Tb927.10.4010_for | GGGGACAAGTTTGTACAAAAAAGCAGGCTATGTCATCAGCACTTCAGGCG | Tb927.10.4010_rev | GGGGACCACTTTGTACAAGAAAGCTGGGTTTATCCAGCAAATGCAGCCGG |
| 9_hypothetical | D9 | Tb927.3.1740_for | GGGGACAAGTTTGTACAAAAAAGCAGGCTATGGCTAAGGGTAAAGGTAAAAAGAAGG | Tb927.3.1740_rev | GGGGACCACTTTGTACAAGAAAGCTGGGTTTACTTAACCAACTCAGGGAGAACTGG |
| 9_hypothetical | D10 | Tb927.11.16630_for | GGGGACAAGTTTGTACAAAAAAGCAGGCTATGGCGTGTTCTCTCCGTCA | Tb927.11.16630_rev | GGGGACCACTTTGTACAAGAAAGCTGGGTCTAAATATTTGCTGCAGAACCACCTGC |
| 9_hypothetical | D11 | Tb927.11.3920_for | GGGGACAAGTTTGTACAAAAAAGCAGGCTATGGGTGAATATCCAGACAACGC | Tb927.11.3920_rev | GGGGACCACTTTGTACAAGAAAGCTGGGTTTACATTCCCTCTTTTTTCCCCTTCG |
| 9_hypothetical | D12 | Tb927.10.15670_for | GGGGACAAGTTTGTACAAAAAAGCAGGCTATGTTCTCACAGCTACGAGTTCAC | Tb927.10.15670_rev | GGGGACCACTTTGTACAAGAAAGCTGGGTTTAAAACACTGTCCGTAGTGGTAATATCAT |
| 9_hypothetical | D13 | Tb927.10.11090_for | GGGGACAAGTTTGTACAAAAAAGCAGGCTATGAGGGTTACTGGGAAATGTTTTGT | Tb927.10.11090_rev | GGGGACCACTTTGTACAAGAAAGCTGGGTCTAGTTTTCTTCACTGTCTGTGTTGGC |
| 9_hypothetical | D14 | Tb927.8.1210_for | GGGGACAAGTTTGTACAAAAAAGCAGGCTATGTCTGCGCCTCTTGCAC | Tb927.8.1210_rev | GGGGACCACTTTGTACAAGAAAGCTGGGTTTAGGCAACGCCGCACTTAC |
| 9_hypothetical | D15 | Tb927.4.660_for | GGGGACAAGTTTGTACAAAAAAGCAGGCTATGCCACCTAAAACTGTGAAAGTGA | Tb927.4.660_rev | GGGGACCACTTTGTACAAGAAAGCTGGGTTTAACCCCAAGTAAGTGGGCGA |
| 9_hypothetical | D16 | Tb927.5.700_for | GGGGACAAGTTTGTACAAAAAAGCAGGCTATGGGTGGTGGAGCAGCG | Tb927.5.700_rev | GGGGACCACTTTGTACAAGAAAGCTGGGTCTACAGCCCACATGAGTACTCCA |
| 9_hypothetical | D17 | Tb927.10.3480_for | GGGGACAAGTTTGTACAAAAAAGCAGGCTATGATCAAAGGGAGTGATCCGCT | Tb927.10.3480_rev | GGGGACCACTTTGTACAAGAAAGCTGGGTTCATTGCCTTCGTCGTTGTTCTT |
| 9_hypothetical | D18 | Tb927.10.530_for | GGGGACAAGTTTGTACAAAAAAGCAGGCTATGGGCAATGAACCTGATGAAGC | Tb927.10.530_rev | GGGGACCACTTTGTACAAGAAAGCTGGGTTTAACCTCTCCTATGTGCGGCC |
| 9_hypothetical | D19 | Tb927.9.15335_for | GGGGACAAGTTTGTACAAAAAAGCAGGCTATGCCGGTACGACCACCC | Tb927.9.15335_rev | GGGGACCACTTTGTACAAGAAAGCTGGGTTTAGGCCTCCACTTGCACGG |
| 9_hypothetical | D20 | Tb927.11.3450_for | GGGGACAAGTTTGTACAAAAAAGCAGGCTATGTGGACCAGGCGTAATGTGT | Tb927.11.3450_rev | GGGGACCACTTTGTACAAGAAAGCTGGGTTCACTTTCCCCACTTGTCAATACCG |
| 9_hypothetical | D21 | Tb927.11.5370_for | GGGGACAAGTTTGTACAAAAAAGCAGGCTATGTTAAGAACGAAAAGCACAACCG | Tb927.11.5370_rev | GGGGACCACTTTGTACAAGAAAGCTGGGTCTACTCCATTGCCGTATCCCAGT |
| 9_hypothetical | D22 | Tb927.11.690_for | GGGGACAAGTTTGTACAAAAAAGCAGGCTATGGAGGCCGCTGTATACGAG | Tb927.11.690_rev | GGGGACCACTTTGTACAAGAAAGCTGGGTTCAAACAACAGAAGGGAAGAAAAAAAAATC |
| 9_hypothetical | D23 | Tb927.10.4100_for | GGGGACAAGTTTGTACAAAAAAGCAGGCTATGCAAACGCCGGTATCTATCAC | Tb927.10.4100_rev | GGGGACCACTTTGTACAAGAAAGCTGGGTCTACACCCCAGGTGAACGCT |
| 9_hypothetical | D24 | Tb927.8.6250_for | GGGGACAAGTTTGTACAAAAAAGCAGGCTATGCGCTCCAATGGCGCA | Tb927.8.6250_rev | GGGGACCACTTTGTACAAGAAAGCTGGGTTCACACCTTACTCTTTTCCTTTCGC |
| 9_hypothetical | E1 | Tb927.8.3300_for | GGGGACAAGTTTGTACAAAAAAGCAGGCTATGAGGCGGTTATTTGTAACCACTG | Tb927.8.3300_rev | GGGGACCACTTTGTACAAGAAAGCTGGGTTTATCCGCTGAGTCTCGATTGAGC |
| 9_hypothetical | E2 | Tb927.11.310_for | GGGGACAAGTTTGTACAAAAAAGCAGGCTATGCTGGTGGCAGTTCGAGA | Tb927.11.310_rev | GGGGACCACTTTGTACAAGAAAGCTGGGTCTAACTGGAAGACTCCACACGGG |
| 9_hypothetical | E3 | Tb927.7.3530_for | GGGGACAAGTTTGTACAAAAAAGCAGGCTATGGGGGAGTCCTTCGTCCC | Tb927.7.3530_rev | GGGGACCACTTTGTACAAGAAAGCTGGGTTTACAAAATGTCGGCATCCCCC |
| 9_hypothetical | E4 | Tb927.10.12860_for | GGGGACAAGTTTGTACAAAAAAGCAGGCTATGCCACTAGAAACAGTTAATTTTATGAAT | Tb927.10.12860_rev | GGGGACCACTTTGTACAAGAAAGCTGGGTCTAATCTATGGTAAAAATATGTTTGTTGCT |
| 9_hypothetical | E5 | Tb927.11.13840_for | GGGGACAAGTTTGTACAAAAAAGCAGGCTATGTCCTTTATCGGCCGCCG | Tb927.11.13840_rev | GGGGACCACTTTGTACAAGAAAGCTGGGTCTACGCCTCGTCTTTTGAGGAAA |
| 9_hypothetical | E6 | Tb927.5.2500_for | GGGGACAAGTTTGTACAAAAAAGCAGGCTATGTCTCGACGTGTATTGTGGGA | Tb927.5.2500_rev | GGGGACCACTTTGTACAAGAAAGCTGGGTTCATACACTTTCCCCTTCCTCTTTGC |
| 9_hypothetical | E7 | Tb927.3.5590_for | GGGGACAAGTTTGTACAAAAAAGCAGGCTATGTCAGTATGGCAAAAATGGAAGCT | Tb927.3.5590_rev | GGGGACCACTTTGTACAAGAAAGCTGGGTTTACGCCTTTTGTCCTCGCAAC |
| 9_hypothetical | E8 | Tb927.10.15170_for | GGGGACAAGTTTGTACAAAAAAGCAGGCTATGTCCTCCCTGACATCCGTC | Tb927.10.15170_rev | GGGGACCACTTTGTACAAGAAAGCTGGGTCTACTGGTTTTCCTCCCTCTGTGG |
| 9_hypothetical | E9 | Tb927.7.3820_for | GGGGACAAGTTTGTACAAAAAAGCAGGCTATGGATTTATCGGAGAAGCGACG | Tb927.7.3820_rev | GGGGACCACTTTGTACAAGAAAGCTGGGTCTATACCGAGGCCGGAGGC |
| 9_hypothetical | E10 | Tb927.7.7210_for | GGGGACAAGTTTGTACAAAAAAGCAGGCTATGTCATCCCTGCCAAGTGGT | Tb927.7.7210_rev | GGGGACCACTTTGTACAAGAAAGCTGGGTTCACAGTCCGTATATACGGGACTTG |
| 9_hypothetical | E11 | Tb927.7.4530_for | GGGGACAAGTTTGTACAAAAAAGCAGGCTATGCCCCGGATCCACATATTCG | Tb927.7.4530_rev | GGGGACCACTTTGTACAAGAAAGCTGGGTTCATCCATTCTTCTCGTTCGTTTTACC |
| 9_hypothetical | E12 | Tb927.11.7130_for | GGGGACAAGTTTGTACAAAAAAGCAGGCTATGCAACGGTATCGTGGGAAGC | Tb927.11.7130_rev | GGGGACCACTTTGTACAAGAAAGCTGGGTCTAGCCTGTACAGCAGCACCG |
| 9_hypothetical | E13 | Tb927.9.8510_for | GGGGACAAGTTTGTACAAAAAAGCAGGCTATGAAAGATCTGCCTTCACCCGA | Tb927.9.8510_rev | GGGGACCACTTTGTACAAGAAAGCTGGGTTCATCCTTCACTGTTCAGCAGGG |
| 9_hypothetical | E14 | Tb927.7.4700_for | GGGGACAAGTTTGTACAAAAAAGCAGGCTATGAGCAAGGAGAAGCTACGCT | Tb927.7.4700_rev | GGGGACCACTTTGTACAAGAAAGCTGGGTTTAATCCTCATTCACGGACCACAGT |
| 9_hypothetical | E15 | Tb927.1.4280_for | GGGGACAAGTTTGTACAAAAAAGCAGGCTATGGAGAAGGTAGCAACAACAGC | Tb927.1.4280_rev | GGGGACCACTTTGTACAAGAAAGCTGGGTTCACATCACTGGTTCAACAATCCTAAC |
| 9_hypothetical | E16 | Tb927.11.2680_for | GGGGACAAGTTTGTACAAAAAAGCAGGCTATGTTCTGCGAAGATGGCTCCC | Tb927.11.2680_rev | GGGGACCACTTTGTACAAGAAAGCTGGGTTTACCCAGCCAATGCATGCTG |
| 9_hypothetical | E17 | Tb927.5.1770_for | GGGGACAAGTTTGTACAAAAAAGCAGGCTATGGACAACCAAGGTGATGCTG | Tb927.5.1770_rev | GGGGACCACTTTGTACAAGAAAGCTGGGTTCAGTGGTTTCCATCCGATCGG |
| 9_hypothetical | E18 | Tb927.3.980_for | GGGGACAAGTTTGTACAAAAAAGCAGGCTATGAATAGCGAAGAAGGCCGGC | Tb927.3.980_rev | GGGGACCACTTTGTACAAGAAAGCTGGGTTTATGACTCCCTACTAGGGAACTCACC |
| 9_hypothetical | E19 | Tb927.6.1750_for | GGGGACAAGTTTGTACAAAAAAGCAGGCTATGAGTGGAGCAACTGAGAAACTTTT | Tb927.6.1750_rev | GGGGACCACTTTGTACAAGAAAGCTGGGTCTACGAAGAACCTCCAAAGAGTTCC |
| 9_hypothetical | E20 | Tb927.7.2790_for | GGGGACAAGTTTGTACAAAAAAGCAGGCTATGGACGGGGATGATGGCA | Tb927.7.2790_rev | GGGGACCACTTTGTACAAGAAAGCTGGGTTCACAGCACGTTTGACGGC |
| 9_hypothetical | E21 | Tb927.10.12180_for | GGGGACAAGTTTGTACAAAAAAGCAGGCTATGGGAAGAAATAAGTCCCATGCG | Tb927.10.12180_rev | GGGGACCACTTTGTACAAGAAAGCTGGGTTCAAAGCTTGAAAAACTTAATACTCAGCTT |
| 9_hypothetical | E22 | Tb927.7.2490_for | GGGGACAAGTTTGTACAAAAAAGCAGGCTATGCTTCAAAAGGCAGGCCG | Tb927.7.2490_rev | GGGGACCACTTTGTACAAGAAAGCTGGGTTCAAACAGCGCTGTTGCCG |
| 9_hypothetical | E23 | Tb927.9.3870_for | GGGGACAAGTTTGTACAAAAAAGCAGGCTATGAAAAAGAGGAGGCCAATACCCA | Tb927.9.3870_rev | GGGGACCACTTTGTACAAGAAAGCTGGGTCTACGACATATCATTCGCAGTCGC |
| 9_hypothetical | E24 | Tb927.10.15640_for | GGGGACAAGTTTGTACAAAAAAGCAGGCTATGAGCTCTACATTTATTATATCCGCTTCT | Tb927.10.15640_rev | GGGGACCACTTTGTACAAGAAAGCTGGGTCTATTGGTTTTTCTTCCTTTCCACATAGAG |
| 9_hypothetical | F1 | Tb927.9.1980_for | GGGGACAAGTTTGTACAAAAAAGCAGGCTATGGTGGTTGTGGACAATGCTC | Tb927.9.1980_rev | GGGGACCACTTTGTACAAGAAAGCTGGGTCTACCGACTCGTCATGTTGTATGAC |
| 9_hypothetical | F2 | Tb927.11.15730_for | GGGGACAAGTTTGTACAAAAAAGCAGGCTATGGACAGCAGGTGGGCG | Tb927.11.15730_rev | GGGGACCACTTTGTACAAGAAAGCTGGGTTTACACCACACTCGACCGCC |
| 9_hypothetical | F3 | Tb927.8.3980_for | GGGGACAAGTTTGTACAAAAAAGCAGGCTATGGATAGTGTCGCGGAGCC | Tb927.8.3980_rev | GGGGACCACTTTGTACAAGAAAGCTGGGTCTAGCGTTTGCTTCGACAATCGC |
| 9_hypothetical | F4 | Tb927.8.8010_for | GGGGACAAGTTTGTACAAAAAAGCAGGCTATGATGAGTGCCAATAACTGCACC | Tb927.8.8010_rev | GGGGACCACTTTGTACAAGAAAGCTGGGTTCAAGTTGGGTTGGCAGCAG |
| 9_hypothetical | F5 | Tb927.9.9190_for | GGGGACAAGTTTGTACAAAAAAGCAGGCTATGACGACTATATCCGCTTTAGAAAAACA | Tb927.9.9190_rev | GGGGACCACTTTGTACAAGAAAGCTGGGTCTAAAACAGTTTCGATAGCAACTCAGACC |
| 9_hypothetical | F6 | Tb927.10.1980_for | GGGGACAAGTTTGTACAAAAAAGCAGGCTATGACCGATAATAAGGTTCTCTACGTTGG | Tb927.10.1980_rev | GGGGACCACTTTGTACAAGAAAGCTGGGTTTACGCTGCTACTGCAGTCCT |
| 9_hypothetical | F7 | Tb927.4.2300_for | GGGGACAAGTTTGTACAAAAAAGCAGGCTATGTCTGCGGTTTCTTCTCCTAAGC | Tb927.4.2300_rev | GGGGACCACTTTGTACAAGAAAGCTGGGTTTATACATCATCGTCCCATCCCCAA |
| 9_hypothetical | F8 | Tb927.3.5120_for | GGGGACAAGTTTGTACAAAAAAGCAGGCTATGTGGAAAACTAAGGGGCGTG | Tb927.3.5120_rev | GGGGACCACTTTGTACAAGAAAGCTGGGTTTATGAATTTTGCGGCGTCTCCA |
| 9_hypothetical | F9 | Tb927.7.5300_for | GGGGACAAGTTTGTACAAAAAAGCAGGCTATGAATGTCGATGGTACTGTCCAAC | Tb927.7.5300_rev | GGGGACCACTTTGTACAAGAAAGCTGGGTTTACATTGTTTCAGCCAACATCAAAGC |
| 9_hypothetical | F10 | Tb927.11.2790_for | GGGGACAAGTTTGTACAAAAAAGCAGGCTATGTACTACCGAACGCAGCAGA | Tb927.11.2790_rev | GGGGACCACTTTGTACAAGAAAGCTGGGTTCACTCTTTCGTCTCTTTTTCATTTGGA |
| 9_hypothetical | F11 | Tb927.10.11880_for | GGGGACAAGTTTGTACAAAAAAGCAGGCTATGACAATAAGCAGCCCGTGC | Tb927.10.11880_rev | GGGGACCACTTTGTACAAGAAAGCTGGGTCTACTCCCAGGTTGGGATTAGCG |
| 9_hypothetical | F12 | Tb927.6.2090_for | GGGGACAAGTTTGTACAAAAAAGCAGGCTATGGGTTTGTGCAATTCTAAAAGTTTTGA | Tb927.6.2090_rev | GGGGACCACTTTGTACAAGAAAGCTGGGTTTAGGCCGCCTTGTTCCAGG |
| 9_hypothetical | F13 | Tb927.8.1950_for | GGGGACAAGTTTGTACAAAAAAGCAGGCTATGCTCTCGTGGAATCCCGA | Tb927.8.1950_rev | GGGGACCACTTTGTACAAGAAAGCTGGGTTTAAGCTGGGATGAGGCGCT |
| 9_hypothetical | F14 | Tb927.5.2310_for | GGGGACAAGTTTGTACAAAAAAGCAGGCTATGCGCGTGCGAACAAAGC | Tb927.5.2310_rev | GGGGACCACTTTGTACAAGAAAGCTGGGTCTACGGGCTATCAACTCTGTTCTCC |
| 9_hypothetical | F15 | Tb927.6.3990_for | GGGGACAAGTTTGTACAAAAAAGCAGGCTATGTCCGACCATGTGGTATTTGGA | Tb927.6.3990_rev | GGGGACCACTTTGTACAAGAAAGCTGGGTTCACTCATCATTTACCTCCCCTTCC |
| 9_hypothetical | F16 | Tb927.4.2680_for | GGGGACAAGTTTGTACAAAAAAGCAGGCTATGCAACCACCTTCACCAACTG | Tb927.4.2680_rev | GGGGACCACTTTGTACAAGAAAGCTGGGTTTAGGATTGCTTGGGGTTGTAGAGC |
| 9_hypothetical | F17 | Tb927.11.3480_for | GGGGACAAGTTTGTACAAAAAAGCAGGCTATGCAGCAGACCGAAAAGCG | Tb927.11.3480_rev | GGGGACCACTTTGTACAAGAAAGCTGGGTTCACTGTCCAATGACATTGCCAC |
| 9_hypothetical | F18 | Tb927.11.14820_for | GGGGACAAGTTTGTACAAAAAAGCAGGCTATGGCTGCTCTGCGTAGTGT | Tb927.11.14820_rev | GGGGACCACTTTGTACAAGAAAGCTGGGTTTACTCTTTACGCAGCCTTTTATAGTCTC |
| 9_hypothetical | F19 | Tb927.10.7810_for | GGGGACAAGTTTGTACAAAAAAGCAGGCTATGAAGACACTACAAAGCTCCTCCC | Tb927.10.7810_rev | GGGGACCACTTTGTACAAGAAAGCTGGGTCTACTCCCTTTTTGCTTTCTTACTCGG |
| 9_hypothetical | F20 | Tb927.1.2710_for | GGGGACAAGTTTGTACAAAAAAGCAGGCTATGGACTCGCTACGCTCGC | Tb927.1.2710_rev | GGGGACCACTTTGTACAAGAAAGCTGGGTTTAATTTTTTTCTTTTGTTGGATCTACAGA |
| 9_hypothetical | F21 | Tb927.10.13380_for | GGGGACAAGTTTGTACAAAAAAGCAGGCTATGAAAGGAGTATTTTGTGAAAGCGGT | Tb927.10.13380_rev | GGGGACCACTTTGTACAAGAAAGCTGGGTTCACACGAGGCTGCGTAATCG |
| 9_hypothetical | F22 | Tb927.8.2710_for | GGGGACAAGTTTGTACAAAAAAGCAGGCTATGGCCACTGTCACGGAGG | Tb927.8.2710_rev | GGGGACCACTTTGTACAAGAAAGCTGGGTTTAGGAAGTCCCCCCCCACG |
| 9_hypothetical | F23 | Tb927.11.6580_for | GGGGACAAGTTTGTACAAAAAAGCAGGCTATGGAAGAAATCATCGAGTTGTGTGC | Tb927.11.6580_rev | GGGGACCACTTTGTACAAGAAAGCTGGGTTCAGCGGGTATGCTGAACAATAC |
| 9_hypothetical | F24 | Tb927.7.910_for | GGGGACAAGTTTGTACAAAAAAGCAGGCTATGAGAAGCTGCTCGCGTCA | Tb927.7.910_rev | GGGGACCACTTTGTACAAGAAAGCTGGGTTTAAACTTGGCCACTTCCGCC |
| 9_hypothetical | G1 | Tb927.2.5200_for | GGGGACAAGTTTGTACAAAAAAGCAGGCTATGGATGAGCAAAACAGTGAGAGC | Tb927.2.5200_rev | GGGGACCACTTTGTACAAGAAAGCTGGGTCTAATTGTCGGAATCGCTATCGCT |
| 9_hypothetical | G2 | Tb927.10.14860_for | GGGGACAAGTTTGTACAAAAAAGCAGGCTATGCTTCGTCGAGCATGCC | Tb927.10.14860_rev | GGGGACCACTTTGTACAAGAAAGCTGGGTCTAAATCTGCGAGACAACCACATTTC |
| 9_hypothetical | G3 | Tb927.10.590_for | GGGGACAAGTTTGTACAAAAAAGCAGGCTATGTCCAGAAACCCCCCTGATAG | Tb927.10.590_rev | GGGGACCACTTTGTACAAGAAAGCTGGGTTTACTCATCATAAGGTAAATACTGCAGTAA |
| 9_hypothetical | G4 | Tb927.11.340_for | GGGGACAAGTTTGTACAAAAAAGCAGGCTATGACTATGTGGTTTCCTGCCGA | Tb927.11.340_rev | GGGGACCACTTTGTACAAGAAAGCTGGGTCTAAGCATCTTCTATATCATATGGTGTTTT |
| 9_hypothetical | G5 | Tb927.10.6110_for | GGGGACAAGTTTGTACAAAAAAGCAGGCTATGAATCGGGGTAAGGGGGG | Tb927.10.6110_rev | GGGGACCACTTTGTACAAGAAAGCTGGGTCTATACCCTACCAGATTCACGCAGT |
| 9_hypothetical | G6 | Tb927.10.1320_for | GGGGACAAGTTTGTACAAAAAAGCAGGCTATGGCCTTCCTCCTGAGGGC | Tb927.10.1320_rev | GGGGACCACTTTGTACAAGAAAGCTGGGTCTACGACAGGCGAGAGGCG |
| 9_hypothetical | G7 | Tb927.8.8130_for | GGGGACAAGTTTGTACAAAAAAGCAGGCTATGGTAACGCAGCAGCGAATG | Tb927.8.8130_rev | GGGGACCACTTTGTACAAGAAAGCTGGGTTTAATCCCATGGAGAAAAGAAGGAGAAAC |
| 9_hypothetical | G8 | Tb927.1.4340_for | GGGGACAAGTTTGTACAAAAAAGCAGGCTATGAGTAGTACTGAAAGCAGCGGT | Tb927.1.4340_rev | GGGGACCACTTTGTACAAGAAAGCTGGGTCTAACCTCTCACTACGGCACTACT |
| 9_hypothetical | G9 | Tb927.5.4060_for | GGGGACAAGTTTGTACAAAAAAGCAGGCTATGTTACAAGGTGTTGATCACTTTCTTGC | Tb927.5.4060_rev | GGGGACCACTTTGTACAAGAAAGCTGGGTCTATACTGCAGGTGTTTTCCTCAAACA |
| 9_hypothetical | G10 | Tb927.10.1190_for | GGGGACAAGTTTGTACAAAAAAGCAGGCTATGGCTTCCCACAGTAGAGGATG | Tb927.10.1190_rev | GGGGACCACTTTGTACAAGAAAGCTGGGTTTATTTCTTCCGGGTCCGACTCT |
| 9_hypothetical | G11 | Tb927.11.1170_for | GGGGACAAGTTTGTACAAAAAAGCAGGCTATGAAGTTTTGGGTGAAGGCAATAACG | Tb927.11.1170_rev | GGGGACCACTTTGTACAAGAAAGCTGGGTTCACGGTTCTGTCGTTAGTGGT |
| 9_hypothetical | G12 | Tb927.10.14570_for | GGGGACAAGTTTGTACAAAAAAGCAGGCTATGGAGAATTCCAGTTCGTTCGAC | Tb927.10.14570_rev | GGGGACCACTTTGTACAAGAAAGCTGGGTCTACTCTACATCCAAGCCCCATCC |
| 9_hypothetical | G13 | Tb927.7.7320_for | GGGGACAAGTTTGTACAAAAAAGCAGGCTATGTACCCACAGCAACAAATGAGC | Tb927.7.7320_rev | GGGGACCACTTTGTACAAGAAAGCTGGGTCTACACTTCATTCAGCATCCTATCGATG |
| 9_hypothetical | G14 | Tb927.10.7370_for | GGGGACAAGTTTGTACAAAAAAGCAGGCTATGGCAGTTGACTGTTTGGCG | Tb927.10.7370_rev | GGGGACCACTTTGTACAAGAAAGCTGGGTTCAAACTTCAGCGACGACAGC |
| 9_hypothetical | G15 | Tb927.6.3710_for | GGGGACAAGTTTGTACAAAAAAGCAGGCTATGCAGGAGCCCGAAGTAGC | Tb927.6.3710_rev | GGGGACCACTTTGTACAAGAAAGCTGGGTTTAAGACACGCGGGTCCGG |
| 9_hypothetical | G16 | Tb927.4.1480_for | GGGGACAAGTTTGTACAAAAAAGCAGGCTATGGCAGGTGGAGCTGCA | Tb927.4.1480_rev | GGGGACCACTTTGTACAAGAAAGCTGGGTTTATTCTGAGTCACTACTCAGGTCAATCA |

| | | | | | |
|---|---|---|---|---|---|
| 9_hypothetical | G17 | Tb927.9.9860_for | GGGGACAAGTTTGTACAAAAAAGCAGGCTATGAGAGTGCTTGCAAATGAAATCAGG | Tb927.9.9860_rev | GGGGACCACTTTGTACAAGAAAGCTGGGTTCACAAGTCTCCCTTGACAGCA |
| 9_hypothetical | G18 | Tb927.11.13320_for | GGGGACAAGTTTGTACAAAAAAGCAGGCTATGCGAATTGTCCGCATTCAGT | Tb927.11.13320_rev | GGGGACCACTTTGTACAAGAAAGCTGGGTTCATCTGCCGTGCGTTAAACG |
| 9_hypothetical | G19 | Tb927.6.2380_for | GGGGACAAGTTTGTACAAAAAAGCAGGCTATGCTTAGCATCACACGGTGC | Tb927.6.2380_rev | GGGGACCACTTTGTACAAGAAAGCTGGGTTCACAATTCACCAAACTGTTCCTGG |
| 9_hypothetical | G20 | Tb927.9.6580_for | GGGGACAAGTTTGTACAAAAAAGCAGGCTATGGTTCGTGTGAAGGAAAAGGC | Tb927.9.6580_rev | GGGGACCACTTTGTACAAGAAAGCTGGGTTCACGCACAAACACCGGG |
| 9_hypothetical | G21 | Tb927.10.2760_for | GGGGACAAGTTTGTACAAAAAAGCAGGCTATGGAAGGGCGCCACGGC | Tb927.10.2760_rev | GGGGACCACTTTGTACAAGAAAGCTGGGTTTAATATTGACGCTCCAACTCCAGTAGT |
| 9_hypothetical | G22 | Tb927.8.1300_for | GGGGACAAGTTTGTACAAAAAAGCAGGCTATGAGTCGAAGAAATTCGGCTTCC | Tb927.8.1300_rev | GGGGACCACTTTGTACAAGAAAGCTGGGTTTACGATGAAGGCCCTGTGGG |
| 9_hypothetical | G23 | Tb927.11.15120_for | GGGGACAAGTTTGTACAAAAAAGCAGGCTATGAGCTCATCAGAGTCAGGGAG | Tb927.11.15120_rev | GGGGACCACTTTGTACAAGAAAGCTGGGTCTAAAAATCTAGCCAGCTGGCACC |
| 9_hypothetical | G24 | Tb927.10.250_for | GGGGACAAGTTTGTACAAAAAAGCAGGCTATGTCAGAACTGACGGCCGG | Tb927.10.250_rev | GGGGACCACTTTGTACAAGAAAGCTGGGTTCACATTTTGTGATAAAGCAGCTTTCC |
| 9_hypothetical | H1 | Tb927.7.5200_for | GGGGACAAGTTTGTACAAAAAAGCAGGCTATGGAAGATGACATTATACAAGGTCGC | Tb927.7.5200_rev | GGGGACCACTTTGTACAAGAAAGCTGGGTCTATGCAACCGGCTGGTCTAAC |
| 9_hypothetical | H2 | Tb927.11.7570_for | GGGGACAAGTTTGTACAAAAAAGCAGGCTATGCCGAAGGAGGAGCAGT | Tb927.11.7570_rev | GGGGACCACTTTGTACAAGAAAGCTGGGTTTACACACCACAGATGAGTTGAGTAGC |
| 9_hypothetical | H3 | Tb927.5.770_for | GGGGACAAGTTTGTACAAAAAAGCAGGCTATGATGCATCGCACTGCAGC | Tb927.5.770_rev | GGGGACCACTTTGTACAAGAAAGCTGGGTTTATGTGTACTTGTGATATACGTCATTCCC |
| 9_hypothetical | H4 | Tb927.10.11420_for | GGGGACAAGTTTGTACAAAAAAGCAGGCTATGCGAGGCCGATTCTTGTC | Tb927.10.11420_rev | GGGGACCACTTTGTACAAGAAAGCTGGGTTTAAATATGAAGCAGCTTCGCAGCC |
| 9_hypothetical | H5 | Tb927.9.10550_for | GGGGACAAGTTTGTACAAAAAAGCAGGCTATGAGTCGTAGCGTAGACACAAGT | Tb927.9.10550_rev | GGGGACCACTTTGTACAAGAAAGCTGGGTTTAGAAGTATGAGCTCCTCACATTGTCA |
| 9_hypothetical | H6 | Tb927.11.13110_for | GGGGACAAGTTTGTACAAAAAAGCAGGCTATGGGGTCTCCAGTGAAAGACAC | Tb927.11.13110_rev | GGGGACCACTTTGTACAAGAAAGCTGGGTTCATATGAAGTACGCGGCAGACA |
| 9_hypothetical | H7 | Tb927.7.6750_for | GGGGACAAGTTTGTACAAAAAAGCAGGCTATGAGCTCCTACTGTGATATTTGCCG | Tb927.7.6750_rev | GGGGACCACTTTGTACAAGAAAGCTGGGTTCACAACCCCGGACATGCC |
| 9_hypothetical | H8 | Tb927.9.2270_for | GGGGACAAGTTTGTACAAAAAAGCAGGCTATGTTACGGCGCTCCTTCATG | Tb927.9.2270_rev | GGGGACCACTTTGTACAAGAAAGCTGGGTTCAGTGCAGAGCAGCCGC |
| 9_hypothetical | H9 | Tb927.2.4140_for | GGGGACAAGTTTGTACAAAAAAGCAGGCTATGAAGCGAGAGCGCTCATTTT | Tb927.2.4140_rev | GGGGACCACTTTGTACAAGAAAGCTGGGTTTAAAAGCGTTTCATTTGTCCATGACC |
| 9_hypothetical | H10 | Tb927.11.3970_for | GGGGACAAGTTTGTACAAAAAAGCAGGCTATGGCAGCTAACCGAAGTATATTTCTTT | Tb927.11.3970_rev | GGGGACCACTTTGTACAAGAAAGCTGGGTCTATTTTCGTTCCCTCTTTTTCGTTGC |
| 9_hypothetical | H11 | Tb927.3.2660_for | GGGGACAAGTTTGTACAAAAAAGCAGGCTATGGGATTTGTCGACCGTGGT | Tb927.3.2660_rev | GGGGACCACTTTGTACAAGAAAGCTGGGTCTACTCAGAAGGGGTTTCACTTACGC |
| 9_hypothetical | H12 | Tb927.10.7890_for | GGGGACAAGTTTGTACAAAAAAGCAGGCTATGTTGCGTCATGGTTCCCAC | Tb927.10.7890_rev | GGGGACCACTTTGTACAAGAAAGCTGGGTCTATATGGCCACATTCAGGTACCCA |
| 9_hypothetical | H13 | Tb927.11.3960_for | GGGGACAAGTTTGTACAAAAAAGCAGGCTATGAATGATCGGCCACATTTTTATGAAG | Tb927.11.3960_rev | GGGGACCACTTTGTACAAGAAAGCTGGGTTCACAATGAGATAAGGTAAGAGTCATTGAG |
| 9_hypothetical | H14 | Tb927.7.5250_for | GGGGACAAGTTTGTACAAAAAAGCAGGCTATGGGCCATTGTTGTGCCAC | Tb927.7.5250_rev | GGGGACCACTTTGTACAAGAAAGCTGGGTTTATGATTGACAGTGTGCCTTTCCG |
| 9_hypothetical | H15 | Tb927.6.630_for | GGGGACAAGTTTGTACAAAAAAGCAGGCTATGATGGAGTTGTTACAACTACCCAC | Tb927.6.630_rev | GGGGACCACTTTGTACAAGAAAGCTGGGTTTATCGGCAGCGCAGCGG |
| 9_hypothetical | H16 | Tb927.9.1730_for | GGGGACAAGTTTGTACAAAAAAGCAGGCTATGGCACGAAGTGAGGGGAA | Tb927.9.1730_rev | GGGGACCACTTTGTACAAGAAAGCTGGGTTTACGCATCCTCTCGAAGGCT |
| 9_hypothetical | H17 | Tb927.10.6160_for | GGGGACAAGTTTGTACAAAAAAGCAGGCTATGTGCACCTCTTTTGACGCC | Tb927.10.6160_rev | GGGGACCACTTTGTACAAGAAAGCTGGGTTCATTCACGTCGTTCCTGCG |
| 9_hypothetical | H18 | Tb927.1.4520_for | GGGGACAAGTTTGTACAAAAAAGCAGGCTATGTACAGCGGGGATCGTGC | Tb927.1.4520_rev | GGGGACCACTTTGTACAAGAAAGCTGGGTTCACGACTTCTCTACTTCCGCC |
| 9_hypothetical | H19 | Tb927.5.3870_for | GGGGACAAGTTTGTACAAAAAAGCAGGCTATGAGGCGCTGCGGGAAT | Tb927.5.3870_rev | GGGGACCACTTTGTACAAGAAAGCTGGGTTTAAGGATACGTCGTAACCAAACTCCC |
| 9_hypothetical | H20 | Tb927.11.210_for | GGGGACAAGTTTGTACAAAAAAGCAGGCTATGGATGTTGATAGTGACAGGCGC | Tb927.11.210_rev | GGGGACCACTTTGTACAAGAAAGCTGGGTCTAGCCCCCAGATCTCTCAGGT |
| 9_hypothetical | H21 | Tb927.10.13610_for | GGGGACAAGTTTGTACAAAAAAGCAGGCTATGCTGTCGTCCCGTAGCA | Tb927.10.13610_rev | GGGGACCACTTTGTACAAGAAAGCTGGGTCTAAAACCGCCTTCGCGGA |
| 9_hypothetical | H22 | Tb927.11.14850_for | GGGGACAAGTTTGTACAAAAAAGCAGGCTATGTCTGATCAGGACGGCAACC | Tb927.11.14850_rev | GGGGACCACTTTGTACAAGAAAGCTGGGTTTAGCGCTCAAACAGGAAGACG |
| 9_hypothetical | H23 | Tb927.9.3860_for | GGGGACAAGTTTGTACAAAAAAGCAGGCTATGGGAGGGACGTCTACGAAG | Tb927.9.3860_rev | GGGGACCACTTTGTACAAGAAAGCTGGGTTTAGTGAGGGAATATCCCCAACGT |
| 9_hypothetical | H24 | Tb927.11.8240_for | GGGGACAAGTTTGTACAAAAAAGCAGGCTATGTCTGATGTTCCGGAGCTTCC | Tb927.11.8240_rev | GGGGACCACTTTGTACAAGAAAGCTGGGTCTACACAAAATATGTTGTAGTCAATCGATC |
| 9_hypothetical | I1 | Tb927.10.11570_for | GGGGACAAGTTTGTACAAAAAAGCAGGCTATGTCGTCGCGCTTAACGC | Tb927.10.11570_rev | GGGGACCACTTTGTACAAGAAAGCTGGGTCTAACCGGCAATATTACGATCTTCAGA |
| 9_hypothetical | I2 | Tb927.7.4640_for | GGGGACAAGTTTGTACAAAAAAGCAGGCTATGCGATTGGGGATTCAACAGAAT | Tb927.7.4640_rev | GGGGACCACTTTGTACAAGAAAGCTGGGTTTACATGGCGCGATGCACG |
| 9_hypothetical | I3 | Tb927.9.1530_for | GGGGACAAGTTTGTACAAAAAAGCAGGCTATGCCACCACGTGTAACTGAGA | Tb927.9.1530_rev | GGGGACCACTTTGTACAAGAAAGCTGGGTTCACCACACTGCCCGATCC |
| 9_hypothetical | I4 | Tb927.7.3570_for | GGGGACAAGTTTGTACAAAAAAGCAGGCTATGGAGGACGAGTGGAACACC | Tb927.7.3570_rev | GGGGACCACTTTGTACAAGAAAGCTGGGTCTAGTTTATCCCAAGTCGCTTCGC |
| 9_hypothetical | I5 | Tb927.11.12870_for | GGGGACAAGTTTGTACAAAAAAGCAGGCTATGGAGAGTAAAACGTGGTTCGGC | Tb927.11.12870_rev | GGGGACCACTTTGTACAAGAAAGCTGGGTCTAAACAATGCCTGCAATGCGC |
| 9_hypothetical | I6 | Tb927.5.1540_for | GGGGACAAGTTTGTACAAAAAAGCAGGCTATGAGTAAGAAGCCGAATCCGCA | Tb927.5.1540_rev | GGGGACCACTTTGTACAAGAAAGCTGGGTCTATACATTGGGGAATTCTATGTCACCTTC |
| 9_hypothetical | I7 | Tb927.7.5730_for | GGGGACAAGTTTGTACAAAAAAGCAGGCTATGATGGAGGGAGCAACGCA | Tb927.7.5730_rev | GGGGACCACTTTGTACAAGAAAGCTGGGTTCAACTCCTTTTCCGTTTTTGTCTAGA |
| 9_hypothetical | I8 | Tb927.5.2320_for | GGGGACAAGTTTGTACAAAAAAGCAGGCTATGTTGTGCGCGTTGGATGA | Tb927.5.2320_rev | GGGGACCACTTTGTACAAGAAAGCTGGGTTCAAGCGGTTGTCCTCTTCCT |
| 9_hypothetical | I9 | Tb927.8.3010_for | GGGGACAAGTTTGTACAAAAAAGCAGGCTATGAAATATAGTGGTGACACTCCTGTGG | Tb927.8.3010_rev | GGGGACCACTTTGTACAAGAAAGCTGGGTTTACGTGGACTTCATATAGCTCAGCC |
| 9_hypothetical | I10 | Tb927.9.15070_for | GGGGACAAGTTTGTACAAAAAAGCAGGCTATGCCGTGGCAGTGGCTC | Tb927.9.15070_rev | GGGGACCACTTTGTACAAGAAAGCTGGGTTTAACTAGGGTTATTGGCTAAATACTCCGC |
| 9_hypothetical | I11 | Tb927.7.1100_for | GGGGACAAGTTTGTACAAAAAAGCAGGCTATGAATACCACCGGGGAGCC | Tb927.7.1100_rev | GGGGACCACTTTGTACAAGAAAGCTGGGTCTATAAAGGTTCAGGTGCTACACCAC |
| 9_hypothetical | I12 | Tb927.2.3750_for | GGGGACAAGTTTGTACAAAAAAGCAGGCTATGAGTCTTCGCAGATTCTACAATGAG | Tb927.2.3750_rev | GGGGACCACTTTGTACAAGAAAGCTGGGTTCAAGTGAAATAAGTCGAATCGTCTCG |
| 9_hypothetical | I13 | Tb927.6.1120_for | GGGGACAAGTTTGTACAAAAAAGCAGGCTATGAGTGATGTTCTCGCTATACTCGA | Tb927.6.1120_rev | GGGGACCACTTTGTACAAGAAAGCTGGGTTCAGCTAAGCGCAAGCAAGG |
| 9_hypothetical | I14 | Tb927.10.13660_for | GGGGACAAGTTTGTACAAAAAAGCAGGCTATGGTATTTGGCGTAAGGCGATG | Tb927.10.13660_rev | GGGGACCACTTTGTACAAGAAAGCTGGGTCTACTTGGGAGGGTTTAGAAGCTTCA |
| 9_hypothetical | I15 | Tb927.5.3880_for | GGGGACAAGTTTGTACAAAAAAGCAGGCTATGGTCGGCAGCGCATCG | Tb927.5.3880_rev | GGGGACCACTTTGTACAAGAAAGCTGGGTTCATCTCCCTCCCGTATCCGC |
| 9_hypothetical | I16 | Tb927.7.6730_for | GGGGACAAGTTTGTACAAAAAAGCAGGCTATGGAGTTTGAGGCCGGGC | Tb927.7.6730_rev | GGGGACCACTTTGTACAAGAAAGCTGGGTCTATGGTGCGTACGTGCTCC |
| 9_hypothetical | I17 | Tb927.10.8080_for | GGGGACAAGTTTGTACAAAAAAGCAGGCTATGAGTGAGTTGGAGCTGAGTGA | Tb927.10.8080_rev | GGGGACCACTTTGTACAAGAAAGCTGGGTCTACGCATCCTCAAGTACAAGCC |
| 9_hypothetical | I18 | Tb927.11.930_for | GGGGACAAGTTTGTACAAAAAAGCAGGCTATGCCTCACAAGGAGGCTGA | Tb927.11.930_rev | GGGGACCACTTTGTACAAGAAAGCTGGGTCTACACCTCCTCTGCTTCTTCAACA |
| 9_hypothetical | I19 | Tb927.7.4920_for | GGGGACAAGTTTGTACAAAAAAGCAGGCTATGTGGATCGTGTTTATGAATGGTCA | Tb927.7.4920_rev | GGGGACCACTTTGTACAAGAAAGCTGGGTCTACAGGAATGCTTCCAGGTCGA |
| 9_hypothetical | I20 | Tb927.9.5380_for | GGGGACAAGTTTGTACAAAAAAGCAGGCTATGATGACAAAAACCTATGGGTCGAAA | Tb927.9.5380_rev | GGGGACCACTTTGTACAAGAAAGCTGGGTTCACTTCGCCATAAACAGAAGGTATTG |
| 9_hypothetical | I21 | Tb927.4.1150_for | GGGGACAAGTTTGTACAAAAAAGCAGGCTATGACATGTAGGAGTGAGGAAACTAACT | Tb927.4.1150_rev | GGGGACCACTTTGTACAAGAAAGCTGGGTTTACTCCAACGTCCATCCCTTCC |
| 9_hypothetical | I22 | Tb927.8.3840_for | GGGGACAAGTTTGTACAAAAAAGCAGGCTATGCCCGAGGCGCATATTG | Tb927.8.3840_rev | GGGGACCACTTTGTACAAGAAAGCTGGGTCTAACTACCAAAGTTGAAAGTAAGCGTG |
| 9_hypothetical | I23 | Tb927.11.16460_for | GGGGACAAGTTTGTACAAAAAAGCAGGCTATGCGCTGGACCACTCTGT | Tb927.11.16460_rev | GGGGACCACTTTGTACAAGAAAGCTGGGTTTAATAGAAGGAAAACCGGGGCTCT |
| 9_hypothetical | I24 | Tb927.11.15310_for | GGGGACAAGTTTGTACAAAAAAGCAGGCTATGGACCCGACCTTACGCA | Tb927.11.15310_rev | GGGGACCACTTTGTACAAGAAAGCTGGGTTCAACTATCAGCCGCTTCCGT |
| 9_hypothetical | J1 | Tb927.10.11550_for | GGGGACAAGTTTGTACAAAAAAGCAGGCTATGGACAATCAGCGCAATACAGAA | Tb927.10.11550_rev | GGGGACCACTTTGTACAAGAAAGCTGGGTCTACATCAACGTAAACTCATCCTCTTCG |
| 9_hypothetical | J2 | Tb927.5.3900_for | GGGGACAAGTTTGTACAAAAAAGCAGGCTATGGGCAAGGGTCGCGAC | Tb927.5.3900_rev | GGGGACCACTTTGTACAAGAAAGCTGGGTTCAACTATTTTCATCTCCTTCACCTTTTTC |
| 9_hypothetical | J3 | Tb927.10.6590_for | GGGGACAAGTTTGTACAAAAAAGCAGGCTATGCCCTCCTCTCAGTGGGT | Tb927.10.6590_rev | GGGGACCACTTTGTACAAGAAAGCTGGGTTCAGGGCTTCAACCTACTTATCGC |
| 9_hypothetical | J4 | Tb927.7.3010_for | GGGGACAAGTTTGTACAAAAAAGCAGGCTATGAAGTTCCACCTTCAAAGCGT | Tb927.7.3010_rev | GGGGACCACTTTGTACAAGAAAGCTGGGTTCATTGAGCAACCAGCCAACC |
| 9_hypothetical | J5 | Tb927.3.2130_for | GGGGACAAGTTTGTACAAAAAAGCAGGCTATGCATCACGCTCGTGGGC | Tb927.3.2130_rev | GGGGACCACTTTGTACAAGAAAGCTGGGTTCACTCACAGGACAGACGAAAAATG |
| 9_hypothetical | J6 | Tb927.8.5050_for | GGGGACAAGTTTGTACAAAAAAGCAGGCTATGGGTGCCTGTGTTTCAGATG | Tb927.8.5050_rev | GGGGACCACTTTGTACAAGAAAGCTGGGTTTATTCAACAGGTCTCGAGATTGTATCCA |
| 9_hypothetical | J7 | Tb927.7.5320_for | GGGGACAAGTTTGTACAAAAAAGCAGGCTATGGAAACACAGGTACCTCAGAATGT | Tb927.7.5320_rev | GGGGACCACTTTGTACAAGAAAGCTGGGTCTACCATCGTCTAATGAACTTAATTTGCAT |
| 9_hypothetical | J8 | Tb927.7.4990_for | GGGGACAAGTTTGTACAAAAAAGCAGGCTATGTCTGGTTCTTTCCGAAAAATATGTGA | Tb927.7.4990_rev | GGGGACCACTTTGTACAAGAAAGCTGGGTTCACCGCTCGGCTAGAAGC |
| 9_hypothetical | J9 | Tb927.10.7130_for | GGGGACAAGTTTGTACAAAAAAGCAGGCTATGGCTTCCGACTTGCAGAAG | Tb927.10.7130_rev | GGGGACCACTTTGTACAAGAAAGCTGGGTTTAGCGCAGCAACTTCCTCAC |
| 9_hypothetical | J10 | Tb927.7.5290_for | GGGGACAAGTTTGTACAAAAAAGCAGGCTATGTCTCACAGCCAGAACAGCA | Tb927.7.5290_rev | GGGGACCACTTTGTACAAGAAAGCTGGGTTCAGTCATCATTCTGCACCAGC |
| 9_hypothetical | J11 | Tb927.7.1650_for | GGGGACAAGTTTGTACAAAAAAGCAGGCTATGGGCAAAAGACGGAGGTCC | Tb927.7.1650_rev | GGGGACCACTTTGTACAAGAAAGCTGGGTCTATCCTTCGCCGTCACTCTTG |
| 9_hypothetical | J12 | Tb927.10.9100_for | GGGGACAAGTTTGTACAAAAAAGCAGGCTATGGCAAAAACACTTAGCGATGC | Tb927.10.9100_rev | GGGGACCACTTTGTACAAGAAAGCTGGGTTCATTCCAATCTGCCTTTCCTCG |
| 9_hypothetical | J13 | Tb927.10.1960_for | GGGGACAAGTTTGTACAAAAAAGCAGGCTATGTCGAACGGGTATAAGCGC | Tb927.10.1960_rev | GGGGACCACTTTGTACAAGAAAGCTGGGTTCAAAATGCTACAGAAGACATACGTTGC |
| 9_hypothetical | J14 | Tb927.9.2070_for | GGGGACAAGTTTGTACAAAAAAGCAGGCTATGAGTCGACAGACGAGGCG | Tb927.9.2070_rev | GGGGACCACTTTGTACAAGAAAGCTGGGTTCATGAGGGTTTGGGCATTCG |
| 9_hypothetical | J15 | Tb927.8.6860_for | GGGGACAAGTTTGTACAAAAAAGCAGGCTATGGGTACATCTGGTTCTACGATAGATG | Tb927.8.6860_rev | GGGGACCACTTTGTACAAGAAAGCTGGGTCTATCGAAACCGGAGCATTTTGATGG |
| 9_hypothetical | J16 | Tb927.2.2400_for | GGGGACAAGTTTGTACAAAAAAGCAGGCTATGCATTATGCCTCGCGTGG | Tb927.2.2400_rev | GGGGACCACTTTGTACAAGAAAGCTGGGTTTAAATCTTACCTTTGCCACACCATCG |
| 9_hypothetical | J17 | Tb927.10.2930_for | GGGGACAAGTTTGTACAAAAAAGCAGGCTATGGGGAATTGCTTGTGCTGC | Tb927.10.2930_rev | GGGGACCACTTTGTACAAGAAAGCTGGGTTTACATCAAACTTTCGTCACCATCTTCT |
| 9_hypothetical | J18 | Tb927.8.880_for | GGGGACAAGTTTGTACAAAAAAGCAGGCTATGACTATCAACATCTTTGCTGAACTGG | Tb927.8.880_rev | GGGGACCACTTTGTACAAGAAAGCTGGGTTTACATAACATGGCAGCATTTCGTTTT |
| 9_hypothetical | J19 | Tb927.11.5590_for | GGGGACAAGTTTGTACAAAAAAGCAGGCTATGAAAATATTGCACACACGCGC | Tb927.11.5590_rev | GGGGACCACTTTGTACAAGAAAGCTGGGTCTAGCCACGGTAAAAATCAACAACAAC |
| 9_hypothetical | J20 | Tb927.8.4100_for | GGGGACAAGTTTGTACAAAAAAGCAGGCTATGCCGTTGTGGAAACAAACCA | Tb927.8.4100_rev | GGGGACCACTTTGTACAAGAAAGCTGGGTTCACTGCTCGACCCTTTCTTTGT |
| 9_hypothetical | J22 | Tb927.6.2920_for | GGGGACAAGTTTGTACAAAAAAGCAGGCTATGCTGCATGAAATTTACGCTCCA | Tb927.6.2920_rev | GGGGACCACTTTGTACAAGAAAGCTGGGTTTAATCGCTAAACTCCTCAATTACCTCTC |
| 9_hypothetical | J23 | Tb927.11.9620.1_for | GGGGACAAGTTTGTACAAAAAAGCAGGCTATGCTGTCTGTTGTGCGCG | Tb927.11.9620.1_rev | GGGGACCACTTTGTACAAGAAAGCTGGGTTCACCAACCCAAACCCTTAAGAAATC |
| 9_hypothetical | J24 | Tb927.5.2680_for | GGGGACAAGTTTGTACAAAAAAGCAGGCTATGGTCGATGAAAAATTACCGACGT | Tb927.5.2680_rev | GGGGACCACTTTGTACAAGAAAGCTGGGTTTAATTCTCTTCTTCTGCGGCTGG |
| 9_hypothetical | K1 | Tb927.11.1430_for | GGGGACAAGTTTGTACAAAAAAGCAGGCTATGGCTTACCAGCAGTCACGG | Tb927.11.1430_rev | GGGGACCACTTTGTACAAGAAAGCTGGGTTTACTTGGATTCACCTGCTTTGATATCC |
| 9_hypothetical | K2 | Tb927.11.5560_for | GGGGACAAGTTTGTACAAAAAAGCAGGCTATGATGCCAGTATTCGGCCCA | Tb927.11.5560_rev | GGGGACCACTTTGTACAAGAAAGCTGGGTTTATTCAACGAAAGCGCGAATTGC |
| 9_hypothetical | K3 | Tb927.5.2360_for | GGGGACAAGTTTGTACAAAAAAGCAGGCTATGGAGAGAAATACTCCTCGTTACCA | Tb927.5.2360_rev | GGGGACCACTTTGTACAAGAAAGCTGGGTTTAACGCACATATGTCTTCGGAGC |
| 9_hypothetical | K4 | Tb927.3.850_for | GGGGACAAGTTTGTACAAAAAAGCAGGCTATGACGTTGGCGAGCGAAG | Tb927.3.850_rev | GGGGACCACTTTGTACAAGAAAGCTGGGTTCACGCACGTTCCCCTACG |
| 9_hypothetical | K5 | Tb927.4.3530_for | GGGGACAAGTTTGTACAAAAAAGCAGGCTATGAGGAGTCGGTTCTTTACACACG | Tb927.4.3530_rev | GGGGACCACTTTGTACAAGAAAGCTGGGTCTAACTAACCCAGTTGTTGGTGCG |
| 9_hypothetical | K6 | Tb927.11.13700_for | GGGGACAAGTTTGTACAAAAAAGCAGGCTATGGCATCAACCGATGAAGTAGTG | Tb927.11.13700_rev | GGGGACCACTTTGTACAAGAAAGCTGGGTCTACTGGGAAAGGTGTGACAAAGTG |
| 9_hypothetical | K7 | Tb927.7.4050_for | GGGGACAAGTTTGTACAAAAAAGCAGGCTATGATCCACAGCACCGCGA | Tb927.7.4050_rev | GGGGACCACTTTGTACAAGAAAGCTGGGTTCAAGTATTACCCTGGCGAATGGT |
| 9_hypothetical | K8 | Tb927.11.12620_for | GGGGACAAGTTTGTACAAAAAAGCAGGCTATGAGAGAACGAAACGCTTCCG | Tb927.11.12620_rev | GGGGACCACTTTGTACAAGAAAGCTGGGTTCACTCCCCCACTTCCCTTCT |
| 9_hypothetical | K9 | Tb927.11.14450_for | GGGGACAAGTTTGTACAAAAAAGCAGGCTATGACTATTTTGCGGCGTGTGA | Tb927.11.14450_rev | GGGGACCACTTTGTACAAGAAAGCTGGGTTCACATAGCTGAGCTGTTGGATCT |
| 9_hypothetical | K10 | Tb927.6.2650_for | GGGGACAAGTTTGTACAAAAAAGCAGGCTATGGACGTGTTTTCAAGACATGGG | Tb927.6.2650_rev | GGGGACCACTTTGTACAAGAAAGCTGGGTTCACGAAATGCCACCCTTCAAA |
| 9_hypothetical | K11 | Tb927.11.840.1_for | GGGGACAAGTTTGTACAAAAAAGCAGGCTATGGCAAAACATCCCGGTGAG | Tb927.11.840.1_rev | GGGGACCACTTTGTACAAGAAAGCTGGGTTTACTTTGCAGCAGCTGGAACA |
| 9_hypothetical | K12 | Tb927.9.14030_for | GGGGACAAGTTTGTACAAAAAAGCAGGCTATGACATCGCTTCCGTCGC | Tb927.9.14030_rev | GGGGACCACTTTGTACAAGAAAGCTGGGTCTATGGACCGTCAGCAAGGTGT |
| 9_hypothetical | K13 | Tb927.7.6480_for | GGGGACAAGTTTGTACAAAAAAGCAGGCTATGAATCAAAGCCCGTTTGATCACC | Tb927.7.6480_rev | GGGGACCACTTTGTACAAGAAAGCTGGGTCTACCTTGTTTCCTGGAAGTGACAG |
| 9_hypothetical | K14 | Tb927.2.3390_for | GGGGACAAGTTTGTACAAAAAAGCAGGCTATGCACGTGGGTGATGGAAC | Tb927.2.3390_rev | GGGGACCACTTTGTACAAGAAAGCTGGGTTCACATGAGCGCAGACAAGGC |
| 9_hypothetical | K15 | Tb927.10.3880_for | GGGGACAAGTTTGTACAAAAAAGCAGGCTATGTTTTGGGCTATCACGGGTTC | Tb927.10.3880_rev | GGGGACCACTTTGTACAAGAAAGCTGGGTCTACGCCAAATCATCACACAATAGAAAG |
| 9_hypothetical | K16 | Tb927.6.2710_for | GGGGACAAGTTTGTACAAAAAAGCAGGCTATGTCAGTTACCGTTGAAGGTGC | Tb927.6.2710_rev | GGGGACCACTTTGTACAAGAAAGCTGGGTTCATGCTGAAAGCAACAAACAATTGA |
| 9_hypothetical | K17 | Tb927.7.6090_for | GGGGACAAGTTTGTACAAAAAAGCAGGCTATGTTGCAAGCGGAAGTAAACTCG | Tb927.7.6090_rev | GGGGACCACTTTGTACAAGAAAGCTGGGTCTACTTCCCTTGTAGGCGCTCT |
| 9_hypothetical | K18 | Tb927.2.1700_for | GGGGACAAGTTTGTACAAAAAAGCAGGCTATGCTTGTCGGGGTGCTTG | Tb927.2.1700_rev | GGGGACCACTTTGTACAAGAAAGCTGGGTTCATTCCTTGTTAGGGAACCCTTCA |
| 9_hypothetical | K19 | Tb927.3.5140_for | GGGGACAAGTTTGTACAAAAAAGCAGGCTATGGATTTGAAGGAGAACCGTAGAAGT | Tb927.3.5140_rev | GGGGACCACTTTGTACAAGAAAGCTGGGTTCATTCCTGAGCTGGGCCTG |
| 9_hypothetical | K20 | Tb927.11.8270_for | GGGGACAAGTTTGTACAAAAAAGCAGGCTATGACGTCACCTCACCTCCTTAC | Tb927.11.8270_rev | GGGGACCACTTTGTACAAGAAAGCTGGGTTCAAAACGTTGCTACACCCGC |
| 9_hypothetical | K21 | Tb927.11.9290_for | GGGGACAAGTTTGTACAAAAAAGCAGGCTATGCTACAAAAGGAGGTATTGGGC | Tb927.11.9290_rev | GGGGACCACTTTGTACAAGAAAGCTGGGTTTACGTCCCCTCAGGCTGTTC |
| 9_hypothetical | K22 | Tb927.6.1720_for | GGGGACAAGTTTGTACAAAAAAGCAGGCTATGCAGTACCTTTACGATGATACGGT | Tb927.6.1720_rev | GGGGACCACTTTGTACAAGAAAGCTGGGTTTAATGTGCAGCGTTCATTAGGCG |
| 9_hypothetical | K23 | Tb927.2.5480_for | GGGGACAAGTTTGTACAAAAAAGCAGGCTATGTCCACGGTAAAGGCAGCC | Tb927.2.5480_rev | GGGGACCACTTTGTACAAGAAAGCTGGGTTTAAACAATGAGGTATCCCTTGGGC |
| 9_hypothetical | K24 | Tb927.10.4650_for | GGGGACAAGTTTGTACAAAAAAGCAGGCTATGAGCGCCAGTGTGTCTCC | Tb927.10.4650_rev | GGGGACCACTTTGTACAAGAAAGCTGGGTTCAACCTCTGACCGCGAGC |
| 9_hypothetical | L1 | Tb927.2.5650_for | GGGGACAAGTTTGTACAAAAAAGCAGGCTATGTCCACGGTACAGGCAGC | Tb927.2.5650_rev | GGGGACCACTTTGTACAAGAAAGCTGGGTTTAAACAATGAGGTATCCCTTGGGC |
| 9_hypothetical | L2 | Tb927.10.8820_for | GGGGACAAGTTTGTACAAAAAAGCAGGCTATGAACATGATATCCCTCAATTTCACCC | Tb927.10.8820_rev | GGGGACCACTTTGTACAAGAAAGCTGGGTCTATCGGCCCGGGATTGGT |
| 9_hypothetical | L3 | Tb927.11.2860_for | GGGGACAAGTTTGTACAAAAAAGCAGGCTATGGAACCCGGAACGGAGG | Tb927.11.2860_rev | GGGGACCACTTTGTACAAGAAAGCTGGGTCTACACGAAAGTCCACACCCG |
| 9_hypothetical | L4 | Tb927.11.15520_for | GGGGACAAGTTTGTACAAAAAAGCAGGCTATGCCGTGGATTCACCTTCAGT | Tb927.11.15520_rev | GGGGACCACTTTGTACAAGAAAGCTGGGTTTAGGAAGTGGCATAGCGCCA |
| 9_hypothetical | L5 | Tb927.11.14760_for | GGGGACAAGTTTGTACAAAAAAGCAGGCTATGGTGCGACTACCACCAATAGT | Tb927.11.14760_rev | GGGGACCACTTTGTACAAGAAAGCTGGGTTTAGATTGGTTGCTCAGTAAGGTAGAATG |
| 9_hypothetical | L6 | Tb927.6.1430_for | GGGGACAAGTTTGTACAAAAAAGCAGGCTATGGATCCCCAGTATTCGCAACC | Tb927.6.1430_rev | GGGGACCACTTTGTACAAGAAAGCTGGGTTCACGACAATCTCTCGAGGATGT |
| 9_hypothetical | L7 | Tb927.11.9120_for | GGGGACAAGTTTGTACAAAAAAGCAGGCTATGCTGCCTTGCCCAAGC | Tb927.11.9120_rev | GGGGACCACTTTGTACAAGAAAGCTGGGTTCACTGATTATCACCATCCTCTTCGT |
| 9_hypothetical | L8 | Tb927.11.5330_for | GGGGACAAGTTTGTACAAAAAAGCAGGCTATGAGTTGGCTTCAGGCTCTCG | Tb927.11.5330_rev | GGGGACCACTTTGTACAAGAAAGCTGGGTTCATGAAACTTCACAATTCCCGGTT |
| 9_hypothetical | L9 | Tb927.10.11040_for | GGGGACAAGTTTGTACAAAAAAGCAGGCTATGGTAGAGGGTCACAGCGAC | Tb927.10.11040_rev | GGGGACCACTTTGTACAAGAAAGCTGGGTTCACTTTGCATCTGCCGGC |
| 9_hypothetical | L10 | Tb927.7.450_for | GGGGACAAGTTTGTACAAAAAAGCAGGCTATGCAGGACGTACCAAATCCAAAC | Tb927.7.450_rev | GGGGACCACTTTGTACAAGAAAGCTGGGTTTATTCTTTCGACTGTCCCTGTTCAG |
| 9_hypothetical | L11 | Tb927.9.11500_for | GGGGACAAGTTTGTACAAAAAAGCAGGCTATGGAAGAAGATGAGGCGTTTGC | Tb927.9.11500_rev | GGGGACCACTTTGTACAAGAAAGCTGGGTTCAACCCTTGGGCAACGTATACA |
| 9_hypothetical | L12 | Tb927.2.2530_for | GGGGACAAGTTTGTACAAAAAAGCAGGCTATGTCAGGGCAACCACACTACG | Tb927.2.2530_rev | GGGGACCACTTTGTACAAGAAAGCTGGGTTCACATAGGGTCCGAAACACCG |
| 9_hypothetical | L13 | Tb927.11.10390_for | GGGGACAAGTTTGTACAAAAAAGCAGGCTATGAGTGGGAAGGGCTTTGTATCA | Tb927.11.10390_rev | GGGGACCACTTTGTACAAGAAAGCTGGGTCTAGTAAAACAACTTCTCACGCCGA |
| 9_hypothetical | L14 | Tb927.7.5330_for | GGGGACAAGTTTGTACAAAAAAGCAGGCTATGCCCATTTCGCCCTTGG | Tb927.7.5330_rev | GGGGACCACTTTGTACAAGAAAGCTGGGTTCAGCGAAAGCGTACCACCA |
| 9_hypothetical | L15 | Tb927.9.7820_for | GGGGACAAGTTTGTACAAAAAAGCAGGCTATGAAGTTGTGTGAACTTAGTCCAATAGC | Tb927.9.7820_rev | GGGGACCACTTTGTACAAGAAAGCTGGGTTCACTCGCTCTTCTCCCTATCTCG |
| 9_hypothetical | L16 | Tb927.3.1860_for | GGGGACAAGTTTGTACAAAAAAGCAGGCTATGTACAGAATAGCAATTCGACGTTTTG | Tb927.3.1860_rev | GGGGACCACTTTGTACAAGAAAGCTGGGTTTATTTTTTACCCCCGTGCCCTTT |
| 9_hypothetical | L17 | Tb927.11.9950_for | GGGGACAAGTTTGTACAAAAAAGCAGGCTATGGACCCCCTTTTACTTTCCAGG | Tb927.11.9950_rev | GGGGACCACTTTGTACAAGAAAGCTGGGTCTACAAGTCATGTCGGCAAAGATAAAC |
| 9_hypothetical | L18 | Tb927.7.560_for | GGGGACAAGTTTGTACAAAAAAGCAGGCTATGTTTGTACTGCACCGGGGC | Tb927.7.560_rev | GGGGACCACTTTGTACAAGAAAGCTGGGTCTAACGTAAAGTCGATCCCACTGC |
| 9_hypothetical | L19 | Tb927.3.1200_for | GGGGACAAGTTTGTACAAAAAAGCAGGCTATGAGTTTGCATGTCGGTAACGG | Tb927.3.1200_rev | GGGGACCACTTTGTACAAGAAAGCTGGGTCTACATTCCGGCAAGGCGAAG |
| 9_hypothetical | L20 | Tb927.10.3020_for | GGGGACAAGTTTGTACAAAAAAGCAGGCTATGTCGGGTCGGCGTGTC | Tb927.10.3020_rev | GGGGACCACTTTGTACAAGAAAGCTGGGTCTACTCACTGACCCCCTCACAC |
| 9_hypothetical | L21 | Tb927.3.4350_for | GGGGACAAGTTTGTACAAAAAAGCAGGCTATGCTTGTTACAACAACAGCGATATCT | Tb927.3.4350_rev | GGGGACCACTTTGTACAAGAAAGCTGGGTTCACGAAGAATCCTTCGGGCG |
| 9_hypothetical | L22 | Tb927.11.16640_for | GGGGACAAGTTTGTACAAAAAAGCAGGCTATGCAGGCGGAGGTGGAC | Tb927.11.16640_rev | GGGGACCACTTTGTACAAGAAAGCTGGGTTCATACGGACACTGCCGACT |
| 9_hypothetical | L23 | Tb927.10.9660_for | GGGGACAAGTTTGTACAAAAAAGCAGGCTATGAAGCGGAACCGTAACCAC | Tb927.10.9660_rev | GGGGACCACTTTGTACAAGAAAGCTGGGTTCATGCCTTTTTCTTCACAAACAACC |
| 9_hypothetical | L24 | Tb927.6.860_for | GGGGACAAGTTTGTACAAAAAAGCAGGCTATGCACATGTACAACCCTGTGG | Tb927.6.860_rev | GGGGACCACTTTGTACAAGAAAGCTGGGTTCATGAACCGCCATTCTGTTGG |
| 9_hypothetical | M1 | Tb927.8.850_for | GGGGACAAGTTTGTACAAAAAAGCAGGCTATGTTTGGGTCTTCGCTAAGCG | Tb927.8.850_rev | GGGGACCACTTTGTACAAGAAAGCTGGGTCTACACGCTCACAGCGCTG |
| 9_hypothetical | M2 | Tb927.1.4410_for | GGGGACAAGTTTGTACAAAAAAGCAGGCTATGCAAAAAACCCCTCGATTAACGA | Tb927.1.4410_rev | GGGGACCACTTTGTACAAGAAAGCTGGGTTCACATGATGCCGTAGTCGTCC |
| 9_hypothetical | M3 | Tb927.5.2950_for | GGGGACAAGTTTGTACAAAAAAGCAGGCTATGAATTCGACGAGTACATCAAGTTACG | Tb927.5.2950_rev | GGGGACCACTTTGTACAAGAAAGCTGGGTCTACGACAGTTGCTGAAGTGTGC |

| | | | | | |
|---|---|---|---|---|---|
| 9_hypothetical | M4 | Tb927.10.15660_for | GGGGACAAGTTTGTACAAAAAAGCAGGCTATGATTCGTTACACTCAATACCTTATCACG | Tb927.10.15660_rev | GGGGACCACTTTGTACAAGAAAGCTGGGTTCAGTTATGCTGTGTGTCGGTTG |
| 9_hypothetical | M5 | Tb927.10.2860_for | GGGGACAAGTTTGTACAAAAAAGCAGGCTATGTCCACTCCGAAGTGCACG | Tb927.10.2860_rev | GGGGACCACTTTGTACAAGAAAGCTGGGTTCAAGCCTTCCGAAGTGCCT |
| 9_hypothetical | M6 | Tb927.11.1220_for | GGGGACAAGTTTGTACAAAAAAGCAGGCTATGTCGGATTTCGTCGATATTAATGTGA | Tb927.11.1220_rev | GGGGACCACTTTGTACAAGAAAGCTGGGTTTACCGTTGCCTACTTCTGTATTCG |
| 9_hypothetical | M7 | Tb927.10.15780_for | GGGGACAAGTTTGTACAAAAAAGCAGGCTATGTCTCTGGGATGTGATCAACACA | Tb927.10.15780_rev | GGGGACCACTTTGTACAAGAAAGCTGGGTTCAACAACCATCAGGCGTTAGC |
| 9_hypothetical | M8 | Tb927.10.11720_for | GGGGACAAGTTTGTACAAAAAAGCAGGCTATGCTAACTCGAAGAGCCGCA | Tb927.10.11720_rev | GGGGACCACTTTGTACAAGAAAGCTGGGTCTATGTCCTCCTCCTTTTAAGCTCCT |
| 9_hypothetical | M9 | Tb927.9.13470_for | GGGGACAAGTTTGTACAAAAAAGCAGGCTATGGATGTGCATTCCCCGTTAC | Tb927.9.13470_rev | GGGGACCACTTTGTACAAGAAAGCTGGGTTTAGCCTATGCGAGCCAAAAATGG |
| 9_hypothetical | M10 | Tb927.3.2630_for | GGGGACAAGTTTGTACAAAAAAGCAGGCTATGCGGGTGTTTAGCTCCCC | Tb927.3.2630_rev | GGGGACCACTTTGTACAAGAAAGCTGGGTTCACAGTGCAGCTGCCTGT |
| 9_hypothetical | M11 | Tb927.11.16470_for | GGGGACAAGTTTGTACAAAAAAGCAGGCTATGACCGAAACTGATGGGGGG | Tb927.11.16470_rev | GGGGACCACTTTGTACAAGAAAGCTGGGTTCAAGCCAATTCACGTTTGTGACC |
| 9_hypothetical | M12 | Tb927.10.12130_for | GGGGACAAGTTTGTACAAAAAAGCAGGCTATGTCTGCGACATCCCCGA | Tb927.10.12130_rev | GGGGACCACTTTGTACAAGAAAGCTGGGTTTACTCTTGCATTACCTCGACGACA |
| 9_hypothetical | M13 | Tb927.11.12560_for | GGGGACAAGTTTGTACAAAAAAGCAGGCTATGGACGTGGAGGAATCAGAGC | Tb927.11.12560_rev | GGGGACCACTTTGTACAAGAAAGCTGGGTCTAGCAACCGTTGTCGGCC |
| 9_hypothetical | M14 | Tb927.10.2260_for | GGGGACAAGTTTGTACAAAAAAGCAGGCTATGAGTTCACGGGAGCCAACA | Tb927.10.2260_rev | GGGGACCACTTTGTACAAGAAAGCTGGGTCTAAATTTGATCGTCGCCGTCATG |
| 9_hypothetical | M15 | Tb927.10.3130_for | GGGGACAAGTTTGTACAAAAAAGCAGGCTATGAATGTTCTCATTGGAAAACGTGAGC | Tb927.10.3130_rev | GGGGACCACTTTGTACAAGAAAGCTGGGTTTACCGTCCTATCACTCCAGGC |
| 9_hypothetical | M16 | Tb927.3.3150_for | GGGGACAAGTTTGTACAAAAAAGCAGGCTATGTCAGCATTTGAAGATAATCCGTTTG | Tb927.3.3150_rev | GGGGACCACTTTGTACAAGAAAGCTGGGTCTACCAACTGCCAAACGGACC |
| 9_hypothetical | M17 | Tb927.10.3000_for | GGGGACAAGTTTGTACAAAAAAGCAGGCTATGAACAGTTTCGTGTCTCATTCCTC | Tb927.10.3000_rev | GGGGACCACTTTGTACAAGAAAGCTGGGTTCATATGGAGTCATCACTGTCATCTCC |
| 9_hypothetical | M18 | Tb927.11.12660_for | GGGGACAAGTTTGTACAAAAAAGCAGGCTATGAAAACCGAGGTGGAGACCA | Tb927.11.12660_rev | GGGGACCACTTTGTACAAGAAAGCTGGGTTCAACAGCAAGTGAGGGATGGC |
| 9_hypothetical | M19 | Tb927.3.1350_for | GGGGACAAGTTTGTACAAAAAAGCAGGCTATGCAACGACGTAAGCGCT | Tb927.3.1350_rev | GGGGACCACTTTGTACAAGAAAGCTGGGTTCAGTTACCGAAGCGTCGCA |
| 9_hypothetical | M20 | Tb927.11.4950_for | GGGGACAAGTTTGTACAAAAAAGCAGGCTATGGAATACAAATCTGTTTTTGAACTTCGA | Tb927.11.4950_rev | GGGGACCACTTTGTACAAGAAAGCTGGGTCTAATGCCGTTTCTGCCGGG |
| 9_hypothetical | M21 | Tb927.7.7270_for | GGGGACAAGTTTGTACAAAAAAGCAGGCTATGCATGTCGGTGCGGAAC | Tb927.7.7270_rev | GGGGACCACTTTGTACAAGAAAGCTGGGTTTAAGACTTCAGGAAGCTTCGTTGG |
| 9_hypothetical | M22 | Tb927.11.2120_for | GGGGACAAGTTTGTACAAAAAAGCAGGCTATGACTCGAGAAGAGGAGGTATTGG | Tb927.11.2120_rev | GGGGACCACTTTGTACAAGAAAGCTGGGTTTACCGCCTTCTGCTCACAAAG |
| 9_hypothetical | M23 | Tb927.3.1660_for | GGGGACAAGTTTGTACAAAAAAGCAGGCTATGACAATGTCGCCTTCTACTCGA | Tb927.3.1660_rev | GGGGACCACTTTGTACAAGAAAGCTGGGTCTAAGACTTCCCTACAACACATTGGAG |
| 9_hypothetical | M24 | Tb927.1.4250_for | GGGGACAAGTTTGTACAAAAAAGCAGGCTATGGTAGGAGGATATTTACGTATGCCT | Tb927.1.4250_rev | GGGGACCACTTTGTACAAGAAAGCTGGGTTTATCGGCTTAGTTTCCGCACATAAG |
| 9_hypothetical | N1 | Tb927.8.3710_for | GGGGACAAGTTTGTACAAAAAAGCAGGCTATGGGACGCCGAAAGTGTTC | Tb927.8.3710_rev | GGGGACCACTTTGTACAAGAAAGCTGGGTTTACTGTTGCTCGCATGCCG |
| 9_hypothetical | N2 | Tb927.4.2780_for | GGGGACAAGTTTGTACAAAAAAGCAGGCTATGACACATTGTGACCCATCGAAC | Tb927.4.2780_rev | GGGGACCACTTTGTACAAGAAAGCTGGGTTCAACGCCCATGAAAGGGGT |
| 9_hypothetical | N3 | Tb927.1.3450_for | GGGGACAAGTTTGTACAAAAAAGCAGGCTATGGCCTCCTTCCTGCCATC | Tb927.1.3450_rev | GGGGACCACTTTGTACAAGAAAGCTGGGTCTACCACAGCCGCACTCTTG |
| 9_hypothetical | N4 | Tb927.10.5920_for | GGGGACAAGTTTGTACAAAAAAGCAGGCTATGGGTGTGCCAGACTTTCTAAAG | Tb927.10.5920_rev | GGGGACCACTTTGTACAAGAAAGCTGGGTTCAGGAAAGTTTTTTCCGTTTTACTTTTTT |
| 9_hypothetical | N5 | Tb927.10.12140_for | GGGGACAAGTTTGTACAAAAAAGCAGGCTATGCCCGGGAGTTTGAATTCC | Tb927.10.12140_rev | GGGGACCACTTTGTACAAGAAAGCTGGGTTCAGTACTTTCCTTGTGAACTGATGTAC |
| 9_hypothetical | N6 | Tb927.10.12250_for | GGGGACAAGTTTGTACAAAAAAGCAGGCTATGGGGTCACCAACCCCAA | Tb927.10.12250_rev | GGGGACCACTTTGTACAAGAAAGCTGGGTTTATTTACCGGCATTTGTTGCACCC |
| 9_hypothetical | N7 | Tb927.11.15800_for | GGGGACAAGTTTGTACAAAAAAGCAGGCTATGTCAAAGCCAACTCGGAAGTC | Tb927.11.15800_rev | GGGGACCACTTTGTACAAGAAAGCTGGGTCTACGATTTGGAAGAAACTGTCGTCC |
| 9_hypothetical | N8 | Tb927.6.4760_for | GGGGACAAGTTTGTACAAAAAAGCAGGCTATGTCGGAGCCGACTACACC | Tb927.6.4760_rev | GGGGACCACTTTGTACAAGAAAGCTGGGTCTAAGCGGACGCCTCCTGC |
| 9_hypothetical | N9 | Tb927.2.4470_for | GGGGACAAGTTTGTACAAAAAAGCAGGCTATGTCACATTCCAGTAGAGATGATGGG | Tb927.2.4470_rev | GGGGACCACTTTGTACAAGAAAGCTGGGTTCATTTCACTGCTTCCCGTAACG |
| 9_hypothetical | N10 | Tb927.8.3170_for | GGGGACAAGTTTGTACAAAAAAGCAGGCTATGCAAGATAAGGCGGTGCAA | Tb927.8.3170_rev | GGGGACCACTTTGTACAAGAAAGCTGGGTTTACAGAGTTTTGCAATACTTAGCCTTAAC |
| 9_hypothetical | N11 | Tb927.11.810_for | GGGGACAAGTTTGTACAAAAAAGCAGGCTATGGAGCCGATGGAGCAACC | Tb927.11.810_rev | GGGGACCACTTTGTACAAGAAAGCTGGGTCTACTGCTCCCAATTAAAGCGGC |
| 9_hypothetical | N12 | Tb927.6.4100_for | GGGGACAAGTTTGTACAAAAAAGCAGGCTATGTCGTCACTAAAAGGGGCGC | Tb927.6.4100_rev | GGGGACCACTTTGTACAAGAAAGCTGGGTTCAAGTGTGAGAAATGCGTTGCA |
| 9_hypothetical | N13 | Tb927.5.3240_for | GGGGACAAGTTTGTACAAAAAAGCAGGCTATGGCGCAACACTCAGAAGC | Tb927.5.3240_rev | GGGGACCACTTTGTACAAGAAAGCTGGGTTTACTGGTCGTACACCGGTGAG |
| 9_hypothetical | N14 | Tb927.11.3490_for | GGGGACAAGTTTGTACAAAAAAGCAGGCTATGGACACCCTTACGTACGTGT | Tb927.11.3490_rev | GGGGACCACTTTGTACAAGAAAGCTGGGTTCATGTCACCAGGAAGTGAATGAGT |
| 9_hypothetical | N15 | Tb927.10.5910_for | GGGGACAAGTTTGTACAAAAAAGCAGGCTATGGGGATTCAAACGATGTCAGAAG | Tb927.10.5910_rev | GGGGACCACTTTGTACAAGAAAGCTGGGTCTATACAACCAACGAGTCCATGTACC |
| 9_hypothetical | N16 | Tb927.11.6260_for | GGGGACAAGTTTGTACAAAAAAGCAGGCTATGGCGGATGGAGCTGAAGG | Tb927.11.6260_rev | GGGGACCACTTTGTACAAGAAAGCTGGGTCTATGCGAGGTTAGTTGGAGCG |
| 9_hypothetical | N17 | Tb927.6.4550_for | GGGGACAAGTTTGTACAAAAAAGCAGGCTATGGCAGCCCAACACACATG | Tb927.6.4550_rev | GGGGACCACTTTGTACAAGAAAGCTGGGTCTACAGGGACTTCCCACAAGTGA |
| 9_hypothetical | N18 | Tb927.1.790_for | GGGGACAAGTTTGTACAAAAAAGCAGGCTATGCGGCTTCATTCGCGT | Tb927.1.790_rev | GGGGACCACTTTGTACAAGAAAGCTGGGTTCACGCCCTTCGACGGAA |
| 9_hypothetical | N19 | Tb927.11.10120_for | GGGGACAAGTTTGTACAAAAAAGCAGGCTATGTCCCTCAACCGCAAGTATAGC | Tb927.11.10120_rev | GGGGACCACTTTGTACAAGAAAGCTGGGTTTATCCCTGTAGTGACTTTTTTCTAGCAAT |
| 9_hypothetical | N20 | Tb927.2.1760_for | GGGGACAAGTTTGTACAAAAAAGCAGGCTATGAGTATTCTGTGGCTTACTTGTATCG | Tb927.2.1760_rev | GGGGACCACTTTGTACAAGAAAGCTGGGTTCACTCACGGTCAGATGAGTGAG |
| 9_hypothetical | N21 | Tb927.10.16090_for | GGGGACAAGTTTGTACAAAAAAGCAGGCTATGCGTAGATCGGGCAGAGG | Tb927.10.16090_rev | GGGGACCACTTTGTACAAGAAAGCTGGGTCTACTTGGCCGCGTGCCC |
| 9_hypothetical | N22 | Tb927.8.1250_for | GGGGACAAGTTTGTACAAAAAAGCAGGCTATGCGACGCCCATTGCGT | Tb927.8.1250_rev | GGGGACCACTTTGTACAAGAAAGCTGGGTTCATGCACCGTGATGCCTTAGT |
| 9_hypothetical | N23 | Tb927.5.3650_for | GGGGACAAGTTTGTACAAAAAAGCAGGCTATGCCGAGCGACCACCAG | Tb927.5.3650_rev | GGGGACCACTTTGTACAAGAAAGCTGGGTTCACAACAAATTCCACGTTCGCT |
| 9_hypothetical | N24 | Tb927.11.12530_for | GGGGACAAGTTTGTACAAAAAAGCAGGCTATGAGCACATCTTTAGTAACGCTTGC | Tb927.11.12530_rev | GGGGACCACTTTGTACAAGAAAGCTGGGTTTATTGCCGGCGTTGCGC |
| 9_hypothetical | O1 | Tb927.7.2050_for | GGGGACAAGTTTGTACAAAAAAGCAGGCTATGAACTGTTCAACTGCAGGCG | Tb927.7.2050_rev | GGGGACCACTTTGTACAAGAAAGCTGGGTCTAACGAGTTTCCCTCTCCCTCC |
| 9_hypothetical | O2 | Tb927.9.11150_for | GGGGACAAGTTTGTACAAAAAAGCAGGCTATGGGTGTAGATAAAAAAATAAGTTCTCTA | Tb927.9.11150_rev | GGGGACCACTTTGTACAAGAAAGCTGGGTTCATATGGTAAAAAATAGAAACACCATTAA |
| 9_hypothetical | O3 | Tb927.7.3270_for | GGGGACAAGTTTGTACAAAAAAGCAGGCTATGCATCGTCGCCTTCGAATT | Tb927.7.3270_rev | GGGGACCACTTTGTACAAGAAAGCTGGGTTCAACTCTCAACAAGAAAAACCATTTTTGC |
| 9_hypothetical | O4 | Tb927.10.3470_for | GGGGACAAGTTTGTACAAAAAAGCAGGCTATGTTGTCACCGATTGTGGACATG | Tb927.10.3470_rev | GGGGACCACTTTGTACAAGAAAGCTGGGTTCAAGCAAGAGGACCTTCCAACA |
| 9_hypothetical | O5 | Tb927.9.2817_for | GGGGACAAGTTTGTACAAAAAAGCAGGCTATGGAAATCCGCATGGTGCC | Tb927.9.2817_rev | GGGGACCACTTTGTACAAGAAAGCTGGGTTCACTCATCAGCTGCTGGAGT |
| 9_hypothetical | O6 | Tb927.10.2590_for | GGGGACAAGTTTGTACAAAAAAGCAGGCTATGCGGATCGCAACGAGTC | Tb927.10.2590_rev | GGGGACCACTTTGTACAAGAAAGCTGGGTTCAGTCAAAGTCCTTCAATTCAAATCGT |
| 9_hypothetical | O7 | Tb927.6.3450_for | GGGGACAAGTTTGTACAAAAAAGCAGGCTATGTATAAGGAAACGGGAGCTACGC | Tb927.6.3450_rev | GGGGACCACTTTGTACAAGAAAGCTGGGTTTACCGTTGCCCCATCAACTG |
| 9_hypothetical | O8 | Tb927.3.3300_for | GGGGACAAGTTTGTACAAAAAAGCAGGCTATGGCCACCCTCAACAACTTAAC | Tb927.3.3300_rev | GGGGACCACTTTGTACAAGAAAGCTGGGTCTAAGAACGCTGACGGTGCG |
| 9_hypothetical | O9 | Tb927.10.1530_for | GGGGACAAGTTTGTACAAAAAAGCAGGCTATGAGTGGCGTTTCTGGTGTTG | Tb927.10.1530_rev | GGGGACCACTTTGTACAAGAAAGCTGGGTCTACAAACACGGTTGAGCTTTACTAATG |
| 9_hypothetical | O10 | Tb927.9.1440_for | GGGGACAAGTTTGTACAAAAAAGCAGGCTATGTCAGCTGCATGTCCGTTG | Tb927.9.1440_rev | GGGGACCACTTTGTACAAGAAAGCTGGGTTCACGCCTTTACAGAACCATGAAA |
| 9_hypothetical | O11 | Tb927.5.4040_for | GGGGACAAGTTTGTACAAAAAAGCAGGCTATGGTTAGACGTAGTCATGTGGTTGC | Tb927.5.4040_rev | GGGGACCACTTTGTACAAGAAAGCTGGGTTTATGGTGCGTTCCACTCAAGC |
| 9_hypothetical | O12 | Tb927.5.1320_for | GGGGACAAGTTTGTACAAAAAAGCAGGCTATGCCGAATTCGGCACCG | Tb927.5.1320_rev | GGGGACCACTTTGTACAAGAAAGCTGGGTCTACGTTGCGTATAGATACGGATGTATC |
| 9_hypothetical | O13 | Tb927.4.590_for | GGGGACAAGTTTGTACAAAAAAGCAGGCTATGCGGCCCACTTTATACTTTCTG | Tb927.4.590_rev | GGGGACCACTTTGTACAAGAAAGCTGGGTTTATCGCCACAGTTGTCTCACAC |
| 9_hypothetical | O14 | Tb927.5.4570_for | GGGGACAAGTTTGTACAAAAAAGCAGGCTATGTGTTTTATATTTGGTGTTGAGATGAGT | Tb927.5.4570_rev | GGGGACCACTTTGTACAAGAAAGCTGGGTTTACGGTGCGGTGGTTCCC |
| 9_hypothetical | O16 | Tb927.6.5090_for | GGGGACAAGTTTGTACAAAAAAGCAGGCTATGGACCATGAGGAAGTTCACGC | Tb927.6.5090_rev | GGGGACCACTTTGTACAAGAAAGCTGGGTTTAGTGCAGCATCCTTCCGTGA |
| 9_hypothetical | O17 | Tb927.11.13340_for | GGGGACAAGTTTGTACAAAAAAGCAGGCTATGCCTCTAATTTTCTTGCAGACGC | Tb927.11.13340_rev | GGGGACCACTTTGTACAAGAAAGCTGGGTCTATTCTTCTTCATTTACAGGCCACTCA |
| 9_hypothetical | O18 | Tb927.7.6950_for | GGGGACAAGTTTGTACAAAAAAGCAGGCTATGCAGCCAACGGGGCGC | Tb927.7.6950_rev | GGGGACCACTTTGTACAAGAAAGCTGGGTCTATCGCGTGTGCCCAATGA |
| 9_hypothetical | O19 | Tb927.5.1340_for | GGGGACAAGTTTGTACAAAAAAGCAGGCTATGGGAACAACATCCACAGATACCC | Tb927.5.1340_rev | GGGGACCACTTTGTACAAGAAAGCTGGGTTCACGTAAGTGCTGGCACCG |
| 9_hypothetical | O20 | Tb927.11.450_for | GGGGACAAGTTTGTACAAAAAAGCAGGCTATGGACGCATATCAATCTTTAAGCGG | Tb927.11.450_rev | GGGGACCACTTTGTACAAGAAAGCTGGGTCTATTTACTGGTGACGGGATAAGGC |
| 9_hypothetical | O21 | Tb927.9.10140_for | GGGGACAAGTTTGTACAAAAAAGCAGGCTATGCAGAAAGCTAAGAACCAAAGCT | Tb927.9.10140_rev | GGGGACCACTTTGTACAAGAAAGCTGGGTCTAATGGGGTGGCTCCGCG |
| 9_hypothetical | O22 | Tb927.8.8160_for | GGGGACAAGTTTGTACAAAAAAGCAGGCTATGAGCGATTCGGAGTCCATTCA | Tb927.8.8160_rev | GGGGACCACTTTGTACAAGAAAGCTGGGTCTATACGGGGTTTTTGAGCAGGG |
| 9_hypothetical | O23 | Tb927.10.10100_for | GGGGACAAGTTTGTACAAAAAAGCAGGCTATGAATAGCAGTGGTCTTGTTTTGGA | Tb927.10.10100_rev | GGGGACCACTTTGTACAAGAAAGCTGGGTCTATTTTTCGGAATTCATTTTCTCGTTACC |
| 9_hypothetical | O24 | Tb927.8.7780_for | GGGGACAAGTTTGTACAAAAAAGCAGGCTATGGGAGCACTGCCGAGC | Tb927.8.7780_rev | GGGGACCACTTTGTACAAGAAAGCTGGGTCTATTCCCTGTAAAATATATTTGACTGCCC |
| 9_hypothetical | P1 | Tb927.11.14440_for | GGGGACAAGTTTGTACAAAAAAGCAGGCTATGATGAGTGCAAAACTTGGGATCA | Tb927.11.14440_rev | GGGGACCACTTTGTACAAGAAAGCTGGGTCTAAAGAAGACCCTTTAGATCCGTTGTT |
| 9_hypothetical | P2 | Tb927.11.14970_for | GGGGACAAGTTTGTACAAAAAAGCAGGCTATGCTTAAGTTGCCTACAATTGCTCA | Tb927.11.14970_rev | GGGGACCACTTTGTACAAGAAAGCTGGGTTTACTCCATCTCCCTTCGCTTAAGA |
| 9_hypothetical | P3 | Tb927.4.4550_for | GGGGACAAGTTTGTACAAAAAAGCAGGCTATGGGAGCACTTCCGAGCA | Tb927.4.4550_rev | GGGGACCACTTTGTACAAGAAAGCTGGGTTTACTCTAATATTAAAGACGGCGCGC |
| 9_hypothetical | P4 | Tb927.7.6910_for | GGGGACAAGTTTGTACAAAAAAGCAGGCTATGTCGGGAAGTGACGCAGA | Tb927.7.6910_rev | GGGGACCACTTTGTACAAGAAAGCTGGGTTTATTCCTTGGTCGCCACAGC |
| 9_hypothetical | P5 | Tb927.7.670_for | GGGGACAAGTTTGTACAAAAAAGCAGGCTATGGAAGATTTTGACTTTCTAGACGATGC | Tb927.7.670_rev | GGGGACCACTTTGTACAAGAAAGCTGGGTTCATGCCAAGCCTTTGGAGAAAC |
| 9_hypothetical | P6 | Tb927.11.10550_for | GGGGACAAGTTTGTACAAAAAAGCAGGCTATGATGGATGCGCAAACGAGT | Tb927.11.10550_rev | GGGGACCACTTTGTACAAGAAAGCTGGGTTCACGCCATGAGCCCTTCG |
| 9_hypothetical | P7 | Tb927.10.1560_for | GGGGACAAGTTTGTACAAAAAAGCAGGCTATGGCACTCAGTTTATCGGCG | Tb927.10.1560_rev | GGGGACCACTTTGTACAAGAAAGCTGGGTCTAGTTGGAAGCCCCTCCAGG |
| 9_hypothetical | P8 | Tb927.2.4990_for | GGGGACAAGTTTGTACAAAAAAGCAGGCTATGACACAGGAACATGGTAGTGAGG | Tb927.2.4990_rev | GGGGACCACTTTGTACAAGAAAGCTGGGTTCAAAACGGTCCGGAGGGC |
| 9_hypothetical | P9 | Tb927.6.3030_for | GGGGACAAGTTTGTACAAAAAAGCAGGCTATGAGCTGGAAATATTCGAATCAGGAG | Tb927.6.3030_rev | GGGGACCACTTTGTACAAGAAAGCTGGGTTCAACAGGTTGCTACACTGCG |
| 9_hypothetical | P10 | Tb927.10.2870_for | GGGGACAAGTTTGTACAAAAAAGCAGGCTATGGATTGGGCCCTCTTTAGCC | Tb927.10.2870_rev | GGGGACCACTTTGTACAAGAAAGCTGGGTTCATTCACTTTGGTACTGTAATTTCGCT |
| 9_hypothetical | P11 | Tb927.7.1410_for | GGGGACAAGTTTGTACAAAAAAGCAGGCTATGCTTCGACCGCACTGC | Tb927.7.1410_rev | GGGGACCACTTTGTACAAGAAAGCTGGGTTTAACGGCACTGAAGAACTATATCCATG |
| 9_hypothetical | P12 | Tb927.3.1180_for | GGGGACAAGTTTGTACAAAAAAGCAGGCTATGCCGGGCTACCCCACG | Tb927.3.1180_rev | GGGGACCACTTTGTACAAGAAAGCTGGGTTTAAAGAATCAGAAAGTAATCCTCCTGTGC |
| 9_hypothetical | P13 | Tb927.11.1630_for | GGGGACAAGTTTGTACAAAAAAGCAGGCTATGTTACGACGCTTTGCCCTC | Tb927.11.1630_rev | GGGGACCACTTTGTACAAGAAAGCTGGGTCTACTCTGCCTCAACGGTTGTTT |
| 9_hypothetical | P14 | Tb927.9.8610_for | GGGGACAAGTTTGTACAAAAAAGCAGGCTATGTCAACTCGGCAGATCCGG | Tb927.9.8610_rev | GGGGACCACTTTGTACAAGAAAGCTGGGTTCACTGCGAACCGCGCGT |
| 9_hypothetical | P15 | Tb927.4.4180_for | GGGGACAAGTTTGTACAAAAAAGCAGGCTATGAGCGACACGGAATCTATTCAGT | Tb927.4.4180_rev | GGGGACCACTTTGTACAAGAAAGCTGGGTCTACACGGGGTGTTTCAGTAGAGA |
| 9_hypothetical | P16 | Tb927.8.6980_for | GGGGACAAGTTTGTACAAAAAAGCAGGCTATGGAGGTTGACTACTCCGTGG | Tb927.8.6980_rev | GGGGACCACTTTGTACAAGAAAGCTGGGTCTAGCAACATTTCTTACTGTTAACCTTGG |
| 9_hypothetical | P17 | Tb927.11.5800_for | GGGGACAAGTTTGTACAAAAAAGCAGGCTATGCCTTTGTTTTCGTATGAGTGTCA | Tb927.11.5800_rev | GGGGACCACTTTGTACAAGAAAGCTGGGTTTATGACACCCCCAACTCTTCGC |
| 9_hypothetical | P18 | Tb927.5.3270_for | GGGGACAAGTTTGTACAAAAAAGCAGGCTATGGGCGACGGCAGCCAT | Tb927.5.3270_rev | GGGGACCACTTTGTACAAGAAAGCTGGGTTCAGCTGCTAGAGCTTTCATCCC |
| 9_hypothetical | P19 | Tb927.10.3090_for | GGGGACAAGTTTGTACAAAAAAGCAGGCTATGAGGTCGTTGGAGGTGCA | Tb927.10.3090_rev | GGGGACCACTTTGTACAAGAAAGCTGGGTTCACTCTTTCCTTGACTCTGCTAATTG |
| 9_hypothetical | P20 | Tb927.8.2620_for | GGGGACAAGTTTGTACAAAAAAGCAGGCTATGGCTCACCCGGTGCCA | Tb927.8.2620_rev | GGGGACCACTTTGTACAAGAAAGCTGGGTCTACTTCCTCTTCTTCGGCACAATAG |
| 9_hypothetical | P21 | Tb927.2.4160_for | GGGGACAAGTTTGTACAAAAAAGCAGGCTATGAGCTCCACACCGAATAATGAC | Tb927.2.4160_rev | GGGGACCACTTTGTACAAGAAAGCTGGGTCTACGATGCAGAGGGACTTGTACG |
| 9_hypothetical | P22 | Tb927.4.5340_for | GGGGACAAGTTTGTACAAAAAAGCAGGCTATGGAGGTCGAAGATTTCATGGC | Tb927.4.5340_rev | GGGGACCACTTTGTACAAGAAAGCTGGGTCTATTGGCAACATTTGTTGCTATTGACC |
| 9_hypothetical | P23 | Tb927.9.14260_for | GGGGACAAGTTTGTACAAAAAAGCAGGCTATGCGGGTAACATTGGAGGAGA | Tb927.9.14260_rev | GGGGACCACTTTGTACAAGAAAGCTGGGTCTACTGCACGTCCTTGTTGCA |
| 9_hypothetical | P24 | Tb927.11.830_for | GGGGACAAGTTTGTACAAAAAAGCAGGCTATGGGCCACGCTTTTTGCC | Tb927.11.830_rev | GGGGACCACTTTGTACAAGAAAGCTGGGTTCACACACTCCTGCCTTCACG |
| 10_hypothetical | A1 | Tb927.7.7130_for | GGGGACAAGTTTGTACAAAAAAGCAGGCTATGGAGCTGCTATCCGTTACGC | Tb927.7.7130_rev | GGGGACCACTTTGTACAAGAAAGCTGGGTTCAGCAGCCGGAAGTAATCACC |
| 10_hypothetical | A2 | Tb927.8.6870_for | GGGGACAAGTTTGTACAAAAAAGCAGGCTATGGCAAGCAGCGTACCAG | Tb927.8.6870_rev | GGGGACCACTTTGTACAAGAAAGCTGGGTTCAATACACTTGTTCCTCCAACCTAAATT |
| 10_hypothetical | A3 | Tb927.11.14470_for | GGGGACAAGTTTGTACAAAAAAGCAGGCTATGGGATCCCTATTCAAGCTACGT | Tb927.11.14470_rev | GGGGACCACTTTGTACAAGAAAGCTGGGTTCACTTCAGGAAGGAGGTCAATGC |
| 10_hypothetical | A4 | Tb927.4.3700_for | GGGGACAAGTTTGTACAAAAAAGCAGGCTATGCGTCGATCACTGACTTTATGG | Tb927.4.3700_rev | GGGGACCACTTTGTACAAGAAAGCTGGGTTTAACCTACTTCAAAGGACCGCCA |
| 10_hypothetical | A5 | Tb927.11.2740_for | GGGGACAAGTTTGTACAAAAAAGCAGGCTATGGCTGTTGTGGAGAAGTTCCT | Tb927.11.2740_rev | GGGGACCACTTTGTACAAGAAAGCTGGGTTCACTTTTTCCCACCTACGGAAGG |
| 10_hypothetical | A6 | Tb927.11.16960_for | GGGGACAAGTTTGTACAAAAAAGCAGGCTATGTGGACAGGGAGCATCAGT | Tb927.11.16960_rev | GGGGACCACTTTGTACAAGAAAGCTGGGTTCACTCAAGGGAAAAACTTTCGGTG |
| 10_hypothetical | A7 | Tb927.7.1230_for | GGGGACAAGTTTGTACAAAAAAGCAGGCTATGAGGCCCACACCAGACC | Tb927.7.1230_rev | GGGGACCACTTTGTACAAGAAAGCTGGGTCTATACAAGGCAGATTTCTTCCTTACTACA |
| 10_hypothetical | A8 | Tb927.7.3300_for | GGGGACAAGTTTGTACAAAAAAGCAGGCTATGGAATGCGTATCACAGTCGC | Tb927.7.3300_rev | GGGGACCACTTTGTACAAGAAAGCTGGGTTCAAATTTCAGCGTCTTCGAGCA |
| 10_hypothetical | A9 | Tb927.10.2130_for | GGGGACAAGTTTGTACAAAAAAGCAGGCTATGAAAACAAACAACTTGGAGCGTC | Tb927.10.2130_rev | GGGGACCACTTTGTACAAGAAAGCTGGGTTTAGGCTTCGGCAGATACCGA |
| 10_hypothetical | A10 | Tb927.7.4460_for | GGGGACAAGTTTGTACAAAAAAGCAGGCTATGAACTGCGAACACGTTGCG | Tb927.7.4460_rev | GGGGACCACTTTGTACAAGAAAGCTGGGTCTACACGCCGGCAAAGACC |
| 10_hypothetical | A11 | Tb927.11.5920_for | GGGGACAAGTTTGTACAAAAAAGCAGGCTATGTTGTCACGGAGCGCC | Tb927.11.5920_rev | GGGGACCACTTTGTACAAGAAAGCTGGGTTCATGGCGTTTGTGCTGCT |
| 10_hypothetical | A12 | Tb927.7.6630_for | GGGGACAAGTTTGTACAAAAAAGCAGGCTATGGACAGTGATACCATTTATTGTGATGAA | Tb927.7.6630_rev | GGGGACCACTTTGTACAAGAAAGCTGGGTTCACCACAACTCAGCGAGGC |
| 10_hypothetical | A13 | Tb927.10.11110_for | GGGGACAAGTTTGTACAAAAAAGCAGGCTATGAATGTTGCAGAGTTGCTGAAAAAC | Tb927.10.11110_rev | GGGGACCACTTTGTACAAGAAAGCTGGGTTTAGCTTCTGGCAGTGAGTCTGA |
| 10_hypothetical | A14 | Tb927.5.4470_for | GGGGACAAGTTTGTACAAAAAAGCAGGCTATGTCCGACGAGGCACCA | Tb927.5.4470_rev | GGGGACCACTTTGTACAAGAAAGCTGGGTTCAGTTGCCTCCACTCCCAA |
| 10_hypothetical | A15 | Tb927.11.1400_for | GGGGACAAGTTTGTACAAAAAAGCAGGCTATGCTGAAGCAATGTCTGCTTCA | Tb927.11.1400_rev | GGGGACCACTTTGTACAAGAAAGCTGGGTTTAGTCGTCAACATTCACCATAATCGATAC |
| 10_hypothetical | A16 | Tb927.10.7200_for | GGGGACAAGTTTGTACAAAAAAGCAGGCTATGCCAGAAATTAATTTGGATTGCAAGG | Tb927.10.7200_rev | GGGGACCACTTTGTACAAGAAAGCTGGGTCTACGAAGCCTGCTGCTGTTC |
| 10_hypothetical | A17 | Tb927.10.10190_for | GGGGACAAGTTTGTACAAAAAAGCAGGCTATGGCTGATCCCATCGCTGT | Tb927.10.10190_rev | GGGGACCACTTTGTACAAGAAAGCTGGGTTCAAAGAACTGAGCAAGCATGCT |
| 10_hypothetical | A18 | Tb927.6.3820_for | GGGGACAAGTTTGTACAAAAAAGCAGGCTATGGAATCCTCCGTTTATTCTGGTACA | Tb927.6.3820_rev | GGGGACCACTTTGTACAAGAAAGCTGGGTCTATAGCGTACATCCCTCCGGTT |
| 10_hypothetical | A19 | Tb927.8.2250_for | GGGGACAAGTTTGTACAAAAAAGCAGGCTATGGACGCGCAAGTTGGAG | Tb927.8.2250_rev | GGGGACCACTTTGTACAAGAAAGCTGGGTTCACCGCACAAACTGAAACACA |
| 10_hypothetical | A20 | Tb927.4.1980_for | GGGGACAAGTTTGTACAAAAAAGCAGGCTATGAAGCTATGGCAACACCTGC | Tb927.4.1980_rev | GGGGACCACTTTGTACAAGAAAGCTGGGTTTACTCACTTATGAGCCGATCGCT |
| 10_hypothetical | A21 | Tb927.7.750_for | GGGGACAAGTTTGTACAAAAAAGCAGGCTATGTTTGAGGGGGCTGGCG | Tb927.7.750_rev | GGGGACCACTTTGTACAAGAAAGCTGGGTCTACTCCGAATAGAAAGAACTTGGTTGC |
| 10_hypothetical | A22 | Tb927.7.3740_for | GGGGACAAGTTTGTACAAAAAAGCAGGCTATGTCGTACGTTGTTGGCGATG | Tb927.7.3740_rev | GGGGACCACTTTGTACAAGAAAGCTGGGTCTATGCACTACTCGGTTTAACATCTTTGT |
| 10_hypothetical | A23 | Tb927.11.16800_for | GGGGACAAGTTTGTACAAAAAAGCAGGCTATGAAGGGTAAATATATGCATAGCACTGTG | Tb927.11.16800_rev | GGGGACCACTTTGTACAAGAAAGCTGGGTCTATACGTTAGCCTCCAAAGTGTTCTC |
| 10_hypothetical | A24 | Tb927.10.12520_for | GGGGACAAGTTTGTACAAAAAAGCAGGCTATGCGCCGATATTCGACTAGGA | Tb927.10.12520_rev | GGGGACCACTTTGTACAAGAAAGCTGGGTTTATGTTTTTTTTCTAAAGAATGATTTCTT |
| 10_hypothetical | B1 | Tb927.11.3510_for | GGGGACAAGTTTGTACAAAAAAGCAGGCTATGTTTGGTGGTTTGGATTCGTCG | Tb927.11.3510_rev | GGGGACCACTTTGTACAAGAAAGCTGGGTTCAATCATCTCCGTCCGTTACTCT |
| 10_hypothetical | B2 | Tb927.10.14880_for | GGGGACAAGTTTGTACAAAAAAGCAGGCTATGGCAGAACATGCTTTTGCGT | Tb927.10.14880_rev | GGGGACCACTTTGTACAAGAAAGCTGGGTCTACTCGGTAATGGGCCCTGA |
| 10_hypothetical | B3 | Tb927.7.4040_for | GGGGACAAGTTTGTACAAAAAAGCAGGCTATGATGGAAAAGCAGTGGATAGTTTCTT | Tb927.7.4040_rev | GGGGACCACTTTGTACAAGAAAGCTGGGTTCATATTTTCATTGAAGAAGCGTCACTCG |
| 10_hypothetical | B4 | Tb927.10.13150_for | GGGGACAAGTTTGTACAAAAAAGCAGGCTATGGACGGCTTCACCTCCAA | Tb927.10.13150_rev | GGGGACCACTTTGTACAAGAAAGCTGGGTTCAAAGTTGAGAAGTACCGAATCTGG |
| 10_hypothetical | B5 | Tb927.7.4890_for | GGGGACAAGTTTGTACAAAAAAGCAGGCTATGAGTACTGGCCCTTCAACCA | Tb927.7.4890_rev | GGGGACCACTTTGTACAAGAAAGCTGGGTCTATGCCGTCACTCCGGTTG |
| 10_hypothetical | B6 | Tb927.6.2520_for | GGGGACAAGTTTGTACAAAAAAGCAGGCTATGGCGGAAATGCTGAACGAA | Tb927.6.2520_rev | GGGGACCACTTTGTACAAGAAAGCTGGGTTCACCTCGTCCCACAGTACAAC |
| 10_hypothetical | B7 | Tb927.10.14510_for | GGGGACAAGTTTGTACAAAAAAGCAGGCTATGGCTGCTTTCAGCGGG | Tb927.10.14510_rev | GGGGACCACTTTGTACAAGAAAGCTGGGTTTAGTCCTCCCCCTTGTCACG |
| 10_hypothetical | B8 | Tb927.10.9350_for | GGGGACAAGTTTGTACAAAAAAGCAGGCTATGTTAACCTCCCACCGCATTTC | Tb927.10.9350_rev | GGGGACCACTTTGTACAAGAAAGCTGGGTCTATCCAATTGAGGCAAGTACTTCTGT |
| 10_hypothetical | B9 | Tb927.11.10540_for | GGGGACAAGTTTGTACAAAAAAGCAGGCTATGCGAGTTAAGCGGAAGAACG | Tb927.11.10540_rev | GGGGACCACTTTGTACAAGAAAGCTGGGTTTAACTGATACGAGGCAACACATAGTG |
| 10_hypothetical | B10 | Tb927.7.4340_for | GGGGACAAGTTTGTACAAAAAAGCAGGCTATGGGAAAGCCGGGCGGC | Tb927.7.4340_rev | GGGGACCACTTTGTACAAGAAAGCTGGGTCTAGTTTGTTACGAAGGAATACGCCT |
| 10_hypothetical | B11 | Tb927.3.4060_for | GGGGACAAGTTTGTACAAAAAAGCAGGCTATGTGTGGCATCGGTTGCG | Tb927.3.4060_rev | GGGGACCACTTTGTACAAGAAAGCTGGGTTCAGGGAGAGCTGCAGCAT |
| 10_hypothetical | B12 | Tb927.4.4130_for | GGGGACAAGTTTGTACAAAAAAGCAGGCTATGAGTCTAGAGGTCGTGGATGC | Tb927.4.4130_rev | GGGGACCACTTTGTACAAGAAAGCTGGGTCTACTCAATTACAAGACGGCTGGC |
| 10_hypothetical | B13 | Tb927.8.8200_for | GGGGACAAGTTTGTACAAAAAAGCAGGCTATGAATCTTGAGGTCGTGGATGC | Tb927.8.8200_rev | GGGGACCACTTTGTACAAGAAAGCTGGGTCTACTCAACTACAAAACGACTGGCC |
| 10_hypothetical | B14 | Tb927.11.17020_for | GGGGACAAGTTTGTACAAAAAAGCAGGCTATGAATTTAAACGAAGTCGCTTCGGG | Tb927.11.17020_rev | GGGGACCACTTTGTACAAGAAAGCTGGGTCTATGAAGATGATGTCTTCATGAGATGAAG |

| | | | | | |
|---|---|---|---|---|---|
| 10_hypothetical | B15 | Tb927.6.410_for | GGGGACAAGTTTGTACAAAAAAGCAGGCTATGACACAACCAGATATATTCATGGGTG | Tb927.6.410_rev | GGGGACCACTTTGTACAAGAAAGCTGGGTTCAGCATCTGTAACGAACTACAACATT |
| 10_hypothetical | B16 | Tb927.4.5000_for | GGGGACAAGTTTGTACAAAAAAGCAGGCTATGTCAGCCCAGTTGCGGC | Tb927.4.5000_rev | GGGGACCACTTTGTACAAGAAAGCTGGGTTTACATGATGCAGCACGGTCC |
| 10_hypothetical | B17 | Tb927.11.12320_for | GGGGACAAGTTTGTACAAAAAAGCAGGCTATGATGCGGTTCACTAGGGATGG | Tb927.11.12320_rev | GGGGACCACTTTGTACAAGAAAGCTGGGTTTATATTCGCCCCTCCACTACATGT |
| 10_hypothetical | B18 | Tb927.8.7420_for | GGGGACAAGTTTGTACAAAAAAGCAGGCTATGTCAGCCCAGCTGCAGC | Tb927.8.7420_rev | GGGGACCACTTTGTACAAGAAAGCTGGGTTTACATGATGCAGCACGGTCC |
| 10_hypothetical | B19 | Tb927.3.5510_for | GGGGACAAGTTTGTACAAAAAAGCAGGCTATGCCACCACCGGCTCTAC | Tb927.3.5510_rev | GGGGACCACTTTGTACAAGAAAGCTGGGTTTACTTCATTTCAGATACTTGCACAGAACG |
| 10_hypothetical | B20 | Tb927.11.1420_for | GGGGACAAGTTTGTACAAAAAAGCAGGCTATGCCGGAAAAACCTGGTTCAC | Tb927.11.1420_rev | GGGGACCACTTTGTACAAGAAAGCTGGGTCTAACCAAATAACACACGACGGGT |
| 10_hypothetical | B21 | Tb927.4.4880_for | GGGGACAAGTTTGTACAAAAAAGCAGGCTATGATTCCGAACCGCTCCATGT | Tb927.4.4880_rev | GGGGACCACTTTGTACAAGAAAGCTGGGTCTAGGGCAACTGTTGCTGCTC |
| 10_hypothetical | B22 | Tb927.5.2660_for | GGGGACAAGTTTGTACAAAAAAGCAGGCTATGACGGAAACAGCGTTTCGA | Tb927.5.2660_rev | GGGGACCACTTTGTACAAGAAAGCTGGGTTTATGCCTTCCATCCACGGCG |
| 10_hypothetical | B23 | Tb927.10.14380_for | GGGGACAAGTTTGTACAAAAAAGCAGGCTATGCACGCGGTAAGCATTTCA | Tb927.10.14380_rev | GGGGACCACTTTGTACAAGAAAGCTGGGTTTACAAATCCACCATAACGCTCGC |
| 10_hypothetical | B24 | Tb927.5.4110_for | GGGGACAAGTTTGTACAAAAAAGCAGGCTATGGTAGGTTTGATCCCGTATGACA | Tb927.5.4110_rev | GGGGACCACTTTGTACAAGAAAGCTGGGTCTATACTTGTCGTTCCTGTGCCTTT |
| 10_hypothetical | C1 | Tb927.3.4580_for | GGGGACAAGTTTGTACAAAAAAGCAGGCTATGCTCTCTCAATTTGATGACAACCC | Tb927.3.4580_rev | GGGGACCACTTTGTACAAGAAAGCTGGGTTTAGGGTACAACTTCATCTCCATCCC |
| 10_hypothetical | C2 | Tb927.5.1180_for | GGGGACAAGTTTGTACAAAAAAGCAGGCTATGCAGTGGCCGTCATGGG | Tb927.5.1180_rev | GGGGACCACTTTGTACAAGAAAGCTGGGTCTACGCTGAACGGCCACAC |
| 10_hypothetical | C3 | Tb927.6.1950_for | GGGGACAAGTTTGTACAAAAAAGCAGGCTATGGCTGGCAGAGTTTCAGATCG | Tb927.6.1950_rev | GGGGACCACTTTGTACAAGAAAGCTGGGTCTAGCGCCTACGGGAGGC |
| 10_hypothetical | C4 | Tb927.11.2280_for | GGGGACAAGTTTGTACAAAAAAGCAGGCTATGACTACATTTGAGCACCTACCTGA | Tb927.11.2280_rev | GGGGACCACTTTGTACAAGAAAGCTGGGTCTAAGATAAGGGATTATGAGGTGCATAGTA |
| 10_hypothetical | C5 | Tb927.6.1070_for | GGGGACAAGTTTGTACAAAAAAGCAGGCTATGTCCCGAGCACTTGAAAGGC | Tb927.6.1070_rev | GGGGACCACTTTGTACAAGAAAGCTGGGTCTACCCTGATGCGATATTTGCCG |
| 10_hypothetical | C6 | Tb927.10.4200_for | GGGGACAAGTTTGTACAAAAAAGCAGGCTATGTCATCCAAGCAGGAGAGAATAGC | Tb927.10.4200_rev | GGGGACCACTTTGTACAAGAAAGCTGGGTTCAAGAATCTGTTGCAGGACGC |
| 10_hypothetical | C7 | Tb927.10.15840_for | GGGGACAAGTTTGTACAAAAAAGCAGGCTATGGATTTCCGAACTATCCGGCA | Tb927.10.15840_rev | GGGGACCACTTTGTACAAGAAAGCTGGGTTTAAAGCGATAGTGCATGTATCAAGTCA |
| 10_hypothetical | C8 | Tb927.10.14480_for | GGGGACAAGTTTGTACAAAAAAGCAGGCTATGGCCATTAAACCAAAGCGTATTGA | Tb927.10.14480_rev | GGGGACCACTTTGTACAAGAAAGCTGGGTTTACAACTTCTTACGCGTGAGACTC |
| 10_hypothetical | C9 | Tb927.1.3560_for | GGGGACAAGTTTGTACAAAAAAGCAGGCTATGTCCGTAGTTCTAGATGTTGTGACG | Tb927.1.3560_rev | GGGGACCACTTTGTACAAGAAAGCTGGGTTTAATGAGCCGATTTACCCACGTAG |
| 10_hypothetical | C10 | Tb927.11.390_for | GGGGACAAGTTTGTACAAAAAAGCAGGCTATGGGTAGTTTACCCTGCGACA | Tb927.11.390_rev | GGGGACCACTTTGTACAAGAAAGCTGGGTCTAAGCAATGGTTGCAGTATCGCC |
| 10_hypothetical | C11 | Tb927.9.12310_for | GGGGACAAGTTTGTACAAAAAAGCAGGCTATGGGATCTAGTGATGATGAGTGGC | Tb927.9.12310_rev | GGGGACCACTTTGTACAAGAAAGCTGGGTTTAGGGTGATTGCGTCGCTG |
| 10_hypothetical | C12 | Tb927.3.1900_for | GGGGACAAGTTTGTACAAAAAAGCAGGCTATGTCGAAGTCGCTCCCCC | Tb927.3.1900_rev | GGGGACCACTTTGTACAAGAAAGCTGGGTTTATGCTTCATCTTCTATCTCATCTTCACC |
| 10_hypothetical | C13 | Tb927.5.2020_for | GGGGACAAGTTTGTACAAAAAAGCAGGCTATGCTGATGGAAACGAAATTTCGC | Tb927.5.2020_rev | GGGGACCACTTTGTACAAGAAAGCTGGGTTTAACCTTCGCGGCGCCG |
| 10_hypothetical | C14 | Tb927.10.8120_for | GGGGACAAGTTTGTACAAAAAAGCAGGCTATGGAGGTTCTTGTTGAAAGGTGTC | Tb927.10.8120_rev | GGGGACCACTTTGTACAAGAAAGCTGGGTCTAAACAACAACAACAACTCCTCGC |
| 10_hypothetical | C15 | Tb927.10.9890_for | GGGGACAAGTTTGTACAAAAAAGCAGGCTATGAGTGTCTCCTTTTTTGTTCAGGAG | Tb927.10.9890_rev | GGGGACCACTTTGTACAAGAAAGCTGGGTTTACAGCCTCGCGTGCGG |
| 10_hypothetical | C16 | Tb927.3.2750_for | GGGGACAAGTTTGTACAAAAAAGCAGGCTATGCAGCGTCAGTGTGACG | Tb927.3.2750_rev | GGGGACCACTTTGTACAAGAAAGCTGGGTTCAACCGCGTTTTGTGTTTCG |
| 10_hypothetical | C17 | Tb927.10.11250_for | GGGGACAAGTTTGTACAAAAAAGCAGGCTATGTCCGACGTACCATACGATGT | Tb927.10.11250_rev | GGGGACCACTTTGTACAAGAAAGCTGGGTTCAGTGCAGAGATGATGCTATATCAGA |
| 10_hypothetical | C18 | Tb927.8.7580_for | GGGGACAAGTTTGTACAAAAAAGCAGGCTATGGAACATCTGAAAATTTTTCCGCG | Tb927.8.7580_rev | GGGGACCACTTTGTACAAGAAAGCTGGGTCTAGGGCAACTGCCGCTG |
| 10_hypothetical | C19 | Tb927.11.2820_for | GGGGACAAGTTTGTACAAAAAAGCAGGCTATGTGGCAGCCACGTCACG | Tb927.11.2820_rev | GGGGACCACTTTGTACAAGAAAGCTGGGTTCAGCGAAACTGGCACGG |
| 10_hypothetical | C20 | Tb927.2.2380_for | GGGGACAAGTTTGTACAAAAAAGCAGGCTATGCGCTCAGAAACACGGC | Tb927.2.2380_rev | GGGGACCACTTTGTACAAGAAAGCTGGGTCTACAATCCACTGCACACTTCTCC |
| 10_hypothetical | C21 | Tb927.11.1610_for | GGGGACAAGTTTGTACAAAAAAGCAGGCTATGCCGGTCCCACATCAGT | Tb927.11.1610_rev | GGGGACCACTTTGTACAAGAAAGCTGGGTTTACGTGTCTAAAACGCTGGCG |
| 10_hypothetical | C22 | Tb927.8.7090_for | GGGGACAAGTTTGTACAAAAAAGCAGGCTATGGATGATGGACCTGACCCTC | Tb927.8.7090_rev | GGGGACCACTTTGTACAAGAAAGCTGGGTTCATTCAACGGCAGTTGCAGAT |
| 10_hypothetical | C23 | Tb927.10.15160_for | GGGGACAAGTTTGTACAAAAAAGCAGGCTATGCTTGCCGTGAAGTACCTTC | Tb927.10.15160_rev | GGGGACCACTTTGTACAAGAAAGCTGGGTCTATGCGATGGGGATGAAGCC |
| 10_hypothetical | C24 | Tb927.8.5750_for | GGGGACAAGTTTGTACAAAAAAGCAGGCTATGCTGTCCTCTGCGCTCG | Tb927.8.5750_rev | GGGGACCACTTTGTACAAGAAAGCTGGGTTTAGAGCAGAGCCGCCTTCA |
| 10_hypothetical | D1 | Tb927.2.5440_for | GGGGACAAGTTTGTACAAAAAAGCAGGCTATGAGTTTCCGAAGACAAAGTTTTAGGG | Tb927.2.5440_rev | GGGGACCACTTTGTACAAGAAAGCTGGGTCTAACGGGGCTGGTTGGC |
| 10_hypothetical | D2 | Tb927.6.2800_for | GGGGACAAGTTTGTACAAAAAAGCAGGCTATGTCAGCCCGAAACAAAATGGG | Tb927.6.2800_rev | GGGGACCACTTTGTACAAGAAAGCTGGGTTTATGAATATCGGTGGCGGAACTG |
| 10_hypothetical | D3 | Tb927.8.610_for | GGGGACAAGTTTGTACAAAAAAGCAGGCTATGGGTCCCAAAAAGGCTGGT | Tb927.8.610_rev | GGGGACCACTTTGTACAAGAAAGCTGGGTTCAAAACTGTGTGCCAACCCC |
| 10_hypothetical | D4 | Tb927.11.16850_for | GGGGACAAGTTTGTACAAAAAAGCAGGCTATGTTTACAGACAGCATTCTTATGAGCC | Tb927.11.16850_rev | GGGGACCACTTTGTACAAGAAAGCTGGGTTCACTTCCCTACCTGGCGATG |
| 10_hypothetical | D5 | Tb927.9.4350_for | GGGGACAAGTTTGTACAAAAAAGCAGGCTATGACCGAATCTGAGAAAGCCCG | Tb927.9.4350_rev | GGGGACCACTTTGTACAAGAAAGCTGGGTTCATCCCCCACTCCATTTCACC |
| 10_hypothetical | D6 | Tb927.11.10580_for | GGGGACAAGTTTGTACAAAAAAGCAGGCTATGTCTCAATTTGCGACCCTGTC | Tb927.11.10580_rev | GGGGACCACTTTGTACAAGAAAGCTGGGTCTACTCGTCAGTTTCGTCAATATCCG |
| 10_hypothetical | D7 | Tb927.10.13470_for | GGGGACAAGTTTGTACAAAAAAGCAGGCTATGCAGCCCAACACGATGC | Tb927.10.13470_rev | GGGGACCACTTTGTACAAGAAAGCTGGGTCTACATGTTGTTAAGAATTTTCCACATTGG |
| 10_hypothetical | D8 | Tb927.4.5330_for | GGGGACAAGTTTGTACAAAAAAGCAGGCTATGAAGGGCCCGCTTTCCA | Tb927.4.5330_rev | GGGGACCACTTTGTACAAGAAAGCTGGGTTCACATCATCGATACCAGAGCACG |
| 10_hypothetical | D9 | Tb927.9.13570_for | GGGGACAAGTTTGTACAAAAAAGCAGGCTATGACTGAAGAGAAGCAGCAGAAGC | Tb927.9.13570_rev | GGGGACCACTTTGTACAAGAAAGCTGGGTCTAATTGGGATGCCGTCTACCCA |
| 10_hypothetical | D10 | Tb927.8.1070_for | GGGGACAAGTTTGTACAAAAAAGCAGGCTATGCACAGTGTTACTGTCCTTTCTC | Tb927.8.1070_rev | GGGGACCACTTTGTACAAGAAAGCTGGGTTCAGCACTTTCTTTCCACACCAC |
| 10_hypothetical | D11 | Tb927.10.8610_for | GGGGACAAGTTTGTACAAAAAAGCAGGCTATGCGCCGTTGCCTCTTC | Tb927.10.8610_rev | GGGGACCACTTTGTACAAGAAAGCTGGGTTCATTTCAACATGTTCCCCCTCG |
| 10_hypothetical | D12 | Tb927.8.6330_for | GGGGACAAGTTTGTACAAAAAAGCAGGCTATGTCGAAGGCCCCTGAGC | Tb927.8.6330_rev | GGGGACCACTTTGTACAAGAAAGCTGGGTTCAAGACCCATCAGCCTCTAGC |
| 10_hypothetical | D13 | Tb927.11.9400_for | GGGGACAAGTTTGTACAAAAAAGCAGGCTATGCAACAGCAAAACCGCAC | Tb927.11.9400_rev | GGGGACCACTTTGTACAAGAAAGCTGGGTTCAATCCCTGCTTCCTTCACTGC |
| 10_hypothetical | D14 | Tb927.4.5320_for | GGGGACAAGTTTGTACAAAAAAGCAGGCTATGGATAATGGCCCTCATCTCTTCA | Tb927.4.5320_rev | GGGGACCACTTTGTACAAGAAAGCTGGGTTCACTCAACAGCAGTTGCGG |
| 10_hypothetical | D15 | Tb927.11.4340_for | GGGGACAAGTTTGTACAAAAAAGCAGGCTATGAACTTTGACGATCTCTTCGCG | Tb927.11.4340_rev | GGGGACCACTTTGTACAAGAAAGCTGGGTCTACGGGTCAAAGGCTTGTGC |
| 10_hypothetical | D16 | Tb927.8.1540_for | GGGGACAAGTTTGTACAAAAAAGCAGGCTATGTCAACTCCATCTGTGGTCGG | Tb927.8.1540_rev | GGGGACCACTTTGTACAAGAAAGCTGGGTCTACTTTGAGTCGGAAACGCTGG |
| 10_hypothetical | D17 | Tb927.7.1580_for | GGGGACAAGTTTGTACAAAAAAGCAGGCTATGATTGCTGATGCGGAATGGAG | Tb927.7.1580_rev | GGGGACCACTTTGTACAAGAAAGCTGGGTCTATCCCTCCTCATCATCATCTTTGC |
| 10_hypothetical | D18 | Tb927.11.16910_for | GGGGACAAGTTTGTACAAAAAAGCAGGCTATGGTGAAAGTTTTATGTCTGCACGG | Tb927.11.16910_rev | GGGGACCACTTTGTACAAGAAAGCTGGGTTCACACCTTTGGCCCCACA |
| 10_hypothetical | D19 | Tb927.5.3850_for | GGGGACAAGTTTGTACAAAAAAGCAGGCTATGGGGATAATAGGTCTGCGCA | Tb927.5.3850_rev | GGGGACCACTTTGTACAAGAAAGCTGGGTTCATCTTTGATGGTCCTGTACCTTCC |
| 10_hypothetical | D20 | Tb927.11.8370_for | GGGGACAAGTTTGTACAAAAAAGCAGGCTATGTCTGATTCGCACCACAGC | Tb927.11.8370_rev | GGGGACCACTTTGTACAAGAAAGCTGGGTTCACCTTTTCTTGTACTGGCTGTG |
| 10_hypothetical | D21 | Tb927.2.4000_for | GGGGACAAGTTTGTACAAAAAAGCAGGCTATGTCACAACGAGTGCCACG | Tb927.2.4000_rev | GGGGACCACTTTGTACAAGAAAGCTGGGTCTAACGCCCTTTTTGTGAAAGAAAGC |
| 10_hypothetical | D22 | Tb927.4.3170_for | GGGGACAAGTTTGTACAAAAAAGCAGGCTATGCAGGGAGACACTAAAACTATGGG | Tb927.4.3170_rev | GGGGACCACTTTGTACAAGAAAGCTGGGTTTACAAAACTGAGCCCATGGAGGG |
| 10_hypothetical | D23 | Tb927.7.3230_for | GGGGACAAGTTTGTACAAAAAAGCAGGCTATGGATGGACAGCGTGCAAC | Tb927.7.3230_rev | GGGGACCACTTTGTACAAGAAAGCTGGGTTCAGTTGCTTGCATGACAAAGGC |
| 10_hypothetical | D24 | Tb927.11.5100_for | GGGGACAAGTTTGTACAAAAAAGCAGGCTATGATATCAGAGAGCCAGATGTTTAGCC | Tb927.11.5100_rev | GGGGACCACTTTGTACAAGAAAGCTGGGTTTACAGAGTTAACGTTAGAGATGTATCAGT |
| 10_hypothetical | E1 | Tb927.7.6610_for | GGGGACAAGTTTGTACAAAAAAGCAGGCTATGTTCTCCGTTGAGTTTGTAAGTGG | Tb927.7.6610_rev | GGGGACCACTTTGTACAAGAAAGCTGGGTTTACTTCACTTTTTGTGGACGATGCG |
| 10_hypothetical | E2 | Tb927.9.15330_for | GGGGACAAGTTTGTACAAAAAAGCAGGCTATGGCGCGAGCAACTGCT | Tb927.9.15330_rev | GGGGACCACTTTGTACAAGAAAGCTGGGTTTAGTATTTATTCAACATGAGAAGTAGATG |
| 10_hypothetical | E3 | Tb927.11.10630_for | GGGGACAAGTTTGTACAAAAAAGCAGGCTATGAAAGTTATTACACATGAGCAAGTAGCG | Tb927.11.10630_rev | GGGGACCACTTTGTACAAGAAAGCTGGGTCTAAAAGCGCCTCTTTATTTCGTTGC |
| 10_hypothetical | E4 | Tb927.8.2860_for | GGGGACAAGTTTGTACAAAAAAGCAGGCTATGTCTGTTAGCGCCCCAACG | Tb927.8.2860_rev | GGGGACCACTTTGTACAAGAAAGCTGGGTCTAAGCAAATGGGTCAAACTCCGC |
| 10_hypothetical | E5 | Tb927.7.5210_for | GGGGACAAGTTTGTACAAAAAAGCAGGCTATGAGCCTGGTACAGAACCAAGT | Tb927.7.5210_rev | GGGGACCACTTTGTACAAGAAAGCTGGGTTCAGTGACTCTCTCCGGTGGC |
| 10_hypothetical | E6 | Tb927.6.1290_for | GGGGACAAGTTTGTACAAAAAAGCAGGCTATGGATCTCTTCCCCAAGCATTTG | Tb927.6.1290_rev | GGGGACCACTTTGTACAAGAAAGCTGGGTTTAAAGTGACTCTATAAAGAGCAGAACTTC |
| 10_hypothetical | E7 | Tb927.8.6360_for | GGGGACAAGTTTGTACAAAAAAGCAGGCTATGTTTTACTCAGGGTGGCAGTTTC | Tb927.8.6360_rev | GGGGACCACTTTGTACAAGAAAGCTGGGTCTAGGCAAAAACCGCCCTATAAACG |
| 10_hypothetical | E8 | Tb927.3.5450_for | GGGGACAAGTTTGTACAAAAAAGCAGGCTATGAGTGATTACGGAGATTCTTTTGATAGC | Tb927.3.5450_rev | GGGGACCACTTTGTACAAGAAAGCTGGGTTTACACATTCCTTGCCTCTTCAACAC |
| 10_hypothetical | E9 | Tb927.8.7760_for | GGGGACAAGTTTGTACAAAAAAGCAGGCTATGGGTGCTGCTCCAAGCA | Tb927.8.7760_rev | GGGGACCACTTTGTACAAGAAAGCTGGGTTTATCTATCTGCGCAATATCTCCAGTCC |
| 10_hypothetical | E10 | Tb927.9.8040_for | GGGGACAAGTTTGTACAAAAAAGCAGGCTATGCAGGTTTTATTGAAACTTCGGC | Tb927.9.8040_rev | GGGGACCACTTTGTACAAGAAAGCTGGGTTCAATGGCATATGAGAAACGACTCC |
| 10_hypothetical | E11 | Tb927.4.4570_for | GGGGACAAGTTTGTACAAAAAAGCAGGCTATGGGTGTTGCACCCAGTCG | Tb927.4.4570_rev | GGGGACCACTTTGTACAAGAAAGCTGGGTTTAGGACTTCCTCCAGTCCCAAATAA |
| 10_hypothetical | E12 | Tb927.10.12720_for | GGGGACAAGTTTGTACAAAAAAGCAGGCTATGGAGACGCACCTCGGC | Tb927.10.12720_rev | GGGGACCACTTTGTACAAGAAAGCTGGGTTTAGCGCCATGGTCGAGGC |
| 10_hypothetical | E13 | Tb927.10.12340_for | GGGGACAAGTTTGTACAAAAAAGCAGGCTATGAGAAGATGGCGCCCTGT | Tb927.10.12340_rev | GGGGACCACTTTGTACAAGAAAGCTGGGTTTAGATGTCCTCCTTAACAGTGCAGT |
| 10_hypothetical | E14 | Tb927.6.4640_for | GGGGACAAGTTTGTACAAAAAAGCAGGCTATGCCCAAGAAAACTCGCGT | Tb927.6.4640_rev | GGGGACCACTTTGTACAAGAAAGCTGGGTTCAAAAACGCATCGGTGGTCC |
| 10_hypothetical | E15 | Tb927.10.5890_for | GGGGACAAGTTTGTACAAAAAAGCAGGCTATGCAAAGGGTAGGTAGTGAAAGGG | Tb927.10.5890_rev | GGGGACCACTTTGTACAAGAAAGCTGGGTTCAATGATTCTTTGACCTTCTCCCCC |
| 10_hypothetical | E16 | Tb927.7.5430_for | GGGGACAAGTTTGTACAAAAAAGCAGGCTATGTCTGCCTCTCGATCTCCC | Tb927.7.5430_rev | GGGGACCACTTTGTACAAGAAAGCTGGGTCTACCTTCTCCCACCCCGC |
| 10_hypothetical | E18 | Tb927.1.3910_for | GGGGACAAGTTTGTACAAAAAAGCAGGCTATGCATCAGCAAGACCCCACC | Tb927.1.3910_rev | GGGGACCACTTTGTACAAGAAAGCTGGGTTCAAGCTCGTTCTCCCTGACG |
| 10_hypothetical | E19 | Tb927.9.13520_for | GGGGACAAGTTTGTACAAAAAAGCAGGCTATGGGCAGCGTGGGACAG | Tb927.9.13520_rev | GGGGACCACTTTGTACAAGAAAGCTGGGTTTACTGGATACCATAGACATGATAGAGACG |
| 10_hypothetical | E20 | Tb927.4.1160_for | GGGGACAAGTTTGTACAAAAAAGCAGGCTATGCGTCATCTCCACACCGC | Tb927.4.1160_rev | GGGGACCACTTTGTACAAGAAAGCTGGGTTCACGACCATTTTTTAAGGAAAGGGT |
| 10_hypothetical | E21 | Tb927.4.1320_for | GGGGACAAGTTTGTACAAAAAAGCAGGCTATGCGCCTTTCCCGCTATG | Tb927.4.1320_rev | GGGGACCACTTTGTACAAGAAAGCTGGGTCTATCTCGAGGCGGATAGAAGCA |
| 10_hypothetical | E22 | Tb927.10.8810_for | GGGGACAAGTTTGTACAAAAAAGCAGGCTATGATCCCCGTGGCAGGC | Tb927.10.8810_rev | GGGGACCACTTTGTACAAGAAAGCTGGGTTTAGTTGAGAGTTGAAGGTGTTGTAAAGT |
| 10_hypothetical | E23 | Tb927.11.3360_for | GGGGACAAGTTTGTACAAAAAAGCAGGCTATGTCTTCGTTGACATGTGAGGAGC | Tb927.11.3360_rev | GGGGACCACTTTGTACAAGAAAGCTGGGTTTATTCCTTCTTCTTGGGTCTCTTCTTAGC |
| 10_hypothetical | E24 | Tb927.10.420_for | GGGGACAAGTTTGTACAAAAAAGCAGGCTATGAACGAAAGTATTATTCAAGTGCCACT | Tb927.10.420_rev | GGGGACCACTTTGTACAAGAAAGCTGGGTTCAATCAGCATCTTCAGCAACTCTG |
| 10_hypothetical | F1 | Tb927.8.4370_for | GGGGACAAGTTTGTACAAAAAAGCAGGCTATGGCTGACAGTAACAACAGCG | Tb927.8.4370_rev | GGGGACCACTTTGTACAAGAAAGCTGGGTTCACTTTGGTGCACTCTGGATCA |
| 10_hypothetical | F2 | Tb927.11.16840.1_for | GGGGACAAGTTTGTACAAAAAAGCAGGCTATGGCCTCGCTACTTCAGCA | Tb927.11.16840.1_rev | GGGGACCACTTTGTACAAGAAAGCTGGGTCTAAGCAACTACCTCCTCACTCCT |
| 10_hypothetical | F3 | Tb927.1.3270_for | GGGGACAAGTTTGTACAAAAAAGCAGGCTATGAGTGCATTTGCGCTGGC | Tb927.1.3270_rev | GGGGACCACTTTGTACAAGAAAGCTGGGTTTAATTTCCCTTCCTTCCGCCTGC |
| 10_hypothetical | F4 | Tb927.7.7220_for | GGGGACAAGTTTGTACAAAAAAGCAGGCTATGAGCACCAGTTCGTCTATATTCTTTG | Tb927.7.7220_rev | GGGGACCACTTTGTACAAGAAAGCTGGGTCTATTGTTCGTGAGAGGCAGGC |
| 10_hypothetical | F5 | Tb927.1.3940_for | GGGGACAAGTTTGTACAAAAAAGCAGGCTATGGCAACACCTCGGCAGG | Tb927.1.3940_rev | GGGGACCACTTTGTACAAGAAAGCTGGGTTCAAAGAATAGACATGGTAGCAAAGCG |
| 10_hypothetical | F6 | Tb927.8.5650_for | GGGGACAAGTTTGTACAAAAAAGCAGGCTATGGATTTTGTTTTCCCAGTATTTGTTCC | Tb927.8.5650_rev | GGGGACCACTTTGTACAAGAAAGCTGGGTCTCACTCCGACGTTAGAGCTTGA |
| 10_hypothetical | F7 | Tb927.8.3950_for | GGGGACAAGTTTGTACAAAAAAGCAGGCTATGGGGAGCGTTGATCATCCG | Tb927.8.3950_rev | GGGGACCACTTTGTACAAGAAAGCTGGGTTTATACCTTCGTAATACCCGACACAGT |
| 10_hypothetical | F8 | Tb927.5.2520_for | GGGGACAAGTTTGTACAAAAAAGCAGGCTATGGTGCCATCCATCCCTCAA | Tb927.5.2520_rev | GGGGACCACTTTGTACAAGAAAGCTGGGTTCAGTCGGCCATAATGCGAGT |
| 10_hypothetical | F9 | Tb927.5.1070_for | GGGGACAAGTTTGTACAAAAAAGCAGGCTATGTCTGCAATCAAAGAACTTGTCGA | Tb927.5.1070_rev | GGGGACCACTTTGTACAAGAAAGCTGGGTCTAATGAGGGCCTCGCAAATCTC |
| 10_hypothetical | F10 | Tb927.3.4270_for | GGGGACAAGTTTGTACAAAAAAGCAGGCTATGGGGCATCCGAGCGAC | Tb927.3.4270_rev | GGGGACCACTTTGTACAAGAAAGCTGGGTCTACATGAGGCTGGCAAGATCCT |
| 10_hypothetical | F11 | Tb927.10.2380_for | GGGGACAAGTTTGTACAAAAAAGCAGGCTATGACTACAGAGGTCAGTAATGACGG | Tb927.10.2380_rev | GGGGACCACTTTGTACAAGAAAGCTGGGTTCACGACTTTCCAAGGGCGG |
| 10_hypothetical | F12 | Tb927.11.6910_for | GGGGACAAGTTTGTACAAAAAAGCAGGCTATGACTATCCTACACCACCCGG | Tb927.11.6910_rev | GGGGACCACTTTGTACAAGAAAGCTGGGTTCAAACGGCAGGTGTTGCTT |
| 10_hypothetical | F13 | Tb927.1.830_for | GGGGACAAGTTTGTACAAAAAAGCAGGCTATGGAACCGGTGGACGGG | Tb927.1.830_rev | GGGGACCACTTTGTACAAGAAAGCTGGGTTTAATTCCCTTTAAGCACCCGTCTC |
| 10_hypothetical | F14 | Tb927.11.16920_for | GGGGACAAGTTTGTACAAAAAAGCAGGCTATGCATCACATACCCATCTCGTCC | Tb927.11.16920_rev | GGGGACCACTTTGTACAAGAAAGCTGGGTTTACGAGCCCACACCTAACCA |
| 10_hypothetical | F15 | Tb927.11.12510_for | GGGGACAAGTTTGTACAAAAAAGCAGGCTATGACGAGTGTTAAGGAAACAAAGGAG | Tb927.11.12510_rev | GGGGACCACTTTGTACAAGAAAGCTGGGTTTAATCTACATTTTTTCTTTGCGCCGC |
| 10_hypothetical | F16 | Tb927.11.5750_for | GGGGACAAGTTTGTACAAAAAAGCAGGCTATGGTGCGGTCCCAGTCTC | Tb927.11.5750_rev | GGGGACCACTTTGTACAAGAAAGCTGGGTTCAAATGATTTCCCCGCTATTTCCA |
| 10_hypothetical | F17 | Tb927.4.2900_for | GGGGACAAGTTTGTACAAAAAAGCAGGCTATGACTGAAAGTGACATGTATCGGGG | Tb927.4.2900_rev | GGGGACCACTTTGTACAAGAAAGCTGGGTTCAATCTGGCAGCTGACTGTCC |
| 10_hypothetical | F18 | Tb927.11.10720_for | GGGGACAAGTTTGTACAAAAAAGCAGGCTATGGTGTACAACAAAGTGCTGCG | Tb927.11.10720_rev | GGGGACCACTTTGTACAAGAAAGCTGGGTTCACCACACAATCTCCCCAGC |
| 10_hypothetical | F19 | Tb927.10.14430_for | GGGGACAAGTTTGTACAAAAAAGCAGGCTATGTGCACCACTAGGTGTCTTCA | Tb927.10.14430_rev | GGGGACCACTTTGTACAAGAAAGCTGGGTTTAACATGTCTCTTTTTCCTTCGCACC |
| 10_hypothetical | F20 | Tb927.3.2430_for | GGGGACAAGTTTGTACAAAAAAGCAGGCTATGAAACGATGCAACAGTGAGTCT | Tb927.3.2430_rev | GGGGACCACTTTGTACAAGAAAGCTGGGTTTATAAAATAGATGCAATAAATGCCGCAGC |
| 10_hypothetical | F21 | Tb927.7.2630_for | GGGGACAAGTTTGTACAAAAAAGCAGGCTATGCCATGGCTAACTCCACTACG | Tb927.7.2630_rev | GGGGACCACTTTGTACAAGAAAGCTGGGTCTACCGACCCTTGACTCGAGG |
| 10_hypothetical | F22 | Tb927.11.4500_for | GGGGACAAGTTTGTACAAAAAAGCAGGCTATGAGTCTCTCCAATCAACTTCCGC | Tb927.11.4500_rev | GGGGACCACTTTGTACAAGAAAGCTGGGTTTATTCCAGGAGTGTGGAGAAAAGC |
| 10_hypothetical | F23 | Tb927.9.7760_for | GGGGACAAGTTTGTACAAAAAAGCAGGCTATGCAGGTTTGCCCATTAGCAC | Tb927.9.7760_rev | GGGGACCACTTTGTACAAGAAAGCTGGGTTTACTGAAAAGGAGGCGGTAGCT |
| 10_hypothetical | F24 | Tb927.6.2350_for | GGGGACAAGTTTGTACAAAAAAGCAGGCTATGCTGTGCAGCGGCCGC | Tb927.6.2350_rev | GGGGACCACTTTGTACAAGAAAGCTGGGTTCACATCCCCGCACCCGT |
| 10_hypothetical | G2 | Tb927.11.14190_for | GGGGACAAGTTTGTACAAAAAAGCAGGCTATGTCTTTTCTTGTGTATGCCGTTGA | Tb927.11.14190_rev | GGGGACCACTTTGTACAAGAAAGCTGGGTTCAGTTATCCTCGTCGCTTTCGT |
| 10_hypothetical | G3 | Tb927.5.2710_for | GGGGACAAGTTTGTACAAAAAAGCAGGCTATGAGCTCTTCACAAGGAGAAATGC | Tb927.5.2710_rev | GGGGACCACTTTGTACAAGAAAGCTGGGTTCACTCGTATTTGACCGCCTCA |
| 10_hypothetical | G4 | Tb927.11.2600_for | GGGGACAAGTTTGTACAAAAAAGCAGGCTATGCTTATTACTGAAAATCTATTTGCCTCA | Tb927.11.2600_rev | GGGGACCACTTTGTACAAGAAAGCTGGGTTCACTTAAATACCACTCCACTACCACC |
| 10_hypothetical | G5 | Tb927.5.4520_for | GGGGACAAGTTTGTACAAAAAAGCAGGCTATGGAGTCTCCTGCGAAAGGT | Tb927.5.4520_rev | GGGGACCACTTTGTACAAGAAAGCTGGGTCTAATTCGATCTTTGCCTCTTTGCAC |
| 10_hypothetical | G6 | Tb927.11.5160_for | GGGGACAAGTTTGTACAAAAAAGCAGGCTATGAAGGAGTTAAACGCCATTCTTCA | Tb927.11.5160_rev | GGGGACCACTTTGTACAAGAAAGCTGGGTTCAGCTTGGACCTAGGCCC |
| 10_hypothetical | G7 | Tb927.10.13200_for | GGGGACAAGTTTGTACAAAAAAGCAGGCTATGACACTGACTCTTCTTAGGAGGG | Tb927.10.13200_rev | GGGGACCACTTTGTACAAGAAAGCTGGGTTCAGACGCGCGACATCGA |
| 10_hypothetical | G8 | Tb927.5.2600_for | GGGGACAAGTTTGTACAAAAAAGCAGGCTATGTGCCACCGTCTCGTCC | Tb927.5.2600_rev | GGGGACCACTTTGTACAAGAAAGCTGGGTTTATTGCGCCACGGGAAAACG |
| 10_hypothetical | G9 | Tb927.10.12910_for | GGGGACAAGTTTGTACAAAAAAGCAGGCTATGTTCCCCCCGGTAGCAC | Tb927.10.12910_rev | GGGGACCACTTTGTACAAGAAAGCTGGGTTCAATCAAGGAAATCGTAGACTTCATTGT |
| 10_hypothetical | G10 | Tb927.11.3300_for | GGGGACAAGTTTGTACAAAAAAGCAGGCTATGGCAGAGGTGAAGCTTCCG | Tb927.11.3300_rev | GGGGACCACTTTGTACAAGAAAGCTGGGTTCAACTTTCGTCAGGGAAAATGGTT |
| 10_hypothetical | G11 | Tb927.11.7690_for | GGGGACAAGTTTGTACAAAAAAGCAGGCTATGCGGCAAATCCATCTAAGTAGTAAC | Tb927.11.7690_rev | GGGGACCACTTTGTACAAGAAAGCTGGGTCTAAACAAAAGTCCTTTTAGCCAATTCCTG |
| 10_hypothetical | G12 | Tb927.4.3120_for | GGGGACAAGTTTGTACAAAAAAGCAGGCTATGGAAATTGAAGGGCATGAGACC | Tb927.4.3120_rev | GGGGACCACTTTGTACAAGAAAGCTGGGTTCACGATATCATGCTAAATAGTTCACTGC |
| 10_hypothetical | G13 | Tb927.10.12590_for | GGGGACAAGTTTGTACAAAAAAGCAGGCTATGGGTGACGACACGACCA | Tb927.10.12590_rev | GGGGACCACTTTGTACAAGAAAGCTGGGTCTACAACCAGAGTTCGTTCCTATCTGG |
| 10_hypothetical | G14 | Tb927.7.480_for | GGGGACAAGTTTGTACAAAAAAGCAGGCTATGCGACCCTCCAGGCGT | Tb927.7.480_rev | GGGGACCACTTTGTACAAGAAAGCTGGGTTCAACTGCCAATGCGGCTC |
| 10_hypothetical | G15 | Tb927.4.1390_for | GGGGACAAGTTTGTACAAAAAAGCAGGCTATGATGTCACCGTCCCTCGC | Tb927.4.1390_rev | GGGGACCACTTTGTACAAGAAAGCTGGGTTCAAGGATTACCTTCGACCTCAGG |
| 10_hypothetical | G16 | Tb927.7.4330_for | GGGGACAAGTTTGTACAAAAAAGCAGGCTATGACGGCACTCGACTTTCAGT | Tb927.7.4330_rev | GGGGACCACTTTGTACAAGAAAGCTGGGTTCATACACTACTACCACCAGAACTCAA |
| 10_hypothetical | G17 | Tb927.5.2420_for | GGGGACAAGTTTGTACAAAAAAGCAGGCTATGAGACACTGCCATATTATTCGGTATG | Tb927.5.2420_rev | GGGGACCACTTTGTACAAGAAAGCTGGGTCTACAGGATGAAGTAATCCTTTTCCTTTGT |
| 10_hypothetical | G18 | Tb927.11.6660_for | GGGGACAAGTTTGTACAAAAAAGCAGGCTATGCTGCGGTACGTTAAAAGAATTG | Tb927.11.6660_rev | GGGGACCACTTTGTACAAGAAAGCTGGGTTCAGAACACAAATACACTTTGAGCATCG |
| 10_hypothetical | G19 | Tb927.11.14370_for | GGGGACAAGTTTGTACAAAAAAGCAGGCTATGTCCAAGGAGGCTCCACAT | Tb927.11.14370_rev | GGGGACCACTTTGTACAAGAAAGCTGGGTCTACTTTCCTCCCTTGTTGAAGCG |
| 10_hypothetical | G20 | Tb927.2.4780_for | GGGGACAAGTTTGTACAAAAAAGCAGGCTATGTCCATCACAGTTGACACTATGAGT | Tb927.2.4780_rev | GGGGACCACTTTGTACAAGAAAGCTGGGTTTACCTCCCGGGGCGTACA |
| 10_hypothetical | G21 | Tb927.11.10610_for | GGGGACAAGTTTGTACAAAAAAGCAGGCTATGCAATATATTATTCACGACCCACACA | Tb927.11.10610_rev | GGGGACCACTTTGTACAAGAAAGCTGGGTTTACTCAATGGGAGGAATGACCAATTC |
| 10_hypothetical | G22 | Tb927.8.8270_for | GGGGACAAGTTTGTACAAAAAAGCAGGCTATGTCTCACAAGCGGGTGGC | Tb927.8.8270_rev | GGGGACCACTTTGTACAAGAAAGCTGGGTTTATTCATCCCCCACGACCCTC |
| 10_hypothetical | G23 | Tb927.11.16330_for | GGGGACAAGTTTGTACAAAAAAGCAGGCTATGGAAAAGAGAACCAACACGTTCA | Tb927.11.16330_rev | GGGGACCACTTTGTACAAGAAAGCTGGGTTTACTGTGCTGTTTCTTTCTGCTGC |
| 10_hypothetical | G24 | Tb927.8.5400_for | GGGGACAAGTTTGTACAAAAAAGCAGGCTATGTCTTTTCACACCGACGTGTT | Tb927.8.5400_rev | GGGGACCACTTTGTACAAGAAAGCTGGGTTTACGTCGCTGATACCCCTAATCT |
| 10_hypothetical | H1 | Tb927.4.4060_for | GGGGACAAGTTTGTACAAAAAAGCAGGCTATGTCTCAAAAACATATCTTCCAGCGG | Tb927.4.4060_rev | GGGGACCACTTTGTACAAGAAAGCTGGGTCTAACCTCCACTTATCTGACTGCTTTC |
| 10_hypothetical | H2 | Tb927.7.4600_for | GGGGACAAGTTTGTACAAAAAAGCAGGCTATGGTGTCGCGAGCAAAGC | Tb927.7.4600_rev | GGGGACCACTTTGTACAAGAAAGCTGGGTTTACGGAACACAACCAGCGTC |

| | | | | | |
|---|---|---|---|---|---|
| 10_hypothetical | H3 | Tb927.10.13590_for | GGGGACAAGTTTGTACAAAAAAGCAGGCTATGATAGACACAAAGCAGTTGTGCT | Tb927.10.13590_rev | GGGGACCACTTTGTACAAGAAAGCTGGGTTCAGCAGCCTGTGATCCCTC |
| 10_hypothetical | H4 | Tb927.10.1310_for | GGGGACAAGTTTGTACAAAAAAGCAGGCTATGCAGACAGCCCCCCCT | Tb927.10.1310_rev | GGGGACCACTTTGTACAAGAAAGCTGGGTTCACCCCAAAAGACCTTCTGCC |
| 10_hypothetical | H5 | Tb927.3.5290_for | GGGGACAAGTTTGTACAAAAAAGCAGGCTATGGGTCGGGCAACAGAGG | Tb927.3.5290_rev | GGGGACCACTTTGTACAAGAAAGCTGGGTTTAACTTCTACACACCCACGATAGTCC |
| 10_hypothetical | H6 | Tb927.10.3310_for | GGGGACAAGTTTGTACAAAAAAGCAGGCTATGAGTACCGAGAACGAGGGC | Tb927.10.3310_rev | GGGGACCACTTTGTACAAGAAAGCTGGGTTCACCGAGTTAGCGCAAAACC |
| 10_hypothetical | H7 | Tb927.4.4480_for | GGGGACAAGTTTGTACAAAAAAGCAGGCTATGAGCACGCGGTGCGCT | Tb927.4.4480_rev | GGGGACCACTTTGTACAAGAAAGCTGGGTTCATCCCTCATGTTGCCGAAGG |
| 10_hypothetical | H8 | Tb927.8.3920_for | GGGGACAAGTTTGTACAAAAAAGCAGGCTATGTCTGTTGAGGAAGTTGATACATTACTT | Tb927.8.3920_rev | GGGGACCACTTTGTACAAGAAAGCTGGGTCTACTTTAATTTATACCACTGATGCCGGC |
| 10_hypothetical | H9 | Tb927.8.6260_for | GGGGACAAGTTTGTACAAAAAAGCAGGCTATGATAGAAGTAGTGAAATCAACCACAAAC | Tb927.8.6260_rev | GGGGACCACTTTGTACAAGAAAGCTGGGTTCAGCGGTATGGCCGGCG |
| 10_hypothetical | H11 | Tb927.11.13520_for | GGGGACAAGTTTGTACAAAAAAGCAGGCTATGGGCTCTGGGTTTGTTACTAGG | Tb927.11.13520_rev | GGGGACCACTTTGTACAAGAAAGCTGGGTCTATTGTTCAGGTTTGCCGGTATGA |
| 10_hypothetical | H12 | Tb927.7.2390_for | GGGGACAAGTTTGTACAAAAAAGCAGGCTATGTATCGGCCTCGTGGCG | Tb927.7.2390_rev | GGGGACCACTTTGTACAAGAAAGCTGGGTTTACTTTATAAGCTGCCGAAGAATAAGCG |
| 10_hypothetical | H13 | Tb927.10.2480_for | GGGGACAAGTTTGTACAAAAAAGCAGGCTATGCAGTCTTGTATGATGCATATGGAA | Tb927.10.2480_rev | GGGGACCACTTTGTACAAGAAAGCTGGGTTCAAGCCCACGTCCGCGT |
| 10_hypothetical | H14 | Tb927.4.4760_for | GGGGACAAGTTTGTACAAAAAAGCAGGCTATGAAAACGAGTCTCCCGCCT | Tb927.4.4760_rev | GGGGACCACTTTGTACAAGAAAGCTGGGTTTACGCTCTGTAAACAAACTGCTCC |
| 10_hypothetical | H15 | Tb927.7.5090_for | GGGGACAAGTTTGTACAAAAAAGCAGGCTATGCAACACACCGAGGCG | Tb927.7.5090_rev | GGGGACCACTTTGTACAAGAAAGCTGGGTCTAGTCTGCATTCACCATGGTCTC |
| 10_hypothetical | H16 | Tb927.11.15890_for | GGGGACAAGTTTGTACAAAAAAGCAGGCTATGACCTTTAGTAGCGCCAAACG | Tb927.11.15890_rev | GGGGACCACTTTGTACAAGAAAGCTGGGTTTACCTGATTTTACGAAGGGACTGTTC |
| 10_hypothetical | H17 | Tb927.11.6460_for | GGGGACAAGTTTGTACAAAAAAGCAGGCTATGTCTCTAGTAGGAGACGTACTTCAAC | Tb927.11.6460_rev | GGGGACCACTTTGTACAAGAAAGCTGGGTTTATTCAAAACGCGGATCATCAACCG |
| 10_hypothetical | H18 | Tb927.2.2950_for | GGGGACAAGTTTGTACAAAAAAGCAGGCTATGCGTGCTTCCAGTGCAA | Tb927.2.2950_rev | GGGGACCACTTTGTACAAGAAAGCTGGGTTCACGCGCCTCCACCATC |
| 10_hypothetical | H19 | Tb927.7.3130_for | GGGGACAAGTTTGTACAAAAAAGCAGGCTATGGACGTGGCAGCATTACG | Tb927.7.3130_rev | GGGGACCACTTTGTACAAGAAAGCTGGGTTCAGCCACCCTTACCCTTCTG |
| 10_hypothetical | H20 | Tb927.11.5640_for | GGGGACAAGTTTGTACAAAAAAGCAGGCTATGCAGCGCTCACCTCAATC | Tb927.11.5640_rev | GGGGACCACTTTGTACAAGAAAGCTGGGTTCAGTGCTGCGCTACCCT |
| 10_hypothetical | H21 | Tb927.10.9280_for | GGGGACAAGTTTGTACAAAAAAGCAGGCTATGGTATCCGCTTCGAAGTTGC | Tb927.10.9280_rev | GGGGACCACTTTGTACAAGAAAGCTGGGTCTAGTAGTAGAGGACCGTCCCGT |
| 10_hypothetical | H22 | Tb927.7.980_for | GGGGACAAGTTTGTACAAAAAAGCAGGCTATGAAAGTCGCAGGTTCAATTCTCG | Tb927.7.980_rev | GGGGACCACTTTGTACAAGAAAGCTGGGTTTAAAGACCCTGTGCATGTAGAATCTC |
| 10_hypothetical | H23 | Tb927.10.12950_for | GGGGACAAGTTTGTACAAAAAAGCAGGCTATGGGGGAGGCCATGGGC | Tb927.10.12950_rev | GGGGACCACTTTGTACAAGAAAGCTGGGTTCACAGGTAGCCATAGTGCCG |
| 10_hypothetical | H24 | Tb927.11.15940_for | GGGGACAAGTTTGTACAAAAAAGCAGGCTATGAGCCGGCCTGATAAGAGAC | Tb927.11.15940_rev | GGGGACCACTTTGTACAAGAAAGCTGGGTTCACTCCCACGGTGGCAG |
| 10_hypothetical | I1 | Tb927.11.10210_for | GGGGACAAGTTTGTACAAAAAAGCAGGCTATGAATCCATCTGGTCCCAGTGG | Tb927.11.10210_rev | GGGGACCACTTTGTACAAGAAAGCTGGGTCTACTCTGCCTTCCTAGCCTTATCC |
| 10_hypothetical | I2 | Tb927.5.670_for | GGGGACAAGTTTGTACAAAAAAGCAGGCTATGCAAGCAGCAAACATTTCGG | Tb927.5.670_rev | GGGGACCACTTTGTACAAGAAAGCTGGGTTTACTCCACTGCTGAGATGAGTCTC |
| 10_hypothetical | I3 | Tb927.11.16700_for | GGGGACAAGTTTGTACAAAAAAGCAGGCTATGTTAACTCGCGTGATTCATGACT | Tb927.11.16700_rev | GGGGACCACTTTGTACAAGAAAGCTGGGTTTATAGGTGGAACTCCTCTACGTCCT |
| 10_hypothetical | I4 | Tb927.3.2490_for | GGGGACAAGTTTGTACAAAAAAGCAGGCTATGGATGCGAAACTTGTGTTCTGT | Tb927.3.2490_rev | GGGGACCACTTTGTACAAGAAAGCTGGGTTCAGTACGGGAGCGGTATGAC |
| 10_hypothetical | I5 | Tb927.9.11890_for | GGGGACAAGTTTGTACAAAAAAGCAGGCTATGCTGACGAACTCTTCTGCCG | Tb927.9.11890_rev | GGGGACCACTTTGTACAAGAAAGCTGGGTTTACACCACGAGAGCACCGC |
| 10_hypothetical | I6 | Tb927.7.7050_for | GGGGACAAGTTTGTACAAAAAAGCAGGCTATGGGCGCACACTCAAAAAGG | Tb927.7.7050_rev | GGGGACCACTTTGTACAAGAAAGCTGGGTTCACCTCATGAACTGCTTTTTGTAGT |
| 10_hypothetical | I7 | Tb927.5.710_for | GGGGACAAGTTTGTACAAAAAAGCAGGCTATGGACAGCGTAGGGGATAAGG | Tb927.5.710_rev | GGGGACCACTTTGTACAAGAAAGCTGGGTTCAGTTCGGTCTCACCCCC |
| 10_hypothetical | I8 | Tb927.11.14180_for | GGGGACAAGTTTGTACAAAAAAGCAGGCTATGAAGACGGGTGCGTCCT | Tb927.11.14180_rev | GGGGACCACTTTGTACAAGAAAGCTGGGTTTACAGGTGAGTGGCATATGGCA |
| 10_hypothetical | I9 | Tb927.9.9310_for | GGGGACAAGTTTGTACAAAAAAGCAGGCTATGTTGCGAGGGTGGCATCC | Tb927.9.9310_rev | GGGGACCACTTTGTACAAGAAAGCTGGGTCTACGACCAAAGCTCCTCCACG |
| 10_hypothetical | I10 | Tb927.4.2920_for | GGGGACAAGTTTGTACAAAAAAGCAGGCTATGACTTGTACGCCGCCCG | Tb927.4.2920_rev | GGGGACCACTTTGTACAAGAAAGCTGGGTCTACAGTCGAATGTCCTTTTGGAGG |
| 10_hypothetical | I11 | Tb927.10.2060_for | GGGGACAAGTTTGTACAAAAAAGCAGGCTATGTTGTCGAACACGGGAGGT | Tb927.10.2060_rev | GGGGACCACTTTGTACAAGAAAGCTGGGTTCACGTTCCGCCCAACTCT |
| 10_hypothetical | I12 | Tb927.11.6830_for | GGGGACAAGTTTGTACAAAAAAGCAGGCTATGTTCCGTAAGAAGGCCCACT | Tb927.11.6830_rev | GGGGACCACTTTGTACAAGAAAGCTGGGTTCATAGCCGGTATTCGCGCG |
| 10_hypothetical | I13 | Tb927.7.1460_for | GGGGACAAGTTTGTACAAAAAAGCAGGCTATGGACGTGGTGTTGGAGGT | Tb927.7.1460_rev | GGGGACCACTTTGTACAAGAAAGCTGGGTCTAAGAAATCGTGCATACGTCAAACAAT |
| 10_hypothetical | I14 | Tb927.7.6390_for | GGGGACAAGTTTGTACAAAAAAGCAGGCTATGAAAGCGCACACTGTCGC | Tb927.7.6390_rev | GGGGACCACTTTGTACAAGAAAGCTGGGTCTAAATCGGCATTTGGGTGGGA |
| 10_hypothetical | I15 | Tb927.8.3540_for | GGGGACAAGTTTGTACAAAAAAGCAGGCTATGAGCTATACTAATCAGTGCGTGAGA | Tb927.8.3540_rev | GGGGACCACTTTGTACAAGAAAGCTGGGTTCATTTGCTTTCGATTGAGGCGT |
| 10_hypothetical | I16 | Tb927.4.1530_for | GGGGACAAGTTTGTACAAAAAAGCAGGCTATGTGGGGCGGTGTGAAGG | Tb927.4.1530_rev | GGGGACCACTTTGTACAAGAAAGCTGGGTTCACCTCTGTAAGTGTCTCTTGAACG |
| 10_hypothetical | I17 | Tb927.7.6810_for | GGGGACAAGTTTGTACAAAAAAGCAGGCTATGAAAAATGAAAGGGTAGTTAGTGCATGG | Tb927.7.6810_rev | GGGGACCACTTTGTACAAGAAAGCTGGGTTTATTCGTTTGCGGCAACCTGA |
| 10_hypothetical | I18 | Tb927.7.5050_for | GGGGACAAGTTTGTACAAAAAAGCAGGCTATGCACGACGGGTCGCAG | Tb927.7.5050_rev | GGGGACCACTTTGTACAAGAAAGCTGGGTTCATGACTGATCATGTGTGGGTATTCC |
| 10_hypothetical | I19 | Tb927.11.16230_for | GGGGACAAGTTTGTACAAAAAAGCAGGCTATGGAGGATGAGCAGAGTAGCC | Tb927.11.16230_rev | GGGGACCACTTTGTACAAGAAAGCTGGGTTTAACACAGTGCGGGGATATTCTCA |
| 10_hypothetical | I20 | Tb927.11.12640_for | GGGGACAAGTTTGTACAAAAAAGCAGGCTATGAGCTCTGCGGTTTCAGC | Tb927.11.12640_rev | GGGGACCACTTTGTACAAGAAAGCTGGGTTCATTCATTCAACTCCTTTTCCTCAGC |
| 10_hypothetical | I21 | Tb927.11.7590_for | GGGGACAAGTTTGTACAAAAAAGCAGGCTATGTGGCACTCTTTTCTGCGG | Tb927.11.7590_rev | GGGGACCACTTTGTACAAGAAAGCTGGGTTTACACTTCCACTTGGGCCCC |
| 10_hypothetical | I22 | Tb927.11.10480_for | GGGGACAAGTTTGTACAAAAAAGCAGGCTATGGAGAGCTTCTCCCCATTTCG | Tb927.11.10480_rev | GGGGACCACTTTGTACAAGAAAGCTGGGTTTAGAAATGTGACCCCGAAAAAGTACC |
| 10_hypothetical | I23 | Tb927.10.11130_for | GGGGACAAGTTTGTACAAAAAAGCAGGCTATGTCATGGGTGGAAGACAAGTCA | Tb927.10.11130_rev | GGGGACCACTTTGTACAAGAAAGCTGGGTCTACAGCCCCACACTGGCT |
| 10_hypothetical | I24 | Tb927.10.9670_for | GGGGACAAGTTTGTACAAAAAAGCAGGCTATGGAGCAGGCACAGCGG | Tb927.10.9670_rev | GGGGACCACTTTGTACAAGAAAGCTGGGTCTATGTCGTTGTGTATTCAACTCCAGA |
| 10_hypothetical | J1 | Tb927.2.2090_for | GGGGACAAGTTTGTACAAAAAAGCAGGCTATGACAATGAAACCCGAAATGGATGA | Tb927.2.2090_rev | GGGGACCACTTTGTACAAGAAAGCTGGGTTCAGCTCCCCGTGTGTGT |
| 10_hypothetical | J2 | Tb927.6.900_for | GGGGACAAGTTTGTACAAAAAAGCAGGCTATGGCAATGGACGAGGTGGA | Tb927.6.900_rev | GGGGACCACTTTGTACAAGAAAGCTGGGTCTATGCCTGGAAAATGTATCGGACA |
| 10_hypothetical | J3 | Tb927.10.7540_for | GGGGACAAGTTTGTACAAAAAAGCAGGCTATGACAGGTGACGTCTGCCC | Tb927.10.7540_rev | GGGGACCACTTTGTACAAGAAAGCTGGGTTTATACAGGAGGGTGCGCTCC |
| 10_hypothetical | J4 | Tb927.7.650_for | GGGGACAAGTTTGTACAAAAAAGCAGGCTATGACGGAGCGCTCGCTG | Tb927.7.650_rev | GGGGACCACTTTGTACAAGAAAGCTGGGTCTAACCATCCGCTTTAGCTCGC |
| 10_hypothetical | J5 | Tb927.9.6110_for | GGGGACAAGTTTGTACAAAAAAGCAGGCTATGATGCAACGTGATACTGAGGGG | Tb927.9.6110_rev | GGGGACCACTTTGTACAAGAAAGCTGGGTTTACCTGTTTGCTTTTCGTGAATCTTTG |
| 10_hypothetical | J6 | Tb927.5.1330_for | GGGGACAAGTTTGTACAAAAAAGCAGGCTATGCCTCTTCAACCATCGTACTTAATG | Tb927.5.1330_rev | GGGGACCACTTTGTACAAGAAAGCTGGGTTCATCGGGCGTTAACAAACTCC |
| 10_hypothetical | J7 | Tb927.9.9640_for | GGGGACAAGTTTGTACAAAAAAGCAGGCTATGAGCAACAAACAAGTGGACGG | Tb927.9.9640_rev | GGGGACCACTTTGTACAAGAAAGCTGGGTTCAGCGCGATTGAAACTTACTTGC |
| 10_hypothetical | J8 | Tb927.11.14240_for | GGGGACAAGTTTGTACAAAAAAGCAGGCTATGACTAGCTTAAATGGCAGCAAGAG | Tb927.11.14240_rev | GGGGACCACTTTGTACAAGAAAGCTGGGTTTACACATTTTCTGCAACATACTTGTCG |
| 10_hypothetical | J9 | Tb927.8.7850_for | GGGGACAAGTTTGTACAAAAAAGCAGGCTATGAATCGCTTCCAACGGAGTG | Tb927.8.7850_rev | GGGGACCACTTTGTACAAGAAAGCTGGGTCTAAAGTTCCTGCGAGTTCACATTATCA |
| 10_hypothetical | J10 | Tb927.1.3290_for | GGGGACAAGTTTGTACAAAAAAGCAGGCTATGGGATGTGCATGCATCCG | Tb927.1.3290_rev | GGGGACCACTTTGTACAAGAAAGCTGGGTTCAGACACTTGACGCACTAATTGAA |
| 10_hypothetical | J11 | Tb927.10.4490_for | GGGGACAAGTTTGTACAAAAAAGCAGGCTATGCCAAAAACAGCGGTGCT | Tb927.10.4490_rev | GGGGACCACTTTGTACAAGAAAGCTGGGTCTACATGATATAGACAATAACATTCATCTT |
| 10_hypothetical | J12 | Tb927.11.11140_for | GGGGACAAGTTTGTACAAAAAAGCAGGCTATGTCAGCTACGAGTTTTCATTTCGC | Tb927.11.11140_rev | GGGGACCACTTTGTACAAGAAAGCTGGGTTCACGCCTTTCTTGCTTTGAGAG |
| 10_hypothetical | J13 | Tb927.7.6740_for | GGGGACAAGTTTGTACAAAAAAGCAGGCTATGATGGACATAGATTGCGCGG | Tb927.7.6740_rev | GGGGACCACTTTGTACAAGAAAGCTGGGTTCAAAAGTCATTCAAGGATGCGGG |
| 10_hypothetical | J14 | Tb927.10.10680_for | GGGGACAAGTTTGTACAAAAAAGCAGGCTATGCTGAGGAAACCGTCAGGG | Tb927.10.10680_rev | GGGGACCACTTTGTACAAGAAAGCTGGGTTCACACAAGCGGAATGGAAAGC |
| 10_hypothetical | J15 | Tb927.7.3850_for | GGGGACAAGTTTGTACAAAAAAGCAGGCTATGCCATTTCCCCAAACACCAC | Tb927.7.3850_rev | GGGGACCACTTTGTACAAGAAAGCTGGGTCTACACTCGCCGAGGCGC |
| 10_hypothetical | J16 | Tb927.5.840_for | GGGGACAAGTTTGTACAAAAAAGCAGGCTATGCCTAAGAAGGATCTATTTTCCTCATCA | Tb927.5.840_rev | GGGGACCACTTTGTACAAGAAAGCTGGGTTTACTCGTGACGAGGCTTAGACG |
| 10_hypothetical | J17 | Tb927.9.12070_for | GGGGACAAGTTTGTACAAAAAAGCAGGCTATGGCTAGTGCCCCCCGC | Tb927.9.12070_rev | GGGGACCACTTTGTACAAGAAAGCTGGGTTTAATCATCGTCATCATCCTCAATGAGTG |
| 10_hypothetical | J18 | Tb927.7.4450_for | GGGGACAAGTTTGTACAAAAAAGCAGGCTATGCTCAGCCAAGATGCCAGA | Tb927.7.4450_rev | GGGGACCACTTTGTACAAGAAAGCTGGGTTTAAAAAAGTGATTCACCGCCGC |
| 10_hypothetical | J19 | Tb927.11.15610_for | GGGGACAAGTTTGTACAAAAAAGCAGGCTATGAAGCAGCAAACGCCGG | Tb927.11.15610_rev | GGGGACCACTTTGTACAAGAAAGCTGGGTCTATCCATCTATTTTCCTGTTAGCAAGCT |
| 10_hypothetical | J20 | Tb927.10.1150_for | GGGGACAAGTTTGTACAAAAAAGCAGGCTATGCTACCGATGGCATCCAACC | Tb927.10.1150_rev | GGGGACCACTTTGTACAAGAAAGCTGGGTTTACCTGCTCGCACGCAAC |
| 10_hypothetical | J21 | Tb927.4.2820_for | GGGGACAAGTTTGTACAAAAAAGCAGGCTATGAATCTCGTGTATGGTTCCCACC | Tb927.4.2820_rev | GGGGACCACTTTGTACAAGAAAGCTGGGTTCAAAGCATACGAACACTTACGCG |
| 10_hypothetical | J22 | Tb927.5.1890_for | GGGGACAAGTTTGTACAAAAAAGCAGGCTATGCGGAGTTCCGACTCACA | Tb927.5.1890_rev | GGGGACCACTTTGTACAAGAAAGCTGGGTTCACCACTTGTAGACTAGTGTCTCAA |
| 10_hypothetical | J23 | Tb927.11.7120_for | GGGGACAAGTTTGTACAAAAAAGCAGGCTATGGGGCGGATAGAGGACCT | Tb927.11.7120_rev | GGGGACCACTTTGTACAAGAAAGCTGGGTCTATAAAGTTCGCTTTAGCGCGGC |
| 10_hypothetical | J24 | Tb927.9.11770_for | GGGGACAAGTTTGTACAAAAAAGCAGGCTATGACAACCCTTAAAGGTGGTGGT | Tb927.9.11770_rev | GGGGACCACTTTGTACAAGAAAGCTGGGTTTAAGCGGTTTTCCTCATGTGATCC |
| 10_hypothetical | K1 | Tb927.11.11650_for | GGGGACAAGTTTGTACAAAAAAGCAGGCTATGGGGGAGCCAGGGGTT | Tb927.11.11650_rev | GGGGACCACTTTGTACAAGAAAGCTGGGTTCAGTCCCTTCCCAAGTGCC |
| 10_hypothetical | K2 | Tb927.8.7360_for | GGGGACAAGTTTGTACAAAAAAGCAGGCTATGCTTCTGTGCCGTCCCG | Tb927.8.7360_rev | GGGGACCACTTTGTACAAGAAAGCTGGGTTCAACGCTCAGTGTCGCTCA |
| 10_hypothetical | K3 | Tb927.5.3540_for | GGGGACAAGTTTGTACAAAAAAGCAGGCTATGAATGCCATCCCACTCGTCC | Tb927.5.3540_rev | GGGGACCACTTTGTACAAGAAAGCTGGGTTTAAATAAAGAACCGATCGTCTATAAGAGT |
| 10_hypothetical | K4 | Tb927.3.1550_for | GGGGACAAGTTTGTACAAAAAAGCAGGCTATGAGTTCAATGGAATTAGCGGATGT | Tb927.3.1550_rev | GGGGACCACTTTGTACAAGAAAGCTGGGTCTAAAGTACTTCCTCGGCCAGCG |
| 10_hypothetical | K5 | Tb927.8.4460_for | GGGGACAAGTTTGTACAAAAAAGCAGGCTATGCTAGAAACTCATTGTCAGTTCAAACA | Tb927.8.4460_rev | GGGGACCACTTTGTACAAGAAAGCTGGGTTCACGTCCTTTCCTTCACACTACT |
| 10_hypothetical | K6 | Tb927.10.7580_for | GGGGACAAGTTTGTACAAAAAAGCAGGCTATGAGTGTCGAAAGGATCGTGC | Tb927.10.7580_rev | GGGGACCACTTTGTACAAGAAAGCTGGGTTTACCGATCACCCTCCTGACC |
| 10_hypothetical | K7 | Tb927.3.3050_for | GGGGACAAGTTTGTACAAAAAAGCAGGCTATGGCGCCGGCAGTAATG | Tb927.3.3050_rev | GGGGACCACTTTGTACAAGAAAGCTGGGTTTATTCATCACACGCTTGTCGTAGC |
| 10_hypothetical | K8 | Tb927.10.12820_for | GGGGACAAGTTTGTACAAAAAAGCAGGCTATGGGAGCCAAACCAGAATCTTTG | Tb927.10.12820_rev | GGGGACCACTTTGTACAAGAAAGCTGGGTTTACTTGGTTACCTTTCGTTTGATGCG |
| 10_hypothetical | K9 | Tb927.4.2580_for | GGGGACAAGTTTGTACAAAAAAGCAGGCTATGCATAATCGTCTATACCACAGGGT | Tb927.4.2580_rev | GGGGACCACTTTGTACAAGAAAGCTGGGTTTACTCGTGCTGCAGTCGTTC |
| 10_hypothetical | K10 | Tb927.7.7290_for | GGGGACAAGTTTGTACAAAAAAGCAGGCTATGGAGCAATATCAATCCCAATCCCA | Tb927.7.7290_rev | GGGGACCACTTTGTACAAGAAAGCTGGGTCTACAATCGAAGGCGGTCAATATCC |
| 10_hypothetical | K11 | Tb927.11.3500_for | GGGGACAAGTTTGTACAAAAAAGCAGGCTATGAGCGTAGAAGAAGTGTCATGTC | Tb927.11.3500_rev | GGGGACCACTTTGTACAAGAAAGCTGGGTTTACAGTTCAAAACACGAATCATCGTG |
| 10_hypothetical | K12 | Tb927.7.6590_for | GGGGACAAGTTTGTACAAAAAAGCAGGCTATGAATCTAGCGAAGAAAATCATCGTGT | Tb927.7.6590_rev | GGGGACCACTTTGTACAAGAAAGCTGGGTTTATTGCACAGACACACCCTCG |
| 10_hypothetical | K13 | Tb927.9.4530_for | GGGGACAAGTTTGTACAAAAAAGCAGGCTATGAAGCGGGATGAACTGTTTGC | Tb927.9.4530_rev | GGGGACCACTTTGTACAAGAAAGCTGGGTCTATGCCTCCACTGGAGCTCC |
| 10_hypothetical | K14 | Tb927.7.2350_for | GGGGACAAGTTTGTACAAAAAAGCAGGCTATGTCCGATGACGCTAAAACCAC | Tb927.7.2350_rev | GGGGACCACTTTGTACAAGAAAGCTGGGTCTAATCTTCTCTTTCCGTTTTCGCATC |
| 10_hypothetical | K15 | Tb927.11.2310_for | GGGGACAAGTTTGTACAAAAAAGCAGGCTATGTATAAGCAGCACACGAACTTTGC | Tb927.11.2310_rev | GGGGACCACTTTGTACAAGAAAGCTGGGTCTAGGCATCGTAATCATCAAAAATAATGTC |
| 10_hypothetical | K16 | Tb927.9.6530_for | GGGGACAAGTTTGTACAAAAAAGCAGGCTATGGGGAACGCACAGGACA | Tb927.9.6530_rev | GGGGACCACTTTGTACAAGAAAGCTGGGTTTACTCCCGGATAATGGTGGCG |
| 10_hypothetical | K17 | Tb927.10.3780_for | GGGGACAAGTTTGTACAAAAAAGCAGGCTATGATATCTCGAAAGGACACCGGG | Tb927.10.3780_rev | GGGGACCACTTTGTACAAGAAAGCTGGGTCTAACTAGCCACAACATCAAAAAGCAAC |
| 10_hypothetical | K18 | Tb927.9.12770_for | GGGGACAAGTTTGTACAAAAAAGCAGGCTATGAAACTACTGCGACGAGGGA | Tb927.9.12770_rev | GGGGACCACTTTGTACAAGAAAGCTGGGTTTAATGTGCACGATCATCTGCACC |
| 10_hypothetical | K19 | Tb927.3.5620_for | GGGGACAAGTTTGTACAAAAAAGCAGGCTATGTCGCGTGTTACCCTTTCC | Tb927.3.5620_rev | GGGGACCACTTTGTACAAGAAAGCTGGGTCTAAAACCGGCGCGGTGG |
| 10_hypothetical | K20 | Tb927.10.16200_for | GGGGACAAGTTTGTACAAAAAAGCAGGCTATGCCGGCCACCTCAACG | Tb927.10.16200_rev | GGGGACCACTTTGTACAAGAAAGCTGGGTTCACAATTCGTAGTGCTCTCGTCG |
| 10_hypothetical | K21 | Tb927.10.9970_for | GGGGACAAGTTTGTACAAAAAAGCAGGCTATGATGGCCAACAAGCGGTT | Tb927.10.9970_rev | GGGGACCACTTTGTACAAGAAAGCTGGGTTCACGCTGATGCTCCCCC |
| 10_hypothetical | K22 | Tb927.3.3800_for | GGGGACAAGTTTGTACAAAAAAGCAGGCTATGGATTGCAAGTTGCAACATGTTC | Tb927.3.3800_rev | GGGGACCACTTTGTACAAGAAAGCTGGGTTTAACAAGTTCGCTTCATGTTACACCG |
| 10_hypothetical | K23 | Tb927.1.2990_for | GGGGACAAGTTTGTACAAAAAAGCAGGCTATGTTAACCCCGACACGGCG | Tb927.1.2990_rev | GGGGACCACTTTGTACAAGAAAGCTGGGTCTATTTATTTCTGTGACGACTTCGTACCT |
| 10_hypothetical | K24 | Tb927.8.780_for | GGGGACAAGTTTGTACAAAAAAGCAGGCTATGGGAGACTGCGTTGCTTCA | Tb927.8.780_rev | GGGGACCACTTTGTACAAGAAAGCTGGGTCTACACAACTGGCGCAACCG |
| 10_hypothetical | L1 | Tb927.11.12460_for | GGGGACAAGTTTGTACAAAAAAGCAGGCTATGTTTTACGCCTCGGAGGAAGG | Tb927.11.12460_rev | GGGGACCACTTTGTACAAGAAAGCTGGGTTCAAGCGACCACTCCCCTG |
| 10_hypothetical | L2 | Tb927.3.1670_for | GGGGACAAGTTTGTACAAAAAAGCAGGCTATGGCGTATTCCACTCGAGTAATACG | Tb927.3.1670_rev | GGGGACCACTTTGTACAAGAAAGCTGGGTCTACTGCAAACTCATTTCCTCTTCCC |
| 10_hypothetical | L3 | Tb927.1.4760_for | GGGGACAAGTTTGTACAAAAAAGCAGGCTATGTCGTCTTACGAGGCCTTTCG | Tb927.1.4760_rev | GGGGACCACTTTGTACAAGAAAGCTGGGTTTACAACACTTCCCGCAATAGGAC |
| 10_hypothetical | L4 | Tb927.7.3090_for | GGGGACAAGTTTGTACAAAAAAGCAGGCTATGTGGAATGAAGTTAGCGCTAGTG | Tb927.7.3090_rev | GGGGACCACTTTGTACAAGAAAGCTGGGTTTACGGCAGTGACGCATCCC |
| 10_hypothetical | L5 | Tb927.8.4900_for | GGGGACAAGTTTGTACAAAAAAGCAGGCTATGTCTTCGGAGGCGGTAAAAAG | Tb927.8.4900_rev | GGGGACCACTTTGTACAAGAAAGCTGGGTTTAAACTTCTAGACCGACAGCAATATCG |
| 10_hypothetical | L6 | Tb927.4.4710_for | GGGGACAAGTTTGTACAAAAAAGCAGGCTATGACTTCAAGTTGGGGCCCG | Tb927.4.4710_rev | GGGGACCACTTTGTACAAGAAAGCTGGGTCTATTGTCCATTTGCGCCAACCA |
| 10_hypothetical | L7 | Tb927.2.3020_for | GGGGACAAGTTTGTACAAAAAAGCAGGCTATGTCTGATGCGTCCGTTAATGAC | Tb927.2.3020_rev | GGGGACCACTTTGTACAAGAAAGCTGGGTTCACTTACCCTCCTCTCCCGG |
| 10_hypothetical | L8 | Tb927.8.1260_for | GGGGACAAGTTTGTACAAAAAAGCAGGCTATGCGACGAAGGCGACAAC | Tb927.8.1260_rev | GGGGACCACTTTGTACAAGAAAGCTGGGTTCAAAAGAGCGCCTCCACAGC |
| 10_hypothetical | L9 | Tb927.8.4350_for | GGGGACAAGTTTGTACAAAAAAGCAGGCTATGCTATATGATGTTGTGGAGTTGTTGG | Tb927.8.4350_rev | GGGGACCACTTTGTACAAGAAAGCTGGGTTCATGCACTGCAATTGGGTAACA |
| 10_hypothetical | L10 | Tb927.10.12920_for | GGGGACAAGTTTGTACAAAAAAGCAGGCTATGAACTGCGACAACCAAGAAGT | Tb927.10.12920_rev | GGGGACCACTTTGTACAAGAAAGCTGGGTCTATCTTCGGCATAGACGCACTG |
| 10_hypothetical | L11 | Tb927.11.2830_for | GGGGACAAGTTTGTACAAAAAAGCAGGCTATGGGAAAGATTCTCCTTTGTGTACAG | Tb927.11.2830_rev | GGGGACCACTTTGTACAAGAAAGCTGGGTTCATTTACCTGAAGAAACTTCTGACTGAG |
| 10_hypothetical | L12 | Tb927.5.2530_for | GGGGACAAGTTTGTACAAAAAAGCAGGCTATGCCACCGCCCACATCT | Tb927.5.2530_rev | GGGGACCACTTTGTACAAGAAAGCTGGGTTCAGCCGCGACGAGTTTTC |
| 10_hypothetical | L13 | Tb927.2.2210_for | GGGGACAAGTTTGTACAAAAAAGCAGGCTATGTGGAGGAGGCACTGCTG | Tb927.2.2210_rev | GGGGACCACTTTGTACAAGAAAGCTGGGTTTACGGCATGGCCACCGT |
| 10_hypothetical | L14 | Tb927.10.14330_for | GGGGACAAGTTTGTACAAAAAAGCAGGCTATGGTCTTACGCCGTGATGGC | Tb927.10.14330_rev | GGGGACCACTTTGTACAAGAAAGCTGGGTCTACACGCGTGTTGTTATCTCACC |
| 10_hypothetical | L15 | Tb927.7.860_for | GGGGACAAGTTTGTACAAAAAAGCAGGCTATGCCAGTGGAATGGCCCA | Tb927.7.860_rev | GGGGACCACTTTGTACAAGAAAGCTGGGTTTACCCAGCAATGCTGGGAACG |
| 10_hypothetical | L16 | Tb927.11.11280_for | GGGGACAAGTTTGTACAAAAAAGCAGGCTATGAATCCGTTTATTTTACCGCCATTAGA | Tb927.11.11280_rev | GGGGACCACTTTGTACAAGAAAGCTGGGTTTACTTCTCGTAGTATGCCTCGCG |
| 10_hypothetical | L17 | Tb927.11.5060_for | GGGGACAAGTTTGTACAAAAAAGCAGGCTATGTTCTTCAGGTTTCATAGCACAACA | Tb927.11.5060_rev | GGGGACCACTTTGTACAAGAAAGCTGGGTTTAACGGCGTCGGTTCCCT |
| 10_hypothetical | L18 | Tb927.9.2550_for | GGGGACAAGTTTGTACAAAAAAGCAGGCTATGCGTCGTGAGGAACGGC | Tb927.9.2550_rev | GGGGACCACTTTGTACAAGAAAGCTGGGTTTAAGCTGCTGTCCGAGAGGC |
| 10_hypothetical | L19 | Tb927.8.6100_for | GGGGACAAGTTTGTACAAAAAAGCAGGCTATGGAAAAGCTCCAACTGGAGC | Tb927.8.6100_rev | GGGGACCACTTTGTACAAGAAAGCTGGGTTCACTGCGAAGTATCCCAACGA |
| 10_hypothetical | L20 | Tb927.9.10530_for | GGGGACAAGTTTGTACAAAAAAGCAGGCTATGAGCCGACGTGGTGGC | Tb927.9.10530_rev | GGGGACCACTTTGTACAAGAAAGCTGGGTCTACCTCGAAGAATAAGAATAACCGCTG |
| 10_hypothetical | L21 | Tb927.11.15970_for | GGGGACAAGTTTGTACAAAAAAGCAGGCTATGGAGGCTCTCCTGCGG | Tb927.11.15970_rev | GGGGACCACTTTGTACAAGAAAGCTGGGTTCACACAAGATTCATGAGCGGC |
| 10_hypothetical | L22 | Tb927.10.13550_for | GGGGACAAGTTTGTACAAAAAAGCAGGCTATGGATTTCATTGCATTTAAGGATGGTG | Tb927.10.13550_rev | GGGGACCACTTTGTACAAGAAAGCTGGGTTTAGGCAGTTCCTGTGCACTGA |
| 10_hypothetical | L23 | Tb927.8.5220_for | GGGGACAAGTTTGTACAAAAAAGCAGGCTATGCTCAGAACTCGCTTGTCAC | Tb927.8.5220_rev | GGGGACCACTTTGTACAAGAAAGCTGGGTTCAAAAGGGTGAGTGAGGACTGG |
| 10_hypothetical | L24 | Tb927.10.4470_for | GGGGACAAGTTTGTACAAAAAAGCAGGCTATGGATTGCCACTTCCGTCAAC | Tb927.10.4470_rev | GGGGACCACTTTGTACAAGAAAGCTGGGTTTAAACGAGAAGAATGCCCCTTATTCC |
| 10_hypothetical | M1 | Tb927.11.240_for | GGGGACAAGTTTGTACAAAAAAGCAGGCTATGTGGCGTTTGTTATTTCTGCG | Tb927.11.240_rev | GGGGACCACTTTGTACAAGAAAGCTGGGTTTATTCATTACCCTTTCCTTCGTAAACGA |
| 10_hypothetical | M2 | Tb927.9.8820_for | GGGGACAAGTTTGTACAAAAAAGCAGGCTATGAAAGCTGCAAGAAGTAACGGA | Tb927.9.8820_rev | GGGGACCACTTTGTACAAGAAAGCTGGGTCTAAACCGCTGAATCTATCAAACCTTCT |
| 10_hypothetical | M3 | Tb927.11.7400_for | GGGGACAAGTTTGTACAAAAAAGCAGGCTATGGAAGACTCTAACGAAGTTGCG | Tb927.11.7400_rev | GGGGACCACTTTGTACAAGAAAGCTGGGTCTAGCTAGACCTTACCAATGTAAATTTATT |
| 10_hypothetical | M4 | Tb927.8.4660_for | GGGGACAAGTTTGTACAAAAAAGCAGGCTATGCTGGGGAGTGTTGATGATGT | Tb927.8.4660_rev | GGGGACCACTTTGTACAAGAAAGCTGGGTTCATGCCAAACAGCGGGTTAC |
| 10_hypothetical | M5 | Tb927.11.3410_for | GGGGACAAGTTTGTACAAAAAAGCAGGCTATGAACAAGGCGGGTGTGTATTC | Tb927.11.3410_rev | GGGGACCACTTTGTACAAGAAAGCTGGGTCTACTCACCGTAAAATCGGTTGAAAGT |
| 10_hypothetical | M6 | Tb927.11.3330_for | GGGGACAAGTTTGTACAAAAAAGCAGGCTATGTGCGGTGTGTCGGGG | Tb927.11.3330_rev | GGGGACCACTTTGTACAAGAAAGCTGGGTTTATCGACCGTTCGCAGAGCT |
| 10_hypothetical | M7 | Tb927.10.2920_for | GGGGACAAGTTTGTACAAAAAAGCAGGCTATGACCAACCCACCCCAAGAC | Tb927.10.2920_rev | GGGGACCACTTTGTACAAGAAAGCTGGGTCTACTTTTTGACATTCCGTGGTGAAAC |
| 10_hypothetical | M8 | Tb927.2.5900_for | GGGGACAAGTTTGTACAAAAAAGCAGGCTATGTGTTGGTGCACTACGCAA | Tb927.2.5900_rev | GGGGACCACTTTGTACAAGAAAGCTGGGTTCAACAGTCTGCAGCAAAAAGCA |
| 10_hypothetical | M9 | Tb927.2.2580_for | GGGGACAAGTTTGTACAAAAAAGCAGGCTATGCCTCGAAAGAGGAATCCGT | Tb927.2.2580_rev | GGGGACCACTTTGTACAAGAAAGCTGGGTCTACGATTGCGCGCTTGGT |
| 10_hypothetical | M10 | Tb927.2.3990_for | GGGGACAAGTTTGTACAAAAAAGCAGGCTATGTGGGACCACATTAATGGTGTTAC | Tb927.2.3990_rev | GGGGACCACTTTGTACAAGAAAGCTGGGTTCAAGTTGTCGAAGCACCAACC |
| 10_hypothetical | M11 | Tb927.1.4740_for | GGGGACAAGTTTGTACAAAAAAGCAGGCTATGACCGCAAACTACCACACG | Tb927.1.4740_rev | GGGGACCACTTTGTACAAGAAAGCTGGGTTCATACTTGCCACAACTCCACAAG |
| 10_hypothetical | M12 | Tb927.9.12800_for | GGGGACAAGTTTGTACAAAAAAGCAGGCTATGGACGGCTGCGTCAAC | Tb927.9.12800_rev | GGGGACCACTTTGTACAAGAAAGCTGGGTTTACATACTCTTCCACATGTGTGAACTC |
| 10_hypothetical | M13 | Tb927.3.3110_for | GGGGACAAGTTTGTACAAAAAAGCAGGCTATGCCGCCTAAAAAGAAGCGA | Tb927.3.3110_rev | GGGGACCACTTTGTACAAGAAAGCTGGGTTTACCGGTCCGTCCTGCC |

| | | | | | |
|---|---|---|---|---|---|
| 10_hypothetical | M14 | Tb927.7.5750_for | GGGGACAAGTTTGTACAAAAAAGCAGGCTATGGTTTCACGTGCACCGTC | Tb927.7.5750_rev | GGGGACCACTTTGTACAAGAAAGCTGGGTTTAGGATGTTTTCAGCACCCTCTTAAG |
| 10_hypothetical | M15 | Tb927.3.4970_for | GGGGACAAGTTTGTACAAAAAAGCAGGCTATGGGAACAATAGAAGGCACAACTG | Tb927.3.4970_rev | GGGGACCACTTTGTACAAGAAAGCTGGGTTCAGCGTGCCGTTTTTACTGT |
| 10_hypothetical | M16 | Tb927.7.1660_for | GGGGACAAGTTTGTACAAAAAAGCAGGCTATGGGGGGTAGAAACACGGG | Tb927.7.1660_rev | GGGGACCACTTTGTACAAGAAAGCTGGGTTCAACTCTCTCGTATTTCCTGCACT |
| 10_hypothetical | M17 | Tb927.5.4130_for | GGGGACAAGTTTGTACAAAAAAGCAGGCTATGCTGCACCCCAATACTCGT | Tb927.5.4130_rev | GGGGACCACTTTGTACAAGAAAGCTGGGTTCAACACTTCTGGTTGTTATCCCAC |
| 10_hypothetical | M18 | Tb927.10.2790_for | GGGGACAAGTTTGTACAAAAAAGCAGGCTATGGACAACTCTGAGGCTGTGA | Tb927.10.2790_rev | GGGGACCACTTTGTACAAGAAAGCTGGGTTTATGGGGTGGTGTCCAAAAGGA |
| 10_hypothetical | M19 | Tb927.11.16530_for | GGGGACAAGTTTGTACAAAAAAGCAGGCTATGATCCGTACCACTCATTTGATGTTG | Tb927.11.16530_rev | GGGGACCACTTTGTACAAGAAAGCTGGGTCTATGGCTTGTGTTTCCCGTGA |
| 10_hypothetical | M20 | Tb927.7.6650_for | GGGGACAAGTTTGTACAAAAAAGCAGGCTATGTCGTACAACGCGTCCAC | Tb927.7.6650_rev | GGGGACCACTTTGTACAAGAAAGCTGGGTTCACACCGCATTGACACCCC |
| 10_hypothetical | M21 | Tb927.11.15370_for | GGGGACAAGTTTGTACAAAAAAGCAGGCTATGAACCAAGAGCAGCTGAACC | Tb927.11.15370_rev | GGGGACCACTTTGTACAAGAAAGCTGGGTTCACATCGCTGCAAGTTGATTTTTT |
| 10_hypothetical | M22 | Tb927.7.1220_for | GGGGACAAGTTTGTACAAAAAAGCAGGCTATGGAGCATCAAAACGACCGC | Tb927.7.1220_rev | GGGGACCACTTTGTACAAGAAAGCTGGGTCTAGTAAATACACGATAGCATCAGCGAC |
| 10_hypothetical | M23 | Tb927.7.6580_for | GGGGACAAGTTTGTACAAAAAAGCAGGCTATGCATGTTCTGTGGTGTATTGTGC | Tb927.7.6580_rev | GGGGACCACTTTGTACAAGAAAGCTGGGTTTATTGCACAGACACACCCTCG |
| 10_hypothetical | M24 | Tb927.11.15070_for | GGGGACAAGTTTGTACAAAAAAGCAGGCTATGAGAAGAGCCTTAAGTGGGCT | Tb927.11.15070_rev | GGGGACCACTTTGTACAAGAAAGCTGGGTTTAAGATTTGTTATCATCTCCTGGACGAAG |
| 10_hypothetical | N1 | Tb927.4.3920_for | GGGGACAAGTTTGTACAAAAAAGCAGGCTATGTCCTCTTTTCTACAGGAACTTGGT | Tb927.4.3920_rev | GGGGACCACTTTGTACAAGAAAGCTGGGTCTACATCTCATGAACCGATTCCTCCA |
| 10_hypothetical | N2 | Tb927.6.3220_for | GGGGACAAGTTTGTACAAAAAAGCAGGCTATGCCAGTTCGACAGAAGTATCCA | Tb927.6.3220_rev | GGGGACCACTTTGTACAAGAAAGCTGGGTTCAATCACTAAGTAACTTTGTCTCATCGC |
| 10_hypothetical | N3 | Tb927.3.4600_for | GGGGACAAGTTTGTACAAAAAAGCAGGCTATGAATATCCTACAACTCTTATCGGTTGC | Tb927.3.4600_rev | GGGGACCACTTTGTACAAGAAAGCTGGGTCTACTGAGTTTGCGCTGCAGC |
| 10_hypothetical | N4 | Tb927.4.4560_for | GGGGACAAGTTTGTACAAAAAAGCAGGCTATGGTTGAGGAGCCAAGCGG | Tb927.4.4560_rev | GGGGACCACTTTGTACAAGAAAGCTGGGTTCAGTTATTGCACTTATCCAGCTCCT |
| 10_hypothetical | N5 | Tb927.2.5820_for | GGGGACAAGTTTGTACAAAAAAGCAGGCTATGGGGCGGCAGCGAGCA | Tb927.2.5820_rev | GGGGACCACTTTGTACAAGAAAGCTGGGTTTAAGCCACTGTCGCGTGC |
| 10_hypothetical | N6 | Tb927.11.2590_for | GGGGACAAGTTTGTACAAAAAAGCAGGCTATGTCAACATCCTTTTCCTCCAAAAAAG | Tb927.11.2590_rev | GGGGACCACTTTGTACAAGAAAGCTGGGTTCAGCTTAATCGTGGAACAGCC |
| 10_hypothetical | N7 | Tb927.5.2980_for | GGGGACAAGTTTGTACAAAAAAGCAGGCTATGCCAGCAGGTAGTGATGACG | Tb927.5.2980_rev | GGGGACCACTTTGTACAAGAAAGCTGGGTTTAGACAGCCCGCAGCAGC |
| 10_hypothetical | N8 | Tb927.8.4230_for | GGGGACAAGTTTGTACAAAAAAGCAGGCTATGCTTCGCCGTAATCTTGTTTCT | Tb927.8.4230_rev | GGGGACCACTTTGTACAAGAAAGCTGGGTTTAACGCCGTACTCTCTCGAACC |
| 10_hypothetical | N9 | Tb927.10.720_for | GGGGACAAGTTTGTACAAAAAAGCAGGCTATGCCAAAGAAGAGTGAACCGG | Tb927.10.720_rev | GGGGACCACTTTGTACAAGAAAGCTGGGTCTACTTACTAATTCGTTTCTGAACAGCAGC |
| 10_hypothetical | N10 | Tb927.4.4580_for | GGGGACAAGTTTGTACAAAAAAGCAGGCTATGGGAGCGGTTCCCAGTC | Tb927.4.4580_rev | GGGGACCACTTTGTACAAGAAAGCTGGGTTCAGTGCTGTAAACCGTGCC |
| 10_hypothetical | N11 | Tb927.8.3490_for | GGGGACAAGTTTGTACAAAAAAGCAGGCTATGAACTTCTTATCCCAGATTTACACCG | Tb927.8.3490_rev | GGGGACCACTTTGTACAAGAAAGCTGGGTCTACGGCGTCAAAACGCAGA |
| 10_hypothetical | N12 | Tb927.5.2800_for | GGGGACAAGTTTGTACAAAAAAGCAGGCTATGTCAGTAGATACCATTCAGCTTGTG | Tb927.5.2800_rev | GGGGACCACTTTGTACAAGAAAGCTGGGTTCATTCACGTAGTCGTTTTGTTGTGC |
| 10_hypothetical | N13 | Tb927.10.13880_for | GGGGACAAGTTTGTACAAAAAAGCAGGCTATGATGGATCAGATGAATATTGGGGTGC | Tb927.10.13880_rev | GGGGACCACTTTGTACAAGAAAGCTGGGTTTACCAAGCCGATAGAGTTTGAACC |
| 10_hypothetical | N14 | Tb927.10.10090_for | GGGGACAAGTTTGTACAAAAAAGCAGGCTATGTCCGTTGCGCTCGAC | Tb927.10.10090_rev | GGGGACCACTTTGTACAAGAAAGCTGGGTCTACCGCGAACAGCGTTGT |
| 10_hypothetical | N15 | Tb927.8.840_for | GGGGACAAGTTTGTACAAAAAAGCAGGCTATGCAGTGCGAGAGATTCAAACA | Tb927.8.840_rev | GGGGACCACTTTGTACAAGAAAGCTGGGTTCAACTCAATGTTATATTAATGCCGCGC |
| 10_hypothetical | N16 | Tb927.10.12870_for | GGGGACAAGTTTGTACAAAAAAGCAGGCTATGACCGAAGCAACCAATGAGG | Tb927.10.12870_rev | GGGGACCACTTTGTACAAGAAAGCTGGGTTTAGTGTTTTTGTTTTCTACGCTTCCG |
| 10_hypothetical | N17 | Tb927.9.2590_for | GGGGACAAGTTTGTACAAAAAAGCAGGCTATGCAGACTAGCAGTGTCTCGG | Tb927.9.2590_rev | GGGGACCACTTTGTACAAGAAAGCTGGGTTTAGGTTCCGAGCCGGCG |
| 10_hypothetical | N18 | Tb927.10.8600_for | GGGGACAAGTTTGTACAAAAAAGCAGGCTATGTCAACAGGTCACCCCGT | Tb927.10.8600_rev | GGGGACCACTTTGTACAAGAAAGCTGGGTCTACGCCTGACCAAAGAACCATG |
| 10_hypothetical | N19 | Tb927.7.5610_for | GGGGACAAGTTTGTACAAAAAAGCAGGCTATGAAGTCAGGAATATCTAGTCGGTCG | Tb927.7.5610_rev | GGGGACCACTTTGTACAAGAAAGCTGGGTTTACCGTGTAGAAACCCTATTGTTCAC |
| 10_hypothetical | N20 | Tb927.10.5170_for | GGGGACAAGTTTGTACAAAAAAGCAGGCTATGCCTACGGCCGATTCAAC | Tb927.10.5170_rev | GGGGACCACTTTGTACAAGAAAGCTGGGTTCAGCGCCGCTTGAATGG |
| 10_hypothetical | N21 | Tb927.11.11810_for | GGGGACAAGTTTGTACAAAAAAGCAGGCTATGAGCAGGGCACTTTTATCTAGGA | Tb927.11.11810_rev | GGGGACCACTTTGTACAAGAAAGCTGGGTTTAAGACGTGGCGCATCCAC |
| 10_hypothetical | N22 | Tb927.8.2580_for | GGGGACAAGTTTGTACAAAAAAGCAGGCTATGCCGAACCATGGAAAGAGTAAG | Tb927.8.2580_rev | GGGGACCACTTTGTACAAGAAAGCTGGGTTCACTTCACAACATCACGAAACTTGT |
| 10_hypothetical | N23 | Tb927.7.2330_for | GGGGACAAGTTTGTACAAAAAAGCAGGCTATGAGATCGCGAAATACGGATGC | Tb927.7.2330_rev | GGGGACCACTTTGTACAAGAAAGCTGGGTTCAGCGGTTACGGATAACCCA |
| 10_hypothetical | N24 | Tb927.9.12780_for | GGGGACAAGTTTGTACAAAAAAGCAGGCTATGTCCATTGCGGTAGAAGAGAGT | Tb927.9.12780_rev | GGGGACCACTTTGTACAAGAAAGCTGGGTCTATCGAAGGGCAGTACACATATCATC |
| 10_hypothetical | O1 | Tb927.4.1060_for | GGGGACAAGTTTGTACAAAAAAGCAGGCTATGCGGTGTTCCGCGCGC | Tb927.4.1060_rev | GGGGACCACTTTGTACAAGAAAGCTGGGTTTATTCTCCTTCACCGTTGGTAGGG |
| 10_hypothetical | O2 | Tb927.3.2780_for | GGGGACAAGTTTGTACAAAAAAGCAGGCTATGTATGTGTTCTCGCCAGCG | Tb927.3.2780_rev | GGGGACCACTTTGTACAAGAAAGCTGGGTTCACACCGCCGATGATGCG |
| 10_hypothetical | O3 | Tb927.10.11850_for | GGGGACAAGTTTGTACAAAAAAGCAGGCTATGATGATCCAGGAGCGTGATGC | Tb927.10.11850_rev | GGGGACCACTTTGTACAAGAAAGCTGGGTTCATCCCTCGTCTTTCCTTTTGAAC |
| 10_hypothetical | O4 | Tb927.8.5240_for | GGGGACAAGTTTGTACAAAAAAGCAGGCTATGGATCCGCACCCAACGC | Tb927.8.5240_rev | GGGGACCACTTTGTACAAGAAAGCTGGGTCTACCGCTGTTCTATGATGCCAC |
| 10_hypothetical | O5 | Tb927.8.7750_for | GGGGACAAGTTTGTACAAAAAAGCAGGCTATGGGGGCTGTGCCCAGC | Tb927.8.7750_rev | GGGGACCACTTTGTACAAGAAAGCTGGGTTCAGTGTTGCAGGCCACGC |
| 10_hypothetical | O6 | Tb927.7.4220_for | GGGGACAAGTTTGTACAAAAAAGCAGGCTATGGTTGGGAAGACGTATCTTCGG | Tb927.7.4220_rev | GGGGACCACTTTGTACAAGAAAGCTGGGTTTAATGCTTCTCTCCTCTTTGCTCTTC |
| 10_hypothetical | O7 | Tb927.8.1290_for | GGGGACAAGTTTGTACAAAAAAGCAGGCTATGAGCCCTTCCAGTTTGACATTG | Tb927.8.1290_rev | GGGGACCACTTTGTACAAGAAAGCTGGGTTCAGCCGTAGAGAAATACGACAACG |
| 10_hypothetical | O8 | Tb927.11.1160_for | GGGGACAAGTTTGTACAAAAAAGCAGGCTATGAAGGCGCTTAGTGACGATTC | Tb927.11.1160_rev | GGGGACCACTTTGTACAAGAAAGCTGGGTCTACAACACTAACCAAGAATCGTCCTC |
| 10_hypothetical | O9 | Tb927.6.2660_for | GGGGACAAGTTTGTACAAAAAAGCAGGCTATGGGTGAAACAGTGAGAAGGTGT | Tb927.6.2660_rev | GGGGACCACTTTGTACAAGAAAGCTGGGTCTAGGCATCCTGCATGAGAAACTG |
| 10_hypothetical | O10 | Tb927.11.3910_for | GGGGACAAGTTTGTACAAAAAAGCAGGCTATGCCTAGTTCCAAGTCGGACA | Tb927.11.3910_rev | GGGGACCACTTTGTACAAGAAAGCTGGGTTTACTTTTCTTCCTCCCCAGGCA |
| 10_hypothetical | O11 | Tb927.8.7540_for | GGGGACAAGTTTGTACAAAAAAGCAGGCTATGCTGGACACCAATGGGACT | Tb927.8.7540_rev | GGGGACCACTTTGTACAAGAAAGCTGGGTTCAAATATTTTTCGCACTCCGTCCA |
| 10_hypothetical | O12 | Tb927.11.16240_for | GGGGACAAGTTTGTACAAAAAAGCAGGCTATGTCATCACGAGAAGAACGCAG | Tb927.11.16240_rev | GGGGACCACTTTGTACAAGAAAGCTGGGTCTACGTGGGTCTGTCGATGAATG |
| 10_hypothetical | O13 | Tb927.9.8620_for | GGGGACAAGTTTGTACAAAAAAGCAGGCTATGGAGGGGTTCAAAACACCAAAC | Tb927.9.8620_rev | GGGGACCACTTTGTACAAGAAAGCTGGGTCTAATGCCCCCGAGTTAAACGG |
| 10_hypothetical | O14 | Tb927.6.3590_for | GGGGACAAGTTTGTACAAAAAAGCAGGCTATGAATTTTGCGACATCACAGCAAG | Tb927.6.3590_rev | GGGGACCACTTTGTACAAGAAAGCTGGGTCTAAATGATCTCATCGCAAAATTGGACG |
| 10_hypothetical | O15 | Tb927.11.2140_for | GGGGACAAGTTTGTACAAAAAAGCAGGCTATGAGCGACTCCATGGGTGA | Tb927.11.2140_rev | GGGGACCACTTTGTACAAGAAAGCTGGGTCTAAGAGCTGTATGCATGCCGG |
| 10_hypothetical | O16 | Tb927.10.14400_for | GGGGACAAGTTTGTACAAAAAAGCAGGCTATGAGCCCCTCTCAATTATCGCG | Tb927.10.14400_rev | GGGGACCACTTTGTACAAGAAAGCTGGGTTCAGTGGCTACAACGGAAAAAAGC |
| 10_hypothetical | O17 | Tb927.11.14650_for | GGGGACAAGTTTGTACAAAAAAGCAGGCTATGATGGAGTTGGCGGAAGGT | Tb927.11.14650_rev | GGGGACCACTTTGTACAAGAAAGCTGGGTTCATGTTGTCAACAACTGGGGG |
| 10_hypothetical | O18 | Tb927.10.13350_for | GGGGACAAGTTTGTACAAAAAAGCAGGCTATGAGTACTCCCCCGCATGC | Tb927.10.13350_rev | GGGGACCACTTTGTACAAGAAAGCTGGGTCTAGCCTGCGGGTGTCGA |
| 10_hypothetical | O19 | Tb927.10.12620_for | GGGGACAAGTTTGTACAAAAAAGCAGGCTATGTCCGTGAAAAGGAACACCC | Tb927.10.12620_rev | GGGGACCACTTTGTACAAGAAAGCTGGGTTTACATACCTCCTTGACCCAAGGG |
| 10_hypothetical | O20 | Tb927.8.7200_for | GGGGACAAGTTTGTACAAAAAAGCAGGCTATGGCTGCCGTGTGGCGG | Tb927.8.7200_rev | GGGGACCACTTTGTACAAGAAAGCTGGGTCTACCACAGCTCTACTGCCTCC |
| 10_hypothetical | O21 | Tb927.9.1940_for | GGGGACAAGTTTGTACAAAAAAGCAGGCTATGACGTTTAGCGCAAGGGAG | Tb927.9.1940_rev | GGGGACCACTTTGTACAAGAAAGCTGGGTTCATCCCATCAAACCAAGAAGCAAC |
| 10_hypothetical | O22 | Tb927.6.2210_for | GGGGACAAGTTTGTACAAAAAAGCAGGCTATGGAAGGTTTGTTTATTCGGGTTGT | Tb927.6.2210_rev | GGGGACCACTTTGTACAAGAAAGCTGGGTTTACTGGCACCTTCTACTATCACCAC |
| 10_hypothetical | O23 | Tb927.8.2880_for | GGGGACAAGTTTGTACAAAAAAGCAGGCTATGTATTCAAGGGCGCTGCC | Tb927.8.2880_rev | GGGGACCACTTTGTACAAGAAAGCTGGGTCTACTGACTGTAACAATGAATCATCCGC |
| 10_hypothetical | O24 | Tb927.10.14320_for | GGGGACAAGTTTGTACAAAAAAGCAGGCTATGTGCTCGCTTTCTCCGAAC | Tb927.10.14320_rev | GGGGACCACTTTGTACAAGAAAGCTGGGTCTAACGCTTTCGTGCACGGG |
| 10_hypothetical | P1 | Tb927.7.6760_for | GGGGACAAGTTTGTACAAAAAAGCAGGCTATGTACCAGTCCGGAGGCC | Tb927.7.6760_rev | GGGGACCACTTTGTACAAGAAAGCTGGGTTCACTCTGCAAATATGGCTCTGC |
| 10_hypothetical | P2 | Tb927.10.5260_for | GGGGACAAGTTTGTACAAAAAAGCAGGCTATGACTGGAGGCAATAGCGGG | Tb927.10.5260_rev | GGGGACCACTTTGTACAAGAAAGCTGGGTTCACTCTTTGGTCGAATCCTTCTTATCG |
| 10_hypothetical | P3 | Tb927.3.2770_for | GGGGACAAGTTTGTACAAAAAAGCAGGCTATGAAGAAAGCATGGGCTCAACTG | Tb927.3.2770_rev | GGGGACCACTTTGTACAAGAAAGCTGGGTTTAGCCTTCGCACCAAACTAAGATATC |
| 10_hypothetical | P4 | Tb927.11.1580_for | GGGGACAAGTTTGTACAAAAAAGCAGGCTATGCCAGCTTTTTCTTCACTCTCTAAG | Tb927.11.1580_rev | GGGGACCACTTTGTACAAGAAAGCTGGGTTCAAGTAGACTTCAAACAGCGCC |
| 10_hypothetical | P5 | Tb927.1.2980_for | GGGGACAAGTTTGTACAAAAAAGCAGGCTATGTTGTTGTCCATCCCGGTAAGA | Tb927.1.2980_rev | GGGGACCACTTTGTACAAGAAAGCTGGGTTTACGGAACAGTTAGTCTCAATCTGTCA |
| 10_hypothetical | P6 | Tb927.4.4530_for | GGGGACAAGTTTGTACAAAAAAGCAGGCTATGTCGGCATCAACTAGTTTCCAC | Tb927.4.4530_rev | GGGGACCACTTTGTACAAGAAAGCTGGGTTCATACAAACTTAAGGGCACCACC |
| 10_hypothetical | P7 | Tb927.11.4010_for | GGGGACAAGTTTGTACAAAAAAGCAGGCTATGGTCAACCAAACAGCGGG | Tb927.11.4010_rev | GGGGACCACTTTGTACAAGAAAGCTGGGTTTATTCGTTAGTCTCCGCAGCAGC |
| 10_hypothetical | P8 | Tb927.9.6160_for | GGGGACAAGTTTGTACAAAAAAGCAGGCTATGACCTCCCTTTTACCAGCGG | Tb927.9.6160_rev | GGGGACCACTTTGTACAAGAAAGCTGGGTTCATTTGGTTGTAGAGGTTATACCGTTC |
| 10_hypothetical | P9 | Tb927.7.5370_for | GGGGACAAGTTTGTACAAAAAAGCAGGCTATGTCAGCGAGGGCAGCA | Tb927.7.5370_rev | GGGGACCACTTTGTACAAGAAAGCTGGGTTTATTTTTTTTGTGTGGTGCTCCCCT |
| 10_hypothetical | P10 | Tb927.3.1320_for | GGGGACAAGTTTGTACAAAAAAGCAGGCTATGAAGTACTTTGCATGTCGTTTGGA | Tb927.3.1320_rev | GGGGACCACTTTGTACAAGAAAGCTGGGTCTATGAAACATGACCGTCCACCG |
| 10_hypothetical | P11 | Tb927.4.2410_for | GGGGACAAGTTTGTACAAAAAAGCAGGCTATGGGGTTTTGGCCGCGT | Tb927.4.2410_rev | GGGGACCACTTTGTACAAGAAAGCTGGGTTTAACTGCAGGGATCACTTGGCG |
| 10_hypothetical | P12 | Tb927.11.16340_for | GGGGACAAGTTTGTACAAAAAAGCAGGCTATGACCACCGCAGCTTCCG | Tb927.11.16340_rev | GGGGACCACTTTGTACAAGAAAGCTGGGTTTAGCAGCAACCGTATCGTTCAA |
| 10_hypothetical | P13 | Tb927.9.13330_for | GGGGACAAGTTTGTACAAAAAAGCAGGCTATGAGTTTTTTGGTTAAGACGTTGCTC | Tb927.9.13330_rev | GGGGACCACTTTGTACAAGAAAGCTGGGTTTAACGCAAATCTGTACTGGGGTC |
| 10_hypothetical | P14 | Tb927.10.7290_for | GGGGACAAGTTTGTACAAAAAAGCAGGCTATGGGGAACGGCCAAATTAATGC | Tb927.10.7290_rev | GGGGACCACTTTGTACAAGAAAGCTGGGTTCATCGTCTTCTATCCGGCGAC |
| 10_hypothetical | P15 | Tb927.3.920_for | GGGGACAAGTTTGTACAAAAAAGCAGGCTATGATTCACCGAGAGTTAGCAGATTC | Tb927.3.920_rev | GGGGACCACTTTGTACAAGAAAGCTGGGTCTATACTTTCAACTCTTCAATCATATATCG |
| 10_hypothetical | P16 | Tb927.11.1340_for | GGGGACAAGTTTGTACAAAAAAGCAGGCTATGAGTTCTGTGAGGCTTCTACTTGG | Tb927.11.1340_rev | GGGGACCACTTTGTACAAGAAAGCTGGGTTTATTCATCAGTGAAGTATAATGGATACCA |
| 10_hypothetical | P17 | Tb927.8.3180_for | GGGGACAAGTTTGTACAAAAAAGCAGGCTATGTCCGGGCGCCGGGGG | Tb927.8.3180_rev | GGGGACCACTTTGTACAAGAAAGCTGGGTTCAAGGAATAAAGGAACCCAACTGGT |
| 10_hypothetical | P18 | Tb927.10.13260_for | GGGGACAAGTTTGTACAAAAAAGCAGGCTATGCAAACAACTACCGCAGCA | Tb927.10.13260_rev | GGGGACCACTTTGTACAAGAAAGCTGGGTTCATTCGCCTTGAGGATACAACCA |
| 10_hypothetical | P19 | Tb927.11.12390_for | GGGGACAAGTTTGTACAAAAAAGCAGGCTATGGCTGACCGAGCAGCT | Tb927.11.12390_rev | GGGGACCACTTTGTACAAGAAAGCTGGGTCTAACCGGAGTCGCCACGT |
| 10_hypothetical | P20 | Tb927.3.3550_for | GGGGACAAGTTTGTACAAAAAAGCAGGCTATGTCACACTCTGCCTTTTTGCA | Tb927.3.3550_rev | GGGGACCACTTTGTACAAGAAAGCTGGGTTTAAAGACTGTAGCAACGGAAATCGG |
| 10_hypothetical | P21 | Tb927.10.11780_for | GGGGACAAGTTTGTACAAAAAAGCAGGCTATGTCCTCGCGTGAGGAGAG | Tb927.10.11780_rev | GGGGACCACTTTGTACAAGAAAGCTGGGTTCATTCCGATGTGGGGATCGT |
| 10_hypothetical | P22 | Tb927.10.6010_for | GGGGACAAGTTTGTACAAAAAAGCAGGCTATGCCCAGGAAGGTGAAGACT | Tb927.10.6010_rev | GGGGACCACTTTGTACAAGAAAGCTGGGTCTATTCGTATTCCATCGTTGCGATGA |
| 10_hypothetical | P23 | Tb927.7.3050_for | GGGGACAAGTTTGTACAAAAAAGCAGGCTATGTTTCGGGTCCGAAGGCT | Tb927.7.3050_rev | GGGGACCACTTTGTACAAGAAAGCTGGGTTCAACTCCATTCTGATTTCTGATCGG |
| 10_hypothetical | P24 | Tb927.10.6930_for | GGGGACAAGTTTGTACAAAAAAGCAGGCTATGGAGGGAGAGGAGTTGCTTT | Tb927.10.6930_rev | GGGGACCACTTTGTACAAGAAAGCTGGGTTCACGCTACATAGATATAAGTGTCCGC |
| hypothetical_las | A1 | Tb927.5.1910_for | GGGGACAAGTTTGTACAAAAAAGCAGGCTATGGAGAATCTCTCTGATGAGGATAAATCC | Tb927.5.1910_rev | GGGGACCACTTTGTACAAGAAAGCTGGGTTCAGAAGGATTGGGCACAGCA |
| hypothetical_las | A2 | Tb927.1.4450_for | GGGGACAAGTTTGTACAAAAAAGCAGGCTATGTACACGCTGTACCCCCG | Tb927.1.4450_rev | GGGGACCACTTTGTACAAGAAAGCTGGGTTTAGCAACCCATTCTGTTATGTGCT |
| hypothetical_las | A3 | Tb927.4.4500_for | GGGGACAAGTTTGTACAAAAAAGCAGGCTATGAGTGAGTGCCCACGCC | Tb927.4.4500_rev | GGGGACCACTTTGTACAAGAAAGCTGGGTTCAGTATTCTAACTTTTTTGCCCACTGT |
| hypothetical_las | A4 | Tb927.8.7830_for | GGGGACAAGTTTGTACAAAAAAGCAGGCTATGAGTGGGCGCCTGTCT | Tb927.8.7830_rev | GGGGACCACTTTGTACAAGAAAGCTGGGTTCAAAGTCCTTGGCATCTTAACTCTTC |
| hypothetical_las | A5 | Tb927.10.14340_for | GGGGACAAGTTTGTACAAAAAAGCAGGCTATGAGTGCTCACACACCGGA | Tb927.10.14340_rev | GGGGACCACTTTGTACAAGAAAGCTGGGTCTATTTGGACTTAACGAGTGAGGCG |
| hypothetical_las | A6 | Tb927.10.15550_for | GGGGACAAGTTTGTACAAAAAAGCAGGCTATGCTGACTGCACTGCATGTAAC | Tb927.10.15550_rev | GGGGACCACTTTGTACAAGAAAGCTGGGTTTACGGTTCTATCCCGTCACCAC |
| hypothetical_las | A7 | Tb927.10.2650_for | GGGGACAAGTTTGTACAAAAAAGCAGGCTATGATATCTCAACCTACTGGAACCACA | Tb927.10.2650_rev | GGGGACCACTTTGTACAAGAAAGCTGGGTCTAGCAGTGCTTCAACAGTAGTAAACT |
| hypothetical_las | A8 | Tb927.5.2440_for | GGGGACAAGTTTGTACAAAAAAGCAGGCTATGGCTGTCATCCCTAAGGTGG | Tb927.5.2440_rev | GGGGACCACTTTGTACAAGAAAGCTGGGTTCATGAGGCAAGTTGCTGTTCAT |
| hypothetical_las | A9 | Tb927.7.5500_for | GGGGACAAGTTTGTACAAAAAAGCAGGCTATGCAAAGTAGTCTCTCTATCTCACCTC | Tb927.7.5500_rev | GGGGACCACTTTGTACAAGAAAGCTGGGTCTAGAACCCCAGCTCCTCACG |
| hypothetical_las | A10 | Tb927.4.960_for | GGGGACAAGTTTGTACAAAAAAGCAGGCTATGAGCCAGTTGACTGCCATTTT | Tb927.4.960_rev | GGGGACCACTTTGTACAAGAAAGCTGGGTTTAGGTGGAGGTGGAAGTTATCCC |
| hypothetical_las | A11 | Tb927.8.2430_for | GGGGACAAGTTTGTACAAAAAAGCAGGCTATGAGTGATTTTGACAACAAATGTCAACC | Tb927.8.2430_rev | GGGGACCACTTTGTACAAGAAAGCTGGGTTTACACTTCTTCATCAGGCTGTTCTG |
| hypothetical_las | A12 | Tb927.10.14920_for | GGGGACAAGTTTGTACAAAAAAGCAGGCTATGACTCAACCGGTGGATGACA | Tb927.10.14920_rev | GGGGACCACTTTGTACAAGAAAGCTGGGTCTACACCCTCAATGGGAAAAGTGAAA |
| hypothetical_las | A13 | Tb927.2.4400_for | GGGGACAAGTTTGTACAAAAAAGCAGGCTATGATTCGGAGACGAGTATGCGA | Tb927.2.4400_rev | GGGGACCACTTTGTACAAGAAAGCTGGGTTCAAAGTTGCACCCATTTGGGG |
| hypothetical_las | A14 | Tb927.11.10710_for | GGGGACAAGTTTGTACAAAAAAGCAGGCTATGCGGCCCCAGGCTGAT | Tb927.11.10710_rev | GGGGACCACTTTGTACAAGAAAGCTGGGTTCATGCAACGAAGCATGATTCCC |
| hypothetical_las | A15 | Tb927.1.3860_for | GGGGACAAGTTTGTACAAAAAAGCAGGCTATGGCGTCAAACCAACAGGAC | Tb927.1.3860_rev | GGGGACCACTTTGTACAAGAAAGCTGGGTTCAAAACCGGGGCTGCAAC |
| hypothetical_las | A16 | Tb927.10.4080_for | GGGGACAAGTTTGTACAAAAAAGCAGGCTATGCGGCGATTTGCACCT | Tb927.10.4080_rev | GGGGACCACTTTGTACAAGAAAGCTGGGTTCACATGCGTGCGGACTG |
| hypothetical_las | A17 | Tb927.7.5190_for | GGGGACAAGTTTGTACAAAAAAGCAGGCTATGGAGCGACTATCTATTATCCACACA | Tb927.7.5190_rev | GGGGACCACTTTGTACAAGAAAGCTGGGTTCAATTTTCGCTGTTTACTAGAGTGGAT |
| hypothetical_las | A18 | Tb927.8.5110_for | GGGGACAAGTTTGTACAAAAAAGCAGGCTATGAGGGGGTTGCTGTCTCAC | Tb927.8.5110_rev | GGGGACCACTTTGTACAAGAAAGCTGGGTTTACTGTAGTATCACAACAGGACTTGTTC |
| hypothetical_las | A19 | Tb927.5.3280_for | GGGGACAAGTTTGTACAAAAAAGCAGGCTATGGCATCCTTGCAGTTGGG | Tb927.5.3280_rev | GGGGACCACTTTGTACAAGAAAGCTGGGTTTACTGCGCAGAAGTATATGGGAGT |
| hypothetical_las | A20 | Tb927.5.3290_for | GGGGACAAGTTTGTACAAAAAAGCAGGCTATGCAAATCACAGGAGTGCGG | Tb927.5.3290_rev | GGGGACCACTTTGTACAAGAAAGCTGGGTTTACTTCTTTGGTGTTGCAACACTTTTT |
| hypothetical_las | A21 | Tb927.8.5170_for | GGGGACAAGTTTGTACAAAAAAGCAGGCTATGAAGGAAGCCGAACAACCAC | Tb927.8.5170_rev | GGGGACCACTTTGTACAAGAAAGCTGGGTTTAATAGACATCTTTGTGATGTTTGATTAC |
| hypothetical_las | A22 | Tb927.10.15280_for | GGGGACAAGTTTGTACAAAAAAGCAGGCTATGCAGCGGGAAGTGATACGT | Tb927.10.15280_rev | GGGGACCACTTTGTACAAGAAAGCTGGGTTTACATTGCACTGAGCCACCC |
| hypothetical_las | A23 | Tb927.8.7800_for | GGGGACAAGTTTGTACAAAAAAGCAGGCTATGTCAACACCAATTGGTTCTCGC | Tb927.8.7800_rev | GGGGACCACTTTGTACAAGAAAGCTGGGTTTACCTGAACTCAATGGCACCAC |
| hypothetical_las | A24 | Tb927.11.4230_for | GGGGACAAGTTTGTACAAAAAAGCAGGCTATGGTGGACGAAGAGACGCT | Tb927.11.4230_rev | GGGGACCACTTTGTACAAGAAAGCTGGGTTCACGAGCTACGTGCTGCA |
| hypothetical_las | B1 | Tb927.10.8180_for | GGGGACAAGTTTGTACAAAAAAGCAGGCTATGAGCGGGTCGCGTCTG | Tb927.10.8180_rev | GGGGACCACTTTGTACAAGAAAGCTGGGTCTAAAGCAGCCGCTCCAAGG |
| hypothetical_las | B2 | Tb927.8.3870_for | GGGGACAAGTTTGTACAAAAAAGCAGGCTATGCCAGTTACTGTCGAGGAAGT | Tb927.8.3870_rev | GGGGACCACTTTGTACAAGAAAGCTGGGTCTATTCACTGTCCTCAAACTGAAACGA |
| hypothetical_las | B3 | Tb927.7.2560_for | GGGGACAAGTTTGTACAAAAAAGCAGGCTATGCAACGCCTGCTTCATGC | Tb927.7.2560_rev | GGGGACCACTTTGTACAAGAAAGCTGGGTTTATTCAATAAGAAAAAATGTGAAGGGCCC |
| hypothetical_las | B4 | Tb927.2.2340_for | GGGGACAAGTTTGTACAAAAAAGCAGGCTATGGCCGCCGTAGTGCAG | Tb927.2.2340_rev | GGGGACCACTTTGTACAAGAAAGCTGGGTTTATGTTGACACCGTCATTCCCCC |
| hypothetical_las | B5 | Tb927.8.5160_for | GGGGACAAGTTTGTACAAAAAAGCAGGCTATGGGTGTGTGCCAGTCAAATC | Tb927.8.5160_rev | GGGGACCACTTTGTACAAGAAAGCTGGGTTCAGCAAGTGAGCGTAACTCCG |
| hypothetical_las | B6 | Tb927.11.10150_for | GGGGACAAGTTTGTACAAAAAAGCAGGCTATGGAGCGAAGGAGGTTGTTGC | Tb927.11.10150_rev | GGGGACCACTTTGTACAAGAAAGCTGGGTTCAGCTGCTATCGTCCCCATC |
| hypothetical_las | B7 | Tb927.3.4030_for | GGGGACAAGTTTGTACAAAAAAGCAGGCTATGACCTTTAGCACGCTTGTGC | Tb927.3.4030_rev | GGGGACCACTTTGTACAAGAAAGCTGGGTCTATTCAAGTGGCGGTAGGTCAGT |
| hypothetical_las | B8 | Tb927.11.16010_for | GGGGACAAGTTTGTACAAAAAAGCAGGCTATGAACCCCACTGATCTTCGTGA | Tb927.11.16010_rev | GGGGACCACTTTGTACAAGAAAGCTGGGTTTATCGCACATCACTAAAAAAGCCCA |
| hypothetical_las | B9 | Tb927.10.15240_for | GGGGACAAGTTTGTACAAAAAAGCAGGCTATGCAGAGCGCGGGAAGC | Tb927.10.15240_rev | GGGGACCACTTTGTACAAGAAAGCTGGGTCTATCGATCGCGCGCACG |
| hypothetical_las | B10 | Tb927.11.9080_for | GGGGACAAGTTTGTACAAAAAAGCAGGCTATGGGAGCCGACGACGAG | Tb927.11.9080_rev | GGGGACCACTTTGTACAAGAAAGCTGGGTTTACTTCCGTGACTTGTGCTGTG |
| hypothetical_las | B11 | Tb927.11.2320_for | GGGGACAAGTTTGTACAAAAAAGCAGGCTATGAGCCAACAAAAGGGGAAGAAA | Tb927.11.2320_rev | GGGGACCACTTTGTACAAGAAAGCTGGGTTCAGCCTTTTCCTTTAGTAACGGGC |
| hypothetical_las | B12 | Tb927.11.15400_for | GGGGACAAGTTTGTACAAAAAAGCAGGCTATGTTCGAAAATTTGATGCGATGGC | Tb927.11.15400_rev | GGGGACCACTTTGTACAAGAAAGCTGGGTTTAGAACCAGCTATTTTTTTGCTTCTGATT |
| hypothetical_las | B13 | Tb927.10.1490_for | GGGGACAAGTTTGTACAAAAAAGCAGGCTATGAGGAAAGCTGGAGCAAATCG | Tb927.10.1490_rev | GGGGACCACTTTGTACAAGAAAGCTGGGTCTAGTGAGGAATCGGTTCTTGGTG |
| hypothetical_las | B14 | Tb927.11.910_for | GGGGACAAGTTTGTACAAAAAAGCAGGCTATGGAGGTGTTCGATCTGGCG | Tb927.11.910_rev | GGGGACCACTTTGTACAAGAAAGCTGGGTTCACGTCACTTTGATTGCTTCTGG |
| hypothetical_las | B15 | Tb927.10.10050_for | GGGGACAAGTTTGTACAAAAAAGCAGGCTATGTTGTTGGGGTTGAAAGATCTTTTTATT | Tb927.10.10050_rev | GGGGACCACTTTGTACAAGAAAGCTGGGTCTACTGATGAACTCCGGGGCT |
| hypothetical_las | B16 | Tb927.11.890_for | GGGGACAAGTTTGTACAAAAAAGCAGGCTATGAAGAAACTCAGATCATTTCTTCCCTG | Tb927.11.890_rev | GGGGACCACTTTGTACAAGAAAGCTGGGTCTACTCTCTTGAAAGCTCACGCAA |
| hypothetical_las | B17 | Tb927.10.3010_for | GGGGACAAGTTTGTACAAAAAAGCAGGCTATGCCATTACCATCCACGCTACC | Tb927.10.3010_rev | GGGGACCACTTTGTACAAGAAAGCTGGGTTTAGTCTTTCGGTTTAGACTGGGCA |
| hypothetical_las | B18 | Tb927.9.11670_for | GGGGACAAGTTTGTACAAAAAAGCAGGCTATGCGGCCGCCGACGTTA | Tb927.9.11670_rev | GGGGACCACTTTGTACAAGAAAGCTGGGTCTAAGAGTGGACAGAAGTCACACG |
| hypothetical_las | B19 | Tb927.3.5160_for | GGGGACAAGTTTGTACAAAAAAGCAGGCTATGAGTGACCGTCCCCGAC | Tb927.3.5160_rev | GGGGACCACTTTGTACAAGAAAGCTGGGTTTATTTGTCCGTCAGGTCCACAAC |
| hypothetical_las | B20 | Tb927.10.13180_for | GGGGACAAGTTTGTACAAAAAAGCAGGCTATGCTCCCTACGCCATACCAT | Tb927.10.13180_rev | GGGGACCACTTTGTACAAGAAAGCTGGGTTTACTTCACTTTTCTCTTCGCTACTTTTTC |
| hypothetical_las | B21 | Tb927.2.4050_for | GGGGACAAGTTTGTACAAAAAAGCAGGCTATGTCTTCGAGTCGTGGAAGTTTTC | Tb927.2.4050_rev | GGGGACCACTTTGTACAAGAAAGCTGGGTTCAATTCTTCTTCGGTGATATCGATGG |
| hypothetical_las | B22 | Tb927.7.3550_for | GGGGACAAGTTTGTACAAAAAAGCAGGCTATGTCTATATTCGGTGAACCACTTCCC | Tb927.7.3550_rev | GGGGACCACTTTGTACAAGAAAGCTGGGTTTAGAAACCTCGCCCAACGGA |
| hypothetical_las | B23 | Tb927.2.5860_for | GGGGACAAGTTTGTACAAAAAAGCAGGCTATGGTTGCCACGGAACAATCAA | Tb927.2.5860_rev | GGGGACCACTTTGTACAAGAAAGCTGGGTTCACGATGGCTCAACCTCACT |

| | | | | | |
|---|---|---|---|---|---|
| hypothetical_las | B24 | Tb927.9.13050_for | GGGGACAAGTTTGTACAAAAAAGCAGGCTATGAAGGATCCTGTAGTTAGCGGT | Tb927.9.13050_rev | GGGGACCACTTTGTACAAGAAAGCTGGGTTCAGATGTGTTTCCACTGCAACC |
| hypothetical_las | C1 | Tb927.8.4160_for | GGGGACAAGTTTGTACAAAAAAGCAGGCTATGGAAAACAGCACAGTAGCCTTC | Tb927.8.4160_rev | GGGGACCACTTTGTACAAGAAAGCTGGGTTTAAAAACAAAGCTTCAGTGCTTTCGC |
| hypothetical_las | C2 | Tb927.11.8960_for | GGGGACAAGTTTGTACAAAAAAGCAGGCTATGTCCGCATTTGAGTTTCTTCGG | Tb927.11.8960_rev | GGGGACCACTTTGTACAAGAAAGCTGGGTTCACGCCATGTCCGGAAGA |
| hypothetical_las | C3 | Tb927.11.14040_for | GGGGACAAGTTTGTACAAAAAAGCAGGCTATGCGTTTTCCTAGCGGCTC | Tb927.11.14040_rev | GGGGACCACTTTGTACAAGAAAGCTGGGTCTAACTTACATTTTCAAGGCAATACCGTAT |
| hypothetical_las | C4 | Tb927.7.4150_for | GGGGACAAGTTTGTACAAAAAAGCAGGCTATGGTCGTTCATCTTACCCCACT | Tb927.7.4150_rev | GGGGACCACTTTGTACAAGAAAGCTGGGTTCATTTCGTCCGCCGAAATTTCT |
| hypothetical_las | C5 | Tb927.10.13070_for | GGGGACAAGTTTGTACAAAAAAGCAGGCTATGACGGATAAGGAAGATGCCAGT | Tb927.10.13070_rev | GGGGACCACTTTGTACAAGAAAGCTGGGTTTAGCTGTGAAACCAGCGGAC |
| hypothetical_las | C6 | Tb927.3.1310_for | GGGGACAAGTTTGTACAAAAAAGCAGGCTATGTTGCTTTGCAAGCAACCAG | Tb927.3.1310_rev | GGGGACCACTTTGTACAAGAAAGCTGGGTTCACACACCACAACGCTCATTT |
| hypothetical_las | C7 | Tb927.2.4520_for | GGGGACAAGTTTGTACAAAAAAGCAGGCTATGAAGCGTGTTCTCGTGCTC | Tb927.2.4520_rev | GGGGACCACTTTGTACAAGAAAGCTGGGTTCAAATCTCCGTTCGCCGC |
| hypothetical_las | C8 | Tb927.11.5910_for | GGGGACAAGTTTGTACAAAAAAGCAGGCTATGCTCACAACCTTCGATTTTGGT | Tb927.11.5910_rev | GGGGACCACTTTGTACAAGAAAGCTGGGTCTAGCCTTCATCTGCATCTGTACCT |
| hypothetical_las | C9 | Tb927.8.4560_for | GGGGACAAGTTTGTACAAAAAAGCAGGCTATGACCTTGACAGGCCTAAGCA | Tb927.8.4560_rev | GGGGACCACTTTGTACAAGAAAGCTGGGTCTACCAGCCGGTGACACTTTC |
| hypothetical_las | C10 | Tb927.7.7100_for | GGGGACAAGTTTGTACAAAAAAGCAGGCTATGTCGCTTGCTTACCTCAAAAGG | Tb927.7.7100_rev | GGGGACCACTTTGTACAAGAAAGCTGGGTTTAAGCAATATTTACACCCCGCCC |
| hypothetical_las | C11 | Tb927.11.6430_for | GGGGACAAGTTTGTACAAAAAAGCAGGCTATGGAGTCCGTGCCGCGC | Tb927.11.6430_rev | GGGGACCACTTTGTACAAGAAAGCTGGGTTCAAAAAGACCACTGCGCAGT |
| hypothetical_las | C12 | Tb927.6.1180_for | GGGGACAAGTTTGTACAAAAAAGCAGGCTATGACTGATTTCGGGCGCG | Tb927.6.1180_rev | GGGGACCACTTTGTACAAGAAAGCTGGGTTCATGCACCCTCCTCCTCCG |
| hypothetical_las | C13 | Tb927.4.2640_for | GGGGACAAGTTTGTACAAAAAAGCAGGCTATGTACGGCGTCACCCATCC | Tb927.4.2640_rev | GGGGACCACTTTGTACAAGAAAGCTGGGTCTAGAAGCAAGAGCCGCGG |
| hypothetical_las | C14 | Tb927.6.1730_for | GGGGACAAGTTTGTACAAAAAAGCAGGCTATGCAGCAGAAGGTACTGATTGC | Tb927.6.1730_rev | GGGGACCACTTTGTACAAGAAAGCTGGGTTCACCAATCACTATCACTGCTATCAAATG |
| hypothetical_las | C15 | Tb927.9.12490_for | GGGGACAAGTTTGTACAAAAAAGCAGGCTATGCAAGACATGCGTATGTTTAATGTATG | Tb927.9.12490_rev | GGGGACCACTTTGTACAAGAAAGCTGGGTCTATGAACTGTCTTGAAGGAATGAGGC |
| hypothetical_las | C16 | Tb927.9.12990_for | GGGGACAAGTTTGTACAAAAAAGCAGGCTATGAAACTCTTCACACCAGAGGAGA | Tb927.9.12990_rev | GGGGACCACTTTGTACAAGAAAGCTGGGTCTACAACACAGTCGCGCCTC |
| hypothetical_las | C17 | Tb927.4.2990_for | GGGGACAAGTTTGTACAAAAAAGCAGGCTATGTCTCTTTCCCTGGGTTGTAGG | Tb927.4.2990_rev | GGGGACCACTTTGTACAAGAAAGCTGGGTTTATCGCTTGAGTAAGGGCAATTTATTCC |
| hypothetical_las | C18 | Tb927.11.2510_for | GGGGACAAGTTTGTACAAAAAAGCAGGCTATGAAACCTCCTAGTGCGGCT | Tb927.11.2510_rev | GGGGACCACTTTGTACAAGAAAGCTGGGTTTATTGGCTTAGATTCACTTTTTGTTTTTT |
| hypothetical_las | C19 | Tb927.9.10450_for | GGGGACAAGTTTGTACAAAAAAGCAGGCTATGGTGGGAAAAAGGGGTCGT | Tb927.9.10450_rev | GGGGACCACTTTGTACAAGAAAGCTGGGTCTACCAGAAACTCAGCAGTGCG |
| hypothetical_las | C20 | Tb927.8.2820_for | GGGGACAAGTTTGTACAAAAAAGCAGGCTATGCCGCTATACAATATCACTATTCATCC | Tb927.8.2820_rev | GGGGACCACTTTGTACAAGAAAGCTGGGTCTATACTGGAAAACGGGCGGC |
| hypothetical_las | C21 | Tb927.11.620_for | GGGGACAAGTTTGTACAAAAAAGCAGGCTATGAGGAACCGGCGGCGT | Tb927.11.620_rev | GGGGACCACTTTGTACAAGAAAGCTGGGTTCAGTGCACAACTAAAATCTCACCTC |
| hypothetical_las | C22 | Tb927.8.790_for | GGGGACAAGTTTGTACAAAAAAGCAGGCTATGACAGGCACAACCGAAAACC | Tb927.8.790_rev | GGGGACCACTTTGTACAAGAAAGCTGGGTCTAAACGTATTTTGCGGGATCAGAAAAG |
| hypothetical_las | C23 | Tb927.8.3080_for | GGGGACAAGTTTGTACAAAAAAGCAGGCTATGCTGAGAAACTTGCAAGCCC | Tb927.8.3080_rev | GGGGACCACTTTGTACAAGAAAGCTGGGTTCATAGAGGCGCTACATCCTCC |
| hypothetical_las | C24 | Tb927.7.1150_for | GGGGACAAGTTTGTACAAAAAAGCAGGCTATGCGAGGATCCAATAGGGCG | Tb927.7.1150_rev | GGGGACCACTTTGTACAAGAAAGCTGGGTTTACCACTTCTTGCCCCCGT |
| hypothetical_las | D1 | Tb927.5.1010_for | GGGGACAAGTTTGTACAAAAAAGCAGGCTATGGAAACTGCCGCAGAGC | Tb927.5.1010_rev | GGGGACCACTTTGTACAAGAAAGCTGGGTTCACCTTCGCATGTCCCCG |
| hypothetical_las | D2 | Tb927.7.7250_for | GGGGACAAGTTTGTACAAAAAAGCAGGCTATGCCACCCAGACCTAGGC | Tb927.7.7250_rev | GGGGACCACTTTGTACAAGAAAGCTGGGTCTACGCTCTGCAATCCCCCC |
| hypothetical_las | D3 | Tb927.7.7190_for | GGGGACAAGTTTGTACAAAAAAGCAGGCTATGGGAAAAAAGAAGGCGGAAAGG | Tb927.7.7190_rev | GGGGACCACTTTGTACAAGAAAGCTGGGTTTACACTCCGACACTGCCGG |
| hypothetical_las | D4 | Tb927.11.6420_for | GGGGACAAGTTTGTACAAAAAAGCAGGCTATGTCTCTTGGGAAAGTATTGCGTT | Tb927.11.6420_rev | GGGGACCACTTTGTACAAGAAAGCTGGGTCTACTTCTCGTTCGCCTCATTGTC |
| hypothetical_las | D5 | Tb927.10.8000_for | GGGGACAAGTTTGTACAAAAAAGCAGGCTATGTTGGATGCGGAGGACTTTG | Tb927.10.8000_rev | GGGGACCACTTTGTACAAGAAAGCTGGGTTCAATCTACATCGATGGCAAATGTCTT |
| hypothetical_las | D6 | Tb927.3.4510_for | GGGGACAAGTTTGTACAAAAAAGCAGGCTATGGGTGAAGTAATGATTTTGCGAGA | Tb927.3.4510_rev | GGGGACCACTTTGTACAAGAAAGCTGGGTTTAGTCAATGTACGGGACGCGG |
| hypothetical_las | D7 | Tb927.2.4810_for | GGGGACAAGTTTGTACAAAAAAGCAGGCTATGGAGGAGAGGAAAGGTCACG | Tb927.2.4810_rev | GGGGACCACTTTGTACAAGAAAGCTGGGTCTACACAAGTGGGGGGAGCG |
| hypothetical_las | D8 | Tb927.9.14190_for | GGGGACAAGTTTGTACAAAAAAGCAGGCTATGAGCGAGTGGGAAGTAGATGAG | Tb927.9.14190_rev | GGGGACCACTTTGTACAAGAAAGCTGGGTTCAGGAAGCAACCATGAGCTCA |
| hypothetical_las | D9 | Tb927.4.2470_for | GGGGACAAGTTTGTACAAAAAAGCAGGCTATGGAGCTACTCAACGTGTTACCG | Tb927.4.2470_rev | GGGGACCACTTTGTACAAGAAAGCTGGGTTCATATTTGCTCAGTCTTTTGAATGGTGC |
| hypothetical_las | D10 | Tb927.9.4630_for | GGGGACAAGTTTGTACAAAAAAGCAGGCTATGCAGAGTTTGGTCTCTTTGCG | Tb927.9.4630_rev | GGGGACCACTTTGTACAAGAAAGCTGGGTTCAGCAAACAAAAAGATCTTCCGCT |
| hypothetical_las | D11 | Tb927.9.5620_for | GGGGACAAGTTTGTACAAAAAAGCAGGCTATGGAGGCGATTTACTTTACTACTAAGC | Tb927.9.5620_rev | GGGGACCACTTTGTACAAGAAAGCTGGGTCTAAATCGCGCAATTTGCTCCTTG |
| hypothetical_las | D12 | Tb927.7.1310_for | GGGGACAAGTTTGTACAAAAAAGCAGGCTATGGAGTCAAAACGCTTTTCGGT | Tb927.7.1310_rev | GGGGACCACTTTGTACAAGAAAGCTGGGTTCACTCCTGTCGACCCTCATTT |
| hypothetical_las | D13 | Tb927.10.870_for | GGGGACAAGTTTGTACAAAAAAGCAGGCTATGAGCGAGCCCAGGGAAG | Tb927.10.870_rev | GGGGACCACTTTGTACAAGAAAGCTGGGTTTAGAGGGCCTCTCGTTGCG |
| hypothetical_las | D14 | Tb927.3.2070_for | GGGGACAAGTTTGTACAAAAAAGCAGGCTATGGGAATGAGACGAAGCCGG | Tb927.3.2070_rev | GGGGACCACTTTGTACAAGAAAGCTGGGTTTAGTCGAGCACATGCCGTG |
| hypothetical_las | D15 | Tb927.10.5420_for | GGGGACAAGTTTGTACAAAAAAGCAGGCTATGCGTGTGACATGTGTAGTTGC | Tb927.10.5420_rev | GGGGACCACTTTGTACAAGAAAGCTGGGTTTATCGCCTGTTGCAACGCA |
| hypothetical_las | D16 | Tb927.10.11470_for | GGGGACAAGTTTGTACAAAAAAGCAGGCTATGTTATGCCGACGTTTCCTCG | Tb927.10.11470_rev | GGGGACCACTTTGTACAAGAAAGCTGGGTCTACTCACGGTGGCATACCCC |
| hypothetical_las | D17 | Tb927.10.5390_for | GGGGACAAGTTTGTACAAAAAAGCAGGCTATGATGCAAAATGGTTTCGTTCTAATCC | Tb927.10.5390_rev | GGGGACCACTTTGTACAAGAAAGCTGGGTTTACCAAAACCAGCGAAGCAAGT |
| hypothetical_las | D18 | Tb927.9.3680_for | GGGGACAAGTTTGTACAAAAAAGCAGGCTATGGCCCAGTGCTTTGGAAGT | Tb927.9.3680_rev | GGGGACCACTTTGTACAAGAAAGCTGGGTTCATTTGTCCTTCGAGGGGATCC |
| hypothetical_las | D19 | Tb927.8.4360_for | GGGGACAAGTTTGTACAAAAAAGCAGGCTATGACTTTCGTTGACCTATTTGACATGA | Tb927.8.4360_rev | GGGGACCACTTTGTACAAGAAAGCTGGGTCTACCACCGCAAGATTTCTTGTTCC |
| hypothetical_las | D20 | Tb927.7.2150_for | GGGGACAAGTTTGTACAAAAAAGCAGGCTATGGAGCGGACGAACCGT | Tb927.7.2150_rev | GGGGACCACTTTGTACAAGAAAGCTGGGTCTACACGCAAACAGGGCGT |
| hypothetical_las | D21 | Tb927.10.14230_for | GGGGACAAGTTTGTACAAAAAAGCAGGCTATGGATGATCCATGCAGCGC | Tb927.10.14230_rev | GGGGACCACTTTGTACAAGAAAGCTGGGTTCAGTCACTCGCGTAGCCTAA |
| hypothetical_las | D22 | Tb927.9.2730_for | GGGGACAAGTTTGTACAAAAAAGCAGGCTATGCTAAACTACACTTACGTGGAGAAG | Tb927.9.2730_rev | GGGGACCACTTTGTACAAGAAAGCTGGGTTCATACGTAGCGCTCAGCAAAC |
| hypothetical_las | D23 | Tb927.4.5120_for | GGGGACAAGTTTGTACAAAAAAGCAGGCTATGGCTGCTCGCCGCCAT | Tb927.4.5120_rev | GGGGACCACTTTGTACAAGAAAGCTGGGTTCATTTCCGGAACACGAACGTC |
| hypothetical_las | D24 | Tb927.9.15290_for | GGGGACAAGTTTGTACAAAAAAGCAGGCTATGAATCCAAGTGCCCAGCCG | Tb927.9.15290_rev | GGGGACCACTTTGTACAAGAAAGCTGGGTTCACGAGATCGGCAGGCC |
| hypothetical_las | E1 | Tb927.10.11340_for | GGGGACAAGTTTGTACAAAAAAGCAGGCTATGATGCGTCGTATGACTGCAAC | Tb927.10.11340_rev | GGGGACCACTTTGTACAAGAAAGCTGGGTTTAGTTGCGGACCTTTCTTGACTTC |
| hypothetical_las | E2 | Tb927.7.4370_for | GGGGACAAGTTTGTACAAAAAAGCAGGCTATGGAGGGTGTTGCTTCATCCA | Tb927.7.4370_rev | GGGGACCACTTTGTACAAGAAAGCTGGGTCTAGTAACCTGCCAAGTCCCAGA |
| hypothetical_las | E3 | Tb927.8.1280_for | GGGGACAAGTTTGTACAAAAAAGCAGGCTATGTCATATGAAAAGATACTATGTATGGAA | Tb927.8.1280_rev | GGGGACCACTTTGTACAAGAAAGCTGGGTCTAGAGAGTCCACGGCACAATATTTC |
| hypothetical_las | E4 | Tb927.2.2370_for | GGGGACAAGTTTGTACAAAAAAGCAGGCTATGTCATCGGAGGAAGCCATTCA | Tb927.2.2370_rev | GGGGACCACTTTGTACAAGAAAGCTGGGTCTACCACAATGAGCCCATCCAC |
| hypothetical_las | E5 | Tb927.5.2510_for | GGGGACAAGTTTGTACAAAAAAGCAGGCTATGAATCGCAAGGGGTATAAGGCT | Tb927.5.2510_rev | GGGGACCACTTTGTACAAGAAAGCTGGGTTTACATGTCGTTTGCGTTAATGTGATAC |
| hypothetical_las | E6 | Tb927.7.3930_for | GGGGACAAGTTTGTACAAAAAAGCAGGCTATGATGTTTTTTTCTCTGTGCCGTAGA | Tb927.7.3930_rev | GGGGACCACTTTGTACAAGAAAGCTGGGTCTAATCGTCTTCACTACCGAGTAGCT |
| hypothetical_las | E7 | Tb927.7.1570_for | GGGGACAAGTTTGTACAAAAAAGCAGGCTATGTTTCGTGTCTCCTTCCCATTG | Tb927.7.1570_rev | GGGGACCACTTTGTACAAGAAAGCTGGGTTTATTCTCCCCTTACCTCCGTGC |
| hypothetical_las | E8 | Tb927.5.3260_for | GGGGACAAGTTTGTACAAAAAAGCAGGCTATGGATCCACTAAATGCAGTTGCG | Tb927.5.3260_rev | GGGGACCACTTTGTACAAGAAAGCTGGGTTTACAATGGCAGCAAGTGAGCA |
| hypothetical_las | E9 | Tb927.11.4520_for | GGGGACAAGTTTGTACAAAAAAGCAGGCTATGTTCCGCGAGGCACTTG | Tb927.11.4520_rev | GGGGACCACTTTGTACAAGAAAGCTGGGTCTACAAGTTTACCACATTCCACGGC |
| hypothetical_las | E10 | Tb927.5.500_for | GGGGACAAGTTTGTACAAAAAAGCAGGCTATGTTACCACCAATACGCCGC | Tb927.5.500_rev | GGGGACCACTTTGTACAAGAAAGCTGGGTCTATAGCTGAACTGAATACTTGTCGGTC |
| hypothetical_las | E11 | Tb927.8.3200_for | GGGGACAAGTTTGTACAAAAAAGCAGGCTATGGTTACCACTGGTCTTAGTCCTG | Tb927.8.3200_rev | GGGGACCACTTTGTACAAGAAAGCTGGGTTCAGCCACGCTGTGGTTCC |
| hypothetical_las | E12 | Tb927.10.1970_for | GGGGACAAGTTTGTACAAAAAAGCAGGCTATGGCCCAACCGGTGGAT | Tb927.10.1970_rev | GGGGACCACTTTGTACAAGAAAGCTGGGTTCAGGCCTCCCCCGGCGG |
| hypothetical_las | E13 | Tb927.10.9650_for | GGGGACAAGTTTGTACAAAAAAGCAGGCTATGTTGAGCACCCTTTTCTTCGAT | Tb927.10.9650_rev | GGGGACCACTTTGTACAAGAAAGCTGGGTCTATAGCCCTGCTAACTTCTTCTGC |
| hypothetical_las | E14 | Tb927.7.5390_for | GGGGACAAGTTTGTACAAAAAAGCAGGCTATGGAGAGAACGGAGGATGTAACA | Tb927.7.5390_rev | GGGGACCACTTTGTACAAGAAAGCTGGGTTCAACCAAACAATGTTTCTGATTTTGGC |
| hypothetical_las | E15 | Tb927.3.5020_for | GGGGACAAGTTTGTACAAAAAAGCAGGCTATGGGAATACAAGGTGATTTGGCTC | Tb927.3.5020_rev | GGGGACCACTTTGTACAAGAAAGCTGGGTTTATTGGCGACCATCAGCTGAAG |
| hypothetical_las | E16 | Tb927.3.4620_for | GGGGACAAGTTTGTACAAAAAAGCAGGCTATGCGCGACATGGAGGTGG | Tb927.3.4620_rev | GGGGACCACTTTGTACAAGAAAGCTGGGTCTAATCTGCGCTCATCATGACCTC |
| hypothetical_las | E17 | Tb927.2.4950_for | GGGGACAAGTTTGTACAAAAAAGCAGGCTATGCTGAAGCCACCTTTTATCCG | Tb927.2.4950_rev | GGGGACCACTTTGTACAAGAAAGCTGGGTTCATTCATGCAGCGGCAACA |
| hypothetical_las | E18 | Tb927.3.1330_for | GGGGACAAGTTTGTACAAAAAAGCAGGCTATGCGAGGAACTGTTTTGATCTTTGG | Tb927.3.1330_rev | GGGGACCACTTTGTACAAGAAAGCTGGGTTCACCAAATAGAATCAGTCGCATTAATGA |
| hypothetical_las | E19 | Tb927.11.4280_for | GGGGACAAGTTTGTACAAAAAAGCAGGCTATGGCAGCACCCTTCATTCCT | Tb927.11.4280_rev | GGGGACCACTTTGTACAAGAAAGCTGGGTTTACTCCATATCATGGCAAAACCCC |
| hypothetical_las | E20 | Tb927.10.7610_for | GGGGACAAGTTTGTACAAAAAAGCAGGCTATGCGTCGCCTACAAATTCGG | Tb927.10.7610_rev | GGGGACCACTTTGTACAAGAAAGCTGGGTCTACTTTGTTTCGAATAGTTCCTTTGTAGA |
| hypothetical_las | E21 | Tb927.11.2290_for | GGGGACAAGTTTGTACAAAAAAGCAGGCTATGACTACATGGCTCTTCAATGTAAGTG | Tb927.11.2290_rev | GGGGACCACTTTGTACAAGAAAGCTGGGTTTATGAAGAGTCACTTTTGGATAGGAACAT |
| hypothetical_las | E22 | Tb927.8.7490_for | GGGGACAAGTTTGTACAAAAAAGCAGGCTATGGAAGCCTTCATCGAAGCG | Tb927.8.7490_rev | GGGGACCACTTTGTACAAGAAAGCTGGGTTTATAGACTACCGAGAATATGACCAAAGTA |
| hypothetical_las | E24 | Tb927.9.10100_for | GGGGACAAGTTTGTACAAAAAAGCAGGCTATGGTAATGAAAGCCAATTTCTGCTTC | Tb927.9.10100_rev | GGGGACCACTTTGTACAAGAAAGCTGGGTTTACTCACCGTCACGAGCGA |
| hypothetical_las | F1 | Tb927.8.6790_for | GGGGACAAGTTTGTACAAAAAAGCAGGCTATGGTGACGATGCGGATGGA | Tb927.8.6790_rev | GGGGACCACTTTGTACAAGAAAGCTGGGTTTAAACAGTCCTCCCCACTGTGG |
| hypothetical_las | F2 | Tb927.11.6410_for | GGGGACAAGTTTGTACAAAAAAGCAGGCTATGAGCTCAAGCGTCCTCCA | Tb927.11.6410_rev | GGGGACCACTTTGTACAAGAAAGCTGGGTCTAGCGACGCTCAGATTCCTGC |
| hypothetical_las | F3 | Tb927.7.5150_for | GGGGACAAGTTTGTACAAAAAAGCAGGCTATGCCGGCAAAAAGCCGG | Tb927.7.5150_rev | GGGGACCACTTTGTACAAGAAAGCTGGGTTCATGTTAGCGGCGCAAGG |
| hypothetical_las | F4 | Tb927.7.6410_for | GGGGACAAGTTTGTACAAAAAAGCAGGCTATGTCTTCGGGGGCTGTACTG | Tb927.7.6410_rev | GGGGACCACTTTGTACAAGAAAGCTGGGTTCAAGCAGTGGAAGCCGCC |
| hypothetical_las | F5 | Tb927.8.4960_for | GGGGACAAGTTTGTACAAAAAAGCAGGCTATGCTGCGGCATACGACG | Tb927.8.4960_rev | GGGGACCACTTTGTACAAGAAAGCTGGGTTTAATTGCAGTGATCAAGCAACAGAAG |
| hypothetical_las | F6 | Tb927.4.5380_for | GGGGACAAGTTTGTACAAAAAAGCAGGCTATGGCAACCAACGCGCGC | Tb927.4.5380_rev | GGGGACCACTTTGTACAAGAAAGCTGGGTTCATCTGGCGGTCCTCCTCG |
| hypothetical_las | F8 | Tb927.3.1800_for | GGGGACAAGTTTGTACAAAAAAGCAGGCTATGAAGCGGGAAAAGCGACAA | Tb927.3.1800_rev | GGGGACCACTTTGTACAAGAAAGCTGGGTTCATTGGCCTCTGTCACGTTTTC |
| hypothetical_las | F9 | Tb927.8.2390_for | GGGGACAAGTTTGTACAAAAAAGCAGGCTATGGCAACGCTTCGGTTCAA | Tb927.8.2390_rev | GGGGACCACTTTGTACAAGAAAGCTGGGTTCACTTCGTCGCCTTCTTTTTTTTG |
| hypothetical_las | F10 | Tb927.8.4320_for | GGGGACAAGTTTGTACAAAAAAGCAGGCTATGGAACGACTTGATGAACTCCAGC | Tb927.8.4320_rev | GGGGACCACTTTGTACAAGAAAGCTGGGTTCACGAGTCTTCTCTGTACGTACAA |
| hypothetical_las | F11 | Tb927.4.2400_for | GGGGACAAGTTTGTACAAAAAAGCAGGCTATGGAGGGAGAGCGTGGC | Tb927.4.2400_rev | GGGGACCACTTTGTACAAGAAAGCTGGGTCTAACTACCAGACATGAGGCTGCC |
| hypothetical_las | F12 | Tb927.7.2170_for | GGGGACAAGTTTGTACAAAAAAGCAGGCTATGTCTGGTTCCTTTGCTCAGC | Tb927.7.2170_rev | GGGGACCACTTTGTACAAGAAAGCTGGGTCTACTCATCCGCGACAGCCG |
| hypothetical_las | F13 | Tb927.6.3560_for | GGGGACAAGTTTGTACAAAAAAGCAGGCTATGGATGATTCGGATCCCCCAC | Tb927.6.3560_rev | GGGGACCACTTTGTACAAGAAAGCTGGGTTCACCTTCTCTGGAGCTCTCTTTT |
| hypothetical_las | F14 | Tb927.1.4480_for | GGGGACAAGTTTGTACAAAAAAGCAGGCTATGGAACATCCCCATTTGCAGG | Tb927.1.4480_rev | GGGGACCACTTTGTACAAGAAAGCTGGGTCTACAAGCCAACGTTGTCATGACT |
| hypothetical_las | F15 | Tb927.8.3700_for | GGGGACAAGTTTGTACAAAAAAGCAGGCTATGAGCGAGTCAGTGGGTACG | Tb927.8.3700_rev | GGGGACCACTTTGTACAAGAAAGCTGGGTTCAGCGAGGTGACATTTTTTCCTT |
| hypothetical_las | F16 | Tb927.10.12060_for | GGGGACAAGTTTGTACAAAAAAGCAGGCTATGCGGGGGAGCTGCAGC | Tb927.10.12060_rev | GGGGACCACTTTGTACAAGAAAGCTGGGTCTACCTACAAAGAACTGAAGCCACG |
| hypothetical_las | F17 | Tb927.8.2220_for | GGGGACAAGTTTGTACAAAAAAGCAGGCTATGGATACAACCAATGGCGGTG | Tb927.8.2220_rev | GGGGACCACTTTGTACAAGAAAGCTGGGTTCAAGCACACTGAACCTTAGTTGC |
| known_last | A1 | Tb927.9.12650_for | GGGGACAAGTTTGTACAAAAAAGCAGGCTATGAGTAAGATGGCTGGTGCTACG | Tb927.9.12650_rev | GGGGACCACTTTGTACAAGAAAGCTGGGTTTATTGTTGGATGCGCAAAAGTATTGT |
| known_last | A2 | Tb927.7.6080_for | GGGGACAAGTTTGTACAAAAAAGCAGGCTATGCCTCACAGAGCAGTTTGG | Tb927.7.6080_rev | GGGGACCACTTTGTACAAGAAAGCTGGGTTCATCCGCTGTTCTTGCGC |
| known_last | A3 | Tb927.1.1890_for | GGGGACAAGTTTGTACAAAAAAGCAGGCTATGCCGTCGCCCAAACTC | Tb927.1.1890_rev | GGGGACCACTTTGTACAAGAAAGCTGGGTCTAATCTATATGTAAAACAACAGTAGACTG |
| known_last | A4 | Tb927.10.10340_for | GGGGACAAGTTTGTACAAAAAAGCAGGCTATGCGCGTGAAGAGCATTGT | Tb927.10.10340_rev | GGGGACCACTTTGTACAAGAAAGCTGGGTCTAACGACTTCTATTTGACTGAATACGAGT |
| known_last | A5 | Tb927.10.9700_for | GGGGACAAGTTTGTACAAAAAAGCAGGCTATGAAGGACAGTTATCGCGATCCC | Tb927.10.9700_rev | GGGGACCACTTTGTACAAGAAAGCTGGGTCTACTGTCCATGATCGCCGTTC |
| known_last | A6 | Tb927.11.16100_for | GGGGACAAGTTTGTACAAAAAAGCAGGCTATGCGGTCGGGGGGGTTC | Tb927.11.16100_rev | GGGGACCACTTTGTACAAGAAAGCTGGGTTCAGCCCTCAACTAACTGCTCG |
| known_last | A7 | Tb927.4.3810_for | GGGGACAAGTTTGTACAAAAAAGCAGGCTATGATGGACGACGAACTACTTCTACA | Tb927.4.3810_rev | GGGGACCACTTTGTACAAGAAAGCTGGGTTTACAACGGCCCTGTGCCA |
| known_last | A8 | Tb927.4.450_for | GGGGACAAGTTTGTACAAAAAAGCAGGCTATGCTGACGAGATTTGATGTTCGC | Tb927.4.450_rev | GGGGACCACTTTGTACAAGAAAGCTGGGTTCACGTACGATTTGCAAGTCCAG |
| known_last | A9 | Tb927.11.10190_for | GGGGACAAGTTTGTACAAAAAAGCAGGCTATGGCACAAATGTTCCCCTCTG | Tb927.11.10190_rev | GGGGACCACTTTGTACAAGAAAGCTGGGTTCATGCAGAGCGCGTCGC |
| known_last | A10 | Tb927.11.10280_for | GGGGACAAGTTTGTACAAAAAAGCAGGCTATGACCTCTCGCATTGAGAAGAGT | Tb927.11.10280_rev | GGGGACCACTTTGTACAAGAAAGCTGGGTTTATTGCTGCAGTTGTTGCTGTTG |
| known_last | A11 | Tb927.11.14070_for | GGGGACAAGTTTGTACAAAAAAGCAGGCTATGACGAAAGAGGATCACGGTGA | Tb927.11.14070_rev | GGGGACCACTTTGTACAAGAAAGCTGGGTTCACAACAAAAATGCATATGAAAGTAAACT |
| known_last | A12 | Tb927.6.3550_for | GGGGACAAGTTTGTACAAAAAAGCAGGCTATGCGAATCGTAGCTCCGTTG | Tb927.6.3550_rev | GGGGACCACTTTGTACAAGAAAGCTGGGTCTACACAGAGTTTGCGTTCCTTATCG |
| known_last | A13 | Tb927.5.3510_for | GGGGACAAGTTTGTACAAAAAAGCAGGCTATGTATATCAAAAATATTCTTATTTCCGGA | Tb927.5.3510_rev | GGGGACCACTTTGTACAAGAAAGCTGGGTTCATTCAACATGCTCACGGACTC |
| known_last | A14 | Tb927.11.10670_for | GGGGACAAGTTTGTACAAAAAAGCAGGCTATGTCGTCCCGCGATTTGAG | Tb927.11.10670_rev | GGGGACCACTTTGTACAAGAAAGCTGGGTTCATTCCAACTGGTCCGGTGG |
| known_last | A15 | Tb927.7.2300_for | GGGGACAAGTTTGTACAAAAAAGCAGGCTATGTTCGCTTTCCTTCCTCCTGC | Tb927.7.2300_rev | GGGGACCACTTTGTACAAGAAAGCTGGGTTCATGACACGAGTCCAGCGG |
| known_last | A16 | Tb927.8.4870_for | GGGGACAAGTTTGTACAAAAAAGCAGGCTATGCCCGCTCCCCATAATTCG | Tb927.8.4870_rev | GGGGACCACTTTGTACAAGAAAGCTGGGTTTACTCCTCCTTCTTCGTGGAGAG |
| known_last | A17 | Tb927.7.7470_for | GGGGACAAGTTTGTACAAAAAAGCAGGCTATGAAAGCACCAGCCTTGCTG | Tb927.7.7470_rev | GGGGACCACTTTGTACAAGAAAGCTGGGTCTAAAAATCACCCAGCTCCCCCA |
| known_last | A18 | Tb927.5.330_for | GGGGACAAGTTTGTACAAAAAAGCAGGCTATGACCTCACTGCTGTCGAAGC | Tb927.5.330_rev | GGGGACCACTTTGTACAAGAAAGCTGGGTTCAGGAAGCATAACGACCAACAAA |
| known_last | A19 | Tb927.10.13040_for | GGGGACAAGTTTGTACAAAAAAGCAGGCTATGAAAGCACCAGCCTTGCTG | Tb927.10.13040_rev | GGGGACCACTTTGTACAAGAAAGCTGGGTTTAAAACTTATCAAAATCCGTGGTCCGA |
| known_last | A20 | Tb927.11.10730_for | GGGGACAAGTTTGTACAAAAAAGCAGGCTATGTCAAGTTCGCGAAAACGC | Tb927.11.10730_rev | GGGGACCACTTTGTACAAGAAAGCTGGGTTTACGAGTTAAATGTTTCCGTAAATAGAGG |
| known_last | A21 | Tb927.6.190_for | GGGGACAAGTTTGTACAAAAAAGCAGGCTATGTGTGTCCTGCAGTGGCG | Tb927.6.190_rev | GGGGACCACTTTGTACAAGAAAGCTGGGTTTATGCCTCATGTTCCGCAAAGT |
| known_last | A22 | Tb927.11.13740_for | GGGGACAAGTTTGTACAAAAAAGCAGGCTATGACCACCACGAAGGCTTCG | Tb927.11.13740_rev | GGGGACCACTTTGTACAAGAAAGCTGGGTTCACCGCTGCGCTTCGGG |
| known_last | A23 | Tb927.1.750_for | GGGGACAAGTTTGTACAAAAAAGCAGGCTATGGCGATGCGCAGCAAT | Tb927.1.750_rev | GGGGACCACTTTGTACAAGAAAGCTGGGTCTACCGTATCATTGGTAAGAAGCGC |
| known_last | A24 | Tb927.2.2240_for | GGGGACAAGTTTGTACAAAAAAGCAGGCTATGTCGCAACTGAGCTCCCC | Tb927.2.2240_rev | GGGGACCACTTTGTACAAGAAAGCTGGGTCTACGCTGCGTAATGTTGCCT |
| known_last | B1 | Tb927.9.5590_for | GGGGACAAGTTTGTACAAAAAAGCAGGCTATGGCGGAGGCACACAAGT | Tb927.9.5590_rev | GGGGACCACTTTGTACAAGAAAGCTGGGTCTAGAACAGCAGACGCGGC |
| known_last | B2 | Tb927.11.6170_for | GGGGACAAGTTTGTACAAAAAAGCAGGCTATGCGCTTAAAGGCCACACC | Tb927.11.6170_rev | GGGGACCACTTTGTACAAGAAAGCTGGGTCTACTGCAGAAACTTGAGGTTATTAATATG |
| known_last | B3 | Tb927.3.3870_for | GGGGACAAGTTTGTACAAAAAAGCAGGCTATGGAGAGGAGACAGGCGGG | Tb927.3.3870_rev | GGGGACCACTTTGTACAAGAAAGCTGGGTTTACATGGCATCCGGCTCGC |
| known_last | B4 | Tb927.10.3650_for | GGGGACAAGTTTGTACAAAAAAGCAGGCTATGCTCTCAACGAAGCAACTTCTC | Tb927.10.3650_rev | GGGGACCACTTTGTACAAGAAAGCTGGGTTCACAGCGTTCCCGACTCG |
| known_last | B5 | Tb927.4.3860_for | GGGGACAAGTTTGTACAAAAAAGCAGGCTATGATTTACCACAGTGAAACTGATGGT | Tb927.4.3860_rev | GGGGACCACTTTGTACAAGAAAGCTGGGTTTAATTCCCGTGAAGCTCACGC |
| known_last | B7 | Tb927.4.3880_for | GGGGACAAGTTTGTACAAAAAAGCAGGCTATGATTTACCACAGTGAAACTGATGGT | Tb927.4.3880_rev | GGGGACCACTTTGTACAAGAAAGCTGGGTTTAATTCCCGTGAAGTTCACGCTC |
| known_last | B8 | Tb927.11.4160_for | GGGGACAAGTTTGTACAAAAAAGCAGGCTATGGCAACACTGACGGTGAC | Tb927.11.4160_rev | GGGGACCACTTTGTACAAGAAAGCTGGGTTCAGTATCCACAAATAAGGGGTTCTCC |
| known_last | B9 | Tb927.5.320_for | GGGGACAAGTTTGTACAAAAAAGCAGGCTATGACCTCACTGCTGTCGAAGC | Tb927.5.320_rev | GGGGACCACTTTGTACAAGAAAGCTGGGTTCAATTAGCATCATTTTCCGTCATTCCG |
| known_last | B10 | Tb927.11.7210_for | GGGGACAAGTTTGTACAAAAAAGCAGGCTATGCACGTAGACGAGGGAAAGT | Tb927.11.7210_rev | GGGGACCACTTTGTACAAGAAAGCTGGGTTTACTTCTTTCTCCGTACGTTGTCCC |
| known_last | B11 | Tb927.4.3750_for | GGGGACAAGTTTGTACAAAAAAGCAGGCTATGTTTTCTTTCACGCCTCGGG | Tb927.4.3750_rev | GGGGACCACTTTGTACAAGAAAGCTGGGTCTATGTGGTACGCGCCTCCC |
| known_last | B12 | Tb927.11.1480_for | GGGGACAAGTTTGTACAAAAAAGCAGGCTATGCCGGCAGCTTTGTGC | Tb927.11.1480_rev | GGGGACCACTTTGTACAAGAAAGCTGGGTTCAAAGCATGGAAGAACATGTATCGA |
| known_last | B13 | Tb927.11.150_for | GGGGACAAGTTTGTACAAAAAAGCAGGCTATGTGCTACAGGGACGTCATTACA | Tb927.11.150_rev | GGGGACCACTTTGTACAAGAAAGCTGGGTCTAACCTTCATCAAGTAAAATTTTGTTATC |
| known_last | B14 | Tb927.10.5380_for | GGGGACAAGTTTGTACAAAAAAGCAGGCTATGCTTAACACTGTGAAATGGGCT | Tb927.10.5380_rev | GGGGACCACTTTGTACAAGAAAGCTGGGTCTACTCCTTTTTCTTCTCCATCGAGGC |
| known_last | B15 | Tb927.10.2430_for | GGGGACAAGTTTGTACAAAAAAGCAGGCTATGCACGAATCGACAGTGGG | Tb927.10.2430_rev | GGGGACCACTTTGTACAAGAAAGCTGGGTTTAAGAGGAAGAACCGCGTGGC |
| known_last | B16 | Tb927.4.3160_for | GGGGACAAGTTTGTACAAAAAAGCAGGCTATGAAACAAGGTAACGGGGCCA | Tb927.4.3160_rev | GGGGACCACTTTGTACAAGAAAGCTGGGTCTACATCCTTGATGCCCGCTTG |
| known_last | B17 | Tb927.5.3670_for | GGGGACAAGTTTGTACAAAAAAGCAGGCTATGCCACAGACTGTTAGCAGC | Tb927.5.3670_rev | GGGGACCACTTTGTACAAGAAAGCTGGGTTTATGTCTGAGTAGGTGGCACAGA |
| known_last | B18 | Tb927.6.770_for | GGGGACAAGTTTGTACAAAAAAGCAGGCTATGTGCCGGCAGGAAGGT | Tb927.6.770_rev | GGGGACCACTTTGTACAAGAAAGCTGGGTTTAGTATTCCTGAAGGTGCATGTCGT |
| known_last | B19 | Tb927.4.4430_for | GGGGACAAGTTTGTACAAAAAAGCAGGCTATGGTTACCTTCATGCGTGAATTGT | Tb927.4.4430_rev | GGGGACCACTTTGTACAAGAAAGCTGGGTTTAGCTGGCAGCGCTAACATTT |

| | | | | | |
|---|---|---|---|---|---|
| known_last | B20 | Tb927.7.7520_for | GGGGACAAGTTTGTACAAAAAAGCAGGCTATGGTTATCTTCATGCGTGAATTGTATGT | Tb927.7.7520_rev | GGGGACCACTTTGTACAAGAAAGCTGGGTTTAGCTGGCGGCGCTAGC |
| known_last | B24 | Tb927.5.4550_for | GGGGACAAGTTTGTACAAAAAAGCAGGCTATGATTTCCATTCATCAGTCGATTCTCTT | Tb927.5.4550_rev | GGGGACCACTTTGTACAAGAAAGCTGGGTCTAAGCTTCAGGGCTCTGGGT |
| known_last | C4 | Tb927.4.4410_for | GGGGACAAGTTTGTACAAAAAAGCAGGCTATGGTTACCTTCATGCGTGAATTGT | Tb927.4.4410_rev | GGGGACCACTTTGTACAAGAAAGCTGGGTTTAGCTGGCAGCGCTAACATTATC |
| known_last | C5 | Tb927.11.5940_for | GGGGACAAGTTTGTACAAAAAAGCAGGCTATGAACGATTACCGTTTGTGTGGT | Tb927.11.5940_rev | GGGGACCACTTTGTACAAGAAAGCTGGGTTCACACCGACACAAAGGGCT |
| known_last | C6 | Tb927.6.430_for | GGGGACAAGTTTGTACAAAAAAGCAGGCTATGAGTTGGCAGGAAGGTGGT | Tb927.6.430_rev | GGGGACCACTTTGTACAAGAAAGCTGGGTTTAAATAGAATAATTTTCACGGCAGTACAG |
| known_last | C7 | Tb927.6.800_for | GGGGACAAGTTTGTACAAAAAAGCAGGCTATGTGCCGGCAGGAAGGT | Tb927.6.800_rev | GGGGACCACTTTGTACAAGAAAGCTGGGTTTAAAAAGAACACATTTCACGGTAGCAAG |
| known_last | C9 | Tb927.3.3860_for | GGGGACAAGTTTGTACAAAAAAGCAGGCTATGCCTATCCCAACGGTAACCA | Tb927.3.3860_rev | GGGGACCACTTTGTACAAGAAAGCTGGGTTTACACGTTCGCAGCTTGGTG |
| known_last | C10 | Tb927.1.4010_for | GGGGACAAGTTTGTACAAAAAAGCAGGCTATGTTTCACCTTCCCAGGGCG | Tb927.1.4010_rev | GGGGACCACTTTGTACAAGAAAGCTGGGTCTAACAACCCGGTACAGAACGC |
| known_last | C11 | Tb927.5.1450_for | GGGGACAAGTTTGTACAAAAAAGCAGGCTATGAACGACAGGACGAGTTTACTCG | Tb927.5.1450_rev | GGGGACCACTTTGTACAAGAAAGCTGGGTTCATAACGTTACCGTAAAGCCATCAG |
| known_last | C12 | Tb927.8.7940_for | GGGGACAAGTTTGTACAAAAAAGCAGGCTATGGCTTCCTTCATGCGTCTTG | Tb927.8.7940_rev | GGGGACCACTTTGTACAAGAAAGCTGGGTCTAATGAGAACACCTTCCAGCATCAT |
| known_last | C13 | Tb927.6.740_for | GGGGACAAGTTTGTACAAAAAAGCAGGCTATGAACCAGAAGCGATGCGAAG | Tb927.6.740_rev | GGGGACCACTTTGTACAAGAAAGCTGGGTTCACTCCTGCGCAGTGTCC |
| known_last | C14 | Tb927.5.3320_for | GGGGACAAGTTTGTACAAAAAAGCAGGCTATGCGCAATGTTTCTTCCCTCG | Tb927.5.3320_rev | GGGGACCACTTTGTACAAGAAAGCTGGGTTCATGCATCAACGGCAACAACG |
| known_last | C15 | Tb927.11.8150_for | GGGGACAAGTTTGTACAAAAAAGCAGGCTATGAACAATTATCATATTTACGACGAAATT | Tb927.11.8150_rev | GGGGACCACTTTGTACAAGAAAGCTGGGTCTACGGACGTAAAAGTTCTTTAATACCCC |
| known_last | C16 | Tb927.9.7150_for | GGGGACAAGTTTGTACAAAAAAGCAGGCTATGTCAGTCGGCAAGGTCGC | Tb927.9.7150_rev | GGGGACCACTTTGTACAAGAAAGCTGGGTTTAAGCACCCAACCCTACCGG |
| known_last | C17 | Tb927.6.200_for | GGGGACAAGTTTGTACAAAAAAGCAGGCTATGTGCTGGCAGGAAGGTGG | Tb927.6.200_rev | GGGGACCACTTTGTACAAGAAAGCTGGGTCTAATGTAACGACGGAAGCGGAC |
| known_last | C18 | Tb927.8.7870_for | GGGGACAAGTTTGTACAAAAAAGCAGGCTATGGCTTCCTTCATGCGTCTTG | Tb927.8.7870_rev | GGGGACCACTTTGTACAAGAAAGCTGGGTTCAATGCTTAGGGGAATATTTGCTGG |
| known_last | C20 | Tb927.1.1910_for | GGGGACAAGTTTGTACAAAAAAGCAGGCTATGTTACCATCCTCTCAGGACCAAC | Tb927.1.1910_rev | GGGGACCACTTTGTACAAGAAAGCTGGGTTCAAACATCTCCGGTATCAGATTGACG |
| known_last | C21 | Tb927.9.4900_for | GGGGACAAGTTTGTACAAAAAAGCAGGCTATGTCCGATCTCCTTGAGTTCTTTGT | Tb927.9.4900_rev | GGGGACCACTTTGTACAAGAAAGCTGGGTTTACAAGCCGTCAGCCGTAGT |
| known_last | C22 | Tb927.11.10520_for | GGGGACAAGTTTGTACAAAAAAGCAGGCTATGTTCAATGTCTCACCAGCGAG | Tb927.11.10520_rev | GGGGACCACTTTGTACAAGAAAGCTGGGTTCATTTCCGGTAGACGTTGCG |
| known_last | C23 | Tb927.5.3030_for | GGGGACAAGTTTGTACAAAAAAGCAGGCTATGCATCTCTACTTAACCAAGCGAG | Tb927.5.3030_rev | GGGGACCACTTTGTACAAGAAAGCTGGGTCTAGTGACGAAGAGGTGTGTGACA |
| known_last | C24 | Tb927.8.650_for | GGGGACAAGTTTGTACAAAAAAGCAGGCTATGATAACAAAAAATTCCATAATCAAAGAG | Tb927.8.650_rev | GGGGACCACTTTGTACAAGAAAGCTGGGTTCAATCATACTCCGTCAGAAGTTTCAG |
| known_last | D1 | Tb927.3.2690_for | GGGGACAAGTTTGTACAAAAAAGCAGGCTATGAATTTCTATGAACGAGTAGCGAGTG | Tb927.3.2690_rev | GGGGACCACTTTGTACAAGAAAGCTGGGTTTACGTGTTCCCAACACCCTCC |
| known_last | D2 | Tb927.8.7920_for | GGGGACAAGTTTGTACAAAAAAGCAGGCTATGGCTTCCTTCATGCGTCTTG | Tb927.8.7920_rev | GGGGACCACTTTGTACAAGAAAGCTGGGTTTATTCTGGCAATAATGGACACATATTACA |
| known_last | D3 | Tb927.8.7900_for | GGGGACAAGTTTGTACAAAAAAGCAGGCTATGGCTTCCTTCATGCGTCTTG | Tb927.8.7900_rev | GGGGACCACTTTGTACAAGAAAGCTGGGTTTATTCTGGCAATAATGGACACATATTGCA |
| known_last | D4 | Tb927.8.7930_for | GGGGACAAGTTTGTACAAAAAAGCAGGCTATGGCTTCCTTCATGCGTCTTG | Tb927.8.7930_rev | GGGGACCACTTTGTACAAGAAAGCTGGGTTTATTCTGGCAATAACGGACACATATTACA |
| known_last | D5 | Tb927.8.7890_for | GGGGACAAGTTTGTACAAAAAAGCAGGCTATGGCTTCCTTCATGCGTCTTG | Tb927.8.7890_rev | GGGGACCACTTTGTACAAGAAAGCTGGGTTTATTCTGGCAATAACGGACGCA |
| known_last | D6 | Tb927.7.6070_for | GGGGACAAGTTTGTACAAAAAAGCAGGCTATGCCTCACAGAGCAGTTTGG | Tb927.7.6070_rev | GGGGACCACTTTGTACAAGAAAGCTGGGTCTAGAGGTCACCCTTCAGCACC |
| known_last | D7 | Tb927.8.7590_for | GGGGACAAGTTTGTACAAAAAAGCAGGCTATGGGCTTTAGGGCTGGCT | Tb927.8.7590_rev | GGGGACCACTTTGTACAAGAAAGCTGGGTTTAAAATGGGCCCTTGCATTGGT |
| known_last | D9 | Tb927.6.330_for | GGGGACAAGTTTGTACAAAAAAGCAGGCTATGAGTTGGCAGGAAGGTGGT | Tb927.6.330_rev | GGGGACCACTTTGTACAAGAAAGCTGGGTTTAACAGGTACTACCAACACTTTCCAGG |
| known_last | D10 | Tb927.6.280_for | GGGGACAAGTTTGTACAAAAAAGCAGGCTATGTGCTGGCAGGAAGGTGG | Tb927.6.280_rev | GGGGACCACTTTGTACAAGAAAGCTGGGTTTAACAGGTACTACCAACACTTTCCAGG |
| known_last | D12 | Tb927.7.6050_for | GGGGACAAGTTTGTACAAAAAAGCAGGCTATGCCTCACAGAGCAGTTTGG | Tb927.7.6050_rev | GGGGACCACTTTGTACAAGAAAGCTGGGTCTAGAGGTCACCCTTCAGCGA |
| known_last | D14 | Tb927.6.5110_for | GGGGACAAGTTTGTACAAAAAAGCAGGCTATGCCGTTTTTGGCGTCGA | Tb927.6.5110_rev | GGGGACCACTTTGTACAAGAAAGCTGGGTCTAGTGAATTCTTTGGATGGTTTGTATAAA |
| known_last | D15 | Tb927.7.6060_for | GGGGACAAGTTTGTACAAAAAAGCAGGCTATGCCTCACAGAGCAGTTTGG | Tb927.7.6060_rev | GGGGACCACTTTGTACAAGAAAGCTGGGTCTAGAGGTCACCCTTCATCGACG |
| known_last | D16 | Tb927.11.6350_for | GGGGACAAGTTTGTACAAAAAAGCAGGCTATGGTTGTGACCCGCTCCC | Tb927.11.6350_rev | GGGGACCACTTTGTACAAGAAAGCTGGGTTCATCCGTTTGAGCTTACCACCT |
| known_last | D17 | Tb927.4.4470_for | GGGGACAAGTTTGTACAAAAAAGCAGGCTATGCTTAAATTACTTCTACTGCCATTACTC | Tb927.4.4470_rev | GGGGACCACTTTGTACAAGAAAGCTGGGTTCAACTATTTTTTCCATTGTCCCCGA |
| known_last | D18 | Tb927.6.310_for | GGGGACAAGTTTGTACAAAAAAGCAGGCTATGAGCTGGCAGGAAGGTGG | Tb927.6.310_rev | GGGGACCACTTTGTACAAGAAAGCTGGGTTCATGAAGGATATTGGCTGGCAC |
| known_last | D19 | Tb927.10.3390_for | GGGGACAAGTTTGTACAAAAAAGCAGGCTATGAGCGGCCGCGGAGGA | Tb927.10.3390_rev | GGGGACCACTTTGTACAAGAAAGCTGGGTCTACTCTCCAACGACAATCACCTCA |
| known_last | D20 | Tb927.6.300_for | GGGGACAAGTTTGTACAAAAAAGCAGGCTATGTGCTGGCAGGAAGGTGG | Tb927.6.300_rev | GGGGACCACTTTGTACAAGAAAGCTGGGTTCATGAAGGATATTGGCTGGCAC |
| known_last | D21 | Tb927.11.5780_for | GGGGACAAGTTTGTACAAAAAAGCAGGCTATGCGGCGGTTGAGTCTGG | Tb927.11.5780_rev | GGGGACCACTTTGTACAAGAAAGCTGGGTCTAAGAGAAAAAATATGGGGAGTTTAGGAC |
| known_last | D22 | Tb927.9.11850_for | GGGGACAAGTTTGTACAAAAAAGCAGGCTATGTCGTCGCGGATAGAGCG | Tb927.9.11850_rev | GGGGACCACTTTGTACAAGAAAGCTGGGTCTAATAACCATTTATGTGTTTATTGCCACC |
| known_last | D23 | Tb927.10.8590_for | GGGGACAAGTTTGTACAAAAAAGCAGGCTATGCAAAGTGGTGTGGGTAACATT | Tb927.10.8590_rev | GGGGACCACTTTGTACAAGAAAGCTGGGTTCACGGAACTGGTGCCGG |
| known_last | D24 | Tb927.5.650_for | GGGGACAAGTTTGTACAAAAAAGCAGGCTATGTTTCTCCTTTCCCGCACTG | Tb927.5.650_rev | GGGGACCACTTTGTACAAGAAAGCTGGGTTCACTCCCCATTCCTTGACGT |
| known_last | E1 | Tb927.11.13360_for | GGGGACAAGTTTGTACAAAAAAGCAGGCTATGCCGAAACGAAGCCGTC | Tb927.11.13360_rev | GGGGACCACTTTGTACAAGAAAGCTGGGTTTAGGCGAACATTGAGTTGTGCG |
| known_last | E2 | Tb927.4.2460_for | GGGGACAAGTTTGTACAAAAAAGCAGGCTATGGGTGCTTGCATAGCCG | Tb927.4.2460_rev | GGGGACCACTTTGTACAAGAAAGCTGGGTCTACACCCTTACATTGGGACTTTCTC |
| known_last | E3 | Tb927.4.770_for | GGGGACAAGTTTGTACAAAAAAGCAGGCTATGTTTCTATCCCTTGATGTCACGC | Tb927.4.770_rev | GGGGACCACTTTGTACAAGAAAGCTGGGTTTACGACAAACTGAAGTTCCTTCTGC |
| known_last | E4 | Tb927.4.4460_for | GGGGACAAGTTTGTACAAAAAAGCAGGCTATGGTTACCTTCATGCGTGAATTGT | Tb927.4.4460_rev | GGGGACCACTTTGTACAAGAAAGCTGGGTTCAACTATTTTTTCCATTGTCCCCGA |
| known_last | E5 | Tb927.10.14410_for | GGGGACAAGTTTGTACAAAAAAGCAGGCTATGAACGTTGATTTATCGAACAAGTCTATC | Tb927.10.14410_rev | GGGGACCACTTTGTACAAGAAAGCTGGGTTTAAATGAAGACTGGGCGCTGAG |
| known_last | E6 | Tb927.11.12110_for | GGGGACAAGTTTGTACAAAAAAGCAGGCTATGGGCCCCAACAGACAAAGG | Tb927.11.12110_rev | GGGGACCACTTTGTACAAGAAAGCTGGGTTCAACGCATGGGGGTCGT |
| known_last | E7 | Tb927.9.4080_for | GGGGACAAGTTTGTACAAAAAAGCAGGCTATGGCATCGGAGGCAGCG | Tb927.9.4080_rev | GGGGACCACTTTGTACAAGAAAGCTGGGTTTATTCGCCCCTAACTGCCTCC |
| known_last | E8 | Tb927.10.8170_for | GGGGACAAGTTTGTACAAAAAAGCAGGCTATGGAAGTACGAGATGATAACCCGA | Tb927.10.8170_rev | GGGGACCACTTTGTACAAGAAAGCTGGGTCTAGAAAGTAAACCTGGAACTGGCAC |
| known_last | E9 | Tb927.9.13610_for | GGGGACAAGTTTGTACAAAAAAGCAGGCTATGCGGAAAGACGAATACCGG | Tb927.9.13610_rev | GGGGACCACTTTGTACAAGAAAGCTGGGTCTATTTGTAGCTCCTTGGGTGCTG |
| known_last | E10 | Tb927.5.2410_for | GGGGACAAGTTTGTACAAAAAAGCAGGCTATGAGCAAAGCTGCGGTTTCA | Tb927.5.2410_rev | GGGGACCACTTTGTACAAGAAAGCTGGGTTTAAGTTCTAACAAGACTTTCCTGGCTAAG |
| known_last | E11 | Tb927.8.4880_for | GGGGACAAGTTTGTACAAAAAAGCAGGCTATGGAGGATTGGGTTAGCTGCC | Tb927.8.4880_rev | GGGGACCACTTTGTACAAGAAAGCTGGGTTCATGGAGTATCGAACAACTTGTCAAG |
| known_last | E12 | Tb927.9.5520_for | GGGGACAAGTTTGTACAAAAAAGCAGGCTATGAATCAGAACAATTTATCTTGCGAGGA | Tb927.9.5520_rev | GGGGACCACTTTGTACAAGAAAGCTGGGTTCACCTTCTTGAAATACGTAACGCC |
| known_last | E13 | Tb927.7.3000_for | GGGGACAAGTTTGTACAAAAAAGCAGGCTATGACGTCTCAAACGTCGGC | Tb927.7.3000_rev | GGGGACCACTTTGTACAAGAAAGCTGGGTTCATCCCTCCCCGTCGATACA |
| known_last | E14 | Tb927.8.6200_for | GGGGACAAGTTTGTACAAAAAAGCAGGCTATGGATGGTGAACAATTAGAAACAGATCC | Tb927.8.6200_rev | GGGGACCACTTTGTACAAGAAAGCTGGGTTCAGTACCCTGCCTCATGTACG |
| known_last | E15 | Tb927.9.12400_for | GGGGACAAGTTTGTACAAAAAAGCAGGCTATGTGTGCCTTAAGGGATATCGCG | Tb927.9.12400_rev | GGGGACCACTTTGTACAAGAAAGCTGGGTTTAGTTTACCATAAAGTCATGGAACAAAAG |
| known_last | E16 | Tb927.10.13330_for | GGGGACAAGTTTGTACAAAAAAGCAGGCTATGGATCAATCATGCAACGAAGCT | Tb927.10.13330_rev | GGGGACCACTTTGTACAAGAAAGCTGGGTCTATCTGCAACCGTCACTGCG |
| known_last | E17 | Tb927.11.8210_for | GGGGACAAGTTTGTACAAAAAAGCAGGCTATGACGAGTATCGAACAGATTGAAATCA | Tb927.11.8210_rev | GGGGACCACTTTGTACAAGAAAGCTGGGTTCAACTCGCGAACAACTGATCG |
| known_last | E18 | Tb927.11.260_for | GGGGACAAGTTTGTACAAAAAAGCAGGCTATGAGCATTCAAAAAAAGTTTGAGCTTACG | Tb927.11.260_rev | GGGGACCACTTTGTACAAGAAAGCTGGGTCTAAAGAAGATGAGTCTTCAAATCAGGATC |
| known_last | E19 | Tb927.9.7720_for | GGGGACAAGTTTGTACAAAAAAGCAGGCTATGCCCTCCTTAATACGTACCAACT | Tb927.9.7720_rev | GGGGACCACTTTGTACAAGAAAGCTGGGTTTAGGGTCGCAGCAAACTGTG |
| known_last | E20 | Tb927.11.14630_for | GGGGACAAGTTTGTACAAAAAAGCAGGCTATGCCAATCTTGTGCGATTGCC | Tb927.11.14630_rev | GGGGACCACTTTGTACAAGAAAGCTGGGTTTACCACCATCCTCTGTGTACCG |
| known_last | E21 | Tb927.10.740_for | GGGGACAAGTTTGTACAAAAAAGCAGGCTATGAGCTGCGAAAGCAACGG | Tb927.10.740_rev | GGGGACCACTTTGTACAAGAAAGCTGGGTTTAGGCATCTTCCTCTACCTTCAAACG |
| known_last | E22 | Tb927.11.9190_for | GGGGACAAGTTTGTACAAAAAAGCAGGCTATGGGAAATCAGCTCGCAGC | Tb927.11.9190_rev | GGGGACCACTTTGTACAAGAAAGCTGGGTTTACTCAGCGTTGAACCACAGTG |
| known_last | E23 | Tb927.11.11880_for | GGGGACAAGTTTGTACAAAAAAGCAGGCTATGGGCACCGTGTCTCCG | Tb927.11.11880_rev | GGGGACCACTTTGTACAAGAAAGCTGGGTTTACTCGGCTGCGGCTACC |
| known_last | E24 | Tb927.8.710_for | GGGGACAAGTTTGTACAAAAAAGCAGGCTATGATAACTCCAGAAACGGCAGGA | Tb927.8.710_rev | GGGGACCACTTTGTACAAGAAAGCTGGGTTTAACGACCATCGTCCATCAACAC |
| known_last | F1 | Tb927.3.3850_for | GGGGACAAGTTTGTACAAAAAAGCAGGCTATGAATACAAACCGGCCGAACAC | Tb927.3.3850_rev | GGGGACCACTTTGTACAAGAAAGCTGGGTTCAGGAGGAGTTAAATGTGTCCACA |
| known_last | F2 | Tb927.3.4230_for | GGGGACAAGTTTGTACAAAAAAGCAGGCTATGTCGTCCAGTCAAAACCATCTG | Tb927.3.4230_rev | GGGGACCACTTTGTACAAGAAAGCTGGGTTCATGGTAGTCTTTCCATCTCAATATATTC |
| known_last | F3 | Tb927.1.370_for | GGGGACAAGTTTGTACAAAAAAGCAGGCTATGTCAGGAAACCAAGGAAGAAAGC | Tb927.1.370_rev | GGGGACCACTTTGTACAAGAAAGCTGGGTTTAACGGATATGCACCCTCTTTTTTCT |
| known_last | F4 | Tb927.1.290_for | GGGGACAAGTTTGTACAAAAAAGCAGGCTATGTCAGGAAACCAAGGAAGAAAGC | Tb927.1.290_rev | GGGGACCACTTTGTACAAGAAAGCTGGGTTTAACGGATATGTACTCTCCTTTTTCTCAG |
| known_last | F5 | Tb927.2.170_for | GGGGACAAGTTTGTACAAAAAAGCAGGCTATGTCAGGAAACCAAGGAAGAAAGC | Tb927.2.170_rev | GGGGACCACTTTGTACAAGAAAGCTGGGTTTAGGTATGTACCCTCCTTTTTCTCAGT |
| known_last | F6 | Tb927.1.480_for | GGGGACAAGTTTGTACAAAAAAGCAGGCTATGTCAGGAAACCAAGGAAGAAAGC | Tb927.1.480_rev | GGGGACCACTTTGTACAAGAAAGCTGGGTTTAACGGATATGTACCCTTCTTTTTCTCAG |
| known_last | F7 | Tb927.2.950_for | GGGGACAAGTTTGTACAAAAAAGCAGGCTATGTCAGGAAACCAAGGAAGAAAGC | Tb927.2.950_rev | GGGGACCACTTTGTACAAGAAAGCTGGGTTTAATATATATGTACCCTCCTTTTTCCCAG |
| known_last | F8 | Tb927.9.4370_for | GGGGACAAGTTTGTACAAAAAAGCAGGCTATGCTGGTCGCCACAGCG | Tb927.9.4370_rev | GGGGACCACTTTGTACAAGAAAGCTGGGTTCAAGTAGGGACCATTTCCACAATG |
| known_last | F9 | Tb927.11.4690_for | GGGGACAAGTTTGTACAAAAAAGCAGGCTATGCGGCTAAATAGCTGCTGG | Tb927.11.4690_rev | GGGGACCACTTTGTACAAGAAAGCTGGGTTTACACCGTAATTTCTACACTGTCAGTAAC |
| known_last | F10 | Tb927.8.6070_for | GGGGACAAGTTTGTACAAAAAAGCAGGCTATGGCACAACTTGACCAGTTGC | Tb927.8.6070_rev | GGGGACCACTTTGTACAAGAAAGCTGGGTTCAATCGACACCTCCAAACATGAAC |
| known_last | F11 | Tb927.11.15750_for | GGGGACAAGTTTGTACAAAAAAGCAGGCTATGTCGGATCCAACGCCCC | Tb927.11.15750_rev | GGGGACCACTTTGTACAAGAAAGCTGGGTTTAAGAATCCCTGCTCTCGTCGT |
| known_last | F12 | Tb927.9.9740_for | GGGGACAAGTTTGTACAAAAAAGCAGGCTATGAGTGTGTCTCTGGCTCCC | Tb927.9.9740_rev | GGGGACCACTTTGTACAAGAAAGCTGGGTCTATCGGTGCAGAGGAGGTAACG |
| known_last | F13 | Tb927.7.4900_for | GGGGACAAGTTTGTACAAAAAAGCAGGCTATGGGTGTTCCAAAATTCTTTCGATGG | Tb927.7.4900_rev | GGGGACCACTTTGTACAAGAAAGCTGGGTTCATCGTTTCCTGTCTCCTGATAAATG |
| known_last | F14 | Tb927.8.5620_for | GGGGACAAGTTTGTACAAAAAAGCAGGCTATGCTTGATCTAGATAGCTGGGTGG | Tb927.8.5620_rev | GGGGACCACTTTGTACAAGAAAGCTGGGTCTACAGTGCCGCTTCACACAT |
| known_last | F15 | Tb927.11.11540_for | GGGGACAAGTTTGTACAAAAAAGCAGGCTATGTCGGGTCGCACCGTT | Tb927.11.11540_rev | GGGGACCACTTTGTACAAGAAAGCTGGGTTCAAAATACGCAGGCATCAAACCC |
| known_last | F16 | Tb927.5.3160_for | GGGGACAAGTTTGTACAAAAAAGCAGGCTATGACCACAACGCCAGGAGG | Tb927.5.3160_rev | GGGGACCACTTTGTACAAGAAAGCTGGGTTCAAACGTTTTTGGAAACTGTCAGTG |
| known_last | F17 | Tb927.8.1130_for | GGGGACAAGTTTGTACAAAAAAGCAGGCTATGAGCCATGAATTCAGTGGCTC | Tb927.8.1130_rev | GGGGACCACTTTGTACAAGAAAGCTGGGTTTAACCAGAATGGTCACCGTCACC |
| known_last | F18 | Tb927.11.11080_for | GGGGACAAGTTTGTACAAAAAAGCAGGCTATGTCTGAAATAGCTCACTCTGCG | Tb927.11.11080_rev | GGGGACCACTTTGTACAAGAAAGCTGGGTTCAATAGGGTTTAGTGAAAGCATTCTCC |
| known_last | F19 | Tb927.4.3020_for | GGGGACAAGTTTGTACAAAAAAGCAGGCTATGTTCACACGCGCTCTCAG | Tb927.4.3020_rev | GGGGACCACTTTGTACAAGAAAGCTGGGTTCACACCTCGTGTAACTCCCAT |
| known_last | F20 | Tb927.1.1880_for | GGGGACAAGTTTGTACAAAAAAGCAGGCTATGAATGAATCCCACATGTGTCATCA | Tb927.1.1880_rev | GGGGACCACTTTGTACAAGAAAGCTGGGTTCACTCTTTCATCATAATAGGCACGC |
| known_last | F21 | Tb927.11.1460_for | GGGGACAAGTTTGTACAAAAAAGCAGGCTATGCGCAGTGCGCTTCCA | Tb927.11.1460_rev | GGGGACCACTTTGTACAAGAAAGCTGGGTTTAAGCCAAACTGCATGAATCCCC |
| known_last | F22 | Tb927.11.14560_for | GGGGACAAGTTTGTACAAAAAAGCAGGCTATGCTGCATGCAACTGTGGC | Tb927.11.14560_rev | GGGGACCACTTTGTACAAGAAAGCTGGGTTCAAAACAAACCACATTCTTCATGCAC |
| known_last | F23 | Tb927.11.11550_for | GGGGACAAGTTTGTACAAAAAAGCAGGCTATGTCGGGTCGCACCGTT | Tb927.11.11550_rev | GGGGACCACTTTGTACAAGAAAGCTGGGTTTAGTCACTGAAGTCAAGAGAGGACTC |
| known_last | F24 | Tb927.8.4950_for | GGGGACAAGTTTGTACAAAAAAGCAGGCTATGGAGTACAAGCGATGCAACTC | Tb927.8.4950_rev | GGGGACCACTTTGTACAAGAAAGCTGGGTCTAGCAGGTCAGAGACTCCTCAG |
| known_last | G1 | Tb927.11.14210_for | GGGGACAAGTTTGTACAAAAAAGCAGGCTATGTCACAAAGATCGAGGAAAGTTCC | Tb927.11.14210_rev | GGGGACCACTTTGTACAAGAAAGCTGGGTTTAGCTCGACATCTTCGAGCGT |
| known_last | G2 | Tb927.11.11090_for | GGGGACAAGTTTGTACAAAAAAGCAGGCTATGTCCCTGGACCCCCGT | Tb927.11.11090_rev | GGGGACCACTTTGTACAAGAAAGCTGGGTCTACTGCGAAAAGACACTTTTAAAATCAGG |
| known_last | G3 | Tb927.11.7290_for | GGGGACAAGTTTGTACAAAAAAGCAGGCTATGATCTCCTCCAGTCAAGGGGG | Tb927.11.7290_rev | GGGGACCACTTTGTACAAGAAAGCTGGGTTCAGGACACATGTTCCTCTGTGT |
| known_last | G4 | Tb927.11.3780_for | GGGGACAAGTTTGTACAAAAAAGCAGGCTATGACATGCATACAGGAAAACATTACAGG | Tb927.11.3780_rev | GGGGACCACTTTGTACAAGAAAGCTGGGTCTACTCAGATGGTTCCACTGCCA |
| known_last | G5 | Tb927.3.5440_for | GGGGACAAGTTTGTACAAAAAAGCAGGCTATGAAAGGAAATGACAACGACTGTAACG | Tb927.3.5440_rev | GGGGACCACTTTGTACAAGAAAGCTGGGTCTACTTGATGACGCGCACCG |
| known_last | G6 | Tb927.11.5190_for | GGGGACAAGTTTGTACAAAAAAGCAGGCTATGGTGTATGAGAGTGAATCTTGCAG | Tb927.11.5190_rev | GGGGACCACTTTGTACAAGAAAGCTGGGTTTAACGGGCACCGTGTAGCC |
