## Supplementary material for "A *Trypanosoma brucei* ORFeome-based Gain-of-Function Library reveals novel genes associated with melarsoprol resistance": SUP. 2

**SUP. 2 - ORFeome PCR Amplification Tracking**

| Plate name | Min length* | Max length* | # Anticipated | # Positive (SYBR) | Missing | % Positive (SYBR) |
|---|---|---|---|---|---|---|
| **0_hypothetical** | 102 | 375 | 373 | 369 | 4 | 99% |
| **0_known** | 147 | 591 | 315 | 308 | 7 | 98% |
| **1_hypothetical** | 375 | 522 | 374 | 362 | 12 | 97% |
| **1_known** | 594 | 849 | 365 | 355 | 10 | 97% |
| **2_hypothetical** | 522 | 666 | 372 | 343 | 29 | 92% |
| **2_known** | 849 | 1056 | 363 | 351 | 12 | 97% |
| **3_hypothetical** | 669 | 822 | 382 | 374 | 8 | 98% |
| **3_known** | 1056 | 1287 | 367 | 333 | 34 | 91% |
| **4_hypothetical** | 822 | 993 | 372 | 344 | 28 | 92% |
| **4_known** | 1287 | 1524 | 348 | 344 | 4 | 99% |
| **5_hypothetical** | 993 | 1155 | 365 | 346 | 19 | 95% |
| **5_known** | 1524 | 1857 | 342 | 283 | 59 | 83% |
| **6_hypothetical** | 1158 | 1365 | 375 | 353 | 22 | 94% |
| **6_known** | 1857 | 2337 | 362 | 331 | 31 | 91% |
| **7_hypothetical** | 1368 | 1635 | 378 | 314 | 64 | 83% |
| **7_known** | 2340 | 3504 | 376 | 310 | 66 | 82% |
| **8_hypothetical** | 1635 | 1953 | 381 | 375 | 6 | 98% |
| **9_hypothetical** | 1953 | 2508 | 381 | 380 | 1 | 100% |
| **10_hypothetical** | 2508 | 3501 | 381 | 378 | 3 | 99% |
| **hypothetical_last** | 3504 | 4488 | 135 | 132 | 3 | 98% |
| **known_last** | 3507 | 4497 | 138 | 135 | 3 | 98% |
| * base pairs | | ORFs | 7245 | 6820 | 425 | 94.3% |

**REDO PCR REACTIONS**

| **NEGATIVE PICKS** | **Total #** | **# Positive** | **# Missing** |
|---|---|---|---|
| NEG_PICKS_#1 | 252 | 130 | 122 |
| NEG_PICKS_#2 | 177 | 98 | 79 |
| Total | 429 | 228 | 201 |

**FINAL TOTALS**

| **TOTAL ORFs** | 7245 |
|---|---|
| Total Missing | 201 |
| Total Positive | 7044 |
| **Percent Positive** | **97.2%** |
