## Supplementary figures and images for "A *Trypanosoma brucei* ORFeome-based Gain-of-Function Library reveals novel genes associated with melarsoprol resistance"

### SUP. 3

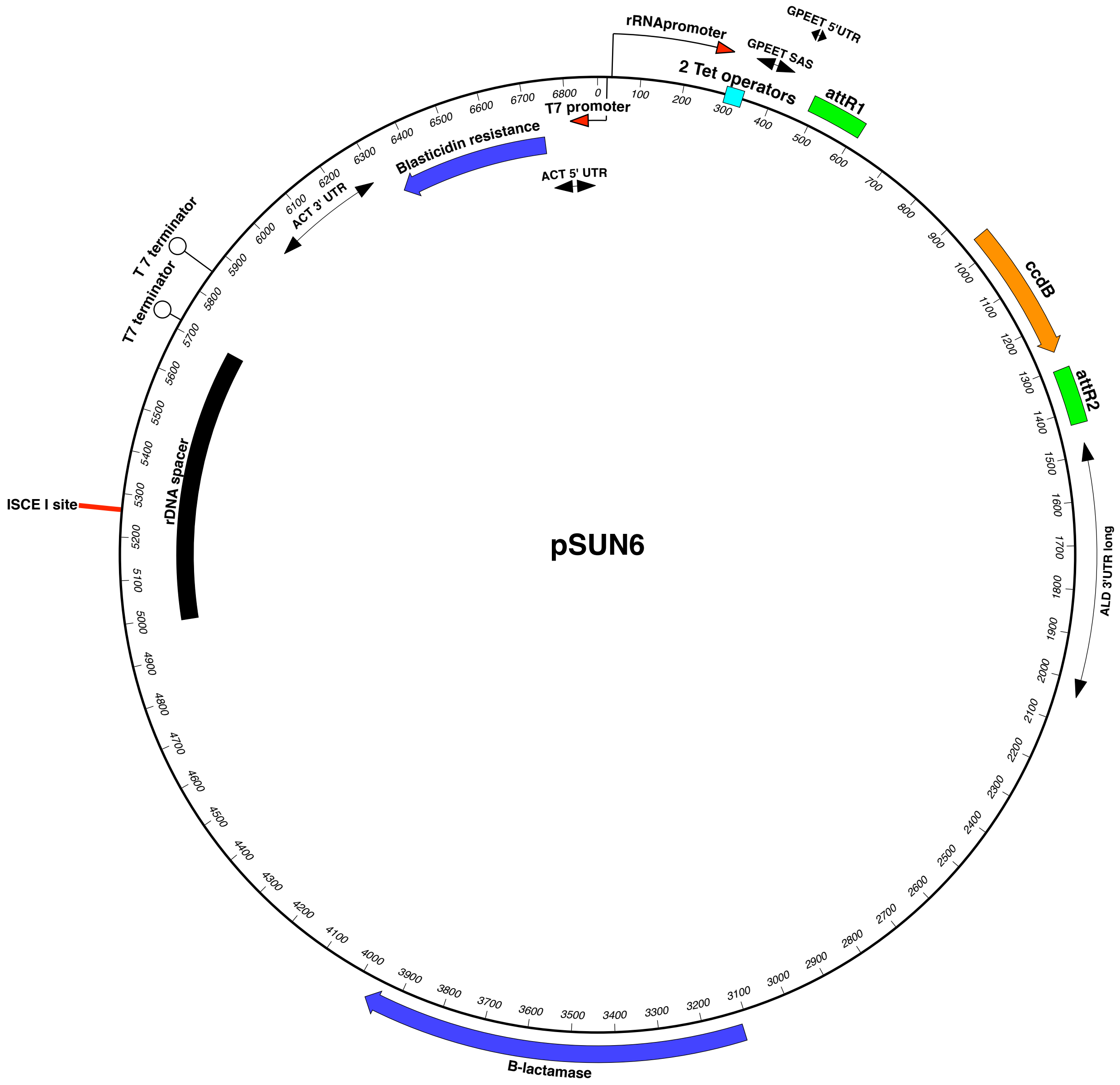

### SUP. 6

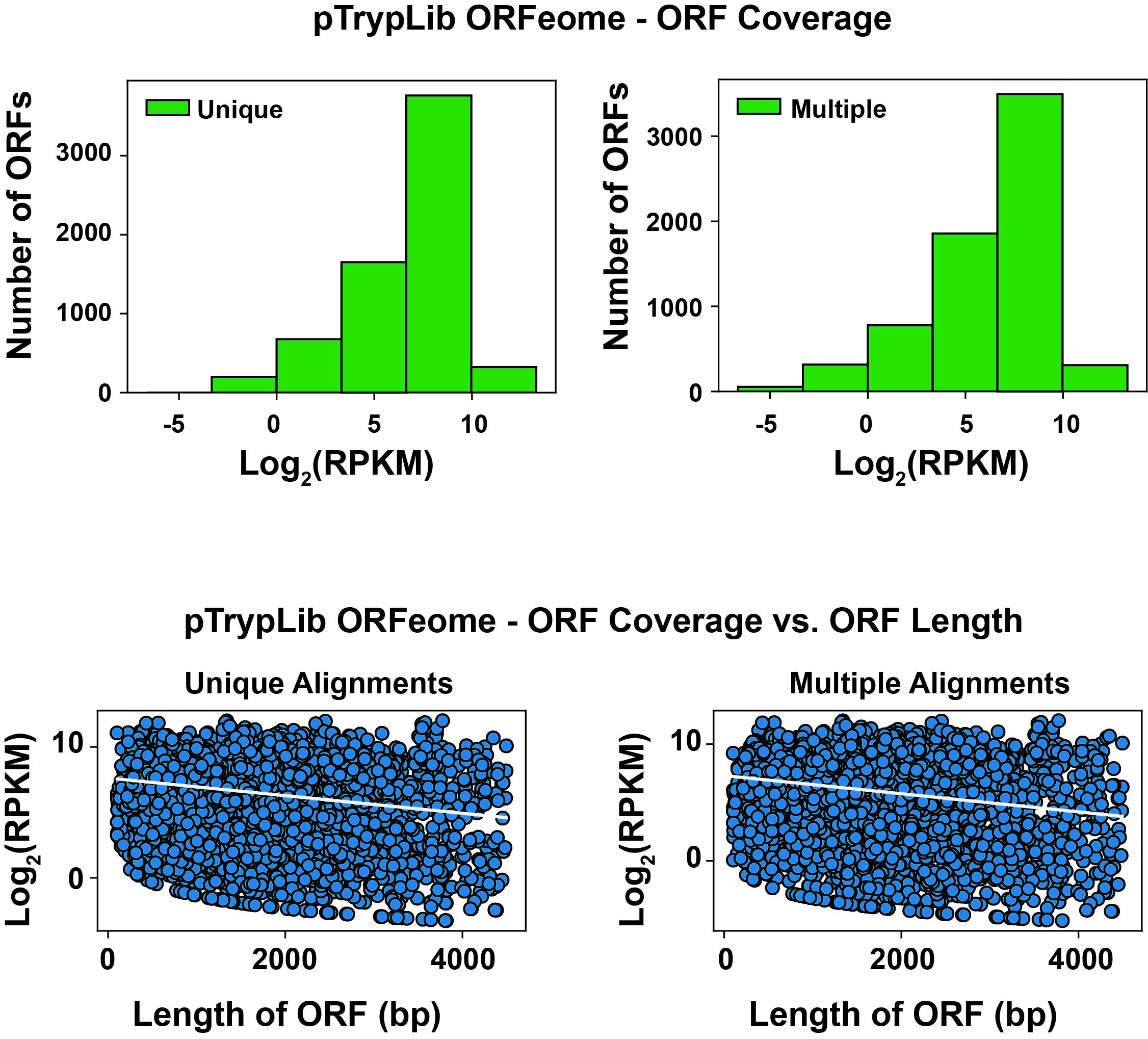

### SUP. 7

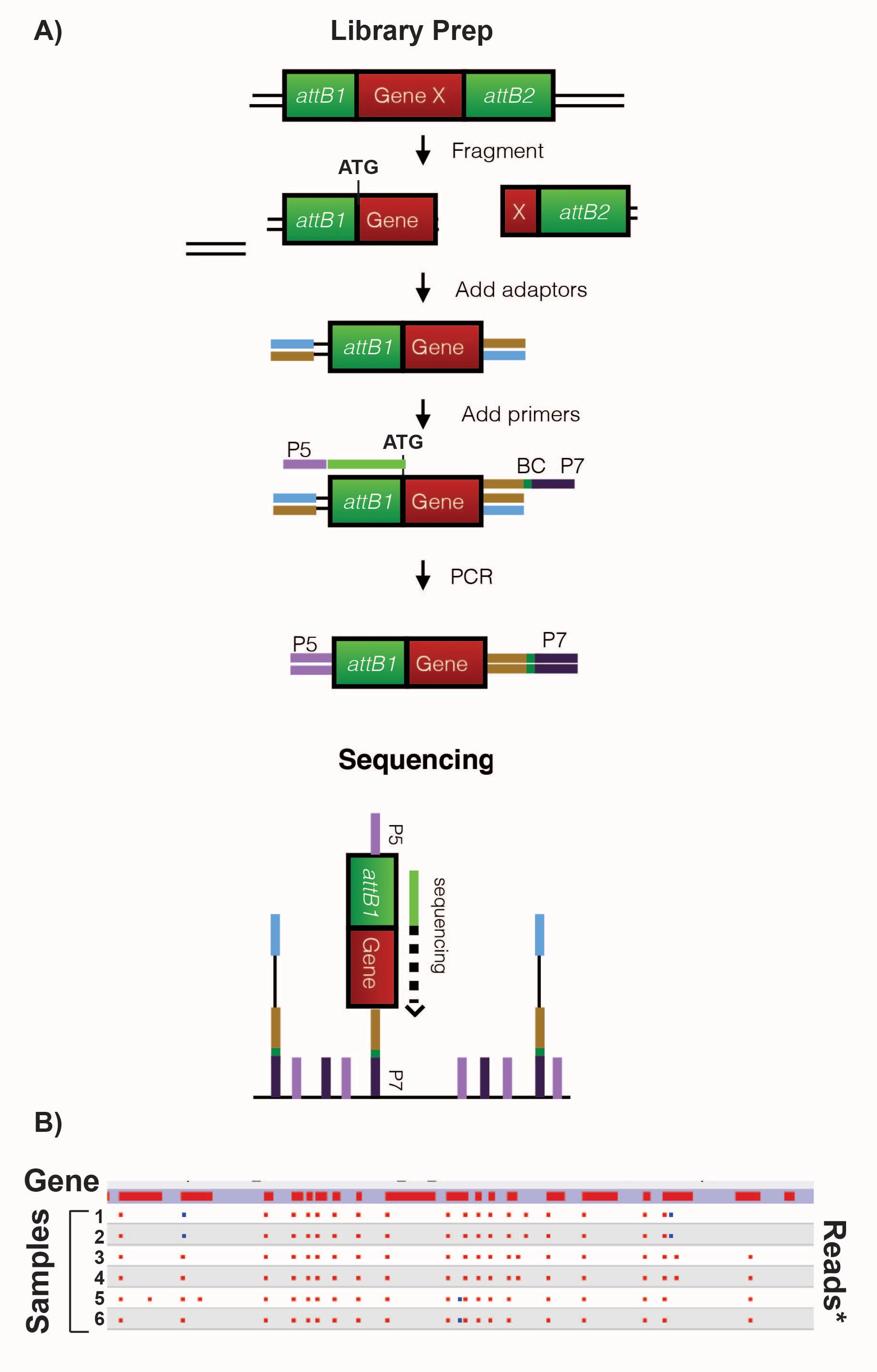

### SUP. 8

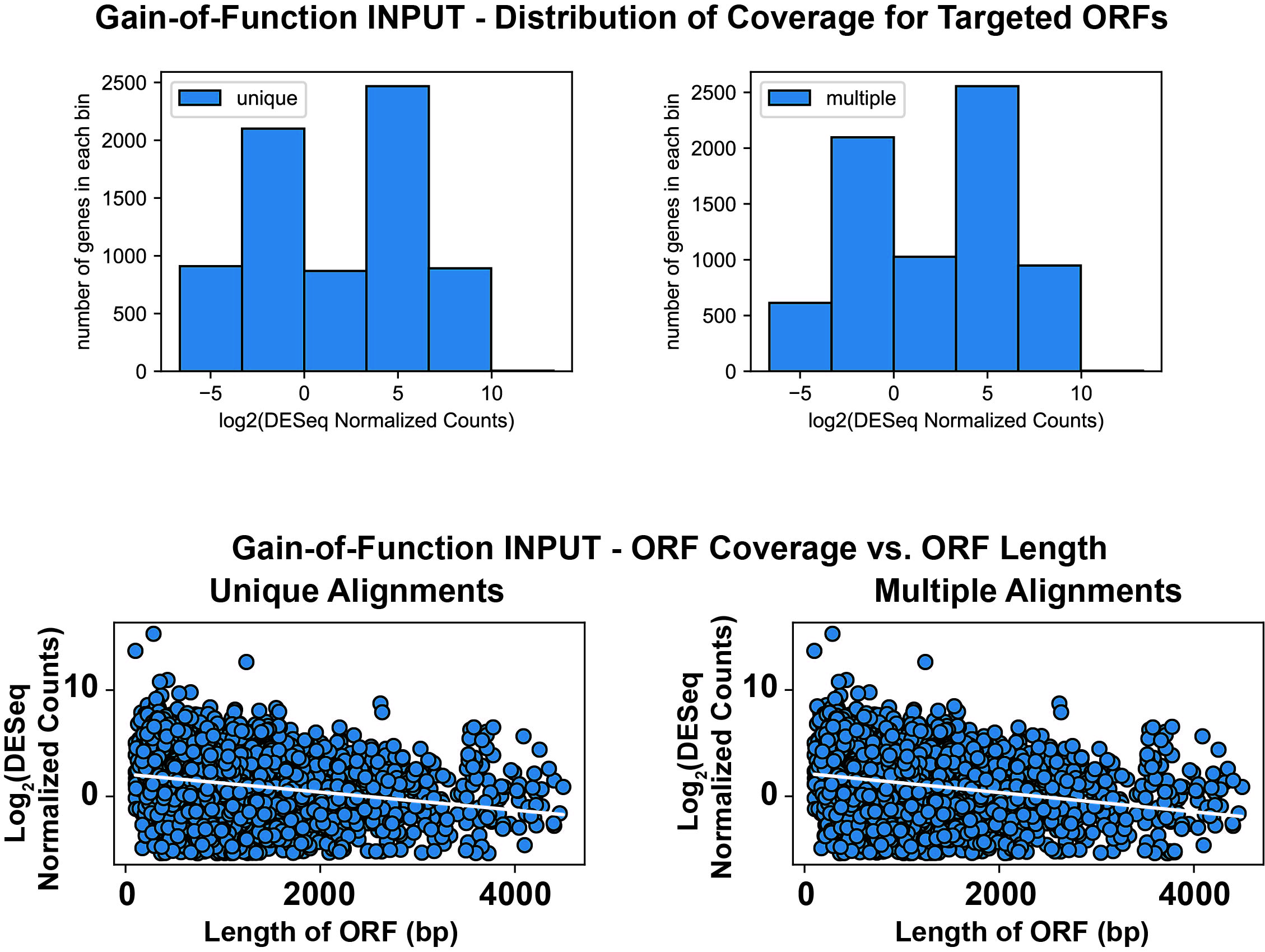
