## Supplementary material for "A *Trypanosoma brucei* ORFeome-based Gain-of-Function Library reveals novel genes associated with melarsoprol resistance": SUP. 4

| Plate name | Min length* | Max length* | Total | missing | #_MISS _PLATE | ORF# / MISS_ Plate |
|---|---|---|---|---|---|---|
| 0_hypothetical | 102 | 375 | 373 | 124 | | |
| 0_known | 147 | 591 | 315 | 51 | | |
| 1_hypothetical | 375 | 522 | 374 | 72 | | |
| 1_known | 522 | 666 | 365 | 64 | **1** | **311** |
| 2_hypothetical | 594 | 849 | 372 | 111 | | |
| 2_known | 849 | 1056 | 363 | 77 | | |
| 3_hypothetical | 669 | 822 | 382 | 89 | **2** | **277** |
| 3_known | 822 | 993 | 367 | 226 | | |
| 4_hypothetical | 993 | 1155 | 372 | 105 | **3** | **331** |
| 4_known | 1056 | 1287 | 348 | 171 | | |
| 5_hypothetical | 1287 | 1524 | 365 | 104 | | |
| 5_known | 1158 | 1365 | 342 | 94 | **4** | **369** |
| 6_hypothetical | 1524 | 1857 | 375 | 158 | | |
| 6_known | 1368 | 1635 | 362 | 173 | **5** | **331** |
| 7_hypothetical | 1857 | 2337 | 378 | 106 | | |
| 7_known | 1635 | 1953 | 376 | 117 | | |
| 8_hypothetical | 1953 | 2508 | 381 | 147 | **6** | **370** |
| 9_hypothetical | 2340 | 3504 | 381 | 195 | | |
| 10_hypothetical | 2508 | 3501 | 381 | 141 | **7** | **336** |
| hypothetical_last | 3504 | 4488 | 135 | 58 | | |
| known_last | 3507 | 4497 | 138 | 67 | **8** | **125** |
| NEG_PICKS_#1 | 102 | 1365 | 252 | 122 | | |
| NEG_PICKS_#2 | 1857 | 4497 | 177 | 79 | | |
| 1_MISS | 102 | 666 | 311 | | | |
| 2_MISS | 594 | 822 | 277 | | | |
| 3_MISS | 822 | 1155 | 331 | | | |
| 4_MISS | 1056 | 1365 | 369 | | | |
| 5_MISS | 1524 | 1635 | 331 | | | |
| 6_MISS | 1857 | 2508 | 370 | | | |
| 7_MISS | 2340 | 3501 | 336 | | | |
| 8_MISS | 3504 | 4497 | 125 | | | |

**21 Pools** of original ORFs from oligo plates.
**2 Pools** of PCR NEG_PICKs_Redo PCR reactions

**8 Pools** of MISSING ORFs from original 21 PCR plates.
Were identified as "Missing" from first ORFeome assesments (pENTR & pDEST). Isolated from PCR plates to form new sets of ORFS for an additional 8 gateway clonings.

**31 Total Pools for final pENTR followed by pDEST (pSUN6) transfections**
