## Supplementary material for "A *Trypanosoma brucei* ORFeome-based Gain-of-Function Library reveals novel genes associated with melarsoprol resistance": SUP. 10

| GENE ID | CATEGORY | DESCRIPTION | LOCALIZATION | FOLD CHANGE | P ADJUSTED |
|---|---|---|---|---|---|
| *Tb927.4.4810* | Endocytic+ | hypothetical protein, conserved | cytoplasm(points, weak), endocytic | 597 | 3.77E-42 |
| *Tb927.11.590* | Mitochondrial* | hypothetical protein, conserved | Strong mitochondrial signal, some enrichment around the kinetoplast | 350 | 1.81E-39 |
| *Tb927.7.2780* | Gene Expression# | hypothetical protein, conserved | ND | 322 | 2.83E-86 |
| *Tb927.10.12370* | Biosynthetic# | gamma-glutamylcysteine synthetase (GCS) | nucleoplasm | 191 | 2.19E-43 |
| *Tb927.5.3450* | Gene Expression# | hypothetical protein, conserved | Strong cytoplasm signal with a tendency to be patchy (quite similar to ribosomal proteins) | 126 | 4.50E-25 |
| *Tb927.11.11435* | Flagellar+ | dynein light chain lc6 | Strong flagellum axoneme signal | 120 | 1.30E-26 |
| *Tb927.7.6190* | Endocytic+ | Ring finger domain containing protein, putative | Strong endocytic system signal | 113 | 3.13E-40 |
| *Tb927.11.12380* | Hypothetical | hypothetical protein, conserved | nuclear, mass spec | 97 | 5.89E-48 |
| *Tb927.6.1280* | Gene Expression# | translation initiation factor EIF-2b alpha subunit, putative | cytoplasm | 81 | 1.14E-48 |
| *Tb927.8.3820* | Stress Granule* | Stress granule protein | Localisation to starvation stress granules | 53 | 2.04E-39 |
| *Tb927.3.5250* | Gene Expression# | Zinc finger CCCH domain-containing protein 8 (ZC3H8) | cytoplasm(points) | 53 | 2.45E-13 |
| *Tb927.9.10930* | Gene Expression# | Mediator of RNA polymerase II transcription subunit 7 (MED-T7) | nucleoplasm(points) | 49 | 1.98E-18 |
| *Tb927.7.770* | Zinc finger | Ring finger domain containing protein, putative | Moderate nuclear lumen signal, weak cytoplasm signal | 47 | 7.14E-27 |
| *Tb927.9.7820* | Hypothetical | hypothetical protein, conserved | Nuclear, mass spec | 39 | 3.88E-14 |
| *Tb927.11.15280* | Biosynthetic | tRNA-sepcific adenosine deaminase (ADAT3) | cytoplasm(reticulated, weak) | 37 | 1.59E-18 |
| *Tb927.4.1910* | Gene Expression# | hypothetical protein, conserved | cytoplasm(points, reticulated) | 35 | 1.10E-22 |
| *Tb927.9.15020* | Flagellar+ | hypothetical protein, conserved | Weak axoneme signal, and weak reticulated signal through the cytoplasm | 31 | 4.45E-20 |
| *Tb927.10.2830* | Endocytic+ | hypothetical protein, conserved | endocytic, cytoplasm | 31 | 9.17E-31 |
| *Tb927.10.12050* | Mitochondrial* | hypothetical protein, conserved | kinetoplast, mitochondrion | 31 | 2.16E-34 |
| *Tb927.9.7200* | Mitochondrial* | hypothetical protein, conserved | kinetoplast, mitochondrion | 31 | 5.61E-28 |
| *Tb927.7.5460* | Endo/ Exo | exosome-associated protein 3,3' exoribonuclease, putative (EAP3) | Nucleoplasm, tendency to be a little punctate, tendency to be around the nucleolar periphery | 28 | 9.55E-27 |
| *Tb927.4.4540* | Zinc finger | zinc finger domain, LSD1 subclass, putative | cytoplasm(reticulated) | 25 | 8.84E-47 |
| *Tb927.5.4370* | Hypothetical | hypothetical protein, conserved | endocytic, cytoplasm | 24 | 3.59E-12 |
| *Tb927.9.7080* | Mitochondrial & ER○ | hypothetical protein, conserved | cytoplasm(patchy, points) | 23 | 1.14E-23 |
| *Tb927.2.5210* | Mitochondrial# | 3-oxoacyl-ACP reductase, putative | mitochondrion, kinetoplast(strong) | 23 | 2.16E-34 |
| *Tb927.11.2910* | Mitochondrial* | phosphoglycerate mutase, putative (iPGAM) | mitochondrion(75%), kinetoplast(75%), cytoplasm(weak, 25%) | 23 | 6.79E-05 |
| *Tb927.11.7475* | Zinc finger# | AN1-like Zinc finger containing protein, putative | Cytoplasm, nuclear lumen and flagellum cytoplasm. Tendency to be enriched in the nucleolus | 20 | 1.50E-30 |
| *Tb927.9.5220* | Mitochondrial○ | conserved protein | endocytic, cytoplasm(weak) | 19 | 2.60E-21 |
| *Tb927.3.4930* | Hypothetical | hypothetical protein, conserved | cytoplasm, flagellar cytoplasm, | 19 | 3.79E-27 |
| *Tb927.8.1930* | Gene Expression# | Isy1-like splicing family | nucleoplasm | 16 | 8.36E-27 |
| *Tb927.10.6850* | Mitochondrial# | Mitochondrial ribosomal protein S18, putative | cytoplasm(reticulated) | 16 | 3.68E-10 |
| *Tb927.11.1810* | Flagellar + | Ring finger domain containing protein, putative | flagellar pocket(ring) | 16 | 1.50E-18 |
| *Tb927.5.2620* | Gene Expression | hypothetical protein, conserved | cytoplasm(points) | 15 | 1.78E-09 |
| *Tb927.10.390* | Flagellar* | hypothetical protein, conserved | Flagellum Matrix Proteome (BSF) | 14 | 2.78E-12 |
| *Tb927.3.1610* | Kinase | protein kinase, putative | nucleoplasm | 14 | 2.82E-35 |
| *Tb927.11.2350* | Hypothetical | hypothetical protein, conserved | nucleus, cytoplasm(reticulated) | 13 | 2.19E-05 |
| *Tb927.8.7790* | Mitochondrial○ | zinc finger domain, LSD1 subclass, putative | ND | 13 | 7.88E-22 |
| *Tb927.5.4150* | Flagellar* | hypothetical protein, conserved | paraflagellar rod | 13 | 6.29E-06 |
| *Tb927.8.3340* | Hypothetical | hypothetical protein, conserved | nucleoplasm | 13 | 1.28E-11 |
| *Tb927.10.1490* | Gene Expression | Temperature dependent protein affecting M2 dsRNA replication, putative | cytoplasm(weak) | 12 | 6.62E-16 |
| *Tb927.4.890* | Gene Expression | small nuclear ribonucleoprotein SmD3, putative (SmD3) | nucleoplasm | 11 | 4.78E-11 |
| *Tb927.7.5360* | Pathogenesis | Haemolysin-III related, putative | cytoplasm | 10 | 8.31E-27 |
| *Tb927.7.710* | Flagellar * | heat shock 70 kDa protein, putative (HSP70) | cell tip(anterior), cytoplasm, flagellar cytoplasm | 10 | 3.15E-26 |
| *Tb927.9.4930* | Metabolic / biosynth | divalent cation transporter, putative | cytoplasm(reticulated) | 9 | 7.16E-54 |
| *Tb927.9.10850* | Gene Expression | Splicing factor 3B subunit 10 (SF3b10), putative | nucleoplasm | 9 | 1.80E-49 |
| *Tb927.11.5600* | Mitochondrial# | Archaic Translocase of outer membrane 14 kDa subunit | mitochondrion | 8 | 9.21E-11 |
| *Tb927.10.9060* | Flagellar + | hypothetical protein, conserved | basal body | 8 | 5.86E-10 |
| *Tb927.1.1020* | Flagellar + | leucine-rich repeat-containing protein | hook complex | 8 | 1.61E-10 |
| *Tb927.2.2130* | Trafficking | small GTP-binding protein RAB6, putative | golgi apparatus | 8 | 7.70E-22 |
| *Tb927.1.3310* | Hypothetical | hypothetical protein, conserved | plasma membrane(posterior) | 8 | 2.28E-09 |
| *Tb927.8.4200* | Gene Expression | hypothetical protein, conserved | cytoplasm | 7 | 7.02E-19 |
| *Tb927.3.5190* | Mitochondrial+ | hypothetical protein, conserved | mitochondrion | 7 | 8.48E-21 |
| *Tb927.9.3480* | Gene Expression | U5Cwc21 small nuclear ribonucleoprotein (CWC21) | nucleoplasm | 7 | 8.83E-10 |
| *Tb927.8.2391* | Hypothetical | hypothetical protein, conserved | ND | 7 | 9.18E-10 |
| *Tb927.2.3780* | Gene Expression | translation initiation factor IF-2, putative | cytoplasm | 6 | 1.12E-11 |
| *Tb927.1.1500* | Gene Expression | conserved protein, unknown function | cytoplasm(reticulated) | 6 | 1.34E-05 |
| *Tb927.6.3980* | Flagellar + | hypothetical protein, conserved | Flagellum, axoneme | 6 | 3.88E-10 |

Key: + From localization data, ○ Proteomic data, * both localization and proteomics, or # published
