## Supplementary material for "A *Trypanosoma brucei* ORFeome-based Gain-of-Function Library reveals novel genes associated with melarsoprol resistance": SUP. 11

| Protein Abb | Gene | INPUT2_1 | INPUT2_2 | INPUT2_3 | INPUT1_1 | INPUT1_2 | INPUT1_3 | MEL-1x_1 | MEL-1x_2 | MEL-1x_3 | MEL-2x_1 | MEL-2x_2 | MEL-2x_3 | INPUT2_AVG | INPUT1_AVG | MEL-1x_AVG | MEL-2x_AVG | Fold MEL-1x over INPUT2 |
|---|---|---|---|---|---|---|---|---|---|---|---|---|---|---|---|---|---|---|
| **TryS** | *Tb927.2.4370* | 0.000 | 1.181 | 0.998 | 0.000 | 0.486 | 0.000 | 2.276 | 0.000 | 0.000 | 0.000 | 0.000 | 0.000 | 0.726 | 0.162 | 0.759 | 0.000 | 1.044 |
| **TR** | *Tb927.10.10390* | 55.661 | 63.200 | 45.892 | 55.116 | 50.107 | 49.140 | 122.885 | 27.305 | 0.827 | 100.379 | 6.929 | 19.339 | 54.918 | 51.454 | 50.339 | 42.216 | 0.917 |
| **GSH1** | *Tb927.10.12370* | 51.637 | 52.568 | 47.887 | 63.503 | 40.378 | 43.433 | 9211.795 | 5615.217 | 14158.814 | 16159.151 | 32023.616 | 7464.779 | 50.698 | 49.105 | 9661.942 | 18549.182 | 190.580 |
| **GSH2** | *Tb927.7.4000* | 0.671 | 0.000 | 0.000 | 0.599 | 0.486 | 0.000 | 2.276 | 2.409 | 0.000 | 0.000 | 0.000 | 0.000 | 0.224 | 0.362 | 1.562 | 0.000 | 6.986 |
| **ODC** | *Tb927.11.13730* | 0.000 | 0.591 | 0.000 | 0.000 | 0.000 | 0.000 | 3.413 | 1.606 | 0.000 | 1.930 | 0.000 | 1.381 | 0.197 | 0.000 | 1.673 | 1.104 | 8.498 |
| **SpS** | *Tb927.9.7770* | 185.090 | 160.657 | 154.636 | 188.113 | 156.160 | 142.347 | 93.301 | 160.618 | 220.042 | 71.424 | 193.999 | 142.278 | 166.794 | 162.207 | 157.987 | 135.900 | 0.947 |
| **Prx** | *Tb927.8.1990* | 63.038 | 53.749 | 58.861 | 64.701 | 54.972 | 64.357 | 103.542 | 44.973 | 0.827 | 169.873 | 0.000 | 52.491 | 58.550 | 61.344 | 49.781 | 74.121 | 0.850 |
| **Px** | *Tb927.11.15920* | 0.000 | 0.591 | 0.000 | 0.000 | 0.000 | 0.000 | 6.827 | 0.000 | 1.654 | 0.000 | 0.000 | 0.000 | 0.197 | 0.000 | 2.827 | 0.000 | 14.359 |
| **RR** | *Tb927.11.7840* | 0.671 | 0.591 | 0.998 | 0.000 | 0.000 | 0.000 | 3.413 | 0.803 | 0.000 | 0.000 | 6.929 | 1.381 | 0.753 | 0.000 | 1.406 | 2.770 | 1.867 |
| **RR** | *Tb927.11.12790* | 0.000 | 0.000 | 0.000 | 0.000 | 0.000 | 0.000 | 0.000 | 0.000 | 0.000 | 0.000 | 0.000 | 0.000 | 0.000 | 0.000 | 0.000 | 0.000 | |
| **UMSBP** | *Tb927.10.6070* | 0.671 | 0.591 | 0.998 | 0.000 | 0.000 | 0.317 | 7.965 | 0.000 | 0.827 | 0.000 | 0.000 | 2.763 | 0.753 | 0.106 | 2.931 | 0.921 | 3.892 |
| **1-c-Grx1** | *Tb927.9.3590* | 145.523 | 145.300 | 156.631 | 162.353 | 169.295 | 173.416 | 92.163 | 105.205 | 36.398 | 167.942 | 83.142 | 93.931 | 149.152 | 168.355 | 77.922 | 115.005 | 0.522 |
| | *Tb927.9.5770* | gene not in data | | | | | | | | | | | | | | | | |
| | *Tb927.9.5860* | gene not in data | | | | | | | | | | | | | | | | |
| | *Tb927.7.1120* | 0.000 | 1.772 | 0.998 | 0.599 | 0.000 | 0.000 | 2.276 | 0.803 | 2.482 | 0.000 | 0.000 | 0.000 | 0.923 | 0.200 | 1.853 | 0.000 | 2.008 |
| | *Tb927.7.1130* | 0.671 | 0.000 | 0.000 | 0.000 | 0.000 | 0.000 | 1.138 | 0.000 | 0.827 | 0.000 | 0.000 | 0.000 | 0.224 | 0.000 | 0.655 | 0.000 | 2.930 |
| | *Tb927.7.1140* | 254.163 | 232.717 | 271.361 | 257.008 | 287.023 | 235.554 | 1030.865 | 774.180 | 392.106 | 1523.064 | 512.710 | 1006.999 | 252.747 | 259.862 | 732.384 | 1014.258 | 2.898 |
| | *Tb927.10.10420* | 0.000 | 0.000 | 0.000 | 0.000 | 0.000 | 0.634 | 1.138 | 1.606 | 0.827 | 0.000 | 6.929 | 0.000 | 0.000 | 0.211 | 1.190 | 2.310 | |
| | *Tb927.5.950* | 4.694 | 8.269 | 8.979 | 4.194 | 4.378 | 6.658 | 5.689 | 0.000 | 0.827 | 1.930 | 0.000 | 0.000 | 7.314 | 5.077 | 2.172 | 0.643 | 0.297 |
| | *Tb927.4.1350* | 23.472 | 18.901 | 17.958 | 16.175 | 14.594 | 15.535 | 25.032 | 6.425 | 0.827 | 19.304 | 0.000 | 5.525 | 20.110 | 15.435 | 10.761 | 8.276 | 0.535 |
| **Trx** | *Tb927.9.3370* | 172.348 | 161.248 | 214.495 | 160.555 | 170.755 | 211.460 | 153.606 | 118.054 | 156.346 | 129.335 | 131.642 | 117.414 | 182.697 | 180.923 | 142.669 | 126.130 | 0.781 |
