## Supplementary material for "A *Trypanosoma brucei* ORFeome-based Gain-of-Function Library reveals novel genes associated with melarsoprol resistance": SUP. 12

| | | Occurance of Melarsoprol GoF Hits in RIT-Seq screens | | | | | |
|---|---|---|---|---|---|---|---|
| **Gene Name** | **RIT-Seq Screens, descriptive** | **RIT-seq PCF** | **RIT-seq BSF** | **PCF& BSF** | **Melarsoprol** | **Nif** | **APOLI** |
| *Tb927.4.4810* | | | | | | | |
| *Tb927.11.590* | | | | | | | |
| *Tb927.7.2780* | significant loss of fitness in all four experiments | 1 | 1 | 1 | | | |
| *Tb927.10.12370* | Identified as lethal in pro. / drug target | 1 | | | | | |
| *Tb927.5.3450* | significant loss of fitness in the DIF experiment only | 1 | 1 | 1 | | | |
| *Tb927.11.11435* | | | | | | | |
| *Tb927.7.6190* | | | | | | | |
| *Tb927.11.12380* | | | | | | | |
| *Tb927.6.1280* | | | | | | | |
| *Tb927.8.3820* | significant loss of fitness in all four experiments | 1 | 1 | 1 | | | |
| *Tb927.3.5250* | significant loss of fitness in BF experiments only/ ZFP Family member | | 1 | | | | |
| *Tb927.9.10930* | | | | | | | |
| *Tb927.7.770* | | | | | | | |
| *Tb927.9.7820* | | | | | | | |
| *Tb927.11.15280* | no significant loss of fitness in any experiment, Sensitivity to APOLI RNAi screen | | | | | | 1 |
| *Tb927.4.1910* | | | | | | | |
| *Tb927.9.15020* | | | | | | | |
| *Tb927.10.2830* | | | | | | | |
| *Tb927.10.12050* | Identified in Mel. Screen/Identified as non-essential, no loss of fitness when KD | | | | 1 | | |
| *Tb927.9.7200* | | | | | | | |
| *Tb927.7.5460* | Identified as lethal in pro./ exosome complex | 1 | | | | | |
| *Tb927.4.4540* | | | | | | | |
| *Tb927.5.4370* | significant loss of fitness in all four experiments | 1 | 1 | 1 | | | |
| *Tb927.9.7080* | | | | | | | |
| *Tb927.2.5210* | | | | | | | |
| *Tb927.11.2910* | | | | | | | |
| *Tb927.11.7475* | | | | | | | |
| *Tb927.9.5220* | significant loss of fitness in the DIF experiment only | 1 | 1 | 1 | | | |
| *Tb927.3.4930* | | | | | | | |
| *Tb927.8.1930* | significant loss of fitness in bloodstream form experiments | | 1 | | | | |
| *Tb927.10.6850* | | | | | | | |
| *Tb927.11.1810* | | | | | | | |
| *Tb927.5.2620* | significant loss of fitness in procyclic form experiments | 1 | | | | | |
| *Tb927.10.390* | Identified in Nif Screen (low sig) | | | | | 1 | |
| *Tb927.3.1610* | Identified in Nif Screen (low sig)/ significant loss in differentiation experiments | 1 | 1 | 1 | | 1 | |
| *Tb927.11.2350* | | | | | | | |
| *Tb927.8.7790* | significant loss in all 4 experiments | 1 | 1 | 1 | | | |
| *Tb927.5.4150* | significant loss in differentiation experiments | 1 | 1 | 1 | | | |
| *Tb927.8.3340* | | | | | | | |
| *Tb927.10.1490* | HIGH melarsoprol hit/ Significant loss in bloodstream form experiments | | 1 | | 1 | | |
| *Tb927.4.890* | | | | | | | |
| *Tb927.7.5360* | | | | | | | |
| *Tb927.7.710* | | | | | | | |
| *Tb927.9.4930* | | | | | | | |
| *Tb927.9.10850* | | | | | | | |
| *Tb927.11.5600* | significant loss of fitness in all four experiments | 1 | 1 | 1 | | | |
| *Tb927.10.9060* | significant loss of fitness in procyclic form experiments | 1 | | | | | |
| *Tb927.1.1020* | significant loss of fitness in differentiation experiments | 1 | 1 | 1 | | | |
| *Tb927.2.2130* | | | | | | | |
| *Tb927.1.3310* | | | | | | | |
| *Tb927.8.4200* | Identified in Nif Screen (low sig) | | | | | 1 | |
| *Tb927.3.5190* | | | | | | | |
| *Tb927.9.3480* | | | | | | | |
| *Tb927.8.2391* | | | | | | | |
| *Tb927.2.3780* | significant loss of fitness in procyclic form experiments | 1 | | | | | |
| *Tb927.1.1500* | | | | | | | |
| *Tb927.6.3980* | Identified in Nif Screen (low sig) | | | | | 1 | |
| | Total = | 15 | 13 | 10 | 2 | 4 | 1 |
| | Percent = | 26.3% | 22.8% | 17.5% | 3.5% | 7.0% | 1.8% |
