## Supplementary material for "A *Trypanosoma brucei* ORFeome-based Gain-of-Function Library reveals novel genes associated with melarsoprol resistance": SUP. 13

| | | *Trypanosoma* | | | | | | | *Leishmania* | | | | | | *Crithidia* |
|---|---|---|---|---|---|---|---|---|---|---|---|---|---|---|---|
| Gene ID | Ortholog Group # | *brucei brucei* | *congolense* | *gambiense* | *cruzi* | *viva1* | *grayi* | *evansi* | *braziliensis* | *major* | *donovani* | *amazonensis* | *infantum* | *me1icana* | *fasciculata* |
| *Tb927.4.4810* | OG5_206657 | 1 | | 1 | | | | 1 | | | | | | | |
| *Tb927.11.590* | OG5_151682 | 1 | 1 | 1 | 1 | 1 | 1 | 1 | 1 | 1 | 1 | 1 | 1 | 1 | 1 |
| *Tb927.7.2780* | OG5_148621 | 1 | 1 | 1 | 1 | 1 | 1 | 1 | 1 | 1 | 1 | 1 | 1 | 1 | 1 |
| *Tb927.10.12370* | OG5_128698 | 1 | 1 | 1 | 1 | 1 | 1 | 1 | 1 | 1 | 1 | 1 | 1 | 1 | 1 |
| *Tb927.5.3450* | OG5_128129 | 1 | 1 | 1 | 1 | | 1 | 1 | 1 | 1 | 1 | 1 | 1 | 1 | 1 |
| *Tb927.11.11435* | OG5_173500 | 1 | | | 1 | | 1 | | 1 | 1 | 1 | 1 | 1 | 1 | 1 |
| *Tb927.7.6190* | OG5_140484 | 1 | 1 | 1 | 1 | 1 | 1 | 1 | 1 | 1 | 1 | 1 | 1 | 1 | 1 |
| *Tb927.11.12380* | OG5_145874 | 1 | 1 | 1 | 1 | 1 | 1 | 1 | 1 | 1 | 1 | 1 | 1 | 1 | 1 |
| *Tb927.6.1280* | OG5_127805 | 1 | 1 | 1 | 1 | 1 | 1 | 1 | 1 | 1 | 1 | 1 | 1 | 1 | 1 |
| *Tb927.8.3820* | OG5_145191 | 1 | 1 | 1 | 1 | 1 | 1 | 1 | 1 | 1 | 1 | 1 | 1 | 1 | 1 |
| *Tb927.3.5250* | OG5_173124 | 1 | 1 | 1 | 1 | 1 | 1 | 1 | | | | | | | |
| *Tb927.9.10930* | OG5_154727 | 1 | | 1 | 1 | | 1 | 1 | 1 | 1 | 1 | 1 | 1 | 1 | 1 |
| *Tb927.7.770* | OG5_148730 | 1 | 1 | 1 | 1 | 1 | 1 | 1 | 1 | 1 | 1 | 1 | 1 | 1 | 1 |
| *Tb927.9.7820* | OG5_162387 | 1 | 1 | 1 | 1 | 1 | 1 | 1 | | | | | | | |
| *Tb927.11.15280* | OG5_149647 | 1 | 1 | 1 | 1 | 1 | 1 | 1 | 1 | 1 | 1 | 1 | 1 | 1 | 1 |
| *Tb927.4.1910* | OG5_162290 | 1 | 1 | 1 | 1 | 1 | 1 | 1 | | | | | | | |
| *Tb927.9.15020* | OG5_162404 | 1 | 1 | 1 | 1 | 1 | 1 | 1 | | | | | | | |
| *Tb927.10.2830* | OG5_154560 | 1 | 1 | 1 | 1 | 1 | 1 | 1 | 1 | 1 | 1 | 1 | 1 | 1 | 1 |
| *Tb927.10.12050* | OG5_148943 | 1 | 1 | 1 | 1 | 1 | 1 | 1 | 1 | 1 | 1 | 1 | 1 | 1 | 1 |
| *Tb927.9.7200* | OG5_151433 | 1 | 1 | 1 | 1 | 1 | 1 | 1 | 1 | 1 | 1 | 1 | 1 | 1 | 1 |
| *Tb927.7.5460* | OG5_147017 | 1 | 1 | 1 | 1 | 1 | 1 | 1 | 1 | 1 | 1 | 1 | 1 | 1 | 1 |
| *Tb927.4.4540* | OG5_137522 | 1 | | 1 | | | | 1 | | | | | | | |
| *Tb927.5.4370* | OG5_151442 | 1 | | 1 | 1 | 1 | 1 | 1 | 1 | 1 | 1 | 1 | 1 | 1 | 1 |
| *Tb927.9.7080* | OG5_154530 | 1 | | 1 | 1 | 1 | 1 | 1 | 1 | 1 | 1 | 1 | 1 | 1 | 1 |
| *Tb927.2.5210* | OG5_126618 | 1 | 1 | 1 | 1 | 1 | 1 | 1 | 1 | 1 | 1 | 1 | 1 | 1 | 1 |
| *Tb927.11.2910* | OG5_147168 | 1 | 1 | | 1 | 1 | 1 | 1 | 1 | 1 | 1 | 1 | 1 | 1 | 1 |
| *Tb927.11.7475* | OG5_146402 | 1 | | | 1 | | | | 1 | 1 | 1 | 1 | 1 | 1 | 1 |
| *Tb927.9.5220* | OG5_148426 | 1 | 1 | 1 | 1 | 1 | 1 | 1 | 1 | 1 | 1 | 1 | 1 | 1 | 1 |
| *Tb927.3.4930* | OG5_174188 | 1 | 1 | 1 | 1 | 1 | | 1 | | | | | | | |
| *Tb927.8.1930* | OG5_128335 | 1 | | 1 | 1 | 1 | 1 | 1 | 1 | 1 | 1 | 1 | 1 | 1 | 1 |
| *Tb927.10.6850* | OG5_151867 | 1 | 1 | 1 | 1 | 1 | 1 | 1 | 1 | 1 | 1 | 1 | 1 | 1 | 1 |
| *Tb927.11.1810* | OG5_133219 | 1 | 1 | 1 | 1 | 1 | | 1 | 1 | 1 | 1 | 1 | 1 | 1 | 1 |
| *Tb927.5.2620* | OG5_145922 | 1 | 1 | 1 | 1 | 1 | 1 | 1 | 1 | 1 | 1 | 1 | 1 | 1 | 1 |
| *Tb927.10.390* | OG5_148610 | 1 | 1 | 1 | 1 | 1 | 1 | 1 | 1 | 1 | 1 | 1 | 1 | 1 | 1 |
| *Tb927.3.1610* | OG5_148710 | 1 | 1 | 1 | 1 | 1 | 1 | 1 | 1 | 1 | 1 | 1 | 1 | 1 | 1 |
| *Tb927.11.2350* | OG5_148508 | 1 | 1 | 1 | 1 | 1 | 1 | 1 | 1 | 1 | 1 | 1 | 1 | 1 | 1 |
| *Tb927.8.7790* | OG5_143050 | 1 | 1 | 1 | 1 | | | 1 | 1 | 1 | 1 | 1 | 1 | 1 | 1 |
| *Tb927.5.4150* | OG5_148907 | 1 | 1 | 1 | 1 | 1 | 1 | 1 | 1 | 1 | 1 | 1 | 1 | 1 | 1 |
| *Tb927.8.3340* | OG5_185219 | 1 | | 1 | 1 | | 1 | 1 | | | | | | | 1 |
| *Tb927.10.1490* | OG5_139047 | 1 | 1 | 1 | 1 | 1 | 1 | 1 | 1 | 1 | 1 | 1 | 1 | 1 | 1 |
| *Tb927.4.890* | OG5_127840 | 1 | 1 | 1 | 1 | 1 | 1 | 1 | 1 | 1 | 1 | 1 | 1 | 1 | 1 |
| *Tb927.7.5360* | OG5_163796 | 1 | 1 | 1 | 1 | | 1 | 1 | | | | | | | |
| *Tb927.7.710* | OG5_126588 | 1 | 1 | 1 | 1 | 1 | 1 | 1 | 1 | 1 | 1 | 1 | 1 | 1 | 1 |
| *Tb927.9.4930* | OG5_146815 | 1 | 1 | 1 | 1 | 1 | 1 | 1 | 1 | 1 | 1 | 1 | 1 | 1 | 1 |
| *Tb927.9.10850* | OG5_128788 | 1 | 1 | 1 | 1 | 1 | 1 | 1 | 1 | 1 | 1 | 1 | 1 | 1 | 1 |
| *Tb927.11.5600* | OG5_148673 | 1 | 1 | 1 | 1 | 1 | 1 | 1 | 1 | 1 | 1 | 1 | 1 | 1 | 1 |
| *Tb927.10.9060* | OG5_173560 | 1 | 1 | 1 | | 1 | | 1 | | | | | | | |
| *Tb927.1.1020* | OG5_15192 | 1 | 1 | 1 | 1 | 1 | 1 | 1 | 1 | 1 | 1 | 1 | 1 | 1 | 1 |
| *Tb927.2.2130* | OG5_127256 | 1 | 1 | 1 | 1 | 1 | 1 | 1 | 1 | 1 | 1 | 1 | 1 | 1 | 1 |
| *Tb927.1.3310* | OG5_157931 | 1 | 1 | 1 | 1 | 1 | 1 | 1 | 1 | 1 | 1 | 1 | 1 | 1 | 1 |
| *Tb927.8.4200* | OG5_185221 | 1 | 1 | 1 | | | | 1 | | | | | | | |
| *Tb927.3.5190* | OG5_151752 | 1 | 1 | 1 | 1 | 1 | 1 | 1 | 1 | 1 | 1 | 1 | 1 | 1 | 1 |
| *Tb927.9.3480* | OG5_153717 | 1 | 1 | 1 | 1 | 1 | 1 | 1 | 1 | 1 | 1 | 1 | 1 | 1 | 1 |
| *Tb927.8.2391* | OG5_tviv_ | 1 | | | | 1 | | | | | | | | | |
| *Tb927.2.3780* | OG5_127312 | 1 | 1 | 1 | 1 | 1 | 1 | 1 | 1 | 1 | 1 | 1 | 1 | 1 | 1 |
| *Tb927.1.1500* | OG5_143923 | 1 | 1 | 1 | 1 | 1 | 1 | 1 | 1 | 1 | 1 | 1 | 1 | 1 | 1 |
| *Tb927.6.3980* | OG5_146038 | 1 | 1 | 1 | 1 | 1 | 1 | 1 | 1 | 1 | 1 | 1 | 1 | 1 | 1 |
| | SUM | 57 | 47 | 53 | 52 | 47 | 48 | 54 | 45 | 45 | 45 | 45 | 45 | 45 | 46 |
| | % | 100% | 82% | 93% | 91% | 82% | 84% | 95% | 79% | 79% | 79% | 79% | 79% | 79% | 81% |
